# Physiological reprogramming *in vivo* mediated by Sox4 pioneer factor activity

**DOI:** 10.1101/2023.02.14.528556

**Authors:** Takeshi Katsuda, Jonathan Sussman, Kenji Ito, Andrew Katznelson, Salina Yuan, Jinyang Li, Allyson J. Merrell, Naomi Takenaka, Hector Cure, Qinglan Li, Reyaz Ur Rasool, Irfan A. Asangani, Kenneth S. Zaret, Ben Z. Stanger

## Abstract

Tissue damage elicits cell fate switching through a process called metaplasia, but how the starting cell fate is silenced and the new cell fate is activated has not been investigated in animals. In cell culture, pioneer transcription factors mediate “reprogramming” by opening new chromatin sites for expression that can attract transcription factors from the starting cell’s enhancers. Here we report that Sox4 is sufficient to initiate hepatobiliary metaplasia in the adult liver. In lineage-traced cells, we assessed the timing of Sox4-mediated opening of enhancer chromatin versus enhancer decommissioning. Initially, Sox4 directly binds to and closes hepatocyte regulatory sequences via a motif it overlaps with Hnf4a, a hepatocyte master regulator. Subsequently, Sox4 exerts pioneer factor activity to open biliary regulatory sequences. The results delineate a hierarchy by which gene networks become reprogrammed under physiological conditions, providing deeper insight into the basis for cell fate transitions in animals.

---

Metaplasia is an adaptive cellular response to tissue injury and involves a variety of organs, including the lung (squamous metaplasia), esophagus (intestinal metaplasia) and pancreas (acinar-ductal metaplasia)^1, 2^. Metaplasia involves the induction of multiple transcription factors^3–6^, and loss-of function studies suggest that such factors function in concert with epigenetic remodeling^3–8^. Whether the induced transcription factors are sufficient to elicit metaplasia, and how they might initiate cell fate changes *in vivo* is not known. Furthermore, how the genetic network of the starting cell fate is suppressed while the new cell fate is activated during metaplasia remains unclear. Pioneer transcription factors, by virtue of their inherent ability to target DNA on a nucleosome, elicit cell fate changes by targeting silent genes in chromatin^9^. Pioneer factors can also enable further chromatin compaction and repression^10^ and are involved in cancer progression and the expression of circadian rhythm genes^11, 12^. Pioneer factors can trigger reprogramming of one cell type to another in cultured cells – notable examples include OCT4, SOX2 and KLF4 in induced pluripotent stem cells (iPSCs)^13–16^, FOXA1/2/3 in induced hepatocytes^17^, and ASCL1 and NEUROD1 in induced neurons^18, 19^. In cultured cells, repression of the starting fate genes is thought to occur by the reprogramming factors, bound at new sites, drawing away the starting cell transcription factors from active enhancers^13, 20^. Transcription factor-mediated enhancer decommissioning has also been reported in the context of cell fate decisions^21, 22^.

We and others have reported that the liver undergoes a hepatobiliary metaplasia (“biliary reprogramming”), wherein hepatocytes are reprogrammed to become biliary epithelial cells as a conserved *in vivo* response to liver injury^23–26^. Here, we use a hepatobiliary metaplasia model to characterize the genetic cascades involved in this physiological cell fate change to understand the gene activation and repression programs responsible for the reprogramming process in animals and how they may differ from those present in cell culture. Notably, our experimental design allows us to trace and isolate the individual cells undergoing metaplasia at different time points of the process.

## Results

### Sox4 induces biliary reprogramming

We recently reported that in response to cholestatic injury induced with a 0.1% 3,5-diethoxycarbonyl-1,4-dihydrocollidine (DDC) diet, adult hepatocytes alter their chromatin landscapes to resemble those of biliary epithelial cells; moreover, newly opened chromatin regions were highly enriched for Sox binding motifs^27^. Given that Sox transcription factors are known to possess pioneer factor activity in cell culture^15, 28^, we hypothesized that one or more Sox factors facilitate biliary reprogramming by directly eliciting chromatin accessibility. To profile the expression of all *Sox* genes during biliary reprogramming, we first performed single cell RNA sequencing (scRNA-Seq) during DDC-induced hepatobiliary metaplasia, including pseudotime analysis^29^ based on known marker genes expressed in hepatocytes and reprogrammed cells at different stages (**Extended Data Fig. 1a-b**). This allowed us to identify Cd24 as a surface marker of cells at the early-to-intermediate stages of reprogramming and Epcam as a surface marker of cells at the intermediate-to-late stages of reprogramming (**Extended Data Fig. 1c-d, Supplementary Fig. 1)**. RNA-sequencing of cells isolated at various stages of reprogramming (**Extended Data Fig. 1e-g, Supplementary Fig. 2-3**) revealed *Sox4* and *Sox9* to be the only *Sox* factors to be expressed (**Extended Data Fig. 2a)**. *Sox9* was weakly expressed in normal hepatocytes, as previously reported in a subpopulation of periportal hepatocytes (**Extended Data Fig. 2b**)^30^, while *Sox4* expression was virtually undetectable in hepatocytes at baseline but rapidly induced during reprogramming (**Extended Data Fig. 2b**).

We then asked whether ectopic expression of *Sox4* and *Sox9* – delivered via an adeno-associated virus (AAV) gene transfer system – could initiate biliary reprogramming of hepatocytes under homeostatic (i.e. non-injury) conditions. To this end, we produced AAV8-TBG-*HA-Sox4-P2A-Cre* and AAV8-TBG-*HA-Sox9-P2A-Cre* and injected viral preps individually or concurrently to LSL-Rosa26-*Cas9-EGFP* mice (**Fig. 1a**). The system enables robust infection of >95% of hepatocytes and, significantly, allows infected cells at different stages of reprogramming to be recognized and isolated by virtue of an EGFP lineage tracer^31^. As a control, mice were injected with AAV8 virus carrying Cre but no additional payload (empty vector; EV).

**Fig. 1.**
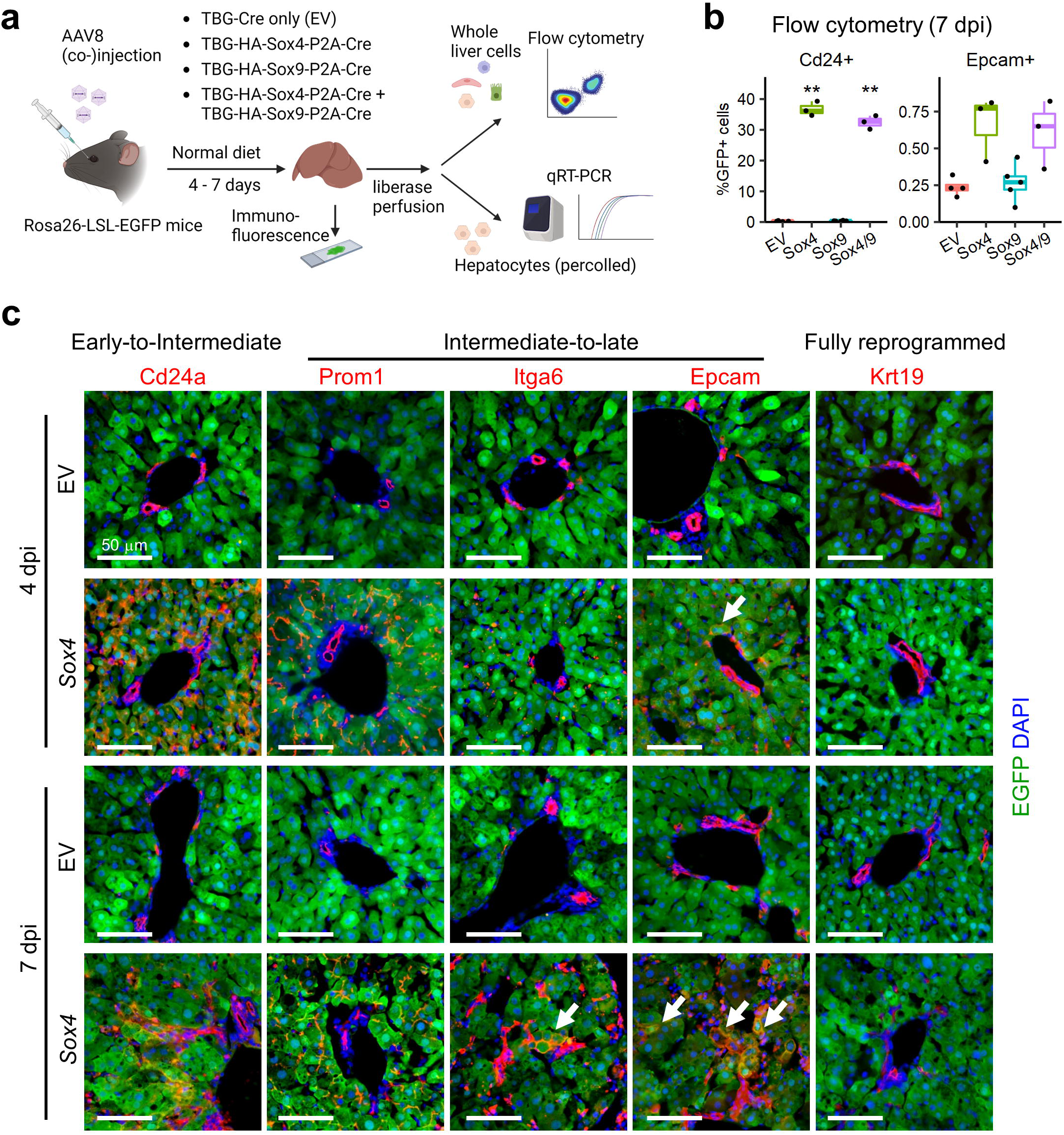
Exogenous expression of Sox4 is sufficient to induce biliary phenotypes in adult hepatocytes. (a) Experimental design. AAV8 packaged with *HA*-tagged *Sox4*, *Sox9*, both (*Sox4/9*), or neither (EV), under the control of the TBG promoter, were injected into Rosa-LSL-*Cas9-EGFP* mice. After being maintained on a normal diet for 4-7d, liver tissue and percoll-enriched hepatocytes were harvested and used for immunofluorescence, flow cytometry and qRT-PCR, respectively. (b) Reprogramming efficiency assessed by flow cytometry at 7 dpi using Cd24 (early-to-intermediate) and Epcam (intermediate-to-late) markers (n = 3-5 per group). Representative flow plots are shown in **Supplementary Fig. 4**. (c) Representative immunofluorescence of a panel of reprogramming markers in liver tissue harvested from mice injected with AAV-empty vector (EV) or AAV-TBG-*HA-Sox4* (Sox4) at either 4 dpi or 7 dpi timepoints. Arrows indicate EGFP+ hepatocyte-derived cells expressing the indicated reprogramming marker. Scale bar = 50 μm.

We harvested hepatocytes at 7 days post injection (dpi) and confirmed increased expression of both *Sox4* (∼1000-1500-fold) and *Sox9* (∼10-30-fold) by qRT-PCR (**Extended Data Fig. 2c**). Importantly, RNA-Seq demonstrated comparable increases in *Sox4* and *Sox9* during DDC-induced reprogramming (**Extended Data Fig. 2d**); therefore, we concluded that our expression system reasonably recapitulated the upregulation of *Sox4* and *Sox9* observed under physiological conditions. Strikingly, flow cytometry demonstrated that ectopic expression of *Sox4* or *Sox4* and *Sox9* together, but not *Sox9* alone, induced robust expression of Cd24 and modest expression of Epcam in EGFP+ hepatocyte-derived cells (**Fig. 1b, Supplementary Fig. 4**). A broader analysis of gene expression by qRT-PCR confirmed the induction of multiple biliary genes and the repression of multiple hepatocyte genes following ectopic expression of *Sox4* and *Sox4/9*, while *Sox9* alone had no effect (**Extended Data Fig. 2e**). Therefore, we focused our attention on *Sox4*.

*Sox4* mice lost weight following viral induction, reaching a nadir at 7-8 dpi before recovering most of the weight by 14 dpi (**Extended Data Fig. 3a**). During this period, the liver became smaller and paler than empty vector-injected controls (**Extended Data Fig. 3b-c**), with atypical ductal cells (**Fig. 3d**), a hallmark of chronic and fulminant liver diseases^32^. *Sox4* mRNA expression reached its maximum level from days 1-4 dpi, began to diminish slightly by 7 dpi, and diminished greatly by 10 dpi as confirmed at the protein level by immunofluorescence (**Extended Data Fig. 3e-f**).

Consistent with the Cd24 to Epcam reprogramming sequence identified at the mRNA level in DDC-treated mice (**Extended Data Fig. 1**), we found that Cd24 was robustly expressed at 4 dpi in *Sox4* hepatocytes at the protein level, whereas Epcam became strongly expressed only at 7 dpi (**Fig. 1c**). Other intermediate-to-late biliary reprogramming markers, such as Prom1 and Itga6, became detectable between 4 dpi and 7 dpi (**Fig. 1c**). We never observed EGFP+ cells that co-expressed Krt19, a marker of fully reprogrammed biliary cells (**Extended Data Fig. 1a-c**), suggesting that Sox4 is unable to induce the final stages of biliary reprogramming (**Fig. 1c**). Consistent with these data, flow cytometry demonstrated that Cd24 expression peaked at 4 dpi, becoming detectable in over 60% of EGFP+ hepatocytes before dropping back to baseline, while Epcam expression in EGFP+ cells continued to increase after 4 dpi (**Extended Data Fig. 3g**).

PCA mapping of RNA-Seq data demonstrated that *Sox4*-expressing hepatocytes exhibited a similar degree of reprogramming compared to DDC-induced early reprogrammed cells (**Extended Data Fig. 3g**). Using gene sets built as differentially expressed genes (fold changes > 2 or < 0.5, p.adj < 0.05, DESeq2^33^) between hepatocytes and Rep_early cells (2,355 upregulated and 1,189 downregulated genes in Rep_early vs. hepatocytes) (**Table S1**), GSEA confirmed that *Sox4*-expressing hepatocytes were significantly enriched for a reprogrammed cell signature and under-represented for the hepatocyte signature (**Extended Data Fig. 3i**). Taken together, the data show that Sox4 is sufficient to induce the initial stages of biliary reprogramming, including the repression of hepatocyte gene expression.

### Sox4 remodels chromatin landscapes

Since Sox4 dominantly initiated a biliary fate conversion *in vivo*, we hypothesized that Sox4 mediates reprogramming by changing chromatin conformation. We performed Assay for Transposase-Accessible Chromatin using sequencing (ATAC-Seq)^34^ with hepatocytes isolated at 4 dpi from AAV-EV or AAV-*HA-Sox4*-injected mice. Differential peak analysis comparing *Sox4*-expressing hepatocytes to control hepatocytes identified 20,329 regions with increased accessibility (i.e. newly opened), 14,564 regions with decreased accessibility (i.e. newly closed), and 92,894 regions exhibiting no change (**Fig. 2a**). Analysis of transposase cleavage patterns in deeply sequenced samples (we had >100 million uniquely mapped tags) enables visualization of transcription factor footprints within ATAC-Seq tags^35^. As predicted, motif analysis of the merged replicates indicated that Sox binding is enriched in newly opened regions, while no difference was observed in the unchanged regions (**Fig. 2b**). Unexpectedly, however, in the control EV hepatocytes, where Sox4 is not expressed, we observed a Sox4 footprint in regions that become closed at 4 dpi (**Extended Data Fig. 4a**). Our analysis of published ChIP-Seq data^36, 37^ indicated that these unexpected footprints colocalized with binding of the hepatocyte transcription factors Hnf4a^38, 39^ and Rxra^40^ (**Extended Data Fig. 4b**). When compared with previously published ATAC-Seq studies^27, 41^, *Sox4*-expressing hepatocytes exhibited open chromatin profiles resembling early reprogrammed cells (**Fig. 2c**), consistent with our findings with RNA-Seq (**Extended Data Fig. 3h**). Thus, ectopic expression of *Sox4* results in a chromatin landscape mirroring that which is present under physiological conditions in the early stages of biliary reprogramming.

**Fig. 2.**
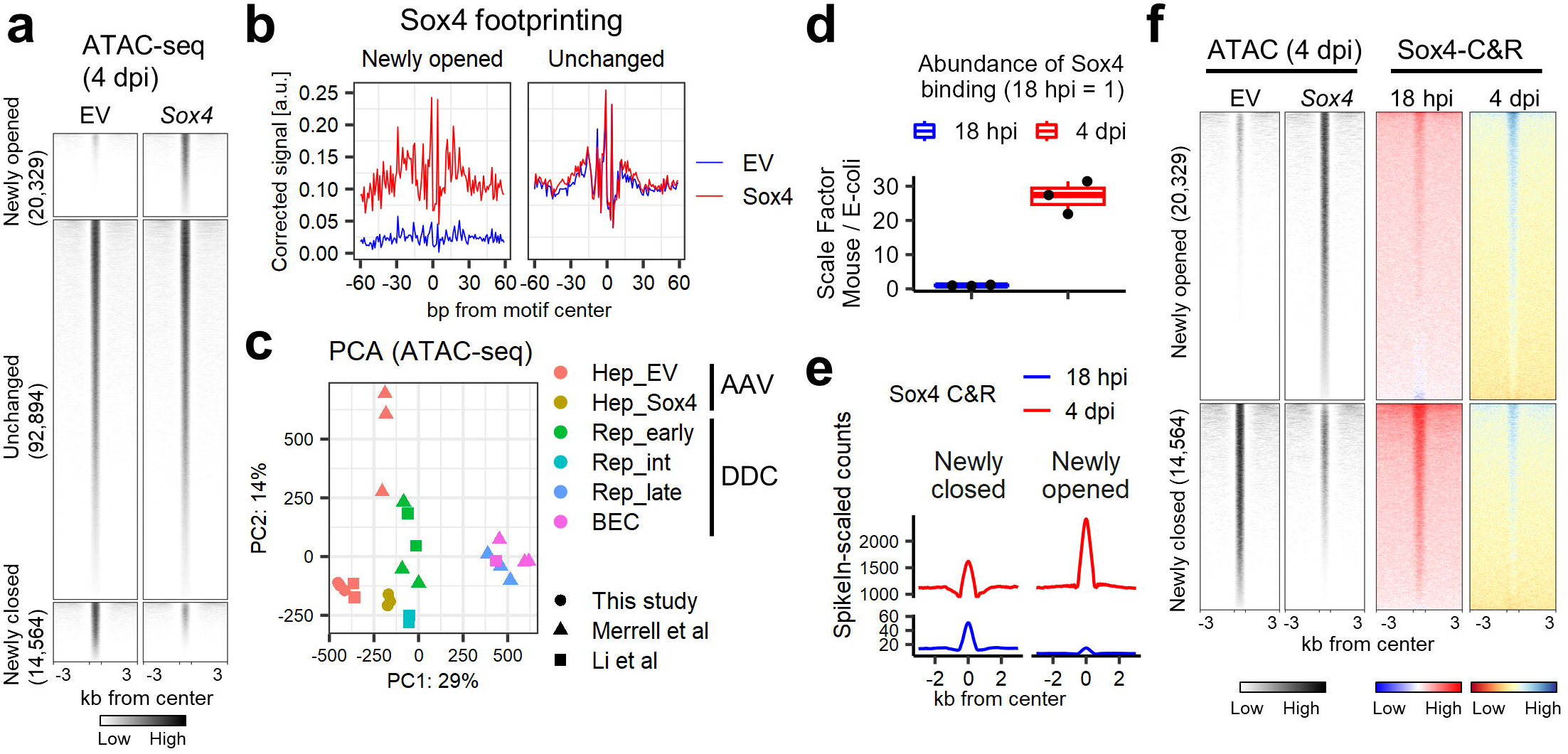
The chromatin landscape following ectopic expression of *Sox4* recapitulates that of partially reprogrammed cells. (a) Differential peak analysis identified 20,329 opened regions and 14,564 closed regions resulting from *Sox4* expression in hepatocytes compared with empty vector (EV). DiffBind and DESeq2 packages were used with an FDR cutoff set to 0.01. (b) Results of DNA footprint analysis for the Sox4 binding motif comparing empty vector and *Sox4* hepatocytes at newly opened and unchanged regions as defined in **(a)**. TOBIAS software was used with the default setting, and motifs assigned “bound” either for empty vector or *Sox4* hepatocytes were used for visualization of the averaged signals. (c) PCA mapping for ATAC-Seq of empty vector (EV) and *Sox4* hepatocytes (n = 3 per group). ATAC-Seq was performed using percoll-enriched hepatocytes at 4 dpi of AAV-empty vector and AAV-*HA-Sox4* injection. The data are shown in comparison with previously published data^27, 41^. The data obtained in this study were downsampled by approximately 1/100-fold to adjust read depth to be comparable to the previous data sets. Note that “Rep_int (intermed)” in this panel does not indicate Cd24+ cells but rather indicates Sox9+ cells sorted from Sox9-RFP reporter mice treated with DDC^41^. (d) Quantification of Sox4 total binding was estimated using E-coli spike-in genomic DNA. Scale factors were calculated as (Total read counts aligned to mouse genome/Total read counts aligned to E-coli genome). Data are shown as fold-change compared to the mean value at 18 hpi data (n = 3). (e) Averaged aggregate plots for spike-in-scaled Sox4 CUT&RUN (C&R)-Seq data at newly closed and newly opened regions at the designated time points. (f) Sox4 CUT&RUN (C&R) signals (right two columns) are visualized as heatmaps for the newly opened and closed regions defined by ATAC-Seq (left two columns). The corresponding averaged aggregate plots for Sox4 CUT&RUN are shown in **Extended Data Fig. 4f**. Note that Sox4 data are normalized genome-wide in each sample (18 hpi and 4 dpi) and do not support quantitative comparison between the two timepoints. The numbers in brackets indicate number of regions.

### Sox4 binding precedes major changes in gene expression during reprogramming

To explore the association between Sox4 binding and chromatin opening, we performed a genome-wide examination of Sox4 binding patterns at the early stage of reprogramming *in vivo*. To this end, we performed Cleavage Under Targets & Release Using Nuclease sequencing (CUT&RUN-Seq)^42, 43^ using hepatocytes isolated at 18 hours post injection (18 hpi) and 4 dpi. Hepatocytes at 18 hpi showed weak HA-Sox4 expression at the protein level (**Extended Data Fig. 4c**) and minimal changes in gene expression (**Extended Data Fig. 4d**), thereby providing a profile of Sox4 binding during the early stages of *Sox4*-induced reprogramming. We obtained 9,463 and 19,362 Sox4 CUT&RUN peaks at 18 hpi and 4 dpi samples respectively. As predicted, the peaks were highly enriched for Sox binding motifs (**Extended Data Fig. 4e**).

Using an E-coli DNA spike-in approach, we quantitated Sox4 binding at each timepoint. As expected, Sox4 binding increased from 18 hpi to 4 dpi in both newly closed and newly opened regions (**Fig. 2d**) in proportions that were comparable to the relative amounts of newly closed and opened peaks, as seen by ATAC-Seq. The increase in binding was greatest in the newly opened regions, with the averaged peak summit (4 dpi/18 hpi) increasing by 160-fold in the newly opened regions and by 32-fold in the newly closed regions (**Fig. 2e**), a trend that was also seen when the data were normalized across the genome (**Fig. 2f, Extended Data Fig. 4f**). Our data indicate that extensive Sox4 binding (18 hpi) precedes major changes in gene expression. Consequently, we set out to understand modulators and consequences of Sox4 binding.

### Sox4 targets genes for silencing or activation during reprogramming

To assess the association of the ATAC peaks with gene expression, we annotated each peak with its nearest gene, associating 19,985 genes in total to all ATAC peaks (n =127,787) identified at any of the newly opened, unchanged, or newly closed regions. The annotated genes were then ranked based on the fold-change between empty vector and *Sox4*-expressing hepatocytes, and the ranked gene list was used as input for GSEA^44^. Consistent with our RNA-Seq results, regions exhibiting decreased chromatin accessibility upon *Sox4* expression were found in proximity to hepatocyte genes (**Fig. 3a**, upper), while regions exhibiting increased chromatin accessibility upon *Sox4* expression were found in proximity to reprogramming-associated genes (**Fig. 3a**, lower). Approximately half of the genes associated with newly closed regions (2,685/5,272; 50.9%) were downregulated during DDC-induced reprogramming, including many known hepatocyte genes (**Fig. 3b, Table S2**). By contrast, more than half of the genes associated with newly opened regions (3,663/5,823; 62.9%) were upregulated during DDC-induced reprogramming, including many known biliary genes (**Fig. 3c, Table S2**). Gene ontology analysis of ATAC-Seq (**Table S3**) and RNA-Seq (**Table S3**) data revealed that terms enriched for newly closed region-associated genes were shared with the terms under-represented in DDC-induced reprogrammed cells, while terms enriched for newly opened region-associated genes were shared with the terms over-represented in DDC-induced reprogrammed cells (**Extended Data Fig. 5a-d**). When we annotated Sox4 peaks at the 18 hpi and 4 dpi timepoints with the nearest gene and performed GSEA, there was enrichment of a hepatocyte signature for genes associated with Sox4 peaks at 18 hpi (**Fig. 3d**, upper) and enrichment of a reprogramming signature for genes associated with Sox4 peaks at 4 dpi (**Fig. 3d**, lower). We conclude that Sox4-induced changes in chromatin accessibility are associated with corresponding decreased expression of hepatocyte genes and increased expression of biliary genes, as occurs early in DDC-induced reprogramming.

**Fig. 3.**
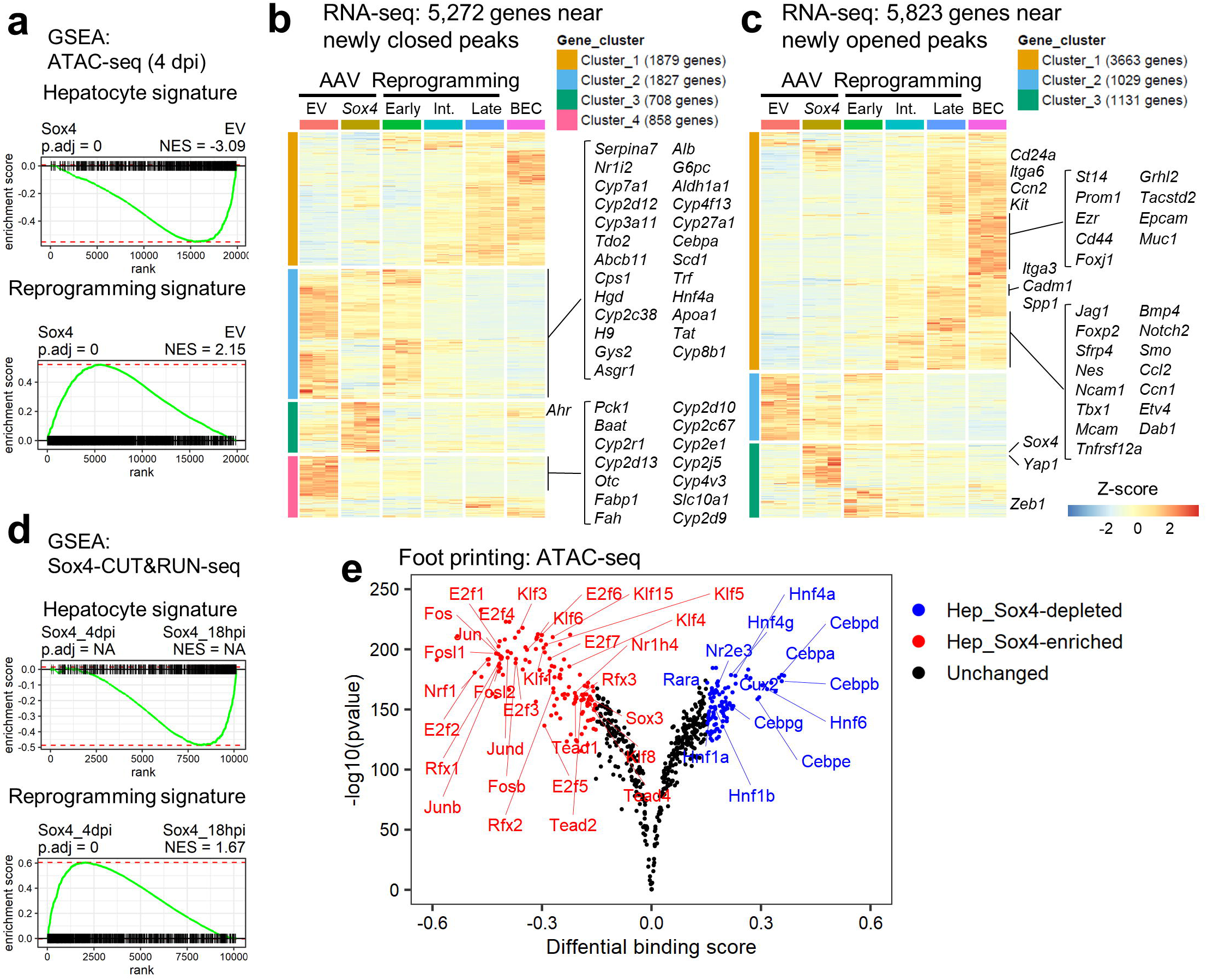
Sox4 binding silences hepatocyte genes while priming biliary genes by altering binding patterns. (a) GSEA was performed with genes annotated to 4dpi ATAC-Seq peaks to compare empty vector and *Sox4* hepatocytes using the hepatocyte-enriched and Rep_early-enriched signatures as described earlier (**Table S1, Extended Data Fig. 3i, Methods**). (b) Heatmap visualization of the 5,272 genes near newly closed peaks using the RNA-Seq data of the DDC-induced reprogramming experiment (**Extended Data Fig. 3h**). Genes were categorized to four clusters by *k*-means clustering. Representative hepatocyte genes (39/50 from a manually curated gene list; **Table S2**) are shown on the right. (c) Heatmap visualization of the 5,823 genes near newly opened peaks using the RNA-Seq data. The genes were categorized to three clusters by *k*-means clustering. Representative biliary/reprogramming genes (33/49 from a manually curated gene **list; Table S2**) are shown on the right. (d) GSEA was performed with genes annotated to the Sox4 CUT&RUN peaks identified either at 18 hpi and 4 dpi using the hepatocyte-enriched and Rep_early-enriched signatures as described earlier (**Table S1, Extended Data Fig. 3i, Methods**). “NA” was assigned when p.adj < 1e-50. (e) Global footprint analysis depicted as a volcano plot. The analysis used all motifs assigned as “bound” by TOBIAS in either empty vector or *Sox4* hepatocytes. Differential footprints are defined as those with Log2(fold-change of footprint score) > 0.15 and log10(pvalue) < -100.

We then performed DNA footprint analysis to identify transcription factor binding sites that were enriched or depleted following *Sox4* expression. This confirmed that most of the transcription factor binding sites previously identified as enriched in newly opened regions during DDC-induced reprogramming (e.g. AP1, E2F, and Tead)^27^ were also enriched in the *Sox4* newly-opened regions (**Fig. 3e**). Interestingly, we also found that transcription factors associated with hepatocyte identity and function (e.g. Hnf4a/g, Rara, and Cebpa/b/d/e/g) lost their footprints following *Sox4* expression (**Fig. 3e**). Hnf4a motifs were also evident within many Sox4-CUT&RUN peaks (**Extended Data Fig. 4e**).

**Fig. 4.**
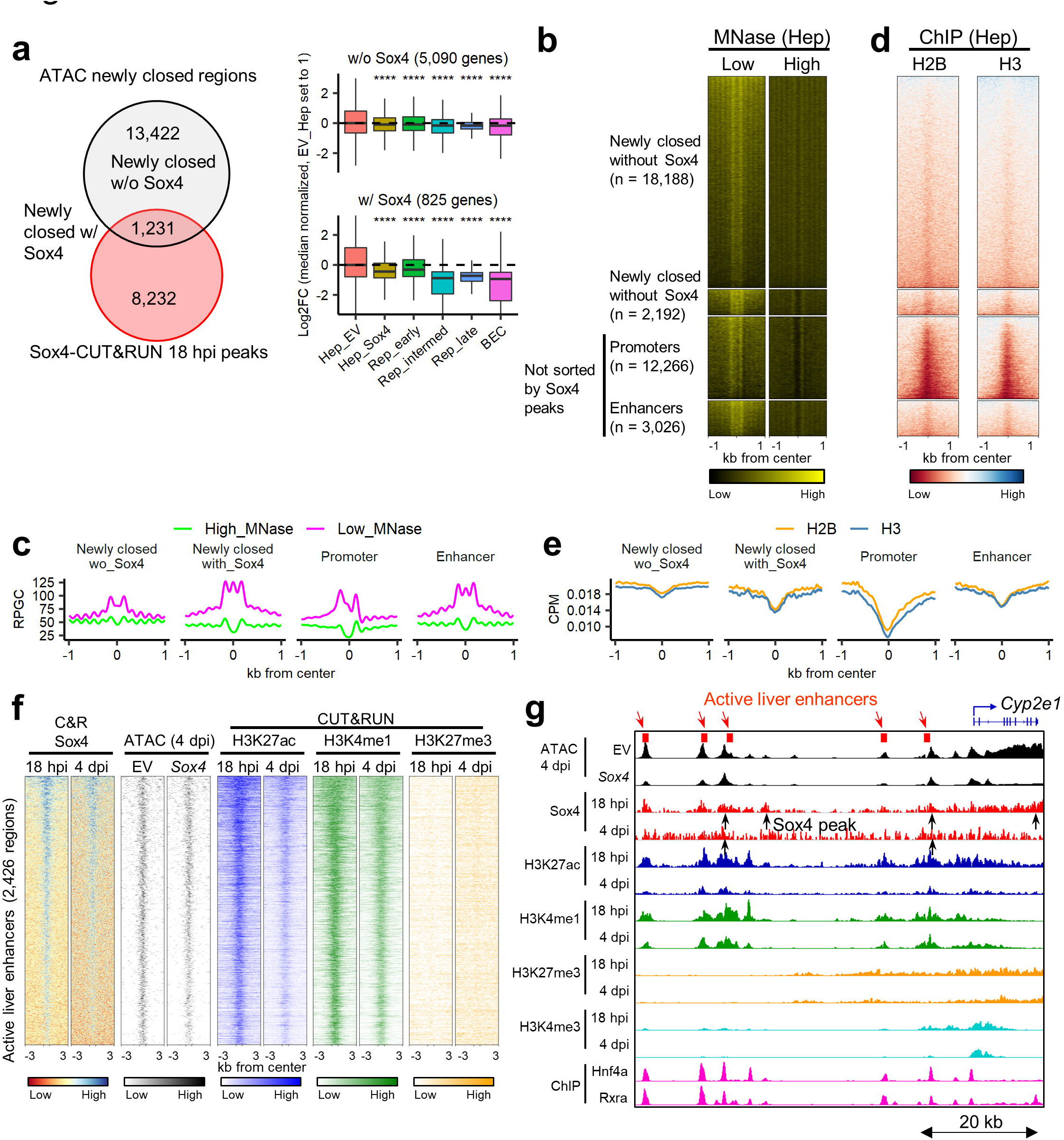
Sox4 suppresses hepatocyte identity by inactivating hepatocyte enhancers. (a) Newly closed regions with or without overlaps with 18hpi-Sox4 peaks were annotated with their nearest genes, and expression of each gene in empty vector or *Sox4*-expressing hepatocytes was compared with the DDC-induced reprogrammed cells. ****p < 0.0001. p-value was calculated by Wilcoxon rank sum test. (b) Heatmap representation of MNase-seq at low and high levels for the newly closed regions with or without overlaps with Sox4 peaks at 18 hpi along with active liver promoters and active liver enhancers. The corresponding averaged aggregate plots are shown in **(c)**. (c) Averaged aggregate plots of hepatocyte MNase-Seq in newly closed regions with or without overlaps with Sox4 peaks (18 hpi) along with active liver promoters and active liver enhancers. (d) Heatmap representation of ChIP-seq for core histone H2B and H3 for the same regions as in **(b)**. The corresponding averaged aggregate plots are shown in **(e)**. (e) Averaged aggregate plots of H2B and H3 ChIP-Seq for the same regions as in **(b)**. (f) ATAC-Seq and CUT&RUN-Seq of Sox4, H3K27ac, H3K4me1 and H3K27me3 signals visualized as heatmaps at active liver enhancer regions. The corresponding averaged aggregate plots are shown in **Extended Data Fig. 6d,e**. (g) Genome browser view of the *Cyp2e1* gene is shown as an example of Sox4-induced closing of a hepatocyte gene-associated region. ChIP-Seq data were obtained from GEO datasets (GSE137066 for Hnf4a; GSE53736 for Rxra) reflecting normal liver cells.

Collectively, these results indicate that changes in chromatin and transcription factor binding landscapes following *Sox4* ectopic expression resemble those of cells undergoing physiological biliary reprogramming.

### Sox4 initially closes active hepatocyte enhancers and evicts Hnf4a

We next assessed whether early Sox4 binding (18 hpi) in regions destined to undergo chromatin closing (**Fig. 2f**, lower) was correlated with a loss of gene expression at 4 dpi (**Fig. 4a**, left). Strikingly, genes associated with newly closed regions that were bound by Sox4 (n=825 genes) exhibited a marked reduction in gene expression during DDC-induced reprogramming (**Fig. 4a**, right bottom). By contrast, genes associated with newly closed regions that were not bound by Sox4 (n=5,090 genes) exhibited much smaller reductions in gene expression (**Fig. 4a**, right top). We observed that newly closed regions were highly enriched in areas distal to the TSS (**Extended Data Fig. 6a-b**), raising the possibility that Sox4 binding modulates the activity of hepatocyte enhancers.

To assess the initial chromatin targeting by Sox4 more deeply, we compared sites of Sox4 binding with regions previously characterized based on their sensitivity to MNase and histone modifications. Upon high-level MNase digestion, labile nucleosomes and free DNA are destroyed and stable nucleosomes are resistant, whereas low-level MNase digestion preserves labile nucleosomes at enhancers^45^ (**Extended Data Fig. 6c**). Open regions initially targeted by Sox4 that later became closed exhibited MNase profiles resembling those of active liver enhancers (**Fig. 4b-c**) and had patterns of H2B and H3 binding (ChIP-Seq) similar to those associated with active liver enhancers (**Fig. 4d-e**). Moreover, *Sox4* expression was associated with reduced accessibility of these enhancers at 4 dpi compared to empty vector (**Fig. 4f-g, Extended Data Fig. 6d**) and resulted in a decrease of the active enhancer marks H3K27ac and H3K4me1 at 4 dpi compared to 18 hpi (**Fig. 4f-g, Extended Data Fig. 6e**). Collectively, these results indicate that shortly after its induction, Sox4 binds to and inactivates active liver enhancers.

We next sought to understand how Sox4 targets active liver enhancers. Given the unexpected finding that the Hnf4a binding motif was enriched in both 18 hpi and 4 dpi Sox4-CUT&RUN peaks (**Extended Data Fig. 4e**), and more so at 18 hpi than 4 dpi (**Fig. 5a**), we considered the possibility that targeted rather than promiscuous binding was responsible for Sox4’s localization to Hnf4a sites. Remarkably, we found that the Sox binding motif (CTTTGT/ACAAAG) overlaps the binding motifs for Hnf4a (CAAAG/CTTTG) and other hepatocyte-enriched transcription factors (**Fig. 5b**). Considering that pioneer factors can bind partial motifs^16^, Sox4 could recognize such partial motifs for Hnf4a. Indeed, using published ChIP-Seq data^37^, we found substantial overlap of Sox4 binding sites with Hnf4a binding sites in adult hepatocytes (**Fig. 4g** bottom, **Extended Data Fig. 6f** left, **g** left), but not in colon epithelial cells (**Extended Data Fig. 6f** right, **g** right)^46^. These results indicate that Sox4 initially binds to regions that are open and occupied by Hnf4a in hepatocytes.

**Fig. 5.**
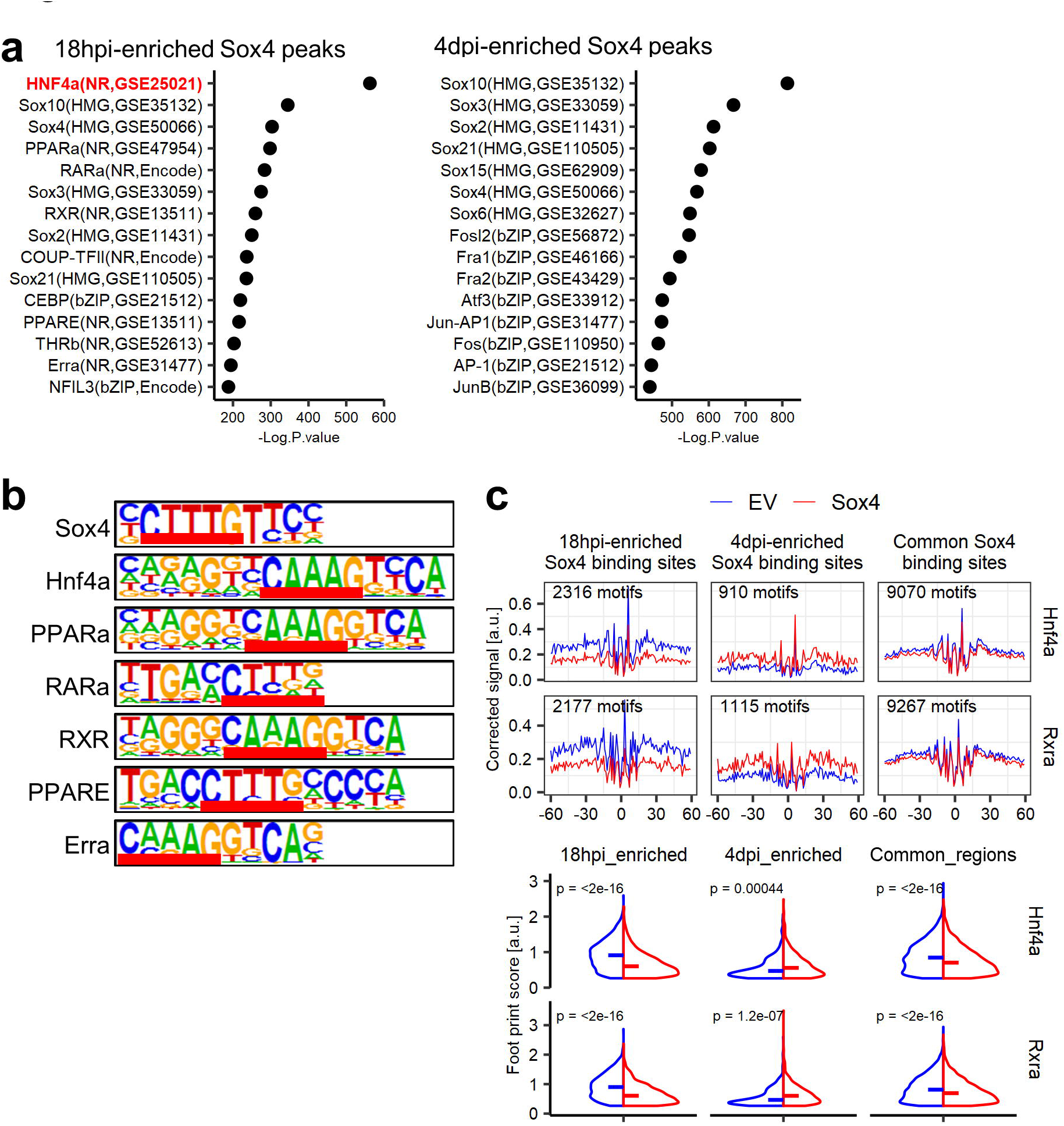
Sox4 evicts hepatocyte transcription factors by hijacking their binding motifs. (a) HOMER motif analysis with known motifs for 18hpi- and 4dpi-enriched Sox4 CUT&RUN peaks. 18hpi-enriched Sox4 peaks (n = 2,327) and 4dpi-enriched Sox4 peaks (n = 3,136) were identified using DiffBind and DESeq2 packages. The Hnf4a motif, highlighted in red, ranked first among 18 hpi-enriched Sox4 binding sites. (b) Consensus binding sequences of several hepatocyte transcription factors exhibiting partial overlap with that of Sox4 (overlapped sequences are underlined). (c) Averaged aggregate plots of DNA footprinting of Hnf4a and Rxra at the 18hpi-enriched Sox4 binding sites, 4dpi-enriched Sox4 binding sites and common Sox4 binding sites (upper panel). The data are shown as the comparison between empty vector and *Sox4* hepatocytes. Motifs assigned “bound” by TOBIAS for either the empty vector or *Sox4* hepatocytes were used for the analysis. The lower violin plots show the comparison of the footprint scores, indicative of the binding probability of the transcription factors, at each region. Bars indicate median values. p-values were calculated by Wilcoxon rank sum test.

To explore the consequence of Sox4 binding, we used ATAC-Seq data to compare the footprints of Hnf4a and another hepatocyte transcription factor, Rxra, with those that were bound by Sox4 more efficiently at 18 hpi than at 4 dpi. Strikingly, *Sox4-*expressing hepatocytes exhibited decreased Hnf4a and Rxra footprints at 4 dpi compared to hepatocytes that received empty vector (**Fig. 5c**). We also confirmed the reduction of footprints of the factors in active liver enhancers (**Extended Data Fig. 6h**). Collectively, the data indicate that Sox4 suppresses hepatocyte identity in part by evicting resident hepatocyte transcription factors and reducing the accessibility and activity of hepatocyte enhancers.

### Sox4 opens chromatin in biliary regions

Next, we returned to our earlier hypothesis that Sox4 primes the biliary phenotype in hepatocytes by acting as a pioneer factor. Based on the strong correlation between Sox4 binding and changes in the chromatin landscape (**Fig. 2f**), we first assessed the ability of Sox4 to bind nucleosomal DNA, as this is a defining feature of pioneer factor activity. To this end, we incubated recombinant Sox4 protein with nucleosome particles assembled on a human *LIN28B* DNA fragment, which was previously demonstrated to bind Sox2^16^. Electrophoretic mobility shift assay (EMSA) confirmed that recombinant Sox4 binds specifically to its target site (**Extended Data Fig. 7a)** and to assembled *LIN28B* nucleosomes (**Fig. 6a**). Given that pioneer factor activity is associated with opening of previously closed chromatin, we divided the regions of newly opened chromatin (based on ATAC-Seq peaks) into two sub-regions: those whose accessibility increased over baseline upon *Sox4* expression (“more opened regions,” or MORs) (**Fig. 6b**, upper left) and those regions that only became accessible when *Sox4* was expressed (“*de novo* opened regions,” or DORs) (**Fig. 6b**, lower left). We visualized Sox4-CUT&RUN signals at these regions and confirmed that both MORs and DORs were bound by Sox4 at 4 dpi (**Fig. 6b**, right heatmaps). DORs, and to a lesser extent MORs, fell into broad domains of chromatin enriched for high-MNase signals, compared to active promoters and enhancers **(Figs. 6c-d)**. Sox4-targeted DORs and MORs were also found in broad domains enriched for core histones (**Figs. 6e-f**). We conclude that Sox4 acts as a pioneer factor by targeting nucleosomal DORs, and to a lesser extent MORs, leading to increased chromatin accessibility.

**Fig. 6.**
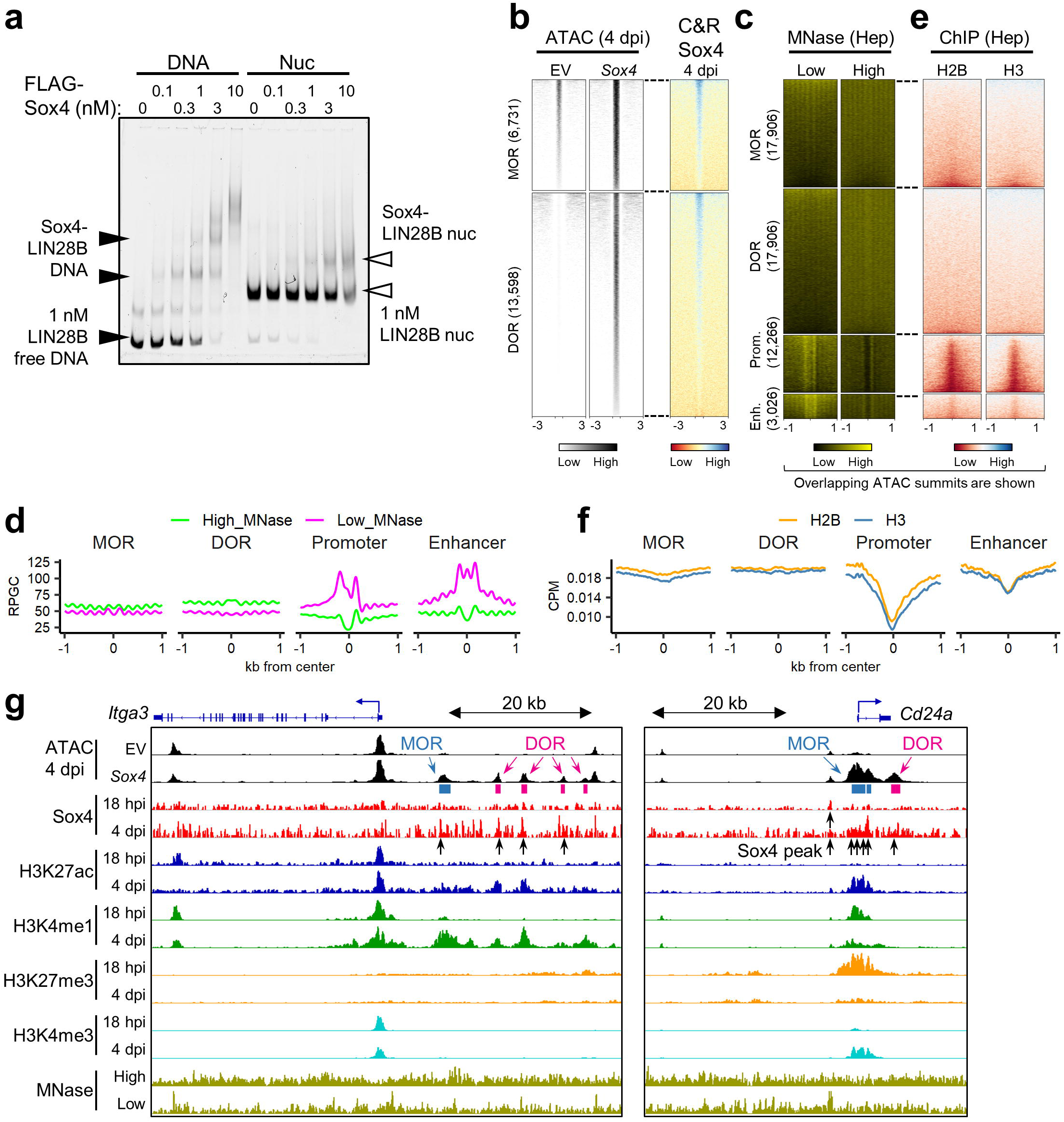
Sox4 opens putative biliary cell enhancers in hepatocytes. (a) EMSA assay to evaluate the binding ability of purified recombinant FLAG-Sox4 to naked *LIN28B* DNA and *in vitro* assembled nucleosomal *LIN28B* DNA. (b) Sox4 CUT&RUN-Seq data (right column) are shown for the more opened regions (MOR) and *de novo* opened regions (DOR), which correspond to newly opened regions with weak ATAC peaks in empty vector and those without ATAC peaks in empty vector (left two columns). Numbers in brackets indicate number of regions. (c) Heatmaps of previously published MNase-Seq data obtained for adult hepatocytes at low and high levels of MNase ^45^. The regions are centered with all the intersectable ATAC peaks of either empty vector or *Sox4*-expressing hepatocytes. Numbers in brackets indicate number of regions. (d) Averaged aggregate plots corresponding to the heatmaps shown in **(c)**. (e) Heatmaps of previously published H2B and H3 ChIP-Seq obtained for adult hepatocytes ^45^. Regions are centered with all intersectable ATAC peaks of either empty vector or *Sox4* hepatocytes. (f) Averaged aggregate plots corresponding to the heatmaps shown in **(d)**. (g) Genome browser views of two examples for Sox4-induced opening at biliary epithelial cell genes: *Itga3* (intermediate-to-late reprogramming marker) and *Cd24a* (early-to-intermediate reprogramming marker). Negative values for Sox4 tracks are not shown. (h)

At DORs, active enhancer and promoter marks were increased from faint or residual backgrounds, and at MORs the active marks were initially more elevated and increased, by 4 dpi, while the H3K27me3 repressive mark was marginal to begin with and reduced at 4 dpi (**Extended Data Fig. 7b-c**). In accordance with our earlier finding that newly opened regions were enriched in proximity to reprogramming-related genes (**Fig. 3a,c**), comparable chromatin/histone mark changes were observed in the regions near biliary genes; concordantly, Sox4 binding was observed either at putative TSS-distal enhancers (**Fig. 6g**, left) or at promoters/gene bodies (**Fig. 6g**, right). Moreover, DOR peaks were relatively enriched in TSS-distal regions compared to MOR peaks and unchanged regions (**Extended Data Fig. 6a-b**), indicating that DORs contain *de novo* primed enhancers. Collectively, these results indicate that Sox4 binding by 4 dpi leads to chromatin opening and the acquisition of active or primed characteristics of cis-regulatory elements associated with biliary phenotypes.

### Sox4 and Sox9 are YAP targets

Finally, we sought to identify a direct link between the signaling pathways known to regulate biliary reprogramming and the expression of Sox4. The transcriptional co-activator Yap, which mediates output from the Hippo signaling pathway, is among the most well-characterized factors driving biliary reprogramming^26, 41^. Thus, we hypothesized that *Sox4* is a direct target of Yap. Using previously published ChIP-Seq data from hepatocytes isolated from control or DDC-treated livers^41, 47^, we observed that Yap binds to the *Sox4* gene in mice fed DDC (**Extended Data Fig. 7e**). Similar observations were made for *Sox9* (**Extended Data Fig. 7e,** right). Concordantly, analysis of a previously published microarray data^26^ revealed that expression of YAPS127A induced expression of *Sox4* and *Sox9* in hepatocytes (**Extended Data Fig. 7f**). Taken together, the findings indicate that liver injury induces Yap-mediated expression of *Sox4*, resulting in subsequent changes in chromatin configuration to facilitate biliary reprogramming.

## Discussion

The epigenetic mechanisms underlying cell fate switching, or reprogramming, have been studied in detail in the context of induced pluripotency, where pioneer factors enable a change in cell identify by reconfiguring chromatin. Our study provides evidence that Sox4 acts as a pioneer factor *in vivo* in a well-defined system of physiological reprogramming: hepatobiliary metaplasia, a hallmark of chronic and fulminant liver disease^32^. Moreover, we found that Sox4 exerts this activity through a sequential process in which enhancers associated with the starting (hepatocyte) cell type are decommissioned prior to the activation of enhancers associated with the acquired (biliary) cell type. Such enhancer reorganization by pioneer factors, including silencing of the starting cell’s enhancers, has been proposed for iPSCs, where early and preferential binding to active somatic enhancers causes the redistribution of associated somatic transcription factors like P300 and the recruitment of enhancer silencing factors like Hdac1^13^. In addition, as we observed for Hnf4a and Rxra, pioneer factors can compete for binding sites in enhancers to influence lineage trajectories^48, 49^.

Our data support and reconcile these models in an *in vivo* setting and offer a possible molecular mechanism to account for the specificity of binding in a physiologically relevant context. Specifically, we observed that key hepatocyte transcription factors – including Hnf4a, a master regulator of hepatocyte identity^38, 39^ – share binding motifs with Sox4, suggesting that Sox4 may hijack these sites when it is expressed. Consistent with this idea, Sox4 preferentially binds to open chromatin regions occupied by Hnf4a and Rxra early in biliary reprogramming; subsequently, these factors are evicted from their native binding sites. While our results do not prove that Sox4 is directly responsible for this eviction, the data suggest that Sox4 competes with hepatocyte transcription factors like Hnf4a and Rxra for binding to hepatocyte enhancers early in the reprogramming process. Thus, our study provides evidence that pioneer factors can coordinately disrupt a starting cell fate while engaging closed chromatin to initiate a new cell fate.

Our findings also provide a molecular link between the signaling pathways that have been identified as mediators of biliary reprogramming and the subsequent epigenetic and transcriptional changes that stabilize the reprogrammed state. Cell-cell signaling following liver injury results in activation of the Notch, Yap, and TGFβ signaling pathways, all of which are required for biliary reprogramming^23, 25, 26^. The pioneer factor Sox4, whose expression is induced directly by Yap, explains at least in part how such exogenous upstream signals converge on the epigenome to alter cell state. Other epigenetic and transcriptional regulatory factors are also likely to participate in the conversion of hepatocytes to a fully reprogrammed state^41^.

The results presented here are thus consistent with a multi-step model in which pioneer factor activity, through both positive and negative effects on chromatin accessibility, is essential for the cell fate changes accompanying tissue metaplasia *in vivo* (summarized in **Extended Data Fig. 8)**. First, signals from the injured liver microenvironment result in the induction of Sox4 through the Hippo/YAP pathway. Sox4 then binds to regions of open chromatin occupied by hepatocyte-specific transcription factors (e.g. Hnf4a and Rxra), causing their displacement. Finally, Sox4 acts as a traditional pioneer factor by binding to canonical Sox binding sites in biliary enhancers in closed chromatin, causing the regions to become open and accessible to other biliary transcription factors. We suspect that this cascade may be employed in other cases of metaplasia, including those leading to human cancers^4^.

## Acknowledgements

This work was supported by NIH grants R01DK083355 and R01GM36477, the Fred and Suzanne Biesecker Pediatric Liver Center, the Abramson Family Cancer Research Institute, The International Medical Research Foundation, The Daiichi Sankyo Foundation of Life Science, The Mochida Memorial Foundation for Medical and Pharmaceutical Research, The Mitsukoshi Health and Welfare Foundation, The Uehara Memorial Foundation, The Kanae Foundation, The Japanese Biochemical Society, and the Osamu Hayaishi Memorial Scholarship for study abroad. We thank the Penn Center for Molecular Studies in Digestive and Liver Disease (P30DK050306) for assistance with tissue processing, Véronique Lefebrvre and Rajan Jain for helpful discussions and comments on the manuscript, and members of the Stanger and Zaret labs for useful suggestions.

## Author Contributions

Conceptualization, T.K. and B.Z.S; methodology, T.K., J.S., K.I., A.K., S.Y., J.L., A.J.M., N.T., Q.L., I.A., K.Z., B.Z.S.; experimental design, T.K., J.S., K.Z., B.Z.S.; conduction of experiments, T.K ., S.Y., H.C., N.T.; resources, R.U.R, I.A.; bioinformatics analysis, T.K., J.S., K.I., A.K.; writing, T.K., J.S., K.Z., B.Z.S.; funding acquisition, K.Z., B.Z.S.; supervision, K.Z., B.Z.S.

## Methods

### Lead Contact

Further information and requests for resources and reagents should be directed to and will be fulfilled by the lead contact, Ben Z. Stanger

### Materials availability

All unique and stable reagents generated in this study of which sufficient quantities exists are available from the Lead Contact with a completed Materials Transfer Agreement.

### Data and code availability

Data that generated during the study have been deposited in Gene Expression Ominubus (GEO) with the accession number GSE221225 (SuperSeries) with SubSeries accession numbers GSE218945 and GSE218947 (RNA-Seq); GSE219052 (ATAC-Seq); GSE221223 and GSE221224 (CUT&RUN-Seq). Detailed scripts and parameters used for each step of the analysis provided by reasonable request to the authors.

### Mice

Rosa-LSL-Cas9-EGFP mice^50^ on a C57BL/6J background were purchased from the Jackson Laboratory (strain #026175) and maintained as homozygotes. All mouse experiment procedures used in this study were performed following the NIH guidelines. All mouse procedure protocols used in this study were in accordance with, and with the approval of, the Institutional Animal Care and Use Committee of the University of Pennsylvania.

For characterization of the reprogramming stage, 4-5 week-old mice were retro-orbitally injected with AAV:ITR-U6-sgRNA(backbone)-TBG-Cre-WPRE-hGHpA-ITR ^31^ (empty vector: EV) with 5 × 10^11^ genome copies/mouse/gene. One week later, induction of biliary reprogramming was started by initiating a 0.1% 3,5-diethoxycarbonyl-1,4-dihydrocollidine (DDC) diet (Envigo). Eight weeks after the DDC challenge, reprogrammed cells and biliary cells were harvested as described below.

For the exogenous expression of *Sox4*, *Sox9* and *Sox4*/*Sox9*, 4- to 8-week-old mice were retro-orbitally injected with 5 × 10^11^ genome copies/mouse/gene. For the Sox4 expression experiments for RNA-Seq, ATAC-Seq, and CUT&RUN-Seq experiments, 1 × 10^12^ genome copies/mouse of AAV8-TBG-*HA-Sox4-P2A-Cre* were retro-orbitally injected.

### Plasmid cloning

All PCR reactions were performed using the Phusion Flash High-Fidelity PCR Master Mix (Thermo) following the manufacturer’s instructions. For all AAV plasmids, endotoxin was eliminated by treating the plasmids with Endozero columns (Zymo Research) before proceeding to AAV production. Transformation was performed using Stbl3 bacteria (Thermo) following the manufacturer’s instruction.

### AAV-HA-Sox4-P2A-Cre plasmid

Mouse *HA-Sox4-P2A* and *P2A-Cre* blocks were PCR-amplified from pLVXT-Sox4 (Addgene, #101121) and AAV:ITR-U6-sgRNA(backbone)-TBG-Cre-WPRE-hGHpA-ITR^31^ respectively, using the primers listed i**n Table S4**. The AAV-TBG backbone was prepared by removing EGFP from pAAV.TBG.PI.eGFP.WPRE.bGH (Addgene, #105535) using NotI-HF (NEB) and BamHI-HF (NEB). The two PCR-amplified DNA blocks were inserted into the linearized AAV-TBG vector by Gibson Assembly using an NEBuilder HiFi DNA assembly kit (NEB) following the manufacturer’s instruction.

### AAV-HA-Sox9-P2A-Cre plasmid

Mouse *HA-Sox9-P2A* and *P2A-Cre* blocks were PCR-amplified from pWPXL-Sox9 (Addgene, #36979) and AAV:ITR-U6-sgRNA(backbone)-TBG-Cre-WPRE-hGHpA-ITR(Katsuda et al., 2022) respectively, using the primers listed **in Table S4**. The two blocks were inserted into the linearized AAV-TBG vector prepared as described above using the NEBuilder assembly kit.

### AAV-FLAG-Sox4-P2A-Cre plasmid

Mouse *FLAG-Sox4-P2A* and *P2A-Cre* blocks were PCR-amplified from pLVXT-Sox4 and AAV:ITR-U6-sgRNA(backbone)-TBG-Cre-WPRE-hGHpA-ITR respectively, using the primers listed **in Table S4**. The two blocks were inserted into the linearized AAV-TBG vector prepared as above using the NEBuilder assembly kit.

### CMV-FLAG-Sox4 plasmid

The *FLAG-Sox4 block was amplified from the AAV-FLAG-Sox4-P2A-Cre* plasmid using a forward primer: 5’-AGTGCTAGCGCCACCATGGACTACAAAGACG, and a reverse primer: 5’-TCGTGTACATCAGTAGGTGAAGACCAGGTTAGAGATGC. The DNA block was then digested with NheI-HF (NEB) and BsrGI-HF (NEB) and cloned into the CMV backbone vector prepared by linearizing the mEGFP-N1-YAPS127A-L318E plasmid (Addgene, #166465) using NheI-HF and BsrGI-HF.

### AAV preparation

90-100% confluent 293T cells in 15 cm dishes were replenished with 15 ml fresh DMEM (Thermo) supplemented with 2% FBS (Thermo) without antibiotics. For a 15 cm plate, 16 μg AAV8-Rep/Cap plasmid (Grompe Lab), 16 μg Ad5-Helper plasmid (Grompe Lab), 16 μg AAV transfer vector, and 144 μl of 1 mg/ml polyethylenimine (PEI) (Polysciences) were mixed in 9 ml OptiMEM (Thermo). After incubation at room temperature (RT) for 15 minutes, plasmid/PEI complex was added to 293T cells in a dropwise manner, and the plates were gently rocked to mix. After incubation in a CO_2_ incubator for 6 days, the cells and culture supernatant were harvested into 50 ml tubes and centrifuged at 1,900 ×g for 15 minutes. The supernatant was transferred to new tubes, and 1/40,000 volume of Benzonase (Sigma-Aldrich) was added and mixed thoroughly by inversion. After digestion of non-viral DNA by incubating at 37°C for 30 minutes, virus medium was centrifuged at 1,900 ×g for 15 minutes, and the supernatant was filtered with a 0.22 μm a filter unit containing a PES membrane (Thermo). Then, 1/4 volume of 40% polyethylene glycol 8000 (PEG8000) in 2.5 M NaCl was added and mixed thoroughly by inversion. Following overnight incubation at 4°C, precipitated AAV was collected by centrifugation at 3,000 ×g for 15 minutes. After removal of the supernatant, precipitate was homogenized in 100 μl PBS per 15 cm dish by through pipetting. Non-AAV precipitate was eliminated by centrifugation at 2,200 ×g for 5 minutes. Smaller debris were further removed by filtrating the eluted AAV with a 0.45 μm filter columns (Corning). This crude AAV was titrated by qPCR using the AAV8-TBG-Cre (Penn Vector Core) as a standard and a forward primer: 5’-GGAACCCCTAGTGATGGAGTT, and a reverse primer: 5’-CGGCCTCAGTGAGCGA and directly used for KO experiments without further purification.

### Immunofluorescence

Frozen sections were used for immunofluorescence. Tissue was fixed in zinc-formalin overnight, equilibriated in 30% sucrose/PBS, embedded in Tissue-Tek® O.C.T. compound (Sakura), and 8 μm sections were prepared. Remnant O.C.T. compound was removed by submerging the slides in PBS for 5 minutes. The specimens were permeabilized with 0.1% Triton X-100 (Fisher) in PBS at RT for 15 minutes. After treatment with the Blocking One Histo (Nacalai) at RT for 10 minutes, the specimens were incubated with primary antibodies (**Table S5**) diluted in 1/20× Blocking One Histo at RT for 1 hour or at 4°C overnight. The sections were then stained using donkey anti-rabbit, rat, or goat antibodies conjugated with AlexaFluor 488, AlexaFluor 594 or AlexaFluor 647 (Invitrogen) **(Table S5)** at 1/300 dilution and DAPI (Thermo) at 1/1000 dilution. After incubation at RT for 1 hour, the specimens were mounted in Aqua-Poly/Mount (Polysciences), and imaged using an Olympus IX71 inverted fluorescent microscope.

### Hematoxylin and Eosin (H&E) staining

H&E staining was performed by Penn Molecular Pathology and Imaging Core (MPIC).

### Hepatocyte isolation

Livers were perfused with 40 ml of HBSS (Thermo), followed by 40 ml HBSS with 1 mM EGTA (Sigma), then 40 ml HBSS with 5 mM CaCl_2_ (Sigma) and 40 µg/ml liberase (Sigma). Following perfusion, livers were mechanically dispersed with tweezers, resuspended in 10 ml wash medium (DMEM supplemented with 5% FBS), and filtrated with a 70 μm cell strainer. The cells were centrifuged at 50 ×*g* at 4°C for 5 minutes. Then, the cells were resuspended in complete percoll solution (10.8 ml percoll (Cytiva), 12.5 ml wash medium, and 1.2 ml 10× HBSS per liver) and centrifuged at 50 ×*g* at 4°C for 10 minutes. After a single wash with 10 ml medium, cells were spun at 50 ×*g* at 4°C for 5 minutes and then used for downstream experiments.

### Whole liver cell isolation from normal mice

Livers were digested by the two-step liberase perfusion as described above. Then, the undigested remaining tissue was transferred to a 1.5 ml tube, minced with surgical scissors, and further digested with 10× concentrated liberase (∼ 430 μl/tube of 400 μg/ml in HBSS with 5 mM CaCl_2_) at 37°C for 30 minutes while vortexing the sample several times intermittently. The digested tissue was filtered with a 70 μm cell strainer and combined with the cell suspension digested previously. The cells were then centrifuged at 300 ×*g* at 4°C for 5 minutes. Then, the cells were suspended in 10 ml ACK lysis buffer (Quality Biological) and incubated on ice for 10 minutes to remove red blood cells. The cells were then collected by centrifugation at 300 ×*g* at 4°C for 5 minutes and used for downstream analyses.

### Whole liver cell isolation from DDC-treated mice

Livers were digested by the two-step liberase perfusion as described above. Following perfusion, livers were submerged in 10 ml fresh HBSS with 5 mM CaCl_2_, 40 µg/ml liberase and 40 µg/ml DNaseI (Millipore) in a C-tube (Miltenyi) and further digested using a gentleMACS Octo dissociator (Miltenyi) with a heating unit using the “37C_m_LIDK_1” protocol. Dissociated tissue was diluted in flow buffer (HBSS, pH 7.4) supplemented with 25 mM HEPES (Thermo), 5 mM MgCl_2_ (MedSupply Partners), 1× Pen/Strep (Thermo), 1× Fungizone (Thermo), 1× NEAA (Thermo), 1× Glutamax (Thermo), 0.3% glucose (Sigma), 1× sodium pyruvate (Thermo) supplemented with 40 µg/ml DNaseI (hereafter flow buffer(+)). Undigested tissue was removed by passing it through a 70 μm cell strainer, and the cells were centrifuged at 300 ×*g* at 4°C for 5 minutes. Then, the cells were suspended in 10 ml ACK lysis buffer and incubated on ice for 10 minutes. The cells were then collected by centrifugation at 300 ×*g* at 4°C for 5 minutes and used for downstream analyses.

### Flow cytometry

Cells were resuspended in 2-3 ml flow buffer(+), and filtered with a 35 μm cell strainer equipped with a FACS tube (BD). The cell suspension was then transferred to a round-bottom 96 well plate at 100-150 μl/well and centrifuged at ∼800 ×*g* at 4°C for 1 minute with a slow brake. The cells were then resuspended in 100 μl/well of flow buffer(+) containing fluorophore-conjugated antibodies (**Table S5**) and incubated on ice for 20 minutes. After two washes in flow buffer(+) (150-200 μl/well, ∼800 ×*g* at 4°C for 1 minute, slow brake), the cells were resuspended in flow buffer(+) containing 1/1,000× TO-PRO-3 (Thermo) and analyzed using an LSR II flow cytometer (BD).

### Fluorescence-activated cell sorting (FACS) of DDC-treated whole liver cells

Cells were resuspended in 5 ml flow buffer(+) by centrifugation at 300 ×*g* at 4°C for 5 minutes. After removal of the supernatant, the volume was increased to 1-1.5 ml with flow buffer(+), and 1/100 volume of rat anti-Cd45, rat-anti-Cd11b and rat anti-Cd31 antibodies (**Table S5)** were added and incubated on ice for 10 minutes. After washing with 2 ml flow buffer(+) (300 ×*g* at 4°C for 5 minutes, slow brake), the cells were resuspended in 5 ml flow buffer(+), 600 μl Dynabeads-anti-rat IgG (Thermo) were added, and the cell/bead mixture was incubated at 4°C for 30 minutes with gentle tilting and rotation. The suspension was transferred to 5 ml FACS tubes and placed on a DynaMag™-5 Magnet (Thermo) for 2 minutes. The supernatant was transferred to a new tube, and the cells were collected by centrifugation at 2,000 rpm (∼ 800×*g*) at 4°C for 2 minutes with a slow brake. The cells were then resuspended in MACS buffer (PBS, 0.5% BSA, 2 mM EDTA) to the final volume of approximately 1.5 ml, and 150 μl CD326 (EpCAM) MicroBeads (Miltenyi) were added. After incubation at 4 °C for 15 minutes, the cells were washed with an equal volume of MACS buffer then centrifuged at 2,000 rpm (∼ 800×*g*) at 4°C for 2 minutes with a slow brake. Cells were resuspended in 2 ml MACS buffer, and Epcam+ cells and Epcam-cells were separated using LS columns (Miltenyi) following the manufacturer’s instruction (4 columns were used per animal; 0.5 ml suspension/column). The cells were then collected by centrifugation at 2,000 rpm (∼ 800×*g*) at 4°C for 2 minutes with a slow brake. The cells were then resuspended in 0.5-1 ml flow buffer(+), and approximately 10-15 μl of each were set aside for fluorescence-minus one (FMO) controls (FMO-Brilliant Violet 421 (BV421): all stained except BV421-Cd24; and FMO-PE/Dazzle 594: all stained except PE/Dazzle594-Epcam), and stained in a 96 well round bottom plate as described earlier. The cells to be used for FACS were stained with BV421-Cd24 (Biolegend), PE/Dazzle594-Epcam (Biolegend), PE/Cy7-Cd11b/Cd31/Cd45 (Table S5) at 1:100 dilution in 15 ml tubes on ice for 20 minutes. After washing in 2 ml flow buffer(+) once by centrifugation at 2,000 rpm (∼ 800×*g*) at 4°C for 2 minutes with a slow brake, the cells were resuspended in 1-3 ml flow buffer(+) with 1/1,000 TO-PRO-3, and the cells were sorted on an Aria II sorter (BD).

### Total RNA isolation and reverse transcription

Total RNA was extracted using the NucleoSpin RNA Kit (Takara) following the manufacturer’s instructions. Approximately 500 ng of RNA was reverse transcribed in 20 μl volume using High Capacity cDNA Reverse Transcription Kit (Thermo). cDNA was diluted at 1:20 ratio in water and used for qPCR.

### qPCR

qPCR was performed at 10 μl/well using the Bio-Rad CFX 384 qPCR machine (Bio-rad). Each well contained: 3 μl diluted cDNA, 0.25 μl each of 10 μM forward and reverse primers (**Table S6**), 1.5 μl H_2_O and 5 μl SsoAdvanced SYBR reagent (Bio-rad).

### RNA-Seq

Library preparation and sequencing were performed by Novogene (Sacramento, CA) using a Novaseq 6000 (Illumina).

### Bioinformatics for RNA-Seq

Reads were aligned to the mouse genome (GRCm39) using STAR aligner with default parameters ^51^. Gene-count matrices were produced by featureCounts^52^. To compare gene expression between samples, expression levels were normalized based on the “median of ratios” method using DESeq2(Love et al., 2014). To compare expression levels between different genes, “Transcripts per million (TPM)” normalization was performed using Salmon^53^. To build the gene sets for gene set enrichment analysis (GSEA), differential gene expression analysis was performed between hepatocytes and Rep_early cells using the median normalized data with the cut-off values of p.adj < 0.05 and |log2(fold-change)| ≥ 1. The generated gene sets are listed in **Table S1**.

### Bioinformatics for single cell RNA-Seq

Data were obtained in our earlier study^27^ and deposited with the accession number GSE157698. Using R Seurat package, Seurat objects were created with arguments “min.cells = 3, min.genes = 200.” The cells were computationally filtered for YFP+ cells using “subset = nFeature_RNA > 200 & nFeature_RNA < 4000 & percent.mt < 0.25 & percent.yfp > 0.” Pesudotime analysis was performed using the monocle3 R package^29^. Briefly, the data were visualized with UMAP, and outlier cells were manually removed using choose_cells function in monocle3. Then, a trajectory was generated using cluster_cells and learn_graph functions, which calculated the pseudotime along the DDC-induced biliary reprogramming of hepatocytes. Each cell was assigned a pseudotime using order_cells function. Gene expression changes along the psuedotime were visualized using the plot_genes_in_pseudotime function.

### ATAC-Seq

50,000 cells were isolated from three AAV8-EV- or AAV8-*HA-Sox4-P2A-Cre*-injected (1 × 10^12^ gc/mouse) liver samples at 4 dpi and used as input for ATAC-Seq library preparation. Libraries were prepared as described^34^ with minor modifications. Briefly, nuclei were isolated from the cells using a solution of 10 mM Tris-HCl pH 7.4, 10 mM NaCl, 3 mM MgCl_2_, 0.1% IGEPAL CA-630. Immediately following isolation, the transposition reaction was conducted using Tn5 transposase (Diagenode) and TD buffer (Illumina) for 30 minutes at 37 °C. Transposed DNA fragments were purified using a Qiagen MinElute Kit, barcoded, and PCR amplified for 7-9 cycles depending on the samples using NEBNext High Fidelity 2× PCR master mix (New England Biolabs). The optimal cycle number was determined empirically each time by qPCR. The libraries were then purified with AMPure XP beads. Paired-end 150 × 2 sequencing was performed by Novogene (Sacramento, CA) using a NovaSeq 6000 (Illumina).

### Bioinformatics for ATAC-Seq

Reads were aligned to the mouse genome (mm10) using Bowtie2 (Langmead and Salzberg, 2012) with options “--very-sensitive -X 1000 --dovetail -1”, and duplicates were removed using Picard (http://broadinstitute.github.io/picard/). Peak calling was performed using MACS2^54^ with an FDR of 0.01 (default setting). Motif analysis was performed using HOMER (http://homer.ucsd.edu/homer/motif/) with the option “-size 300 –mask”. Differential peak analysis was performed using triplicate samples (BAM files and peaks called independently for each replicate) using DiffBind^55^ and DESeq2. Differential peaks were then annotated to the nearest genes using the annotatePeak function of ChIPSeeker R package^56^, and the expression of these genes during DDC-induced reprogramming was analyzed using the RNA-Seq data as described above (GSE218945). Using the genes enriched in the newly closed/opened regions, gene ontology analysis was performed using the “enrichGO” function in clusterProfiler R package^57^. Analysis of the genomic distribution of the ATAC peaks was performed using plotAnnoBar and plotDistToTSS functions in ChIPSeeker. For GSEA, the annotated genes were ranked based on the log-fold change between EV and *Sox4*-expressing hepatocytes calculated by DESeq2, and the ranked gene list was used as input for the fgsea R package (fast preranked GSEA)^44^. For visualization using the Integrative Genomics Viewer (IGV) track browser^58^ and deepTools (for generation of heatmaps)^59^, BAM files were converted to bigwig files using bamCoverage with the “reads per genome coverage (RPGC)” normalization method.

Footprinting analysis was performed on replicate-merged BAM files using TOBIAS^35^ with the default settings. For global foot printing analysis, all the foot prints that were assigned “bound” either in EV_Hep or Sox4_Hep were used as the input for the analysis (default setting of the TOBIAS pipeline). The output “bindetect_results” file was imported to R for visualization. For visualization of the aggregate footprints, corrected bigwig signals were retrieved using the ScoreBed function, and plotted using the ggplot2 R package. TOBIAS scores were retrieved using the “PlotAggregate” function with the option, “--output-txt”, and visualized using ggplot2.

### CUT&RUN-Seq

CUT&RUN DNA was prepared following EpiCypher® CUTANA™ CUT&RUN Protocol v2.0 with minor modification. Briefly, 500,000 cells were isolated from AAV8-*HA-Sox4-P2A-Cre*-injected (1 × 10^12^ gc/mouse) livers at 18 hpi and 4 dpi (n = 3, each timepoint). The cells were washed in wash buffer (20 mM HEPES, pH 7.5, 150 mM NaCl (Sigma), 0.5 mM spermidine (Sigma) and EDTA-free protease inhibitors (Roche)) twice followed by centrifugation at 600 ×*g* at 4 °C for 3 minutes. The cells were resuspended in 150 μl wash buffer, 15 μl Concanavalin A-coated magnetic beads (EpiCypher) were added, and the cells/bead conjugate was bound to a magnet. After removal of the supernatant, the cells were resuspended in 50 μl antibody reaction buffer (wash buffer with 5% digitonin and 0.5 M EDTA), 0.5 μl antibody was added (**Table S5**) and incubated at 4°C overnight. Following two washes in 200 μl permeabilization buffer (wash buffer with 5% digitonin), beads were resuspended in 50 μl permeabilization buffer. Then, 2.5 μl pAG-MNase (EpiCypher) was added, and the samples were incubated at RT for 10 minutes. While on the magnet, the supernatant was removed, and the samples were washed twice in 200 μl cold permeabilization buffer. Following resuspension in 50 μl permeabilization buffer, 1 μl 100 mM CaCl_2_ was added to activate MNase, and MNase digestion was performed at 4°C for 2 hours. The reaction was stopped by adding 33 μl STOP buffer (340 mM NaCl, 20 mM EDTA, 4 mM EGTA, 10 μg/ml RNase A, 50 μg/ml glycogen), and 1 μl E-Coli spike-in DNA (0.5 ng/μl) (EpiCypher) to each tube. After incubation at 37°C for 10 minutes, beads were bound to the magnet, and the supernatant was transferred to a new tube. The DNA was cleaned up with the NEB Monarch kit (NEB), eluted in 12 μl elution buffer, and used as CUT&RUN DNA. Library preparation was performed using NEBNext Ultra II End Prep kit (NEB) with a slightly modified protocol. Briefly, after adaptor ligation, HA-Sox4 CUT&RUN DNA samples were purified with 1.75× AMPure XP beads (Beckman), while histone and IgG CUT&RUN samples were purified with 1.1× AMPure XP beads. 14-cycle PCR with index primers (NEB) was performed (initial denaturation: 1 cycle of 98 °C for 45 seconds; annealing/extension: 14 cycles of 98°C for 15 seconds and 60°C for 10 seconds; final extension: 72°C for 1 minute). Finally, the libraries were purified with AMPure XP beads (for HA-Sox4 0.8× followed by 1.2×; for histone and isotype control 0.9×). For samples with adaptor contamination, further AMPure bead cleanup or gel extraction was performed. Sequencing was performed using an Illumina NextSeq500 with a 150 cycle mid-output reagent kit (75-bp paired end) and an Illumina NextSeq2000 with 100 cycle S2 reagent kit (65-bp paired end).

### Bioinformatics for Sox4 CUT&RUN-Seq

Sox4 and isotype control CUT&RUN-Seq data were obtained in two separate sequencing experiments. The 75-bp data obtained from the NextSeq500 experiment were first trimmed with Cutadapt (ver. 4.1) shorten the reads to 65-bp in order to merge into single 65-bp fastq files. Reads were aligned to the mouse genome (mm10) using Bowtie2 with the option “-X 1000”, and duplicates were removed using Picard. For Sox4 differential peak analysis, peak calling was performed for each of the three biological replicates using MACS2 with each of the Sox4 BAM files and the combined isotype control BAM file for each timepoint. The FDR was set to 0.1 for peak calling of individual samples, and the called peaks showed robust enrichment of Sox binding motifs as confirmed by HOMER. This analysis generated 13,081 ± 5,151 and 28,160 ± 13,294 peaks for 18 hpi and 4 dpi samples, respectively. Using these peaks and BAM files for each replicate, differential peak analysis was performed using DiffBind and DESeq2 with the FDR cut-off set to 0.05 (default setting). By this analysis, we obtained 2,327 peaks enriched for 18 hpi and 3,136 peaks enriched for 4 dpi. For GSEA, all the Sox4 peaks were annotated to the nearest genes using the annotatePeak function of ChIPSeeker R package(Yu et al., 2015), and the annotated genes were ranked based on the log-fold change between 18 hpi and 4dpi samples calculated by DESeq2, and the ranked gene list was used as input for fgsea R package (fast preranked GSEA)^44^. After confirmation of reproducibility across the three biological replicates by PCA mapping (**Supplementary Fig. 5**), we re-performed peak calling using MACS2 with replicate-merged Sox4 samples and the corresponding replicated-merged isotype control without FDR filtering, which generated 9,463 and 19,362 Sox4 peaks at 18 hpi and 4 dpi samples respectively. Motif analysis was performed using HOMER with the option “-size 300 –mask.” For visualization with IGV and deepTools, Sox4 BAM files were normalized by subtracting the isotype control signals using the bamCompare function.

Quantification of total Sox4 binding was performed by calculating scale factors for each Sox4 sample using the E-coli spike-in controls. Briefly, the reads were aligned to E-coli genome (K12_MG1655) using Bowtie2, and the scale factors were calculated as the ratios of “(Number of mouse-aligned reads)/(#Number of E-coli-aligned reads)”.

### Bioinformatics for histone post-translational modification CUT&RUN-Seq

CUT&RUN-Seq of histone post-translational modification was performed with either NextSeq500 (75-bp) or NextSeq2000 (65-bp) using the same samples as used for Sox4 CUT&RUN-Seq. Reads were aligned to the mouse genome (mm10) using Bowtie2 with the option “-X 1000,” and duplicates were removed using Picard. For visualization using IGV and deepTools, BAM files were converted to bigwig files using bamCoverage with the “read counts per million (CPM)” normalization method. Once we confirmed the reproducibility across the three biological replicates by PCA mapping (**Supplementary Fig. 5**), we combined the fastq files (75-bp data were shortened to 65-bp using Cutadapt) and obtained single bigwig files for each group. Unless otherwise mentioned, all the heatmaps and aggregate plots in this manuscript are shown for the replicate-merged data.

### Immunocytochemistry of HA-Sox4 prior to CUT&RUN-Seq

After the overnight primary antibody reaction as described above, the cells were incubated with 1/300 AlexaFluor 594-conjugated anti-rabbit IgG (Invitrogen) diluted in permeabilization buffer at RT for 1 hour. Then, the cells were spread onto a 24 well plate for imaging.

### Protein Expression and Purification

CMV-FLAG-Sox4 was produced using 10× 15 cm plates of 293T cells. Approximately 70% confluent plates were replenished with 15 ml/plate fresh DMEM (Thermo) supplemented with 2% FBS (Thermo) without antibiotics. For a 15 cm plate, 48 μg CMV-FLAG-Sox4 plasmid and 144

μl of 1 mg/ml PEI were mixed in 9 ml OptiMEM. After incubation at RT for 15 minutes, plasmid/PEI complex was added to 293T cells in a dropwise manner, and the plates were gently shaken back and forth to mix the medium evenly. After incubation in a CO_2_ incubator for 2 days, the cells were harvested by standard trypsinization. After washing twice in 10 ml PBS by centrifugation at 800×g for 2 minutes, the cells were resuspended in lysis buffer (50 mM Tris-HCl, pH 7.4, 150 mM NaCl, 1 mM EDTA, 1% TritonX-100, 1x protease/phosphatase inhibitor (Pierce)) to the final volume of approximately 12 ml. After incubation at RT on a rotator for 30 minutes, the suspension was split into 300 μl aliquots, and sonicated using the Bioruptor Plus Sonicator (Diagenode) with High Power for 5 cycles (30 seconds ON and 30 seconds OFF for 5 minutes per cycle). Sonicated samples were centrifuged at 4°C at 14,000×g for 15 minutes, and the supernatant was collected and combined into one tube. Small debris were further removed by passing the samples through a 0.45 μm PVDF filter (Millipore). An affinity chromatography column was prepared with ∼0.6 ml anti-FLAG M2 beads (Sigma) following the manufacturer’s instructions. The lysate was loaded onto the column under gravity flow. The column was washed with 12 ml (∼20× volume) TBS (50 mM Tris-HCl, 150 mM NaCl, pH 7.4), and FLAG-Sox4 protein was eluted by 5 rounds of competitive elution with 1× column volume (0.6 ml) of 100 μg/ml FLAG peptide (Sigma) in TBS. Eluted FLAG-Sox4 was concentrated to ∼50 μl using 30K MWCO columns (Thermo). Finally, the buffer was exchanged with 20 mM Tris-HCl, pH 7.5, 150 mM NaCl, 1 mM DTT using 7K MWCO Pierce Zeba™ Desalt Spin Columns (Thermo). FLAG-Sox4 concentration was estimated by quantifying the band densities corresponding to ∼65 kDa by SDS-PAGE using BSA standards (**Supplementary Fig. 6**).

### Nucleosome Preparation

The 162 bp *LIN28B* DNA fragment corresponds to the genomic location: hg18-chr6: 105,638,004-105,638,165 AGTGGTATTAACATATCCTCAGTGGTGAGTATTAACATGGAACTTACTCCAACAATA CAGATGCTGAATAAATGTAGTCTAAGTGAAGGAAGAAGGAAAGGTGGGAGCTGCCA TCACTCAGAATTGTCCAGCAGGGATTGTGCAA GCTTGTGAATAAAGACA The DNA sequence was created by PCR with end-labeled primers: Cy5-*LIN28B*-Fw: 5Cy5/AGTGGTATTAACATATCCTCAGTGGTG; *LIN28B*-Rv: TGTCTTTATTCACAAGCTTGCACAA. The 162 bp fluorescent-tagged DNA fragments were gel extracted. The nucleosomes were reconstituted by salt dilution. Briefly, a reaction mixture was prepared with 10-20 pM labeled DNA fragment and octamer at a 1:1 DNA:histone ratio and diluted to 10 μl in 10 mM Tris-HCl, pH 8.0, 2 M NaCl, and then incubated at RT for 30 minutes. Then, 3.5, 6.5, 13.5, and 46.5 μl of 10 mM Tris-HCl pH 8.0 was added at 30 minutes intervals, which brought the reactions to 1.48, 1.0, 0.6 and 0.25 M NaCl (80 μl total volume). The mononucleosomes were further purified by 10 – 30 % glycerol gradient followed by dialysis with 10 mM Tris-HCl, pH 8.0, and 1 mM BME. The reconstituted nucleosomes were heat-shifted by incubating at 37 °C for 30 min.

### Binding Reactions

The end-labeled oligonucleotides containing specific or non-specific sites, *LIN28B*-DNA, and *LIN28B*-nucleosomes were incubated with recombinant proteins in DNA-binding buffer (10 mM Tris-HCl (pH7.5), 1 mM MgCl2, 1 mM DTT, 50 mM KCl, 0.3 mg/ml BSA, 5% Glycerol) at RT for 30 minutes. Free and bound DNA were separated on 5% non-denaturing polyacrylamide gels run in 0.5× Tris–borate–EDTA and visualized using a PhosphorImager using Cy5 fluorescence setting (excitation at 633 nm and emission filter 670 BP 30) and high sensitivity setting.

### Bioinformatics for ChIP-Seq

ChIP-Seq data were downloaded from either the GEO database with the accession numbers GSE57559 for H2B and H3^45^; GSE137066 for adult liver Hnf4a^37^; GSE167287 for colon epithelial cell Hnf4a^46^; GSE53736 for the adult liver Rxra data^36^; GSE29184 for the mouse adult liver H3K4me3^60^; GSE111502 for Yap of DDC-treated hepatocytes^41^.

When biological replicates were available, the downloaded fastq files were first combined. Reads were aligned to the mouse genome (mm10) using Bowtie2 with the option “-X 1000,” and duplicates were removed using Picard. For visualization using the IGV and deepTools, BAM files were converted to bigwig files using the bamCoverage with the CPM normalization method. Peak calling was performed using MACS2^54^ with an FDR of 0.01 (default setting) without control inputs for H2B, H3 and H3K4me3 and with control inputs for liver/colon Hnf4a and Rxra.

The bigwig file for the Yap-ChIP-Seq data was directly downloaded from GEO and used for IGV visualization.

### Bioinformatics for MNase-Seq

Low- and high-level MNase-Seq data were downloaded from GEO with the accession number GSE57559^45^. Reads were aligned to the mouse genome (mm10) using Bowtie2 with the option “-X 1000,” and duplicates were removed using Picard. BAM files of biological replicates were combined using “samtools merge,” and inputted into DANPOS3^61^ to calculate the nucleosome occupancy. For visualization using the IGV and deepTools, the DANPOS3-generated BAM files were converted to bigwig files using the bamCoverage with the RPGC normalization method.

### Generation of the list of active liver enhancer loci

Liver-specific active enhancers in this study are defined as genomic regions which are p300+ H3K4me1+ H3K4me3-H3K27ac+ with DNA hypersensitivity (DHS). To obtain the list of these regions, we used a previously published adult mouse liver-specific enhancer list that was identified by Shen and colleagues based on ChIP-Seq data of p300, H3K4me1, H3K4me3 (ENCODE, GSE29184)(Shen et al., 2012). Since H3K27ac predominately marks active enhancers, we filtered these enhancers so that their central 1 kb regions have 300 or more base-pair overlap with adult mice liver H3K27ac peaks that were identified in the same study (ENCODE, GSE29184)^60^. For filtering DHS-positive adult liver-specific enhancers, we further filtered them so that their central 1kb regions have 300 or more base-pair overlap with adult mice liver DHS peaks (ENCODE, GSM1014195).

### Generating a list of active liver promoter loci

Active promoters in this study are defined as H3K4me3+ transcription start sites (TSSs) in the adult liver in the proximity of highly expressed genes. H3K4me3+ regions were defined as H3K4me3 ChIP-Seq peaks (GSE29184) with 0.5 kb extension bilaterally. Highly expressed genes in hepatocytes were defined as the genes ranked within the top 25% in normal hepatocytes using the TPM-normalized RNA-Seq data obtained above (GSE218947). The H3K4me3+ regions were annotated with the nearest genes (output of MACS2), and then filtered with the list of hepatocyte-highly expressed genes. TSSs included in these regions were regarded as active liver promoter loci (n = 13,458).

### Bioinformatics for microarray of YAPS127A-expressed hepatocytes

Microarray data of YAPS127A-expressed mouse hepatocytes were downloaded from GEO with the accession number GSE55560^26^. We used a pipeline provided by Klaus and Reisernauer (https://bioconductor.org/packages/release/workflows/vignettes/maEndToEnd/inst/doc/MA-Workflow.html). Briefly, the “robust multichip average (RMA)” algorithm was used for background correction, quantile normalization and data summarization using the oligo R package. The probes were filtered with a cut-off median signal intensity > 4.

