## Extended_Data_Figures for "Physiological reprogramming *in vivo* mediated by Sox4 pioneer factor activity"

### Extended Data Fig.1

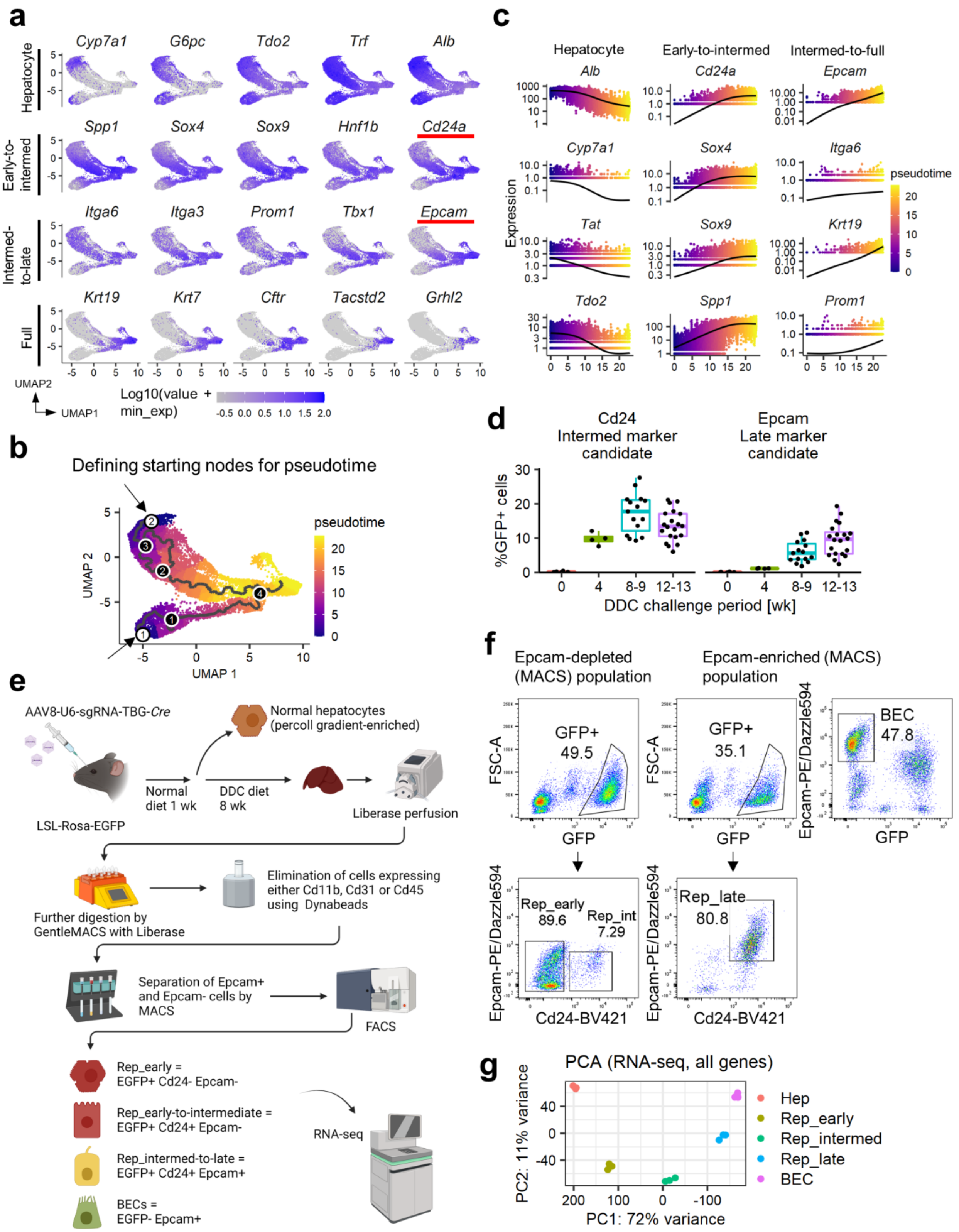

**Extended Data Fig. 1. Surface markers Cd24 and Epcam characterize reprogrammed cells at early-to-intermediate and late stages respectively.**

(a) Single cell RNA-Seq (scRNA-Seq) using YFP+ cells isolated from Rosa26-LSL-YFP mice challenged with 0.1% DDC (n = 2 wild type mice)[1]. Hepatocyte-derived cells (those with at least one *YFP* read count) were visualized by UMAP projection for hepatocyte genes and biliary reprogramming genes at different stages. *Cd24a* and *Epcam*, nominated as candidate surface markers of early-to-intermediate and intermediate-to-late stages of reprogramming, are highlighted with red underscores. Data were downloaded from GEO (accession number GSE157698).

(b) Pseudotime was defined using the monocle3 R package based on the expression pattern of known hepatocyte and biliary/reprogramming marker genes as shown in **(a)** (see **Methods** for a detail analytical pipeline). The initial points of the trajectory highlighted as white nodes were determined based on the expression patterns of the above mentioned known hepatocyte and biliary reprogramming marker genes.

(c) Pseudotime analyses nominated *Cd24* and *Epcam* as candidate surface markers for early-to-intermediate and intermediate-to-late reprogrammed cells, respectively. The relevant gating strategy and representative flow plots are described in **Supplementary Fig. 1.**

(d) Flow cytometry confirms that Cd24 is expressed at the protein level earlier than Epcam in the weeks post injection (wpi) following DDC challenge (n = 4 for 0 wpi; n = 4 for 4 wpi; n = 15 for 8-9 wpi; n = 21 for 12-13 wpi).

(e) Schematic showing the strategy for isolating early reprogrammed cells (EGFP+ Cd24- Epcam-), early-to-intermediately reprogrammed cells (EGFP+ Cd24+ Epcam-), and intermediate-to-late reprogrammed cells (EGFP+ Cd24+ Epcam+).

(f) Representative flow cytometry plots and gating used for isolating cell populations by FACS. The relevant gating strategy is described in **Supplementary Fig. 2 and 3**.

(g) PCA of whole transcriptomes (RNA-Seq) for the indicated cell populations along the hepatocyte-to-biliary continuum (n = 3 per group).

### Extended Data Fig. 2

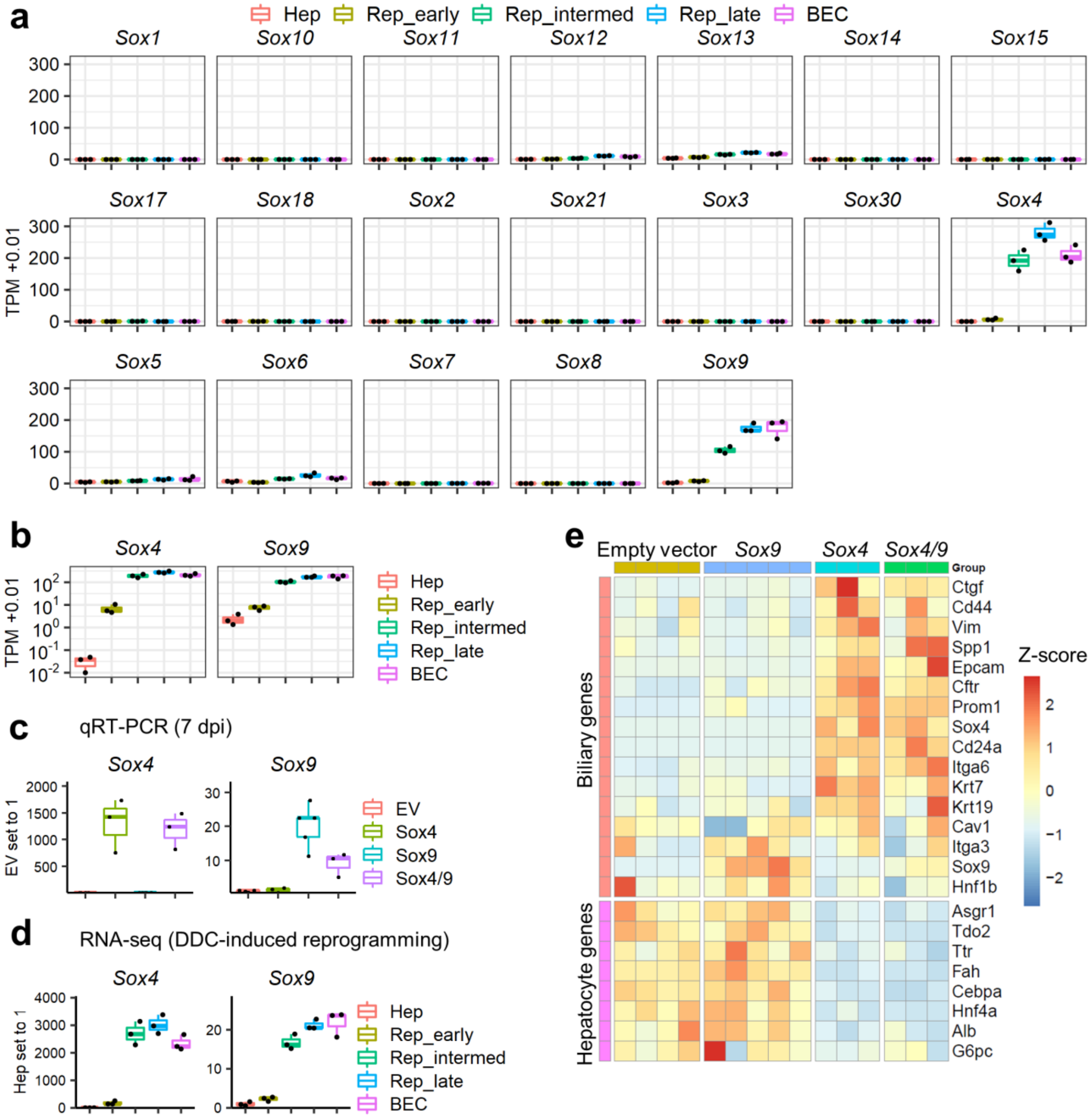

**Extended Data Fig. 2. *Sox4* and *Sox9* are the most abundantly expressed *Sox* genes in the liver upon DDC-induced reprogramming.**

(a) Expression profiles for all Sox genes as quantified by RNA-Seq. Expression levels are represented by TPM normalization (n = 3).

(b) Data for *Sox4* and *Sox9* from panel **(a)** with the y-axis transformed to a log scale to illustrate differences in baseline expression in hepatocytes (Hep) (n = 3).

(c) qRT-PCR of *Sox4* and *Sox9* at 7 dpi. Expression levels are normalized to *Actb*, with empty vector (EV) hepatocyte expression levels set to one (n = 3-5).

(d) Results of RNA-Seq showing the expression of *Sox4* and *Sox9* during DDC-induced biliary reprogramming. Expression levels are normalized to the median expression for each sample, with the expression level of hepatocytes (Hep) set to one (n = 3).

(e) Heatmap of qRT-PCR data obtained at 7 dpi for biliary genes and hepatocyte genes. Expression levels are normalized to *Actb* and shown as z-scores for each gene (row) (n = 3-5).

### Extended Data Fig. 3

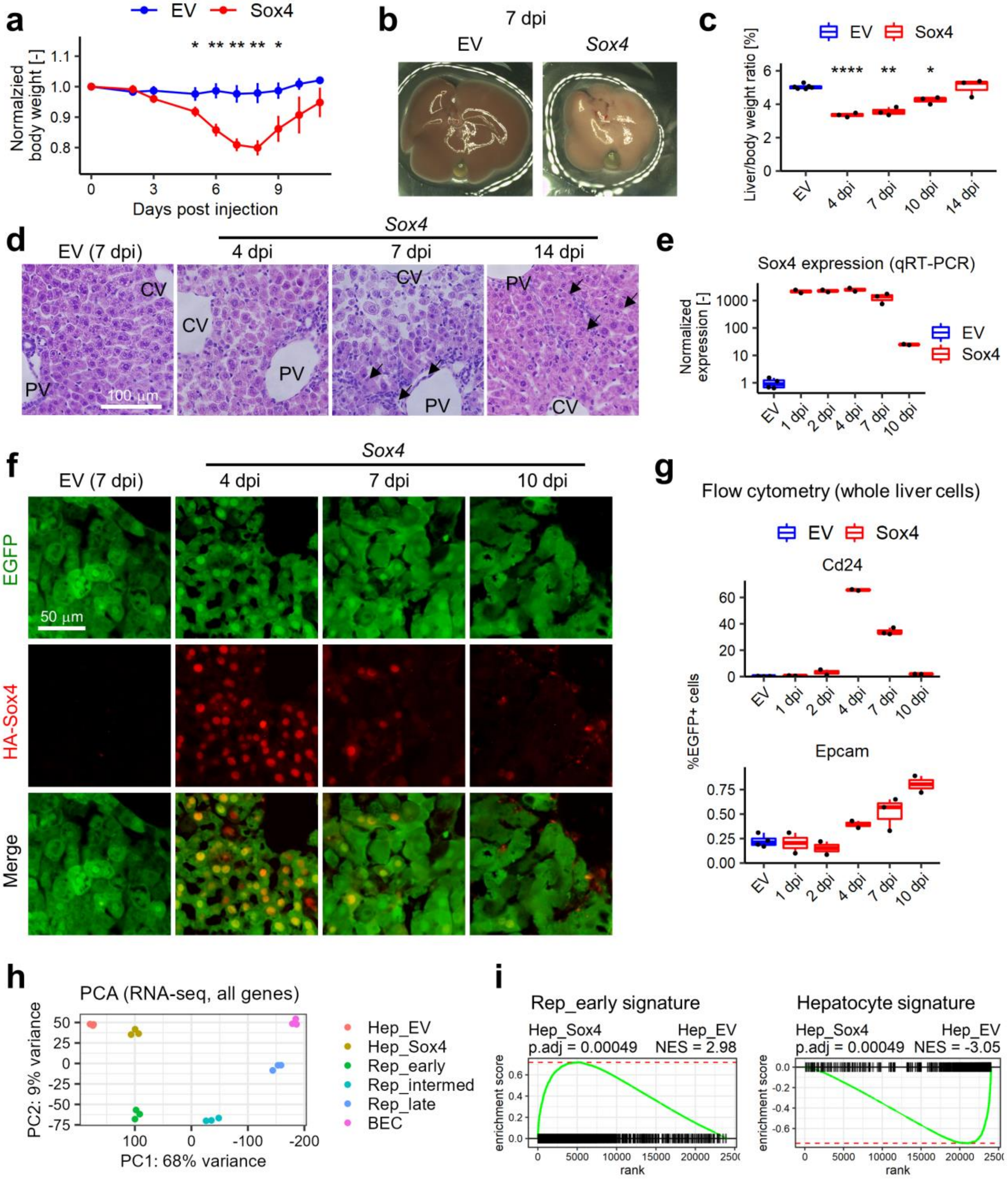

**Extended Data Fig. 3. Characterization of phenotypic changes in *Sox4* expressing mice and kinetics of AAV-mediated *Sox4* expression in hepatocytes.**

(a) Body weight changes following *Sox4* expression in hepatocytes. Data are shown as mean ± SE (n = 7 per group).

(b) Representative macroscopic liver images of an AAV-*HA-Sox4*-injected mouse and a control empty vector-injected mouse at 7 dpi.

(c) Liver-to-body weight ratios are shown for the designated time points after AAV-*HA-Sox4* injection (n = 3 per time point). Data points for the empty vector (EV)-injected control are combined from 4, 7, 10, and 14 dpi (n = 6 total).

(d) HE staining of AAV-*HA-Sox4*-injected livers and control livers (empty vector, EV) at the designated time points. Arrows indicate atypically proliferating ductal cells extending from the portal area to the parenchymal regions. PV and CV indicate peri-portal vein area and peri-central vein area, respectively. Scale bar = 100 μm.

(e) Kinetics of *Sox4* mRNA expression as assessed by qRT-PCR following AAV-*HA-Sox4* injection (n = 2-3 per time point for *Sox4* samples). Data points for the empty vector control are combined from 4, 7 and 10 dpi (n = 4 total). Expression levels are normalized to *Actb*, and the expression level of the empty vector control is set to one.

(f) Kinetics of Sox4 protein expression as assessed by immunofluorescence using an anti-HA-tag antibody. Scale bar = 50 μm.

(g) Kinetics of protein expression of the early-to-intermediate reprogramming marker Cd24 and the intermediate-to-late reprogramming marker Epcam as assessed by flow cytometry using the whole liver cells isolated at the designated time points (n = 2-3 per time point). Data points for the empty vector control are combined from 4, 7 and 10 dpi (n = 4 total).

(h) PCA plot of RNA-Seq data from AAV-empty vector- and AAV-*HA-Sox4*-injected mice, along with DDC-induced reprogrammed cells (n = 3 per group).

(i) GSEA using RNA-Seq data as input for comparison of empty vector and *Sox4* hepatocytes, using the hepatocyte-enriched and early reprogrammed cell (Rep_early)-enriched signatures.

Statistical differences were calculated by t-test. *p $< 0.05$, **p $< 0.01$, ****p$<0.0001$

### Extended Data Fig. 4

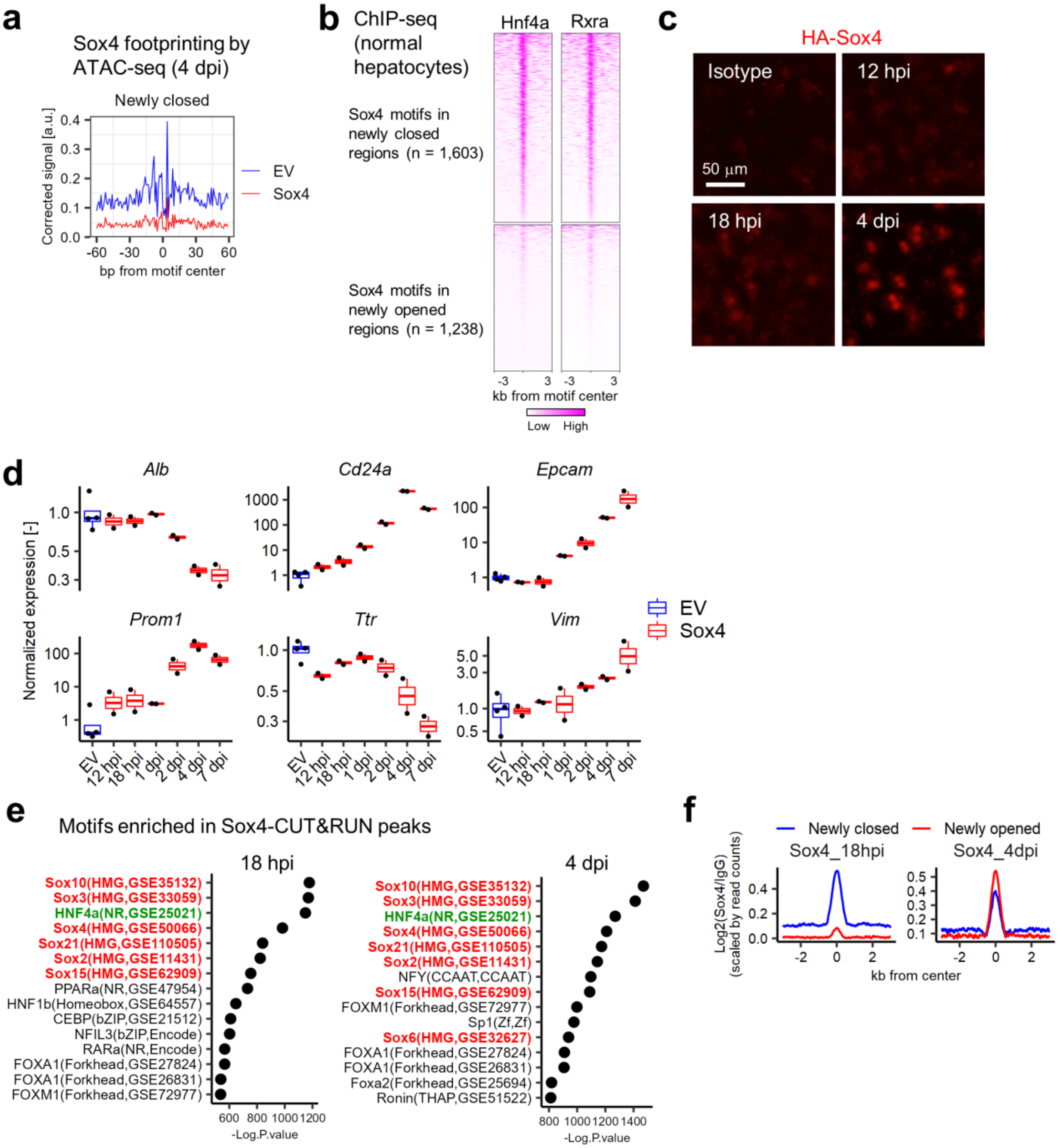

**Extended Data Fig. 4. Kinetic analysis of phenotypic changes following *Sox4* expression and profiling the Sox4 binding sites in hepatocytes by CUT&RUN-Seq.**

(a) Results of DNA footprint analysis for the Sox4 binding motif comparing empty vector and *Sox4* hepatocytes at newly closed regions as defined in **Fig. 2a**. TOBIAS software was used with the default setting, and motifs assigned "bound" either for empty vector or *Sox4* hepatocytes were used for visualization of the averaged signals (see **Methods** for a detailed analytical pipeline).

(b) Published data from ChIP-Seq of hepatocyte transcription factors Hnf4a[2] and Rxra[3], visualized for Sox4 motifs assigned as "bound" by TOBIAS in either empty vector or *Sox4* hepatocytes found in newly closed and newly opened regions.

(c) Immunofluorescence staining for HA-Sox4 protein in primary hepatocytes at the indicated timepoints following infection. Percoll-enriched hepatocytes were treated with digitonin for permeabilization, captured on a magnet using concanavalin A-conjugated beads, and then incubated with an anti-HA antibody overnight. On the next day, HA-Sox4 was detected using an Alexa fluor594-conjugated secondary antibody.

(d) Kinetic analysis of phenotypic changes following *Sox4* expression as assessed by qRT-PCR (n = 2 per time point for *Sox4* expressing samples). Data points for the empty vector (EV) control are combined from 4, 7 and 10 dpi (n = 4 total). Expression levels are normalized to *Actb*, with the expression level of the empty vector control set to one.

(e) Results of HOMER motif analysis using all Sox4 peaks from CUT&RUN-Seq (n = 9,463 at 18 hpi; n = 19,362 at 4 dpi). The top 15 motifs from each timepoint ranked as p-values are shown. Peak calling was performed with the MACS2 peak caller along with Isotype IgG control, with FDR cutoffs set to 0.99, the most relaxed condition. Sox motifs, highlighted in red, were highly ranked. The Hnf4a motif, highlighted in green, was the third most significantly enriched motif at both 18 hpi and 4 dpi.

(f) Averaged aggregate plots for Sox4 CUT&RUN-Seq which are represented as the log2 ratio of Sox4 signals / IgG signals. The plots correspond to the Sox4 heatmaps shown in **Fig. 2f,** right two columns.

#### Extended Data Fig. 5

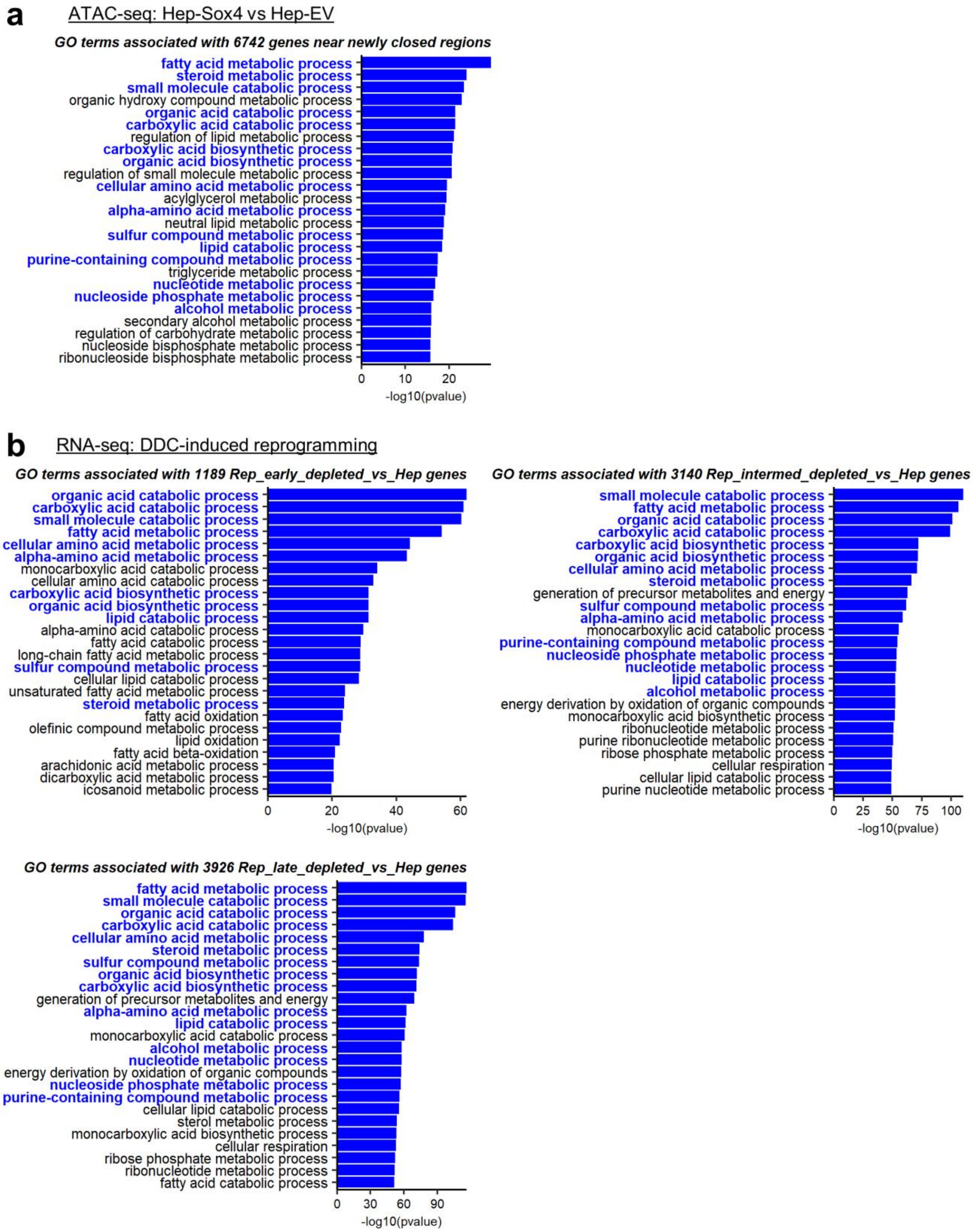

### Extended Data Fig. 5 (continued)

**c** ATAC-seq: Hep-Sox4 vs Hep-EV

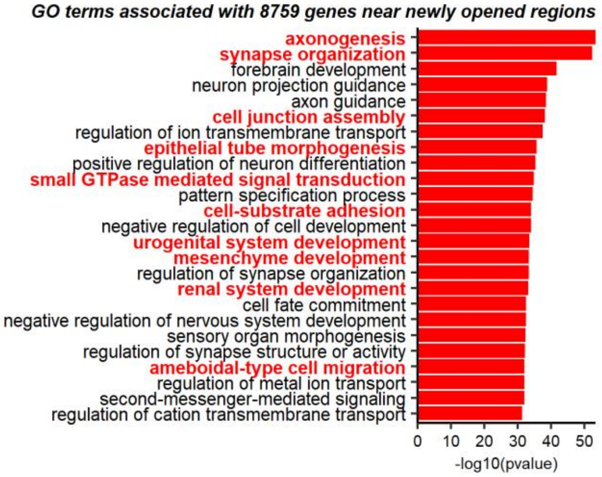


**d** RNA-seq: DDC-induced reprogramming

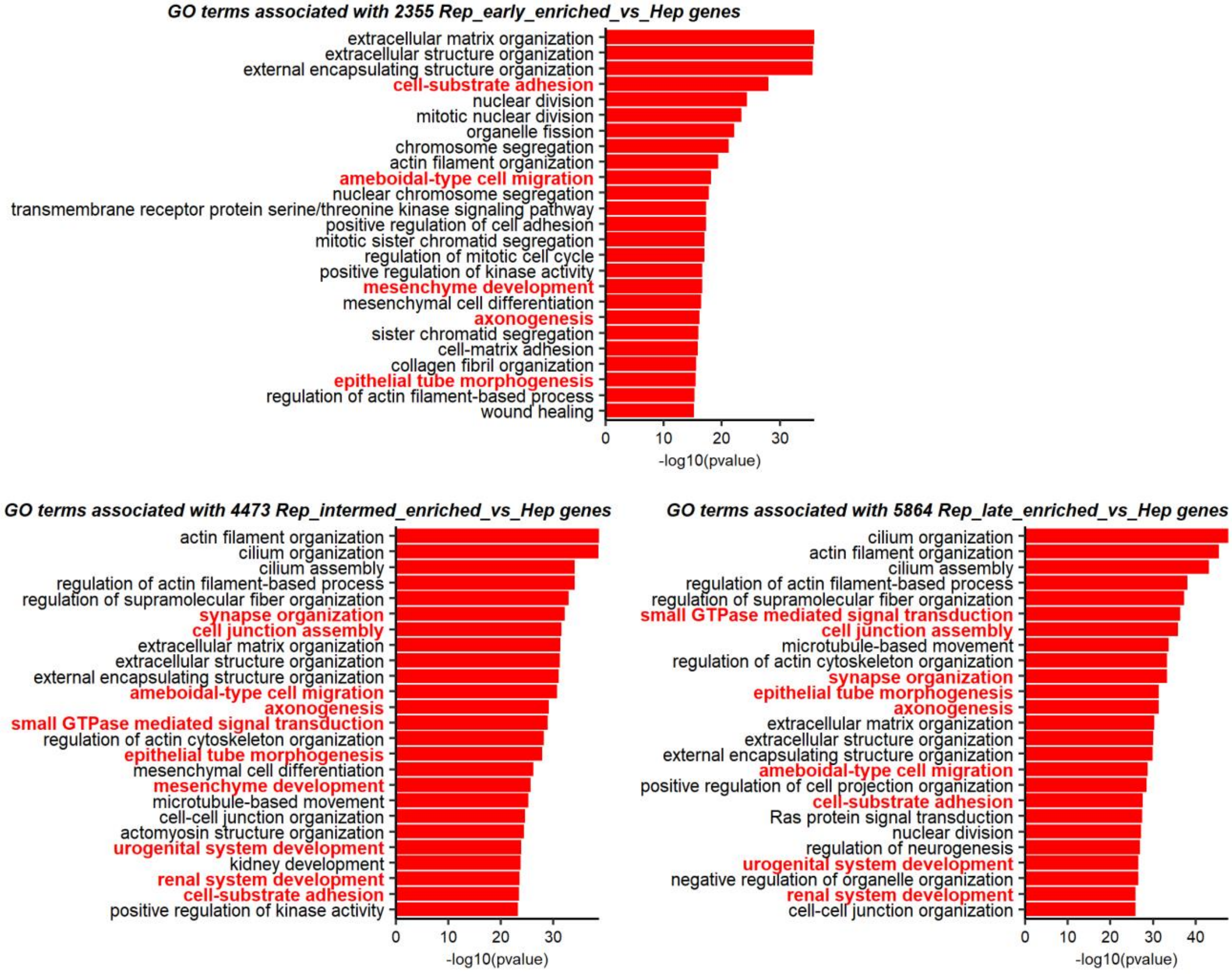

**Extended Data Fig. 5. Gene ontology (GO) analysis for genes associated with newly closed and opened regions following *Sox4* expression in hepatocytes.**

(a) Newly closed regions (n = 14,564, **Fig. 2a**) in *Sox4* expressing hepatocytes compared with empty vector hepatocytes were annotated with the nearest genes, and this gene list was used as the input for GO enrichment analysis. Top 25 GO terms are shown.

(b) GO analysis was performed using genes downregulated during DDC-induced reprogramming at different stages (RNA-Seq), namely Rep_early vs Hep (upper left), Rep_intermed vs Hep (upper right), and Rep_late vs Hep (lower). Top 25 GO terms are shown. GO terms shared between the newly-closed region-associated gene set and DDC-induced reprogramming context at any reprogramming stages are highlighted in bold blue texts.

(c) Newly-opened regions (n = 20,329, **Fig. 2a**) in *Sox4* expressing hepatocytes compared with empty vector hepatocytes were annotated with the nearest genes, and this gene list was used as the input for GO analysis. Top 25 GO terms are shown.

(d) GO analysis was performed using genes upregulated during DDC-induced reprogramming at different stages (RNA-Seq), namely Rep_early vs Hep (upper), Rep_intermed vs Hep (lower left), and Rep_late vs Hep (lower right). Top 25 GO terms are shown. GO terms shared between the newly-opened region-associated gene set and DDC-induced reprogramming context at any reprogramming stages are highlighted in bold red texts.

### Extended Data Fig. 6

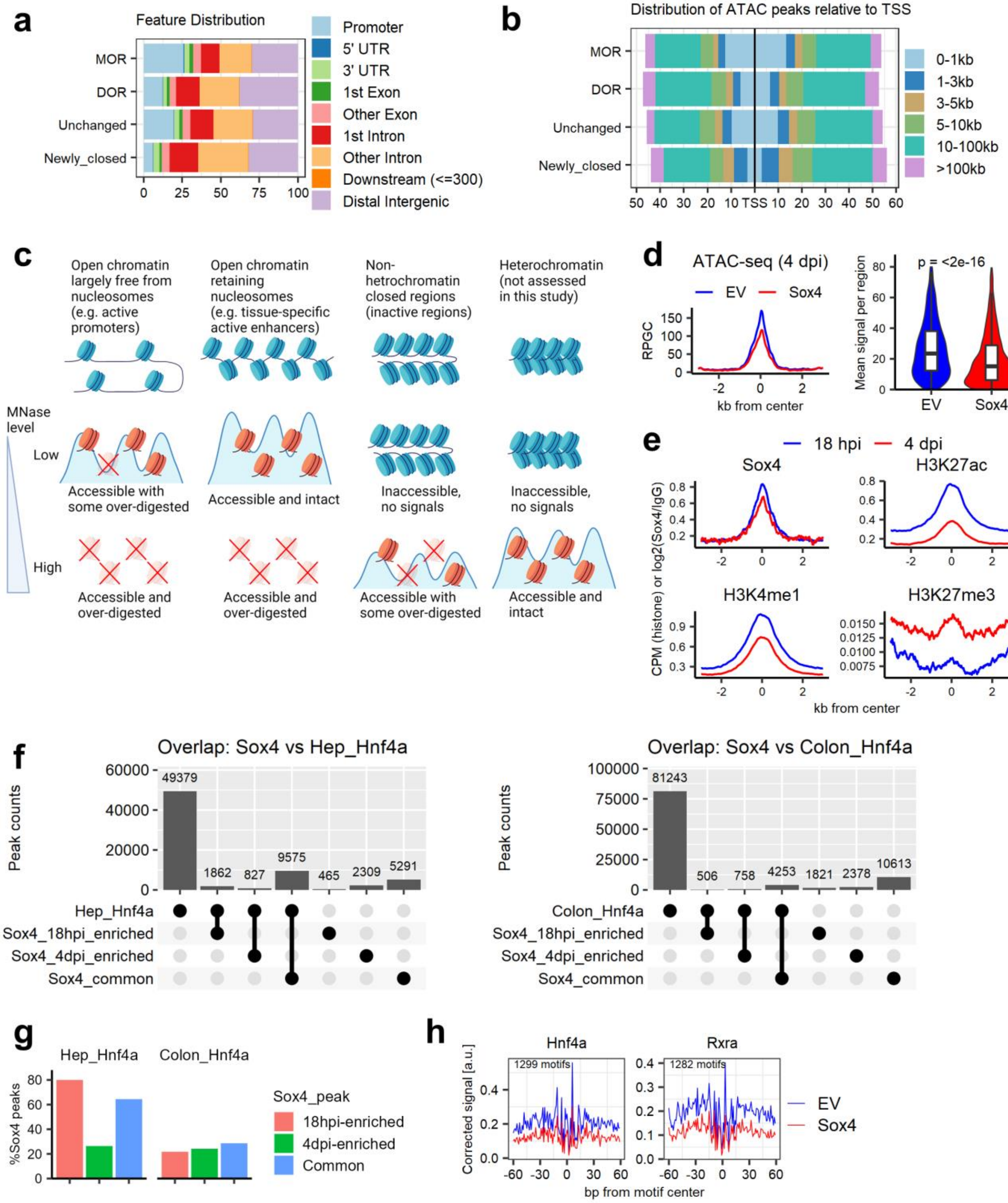

**Extended Data Fig. 6. Genomic distribution of Sox4-induced changes in chromatin accessibility and characterization of newly closed regions.**

(a) Feature distribution of ATAC-seq peaks at MOR, DOR, unchanged regions and newly closed regions. Promoter regions are defined as TSS ± 1 kb.

(b) Distribution of the distance of ATAC-seq peaks relative to nearest transcription start sites (TSSs).

(c) Nucleosomal states are roughly categorized into four groups, and how these states can be detected in terms of the output of MNase-low-seq and MNase-high-seq.

(d) Aggregate plots corresponding to the ATAC-seq data as shown in **Fig. 4f** on the left, and the comparison of the average signals in each region between empty vector and *Sox4* expressing hepatocytes are shown on the right. P-value is calculated by Wilcoxon rank sum test.

(e) Averaged aggregate plots corresponding to the CUT&RUN-seq data as shown in **Fig. 4f**.

(f) Peak overlaps are quantified between Sox4 peaks and Hnf4a peaks obtained from hepatocytes[2] or colon epithelial cells[4].

(g) Summary of quantification of peak overlap shown in **(f)**.

(h) ATAC-seq footpriting analysis of hepatocyte transcription factors Hnf4a and Rxra at motifs overlapping the active liver enhancers. The data are shown as the comparison between empty vector and Sox4 hepatocytes. Motifs assigned as “bound” by TOBIAS for the empty vector and *Sox4* expressing samples are combined and used for the analysis.

### Extended Data Fig. 7

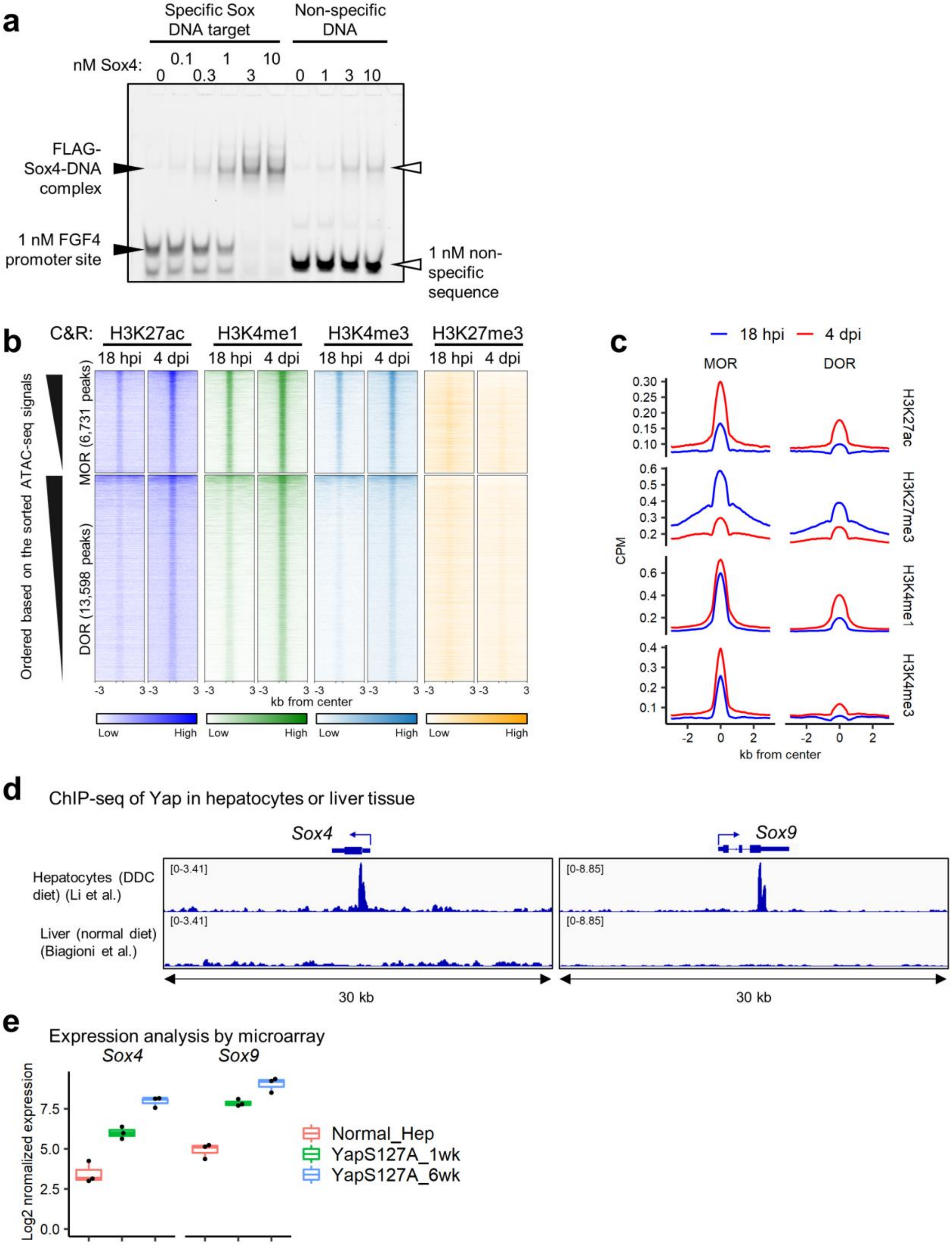

**Extended Data Fig. 7. Specificity of *in vitro* DNA binding activity of recombinant Sox4, and profiling histone post-translational modifications at the newly opened regions.**

(a) EMSA showing the affinity of increasing amounts of recombinant Sox4 protein to Cy5-labeled DNA fragment containing the Sox motif or non-specific sequences. The concentrations used for each protein (nM) are indicated above each lane. DNA sequences of the Cy5-labeled probes are shown in the table at the bottom.

(b) CUT&RUN signals for H3K27ac, H3K4me1, H3K4me3 and H3K27me3 visualized as heatmaps for MORs and DORs. The rows are reordered according to the signal intensities from ATAC-Seq.

(c) Averaged aggregate plots for **(c)**.

(d) Genome browser view of Yap ChIP-Seq at the *Sox4* and *Sox9* regions. The analysis was performed on hepatocytes isolated from DDC-treated animals (upper panels) or liver from animals fed a normal diet (lower panels). Data were downloaded from GEO[5,6].

(e) *Sox4* and *Sox9* expression was assessed by microarray following expression of a constitutively active human YAP (YAPS127A) in adult hepatocytes. Data were downloaded from GEO[7].

Extended Data Fig. 8

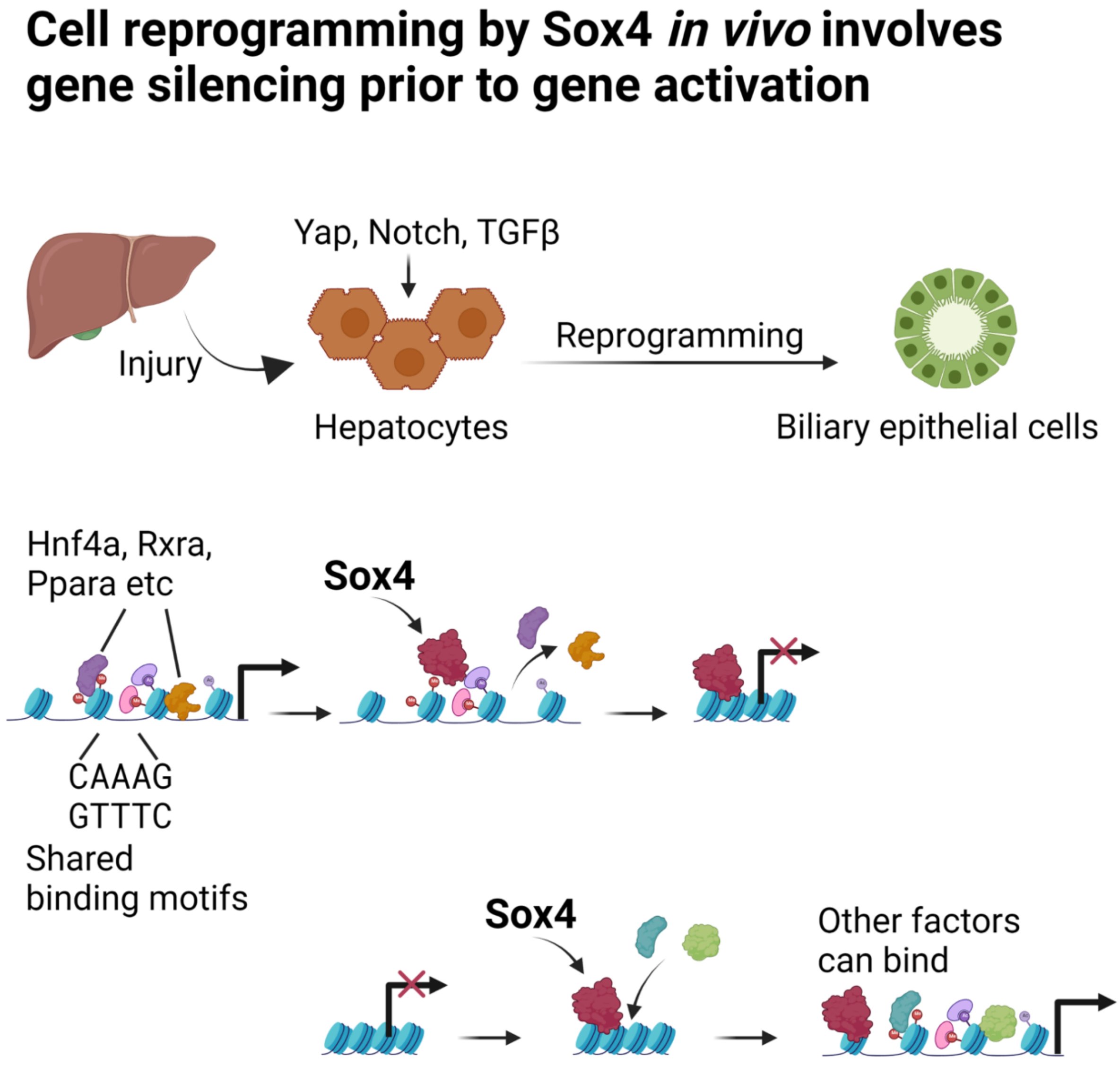

**Extended Data Fig. 8. Schematic for the model of the regulatory mechanism of hepatobiliary reprogramming mediated by Sox4.**
