## Supplementary_Figures for "Physiological reprogramming *in vivo* mediated by Sox4 pioneer factor activity"

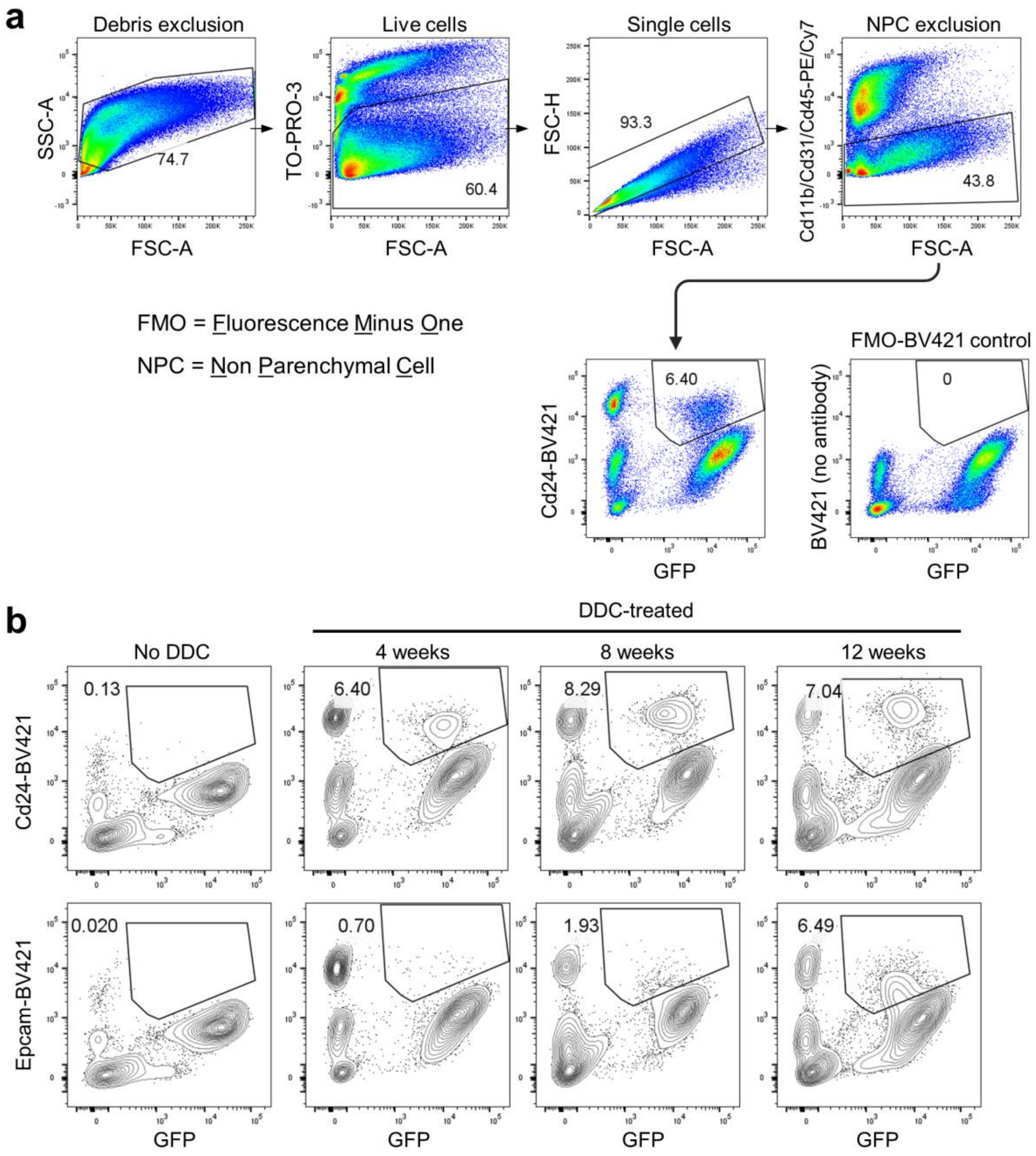


**Supplementary Fig. 1. Gating strategy for flow cytometry of DDC-induced reprogrammed cells and the representative results of each time point, related to Extended Data Fig. 1c.**

(a) Representative plots of gating strategy for flow cytometry of whole liver cells challenged with DDC for the designated time points (**Extended Data Fig. 1c**). Cells harvested from normal livers (No DDC) serve as the week 0 control.

(b) Representative plots indicating the reprogrammed cell fractions at the designated time points. The rectangle gates indicate the GFP/BV421 double positive cells with percentage of the total parental population. The percentages shown in **Extended Data Fig. 1c** are calculated as (GFP+/BV421+)/(GFP+) ratios.

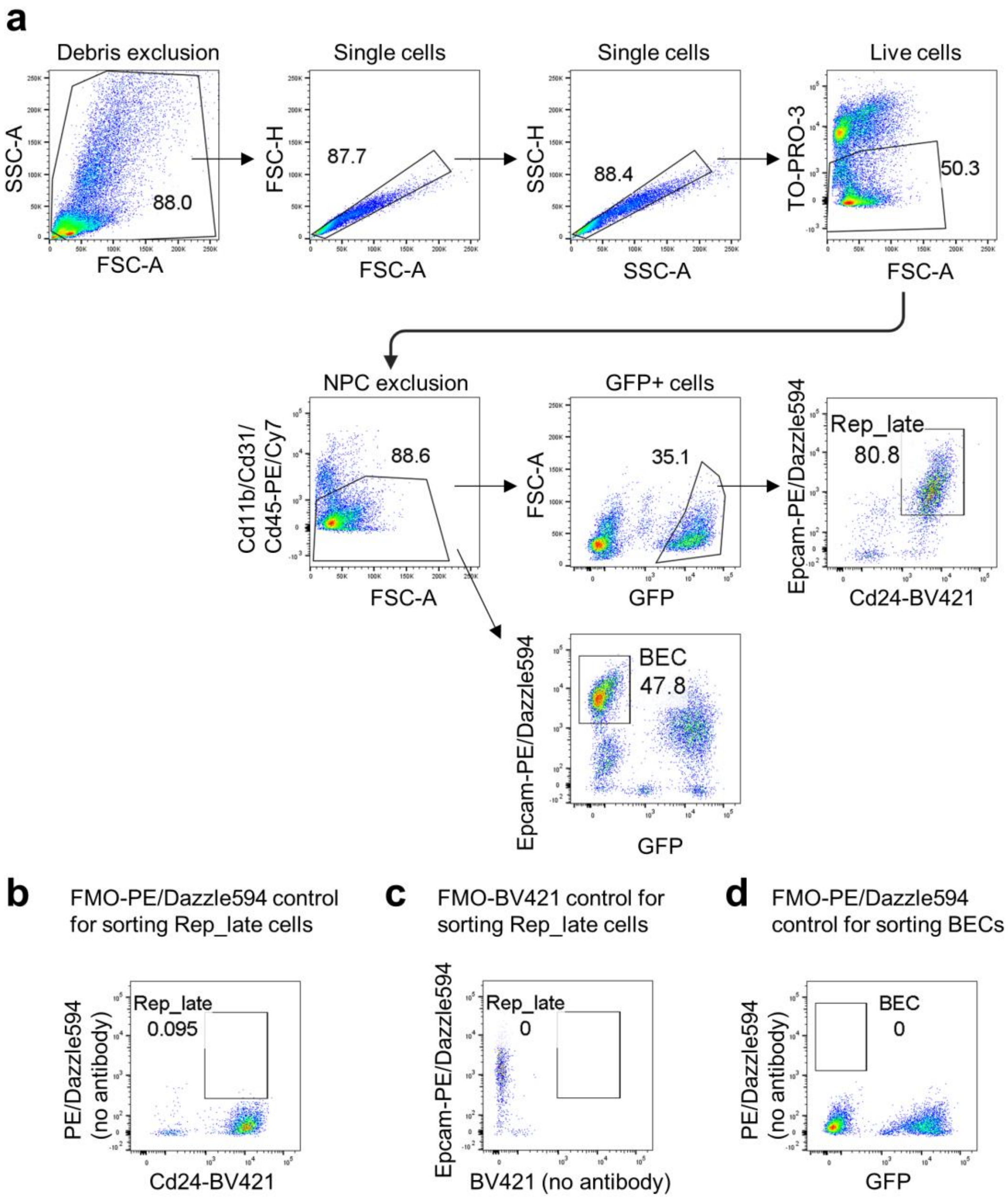


**Supplementary Fig. 2. Gating strategy for FACS sorting of Rep_late and biliary epithelial cells, related to Extended Data Fig. 1e, f.**

(a) Representative plots of gating strategy for FACS sorting of Rep_late and biliary epithelial cells using Epcam+ fraction which was enriched by MACS (**Extended Data Fig. 1e**).

(b) Representative plots of PE/Dazzle594-fluorescence minus one (FMO) control, which was used to determine the threshold of Epcam expression of Rep_late cells.

(c) Representative plots of BV421-FMO control, which was used to determine the threshold of Cd24 expression of Rep_late cells.

(d) Representative plots of PE/Dazzle594-FMO control, which was used to determine the threshold of Cd24 expression of biliary epithelial cells.

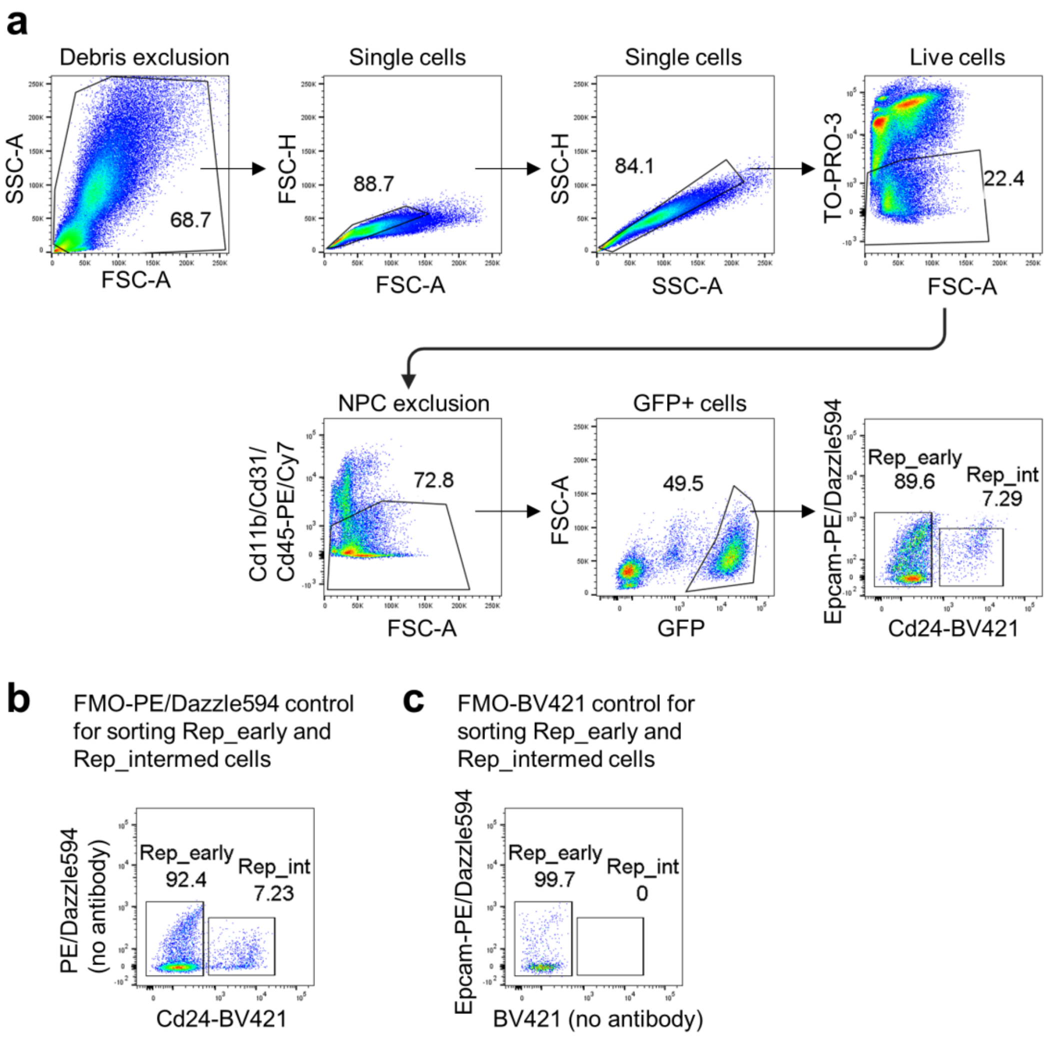


**Supplementary Fig. 3. Gating strategy for FACS sorting of Rep_early and Rep_intermed cells, related to Extended Data Fig. 1e, f.**

(a) Representative plots of gating strategy for FACS sorting of Rep_early and Rep_intermed cells using Epcam- fraction which was enriched by MACS (**Extended Data Fig. 1e**).

(b) Representative plots of PE/Dazzle594-FMO control, which was used to determine the threshold of Epcam expression of Rep_early and Rep_intermed cells.

(c) Representative plots of BV421-FMO control, which was used to determine the threshold of Cd24 expression of Rep_early and Rep_intermed cells.

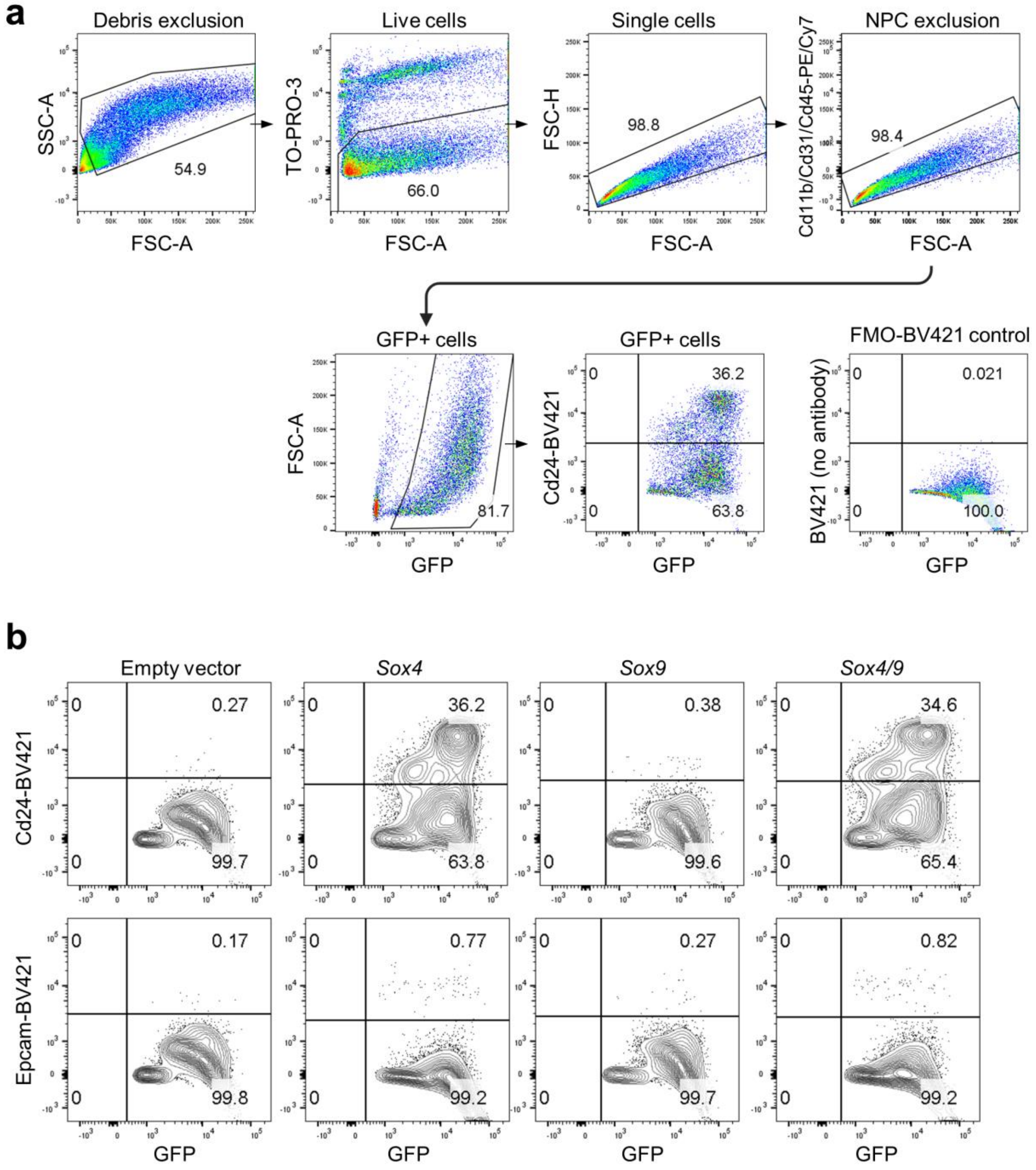


**Supplementary Fig. 4. Gating strategy for determination of reprogramming efficiency of hepatocytes expressing *Sox4*, *Sox9* or both, related to Fig. 1b, Extended Data Fig. 3g.**

(a) Representative plots of gating strategy for flow cytometry of whole liver cells expressing Sox4, Sox9 or both at 7 dpi (experimental design described in **Fig. 1a**). Empty vector-injected cells serve as a control.

(b) Representative plots indicating the reprogrammed cell fractions in the early (Cd24) and late (Epcam) stages. The percentages shown in the right top quadrant indicates GFP/BV421 double positive ratios based on the total GFP positive cells.

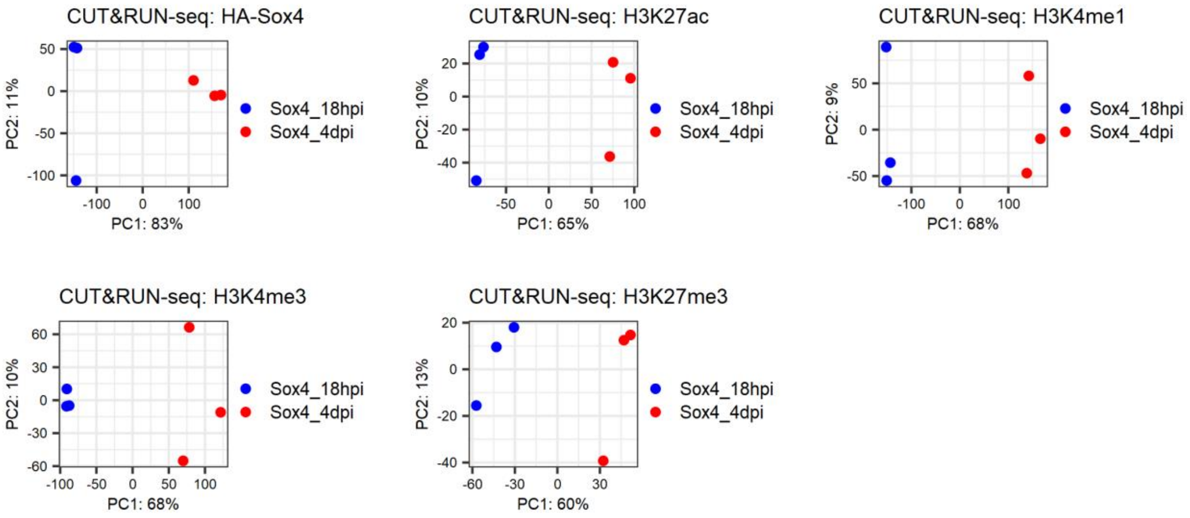


**Supplementary Figure. 5. Validation of consistency in CUT&RUN-seq data among three replicates in terms of PCA mapping.**

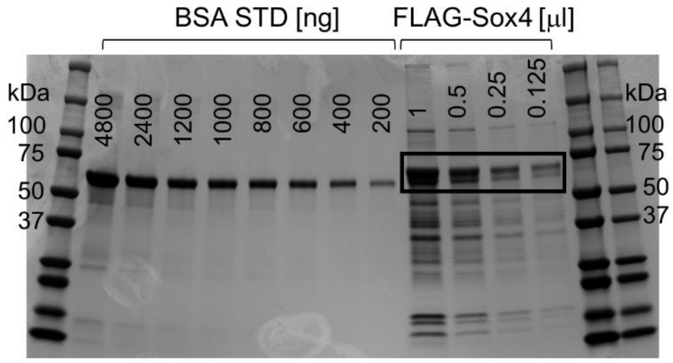


**Supplementary Figure 6. Quantification of purified FLAG-Sox4 by SDS-PAGE using a BSA standard. Rectangle region corresponds to the typical Sox4 size, and used for quantification.**
