## Supplementary material for "Physiological reprogramming *in vivo* mediated by Sox4 pioneer factor activity": Table S1

**Table S1. Differentially expressed genes between hepatocytes and early reprogrammed cells.**

**Upregulated genes in Rep_early v.s. Hep**

| Gene_id | baseMean | log2FoldChange | lfcSE | pvalue | padj |
|---|---|---|---|---|---|
| Mfge8 | 6902.877868 | 6.065226137 | 0.136211178 | 0 | 0 |
| Tagln2 | 5569.887589 | 5.300897757 | 0.150242574 | 3.27E-273 | 2.09E-269 |
| Nid1 | 3343.827712 | 5.7155129 | 0.174352408 | 8.72E-237 | 4.19E-233 |
| Anxa2 | 5627.70021 | 7.162708988 | 0.237551702 | 6.37E-200 | 2.44E-196 |
| Anxa5 | 16437.78852 | 2.385633882 | 0.080287084 | 3.16E-194 | 1.01E-190 |
| Serpinb6a | 1972.53765 | 5.36032416 | 0.189417256 | 1.25E-176 | 3.43E-173 |
| Flna | 12926.99246 | 4.106605221 | 0.146557837 | 3.31E-173 | 7.95E-170 |
| Cd63 | 2860.789729 | 6.72470182 | 0.24159987 | 3.79E-171 | 8.08E-168 |
| Ahnak | 4276.591661 | 4.789840727 | 0.178770851 | 3.95E-159 | 7.58E-156 |
| Spp1 | 104686.7999 | 7.040125069 | 0.263957036 | 5.64E-158 | 9.83E-155 |
| Osbpl3 | 3492.147106 | 6.595340804 | 0.254853038 | 8.76E-148 | 1.40E-144 |
| Dmpk | 2014.690643 | 5.961753319 | 0.241753759 | 1.91E-135 | 2.82E-132 |
| Pygb | 2106.44206 | 4.071633822 | 0.168784115 | 1.35E-129 | 1.85E-126 |
| Cystm1 | 913.1600743 | 3.198601854 | 0.134338378 | 3.73E-126 | 4.77E-123 |
| Emp2 | 1701.891287 | 3.553394165 | 0.15337867 | 1.57E-119 | 1.89E-116 |
| Fads3 | 1438.431683 | 2.858082654 | 0.124984406 | 3.19E-116 | 3.60E-113 |
| Bicc1 | 7310.942721 | 5.707049078 | 0.250390487 | 5.86E-115 | 6.25E-112 |
| S100a11 | 2499.42336 | 6.44429242 | 0.285647586 | 1.32E-112 | 1.27E-109 |
| Cstb | 2245.495281 | 2.695056312 | 0.120281901 | 3.81E-111 | 3.48E-108 |
| Ubd | 999.3194965 | 7.071871326 | 0.327565344 | 2.35E-103 | 1.96E-100 |
| Lama5 | 7209.224019 | 6.125436108 | 0.285874083 | 2.03E-102 | 1.62E-99 |
| Ddr1 | 4937.617359 | 5.842408139 | 0.278226491 | 6.95E-98 | 5.13E-95 |
| Ly6d | 1741.198405 | 6.245572843 | 0.302363885 | 8.25E-96 | 5.86E-93 |
| Abcc4 | 1273.192084 | 6.039638554 | 0.303929939 | 1.82E-88 | 1.13E-85 |
| Npdc1 | 1128.275668 | 4.460223271 | 0.225142566 | 3.37E-88 | 2.02E-85 |
| Iqgap1 | 3693.490198 | 3.518485929 | 0.17764993 | 3.95E-88 | 2.30E-85 |
| Prelp | 1636.616549 | 5.253405862 | 0.264935555 | 4.83E-88 | 2.72E-85 |
| Cybb | 849.2809416 | 6.131587285 | 0.311438476 | 4.42E-87 | 2.43E-84 |
| Krt18 | 21172.22313 | 2.281766701 | 0.116067439 | 6.20E-87 | 3.30E-84 |
| Dusp4 | 769.6146092 | 3.905223968 | 0.200453407 | 5.03E-85 | 2.61E-82 |
| Entpd2 | 539.9128162 | 3.064079829 | 0.158924728 | 1.43E-83 | 7.02E-81 |
| Unc13b | 1376.866657 | 3.346520376 | 0.174792871 | 1.19E-82 | 5.71E-80 |
| Pam | 1955.879668 | 4.265502789 | 0.222306528 | 2.77E-82 | 1.30E-79 |
| Steap2 | 1723.323568 | 3.255639626 | 0.172812814 | 4.13E-80 | 1.89E-77 |
| Golm1 | 1057.67312 | 3.412252829 | 0.18159298 | 1.40E-79 | 6.10E-77 |
| Rragd | 3357.50487 | 8.533547168 | 0.449484709 | 3.86E-79 | 1.64E-76 |
| Lamc1 | 1937.260399 | 2.405785597 | 0.130429419 | 7.81E-77 | 3.26E-74 |
| Smpd3 | 1931.635961 | 3.219283797 | 0.178156907 | 5.54E-74 | 2.21E-71 |
| Mtmr11 | 638.9640192 | 2.711456442 | 0.149776292 | 5.75E-74 | 2.25E-71 |

| | | | | | |
|---|---|---|---|---|---|
| Tpm1 | 3669.779031 | 1.969787971 | 0.109380433 | 1.95E-73 | 7.49E-71 |
| Pak6 | 1667.587799 | 6.796154113 | 0.376401661 | 5.30E-72 | 1.99E-69 |
| Pitpnm1 | 915.804418 | 3.49999021 | 0.199764463 | 1.59E-69 | 5.75E-67 |
| Sh3pxd2b | 1590.483526 | 4.097549805 | 0.235924959 | 3.41E-68 | 1.21E-65 |
| Trim47 | 1675.333833 | 4.766001389 | 0.275350312 | 9.91E-68 | 3.40E-65 |
| Rhbdf1 | 1328.09275 | 2.415335991 | 0.139989623 | 1.50E-67 | 4.98E-65 |
| Krt8 | 33827.50579 | 2.421114881 | 0.140276161 | 1.85E-67 | 6.00E-65 |
| Ets2 | 3247.801541 | 2.476794935 | 0.144702216 | 1.02E-66 | 3.28E-64 |
| Epb41l1 | 1148.814833 | 6.10598282 | 0.354674979 | 5.77E-66 | 1.82E-63 |
| Itga6 | 830.2633331 | 2.929508393 | 0.172405414 | 2.04E-65 | 6.31E-63 |
| Vcam1 | 4509.077815 | 4.452857942 | 0.263563753 | 1.17E-64 | 3.50E-62 |
| Gsta1 | 510.5820908 | 4.384058327 | 0.263775244 | 3.56E-63 | 9.76E-61 |
| Ifi27l2b | 869.8349867 | 2.833053728 | 0.172834823 | 1.66E-61 | 4.09E-59 |
| Rras | 1192.43813 | 1.956790621 | 0.119527855 | 5.13E-61 | 1.25E-58 |
| Slc43a2 | 4679.899199 | 2.927934201 | 0.179091129 | 8.12E-61 | 1.95E-58 |
| Myl12a | 2480.199164 | 1.582518612 | 0.098195289 | 4.35E-59 | 1.02E-56 |
| Tgfbr2 | 1823.698814 | 1.729527716 | 0.107950139 | 1.73E-58 | 4.00E-56 |
| Itpr3 | 1859.686378 | 6.32988895 | 0.391653212 | 3.79E-58 | 8.66E-56 |
| Agrn | 18322.92867 | 1.354391997 | 0.084964472 | 9.15E-58 | 2.06E-55 |
| Cav2 | 808.5762741 | 3.934850365 | 0.247073595 | 9.25E-58 | 2.06E-55 |
| Cd9 | 3123.118331 | 3.227552883 | 0.203971503 | 1.90E-57 | 4.14E-55 |
| Arrb1 | 1004.049177 | 4.435945078 | 0.278474054 | 1.88E-57 | 4.14E-55 |
| Sprr1a | 552.0048189 | 7.873098531 | 0.49176401 | 4.70E-57 | 1.01E-54 |
| Myo1c | 2795.299184 | 1.643693798 | 0.10421127 | 7.70E-57 | 1.64E-54 |
| Abcc5 | 1363.308835 | 4.307275423 | 0.272732737 | 9.72E-57 | 2.05E-54 |
| Pkd2 | 2618.315516 | 1.299077417 | 0.082403876 | 2.31E-56 | 4.71E-54 |
| Tm4sf4 | 12492.0944 | 2.254533345 | 0.143880083 | 3.10E-56 | 6.26E-54 |
| Uap1l1 | 1267.481248 | 3.230985037 | 0.206471359 | 4.39E-56 | 8.78E-54 |
| F11r | 6594.67141 | 1.065004901 | 0.06785749 | 5.20E-56 | 1.02E-53 |
| Slpi | 513.3845618 | 7.841733221 | 0.495750681 | 8.04E-56 | 1.56E-53 |
| Ccdc120 | 701.478924 | 2.7353855 | 0.175348676 | 1.02E-55 | 1.96E-53 |
| Chmp4c | 1792.032984 | 2.57072206 | 0.165199459 | 1.91E-55 | 3.60E-53 |
| Specc1 | 764.9099083 | 2.560918827 | 0.164213717 | 2.02E-55 | 3.76E-53 |
| Gm3776 | 443.2839314 | 4.882928116 | 0.316917903 | 9.28E-55 | 1.71E-52 |
| Plet1 | 2059.923758 | 4.378116434 | 0.284634133 | 1.71E-54 | 3.09E-52 |
| Cbr3 | 476.737001 | 6.886760472 | 0.444770273 | 2.34E-54 | 4.20E-52 |
| Myo7a | 1894.705031 | 2.466126278 | 0.160222232 | 2.47E-54 | 4.38E-52 |
| Ezr | 2894.776709 | 2.87829062 | 0.188868377 | 2.30E-53 | 4.06E-51 |
| Cdc42se1 | 1443.370978 | 1.835668448 | 0.12036433 | 2.98E-53 | 5.20E-51 |
| Lgals1 | 1864.264472 | 3.175279343 | 0.210624093 | 1.82E-52 | 3.11E-50 |
| Pakap | 831.1717092 | 3.800837076 | 0.251035397 | 1.99E-52 | 3.38E-50 |
| Slc25a24 | 2130.409856 | 8.482650468 | 0.549972335 | 2.85E-52 | 4.76E-50 |
| Fmo2 | 1337.029237 | 4.824042454 | 0.322467263 | 1.36E-51 | 2.21E-49 |

| | | | | | |
|---|---|---|---|---|---|
| Tmsb10 | 3903.761089 | 6.517295322 | 0.440652378 | 1.17E-49 | 1.87E-47 |
| Tes | 1160.343998 | 3.126335362 | 0.212722772 | 1.19E-49 | 1.89E-47 |
| Rhoc | 1087.529876 | 3.559151763 | 0.243687322 | 2.41E-49 | 3.77E-47 |
| Rgs5 | 21587.48284 | 4.769915975 | 0.325203791 | 2.63E-49 | 4.06E-47 |
| Clcf1 | 3063.556035 | 5.499211505 | 0.375927555 | 4.71E-49 | 7.17E-47 |
| Rab3d | 893.8767309 | 1.757507478 | 0.120002228 | 5.80E-49 | 8.77E-47 |
| Nck2 | 675.8478451 | 3.586077168 | 0.247280689 | 4.34E-48 | 6.45E-46 |
| Abca5 | 846.7686621 | 1.859706203 | 0.129034818 | 7.14E-48 | 1.05E-45 |
| Plxna3 | 616.2926403 | 4.664045333 | 0.322244045 | 1.68E-47 | 2.46E-45 |
| Mapk3 | 2336.977053 | 1.35331405 | 0.094069505 | 1.81E-47 | 2.64E-45 |
| Alpk1 | 590.6005021 | 3.490807743 | 0.244233093 | 6.67E-47 | 9.47E-45 |
| Fkbp1a | 2034.011696 | 1.141112239 | 0.079713842 | 6.77E-47 | 9.56E-45 |
| Gipc2 | 518.4771754 | 7.929229395 | 0.545405083 | 1.22E-46 | 1.70E-44 |
| Rtn4 | 8745.166298 | 1.663808646 | 0.117022187 | 1.50E-46 | 2.08E-44 |
| Id1 | 742.5083659 | 4.768532588 | 0.337546186 | 1.85E-46 | 2.54E-44 |
| Klc1 | 939.6043412 | 1.453668533 | 0.102702742 | 4.93E-46 | 6.61E-44 |
| Tnfrsf12a | 1602.336135 | 3.313808985 | 0.235692302 | 5.60E-46 | 7.46E-44 |
| Unc5b | 1262.892053 | 4.190946346 | 0.297046843 | 6.17E-46 | 8.11E-44 |
| Vim | 1716.491899 | 4.364972921 | 0.308692278 | 7.14E-46 | 9.32E-44 |
| Haus8 | 512.4162491 | 3.189711041 | 0.226988403 | 7.50E-46 | 9.73E-44 |
| Tax1bp3 | 722.2085877 | 2.283144067 | 0.162700463 | 1.69E-45 | 2.18E-43 |
| Ano6 | 1739.712585 | 2.174876743 | 0.154955357 | 2.23E-45 | 2.85E-43 |
| Isyna1 | 933.0269572 | 2.20292636 | 0.157253905 | 2.55E-45 | 3.24E-43 |
| Lgals3 | 560.0719328 | 8.660884979 | 0.60497043 | 2.73E-45 | 3.44E-43 |
| Rcan2 | 244.3838137 | 3.558365153 | 0.255039958 | 3.93E-45 | 4.90E-43 |
| Maoa | 483.5631152 | 2.099010841 | 0.150783447 | 7.58E-45 | 9.21E-43 |
| Spats2l | 390.0388806 | 4.905438452 | 0.351016692 | 2.74E-44 | 3.23E-42 |
| Bmp6 | 1444.196603 | 7.760724324 | 0.553839089 | 4.82E-44 | 5.60E-42 |
| Arhgap27 | 584.3936866 | 4.89693387 | 0.353199544 | 5.20E-44 | 5.98E-42 |
| Card10 | 3561.730514 | 2.165259751 | 0.157795172 | 1.03E-43 | 1.17E-41 |
| Lpl | 1437.187349 | 3.226958871 | 0.23610241 | 1.57E-43 | 1.76E-41 |
| Vill | 1270.715557 | 5.545270807 | 0.401338377 | 3.20E-43 | 3.57E-41 |
| App | 11500.26785 | 1.941963033 | 0.143049308 | 7.47E-43 | 8.14E-41 |
| Csf1 | 2009.083438 | 3.34209467 | 0.247487286 | 1.23E-42 | 1.34E-40 |
| Flt4 | 577.5153147 | 3.576803115 | 0.265414499 | 2.61E-42 | 2.77E-40 |
| Tmem176b | 12340.78038 | 1.430152547 | 0.105844708 | 3.29E-42 | 3.45E-40 |
| Pdlim7 | 626.095726 | 2.313363721 | 0.17133085 | 3.35E-42 | 3.50E-40 |
| Crtap | 533.1823875 | 2.623322816 | 0.195054637 | 5.80E-42 | 5.95E-40 |
| Arpc1b | 2020.229584 | 2.074927896 | 0.154667044 | 6.65E-42 | 6.75E-40 |
| Nfkbiz | 2187.735202 | 2.593001157 | 0.195162358 | 2.24E-41 | 2.25E-39 |
| Alpl | 901.7863714 | 1.65785534 | 0.125012645 | 8.90E-41 | 8.89E-39 |
| Rere | 2387.41737 | 1.181615955 | 0.088935879 | 1.05E-40 | 1.04E-38 |
| Slc25a4 | 1174.065257 | 5.018113948 | 0.380332792 | 1.79E-40 | 1.73E-38 |

| | | | | | |
|---|---|---|---|---|---|
| Mvp | 3555.029373 | 2.550804481 | 0.194824204 | 3.79E-40 | 3.65E-38 |
| Igsf8 | 870.8610881 | 2.366784767 | 0.180538664 | 6.26E-40 | 6.01E-38 |
| Arhgap44 | 513.1554464 | 2.774073505 | 0.211867739 | 6.99E-40 | 6.64E-38 |
| Sptan1 | 8904.858141 | 1.067579888 | 0.081280901 | 7.70E-40 | 7.28E-38 |
| Tmem176a | 7915.062951 | 1.281894443 | 0.098207905 | 1.58E-39 | 1.47E-37 |
| Synpo | 2028.95591 | 2.415900575 | 0.186481128 | 2.57E-39 | 2.37E-37 |
| Miga1 | 644.6403984 | 2.776972734 | 0.214605272 | 5.58E-39 | 5.12E-37 |
| Rab34 | 937.8339107 | 2.199872603 | 0.170866369 | 1.40E-38 | 1.28E-36 |
| Src | 1239.87662 | 2.261548031 | 0.176239468 | 1.81E-38 | 1.64E-36 |
| Tcaf1 | 653.8295039 | 2.628778477 | 0.205245143 | 3.26E-38 | 2.94E-36 |
| Ankrd10 | 1032.873478 | 1.838440966 | 0.143854252 | 4.03E-38 | 3.60E-36 |
| Serpine1 | 310.4900576 | 5.338339815 | 0.419646556 | 6.42E-38 | 5.67E-36 |
| Pak1 | 808.1342862 | 2.848483587 | 0.224014514 | 9.20E-38 | 8.03E-36 |
| Gls | 777.5880481 | 2.868965432 | 0.226553971 | 1.13E-37 | 9.75E-36 |
| Stmn1 | 794.0286794 | 2.986853977 | 0.23596896 | 1.13E-37 | 9.75E-36 |
| Plin4 | 464.4996555 | 4.865574617 | 0.385960264 | 1.41E-37 | 1.21E-35 |
| Anxa4 | 5265.124635 | 1.064128906 | 0.083836898 | 2.61E-37 | 2.21E-35 |
| Rbbp8 | 559.8969069 | 2.177598556 | 0.172964754 | 3.66E-37 | 3.07E-35 |
| Abcb1a | 580.5758382 | 3.337000092 | 0.267716311 | 1.00E-36 | 8.31E-35 |
| Scn1b | 411.1688752 | 3.07038867 | 0.247620674 | 2.93E-36 | 2.33E-34 |
| Sh3bp4 | 768.7533521 | 2.627110222 | 0.211164951 | 3.78E-36 | 3.00E-34 |
| H2-Q1 | 166.7637522 | 4.036511237 | 0.326807556 | 3.93E-36 | 3.10E-34 |
| Adamts7 | 987.7749654 | 2.387136401 | 0.193789322 | 8.29E-36 | 6.41E-34 |
| Epb41l2 | 992.8211022 | 2.727838102 | 0.220305086 | 1.06E-35 | 8.17E-34 |
| Nedd9 | 1243.650118 | 2.354471723 | 0.191081015 | 1.10E-35 | 8.41E-34 |
| Adam11 | 957.3567386 | 2.472978795 | 0.203073 | 5.08E-35 | 3.84E-33 |
| Col4a1 | 3848.690724 | 2.598377989 | 0.215274437 | 1.53E-34 | 1.14E-32 |
| Aldoa | 4625.542072 | 1.096374005 | 0.090270359 | 2.25E-34 | 1.67E-32 |
| Klhdc8b | 721.2995738 | 1.968671887 | 0.162985528 | 2.27E-34 | 1.67E-32 |
| Map4k4 | 2315.772347 | 2.704978691 | 0.224434 | 2.42E-34 | 1.78E-32 |
| Lrtm2 | 254.542963 | 6.209269919 | 0.513525314 | 2.57E-34 | 1.88E-32 |
| Ppl | 4769.684527 | 3.499315563 | 0.292544153 | 4.25E-34 | 3.09E-32 |
| Cavin2 | 553.8028122 | 2.897721863 | 0.240800824 | 5.17E-34 | 3.73E-32 |
| Pea15a | 1016.954419 | 2.088163857 | 0.174333388 | 7.19E-34 | 5.17E-32 |
| Lysmd2 | 189.4115612 | 3.138069751 | 0.264448395 | 2.50E-33 | 1.76E-31 |
| Fhod1 | 756.0451718 | 2.694196099 | 0.227648404 | 5.00E-33 | 3.49E-31 |
| Maged2 | 781.4167773 | 1.73423385 | 0.146505414 | 5.81E-33 | 4.03E-31 |
| Phldb1 | 1087.579966 | 2.600419711 | 0.219823103 | 7.31E-33 | 5.02E-31 |
| 1700017B05Rik | 1170.847373 | 1.349801579 | 0.114172266 | 8.00E-33 | 5.48E-31 |
| Ptgr1 | 622.257811 | 2.007667132 | 0.170338596 | 8.32E-33 | 5.68E-31 |
| Tmem200b | 172.8857394 | 3.165547408 | 0.270147316 | 1.47E-32 | 9.97E-31 |
| St14 | 1708.65254 | 6.203218893 | 0.521202728 | 1.70E-32 | 1.15E-30 |
| Btg3 | 340.3712981 | 2.110539816 | 0.180646174 | 2.47E-32 | 1.66E-30 |

| | | | | | |
|---|---|---|---|---|---|
| B4galt5 | 1641.815408 | 1.697711819 | 0.145164392 | 2.69E-32 | 1.80E-30 |
| Pcolce | 380.9320447 | 2.339844832 | 0.200186498 | 2.87E-32 | 1.90E-30 |
| Amdhd2 | 922.0458705 | 1.46192043 | 0.125340278 | 4.86E-32 | 3.21E-30 |
| Mcm6 | 958.1815709 | 2.726722362 | 0.236289665 | 9.40E-32 | 6.14E-30 |
| Tbc1d24 | 1001.858609 | 1.226433491 | 0.105501113 | 9.93E-32 | 6.46E-30 |
| Ankrd13b | 647.7782181 | 1.936814313 | 0.167185761 | 1.05E-31 | 6.79E-30 |
| Arhgef16 | 1015.504197 | 1.183540664 | 0.101941558 | 1.46E-31 | 9.39E-30 |
| Them6 | 404.6603918 | 1.872701574 | 0.162873261 | 2.35E-31 | 1.50E-29 |
| Nrg1 | 434.418342 | 3.760777178 | 0.330709076 | 5.14E-31 | 3.23E-29 |
| Sirpa | 1056.623764 | 2.014559228 | 0.176645922 | 8.94E-31 | 5.54E-29 |
| Klf6 | 3236.996363 | 2.931819051 | 0.259353832 | 1.11E-30 | 6.87E-29 |
| Osbpl7 | 397.1441368 | 3.242965085 | 0.284589123 | 1.19E-30 | 7.33E-29 |
| Galns | 510.338008 | 1.87579659 | 0.165419099 | 1.41E-30 | 8.59E-29 |
| Nagk | 612.1433348 | 1.863566791 | 0.164630223 | 1.84E-30 | 1.11E-28 |
| Gltp | 363.8767919 | 3.31169326 | 0.292703977 | 2.46E-30 | 1.47E-28 |
| Smoc2 | 255.4244332 | 2.492784852 | 0.221829867 | 3.26E-30 | 1.94E-28 |
| Cavin1 | 1055.494728 | 2.877739709 | 0.254545549 | 3.33E-30 | 1.97E-28 |
| Cxcl16 | 1465.568382 | 3.230345683 | 0.287174911 | 3.45E-30 | 2.03E-28 |
| Cul7 | 1027.888616 | 1.441612402 | 0.127745022 | 4.44E-30 | 2.59E-28 |
| Wnk4 | 352.5355701 | 2.753561244 | 0.245218663 | 4.63E-30 | 2.69E-28 |
| Chrm3 | 305.1106829 | 2.9719887 | 0.264553574 | 5.37E-30 | 3.10E-28 |
| Cul9 | 988.5653489 | 1.50595628 | 0.13407224 | 6.55E-30 | 3.76E-28 |
| Lamb2 | 2218.309935 | 5.284396506 | 0.463143935 | 6.88E-30 | 3.93E-28 |
| Espn | 100.1849283 | 4.141020633 | 0.367439205 | 7.75E-30 | 4.42E-28 |
| Ankrd27 | 1384.88522 | 1.044874906 | 0.092851758 | 9.43E-30 | 5.34E-28 |
| Sox4 | 6011.501005 | 7.541602947 | 0.656659124 | 1.35E-29 | 7.59E-28 |
| Hook2 | 1043.872935 | 1.45946583 | 0.130809238 | 1.63E-29 | 9.12E-28 |
| St3gal2 | 772.6672223 | 3.483148869 | 0.311650773 | 1.75E-29 | 9.79E-28 |
| Rab31 | 697.802115 | 4.538106139 | 0.407706248 | 2.61E-29 | 1.45E-27 |
| Mthfd1l | 302.4856827 | 2.477359287 | 0.22401344 | 3.04E-29 | 1.68E-27 |
| Slc20a1 | 1066.056153 | 2.218991498 | 0.201089422 | 3.52E-29 | 1.93E-27 |
| Rap1gap | 1801.710336 | 1.693510931 | 0.15296843 | 3.55E-29 | 1.94E-27 |
| Plxnd1 | 1309.856131 | 2.73995077 | 0.248387086 | 3.88E-29 | 2.12E-27 |
| Olfm2 | 234.8928339 | 2.999580844 | 0.272766885 | 4.66E-29 | 2.53E-27 |
| Ppp1r9b | 1239.144456 | 3.066432383 | 0.277055515 | 4.85E-29 | 2.63E-27 |
| Bcam | 2672.586742 | 3.647534805 | 0.328783173 | 4.97E-29 | 2.69E-27 |
| Large1 | 622.4581527 | 1.891168589 | 0.171639438 | 5.24E-29 | 2.83E-27 |
| Abcc12 | 98.95334943 | 3.944793818 | 0.358759024 | 5.35E-29 | 2.87E-27 |
| Cpe | 3309.694741 | 14.11836314 | 2.496479642 | 5.81E-29 | 3.08E-27 |
| Serpina7 | 1702.527086 | 3.449701963 | 0.315735902 | 6.29E-29 | 3.33E-27 |
| Gata6 | 1445.944863 | 1.479491087 | 0.134270128 | 7.72E-29 | 4.07E-27 |
| Pld2 | 760.4973709 | 1.669087884 | 0.151745915 | 8.46E-29 | 4.45E-27 |
| Sorl1 | 796.2105098 | 2.866406362 | 0.26164578 | 9.16E-29 | 4.79E-27 |

| | | | | | |
|---|---|---|---|---|---|
| Nfkb2 | 1428.58115 | 2.186380285 | 0.200435645 | 1.41E-28 | 7.32E-27 |
| Renbp | 183.3495117 | 4.841444901 | 0.44240731 | 1.75E-28 | 9.01E-27 |
| Gstm5 | 824.4358722 | 1.599790126 | 0.146221103 | 1.87E-28 | 9.60E-27 |
| Cd44 | 5217.688761 | 5.853114876 | 0.537569429 | 1.88E-28 | 9.61E-27 |
| Ptpn21 | 1813.38248 | 1.261775269 | 0.115330554 | 2.22E-28 | 1.13E-26 |
| Frmd6 | 1107.315075 | 2.086454107 | 0.191654133 | 2.46E-28 | 1.25E-26 |
| Dennd11 | 778.6740688 | 1.332412419 | 0.122407369 | 3.82E-28 | 1.92E-26 |
| Stk10 | 198.7867947 | 2.608680636 | 0.241345208 | 4.08E-28 | 2.04E-26 |
| Tnfaip3 | 884.3747103 | 2.198027528 | 0.203187873 | 4.34E-28 | 2.16E-26 |
| Usp20 | 584.2091082 | 1.686670957 | 0.155878044 | 5.74E-28 | 2.85E-26 |
| Atpif1 | 624.1112407 | 1.636047827 | 0.151269622 | 7.21E-28 | 3.56E-26 |
| Aldh3b1 | 313.3033028 | 2.382088236 | 0.221000269 | 8.75E-28 | 4.29E-26 |
| Rasal1 | 1177.640221 | 7.839031057 | 0.700945993 | 1.12E-27 | 5.49E-26 |
| Mgat4b | 2191.490491 | 1.726584121 | 0.161090059 | 1.89E-27 | 9.09E-26 |
| Flot1 | 1403.780142 | 1.738779532 | 0.162566765 | 2.00E-27 | 9.57E-26 |
| Ptprs | 1970.867571 | 1.890543076 | 0.17662053 | 2.07E-27 | 9.90E-26 |
| Parp3 | 1163.188464 | 1.052847382 | 0.098028661 | 2.67E-27 | 1.27E-25 |
| Gm20559 | 371.8978432 | 2.929219903 | 0.27444 | 3.50E-27 | 1.65E-25 |
| Mcm2 | 443.1561198 | 2.590796837 | 0.243541365 | 3.53E-27 | 1.66E-25 |
| Map3k1 | 3685.649888 | 1.529283646 | 0.143008937 | 5.99E-27 | 2.80E-25 |
| Acot9 | 396.9432547 | 2.753169934 | 0.260926837 | 6.10E-27 | 2.84E-25 |
| Itpripl2 | 712.9640184 | 4.377999239 | 0.41312416 | 6.32E-27 | 2.94E-25 |
| Fam102b | 539.5275745 | 6.097952841 | 0.566588207 | 1.05E-26 | 4.85E-25 |
| Lcn2 | 9438.525141 | 4.258323722 | 0.409637172 | 1.17E-26 | 5.40E-25 |
| Epn2 | 669.5473783 | 1.769305152 | 0.168247832 | 1.42E-26 | 6.48E-25 |
| Sestd1 | 1132.164538 | 2.585101138 | 0.246541438 | 1.67E-26 | 7.60E-25 |
| Arhgdia | 4321.844229 | 1.142602011 | 0.108496148 | 1.79E-26 | 8.10E-25 |
| Slc5a6 | 367.1006678 | 1.660326891 | 0.15823807 | 1.88E-26 | 8.49E-25 |
| Dtx3 | 837.0699689 | 2.471236291 | 0.236711702 | 2.12E-26 | 9.55E-25 |
| Rad51b | 414.2208351 | 5.54381123 | 0.521719019 | 2.25E-26 | 1.01E-24 |
| Tmem106a | 256.2543295 | 1.942280267 | 0.185852239 | 2.55E-26 | 1.14E-24 |
| Sgce | 311.2840929 | 2.476255068 | 0.23735701 | 2.84E-26 | 1.27E-24 |
| Lpar2 | 298.2543136 | 3.217060493 | 0.307304524 | 3.31E-26 | 1.47E-24 |
| Trip10 | 650.5203082 | 1.670836826 | 0.160426722 | 4.75E-26 | 2.10E-24 |
| Clip2 | 531.2050821 | 2.91459061 | 0.28267509 | 6.94E-26 | 3.04E-24 |
| Kctd17 | 347.6478245 | 2.276255469 | 0.219738524 | 6.92E-26 | 3.04E-24 |
| Ppp1r18 | 407.6433091 | 4.450850917 | 0.426988059 | 7.41E-26 | 3.23E-24 |
| Gss | 2150.458779 | 1.089322957 | 0.104383037 | 8.41E-26 | 3.64E-24 |
| Mbp | 759.4911749 | 4.925468889 | 0.470011399 | 1.16E-25 | 5.03E-24 |
| Rai1 | 1483.517215 | 1.132202026 | 0.109099461 | 1.24E-25 | 5.33E-24 |
| Cd36 | 1639.383397 | 2.34095487 | 0.228477291 | 1.43E-25 | 6.12E-24 |
| Ctps | 712.0956619 | 1.659065908 | 0.161210377 | 1.68E-25 | 7.11E-24 |
| Tmc6 | 564.392668 | 1.59500117 | 0.155043791 | 1.78E-25 | 7.52E-24 |

| | | | | | |
|---|---|---|---|---|---|
| Ccdc88c | 847.9117176 | 3.62146795 | 0.350830778 | 1.90E-25 | 7.99E-24 |
| Ier3 | 1501.956101 | 5.786743638 | 0.558972517 | 2.11E-25 | 8.82E-24 |
| Cln6 | 255.0913744 | 3.353779228 | 0.329637397 | 2.34E-25 | 9.78E-24 |
| Efhd2 | 2328.641903 | 1.653273095 | 0.161371391 | 2.60E-25 | 1.08E-23 |
| Ywhah | 2545.35393 | 1.256148445 | 0.122391085 | 3.35E-25 | 1.38E-23 |
| Pip4k2a | 149.09379 | 4.18435387 | 0.40637381 | 3.37E-25 | 1.39E-23 |
| Crim1 | 956.6138562 | 1.240846181 | 0.121008723 | 3.64E-25 | 1.49E-23 |
| Myo9b | 3108.436524 | 1.339131184 | 0.131077232 | 4.55E-25 | 1.86E-23 |
| Map3k14 | 444.9455037 | 2.20341747 | 0.216496263 | 4.84E-25 | 1.97E-23 |
| Pawr | 1259.389142 | 1.776298506 | 0.175042291 | 5.37E-25 | 2.18E-23 |
| Plpp2 | 1312.809125 | 1.766536982 | 0.174319988 | 7.04E-25 | 2.83E-23 |
| Epdr1 | 380.2303894 | 2.588141523 | 0.255375465 | 7.63E-25 | 3.06E-23 |
| Ahnak2 | 1008.818222 | 12.92373979 | 2.502389971 | 8.20E-25 | 3.28E-23 |
| Gal3st1 | 171.7064654 | 3.470761588 | 0.343109497 | 9.22E-25 | 3.68E-23 |
| Als2cl | 2160.895612 | 1.387127881 | 0.137197139 | 1.10E-24 | 4.39E-23 |
| Septin6 | 240.672614 | 3.445001111 | 0.340123648 | 1.49E-24 | 5.90E-23 |
| Cgref1 | 117.1709408 | 3.12736073 | 0.309239645 | 1.49E-24 | 5.90E-23 |
| Cd14 | 1958.505159 | 6.13513578 | 0.619672604 | 2.55E-24 | 9.94E-23 |
| Ajuba | 1944.842221 | 2.023153766 | 0.202696518 | 2.58E-24 | 1.00E-22 |
| Pkmyt1 | 249.9166312 | 3.872474487 | 0.38522058 | 2.91E-24 | 1.13E-22 |
| Sstr2 | 166.7671628 | 3.125101211 | 0.313632421 | 2.92E-24 | 1.13E-22 |
| Chpf | 458.4690335 | 4.026061561 | 0.399408367 | 3.21E-24 | 1.24E-22 |
| Aldh18a1 | 365.8207181 | 3.170255093 | 0.317836464 | 3.58E-24 | 1.38E-22 |
| Sh2d4a | 967.132892 | 1.241274358 | 0.12391401 | 4.12E-24 | 1.57E-22 |
| Tead1 | 1920.464134 | 1.492962968 | 0.150009209 | 6.54E-24 | 2.48E-22 |
| Scamp5 | 838.6320103 | 1.095167426 | 0.109741122 | 7.75E-24 | 2.93E-22 |
| Mmp14 | 4077.948657 | 1.486992406 | 0.15047288 | 1.08E-23 | 4.04E-22 |
| Prr15l | 824.9309506 | 6.004080608 | 0.592554657 | 1.15E-23 | 4.29E-22 |
| Rtkn | 415.7613345 | 1.726990929 | 0.175022595 | 1.24E-23 | 4.61E-22 |
| Zmynd15 | 465.3682816 | 3.31301414 | 0.33582341 | 1.25E-23 | 4.64E-22 |
| Cpeb1 | 455.0611438 | 2.66780218 | 0.272568397 | 1.35E-23 | 4.97E-22 |
| Clu | 256025.3423 | 1.347028038 | 0.136283725 | 1.38E-23 | 5.09E-22 |
| Hdac7 | 1556.245244 | 4.024136923 | 0.408504548 | 1.55E-23 | 5.70E-22 |
| H19 | 160.3576547 | 7.994986 | 0.794896445 | 1.68E-23 | 6.16E-22 |
| Slc6a8 | 1337.444188 | 1.608487242 | 0.163718116 | 2.09E-23 | 7.62E-22 |
| Pgd | 2587.353257 | 1.467658954 | 0.149643188 | 2.45E-23 | 8.88E-22 |
| Spaca6 | 362.1219917 | 3.352226337 | 0.343799782 | 2.60E-23 | 9.40E-22 |
| Trip6 | 957.7624527 | 1.273577305 | 0.130349684 | 3.42E-23 | 1.23E-21 |
| Pafah1b3 | 461.3010025 | 2.356397054 | 0.241845088 | 3.56E-23 | 1.28E-21 |
| Epha1 | 966.5023088 | 1.083896703 | 0.110661004 | 3.77E-23 | 1.34E-21 |
| Maff | 732.9096626 | 2.493058284 | 0.257037216 | 3.76E-23 | 1.34E-21 |
| Gm11843 | 224.6431216 | 3.417844363 | 0.353864655 | 4.04E-23 | 1.44E-21 |
| Rcn1 | 668.736671 | 4.230281686 | 0.428845171 | 4.44E-23 | 1.57E-21 |

| | | | | | |
|---|---|---|---|---|---|
| Pfkp | 497.9061665 | 4.634677078 | 0.475523031 | 4.55E-23 | 1.60E-21 |
| Arsa | 1561.34212 | 1.37777063 | 0.141187316 | 4.64E-23 | 1.63E-21 |
| Il3ra | 300.73524 | 3.637347603 | 0.372771696 | 4.78E-23 | 1.67E-21 |
| Numbl | 253.1200083 | 4.858491272 | 0.491482438 | 5.42E-23 | 1.89E-21 |
| Pdgfb | 1215.488821 | 5.185749978 | 0.52466783 | 6.21E-23 | 2.16E-21 |
| Ssbp4 | 537.9822375 | 3.564507303 | 0.368848017 | 6.61E-23 | 2.30E-21 |
| Mapk7 | 311.1175863 | 2.903077475 | 0.299707351 | 7.66E-23 | 2.66E-21 |
| Lynx1 | 827.3898248 | 3.105128967 | 0.320365456 | 8.82E-23 | 3.05E-21 |
| Foxp4 | 2932.029124 | 1.151870181 | 0.118905114 | 1.21E-22 | 4.19E-21 |
| Gsta2 | 1064.69359 | 1.986691795 | 0.207499922 | 1.59E-22 | 5.44E-21 |
| Elovl7 | 741.0674179 | 8.09463339 | 0.804774181 | 2.12E-22 | 7.22E-21 |
| Plekhg2 | 926.1263878 | 1.57372618 | 0.164356991 | 2.31E-22 | 7.86E-21 |
| Vcl | 1443.456179 | 1.005088541 | 0.104345627 | 2.57E-22 | 8.70E-21 |
| Fbf1 | 763.0083194 | 1.573624386 | 0.164905234 | 3.41E-22 | 1.14E-20 |
| Abhd2 | 9356.064568 | 1.068066736 | 0.111376937 | 3.92E-22 | 1.31E-20 |
| Micall2 | 254.2785188 | 3.435031812 | 0.358469558 | 4.18E-22 | 1.40E-20 |
| Efemp1 | 1823.420415 | 4.752561098 | 0.492800016 | 5.47E-22 | 1.82E-20 |
| Galnt10 | 665.5387705 | 1.119192041 | 0.117403734 | 5.89E-22 | 1.96E-20 |
| Tspan17 | 229.8664152 | 3.067068087 | 0.325027156 | 6.16E-22 | 2.04E-20 |
| Amot | 1461.64766 | 1.202611047 | 0.126710513 | 7.28E-22 | 2.40E-20 |
| Msrb3 | 486.8743528 | 1.192098674 | 0.125464685 | 7.42E-22 | 2.44E-20 |
| Ccn1 | 15724.42294 | 3.034057867 | 0.3247315 | 8.55E-22 | 2.80E-20 |
| Pqlc3 | 189.9758091 | 3.835245686 | 0.408993301 | 1.50E-21 | 4.86E-20 |
| Rarg | 572.9018362 | 2.183394641 | 0.233951344 | 1.50E-21 | 4.86E-20 |
| Ttc39a | 267.6521604 | 2.202105938 | 0.235359508 | 1.51E-21 | 4.86E-20 |
| Tpm4 | 2393.122075 | 1.720202672 | 0.184003563 | 1.65E-21 | 5.32E-20 |
| Synj2 | 884.6744228 | 2.071044236 | 0.221934015 | 1.68E-21 | 5.41E-20 |
| Vopp1 | 335.0683253 | 1.750613213 | 0.186997586 | 1.99E-21 | 6.39E-20 |
| Smad7 | 783.8921838 | 1.773309471 | 0.190536926 | 2.61E-21 | 8.29E-20 |
| Rasa2 | 307.3882813 | 1.825850499 | 0.19716163 | 5.10E-21 | 1.59E-19 |
| Shc2 | 1090.548853 | 9.387457823 | 0.943881129 | 5.45E-21 | 1.69E-19 |
| Spry1 | 453.7389473 | 4.097251714 | 0.446173455 | 8.19E-21 | 2.51E-19 |
| Vars | 2957.296763 | 1.282576065 | 0.139653887 | 1.25E-20 | 3.78E-19 |
| Sall1 | 1215.771741 | 1.077904565 | 0.117269692 | 1.33E-20 | 4.02E-19 |
| Vat1 | 2251.366067 | 1.566495578 | 0.171605204 | 1.53E-20 | 4.61E-19 |
| Ypel4 | 147.3591434 | 5.447297918 | 0.592308986 | 1.68E-20 | 5.02E-19 |
| Jag1 | 2747.116342 | 2.145327946 | 0.235969225 | 1.71E-20 | 5.12E-19 |
| Cd68 | 66.96384398 | 3.577859007 | 0.394519638 | 1.73E-20 | 5.15E-19 |
| Casp12 | 818.8026255 | 7.780384341 | 0.817313033 | 2.45E-20 | 7.20E-19 |
| Mgst3 | 503.0049499 | 1.765362149 | 0.195095197 | 2.74E-20 | 8.05E-19 |
| Pgm2 | 331.3203627 | 1.480459325 | 0.163203587 | 3.08E-20 | 9.01E-19 |
| Myo15b | 651.916575 | 6.549127312 | 0.694702291 | 3.39E-20 | 9.89E-19 |
| Ugdh | 5793.248318 | 1.156439417 | 0.127142443 | 3.82E-20 | 1.11E-18 |

| | | | | | |
|---|---|---|---|---|---|
| Tgfb2 | 1009.420013 | 6.585584989 | 0.701378901 | 4.02E-20 | 1.16E-18 |
| Mapt | 70.6686372 | 5.407855072 | 0.590224047 | 4.17E-20 | 1.20E-18 |
| Zfp704 | 1516.746185 | 1.591581278 | 0.176351649 | 4.62E-20 | 1.33E-18 |
| Mapk8ip3 | 1372.431787 | 1.229733391 | 0.13604765 | 5.08E-20 | 1.45E-18 |
| Agfg1 | 1702.21483 | 1.05120464 | 0.116032068 | 5.30E-20 | 1.52E-18 |
| Adamts1 | 1448.208397 | 1.652299992 | 0.184740049 | 7.47E-20 | 2.11E-18 |
| Syt12 | 261.8848708 | 3.240875284 | 0.365354021 | 7.86E-20 | 2.21E-18 |
| Hkdc1 | 820.3382632 | 11.58168327 | 2.444787278 | 9.86E-20 | 2.75E-18 |
| Rrs1 | 600.3031398 | 1.391446468 | 0.155724025 | 1.09E-19 | 3.02E-18 |
| Vsig10 | 1193.512276 | 2.377438714 | 0.267838541 | 1.16E-19 | 3.21E-18 |
| Zc2hc1a | 370.0259412 | 3.591245472 | 0.400648149 | 1.20E-19 | 3.32E-18 |
| Akr1b3 | 295.984315 | 1.576617875 | 0.177225425 | 1.30E-19 | 3.59E-18 |
| Ubqln2 | 1997.223593 | 1.133147721 | 0.126699218 | 1.36E-19 | 3.74E-18 |
| Cachd1 | 202.4523962 | 6.909570749 | 0.755197715 | 1.51E-19 | 4.14E-18 |
| Sertad1 | 208.1011079 | 1.929759632 | 0.217910256 | 1.60E-19 | 4.38E-18 |
| Gm30692 | 59.74695032 | 5.467056215 | 0.609375502 | 1.61E-19 | 4.39E-18 |
| Col1a1 | 287.0758492 | 7.08515035 | 0.797604679 | 1.65E-19 | 4.50E-18 |
| Lrrc8b | 514.4137737 | 2.853441122 | 0.322772026 | 1.89E-19 | 5.11E-18 |
| Arl6ip5 | 1454.711163 | 1.092037525 | 0.123242007 | 3.17E-19 | 8.50E-18 |
| Pacs1 | 485.1246709 | 1.858731294 | 0.21149142 | 3.20E-19 | 8.57E-18 |
| Tlnrd1 | 768.2721929 | 1.962854899 | 0.224062787 | 3.52E-19 | 9.41E-18 |
| Adamtsl4 | 372.6859035 | 4.572801291 | 0.521882131 | 4.78E-19 | 1.27E-17 |
| Kifc3 | 5117.250248 | 1.517249816 | 0.172638169 | 5.26E-19 | 1.39E-17 |
| Cpne8 | 1053.091013 | 2.746020626 | 0.318099222 | 5.78E-19 | 1.52E-17 |
| Sik1 | 2631.830037 | 2.176952343 | 0.251859412 | 6.87E-19 | 1.80E-17 |
| Per3 | 920.3512384 | 2.248823213 | 0.259780037 | 7.13E-19 | 1.86E-17 |
| Rbpms | 2343.026099 | 1.057972039 | 0.120641761 | 7.14E-19 | 1.86E-17 |
| Jpt1 | 993.5537819 | 1.340788871 | 0.153598063 | 7.29E-19 | 1.90E-17 |
| Slx4 | 295.4564903 | 1.73294193 | 0.19929581 | 7.54E-19 | 1.96E-17 |
| Lmna | 3808.725539 | 1.310368398 | 0.150269889 | 7.73E-19 | 2.01E-17 |
| Nfkbie | 825.1573438 | 2.115699006 | 0.244664644 | 7.83E-19 | 2.03E-17 |
| Gpx3 | 580.382517 | 7.530804759 | 0.829499765 | 8.57E-19 | 2.21E-17 |
| Scd2 | 2893.259598 | 4.541717746 | 0.533527927 | 9.26E-19 | 2.38E-17 |
| Pvr | 1029.550134 | 1.127761245 | 0.129307862 | 9.30E-19 | 2.39E-17 |
| Icam1 | 1380.017496 | 1.786781949 | 0.206611116 | 9.61E-19 | 2.46E-17 |
| Sparc | 3332.374321 | 2.58709178 | 0.301478489 | 1.01E-18 | 2.57E-17 |
| Tox | 167.0679494 | 1.978142194 | 0.228968598 | 1.02E-18 | 2.61E-17 |
| Arhgef2 | 883.8819265 | 5.799117381 | 0.649830334 | 1.05E-18 | 2.67E-17 |
| Slc11a1 | 90.88817745 | 2.731701884 | 0.318191242 | 1.15E-18 | 2.91E-17 |
| Ttll7 | 196.8109785 | 5.233184295 | 0.591497142 | 1.36E-18 | 3.43E-17 |
| Ngf | 821.5664978 | 10.9723711 | 2.400109807 | 1.83E-18 | 4.58E-17 |
| Osmr | 728.5720933 | 4.94686219 | 0.568965671 | 1.86E-18 | 4.66E-17 |
| Ccdc9b | 488.6145843 | 3.435218431 | 0.398027382 | 1.95E-18 | 4.87E-17 |

| | | | | | |
|---|---|---|---|---|---|
| Tspan8 | 2652.598426 | 10.90850171 | 2.321897702 | 1.97E-18 | 4.90E-17 |
| Ehd2 | 874.5795891 | 3.652187695 | 0.420982198 | 2.02E-18 | 5.03E-17 |
| Smtn | 179.215168 | 2.979542859 | 0.348917853 | 2.11E-18 | 5.24E-17 |
| Panx1 | 187.9800615 | 2.865435423 | 0.333356699 | 2.31E-18 | 5.73E-17 |
| Tnip1 | 1633.710541 | 1.267445407 | 0.147652285 | 2.45E-18 | 6.06E-17 |
| Map3k6 | 206.4725233 | 3.282308149 | 0.386254895 | 2.79E-18 | 6.88E-17 |
| Wfdc2 | 803.7245441 | 2.8590404 | 0.339042971 | 3.45E-18 | 8.46E-17 |
| Nol4l | 351.6902361 | 3.098056482 | 0.36438619 | 3.56E-18 | 8.71E-17 |
| Elf3 | 545.9594396 | 2.666579422 | 0.315361136 | 4.17E-18 | 1.02E-16 |
| Apobec3 | 1030.778116 | 5.700670243 | 0.652291516 | 4.84E-18 | 1.18E-16 |
| AI506816 | 838.3826187 | 1.369974026 | 0.161364509 | 5.34E-18 | 1.30E-16 |
| Iqce | 668.231522 | 1.323358055 | 0.155797112 | 5.74E-18 | 1.39E-16 |
| Myl12b | 270.2849819 | 2.396116196 | 0.283430787 | 6.09E-18 | 1.47E-16 |
| Vegfb | 442.2222335 | 1.460852362 | 0.172524717 | 6.44E-18 | 1.55E-16 |
| Slc7a11 | 289.7717956 | 12.44780641 | 2.714338604 | 6.81E-18 | 1.64E-16 |
| Ttyh3 | 1151.164908 | 1.492815384 | 0.177437849 | 1.01E-17 | 2.40E-16 |
| Flt1 | 91.95648371 | 4.593917652 | 0.544980558 | 1.04E-17 | 2.47E-16 |
| Adamts10 | 323.1297261 | 2.71577664 | 0.324269601 | 1.27E-17 | 2.98E-16 |
| Nenf | 428.8913011 | 1.233511162 | 0.1470308 | 1.78E-17 | 4.13E-16 |
| Nrg4 | 319.757649 | 1.859741158 | 0.22425002 | 1.86E-17 | 4.32E-16 |
| Tnks1bp1 | 3117.175824 | 1.016201972 | 0.121632563 | 2.02E-17 | 4.68E-16 |
| Syne4 | 206.0681172 | 2.17355425 | 0.263314209 | 2.55E-17 | 5.87E-16 |
| Ccdc88a | 262.5824066 | 3.46820428 | 0.418176334 | 2.68E-17 | 6.16E-16 |
| Enc1 | 2788.519373 | 1.350095108 | 0.162505868 | 2.69E-17 | 6.16E-16 |
| Gstm3 | 9891.051661 | 1.637389148 | 0.197998608 | 2.83E-17 | 6.47E-16 |
| Klf7 | 669.0837074 | 1.529097258 | 0.184087438 | 2.85E-17 | 6.49E-16 |
| Spink1 | 106.7319113 | 6.455879152 | 0.759897383 | 2.86E-17 | 6.50E-16 |
| Nt5c3b | 304.9629369 | 1.733738121 | 0.209372396 | 3.03E-17 | 6.87E-16 |
| Dpysl3 | 1541.559428 | 5.558369038 | 0.652995278 | 3.18E-17 | 7.22E-16 |
| 2010003K11Rik | 269.8857951 | 2.094390797 | 0.255146057 | 3.26E-17 | 7.39E-16 |
| Tgif1 | 2104.999954 | 1.126115295 | 0.136493887 | 3.40E-17 | 7.70E-16 |
| Cand2 | 348.2391153 | 2.746722163 | 0.334988936 | 3.56E-17 | 8.04E-16 |
| Tmprss2 | 865.3803796 | 1.679572724 | 0.204034333 | 3.98E-17 | 8.95E-16 |
| Fgd1 | 263.1404083 | 1.787635711 | 0.217024614 | 4.20E-17 | 9.40E-16 |
| Defb1 | 80.18627089 | 3.745101019 | 0.461309305 | 4.34E-17 | 9.69E-16 |
| Col4a5 | 1479.027157 | 2.187206002 | 0.267866432 | 4.45E-17 | 9.94E-16 |
| Stxbp1 | 365.8815636 | 3.605812953 | 0.433978882 | 4.51E-17 | 1.01E-15 |
| Lig1 | 583.7963409 | 2.411929002 | 0.296397984 | 4.68E-17 | 1.04E-15 |
| Myo5c | 899.6359621 | 10.54674543 | 2.369988354 | 5.05E-17 | 1.12E-15 |
| Pdgfa | 613.5389746 | 1.063220867 | 0.128675528 | 5.64E-17 | 1.24E-15 |
| Pcgf2 | 515.508143 | 1.513284696 | 0.184502398 | 5.80E-17 | 1.28E-15 |
| Zfand2a | 394.3956612 | 1.256052281 | 0.152603884 | 5.85E-17 | 1.29E-15 |
| Prrg4 | 366.0548761 | 2.009109165 | 0.245392679 | 5.96E-17 | 1.31E-15 |

| | | | | | |
|---|---|---|---|---|---|
| 9830144P21Rik | 62.84849137 | 3.430372598 | 0.422370203 | 6.10E-17 | 1.34E-15 |
| Cep170 | 942.8738129 | 1.262028169 | 0.153518958 | 6.57E-17 | 1.44E-15 |
| Id3 | 1361.904331 | 2.231918914 | 0.276239675 | 7.77E-17 | 1.69E-15 |
| B4galt6 | 2330.660997 | 9.493170406 | 1.236019743 | 8.16E-17 | 1.77E-15 |
| Limd2 | 808.6726302 | 1.020268841 | 0.124025935 | 8.26E-17 | 1.79E-15 |
| Apoa4 | 1084.809155 | 2.574369734 | 0.319829303 | 8.53E-17 | 1.84E-15 |
| Alcam | 7033.895068 | 1.144314154 | 0.139321968 | 9.59E-17 | 2.06E-15 |
| Marcksl1 | 1259.36562 | 2.898905894 | 0.357032033 | 1.08E-16 | 2.32E-15 |
| Camk1 | 677.6305656 | 1.353527217 | 0.166009218 | 1.26E-16 | 2.69E-15 |
| Homer3 | 249.8333661 | 3.713452702 | 0.452521862 | 1.30E-16 | 2.77E-15 |
| Gsta4 | 1446.511618 | 1.264161554 | 0.1554045 | 1.36E-16 | 2.89E-15 |
| Per1 | 1792.911099 | 2.420168978 | 0.302538558 | 1.39E-16 | 2.96E-15 |
| Slc1a5 | 134.5869544 | 2.419238875 | 0.299707885 | 1.44E-16 | 3.07E-15 |
| S100a6 | 1433.97565 | 9.441068076 | 1.242079158 | 1.49E-16 | 3.17E-15 |
| Mex3d | 432.687381 | 1.806657029 | 0.223485285 | 1.55E-16 | 3.27E-15 |
| Pck2 | 692.4583247 | 1.498686599 | 0.185544987 | 1.65E-16 | 3.48E-15 |
| Myo7b | 162.8584993 | 3.167986075 | 0.392831138 | 1.82E-16 | 3.83E-15 |
| Cdc42ep1 | 2642.241788 | 1.06530294 | 0.131366749 | 1.92E-16 | 4.03E-15 |
| Cyp4f16 | 359.3505711 | 1.27443297 | 0.157697232 | 1.92E-16 | 4.04E-15 |
| Bin1 | 559.9559464 | 1.006838035 | 0.123977285 | 2.08E-16 | 4.36E-15 |
| Card19 | 422.0498646 | 1.564645143 | 0.194741138 | 2.09E-16 | 4.38E-15 |
| Cd2ap | 6195.42875 | 1.141804536 | 0.141378671 | 2.47E-16 | 5.14E-15 |
| Dip2a | 536.4515405 | 1.173703506 | 0.145333731 | 2.57E-16 | 5.35E-15 |
| Plscr1 | 2805.771494 | 1.752765057 | 0.219933191 | 3.18E-16 | 6.58E-15 |
| Prmt2 | 389.0165 | 4.001318137 | 0.493296808 | 3.25E-16 | 6.72E-15 |
| Cfb | 373.2758448 | 1.25846604 | 0.156539638 | 3.25E-16 | 6.72E-15 |
| Ankrd12 | 565.6495619 | 1.762030938 | 0.221709883 | 3.55E-16 | 7.32E-15 |
| Zdhhc2 | 235.6616781 | 2.229451334 | 0.280825042 | 3.84E-16 | 7.90E-15 |
| Fndc10 | 160.4928545 | 7.059745241 | 0.852976429 | 3.86E-16 | 7.93E-15 |
| Gpx2 | 225.5354571 | 5.416038674 | 0.663856197 | 4.18E-16 | 8.56E-15 |
| Il17rb | 482.2129692 | 1.307884867 | 0.163828229 | 4.26E-16 | 8.73E-15 |
| Zyx | 2597.269208 | 1.382143599 | 0.173642437 | 4.56E-16 | 9.31E-15 |
| Pparg | 190.8325974 | 1.659186094 | 0.209336399 | 4.57E-16 | 9.31E-15 |
| Megf8 | 1795.980567 | 1.105714762 | 0.138747675 | 4.68E-16 | 9.53E-15 |
| Ntn4 | 703.8153015 | 2.844129568 | 0.357252906 | 4.78E-16 | 9.71E-15 |
| Patz1 | 628.759567 | 1.238664255 | 0.154729579 | 4.81E-16 | 9.76E-15 |
| Plscr3 | 1050.621523 | 1.272575947 | 0.159631416 | 5.00E-16 | 1.01E-14 |
| Inava | 848.6684542 | 8.097525449 | 0.948546344 | 5.09E-16 | 1.03E-14 |
| Naip2 | 377.4846798 | 2.028465016 | 0.257831797 | 5.22E-16 | 1.06E-14 |
| Tspan15 | 164.8650929 | 3.304526894 | 0.417084546 | 5.36E-16 | 1.08E-14 |
| Sema3b | 343.1791066 | 3.184715238 | 0.406138204 | 6.05E-16 | 1.22E-14 |
| Prom1 | 1700.99516 | 5.846588177 | 0.71229252 | 6.22E-16 | 1.25E-14 |
| Rbp1 | 1190.235878 | 1.314084798 | 0.165993615 | 7.23E-16 | 1.45E-14 |

| | | | | | |
|---|---|---|---|---|---|
| 2200002D01Rik | 422.0612183 | 1.356453845 | 0.171434288 | 7.36E-16 | 1.47E-14 |
| Sat1 | 2158.541359 | 1.549654307 | 0.196797014 | 7.65E-16 | 1.53E-14 |
| Igdcc4 | 576.4080511 | 10.08013172 | 2.367457897 | 8.23E-16 | 1.64E-14 |
| Caskin2 | 917.0766179 | 1.142152281 | 0.144343236 | 8.41E-16 | 1.68E-14 |
| Cldn4 | 2945.534617 | 10.17558507 | 2.325285256 | 8.56E-16 | 1.71E-14 |
| Cys1 | 897.4977421 | 2.234828933 | 0.286123684 | 8.57E-16 | 1.71E-14 |
| Ccn5 | 429.3534084 | 10.19206185 | 2.393076996 | 8.90E-16 | 1.77E-14 |
| Fgfr2 | 2376.307682 | 1.022390816 | 0.128867786 | 9.18E-16 | 1.82E-14 |
| Ihh | 144.1151708 | 2.907589523 | 0.376446988 | 9.91E-16 | 1.96E-14 |
| Tmem237 | 245.9704624 | 1.511901485 | 0.19236502 | 1.09E-15 | 2.16E-14 |
| Il10rb | 334.0185372 | 1.892494164 | 0.241750512 | 1.11E-15 | 2.19E-14 |
| Adrb2 | 115.8926575 | 4.649546714 | 0.590003496 | 1.16E-15 | 2.29E-14 |
| H2-T-ps | 65.49388414 | 4.871006565 | 0.628917792 | 1.29E-15 | 2.53E-14 |
| Plp2 | 422.2736418 | 3.283069555 | 0.41926012 | 1.32E-15 | 2.58E-14 |
| Elmo1 | 189.5489611 | 1.970488318 | 0.253282592 | 1.37E-15 | 2.67E-14 |
| Irak3 | 201.5284826 | 4.037228502 | 0.524647102 | 1.40E-15 | 2.72E-14 |
| Gas6 | 8246.924178 | 1.37541654 | 0.176142238 | 1.51E-15 | 2.92E-14 |
| Myof | 3592.60382 | 7.899529148 | 0.93788107 | 1.62E-15 | 3.12E-14 |
| Gmip | 381.6356426 | 1.329280952 | 0.170079018 | 1.72E-15 | 3.30E-14 |
| Llgl1 | 950.5135454 | 1.006586729 | 0.128184249 | 1.80E-15 | 3.45E-14 |
| Pofut2 | 689.6872368 | 1.0793335 | 0.137564519 | 1.87E-15 | 3.59E-14 |
| Dusp10 | 443.5759271 | 1.62355643 | 0.209698213 | 2.14E-15 | 4.08E-14 |
| Usp43 | 282.5132071 | 9.936082669 | 2.378376387 | 2.14E-15 | 4.08E-14 |
| Itga3 | 4405.744063 | 1.05445371 | 0.134799658 | 2.16E-15 | 4.12E-14 |
| Pmepa1 | 571.701155 | 3.453832681 | 0.444822411 | 2.43E-15 | 4.62E-14 |
| Ldb1 | 1551.88774 | 1.183519308 | 0.151944758 | 2.44E-15 | 4.65E-14 |
| Slc48a1 | 1834.725727 | 1.335626303 | 0.171711154 | 2.55E-15 | 4.85E-14 |
| Eif6 | 2382.439995 | 1.403160481 | 0.181630495 | 2.61E-15 | 4.95E-14 |
| Phlda2 | 49.27107646 | 3.134135861 | 0.408665469 | 2.80E-15 | 5.30E-14 |
| Sdk1 | 46.62161858 | 4.518592074 | 0.577819945 | 3.28E-15 | 6.19E-14 |
| Cfap44 | 169.760744 | 9.937909651 | 2.418428301 | 3.46E-15 | 6.52E-14 |
| Arrb2 | 557.5798128 | 1.342413208 | 0.173976743 | 3.93E-15 | 7.39E-14 |
| Cbr1 | 2163.358895 | 1.400926063 | 0.182053447 | 4.16E-15 | 7.81E-14 |
| Prkx | 470.6073335 | 1.247864092 | 0.162230723 | 4.30E-15 | 8.05E-14 |
| Rph3al | 222.2133515 | 1.866409109 | 0.244357763 | 4.48E-15 | 8.37E-14 |
| Filip1l | 853.4650318 | 3.08632359 | 0.401652774 | 5.11E-15 | 9.50E-14 |
| Glis3 | 1805.987742 | 8.841594063 | 1.225152713 | 5.20E-15 | 9.66E-14 |
| Nt5c | 624.4954379 | 1.140433676 | 0.148338593 | 5.68E-15 | 1.05E-13 |
| Gsn | 204.4074316 | 3.857084161 | 0.502755594 | 5.82E-15 | 1.07E-13 |
| Capg | 357.6854752 | 3.7082518 | 0.482248863 | 6.56E-15 | 1.21E-13 |
| Sfi1 | 715.685044 | 1.560446439 | 0.205428087 | 6.75E-15 | 1.24E-13 |
| Ccl2 | 1991.525862 | 9.041125329 | 1.26222352 | 7.04E-15 | 1.29E-13 |
| Arrdc1 | 598.5550887 | 2.471477818 | 0.327419114 | 7.07E-15 | 1.29E-13 |

| | | | | | |
|---|---|---|---|---|---|
| Smpdl3b | 85.33293135 | 4.15930076 | 0.563881611 | 1.30E-14 | 2.35E-13 |
| Sema6b | 136.7105071 | 2.093610489 | 0.281300481 | 1.39E-14 | 2.48E-13 |
| Wls | 710.2318948 | 4.071127002 | 0.539510151 | 1.62E-14 | 2.88E-13 |
| Gramd1b | 1296.929223 | 5.409544651 | 0.695736507 | 2.05E-14 | 3.61E-13 |
| Pwwp2b | 389.8822049 | 1.777423812 | 0.238556101 | 2.06E-14 | 3.63E-13 |
| Ift57 | 380.7265344 | 2.608226327 | 0.349150272 | 2.08E-14 | 3.65E-13 |
| Myzap | 218.2572142 | 4.970773497 | 0.648445325 | 2.17E-14 | 3.80E-13 |
| Wtip | 322.9003128 | 3.515185108 | 0.468978906 | 2.45E-14 | 4.26E-13 |
| Tubb2b | 799.4274759 | 2.87809359 | 0.387469377 | 2.77E-14 | 4.79E-13 |
| Cntln | 203.4023102 | 2.935633569 | 0.394364355 | 2.94E-14 | 5.09E-13 |
| Cyp21a1 | 42.28162249 | 5.49506169 | 0.734014519 | 3.53E-14 | 6.05E-13 |
| Igsf1 | 246.8485605 | 2.415866465 | 0.329061509 | 4.21E-14 | 7.18E-13 |
| Sntb2 | 658.9291161 | 1.233292766 | 0.167740926 | 4.27E-14 | 7.28E-13 |
| Gm10382 | 298.4206319 | 2.226799275 | 0.306080665 | 4.36E-14 | 7.41E-13 |
| Kif21b | 199.0264851 | 4.567312151 | 0.620614962 | 4.53E-14 | 7.69E-13 |
| Ralgds | 504.2243016 | 2.523327932 | 0.343992567 | 4.53E-14 | 7.69E-13 |
| Plekha1 | 577.5528988 | 1.091967932 | 0.147256012 | 4.83E-14 | 8.17E-13 |
| Dnah1 | 50.18948693 | 3.666703906 | 0.505082562 | 4.93E-14 | 8.33E-13 |
| Tbc1d1 | 937.9133007 | 1.729766156 | 0.236403502 | 5.27E-14 | 8.86E-13 |
| Hspb1 | 1610.709725 | 1.67880924 | 0.230132431 | 5.26E-14 | 8.86E-13 |
| Myc | 751.0874706 | 2.394666373 | 0.331499956 | 5.47E-14 | 9.17E-13 |
| Adcy6 | 1714.314775 | 1.043379418 | 0.141058317 | 5.50E-14 | 9.22E-13 |
| Neurl1a | 470.1662382 | 6.289350378 | 0.819773881 | 5.86E-14 | 9.80E-13 |
| Ccdc34 | 319.247354 | 1.526609487 | 0.207988375 | 6.81E-14 | 1.14E-12 |
| Col1a2 | 210.6582226 | 5.423233876 | 0.761530567 | 6.99E-14 | 1.16E-12 |
| Scube3 | 1894.471044 | 6.101474677 | 0.794659842 | 7.96E-14 | 1.32E-12 |
| Klf16 | 267.4824656 | 1.593769781 | 0.219776548 | 8.10E-14 | 1.34E-12 |
| Ppp1r14b | 821.5301011 | 1.586678494 | 0.218453495 | 8.12E-14 | 1.34E-12 |
| Arhgap22 | 127.3198972 | 9.407839398 | 2.396282636 | 8.27E-14 | 1.37E-12 |
| Iffo2 | 802.0168097 | 1.352752527 | 0.185704598 | 8.79E-14 | 1.45E-12 |
| Gpnmb | 193.3997859 | 8.41311897 | 1.116122528 | 8.99E-14 | 1.48E-12 |
| Sinhcaf | 306.9271048 | 1.965234792 | 0.270704947 | 9.15E-14 | 1.51E-12 |
| Smc4 | 580.4512356 | 1.0567421 | 0.144232034 | 9.66E-14 | 1.59E-12 |
| Nlgn2 | 821.4250603 | 2.32958015 | 0.323903456 | 9.92E-14 | 1.63E-12 |
| Evpl | 386.3438727 | 3.968232614 | 0.540819763 | 1.03E-13 | 1.69E-12 |
| Evc2 | 494.090182 | 2.650587829 | 0.366515433 | 1.11E-13 | 1.81E-12 |
| Hsf4 | 173.3766554 | 1.301727075 | 0.179120715 | 1.13E-13 | 1.84E-12 |
| Bhlhe40 | 4417.571025 | 1.27399246 | 0.175508923 | 1.14E-13 | 1.86E-12 |
| Cdc42ep5 | 191.1716294 | 2.144607477 | 0.299182939 | 1.14E-13 | 1.86E-12 |
| Dvl2 | 358.4362429 | 1.294864977 | 0.178430315 | 1.20E-13 | 1.95E-12 |
| Hmox1 | 397.8150982 | 2.890228123 | 0.407621006 | 1.21E-13 | 1.97E-12 |
| Ssh3 | 1102.583597 | 1.11789478 | 0.153664256 | 1.33E-13 | 2.16E-12 |
| Slc4a9 | 108.0643503 | 4.824545097 | 0.663537731 | 1.38E-13 | 2.23E-12 |

| | | | | | |
|---|---|---|---|---|---|
| Plekhh1 | 484.7890274 | 2.536306974 | 0.352099695 | 1.52E-13 | 2.44E-12 |
| Mturn | 357.8250301 | 2.132234429 | 0.299701922 | 1.82E-13 | 2.89E-12 |
| Nob1 | 296.8376211 | 1.114670298 | 0.154150323 | 1.89E-13 | 3.00E-12 |
| Aqp7 | 53.23588197 | 3.293284335 | 0.467614718 | 2.14E-13 | 3.40E-12 |
| Sgk1 | 893.018067 | 1.550121763 | 0.216346617 | 2.16E-13 | 3.42E-12 |
| Tspo | 2103.880392 | 1.088771587 | 0.151299916 | 2.26E-13 | 3.56E-12 |
| Tinag | 63.22689619 | 9.360824484 | 2.441283056 | 2.30E-13 | 3.62E-12 |
| Rhno1 | 221.0462134 | 1.284141767 | 0.178941597 | 2.41E-13 | 3.80E-12 |
| Arntl2 | 239.926964 | 2.071478919 | 0.291970425 | 2.46E-13 | 3.86E-12 |
| Abi2 | 345.2022163 | 1.723331505 | 0.242872308 | 2.83E-13 | 4.43E-12 |
| Wfdc3 | 166.8394908 | 8.266362177 | 1.240793664 | 2.92E-13 | 4.55E-12 |
| Gadd45b | 590.3505394 | 2.331339284 | 0.330751167 | 2.96E-13 | 4.60E-12 |
| Nop56 | 712.0553827 | 1.060775713 | 0.147933665 | 3.02E-13 | 4.70E-12 |
| Gm47163 | 168.2346981 | 9.409919651 | 2.397409148 | 3.02E-13 | 4.70E-12 |
| Spire2 | 89.62898808 | 5.543209872 | 0.760937436 | 3.09E-13 | 4.80E-12 |
| Slc10a2 | 1044.596959 | 1.74876222 | 0.247186736 | 3.15E-13 | 4.89E-12 |
| Ncoa7 | 222.6560308 | 3.342814474 | 0.467941696 | 3.17E-13 | 4.91E-12 |
| Cbarp | 627.3869157 | 1.75722071 | 0.248660038 | 3.32E-13 | 5.13E-12 |
| Pisd-ps1 | 346.6669703 | 1.15240251 | 0.16135603 | 3.42E-13 | 5.27E-12 |
| Cdk1 | 134.3869496 | 3.015381815 | 0.431957898 | 3.81E-13 | 5.86E-12 |
| Eln | 69.43323387 | 5.286429256 | 0.741216175 | 3.90E-13 | 5.99E-12 |
| Wipf3 | 295.4003071 | 1.413399144 | 0.199936929 | 4.01E-13 | 6.16E-12 |
| Gar1 | 254.5375941 | 1.396639205 | 0.197694868 | 4.48E-13 | 6.83E-12 |
| Srgap2 | 1732.919358 | 1.073052406 | 0.15083579 | 4.49E-13 | 6.84E-12 |
| Arhgap11a | 186.5055447 | 2.553012091 | 0.36783346 | 4.68E-13 | 7.12E-12 |
| Wee1 | 633.3205705 | 2.022429872 | 0.290956551 | 5.21E-13 | 7.87E-12 |
| AC120557.1 | 465.4717434 | 1.176882359 | 0.16619793 | 5.44E-13 | 8.21E-12 |
| 2410002F23Rik | 543.5089778 | 1.328471673 | 0.18861521 | 5.49E-13 | 8.28E-12 |
| Glis2 | 1730.018636 | 8.809953528 | 2.237531616 | 6.06E-13 | 9.11E-12 |
| Piezo1 | 568.0378654 | 1.502228805 | 0.214289837 | 6.29E-13 | 9.44E-12 |
| Cldn7 | 1159.772996 | 8.068632723 | 1.222013888 | 6.83E-13 | 1.02E-11 |
| Nrm | 108.2153474 | 3.38671312 | 0.480545466 | 7.23E-13 | 1.08E-11 |
| Orm2 | 812.1729721 | 2.562952358 | 0.375057217 | 8.87E-13 | 1.31E-11 |
| Sh3bgrl3 | 845.8108824 | 2.235424199 | 0.321804299 | 8.91E-13 | 1.32E-11 |
| Chst14 | 157.8270391 | 2.026879686 | 0.292547027 | 1.03E-12 | 1.52E-11 |
| Crybg2 | 82.83462426 | 1.7847412 | 0.258483513 | 1.12E-12 | 1.65E-11 |
| Zfp408 | 282.5098689 | 1.122015948 | 0.160297596 | 1.13E-12 | 1.66E-11 |
| Mlph | 956.576832 | 5.745949519 | 0.786308039 | 1.14E-12 | 1.67E-11 |
| Dhrs13 | 175.7217994 | 1.67164684 | 0.242707753 | 1.15E-12 | 1.68E-11 |
| Fam110c | 159.0920429 | 1.804321355 | 0.262003058 | 1.29E-12 | 1.89E-11 |
| Gimap8 | 199.25466 | 1.874356537 | 0.272025847 | 1.43E-12 | 2.09E-11 |
| Stab1 | 192.5229586 | 1.833407689 | 0.269085332 | 1.58E-12 | 2.31E-11 |
| Ccnb2 | 93.98681835 | 5.41486695 | 0.770585981 | 1.71E-12 | 2.49E-11 |

| | | | | | |
|---|---|---|---|---|---|
| S100g | 48.57264044 | 9.313432101 | 2.519951141 | 1.80E-12 | 2.61E-11 |
| Relb | 949.3516041 | 3.176315277 | 0.475024541 | 1.83E-12 | 2.65E-11 |
| Vasp | 1151.082075 | 1.371632556 | 0.19993795 | 1.84E-12 | 2.65E-11 |
| Ppan | 430.995834 | 1.154335246 | 0.167510102 | 2.04E-12 | 2.94E-11 |
| Anxa9 | 147.648838 | 2.408113369 | 0.35515959 | 2.29E-12 | 3.28E-11 |
| F2rl1 | 687.9524739 | 8.617552664 | 2.262344641 | 2.39E-12 | 3.42E-11 |
| Cx3cl1 | 553.4927526 | 8.666055885 | 2.285650701 | 2.61E-12 | 3.72E-11 |
| Col6a1 | 131.2314135 | 4.429895782 | 0.643249825 | 2.65E-12 | 3.78E-11 |
| Zfp579 | 418.3026074 | 1.209827611 | 0.177423212 | 2.80E-12 | 3.99E-11 |
| Tpd52 | 1203.018971 | 1.004409371 | 0.146491308 | 3.12E-12 | 4.44E-11 |
| Ccdc80 | 80.17158069 | 3.601156 | 0.540525792 | 3.17E-12 | 4.50E-11 |
| Fmnl3 | 172.9649336 | 4.569497886 | 0.658031552 | 3.20E-12 | 4.54E-11 |
| Ltbp2 | 2001.585713 | 3.982499634 | 0.596264017 | 3.33E-12 | 4.71E-11 |
| Tuft1 | 1638.192115 | 1.601015198 | 0.237840347 | 3.44E-12 | 4.86E-11 |
| Hes1 | 3064.428263 | 2.678215214 | 0.406414655 | 3.75E-12 | 5.28E-11 |
| Fam124a | 562.6914112 | 1.821034351 | 0.272036956 | 3.86E-12 | 5.43E-11 |
| Nek8 | 575.843703 | 1.225468334 | 0.180629845 | 3.94E-12 | 5.53E-11 |
| Fam83g | 249.1450262 | 1.416076534 | 0.209666376 | 3.98E-12 | 5.59E-11 |
| Fyb2 | 313.8812169 | 1.424998137 | 0.211473754 | 4.01E-12 | 5.63E-11 |
| Zfp7 | 227.0214703 | 1.744920581 | 0.259467489 | 4.24E-12 | 5.93E-11 |
| Mfsd10 | 530.4308295 | 1.637285965 | 0.244682387 | 4.76E-12 | 6.63E-11 |
| Plekha2 | 518.5000918 | 2.961048389 | 0.437768795 | 4.85E-12 | 6.74E-11 |
| Tlr2 | 197.9122868 | 2.040330664 | 0.307392512 | 4.99E-12 | 6.92E-11 |
| Fes | 209.5557772 | 1.632999609 | 0.24477706 | 5.14E-12 | 7.10E-11 |
| Loxl2 | 261.5046318 | 3.874581135 | 0.569524814 | 5.14E-12 | 7.10E-11 |
| Hp | 63465.89234 | 1.186893633 | 0.176629469 | 6.34E-12 | 8.69E-11 |
| Mki67 | 612.7062076 | 7.531150431 | 1.086438786 | 7.29E-12 | 9.93E-11 |
| Plekhg5 | 368.7457472 | 1.335803242 | 0.200747646 | 7.90E-12 | 1.07E-10 |
| Neurl3 | 937.2942732 | 3.956371852 | 0.611967006 | 7.94E-12 | 1.08E-10 |
| Nol3 | 60.02810625 | 2.187199064 | 0.33337137 | 8.46E-12 | 1.14E-10 |
| Guca2b | 85.09988825 | 8.649640188 | 2.365316634 | 8.80E-12 | 1.19E-10 |
| Snta1 | 1188.451695 | 1.010995639 | 0.150850069 | 8.90E-12 | 1.20E-10 |
| Otud1 | 549.4273923 | 1.689784833 | 0.257044216 | 9.09E-12 | 1.22E-10 |
| Car13 | 246.9664239 | 5.543975998 | 0.795994561 | 9.29E-12 | 1.25E-10 |
| Gcnt2 | 860.2779428 | 1.042873812 | 0.156473359 | 9.64E-12 | 1.29E-10 |
| Syt14 | 302.476795 | 5.596505878 | 0.806526088 | 9.73E-12 | 1.30E-10 |
| Tcim | 1708.433264 | 1.729479867 | 0.264314766 | 1.08E-11 | 1.44E-10 |
| Col6a3 | 98.86697856 | 9.844594406 | 2.57178346 | 1.24E-11 | 1.64E-10 |
| Knstrn | 85.9622077 | 4.31841862 | 0.657242429 | 1.27E-11 | 1.68E-10 |
| Casp4 | 284.5309296 | 9.397646294 | 2.451985561 | 1.29E-11 | 1.70E-10 |
| Morc4 | 210.0469946 | 1.463428801 | 0.2228235 | 1.29E-11 | 1.71E-10 |
| Exoc6b | 566.5901023 | 1.039320969 | 0.156725978 | 1.40E-11 | 1.85E-10 |
| Lrrc8e | 194.3214274 | 3.977729113 | 0.59329945 | 1.41E-11 | 1.85E-10 |

| | | | | | |
|---|---|---|---|---|---|
| Abcc1 | 531.2414062 | 5.821230627 | 0.854936967 | 1.47E-11 | 1.93E-10 |
| Rdh13 | 280.8483168 | 1.078458977 | 0.163321473 | 1.51E-11 | 1.99E-10 |
| Ankrd6 | 231.7641769 | 4.763687262 | 0.700734425 | 1.52E-11 | 1.99E-10 |
| Dqx1 | 286.2680548 | 1.467582427 | 0.224672525 | 1.57E-11 | 2.05E-10 |
| Gm4673 | 242.5809454 | 1.514772367 | 0.23239163 | 1.60E-11 | 2.09E-10 |
| Fblim1 | 1361.651429 | 7.512501745 | 1.201124075 | 1.65E-11 | 2.15E-10 |
| Herc3 | 481.0671561 | 1.307929978 | 0.199814343 | 1.71E-11 | 2.22E-10 |
| Tbc1d4 | 316.2216909 | 1.521537964 | 0.23334522 | 1.96E-11 | 2.54E-10 |
| Serpinb8 | 97.19369215 | 1.759521464 | 0.272357401 | 1.98E-11 | 2.57E-10 |
| Agpat1 | 579.3948924 | 1.553560255 | 0.239771306 | 2.00E-11 | 2.60E-10 |
| Slc5a1 | 1330.675187 | 6.67838095 | 0.956184566 | 2.09E-11 | 2.69E-10 |
| Ehbp1l1 | 179.9912164 | 1.480174276 | 0.228184644 | 2.12E-11 | 2.73E-10 |
| Anks6 | 165.4853608 | 2.625652071 | 0.408050474 | 2.34E-11 | 3.01E-10 |
| C1qtnf6 | 270.8704403 | 3.021773557 | 0.466315968 | 2.35E-11 | 3.03E-10 |
| Csf2ra | 283.1402822 | 1.805228092 | 0.279063563 | 2.43E-11 | 3.12E-10 |
| Cxcl10 | 146.0387893 | 2.367615842 | 0.372548664 | 2.44E-11 | 3.13E-10 |
| Baiap2 | 1499.209392 | 1.246869164 | 0.192102899 | 2.52E-11 | 3.23E-10 |
| Nrip2 | 99.37550179 | 4.004308238 | 0.6081303 | 2.54E-11 | 3.25E-10 |
| Insyn1 | 111.7776807 | 6.826303978 | 0.996311023 | 2.77E-11 | 3.54E-10 |
| Slc12a4 | 914.6871322 | 1.110268608 | 0.170521423 | 2.87E-11 | 3.65E-10 |
| Trim35 | 1506.964685 | 2.750828434 | 0.425290412 | 3.08E-11 | 3.91E-10 |
| Neu3 | 117.7387385 | 1.58643996 | 0.24865844 | 3.31E-11 | 4.20E-10 |
| Nampt | 1349.240771 | 1.501199785 | 0.233917478 | 3.36E-11 | 4.26E-10 |
| Itpkc | 261.729967 | 1.258166003 | 0.195063276 | 3.53E-11 | 4.46E-10 |
| Tlcd5 | 236.7492689 | 3.306970822 | 0.521351977 | 3.55E-11 | 4.48E-10 |
| Enoph1 | 220.9560518 | 1.130364816 | 0.174772195 | 4.05E-11 | 5.06E-10 |
| Ctdspl | 482.493607 | 1.120809157 | 0.173295756 | 4.15E-11 | 5.17E-10 |
| Sdsl | 456.2238763 | 1.144650673 | 0.177617915 | 4.36E-11 | 5.43E-10 |
| Bcl9l | 2085.715665 | 1.047034324 | 0.16225093 | 4.47E-11 | 5.57E-10 |
| Fam131c | 129.5904495 | 1.494171255 | 0.234381066 | 4.60E-11 | 5.71E-10 |
| H2-Q5 | 311.7463863 | 1.491542463 | 0.234035268 | 4.63E-11 | 5.74E-10 |
| Gm45819 | 82.04792913 | 3.189830214 | 0.509479561 | 4.66E-11 | 5.78E-10 |
| Csnk1e | 773.958128 | 1.074707687 | 0.166726467 | 4.76E-11 | 5.90E-10 |
| Lmcd1 | 250.8541308 | 5.20931328 | 0.806036107 | 5.19E-11 | 6.40E-10 |
| Cchcr1 | 625.7546817 | 1.132003158 | 0.176356224 | 5.30E-11 | 6.54E-10 |
| Tpcn2 | 203.2300929 | 1.464675702 | 0.23033684 | 5.35E-11 | 6.59E-10 |
| Jrk | 125.6152616 | 1.781513381 | 0.28134357 | 5.38E-11 | 6.61E-10 |
| Tulp3 | 339.4172723 | 5.321162943 | 0.796582922 | 5.42E-11 | 6.66E-10 |
| Gm49172 | 178.1896256 | 3.135176596 | 0.490665667 | 5.52E-11 | 6.78E-10 |
| Snrpf | 163.218997 | 1.833368849 | 0.291343018 | 5.90E-11 | 7.23E-10 |
| Ephb6 | 336.9424549 | 3.022673894 | 0.476006376 | 6.06E-11 | 7.41E-10 |
| Dbf4 | 120.8693932 | 1.813587748 | 0.288508863 | 6.38E-11 | 7.79E-10 |
| Cdkn2c | 312.9823092 | 1.358814889 | 0.214454637 | 6.41E-11 | 7.82E-10 |

| | | | | | |
|---|---|---|---|---|---|
| Cav1 | 1004.066112 | 3.040962185 | 0.484336783 | 6.48E-11 | 7.90E-10 |
| Nfatc2 | 327.7043917 | 7.308322628 | 1.224124474 | 6.61E-11 | 8.04E-10 |
| Eps8l3 | 66.40454041 | 8.821691834 | 2.454717166 | 6.83E-11 | 8.31E-10 |
| Gpx7 | 100.0423599 | 1.691048634 | 0.268413386 | 7.47E-11 | 9.05E-10 |
| Fgf21 | 40.13117571 | 3.860698577 | 0.62677586 | 7.89E-11 | 9.53E-10 |
| Apbb1 | 340.3294945 | 3.610176224 | 0.565181044 | 7.97E-11 | 9.62E-10 |
| Sfn | 352.3405021 | 3.963015761 | 0.634177483 | 8.00E-11 | 9.65E-10 |
| Gabbr1 | 495.1106324 | 2.856661874 | 0.452631501 | 8.03E-11 | 9.67E-10 |
| Comtd1 | 59.46361853 | 2.570726588 | 0.416075375 | 8.07E-11 | 9.72E-10 |
| Prdm16 | 278.5639074 | 3.897578806 | 0.60827606 | 8.08E-11 | 9.72E-10 |
| Dsn1 | 106.9846128 | 2.206328295 | 0.352638828 | 8.82E-11 | 1.06E-09 |
| Ccdc82 | 468.2548225 | 1.0470342 | 0.164728752 | 8.83E-11 | 1.06E-09 |
| Dpysl2 | 148.3984993 | 2.394409612 | 0.384202671 | 9.04E-11 | 1.08E-09 |
| Sybu | 37.60756693 | 5.034609742 | 0.790208496 | 9.61E-11 | 1.15E-09 |
| Marchf9 | 316.2670096 | 1.993360951 | 0.318745554 | 9.94E-11 | 1.18E-09 |
| Rgs2 | 502.9169095 | 7.095573751 | 1.089254356 | 1.03E-10 | 1.23E-09 |
| H2-Q2 | 124.1725493 | 3.669721921 | 0.581715287 | 1.09E-10 | 1.30E-09 |
| Cdc20 | 133.4927589 | 3.402912703 | 0.553887809 | 1.10E-10 | 1.30E-09 |
| Tmem229a | 684.456143 | 7.860531943 | 2.209715479 | 1.14E-10 | 1.36E-09 |
| Glipr2 | 242.2080239 | 3.277315653 | 0.533939217 | 1.15E-10 | 1.37E-09 |
| Sytl5 | 513.7245561 | 7.227549904 | 1.227715004 | 1.19E-10 | 1.41E-09 |
| Rdh9 | 144.3379357 | 1.772722975 | 0.287500498 | 1.27E-10 | 1.50E-09 |
| Eif4e3 | 274.3497036 | 1.8954037 | 0.308814547 | 1.32E-10 | 1.56E-09 |
| Tspan4 | 886.0963286 | 1.08119825 | 0.17217918 | 1.33E-10 | 1.56E-09 |
| Rbm3 | 1762.268476 | 1.348716725 | 0.216911684 | 1.37E-10 | 1.61E-09 |
| Bex3 | 306.8733245 | 3.723316377 | 0.594707423 | 1.37E-10 | 1.61E-09 |
| Pprc1 | 570.1787801 | 1.403734303 | 0.226495909 | 1.48E-10 | 1.73E-09 |
| Cxcl14 | 137.5969544 | 2.430300815 | 0.401219173 | 1.58E-10 | 1.84E-09 |
| Timp3 | 835.1460626 | 2.688924084 | 0.443664895 | 1.71E-10 | 1.99E-09 |
| Rhoq | 728.9108823 | 1.102072961 | 0.176832204 | 1.79E-10 | 2.08E-09 |
| Ift43 | 203.7368131 | 1.516401515 | 0.24541028 | 1.79E-10 | 2.08E-09 |
| Lrrc45 | 516.0403133 | 1.079780555 | 0.173351956 | 1.86E-10 | 2.16E-09 |
| Ccdc57 | 205.859654 | 1.253392004 | 0.201928563 | 1.89E-10 | 2.19E-09 |
| Cyp2a5 | 16867.11266 | 1.475141434 | 0.239877959 | 1.90E-10 | 2.20E-09 |
| Msantd3 | 155.5618807 | 7.102979291 | 1.224326422 | 1.97E-10 | 2.28E-09 |
| Baiap2l2 | 96.56837681 | 7.974499056 | 2.306392867 | 2.02E-10 | 2.33E-09 |
| Cdh17 | 307.0721841 | 7.903678765 | 2.254752119 | 2.09E-10 | 2.41E-09 |
| Rasl10b | 46.12216366 | 3.396238993 | 0.566703875 | 2.13E-10 | 2.44E-09 |
| Ifngr1 | 1341.758505 | 1.259200985 | 0.2037477 | 2.13E-10 | 2.45E-09 |
| Col16a1 | 1913.69806 | 5.318405537 | 0.833094537 | 2.25E-10 | 2.57E-09 |
| Srd5a3 | 345.3488126 | 1.026748029 | 0.165433597 | 2.34E-10 | 2.67E-09 |
| Crip1 | 289.1873147 | 5.412603141 | 0.853545344 | 2.41E-10 | 2.74E-09 |
| Col5a1 | 79.65745786 | 4.293633439 | 0.702054912 | 2.49E-10 | 2.82E-09 |

| | | | | | |
|---|---|---|---|---|---|
| Kcnj8 | 108.4920928 | 5.22573058 | 0.817825576 | 2.52E-10 | 2.87E-09 |
| Ifrd1 | 911.3777563 | 1.68362181 | 0.27838932 | 2.68E-10 | 3.03E-09 |
| Cdt1 | 223.480793 | 1.385042377 | 0.226569178 | 2.69E-10 | 3.04E-09 |
| Dapp1 | 411.7185136 | 6.17312927 | 0.937658028 | 2.73E-10 | 3.09E-09 |
| Srxn1 | 2117.593055 | 1.14585438 | 0.186174206 | 2.74E-10 | 3.10E-09 |
| Usp35 | 163.7154416 | 1.673264214 | 0.274706143 | 2.75E-10 | 3.10E-09 |
| Btc | 328.365126 | 1.454405186 | 0.237725209 | 2.80E-10 | 3.16E-09 |
| Racgap1 | 167.0399256 | 3.305965073 | 0.552806968 | 2.83E-10 | 3.19E-09 |
| Zfp791 | 86.93940525 | 1.582360176 | 0.260002829 | 2.94E-10 | 3.31E-09 |
| Timp1 | 96.65089006 | 7.964159145 | 2.311178285 | 2.95E-10 | 3.32E-09 |
| Grk3 | 432.1468736 | 1.16698008 | 0.190578456 | 3.18E-10 | 3.55E-09 |
| Cdca3 | 109.9265191 | 2.515626774 | 0.423534952 | 3.31E-10 | 3.68E-09 |
| A330074K22Rik | 125.2870763 | 7.785046189 | 2.272554832 | 3.42E-10 | 3.80E-09 |
| Syde1 | 471.9481987 | 6.151342545 | 0.939992732 | 3.46E-10 | 3.84E-09 |
| Atp6v0d2 | 83.47036861 | 2.373746714 | 0.3998361 | 3.47E-10 | 3.85E-09 |
| Ccn2 | 5388.782483 | 6.962097689 | 1.205158124 | 3.52E-10 | 3.90E-09 |
| Phactr2 | 475.4612223 | 1.278472553 | 0.209478388 | 3.54E-10 | 3.92E-09 |
| Pls1 | 575.5978219 | 1.0979144 | 0.179407933 | 3.60E-10 | 3.98E-09 |
| Gm20008 | 148.508298 | 1.890463673 | 0.315266496 | 3.61E-10 | 4.00E-09 |
| Sptlc2 | 1670.012782 | 1.088715269 | 0.177874813 | 3.62E-10 | 4.01E-09 |
| Ap3m2 | 167.7451292 | 1.90531106 | 0.315569259 | 3.66E-10 | 4.04E-09 |
| Lgmn | 1085.643284 | 1.499850032 | 0.248603896 | 3.68E-10 | 4.06E-09 |
| Top2a | 483.1935635 | 4.006513094 | 0.686057692 | 3.71E-10 | 4.09E-09 |
| Cep192 | 492.0943633 | 1.197429718 | 0.196724864 | 4.00E-10 | 4.40E-09 |
| Zfp651 | 409.0902105 | 3.977342632 | 0.641495264 | 4.11E-10 | 4.51E-09 |
| Lipn | 34.2203007 | 8.399919902 | 2.499241632 | 4.17E-10 | 4.56E-09 |
| Hid1 | 442.212484 | 3.700815549 | 0.632114235 | 4.21E-10 | 4.60E-09 |
| Samd1 | 361.15989 | 1.182504675 | 0.194910322 | 4.37E-10 | 4.77E-09 |
| Dsg1c | 553.2878547 | 2.782065038 | 0.475841261 | 4.44E-10 | 4.84E-09 |
| Baz1a | 555.3098189 | 1.112946318 | 0.1828369 | 4.65E-10 | 5.05E-09 |
| Foxm1 | 141.6650028 | 4.740969834 | 0.772900801 | 4.89E-10 | 5.29E-09 |
| Cdc25a | 362.6038333 | 1.007835625 | 0.165344449 | 4.96E-10 | 5.37E-09 |
| Jade2 | 953.8595446 | 1.712156335 | 0.288721838 | 5.08E-10 | 5.49E-09 |
| Ciart | 136.3040224 | 2.905502758 | 0.500185142 | 5.53E-10 | 5.95E-09 |
| Krt8-ps | 71.82928407 | 1.529486908 | 0.255806433 | 5.96E-10 | 6.41E-09 |
| Uhrf1 | 378.206148 | 4.951064796 | 0.854853891 | 6.00E-10 | 6.44E-09 |
| Slc5a9 | 47.01619117 | 7.781077733 | 2.322663943 | 6.05E-10 | 6.48E-09 |
| Hcn3 | 136.3199744 | 1.380371401 | 0.231333827 | 6.49E-10 | 6.95E-09 |
| Rassf1 | 204.8080249 | 1.248318574 | 0.207691604 | 6.55E-10 | 7.01E-09 |
| Col3a1 | 373.5077786 | 4.848008469 | 0.85391695 | 6.83E-10 | 7.29E-09 |
| Coq8b | 575.2329452 | 1.017488593 | 0.168674992 | 6.97E-10 | 7.43E-09 |
| Gm16505 | 92.69693653 | 7.902156887 | 2.323500362 | 7.46E-10 | 7.92E-09 |
| Adap1 | 133.0891506 | 3.493769474 | 0.583072886 | 7.56E-10 | 8.03E-09 |

| | | | | | |
|---|---|---|---|---|---|
| Csn3 | 23.81410256 | 8.359174399 | 2.527512328 | 8.37E-10 | 8.84E-09 |
| Actg1 | 14618.63977 | 1.295591435 | 0.219214096 | 8.72E-10 | 9.20E-09 |
| Angptl2 | 57.6142951 | 3.523627929 | 0.59771543 | 8.76E-10 | 9.25E-09 |
| Timp2 | 2446.47702 | 3.554942833 | 0.612963769 | 8.81E-10 | 9.29E-09 |
| Cit | 163.2408144 | 6.084396343 | 0.960767602 | 8.92E-10 | 9.39E-09 |
| Gas8 | 300.9570558 | 1.343207404 | 0.226409954 | 9.06E-10 | 9.53E-09 |
| Ccnd1 | 2705.489028 | 1.714465814 | 0.293556489 | 9.34E-10 | 9.82E-09 |
| Tprn | 347.3203135 | 1.286483145 | 0.217176249 | 9.38E-10 | 9.85E-09 |
| Nrp2 | 304.8150418 | 1.39995417 | 0.23646924 | 9.45E-10 | 9.92E-09 |
| Heg1 | 3677.045021 | 1.412927446 | 0.24003737 | 9.89E-10 | 1.04E-08 |
| AA986860 | 446.68068 | 3.166529081 | 0.557446514 | 1.00E-09 | 1.05E-08 |
| Ift122 | 507.7082696 | 1.031284792 | 0.172676992 | 1.08E-09 | 1.12E-08 |
| Ildr2 | 1117.511288 | 1.23310175 | 0.208890304 | 1.13E-09 | 1.18E-08 |
| Wnt4 | 683.4328774 | 5.966623124 | 0.94489678 | 1.21E-09 | 1.25E-08 |
| Zfp90 | 224.9905554 | 1.795027893 | 0.308107943 | 1.33E-09 | 1.38E-08 |
| Mcm3 | 349.9815276 | 1.886556264 | 0.327896287 | 1.41E-09 | 1.45E-08 |
| Gm47283 | 340.8689251 | 1.273843174 | 0.217686942 | 1.46E-09 | 1.50E-08 |
| 4930503L19Rik | 179.7783678 | 3.278401566 | 0.561170547 | 1.47E-09 | 1.51E-08 |
| Tmem71 | 118.1168895 | 1.720382647 | 0.298392941 | 1.56E-09 | 1.60E-08 |
| Gm16559 | 139.9860421 | 2.018156177 | 0.353273559 | 1.58E-09 | 1.62E-08 |
| Icosl | 55.80391743 | 2.667548595 | 0.471736772 | 1.70E-09 | 1.73E-08 |
| Afap1 | 518.9602312 | 4.24062621 | 0.705228652 | 1.71E-09 | 1.74E-08 |
| Elapor1 | 640.0389491 | 1.896868245 | 0.331089805 | 1.72E-09 | 1.75E-08 |
| Styk1 | 38.49326863 | 7.73363198 | 2.35315541 | 1.75E-09 | 1.78E-08 |
| CAAA01147332.1 | 282.2392826 | 1.958146069 | 0.343553841 | 1.77E-09 | 1.80E-08 |
| Cerk | 281.5153262 | 1.966432793 | 0.339976372 | 1.78E-09 | 1.80E-08 |
| Trpm4 | 296.6896639 | 2.556860285 | 0.44642365 | 1.84E-09 | 1.87E-08 |
| Tchp | 218.0761285 | 1.39568539 | 0.239854479 | 1.86E-09 | 1.89E-08 |
| Emp1 | 41.72783695 | 7.822141134 | 2.362101541 | 1.88E-09 | 1.90E-08 |
| Trpv4 | 353.5537042 | 7.50176785 | 2.237050229 | 1.89E-09 | 1.91E-08 |
| Gm49599 | 86.69629351 | 1.831574884 | 0.318687147 | 1.89E-09 | 1.91E-08 |
| Zfp28 | 153.3318207 | 1.857848205 | 0.32376935 | 2.00E-09 | 2.01E-08 |
| Fhl3 | 270.4051534 | 6.748173865 | 1.239932795 | 2.03E-09 | 2.04E-08 |
| Lrp8 | 50.6908175 | 5.021881166 | 0.837078462 | 2.07E-09 | 2.08E-08 |
| Taf1d | 544.3843068 | 1.212172664 | 0.209015097 | 2.08E-09 | 2.09E-08 |
| Npc1l1 | 24.68061548 | 7.696588746 | 2.372063678 | 2.10E-09 | 2.10E-08 |
| Bmyc | 120.5819691 | 2.003104071 | 0.348241175 | 2.13E-09 | 2.13E-08 |
| Gm42892 | 122.7300029 | 2.035904155 | 0.356371041 | 2.14E-09 | 2.13E-08 |
| Clcn5 | 442.4630899 | 1.070898514 | 0.183598318 | 2.18E-09 | 2.18E-08 |
| Fosl2 | 1002.982682 | 1.448841275 | 0.252296472 | 2.27E-09 | 2.26E-08 |
| Kctd12 | 316.8826791 | 1.256036186 | 0.217002904 | 2.40E-09 | 2.38E-08 |
| Rnf145 | 257.9462439 | 1.246347952 | 0.215827478 | 2.43E-09 | 2.41E-08 |
| Adgrg1 | 2979.999731 | 1.213633239 | 0.209602197 | 2.44E-09 | 2.42E-08 |

| | | | | | |
|---|---|---|---|---|---|
| Gbp8 | 195.621099 | 2.231227894 | 0.391121725 | 2.57E-09 | 2.54E-08 |
| Cers6 | 759.3318803 | 1.470205147 | 0.258581005 | 2.78E-09 | 2.73E-08 |
| Tmem86a | 403.5285835 | 3.079321767 | 0.559883227 | 2.84E-09 | 2.79E-08 |
| Tle6 | 219.2206023 | 1.602653819 | 0.281056618 | 2.85E-09 | 2.79E-08 |
| Fam83a | 45.01744728 | 2.717663183 | 0.489039279 | 2.96E-09 | 2.89E-08 |
| Osbpl10 | 69.4691096 | 7.52703269 | 2.296801331 | 3.02E-09 | 2.95E-08 |
| Evc | 678.8361527 | 1.818137863 | 0.322330442 | 3.42E-09 | 3.32E-08 |
| Smad6 | 231.3298228 | 1.335425952 | 0.234861498 | 3.78E-09 | 3.66E-08 |
| Dnttip1 | 433.5773599 | 1.003711247 | 0.174487433 | 3.88E-09 | 3.75E-08 |
| Col6a2 | 77.93271812 | 3.047846075 | 0.547570472 | 4.02E-09 | 3.88E-08 |
| Irx3 | 142.4074215 | 3.23355851 | 0.58929251 | 4.08E-09 | 3.93E-08 |
| Mcm5 | 323.9799471 | 2.368238768 | 0.430425769 | 4.32E-09 | 4.16E-08 |
| Ctnnal1 | 289.8493947 | 1.20037026 | 0.210985147 | 4.48E-09 | 4.31E-08 |
| Il1r1 | 1044.662453 | 1.234362472 | 0.218001548 | 4.60E-09 | 4.41E-08 |
| Hspa1b | 1231.268256 | 1.750587224 | 0.316109388 | 5.16E-09 | 4.90E-08 |
| Scai | 499.9084881 | 1.105576612 | 0.194783123 | 5.29E-09 | 5.01E-08 |
| Rcan3 | 288.8356164 | 3.198483721 | 0.565109106 | 5.86E-09 | 5.54E-08 |
| Acot3 | 44.61349219 | 2.540078691 | 0.464866717 | 6.08E-09 | 5.73E-08 |
| Tacc3 | 166.9694321 | 1.828869275 | 0.329805099 | 6.08E-09 | 5.73E-08 |
| C2cd4c | 122.5220656 | 6.467225411 | 1.229729411 | 6.13E-09 | 5.77E-08 |
| Phgdh | 206.8520836 | 4.661905922 | 0.797399526 | 6.81E-09 | 6.38E-08 |
| Klhl13 | 455.4124823 | 1.173876326 | 0.209167462 | 6.89E-09 | 6.45E-08 |
| Tuba1a | 638.7868892 | 2.441128578 | 0.441798078 | 6.91E-09 | 6.46E-08 |
| Fbn1 | 40.7176792 | 4.058631318 | 0.751045631 | 8.41E-09 | 7.79E-08 |
| Fosb | 3546.966915 | 4.131943617 | 0.789453147 | 8.42E-09 | 7.80E-08 |
| CT010467.1 | 1492.916346 | 2.87244282 | 0.540332501 | 8.50E-09 | 7.86E-08 |
| Il34 | 213.3188316 | 6.517460726 | 1.255391162 | 9.02E-09 | 8.32E-08 |
| Nrxn2 | 33.09278625 | 4.431649255 | 0.824590045 | 9.13E-09 | 8.41E-08 |
| Ccne1 | 49.27501703 | 2.303241181 | 0.423126327 | 9.18E-09 | 8.45E-08 |
| Acnat2 | 1061.621396 | 1.417594425 | 0.257184557 | 9.21E-09 | 8.48E-08 |
| Tpx2 | 124.6648931 | 4.58273023 | 0.827404269 | 9.25E-09 | 8.52E-08 |
| 1810064F22Rik | 163.7644254 | 1.718834233 | 0.314828284 | 9.27E-09 | 8.53E-08 |
| Mlkl | 292.4996137 | 1.125287891 | 0.202525795 | 1.00E-08 | 9.19E-08 |
| Igfbp6 | 53.5334736 | 7.139954661 | 2.273885654 | 1.02E-08 | 9.35E-08 |
| Kif5c | 79.68858742 | 7.127334382 | 2.244175972 | 1.03E-08 | 9.46E-08 |
| Arl10 | 177.8805636 | 1.143709676 | 0.205655413 | 1.04E-08 | 9.50E-08 |
| Tead4 | 177.9536918 | 6.939960359 | 2.173175355 | 1.10E-08 | 1.00E-07 |
| Adarb1 | 407.6972947 | 1.103252161 | 0.198812778 | 1.10E-08 | 1.01E-07 |
| Sap30 | 117.8815048 | 1.294601031 | 0.234482141 | 1.11E-08 | 1.01E-07 |
| Gab2 | 171.2554556 | 3.019303413 | 0.548520646 | 1.13E-08 | 1.03E-07 |
| Gas2l1 | 1124.24699 | 1.100628635 | 0.198905854 | 1.14E-08 | 1.04E-07 |
| Fstl1 | 360.0454495 | 5.716005802 | 0.981557518 | 1.18E-08 | 1.07E-07 |
| Plat | 563.8693799 | 7.06459065 | 2.187768168 | 1.18E-08 | 1.07E-07 |

| | | | | | |
|---|---|---|---|---|---|
| Chrnb1 | 999.15239 | 6.24009944 | 1.202616745 | 1.20E-08 | 1.09E-07 |
| Zfp251 | 198.7566525 | 1.486118771 | 0.270778644 | 1.23E-08 | 1.12E-07 |
| Zdhhc8 | 586.9771956 | 1.027586622 | 0.185850464 | 1.26E-08 | 1.14E-07 |
| Serpinh1 | 630.7138437 | 2.38677323 | 0.444386775 | 1.29E-08 | 1.16E-07 |
| Cfap69 | 284.8246777 | 3.985429599 | 0.706291958 | 1.29E-08 | 1.17E-07 |
| Pgap4 | 192.0518371 | 6.931632728 | 2.184489631 | 1.30E-08 | 1.17E-07 |
| Elf4 | 469.0925564 | 3.980524912 | 0.705250893 | 1.31E-08 | 1.18E-07 |
| Bgn | 567.8177548 | 3.130807726 | 0.600073643 | 1.34E-08 | 1.20E-07 |
| C230037L18Rik | 28.31317739 | 3.653558701 | 0.670039307 | 1.34E-08 | 1.21E-07 |
| Abr | 933.1223191 | 1.322727226 | 0.241227333 | 1.35E-08 | 1.21E-07 |
| C1qtnf1 | 105.6272893 | 1.798016656 | 0.332669027 | 1.39E-08 | 1.24E-07 |
| AI314278 | 17.25479758 | 7.59677328 | 2.433145305 | 1.40E-08 | 1.25E-07 |
| Fbn2 | 20.35124697 | 7.514174945 | 2.402986024 | 1.47E-08 | 1.31E-07 |
| Asb11 | 17.31992749 | 7.639602063 | 2.460162984 | 1.53E-08 | 1.36E-07 |
| Zfp984 | 144.0600448 | 1.572075644 | 0.291177167 | 1.53E-08 | 1.37E-07 |
| Mt1 | 3292.755801 | 3.36226035 | 0.649187764 | 1.55E-08 | 1.38E-07 |
| Cenpt | 190.8069437 | 1.507948342 | 0.278730559 | 1.56E-08 | 1.39E-07 |
| Kntc1 | 117.5399113 | 7.376013393 | 2.322164615 | 1.72E-08 | 1.52E-07 |
| Mrc2 | 22.41579253 | 7.291806068 | 2.350416042 | 1.78E-08 | 1.58E-07 |
| Dapk2 | 189.1217962 | 1.397756836 | 0.259656524 | 1.88E-08 | 1.66E-07 |
| Plaur | 449.2057924 | 5.527077153 | 0.957556644 | 1.93E-08 | 1.69E-07 |
| Itga2 | 1252.831031 | 3.311275372 | 0.61594128 | 1.93E-08 | 1.69E-07 |
| Anln | 148.3797662 | 4.075848628 | 0.740664493 | 2.10E-08 | 1.83E-07 |
| Sparcl1 | 46.78317022 | 6.654295351 | 1.33233663 | 2.16E-08 | 1.89E-07 |
| Gm9844 | 118.5485654 | 6.880603531 | 2.208200567 | 2.18E-08 | 1.90E-07 |
| Ccnb1 | 77.85568271 | 7.549456781 | 2.374418192 | 2.21E-08 | 1.93E-07 |
| Slc25a27 | 153.1267644 | 2.034625069 | 0.383336508 | 2.24E-08 | 1.96E-07 |
| Grhl1 | 84.43651385 | 1.737734247 | 0.328173595 | 2.29E-08 | 2.00E-07 |
| Hbegf | 1056.484044 | 3.500870582 | 0.687566095 | 2.32E-08 | 2.02E-07 |
| Sema4c | 375.3262674 | 2.15164622 | 0.404274078 | 2.36E-08 | 2.05E-07 |
| Bmp8b | 48.836341 | 2.901572583 | 0.54400262 | 2.38E-08 | 2.07E-07 |
| Ltbp3 | 2784.312219 | 3.151670151 | 0.588911876 | 2.45E-08 | 2.12E-07 |
| Ust | 125.2792792 | 6.794974917 | 2.19431097 | 2.48E-08 | 2.14E-07 |
| Chml | 189.291024 | 1.095073286 | 0.20305718 | 2.55E-08 | 2.19E-07 |
| Samd10 | 337.0175874 | 1.210618677 | 0.222325821 | 2.60E-08 | 2.24E-07 |
| Ptgds | 115.6210816 | 2.289328045 | 0.436580625 | 2.69E-08 | 2.31E-07 |
| Incenp | 238.8731454 | 1.739904053 | 0.330862405 | 2.73E-08 | 2.34E-07 |
| Mllt3 | 244.0864129 | 1.223670007 | 0.228955269 | 2.73E-08 | 2.34E-07 |
| 2700081O15Rik | 611.4329317 | 5.404332328 | 0.943055019 | 2.75E-08 | 2.36E-07 |
| Mansc1 | 327.2258063 | 1.439937087 | 0.270636576 | 2.81E-08 | 2.41E-07 |
| Slc51b | 20.9999369 | 4.21211864 | 0.782381579 | 2.81E-08 | 2.41E-07 |
| Hmmr | 76.57121725 | 4.896085568 | 0.902497931 | 2.83E-08 | 2.42E-07 |
| Abcc9 | 115.2394672 | 2.316644214 | 0.444226049 | 3.11E-08 | 2.65E-07 |

| | | | | | |
|---|---|---|---|---|---|
| Cmtm7 | 121.3594006 | 2.676241441 | 0.509864618 | 3.25E-08 | 2.75E-07 |
| Kif22 | 82.95101749 | 2.361109716 | 0.452050077 | 3.26E-08 | 2.76E-07 |
| Junb | 9983.881613 | 1.043911932 | 0.19594229 | 3.41E-08 | 2.88E-07 |
| Prrg1 | 122.3684907 | 1.760140088 | 0.333345758 | 3.41E-08 | 2.89E-07 |
| Tsku | 2146.539779 | 1.636502397 | 0.313448814 | 3.58E-08 | 3.02E-07 |
| Adamts14 | 236.3056715 | 4.401431761 | 0.798145314 | 3.88E-08 | 3.25E-07 |
| Adamts2 | 80.26977386 | 3.413832767 | 0.65656357 | 4.02E-08 | 3.37E-07 |
| Loxl1 | 116.9155828 | 6.885479939 | 2.238636539 | 4.13E-08 | 3.44E-07 |
| Lrrc1 | 276.4700916 | 2.802733493 | 0.532418406 | 4.63E-08 | 3.84E-07 |
| Ckap2l | 65.8260631 | 6.896433525 | 2.266351276 | 4.83E-08 | 3.99E-07 |
| Dnase1l1 | 162.9690528 | 1.02476575 | 0.193101733 | 5.04E-08 | 4.15E-07 |
| Mmp2 | 40.60420078 | 4.557750416 | 0.896028938 | 5.16E-08 | 4.24E-07 |
| Gm50475 | 47.08808995 | 2.819677072 | 0.556813122 | 5.33E-08 | 4.38E-07 |
| Olfr1372-ps1 | 1136.60357 | 3.569965703 | 0.678724906 | 5.56E-08 | 4.55E-07 |
| Mmp24 | 104.7642708 | 6.685015543 | 2.208798232 | 5.88E-08 | 4.80E-07 |
| Gins2 | 77.07097145 | 3.250999347 | 0.621942447 | 6.02E-08 | 4.90E-07 |
| Mvb12b | 509.0379139 | 1.136112925 | 0.216262508 | 6.12E-08 | 4.97E-07 |
| Klf4 | 96.95380269 | 4.486096548 | 0.83764774 | 6.12E-08 | 4.97E-07 |
| Capn5 | 751.634702 | 1.105607076 | 0.21139209 | 6.13E-08 | 4.97E-07 |
| Areg | 17.28864923 | 3.531640551 | 0.69518728 | 6.17E-08 | 5.01E-07 |
| Ptpre | 506.8055087 | 1.368809114 | 0.264388226 | 6.27E-08 | 5.08E-07 |
| Ifnlr1 | 119.579872 | 4.333658598 | 0.802091167 | 6.51E-08 | 5.26E-07 |
| Necap2 | 440.021486 | 1.049215578 | 0.200135253 | 6.58E-08 | 5.31E-07 |
| Pfkfb2 | 390.6373624 | 1.134220401 | 0.217458614 | 6.63E-08 | 5.34E-07 |
| Ccna2 | 142.806624 | 3.066567531 | 0.614280122 | 6.95E-08 | 5.58E-07 |
| Il1rn | 1510.476814 | 1.433600222 | 0.278876903 | 7.27E-08 | 5.83E-07 |
| Map2k6 | 324.2190256 | 1.035894431 | 0.199252118 | 7.55E-08 | 6.04E-07 |
| Lamc2 | 1703.188094 | 6.429480123 | 2.104919827 | 8.17E-08 | 6.52E-07 |
| Eif4ebp1 | 774.164529 | 1.182089283 | 0.22898844 | 8.74E-08 | 6.97E-07 |
| Mapk4 | 63.89398407 | 2.207414274 | 0.436301932 | 9.37E-08 | 7.44E-07 |
| Gm609 | 1525.888356 | 5.20713071 | 0.953000926 | 9.39E-08 | 7.45E-07 |
| Snhg18 | 191.3589224 | 2.288502093 | 0.451047654 | 9.45E-08 | 7.50E-07 |
| Cx3cr1 | 20.23214343 | 4.665202892 | 0.89969161 | 9.47E-08 | 7.51E-07 |
| Dcbld1 | 251.2185711 | 1.187531944 | 0.230675045 | 9.53E-08 | 7.55E-07 |
| Diaph3 | 65.97491546 | 3.338215741 | 0.662727536 | 9.60E-08 | 7.60E-07 |
| Gm13387 | 100.6394888 | 6.518476087 | 2.168480682 | 9.64E-08 | 7.63E-07 |
| Zbtb12 | 225.3564426 | 1.3595333 | 0.266619331 | 9.73E-08 | 7.69E-07 |
| Igfbp7 | 7378.973716 | 1.60169106 | 0.319100854 | 9.91E-08 | 7.82E-07 |
| Gm31522 | 31.12292373 | 2.482419309 | 0.50429319 | 1.04E-07 | 8.16E-07 |
| Kif20a | 89.27884143 | 2.387981151 | 0.481072953 | 1.08E-07 | 8.52E-07 |
| 4933431E20Rik | 482.1316081 | 1.006471041 | 0.19517724 | 1.11E-07 | 8.70E-07 |
| Mogat2 | 53.67526517 | 2.310425265 | 0.465751729 | 1.13E-07 | 8.85E-07 |
| Slc24a3 | 37.75884176 | 4.365768009 | 0.833642101 | 1.13E-07 | 8.86E-07 |

| | | | | | |
|---|---|---|---|---|---|
| Ephb2 | 132.6122714 | 6.540414218 | 2.204565122 | 1.13E-07 | 8.89E-07 |
| Septin8 | 893.075375 | 2.770334848 | 0.55304064 | 1.16E-07 | 9.07E-07 |
| Adamtsl2 | 34.3794196 | 5.608688058 | 1.071589266 | 1.27E-07 | 9.85E-07 |
| Prc1 | 183.748832 | 4.208209551 | 0.866441091 | 1.29E-07 | 1.00E-06 |
| Dusp5 | 545.1608575 | 4.613140124 | 0.984041488 | 1.31E-07 | 1.01E-06 |
| Bcl2 | 255.0492226 | 5.113148082 | 0.945954379 | 1.32E-07 | 1.02E-06 |
| Gm42679 | 48.17681847 | 6.694781277 | 2.263129577 | 1.34E-07 | 1.04E-06 |
| Pbx4 | 56.05103632 | 2.164990974 | 0.438871015 | 1.34E-07 | 1.04E-06 |
| Serpinb6b | 1144.125425 | 2.220252349 | 0.457930102 | 1.40E-07 | 1.09E-06 |
| Kif20b | 118.4769305 | 2.486285275 | 0.506859302 | 1.43E-07 | 1.11E-06 |
| Ttc12 | 183.9360617 | 2.670952844 | 0.528140216 | 1.43E-07 | 1.11E-06 |
| Sema5a | 1725.479122 | 5.154471542 | 0.96545607 | 1.53E-07 | 1.18E-06 |
| Fbxw17 | 162.9102618 | 5.053957283 | 0.937555151 | 1.53E-07 | 1.18E-06 |
| Gm49405 | 56.69017637 | 2.280571317 | 0.458809115 | 1.54E-07 | 1.19E-06 |
| Mis18a | 82.19128069 | 1.473937871 | 0.294718737 | 1.56E-07 | 1.20E-06 |
| Fgr | 115.1481992 | 2.023874308 | 0.407295338 | 1.61E-07 | 1.24E-06 |
| Parm1 | 343.1141576 | 5.032063006 | 0.93475696 | 1.63E-07 | 1.26E-06 |
| Syn2 | 50.11048499 | 6.563305452 | 2.247170584 | 1.69E-07 | 1.29E-06 |
| Ccl7 | 119.4539432 | 6.641701519 | 2.237663793 | 1.69E-07 | 1.29E-06 |
| Inpp5j | 66.75223651 | 5.751795815 | 1.230111497 | 1.76E-07 | 1.34E-06 |
| Crym | 115.0287656 | 1.45736888 | 0.293844989 | 1.86E-07 | 1.42E-06 |
| Calml4 | 19.65055435 | 6.6560405 | 2.299605264 | 1.87E-07 | 1.42E-06 |
| Rasgrp3 | 69.87697963 | 1.929287025 | 0.394001144 | 1.90E-07 | 1.44E-06 |
| Card14 | 29.6980435 | 3.18483262 | 0.640362849 | 1.90E-07 | 1.45E-06 |
| Tmem159 | 169.569811 | 1.495855019 | 0.301108491 | 1.92E-07 | 1.46E-06 |
| A930004J17Rik | 117.7716395 | 1.115000009 | 0.22184277 | 1.93E-07 | 1.46E-06 |
| Bex2 | 29.23493238 | 4.040013231 | 0.812836954 | 1.96E-07 | 1.48E-06 |
| Tmem253 | 45.00653487 | 1.84941934 | 0.378567127 | 1.98E-07 | 1.50E-06 |
| Iqgap3 | 168.6417664 | 6.277448294 | 1.369790988 | 1.99E-07 | 1.51E-06 |
| Il17rd | 376.2844544 | 2.837973601 | 0.568774882 | 2.00E-07 | 1.52E-06 |
| Rpl38 | 1555.587897 | 1.051516486 | 0.209597223 | 2.12E-07 | 1.59E-06 |
| Ninl | 391.144416 | 1.749573211 | 0.354556917 | 2.15E-07 | 1.62E-06 |
| Kifc5b | 55.79256431 | 2.777947365 | 0.573442202 | 2.21E-07 | 1.66E-06 |
| Slc52a3 | 240.9556715 | 4.977133505 | 0.937232689 | 2.24E-07 | 1.68E-06 |
| Slc7a1 | 304.9468346 | 3.635141273 | 0.717820085 | 2.26E-07 | 1.69E-06 |
| 2310034O05Rik | 18.68221474 | 3.667325731 | 0.766960857 | 2.27E-07 | 1.70E-06 |
| Kctd10 | 489.3811112 | 1.155985559 | 0.232497665 | 2.33E-07 | 1.74E-06 |
| Nmrk1 | 366.0884245 | 1.180117305 | 0.237927252 | 2.36E-07 | 1.76E-06 |
| Il7 | 63.06247507 | 2.840581078 | 0.574013927 | 2.41E-07 | 1.80E-06 |
| Igf2bp2 | 124.9533154 | 1.784463775 | 0.369256773 | 2.49E-07 | 1.85E-06 |
| Rad51 | 87.02254047 | 2.76109372 | 0.575642957 | 2.49E-07 | 1.85E-06 |
| Ifit2 | 209.9903753 | 1.201110615 | 0.242340531 | 2.72E-07 | 2.01E-06 |
| Mpp2 | 194.2380619 | 6.269401943 | 2.156360812 | 2.78E-07 | 2.05E-06 |

| | | | | | |
|---|---|---|---|---|---|
| Tceal5 | 37.35941911 | 3.841812315 | 0.780259814 | 2.91E-07 | 2.15E-06 |
| Gm38357 | 39.04917212 | 3.415816085 | 0.735360887 | 2.92E-07 | 2.15E-06 |
| Nudt18 | 428.7179947 | 1.098812179 | 0.222137141 | 2.92E-07 | 2.15E-06 |
| Kif4 | 58.56447958 | 6.73298051 | 2.314255068 | 3.05E-07 | 2.24E-06 |
| Sirt6 | 243.4401286 | 1.169449432 | 0.237695918 | 3.15E-07 | 2.31E-06 |
| Paqr4 | 174.4009437 | 2.456454152 | 0.502182911 | 3.16E-07 | 2.32E-06 |
| Gspt2 | 109.016343 | 1.635850026 | 0.336359836 | 3.20E-07 | 2.34E-06 |
| Flvcr2 | 109.0095871 | 1.571334406 | 0.327149548 | 3.21E-07 | 2.35E-06 |
| Olfml3 | 24.33583492 | 5.85521851 | 1.295637154 | 3.23E-07 | 2.36E-06 |
| Pdzk1ip1 | 816.5625522 | 1.273689167 | 0.26107294 | 3.23E-07 | 2.36E-06 |
| Plcd3 | 694.4379766 | 4.926945034 | 0.944047585 | 3.24E-07 | 2.36E-06 |
| Stambpl1 | 107.1529584 | 2.612150789 | 0.541877776 | 3.33E-07 | 2.43E-06 |
| Coro6 | 147.3621 | 5.604309865 | 1.22864057 | 3.38E-07 | 2.46E-06 |
| Rasgrf2 | 28.76391353 | 3.235811292 | 0.671442935 | 3.44E-07 | 2.50E-06 |
| Cnbd2 | 79.13870619 | 3.313549832 | 0.705481195 | 3.47E-07 | 2.52E-06 |
| Eid2 | 118.0403526 | 2.594163664 | 0.541707494 | 3.49E-07 | 2.53E-06 |
| Prelid2 | 35.53598776 | 3.382396595 | 0.710092059 | 3.67E-07 | 2.66E-06 |
| Thbs2 | 26.43254022 | 6.446489606 | 2.26873503 | 3.76E-07 | 2.72E-06 |
| Cd63-ps | 16.8973978 | 6.551076311 | 2.30878993 | 3.87E-07 | 2.80E-06 |
| Ddah2 | 682.6662607 | 4.029150719 | 0.802094475 | 4.08E-07 | 2.93E-06 |
| Dlgap5 | 59.25108218 | 6.654183735 | 2.307358095 | 4.13E-07 | 2.97E-06 |
| Mroh3 | 28.90483728 | 6.419422108 | 2.262702434 | 4.44E-07 | 3.18E-06 |
| Dcdc2a | 2247.349072 | 1.811259525 | 0.378209745 | 4.59E-07 | 3.28E-06 |
| Cbx6 | 2013.689666 | 1.752295449 | 0.369299044 | 4.63E-07 | 3.31E-06 |
| Dok4 | 84.57630924 | 2.768796871 | 0.575652295 | 4.65E-07 | 3.32E-06 |
| Tctn3 | 174.3446301 | 1.490256308 | 0.310599701 | 4.89E-07 | 3.48E-06 |
| Pamr1 | 2420.952237 | 5.401551386 | 1.19219528 | 4.92E-07 | 3.50E-06 |
| Cpxm1 | 14.83367858 | 6.379284805 | 2.282021712 | 4.99E-07 | 3.54E-06 |
| Pdgfrb | 31.44606993 | 5.915185109 | 1.340853409 | 5.05E-07 | 3.58E-06 |
| Trim59 | 100.6685281 | 6.065686221 | 2.15431686 | 5.24E-07 | 3.71E-06 |
| AU020206 | 151.1958078 | 1.339263899 | 0.279793567 | 5.46E-07 | 3.86E-06 |
| Adgrg2 | 198.8521732 | 2.974046135 | 0.621116993 | 5.47E-07 | 3.86E-06 |
| Cacna1b | 17.66387989 | 4.255074816 | 0.874437255 | 5.48E-07 | 3.87E-06 |
| Bok | 865.7545512 | 1.217387136 | 0.254629066 | 5.51E-07 | 3.89E-06 |
| Cyb561 | 990.9136657 | 1.034203274 | 0.214086187 | 5.52E-07 | 3.90E-06 |
| Hells | 177.1201959 | 1.995949466 | 0.432591898 | 5.54E-07 | 3.91E-06 |
| Gm42928 | 45.85838748 | 3.263115795 | 0.696916957 | 5.63E-07 | 3.96E-06 |
| H2-T10 | 114.2765341 | 1.1108225 | 0.230397459 | 5.63E-07 | 3.96E-06 |
| Erf | 1483.739293 | 1.002092594 | 0.20755973 | 5.64E-07 | 3.97E-06 |
| Cdkn3 | 24.09208298 | 5.86145715 | 1.334629818 | 5.67E-07 | 3.99E-06 |
| Cers5 | 280.2544323 | 1.947577649 | 0.412726689 | 5.74E-07 | 4.04E-06 |
| Scn8a | 226.5834904 | 1.617171334 | 0.345467546 | 5.76E-07 | 4.05E-06 |
| Fam107a | 234.8449596 | 4.118637166 | 0.914796051 | 5.79E-07 | 4.07E-06 |

| | | | | | |
|---|---|---|---|---|---|
| Nucb2 | 1194.176929 | 1.051702334 | 0.218526308 | 5.85E-07 | 4.10E-06 |
| Plekhh3 | 372.2277056 | 1.007746427 | 0.208508856 | 5.89E-07 | 4.13E-06 |
| Spice1 | 247.1774444 | 1.1462832 | 0.239579389 | 5.94E-07 | 4.16E-06 |
| Mrgbp | 256.963942 | 1.000226856 | 0.207514735 | 6.03E-07 | 4.22E-06 |
| Ikbip | 283.2262314 | 1.031158333 | 0.213518722 | 6.09E-07 | 4.25E-06 |
| Col5a2 | 41.54715246 | 2.724628788 | 0.601872366 | 6.12E-07 | 4.27E-06 |
| Cenpf | 188.2855359 | 6.762917018 | 2.325610033 | 6.15E-07 | 4.30E-06 |
| Camk2n2 | 81.7164672 | 1.727503465 | 0.371731598 | 6.19E-07 | 4.32E-06 |
| Cd24a | 7914.647318 | 5.420387893 | 1.213440278 | 6.36E-07 | 4.43E-06 |
| Gm30177 | 32.55434467 | 2.166411947 | 0.47332882 | 6.39E-07 | 4.45E-06 |
| Guca2a | 14.58698859 | 6.498904635 | 2.325196717 | 6.48E-07 | 4.51E-06 |
| Rpl38-ps2 | 91.22699044 | 1.310724941 | 0.277131912 | 6.58E-07 | 4.57E-06 |
| Gm43361 | 47.76247661 | 3.265722338 | 0.734726105 | 6.61E-07 | 4.59E-06 |
| Ica1 | 157.4927787 | 1.385839708 | 0.293632005 | 6.64E-07 | 4.61E-06 |
| Hddc2 | 112.5066194 | 1.748588671 | 0.37428144 | 6.64E-07 | 4.61E-06 |
| Nqo1 | 307.819888 | 1.008314025 | 0.209836857 | 6.95E-07 | 4.80E-06 |
| Basp1 | 1021.738777 | 6.838959823 | 2.313480943 | 7.20E-07 | 4.95E-06 |
| Taf1c | 170.1973506 | 1.348845142 | 0.285836701 | 7.24E-07 | 4.98E-06 |
| Misp | 318.2898526 | 1.780824828 | 0.378770999 | 7.24E-07 | 4.98E-06 |
| Cavin3 | 114.724337 | 5.967181179 | 2.141662243 | 7.57E-07 | 5.20E-06 |
| Mgp | 11.72916205 | 6.7965482 | 2.427076831 | 7.59E-07 | 5.21E-06 |
| Ctxn1 | 427.1688268 | 3.091021297 | 0.647899757 | 7.68E-07 | 5.27E-06 |
| AC110211.1 | 550.5848049 | 1.11764698 | 0.235624651 | 7.70E-07 | 5.28E-06 |
| Zscan2 | 273.3660954 | 1.08074573 | 0.226834848 | 7.70E-07 | 5.28E-06 |
| Hspa1a | 340.0292337 | 2.084041139 | 0.454571605 | 7.91E-07 | 5.41E-06 |
| Rrm2 | 220.1373501 | 2.089879035 | 0.463281777 | 7.92E-07 | 5.41E-06 |
| Tedc1 | 70.58738618 | 6.095145459 | 2.189290356 | 7.95E-07 | 5.43E-06 |
| Spns3 | 97.22993108 | 4.776205028 | 0.955147743 | 8.04E-07 | 5.48E-06 |
| Fry | 397.0166883 | 1.070835555 | 0.225811948 | 8.55E-07 | 5.81E-06 |
| Mamld1 | 384.3744535 | 5.826664467 | 2.085372876 | 8.57E-07 | 5.82E-06 |
| Kcnb2 | 14.68221559 | 6.393320019 | 2.349520465 | 8.72E-07 | 5.92E-06 |
| Mid1 | 149.4161712 | 1.252899593 | 0.26654996 | 9.19E-07 | 6.22E-06 |
| Clba1 | 95.96405526 | 1.560875032 | 0.337700109 | 9.22E-07 | 6.23E-06 |
| Poc1a | 62.45623405 | 1.326340548 | 0.284795279 | 9.55E-07 | 6.44E-06 |
| Ddit3 | 209.6482995 | 1.680055387 | 0.369326999 | 9.67E-07 | 6.51E-06 |
| Vldlr | 740.5016774 | 1.116832651 | 0.237410609 | 9.76E-07 | 6.56E-06 |
| Zfp469 | 396.8982863 | 5.797120226 | 2.087195056 | 9.88E-07 | 6.63E-06 |
| Odad1 | 306.2584663 | 3.895122241 | 0.809089279 | 1.01E-06 | 6.79E-06 |
| Tgfb1 | 235.723318 | 1.451585123 | 0.313268444 | 1.02E-06 | 6.82E-06 |
| Ltb | 613.3930778 | 4.036881911 | 0.84881222 | 1.02E-06 | 6.84E-06 |
| Gm13751 | 9.869464431 | 6.697344673 | 2.451626066 | 1.05E-06 | 7.04E-06 |
| Gm14964 | 9.575115214 | 6.589986639 | 2.416964692 | 1.09E-06 | 7.30E-06 |
| Zfp429 | 81.41093935 | 1.539041199 | 0.334393758 | 1.11E-06 | 7.42E-06 |

| | | | | | |
|---|---|---|---|---|---|
| Gm44639 | 19.74551052 | 6.416915824 | 2.324711174 | 1.12E-06 | 7.47E-06 |
| 1110051M20Rik | 220.5302099 | 1.237421897 | 0.266716907 | 1.13E-06 | 7.54E-06 |
| Magohb | 78.62208101 | 1.52193291 | 0.333265885 | 1.18E-06 | 7.81E-06 |
| Tekt5 | 15.40230972 | 4.858936624 | 1.005117822 | 1.18E-06 | 7.85E-06 |
| Lum | 43.07692162 | 3.899092423 | 0.898997318 | 1.19E-06 | 7.88E-06 |
| Gm36283 | 60.01923427 | 1.728347274 | 0.381465863 | 1.19E-06 | 7.88E-06 |
| Gm37422 | 70.97964144 | 1.57656986 | 0.345159407 | 1.22E-06 | 8.07E-06 |
| Gas2l3 | 197.406415 | 1.644034387 | 0.364375961 | 1.30E-06 | 8.52E-06 |
| Veph1 | 205.4741235 | 1.694207912 | 0.37629996 | 1.42E-06 | 9.27E-06 |
| Clstn1 | 1488.206091 | 2.589768745 | 0.564592946 | 1.43E-06 | 9.35E-06 |
| Ccdc68 | 72.84594249 | 1.886774027 | 0.418269057 | 1.46E-06 | 9.48E-06 |
| Eda | 64.36601547 | 1.289054945 | 0.281437421 | 1.48E-06 | 9.61E-06 |
| Plet1os | 77.76262449 | 2.339742726 | 0.524608878 | 1.48E-06 | 9.63E-06 |
| Gm26532 | 19.68934037 | 6.280488859 | 2.30132031 | 1.53E-06 | 9.94E-06 |
| Rims2 | 147.3124243 | 4.091503655 | 0.889158316 | 1.56E-06 | 1.01E-05 |
| Epb41l4a | 490.01875 | 5.641944996 | 2.065620117 | 1.56E-06 | 1.01E-05 |
| Enkd1 | 98.5784875 | 2.487150405 | 0.548799566 | 1.59E-06 | 1.03E-05 |
| B9d1 | 123.0617653 | 2.220110217 | 0.489236412 | 1.59E-06 | 1.03E-05 |
| Kcnq1ot1 | 171.0707925 | 1.528452193 | 0.340905242 | 1.60E-06 | 1.03E-05 |
| Gm37352 | 23.01723207 | 6.381250512 | 2.331581613 | 1.65E-06 | 1.07E-05 |
| Nin | 271.069687 | 1.061500496 | 0.229890538 | 1.66E-06 | 1.07E-05 |
| Gm43360 | 32.79507819 | 2.742360944 | 0.634135996 | 1.66E-06 | 1.07E-05 |
| Gm45457 | 158.8485913 | 1.750143389 | 0.39452483 | 1.69E-06 | 1.09E-05 |
| Ercc1 | 171.0147798 | 1.025928234 | 0.221773755 | 1.69E-06 | 1.09E-05 |
| Trim62 | 91.27831857 | 1.598035053 | 0.356366523 | 1.70E-06 | 1.09E-05 |
| 9530077C05Rik | 113.1858101 | 3.353490793 | 0.725504085 | 1.71E-06 | 1.10E-05 |
| Shisa4 | 490.9216844 | 5.60012464 | 2.059433666 | 1.78E-06 | 1.14E-05 |
| Hcar2 | 54.58956443 | 1.67547259 | 0.378554186 | 1.82E-06 | 1.16E-05 |
| Creb5 | 259.0509532 | 5.669887973 | 2.092213195 | 1.85E-06 | 1.18E-05 |
| Psd | 180.1177899 | 1.794823721 | 0.406695231 | 1.86E-06 | 1.19E-05 |
| Krt5 | 7.169073494 | 6.311278124 | 2.398361671 | 2.00E-06 | 1.27E-05 |
| Oxct1 | 696.4409434 | 1.848885966 | 0.416536721 | 2.01E-06 | 1.28E-05 |
| 1500011B03Rik | 110.7294255 | 2.003541958 | 0.452573774 | 2.03E-06 | 1.29E-05 |
| Trim24 | 1029.7912 | 1.140546694 | 0.251757772 | 2.07E-06 | 1.31E-05 |
| Plk1 | 69.1291971 | 4.937527612 | 1.067104272 | 2.08E-06 | 1.31E-05 |
| Airn | 113.8688522 | 1.829342446 | 0.413913847 | 2.14E-06 | 1.35E-05 |
| Fam83d | 43.90729814 | 6.086997123 | 2.258090991 | 2.18E-06 | 1.37E-05 |
| Fkbp5 | 1265.795796 | 1.198004249 | 0.266126602 | 2.19E-06 | 1.38E-05 |
| Atn1 | 147.1075151 | 1.239309075 | 0.275592724 | 2.20E-06 | 1.38E-05 |
| Eid3 | 18.31802696 | 2.626024859 | 0.606745182 | 2.25E-06 | 1.41E-05 |
| Btbd19 | 44.23332552 | 1.901600389 | 0.439437317 | 2.27E-06 | 1.42E-05 |
| Gpat3 | 142.7667544 | 1.012571205 | 0.222793909 | 2.30E-06 | 1.44E-05 |
| Frmd4a | 522.0092215 | 1.567292154 | 0.356219409 | 2.39E-06 | 1.49E-05 |

| | | | | | |
|---|---|---|---|---|---|
| Ank1 | 96.47599374 | 5.677405476 | 2.119114902 | 2.40E-06 | 1.50E-05 |
| Psat1 | 122.1402971 | 1.763176857 | 0.402941759 | 2.50E-06 | 1.56E-05 |
| Bicd1 | 254.2985166 | 5.541822658 | 2.07840897 | 2.55E-06 | 1.58E-05 |
| Foxj1 | 1158.630639 | 5.533364813 | 2.048482894 | 2.55E-06 | 1.59E-05 |
| Gm36638 | 17.66924104 | 3.381081098 | 0.776750737 | 2.67E-06 | 1.65E-05 |
| Cdca8 | 74.4217961 | 1.780856938 | 0.40870706 | 2.67E-06 | 1.65E-05 |
| Lrmda | 28.40672573 | 2.452215131 | 0.57592115 | 2.69E-06 | 1.66E-05 |
| Dab1 | 428.2911511 | 4.528700013 | 0.962067532 | 2.76E-06 | 1.70E-05 |
| Krt23 | 775.5902267 | 2.427149613 | 0.579895225 | 2.77E-06 | 1.71E-05 |
| Smim3 | 170.244785 | 1.632552605 | 0.371832846 | 2.81E-06 | 1.74E-05 |
| Cxcl1 | 1410.42543 | 1.197292228 | 0.269265261 | 2.84E-06 | 1.75E-05 |
| Ager | 70.79660825 | 5.597680676 | 2.126525825 | 2.86E-06 | 1.76E-05 |
| Fignl1 | 63.27404966 | 3.359627027 | 0.790860909 | 2.92E-06 | 1.80E-05 |
| Cyp2b10 | 495.0599361 | 2.708062491 | 0.654499787 | 2.99E-06 | 1.84E-05 |
| Krtcap3 | 182.9199105 | 1.452092376 | 0.327944798 | 3.00E-06 | 1.84E-05 |
| Lmntd2 | 142.4642597 | 1.212896734 | 0.273353252 | 3.04E-06 | 1.86E-05 |
| Cd200 | 290.627671 | 5.482876447 | 2.062716617 | 3.05E-06 | 1.87E-05 |
| Meltf | 168.9532373 | 5.187468139 | 1.267902244 | 3.05E-06 | 1.87E-05 |
| Bub1 | 40.9483308 | 5.757220829 | 2.19311505 | 3.09E-06 | 1.89E-05 |
| Dzip1l | 765.3826729 | 4.424719531 | 0.939359696 | 3.15E-06 | 1.92E-05 |
| Ier5l | 202.692531 | 4.464321126 | 0.954128976 | 3.20E-06 | 1.96E-05 |
| Pbk | 51.82096269 | 6.147589683 | 2.302068933 | 3.27E-06 | 1.99E-05 |
| Coprs | 78.59748785 | 1.402646686 | 0.320755881 | 3.29E-06 | 2.00E-05 |
| Spag5 | 98.07731213 | 2.235902853 | 0.53164572 | 3.30E-06 | 2.01E-05 |
| Rnase1 | 18.03063134 | 6.219351567 | 2.344416269 | 3.30E-06 | 2.01E-05 |
| Plch1 | 155.1805093 | 4.946187649 | 1.199196144 | 3.30E-06 | 2.01E-05 |
| Aoc2 | 78.73395579 | 1.727015238 | 0.397935032 | 3.31E-06 | 2.01E-05 |
| Sgcb | 72.74187539 | 2.074149292 | 0.477652411 | 3.42E-06 | 2.08E-05 |
| A230083G16Rik | 84.11065141 | 1.26034423 | 0.28723823 | 3.44E-06 | 2.09E-05 |
| Adamtsl5 | 21.26029068 | 3.281355351 | 0.763390627 | 3.45E-06 | 2.09E-05 |
| Timp4 | 11.93167421 | 6.215124545 | 2.360865337 | 3.64E-06 | 2.20E-05 |
| Mab21l4 | 94.28698676 | 1.830515728 | 0.425798664 | 3.67E-06 | 2.22E-05 |
| Plk3 | 1881.600877 | 1.327117813 | 0.305047741 | 3.77E-06 | 2.27E-05 |
| Thbs1 | 3384.490382 | 4.78680403 | 1.159180156 | 3.80E-06 | 2.28E-05 |
| Neurl1b | 148.6441284 | 5.199983919 | 1.202959658 | 3.99E-06 | 2.39E-05 |
| Gata3 | 18.3510448 | 3.735672986 | 0.879099884 | 4.07E-06 | 2.43E-05 |
| Gm49327 | 73.64457218 | 1.714605192 | 0.399102674 | 4.15E-06 | 2.48E-05 |
| Arhgef6 | 72.6600184 | 2.292753299 | 0.544965387 | 4.36E-06 | 2.60E-05 |
| Srd5a2 | 311.6276207 | 1.069747558 | 0.243248271 | 4.37E-06 | 2.60E-05 |
| Cdc6 | 46.05270897 | 5.8418265 | 2.235959205 | 4.39E-06 | 2.61E-05 |
| Apela | 11.29400117 | 6.787019728 | 2.598543123 | 4.40E-06 | 2.61E-05 |
| Dtl | 105.5778503 | 2.673504057 | 0.64577672 | 4.40E-06 | 2.62E-05 |
| Cidec | 51.46699202 | 1.647814473 | 0.388464468 | 4.42E-06 | 2.63E-05 |

| | | | | | |
|---|---|---|---|---|---|
| Ska2 | 113.6722477 | 1.75947524 | 0.409665741 | 4.56E-06 | 2.70E-05 |
| Thsd7a | 160.8625997 | 5.488184273 | 2.10004551 | 4.59E-06 | 2.72E-05 |
| Arid5a | 221.6635287 | 1.718997979 | 0.407209952 | 4.59E-06 | 2.72E-05 |
| Gpc3 | 30.3801853 | 5.1871583 | 1.307265221 | 4.82E-06 | 2.84E-05 |
| Opn3 | 29.28124358 | 1.982829834 | 0.471492462 | 4.86E-06 | 2.86E-05 |
| Gpsm1 | 252.0806411 | 4.332374266 | 0.941202921 | 4.87E-06 | 2.87E-05 |
| Itgb3 | 33.89830086 | 4.276056598 | 1.050663403 | 4.96E-06 | 2.92E-05 |
| Grip1 | 88.74446334 | 2.768498687 | 0.64962761 | 4.99E-06 | 2.93E-05 |
| Tmod1 | 116.5443777 | 5.466932369 | 2.117613015 | 5.03E-06 | 2.95E-05 |
| Tmem132a | 566.3561289 | 3.116685412 | 0.708552135 | 5.11E-06 | 3.00E-05 |
| Tnfrsf11a | 85.1337696 | 3.656744739 | 0.827062702 | 5.11E-06 | 3.00E-05 |
| Nradd | 310.3414058 | 1.520342326 | 0.354692646 | 5.26E-06 | 3.08E-05 |
| AC156638.1 | 14.4341119 | 2.901294846 | 0.69329885 | 5.81E-06 | 3.38E-05 |
| Slc1a1 | 42.56718334 | 4.976498462 | 1.259949482 | 5.86E-06 | 3.41E-05 |
| Prkab2 | 176.5981521 | 1.114822692 | 0.256824197 | 5.88E-06 | 3.42E-05 |
| Kbtbd8 | 41.077784 | 2.097666941 | 0.51306654 | 5.95E-06 | 3.45E-05 |
| Ltbp4 | 477.6364358 | 1.460915624 | 0.347990697 | 6.03E-06 | 3.50E-05 |
| Nusap1 | 103.2291263 | 1.865363607 | 0.449285706 | 6.15E-06 | 3.56E-05 |
| Myo5a | 131.6653019 | 1.783379346 | 0.421179515 | 6.21E-06 | 3.59E-05 |
| Fgd3 | 516.9420966 | 4.232429063 | 0.928299947 | 6.31E-06 | 3.65E-05 |
| Gm31084 | 64.30040656 | 1.277049089 | 0.298612945 | 6.44E-06 | 3.72E-05 |
| Clspn | 67.72578872 | 6.453016153 | 2.426119804 | 6.49E-06 | 3.74E-05 |
| Gm42929 | 51.01133385 | 1.983599048 | 0.484041624 | 6.50E-06 | 3.75E-05 |
| Pkia | 312.5624011 | 4.228968874 | 0.930553335 | 6.58E-06 | 3.79E-05 |
| Cyp2c55 | 164.3975735 | 1.469794629 | 0.351862437 | 6.69E-06 | 3.85E-05 |
| Kazn | 95.36966977 | 2.619690718 | 0.618477668 | 6.83E-06 | 3.93E-05 |
| Krt20 | 69.46770226 | 5.785304751 | 2.240382092 | 6.83E-06 | 3.93E-05 |
| Cldn23 | 84.22403011 | 2.926135344 | 0.693831376 | 6.91E-06 | 3.97E-05 |
| Pdgfra | 23.43576109 | 4.035270894 | 0.968256176 | 7.04E-06 | 4.03E-05 |
| Tmem45b | 118.8257564 | 4.809373722 | 1.220847869 | 7.04E-06 | 4.03E-05 |
| Stil | 69.22256311 | 4.325037761 | 0.969454336 | 7.11E-06 | 4.07E-05 |
| Clca3a1 | 42.1005888 | 3.518865187 | 0.850935009 | 7.19E-06 | 4.11E-05 |
| Rad54l | 52.05831429 | 5.466114035 | 2.160300864 | 7.50E-06 | 4.28E-05 |
| Tppp3 | 90.84479318 | 5.336603104 | 2.107607969 | 7.58E-06 | 4.32E-05 |
| Slc9a7 | 104.8055513 | 1.648736613 | 0.397363235 | 7.69E-06 | 4.38E-05 |
| Tmem51 | 317.5507215 | 1.090466088 | 0.256893616 | 7.77E-06 | 4.43E-05 |
| Ptgfr | 404.575504 | 4.777509347 | 1.217560493 | 7.79E-06 | 4.44E-05 |
| Tcf19 | 209.2972139 | 1.482470437 | 0.35842119 | 7.81E-06 | 4.45E-05 |
| H2ax | 330.2827875 | 1.029131497 | 0.240937524 | 7.87E-06 | 4.47E-05 |
| Gm43980 | 118.8122645 | 1.551092057 | 0.371003283 | 8.04E-06 | 4.56E-05 |
| Ntf3 | 25.71582979 | 5.525211208 | 1.464831714 | 8.08E-06 | 4.58E-05 |
| Tonsl | 222.5322796 | 1.08891905 | 0.256166782 | 8.16E-06 | 4.62E-05 |
| Kif3c | 322.9603925 | 2.223669519 | 0.53085084 | 8.27E-06 | 4.68E-05 |

| | | | | | |
|---|---|---|---|---|---|
| Plod2 | 169.7300511 | 5.357262671 | 2.118261367 | 8.47E-06 | 4.78E-05 |
| Hmgcll1 | 64.0633418 | 5.272578315 | 2.103657536 | 8.55E-06 | 4.82E-05 |
| Zfp40 | 108.3734807 | 1.0450924 | 0.24544268 | 8.69E-06 | 4.89E-05 |
| Snhg12 | 158.070228 | 1.064533015 | 0.250861354 | 8.80E-06 | 4.95E-05 |
| Gm47528 | 13.4638242 | 2.696997802 | 0.680743312 | 8.93E-06 | 5.02E-05 |
| Spata6 | 279.0047109 | 4.219084762 | 0.953543084 | 8.98E-06 | 5.04E-05 |
| Dpt | 12.71434071 | 6.370245147 | 2.470051311 | 9.01E-06 | 5.06E-05 |
| Tpbgl | 34.86131448 | 1.605213787 | 0.392277194 | 9.08E-06 | 5.09E-05 |
| Aebp1 | 176.1054284 | 3.022040163 | 0.74066663 | 9.26E-06 | 5.18E-05 |
| Ap1s2 | 23.28046215 | 2.313877289 | 0.566612724 | 9.34E-06 | 5.22E-05 |
| Gm42793 | 566.2139603 | 5.103539087 | 2.021151174 | 9.39E-06 | 5.25E-05 |
| Sox9 | 3447.10055 | 1.129306224 | 0.269384333 | 9.64E-06 | 5.38E-05 |
| Slc41a3 | 133.3096149 | 1.748075087 | 0.434720101 | 9.69E-06 | 5.41E-05 |
| Wdr72 | 204.2332789 | 5.186832959 | 2.071526921 | 9.81E-06 | 5.47E-05 |
| Fkbp1b | 31.87372976 | 5.281791428 | 2.129185071 | 9.84E-06 | 5.48E-05 |
| Zfp599 | 94.56903607 | 2.553227487 | 0.614727607 | 9.89E-06 | 5.51E-05 |
| Scube1 | 10.99974653 | 5.900974115 | 2.345862511 | 1.01E-05 | 5.60E-05 |
| Cacna2d4 | 15.39788686 | 3.668014638 | 0.87700017 | 1.02E-05 | 5.67E-05 |
| Chaf1b | 172.7907734 | 1.470577803 | 0.35979666 | 1.03E-05 | 5.73E-05 |
| Il20rb | 44.7960398 | 2.666399297 | 0.671588238 | 1.04E-05 | 5.79E-05 |
| Tbx1 | 580.1924842 | 6.52604175 | 2.453957738 | 1.05E-05 | 5.80E-05 |
| Pde10a | 38.64573954 | 3.648433073 | 0.872933957 | 1.05E-05 | 5.84E-05 |
| Fam171b | 48.01591245 | 5.379853316 | 2.162141477 | 1.07E-05 | 5.90E-05 |
| Lox | 8.969308205 | 5.62088054 | 2.274016058 | 1.08E-05 | 5.96E-05 |
| Egflam | 1011.28862 | 5.07215619 | 2.013230905 | 1.08E-05 | 5.97E-05 |
| Tmem54 | 68.25101052 | 5.288468257 | 2.141059525 | 1.10E-05 | 6.07E-05 |
| Ets1 | 789.878056 | 1.189127219 | 0.286188242 | 1.11E-05 | 6.12E-05 |
| Mtcl1 | 872.6430068 | 5.033686639 | 2.006110637 | 1.12E-05 | 6.19E-05 |
| Fchsd1 | 471.3226952 | 1.490974401 | 0.366693345 | 1.13E-05 | 6.21E-05 |
| Adam15 | 279.7673447 | 2.214774758 | 0.53884825 | 1.13E-05 | 6.23E-05 |
| Mok | 55.98178475 | 5.165339038 | 2.088991851 | 1.13E-05 | 6.24E-05 |
| Vil1 | 365.724313 | 4.131215971 | 0.94091626 | 1.14E-05 | 6.29E-05 |
| Ube2c | 73.83491087 | 3.036677985 | 0.78796265 | 1.15E-05 | 6.30E-05 |
| Dynlt1f | 121.9569769 | 1.819596348 | 0.451013275 | 1.15E-05 | 6.31E-05 |
| Gm16011 | 838.1269171 | 5.073602197 | 2.020593866 | 1.15E-05 | 6.31E-05 |
| Rad18 | 107.0307597 | 1.325354069 | 0.322977409 | 1.15E-05 | 6.33E-05 |
| Nebl | 109.6906408 | 2.759198772 | 0.668466792 | 1.16E-05 | 6.34E-05 |
| Cfap54 | 86.12145659 | 3.073273383 | 0.740709761 | 1.17E-05 | 6.40E-05 |
| Slco1a6 | 35.08598132 | 5.291378177 | 2.145761021 | 1.17E-05 | 6.41E-05 |
| Tnfsf12 | 179.1421571 | 1.234735425 | 0.299911553 | 1.19E-05 | 6.48E-05 |
| Gm44950 | 93.05090018 | 1.242152584 | 0.301032642 | 1.22E-05 | 6.68E-05 |
| Mamdc4 | 65.2279345 | 1.639664944 | 0.408687175 | 1.23E-05 | 6.73E-05 |
| Cib3 | 873.5887824 | 1.053118916 | 0.253590018 | 1.24E-05 | 6.74E-05 |

| | | | | | |
|---|---|---|---|---|---|
| Atp2b2 | 133.9032802 | 1.104102488 | 0.266975815 | 1.27E-05 | 6.93E-05 |
| Ncapg | 58.65207334 | 3.015850385 | 0.769338174 | 1.28E-05 | 6.94E-05 |
| Usp11 | 325.6033467 | 1.596630077 | 0.393344884 | 1.28E-05 | 6.95E-05 |
| Sh2d4b | 143.4454641 | 5.0710651 | 2.048795179 | 1.29E-05 | 6.98E-05 |
| Gm17024 | 91.74932521 | 2.596917498 | 0.684850091 | 1.32E-05 | 7.15E-05 |
| Gm42937 | 35.817212 | 2.671703941 | 0.6979712 | 1.32E-05 | 7.17E-05 |
| 4930579G18Rik | 93.24855243 | 1.347684514 | 0.330137212 | 1.33E-05 | 7.21E-05 |
| Gpr153 | 21.26597132 | 5.332790754 | 2.180003004 | 1.35E-05 | 7.31E-05 |
| Spata24 | 46.40203174 | 1.929925115 | 0.484015135 | 1.38E-05 | 7.42E-05 |
| Ntrk2 | 431.6986098 | 2.775914337 | 0.744290111 | 1.40E-05 | 7.53E-05 |
| Ccdc18 | 66.93967832 | 2.714133786 | 0.664223547 | 1.42E-05 | 7.62E-05 |
| Upk3b | 9.530091929 | 5.441419047 | 2.236923301 | 1.42E-05 | 7.63E-05 |
| Usp2 | 403.8005691 | 2.67823653 | 0.715361082 | 1.43E-05 | 7.66E-05 |
| Wdr62 | 79.69927961 | 1.084292711 | 0.262966596 | 1.46E-05 | 7.84E-05 |
| Fam241a | 77.1853476 | 1.202865165 | 0.294844694 | 1.47E-05 | 7.88E-05 |
| Ntn1 | 103.2084884 | 2.815578033 | 0.699934569 | 1.48E-05 | 7.90E-05 |
| Bub1b | 113.7057321 | 2.559407749 | 0.656292665 | 1.49E-05 | 7.96E-05 |
| Noct | 758.1771916 | 2.099979863 | 0.549172296 | 1.51E-05 | 8.04E-05 |
| Tmc5 | 43.86023151 | 5.316177924 | 2.176321236 | 1.52E-05 | 8.10E-05 |
| Hcn2 | 58.37663793 | 1.660608709 | 0.417106609 | 1.52E-05 | 8.13E-05 |
| Atf3 | 3243.267135 | 3.501087967 | 0.970810065 | 1.53E-05 | 8.18E-05 |
| Ccdc3 | 8.914326112 | 5.426077422 | 2.242072199 | 1.53E-05 | 8.18E-05 |
| Ankrd55 | 8.762867722 | 4.851581873 | 1.312593033 | 1.55E-05 | 8.26E-05 |
| Tnfsfm13 | 29.82880007 | 2.48014418 | 0.63925822 | 1.55E-05 | 8.27E-05 |
| Gm16091 | 37.80693799 | 4.226320648 | 1.005008673 | 1.58E-05 | 8.42E-05 |
| Gm49590 | 37.39943702 | 2.988705338 | 0.759187315 | 1.60E-05 | 8.49E-05 |
| Nup62cl | 19.01550381 | 2.968428803 | 0.793025116 | 1.63E-05 | 8.65E-05 |
| Rgl1 | 100.54401 | 1.389878005 | 0.347811371 | 1.68E-05 | 8.91E-05 |
| Crocc | 215.0210224 | 1.028986459 | 0.250512513 | 1.70E-05 | 8.98E-05 |
| Napepld | 101.6244337 | 1.103767036 | 0.270662463 | 1.70E-05 | 8.98E-05 |
| Nupr1 | 99.07289363 | 2.748898562 | 0.68474329 | 1.72E-05 | 9.05E-05 |
| Pkhd1 | 4275.810134 | 1.034782472 | 0.253481084 | 1.73E-05 | 9.12E-05 |
| Gm43481 | 75.31890678 | 1.427438807 | 0.357094666 | 1.74E-05 | 9.16E-05 |
| Zfp248 | 87.29342852 | 1.519682955 | 0.380375953 | 1.75E-05 | 9.24E-05 |
| Melk | 62.82479422 | 5.104031128 | 2.116564577 | 1.81E-05 | 9.54E-05 |
| Espl1 | 82.51814812 | 3.81367336 | 0.969287739 | 1.82E-05 | 9.59E-05 |
| Fmod | 9.003728375 | 5.586354399 | 2.305855697 | 1.83E-05 | 9.62E-05 |
| Phf11d | 136.5903438 | 1.650966081 | 0.415427875 | 1.86E-05 | 9.75E-05 |
| Pask | 113.5348826 | 1.388149427 | 0.345006105 | 1.87E-05 | 9.78E-05 |
| Nek5 | 134.9824918 | 2.441587593 | 0.609011787 | 1.90E-05 | 9.95E-05 |
| Cip2a | 129.6753678 | 1.126413764 | 0.277913706 | 1.91E-05 | 0.000100074 |
| Enkur | 81.68257409 | 4.524795494 | 1.219060433 | 1.92E-05 | 0.000100349 |
| Ophn1 | 153.9617263 | 1.033753489 | 0.25303709 | 1.95E-05 | 0.000101776 |

| | | | | | |
|---|---|---|---|---|---|
| Scara3 | 5836.559404 | 4.044353303 | 0.9613204 | 1.96E-05 | 0.000102367 |
| Tmem154 | 53.94579418 | 3.644957655 | 0.917402278 | 1.98E-05 | 0.000103097 |
| Arhgap4 | 82.53477645 | 1.32805815 | 0.334767804 | 1.98E-05 | 0.000103187 |
| Syn3 | 79.4794392 | 1.521475167 | 0.385892623 | 2.00E-05 | 0.000104028 |
| 9130017K11Rik | 40.95060515 | 5.074228847 | 2.119481366 | 2.06E-05 | 0.000107217 |
| Trim16 | 136.1153518 | 4.954477951 | 2.072727104 | 2.15E-05 | 0.000111121 |
| Fhl1 | 84.42554324 | 2.561032531 | 0.667033763 | 2.19E-05 | 0.000113244 |
| Rhpn1 | 60.97190403 | 4.944660878 | 2.073196871 | 2.22E-05 | 0.000114649 |
| Rnf180 | 214.8265372 | 4.89799181 | 2.036346935 | 2.23E-05 | 0.000115263 |
| 2410006H16Rik | 204.3508789 | 1.1891509 | 0.299241858 | 2.26E-05 | 0.000116802 |
| A930033H14Rik | 65.19165595 | 2.453203713 | 0.667991037 | 2.29E-05 | 0.000118047 |
| Emilin1 | 63.80878955 | 1.690573447 | 0.442470521 | 2.31E-05 | 0.000119271 |
| H1f5 | 10.73173961 | 5.020664461 | 1.415599102 | 2.33E-05 | 0.000120065 |
| Flrt3 | 300.4607345 | 4.996895454 | 2.075754312 | 2.35E-05 | 0.000121195 |
| Sh3d21 | 149.3586842 | 1.679767542 | 0.433302031 | 2.43E-05 | 0.000124626 |
| Sync | 32.68998573 | 4.174756707 | 1.028589562 | 2.44E-05 | 0.000125305 |
| Cygb | 21.58253731 | 3.302836487 | 0.886898706 | 2.50E-05 | 0.000127977 |
| Cuedc1 | 613.8840144 | 2.688356111 | 0.688076924 | 2.51E-05 | 0.000128326 |
| Flt3l | 83.04877609 | 1.178931078 | 0.298044703 | 2.51E-05 | 0.000128597 |
| Anxa3 | 1721.356436 | 2.92662012 | 0.743571103 | 2.54E-05 | 0.000130125 |
| Mroh6 | 20.6931891 | 1.880626774 | 0.493719806 | 2.61E-05 | 0.000133321 |
| Gm11639 | 11.88286842 | 4.213299371 | 1.050276979 | 2.66E-05 | 0.000135508 |
| Nemp2 | 99.53917871 | 3.998026528 | 0.971806659 | 2.71E-05 | 0.000137679 |
| Pdp1 | 126.881524 | 1.249560049 | 0.318314001 | 2.71E-05 | 0.000137888 |
| Id4 | 12.35752963 | 4.724417717 | 1.233545431 | 2.72E-05 | 0.000138392 |
| Atp10d | 151.5524357 | 1.094222404 | 0.275781901 | 2.78E-05 | 0.00014103 |
| Zfp689 | 93.77918479 | 1.055554862 | 0.263872592 | 2.80E-05 | 0.000141905 |
| Tcf24 | 347.3894377 | 1.386413189 | 0.359639989 | 2.88E-05 | 0.000145331 |
| Aspm | 105.6446613 | 2.523102932 | 0.684025874 | 2.95E-05 | 0.000148729 |
| Msln | 683.1667915 | 4.375112494 | 1.21165931 | 3.00E-05 | 0.000151019 |
| Mbnl3 | 26.88781988 | 2.131658328 | 0.569326604 | 3.01E-05 | 0.000151271 |
| Arhgap36 | 56.66891399 | 1.69276794 | 0.445812798 | 3.01E-05 | 0.000151366 |
| Efemp2 | 333.6761589 | 2.418444235 | 0.62643162 | 3.05E-05 | 0.000153097 |
| Spon1 | 401.3325519 | 4.058606435 | 1.008009381 | 3.07E-05 | 0.000153955 |
| Ankrd1 | 2770.215599 | 3.639268681 | 0.968069255 | 3.08E-05 | 0.000154746 |
| Adamts15 | 163.6299297 | 2.591399636 | 0.70375158 | 3.10E-05 | 0.000155428 |
| Gm19221 | 41.19545013 | 1.779400085 | 0.462743487 | 3.12E-05 | 0.000156507 |
| Smim24 | 22.60151548 | 2.542057895 | 0.667556699 | 3.15E-05 | 0.000157586 |
| Spindoc | 186.573418 | 1.007092956 | 0.254006188 | 3.24E-05 | 0.000161874 |
| Gm46516 | 90.23536574 | 1.527192985 | 0.404334704 | 3.24E-05 | 0.000161939 |
| Azin2 | 115.3553129 | 1.662815524 | 0.441757641 | 3.32E-05 | 0.000165312 |
| Selenom | 95.91489829 | 4.812880506 | 2.062683533 | 3.38E-05 | 0.00016831 |
| Gm47950 | 24.94691639 | 3.274789592 | 0.8319668 | 3.40E-05 | 0.00016934 |

| | | | | | |
|---|---|---|---|---|---|
| Fkbp10 | 123.6029324 | 4.805247581 | 2.067422692 | 3.70E-05 | 0.0001834 |
| Palm | 760.8906651 | 1.395376363 | 0.368843553 | 3.72E-05 | 0.00018398 |
| Plxna4 | 14.42372022 | 4.561309782 | 1.309622286 | 3.75E-05 | 0.00018555 |
| Cdc25c | 11.88814144 | 5.089224713 | 2.216911871 | 3.77E-05 | 0.000186243 |
| Cdca7l | 107.6290371 | 3.252087819 | 0.834259485 | 3.83E-05 | 0.000189155 |
| Cenpe | 89.47404783 | 4.04798151 | 1.030247002 | 3.88E-05 | 0.000191165 |
| 5930420M18Rik | 34.9787551 | 1.91632513 | 0.514576957 | 3.89E-05 | 0.000191608 |
| Gm10030 | 35.15347142 | 1.53955992 | 0.411733163 | 3.98E-05 | 0.000195637 |
| Adgra2 | 71.92095866 | 1.958842881 | 0.529359291 | 4.01E-05 | 0.000197114 |
| Mcm4 | 383.1632486 | 1.148698202 | 0.298493902 | 4.04E-05 | 0.000198439 |
| Oacyl | 29.06736327 | 5.03947488 | 2.174058113 | 4.06E-05 | 0.000199198 |
| Svep1 | 11.10221346 | 5.568966754 | 2.370495912 | 4.08E-05 | 0.000199878 |
| Pkm | 452.1244789 | 1.127151066 | 0.29123535 | 4.11E-05 | 0.000201334 |
| Ampd3 | 249.1452166 | 2.084586597 | 0.574765894 | 4.19E-05 | 0.000204965 |
| Dcn | 140.1508072 | 2.008684321 | 0.561717727 | 4.24E-05 | 0.000207356 |
| B4galt2 | 210.7762572 | 4.639218108 | 2.006973524 | 4.34E-05 | 0.000211628 |
| Ccdc74a | 30.78210296 | 3.042114664 | 0.816264268 | 4.37E-05 | 0.000213118 |
| Fam189b | 109.872702 | 2.513136276 | 0.659432683 | 4.38E-05 | 0.000213358 |
| Cklf | 86.76632974 | 1.888388191 | 0.50564954 | 4.52E-05 | 0.000220038 |
| Magi2 | 24.37552289 | 4.862626508 | 2.127022844 | 4.54E-05 | 0.000220581 |
| Dhrs9 | 150.1315802 | 1.232848343 | 0.325281558 | 4.56E-05 | 0.000221476 |
| Zdhhc1 | 360.0125597 | 1.103380682 | 0.285249034 | 4.59E-05 | 0.000222963 |
| Sapcd2 | 24.14546453 | 4.844139792 | 2.13411354 | 4.60E-05 | 0.000223314 |
| Gnb5 | 138.5540797 | 4.669622579 | 2.036292785 | 4.62E-05 | 0.00022413 |
| Sod3 | 498.9041449 | 1.756315711 | 0.478948662 | 4.70E-05 | 0.000227578 |
| Dbp | 1420.647959 | 1.758304453 | 0.487625981 | 4.70E-05 | 0.000227774 |
| Pclaf | 73.92880739 | 3.447311665 | 0.967715717 | 4.70E-05 | 0.000227825 |
| Cep112 | 99.60240558 | 2.784584294 | 0.731445618 | 4.75E-05 | 0.000229687 |
| Gm12498 | 13.03505867 | 2.204678947 | 0.617240854 | 4.78E-05 | 0.000231015 |
| Gm16170 | 103.5760646 | 1.325275517 | 0.351746826 | 5.02E-05 | 0.000241957 |
| Lrrc39 | 32.42581701 | 1.539705894 | 0.41786769 | 5.02E-05 | 0.000242141 |
| Zc3hav1l | 191.1381486 | 3.121407119 | 0.808182675 | 5.09E-05 | 0.000245088 |
| Akr1b10 | 104.6207861 | 1.742579447 | 0.474168298 | 5.18E-05 | 0.000249191 |
| Sh3tc1 | 77.0993071 | 1.812977071 | 0.487282658 | 5.22E-05 | 0.000250872 |
| Meis3 | 184.4185266 | 4.697316273 | 2.054632015 | 5.23E-05 | 0.000251206 |
| Cep85 | 6002.535814 | 1.046538291 | 0.27348913 | 5.27E-05 | 0.000253378 |
| Lbh | 256.5669059 | 1.638200518 | 0.441423655 | 5.28E-05 | 0.000253825 |
| Kif18b | 46.11416149 | 5.055764474 | 2.211184598 | 5.39E-05 | 0.000258358 |
| Dusp6 | 2465.49765 | 1.038256222 | 0.272125173 | 5.43E-05 | 0.000260244 |
| Mapk8ip1 | 662.1728763 | 1.544379543 | 0.410904713 | 5.46E-05 | 0.000261306 |
| Ticrr | 34.79120974 | 5.094563142 | 2.228538682 | 5.54E-05 | 0.000265311 |
| Fa2h | 18.45245771 | 4.919936871 | 2.178988557 | 5.55E-05 | 0.000265679 |
| Gsta5 | 9.110892409 | 3.417843344 | 0.97259663 | 5.57E-05 | 0.00026649 |

| | | | | | |
|---|---|---|---|---|---|
| Birc5 | 61.90512381 | 1.291859272 | 0.345881608 | 5.63E-05 | 0.000269076 |
| Atp10a | 571.1271433 | 2.119111531 | 0.585367589 | 5.73E-05 | 0.000273628 |
| Fanca | 103.4861522 | 1.178477277 | 0.313035916 | 5.78E-05 | 0.000275244 |
| Gadd45a | 297.0520095 | 1.390108355 | 0.379793351 | 6.01E-05 | 0.000285356 |
| Gm13056 | 26.31396775 | 3.800867541 | 0.977123568 | 6.05E-05 | 0.000287057 |
| Dusp13 | 4.426200121 | 5.318768659 | 2.369982295 | 6.07E-05 | 0.000287613 |
| Gm42941 | 16.39181064 | 4.430192874 | 1.320529003 | 6.09E-05 | 0.000288344 |
| Sbsn | 49.57013045 | 2.523571676 | 0.683723213 | 6.11E-05 | 0.000289253 |
| Ttk | 33.32754751 | 4.377312291 | 1.300585031 | 6.15E-05 | 0.000290878 |
| Smim22 | 101.2672811 | 1.541051475 | 0.427776008 | 6.39E-05 | 0.000301935 |
| Nuf2 | 45.99483259 | 4.505041867 | 1.35904702 | 6.61E-05 | 0.000311198 |
| Ggt1 | 88.49727745 | 4.105732854 | 1.200575257 | 6.66E-05 | 0.000313278 |
| Zfp9 | 119.4940582 | 1.067545975 | 0.284897678 | 6.90E-05 | 0.000323758 |
| Matn2 | 28.0783044 | 4.203870242 | 1.245819103 | 6.93E-05 | 0.000324981 |
| Gm19705 | 100.0284431 | 1.332302884 | 0.365380129 | 7.18E-05 | 0.000335841 |
| Adamts5 | 143.7100283 | 2.465060951 | 0.674839633 | 7.20E-05 | 0.000336288 |
| Celsr2 | 469.216707 | 1.356561622 | 0.365215887 | 7.26E-05 | 0.000338644 |
| Peg13 | 293.5557734 | 1.25390951 | 0.342633138 | 7.30E-05 | 0.000340279 |
| Klc3 | 225.384967 | 3.671962271 | 0.949307687 | 7.35E-05 | 0.000342356 |
| Frzb | 7.619930775 | 5.514693076 | 2.435586588 | 7.41E-05 | 0.000345216 |
| Gm45540 | 29.42838719 | 1.696265539 | 0.4791364 | 7.44E-05 | 0.000346286 |
| Ect2 | 151.2672817 | 1.475188691 | 0.411805496 | 7.57E-05 | 0.000352079 |
| Bambi | 173.723106 | 1.102851415 | 0.296841063 | 7.58E-05 | 0.000352463 |
| Lockd | 11.66896182 | 4.947936464 | 2.235588734 | 7.60E-05 | 0.000353204 |
| Gm9821 | 113.6853601 | 2.047164678 | 0.570309258 | 7.70E-05 | 0.000357518 |
| Col8a1 | 6.672028532 | 5.198338922 | 2.333610459 | 7.70E-05 | 0.000357766 |
| Marcks | 2298.459478 | 1.283776152 | 0.349639126 | 7.73E-05 | 0.000358898 |
| BC016579 | 79.32058624 | 2.759563182 | 0.757081436 | 7.73E-05 | 0.000358898 |
| Cdca5 | 31.9065435 | 2.790616995 | 0.796107992 | 7.80E-05 | 0.000361704 |
| Snn | 228.0071003 | 2.700528781 | 0.742194115 | 8.30E-05 | 0.000382974 |
| Notch3 | 20.13278178 | 4.334382087 | 1.325395124 | 8.34E-05 | 0.000384936 |
| Sulf1 | 14.0474772 | 5.212491051 | 2.337376205 | 8.60E-05 | 0.000396015 |
| Tcea2 | 81.58101803 | 1.634444584 | 0.457148174 | 8.62E-05 | 0.000396841 |
| Crb2 | 43.57258625 | 4.641396163 | 2.115706026 | 8.63E-05 | 0.000397224 |
| Adam32 | 16.57665323 | 3.328520108 | 0.936649261 | 8.71E-05 | 0.000400687 |
| Lrrc75b | 146.8020911 | 4.44132203 | 2.022347134 | 8.76E-05 | 0.000402621 |
| Col6a5 | 5.27860694 | 5.336946052 | 2.410794677 | 8.88E-05 | 0.00040764 |
| Lpcat4 | 351.3912223 | 2.642126514 | 0.726976157 | 9.15E-05 | 0.000419703 |
| Ttc22 | 103.1412851 | 4.454241054 | 2.043216439 | 9.35E-05 | 0.000427996 |
| Cenpi | 25.55071118 | 4.605654407 | 2.117653678 | 9.46E-05 | 0.000432873 |
| Mis18bp1 | 53.623487 | 1.884096064 | 0.536832277 | 9.50E-05 | 0.000433724 |
| Nbea | 139.2233913 | 1.514836417 | 0.42309152 | 9.56E-05 | 0.000436315 |
| Kif2c | 46.20484629 | 2.194906891 | 0.633195654 | 9.62E-05 | 0.000438689 |

| | | | | | |
|---|---|---|---|---|---|
| Spsb1 | 115.0903515 | 4.037087136 | 1.220679079 | 9.62E-05 | 0.000438689 |
| Ttc16 | 26.38531768 | 2.291194574 | 0.667647435 | 9.71E-05 | 0.000442314 |
| 9330162G02Rik | 47.41581936 | 1.549350474 | 0.441743903 | 9.73E-05 | 0.000443218 |
| Ubtd2 | 107.105615 | 1.196949505 | 0.328895564 | 9.82E-05 | 0.000446953 |
| Spa17 | 49.73769875 | 1.812983628 | 0.514927607 | 9.96E-05 | 0.000452692 |
| 1700016C15Rik | 8.044629714 | 4.948298898 | 2.271653566 | 0.000100195 | 0.000455325 |
| Ndc80 | 42.83197799 | 4.747139156 | 2.17572357 | 0.000100536 | 0.000456554 |
| Cyp1b1 | 20.68097439 | 4.57684133 | 2.12264414 | 0.000100746 | 0.000457289 |
| Gm17415 | 11.77397772 | 3.178763143 | 0.891040212 | 0.000101242 | 0.000459103 |
| Plppr3 | 87.84527235 | 1.433314874 | 0.407133111 | 0.000101998 | 0.000462207 |
| Lrrn4 | 7.565095521 | 4.54440529 | 1.441773115 | 0.000103822 | 0.000469915 |
| Gm15895 | 30.23636599 | 4.739754909 | 2.172001399 | 0.000104349 | 0.000472081 |
| Parvb | 16.27627889 | 2.351840247 | 0.698821571 | 0.000104932 | 0.00047443 |
| Fut10 | 98.96711906 | 1.30026216 | 0.363210066 | 0.000105074 | 0.000474913 |
| Gm43305 | 2986.452655 | 2.847499225 | 0.903791098 | 0.000105829 | 0.000477986 |
| Depdc1a | 21.46708159 | 4.744392009 | 2.188530546 | 0.000105886 | 0.00047813 |
| Per2 | 1000.550869 | 1.529209143 | 0.444074933 | 0.000107229 | 0.000483626 |
| Gm44284 | 23.56829357 | 4.409002928 | 1.396226324 | 0.000107523 | 0.000484837 |
| Gpm6b | 96.80940341 | 4.373673639 | 2.028448829 | 0.000108789 | 0.000490317 |
| Pdlim2 | 73.82258096 | 3.583402941 | 0.958636801 | 0.00011065 | 0.00049777 |
| Gm36026 | 15.10109659 | 3.149752772 | 0.88829703 | 0.000111963 | 0.000502969 |
| Gm50232 | 19.89243577 | 4.560505768 | 2.127111527 | 0.000115555 | 0.000517528 |
| Gm38393 | 50.05274479 | 1.321587778 | 0.372370135 | 0.000115621 | 0.000517705 |
| Gm45774 | 8.920323414 | 4.700211091 | 1.525704843 | 0.000115892 | 0.000518557 |
| St6galnac2 | 262.5772668 | 4.418576657 | 2.047050308 | 0.000118995 | 0.000531323 |
| Chtf18 | 70.2161767 | 1.951786679 | 0.574200332 | 0.000121612 | 0.000542504 |
| Dio3os | 128.3888857 | 1.794087228 | 0.520845071 | 0.000121909 | 0.000543701 |
| Zscan30 | 39.37612708 | 2.228497499 | 0.638811649 | 0.0001224 | 0.000545386 |
| Gm3788 | 51.63671794 | 4.38484346 | 2.053363012 | 0.000122997 | 0.000547665 |
| Trmt13 | 125.3649653 | 1.028019608 | 0.290629983 | 0.000123061 | 0.000547822 |
| Gm31814 | 27.73860891 | 1.581301433 | 0.46171801 | 0.000123852 | 0.000551214 |
| Asf1b | 32.78993591 | 2.150573822 | 0.638934885 | 0.000124803 | 0.000555191 |
| Galnt12 | 115.1086543 | 4.299563011 | 2.015229114 | 0.000125597 | 0.000558334 |
| Fam72a | 25.79667033 | 1.859612221 | 0.551547261 | 0.000126569 | 0.000562264 |
| Angptl8 | 1888.43215 | 2.566213761 | 0.813765313 | 0.000128146 | 0.000569009 |
| Gm5148 | 80.88554426 | 1.58037271 | 0.45214093 | 0.000128294 | 0.000569366 |
| Nkd2 | 5.213609525 | 4.93956301 | 2.312316746 | 0.000128316 | 0.000569366 |
| Gm43127 | 35.73598559 | 2.728310684 | 0.789722083 | 0.000130001 | 0.000576045 |
| Rem2 | 130.7020789 | 2.350168162 | 0.689238099 | 0.000130332 | 0.000577377 |
| Trpc1 | 71.14294356 | 2.909527067 | 0.808461921 | 0.000131356 | 0.000581376 |
| 4732463B04Rik | 26.17044548 | 1.622573644 | 0.476397612 | 0.000131648 | 0.000582266 |
| Mark1 | 55.00052415 | 4.316740209 | 2.038171077 | 0.000132628 | 0.000586201 |
| Gm37558 | 6.569173257 | 4.934093029 | 2.306341382 | 0.000132608 | 0.000586201 |

| | | | | | |
|---|---|---|---|---|---|
| Mast1 | 97.858653 | 1.406320962 | 0.409173263 | 0.000138107 | 0.000607583 |
| Npb | 27.61570628 | 1.89955837 | 0.575384208 | 0.00013824 | 0.000607922 |
| Wdr31 | 110.092143 | 2.322090029 | 0.66483858 | 0.000138454 | 0.000608583 |
| Scn2a | 11.20260729 | 4.181356306 | 1.334667366 | 0.000138512 | 0.000608697 |
| Plcg2 | 222.8589234 | 3.495122816 | 0.949988045 | 0.000141091 | 0.000618754 |
| Cdsn | 164.7328046 | 4.257224853 | 2.004960323 | 0.00014149 | 0.000620366 |
| Arhgef39 | 24.56724148 | 4.10671061 | 1.308602119 | 0.000141776 | 0.000621335 |
| Gpr137b | 91.55136536 | 3.517470816 | 0.962538692 | 0.000143876 | 0.000629533 |
| 6330403L08Rik | 85.06653915 | 1.287983631 | 0.371464071 | 0.000144776 | 0.000633326 |
| Garnl3 | 107.4779596 | 2.347902753 | 0.678044484 | 0.000145536 | 0.00063578 |
| Ptpn14 | 1536.555308 | 2.196393577 | 0.685148024 | 0.000147431 | 0.000642449 |
| Cdr2l | 152.11532 | 3.449164652 | 0.938566671 | 0.000151945 | 0.000659723 |
| Oas1g | 50.4381167 | 2.2548192 | 0.66445428 | 0.00015267 | 0.000662423 |
| Large2 | 252.7174892 | 1.331213503 | 0.384648762 | 0.000155474 | 0.000673675 |
| Cenps | 15.31394597 | 4.639400368 | 2.199704832 | 0.000155664 | 0.000674345 |
| Spdl1 | 52.38987669 | 1.583359202 | 0.464457238 | 0.000158146 | 0.000683707 |
| Steap1 | 60.49469526 | 2.358104293 | 0.689547222 | 0.000159736 | 0.00068965 |
| Gm11944 | 34.30504677 | 1.964514097 | 0.578184362 | 0.000162742 | 0.000701996 |
| Gm43808 | 17.80249199 | 1.891615271 | 0.565271052 | 0.000167708 | 0.000720986 |
| Gdpd5 | 486.8092151 | 2.267943333 | 0.657527448 | 0.00016833 | 0.000723116 |
| Snhg16 | 54.79123552 | 1.203635608 | 0.34466447 | 0.000168436 | 0.000723306 |
| Prph | 21.41619843 | 3.579775318 | 1.009880463 | 0.000169019 | 0.000725647 |
| Gm38228 | 24.01816397 | 4.357292847 | 2.0930472 | 0.000169061 | 0.000725666 |
| Aurkb | 72.99132965 | 2.366192265 | 0.744734999 | 0.000170316 | 0.000730725 |
| Thsd4 | 1093.608678 | 4.119139809 | 1.957947268 | 0.000172202 | 0.000737169 |
| Nkain1 | 86.4395535 | 4.194625175 | 2.014476342 | 0.000172918 | 0.000739571 |
| Kctd11 | 141.6557435 | 1.074227041 | 0.306066706 | 0.000177011 | 0.000755899 |
| Ly6k | 6.819531188 | 5.264081168 | 2.478612815 | 0.000177533 | 0.000757961 |
| 6530402F18Rik | 150.7293879 | 3.785671412 | 1.199593014 | 0.000177574 | 0.000757966 |
| Slc44a2 | 994.903984 | 1.490102595 | 0.433630126 | 0.00017829 | 0.000760855 |
| Cacnb3 | 476.2427147 | 3.742243613 | 1.180777891 | 0.000178714 | 0.000762331 |
| Plk4 | 99.29275816 | 1.125724562 | 0.322471137 | 0.000178965 | 0.000763058 |
| Ptrh1 | 72.44651728 | 1.027286409 | 0.292088967 | 0.000179637 | 0.000765751 |
| Pnldc1 | 15.22607229 | 1.980428525 | 0.613730608 | 0.0001826 | 0.000777862 |
| Atad2 | 306.9481756 | 1.097049113 | 0.315885449 | 0.000182658 | 0.00077794 |
| Dusp9 | 43.86067883 | 2.895207149 | 0.836294974 | 0.000184749 | 0.000786322 |
| A4galt | 97.65944013 | 4.225193843 | 2.030577564 | 0.000185407 | 0.00078877 |
| Nr4a1 | 1954.582923 | 2.788735402 | 0.942083911 | 0.00018571 | 0.000789884 |
| 2510016D11Rik | 37.99216227 | 1.349304227 | 0.394875491 | 0.00018803 | 0.000799045 |
| Fhdc1 | 81.407836 | 3.807565433 | 1.217906186 | 0.000188674 | 0.000801602 |
| Ydjc | 39.60354317 | 1.886485074 | 0.570029133 | 0.000189047 | 0.000803012 |
| Ptk7 | 626.2119198 | 3.733561695 | 1.1870352 | 0.000189419 | 0.000804338 |
| P3h3 | 160.2122771 | 2.835271768 | 0.815784886 | 0.000192404 | 0.000815283 |

| | | | | | |
|---|---|---|---|---|---|
| Atp8a1 | 174.8818103 | 1.0838655 | 0.312229356 | 0.000196009 | 0.000828913 |
| Boc | 14.34893349 | 4.043758977 | 1.335636512 | 0.000196722 | 0.000831744 |
| Cebpd | 140.1707693 | 1.021620137 | 0.293730865 | 0.000199032 | 0.000840544 |
| Hand2 | 6.578559141 | 5.032955692 | 2.406970081 | 0.000199379 | 0.00084168 |
| Gm29585 | 10.59441738 | 4.364099295 | 2.137643924 | 0.000208264 | 0.000876451 |
| Chst8 | 81.45846187 | 4.292007321 | 2.093078685 | 0.000212488 | 0.000893484 |
| Atp6v0e2 | 282.3084529 | 2.467473428 | 0.729077304 | 0.000219482 | 0.000920474 |
| 1700003F12Rik | 21.4230467 | 3.040561062 | 0.922225508 | 0.000221144 | 0.000927037 |
| Zfp677 | 79.17358079 | 1.509598263 | 0.450530773 | 0.000223076 | 0.000934727 |
| Kif15 | 56.74713637 | 3.473541805 | 1.004609571 | 0.000224224 | 0.000938924 |
| Muc13 | 6.186834385 | 4.51198505 | 2.219190748 | 0.00022468 | 0.000940627 |
| Gm28172 | 20.36134357 | 1.861197377 | 0.57677776 | 0.000227106 | 0.000949748 |
| Prr11 | 45.58237529 | 3.8174744 | 1.254586628 | 0.000229108 | 0.000957692 |
| Nmb | 19.11236823 | 3.439289839 | 0.989778937 | 0.000229851 | 0.000960179 |
| Nr4a2 | 47.60476213 | 2.166437593 | 0.673827259 | 0.000235309 | 0.00098107 |
| Gm20548 | 16.93634852 | 2.420750813 | 0.747290401 | 0.000238431 | 0.000992997 |
| Rbms3 | 396.3315416 | 3.336771051 | 0.948807727 | 0.000239048 | 0.000995133 |
| Ncaph | 125.5472099 | 1.525362395 | 0.467473513 | 0.00023953 | 0.000996709 |
| Gm48427 | 22.69402501 | 4.352576182 | 2.146063481 | 0.000243518 | 0.00101111 |
| Parpbp | 28.47248195 | 3.921106439 | 1.31709818 | 0.000243739 | 0.001011689 |
| Gem | 69.64046124 | 4.083679592 | 2.02029354 | 0.000244984 | 0.0010161 |
| C230096K16Rik | 29.55870902 | 1.949012737 | 0.62181121 | 0.00024768 | 0.001025951 |
| Bicdl1 | 120.5742969 | 1.141034061 | 0.339660993 | 0.000248227 | 0.001027551 |
| Trim6 | 32.17250932 | 3.059169581 | 0.949664181 | 0.000257373 | 0.001062888 |
| Cdc7 | 29.52995471 | 4.308817769 | 2.136197697 | 0.000258296 | 0.00106647 |
| Tnc | 12.4546839 | 4.210448024 | 1.466721768 | 0.000258844 | 0.001068506 |
| Ercc6l | 41.86769784 | 1.946055722 | 0.599582996 | 0.000260341 | 0.001073991 |
| Tcf21 | 5.002477697 | 4.773639129 | 2.355671774 | 0.000260754 | 0.00107523 |
| Egr3 | 456.9616079 | 5.800917185 | 2.764170962 | 0.000266838 | 0.00109843 |
| Gm45853 | 12.19484937 | 4.287710069 | 2.14278607 | 0.000267863 | 0.001102417 |
| 3010003L21Rik | 29.82646201 | 2.430314948 | 0.762047795 | 0.000268298 | 0.001103968 |
| Rmi2 | 11.1887712 | 4.396994049 | 2.195117233 | 0.000272109 | 0.001118451 |
| Mthfd2 | 70.40004096 | 1.184509215 | 0.355695284 | 0.000275921 | 0.001132178 |
| Gm11419 | 30.20609575 | 1.469205525 | 0.455974319 | 0.000277334 | 0.001137734 |
| Gm44321 | 27.87095241 | 2.510529602 | 0.768348555 | 0.000280299 | 0.001149159 |
| Pgm5 | 36.6665419 | 3.696045059 | 1.23416183 | 0.000286036 | 0.001170928 |
| Nbl1 | 36.84218494 | 4.449931653 | 2.2138775 | 0.000289879 | 0.001184451 |
| Ace | 88.97615753 | 2.287687683 | 0.734055988 | 0.000293189 | 0.001197148 |
| Mt2 | 13.09228646 | 3.20083719 | 1.075183315 | 0.000297118 | 0.001211643 |
| Scel | 34.66803291 | 4.018410485 | 2.028182436 | 0.000297627 | 0.001213204 |
| 1700007J10Rik | 12.75055289 | 4.131833476 | 2.095759956 | 0.000303867 | 0.001237062 |
| Nr2c2ap | 24.6422567 | 1.796407508 | 0.574415994 | 0.000304025 | 0.001237256 |
| Arhgef25 | 326.3183505 | 2.261424631 | 0.703408279 | 0.000304084 | 0.001237256 |

| | | | | | |
|---|---|---|---|---|---|
| Upp1 | 82.66157653 | 1.149523673 | 0.346030464 | 0.00031146 | 0.001265292 |
| B130055M24Rik | 43.2211138 | 1.753339593 | 0.553970968 | 0.000312596 | 0.00126883 |
| Siglecf | 4.047078919 | 4.879502896 | 2.442061021 | 0.000313604 | 0.001272114 |
| Neil3 | 28.39842296 | 4.779099327 | 2.373014265 | 0.000314961 | 0.001277079 |
| B3glct | 159.1523964 | 1.030114007 | 0.306091039 | 0.000316001 | 0.001281025 |
| Rcan1 | 613.603878 | 1.209300018 | 0.370152873 | 0.000319722 | 0.001295289 |
| Polq | 44.58854552 | 3.720043565 | 1.26898555 | 0.000320888 | 0.001299192 |
| Bard1 | 44.89485022 | 3.441873466 | 1.044041626 | 0.000322388 | 0.001304163 |
| Abcb1b | 71.35472353 | 1.415740237 | 0.443232108 | 0.000322534 | 0.001304478 |
| Fam131a | 54.12628782 | 3.631582854 | 1.225959338 | 0.000323209 | 0.001306932 |
| Gm43197 | 66.11778939 | 3.605310785 | 1.214086188 | 0.000325958 | 0.001316661 |
| Fst | 416.2013913 | 1.073652265 | 0.323528356 | 0.000326523 | 0.00131839 |
| Gm44164 | 19.09565313 | 3.377697696 | 1.013776513 | 0.000326937 | 0.001319781 |
| Lif | 633.8391944 | 1.679832 | 0.545848409 | 0.000328739 | 0.001326499 |
| Mllt11 | 89.3895556 | 1.151215389 | 0.346930165 | 0.000331691 | 0.001337284 |
| Gm42659 | 25.12638908 | 1.652249801 | 0.519459004 | 0.000331633 | 0.001337284 |
| Dkk3 | 77.99882565 | 3.268664269 | 0.96466029 | 0.0003339 | 0.001345798 |
| Cyp2j9 | 248.4499369 | 1.058869196 | 0.31896594 | 0.000336516 | 0.001355029 |
| Troap | 31.33847961 | 4.188332204 | 2.131107652 | 0.000340532 | 0.00136919 |
| Gm15567 | 16.48719522 | 2.129968966 | 0.688402158 | 0.000341924 | 0.001373925 |
| Chsy1 | 142.2913045 | 1.325597505 | 0.412665385 | 0.000343778 | 0.00137964 |
| Jdp2 | 98.06765882 | 2.419536176 | 0.753329795 | 0.000349357 | 0.001399811 |
| Itga8 | 15.07959948 | 3.338958269 | 1.118376631 | 0.000350295 | 0.001402568 |
| Ncapd2 | 462.4268365 | 1.052547442 | 0.31848714 | 0.000350538 | 0.001403248 |
| Ksr1 | 84.52119396 | 2.073329986 | 0.655307812 | 0.000353379 | 0.001412852 |
| D130040H23Rik | 62.39691195 | 3.237048187 | 0.954997999 | 0.00035369 | 0.001413804 |
| Dnajb1 | 703.3704621 | 1.096400512 | 0.333845878 | 0.000356297 | 0.001423334 |
| Aldh1a2 | 206.6968837 | 3.65604912 | 1.260043936 | 0.000357532 | 0.001427675 |
| Lama2 | 14.06551884 | 3.712463857 | 1.202108199 | 0.000363631 | 0.00145082 |
| Pole2 | 45.15455352 | 2.445888777 | 0.770663329 | 0.000374523 | 0.001490867 |
| Iqank1 | 53.56789908 | 1.390456655 | 0.432397866 | 0.00038238 | 0.001519305 |
| Mst1r | 60.31536141 | 1.551694049 | 0.496795195 | 0.000382473 | 0.001519361 |
| Mex3b | 87.36450112 | 2.708010662 | 0.836830615 | 0.000382721 | 0.001520032 |
| Sftpa1 | 26.40572689 | 1.521896962 | 0.489947769 | 0.000385301 | 0.001529645 |
| G930009F23Rik | 214.0800398 | 3.514190947 | 1.197949074 | 0.000387544 | 0.001537916 |
| Kif23 | 174.949481 | 3.216002648 | 0.959742291 | 0.000387663 | 0.001538069 |
| Gm12592 | 13.80618582 | 4.112772623 | 2.129258331 | 0.00038963 | 0.001545553 |
| H1f10 | 92.64661293 | 2.471432532 | 0.787186966 | 0.000391891 | 0.001552599 |
| Ar | 47.5156975 | 1.459913014 | 0.469019244 | 0.000393407 | 0.001557639 |
| Optc | 3.987759323 | 4.347394202 | 2.254499592 | 0.000395473 | 0.001563883 |
| Chek1 | 71.09400761 | 1.660357589 | 0.539235086 | 0.00039729 | 0.001569773 |
| Ablim2 | 64.02134382 | 1.005469867 | 0.304024667 | 0.000397563 | 0.001570528 |
| Carmil3 | 73.0246665 | 3.257980006 | 0.988408669 | 0.000400386 | 0.001581031 |

| | | | | | |
|---|---|---|---|---|---|
| Scarf2 | 441.453877 | 3.396920372 | 1.06679351 | 0.000402558 | 0.001588625 |
| Depdc1b | 11.88848391 | 4.089841133 | 2.134316432 | 0.000413348 | 0.001628528 |
| Phf21b | 24.1227677 | 3.264435191 | 0.993623823 | 0.000416879 | 0.001641092 |
| Cdca7 | 70.59290676 | 1.27767643 | 0.401706427 | 0.000421828 | 0.001658392 |
| B3gnt2 | 75.77479875 | 1.009550206 | 0.307172344 | 0.00042238 | 0.001660024 |
| Ankrd9 | 115.9023408 | 1.094610146 | 0.338022223 | 0.000423418 | 0.00166342 |
| Bex1 | 110.4057192 | 3.859801255 | 2.007966645 | 0.000423665 | 0.001664051 |
| Dbn1 | 703.0422279 | 3.449008605 | 1.185078544 | 0.000424392 | 0.001666567 |
| Dubr | 99.05120722 | 2.016518102 | 0.645114569 | 0.000427153 | 0.001676035 |
| Sema6c | 85.88822162 | 2.465511597 | 0.797026152 | 0.000430033 | 0.001686304 |
| 9530034E10Rik | 9.284664292 | 2.390121399 | 0.818819893 | 0.000433379 | 0.001698728 |
| Mmgt2 | 103.1822023 | 1.133796904 | 0.348850746 | 0.000434842 | 0.00170342 |
| Gm17690 | 48.54150409 | 1.480674357 | 0.478903197 | 0.000435754 | 0.001706642 |
| Hamp | 4920.090558 | 1.291498587 | 0.409917364 | 0.000439904 | 0.00172009 |
| Gm16617 | 16.29209584 | 3.921248534 | 2.061523407 | 0.000441913 | 0.001727595 |
| Gm37856 | 8.856355769 | 4.188515577 | 2.194537395 | 0.000442395 | 0.001729126 |
| Kirrel | 803.298023 | 3.47667109 | 1.209375451 | 0.000443875 | 0.001734555 |
| Sgo1 | 33.78505758 | 3.347255018 | 1.053034736 | 0.000451017 | 0.001761749 |
| Ltbp1 | 91.28606414 | 3.513683026 | 1.230046897 | 0.000452972 | 0.001769025 |
| Pcdh19 | 127.6479121 | 3.795037163 | 1.995461242 | 0.000456812 | 0.001783296 |
| BB123696 | 4.212275409 | 4.607753136 | 2.408400891 | 0.00045706 | 0.0017839 |
| Peg10 | 20.65387201 | 1.613964069 | 0.528885693 | 0.000458507 | 0.001789185 |
| Ranbp17 | 84.76697498 | 1.931145519 | 0.626543155 | 0.000462608 | 0.001804088 |
| C1qtnf12 | 64.81150577 | 1.715607183 | 0.56178119 | 0.000463183 | 0.001805716 |
| D430020J02Rik | 69.9641887 | 2.663264382 | 0.839235204 | 0.000464759 | 0.001811369 |
| Auts2 | 442.3884988 | 3.724642788 | 1.946367405 | 0.000466038 | 0.001815985 |
| Gm26586 | 14.39900978 | 2.833279619 | 0.921136734 | 0.000470829 | 0.001831371 |
| Misp3 | 12.44467478 | 2.752171788 | 0.944456219 | 0.000476177 | 0.001850235 |
| Zfp37 | 74.42362999 | 3.488630142 | 1.228405876 | 0.000480404 | 0.001864772 |
| Artn | 15.14717652 | 4.242503633 | 2.234407233 | 0.000482556 | 0.001872368 |
| 2900089D17Rik | 27.76272719 | 1.98737669 | 0.674141967 | 0.000485619 | 0.001883111 |
| Slc4a3 | 871.2598589 | 3.430091056 | 1.200364186 | 0.000486174 | 0.001884503 |
| A630001O12Rik | 7.320688426 | 3.704038181 | 1.341745848 | 0.000486131 | 0.001884503 |
| Rasl10a | 91.96391447 | 1.841673538 | 0.602756181 | 0.00049893 | 0.001929271 |
| Hspa12a | 46.63508459 | 1.628114205 | 0.531862259 | 0.000503582 | 0.001945691 |
| Lhfp | 342.1030529 | 2.416698601 | 0.880802991 | 0.00050415 | 0.001947126 |
| Plekha4 | 81.79254494 | 1.272039609 | 0.402504711 | 0.000504157 | 0.001947126 |
| Astn2 | 3.85465702 | 4.382164101 | 2.324150612 | 0.000504265 | 0.001947154 |
| Chrdl2 | 38.3528237 | 3.466596898 | 1.226052403 | 0.000506379 | 0.001954356 |
| Ankle1 | 32.154782 | 3.613874928 | 1.306800411 | 0.000506864 | 0.001955616 |
| Tnfaip8 | 301.0220114 | 3.109903551 | 0.95122495 | 0.000511191 | 0.001970723 |
| Tmem130 | 134.6827915 | 3.738716791 | 1.982608101 | 0.000511435 | 0.00197127 |
| B430203G13Rik | 8.558409291 | 4.045823071 | 2.155934415 | 0.00051821 | 0.001995377 |

| | | | | | |
|---|---|---|---|---|---|
| 1700058P15Rik | 21.23741341 | 2.679941397 | 0.861338575 | 0.000521019 | 0.002005792 |
| Knl1 | 59.67721292 | 4.108122756 | 1.589763836 | 0.000534328 | 0.002054144 |
| Pde6c | 11.66684245 | 3.008855076 | 1.031572644 | 0.000539256 | 0.002071865 |
| Gucy1a1 | 10.65437665 | 3.50292482 | 1.176964352 | 0.000558834 | 0.002141061 |
| Plscr4 | 45.94166742 | 2.686415134 | 0.877161124 | 0.000564846 | 0.002161938 |
| Wnt5a | 178.9172445 | 1.210647404 | 0.386608535 | 0.000568918 | 0.002176655 |
| Gm45221 | 51.30545664 | 1.227740685 | 0.393199567 | 0.000573279 | 0.00219072 |
| Tgfb3 | 391.4622532 | 3.626623046 | 1.945069553 | 0.000575856 | 0.002199769 |
| Gdf10 | 13.69962413 | 2.744937691 | 0.996773593 | 0.000576393 | 0.00220043 |
| Pwwp4c | 12.896929 | 3.506202603 | 1.273099284 | 0.000580531 | 0.002215787 |
| D730005E14Rik | 39.29941939 | 3.85075685 | 2.074776825 | 0.000581981 | 0.002220436 |
| Gm45244 | 6.694165504 | 2.877844892 | 0.976617744 | 0.000585267 | 0.0022312 |
| Colca2 | 45.28046216 | 2.666956803 | 0.874418504 | 0.000587747 | 0.00224021 |
| Zfp185 | 247.3816632 | 3.625537447 | 1.953010856 | 0.00058955 | 0.002246634 |
| Lrrc75a | 30.08233169 | 2.660414309 | 0.869872464 | 0.000590944 | 0.002251054 |
| Rptoros | 8.135022389 | 3.45880598 | 1.160261709 | 0.000596357 | 0.002270319 |
| Cox7a1 | 30.53113275 | 1.441669585 | 0.478060755 | 0.000601899 | 0.002290509 |
| Gm16184 | 56.58035589 | 2.287484768 | 0.74749046 | 0.000602983 | 0.002294179 |
| Map1a | 24.78475105 | 3.890268459 | 2.11139732 | 0.000609415 | 0.00231544 |
| Socs3 | 416.64382 | 1.96173464 | 0.708915567 | 0.000613148 | 0.002328699 |
| Zfp462 | 599.0929267 | 1.793595923 | 0.589455444 | 0.000615332 | 0.002336069 |
| 2310040G24Rik | 80.17449762 | 1.009887865 | 0.318543078 | 0.000622617 | 0.002361859 |
| Cldn34c1 | 41.91100176 | 1.75488273 | 0.581466347 | 0.000627606 | 0.002379844 |
| Ubxn11 | 90.05028327 | 3.651332742 | 1.985447912 | 0.000629273 | 0.002385692 |
| Fut1 | 14.15410487 | 3.573236467 | 1.329876423 | 0.00063157 | 0.002392039 |
| Rin1 | 181.2400013 | 2.249603125 | 0.735986957 | 0.000633059 | 0.002396261 |
| Rcn3 | 66.14532956 | 2.389016253 | 0.804202176 | 0.000634786 | 0.002401377 |
| Scn7a | 701.9877249 | 3.586881918 | 1.94559045 | 0.000643911 | 0.002434455 |
| Smtnl2 | 266.6309909 | 1.191404791 | 0.380090473 | 0.000645986 | 0.002440857 |
| Fanci | 41.16299358 | 2.751759434 | 0.937223708 | 0.00065636 | 0.002477131 |
| Gm44899 | 10.47805105 | 3.927268544 | 2.147069522 | 0.000657465 | 0.002480812 |
| Plxdc2 | 199.3478982 | 3.996729844 | 2.169430284 | 0.000658056 | 0.002482065 |
| Adm | 12.13534652 | 2.750375068 | 0.934248518 | 0.000664367 | 0.002504392 |
| Mfap4 | 4.013781849 | 4.302855089 | 2.344912698 | 0.000667929 | 0.00251535 |
| Cenph | 30.96775385 | 2.173733869 | 0.740486137 | 0.000673672 | 0.002534989 |
| Mbd4 | 101.9419594 | 1.184206411 | 0.380691958 | 0.000676799 | 0.002546254 |
| Snai3 | 5.707285013 | 4.591066168 | 2.485037807 | 0.000680573 | 0.002558951 |
| Nrarp | 30.05384366 | 3.753444103 | 2.061078192 | 0.000681645 | 0.002562478 |
| Sarm1 | 77.55475636 | 1.720647581 | 0.593420207 | 0.000683375 | 0.002567471 |
| Cmya5 | 7.184232572 | 3.888521548 | 2.148685967 | 0.000703403 | 0.002638588 |
| Gm44949 | 24.90828829 | 1.991991512 | 0.70196196 | 0.000707877 | 0.002653775 |
| Ccsap | 22.63345117 | 2.212806074 | 0.762674306 | 0.000708005 | 0.002653775 |
| Gm17359 | 32.00249849 | 1.885467504 | 0.656048129 | 0.000708172 | 0.002653882 |

| | | | | | |
|---|---|---|---|---|---|
| Htra3 | 41.1353314 | 3.035577171 | 0.968061471 | 0.000715569 | 0.002678987 |
| Gm7628 | 45.33476507 | 1.279408742 | 0.418554064 | 0.000717058 | 0.002684036 |
| Rgs11 | 123.9733222 | 3.068821298 | 0.990800126 | 0.000725884 | 0.002714958 |
| Rbm38 | 93.73252033 | 1.148969037 | 0.372920227 | 0.000729701 | 0.002726577 |
| Dnah7b | 249.2872674 | 3.272020142 | 1.197509464 | 0.000730056 | 0.002727369 |
| Gm37818 | 10.70577781 | 3.517524119 | 1.33553252 | 0.000741078 | 0.002764781 |
| Gm15911 | 7.192873984 | 3.985535775 | 2.210174212 | 0.000750851 | 0.00279806 |
| Kif14 | 36.35849824 | 3.361800028 | 1.252236111 | 0.000751157 | 0.002798576 |
| Prtg | 145.4384458 | 3.521488407 | 1.949911529 | 0.000751612 | 0.002798641 |
| Eme1 | 16.47134209 | 3.839561993 | 2.132899902 | 0.00075141 | 0.002798641 |
| Spire1 | 47.906106 | 1.824673034 | 0.619366174 | 0.000764207 | 0.002838928 |
| Sfxn3 | 763.4394252 | 2.499499007 | 0.83161615 | 0.00077763 | 0.002883767 |
| Atp8b2 | 593.136272 | 1.28136368 | 0.421555574 | 0.000779666 | 0.002890759 |
| Antxr1 | 37.66410249 | 1.940398789 | 0.663172029 | 0.00079631 | 0.002943939 |
| Gm29361 | 23.06675745 | 1.390265069 | 0.464864885 | 0.000798722 | 0.002951719 |
| Tekt2 | 12.19176898 | 3.950652773 | 2.206458154 | 0.000805758 | 0.002975429 |
| Doc2g | 42.89293358 | 1.087826841 | 0.352673309 | 0.0008077 | 0.002982028 |
| Hyls1 | 52.89767621 | 1.011371975 | 0.326360108 | 0.000815255 | 0.003007316 |
| Slc9a5 | 110.2848241 | 3.232383822 | 1.198778237 | 0.0008201 | 0.003023152 |
| Tnxb | 33.73380439 | 1.570910683 | 0.553024636 | 0.000822493 | 0.003031199 |
| Lor | 22.55616014 | 1.558363134 | 0.535324512 | 0.000831152 | 0.003056262 |
| Chaf1a | 127.8561661 | 1.090152746 | 0.358333794 | 0.000835002 | 0.003068658 |
| Fbln1 | 14.84826511 | 2.435478475 | 0.897370998 | 0.000836131 | 0.003072219 |
| Tox3 | 113.4933192 | 3.505747748 | 1.97308373 | 0.000851064 | 0.003119918 |
| Kitl | 1002.121375 | 1.623172765 | 0.556824509 | 0.000858666 | 0.003143582 |
| Gm15347 | 27.08999816 | 3.554361676 | 2.010340627 | 0.000859382 | 0.003145568 |
| Gm44202 | 9.146276607 | 1.89737111 | 0.682351853 | 0.000860312 | 0.003147207 |
| Cdca2 | 48.40059752 | 2.39929546 | 0.912361745 | 0.00086521 | 0.003163314 |
| 4833438C02Rik | 49.28965929 | 1.43245428 | 0.4937364 | 0.00087748 | 0.003203293 |
| Sorbs2os | 45.74983092 | 1.780675004 | 0.611156667 | 0.000880729 | 0.003212712 |
| Gm18228 | 31.06247993 | 1.708564916 | 0.605744959 | 0.000882002 | 0.003216741 |
| Srpx2 | 4.138825762 | 4.372707776 | 2.453295238 | 0.000885398 | 0.0032279 |
| Gm5577 | 15.21096952 | 3.714293144 | 2.10681614 | 0.000890417 | 0.003244965 |
| Orc1 | 17.1672895 | 3.644918036 | 2.074453072 | 0.000900186 | 0.003279321 |
| Pmaip1 | 184.9748214 | 2.485884204 | 0.841084499 | 0.000901339 | 0.003282277 |
| 2610020C07Rik | 24.42932108 | 1.864766313 | 0.677068864 | 0.000920472 | 0.003343703 |
| Fos | 8435.534606 | 1.229709011 | 0.420985756 | 0.00092466 | 0.003357645 |
| Pkhd1l1 | 12.58094383 | 3.930543532 | 2.238599619 | 0.000925201 | 0.003358337 |
| Zgrf1 | 75.24032242 | 1.006605419 | 0.327214264 | 0.000931883 | 0.003378759 |
| 5430405H02Rik | 21.11507078 | 1.884269282 | 0.668107369 | 0.000948735 | 0.003431429 |
| Ccdc116 | 11.65287636 | 3.745154944 | 2.145425389 | 0.000953083 | 0.003444559 |
| Synm | 50.19422357 | 1.373094735 | 0.4701663 | 0.000957617 | 0.003458062 |
| Mxd3 | 33.93526848 | 2.097844141 | 0.740067387 | 0.000962731 | 0.00347289 |

| | | | | | |
|---|---|---|---|---|---|
| Barx2 | 11.63264111 | 4.223760957 | 1.820495341 | 0.000964373 | 0.003478158 |
| Gpr17 | 50.16404841 | 1.163872611 | 0.392283438 | 0.000965317 | 0.003480909 |
| Il15 | 62.20420724 | 1.042596004 | 0.342841401 | 0.000982407 | 0.003537883 |
| Mex3a | 576.340449 | 2.884834276 | 0.946170168 | 0.000988272 | 0.003555413 |
| Hdx | 69.25697737 | 3.489501557 | 2.001147098 | 0.000992175 | 0.003567033 |
| Gm17300 | 15.38577159 | 2.641759987 | 1.002956119 | 0.001009552 | 0.003624753 |
| Nbdy | 58.24988229 | 1.049426055 | 0.346608513 | 0.001012095 | 0.003632527 |
| Flrt1 | 43.12652168 | 2.516354136 | 0.877283579 | 0.001015039 | 0.003642411 |
| 6230400D17Rik | 37.48373846 | 1.291358675 | 0.443285417 | 0.001024481 | 0.003674234 |
| Kif11 | 135.2306519 | 1.39910746 | 0.493088327 | 0.001047027 | 0.003744594 |
| R74862 | 31.01268452 | 1.872080323 | 0.656947953 | 0.001052967 | 0.003761633 |
| Atp1a2 | 5.776526169 | 3.806817459 | 2.209796603 | 0.001060922 | 0.00378723 |
| Gm830 | 656.8243657 | 3.099701072 | 1.186349281 | 0.001069299 | 0.003811465 |
| Sh3bgr | 16.84616092 | 3.716590303 | 2.154657504 | 0.001072079 | 0.003819952 |
| Noxa1 | 59.2643073 | 1.278163459 | 0.441250324 | 0.001080331 | 0.003845786 |
| Ak7 | 13.38465702 | 1.977907005 | 0.738008858 | 0.001082961 | 0.003853718 |
| Cd34 | 9.532041398 | 3.406300612 | 1.379486329 | 0.001084651 | 0.003859016 |
| Gm12207 | 7.288773403 | 3.815767398 | 2.223290042 | 0.001089415 | 0.003872631 |
| Ntng2 | 276.1655324 | 2.87912133 | 0.963187631 | 0.001099673 | 0.003905946 |
| Myom1 | 58.95770212 | 1.68136818 | 0.594661309 | 0.001115339 | 0.003956463 |
| Greb1l | 110.5681478 | 1.293193722 | 0.453616453 | 0.00112077 | 0.003974261 |
| Vcan | 1058.375989 | 2.876690311 | 0.972756657 | 0.001130398 | 0.004004702 |
| Fancd2 | 54.80600656 | 1.830425957 | 0.644078847 | 0.001133968 | 0.004016606 |
| Gm43313 | 6.554451951 | 3.642804881 | 2.1451559 | 0.001162365 | 0.004105068 |
| Fam83c | 21.92192279 | 2.119311828 | 0.775140801 | 0.001171986 | 0.004136003 |
| Ccdc146 | 27.77701718 | 1.700311016 | 0.615871444 | 0.001176643 | 0.004150147 |
| Tead2 | 505.5702706 | 1.449913856 | 0.497608436 | 0.001188807 | 0.004188434 |
| Shcbp1 | 42.82312221 | 2.0666411 | 0.754495944 | 0.001206595 | 0.004244092 |
| Gm41442 | 13.48279305 | 2.700362582 | 1.016106174 | 0.001215537 | 0.004271631 |
| Dok1 | 91.43570034 | 2.374446533 | 0.82442228 | 0.001217334 | 0.004277162 |
| Tmem52 | 25.79017284 | 3.179255524 | 1.267175323 | 0.001222969 | 0.004294602 |
| Mir7678 | 2.548734967 | 4.173578512 | 2.459992357 | 0.001228704 | 0.004313164 |
| Cfap52 | 17.88567296 | 3.421088285 | 2.034532655 | 0.001249364 | 0.004376081 |
| Pnpla3 | 16.08431259 | 1.894386803 | 0.726004531 | 0.001258799 | 0.004406714 |
| Gm26873 | 48.12395484 | 1.10053698 | 0.37742222 | 0.001259235 | 0.004407437 |
| Pygm | 8.289811685 | 3.598207595 | 2.142495793 | 0.001267699 | 0.004435445 |
| Vps37d | 141.9612825 | 1.459341127 | 0.52056105 | 0.001304276 | 0.004546841 |
| Dusp8 | 222.6902988 | 1.159139455 | 0.405409032 | 0.001304092 | 0.004546841 |
| Kcnk4 | 3.900643381 | 4.598202043 | 2.713815748 | 0.001311128 | 0.004566579 |
| Pclo | 35.60066956 | 1.344550415 | 0.475699706 | 0.00131232 | 0.004569903 |
| Gm31224 | 39.33679131 | 2.414107966 | 0.862987816 | 0.001319967 | 0.004591533 |
| Sptb | 177.5237466 | 1.051887163 | 0.357286474 | 0.001321759 | 0.004596932 |
| Cdh3 | 298.4689924 | 4.204766578 | 2.522657955 | 0.001323989 | 0.004603022 |

| | | | | | |
|---|---|---|---|---|---|
| Zfp61 | 69.43787509 | 1.799539129 | 0.644171799 | 0.001325756 | 0.00460833 |
| Gm38077 | 40.23889417 | 1.023521113 | 0.347440142 | 0.001329592 | 0.004619153 |
| Gm37893 | 51.22166638 | 1.018755206 | 0.34907233 | 0.001373102 | 0.004755675 |
| Paqr8 | 84.49051886 | 3.266313615 | 1.962653092 | 0.00138026 | 0.004778742 |
| 2010016I18Rik | 23.22370327 | 1.694827166 | 0.619882361 | 0.001382701 | 0.004786331 |
| Slc22a17 | 272.2368661 | 2.403979319 | 0.870947085 | 0.00138824 | 0.004802906 |
| 2700099C18Rik | 22.49674793 | 2.160055126 | 0.790422869 | 0.001403569 | 0.004848947 |
| Ciita | 27.92394648 | 1.453957789 | 0.521381094 | 0.00140641 | 0.004855264 |
| Cep55 | 42.01876417 | 3.080370888 | 1.24256857 | 0.001418815 | 0.004891052 |
| Mmp11 | 278.7296515 | 1.88089839 | 0.674586608 | 0.001424478 | 0.004908809 |
| Zfp57 | 176.9971555 | 3.02590955 | 1.212563032 | 0.001424226 | 0.004908809 |
| Fxyd3 | 331.782777 | 3.226110266 | 1.950147261 | 0.001429695 | 0.004925018 |
| Tmem151a | 80.14549747 | 1.918866638 | 0.695943153 | 0.001434899 | 0.004940285 |
| Msn | 1386.492042 | 1.441152523 | 0.537970197 | 0.001440552 | 0.004957079 |
| S100b | 78.17816672 | 3.254658818 | 1.96548699 | 0.00144113 | 0.004958181 |
| Gm37309 | 31.13839739 | 1.440887916 | 0.530490418 | 0.001473503 | 0.005059582 |
| Gm10130 | 51.61521536 | 1.174845095 | 0.413921903 | 0.001485346 | 0.005096597 |
| Abcc8 | 6.264116068 | 3.923184828 | 2.383825028 | 0.001491868 | 0.005114405 |
| Sult2b1 | 100.1641722 | 3.229511637 | 1.963560529 | 0.001503464 | 0.005147723 |
| Agap2 | 179.0883124 | 1.06450003 | 0.37235435 | 0.001504861 | 0.005150666 |
| Mdh1b | 5.578590861 | 3.478317381 | 1.533965233 | 0.00150967 | 0.005166206 |
| Galnt15 | 87.77506188 | 3.832053941 | 2.349281872 | 0.001513121 | 0.00517709 |
| Smpd5 | 46.5935142 | 3.026846097 | 1.230216907 | 0.001556441 | 0.005312996 |
| Tmem198 | 10.13745932 | 3.448706892 | 2.116164017 | 0.001562589 | 0.005332086 |
| Gm12454 | 13.74532642 | 3.479093891 | 2.141282905 | 0.001570751 | 0.005358983 |
| Gm37979 | 4.80373724 | 3.593070552 | 2.211749792 | 0.001573261 | 0.005366594 |
| Gm42664 | 69.00238182 | 1.576141938 | 0.589093958 | 0.00159549 | 0.00543662 |
| Cdr2 | 33.47884048 | 3.085773373 | 1.288307128 | 0.00160139 | 0.005451884 |
| Adam12 | 17.45693238 | 3.36131193 | 2.072476489 | 0.00160498 | 0.005459749 |
| AF357399 | 7.429024261 | 3.029065805 | 1.146360579 | 0.001606832 | 0.005464595 |
| Gm43289 | 5.28696424 | 2.744662016 | 1.097263675 | 0.001615837 | 0.005491328 |
| Gm39556 | 6.661985151 | 3.310080616 | 1.438220835 | 0.001625894 | 0.005522571 |
| Clvs2 | 3.291105887 | 4.21658955 | 2.592659331 | 0.001635962 | 0.005553819 |
| Mir3058 | 19.7724729 | 1.471316821 | 0.543565411 | 0.001671852 | 0.00566464 |
| Gm44200 | 7.713401625 | 3.592631596 | 2.232284623 | 0.001685037 | 0.00570327 |
| Gm17501 | 37.0080403 | 2.543293164 | 1.0986391 | 0.001690671 | 0.005720323 |
| Zswim3 | 58.35830633 | 1.02385143 | 0.357461496 | 0.001691953 | 0.005722641 |
| 2810408A11Rik | 79.58755612 | 2.963878335 | 1.216634903 | 0.001693014 | 0.005725221 |
| Capn6 | 16.05002096 | 3.232806098 | 1.405201312 | 0.001699113 | 0.005744835 |
| Plpp7 | 33.67716631 | 1.345972131 | 0.496017946 | 0.001704953 | 0.005762549 |
| Niban3 | 36.70527066 | 1.909679787 | 0.735339717 | 0.001711487 | 0.005779542 |
| Gm19265 | 13.38801451 | 3.022688447 | 1.164866098 | 0.001736893 | 0.005857694 |
| Gm38253 | 37.44126495 | 1.393856399 | 0.513924282 | 0.001768229 | 0.005955438 |

| | | | | | |
|---|---|---|---|---|---|
| Gm29438 | 20.22954277 | 2.390317773 | 0.90097401 | 0.001768913 | 0.005956696 |
| Kcnab1 | 346.1624048 | 2.046237004 | 0.76156073 | 0.001771167 | 0.005961147 |
| Traip | 26.9616821 | 1.836059226 | 0.696361738 | 0.001772601 | 0.005963882 |
| Gm17971 | 15.02317223 | 1.99865243 | 0.813620426 | 0.001793828 | 0.00603107 |
| Tmem138 | 79.62277757 | 1.051055885 | 0.370797459 | 0.001801004 | 0.006053077 |
| Mapk11 | 128.9754024 | 2.702015206 | 0.960619562 | 0.001813172 | 0.006085448 |
| Gm44167 | 4.278832306 | 3.758563314 | 1.75994623 | 0.001814009 | 0.00608719 |
| Art4 | 8.077303282 | 3.526919454 | 1.655448816 | 0.001847061 | 0.006185125 |
| Mustn1 | 9.138583595 | 2.949721587 | 1.136912206 | 0.00185898 | 0.006221779 |
| Gm42986 | 10.74769537 | 2.46163304 | 0.959512013 | 0.001865279 | 0.006239596 |
| Gm27042 | 20.15360012 | 3.001325951 | 1.272528922 | 0.00186809 | 0.006246822 |
| Ggn | 26.65313382 | 3.020458853 | 1.286589715 | 0.001872096 | 0.006259127 |
| Gm26797 | 28.2310719 | 2.308377051 | 0.862611522 | 0.001898728 | 0.00633823 |
| Tyro3 | 92.90752832 | 1.712314661 | 0.639061575 | 0.001909265 | 0.006371188 |
| Nfkbid | 57.95808475 | 2.698345945 | 0.970688643 | 0.001921648 | 0.006409168 |
| Gm19337 | 5.318327064 | 4.074453521 | 2.596631978 | 0.00193314 | 0.006443017 |
| Exo1 | 47.0714357 | 2.846277495 | 1.076654806 | 0.001945632 | 0.006477902 |
| Efna4 | 171.0829815 | 3.041243056 | 1.925855709 | 0.001969353 | 0.006550061 |
| Dscc1 | 19.53961521 | 3.483885872 | 2.224177321 | 0.001982534 | 0.006590476 |
| Gm45033 | 5.232767472 | 3.431093305 | 2.193604662 | 0.001992077 | 0.006617612 |
| Gm10791 | 25.0397129 | 2.935864481 | 1.246334942 | 0.002002672 | 0.006648207 |
| Gldn | 8.604034614 | 2.014472934 | 0.843535288 | 0.002031157 | 0.00673704 |
| Gm19409 | 6.266217993 | 3.365549204 | 1.575693416 | 0.002048818 | 0.006789651 |
| Ceacam16 | 59.87706592 | 3.071692629 | 1.964065027 | 0.002058957 | 0.006819717 |
| Gm38372 | 53.31086977 | 1.031462741 | 0.370037823 | 0.002062137 | 0.006827893 |
| Gm26203 | 3.787269938 | 3.785150689 | 2.437698557 | 0.002067041 | 0.00684039 |
| Anxa1 | 177.3922241 | 2.727367304 | 1.010620205 | 0.002068504 | 0.006843071 |
| Gm33782 | 14.5099141 | 3.386729613 | 2.185765133 | 0.002080296 | 0.006877341 |
| Gm43462 | 7.257643157 | 3.407481293 | 2.19684089 | 0.00208111 | 0.006878847 |
| Ncapg2 | 193.4186314 | 1.020852071 | 0.367152365 | 0.002096433 | 0.006919757 |
| Pmp22 | 32.38315648 | 2.275045954 | 0.860195536 | 0.002099734 | 0.006928473 |
| Cyp2s1 | 166.3646658 | 3.063039752 | 1.964651349 | 0.002106628 | 0.006948829 |
| Gm43328 | 24.87482657 | 1.266386797 | 0.472465641 | 0.002127833 | 0.007010341 |
| Gm43857 | 19.55026254 | 1.681726929 | 0.657145645 | 0.00215015 | 0.007075993 |
| Fam180a | 4.565605533 | 4.133918483 | 2.695645263 | 0.002165054 | 0.007117085 |
| Gm10605 | 4.343357819 | 3.151390829 | 1.43240056 | 0.002168374 | 0.007125559 |
| Ephb3 | 96.56682762 | 3.077551609 | 1.986061672 | 0.00216968 | 0.007127409 |
| P4htm | 58.75262553 | 3.054567072 | 1.970908354 | 0.002173305 | 0.007138095 |
| Cdc25b | 96.33583443 | 1.324273464 | 0.493690578 | 0.002178933 | 0.007155358 |
| Gm6483 | 31.26319663 | 1.285087905 | 0.478438734 | 0.002190417 | 0.007186919 |
| Tnfrsf9 | 23.39224724 | 2.7728806 | 1.063979257 | 0.002202908 | 0.007224199 |
| Casc1 | 34.67512958 | 3.076537876 | 1.992604087 | 0.002204605 | 0.00722606 |
| Gm31036 | 42.07344191 | 1.457278766 | 0.554606135 | 0.002234895 | 0.007316594 |

| | | | | | |
|---|---|---|---|---|---|
| Hmgn3 | 62.0266755 | 1.661319322 | 0.627183713 | 0.002249537 | 0.007359505 |
| Pwwp3b | 134.36951 | 2.627730786 | 0.962284985 | 0.002254171 | 0.007372152 |
| Tbx20 | 7.83135407 | 2.880557821 | 1.146481898 | 0.002274329 | 0.007433011 |
| 2610306O10Rik | 12.92297495 | 2.741592506 | 1.04492218 | 0.002277474 | 0.007440758 |
| Gm22009 | 18.46412462 | 2.43525994 | 1.154484579 | 0.002287813 | 0.007469449 |
| A730063M14Rik | 28.3071877 | 1.165413025 | 0.43416675 | 0.002299645 | 0.00750425 |
| Gm17146 | 4.039816762 | 3.744415625 | 2.458909262 | 0.002302623 | 0.007512688 |
| Dync2li1 | 358.8915966 | 1.146291775 | 0.415286744 | 0.002316379 | 0.007552435 |
| Pde3a | 4.467158648 | 3.243792478 | 1.53700232 | 0.002317516 | 0.007554859 |
| Gm31718 | 26.08707724 | 1.426305797 | 0.561056651 | 0.002323718 | 0.007571218 |
| 4930517G19Rik | 5.365800659 | 3.520268411 | 2.322168949 | 0.002339467 | 0.007618652 |
| Gm39041 | 12.58055945 | 3.220993763 | 1.542564152 | 0.002380311 | 0.00773853 |
| Bnip5 | 37.92618617 | 1.594044828 | 0.652256343 | 0.002389368 | 0.007765345 |
| 1700126G02Rik | 10.37224857 | 2.701851323 | 1.02889172 | 0.002400023 | 0.007795362 |
| Epha3 | 3.546563899 | 3.814521749 | 2.533159657 | 0.002400229 | 0.007795362 |
| Fam171a2 | 180.1094731 | 1.913440556 | 0.7352084 | 0.002401096 | 0.007796859 |
| Gm43080 | 29.19929072 | 1.179957313 | 0.440549255 | 0.002421698 | 0.007857111 |
| Fstl3 | 140.4054707 | 2.185397937 | 0.833796518 | 0.002451358 | 0.007939916 |
| Tmem252 | 74.90748319 | 3.031258127 | 2.005645788 | 0.002461541 | 0.007970209 |
| Rab7b | 22.00571704 | 3.063181064 | 2.028432504 | 0.002479598 | 0.00802191 |
| Fam81b | 3.600792726 | 3.329050835 | 1.608906881 | 0.002479332 | 0.00802191 |
| Rftn1 | 4.25952445 | 3.499793674 | 2.333758373 | 0.002489159 | 0.008051482 |
| Gm47258 | 8.562933854 | 3.161945135 | 2.099349816 | 0.002498732 | 0.008079115 |
| H2-DMa | 18.03442717 | 1.770552428 | 0.718188869 | 0.002501422 | 0.008082975 |
| Megf6 | 426.8945823 | 1.830283483 | 0.718515935 | 0.002503749 | 0.008088396 |
| AC132288.1 | 21.70584802 | 3.038385467 | 2.013531127 | 0.002516638 | 0.008122569 |
| Gm26542 | 6.722667478 | 3.333280365 | 2.226883453 | 0.002531424 | 0.008166173 |
| 1810062O18Rik | 11.53635832 | 2.281287651 | 0.901019018 | 0.002536384 | 0.008176674 |
| 4933440N22Rik | 20.83218877 | 1.560920407 | 0.633553222 | 0.002541553 | 0.008191963 |
| 5330438I03Rik | 3.174904464 | 3.476048799 | 2.328471436 | 0.002554844 | 0.008230654 |
| Cdh11 | 3.77104791 | 3.665208747 | 2.468108384 | 0.002564902 | 0.008258899 |
| Rasd2 | 15.43831834 | 3.265452119 | 2.187743598 | 0.002586013 | 0.0083185 |
| Lgals2 | 27.41397135 | 2.847255241 | 1.270368879 | 0.002592716 | 0.008338665 |
| Sgo2a | 26.88238683 | 2.7456485 | 1.09212409 | 0.002602359 | 0.008366975 |
| Pcdh12 | 16.16690976 | 1.345238117 | 0.528193335 | 0.00262546 | 0.008436905 |
| Fxyd7 | 11.92970501 | 1.907914951 | 0.771069134 | 0.002628014 | 0.008443699 |
| A430019L02Rik | 7.166400337 | 3.512674033 | 2.384580763 | 0.002629815 | 0.008446658 |
| Snord3a | 2.611901433 | 3.958123579 | 2.673950305 | 0.00264191 | 0.008484084 |
| Gm42798 | 9.679139786 | 2.746470127 | 1.099884763 | 0.002649682 | 0.008506196 |
| Gm38155 | 32.54138812 | 1.005179247 | 0.36895357 | 0.002655366 | 0.008523018 |
| Rpl24 | 12.63447043 | 1.729028686 | 0.702024618 | 0.002665738 | 0.008554879 |
| Ccdc33 | 8.690959668 | 3.430568501 | 2.320877815 | 0.002671608 | 0.00857085 |
| Cercam | 130.8035382 | 2.874501841 | 1.919895351 | 0.002685179 | 0.008612949 |

| | | | | | |
|---|---|---|---|---|---|
| Ggt7 | 100.5594811 | 2.881830836 | 1.927403607 | 0.002699266 | 0.00865524 |
| Wnt7a | 247.4432558 | 2.850182858 | 1.908249804 | 0.002719595 | 0.008713147 |
| Gm43654 | 12.26695543 | 2.122730439 | 0.884672396 | 0.002724694 | 0.008726569 |
| Zfp566 | 30.87606071 | 1.604670484 | 0.635095055 | 0.00277442 | 0.008865117 |
| Gm38248 | 27.16281525 | 1.58917582 | 0.642177444 | 0.002776919 | 0.008871626 |
| Cenpw | 33.22837866 | 1.21147708 | 0.467966717 | 0.002783115 | 0.008885504 |
| Sema3c | 808.0787374 | 2.835758222 | 1.908535034 | 0.002784177 | 0.008887415 |
| Cyp3a57 | 4.307190447 | 3.58317541 | 1.849638934 | 0.002798352 | 0.008923759 |
| Cacna1c | 17.17773824 | 2.733680886 | 1.109588144 | 0.002804852 | 0.008942998 |
| Colec11 | 18.86180813 | 1.660775349 | 0.704235996 | 0.002840675 | 0.009051204 |
| Gm16144 | 7.251655938 | 3.229751392 | 2.216061732 | 0.002866493 | 0.009127404 |
| Gm44669 | 12.14047858 | 1.739032575 | 0.734712435 | 0.002890149 | 0.009193679 |
| Gm19510 | 4.773269998 | 3.063555227 | 1.519279265 | 0.003021495 | 0.009562256 |
| Gm47694 | 17.57766441 | 1.759700678 | 0.74124864 | 0.003042553 | 0.009624136 |
| Gm13073 | 5.237082 | 3.254589116 | 2.260063296 | 0.003075009 | 0.009713992 |
| Fermt1 | 704.7645499 | 1.36107847 | 0.526995722 | 0.003103493 | 0.009800746 |
| 1700112D23Rik | 11.43709138 | 1.611565431 | 0.675853719 | 0.003106187 | 0.009807641 |
| Ttyh1 | 34.00293444 | 1.409940229 | 0.559496168 | 0.003181134 | 0.01001298 |
| Zfp382 | 23.0605672 | 2.717515691 | 1.243833885 | 0.003195673 | 0.010052148 |
| Ncs1 | 179.9732706 | 2.810471968 | 1.947270909 | 0.00320911 | 0.010092761 |
| Gm49204 | 33.81194715 | 1.312350587 | 0.519991953 | 0.003224676 | 0.010136744 |
| Gm42940 | 7.250312264 | 2.856314396 | 1.3708083 | 0.00325148 | 0.010209286 |
| Fgf9 | 12.49361046 | 3.069066423 | 2.158654432 | 0.003276922 | 0.01027572 |
| Gm21451 | 11.04033359 | 3.010061715 | 2.11096758 | 0.00328661 | 0.010299367 |
| Gm15344 | 52.92045438 | 1.487687794 | 0.632264836 | 0.003311255 | 0.010363061 |
| 6820402A03Rik | 16.0052439 | 3.024251691 | 2.125492035 | 0.003314246 | 0.010370731 |
| Ccdc180 | 7.860542593 | 3.06675288 | 2.152471726 | 0.003358538 | 0.010502477 |
| Septin4 | 69.74843512 | 1.439045765 | 0.593115717 | 0.003371301 | 0.010537238 |
| Cd28 | 4.103774605 | 3.178001146 | 2.253316598 | 0.003383818 | 0.010567754 |
| Gm45223 | 24.7620963 | 1.725022982 | 0.770315064 | 0.003389765 | 0.010582882 |
| Tnfrsf18 | 38.84963068 | 1.64481998 | 0.696554402 | 0.003395879 | 0.010600248 |
| Nav1 | 156.9112604 | 1.031081432 | 0.394278009 | 0.003421309 | 0.010669215 |
| Gm48146 | 5.572446707 | 3.09727543 | 2.207316355 | 0.003436556 | 0.0107098 |
| Xrra1 | 12.72276311 | 2.641362577 | 1.103287541 | 0.003450689 | 0.010748607 |
| Gm28988 | 5.973808944 | 3.316650265 | 2.398826421 | 0.003475815 | 0.010821603 |
| 1700109K24Rik | 33.96624116 | 1.031115226 | 0.393459016 | 0.003524688 | 0.010954217 |
| Gm25788 | 3.102578659 | 3.177490256 | 2.275659154 | 0.003533134 | 0.010975134 |
| Gm47690 | 14.34442471 | 1.971593798 | 0.831745111 | 0.003546626 | 0.011013479 |
| Gm26724 | 31.13228853 | 1.273967891 | 0.503144992 | 0.003548982 | 0.011019013 |
| Zfp69 | 27.76231811 | 1.271657905 | 0.512972293 | 0.003561328 | 0.011053767 |
| Dmtn | 47.84883153 | 2.63761492 | 1.226529684 | 0.003565492 | 0.011064903 |
| Htra1 | 11.64470605 | 2.252489971 | 0.968792359 | 0.003578814 | 0.011102654 |
| E330037G11Rik | 8.224862645 | 2.204570773 | 0.931256293 | 0.003603179 | 0.01117463 |

| | | | | | |
|---|---|---|---|---|---|
| Gm48137 | 8.980195408 | 2.6160181 | 1.098642856 | 0.003608301 | 0.011188707 |
| Gm49085 | 3.623409313 | 3.083536556 | 2.217524757 | 0.00362961 | 0.011249328 |
| 9430062P05Rik | 30.52276144 | 1.572347779 | 0.647884944 | 0.003630554 | 0.011250438 |
| Cux2 | 42.65236181 | 3.286771244 | 1.863066964 | 0.003646635 | 0.011292976 |
| Gprc5a | 291.4088848 | 2.736533453 | 1.954442245 | 0.003682161 | 0.011388903 |
| Cdhr1 | 8.286868632 | 3.246483943 | 2.380396708 | 0.003734361 | 0.011529298 |
| Fhl2 | 28.85805277 | 2.822362178 | 2.023802947 | 0.00374007 | 0.011543734 |
| Gm15512 | 3.016083546 | 3.395223527 | 2.492421995 | 0.003752594 | 0.011574417 |
| Dsel | 88.43937983 | 2.621019961 | 1.234371869 | 0.003762242 | 0.011600447 |
| Zfp52 | 62.16139758 | 1.232937409 | 0.486546997 | 0.003787846 | 0.011668146 |
| Mtss2 | 276.92488 | 1.241596668 | 0.492961937 | 0.003788675 | 0.011668828 |
| Pla2g3 | 6.076133202 | 3.104556903 | 2.262322378 | 0.003790097 | 0.011671333 |
| Pla2r1 | 111.4959879 | 2.074616269 | 0.849340846 | 0.003803954 | 0.011704614 |
| Gm19144 | 15.3883404 | 1.890735879 | 0.830826967 | 0.003818964 | 0.011741388 |
| 4930533B01Rik | 3.330729411 | 3.373406563 | 2.496428129 | 0.003822395 | 0.011748172 |
| Mfap2 | 4.987026876 | 3.056765338 | 2.232117494 | 0.003838962 | 0.011789649 |
| Snhg4 | 35.85122458 | 1.067283398 | 0.414100811 | 0.003852299 | 0.011825984 |
| Lamc3 | 489.9696401 | 2.797666517 | 2.066425126 | 0.003880484 | 0.011901929 |
| Actc1 | 7.191257163 | 2.886459638 | 1.368746974 | 0.003901418 | 0.0119604 |
| Gm49961 | 3.999913995 | 2.824569618 | 1.432789565 | 0.003927045 | 0.012033197 |
| Gm42726 | 3.677621472 | 3.481736121 | 2.629018995 | 0.003939659 | 0.012066071 |
| Gm49322 | 11.8910708 | 2.188998509 | 0.951879273 | 0.003977795 | 0.012167337 |
| A430057M04Rik | 13.99972847 | 2.629281442 | 1.255139004 | 0.003981517 | 0.012176779 |
| Prkg2 | 7.563089527 | 2.708642147 | 1.331588275 | 0.003985288 | 0.01218637 |
| Hand2os1 | 6.42301618 | 3.038691118 | 1.675054816 | 0.003988325 | 0.012193714 |
| Col4a6 | 492.2254006 | 2.618116473 | 1.893442568 | 0.003997448 | 0.012215768 |
| Arl4c | 1434.469653 | 2.187303809 | 0.981505262 | 0.004020408 | 0.012280064 |
| Nnat | 39.03537696 | 1.075823186 | 0.423330752 | 0.004043248 | 0.012343052 |
| Upk1a | 4.949891591 | 3.079021257 | 2.283694414 | 0.004056591 | 0.012374826 |
| Gm16153 | 40.57858706 | 1.064458139 | 0.4204974 | 0.004092552 | 0.012468495 |
| Gabra3 | 3.306427611 | 3.454481298 | 2.630394893 | 0.004093144 | 0.012468495 |
| Folh1 | 3.747788327 | 3.530304493 | 2.656674306 | 0.00415382 | 0.012635276 |
| Igsf23 | 26.5883022 | 1.802004265 | 0.809128223 | 0.004176652 | 0.012696679 |
| Smim38 | 10.62981048 | 2.873379201 | 2.134049339 | 0.004242142 | 0.012877404 |
| Lekr1 | 32.10689104 | 2.608384597 | 1.280316239 | 0.004244635 | 0.012882933 |
| Acta2 | 4.015993473 | 3.448144853 | 2.663619524 | 0.004247983 | 0.012891056 |
| Rnf122 | 127.1009787 | 1.808047934 | 0.755543828 | 0.00429743 | 0.013020521 |
| Olfml2b | 95.59418086 | 2.633212271 | 1.93735774 | 0.004310724 | 0.013053934 |
| Stc2 | 10.35118102 | 2.895930568 | 2.168484599 | 0.004322937 | 0.013079218 |
| Gm45342 | 5.795273633 | 2.884950369 | 2.155683495 | 0.004331776 | 0.013103895 |
| Gabrb3 | 8.176214594 | 1.987386892 | 0.960223523 | 0.004377011 | 0.013228223 |
| Cfap157 | 30.2241518 | 1.316034959 | 0.544738694 | 0.004398336 | 0.013286396 |
| Gnai1 | 39.75036631 | 2.688961537 | 1.387881217 | 0.004421125 | 0.013348932 |

| | | | | | |
|---|---|---|---|---|---|
| Zmynd12 | 12.25651761 | 1.84480821 | 0.82832002 | 0.004424062 | 0.013355699 |
| Gm36423 | 4.42713105 | 3.161443345 | 2.439886421 | 0.004426428 | 0.013360738 |
| Adgrd1 | 3.607110523 | 3.119977002 | 2.377212744 | 0.004433759 | 0.01337866 |
| Gpm6a | 168.999635 | 2.634150149 | 1.951809639 | 0.00445365 | 0.013434455 |
| Gm43379 | 24.30277837 | 1.393460374 | 0.594247209 | 0.004494402 | 0.013548865 |
| Sntg1 | 3.33253655 | 3.084588942 | 2.362940556 | 0.004500503 | 0.013562994 |
| Gm44280 | 6.745677236 | 2.661972164 | 1.367329309 | 0.004604797 | 0.01385337 |
| Plvap | 38.10282421 | 1.103659759 | 0.449666818 | 0.004612879 | 0.013873338 |
| Gm15774 | 7.440708877 | 2.629181077 | 1.33740674 | 0.004652169 | 0.013971797 |
| Gm14239 | 4.197925734 | 3.138199253 | 2.461073694 | 0.004661529 | 0.013993336 |
| Spint1 | 1028.786132 | 2.547793559 | 1.925425725 | 0.004726944 | 0.014167543 |
| Scn5a | 37.09287596 | 2.974949343 | 2.347710598 | 0.004759653 | 0.014247776 |
| Gm22748 | 13.74215579 | 2.806539143 | 1.568535948 | 0.004787392 | 0.014317412 |
| Gm48342 | 15.08042945 | 2.202001582 | 1.22523806 | 0.004797756 | 0.014343938 |
| Gm43256 | 5.569577246 | 2.655679761 | 1.378620951 | 0.004817635 | 0.014398883 |
| Gprin2 | 52.00996557 | 2.599959517 | 1.968528159 | 0.004819061 | 0.014400904 |
| Zfp827 | 327.1924531 | 2.451617929 | 1.199818243 | 0.004887459 | 0.014578059 |
| Rad51ap1 | 31.72731454 | 1.990728704 | 0.926531603 | 0.004901701 | 0.014613725 |
| Rasl12 | 24.8138623 | 2.716544429 | 2.095435419 | 0.004904668 | 0.0146203 |
| Islr | 7.516523445 | 2.299326606 | 1.114778762 | 0.00490771 | 0.014624823 |
| Gm37233 | 13.89206539 | 2.636918923 | 2.012514644 | 0.004931917 | 0.014690116 |
| Gm38319 | 31.95420162 | 1.042144434 | 0.42330653 | 0.005010205 | 0.014897869 |
| Nyap1 | 27.84093236 | 1.523497191 | 0.660706773 | 0.005024483 | 0.01493801 |
| Gm7525 | 3.792390722 | 2.910360231 | 2.271440279 | 0.005025701 | 0.014939316 |
| Ypel1 | 115.1976185 | 1.084340449 | 0.43409775 | 0.005067416 | 0.015047002 |
| Lca5l | 25.93240982 | 2.600374678 | 1.990459146 | 0.005069001 | 0.015048778 |
| Slco3a1 | 2517.869175 | 1.458400718 | 0.624747717 | 0.005077111 | 0.015064136 |
| Mmp23 | 96.6510357 | 2.015129234 | 0.886414391 | 0.005098857 | 0.015120767 |
| Gm42658 | 16.93035635 | 1.934524958 | 0.898121061 | 0.005128711 | 0.015191403 |
| Cd248 | 3.136082681 | 3.220975011 | 2.629525665 | 0.005155875 | 0.015260176 |
| Adamts12 | 8.718033689 | 2.621588805 | 1.287852497 | 0.005252303 | 0.015509609 |
| 2010310C07Rik | 3.732800427 | 2.848629756 | 1.621579378 | 0.005277725 | 0.015570302 |
| Gm44171 | 4.056537807 | 3.055206571 | 2.48533655 | 0.005321693 | 0.015680731 |
| Gm48062 | 26.56335601 | 1.473564479 | 0.636459402 | 0.005321504 | 0.015680731 |
| Fxyd6 | 66.25037056 | 2.446893993 | 1.225852364 | 0.005325739 | 0.015690245 |
| Rps10 | 38.58873571 | 1.043458697 | 0.427191558 | 0.005348786 | 0.015743644 |
| Wdr66 | 7.95880314 | 2.729303842 | 2.16185516 | 0.005411177 | 0.015912643 |
| Ppp1r42 | 10.94565539 | 1.652101548 | 0.77422424 | 0.005413961 | 0.01591595 |
| Gm50462 | 19.34308168 | 1.891630992 | 0.875252141 | 0.005426124 | 0.015946822 |
| Akr1b8 | 262.9965729 | 1.665367357 | 0.739315386 | 0.005440962 | 0.015985534 |
| Gm49838 | 4.429623233 | 2.929220985 | 1.860436379 | 0.005450176 | 0.016010153 |
| Gm15718 | 8.324807322 | 1.953166598 | 0.919051553 | 0.005464639 | 0.016047726 |
| Gm31763 | 12.17222518 | 1.609282285 | 0.731960955 | 0.005472097 | 0.016067168 |

| | | | | | |
|---|---|---|---|---|---|
| Itgbl1 | 2.580473122 | 3.19025557 | 2.66350919 | 0.005499009 | 0.016141248 |
| Stx11 | 183.0128873 | 1.478569083 | 0.631193858 | 0.005529866 | 0.016221898 |
| Palb2 | 55.5130709 | 1.041723795 | 0.427677486 | 0.005588234 | 0.016378099 |
| Ccdc148 | 135.7895271 | 1.0599838 | 0.43031839 | 0.00563335 | 0.016497728 |
| Fbxl12os | 21.97974478 | 1.657570088 | 0.735594123 | 0.005655797 | 0.016553362 |
| Gm42548 | 4.894239027 | 2.799843723 | 2.275328234 | 0.005660854 | 0.016565636 |
| Gm25835 | 16.48387159 | 1.567672777 | 0.736699417 | 0.005672267 | 0.016591447 |
| Gm11839 | 2.912797392 | 2.883923139 | 2.357902134 | 0.005744129 | 0.016781182 |
| Mgst2 | 29.24544916 | 2.603738224 | 2.095332094 | 0.005801892 | 0.016937041 |
| Lrrc27 | 29.65200797 | 2.310607967 | 1.009140099 | 0.005841643 | 0.017032919 |
| Egr2 | 373.9044641 | 2.401209719 | 1.975687392 | 0.005862482 | 0.017077549 |
| Gm17092 | 24.59156564 | 1.445104298 | 0.640308611 | 0.005870278 | 0.017095069 |
| Sfmbt2 | 22.23386178 | 2.494355794 | 1.337620314 | 0.005876779 | 0.017111405 |
| 4930533K18Rik | 14.03039637 | 2.564267852 | 2.060585489 | 0.005895106 | 0.017154356 |
| Rap1gap2 | 25.59721036 | 1.956813256 | 0.870320017 | 0.005919692 | 0.017215459 |
| Srf | 830.0578623 | 1.085961661 | 0.460794002 | 0.0059299 | 0.01723992 |
| Lrrc9 | 210.0672633 | 2.397521182 | 1.898791358 | 0.005940216 | 0.017267297 |
| Fosl1 | 7.67853594 | 2.721632913 | 2.237543834 | 0.005965978 | 0.017336929 |
| Trdc | 3.659248734 | 2.981106764 | 2.539360409 | 0.005972397 | 0.01734714 |
| Zfp850 | 47.88739691 | 1.142609023 | 0.478642141 | 0.005984367 | 0.017377211 |
| Nat14 | 63.57177795 | 2.297690876 | 1.017461115 | 0.006017686 | 0.017460751 |
| 9930120I10Rik | 10.12155039 | 1.733788004 | 0.812076696 | 0.006101007 | 0.017681126 |
| Ovgp1 | 15.60259825 | 1.737183752 | 0.81876182 | 0.00616195 | 0.017833506 |
| Slc27a3 | 190.5969161 | 1.084557855 | 0.45640247 | 0.00620041 | 0.017936701 |
| Gm23608 | 2.75310151 | 2.772213296 | 2.304540018 | 0.006203956 | 0.017944252 |
| Tlr1 | 8.113039658 | 2.629920857 | 2.170060967 | 0.006227277 | 0.018006278 |
| Gm7665 | 10.29634377 | 2.618711045 | 2.159220017 | 0.006230303 | 0.018012315 |
| Heph | 61.35622795 | 2.418252688 | 1.943776989 | 0.006255569 | 0.018079914 |
| Gm22107 | 10.72992931 | 2.135768206 | 1.064897359 | 0.006323447 | 0.018240385 |
| Gm47430 | 12.59300245 | 2.340997916 | 1.0875417 | 0.006472466 | 0.018639421 |
| Ska1 | 10.33649996 | 2.611716315 | 2.17487603 | 0.006486149 | 0.018673221 |
| Slco5a1 | 52.26001801 | 2.390829447 | 1.94266983 | 0.00649724 | 0.018693933 |
| Gm43389 | 4.852571574 | 2.691338338 | 2.28532815 | 0.006503489 | 0.018706302 |
| Gm37860 | 6.381536231 | 2.418845334 | 1.179210557 | 0.00655457 | 0.018841933 |
| Aoc1 | 43.49828331 | 2.35505531 | 1.236694708 | 0.006571904 | 0.018877626 |
| 4930483K19Rik | 10.08886839 | 1.650358484 | 0.825772976 | 0.006571265 | 0.018877626 |
| Gm30881 | 8.991270544 | 2.44220307 | 1.338327723 | 0.006587063 | 0.018915507 |
| Txndc2 | 3.02333417 | 2.913958467 | 2.553038695 | 0.00662229 | 0.018999837 |
| Fgl2 | 11.74524803 | 1.655937402 | 0.806120286 | 0.006629441 | 0.019017281 |
| Abhd16b | 9.931723088 | 1.994841975 | 1.019470745 | 0.006642347 | 0.019040927 |
| Orai2 | 152.4521594 | 2.202317137 | 0.956597691 | 0.00665473 | 0.019067025 |
| Pcdhga12 | 22.9081637 | 1.910043943 | 0.863881089 | 0.006662969 | 0.01908778 |
| 4933427E11Rik | 3.488329757 | 2.845233903 | 2.511771317 | 0.006669145 | 0.019096921 |

| | | | | | |
|---|---|---|---|---|---|
| Gm19345 | 24.77349442 | 2.39340649 | 1.96210649 | 0.006709005 | 0.019199598 |
| Gm49396 | 30.74948687 | 1.094109005 | 0.465357865 | 0.006729287 | 0.01924903 |
| Gm26628 | 18.88031369 | 1.271443937 | 0.561594515 | 0.006748681 | 0.019290131 |
| Gm5526 | 17.67386057 | 1.233524383 | 0.552803686 | 0.006757732 | 0.019313126 |
| Orm3 | 60.33650209 | 1.054602592 | 0.451029889 | 0.006766309 | 0.019334758 |
| Asb5 | 7.405377834 | 2.019686928 | 0.974012105 | 0.006769325 | 0.019340496 |
| Inhba | 1380.10769 | 1.094137799 | 0.476942326 | 0.006789403 | 0.019394974 |
| Slc10a6 | 3.110259815 | 3.616464827 | 3.553227715 | 0.006801996 | 0.019425165 |
| Gm42672 | 8.582683681 | 2.037486682 | 0.995531965 | 0.006821262 | 0.019474388 |
| Gm6583 | 3.719172248 | 2.701257177 | 2.350976889 | 0.00687218 | 0.019605173 |
| Gm12870 | 3.158036353 | 2.745192783 | 2.388648877 | 0.006883471 | 0.019631548 |
| Gm37660 | 4.679421993 | 2.801156074 | 2.529172286 | 0.006926283 | 0.019730191 |
| Rasip1 | 24.05769303 | 1.954618674 | 0.927744517 | 0.006970881 | 0.019842505 |
| Mettl24 | 8.23009628 | 2.660194558 | 2.349966856 | 0.006979945 | 0.019862415 |
| Krt7 | 1733.540362 | 2.344290749 | 2.006134001 | 0.006984693 | 0.019872531 |
| Gm20492 | 8.699766359 | 1.895677357 | 0.952669163 | 0.007040735 | 0.020011663 |
| 4933401D09Rik | 6.418892664 | 2.636469436 | 2.308384129 | 0.007079959 | 0.020105281 |
| Gm18935 | 7.325786067 | 2.55768158 | 2.204018482 | 0.007082534 | 0.020109618 |
| Gm49783 | 16.4405106 | 1.400383428 | 0.647145378 | 0.007172856 | 0.020335984 |
| Gm45718 | 22.41987685 | 1.193770011 | 0.529642396 | 0.007199752 | 0.020406209 |
| Haspin | 19.60890872 | 1.769716476 | 0.892196337 | 0.007229254 | 0.020483775 |
| Mypopos | 23.12913482 | 1.231704555 | 0.539779568 | 0.007231362 | 0.020486725 |
| Gm17816 | 6.664100445 | 2.555959193 | 2.267177844 | 0.007387291 | 0.020882237 |
| 4933413C19Rik | 11.13654784 | 2.439297326 | 2.101343119 | 0.007454637 | 0.0210571 |
| Mxra8 | 46.19668268 | 1.251718631 | 0.578817071 | 0.007514746 | 0.021208159 |
| Pcdhgb8 | 6.684915383 | 2.537687211 | 2.259302739 | 0.00757213 | 0.021351271 |
| 9130230N09Rik | 13.99822747 | 1.228871397 | 0.556302966 | 0.007575289 | 0.02135704 |
| Tmem17 | 29.41035105 | 1.150749253 | 0.509629801 | 0.00758146 | 0.021371297 |
| Gins3 | 39.21095441 | 1.031971412 | 0.44114145 | 0.007600745 | 0.02141308 |
| Kbtbd11 | 11.75345388 | 2.348532957 | 1.332376272 | 0.007617528 | 0.021454062 |
| Gm4202 | 16.47243123 | 1.534973003 | 0.730151546 | 0.007699006 | 0.021664461 |
| Sema3d | 2.952288737 | 2.713813223 | 2.512232249 | 0.007701372 | 0.021667942 |
| Armcx4 | 66.7184097 | 1.53870181 | 0.708987147 | 0.007742129 | 0.021763467 |
| Gm30363 | 2.261207557 | 2.765145616 | 2.601943496 | 0.00778498 | 0.021874313 |
| Slc2a4 | 11.57222567 | 1.55242221 | 0.759538942 | 0.007817525 | 0.021946478 |
| Arhgef4 | 15.84248574 | 2.344889319 | 2.044812126 | 0.007969155 | 0.022336215 |
| Dio3 | 27.50947446 | 1.898143196 | 0.920633016 | 0.008010034 | 0.022437681 |
| Gm10676 | 7.515737432 | 2.332984899 | 1.352894387 | 0.008022166 | 0.022465108 |
| Gm38380 | 3.289518435 | 3.015063812 | 3.566171613 | 0.008031478 | 0.022484572 |
| Osbpl5 | 222.0661148 | 1.634217576 | 0.763181619 | 0.008188461 | 0.022854094 |
| Arhgap27os1 | 5.576378585 | 2.665043581 | 2.636842721 | 0.008242834 | 0.02299157 |
| Meg3 | 151.6559593 | 1.532446103 | 0.812356011 | 0.008252733 | 0.023010053 |
| Ecscr | 215.56994 | 1.410977494 | 0.646512332 | 0.008251881 | 0.023010053 |

| | | | | | |
|---|---|---|---|---|---|
| Csrnp1 | 687.2792817 | 1.123102948 | 0.513299124 | 0.008339439 | 0.023211338 |
| Cenpp | 12.92350135 | 1.408777656 | 0.659119291 | 0.008350273 | 0.023228017 |
| Itgb8 | 868.3872494 | 1.583192024 | 0.738133514 | 0.008365012 | 0.023258901 |
| Uckl1os | 20.15096387 | 1.317176962 | 0.612858304 | 0.008453001 | 0.023476344 |
| Foxf1 | 6.762725972 | 2.50228593 | 2.357941871 | 0.008458476 | 0.02348336 |
| 5830432E09Rik | 5.129791833 | 2.644624532 | 2.652357418 | 0.008462122 | 0.023488079 |
| Ggta1 | 148.3715572 | 1.625808453 | 0.76896027 | 0.008478152 | 0.023522366 |
| Fxyd5 | 70.55527254 | 1.129577018 | 0.505000093 | 0.008480765 | 0.023526216 |
| Zfp991 | 28.85866783 | 1.74572593 | 0.876381358 | 0.008537634 | 0.023650478 |
| Ms4a4d | 113.1788769 | 1.636913463 | 0.775298815 | 0.008591323 | 0.023771061 |
| Gm44008 | 5.844467025 | 2.375192367 | 2.161626405 | 0.008602801 | 0.023789099 |
| Dnajc27 | 62.92121303 | 1.057391117 | 0.463803032 | 0.008611981 | 0.023811055 |
| Gm28809 | 7.204385908 | 2.494590294 | 2.432251987 | 0.008697328 | 0.024026491 |
| Msantd1 | 15.23182293 | 1.526337869 | 0.722443319 | 0.008709981 | 0.024049179 |
| Gm4131 | 6.573778702 | 1.790794012 | 0.920687063 | 0.008735155 | 0.024109934 |
| 4933413J09Rik | 6.189738237 | 2.368581688 | 2.170539486 | 0.008746866 | 0.024138785 |
| Dusp18 | 167.9395725 | 2.131183483 | 1.867012224 | 0.008767316 | 0.024191743 |
| Gm49971 | 8.931135157 | 2.18385392 | 1.064811338 | 0.008793584 | 0.024250275 |
| Cspg4 | 179.356123 | 2.128860462 | 1.869385878 | 0.008836678 | 0.024344631 |
| Lhx2 | 2.875511283 | 2.683284681 | 2.82480236 | 0.008853722 | 0.024384583 |
| Gm7575 | 14.76773412 | 2.269899315 | 2.075793465 | 0.008943583 | 0.024601426 |
| 1520401A03Rik | 3.16923942 | 2.578396734 | 2.605270372 | 0.008956095 | 0.024627656 |
| Gm27241 | 3.773819224 | 2.542600436 | 2.535917289 | 0.008988509 | 0.024706171 |
| Pif1 | 16.98965703 | 2.238154993 | 1.306593602 | 0.009026495 | 0.024791617 |
| Tcp10b | 5.197854709 | 2.320958841 | 1.260022878 | 0.009093659 | 0.024966584 |
| Rnasel | 44.06982505 | 1.094887186 | 0.489404381 | 0.009098386 | 0.024973951 |
| Cldn6 | 552.9833547 | 2.054571696 | 0.945064362 | 0.009098946 | 0.024973951 |
| Gpr55 | 7.246611866 | 1.937668593 | 1.009396673 | 0.009120876 | 0.02501983 |
| Hrob | 14.98934867 | 2.229211693 | 1.159086641 | 0.009136924 | 0.02505459 |
| Gm10399 | 9.633103791 | 2.253823003 | 2.052672021 | 0.009179084 | 0.025154333 |
| Gm8234 | 8.025496987 | 2.268793323 | 2.095604848 | 0.009230018 | 0.025272261 |
| Nes | 206.388938 | 2.087777944 | 0.997467669 | 0.009303488 | 0.025433511 |
| Zfr2 | 73.37308608 | 1.037391682 | 0.455891289 | 0.009338922 | 0.025523105 |
| Gm45104 | 17.19345097 | 2.275904911 | 1.251440107 | 0.009396868 | 0.025666852 |
| Adcy3 | 107.2905637 | 1.020327641 | 0.452834504 | 0.009414109 | 0.025706625 |
| Gm15247 | 6.029789644 | 2.378618098 | 2.323432579 | 0.009432191 | 0.025752337 |
| Gpc6 | 20.7949628 | 1.844490102 | 0.929454601 | 0.00946665 | 0.025835393 |
| Cpz | 2.85900685 | 2.724859599 | 4.285009086 | 0.009485455 | 0.025879353 |
| Osbpl6 | 41.5480278 | 1.592056977 | 0.758962373 | 0.009531911 | 0.025987628 |
| Gm43375 | 7.996667041 | 2.29244476 | 1.464047105 | 0.009531807 | 0.025987628 |
| Gm50383 | 7.469388308 | 1.871113435 | 1.056522288 | 0.0095721 | 0.026082374 |
| Tssk4 | 7.553674766 | 2.354566644 | 2.347163209 | 0.009672721 | 0.026311721 |
| Gm15903 | 5.14904422 | 2.257507131 | 1.414732689 | 0.009684439 | 0.026336131 |

| | | | | | |
|---|---|---|---|---|---|
| Cfap100 | 11.67333179 | 1.489373224 | 0.778236858 | 0.00971882 | 0.026418396 |
| Gm48624 | 9.394557583 | 2.231810854 | 1.385652335 | 0.009802375 | 0.026615362 |
| Slc16a4 | 27.37380509 | 1.110580776 | 0.510447035 | 0.009866884 | 0.026752667 |
| BC034090 | 243.2802067 | 2.105413367 | 1.212024642 | 0.009873617 | 0.02676714 |
| Spock2 | 7.957970366 | 2.21903341 | 2.091828962 | 0.00994812 | 0.02695008 |
| Aunip | 8.520587304 | 2.235464184 | 1.40300729 | 0.00995689 | 0.026970032 |
| Csdc2 | 6.220364987 | 2.21591915 | 1.192893601 | 0.009959983 | 0.026970796 |
| Lrat | 16.74285981 | 1.400501385 | 0.724666328 | 0.009998815 | 0.027064493 |
| Plekhh2 | 204.7882611 | 2.059525951 | 1.906849832 | 0.010007377 | 0.02708385 |
| Xylt1 | 67.43737179 | 2.081477078 | 1.912077007 | 0.010026916 | 0.027129078 |
| Plac9a | 13.8860135 | 1.826651255 | 0.930297143 | 0.010130876 | 0.027387187 |
| AC109619.1 | 5.658209907 | 2.326676001 | 2.368793071 | 0.010153135 | 0.027435766 |
| Lime1 | 4.516010185 | 2.288491866 | 1.510223461 | 0.010167098 | 0.02746963 |
| Gm15411 | 16.52105336 | 1.342250895 | 0.664765745 | 0.0101986 | 0.02754311 |
| Gfpt2 | 35.4402724 | 2.088149548 | 1.932906996 | 0.010214724 | 0.027578893 |
| Gm49733 | 2.33498306 | 2.594734889 | 3.174375739 | 0.010224342 | 0.027597094 |
| Gm4430 | 24.68542368 | 1.202633329 | 0.584512389 | 0.010283974 | 0.027730749 |
| Emp3 | 21.37145743 | 1.475090134 | 0.716974566 | 0.010294039 | 0.027753989 |
| Gm16892 | 8.836143211 | 2.100794375 | 1.067997716 | 0.01047482 | 0.028196791 |
| Gm48027 | 9.020376118 | 2.133679736 | 1.118069345 | 0.010528451 | 0.028297015 |
| Slc25a43 | 14.03718716 | 2.09088823 | 1.947124927 | 0.010529362 | 0.028297015 |
| Gm16061 | 5.770378399 | 1.767852494 | 1.021260508 | 0.010549049 | 0.028338018 |
| Nfe2l3 | 152.4786213 | 1.902675551 | 1.080485881 | 0.010599034 | 0.0284484 |
| Gm50012 | 12.87070889 | 1.56770967 | 0.80886659 | 0.010668432 | 0.028622659 |
| B3galt2 | 4.549389957 | 2.260578626 | 1.575174988 | 0.01067596 | 0.028638851 |
| Prr15 | 34.80369642 | 2.087732572 | 1.081370717 | 0.01070163 | 0.028695681 |
| Fbln2 | 14.82273622 | 2.123318687 | 1.115083877 | 0.010749075 | 0.028814847 |
| Tesmin | 17.81329935 | 1.090335005 | 0.512112541 | 0.010753149 | 0.028821744 |
| Gm37314 | 7.667613238 | 2.145793457 | 1.315750516 | 0.010785119 | 0.028903395 |
| Frem1 | 3.40353421 | 2.40428144 | 2.988718006 | 0.010918055 | 0.029206634 |
| Dclk2 | 34.67781075 | 2.052957209 | 1.94630782 | 0.010921928 | 0.029208851 |
| Amd-ps3 | 5.786207934 | 2.155309668 | 2.125575278 | 0.011015597 | 0.029426549 |
| Gm50322 | 5.209072474 | 2.213254902 | 2.257939478 | 0.011060782 | 0.029530812 |
| Gm37606 | 4.604368729 | 2.19776313 | 1.489402673 | 0.011231988 | 0.02995457 |
| B3gat2 | 25.39900087 | 1.01214861 | 0.463312911 | 0.011266118 | 0.03003307 |
| Gm28085 | 11.23016953 | 2.07004027 | 1.082866468 | 0.011292927 | 0.030091997 |
| Angpt1 | 193.3652626 | 1.528709211 | 0.753441171 | 0.011321998 | 0.030156899 |
| Ikzf2 | 111.5199932 | 1.976546558 | 0.955301537 | 0.011363961 | 0.03026027 |
| Lrrc4 | 9.307451823 | 2.116760888 | 2.143255387 | 0.011380044 | 0.030290489 |
| B130046B21Rik | 12.63929454 | 1.631798365 | 0.855530291 | 0.011377339 | 0.030290489 |
| Ms4a6b | 8.732021075 | 2.119147188 | 1.332237346 | 0.011410176 | 0.030362268 |
| Gprc5b | 601.8103835 | 1.699338239 | 0.859647544 | 0.011426308 | 0.030396766 |
| Mirg | 6.077197318 | 1.714370919 | 0.988588298 | 0.011506885 | 0.030587955 |

| | | | | | |
|---|---|---|---|---|---|
| Nhs | 19.30302858 | 2.027792217 | 1.976826778 | 0.011513561 | 0.030594952 |
| Serpini1 | 5.246819522 | 2.118541667 | 2.14209677 | 0.011536469 | 0.030651581 |
| Adora2b | 4.817169122 | 2.186310523 | 2.344849547 | 0.01155387 | 0.030685071 |
| D230049E03Rik | 6.568291606 | 2.172661003 | 1.496662642 | 0.011643337 | 0.030903391 |
| Gm26109 | 5.626024752 | 2.175329479 | 1.533370447 | 0.011738888 | 0.031133361 |
| Ednra | 6.150389198 | 2.203429916 | 1.363070267 | 0.011915881 | 0.031554806 |
| Gm17055 | 9.685906737 | 1.361969598 | 0.690192218 | 0.011948476 | 0.031632391 |
| Pdk3 | 127.8046482 | 1.99559426 | 1.198503833 | 0.011961233 | 0.031661798 |
| Pp2d1 | 3.642966945 | 2.204640844 | 1.626582709 | 0.011983848 | 0.031708538 |
| Gm6525 | 3.522392175 | 2.202739186 | 2.451969916 | 0.012017528 | 0.031775747 |
| Chst11 | 22.62992807 | 2.062343693 | 1.289595136 | 0.012022026 | 0.031783263 |
| Oscp1 | 107.134457 | 2.009254497 | 1.209483202 | 0.0120393 | 0.031824547 |
| Gm43196 | 7.514845341 | 2.075915273 | 2.168696856 | 0.012088856 | 0.03192476 |
| Oip5 | 30.34520914 | 1.080690728 | 0.517979189 | 0.012139198 | 0.032053296 |
| Pdgfrl | 17.03264509 | 1.995740912 | 1.973412802 | 0.012156822 | 0.032091 |
| Gm16685 | 19.05921856 | 2.099454992 | 2.360650515 | 0.012196874 | 0.032179022 |
| Klf10 | 1032.178549 | 1.06379289 | 0.515208029 | 0.012226206 | 0.032247541 |
| LXEJ02004842.1 | 6.846537566 | 2.12670562 | 1.486694949 | 0.012246444 | 0.032278741 |
| Dctd | 72.43420448 | 1.956657097 | 0.974367494 | 0.012245542 | 0.032278741 |
| Ttc28 | 195.9291395 | 1.236091039 | 0.593658142 | 0.012251158 | 0.032286731 |
| Cyp2a4 | 1597.00709 | 1.563997261 | 0.961775814 | 0.012277285 | 0.032333202 |
| Tnfsf13 | 14.00877902 | 1.550401407 | 0.793671011 | 0.012338345 | 0.032467458 |
| Gm10941 | 8.461388809 | 1.665546967 | 0.925688336 | 0.012485223 | 0.032808971 |
| Angptl7 | 3.703649572 | 2.113259498 | 1.448578865 | 0.012506538 | 0.032854204 |
| Ppic | 89.98069139 | 1.333122139 | 0.679420879 | 0.012517407 | 0.032875537 |
| Sstr3 | 8.594621682 | 1.705323995 | 0.984520102 | 0.012537551 | 0.032919434 |
| 4930458D05Rik | 16.45733746 | 1.219779565 | 0.622728044 | 0.012558813 | 0.032961731 |
| Pacrg | 44.73442885 | 1.66543905 | 0.838229519 | 0.012573463 | 0.032991158 |
| Gm47644 | 5.114632496 | 1.686231421 | 0.995935737 | 0.012576269 | 0.032993649 |
| Trib2 | 31.55208516 | 1.44470454 | 0.743231443 | 0.012577851 | 0.032993649 |
| Gm43920 | 2.674949527 | 2.177245237 | 3.416797226 | 0.012593151 | 0.033024755 |
| Gm34408 | 2.062744495 | 2.185380999 | 2.507779283 | 0.012630523 | 0.033100147 |
| Mir3100 | 2.261605798 | 1.822323053 | 5.333204845 | 0.012636946 | 0.033112457 |
| Gm13199 | 11.53778192 | 1.469618856 | 0.76427744 | 0.012679219 | 0.033218688 |
| Ehf | 2155.407858 | 2.071610557 | 1.285472075 | 0.01272191 | 0.033321439 |
| Gm43912 | 12.87209034 | 1.450462229 | 0.773109708 | 0.012853522 | 0.033620273 |
| Gm9316 | 6.173257469 | 2.078919315 | 1.413007015 | 0.012869798 | 0.03365367 |
| Gm13590 | 7.558547287 | 1.978782434 | 2.016558686 | 0.012916065 | 0.033760855 |
| Cr2 | 6.630804246 | 1.477331493 | 0.799581639 | 0.012958145 | 0.03385016 |
| Cdh9 | 8.48032186 | 1.97953677 | 2.017175154 | 0.013014461 | 0.033981023 |
| Arhgap27os3 | 7.581260791 | 2.095487334 | 1.490498839 | 0.013049054 | 0.034052815 |
| Gm12043 | 3.321890875 | 2.114949078 | 1.54068813 | 0.013106941 | 0.034185281 |
| Gm22728 | 5.987232346 | 2.015660593 | 2.186319883 | 0.013182371 | 0.034340013 |

| | | | | | |
|---|---|---|---|---|---|
| 1700001K19Rik | 6.608053545 | 1.967206455 | 2.016952475 | 0.013194109 | 0.034361264 |
| Mybl2 | 47.36640821 | 1.025146516 | 0.478312636 | 0.013235769 | 0.034451058 |
| Krt17 | 81.13175066 | 1.889216684 | 1.942816545 | 0.013288341 | 0.034573829 |
| Gm10277 | 16.77985667 | 1.228434959 | 0.630483214 | 0.013316777 | 0.034633726 |
| Ranbp3l | 15.24178826 | 1.202258367 | 0.617719774 | 0.013319536 | 0.034636207 |
| Nkain2 | 28.28272803 | 1.910133454 | 1.959505569 | 0.013382432 | 0.034776203 |
| Stard6 | 21.42168109 | 1.678628929 | 0.887357884 | 0.013432991 | 0.034893411 |
| Gm49493 | 3.811999055 | 2.021332801 | 2.17734286 | 0.013449386 | 0.034931272 |
| Sult1e1 | 13.16924194 | 1.775503573 | 1.329845161 | 0.013513178 | 0.035068478 |
| D430013B06Rik | 15.07799609 | 1.259837283 | 0.656177283 | 0.013610284 | 0.035291847 |
| Psmc3ip | 39.81813451 | 1.179043872 | 0.585139398 | 0.013742399 | 0.035586343 |
| Gm45495 | 22.55597297 | 1.021985487 | 0.490175519 | 0.013746941 | 0.035593301 |
| Plcd1 | 80.30223565 | 1.815185077 | 1.813252912 | 0.013814946 | 0.035754906 |
| Plekho1 | 48.68357418 | 1.030424813 | 0.489907703 | 0.013840188 | 0.035815404 |
| AI854703 | 13.46673192 | 1.900274404 | 1.997253673 | 0.013908994 | 0.035959519 |
| Sgpp2 | 249.1210363 | 1.801777605 | 1.811443847 | 0.01410717 | 0.036427705 |
| Gm128 | 2.302288517 | 2.027088926 | 2.374655139 | 0.014199512 | 0.036626724 |
| Calr4 | 82.53854361 | 1.8017779 | 1.820447167 | 0.014261536 | 0.036766944 |
| Prkar2b | 63.04920691 | 1.933275842 | 1.269127172 | 0.014411514 | 0.037103753 |
| 4833422C13Rik | 30.00353163 | 1.034008201 | 0.505836127 | 0.014487764 | 0.037255082 |
| Dtna | 100.0986315 | 1.778169074 | 1.809995696 | 0.014528382 | 0.037349523 |
| Tril | 71.6891821 | 1.778009855 | 1.815827702 | 0.014622757 | 0.037556924 |
| Gm19552 | 2.543286262 | 1.979168578 | 2.785918681 | 0.014775922 | 0.037909721 |
| Amn | 8.829127038 | 1.613306458 | 1.02988577 | 0.014819641 | 0.038004069 |
| Gm14137 | 10.90735675 | 1.665313628 | 0.907790126 | 0.014826985 | 0.03801532 |
| Fbln5 | 7.237044337 | 1.612545948 | 0.938451272 | 0.014866287 | 0.038105906 |
| Pkdrej | 13.84946535 | 1.350800161 | 0.765668856 | 0.014920996 | 0.038235924 |
| Gjc2 | 3.659464191 | 2.032741907 | 1.695380475 | 0.014997815 | 0.038422515 |
| Gm49730 | 4.088195752 | 1.88733596 | 2.044109436 | 0.014997777 | 0.038422515 |
| Gm42791 | 3.775862461 | 1.932914831 | 2.395401895 | 0.015047033 | 0.038522895 |
| Best1 | 13.94106333 | 1.912823988 | 1.275987021 | 0.01507517 | 0.038574346 |
| Dipk1b | 47.91101185 | 1.767672668 | 1.834538345 | 0.01511665 | 0.038665019 |
| Gm43445 | 6.677196898 | 1.863281904 | 2.01971939 | 0.015120313 | 0.038669233 |
| Tmem171 | 89.03939086 | 1.953383001 | 1.626834771 | 0.015181532 | 0.038820249 |
| Gm5406 | 287.4628938 | 1.585579259 | 0.831565127 | 0.015208283 | 0.038852791 |
| Mir7664 | 3.324430451 | 1.912284957 | 2.761165762 | 0.015215225 | 0.038860179 |
| Scg5 | 6.744962052 | 1.855813001 | 2.032990417 | 0.015238169 | 0.038903247 |
| Tmem231 | 201.3557733 | 1.858124216 | 1.197089569 | 0.01531784 | 0.039075456 |
| E130317F20Rik | 29.44808241 | 1.051836524 | 0.5267478 | 0.01541244 | 0.039285447 |
| Akt3 | 106.3921158 | 1.356457342 | 0.713214678 | 0.015475854 | 0.039402176 |
| Paqr6 | 26.03897718 | 1.765484168 | 1.853431253 | 0.015486087 | 0.039420811 |
| Ggt5 | 2.927429784 | 1.871892939 | 2.886705309 | 0.015518336 | 0.039492426 |
| 2610028D06Rik | 13.84294097 | 1.700676602 | 0.999213909 | 0.015626016 | 0.039729581 |

| | | | | | |
|---|---|---|---|---|---|
| Agpat4 | 102.1801253 | 1.728700806 | 1.812159623 | 0.015633653 | 0.039743732 |
| Rps2-ps9 | 4.394210282 | 1.86931382 | 2.223348449 | 0.015678355 | 0.039846815 |
| 9230116L04Rik | 3.564384409 | 1.87342925 | 2.583291384 | 0.015695624 | 0.03987746 |
| 4833408A19Rik | 8.264159065 | 1.672345634 | 0.952766558 | 0.015696646 | 0.03987746 |
| 4930546K05Rik | 2.509753153 | 1.759761263 | 3.350774643 | 0.015804111 | 0.040099279 |
| Gm3235 | 2.907558384 | 1.41758886 | 2.937905316 | 0.015906735 | 0.040331106 |
| Col14a1 | 122.6544386 | 1.263225863 | 0.716704865 | 0.015916588 | 0.040340101 |
| Dram1 | 189.4635097 | 1.444353165 | 0.768409376 | 0.015921487 | 0.040347187 |
| Insyn2a | 7.716941292 | 1.921317265 | 1.362443197 | 0.016107138 | 0.040774749 |
| Igsf3 | 277.1340391 | 1.407765972 | 0.738849838 | 0.016171111 | 0.040914937 |
| Gm43745 | 3.510881433 | 1.821112318 | 2.707724017 | 0.016208154 | 0.040981645 |
| Pkp1 | 24.83659901 | 1.668995047 | 0.953212464 | 0.016233583 | 0.041035128 |
| Mdk | 426.8659394 | 1.823691493 | 0.985743893 | 0.016238347 | 0.041041764 |
| Gm13003 | 8.82700094 | 1.807247458 | 2.042813855 | 0.016248183 | 0.041061218 |
| Gm38411 | 3.525215377 | 1.834926503 | 2.402485146 | 0.016351944 | 0.041296245 |
| Gm42462 | 21.58282573 | 1.878294071 | 1.328211663 | 0.016359327 | 0.041309453 |
| B230377A18Rik | 3.884645739 | 1.820160536 | 2.220450546 | 0.016593563 | 0.041801924 |
| 4921507P07Rik | 52.29781034 | 1.107688739 | 0.567141769 | 0.016738169 | 0.042121978 |
| Efcab12 | 37.74578346 | 1.840645364 | 1.239111979 | 0.016794853 | 0.042242467 |
| Etv4 | 263.4641707 | 1.819339961 | 0.997147044 | 0.016803964 | 0.042254307 |
| Gm15527 | 2.426759365 | 1.128953113 | 2.173914055 | 0.016922083 | 0.042515362 |
| Gm26770 | 15.52200165 | 1.485999981 | 0.841689573 | 0.017064821 | 0.042854094 |
| Gm29036 | 1.933820075 | 1.650274036 | 2.952681818 | 0.017274264 | 0.043317724 |
| Plk-ps1 | 9.86103978 | 1.415066299 | 0.804917231 | 0.017317431 | 0.043397599 |
| Gm20400 | 6.587290003 | 1.735821234 | 2.251263362 | 0.017348018 | 0.043451554 |
| Gm21781 | 37.85804223 | 1.009343172 | 0.513354147 | 0.017388314 | 0.043535437 |
| Mir199a-1 | 3.720406918 | 1.743996109 | 2.422253572 | 0.017399067 | 0.043550996 |
| Ppfia3 | 3.087612223 | 1.786831829 | 2.197082698 | 0.017403275 | 0.043555848 |
| Gucy1b1 | 5.84592864 | 1.581779932 | 0.978038405 | 0.017417722 | 0.043586321 |
| Gm15910 | 5.031832868 | 1.752808257 | 2.05298509 | 0.017465319 | 0.043699732 |
| Gm45822 | 2.220171202 | 1.798616531 | 2.247390223 | 0.017468737 | 0.043702586 |
| Slc6a19 | 19.52204303 | 1.586417562 | 0.899955657 | 0.017661351 | 0.044115448 |
| Gm16059 | 6.518785437 | 1.730931342 | 2.022593579 | 0.017739056 | 0.044269209 |
| Ssc5d | 8.781582222 | 1.691800716 | 1.882683219 | 0.017780253 | 0.044360482 |
| Gm40309 | 4.833785372 | 1.949110469 | 1.364582114 | 0.017906804 | 0.044624484 |
| Cenpu | 23.59966192 | 1.107746464 | 0.573996902 | 0.017972962 | 0.044765622 |
| Tmprss15 | 3.734876872 | 2.043540211 | 1.651543727 | 0.018013083 | 0.044853909 |
| Gm39460 | 10.97686744 | 1.182421811 | 0.646402213 | 0.018038791 | 0.04490627 |
| Gm37876 | 5.529903367 | 1.702184502 | 2.224131634 | 0.018144278 | 0.045139595 |
| Gm19721 | 8.019660911 | 1.681878089 | 1.881553835 | 0.018159398 | 0.04516467 |
| Gm12977 | 3.552256092 | 1.730459807 | 2.20738105 | 0.018168147 | 0.0451697 |
| Gm5224 | 4.196724465 | 1.709788337 | 2.331401496 | 0.018275185 | 0.045412857 |
| Adamts9 | 1474.426207 | 1.245982774 | 0.666436654 | 0.018358708 | 0.045602118 |

| | | | | | |
|---|---|---|---|---|---|
| Rerg | 3.250120917 | 1.540944 | 2.630218305 | 0.018371736 | 0.045628572 |
| Capsl | 112.9285285 | 1.148500366 | 0.59143535 | 0.018443725 | 0.045766579 |
| Gm7809 | 6.347204834 | 1.694408822 | 2.152967621 | 0.018480962 | 0.04584645 |
| 8430423G03Rik | 2.506559871 | 1.708629972 | 2.373083074 | 0.018533426 | 0.045958778 |
| Duox2 | 27.63473952 | 1.811253197 | 1.02828046 | 0.018681076 | 0.046277078 |
| Edn1 | 100.9704134 | 1.60806293 | 1.81486893 | 0.018703785 | 0.046321375 |
| Hesx1 | 3.476572002 | 1.634945448 | 2.468487805 | 0.018763467 | 0.046463188 |
| Gm16234 | 4.646105372 | 1.690827502 | 2.154414211 | 0.018786529 | 0.046502293 |
| 2810455O05Rik | 11.04567627 | 1.454205374 | 0.839251775 | 0.018827964 | 0.046592838 |
| Gm43362 | 11.56862139 | 1.444533329 | 0.88600621 | 0.01886351 | 0.046656737 |
| Gm44014 | 5.470627906 | 1.562951756 | 1.011190581 | 0.018876701 | 0.046679928 |
| 2610318M16Rik | 9.430813731 | 1.648589274 | 1.884356321 | 0.018887993 | 0.046699237 |
| Gm47071 | 30.30340303 | 1.001549772 | 0.511474519 | 0.018974141 | 0.046882031 |
| Gm21112 | 6.24910811 | 1.810433609 | 1.45646827 | 0.018980665 | 0.046892115 |
| Macroh2a2 | 128.8549198 | 1.096890989 | 0.572355893 | 0.01910822 | 0.047158675 |
| Gm48838 | 11.70605484 | 1.61045187 | 0.985438007 | 0.019120589 | 0.047183133 |
| Fbxo48 | 11.79454991 | 1.787375298 | 1.334450527 | 0.019175381 | 0.047287939 |
| Mchr1 | 30.92040084 | 1.830535497 | 1.165292309 | 0.01920525 | 0.047343347 |
| Gm42615 | 3.441358014 | 1.821869102 | 1.654172518 | 0.019313722 | 0.047574078 |
| Myl9 | 30.57591637 | 1.764051484 | 1.279618969 | 0.019424597 | 0.047828772 |
| Esco2 | 66.82551036 | 1.054991191 | 0.566032707 | 0.019536036 | 0.048072327 |
| Gm36787 | 12.68743537 | 1.114443455 | 0.609635762 | 0.019588644 | 0.048171293 |
| Gm42587 | 3.286162191 | 1.648602599 | 2.193389351 | 0.019603113 | 0.048194125 |
| Gm26413 | 6.141299459 | 1.628547524 | 2.057627811 | 0.019607264 | 0.04819726 |
| Zfp503 | 9.461740171 | 1.588158706 | 0.966125526 | 0.019609411 | 0.04819726 |
| Serpina3d-ps | 2.280879263 | 1.169170093 | 2.289104787 | 0.019643924 | 0.048257365 |
| Gm42819 | 4.434196752 | 1.640606467 | 2.049523411 | 0.019656252 | 0.048281471 |
| Ccl5 | 10.20942871 | 1.786896487 | 1.421070884 | 0.019684409 | 0.048338259 |
| Greb1 | 24.27093453 | 1.844112887 | 1.320360341 | 0.019809811 | 0.048590245 |
| Col23a1 | 21.68003973 | 1.748494381 | 1.299070461 | 0.019821541 | 0.048607864 |
| Gm49959 | 5.987021957 | 1.557779279 | 2.29360958 | 0.019912551 | 0.048786131 |
| H2-DMb1 | 44.3842243 | 1.40374117 | 0.800275171 | 0.019940786 | 0.048831347 |
| Gm22362 | 2.746342044 | 1.508904723 | 2.45651245 | 0.020057474 | 0.049041023 |
| Nsl1 | 35.0112321 | 1.482764734 | 0.931191848 | 0.020079322 | 0.049088186 |
| Gm24430 | 3.097292663 | 1.591408657 | 2.300550139 | 0.020132285 | 0.04920513 |
| Gm44085 | 3.482229855 | 1.579076731 | 2.283730455 | 0.020212293 | 0.049362956 |
| Gm47218 | 12.28044485 | 1.241016282 | 0.708057835 | 0.020369084 | 0.049726892 |

### Downregulated genes in Rep_early v.s. Hep

| Gene_id | baseMean | log2FoldChange | lfcSE | pvalue | padj |
|---|---|---|---|---|---|
| Slco1a1 | 3231.833501 | -8.155338771 | 0.228416059 | 5.37E-280 | 5.15E-276 |
| Cyp4a14 | 957.4810408 | -6.660245808 | 0.293704887 | 8.24E-115 | 8.32E-112 |
| Cyp4a10 | 1455.843269 | -6.473524164 | 0.293131506 | 3.38E-109 | 2.95E-106 |
| Exoc4 | 1240.354355 | -1.973229567 | 0.094071177 | 1.59E-98 | 1.22E-95 |
| Ccng1 | 1756.337645 | -2.539124388 | 0.123349334 | 8.50E-95 | 5.83E-92 |
| Mdh1 | 6692.928416 | -1.58725566 | 0.078256822 | 3.90E-92 | 2.58E-89 |
| Dcxr | 970.2880377 | -2.650249035 | 0.131622353 | 3.39E-91 | 2.17E-88 |
| Rdh16f2 | 901.8944892 | -3.181173177 | 0.165102128 | 7.90E-84 | 3.99E-81 |
| Lonp2 | 4964.667558 | -1.286337475 | 0.068299289 | 1.16E-79 | 5.17E-77 |
| Retsat | 1990.878222 | -2.721065226 | 0.15047467 | 4.82E-74 | 1.97E-71 |
| Pon1 | 5457.19043 | -2.455286053 | 0.139929937 | 4.98E-70 | 1.84E-67 |
| Kyat1 | 1548.660396 | -2.432459765 | 0.140721374 | 5.81E-68 | 2.03E-65 |
| Cyp2c54 | 2045.687488 | -3.977255698 | 0.231031472 | 1.09E-67 | 3.68E-65 |
| Ugt3a1 | 341.8739437 | -4.024371478 | 0.238947849 | 8.08E-65 | 2.46E-62 |
| Aldh6a1 | 4088.091357 | -1.421427248 | 0.084165193 | 1.45E-64 | 4.28E-62 |
| Got1 | 3111.016453 | -1.917838855 | 0.113844595 | 1.58E-64 | 4.58E-62 |
| Ces1b | 825.8355939 | -2.153339213 | 0.12806553 | 2.99E-64 | 8.56E-62 |
| Ddias | 133.2315969 | -3.456027597 | 0.206671035 | 1.52E-63 | 4.28E-61 |
| Pcca | 1130.442133 | -1.265951995 | 0.075726352 | 3.30E-63 | 9.18E-61 |
| Mup3 | 51356.20701 | -6.86420456 | 0.41331084 | 3.67E-63 | 9.92E-61 |
| Il1rap | 1299.288952 | -2.116105253 | 0.127224957 | 4.24E-63 | 1.13E-60 |
| Acat1 | 3064.272365 | -1.548922763 | 0.093133673 | 9.32E-63 | 2.45E-60 |
| Hibadh | 3140.879133 | -1.566534891 | 0.094339704 | 1.25E-62 | 3.23E-60 |
| Phyh | 14242.3588 | -1.767557922 | 0.106682636 | 2.02E-62 | 5.16E-60 |
| Mgst1 | 26265.261 | -1.454535002 | 0.087766101 | 2.45E-62 | 6.18E-60 |
| Acox1 | 20200.0262 | -1.513248965 | 0.092215809 | 1.16E-61 | 2.90E-59 |
| C8g | 2198.496907 | -1.908470697 | 0.118189736 | 1.71E-59 | 4.05E-57 |
| Decr1 | 2010.108183 | -1.252512104 | 0.079403515 | 1.45E-56 | 3.03E-54 |
| Ppm1k | 619.0194521 | -2.561785499 | 0.16317743 | 1.69E-56 | 3.48E-54 |
| Nudt7 | 3653.312666 | -3.762544025 | 0.241137378 | 5.11E-56 | 1.01E-53 |
| Cyp2c37 | 769.3492101 | -2.058279292 | 0.131981768 | 1.13E-55 | 2.14E-53 |
| Sucla2 | 1769.988243 | -1.016601407 | 0.065631381 | 1.44E-54 | 2.64E-52 |
| Cyp8b1 | 2480.616184 | -6.574614671 | 0.435307804 | 9.28E-53 | 1.61E-50 |
| Gm31583 | 130.9256102 | -4.591506223 | 0.30479088 | 2.33E-52 | 3.93E-50 |
| Glo1 | 2621.082912 | -1.388159331 | 0.091875224 | 3.75E-52 | 6.21E-50 |
| Scnn1a | 247.9029112 | -2.837713746 | 0.188857515 | 4.04E-52 | 6.63E-50 |
| Vwa8 | 2535.465299 | -1.183980762 | 0.080298925 | 1.07E-49 | 1.73E-47 |
| Acaa2 | 7491.870667 | -1.292726579 | 0.087855335 | 1.52E-49 | 2.39E-47 |
| Oat | 4452.204195 | -4.100741006 | 0.281378695 | 2.97E-49 | 4.57E-47 |
| Etfrf1 | 456.0244435 | -1.742800137 | 0.120148118 | 1.97E-48 | 2.95E-46 |
| Cyp4a12b | 309.6514245 | -6.491080274 | 0.455461609 | 2.07E-47 | 2.98E-45 |

| | | | | | |
|---|---|---|---|---|---|
| Kynu | 2213.742199 | -1.654919413 | 0.115683981 | 5.83E-47 | 8.35E-45 |
| Cyp2e1 | 26217.94174 | -2.834597429 | 0.199842971 | 1.02E-46 | 1.43E-44 |
| Zadh2 | 1599.427487 | -1.012343532 | 0.071341838 | 4.62E-46 | 6.29E-44 |
| Sdr9c7 | 474.6272133 | -3.193574604 | 0.227630165 | 4.70E-46 | 6.35E-44 |
| Fah | 5816.048037 | -1.35771972 | 0.096010354 | 5.68E-46 | 7.52E-44 |
| Acadsb | 1773.716809 | -1.642687856 | 0.117301234 | 3.01E-45 | 3.78E-43 |
| Pdcd4 | 1660.903775 | -1.529983886 | 0.10936494 | 4.09E-45 | 5.06E-43 |
| Rmnd5a | 2347.188286 | -1.164718207 | 0.08307382 | 5.10E-45 | 6.27E-43 |
| Lgals9 | 6484.278941 | -1.281578178 | 0.091707244 | 6.27E-45 | 7.66E-43 |
| Atp5pb | 3803.614256 | -1.003432967 | 0.07168266 | 1.07E-44 | 1.29E-42 |
| Suox | 1211.369839 | -1.784363926 | 0.12858541 | 1.46E-44 | 1.75E-42 |
| Glul | 3131.906545 | -2.679792055 | 0.194128622 | 1.76E-44 | 2.09E-42 |
| Mmut | 2419.140154 | -1.451865312 | 0.104671996 | 2.19E-44 | 2.59E-42 |
| Gpld1 | 3098.770974 | -1.140681867 | 0.082170646 | 3.01E-44 | 3.52E-42 |
| Cmbl | 1582.215554 | -1.746297331 | 0.126627982 | 5.11E-44 | 5.90E-42 |
| Dpy19l3 | 414.095172 | -2.253034397 | 0.163856361 | 6.36E-44 | 7.26E-42 |
| Zfp874a | 373.1033332 | -2.018231254 | 0.146669151 | 6.58E-44 | 7.47E-42 |
| Ces2e | 4018.950125 | -2.356748062 | 0.173210025 | 3.72E-43 | 4.13E-41 |
| Rmdn2 | 619.4964329 | -1.423596563 | 0.104139246 | 3.93E-43 | 4.33E-41 |
| Sdhb | 2021.044614 | -1.22440423 | 0.089713697 | 6.35E-43 | 6.97E-41 |
| Sulf2 | 737.8486678 | -3.405075229 | 0.252835417 | 1.60E-42 | 1.73E-40 |
| Dglucy | 996.5419743 | -1.530209742 | 0.112979438 | 1.85E-42 | 1.99E-40 |
| Ces2a | 2508.586908 | -2.561341509 | 0.190100434 | 2.08E-42 | 2.22E-40 |
| Adtrp | 584.4179189 | -1.750152402 | 0.1293789 | 2.71E-42 | 2.86E-40 |
| Dhrs1 | 1686.328456 | -1.156801121 | 0.085533304 | 3.95E-42 | 4.10E-40 |
| Blmh | 878.6867204 | -1.233203009 | 0.091225594 | 4.09E-42 | 4.22E-40 |
| Etfdh | 3039.980827 | -1.412518694 | 0.104913147 | 6.09E-42 | 6.22E-40 |
| Decr2 | 1912.30249 | -1.417049048 | 0.105707731 | 7.77E-42 | 7.84E-40 |
| Amy1 | 1792.166186 | -1.33907014 | 0.10091992 | 9.02E-41 | 8.97E-39 |
| Sdr42e1 | 1075.628772 | -1.544847916 | 0.1167433 | 1.24E-40 | 1.22E-38 |
| Cyp4a32 | 174.0791167 | -3.609391447 | 0.275295684 | 1.68E-40 | 1.64E-38 |
| Mmd2 | 124.1383738 | -3.762866366 | 0.288808929 | 6.39E-40 | 6.10E-38 |
| Lrtm1 | 138.4370808 | -4.776193412 | 0.36755822 | 1.11E-39 | 1.04E-37 |
| Lman2 | 3667.76517 | -1.010167461 | 0.077494313 | 1.12E-39 | 1.05E-37 |
| Gm3734 | 77.73279874 | -4.133840328 | 0.31908453 | 1.71E-39 | 1.58E-37 |
| Cd1d1 | 2649.734967 | -1.181738938 | 0.091671 | 1.72E-38 | 1.56E-36 |
| Serpina1b | 48559.93478 | -1.442946183 | 0.112628622 | 3.43E-38 | 3.07E-36 |
| B630019A10Rik | 383.3661657 | -1.712065482 | 0.134171978 | 5.22E-38 | 4.64E-36 |
| Gcdh | 4483.33069 | -1.528541691 | 0.119739788 | 6.44E-38 | 5.67E-36 |
| Acad11 | 3179.576997 | -1.282824485 | 0.100507098 | 6.57E-38 | 5.76E-36 |
| Hsd17b4 | 4942.699386 | -1.101179157 | 0.086518225 | 1.52E-37 | 1.30E-35 |
| Rtn4ip1 | 375.6156007 | -1.250075949 | 0.098422992 | 1.76E-37 | 1.50E-35 |
| Ceacam1 | 1774.233113 | -1.692091098 | 0.133774411 | 1.94E-37 | 1.65E-35 |

| | | | | | |
|---|---|---|---|---|---|
| Amt | 665.4295758 | -1.472989033 | 0.116604755 | 3.29E-37 | 2.77E-35 |
| Crot | 8715.258427 | -1.908630319 | 0.152514906 | 9.62E-37 | 7.99E-35 |
| Cpox | 1623.956429 | -2.189605324 | 0.175232736 | 1.03E-36 | 8.50E-35 |
| Gm10319 | 468.3267566 | -2.3544645 | 0.188220823 | 1.11E-36 | 9.14E-35 |
| Commd3 | 1397.901892 | -1.088953569 | 0.086643171 | 1.23E-36 | 1.00E-34 |
| Ei24 | 2834.461251 | -1.112020997 | 0.088605821 | 1.48E-36 | 1.20E-34 |
| Tuba4a | 2315.478556 | -1.725202486 | 0.138411437 | 1.71E-36 | 1.39E-34 |
| Marchf6 | 2726.655581 | -1.03606771 | 0.082600965 | 1.81E-36 | 1.46E-34 |
| Aldh7a1 | 6689.508536 | -1.346483679 | 0.107817829 | 2.25E-36 | 1.81E-34 |
| Scp2 | 35011.92052 | -1.708716976 | 0.137037618 | 2.56E-36 | 2.05E-34 |
| Lifr | 2127.39759 | -2.177552772 | 0.175930126 | 4.45E-36 | 3.50E-34 |
| Gm3839 | 231.9304227 | -4.682745767 | 0.380804438 | 4.72E-36 | 3.69E-34 |
| Acpp | 101.0676281 | -5.564999398 | 0.450012723 | 5.31E-36 | 4.13E-34 |
| Rnf103 | 1672.8662 | -1.244072505 | 0.100752617 | 1.58E-35 | 1.21E-33 |
| Cmtm6 | 1799.804743 | -1.145183347 | 0.093192647 | 3.67E-35 | 2.79E-33 |
| Bet1 | 721.1215644 | -1.34285523 | 0.109576724 | 4.39E-35 | 3.33E-33 |
| Cyp2c50 | 5996.0609 | -2.904267997 | 0.238920963 | 5.54E-35 | 4.17E-33 |
| Fetub | 6598.019926 | -2.097107324 | 0.172533539 | 7.45E-35 | 5.58E-33 |
| Ahcy | 11950.41057 | -1.444425856 | 0.119178137 | 2.25E-34 | 1.67E-32 |
| Acsl1 | 6853.524814 | -2.155042533 | 0.179837264 | 3.62E-34 | 2.64E-32 |
| Arhgef37 | 162.4489336 | -2.640792894 | 0.220584051 | 4.75E-34 | 3.44E-32 |
| Rgn | 6180.406756 | -2.749865054 | 0.230263686 | 7.28E-34 | 5.21E-32 |
| Pnpla8 | 2023.679281 | -1.12968343 | 0.09409742 | 1.24E-33 | 8.76E-32 |
| Cyp4v3 | 5061.225712 | -1.550183594 | 0.130322819 | 3.39E-33 | 2.38E-31 |
| Endog | 510.526049 | -1.964108923 | 0.166302067 | 5.33E-33 | 3.70E-31 |
| Atp8b5 | 91.11542403 | -5.132207148 | 0.436548214 | 8.35E-33 | 5.68E-31 |
| Hectd2os | 147.589944 | -4.211624709 | 0.362516209 | 2.12E-32 | 1.43E-30 |
| Cpt2 | 1647.538098 | -1.413094992 | 0.12063766 | 2.75E-32 | 1.83E-30 |
| Lama3 | 384.3308152 | -3.68321651 | 0.317103575 | 5.17E-32 | 3.41E-30 |
| Serpina1d | 23703.15904 | -2.518734662 | 0.218750786 | 6.61E-32 | 4.33E-30 |
| Ndufb5 | 909.5620495 | -1.19925546 | 0.103220745 | 1.12E-31 | 7.22E-30 |
| Pnpla7 | 1316.832108 | -1.108179886 | 0.095769084 | 1.82E-31 | 1.16E-29 |
| Cyp2c67 | 1062.441841 | -3.27262527 | 0.286481934 | 2.42E-31 | 1.54E-29 |
| Tars | 1991.618392 | -1.422392885 | 0.123745552 | 3.38E-31 | 2.14E-29 |
| Aadat | 705.0290745 | -1.927257261 | 0.168748185 | 4.24E-31 | 2.68E-29 |
| Slc30a10 | 412.4097797 | -2.77313028 | 0.244147895 | 5.40E-31 | 3.38E-29 |
| Nars2 | 324.4498423 | -1.758378987 | 0.154882842 | 1.30E-30 | 7.97E-29 |
| Abcd3 | 6075.360782 | -1.624408421 | 0.143001771 | 1.37E-30 | 8.37E-29 |
| Dpyd | 5085.169895 | -1.526509209 | 0.134091568 | 1.71E-30 | 1.04E-28 |
| Pxmp2 | 769.3839394 | -1.504301377 | 0.132772851 | 2.08E-30 | 1.26E-28 |
| Ncald | 313.2759059 | -1.871845423 | 0.165654934 | 2.18E-30 | 1.31E-28 |
| Agmat | 1547.729597 | -1.6670846 | 0.148165458 | 2.85E-30 | 1.70E-28 |
| Thnsl2 | 904.7990693 | -1.00484842 | 0.08855091 | 3.25E-30 | 1.94E-28 |

| | | | | | |
|---|---|---|---|---|---|
| Mccc2 | 854.8531618 | -1.168674561 | 0.103133714 | 3.41E-30 | 2.02E-28 |
| Cyp7b1 | 930.2511745 | -3.292929138 | 0.294404845 | 4.13E-30 | 2.41E-28 |
| Cox7a2 | 1252.534603 | -1.015613353 | 0.089666415 | 4.13E-30 | 2.41E-28 |
| Aldh9a1 | 3308.955386 | -1.143234836 | 0.100873835 | 5.27E-30 | 3.05E-28 |
| Tubb4b | 3591.948875 | -1.3887924 | 0.12394216 | 9.43E-30 | 5.34E-28 |
| Adh6-ps1 | 195.9580178 | -3.370270309 | 0.304430523 | 1.32E-29 | 7.44E-28 |
| Irgm1 | 911.9365283 | -1.296067284 | 0.11615905 | 1.90E-29 | 1.06E-27 |
| Hal | 4532.346834 | -2.626095942 | 0.238100636 | 2.08E-29 | 1.16E-27 |
| Gstz1 | 5725.168937 | -1.361283779 | 0.122594827 | 3.17E-29 | 1.74E-27 |
| Serpina1c | 11044.97746 | -1.539334727 | 0.139476588 | 5.31E-29 | 2.86E-27 |
| Paip2b | 435.6410582 | -1.526052505 | 0.138204026 | 5.60E-29 | 3.00E-27 |
| Slc10a1 | 4469.959654 | -2.251048621 | 0.205136552 | 5.67E-29 | 3.02E-27 |
| Serpina1e | 33093.53306 | -4.746396576 | 0.434814696 | 5.69E-29 | 3.02E-27 |
| Srd5a1 | 566.4992275 | -2.620386367 | 0.240030131 | 9.15E-29 | 4.79E-27 |
| Atp11c | 2915.515123 | -1.302392418 | 0.118574038 | 1.06E-28 | 5.53E-27 |
| Sec24d | 1963.099585 | -1.104401679 | 0.100296646 | 1.29E-28 | 6.70E-27 |
| Adh4 | 927.309734 | -1.750574047 | 0.160198199 | 1.48E-28 | 7.67E-27 |
| Cbs | 5761.136914 | -1.518435039 | 0.13904252 | 1.80E-28 | 9.26E-27 |
| Pecr | 1576.428862 | -1.553762476 | 0.142568879 | 2.67E-28 | 1.36E-26 |
| Ephx2 | 5638.831254 | -1.221048587 | 0.112021293 | 3.01E-28 | 1.52E-26 |
| Clec2h | 140.0322225 | -3.802940333 | 0.353446548 | 3.25E-28 | 1.64E-26 |
| Ethe1 | 1090.014811 | -1.847738526 | 0.170353764 | 3.37E-28 | 1.70E-26 |
| Adh5 | 4083.387451 | -1.034252913 | 0.094792728 | 4.24E-28 | 2.12E-26 |
| Fermt2 | 1856.403137 | -1.036901341 | 0.095344441 | 6.17E-28 | 3.06E-26 |
| Alas2 | 630.1115107 | -2.929192705 | 0.273228144 | 6.52E-28 | 3.23E-26 |
| Gm4952 | 808.0923256 | -2.511536321 | 0.234157676 | 7.35E-28 | 3.62E-26 |
| Nbas | 835.6906295 | -1.251433267 | 0.115966904 | 1.19E-27 | 5.82E-26 |
| Mindy3 | 577.7466443 | -1.161390654 | 0.107631601 | 1.35E-27 | 6.56E-26 |
| Mdm2 | 1354.690223 | -1.254709691 | 0.116749335 | 1.83E-27 | 8.85E-26 |
| Aldh1l1 | 17153.78296 | -1.711813156 | 0.160273956 | 1.87E-27 | 9.01E-26 |
| Tspan33 | 253.4158232 | -2.258691877 | 0.212080697 | 2.16E-27 | 1.03E-25 |
| Mbl2 | 6891.189503 | -4.533134575 | 0.428568858 | 2.26E-27 | 1.08E-25 |
| Fpgs | 1692.398453 | -1.555621757 | 0.146429741 | 4.56E-27 | 2.14E-25 |
| Ivd | 2440.98636 | -1.419968506 | 0.133705311 | 5.83E-27 | 2.73E-25 |
| Gbp10 | 43.84532083 | -4.327395819 | 0.412992023 | 1.25E-26 | 5.76E-25 |
| Cisd1 | 1209.97067 | -1.099060832 | 0.10378337 | 1.26E-26 | 5.79E-25 |
| Pgpep1 | 1420.750066 | -1.273966274 | 0.12065596 | 1.26E-26 | 5.80E-25 |
| Trp53inp1 | 4879.480625 | -1.802949001 | 0.17149749 | 1.27E-26 | 5.82E-25 |
| Lamp2 | 10509.48394 | -1.092099902 | 0.103448383 | 1.80E-26 | 8.16E-25 |
| Uso1 | 2150.984933 | -1.0515924 | 0.100014392 | 2.96E-26 | 1.32E-24 |
| Nit2 | 1454.790917 | -1.012184198 | 0.096803788 | 5.78E-26 | 2.55E-24 |
| Igfals | 1327.981817 | -1.651826314 | 0.159924914 | 6.95E-26 | 3.04E-24 |
| Car1 | 65.05378439 | -2.685374047 | 0.260899355 | 8.41E-26 | 3.64E-24 |

| | | | | | |
|---|---|---|---|---|---|
| Bbox1 | 1261.82703 | -1.110227937 | 0.107105023 | 1.31E-25 | 5.64E-24 |
| Clock | 1151.18612 | -1.110334802 | 0.10714877 | 1.36E-25 | 5.82E-24 |
| Klf9 | 2054.218084 | -1.432349905 | 0.138797788 | 1.41E-25 | 6.04E-24 |
| Sort1 | 1877.760543 | -1.560391304 | 0.15149648 | 1.63E-25 | 6.95E-24 |
| Ubr3 | 2560.489468 | -1.068619521 | 0.103189687 | 1.72E-25 | 7.27E-24 |
| Sephs2 | 7898.028626 | -1.264942124 | 0.122907036 | 1.86E-25 | 7.85E-24 |
| Etfa | 3659.776315 | -1.045799093 | 0.101222006 | 2.04E-25 | 8.58E-24 |
| Mpv17l | 340.9810783 | -2.335271168 | 0.229132235 | 2.45E-25 | 1.02E-23 |
| Tfpi2 | 590.9337248 | -2.067051381 | 0.202806119 | 2.86E-25 | 1.18E-23 |
| Cyp2d40 | 288.3421865 | -2.558826117 | 0.251723602 | 2.86E-25 | 1.18E-23 |
| 0610030E20Rik | 550.7116438 | -1.082782673 | 0.10539226 | 3.51E-25 | 1.44E-23 |
| Slc31a1 | 2630.225584 | -1.10986936 | 0.109193373 | 4.63E-25 | 1.89E-23 |
| Ldhd | 2174.866764 | -1.219324917 | 0.119582169 | 5.40E-25 | 2.19E-23 |
| Tex2 | 2484.501089 | -1.027672342 | 0.100527501 | 5.73E-25 | 2.32E-23 |
| Ttc39c | 4446.110703 | -2.183785322 | 0.21585402 | 6.36E-25 | 2.56E-23 |
| Sesn2 | 418.6599438 | -1.667181304 | 0.165508425 | 1.45E-24 | 5.77E-23 |
| Cd302 | 2259.739564 | -1.388967565 | 0.137820649 | 2.00E-24 | 7.81E-23 |
| Acsl5 | 5281.400977 | -1.131016153 | 0.112241736 | 2.27E-24 | 8.86E-23 |
| Cyp2j5 | 6277.69769 | -1.56333438 | 0.156196391 | 2.44E-24 | 9.52E-23 |
| F11 | 1226.561025 | -1.525249948 | 0.152834122 | 3.46E-24 | 1.33E-22 |
| Csrp3 | 103.5276539 | -2.712066808 | 0.273743682 | 4.02E-24 | 1.54E-22 |
| Cat | 37763.41738 | -1.096861853 | 0.109329042 | 4.11E-24 | 1.57E-22 |
| Enho | 174.0155649 | -2.027311278 | 0.204846142 | 6.05E-24 | 2.30E-22 |
| Slc27a5 | 7327.457284 | -2.128305617 | 0.215208446 | 6.50E-24 | 2.47E-22 |
| Echs1 | 3041.637631 | -1.037964323 | 0.104082791 | 8.23E-24 | 3.11E-22 |
| Aen | 484.0944922 | -1.500743548 | 0.151622406 | 9.46E-24 | 3.57E-22 |
| Adipor2 | 6140.491204 | -1.258898085 | 0.126766193 | 9.69E-24 | 3.65E-22 |
| Gm53019 | 57.0915265 | -5.254273301 | 0.534651364 | 1.01E-23 | 3.79E-22 |
| Bscl2 | 676.9304846 | -1.025194761 | 0.10310531 | 1.16E-23 | 4.31E-22 |
| Bach2 | 164.4623929 | -1.728839575 | 0.175242471 | 1.18E-23 | 4.38E-22 |
| Hagh | 2232.536391 | -1.297625048 | 0.131302905 | 1.50E-23 | 5.53E-22 |
| Slc22a7 | 142.8917039 | -2.904826528 | 0.297846731 | 1.62E-23 | 5.95E-22 |
| Zap70 | 58.67807472 | -3.745911459 | 0.38521894 | 2.09E-23 | 7.62E-22 |
| Mup21 | 854.3786608 | -6.060284584 | 0.626850192 | 2.24E-23 | 8.17E-22 |
| Qdpr | 2862.326327 | -1.337703446 | 0.136028567 | 2.62E-23 | 9.44E-22 |
| Bhmt-ps1 | 131.2916615 | -3.274101033 | 0.338354383 | 2.95E-23 | 1.06E-21 |
| Clpx | 2445.584159 | -1.814741441 | 0.186672023 | 4.02E-23 | 1.43E-21 |
| 2010315B03Rik | 389.9599704 | -1.049330356 | 0.107038412 | 4.36E-23 | 1.55E-21 |
| Cyp2d10 | 9162.501461 | -1.35692663 | 0.138093915 | 4.39E-23 | 1.55E-21 |
| Susd4 | 154.3446031 | -8.848544192 | 0.882111044 | 4.78E-23 | 1.67E-21 |
| Aox3 | 5233.849104 | -1.827934952 | 0.188585085 | 5.32E-23 | 1.86E-21 |
| Mpst | 1160.581906 | -1.50108405 | 0.154969344 | 7.94E-23 | 2.75E-21 |
| Clec2d | 770.1801628 | -1.559627569 | 0.162388599 | 1.59E-22 | 5.44E-21 |

| | | | | | |
|---|---|---|---|---|---|
| Slc19a2 | 1680.009348 | -1.24478646 | 0.129897468 | 2.85E-22 | 9.63E-21 |
| Ghr | 4664.985053 | -1.671448568 | 0.175571571 | 3.29E-22 | 1.11E-20 |
| Gramd1c | 524.3665782 | -1.304223405 | 0.136341416 | 3.39E-22 | 1.14E-20 |
| Ehhadh | 2941.345095 | -1.543835382 | 0.162098978 | 3.42E-22 | 1.15E-20 |
| Elovl5 | 3343.903538 | -1.533803995 | 0.161887916 | 6.01E-22 | 2.00E-20 |
| Dbt | 1287.253009 | -1.508219802 | 0.159204857 | 6.09E-22 | 2.02E-20 |
| Adhfe1 | 1281.546138 | -1.350557739 | 0.142739609 | 7.82E-22 | 2.57E-20 |
| Lap3 | 3972.861554 | -1.055355521 | 0.111098649 | 8.10E-22 | 2.66E-20 |
| Dnajc28 | 155.5029066 | -1.379688786 | 0.14627217 | 1.07E-21 | 3.49E-20 |
| Phlda3 | 436.4539258 | -2.609378674 | 0.279822194 | 1.12E-21 | 3.65E-20 |
| Albfm1 | 327.0383192 | -1.542737239 | 0.164068516 | 1.16E-21 | 3.77E-20 |
| Tent5c | 81.28225421 | -4.335161049 | 0.471057744 | 2.39E-21 | 7.63E-20 |
| Slc33a1 | 804.8745948 | -1.239101155 | 0.132504527 | 2.73E-21 | 8.66E-20 |
| Sgpp1 | 1001.215748 | -1.117628597 | 0.119707973 | 3.63E-21 | 1.14E-19 |
| Cyp4a12a | 2016.484184 | -6.516189965 | 0.71541282 | 4.31E-21 | 1.35E-19 |
| Erbb4 | 257.6337189 | -2.512198723 | 0.273447401 | 4.35E-21 | 1.36E-19 |
| Hsd17b10 | 1224.27121 | -1.197546325 | 0.128974663 | 5.00E-21 | 1.56E-19 |
| Abhd6 | 379.1311588 | -1.216090513 | 0.131214375 | 6.11E-21 | 1.89E-19 |
| Gstk1 | 901.233392 | -1.140158469 | 0.123092331 | 7.21E-21 | 2.22E-19 |
| Abat | 4071.826158 | -2.160401787 | 0.236152286 | 7.84E-21 | 2.41E-19 |
| Gm20319 | 240.7847704 | -1.93171514 | 0.211103383 | 8.30E-21 | 2.54E-19 |
| Cldn12 | 1341.156379 | -1.018006367 | 0.109966541 | 8.75E-21 | 2.67E-19 |
| Tmem205 | 3585.068176 | -1.11567146 | 0.121048155 | 8.77E-21 | 2.68E-19 |
| Amdhd1 | 2285.910595 | -1.10305088 | 0.119244178 | 8.84E-21 | 2.69E-19 |
| Gfm2 | 749.4885873 | -1.160580437 | 0.125965867 | 1.12E-20 | 3.41E-19 |
| Hsd11b1 | 4030.393339 | -1.989185347 | 0.21932998 | 1.67E-20 | 5.01E-19 |
| Bhmt | 28291.84794 | -1.945012793 | 0.215785599 | 2.00E-20 | 5.97E-19 |
| C130083M11Rik | 64.46695987 | -2.304936649 | 0.254985198 | 2.04E-20 | 6.08E-19 |
| Oxnad1 | 403.4284322 | -1.09235833 | 0.119251034 | 2.07E-20 | 6.16E-19 |
| Sult1b1 | 230.8859947 | -2.385750459 | 0.264690562 | 2.08E-20 | 6.17E-19 |
| Gstp1 | 20998.42207 | -1.384083612 | 0.151996623 | 2.14E-20 | 6.32E-19 |
| Gria3 | 237.8418521 | -2.186399651 | 0.242140915 | 2.21E-20 | 6.52E-19 |
| Chpt1 | 1652.0309 | -1.077029892 | 0.118041331 | 2.81E-20 | 8.24E-19 |
| Slco2a1 | 1684.378736 | -1.444612187 | 0.159118998 | 3.20E-20 | 9.36E-19 |
| Agxt | 2408.901472 | -1.960107522 | 0.218766188 | 3.37E-20 | 9.86E-19 |
| Hacl1 | 271.9805219 | -1.933759612 | 0.215315026 | 3.83E-20 | 1.11E-18 |
| Serpina3k | 92629.62925 | -2.615991586 | 0.292892207 | 4.00E-20 | 1.16E-18 |
| Sucnr1 | 286.146724 | -2.93376756 | 0.330041426 | 5.01E-20 | 1.44E-18 |
| Ireb2 | 1947.671418 | -1.096294032 | 0.121470394 | 6.89E-20 | 1.96E-18 |
| Cpsf4l | 47.67400905 | -2.869768738 | 0.322458943 | 7.40E-20 | 2.10E-18 |
| Ebpl | 725.6166904 | -1.053482133 | 0.116840232 | 7.85E-20 | 2.21E-18 |
| Gm5424 | 294.2463579 | -2.167953737 | 0.243931411 | 8.07E-20 | 2.27E-18 |
| Hsd17b2 | 1928.589796 | -1.574232718 | 0.176107721 | 8.29E-20 | 2.32E-18 |

| | | | | | |
|---|---|---|---|---|---|
| Slc22a15 | 442.4549186 | -1.508328188 | 0.168822535 | 9.56E-20 | 2.67E-18 |
| Tmem254a | 3434.371436 | -1.060612444 | 0.11826947 | 1.02E-19 | 2.83E-18 |
| Grhpr | 2484.963076 | -1.187574987 | 0.132446749 | 1.06E-19 | 2.94E-18 |
| Mup14 | 220.2115739 | -11.06048669 | 1.369415625 | 1.08E-19 | 2.99E-18 |
| C9 | 5062.348322 | -1.845687865 | 0.208570976 | 1.39E-19 | 3.82E-18 |
| Ero1b | 937.4064343 | -1.157874689 | 0.129679986 | 1.52E-19 | 4.16E-18 |
| Pdk1 | 856.9702466 | -1.447036961 | 0.163312262 | 1.93E-19 | 5.24E-18 |
| Mug1 | 47630.71591 | -1.966531893 | 0.223865745 | 2.22E-19 | 5.99E-18 |
| Plxna2 | 794.7845082 | -1.760155517 | 0.200588844 | 3.27E-19 | 8.76E-18 |
| Sult2a8 | 4597.287247 | -8.432442461 | 0.978509713 | 3.44E-19 | 9.18E-18 |
| Pbld2 | 1290.443319 | -1.578295277 | 0.180103561 | 3.82E-19 | 1.02E-17 |
| Aadac | 5221.332697 | -1.634676067 | 0.186826131 | 4.10E-19 | 1.09E-17 |
| Chic1 | 76.62433719 | -7.525369377 | 0.830090347 | 5.69E-19 | 1.50E-17 |
| Tlcd2 | 949.7024691 | -1.64989136 | 0.18944809 | 5.88E-19 | 1.55E-17 |
| Slc25a13 | 2886.356758 | -1.043551198 | 0.119146024 | 6.86E-19 | 1.80E-17 |
| Slc25a42 | 1176.236855 | -1.042468809 | 0.118946477 | 7.44E-19 | 1.93E-17 |
| Enpep | 1055.792282 | -1.122359445 | 0.1287227 | 8.27E-19 | 2.14E-17 |
| Gm36419 | 51.08797354 | -3.276748686 | 0.38193606 | 8.73E-19 | 2.25E-17 |
| Slc46a1 | 590.6467949 | -1.002634201 | 0.11458278 | 9.30E-19 | 2.39E-17 |
| Gm2788 | 73.24903233 | -3.190205448 | 0.373007903 | 1.00E-18 | 2.56E-17 |
| Tsc22d1 | 3636.47621 | -2.135203582 | 0.248811637 | 1.02E-18 | 2.61E-17 |
| Wdtc1 | 1945.6282 | -1.005742075 | 0.115878934 | 1.69E-18 | 4.25E-17 |
| Fam210a | 839.9343699 | -1.010876333 | 0.117020768 | 2.38E-18 | 5.89E-17 |
| Ccdc162 | 182.4211288 | -1.517347678 | 0.177128261 | 2.58E-18 | 6.38E-17 |
| Ppdpf | 814.6673797 | -1.002919085 | 0.116610603 | 2.99E-18 | 7.34E-17 |
| Fam214a | 1187.745869 | -1.096318189 | 0.127599263 | 3.18E-18 | 7.81E-17 |
| Aaas | 378.9971068 | -1.162736962 | 0.136168006 | 4.74E-18 | 1.15E-16 |
| Hrg | 9857.429302 | -1.35650863 | 0.159326731 | 5.24E-18 | 1.27E-16 |
| Gm4756 | 551.3869792 | -1.531727044 | 0.181542433 | 7.72E-18 | 1.85E-16 |
| Slc45a4 | 462.5715167 | -1.099657439 | 0.129707703 | 8.70E-18 | 2.08E-16 |
| Pm20d1 | 1373.939635 | -1.706184579 | 0.203385638 | 8.81E-18 | 2.10E-16 |
| Pnkd | 1949.121618 | -1.270433715 | 0.150505674 | 9.33E-18 | 2.22E-16 |
| Glyat | 2344.817 | -1.324979021 | 0.157166532 | 9.55E-18 | 2.27E-16 |
| Abhd3 | 598.7206968 | -1.104845259 | 0.130761933 | 1.07E-17 | 2.54E-16 |
| Urad | 520.5567161 | -1.510715927 | 0.1799979 | 1.09E-17 | 2.57E-16 |
| Insig2 | 2707.844627 | -2.178644842 | 0.262670351 | 1.30E-17 | 3.04E-16 |
| AU022252 | 856.6742588 | -1.03847757 | 0.12310557 | 1.39E-17 | 3.26E-16 |
| Gbf1 | 1660.151244 | -1.058315763 | 0.125860693 | 1.60E-17 | 3.73E-16 |
| Elac1 | 188.7118656 | -1.369324979 | 0.163828587 | 1.72E-17 | 4.01E-16 |
| Cyp2u1 | 181.4334045 | -2.137142369 | 0.259283355 | 2.14E-17 | 4.94E-16 |
| Atg5 | 665.5256706 | -1.202649826 | 0.144118052 | 2.33E-17 | 5.37E-16 |
| Slc39a8 | 361.1238199 | -1.190460088 | 0.142929666 | 2.69E-17 | 6.16E-16 |
| Scrn3 | 592.1068952 | -1.41623237 | 0.170773649 | 2.71E-17 | 6.19E-16 |

| | | | | | |
|---|---|---|---|---|---|
| Slc22a28 | 30.51193808 | -6.060618739 | 0.721808945 | 2.85E-17 | 6.49E-16 |
| Slc25a15 | 4594.865544 | -1.1291272 | 0.136050027 | 3.90E-17 | 8.78E-16 |
| Sardh | 8546.791883 | -1.56303958 | 0.189696008 | 4.00E-17 | 8.99E-16 |
| Gramd3 | 581.0196127 | -1.051744341 | 0.126789506 | 4.17E-17 | 9.36E-16 |
| Ppp1r3c | 791.1485404 | -1.786775025 | 0.218125354 | 4.19E-17 | 9.38E-16 |
| Gm30262 | 110.6788181 | -2.086815126 | 0.256556604 | 5.26E-17 | 1.17E-15 |
| Cyb5b | 6724.019993 | -1.231557749 | 0.148826503 | 5.40E-17 | 1.20E-15 |
| Dgka | 268.7516958 | -1.262506747 | 0.153579144 | 5.82E-17 | 1.28E-15 |
| 4931406C07Rik | 2753.613811 | -1.051192492 | 0.12721332 | 5.91E-17 | 1.30E-15 |
| Ulk2 | 1403.785086 | -1.038946368 | 0.126733611 | 9.35E-17 | 2.01E-15 |
| Cdkn1a | 1294.551865 | -2.330747782 | 0.290348643 | 1.09E-16 | 2.34E-15 |
| Paqr9 | 2312.45477 | -3.741789312 | 0.470304753 | 1.15E-16 | 2.46E-15 |
| Plbd1 | 203.1555263 | -2.442609093 | 0.306107279 | 1.57E-16 | 3.31E-15 |
| Acot1 | 646.2609797 | -2.833598381 | 0.357534104 | 2.02E-16 | 4.23E-15 |
| Scp2-ps2 | 87.58193697 | -1.947734789 | 0.244056424 | 2.22E-16 | 4.64E-15 |
| Alb | 764512.0377 | -1.231695404 | 0.152860701 | 2.39E-16 | 4.98E-15 |
| Coq8a | 4650.846139 | -1.674665186 | 0.208911252 | 2.40E-16 | 5.00E-15 |
| Slc6a12 | 829.19693 | -1.528991589 | 0.191737429 | 2.79E-16 | 5.79E-15 |
| Igf1 | 5603.819491 | -1.483795387 | 0.186019925 | 3.42E-16 | 7.06E-15 |
| Slc34a2 | 355.0723003 | -3.209464554 | 0.403713718 | 3.66E-16 | 7.54E-15 |
| Thada | 378.6913192 | -1.066921558 | 0.133589356 | 5.44E-16 | 1.10E-14 |
| Gm4951 | 1130.968667 | -1.359542978 | 0.172046826 | 6.93E-16 | 1.39E-14 |
| Gm16157 | 72.67797902 | -2.265889971 | 0.290279794 | 6.99E-16 | 1.41E-14 |
| Ccnc | 440.5525056 | -1.03656063 | 0.130689977 | 8.96E-16 | 1.78E-14 |
| Ppp1r9a | 796.1139589 | -1.572235671 | 0.200377031 | 8.98E-16 | 1.78E-14 |
| Tmem19 | 1666.704453 | -1.384664309 | 0.176004205 | 9.08E-16 | 1.80E-14 |
| Ndrg2 | 10572.0384 | -1.248086008 | 0.158231276 | 9.30E-16 | 1.84E-14 |
| Paox | 467.222011 | -1.823501685 | 0.234093977 | 1.11E-15 | 2.18E-14 |
| Rnf125 | 1114.113492 | -2.427429624 | 0.313347681 | 1.18E-15 | 2.31E-14 |
| Klhl22 | 440.1582846 | -1.080858587 | 0.137422515 | 1.38E-15 | 2.68E-14 |
| Sult5a1 | 69.8770676 | -8.442070724 | 1.031999217 | 1.39E-15 | 2.70E-14 |
| Msrb1 | 2494.346088 | -1.002363559 | 0.127287096 | 1.44E-15 | 2.79E-14 |
| Myo1b | 5169.108824 | -1.023618871 | 0.130866519 | 1.71E-15 | 3.29E-14 |
| Mfsd4b3-ps | 164.9892399 | -1.182404271 | 0.151318922 | 1.89E-15 | 3.63E-14 |
| Hpd | 11526.28301 | -1.940059718 | 0.252192454 | 2.04E-15 | 3.91E-14 |
| Zfp958 | 172.4961054 | -1.408931858 | 0.181804722 | 2.35E-15 | 4.49E-14 |
| Iigp1 | 3602.198393 | -1.392122722 | 0.180237291 | 2.55E-15 | 4.85E-14 |
| Crem | 318.6324204 | -1.15703146 | 0.148707938 | 2.61E-15 | 4.95E-14 |
| Sugct | 300.8437445 | -1.030843991 | 0.132927029 | 3.65E-15 | 6.88E-14 |
| Ebp | 1992.520831 | -1.164280871 | 0.150907167 | 4.09E-15 | 7.69E-14 |
| Slc38a3 | 18171.6763 | -1.202916303 | 0.155989771 | 4.26E-15 | 7.99E-14 |
| Mgll | 559.5147826 | -1.504583824 | 0.196852916 | 4.86E-15 | 9.05E-14 |
| Pde9a | 226.6531847 | -1.449744293 | 0.190688258 | 6.83E-15 | 1.25E-13 |

| | | | | | |
|---|---|---|---|---|---|
| Nr0b2 | 428.6848339 | -1.891639451 | 0.250939744 | 7.35E-15 | 1.34E-13 |
| Gm5096 | 303.7068556 | -2.766172999 | 0.37047435 | 8.17E-15 | 1.49E-13 |
| Pigr | 15895.96808 | -1.566450085 | 0.207538595 | 8.95E-15 | 1.63E-13 |
| Rida | 5117.22602 | -1.133237282 | 0.149466345 | 1.15E-14 | 2.08E-13 |
| Cyp4f17 | 360.5819921 | -1.299350291 | 0.171834913 | 1.17E-14 | 2.12E-13 |
| Ifit3 | 105.2919954 | -1.699621227 | 0.226796275 | 1.24E-14 | 2.24E-13 |
| Bdh2 | 151.865255 | -2.083118718 | 0.280214606 | 1.38E-14 | 2.48E-13 |
| Nat8f1 | 1071.00527 | -1.548307151 | 0.206948616 | 1.53E-14 | 2.73E-13 |
| Lmbrd2 | 1412.847547 | -1.008211514 | 0.133501035 | 1.79E-14 | 3.18E-13 |
| Tmem86b | 739.1731778 | -1.575900301 | 0.21163183 | 1.94E-14 | 3.43E-13 |
| Ube2e1 | 323.423937 | -1.023029299 | 0.135507018 | 1.97E-14 | 3.47E-13 |
| Gbp11 | 38.11692445 | -2.554886431 | 0.346695165 | 2.21E-14 | 3.88E-13 |
| Chid1 | 578.5841363 | -1.056366871 | 0.14078556 | 2.51E-14 | 4.37E-13 |
| Mgmt | 316.5917011 | -1.944793689 | 0.264396532 | 2.72E-14 | 4.73E-13 |
| Slc15a5 | 31.47402286 | -6.053374705 | 0.812925644 | 2.77E-14 | 4.79E-13 |
| Slc22a1 | 2885.581621 | -1.521448171 | 0.205374227 | 2.76E-14 | 4.79E-13 |
| Mup-ps13 | 30.41902683 | -10.40536243 | 2.998579835 | 3.06E-14 | 5.28E-13 |
| Cfhr1 | 1272.927003 | -1.764981543 | 0.241044512 | 3.06E-14 | 5.29E-13 |
| Klhl7 | 227.559492 | -1.095433038 | 0.146668184 | 3.09E-14 | 5.32E-13 |
| Cyp4f14 | 3521.861278 | -1.809710029 | 0.246174708 | 3.09E-14 | 5.32E-13 |
| Nxt2 | 399.4182943 | -1.389691644 | 0.187479086 | 3.09E-14 | 5.32E-13 |
| BC024386 | 1569.127979 | -1.470404029 | 0.199267577 | 3.34E-14 | 5.73E-13 |
| Sec23b | 1351.920018 | -1.071781252 | 0.143582087 | 3.55E-14 | 6.08E-13 |
| Cyp2d9 | 12943.9436 | -1.809794208 | 0.246808768 | 3.58E-14 | 6.12E-13 |
| Extl1 | 107.5850365 | -3.10173759 | 0.428639657 | 4.04E-14 | 6.90E-13 |
| Lhpp | 353.9288928 | -1.99488667 | 0.273642952 | 4.32E-14 | 7.35E-13 |
| Kmo | 2428.998225 | -1.578339522 | 0.215794799 | 5.22E-14 | 8.81E-13 |
| Abhd18 | 271.6325461 | -1.405862665 | 0.191725191 | 5.51E-14 | 9.22E-13 |
| Gnat1 | 380.1575523 | -1.36859185 | 0.186912177 | 6.70E-14 | 1.12E-12 |
| Ifi47 | 428.2855406 | -1.178265446 | 0.160543559 | 7.07E-14 | 1.18E-12 |
| Slc1a2 | 607.4669923 | -3.295359641 | 0.464289965 | 8.35E-14 | 1.38E-12 |
| Casd1 | 601.5378374 | -1.139499792 | 0.156084838 | 1.02E-13 | 1.67E-12 |
| Shmt1 | 2007.223139 | -1.044475438 | 0.142895578 | 1.07E-13 | 1.75E-12 |
| Camk1d | 872.5918136 | -1.613423265 | 0.223742189 | 1.08E-13 | 1.76E-12 |
| Aqp4 | 36.54572955 | -3.834567953 | 0.544881772 | 1.43E-13 | 2.31E-12 |
| Bco2 | 233.3934605 | -1.290899421 | 0.178959457 | 1.54E-13 | 2.46E-12 |
| Slc17a3 | 598.3860062 | -1.366358091 | 0.189321331 | 1.56E-13 | 2.50E-12 |
| Magt1 | 1463.989814 | -1.21847158 | 0.169065037 | 1.81E-13 | 2.88E-12 |
| Gm16551 | 158.3694224 | -2.223388884 | 0.314185608 | 2.15E-13 | 3.41E-12 |
| Pank1 | 2084.751402 | -1.278228916 | 0.178899004 | 2.62E-13 | 4.12E-12 |
| Etnppl | 1624.763176 | -1.678792451 | 0.237720976 | 2.85E-13 | 4.45E-12 |
| Gm9826 | 59.4033759 | -2.43846031 | 0.349566323 | 3.16E-13 | 4.91E-12 |
| Gstt3 | 411.7071088 | -1.3222338 | 0.18593062 | 3.19E-13 | 4.94E-12 |

| | | | | | |
|---|---|---|---|---|---|
| Acsm1 | 4203.972657 | -1.434672031 | 0.203120025 | 3.87E-13 | 5.94E-12 |
| Cadm4 | 235.0068656 | -2.465215915 | 0.354357922 | 4.04E-13 | 6.19E-12 |
| Lipc | 1259.221237 | -1.031292917 | 0.144537595 | 4.25E-13 | 6.51E-12 |
| Blm | 139.4126154 | -1.389109789 | 0.196825822 | 4.38E-13 | 6.69E-12 |
| Afm | 2380.799946 | -1.993233325 | 0.28504529 | 4.43E-13 | 6.76E-12 |
| Fn3krp | 226.3381668 | -1.134411511 | 0.159850714 | 4.56E-13 | 6.94E-12 |
| Sult2a7 | 23.50816231 | -9.956504639 | 2.951813459 | 4.64E-13 | 7.05E-12 |
| AI182371 | 3605.079203 | -1.50079906 | 0.213458275 | 4.71E-13 | 7.14E-12 |
| Gnmt | 12382.48397 | -1.69947121 | 0.243716648 | 5.03E-13 | 7.62E-12 |
| Tln2 | 157.4549379 | -1.198013778 | 0.16964355 | 5.32E-13 | 8.03E-12 |
| Cplane1 | 772.9753829 | -1.27780983 | 0.181511511 | 5.79E-13 | 8.71E-12 |
| Tm7sf2 | 1431.894175 | -1.323814902 | 0.188713959 | 6.30E-13 | 9.45E-12 |
| Spout1 | 326.1234874 | -1.103332324 | 0.156548304 | 6.74E-13 | 1.01E-11 |
| Magix | 145.2263832 | -1.509991728 | 0.216469012 | 6.92E-13 | 1.03E-11 |
| Adgrf1 | 130.9530229 | -3.54573212 | 0.519724717 | 7.60E-13 | 1.13E-11 |
| Rdh16 | 461.4338517 | -2.070121064 | 0.302301229 | 8.75E-13 | 1.30E-11 |
| Mfsd4b1 | 435.8614984 | -1.102750951 | 0.157668988 | 9.86E-13 | 1.45E-11 |
| Acot11 | 103.345663 | -2.331837656 | 0.341108384 | 9.95E-13 | 1.47E-11 |
| Aldh2 | 26489.0011 | -1.183834361 | 0.169155191 | 1.06E-12 | 1.57E-11 |
| Tymp | 264.3628247 | -1.897303044 | 0.278472344 | 1.43E-12 | 2.09E-11 |
| Gm45425 | 28.50175919 | -6.084059487 | 0.889268324 | 1.55E-12 | 2.26E-11 |
| Mapk15 | 381.841014 | -1.625579215 | 0.238339337 | 1.81E-12 | 2.62E-11 |
| Mreg | 392.8654681 | -1.740121022 | 0.255768953 | 1.93E-12 | 2.79E-11 |
| G6pc | 2487.090987 | -1.878114966 | 0.278221714 | 2.40E-12 | 3.44E-11 |
| Celf2 | 394.3433022 | -2.217946188 | 0.330138913 | 2.45E-12 | 3.51E-11 |
| Slc25a21 | 125.0229346 | -1.841809788 | 0.273731439 | 2.69E-12 | 3.84E-11 |
| Gm35696 | 32.45528046 | -2.732339341 | 0.409110354 | 2.71E-12 | 3.86E-11 |
| Ifit1 | 179.401872 | -1.673887975 | 0.248259951 | 2.96E-12 | 4.21E-11 |
| Afmid | 436.4352751 | -1.419208296 | 0.209587158 | 3.11E-12 | 4.41E-11 |
| Ddit4 | 877.0347367 | -1.651004447 | 0.245006316 | 3.15E-12 | 4.47E-11 |
| Ugt2a3 | 2492.393957 | -1.797857339 | 0.26917212 | 3.76E-12 | 5.29E-11 |
| Zbtb5 | 239.0530269 | -1.182575621 | 0.174380449 | 4.09E-12 | 5.73E-11 |
| Ces1e | 711.5684012 | -1.619295417 | 0.242279398 | 4.29E-12 | 5.99E-11 |
| Gtse1 | 93.65492005 | -1.706034081 | 0.25537884 | 4.52E-12 | 6.29E-11 |
| Nxpe2 | 148.1812551 | -1.632884178 | 0.244224679 | 4.73E-12 | 6.59E-11 |
| Xk | 89.35518604 | -1.620381713 | 0.242830439 | 5.01E-12 | 6.93E-11 |
| Mup12 | 37.30375135 | -8.374178847 | 1.36717084 | 5.26E-12 | 7.27E-11 |
| Comt | 10115.36535 | -1.148553849 | 0.169900741 | 5.33E-12 | 7.36E-11 |
| Ccnf | 268.1785076 | -1.631114052 | 0.245207684 | 5.60E-12 | 7.72E-11 |
| Nlrp12 | 205.4374206 | -3.889419547 | 0.601120168 | 5.85E-12 | 8.03E-11 |
| Ndufaf1 | 380.0744169 | -1.105724106 | 0.164403068 | 6.57E-12 | 9.00E-11 |
| Dock4 | 707.5407348 | -1.02509996 | 0.152268213 | 6.77E-12 | 9.26E-11 |
| Nnmt | 860.953606 | -2.044747924 | 0.311153199 | 6.78E-12 | 9.27E-11 |

| | | | | | |
|---|---|---|---|---|---|
| Arl4a | 402.0568203 | -1.113983764 | 0.16599853 | 7.03E-12 | 9.59E-11 |
| Gulo | 4146.630256 | -1.369150228 | 0.205715636 | 7.57E-12 | 1.03E-10 |
| Mcm10 | 407.1965466 | -2.189715637 | 0.335342724 | 7.87E-12 | 1.07E-10 |
| Tspyl4 | 70.44859373 | -2.996293197 | 0.45936199 | 8.06E-12 | 1.09E-10 |
| Tfb1m | 142.9438176 | -1.283150064 | 0.193649733 | 9.69E-12 | 1.30E-10 |
| Neb | 369.7871637 | -1.530422731 | 0.23309829 | 1.10E-11 | 1.47E-10 |
| Cth | 4751.305475 | -1.680340982 | 0.257599257 | 1.11E-11 | 1.48E-10 |
| Cyp3a25 | 4180.551106 | -1.621648835 | 0.247660521 | 1.16E-11 | 1.54E-10 |
| Arhgap26 | 307.9492233 | -1.353005252 | 0.20591908 | 1.33E-11 | 1.76E-10 |
| 2310001H17Rik | 95.95978737 | -1.761449345 | 0.270727985 | 1.34E-11 | 1.76E-10 |
| Cyp27a1 | 3585.908062 | -1.156591098 | 0.175607241 | 1.53E-11 | 2.01E-10 |
| Eda2r | 190.2811247 | -2.04596097 | 0.317565229 | 1.62E-11 | 2.11E-10 |
| Rhobtb3 | 395.3741534 | -1.03600393 | 0.156995358 | 1.82E-11 | 2.37E-10 |
| Abca8b | 1247.531189 | -1.05266084 | 0.160094855 | 2.04E-11 | 2.65E-10 |
| Srgap3 | 49.21790938 | -6.926701216 | 1.011940735 | 2.06E-11 | 2.66E-10 |
| Tanc2 | 226.7021055 | -1.304012968 | 0.200532408 | 2.16E-11 | 2.78E-10 |
| Galt | 608.0470576 | -1.012779624 | 0.15500227 | 2.70E-11 | 3.45E-10 |
| Gm45941 | 27.03596676 | -7.902243389 | 1.336212488 | 2.84E-11 | 3.62E-10 |
| Serpina3n | 6404.26128 | -1.525045953 | 0.237006247 | 2.87E-11 | 3.65E-10 |
| Igfbp2 | 2614.063083 | -1.439906254 | 0.224686878 | 3.34E-11 | 4.23E-10 |
| Mfsd8 | 188.7976271 | -1.017509355 | 0.156881782 | 3.64E-11 | 4.58E-10 |
| Npr2 | 507.7869207 | -1.256461047 | 0.195578615 | 3.95E-11 | 4.95E-10 |
| Gpcpd1 | 1025.645231 | -1.75502162 | 0.278506742 | 4.95E-11 | 6.12E-10 |
| Dapk1 | 2039.317286 | -1.068430626 | 0.166663974 | 4.98E-11 | 6.16E-10 |
| Odf3b | 120.9933682 | -1.339665737 | 0.210217945 | 5.17E-11 | 6.38E-10 |
| Hyou1 | 3725.334432 | -1.427315633 | 0.22539155 | 5.98E-11 | 7.32E-10 |
| Ugt2b5 | 11161.03453 | -1.593729243 | 0.253628819 | 6.41E-11 | 7.82E-10 |
| Ces1f | 3502.330601 | -3.503923322 | 0.576301842 | 7.67E-11 | 9.28E-10 |
| Gmppb | 355.6339127 | -1.133160655 | 0.17864389 | 7.89E-11 | 9.53E-10 |
| Dmgdh | 4270.370996 | -1.459233393 | 0.232238645 | 8.02E-11 | 9.66E-10 |
| Gpam | 5231.991526 | -1.247570008 | 0.197750343 | 8.41E-11 | 1.01E-09 |
| Tmie | 290.5164944 | -1.4979633 | 0.239433204 | 8.61E-11 | 1.03E-09 |
| Rfx4 | 17.92621913 | -3.812527227 | 0.624740685 | 1.07E-10 | 1.27E-09 |
| Dct | 14.13264872 | -9.060130993 | 2.867492884 | 1.17E-10 | 1.38E-09 |
| 4732465J04Rik | 135.2761788 | -1.485013187 | 0.239055635 | 1.27E-10 | 1.50E-09 |
| Agbl3 | 127.0944135 | -1.359954194 | 0.218340225 | 1.32E-10 | 1.56E-09 |
| Pxmp4 | 1264.234575 | -1.006066441 | 0.159752264 | 1.35E-10 | 1.59E-09 |
| Asl | 5484.97917 | -1.055247794 | 0.168842741 | 1.44E-10 | 1.69E-09 |
| Hao1 | 2381.38118 | -1.893178006 | 0.310653311 | 1.56E-10 | 1.83E-09 |
| Maob | 2149.151945 | -1.217205304 | 0.196259799 | 1.56E-10 | 1.83E-09 |
| Ston2 | 535.3285869 | -1.203681906 | 0.193743996 | 1.57E-10 | 1.83E-09 |
| Lactb2 | 3890.475783 | -1.089285886 | 0.174520699 | 1.59E-10 | 1.85E-09 |
| Slc25a45 | 295.066893 | -1.32902454 | 0.214947266 | 1.73E-10 | 2.02E-09 |

| | | | | | |
|---|---|---|---|---|---|
| Abca8a | 1399.104272 | -2.021831333 | 0.333243805 | 1.82E-10 | 2.11E-09 |
| Cdcp1 | 1036.676466 | -1.178944995 | 0.190651532 | 2.05E-10 | 2.37E-09 |
| Gm11844 | 44.05439073 | -2.016056751 | 0.334261734 | 2.20E-10 | 2.52E-09 |
| Apoa2 | 29949.05955 | -1.291313724 | 0.215692367 | 2.31E-10 | 2.64E-09 |
| Hdhd3 | 253.4069346 | -1.031330394 | 0.16683294 | 2.37E-10 | 2.70E-09 |
| Ahcyl2 | 561.6757661 | -1.06874805 | 0.17310701 | 2.43E-10 | 2.76E-09 |
| Acmsd | 296.9193473 | -1.559356915 | 0.256503658 | 2.57E-10 | 2.91E-09 |
| Ptk6 | 28.12952245 | -3.932598463 | 0.668502717 | 2.59E-10 | 2.93E-09 |
| Mtarc1 | 4417.622663 | -1.106747907 | 0.179645401 | 2.63E-10 | 2.98E-09 |
| Leap2 | 123.6363937 | -2.726980462 | 0.460354727 | 2.65E-10 | 3.00E-09 |
| Plaat3 | 217.2705145 | -1.494136206 | 0.245818095 | 2.82E-10 | 3.17E-09 |
| Pithd1 | 340.2520987 | -1.000470116 | 0.162447374 | 3.08E-10 | 3.45E-09 |
| Capn8 | 21.36772011 | -4.088464325 | 0.693902246 | 3.13E-10 | 3.51E-09 |
| Cyp4f15 | 1617.125215 | -1.223865134 | 0.20060506 | 3.15E-10 | 3.52E-09 |
| Atp5g3 | 3736.266072 | -1.034609517 | 0.168610886 | 3.20E-10 | 3.57E-09 |
| Cfhr2 | 7372.005222 | -1.125502293 | 0.184157306 | 3.43E-10 | 3.82E-09 |
| Gm28438 | 95.97393006 | -2.726438533 | 0.46464712 | 3.59E-10 | 3.97E-09 |
| Cela1 | 177.1386406 | -1.356107854 | 0.223862035 | 3.63E-10 | 4.01E-09 |
| Ugt2b1 | 5937.994386 | -3.045600475 | 0.521443887 | 3.71E-10 | 4.09E-09 |
| Gm37795 | 57.84462543 | -2.03906616 | 0.341852069 | 3.79E-10 | 4.18E-09 |
| mt-Tn | 102.0591981 | -1.339532619 | 0.221319058 | 3.82E-10 | 4.20E-09 |
| Tbkbp1 | 141.1041512 | -1.263836683 | 0.208539451 | 4.00E-10 | 4.39E-09 |
| Necab1 | 535.5719681 | -1.062930651 | 0.17406169 | 4.13E-10 | 4.53E-09 |
| Slc13a2 | 20.44750741 | -5.713360829 | 0.929515461 | 4.17E-10 | 4.56E-09 |
| Sphk2 | 1001.934489 | -1.205584096 | 0.199031688 | 4.44E-10 | 4.84E-09 |
| BC049987 | 73.19393298 | -2.333943196 | 0.398424011 | 4.81E-10 | 5.22E-09 |
| Ccdc30 | 146.632766 | -1.680709699 | 0.282203212 | 4.85E-10 | 5.25E-09 |
| Ropn1l | 119.2868843 | -1.14394496 | 0.188892214 | 4.86E-10 | 5.26E-09 |
| Ube2u | 57.71230701 | -1.766010533 | 0.297367868 | 4.97E-10 | 5.37E-09 |
| D3Ertd751e | 181.7250643 | -1.126427281 | 0.186958483 | 6.20E-10 | 6.65E-09 |
| 4732419C18Rik | 63.37222025 | -1.469915351 | 0.247772708 | 6.85E-10 | 7.31E-09 |
| Serpina12 | 440.3608179 | -1.377441427 | 0.231610957 | 6.96E-10 | 7.42E-09 |
| Ifit3b | 30.71031413 | -2.220563342 | 0.380433645 | 7.48E-10 | 7.94E-09 |
| Isoc2a | 1097.025947 | -1.135151576 | 0.189735513 | 7.61E-10 | 8.07E-09 |
| Apoo | 266.4363373 | -1.01118389 | 0.16842001 | 8.01E-10 | 8.48E-09 |
| Gm16548 | 29.03563962 | -2.18789378 | 0.377737798 | 8.66E-10 | 9.15E-09 |
| Lamb3 | 921.3770423 | -1.76319929 | 0.300415071 | 9.12E-10 | 9.59E-09 |
| Gna14 | 101.152346 | -8.074602241 | 2.703678642 | 9.80E-10 | 1.03E-08 |
| Gm48633 | 155.1112973 | -1.289445412 | 0.21845531 | 1.01E-09 | 1.05E-08 |
| Tbc1d30 | 195.1130102 | -1.538811047 | 0.262868922 | 1.03E-09 | 1.08E-08 |
| Cfh | 18497.17179 | -1.685737067 | 0.289534381 | 1.06E-09 | 1.11E-08 |
| N4bp2 | 626.9326672 | -1.089411876 | 0.183432126 | 1.07E-09 | 1.12E-08 |
| Npas2 | 70.79666511 | -4.119782264 | 0.727317867 | 1.10E-09 | 1.15E-08 |

| | | | | | |
|---|---|---|---|---|---|
| Madd | 487.8038817 | -1.076284577 | 0.181670876 | 1.19E-09 | 1.24E-08 |
| Acss2 | 2949.095796 | -1.714870141 | 0.296560895 | 1.23E-09 | 1.27E-08 |
| Rdh7 | 9274.605396 | -1.130826092 | 0.192891585 | 1.37E-09 | 1.41E-08 |
| Asic5 | 38.89485239 | -2.171238563 | 0.379476294 | 1.43E-09 | 1.48E-08 |
| Gls2 | 4573.752992 | -1.497335333 | 0.259117028 | 1.68E-09 | 1.72E-08 |
| Hapln4 | 178.2555074 | -1.399628434 | 0.241823794 | 1.70E-09 | 1.73E-08 |
| Esrrg | 11.23201636 | -8.421875994 | 2.797439887 | 1.84E-09 | 1.87E-08 |
| Gm4737 | 51.83017482 | -1.933099547 | 0.33996069 | 1.87E-09 | 1.89E-08 |
| Syt11 | 182.827561 | -1.327836027 | 0.229238752 | 1.97E-09 | 1.98E-08 |
| 5730414N17Rik | 42.67015591 | -2.019333902 | 0.356506184 | 2.04E-09 | 2.04E-08 |
| Otc | 4505.54963 | -1.544743242 | 0.269776574 | 2.08E-09 | 2.08E-08 |
| Cpeb2 | 1145.829352 | -1.289177194 | 0.223448905 | 2.18E-09 | 2.17E-08 |
| 9130409I23Rik | 192.8731022 | -1.366192475 | 0.237882991 | 2.35E-09 | 2.33E-08 |
| Denn2b | 1237.020607 | -1.354612141 | 0.235646135 | 2.44E-09 | 2.42E-08 |
| Nat8f2 | 11488.31915 | -1.463874985 | 0.2566412 | 2.60E-09 | 2.57E-08 |
| Gm13456 | 219.0911845 | -1.141414425 | 0.197652787 | 2.62E-09 | 2.59E-08 |
| St3gal3 | 874.0343593 | -1.048226922 | 0.18070517 | 2.62E-09 | 2.59E-08 |
| Amer2 | 58.93190936 | -4.954170543 | 0.8373534 | 2.83E-09 | 2.78E-08 |
| Igtp | 410.365261 | -1.009129237 | 0.174285094 | 2.91E-09 | 2.85E-08 |
| Impg2 | 38.99951935 | -1.969105716 | 0.35069631 | 2.92E-09 | 2.86E-08 |
| Dock8 | 727.0900506 | -1.053611992 | 0.18200894 | 2.96E-09 | 2.89E-08 |
| Tmem8b | 156.0274911 | -1.770876184 | 0.313976329 | 2.99E-09 | 2.92E-08 |
| Cspg5 | 18.946972 | -3.087276554 | 0.556753556 | 3.11E-09 | 3.03E-08 |
| Vmn1r90 | 13.82799156 | -8.228141244 | 2.77154793 | 3.19E-09 | 3.11E-08 |
| Asap2 | 484.8955606 | -1.426041717 | 0.251683117 | 3.47E-09 | 3.37E-08 |
| Aass | 3323.141373 | -1.145107886 | 0.2002433 | 3.56E-09 | 3.45E-08 |
| Nfil3 | 345.3377128 | -1.659791179 | 0.296170896 | 3.85E-09 | 3.72E-08 |
| Gm36264 | 14.7572191 | -2.920302065 | 0.534842699 | 4.74E-09 | 4.53E-08 |
| Fut8 | 152.5306864 | -1.52489724 | 0.273780148 | 5.79E-09 | 5.48E-08 |
| Srms | 18.07701278 | -4.350919648 | 0.790067088 | 5.86E-09 | 5.54E-08 |
| Sc5d | 6522.735873 | -1.192082158 | 0.21252338 | 6.35E-09 | 5.97E-08 |
| Gm8883 | 156.9433376 | -1.496838915 | 0.271763972 | 6.49E-09 | 6.09E-08 |
| Hgd | 5767.565109 | -1.443967608 | 0.260643051 | 6.86E-09 | 6.42E-08 |
| Atp8b4 | 17.3531573 | -2.850738248 | 0.528514228 | 6.91E-09 | 6.46E-08 |
| Lipe | 230.9963135 | -1.122614009 | 0.199947309 | 7.04E-09 | 6.57E-08 |
| Cyp3a11 | 34493.26965 | -1.422145996 | 0.256789472 | 7.10E-09 | 6.62E-08 |
| Ss18l2 | 223.317077 | -1.161608131 | 0.207513238 | 7.22E-09 | 6.73E-08 |
| Dio1 | 416.1815905 | -1.484782037 | 0.26989159 | 8.32E-09 | 7.71E-08 |
| Isoc2b | 177.3147908 | -1.246569411 | 0.22468525 | 8.86E-09 | 8.18E-08 |
| Pyurf | 268.2008094 | -1.008737571 | 0.180571719 | 9.71E-09 | 8.92E-08 |
| Tmem14a | 106.0498655 | -1.051476831 | 0.188886868 | 9.85E-09 | 9.04E-08 |
| Slc17a8 | 109.4741187 | -1.846261529 | 0.341984154 | 1.02E-08 | 9.32E-08 |
| Sgk2 | 693.6257907 | -1.27059685 | 0.230304532 | 1.09E-08 | 9.97E-08 |

| | | | | | |
|---|---|---|---|---|---|
| Pltp | 76.11284702 | -2.758702513 | 0.522817179 | 1.10E-08 | 1.00E-07 |
| Gbp6 | 97.31930268 | -1.517105106 | 0.279293458 | 1.25E-08 | 1.13E-07 |
| Spns2 | 2975.431914 | -1.042041173 | 0.188895566 | 1.29E-08 | 1.16E-07 |
| Svip | 306.4687993 | -1.063770727 | 0.192764536 | 1.32E-08 | 1.19E-07 |
| Prex1 | 47.6812679 | -2.639228167 | 0.500541275 | 1.47E-08 | 1.32E-07 |
| Fmc1 | 165.5128956 | -1.120857648 | 0.204516265 | 1.51E-08 | 1.34E-07 |
| Ugt1a9 | 671.7909745 | -1.351691045 | 0.250469244 | 1.51E-08 | 1.35E-07 |
| Cyp2d37-ps | 575.130604 | -1.40373046 | 0.259652766 | 1.56E-08 | 1.38E-07 |
| Ces3b | 2566.255037 | -4.84463329 | 0.951094314 | 1.57E-08 | 1.39E-07 |
| Arntl | 192.6849049 | -3.287463827 | 0.638549238 | 1.76E-08 | 1.56E-07 |
| Ugt2b38 | 48.51642462 | -3.62182585 | 0.705027926 | 1.77E-08 | 1.56E-07 |
| Hsd3b3 | 1558.612985 | -1.657230719 | 0.310738552 | 1.78E-08 | 1.57E-07 |
| Slc27a2 | 7255.737939 | -1.110278951 | 0.204065767 | 1.88E-08 | 1.66E-07 |
| Fgf1 | 737.3589299 | -1.558734802 | 0.292415021 | 1.95E-08 | 1.71E-07 |
| Aldh4a1 | 2507.573566 | -1.003584934 | 0.183757572 | 1.97E-08 | 1.73E-07 |
| Apol9b | 681.1258834 | -1.66424964 | 0.313601832 | 1.98E-08 | 1.74E-07 |
| Gm45871 | 110.4961469 | -1.199708549 | 0.221605075 | 2.01E-08 | 1.76E-07 |
| Zfp951 | 62.23078478 | -1.652736036 | 0.310980701 | 2.16E-08 | 1.89E-07 |
| Abca9 | 57.00469553 | -1.862579224 | 0.354144353 | 2.25E-08 | 1.96E-07 |
| Hsbp1l1 | 17.9627834 | -2.338499316 | 0.449203209 | 2.42E-08 | 2.09E-07 |
| Bbc3 | 330.5124915 | -1.946413906 | 0.372752075 | 2.45E-08 | 2.12E-07 |
| Trmt9b | 78.38364621 | -1.987449728 | 0.381027866 | 2.52E-08 | 2.17E-07 |
| Gm5621 | 27.25329181 | -2.017616336 | 0.386980137 | 2.67E-08 | 2.30E-07 |
| Gm44243 | 31.37297085 | -2.266665225 | 0.437968404 | 2.80E-08 | 2.40E-07 |
| Med12l | 94.44285171 | -2.052276416 | 0.395500979 | 2.91E-08 | 2.49E-07 |
| Tmigd1 | 17.10750995 | -3.320607935 | 0.649632168 | 2.92E-08 | 2.49E-07 |
| mt-Te | 182.7793095 | -1.470773425 | 0.27943121 | 3.17E-08 | 2.69E-07 |
| Tert | 316.1095245 | -1.213634843 | 0.22883045 | 3.51E-08 | 2.96E-07 |
| Glo1-ps | 22.6710828 | -2.134269523 | 0.416152523 | 3.69E-08 | 3.10E-07 |
| Pex11g | 294.4908088 | -1.12141374 | 0.210988525 | 3.74E-08 | 3.14E-07 |
| Ddc | 1020.346508 | -1.134432505 | 0.213854294 | 3.87E-08 | 3.24E-07 |
| Gm5529 | 26.35256226 | -2.115527092 | 0.413641034 | 4.01E-08 | 3.36E-07 |
| mt-Nd3 | 350.1705923 | -1.499762742 | 0.288178885 | 4.13E-08 | 3.45E-07 |
| Ass1 | 40142.72455 | -1.401246703 | 0.268652216 | 4.33E-08 | 3.60E-07 |
| Apon | 3468.369066 | -1.125799519 | 0.212811544 | 4.32E-08 | 3.60E-07 |
| 9230114K14Rik | 40.92338174 | -1.79991678 | 0.349830449 | 4.34E-08 | 3.61E-07 |
| 1300017J02Rik | 6513.128722 | -1.248532776 | 0.238936871 | 4.82E-08 | 3.98E-07 |
| Fam151b | 50.57162547 | -1.469475356 | 0.283381726 | 4.94E-08 | 4.07E-07 |
| Gas1 | 107.4944131 | -1.670518573 | 0.325963618 | 5.28E-08 | 4.34E-07 |
| Gm50136 | 18.85028508 | -4.266012526 | 0.861203061 | 5.55E-08 | 4.55E-07 |
| Echdc1 | 299.6291448 | -1.116756489 | 0.213008911 | 5.61E-08 | 4.60E-07 |
| Fbp1 | 9434.908133 | -1.250957113 | 0.240378078 | 5.66E-08 | 4.63E-07 |
| Slco2b1 | 1372.102266 | -1.380235322 | 0.269630573 | 5.69E-08 | 4.66E-07 |

| | | | | | |
|---|---|---|---|---|---|
| Cmah | 1943.234371 | -2.226498203 | 0.444788463 | 5.90E-08 | 4.82E-07 |
| Gm5524 | 115.7659489 | -1.374618085 | 0.267199585 | 6.44E-08 | 5.20E-07 |
| Inca1 | 167.934322 | -1.188738044 | 0.2294532 | 7.08E-08 | 5.68E-07 |
| Slco1b2 | 8918.895204 | -1.366999714 | 0.266710746 | 7.33E-08 | 5.87E-07 |
| Pde7b | 33.4619523 | -2.332970473 | 0.470705071 | 7.89E-08 | 6.30E-07 |
| 1700007K13Rik | 13.80336685 | -7.291116065 | 2.646121654 | 8.47E-08 | 6.75E-07 |
| Gm45792 | 25.65472788 | -2.184773802 | 0.442614191 | 9.34E-08 | 7.42E-07 |
| Slc43a1 | 77.66526007 | -2.954706704 | 0.610170002 | 9.62E-08 | 7.61E-07 |
| Xirp1 | 17.77658105 | -2.983696227 | 0.613354013 | 9.63E-08 | 7.62E-07 |
| 5033403H07Rik | 135.1495714 | -1.204424024 | 0.235304149 | 9.83E-08 | 7.76E-07 |
| Haao | 3495.218388 | -1.216669071 | 0.238336362 | 9.99E-08 | 7.88E-07 |
| Gm36041 | 64.21392594 | -1.896534318 | 0.3859747 | 1.26E-07 | 9.78E-07 |
| Plk2 | 1956.833191 | -1.277811534 | 0.253047099 | 1.26E-07 | 9.84E-07 |
| Gm26684 | 69.93434537 | -1.272635156 | 0.251938534 | 1.28E-07 | 9.94E-07 |
| Slc23a1 | 875.5246885 | -1.025914668 | 0.20138954 | 1.42E-07 | 1.10E-06 |
| Ugt2b36 | 6904.458939 | -1.406881605 | 0.282092654 | 1.45E-07 | 1.12E-06 |
| Gm32461 | 65.38232439 | -1.506897801 | 0.303978605 | 1.48E-07 | 1.14E-06 |
| Slc22a30 | 1329.295779 | -2.341307611 | 0.488692677 | 1.60E-07 | 1.23E-06 |
| Gm6170 | 35.06236282 | -1.717044001 | 0.348932888 | 1.62E-07 | 1.25E-06 |
| Ces1d | 4846.406319 | -1.26666964 | 0.253557755 | 1.66E-07 | 1.27E-06 |
| Gm30784 | 27.53769173 | -2.609795115 | 0.549304341 | 1.71E-07 | 1.31E-06 |
| Gm15590 | 84.10377704 | -1.314453287 | 0.26420693 | 1.74E-07 | 1.33E-06 |
| Lgalsl | 539.7969122 | -1.209761524 | 0.241944347 | 1.79E-07 | 1.37E-06 |
| Fam81a | 308.2898849 | -1.062394098 | 0.210115983 | 1.80E-07 | 1.37E-06 |
| Gm40770 | 29.03301413 | -1.84747738 | 0.380683929 | 1.86E-07 | 1.42E-06 |
| Apol7a | 1459.177259 | -1.586641378 | 0.324853109 | 2.02E-07 | 1.53E-06 |
| Gm49463 | 49.70881723 | -1.820356233 | 0.378472126 | 2.37E-07 | 1.77E-06 |
| Zfp760 | 168.7595934 | -1.039221805 | 0.208093571 | 2.40E-07 | 1.79E-06 |
| Gvin1 | 26.21680248 | -2.902927533 | 0.623759124 | 2.48E-07 | 1.84E-06 |
| Htra4 | 10.31369696 | -4.5010984 | 0.939741479 | 2.52E-07 | 1.87E-06 |
| Acaa1b | 4212.932469 | -1.129261474 | 0.2303716 | 2.61E-07 | 1.94E-06 |
| Cyp2f2 | 8750.464772 | -2.044032122 | 0.432446979 | 2.70E-07 | 2.00E-06 |
| 0610031O16Rik | 118.6314505 | -1.395647364 | 0.28830426 | 3.10E-07 | 2.27E-06 |
| Ftcd | 5706.832529 | -1.139007029 | 0.232675539 | 3.36E-07 | 2.45E-06 |
| Polr3g | 125.3474313 | -1.117757758 | 0.228167129 | 3.39E-07 | 2.47E-06 |
| Irf5 | 157.2545146 | -1.771578887 | 0.373620156 | 3.48E-07 | 2.53E-06 |
| Gm35164 | 11.3277361 | -6.419631631 | 1.467072068 | 3.56E-07 | 2.58E-06 |
| B4galnt3 | 33.22448072 | -2.126087534 | 0.456367127 | 3.92E-07 | 2.83E-06 |
| Ctcflos | 51.00435398 | -1.820590451 | 0.39038017 | 4.50E-07 | 3.22E-06 |
| Rnf144a | 54.43551313 | -2.07343435 | 0.447535614 | 4.54E-07 | 3.24E-06 |
| Fam110a | 206.2262549 | -1.011498244 | 0.207537962 | 4.60E-07 | 3.28E-06 |
| Gm12909 | 22.97243683 | -3.316328359 | 0.737162213 | 4.84E-07 | 3.45E-06 |
| Cyp4a31 | 93.13873315 | -1.248569911 | 0.260635616 | 4.93E-07 | 3.50E-06 |

| | | | | | |
|---|---|---|---|---|---|
| C8b | 3746.507413 | -1.852659987 | 0.398559426 | 4.94E-07 | 3.51E-06 |
| Sp4 | 173.6313221 | -1.185915764 | 0.247095976 | 5.10E-07 | 3.62E-06 |
| Selenbp2 | 4616.839794 | -2.49718438 | 0.556492227 | 6.22E-07 | 4.34E-06 |
| Gm49431 | 29.19339011 | -1.930807124 | 0.420901566 | 6.23E-07 | 4.34E-06 |
| Tdo2 | 14666.53154 | -1.851407125 | 0.405290267 | 6.87E-07 | 4.76E-06 |
| F9 | 2549.301488 | -1.015947111 | 0.211897321 | 6.90E-07 | 4.77E-06 |
| Egfr | 6070.606277 | -2.189205139 | 0.485656438 | 7.00E-07 | 4.83E-06 |
| BC023105 | 50.65969244 | -2.025986993 | 0.4459897 | 7.03E-07 | 4.84E-06 |
| Acss3 | 235.27738 | -1.696303887 | 0.369997503 | 7.71E-07 | 5.28E-06 |
| Gpr155 | 680.0996554 | -1.032733407 | 0.217563034 | 8.19E-07 | 5.58E-06 |
| Hmgcs2 | 18355.00074 | -1.085027995 | 0.230294122 | 8.95E-07 | 6.06E-06 |
| Slc25a35 | 21.6861406 | -3.764753117 | 0.845727995 | 9.80E-07 | 6.58E-06 |
| Cyp2d26 | 7786.721027 | -1.031610584 | 0.217141402 | 9.88E-07 | 6.63E-06 |
| Depp1 | 2367.456344 | -1.470427617 | 0.321716999 | 1.06E-06 | 7.11E-06 |
| Nat8f5 | 79.78016015 | -2.840544356 | 0.65634373 | 1.10E-06 | 7.32E-06 |
| St6galnac4 | 51.12053296 | -2.668429318 | 0.606145291 | 1.19E-06 | 7.89E-06 |
| Slc3a1 | 607.9477193 | -1.513876524 | 0.334112941 | 1.20E-06 | 7.94E-06 |
| Rai2 | 45.47658153 | -1.825038352 | 0.40712445 | 1.25E-06 | 8.21E-06 |
| Lrit1 | 117.4120306 | -1.065532447 | 0.229938677 | 1.33E-06 | 8.73E-06 |
| Mup22 | 187.6972937 | -8.599717388 | 2.115979779 | 1.38E-06 | 9.00E-06 |
| Nlrp6 | 1473.457438 | -1.212310835 | 0.264737618 | 1.39E-06 | 9.08E-06 |
| Gm11454 | 13.77466195 | -3.63697022 | 0.846983616 | 1.42E-06 | 9.30E-06 |
| Mup-ps20 | 11.21091451 | -6.349180254 | 1.552436821 | 1.43E-06 | 9.31E-06 |
| Prodh2 | 1580.196679 | -1.358790823 | 0.300703053 | 1.53E-06 | 9.93E-06 |
| Pck1 | 20769.08758 | -1.456535821 | 0.324854766 | 1.54E-06 | 1.00E-05 |
| Celsr1 | 1462.370175 | -1.368046041 | 0.303605016 | 1.57E-06 | 1.02E-05 |
| Apobec1 | 588.6962835 | -1.034948375 | 0.224629384 | 1.64E-06 | 1.06E-05 |
| Ptgs1 | 42.54643769 | -2.098311054 | 0.47506441 | 1.64E-06 | 1.06E-05 |
| Gbp7 | 217.4561944 | -1.100346375 | 0.240407935 | 1.68E-06 | 1.08E-05 |
| Hnf4aos | 28.66882509 | -1.955908115 | 0.44905577 | 1.70E-06 | 1.09E-05 |
| Pstpip2 | 313.0269103 | -1.1853394 | 0.261841709 | 1.89E-06 | 1.20E-05 |
| Erbb3 | 2855.224645 | -1.052026882 | 0.230435971 | 1.89E-06 | 1.21E-05 |
| Gm30906 | 11.38690708 | -3.986379824 | 0.950902249 | 1.91E-06 | 1.22E-05 |
| Efna3 | 9.071597771 | -3.079098566 | 0.71517616 | 1.99E-06 | 1.26E-05 |
| Gm32281 | 10.19832376 | -3.533422965 | 0.834979495 | 2.04E-06 | 1.29E-05 |
| Plg | 23149.35435 | -1.080329168 | 0.238063004 | 2.06E-06 | 1.30E-05 |
| Nlrp5-ps | 38.30392892 | -6.225523891 | 2.505060972 | 2.16E-06 | 1.36E-05 |
| Eef2-ps2 | 35.16473167 | -1.616840508 | 0.368761514 | 2.27E-06 | 1.42E-05 |
| mt-Ty | 49.72062794 | -1.856784131 | 0.430653568 | 2.31E-06 | 1.45E-05 |
| Ddit4l | 493.8019112 | -1.043590974 | 0.23119546 | 2.45E-06 | 1.53E-05 |
| Lyrm7 | 54.67606616 | -1.304310198 | 0.294396762 | 2.57E-06 | 1.59E-05 |
| Sh2d3c | 103.3775141 | -1.365931054 | 0.311335765 | 2.91E-06 | 1.79E-05 |
| Gm6767 | 26.18073403 | -1.547837844 | 0.357309209 | 3.09E-06 | 1.89E-05 |

| | | | | | |
|---|---|---|---|---|---|
| BC024139 | 49.35503527 | -1.349122534 | 0.308800184 | 3.21E-06 | 1.96E-05 |
| Gm15417 | 64.00783042 | -1.130855384 | 0.255239046 | 3.22E-06 | 1.97E-05 |
| Serpina4-ps1 | 160.3037644 | -5.18320734 | 1.334156839 | 3.25E-06 | 1.98E-05 |
| Ugt3a2 | 2728.651214 | -2.644524496 | 0.648995842 | 3.40E-06 | 2.06E-05 |
| Gm5869 | 38.35883434 | -1.421770124 | 0.327078807 | 3.43E-06 | 2.08E-05 |
| Cphx1 | 28.55057676 | -1.626864699 | 0.379637323 | 3.48E-06 | 2.11E-05 |
| Izumo4 | 129.1480145 | -1.131606534 | 0.256499397 | 3.52E-06 | 2.13E-05 |
| Zfp125 | 138.7018529 | -1.300241418 | 0.298346202 | 3.66E-06 | 2.21E-05 |
| B3galt1 | 208.2230636 | -1.808404708 | 0.431212665 | 3.72E-06 | 2.24E-05 |
| Glt28d2 | 80.44252821 | -1.11524182 | 0.253346207 | 3.74E-06 | 2.26E-05 |
| Gm32063 | 32.64284349 | -1.826264139 | 0.433448919 | 3.76E-06 | 2.26E-05 |
| Gm38416 | 33.47614665 | -2.779991291 | 0.683078013 | 3.77E-06 | 2.27E-05 |
| Obox4-ps3 | 4.805751938 | -6.878589693 | 2.731500298 | 3.80E-06 | 2.29E-05 |
| Cyp2c68 | 1720.61075 | -1.216304998 | 0.280381631 | 4.11E-06 | 2.46E-05 |
| Adrb3 | 53.08530868 | -1.262207516 | 0.292810755 | 4.64E-06 | 2.74E-05 |
| Asb2 | 9.078914379 | -4.613268452 | 1.118452063 | 4.75E-06 | 2.80E-05 |
| Gm4956 | 58.92144837 | -5.52751578 | 1.470334446 | 5.29E-06 | 3.10E-05 |
| Clstn3 | 187.6726368 | -1.209400752 | 0.281618424 | 5.39E-06 | 3.15E-05 |
| Fam83f | 49.37687942 | -2.51886035 | 0.631877965 | 5.78E-06 | 3.37E-05 |
| Gm38102 | 20.10008434 | -1.720286468 | 0.413302425 | 5.85E-06 | 3.40E-05 |
| Olfr1033 | 137.8870896 | -1.066768742 | 0.247176983 | 5.90E-06 | 3.42E-05 |
| H2bc21 | 61.0141481 | -1.308659909 | 0.308379493 | 5.91E-06 | 3.43E-05 |
| Aldh8a1 | 3131.058374 | -1.132900889 | 0.26526503 | 6.48E-06 | 3.74E-05 |
| Cfhr4 | 3571.114705 | -1.016146516 | 0.235805877 | 6.74E-06 | 3.88E-05 |
| Akr1c6 | 10358.50332 | -1.290656955 | 0.306695289 | 6.80E-06 | 3.91E-05 |
| mt-Atp8 | 55.21354411 | -1.162791728 | 0.27391972 | 7.03E-06 | 4.03E-05 |
| Gm13340 | 1442.457308 | -1.14938518 | 0.271106427 | 7.29E-06 | 4.17E-05 |
| Mycn | 42.32018168 | -2.641287552 | 0.655370665 | 9.95E-06 | 5.53E-05 |
| Tpd52l1 | 102.418502 | -1.066724511 | 0.253706296 | 9.97E-06 | 5.54E-05 |
| Kyat3 | 1641.580225 | -1.339983834 | 0.326938973 | 1.01E-05 | 5.64E-05 |
| Ces2c | 554.1124305 | -1.058875236 | 0.25355058 | 1.09E-05 | 6.00E-05 |
| C8a | 8765.084562 | -1.22477511 | 0.297892686 | 1.14E-05 | 6.25E-05 |
| Bbs10 | 76.27645969 | -1.327253764 | 0.325019852 | 1.17E-05 | 6.38E-05 |
| 1700018L02Rik | 21.74536913 | -2.096097502 | 0.53770469 | 1.18E-05 | 6.43E-05 |
| Rps23rg1 | 36.73361085 | -1.957323021 | 0.501065788 | 1.19E-05 | 6.49E-05 |
| Amigo2 | 62.59737571 | -1.199921796 | 0.292057017 | 1.20E-05 | 6.53E-05 |
| Gstp-ps | 57.66899655 | -1.671885757 | 0.422793921 | 1.25E-05 | 6.83E-05 |
| Zdhhc14 | 56.71879898 | -1.176163936 | 0.286845033 | 1.33E-05 | 7.18E-05 |
| Prlr | 901.7496998 | -1.243718395 | 0.305902158 | 1.34E-05 | 7.23E-05 |
| Gpr12 | 24.5235708 | -1.514704922 | 0.381964458 | 1.45E-05 | 7.80E-05 |
| Bmp7 | 103.0541266 | -1.551016411 | 0.390102445 | 1.56E-05 | 8.29E-05 |
| Igkc | 162.6583523 | -2.865748096 | 0.776423999 | 1.66E-05 | 8.77E-05 |
| Gm34667 | 55.09359705 | -1.582803349 | 0.404483944 | 1.69E-05 | 8.93E-05 |

| | | | | | |
|---|---|---|---|---|---|
| Gm13522 | 39.39107079 | -2.035287636 | 0.537419783 | 1.79E-05 | 9.41E-05 |
| Hhipl1 | 34.58833831 | -1.898909864 | 0.494019932 | 1.87E-05 | 9.82E-05 |
| Wfdc21 | 2259.028558 | -1.267860098 | 0.321225802 | 1.97E-05 | 0.000102551 |
| Slc10a5 | 528.1282918 | -1.164491122 | 0.291042718 | 2.00E-05 | 0.000104024 |
| Gm2814 | 12.6303219 | -2.607079623 | 0.711918645 | 2.16E-05 | 0.000111919 |
| mt-Tw | 61.98009368 | -1.711027761 | 0.4490209 | 2.18E-05 | 0.000112933 |
| Gm15998 | 20.91827346 | -1.933723838 | 0.514531888 | 2.25E-05 | 0.000116429 |
| 1700061I17Rik | 11.01779263 | -2.463938007 | 0.656660488 | 2.39E-05 | 0.000123006 |
| Cep41 | 112.9931148 | -1.01413815 | 0.252721532 | 2.43E-05 | 0.000124779 |
| Clic5 | 114.532471 | -1.199456387 | 0.303549121 | 2.49E-05 | 0.000127618 |
| Odad3 | 71.34345341 | -1.225909118 | 0.311746682 | 2.52E-05 | 0.000129012 |
| Gm19522 | 18.02141761 | -2.992743086 | 0.842442565 | 2.56E-05 | 0.000130644 |
| Gm40438 | 21.24035268 | -1.852882587 | 0.497736165 | 2.75E-05 | 0.000139729 |
| 9330175E14Rik | 11.64435636 | -2.443181604 | 0.664743207 | 2.80E-05 | 0.000141905 |
| Cyp26b1 | 12.82435546 | -3.185088032 | 0.895973238 | 2.92E-05 | 0.000147359 |
| Gm8566 | 31.54511448 | -1.51385062 | 0.398565485 | 2.93E-05 | 0.00014764 |
| Abcb11 | 5414.293607 | -1.175782571 | 0.301886161 | 2.97E-05 | 0.000149513 |
| Gm44911 | 4.404663182 | -6.551785961 | 2.812762466 | 2.98E-05 | 0.000150208 |
| Osbp2 | 9.973837542 | -2.30271002 | 0.635331272 | 3.12E-05 | 0.000156507 |
| 1700028E10Rik | 48.09402262 | -1.187590187 | 0.306830546 | 3.18E-05 | 0.000159292 |
| AI480526 | 91.67138818 | -1.227957814 | 0.318027403 | 3.30E-05 | 0.000164483 |
| Rpgrip1 | 18.91572733 | -1.890433883 | 0.508881144 | 3.31E-05 | 0.000165083 |
| Tnik | 142.4327525 | -1.46512248 | 0.384679698 | 3.38E-05 | 0.00016827 |
| Camkk2 | 295.0702932 | -1.048967062 | 0.268379927 | 3.44E-05 | 0.000170984 |
| Fam47e | 149.9210212 | -1.4629926 | 0.389297371 | 3.57E-05 | 0.000177387 |
| Sspo | 17.79419403 | -1.722237206 | 0.465675981 | 3.85E-05 | 0.000189733 |
| Etfbkmt | 485.1689736 | -1.093868737 | 0.283858698 | 4.02E-05 | 0.000197414 |
| Gm26588 | 23.99824494 | -1.557358313 | 0.420663833 | 4.04E-05 | 0.000198049 |
| Tcp11l2 | 420.1748451 | -1.358331576 | 0.362007655 | 4.28E-05 | 0.000209084 |
| 1110020A21Rik | 17.00214535 | -1.597599128 | 0.434052987 | 4.44E-05 | 0.000216507 |
| Gm1600 | 27.10672324 | -2.147561337 | 0.610588402 | 4.47E-05 | 0.000217629 |
| Sned1 | 246.1928423 | -1.058572259 | 0.27541822 | 4.50E-05 | 0.000218954 |
| Fn3k | 113.2314708 | -1.231184234 | 0.326463771 | 4.68E-05 | 0.000227185 |
| Gbp3 | 66.83842882 | -1.38374793 | 0.375196271 | 5.30E-05 | 0.000254367 |
| Lin7a | 128.7885007 | -1.754826507 | 0.491136048 | 5.30E-05 | 0.000254367 |
| F13b | 2285.429669 | -1.10916277 | 0.293487232 | 5.37E-05 | 0.000257671 |
| Gabbr2 | 212.778895 | -1.48633402 | 0.410276416 | 5.72E-05 | 0.000273031 |
| mt-Rnr1 | 5026.638486 | -1.018481365 | 0.265979537 | 5.73E-05 | 0.000273308 |
| Hc | 14296.95759 | -1.026135281 | 0.27071618 | 5.76E-05 | 0.000274676 |
| Gm43597 | 46.29901883 | -1.382479504 | 0.375937722 | 5.89E-05 | 0.000280284 |
| 4833411C07Rik | 77.52395679 | -1.434891476 | 0.395105506 | 6.09E-05 | 0.000288344 |
| Hhat | 87.15673383 | -1.296589293 | 0.350528226 | 6.13E-05 | 0.000290035 |
| Gm12669 | 11.90941133 | -1.889705989 | 0.537390712 | 6.45E-05 | 0.000304459 |

| | | | | | |
|---|---|---|---|---|---|
| Fitm1 | 46.20192866 | -1.57225439 | 0.442249759 | 6.79E-05 | 0.000319511 |
| Notch1 | 929.5893766 | -1.065811318 | 0.285398546 | 6.83E-05 | 0.000321081 |
| Gm50432 | 4.630030227 | -5.867088108 | 2.678861987 | 6.88E-05 | 0.000323279 |
| Ncmap | 229.5432408 | -5.041212682 | 2.379706113 | 6.95E-05 | 0.000325839 |
| Vtcn1 | 294.8244656 | -1.277230842 | 0.347782902 | 7.07E-05 | 0.000331113 |
| Ypel2 | 930.4189795 | -1.270606757 | 0.348827238 | 7.25E-05 | 0.000338455 |
| Igha | 258.8135344 | -2.352599975 | 0.701778703 | 7.44E-05 | 0.00034649 |
| Sh2b2 | 42.59318919 | -1.370631311 | 0.379378444 | 7.47E-05 | 0.000347427 |
| Ighm | 4.44484961 | -5.761381877 | 2.663235608 | 8.02E-05 | 0.00037117 |
| Ido2 | 930.1584952 | -1.529319435 | 0.436365038 | 8.23E-05 | 0.000380254 |
| Crybb3 | 29.81360966 | -2.911680636 | 0.90009322 | 8.44E-05 | 0.000389144 |
| Cd83 | 45.87680581 | -2.460274718 | 0.705897376 | 8.62E-05 | 0.000396841 |
| Gm6206 | 15.42147281 | -1.856655511 | 0.538283181 | 8.85E-05 | 0.000406588 |
| Sec14l5 | 11.21496041 | -3.501365874 | 1.105419877 | 9.20E-05 | 0.000421403 |
| Hamp2 | 121.1923577 | -2.337349593 | 0.720043632 | 9.36E-05 | 0.000428233 |
| Pcp4l1 | 28.03302923 | -1.354361188 | 0.381893163 | 9.44E-05 | 0.000431812 |
| Selp | 8.243238668 | -2.971072677 | 0.912885964 | 9.64E-05 | 0.000439708 |
| Adra1b | 930.7245795 | -1.109674722 | 0.307298249 | 0.000100779 | 0.00045733 |
| Cd59b | 38.97008927 | -1.376087194 | 0.392131344 | 0.000103192 | 0.000467288 |
| Gm45727 | 7.6233824 | -2.682461087 | 0.829533917 | 0.000104121 | 0.000471159 |
| Nrros | 18.05988364 | -1.926409724 | 0.566352991 | 0.000107014 | 0.000482884 |
| Cps1 | 70214.499 | -1.50687687 | 0.437617186 | 0.000107193 | 0.00048358 |
| Cfhr3 | 113.9317049 | -1.025716812 | 0.283341227 | 0.000111082 | 0.000499243 |
| Gja10 | 4.691663376 | -4.797467652 | 1.583107997 | 0.000112249 | 0.000504136 |
| Clec4f | 54.60471096 | -2.253520422 | 0.698948745 | 0.000114702 | 0.00051443 |
| Tmem189 | 71.32148529 | -1.079282428 | 0.300200657 | 0.0001157 | 0.000517938 |
| Gm45338 | 55.25311974 | -1.015752379 | 0.281309183 | 0.000116056 | 0.000519047 |
| Lrrc8c | 111.2490945 | -1.887726315 | 0.551048545 | 0.000117895 | 0.00052666 |
| Gm14097 | 27.79823483 | -2.078699213 | 0.639229756 | 0.00012129 | 0.000541195 |
| Prox1os | 52.47955334 | -1.542848162 | 0.454565219 | 0.000125134 | 0.000556533 |
| B930025P03Rik | 13.09187764 | -3.183173177 | 1.039307243 | 0.000126216 | 0.000560959 |
| Mrc1 | 32.0589646 | -1.770296857 | 0.533213954 | 0.000128565 | 0.000569951 |
| Obox4-ps2 | 4.11096817 | -6.173120329 | 2.888208974 | 0.000131556 | 0.000581996 |
| Atp5pb-ps | 12.08419572 | -1.750009891 | 0.523159496 | 0.000135113 | 0.000595672 |
| Drc1 | 68.58000619 | -1.527920691 | 0.448648253 | 0.000135609 | 0.000597581 |
| Upp2 | 1589.728856 | -2.003925636 | 0.624219477 | 0.000140399 | 0.000616197 |
| 1700001C19Rik | 71.9341554 | -1.261644707 | 0.362731544 | 0.000140919 | 0.000618143 |
| Slc22a3 | 21.26046218 | -2.022815598 | 0.628902242 | 0.000141892 | 0.000621703 |
| Jchain | 55.77767149 | -2.408654704 | 0.750796546 | 0.000142788 | 0.00062534 |
| Gm6789 | 94.99565075 | -1.036209726 | 0.291686336 | 0.000145836 | 0.000636801 |
| Mrgprb11-ps | 5.197168801 | -3.328077099 | 1.05043512 | 0.000147108 | 0.000641481 |
| Caln1 | 26.24344278 | -1.463498014 | 0.433189292 | 0.0001485 | 0.000645934 |
| Car5b | 56.19520193 | -1.096956469 | 0.311599104 | 0.000151663 | 0.000658937 |

| | | | | | |
|---|---|---|---|---|---|
| Gm49024 | 9.100791611 | -2.927127332 | 0.926784078 | 0.000152004 | 0.000659831 |
| Aspa | 107.0407348 | -1.382612488 | 0.407462383 | 0.000156352 | 0.000677018 |
| 4933404O12Rik | 39.20438779 | -1.342999058 | 0.394778813 | 0.000165284 | 0.000711999 |
| Wdr89 | 48.13687689 | -1.010491989 | 0.286896304 | 0.000171651 | 0.000735304 |
| Siglec1 | 10.66868237 | -3.350971756 | 1.110297766 | 0.000184222 | 0.00078425 |
| 1810046K07Rik | 15.03150161 | -2.590165367 | 0.861224276 | 0.000189443 | 0.000804338 |
| Ackr2 | 12.13444201 | -2.221546882 | 0.721199699 | 0.000199598 | 0.000842419 |
| Gm15746 | 16.45052063 | -1.671110342 | 0.520065515 | 0.00020432 | 0.000860834 |
| Gm15611 | 7.473531536 | -3.701901336 | 1.273448635 | 0.000212536 | 0.000893489 |
| Gm41556 | 4.992043749 | -5.304788902 | 2.634219569 | 0.000213944 | 0.000898819 |
| A630076J17Rik | 4.819917767 | -3.943787223 | 1.223926606 | 0.000216156 | 0.000907513 |
| Cyp2c23 | 2472.217608 | -1.807660572 | 0.581407493 | 0.000229689 | 0.000959712 |
| Gm12164 | 18.36929233 | -1.582728468 | 0.491623801 | 0.00023003 | 0.000960718 |
| A930007A09Rik | 88.24952887 | -1.608458691 | 0.4925813 | 0.000235635 | 0.000982203 |
| Gm42568 | 9.017933735 | -2.010017455 | 0.648181029 | 0.000243763 | 0.001011689 |
| Gm45470 | 46.73531627 | -1.18944949 | 0.355236081 | 0.000245115 | 0.001016422 |
| Chp2 | 7.154584186 | -2.468790374 | 0.81570427 | 0.000246691 | 0.001022293 |
| 1810008I18Rik | 445.7443689 | -1.846393954 | 0.600365402 | 0.000254751 | 0.001052965 |
| Slc18a2 | 31.89179853 | -1.263045429 | 0.385505269 | 0.000286248 | 0.001171548 |
| Gm4332 | 25.17308006 | -1.320614431 | 0.406564472 | 0.000293988 | 0.001199899 |
| Stard8 | 132.1533907 | -1.09802691 | 0.328789569 | 0.000296546 | 0.001209568 |
| Gm7895 | 8.349603608 | -2.528926314 | 0.849926914 | 0.000313833 | 0.001272775 |
| Tox2 | 124.0917439 | -1.782428902 | 0.577904674 | 0.000316103 | 0.001281168 |
| Lrit2 | 25.7335633 | -1.315005861 | 0.4106021 | 0.000320465 | 0.001298027 |
| Slc35f1 | 90.30474071 | -4.417135134 | 2.330719096 | 0.000320794 | 0.001299084 |
| Gm40787 | 7.533144799 | -2.716903412 | 0.957915536 | 0.000336355 | 0.001354667 |
| Grem2 | 128.2819926 | -1.085192336 | 0.32916105 | 0.000341524 | 0.001372606 |
| Gm12183 | 39.77022469 | -1.209772025 | 0.374084509 | 0.000353299 | 0.00141283 |
| Tgtp1 | 189.0783086 | -1.100854282 | 0.336486951 | 0.000365359 | 0.001457107 |
| mt-Tl2 | 78.53900205 | -1.463144952 | 0.471669016 | 0.000375213 | 0.001492996 |
| Gm4963 | 26.87866694 | -1.299726823 | 0.408758765 | 0.000376717 | 0.001498048 |
| Gm6542 | 8.838768437 | -2.113123608 | 0.713491617 | 0.000377697 | 0.001501635 |
| 5730435O14Rik | 7.122775296 | -2.645775156 | 0.933784743 | 0.000380694 | 0.001513235 |
| Plag1 | 13.74807921 | -2.28168917 | 0.758374988 | 0.000394562 | 0.001561569 |
| Lrig3 | 180.1754024 | -1.028735152 | 0.311482867 | 0.000394829 | 0.001561978 |
| Nat8 | 52.55109667 | -1.669001015 | 0.555261818 | 0.000397921 | 0.001571619 |
| Gm14279 | 6.343731913 | -2.507405631 | 0.860942466 | 0.000414454 | 0.001632552 |
| Gm5431 | 12.428755 | -4.374155235 | 2.362810755 | 0.000416816 | 0.001641092 |
| Inka2 | 56.42789364 | -1.14813249 | 0.356308551 | 0.000419372 | 0.001649554 |
| Nfam1 | 5.709927187 | -4.249780581 | 1.59720696 | 0.000424687 | 0.001667043 |
| Col25a1 | 5.17946948 | -3.493932451 | 1.253697492 | 0.000428224 | 0.001679897 |
| Adra1a | 51.51763476 | -1.262193516 | 0.403701615 | 0.000469333 | 0.001826971 |
| Brdt | 51.18042461 | -1.069490237 | 0.332632046 | 0.000471102 | 0.001832001 |

| | | | | | |
|---|---|---|---|---|---|
| Klhl24 | 1994.956075 | -1.006900033 | 0.311763063 | 0.000485291 | 0.001882601 |
| Gm43611 | 10.74804082 | -1.867834216 | 0.647036553 | 0.000485393 | 0.001882614 |
| Rps18-ps5 | 21.47443429 | -1.267595992 | 0.405035525 | 0.000489783 | 0.001897725 |
| Gm11832 | 12.22925373 | -2.144914221 | 0.76674463 | 0.000509283 | 0.001964159 |
| Gm7466 | 5.152819906 | -2.896660578 | 1.057470988 | 0.00052914 | 0.002035017 |
| Gm15564 | 41.73883714 | -1.703124111 | 0.587098798 | 0.000537755 | 0.002066905 |
| A330023F24Rik | 20.64642534 | -1.44019442 | 0.478831038 | 0.000542946 | 0.002085604 |
| Rnft2 | 56.9558448 | -3.580205582 | 1.303961886 | 0.000548138 | 0.002104706 |
| Rpl39-ps | 15.86546197 | -1.57112593 | 0.531474579 | 0.000575876 | 0.002199769 |
| Acsm2 | 27.15450323 | -1.694594099 | 0.589068756 | 0.00057614 | 0.00219991 |
| H2bc6 | 16.65272505 | -1.322146476 | 0.438000162 | 0.000623235 | 0.002363737 |
| Insig1 | 6344.08681 | -1.237844804 | 0.408810278 | 0.000657831 | 0.002481703 |
| Srcin1 | 23.17694225 | -2.982754158 | 1.046451865 | 0.000662495 | 0.002497829 |
| P2rx3 | 20.71700875 | -1.169816574 | 0.382372483 | 0.000679314 | 0.002555218 |
| Sox7 | 8.07751101 | -3.616704408 | 1.37689903 | 0.000681856 | 0.002562768 |
| Gtpbp4-ps1 | 3.955225657 | -5.878579997 | 3.229630729 | 0.000693565 | 0.00260321 |
| H3c15 | 14.72232009 | -2.934151177 | 1.195214171 | 0.000697794 | 0.002618568 |
| Gm2996 | 4.792976417 | -2.746614928 | 1.039153321 | 0.000707818 | 0.002653775 |
| 9030616G12Rik | 34.62347187 | -1.785577554 | 0.646456652 | 0.000726618 | 0.002716642 |
| Gng7 | 57.90539219 | -1.427920299 | 0.480646966 | 0.000734955 | 0.002744068 |
| Gm4366 | 37.83177618 | -1.002535872 | 0.321312861 | 0.000738303 | 0.002756035 |
| Gm48768 | 18.94616329 | -1.710349002 | 0.596259303 | 0.00073969 | 0.002760676 |
| Slc26a4 | 3.810237697 | -3.956801353 | 1.598967833 | 0.000750277 | 0.002796997 |
| Slc28a1 | 8.223945145 | -2.006716803 | 0.742425015 | 0.000750873 | 0.00279806 |
| Ces3a | 13722.8997 | -1.866657972 | 0.692981161 | 0.000760984 | 0.002830241 |
| Wfikkn2 | 11.76353479 | -3.515640099 | 1.350645523 | 0.000761794 | 0.002831695 |
| Prok1 | 35.54083115 | -2.335695178 | 0.917028073 | 0.000765827 | 0.002844394 |
| Gm5822 | 20.5369602 | -1.298138805 | 0.4364409 | 0.000766129 | 0.002844966 |
| Tubb4b-ps2 | 5.698631028 | -3.251552536 | 1.241518568 | 0.000775005 | 0.00287459 |
| Gm6192 | 43.18435558 | -1.038793106 | 0.337962657 | 0.000783264 | 0.002901856 |
| Gm7105 | 15.78611848 | -1.492926087 | 0.521299088 | 0.000812152 | 0.002997308 |
| Gm5560 | 16.55888807 | -1.663092152 | 0.597806358 | 0.000815933 | 0.003008948 |
| Gm37499 | 15.78804942 | -1.792608342 | 0.638230652 | 0.000832474 | 0.003060538 |
| Dnase1l3 | 34.83746134 | -1.772770867 | 0.653318214 | 0.000838685 | 0.003079832 |
| Lama4 | 1042.066869 | -2.096922555 | 0.747226889 | 0.000841131 | 0.003087633 |
| Fam228b | 27.5443975 | -1.38448318 | 0.478571774 | 0.000846609 | 0.003105958 |
| Gm16418 | 6.003160582 | -2.349737716 | 0.907460223 | 0.000853837 | 0.003126501 |
| Rpsa-ps7 | 9.861355953 | -1.844268544 | 0.68023298 | 0.00093056 | 0.003374598 |
| Zfp-ps | 3.088476683 | -4.799431888 | 2.762921235 | 0.000934295 | 0.003385182 |
| Pim1 | 357.6627216 | -1.434213318 | 0.508745079 | 0.000945682 | 0.003422322 |
| Tbx3os1 | 12.29936202 | -2.323325966 | 0.938207277 | 0.000959881 | 0.003463912 |
| Cd163 | 8.136828457 | -3.380379305 | 1.434842357 | 0.000974923 | 0.003512908 |
| Map9 | 133.7705127 | -1.224027305 | 0.41660485 | 0.000976945 | 0.003519532 |

| | | | | | |
|---|---|---|---|---|---|
| Gm10433 | 8.677071695 | -2.068726496 | 0.7983805 | 0.000993984 | 0.003572868 |
| Olfr1034 | 29.41337433 | -1.170022072 | 0.399070766 | 0.00099949 | 0.003591986 |
| Gm20632 | 31.8846595 | -1.256973253 | 0.435533274 | 0.001029164 | 0.003690339 |
| Gm6900 | 26.03911609 | -1.000788994 | 0.332839315 | 0.001031298 | 0.003695917 |
| Ighg2b | 13.01932977 | -3.451457513 | 1.471341494 | 0.001040294 | 0.003723292 |
| Gm32511 | 9.0475539 | -1.996481469 | 0.778989262 | 0.00104201 | 0.003728042 |
| Gm45716 | 32.43482489 | -1.164341335 | 0.398457713 | 0.001044573 | 0.003736516 |
| Gm41231 | 9.056620413 | -1.928523356 | 0.740072823 | 0.001064874 | 0.003799217 |
| Gm6988 | 8.431371036 | -1.953143053 | 0.742007243 | 0.001092165 | 0.003880716 |
| Gm6580 | 5.633104267 | -2.986605421 | 1.245699826 | 0.001105979 | 0.003925437 |
| Pfn4 | 6.808110666 | -2.235849194 | 0.86512369 | 0.001124423 | 0.003985742 |
| Capn3 | 3.927502696 | -3.02963013 | 1.208420909 | 0.001142085 | 0.004043119 |
| Loxl4 | 268.2265955 | -1.064176298 | 0.361536689 | 0.001145508 | 0.004052994 |
| Hnf4g | 40.52186957 | -2.292820333 | 0.837526499 | 0.001149608 | 0.004065951 |
| Ugt1a5 | 138.0769787 | -1.139277713 | 0.39409326 | 0.0011538 | 0.004078573 |
| Gm37691 | 12.28615943 | -1.707054425 | 0.639754332 | 0.001163416 | 0.004108025 |
| Tex12 | 6.91275814 | -2.158427035 | 0.861398067 | 0.001193323 | 0.0042028 |
| Clec12a | 3.367056039 | -4.140598807 | 2.502484559 | 0.001202824 | 0.004232379 |
| Lrrc51 | 24.88111659 | -1.262903326 | 0.444961875 | 0.001232387 | 0.004325301 |
| Eif4ebp3 | 19.18865728 | -1.530757542 | 0.568578414 | 0.001254519 | 0.004392532 |
| Slc46a3 | 511.7793038 | -1.314172522 | 0.474323354 | 0.001271668 | 0.004448129 |
| Cib2 | 18.19448312 | -1.477895107 | 0.543625787 | 0.001284979 | 0.004492629 |
| 3930402G23Rik | 4.883040736 | -2.545121067 | 1.03743184 | 0.001306806 | 0.004554008 |
| Nhsl2 | 16.64455973 | -3.782571472 | 2.329244599 | 0.001306638 | 0.004554008 |
| Aplp1 | 281.9969343 | -1.23169647 | 0.430729087 | 0.001309648 | 0.004562254 |
| Chac1 | 57.75495605 | -1.319063153 | 0.477206235 | 0.001323681 | 0.004602783 |
| Ehd3 | 53.85588162 | -1.614765733 | 0.618726289 | 0.001351646 | 0.004687284 |
| Cox5b-ps | 28.06023937 | -1.396363482 | 0.514873484 | 0.001370671 | 0.004748971 |
| Gm5540 | 17.50603181 | -1.43222219 | 0.530043312 | 0.001392684 | 0.004816543 |
| Rsad2 | 115.2819557 | -1.196924286 | 0.428383913 | 0.001457159 | 0.00500794 |
| Gm8798 | 9.005877727 | -1.951978004 | 0.790304964 | 0.001499112 | 0.00513557 |
| Gm49437 | 13.47180278 | -1.436656878 | 0.537187882 | 0.001500914 | 0.005140824 |
| Cct6b | 13.51067082 | -1.425402104 | 0.530493409 | 0.001504617 | 0.005150666 |
| Gm21399 | 36.81898051 | -1.067431121 | 0.375114315 | 0.00151962 | 0.005198399 |
| Irf8 | 24.5113928 | -1.323520116 | 0.48093156 | 0.001527359 | 0.005223945 |
| Mup6 | 4.235855954 | -4.068779532 | 2.0943264 | 0.001540333 | 0.005265503 |
| 9930111J21Rik2 | 16.82877235 | -3.737023654 | 2.355577495 | 0.001543461 | 0.005275256 |
| Insc | 200.0343406 | -1.125358373 | 0.400836921 | 0.001545487 | 0.005281239 |
| Hsd17b6 | 1423.229486 | -1.854853703 | 0.769142618 | 0.001599581 | 0.005447593 |
| Ighg2c | 8.914419762 | -3.742191786 | 1.685594552 | 0.001611224 | 0.00547856 |
| Gm7331 | 19.14797796 | -1.282949803 | 0.47061012 | 0.001646556 | 0.005587808 |
| Gm8451 | 21.85094826 | -1.74591801 | 0.703167457 | 0.001660872 | 0.005630418 |
| Hspa9-ps1 | 3.021863686 | -3.548138904 | 1.642329658 | 0.001738934 | 0.005861916 |

| | | | | | |
|---|---|---|---|---|---|
| Corin | 8.00937149 | -1.801321719 | 0.730621204 | 0.001760962 | 0.005933043 |
| Gm37736 | 15.0241769 | -1.786364535 | 0.738314698 | 0.001791644 | 0.006024783 |
| 2900052L18Rik | 82.86557388 | -1.403905696 | 0.539839198 | 0.001841898 | 0.006171067 |
| Gm14270 | 6.726370858 | -1.908721674 | 0.790583262 | 0.001915442 | 0.00639069 |
| Gm21885 | 17.2224576 | -1.499418782 | 0.589275037 | 0.001915824 | 0.006390854 |
| Cd4 | 3.735848116 | -3.910439528 | 2.009334121 | 0.001936862 | 0.006453179 |
| Rpl17-ps5 | 10.01682001 | -1.887983396 | 0.780541516 | 0.00194188 | 0.006468516 |
| Chchd2-ps | 8.305355305 | -1.754967459 | 0.718232801 | 0.001968031 | 0.0065468 |
| Gm5835 | 33.60748241 | -1.11643606 | 0.410143415 | 0.002013045 | 0.006679175 |
| Gm45424 | 7.034473134 | -3.309308298 | 1.537756694 | 0.002039674 | 0.006764019 |
| Rpsa-ps9 | 6.425089595 | -1.997794458 | 0.843678786 | 0.002067337 | 0.00684039 |
| Fras1 | 191.4035981 | -1.003795259 | 0.362061989 | 0.002082518 | 0.006879945 |
| Klf1 | 33.83578837 | -1.171824047 | 0.436570434 | 0.002093501 | 0.006913848 |
| Gm35315 | 34.9068849 | -1.033827182 | 0.375078379 | 0.002132276 | 0.00702081 |
| Gm10157 | 4.733434348 | -2.216952241 | 0.954199446 | 0.002289336 | 0.007471879 |
| Gm43254 | 23.12992486 | -1.350407458 | 0.522993784 | 0.00233864 | 0.007617252 |
| Gm50454 | 3.972542918 | -2.669146439 | 1.160888127 | 0.002369868 | 0.007708498 |
| Inmt | 2946.144281 | -1.522570748 | 0.627459924 | 0.002389785 | 0.007765384 |
| Gm34333 | 20.04864502 | -1.228811246 | 0.47271315 | 0.002410079 | 0.007823381 |
| 4932422M17Rik | 30.09553161 | -1.089988983 | 0.406236249 | 0.002435615 | 0.007896924 |
| Gm4890 | 23.69931967 | -1.292500606 | 0.500839187 | 0.002438624 | 0.00790533 |
| Ldhb | 209.098389 | -1.144262676 | 0.427031879 | 0.002441002 | 0.007909045 |
| 0610040B10Rik | 22.45368176 | -1.328123086 | 0.52240009 | 0.002499718 | 0.008079115 |
| Gm7392 | 7.007407282 | -1.929022804 | 0.816187806 | 0.002518888 | 0.008128465 |
| Cyp2c40 | 55.53062399 | -2.730534091 | 1.551327485 | 0.00253277 | 0.008169141 |
| E030037K01Rik | 6.79876147 | -2.060610331 | 0.922781694 | 0.002557638 | 0.008238273 |
| Hadhb-ps | 17.66734585 | -1.152279656 | 0.442375113 | 0.002768598 | 0.008849463 |
| Gm7429 | 14.25511815 | -1.299400897 | 0.51424469 | 0.002780899 | 0.008881382 |
| Trp53cor1 | 14.67722479 | -1.768215257 | 0.770457473 | 0.002822316 | 0.008995694 |
| Cacng7 | 4.724953302 | -2.556241878 | 1.157244124 | 0.0028575 | 0.009103299 |
| Adgre1 | 9.47130916 | -2.804413028 | 1.476020359 | 0.002891506 | 0.009196369 |
| Rpl28-ps3 | 22.25508918 | -1.048829022 | 0.396303848 | 0.002897523 | 0.009213978 |
| Gm12247 | 11.59624442 | -1.402922778 | 0.574342187 | 0.002932333 | 0.009310787 |
| Gm50063 | 5.946494593 | -2.260240508 | 1.097431292 | 0.002955389 | 0.009379342 |
| Serpina3c | 5.155331749 | -2.677669721 | 1.351910387 | 0.00297251 | 0.009428997 |
| Gm6781 | 6.521443027 | -2.453667497 | 1.187754243 | 0.002989335 | 0.009474538 |
| 9230111E07Rik | 5.203356706 | -3.011922136 | 1.359700768 | 0.003045996 | 0.009633438 |
| Shisa7 | 17.35976496 | -2.884892021 | 1.374200666 | 0.003094442 | 0.009773772 |
| Eif3j2 | 24.70260858 | -1.014730518 | 0.385102447 | 0.003140284 | 0.009902264 |
| Spr-ps1 | 6.818730786 | -3.023307882 | 1.521142115 | 0.003142794 | 0.009908553 |
| Gm42427 | 4.708310653 | -2.266180672 | 1.027126883 | 0.003161326 | 0.009962072 |
| Gm30117 | 7.717643251 | -2.061667634 | 0.989062833 | 0.003163683 | 0.009966224 |
| Ces2b | 2.260503465 | -4.347746574 | 3.218499323 | 0.003175369 | 0.009999755 |

| | | | | | |
|---|---|---|---|---|---|
| Gm5425 | 7.871936726 | -1.789490052 | 0.803537131 | 0.003176451 | 0.010001394 |
| Keg1 | 1201.063923 | -1.740804992 | 0.80087723 | 0.003184422 | 0.010021687 |
| Gm13855 | 3.685190896 | -3.524046037 | 2.06736647 | 0.003268128 | 0.010258202 |
| Cd37 | 3.568173229 | -3.63470317 | 2.631677515 | 0.00327864 | 0.010279428 |
| Ptprc | 10.3577448 | -2.06003739 | 0.986378792 | 0.003284656 | 0.010296609 |
| Xlr3a | 101.2307206 | -1.460411072 | 0.623398924 | 0.003302315 | 0.010341828 |
| Gm31717 | 5.065401613 | -2.512936446 | 1.330247792 | 0.003418805 | 0.010663138 |
| Oaz1-ps | 43.71968148 | -1.099813277 | 0.430744479 | 0.003418374 | 0.010663138 |
| Rpl7a-ps3 | 1.971937308 | -4.059065665 | 3.073897183 | 0.003516278 | 0.010931619 |
| Gm30931 | 4.827639772 | -2.158568353 | 1.081098435 | 0.003542418 | 0.011002192 |
| 1700029J07Rik | 59.74442814 | -1.431870721 | 0.594004776 | 0.003550191 | 0.011020983 |
| Cyp2c69 | 13.83888871 | -2.438875656 | 1.381411324 | 0.003567198 | 0.011068407 |
| BC048644 | 39.21743241 | -1.012784954 | 0.395032794 | 0.003709976 | 0.011466926 |
| Itgb2 | 7.431826891 | -2.437599879 | 1.264569274 | 0.003710777 | 0.011467556 |
| C1qc | 23.46457007 | -1.835908278 | 0.874712289 | 0.003740841 | 0.011543734 |
| Espnl | 3.576752508 | -3.719231718 | 2.82493902 | 0.003802944 | 0.011704348 |
| D030034A15Rik | 2.653942049 | -4.303422269 | 3.487013265 | 0.003883081 | 0.011907992 |
| Gm4013 | 25.85581498 | -1.066255255 | 0.423457708 | 0.003966086 | 0.01213926 |
| Afap1l1 | 19.95070606 | -1.806884635 | 0.824253125 | 0.003990656 | 0.012198897 |
| Gm4742 | 2.862441683 | -2.94609766 | 1.596022305 | 0.004115779 | 0.01253148 |
| Vmn2r20 | 3.06884068 | -3.3481993 | 2.26687804 | 0.004277864 | 0.01297353 |
| Glns-ps1 | 4.853892041 | -2.394906479 | 1.291388259 | 0.004311179 | 0.013053934 |
| Ccn4 | 27.00820762 | -1.353940946 | 0.574540765 | 0.0043182 | 0.013069457 |
| Gm12350 | 15.32678218 | -1.46010015 | 0.652277676 | 0.004352624 | 0.013162813 |
| Gm49602 | 34.96009954 | -1.009266035 | 0.401370867 | 0.004524566 | 0.013629091 |
| Ighv1-80 | 2.438449771 | -3.759561193 | 3.117619555 | 0.004564889 | 0.013741928 |
| Coro2a | 44.55569317 | -1.196720022 | 0.495750084 | 0.004585267 | 0.013798944 |
| Cyp4f39 | 6.342779342 | -2.062213731 | 1.100109818 | 0.004587091 | 0.013802266 |
| Avil | 7.388813982 | -2.765799181 | 1.473026923 | 0.004636494 | 0.013934813 |
| Gm6055 | 7.417165273 | -1.56314027 | 0.711761483 | 0.004696198 | 0.014088594 |
| Marchf1 | 5.872799975 | -2.73025959 | 1.320068618 | 0.004706481 | 0.014115029 |
| Saa4 | 1985.288536 | -1.353903913 | 0.603644119 | 0.004769979 | 0.014276462 |
| Obox4-ps1 | 1.945293539 | -3.927963141 | 3.554741328 | 0.004882077 | 0.014566534 |
| Ndufab1-ps | 7.053353695 | -1.493285632 | 0.683316154 | 0.004936955 | 0.01470081 |
| Zfp36l1-ps | 5.994162274 | -2.23118269 | 1.201338607 | 0.005010131 | 0.014897869 |
| Slc16a13 | 72.38522849 | -1.568245661 | 0.696439564 | 0.005035517 | 0.014966177 |
| Gm2962 | 6.568687773 | -1.744327203 | 0.854810518 | 0.005043221 | 0.014986755 |
| Ikzf4 | 63.63816012 | -1.422966933 | 0.620883631 | 0.005069582 | 0.015048778 |
| Tubb4b-ps1 | 14.83640429 | -1.485839655 | 0.691568322 | 0.00511097 | 0.015150545 |
| Lrrc19 | 2.74631201 | -3.25137289 | 2.683548526 | 0.00511803 | 0.01516913 |
| Gm7208 | 4.422769583 | -2.530551029 | 1.549944302 | 0.005147151 | 0.015241316 |
| Gm5644 | 6.207225769 | -1.811923299 | 0.902468658 | 0.005188163 | 0.015346182 |
| Gm13651 | 2.377901949 | -3.686775087 | 3.235149348 | 0.005224718 | 0.015440031 |

| | | | | | |
|---|---|---|---|---|---|
| Sh3tc2 | 32.72765566 | -2.874259975 | 2.345260413 | 0.005228015 | 0.015447396 |
| Gm12435 | 10.22639307 | -1.282108567 | 0.558513396 | 0.005247343 | 0.015497347 |
| Rpl9-ps7 | 12.35370604 | -1.279261262 | 0.559656664 | 0.005268116 | 0.015546735 |
| Gm11769 | 2.83371162 | -3.217778553 | 2.694793688 | 0.005327577 | 0.015690843 |
| Gm10320 | 6.053480417 | -1.756453791 | 0.872449869 | 0.005341902 | 0.015727129 |
| Gm17040 | 7.108690824 | -1.955968549 | 1.067212161 | 0.005368978 | 0.015800652 |
| Gchfr | 532.5179654 | -1.345277363 | 0.613748493 | 0.005406618 | 0.015901672 |
| Gm10069 | 14.94003861 | -1.065917235 | 0.44257198 | 0.005413404 | 0.01591595 |
| Gm48488 | 3.900775578 | -2.639493033 | 1.320758891 | 0.005482645 | 0.016095677 |
| Acsl6 | 4.668498167 | -3.212765652 | 2.719918578 | 0.005507894 | 0.016164857 |
| Gm4853 | 2.999611076 | -3.097824695 | 2.315073172 | 0.005598079 | 0.016404448 |
| Gm13835 | 16.81999965 | -1.045115915 | 0.435160007 | 0.00564747 | 0.016534035 |
| Gm5461 | 2.230800024 | -3.32402582 | 2.900418218 | 0.005667206 | 0.016579168 |
| Gm7652 | 22.64085239 | -1.451594286 | 0.687603646 | 0.005697771 | 0.01665589 |
| Plxnc1 | 21.54129924 | -1.377727429 | 0.634440819 | 0.005791168 | 0.016908306 |
| Gm6807 | 12.36664678 | -1.245697199 | 0.545835919 | 0.005819311 | 0.016985308 |
| Cyp46a1 | 26.66361283 | -1.191866531 | 0.521397061 | 0.005856441 | 0.017063462 |
| Myom3 | 44.89391743 | -1.377558556 | 0.643770467 | 0.005969948 | 0.017343213 |
| Gm43444 | 4.159091746 | -2.487476712 | 1.47015164 | 0.006247847 | 0.018060316 |
| Rpl36-ps12 | 30.03118728 | -1.048721307 | 0.445404137 | 0.006265338 | 0.01809997 |
| Gm14288 | 2.819391318 | -3.115558325 | 2.790313309 | 0.006291615 | 0.018166113 |
| Gm7476 | 3.22583764 | -2.566670331 | 1.352520746 | 0.006296569 | 0.018176512 |
| Gm10136 | 13.86804738 | -1.177101342 | 0.515669123 | 0.006304952 | 0.018197975 |
| Commd5 | 3.689674459 | -2.610428202 | 1.565190732 | 0.006317616 | 0.018231785 |
| Gm12185 | 3.546828791 | -2.944565047 | 2.610382586 | 0.006452551 | 0.018584858 |
| Gata4 | 105.6331586 | -1.036589928 | 0.443866627 | 0.006522546 | 0.018755496 |
| Fcna | 20.62619477 | -1.901554221 | 1.134498756 | 0.006852463 | 0.01955474 |
| Spink5 | 9.389982952 | -1.380879845 | 0.664143667 | 0.006941006 | 0.019766263 |
| Muc1 | 1063.430292 | -2.735687501 | 2.449547819 | 0.00707682 | 0.020102316 |
| Gm42517 | 20.17534814 | -2.716844298 | 2.473190214 | 0.00708733 | 0.020119012 |
| Gm7079 | 3.762950839 | -2.622933969 | 1.517095279 | 0.007103173 | 0.020156291 |
| Gm46411 | 8.397250568 | -1.729859003 | 0.96067598 | 0.007304806 | 0.020673431 |
| Gm11992 | 38.21948997 | -1.097830154 | 0.483450989 | 0.007488903 | 0.021144555 |
| Gm14044 | 4.155798067 | -2.041548829 | 1.137167816 | 0.007532946 | 0.0212564 |
| 4933431K14Rik | 14.56607374 | -1.15898849 | 0.522028201 | 0.007804153 | 0.021919948 |
| Gm28720 | 7.079678192 | -1.698597177 | 0.941404746 | 0.008138223 | 0.022743648 |
| Gm7461 | 3.047577085 | -2.765585699 | 2.748370796 | 0.008160225 | 0.022795179 |
| Gm15634 | 2.553534082 | -2.657891246 | 2.437676132 | 0.008261409 | 0.023030897 |
| Cdnf | 7.970934097 | -1.613342336 | 0.863041218 | 0.008292088 | 0.023092942 |
| Psma5-ps | 3.940272259 | -1.924219775 | 1.064129047 | 0.008345969 | 0.023226144 |
| Gm12070 | 6.939441509 | -1.446203228 | 0.732895775 | 0.008390715 | 0.023323611 |
| C1qtnf4 | 23.25793332 | -1.321906726 | 0.632268833 | 0.00842137 | 0.023402042 |
| Gm12989 | 6.330060436 | -1.854522551 | 1.09732285 | 0.008455543 | 0.023480006 |

| | | | | | |
|---|---|---|---|---|---|
| C2cd4d | 17.81360803 | -1.018596592 | 0.45210683 | 0.008505362 | 0.023584222 |
| Gm15962 | 7.877273386 | -1.429084551 | 0.734241908 | 0.008504175 | 0.023584222 |
| Gm45301 | 5.787820002 | -1.672017314 | 0.956360416 | 0.008537883 | 0.023650478 |
| Gm14176 | 5.735929577 | -1.727093724 | 0.918554545 | 0.008569794 | 0.023725175 |
| Gm13420 | 41.95592989 | -1.637379753 | 0.841149981 | 0.008574046 | 0.023733523 |
| Ctcfl | 14.99400641 | -1.189236141 | 0.565274836 | 0.008697412 | 0.024026491 |
| Gngt1 | 6.901801064 | -1.565584957 | 0.857091863 | 0.008798598 | 0.024257718 |
| Scd3 | 8.515486382 | -1.825295438 | 0.974498317 | 0.008806882 | 0.024272996 |
| Gm13904 | 4.396824531 | -2.149562694 | 1.320930951 | 0.008880188 | 0.024446949 |
| Nfasc | 10.49046021 | -1.683725059 | 0.981551501 | 0.008881958 | 0.024448316 |
| Gm6392 | 4.641477395 | -1.964264058 | 1.221940134 | 0.008929669 | 0.024569072 |
| Car5a | 454.7033229 | -1.418462335 | 0.752942623 | 0.009023085 | 0.024790554 |
| Gm6740 | 5.426895596 | -1.905494967 | 1.23189883 | 0.009024593 | 0.024791148 |
| Rasl2-9 | 6.951730154 | -1.805532186 | 1.117067777 | 0.009102993 | 0.024979284 |
| Nwd2 | 3.654983315 | -1.799151565 | 1.049040682 | 0.009182002 | 0.025158738 |
| Gm2387 | 10.31620413 | -1.354919856 | 0.684575461 | 0.009223311 | 0.025261103 |
| Gm44744 | 10.9894217 | -1.372129366 | 0.70344896 | 0.009238987 | 0.02529321 |
| Pspn | 3.248545738 | -2.438940794 | 1.717415052 | 0.009253795 | 0.025315692 |
| Gm35665 | 6.216863856 | -1.605866411 | 0.919781698 | 0.009328803 | 0.025499083 |
| Fermt3 | 16.31082138 | -1.024438798 | 0.462822363 | 0.009414049 | 0.025706625 |
| Gm10039 | 8.10734532 | -1.337591282 | 0.674834022 | 0.009443521 | 0.025775937 |
| 1700023B13Rik | 10.79830285 | -1.500948471 | 0.817323973 | 0.009546098 | 0.026020231 |
| Gm16731 | 5.702263737 | -1.943645126 | 1.366042112 | 0.00958707 | 0.026112044 |
| Gm5171 | 3.325419679 | -2.410680787 | 1.730142407 | 0.009663938 | 0.026291556 |
| Rpl17-ps4 | 19.31345191 | -1.082892894 | 0.502519824 | 0.009694736 | 0.026360397 |
| Gm20768 | 2.838797686 | -2.443904323 | 1.857134504 | 0.009742854 | 0.026472479 |
| Gm22146 | 27.49731258 | -1.196497051 | 0.584409171 | 0.00980921 | 0.02662262 |
| C330013E15Rik | 22.9871098 | -1.211068284 | 0.591534276 | 0.009895124 | 0.026817869 |
| Nat8f6 | 5.004872258 | -2.128443415 | 1.952976287 | 0.009934702 | 0.026921331 |
| Ccng2 | 454.845879 | -1.151639871 | 0.553301744 | 0.009947723 | 0.02695008 |
| Gm12216 | 23.91178644 | -1.050136516 | 0.481426803 | 0.009958829 | 0.026970796 |
| Cd300lg | 7.296732644 | -1.562094121 | 0.88288769 | 0.010072958 | 0.027242128 |
| Gm7658 | 4.624602457 | -1.918988835 | 1.149196717 | 0.010125466 | 0.027376419 |
| Gm10819 | 6.294443844 | -1.505918074 | 0.819812581 | 0.010476097 | 0.028196791 |
| Cyp2b9 | 126.104431 | -1.873082387 | 1.830678015 | 0.010488144 | 0.028217856 |
| Prdx2-ps1 | 6.342012233 | -1.547466876 | 0.846782102 | 0.010495091 | 0.028224677 |
| Gdf11 | 18.07229697 | -2.144814462 | 1.138896904 | 0.010522001 | 0.028285155 |
| Psma8 | 10.80323587 | -1.274610249 | 0.645554207 | 0.010531891 | 0.02829985 |
| Ly6a | 120.8567282 | -1.069114527 | 0.501256849 | 0.010579375 | 0.028407553 |
| Nat1 | 13.29458659 | -1.273183393 | 0.653406243 | 0.010895498 | 0.029158485 |
| Gm13394 | 5.195303177 | -1.840497723 | 1.274787202 | 0.010901437 | 0.029170311 |
| Gm10053 | 7.686813391 | -1.556024584 | 0.901268701 | 0.010920096 | 0.029208024 |
| Gm6733 | 7.764481761 | -1.407421543 | 0.749069164 | 0.010968893 | 0.029322194 |

| | | | | | |
|---|---|---|---|---|---|
| Gm26616 | 13.2352968 | -1.044627639 | 0.493584715 | 0.011203222 | 0.029882007 |
| Pcolce2 | 3.594650114 | -2.305210742 | 1.785003797 | 0.011269639 | 0.030034113 |
| Fut4 | 46.73384781 | -2.159844182 | 1.415114919 | 0.011329811 | 0.030173523 |
| A530083I20Rik | 2.965048011 | -2.313158503 | 1.943947039 | 0.011489164 | 0.030551275 |
| Spata33 | 23.5128086 | -1.648254303 | 0.893829995 | 0.011574863 | 0.030736571 |
| Gm5457 | 3.216130893 | -1.823191389 | 1.203114361 | 0.011673421 | 0.030972575 |
| Tspan2 | 48.63978492 | -2.143502934 | 1.426640369 | 0.011712255 | 0.031071314 |
| P2ry4 | 85.58427689 | -1.740150626 | 1.363347844 | 0.01175469 | 0.031166657 |
| Gm45609 | 4.394684989 | -1.83836684 | 1.294944107 | 0.01178365 | 0.031234811 |
| Cd180 | 2.49383644 | -2.260288876 | 4.224480198 | 0.011821665 | 0.031318273 |
| Rpl19-ps11 | 103.5214918 | -1.123698491 | 0.558124237 | 0.011964626 | 0.031666409 |
| Gm9575 | 5.83073288 | -1.571565413 | 0.923432727 | 0.011969107 | 0.031673902 |
| Atad3aos | 8.491836848 | -1.443655895 | 0.800801228 | 0.012085865 | 0.031921254 |
| Gm19935 | 64.47491711 | -2.11379414 | 2.384780707 | 0.012220293 | 0.032236377 |
| Ndufa4l2 | 11.04792971 | -1.17895605 | 0.601423136 | 0.01222883 | 0.032250025 |
| Gm12254 | 14.68593975 | -1.308669651 | 0.704419704 | 0.012297987 | 0.032374578 |
| Mboat1 | 387.7333064 | -1.748089228 | 0.940241344 | 0.012348045 | 0.032488529 |
| Gm43429 | 13.58241762 | -1.089426433 | 0.534092946 | 0.012386921 | 0.032577415 |
| Fabp1 | 5665.047044 | -1.287708626 | 0.702901587 | 0.012507572 | 0.032854204 |
| Cyp3a59 | 243.0040363 | -1.243111817 | 0.664501371 | 0.012595888 | 0.03302742 |
| 9530080O11Rik | 7.685260137 | -1.438876393 | 0.830109532 | 0.012695508 | 0.033256823 |
| Slc4a8 | 23.06545903 | -2.052949786 | 2.404920863 | 0.012913553 | 0.033758887 |
| Hsd3b2 | 424.6461296 | -1.242324088 | 0.670657825 | 0.012986938 | 0.033918389 |
| Fam167a | 311.7179687 | -2.006513451 | 1.093846234 | 0.013124942 | 0.03422293 |
| Gm47602 | 10.20789157 | -1.613415464 | 0.909890404 | 0.013249588 | 0.03448235 |
| 9330121J05Rik | 6.751324277 | -1.388568147 | 0.796267457 | 0.013292434 | 0.034577153 |
| Oas3 | 40.17653559 | -2.005994138 | 2.425023921 | 0.013896965 | 0.035938102 |
| Flicr | 3.600762049 | -2.056056024 | 1.633656901 | 0.014001887 | 0.036177954 |
| Rps15-ps2 | 7.657203354 | -1.336013039 | 0.745101382 | 0.014045433 | 0.036278049 |
| Tmem132e | 2.479152124 | -1.704278994 | 2.976719441 | 0.01411146 | 0.036429704 |
| Car3 | 11777.77454 | -1.555610848 | 1.127932769 | 0.014218107 | 0.036664833 |
| Gm2058 | 4.451247136 | -1.812883206 | 1.191847165 | 0.014517382 | 0.037326242 |
| Gm42777 | 4.484898925 | -1.760378149 | 1.257562145 | 0.014577684 | 0.037466229 |
| Rab40b | 11.48975692 | -1.290039121 | 0.712262111 | 0.014653121 | 0.037624839 |
| Gm11335 | 5.503070163 | -2.001103904 | 1.653577163 | 0.01519802 | 0.038842084 |
| Gm12250 | 15.48408481 | -1.111556295 | 0.576988052 | 0.015203436 | 0.03885075 |
| Gm19196 | 9.10640304 | -1.447529648 | 0.834991701 | 0.015383341 | 0.039221696 |
| Gm20900 | 10.69258239 | -1.307326069 | 0.753175446 | 0.015469519 | 0.039394312 |
| P2ry6 | 20.07441305 | -1.516208168 | 0.903102505 | 0.015825602 | 0.040146613 |
| Vsig4 | 19.12971314 | -1.5791401 | 1.186981345 | 0.015830737 | 0.040154332 |
| Cd300ld | 3.800140866 | -2.0936462 | 1.848280279 | 0.015926358 | 0.040354205 |
| Gm15608 | 14.61158076 | -1.175865646 | 0.634144979 | 0.015942787 | 0.040385166 |
| Tagap | 7.892845261 | -1.84488439 | 2.428911784 | 0.01595813 | 0.040413365 |

| | | | | | |
|---|---|---|---|---|---|
| Cyp2d38-ps | 135.1252822 | -1.291624794 | 0.773150057 | 0.016107205 | 0.040774749 |
| Cdh5 | 22.75661424 | -1.302389371 | 0.769731423 | 0.016193545 | 0.040960896 |
| E230029C05Rik | 3.428990898 | -1.978125744 | 1.700082752 | 0.016228595 | 0.041027923 |
| Got1l1 | 3.303518296 | -1.789192602 | 2.058014964 | 0.016347429 | 0.041290274 |
| Slc30a2 | 2.699828127 | -1.819052915 | 2.799433108 | 0.016386976 | 0.041357503 |
| Serpina9 | 4.99340209 | -1.657450477 | 1.971140062 | 0.01645809 | 0.041531519 |
| Cdkl1 | 3.725095484 | -1.709062636 | 3.142923791 | 0.01656009 | 0.041734037 |
| Ksr2 | 16.95337411 | -1.814715824 | 2.420250454 | 0.016591414 | 0.041801924 |
| Gm5512 | 2.63531813 | -2.002437391 | 1.565510832 | 0.016613612 | 0.041841447 |
| Gm10232 | 6.863455904 | -1.299578504 | 0.752566252 | 0.016707676 | 0.042056271 |
| B230206L02Rik | 14.32575145 | -1.085452216 | 0.571703824 | 0.01683118 | 0.042317198 |
| 5830444B04Rik | 12.4705625 | -1.555667466 | 1.123591208 | 0.016866154 | 0.04238847 |
| Igkv6-15 | 2.966492718 | -1.731753522 | 2.738995294 | 0.017073973 | 0.042871465 |
| Gm4887 | 2.07898295 | -1.917803921 | 1.881456709 | 0.017176805 | 0.043107108 |
| Gm37419 | 3.488445271 | -1.73939124 | 1.167191132 | 0.017315886 | 0.043397599 |
| Cyfip2 | 219.6450768 | -1.650337786 | 1.091078561 | 0.017322878 | 0.043399914 |
| Smarce1-ps1 | 5.030634679 | -1.616024416 | 1.089931415 | 0.017885121 | 0.044587342 |
| Gm10644 | 16.49254918 | -1.219396231 | 0.711528742 | 0.018104048 | 0.045057033 |
| Chrm1 | 8.474492602 | -1.883989402 | 1.764450089 | 0.018139754 | 0.045139595 |
| Gm29863 | 2.282948114 | -1.501087434 | 2.379640567 | 0.018376485 | 0.045634463 |
| Wfdc17 | 8.095122121 | -1.424667013 | 0.950495744 | 0.018432743 | 0.045750487 |
| Kiss1r | 10.92723229 | -1.194624793 | 0.680464421 | 0.018615171 | 0.046143599 |
| Gm17080 | 21.7268911 | -1.095883609 | 0.605368654 | 0.019032097 | 0.047001025 |
| Gm13268 | 6.858668112 | -1.347325342 | 0.845020115 | 0.019165692 | 0.047270119 |
| Gm17802 | 4.312534003 | -1.47637756 | 1.008268694 | 0.019280326 | 0.04750401 |
| Gm8518 | 3.648777173 | -1.679787068 | 1.200874432 | 0.019543323 | 0.048077926 |
| Gm47761 | 17.34881042 | -1.338343661 | 0.88332782 | 0.019622117 | 0.048222315 |
| Folr2 | 2.972787477 | -1.896693923 | 1.423466781 | 0.01963256 | 0.048240794 |
| Gm3362 | 4.03335458 | -1.780287463 | 1.599512079 | 0.01972955 | 0.048431425 |
| Asb14 | 7.824817107 | -1.396581384 | 0.925671159 | 0.020250085 | 0.04944896 |
