## Supplementary material for "Physiological reprogramming *in vivo* mediated by Sox4 pioneer factor activity": Table S2

**Table S2. List of known hepatocyte and biliary/reprogrammed cell genes.**

| Hepatocyte genes (source) | | Biliary/Reprogrammed cell genes (source) | |
|---|---|---|---|
| *Alb* | (Tarlow et al., 2014) | *Ccl2* | (Tarlow *et al.*, 2014) |
| *Cyp7a1* | (Tarlow *et al.*, 2014) | *Zeb1* | (Tarlow *et al.*, 2014) |
| *F9* | (Tarlow *et al.*, 2014) | *Mst1r* | (Tarlow *et al.*, 2014) |
| *Fah* | (Tarlow *et al.*, 2014) | *Bmp4* | (Tarlow *et al.*, 2014) |
| *Hgd* | (Tarlow *et al.*, 2014) | *Vim* | (Tarlow *et al.*, 2014) |
| *Hnf4a* | (Tarlow *et al.*, 2014) | *Bmp1* | (Tarlow *et al.*, 2014) |
| *Tdo2* | (Tarlow *et al.*, 2014) | *Foxp2* | (Tarlow *et al.*, 2014) |
| *Cps1* | (Schaub et al., 2018) | *Sfrp4* | (Tarlow *et al.*, 2014) |
| *Cyp26a1* | (Schaub *et al.*, 2018) | *Etv4* | (Tarlow *et al.*, 2014) |
| *Cyp27a1* | (Schaub *et al.*, 2018) | *Dab1* | (Tarlow *et al.*, 2014) |
| *Cyp2c38* | (Schaub *et al.*, 2018) | *Kit* | (Tarlow *et al.*, 2014) |
| *Cyp2c67* | (Schaub *et al.*, 2018) | *Nes* | (Tarlow *et al.*, 2014) |
| *Cyp2d10* | (Schaub *et al.*, 2018) | *Ncam1* | (Tarlow *et al.*, 2014) |
| *Cyp2d11* | (Schaub *et al.*, 2018) | *Tbx1* | (Tarlow *et al.*, 2014) |
| *Cyp2d12* | (Schaub *et al.*, 2018) | *Mcam* | (Tarlow *et al.*, 2014) |
| *Cyp2d13* | (Schaub *et al.*, 2018) | *Tgfb1* | (Tarlow *et al.*, 2014) |
| *Cyp2d40* | (Schaub *et al.*, 2018) | *Fzd10* | (Tarlow *et al.*, 2014) |
| *Cyp2d9* | (Schaub *et al.*, 2018) | *Cav1* | (Tarlow *et al.*, 2014) |
| *Cyp2j5* | (Schaub *et al.*, 2018) | *Yap1* | (Tarlow *et al.*, 2014) |
| *Cyp2r1* | (Schaub *et al.*, 2018) | *Cd44* | (Tarlow *et al.*, 2014) |
| *Cyp3a11* | (Schaub *et al.*, 2018) | *Jag1* | (Tarlow *et al.*, 2014) |
| *Cyp4f13* | (Schaub *et al.*, 2018) | *Hnf1b* | (Tarlow *et al.*, 2014) |
| *Cyp4v3* | (Schaub *et al.*, 2018) | *Spp1* | (Tarlow *et al.*, 2014) |
| *Cyp8b1* | (Schaub *et al.*, 2018) | *Smo* | (Tarlow *et al.*, 2014) |
| *Foxa3* | (Schaub *et al.*, 2018) | *Itga3* | (Tarlow *et al.*, 2014) |
| *Hnf1a* | (Schaub *et al.*, 2018) | *Notch2* | (Tarlow *et al.*, 2014) |
| *Mup1* | (Schaub *et al.*, 2018) | *Foxj1* | (Tarlow *et al.*, 2014) |
| *Otc* | (Schaub *et al.*, 2018) | *Sox9* | (Tarlow *et al.*, 2014) |
| *Tat* | (Schaub *et al.*, 2018) | *Ccn1* | (Tarlow *et al.*, 2014) |
| *Trf* | (Schaub *et al.*, 2018) | *Tnfrsf12a* | (Tarlow *et al.*, 2014) |
| *Ttr* | (Schaub *et al.*, 2018) | *Krt19* | (Tarlow *et al.*, 2014) |
| *Abcb11* | (Katsuda et al., 2020) | *Grhl2* | (Tarlow *et al.*, 2014) |
| *Ahr* | (Katsuda *et al.*, 2020) | *Cftr* | (Tarlow *et al.*, 2014) |
| *Aldh1a1* | (Katsuda *et al.*, 2020) | *Epcam* | (Tarlow *et al.*, 2014) |
| *Ass1* | (Katsuda *et al.*, 2020) | *Krt7* | (Tarlow *et al.*, 2014) |
| *Baat* | (Katsuda *et al.*, 2020) | *Tacstd2* | (Tarlow *et al.*, 2014) |
| *Cyp1a2* | (Katsuda *et al.*, 2020) | *Ccn2* | (Tarlow *et al.*, 2014) |
| *Cyp2e1* | (Katsuda *et al.*, 2020) | *Krt17* | (Schaub *et al.*, 2018) |
| *G6pc* | (Katsuda *et al.*, 2020) | *Prom1* | (Schaub *et al.*, 2018) |
| *Gsta1* | (Katsuda *et al.*, 2020) | *Sstr2* | (Schaub *et al.*, 2018) |
| *Gys2* | (Katsuda *et al.*, 2020) | *St14* | (Schaub *et al.*, 2018) |
| *Nr1i2* | (Katsuda *et al.*, 2020) | *Cldn4* | (Schaub *et al.*, 2018) |

| | | | |
|---|---|---|---|
| *Nr1i3* | (Katsuda *et al.*, 2020) | *Muc1* | (Schaub *et al.*, 2018) |
| *Pck1* | (Katsuda *et al.*, 2020) | *Sox4* | (Poncy et al., 2015) |
| *Slc10a1* | (Katsuda *et al.*, 2020) | *Ezr* | (Merrell et al., 2021) |
| *Serpina7* | (Katsuda *et al.*, 2020) | *Cd24a* | (Merrell et al., 2021) |
| *Cebpa* | (Jakobsen et al., 2013) | *Igfbp7* | (Merrell et al., 2021) |
| *Cebpb* | (Jakobsen *et al.*, 2013) | *Itga6* | (Yanger et al., 2013) |
| *Asgr1* | (Peters et al., 2016) | *Cadm1* | (Ito et al., 2007) |
| *Fabp1* | (Huang et al., 2016) | | |
