## Supplementary material for "Physiological reprogramming *in vivo* mediated by Sox4 pioneer factor activity": Table S3

**Table S3. GO terms enriched in differentially opened regions in Sox4-expressing hepatocytes compared to empty virus-injected hepatocytes, and GO terms enriched for differentially expressed genes between hepatocytes and reprogrammed cells.**

**GO terms associated with 6742 genes near newly closed regions**

| ID | Description | GeneRatio | pvalue | p.adjust |
|---|---|---|---|---|
| GO:0006631 | fatty acid metabolic process | 210/5895 | 2.69E-30 | 1.70E-26 |
| GO:0008202 | steroid metabolic process | 162/5895 | 8.88E-25 | 2.80E-21 |
| GO:0044282 | small molecule catabolic process | 165/5895 | 3.49E-24 | 7.34E-21 |
| GO:1901615 | organic hydroxy compound metabolic process | 228/5895 | 1.07E-23 | 1.68E-20 |
| GO:0016054 | organic acid catabolic process | 124/5895 | 3.60E-22 | 3.79E-19 |
| GO:0046395 | carboxylic acid catabolic process | 124/5895 | 3.60E-22 | 3.79E-19 |
| GO:0019216 | regulation of lipid metabolic process | 176/5895 | 7.25E-22 | 6.53E-19 |
| GO:0046394 | carboxylic acid biosynthetic process | 152/5895 | 1.35E-21 | 1.07E-18 |
| GO:0016053 | organic acid biosynthetic process | 152/5895 | 2.07E-21 | 1.45E-18 |
| GO:0062012 | regulation of small molecule metabolic process | 173/5895 | 2.54E-21 | 1.60E-18 |
| GO:0006520 | cellular amino acid metabolic process | 131/5895 | 2.82E-20 | 1.62E-17 |
| GO:0006639 | acylglycerol metabolic process | 86/5895 | 3.66E-20 | 1.92E-17 |
| GO:1901605 | alpha-amino acid metabolic process | 103/5895 | 6.82E-20 | 3.31E-17 |
| GO:0006638 | neutral lipid metabolic process | 86/5895 | 1.41E-19 | 6.36E-17 |

| | | | | |
|---|---|---|---|---|
| GO:0006790 | sulfur compound metabolic process | 140/5895 | 2.40E-19 | 1.01E-16 |
| GO:0016042 | lipid catabolic process | 156/5895 | 3.60E-19 | 1.42E-16 |
| GO:0072521 | purine-containing compound metabolic process | 179/5895 | 3.52E-18 | 1.31E-15 |
| GO:0006641 | triglyceride metabolic process | 70/5895 | 4.98E-18 | 1.74E-15 |
| GO:0009117 | nucleotide metabolic process | 188/5895 | 1.50E-17 | 4.98E-15 |
| GO:0006753 | nucleoside phosphate metabolic process | 190/5895 | 3.70E-17 | 1.17E-14 |
| GO:0006066 | alcohol metabolic process | 150/5895 | 1.02E-16 | 3.01E-14 |
| GO:1902652 | secondary alcohol metabolic process | 82/5895 | 1.05E-16 | 3.01E-14 |
| GO:0006109 | regulation of carbohydrate metabolic process | 103/5895 | 1.46E-16 | 3.88E-14 |
| GO:0033865 | nucleoside bisphosphate metabolic process | 64/5895 | 1.60E-16 | 3.88E-14 |
| GO:0033875 | ribonucleoside bisphosphate metabolic process | 64/5895 | 1.60E-16 | 3.88E-14 |
| GO:0034032 | purine nucleoside bisphosphate metabolic process | 64/5895 | 1.60E-16 | 3.88E-14 |
| GO:0015711 | organic anion transport | 197/5895 | 2.04E-16 | 4.54E-14 |
| GO:0031589 | cell-substrate adhesion | 159/5895 | 2.09E-16 | 4.54E-14 |
| GO:0008203 | cholesterol metabolic process | 77/5895 | 2.14E-16 | 4.54E-14 |
| GO:0031669 | cellular response to nutrient levels | 100/5895 | 2.16E-16 | 4.54E-14 |
| GO:0010876 | lipid localization | 180/5895 | 2.47E-16 | 5.03E-14 |
| GO:0044262 | cellular carbohydrate metabolic process | 138/5895 | 2.84E-16 | 5.60E-14 |

| | | | | |
|---|---|---|---|---|
| GO:1901652 | response to peptide | 175/5895 | 4.13E-16 | 7.90E-14 |
| GO:0019693 | ribose phosphate metabolic process | 156/5895 | 4.88E-16 | 8.91E-14 |
| GO:0042493 | response to drug | 126/5895 | 4.94E-16 | 8.91E-14 |
| GO:0046486 | glycerolipid metabolic process | 161/5895 | 5.68E-16 | 9.91E-14 |
| GO:0009259 | ribonucleotide metabolic process | 152/5895 | 5.81E-16 | 9.91E-14 |
| GO:0009150 | purine ribonucleotide metabolic process | 148/5895 | 9.48E-16 | 1.57E-13 |
| GO:0055088 | lipid homeostasis | 85/5895 | 1.27E-15 | 2.06E-13 |
| GO:0016125 | sterol metabolic process | 79/5895 | 2.28E-15 | 3.59E-13 |
| GO:0045834 | positive regulation of lipid metabolic process | 90/5895 | 2.34E-15 | 3.60E-13 |
| GO:0072330 | monocarboxylic acid biosynthetic process | 101/5895 | 2.48E-15 | 3.65E-13 |
| GO:0030111 | regulation of Wnt signaling pathway | 140/5895 | 2.49E-15 | 3.65E-13 |
| GO:0006163 | purine nucleotide metabolic process | 157/5895 | 3.20E-15 | 4.59E-13 |
| GO:0006637 | acyl-CoA metabolic process | 55/5895 | 4.29E-15 | 5.89E-13 |
| GO:0035383 | thioester metabolic process | 55/5895 | 4.29E-15 | 5.89E-13 |
| GO:0009896 | positive regulation of catabolic process | 193/5895 | 4.86E-15 | 6.52E-13 |
| GO:0090407 | organophosphate biosynthetic process | 181/5895 | 5.24E-15 | 6.88E-13 |
| GO:0005996 | monosaccharide metabolic process | 124/5895 | 5.56E-15 | 7.16E-13 |
| GO:0031331 | positive regulation of cellular catabolic process | 170/5895 | 8.93E-15 | 1.11E-12 |

| | | | | |
|---|---|---|---|---|
| GO:0010675 | regulation of cellular carbohydrate metabolic process | 87/5895 | 8.94E-15 | 1.11E-12 |
| GO:0044242 | cellular lipid catabolic process | 108/5895 | 1.35E-14 | 1.64E-12 |
| GO:0042180 | cellular ketone metabolic process | 105/5895 | 1.46E-14 | 1.73E-12 |
| GO:0007264 | small GTPase mediated signal transduction | 183/5895 | 1.48E-14 | 1.73E-12 |
| GO:0031668 | cellular response to extracellular stimulus | 107/5895 | 1.81E-14 | 2.07E-12 |
| GO:0006694 | steroid biosynthetic process | 86/5895 | 1.92E-14 | 2.17E-12 |
| GO:0009267 | cellular response to starvation | 84/5895 | 2.11E-14 | 2.33E-12 |
| GO:0033500 | carbohydrate homeostasis | 127/5895 | 2.68E-14 | 2.92E-12 |
| GO:0006914 | autophagy | 178/5895 | 3.34E-14 | 3.51E-12 |
| GO:0061919 | process utilizing autophagic mechanism | 178/5895 | 3.34E-14 | 3.51E-12 |
| GO:0048732 | gland development | 193/5895 | 3.55E-14 | 3.68E-12 |
| GO:0050673 | epithelial cell proliferation | 182/5895 | 4.15E-14 | 4.22E-12 |
| GO:0042593 | glucose homeostasis | 126/5895 | 4.99E-14 | 5.00E-12 |
| GO:0007623 | circadian rhythm | 106/5895 | 5.27E-14 | 5.19E-12 |
| GO:0016055 | Wnt signaling pathway | 179/5895 | 5.52E-14 | 5.36E-12 |
| GO:0060537 | muscle tissue development | 192/5895 | 5.86E-14 | 5.61E-12 |
| GO:0010906 | regulation of glucose metabolic process | 72/5895 | 7.98E-14 | 7.52E-12 |
| GO:0198738 | cell-cell signaling by wnt | 179/5895 | 9.14E-14 | 8.48E-12 |

| | | | | |
|---|---|---|---|---|
| GO:0043434 | response to peptide hormone | 147/5895 | 9.29E-14 | 8.49E-12 |
| GO:0007517 | muscle organ development | 175/5895 | 9.55E-14 | 8.61E-12 |
| GO:0009063 | cellular amino acid catabolic process | 56/5895 | 1.37E-13 | 1.22E-11 |
| GO:0045666 | positive regulation of neuron differentiation | 191/5895 | 1.96E-13 | 1.72E-11 |
| GO:0006869 | lipid transport | 155/5895 | 3.01E-13 | 2.60E-11 |
| GO:1901653 | cellular response to peptide | 133/5895 | 3.14E-13 | 2.67E-11 |
| GO:0042594 | response to starvation | 90/5895 | 3.36E-13 | 2.83E-11 |
| GO:0072329 | monocarboxylic acid catabolic process | 68/5895 | 4.13E-13 | 3.42E-11 |
| GO:0046890 | regulation of lipid biosynthetic process | 92/5895 | 4.28E-13 | 3.51E-11 |
| GO:0071496 | cellular response to external stimulus | 126/5895 | 4.79E-13 | 3.87E-11 |
| GO:0048511 | rhythmic process | 133/5895 | 5.72E-13 | 4.57E-11 |
| GO:0006006 | glucose metabolic process | 98/5895 | 7.87E-13 | 6.21E-11 |
| GO:0071375 | cellular response to peptide hormone stimulus | 116/5895 | 8.62E-13 | 6.71E-11 |
| GO:0007265 | Ras protein signal transduction | 149/5895 | 8.84E-13 | 6.80E-11 |
| GO:0062013 | positive regulation of small molecule metabolic process | 81/5895 | 1.27E-12 | 9.66E-11 |
| GO:0001889 | liver development | 69/5895 | 1.49E-12 | 1.12E-10 |
| GO:0014706 | striated muscle tissue development | 180/5895 | 1.61E-12 | 1.19E-10 |
| GO:0042176 | regulation of protein catabolic process | 163/5895 | 1.74E-12 | 1.27E-10 |

| | | | | |
|---|---|---|---|---|
| GO:0051056 | regulation of small GTPase mediated signal transduction | 121/5895 | 2.65E-12 | 1.92E-10 |
| GO:0019395 | fatty acid oxidation | 61/5895 | 2.82E-12 | 2.02E-10 |
| GO:0019318 | hexose metabolic process | 107/5895 | 2.97E-12 | 2.10E-10 |
| GO:0031667 | response to nutrient levels | 146/5895 | 3.37E-12 | 2.36E-10 |
| GO:0001667 | ameboidal-type cell migration | 172/5895 | 4.86E-12 | 3.37E-10 |
| GO:0010976 | positive regulation of neuron projection development | 154/5895 | 5.27E-12 | 3.61E-10 |
| GO:0009062 | fatty acid catabolic process | 58/5895 | 6.35E-12 | 4.31E-10 |
| GO:0061008 | hepaticobiliary system development | 69/5895 | 6.52E-12 | 4.38E-10 |
| GO:0010810 | regulation of cell-substrate adhesion | 100/5895 | 7.92E-12 | 5.26E-10 |
| GO:0046578 | regulation of Ras protein signal transduction | 109/5895 | 8.67E-12 | 5.70E-10 |
| GO:0009895 | negative regulation of catabolic process | 133/5895 | 9.93E-12 | 6.46E-10 |
| GO:0051090 | regulation of DNA-binding transcription factor activity | 167/5895 | 1.09E-11 | 7.01E-10 |
| GO:0048638 | regulation of developmental growth | 165/5895 | 1.14E-11 | 7.26E-10 |
| GO:1901606 | alpha-amino acid catabolic process | 47/5895 | 1.70E-11 | 1.07E-09 |
| GO:0060560 | developmental growth involved in morphogenesis | 120/5895 | 2.08E-11 | 1.30E-09 |
| GO:0051098 | regulation of binding | 158/5895 | 2.11E-11 | 1.30E-09 |
| GO:0009991 | response to extracellular stimulus | 154/5895 | 2.28E-11 | 1.40E-09 |
| GO:0031346 | positive regulation of cell projection organization | 186/5895 | 2.34E-11 | 1.42E-09 |

| | | | | |
|---|---|---|---|---|
| GO:0032868 | response to insulin | 106/5895 | 2.78E-11 | 1.67E-09 |
| GO:0090132 | epithelium migration | 127/5895 | 2.91E-11 | 1.73E-09 |
| GO:0034440 | lipid oxidation | 62/5895 | 2.95E-11 | 1.74E-09 |
| GO:0010631 | epithelial cell migration | 126/5895 | 3.82E-11 | 2.23E-09 |
| GO:0055092 | sterol homeostasis | 53/5895 | 4.32E-11 | 2.50E-09 |
| GO:1901617 | organic hydroxy compound biosynthetic process | 102/5895 | 4.55E-11 | 2.61E-09 |
| GO:0090130 | tissue migration | 127/5895 | 5.00E-11 | 2.84E-09 |
| GO:0007160 | cell-matrix adhesion | 98/5895 | 5.37E-11 | 3.02E-09 |
| GO:0010565 | regulation of cellular ketone metabolic process | 71/5895 | 5.53E-11 | 3.08E-09 |
| GO:0016051 | carbohydrate biosynthetic process | 92/5895 | 5.57E-11 | 3.08E-09 |
| GO:0034329 | cell junction assembly | 168/5895 | 6.34E-11 | 3.48E-09 |
| GO:0032869 | cellular response to insulin stimulus | 93/5895 | 8.25E-11 | 4.49E-09 |
| GO:1901342 | regulation of vasculature development | 145/5895 | 8.86E-11 | 4.78E-09 |
| GO:0042632 | cholesterol homeostasis | 52/5895 | 9.95E-11 | 5.32E-09 |
| GO:0050678 | regulation of epithelial cell proliferation | 150/5895 | 1.01E-10 | 5.37E-09 |
| GO:0000209 | protein polyubiquitination | 105/5895 | 1.26E-10 | 6.60E-09 |
| GO:0043542 | endothelial cell migration | 96/5895 | 1.80E-10 | 9.40E-09 |
| GO:0046889 | positive regulation of lipid biosynthetic process | 55/5895 | 2.14E-10 | 1.11E-08 |

| | | | | |
|---|---|---|---|---|
| GO:0043648 | dicarboxylic acid metabolic process | 49/5895 | 2.24E-10 | 1.14E-08 |
| GO:0046364 | monosaccharide biosynthetic process | 49/5895 | 2.24E-10 | 1.14E-08 |
| GO:0030522 | intracellular receptor signaling pathway | 94/5895 | 2.31E-10 | 1.16E-08 |
| GO:0001655 | urogenital system development | 150/5895 | 2.52E-10 | 1.26E-08 |
| GO:0006575 | cellular modified amino acid metabolic process | 79/5895 | 2.81E-10 | 1.40E-08 |
| GO:0001676 | long-chain fatty acid metabolic process | 59/5895 | 2.98E-10 | 1.47E-08 |
| GO:0006635 | fatty acid beta-oxidation | 43/5895 | 3.22E-10 | 1.57E-08 |
| GO:0001678 | cellular glucose homeostasis | 78/5895 | 3.76E-10 | 1.82E-08 |
| GO:0045785 | positive regulation of cell adhesion | 168/5895 | 4.20E-10 | 2.02E-08 |
| GO:0010769 | regulation of cell morphogenesis involved in differentiation | 141/5895 | 4.95E-10 | 2.36E-08 |
| GO:0019319 | hexose biosynthetic process | 46/5895 | 5.02E-10 | 2.38E-08 |
| GO:0060070 | canonical Wnt signaling pathway | 119/5895 | 5.50E-10 | 2.59E-08 |
| GO:0046942 | carboxylic acid transport | 135/5895 | 5.64E-10 | 2.63E-08 |
| GO:0120254 | olefinic compound metabolic process | 64/5895 | 6.50E-10 | 3.00E-08 |
| GO:0043547 | positive regulation of GTPase activity | 98/5895 | 6.51E-10 | 3.00E-08 |
| GO:0043087 | regulation of GTPase activity | 125/5895 | 6.67E-10 | 3.05E-08 |
| GO:0042445 | hormone metabolic process | 89/5895 | 6.88E-10 | 3.05E-08 |
| GO:0006695 | cholesterol biosynthetic process | 33/5895 | 6.92E-10 | 3.05E-08 |

| | | | | |
|---|---|---|---|---|
| GO:0090207 | regulation of triglyceride metabolic process | 33/5895 | 6.92E-10 | 3.05E-08 |
| GO:1902653 | secondary alcohol biosynthetic process | 33/5895 | 6.92E-10 | 3.05E-08 |
| GO:0060828 | regulation of canonical Wnt signaling pathway | 102/5895 | 6.95E-10 | 3.05E-08 |
| GO:0015849 | organic acid transport | 136/5895 | 6.97E-10 | 3.05E-08 |
| GO:0070371 | ERK1 and ERK2 cascade | 132/5895 | 7.57E-10 | 3.29E-08 |
| GO:1901568 | fatty acid derivative metabolic process | 74/5895 | 8.28E-10 | 3.58E-08 |
| GO:0010256 | endomembrane system organization | 159/5895 | 8.65E-10 | 3.71E-08 |
| GO:1990778 | protein localization to cell periphery | 144/5895 | 9.00E-10 | 3.84E-08 |
| GO:0048193 | Golgi vesicle transport | 113/5895 | 9.56E-10 | 4.03E-08 |
| GO:0040013 | negative regulation of locomotion | 132/5895 | 9.58E-10 | 4.03E-08 |
| GO:0045765 | regulation of angiogenesis | 130/5895 | 9.96E-10 | 4.16E-08 |
| GO:0019218 | regulation of steroid metabolic process | 58/5895 | 1.00E-09 | 4.16E-08 |
| GO:0014855 | striated muscle cell proliferation | 45/5895 | 1.17E-09 | 4.82E-08 |
| GO:0071322 | cellular response to carbohydrate stimulus | 74/5895 | 1.20E-09 | 4.89E-08 |
| GO:0007409 | axonogenesis | 181/5895 | 1.20E-09 | 4.89E-08 |
| GO:0033002 | muscle cell proliferation | 102/5895 | 1.23E-09 | 4.97E-08 |
| GO:0106106 | cold-induced thermogenesis | 70/5895 | 1.25E-09 | 4.99E-08 |
| GO:0120161 | regulation of cold-induced thermogenesis | 70/5895 | 1.25E-09 | 4.99E-08 |

| | | | | |
|---|---|---|---|---|
| GO:0051961 | negative regulation of nervous system development | 144/5895 | 1.39E-09 | 5.53E-08 |
| GO:0006094 | gluconeogenesis | 44/5895 | 1.53E-09 | 6.01E-08 |
| GO:0044272 | sulfur compound biosynthetic process | 48/5895 | 1.53E-09 | 6.01E-08 |
| GO:1990845 | adaptive thermogenesis | 75/5895 | 1.83E-09 | 7.14E-08 |
| GO:0016236 | macroautophagy | 91/5895 | 1.86E-09 | 7.19E-08 |
| GO:0009410 | response to xenobiotic stimulus | 57/5895 | 2.10E-09 | 8.07E-08 |
| GO:0010632 | regulation of epithelial cell migration | 101/5895 | 2.13E-09 | 8.15E-08 |
| GO:0030258 | lipid modification | 106/5895 | 2.15E-09 | 8.18E-08 |
| GO:0070372 | regulation of ERK1 and ERK2 cascade | 125/5895 | 2.23E-09 | 8.38E-08 |
| GO:0019217 | regulation of fatty acid metabolic process | 52/5895 | 2.23E-09 | 8.38E-08 |
| GO:0072001 | renal system development | 133/5895 | 2.35E-09 | 8.76E-08 |
| GO:0009743 | response to carbohydrate | 90/5895 | 2.45E-09 | 9.11E-08 |
| GO:0071559 | response to transforming growth factor beta | 93/5895 | 2.60E-09 | 9.59E-08 |
| GO:0043255 | regulation of carbohydrate biosynthetic process | 56/5895 | 2.81E-09 | 1.03E-07 |
| GO:0034284 | response to monosaccharide | 85/5895 | 2.92E-09 | 1.06E-07 |
| GO:0071466 | cellular response to xenobiotic stimulus | 54/5895 | 3.20E-09 | 1.16E-07 |
| GO:0048588 | developmental cell growth | 111/5895 | 3.52E-09 | 1.27E-07 |
| GO:0042063 | gliogenesis | 134/5895 | 3.57E-09 | 1.28E-07 |

| | | | | |
|---|---|---|---|---|
| GO:0048738 | cardiac muscle tissue development | 108/5895 | 3.61E-09 | 1.29E-07 |
| GO:0030099 | myeloid cell differentiation | 158/5895 | 3.67E-09 | 1.30E-07 |
| GO:0071326 | cellular response to monosaccharide stimulus | 70/5895 | 3.80E-09 | 1.34E-07 |
| GO:0002573 | myeloid leukocyte differentiation | 95/5895 | 4.72E-09 | 1.66E-07 |
| GO:0033559 | unsaturated fatty acid metabolic process | 60/5895 | 4.79E-09 | 1.67E-07 |
| GO:0006644 | phospholipid metabolic process | 145/5895 | 4.81E-09 | 1.67E-07 |
| GO:0090181 | regulation of cholesterol metabolic process | 25/5895 | 4.86E-09 | 1.68E-07 |
| GO:0006633 | fatty acid biosynthetic process | 69/5895 | 5.08E-09 | 1.74E-07 |
| GO:0009749 | response to glucose | 83/5895 | 5.12E-09 | 1.74E-07 |
| GO:0010721 | negative regulation of cell development | 149/5895 | 5.12E-09 | 1.74E-07 |
| GO:0009746 | response to hexose | 84/5895 | 5.26E-09 | 1.78E-07 |
| GO:0008286 | insulin receptor signaling pathway | 61/5895 | 5.35E-09 | 1.79E-07 |
| GO:0001822 | kidney development | 127/5895 | 5.36E-09 | 1.79E-07 |
| GO:0030336 | negative regulation of cell migration | 114/5895 | 5.55E-09 | 1.84E-07 |
| GO:0042326 | negative regulation of phosphorylation | 175/5895 | 5.61E-09 | 1.85E-07 |
| GO:0045927 | positive regulation of growth | 125/5895 | 5.63E-09 | 1.85E-07 |
| GO:0006805 | xenobiotic metabolic process | 52/5895 | 5.71E-09 | 1.86E-07 |
| GO:1901361 | organic cyclic compound catabolic process | 166/5895 | 5.72E-09 | 1.86E-07 |

| | | | | |
|---|---|---|---|---|
| GO:0022612 | gland morphogenesis | 72/5895 | 6.16E-09 | 1.99E-07 |
| GO:0048545 | response to steroid hormone | 94/5895 | 6.20E-09 | 2.00E-07 |
| GO:0071331 | cellular response to hexose stimulus | 69/5895 | 7.26E-09 | 2.32E-07 |
| GO:0009611 | response to wounding | 181/5895 | 7.29E-09 | 2.32E-07 |
| GO:0051271 | negative regulation of cellular component movement | 129/5895 | 7.93E-09 | 2.51E-07 |
| GO:0045444 | fat cell differentiation | 105/5895 | 7.96E-09 | 2.51E-07 |
| GO:0001659 | temperature homeostasis | 81/5895 | 8.97E-09 | 2.82E-07 |
| GO:0001558 | regulation of cell growth | 164/5895 | 9.02E-09 | 2.82E-07 |
| GO:0071333 | cellular response to glucose stimulus | 68/5895 | 9.69E-09 | 3.01E-07 |
| GO:0045017 | glycerolipid biosynthetic process | 75/5895 | 1.00E-08 | 3.09E-07 |
| GO:0071900 | regulation of protein serine/threonine kinase activity | 165/5895 | 1.04E-08 | 3.20E-07 |
| GO:0071560 | cellular response to transforming growth factor beta stimulus | 90/5895 | 1.05E-08 | 3.23E-07 |
| GO:0016126 | sterol biosynthetic process | 34/5895 | 1.08E-08 | 3.30E-07 |
| GO:0050768 | negative regulation of neurogenesis | 132/5895 | 1.12E-08 | 3.39E-07 |
| GO:0010770 | positive regulation of cell morphogenesis involved in differentiation | 83/5895 | 1.27E-08 | 3.85E-07 |
| GO:0010811 | positive regulation of cell-substrate adhesion | 62/5895 | 1.28E-08 | 3.85E-07 |
| GO:2000146 | negative regulation of cell motility | 117/5895 | 1.35E-08 | 4.03E-07 |
| GO:0032922 | circadian regulation of gene expression | 40/5895 | 1.41E-08 | 4.19E-07 |

| | | | | |
|---|---|---|---|---|
| GO:0043161 | proteasome-mediated ubiquitin-dependent protein catabolic process | 153/5895 | 1.42E-08 | 4.20E-07 |
| GO:0006084 | acetyl-CoA metabolic process | 24/5895 | 1.42E-08 | 4.20E-07 |
| GO:0010498 | proteasomal protein catabolic process | 173/5895 | 1.49E-08 | 4.37E-07 |
| GO:0045913 | positive regulation of carbohydrate metabolic process | 47/5895 | 1.52E-08 | 4.44E-07 |
| GO:0008643 | carbohydrate transport | 70/5895 | 1.54E-08 | 4.47E-07 |
| GO:0072659 | protein localization to plasma membrane | 118/5895 | 1.65E-08 | 4.79E-07 |
| GO:0010594 | regulation of endothelial cell migration | 75/5895 | 1.90E-08 | 5.47E-07 |
| GO:0060419 | heart growth | 58/5895 | 1.91E-08 | 5.47E-07 |
| GO:0030177 | positive regulation of Wnt signaling pathway | 64/5895 | 2.16E-08 | 6.16E-07 |
| GO:0007266 | Rho protein signal transduction | 80/5895 | 2.18E-08 | 6.19E-07 |
| GO:0035023 | regulation of Rho protein signal transduction | 65/5895 | 2.30E-08 | 6.49E-07 |
| GO:0046165 | alcohol biosynthetic process | 65/5895 | 2.30E-08 | 6.49E-07 |
| GO:0016358 | dendrite development | 121/5895 | 2.39E-08 | 6.69E-07 |
| GO:0030178 | negative regulation of Wnt signaling pathway | 73/5895 | 2.44E-08 | 6.80E-07 |
| GO:0015748 | organophosphate ester transport | 52/5895 | 3.29E-08 | 9.13E-07 |
| GO:1905952 | regulation of lipid localization | 74/5895 | 3.45E-08 | 9.54E-07 |
| GO:0072594 | establishment of protein localization to organelle | 158/5895 | 3.48E-08 | 9.59E-07 |
| GO:0042060 | wound healing | 134/5895 | 3.60E-08 | 9.87E-07 |

| | | | | |
|---|---|---|---|---|
| GO:0019915 | lipid storage | 44/5895 | 3.61E-08 | 9.87E-07 |
| GO:1903008 | organelle disassembly | 50/5895 | 3.87E-08 | 1.05E-06 |
| GO:1903034 | regulation of response to wounding | 81/5895 | 3.98E-08 | 1.08E-06 |
| GO:0010506 | regulation of autophagy | 111/5895 | 4.86E-08 | 1.31E-06 |
| GO:0002064 | epithelial cell development | 104/5895 | 5.70E-08 | 1.53E-06 |
| GO:2001233 | regulation of apoptotic signaling pathway | 153/5895 | 6.20E-08 | 1.66E-06 |
| GO:0051235 | maintenance of location | 124/5895 | 6.28E-08 | 1.67E-06 |
| GO:0006111 | regulation of gluconeogenesis | 32/5895 | 6.54E-08 | 1.73E-06 |
| GO:0001503 | ossification | 147/5895 | 6.73E-08 | 1.78E-06 |
| GO:0003018 | vascular process in circulatory system | 93/5895 | 6.79E-08 | 1.78E-06 |
| GO:0043122 | regulation of I-kappaB kinase/NF-kappaB signaling | 80/5895 | 6.95E-08 | 1.82E-06 |
| GO:0007249 | I-kappaB kinase/NF-kappaB signaling | 90/5895 | 6.97E-08 | 1.82E-06 |
| GO:1901293 | nucleoside phosphate biosynthetic process | 88/5895 | 7.05E-08 | 1.83E-06 |
| GO:0051099 | positive regulation of binding | 85/5895 | 7.10E-08 | 1.84E-06 |
| GO:0045860 | positive regulation of protein kinase activity | 161/5895 | 7.29E-08 | 1.88E-06 |
| GO:0022407 | regulation of cell-cell adhesion | 157/5895 | 7.37E-08 | 1.88E-06 |
| GO:0050994 | regulation of lipid catabolic process | 36/5895 | 7.39E-08 | 1.88E-06 |
| GO:0060038 | cardiac muscle cell proliferation | 36/5895 | 7.39E-08 | 1.88E-06 |

| | | | | |
|---|---|---|---|---|
| GO:0042391 | regulation of membrane potential | 164/5895 | 7.51E-08 | 1.90E-06 |
| GO:0003007 | heart morphogenesis | 110/5895 | 7.80E-08 | 1.97E-06 |
| GO:0034504 | protein localization to nucleus | 115/5895 | 8.40E-08 | 2.11E-06 |
| GO:0018212 | peptidyl-tyrosine modification | 125/5895 | 9.10E-08 | 2.28E-06 |
| GO:0035265 | organ growth | 87/5895 | 9.20E-08 | 2.29E-06 |
| GO:0009165 | nucleotide biosynthetic process | 86/5895 | 9.24E-08 | 2.30E-06 |
| GO:0090287 | regulation of cellular response to growth factor stimulus | 111/5895 | 9.45E-08 | 2.34E-06 |
| GO:0008652 | cellular amino acid biosynthetic process | 35/5895 | 9.73E-08 | 2.40E-06 |
| GO:0046777 | protein autophosphorylation | 98/5895 | 1.03E-07 | 2.53E-06 |
| GO:0034446 | substrate adhesion-dependent cell spreading | 52/5895 | 1.10E-07 | 2.69E-06 |
| GO:0006091 | generation of precursor metabolites and energy | 148/5895 | 1.11E-07 | 2.69E-06 |
| GO:0048639 | positive regulation of developmental growth | 90/5895 | 1.16E-07 | 2.81E-06 |
| GO:0050808 | synapse organization | 172/5895 | 1.17E-07 | 2.82E-06 |
| GO:1904375 | regulation of protein localization to cell periphery | 64/5895 | 1.19E-07 | 2.86E-06 |
| GO:1903362 | regulation of cellular protein catabolic process | 102/5895 | 1.19E-07 | 2.86E-06 |
| GO:0006892 | post-Golgi vesicle-mediated transport | 49/5895 | 1.19E-07 | 2.86E-06 |
| GO:0015914 | phospholipid transport | 43/5895 | 1.24E-07 | 2.94E-06 |
| GO:0048872 | homeostasis of number of cells | 126/5895 | 1.31E-07 | 3.10E-06 |

| | | | | |
|---|---|---|---|---|
| GO:0006913 | nucleocytoplasmic transport | 119/5895 | 1.38E-07 | 3.23E-06 |
| GO:0051169 | nuclear transport | 119/5895 | 1.38E-07 | 3.23E-06 |
| GO:0046460 | neutral lipid biosynthetic process | 29/5895 | 1.38E-07 | 3.23E-06 |
| GO:0046463 | acylglycerol biosynthetic process | 29/5895 | 1.38E-07 | 3.23E-06 |
| GO:0098742 | cell-cell adhesion via plasma-membrane adhesion molecules | 87/5895 | 1.54E-07 | 3.59E-06 |
| GO:0010038 | response to metal ion | 108/5895 | 1.59E-07 | 3.69E-06 |
| GO:0001933 | negative regulation of protein phosphorylation | 155/5895 | 1.61E-07 | 3.71E-06 |
| GO:1903320 | regulation of protein modification by small protein conjugation or removal | 97/5895 | 1.68E-07 | 3.87E-06 |
| GO:0010907 | positive regulation of glucose metabolic process | 28/5895 | 1.76E-07 | 4.04E-06 |
| GO:0018108 | peptidyl-tyrosine phosphorylation | 123/5895 | 1.78E-07 | 4.07E-06 |
| GO:0030900 | forebrain development | 145/5895 | 1.80E-07 | 4.10E-06 |
| GO:0016569 | covalent chromatin modification | 166/5895 | 1.84E-07 | 4.17E-06 |
| GO:0046323 | glucose import | 43/5895 | 1.94E-07 | 4.38E-06 |
| GO:0019369 | arachidonic acid metabolic process | 38/5895 | 2.03E-07 | 4.58E-06 |
| GO:0032102 | negative regulation of response to external stimulus | 142/5895 | 2.06E-07 | 4.62E-06 |
| GO:0001952 | regulation of cell-matrix adhesion | 56/5895 | 2.18E-07 | 4.88E-06 |
| GO:0050810 | regulation of steroid biosynthetic process | 41/5895 | 2.21E-07 | 4.93E-06 |
| GO:0051091 | positive regulation of DNA-binding transcription factor activity | 103/5895 | 2.28E-07 | 5.08E-06 |

| | | | | |
|---|---|---|---|---|
| GO:0055017 | cardiac muscle tissue growth | 52/5895 | 2.36E-07 | 5.22E-06 |
| GO:1904659 | glucose transmembrane transport | 53/5895 | 2.55E-07 | 5.62E-06 |
| GO:0050679 | positive regulation of epithelial cell proliferation | 86/5895 | 2.58E-07 | 5.66E-06 |
| GO:0048771 | tissue remodeling | 82/5895 | 2.68E-07 | 5.87E-06 |
| GO:0051052 | regulation of DNA metabolic process | 140/5895 | 2.69E-07 | 5.87E-06 |
| GO:0015749 | monosaccharide transmembrane transport | 54/5895 | 2.73E-07 | 5.94E-06 |
| GO:0070936 | protein K48-linked ubiquitination | 35/5895 | 2.88E-07 | 6.23E-06 |
| GO:0042738 | exogenous drug catabolic process | 31/5895 | 2.88E-07 | 6.23E-06 |
| GO:1901888 | regulation of cell junction assembly | 88/5895 | 3.20E-07 | 6.89E-06 |
| GO:0007162 | negative regulation of cell adhesion | 112/5895 | 3.25E-07 | 6.97E-06 |
| GO:0002040 | sprouting angiogenesis | 57/5895 | 3.26E-07 | 6.97E-06 |
| GO:0060538 | skeletal muscle organ development | 85/5895 | 3.35E-07 | 7.13E-06 |
| GO:0043409 | negative regulation of MAPK cascade | 74/5895 | 3.56E-07 | 7.55E-06 |
| GO:0048813 | dendrite morphogenesis | 77/5895 | 3.57E-07 | 7.55E-06 |
| GO:0045665 | negative regulation of neuron differentiation | 102/5895 | 3.64E-07 | 7.68E-06 |
| GO:0008645 | hexose transmembrane transport | 53/5895 | 3.65E-07 | 7.68E-06 |
| GO:0034219 | carbohydrate transmembrane transport | 54/5895 | 3.89E-07 | 8.15E-06 |
| GO:0031330 | negative regulation of cellular catabolic process | 99/5895 | 3.97E-07 | 8.28E-06 |

| | | | | |
|---|---|---|---|---|
| GO:0030072 | peptide hormone secretion | 110/5895 | 4.27E-07 | 8.89E-06 |
| GO:0150115 | cell-substrate junction organization | 46/5895 | 4.31E-07 | 8.94E-06 |
| GO:0043405 | regulation of MAP kinase activity | 115/5895 | 4.35E-07 | 9.00E-06 |
| GO:0030856 | regulation of epithelial cell differentiation | 68/5895 | 4.51E-07 | 9.30E-06 |
| GO:0090090 | negative regulation of canonical Wnt signaling pathway | 58/5895 | 4.75E-07 | 9.77E-06 |
| GO:0009064 | glutamine family amino acid metabolic process | 35/5895 | 4.82E-07 | 9.87E-06 |
| GO:0042737 | drug catabolic process | 31/5895 | 5.13E-07 | 1.05E-05 |
| GO:0022411 | cellular component disassembly | 140/5895 | 5.39E-07 | 1.10E-05 |
| GO:0050680 | negative regulation of epithelial cell proliferation | 65/5895 | 5.78E-07 | 1.17E-05 |
| GO:0043534 | blood vessel endothelial cell migration | 55/5895 | 5.79E-07 | 1.17E-05 |
| GO:0014013 | regulation of gliogenesis | 67/5895 | 5.94E-07 | 1.20E-05 |
| GO:1903828 | negative regulation of cellular protein localization | 56/5895 | 6.06E-07 | 1.22E-05 |
| GO:1905954 | positive regulation of lipid localization | 50/5895 | 6.09E-07 | 1.22E-05 |
| GO:0007179 | transforming growth factor beta receptor signaling pathway | 72/5895 | 6.12E-07 | 1.22E-05 |
| GO:0008654 | phospholipid biosynthetic process | 73/5895 | 6.12E-07 | 1.22E-05 |
| GO:0034754 | cellular hormone metabolic process | 51/5895 | 6.52E-07 | 1.29E-05 |
| GO:0007030 | Golgi organization | 58/5895 | 6.56E-07 | 1.30E-05 |
| GO:0072576 | liver morphogenesis | 22/5895 | 6.58E-07 | 1.30E-05 |

| | | | | |
|---|---|---|---|---|
| GO:0017144 | drug metabolic process | 37/5895 | 7.11E-07 | 1.40E-05 |
| GO:0019439 | aromatic compound catabolic process | 148/5895 | 7.17E-07 | 1.40E-05 |
| GO:0099173 | postsynapse organization | 82/5895 | 7.30E-07 | 1.43E-05 |
| GO:0009914 | hormone transport | 136/5895 | 7.70E-07 | 1.50E-05 |
| GO:0009755 | hormone-mediated signaling pathway | 69/5895 | 7.99E-07 | 1.55E-05 |
| GO:0050730 | regulation of peptidyl-tyrosine phosphorylation | 102/5895 | 8.48E-07 | 1.64E-05 |
| GO:0035773 | insulin secretion involved in cellular response to glucose stimulus | 45/5895 | 8.55E-07 | 1.65E-05 |
| GO:0006979 | response to oxidative stress | 147/5895 | 8.85E-07 | 1.70E-05 |
| GO:0006690 | icosanoid metabolic process | 58/5895 | 9.01E-07 | 1.72E-05 |
| GO:0019432 | triglyceride biosynthetic process | 24/5895 | 9.03E-07 | 1.72E-05 |
| GO:0045940 | positive regulation of steroid metabolic process | 24/5895 | 9.03E-07 | 1.72E-05 |
| GO:0034599 | cellular response to oxidative stress | 106/5895 | 9.04E-07 | 1.72E-05 |
| GO:0045165 | cell fate commitment | 111/5895 | 9.12E-07 | 1.73E-05 |
| GO:0060541 | respiratory system development | 99/5895 | 9.37E-07 | 1.77E-05 |
| GO:0000422 | autophagy of mitochondrion | 36/5895 | 9.52E-07 | 1.79E-05 |
| GO:0061726 | mitochondrion disassembly | 36/5895 | 9.52E-07 | 1.79E-05 |
| GO:0030073 | insulin secretion | 91/5895 | 9.58E-07 | 1.79E-05 |
| GO:0034248 | regulation of cellular amide metabolic process | 147/5895 | 1.04E-06 | 1.94E-05 |

| | | | | |
|---|---|---|---|---|
| GO:0046326 | positive regulation of glucose import | 26/5895 | 1.08E-06 | 2.00E-05 |
| GO:0050770 | regulation of axonogenesis | 84/5895 | 1.13E-06 | 2.10E-05 |
| GO:0043470 | regulation of carbohydrate catabolic process | 32/5895 | 1.14E-06 | 2.11E-05 |
| GO:0007044 | cell-substrate junction assembly | 44/5895 | 1.15E-06 | 2.12E-05 |
| GO:0062197 | cellular response to chemical stress | 123/5895 | 1.16E-06 | 2.14E-05 |
| GO:0010883 | regulation of lipid storage | 28/5895 | 1.17E-06 | 2.14E-05 |
| GO:0007178 | transmembrane receptor protein serine/threonine kinase signaling pathway | 134/5895 | 1.19E-06 | 2.17E-05 |
| GO:0031647 | regulation of protein stability | 109/5895 | 1.20E-06 | 2.18E-05 |
| GO:0042752 | regulation of circadian rhythm | 58/5895 | 1.23E-06 | 2.23E-05 |
| GO:0001654 | eye development | 149/5895 | 1.29E-06 | 2.34E-05 |
| GO:1901343 | negative regulation of vasculature development | 61/5895 | 1.31E-06 | 2.36E-05 |
| GO:0061572 | actin filament bundle organization | 72/5895 | 1.36E-06 | 2.43E-05 |
| GO:0060021 | roof of mouth development | 46/5895 | 1.36E-06 | 2.43E-05 |
| GO:0006898 | receptor-mediated endocytosis | 94/5895 | 1.36E-06 | 2.43E-05 |
| GO:0051017 | actin filament bundle assembly | 71/5895 | 1.36E-06 | 2.44E-05 |
| GO:0071902 | positive regulation of protein serine/threonine kinase activity | 110/5895 | 1.39E-06 | 2.48E-05 |
| GO:0030301 | cholesterol transport | 42/5895 | 1.40E-06 | 2.48E-05 |
| GO:0009074 | aromatic amino acid family catabolic process | 15/5895 | 1.43E-06 | 2.51E-05 |

| | | | | |
|---|---|---|---|---|
| GO:0032239 | regulation of nucleobase-containing compound transport | 15/5895 | 1.43E-06 | 2.51E-05 |
| GO:0045540 | regulation of cholesterol biosynthetic process | 15/5895 | 1.43E-06 | 2.51E-05 |
| GO:0106118 | regulation of sterol biosynthetic process | 15/5895 | 1.43E-06 | 2.51E-05 |
| GO:0031098 | stress-activated protein kinase signaling cascade | 104/5895 | 1.45E-06 | 2.54E-05 |
| GO:0031396 | regulation of protein ubiquitination | 83/5895 | 1.46E-06 | 2.56E-05 |
| GO:2001234 | negative regulation of apoptotic signaling pathway | 91/5895 | 1.49E-06 | 2.59E-05 |
| GO:0048880 | sensory system development | 151/5895 | 1.59E-06 | 2.77E-05 |
| GO:0045732 | positive regulation of protein catabolic process | 88/5895 | 1.62E-06 | 2.81E-05 |
| GO:0003279 | cardiac septum development | 57/5895 | 1.63E-06 | 2.81E-05 |
| GO:0001763 | morphogenesis of a branching structure | 94/5895 | 1.68E-06 | 2.89E-05 |
| GO:0007369 | gastrulation | 73/5895 | 1.73E-06 | 2.97E-05 |
| GO:0010977 | negative regulation of neuron projection development | 73/5895 | 1.73E-06 | 2.97E-05 |
| GO:0018105 | peptidyl-serine phosphorylation | 122/5895 | 1.74E-06 | 2.97E-05 |
| GO:0051403 | stress-activated MAPK cascade | 99/5895 | 1.74E-06 | 2.97E-05 |
| GO:0007034 | vacuolar transport | 61/5895 | 1.75E-06 | 2.97E-05 |
| GO:0008206 | bile acid metabolic process | 21/5895 | 1.75E-06 | 2.97E-05 |
| GO:0060977 | coronary vasculature morphogenesis | 21/5895 | 1.75E-06 | 2.97E-05 |
| GO:0061041 | regulation of wound healing | 63/5895 | 1.78E-06 | 3.00E-05 |

| | | | | |
|---|---|---|---|---|
| GO:0070302 | regulation of stress-activated protein kinase signaling cascade | 92/5895 | 1.79E-06 | 3.00E-05 |
| GO:0090276 | regulation of peptide hormone secretion | 92/5895 | 1.79E-06 | 3.00E-05 |
| GO:0001935 | endothelial cell proliferation | 64/5895 | 1.79E-06 | 3.00E-05 |
| GO:0046879 | hormone secretion | 132/5895 | 1.82E-06 | 3.04E-05 |
| GO:0007015 | actin filament organization | 154/5895 | 1.85E-06 | 3.09E-05 |
| GO:0120162 | positive regulation of cold-induced thermogenesis | 46/5895 | 1.95E-06 | 3.23E-05 |
| GO:1903706 | regulation of hemopoiesis | 150/5895 | 1.95E-06 | 3.23E-05 |
| GO:0030307 | positive regulation of cell growth | 80/5895 | 1.97E-06 | 3.25E-05 |
| GO:1990138 | neuron projection extension | 80/5895 | 1.97E-06 | 3.25E-05 |
| GO:0010676 | positive regulation of cellular carbohydrate metabolic process | 35/5895 | 2.02E-06 | 3.32E-05 |
| GO:0015918 | sterol transport | 42/5895 | 2.06E-06 | 3.38E-05 |
| GO:0071692 | protein localization to extracellular region | 166/5895 | 2.08E-06 | 3.39E-05 |
| GO:0016570 | histone modification | 157/5895 | 2.14E-06 | 3.49E-05 |
| GO:0070373 | negative regulation of ERK1 and ERK2 cascade | 39/5895 | 2.26E-06 | 3.67E-05 |
| GO:0006865 | amino acid transport | 68/5895 | 2.34E-06 | 3.79E-05 |
| GO:0010508 | positive regulation of autophagy | 62/5895 | 2.35E-06 | 3.79E-05 |
| GO:0071383 | cellular response to steroid hormone stimulus | 67/5895 | 2.35E-06 | 3.80E-05 |
| GO:0010634 | positive regulation of epithelial cell migration | 64/5895 | 2.36E-06 | 3.80E-05 |

| | | | | |
|---|---|---|---|---|
| GO:0150063 | visual system development | 149/5895 | 2.39E-06 | 3.84E-05 |
| GO:1901607 | alpha-amino acid biosynthetic process | 31/5895 | 2.52E-06 | 4.03E-05 |
| GO:0038127 | ERBB signaling pathway | 51/5895 | 2.53E-06 | 4.04E-05 |
| GO:0007519 | skeletal muscle tissue development | 79/5895 | 2.54E-06 | 4.05E-05 |
| GO:1903050 | regulation of proteolysis involved in cellular protein catabolic process | 87/5895 | 2.58E-06 | 4.10E-05 |
| GO:0002791 | regulation of peptide secretion | 144/5895 | 2.64E-06 | 4.19E-05 |
| GO:1904019 | epithelial cell apoptotic process | 53/5895 | 2.71E-06 | 4.29E-05 |
| GO:0045833 | negative regulation of lipid metabolic process | 46/5895 | 2.78E-06 | 4.39E-05 |
| GO:0009306 | protein secretion | 163/5895 | 2.80E-06 | 4.40E-05 |
| GO:0033866 | nucleoside bisphosphate biosynthetic process | 22/5895 | 2.91E-06 | 4.54E-05 |
| GO:0034030 | ribonucleoside bisphosphate biosynthetic process | 22/5895 | 2.91E-06 | 4.54E-05 |
| GO:0034033 | purine nucleoside bisphosphate biosynthetic process | 22/5895 | 2.91E-06 | 4.54E-05 |
| GO:0060562 | epithelial tube morphogenesis | 138/5895 | 3.05E-06 | 4.76E-05 |
| GO:0046324 | regulation of glucose import | 35/5895 | 3.16E-06 | 4.91E-05 |
| GO:0050773 | regulation of dendrite development | 79/5895 | 3.20E-06 | 4.95E-05 |
| GO:0061138 | morphogenesis of a branching epithelium | 87/5895 | 3.20E-06 | 4.95E-05 |
| GO:0035592 | establishment of protein localization to extracellular region | 163/5895 | 3.22E-06 | 4.97E-05 |
| GO:0048754 | branching morphogenesis of an epithelial tube | 75/5895 | 3.50E-06 | 5.38E-05 |

| | | | | |
|---|---|---|---|---|
| GO:0034332 | adherens junction organization | 28/5895 | 3.56E-06 | 5.47E-05 |
| GO:0042149 | cellular response to glucose starvation | 26/5895 | 3.59E-06 | 5.50E-05 |
| GO:0050708 | regulation of protein secretion | 131/5895 | 3.68E-06 | 5.62E-05 |
| GO:0031929 | TOR signaling | 54/5895 | 3.78E-06 | 5.74E-05 |
| GO:0090101 | negative regulation of transmembrane receptor protein serine/threonine kinase signaling pathway | 54/5895 | 3.78E-06 | 5.74E-05 |
| GO:0055074 | calcium ion homeostasis | 169/5895 | 4.08E-06 | 6.19E-05 |
| GO:0051054 | positive regulation of DNA metabolic process | 85/5895 | 4.22E-06 | 6.39E-05 |
| GO:0035337 | fatty-acyl-CoA metabolic process | 13/5895 | 4.27E-06 | 6.43E-05 |
| GO:0046831 | regulation of RNA export from nucleus | 13/5895 | 4.27E-06 | 6.43E-05 |
| GO:0021537 | telencephalon development | 96/5895 | 4.32E-06 | 6.49E-05 |
| GO:0006699 | bile acid biosynthetic process | 15/5895 | 4.36E-06 | 6.53E-05 |
| GO:0045446 | endothelial cell differentiation | 49/5895 | 4.48E-06 | 6.68E-05 |
| GO:0015850 | organic hydroxy compound transport | 95/5895 | 4.49E-06 | 6.68E-05 |
| GO:0034976 | response to endoplasmic reticulum stress | 95/5895 | 4.49E-06 | 6.68E-05 |
| GO:0019373 | epoxygenase P450 pathway | 20/5895 | 4.61E-06 | 6.75E-05 |
| GO:0032373 | positive regulation of sterol transport | 20/5895 | 4.61E-06 | 6.75E-05 |
| GO:0032376 | positive regulation of cholesterol transport | 20/5895 | 4.61E-06 | 6.75E-05 |
| GO:0072574 | hepatocyte proliferation | 20/5895 | 4.61E-06 | 6.75E-05 |

| | | | | |
|---|---|---|---|---|
| GO:0072575 | epithelial cell proliferation involved in liver morphogenesis | 20/5895 | 4.61E-06 | 6.75E-05 |
| GO:0007219 | Notch signaling pathway | 73/5895 | 4.61E-06 | 6.75E-05 |
| GO:1901861 | regulation of muscle tissue development | 73/5895 | 4.61E-06 | 6.75E-05 |
| GO:0030316 | osteoclast differentiation | 50/5895 | 4.64E-06 | 6.77E-05 |
| GO:0016202 | regulation of striated muscle tissue development | 72/5895 | 4.70E-06 | 6.85E-05 |
| GO:0045216 | cell-cell junction organization | 81/5895 | 4.76E-06 | 6.92E-05 |
| GO:1903076 | regulation of protein localization to plasma membrane | 51/5895 | 4.77E-06 | 6.92E-05 |
| GO:0048041 | focal adhesion assembly | 39/5895 | 4.99E-06 | 7.22E-05 |
| GO:0018209 | peptidyl-serine modification | 128/5895 | 5.01E-06 | 7.23E-05 |
| GO:0072522 | purine-containing compound biosynthetic process | 68/5895 | 5.02E-06 | 7.23E-05 |
| GO:2000181 | negative regulation of blood vessel morphogenesis | 55/5895 | 5.16E-06 | 7.42E-05 |
| GO:0016311 | dephosphorylation | 133/5895 | 5.36E-06 | 7.68E-05 |
| GO:0061178 | regulation of insulin secretion involved in cellular response to glucose stimulus | 40/5895 | 5.43E-06 | 7.76E-05 |
| GO:0050796 | regulation of insulin secretion | 76/5895 | 5.44E-06 | 7.76E-05 |
| GO:0032006 | regulation of TOR signaling | 46/5895 | 5.51E-06 | 7.85E-05 |
| GO:0043112 | receptor metabolic process | 83/5895 | 5.56E-06 | 7.90E-05 |
| GO:0043393 | regulation of protein binding | 89/5895 | 5.60E-06 | 7.94E-05 |
| GO:0090092 | regulation of transmembrane receptor protein serine/threonine kinase signaling pathway | 94/5895 | 5.67E-06 | 8.03E-05 |

| | | | | |
|---|---|---|---|---|
| GO:0010893 | positive regulation of steroid biosynthetic process | 19/5895 | 5.73E-06 | 8.06E-05 |
| GO:0090208 | positive regulation of triglyceride metabolic process | 19/5895 | 5.73E-06 | 8.06E-05 |
| GO:0007173 | epidermal growth factor receptor signaling pathway | 47/5895 | 5.75E-06 | 8.06E-05 |
| GO:0010595 | positive regulation of endothelial cell migration | 47/5895 | 5.75E-06 | 8.06E-05 |
| GO:0048634 | regulation of muscle organ development | 73/5895 | 5.83E-06 | 8.16E-05 |
| GO:0003012 | muscle system process | 143/5895 | 5.91E-06 | 8.25E-05 |
| GO:0051057 | positive regulation of small GTPase mediated signal transduction | 37/5895 | 6.09E-06 | 8.48E-05 |
| GO:0030278 | regulation of ossification | 92/5895 | 6.12E-06 | 8.51E-05 |
| GO:0006874 | cellular calcium ion homeostasis | 163/5895 | 6.42E-06 | 8.91E-05 |
| GO:0060249 | anatomical structure homeostasis | 147/5895 | 6.44E-06 | 8.91E-05 |
| GO:0006164 | purine nucleotide biosynthetic process | 66/5895 | 6.61E-06 | 9.13E-05 |
| GO:0046390 | ribose phosphate biosynthetic process | 65/5895 | 6.70E-06 | 9.23E-05 |
| GO:0032259 | methylation | 125/5895 | 6.81E-06 | 9.36E-05 |
| GO:0032872 | regulation of stress-activated MAPK cascade | 89/5895 | 6.85E-06 | 9.40E-05 |
| GO:1900180 | regulation of protein localization to nucleus | 59/5895 | 6.98E-06 | 9.55E-05 |
| GO:0034612 | response to tumor necrosis factor | 75/5895 | 7.00E-06 | 9.55E-05 |
| GO:0021675 | nerve development | 39/5895 | 7.28E-06 | 9.93E-05 |
| GO:0006766 | vitamin metabolic process | 35/5895 | 7.36E-06 | 9.99E-05 |

| | | | | |
|---|---|---|---|---|
| GO:0014009 | glial cell proliferation | 29/5895 | 7.36E-06 | 9.99E-05 |
| GO:0007041 | lysosomal transport | 46/5895 | 7.65E-06 | 0.000103388 |
| GO:0032092 | positive regulation of protein binding | 46/5895 | 7.65E-06 | 0.000103388 |
| GO:0032970 | regulation of actin filament-based process | 143/5895 | 7.92E-06 | 0.000106683 |
| GO:0046849 | bone remodeling | 47/5895 | 7.93E-06 | 0.000106683 |
| GO:0007005 | mitochondrion organization | 164/5895 | 7.96E-06 | 0.000106908 |
| GO:0048017 | inositol lipid-mediated signaling | 68/5895 | 8.18E-06 | 0.000109238 |
| GO:0030518 | intracellular steroid hormone receptor signaling pathway | 48/5895 | 8.18E-06 | 0.000109238 |
| GO:0017038 | protein import | 84/5895 | 8.19E-06 | 0.000109238 |
| GO:0050731 | positive regulation of peptidyl-tyrosine phosphorylation | 77/5895 | 8.28E-06 | 0.000110151 |
| GO:0010001 | glial cell differentiation | 94/5895 | 8.35E-06 | 0.000110863 |
| GO:0046700 | heterocycle catabolic process | 139/5895 | 8.42E-06 | 0.000111567 |
| GO:0035384 | thioester biosynthetic process | 17/5895 | 8.50E-06 | 0.000112232 |
| GO:0071616 | acyl-CoA biosynthetic process | 17/5895 | 8.50E-06 | 0.000112232 |
| GO:0046503 | glycerolipid catabolic process | 33/5895 | 8.78E-06 | 0.000115597 |
| GO:0044403 | symbiotic process | 132/5895 | 8.93E-06 | 0.000117358 |
| GO:0090205 | positive regulation of cholesterol metabolic process | Oct-95 | 8.95E-06 | 0.000117358 |
| GO:0003158 | endothelium development | 53/5895 | 8.99E-06 | 0.000117358 |

| | | | | |
|---|---|---|---|---|
| GO:0016525 | negative regulation of angiogenesis | 53/5895 | 8.99E-06 | 0.000117358 |
| GO:0045598 | regulation of fat cell differentiation | 61/5895 | 9.00E-06 | 0.000117358 |
| GO:0001666 | response to hypoxia | 81/5895 | 9.06E-06 | 0.000117635 |
| GO:1904018 | positive regulation of vasculature development | 81/5895 | 9.06E-06 | 0.000117635 |
| GO:0044270 | cellular nitrogen compound catabolic process | 137/5895 | 9.33E-06 | 0.000120876 |
| GO:0048668 | collateral sprouting | 20/5895 | 9.36E-06 | 0.000121064 |
| GO:0032103 | positive regulation of response to external stimulus | 145/5895 | 9.51E-06 | 0.00012276 |
| GO:0035150 | regulation of tube size | 71/5895 | 9.70E-06 | 0.000124907 |
| GO:0010828 | positive regulation of glucose transmembrane transport | 28/5895 | 9.87E-06 | 0.000126537 |
| GO:0060612 | adipose tissue development | 28/5895 | 9.87E-06 | 0.000126537 |
| GO:1901264 | carbohydrate derivative transport | 34/5895 | 9.90E-06 | 0.000126647 |
| GO:0036293 | response to decreased oxygen levels | 89/5895 | 1.02E-05 | 0.000129893 |
| GO:0062014 | negative regulation of small molecule metabolic process | 45/5895 | 1.02E-05 | 0.000129893 |
| GO:1901616 | organic hydroxy compound catabolic process | 31/5895 | 1.03E-05 | 0.000130959 |
| GO:0006606 | protein import into nucleus | 67/5895 | 1.06E-05 | 0.0001341 |
| GO:2001236 | regulation of extrinsic apoptotic signaling pathway | 67/5895 | 1.06E-05 | 0.0001341 |
| GO:0009072 | aromatic amino acid family metabolic process | 22/5895 | 1.07E-05 | 0.000135004 |
| GO:0016032 | viral process | 112/5895 | 1.09E-05 | 0.00013714 |

| | | | | |
|---|---|---|---|---|
| GO:0050727 | regulation of inflammatory response | 131/5895 | 1.09E-05 | 0.000137686 |
| GO:0006650 | glycerophospholipid metabolic process | 96/5895 | 1.12E-05 | 0.000140435 |
| GO:0009260 | ribonucleotide biosynthetic process | 62/5895 | 1.15E-05 | 0.000144029 |
| GO:0019221 | cytokine-mediated signaling pathway | 130/5895 | 1.15E-05 | 0.000144029 |
| GO:0005976 | polysaccharide metabolic process | 50/5895 | 1.16E-05 | 0.000144241 |
| GO:0043401 | steroid hormone mediated signaling pathway | 50/5895 | 1.16E-05 | 0.000144241 |
| GO:0009152 | purine ribonucleotide biosynthetic process | 59/5895 | 1.18E-05 | 0.000147442 |
| GO:0010827 | regulation of glucose transmembrane transport | 41/5895 | 1.19E-05 | 0.000147592 |
| GO:0050996 | positive regulation of lipid catabolic process | 19/5895 | 1.19E-05 | 0.000147592 |
| GO:0043467 | regulation of generation of precursor metabolites and energy | 57/5895 | 1.20E-05 | 0.000148106 |
| GO:0046822 | regulation of nucleocytoplasmic transport | 54/5895 | 1.20E-05 | 0.000148106 |
| GO:0010959 | regulation of metal ion transport | 147/5895 | 1.30E-05 | 0.000160656 |
| GO:0043535 | regulation of blood vessel endothelial cell migration | 43/5895 | 1.31E-05 | 0.000160656 |
| GO:1904950 | negative regulation of establishment of protein localization | 68/5895 | 1.31E-05 | 0.000160885 |
| GO:0008361 | regulation of cell size | 82/5895 | 1.32E-05 | 0.000162118 |
| GO:0097006 | regulation of plasma lipoprotein particle levels | 33/5895 | 1.33E-05 | 0.000162765 |
| GO:0051896 | regulation of protein kinase B signaling | 67/5895 | 1.34E-05 | 0.000163565 |
| GO:0007416 | synapse assembly | 75/5895 | 1.36E-05 | 0.000165262 |

| | | | | |
|---|---|---|---|---|
| GO:0007031 | peroxisome organization | 21/5895 | 1.39E-05 | 0.000168722 |
| GO:0055090 | acylglycerol homeostasis | 21/5895 | 1.39E-05 | 0.000168722 |
| GO:0070328 | triglyceride homeostasis | 21/5895 | 1.39E-05 | 0.000168722 |
| GO:0031345 | negative regulation of cell projection organization | 80/5895 | 1.42E-05 | 0.000171713 |
| GO:0044264 | cellular polysaccharide metabolic process | 46/5895 | 1.44E-05 | 0.000173952 |
| GO:0048592 | eye morphogenesis | 72/5895 | 1.48E-05 | 0.000178504 |
| GO:0006536 | glutamate metabolic process | 18/5895 | 1.50E-05 | 0.000180517 |
| GO:0090596 | sensory organ morphogenesis | 112/5895 | 1.51E-05 | 0.000181263 |
| GO:0007009 | plasma membrane organization | 49/5895 | 1.53E-05 | 0.000183033 |
| GO:0016052 | carbohydrate catabolic process | 59/5895 | 1.53E-05 | 0.000183033 |
| GO:0035296 | regulation of tube diameter | 70/5895 | 1.57E-05 | 0.000186492 |
| GO:0097746 | regulation of blood vessel diameter | 70/5895 | 1.57E-05 | 0.000186492 |
| GO:0042177 | negative regulation of protein catabolic process | 55/5895 | 1.58E-05 | 0.000187139 |
| GO:0061912 | selective autophagy | 31/5895 | 1.59E-05 | 0.000188391 |
| GO:0045787 | positive regulation of cell cycle | 126/5895 | 1.64E-05 | 0.000194157 |
| GO:0048015 | phosphatidylinositol-mediated signaling | 66/5895 | 1.73E-05 | 0.000204069 |
| GO:0051224 | negative regulation of protein transport | 66/5895 | 1.73E-05 | 0.000204069 |
| GO:0050920 | regulation of chemotaxis | 85/5895 | 1.76E-05 | 0.000206866 |

| | | | | |
|---|---|---|---|---|
| GO:1903670 | regulation of sprouting angiogenesis | 36/5895 | 1.76E-05 | 0.000206872 |
| GO:0000266 | mitochondrial fission | 26/5895 | 1.77E-05 | 0.00020777 |
| GO:0050803 | regulation of synapse structure or activity | 105/5895 | 1.79E-05 | 0.000209795 |
| GO:0042692 | muscle cell differentiation | 138/5895 | 1.82E-05 | 0.000212771 |
| GO:0098656 | anion transmembrane transport | 84/5895 | 1.83E-05 | 0.000213162 |
| GO:0001954 | positive regulation of cell-matrix adhesion | 29/5895 | 1.86E-05 | 0.000216195 |
| GO:0042446 | hormone biosynthetic process | 29/5895 | 1.86E-05 | 0.000216195 |
| GO:1901657 | glycosyl compound metabolic process | 44/5895 | 1.86E-05 | 0.000216195 |
| GO:0046339 | diacylglycerol metabolic process | 17/5895 | 1.88E-05 | 0.000217207 |
| GO:0060976 | coronary vasculature development | 37/5895 | 1.89E-05 | 0.00021853 |
| GO:0032007 | negative regulation of TOR signaling | 24/5895 | 1.90E-05 | 0.000219205 |
| GO:0007033 | vacuole organization | 69/5895 | 2.01E-05 | 0.000231726 |
| GO:0046883 | regulation of hormone secretion | 109/5895 | 2.06E-05 | 0.000236499 |
| GO:0060759 | regulation of response to cytokine stimulus | 51/5895 | 2.07E-05 | 0.000237146 |
| GO:0045807 | positive regulation of endocytosis | 52/5895 | 2.07E-05 | 0.000237153 |
| GO:0030323 | respiratory tube development | 85/5895 | 2.13E-05 | 0.000243575 |
| GO:0060043 | regulation of cardiac muscle cell proliferation | 27/5895 | 2.13E-05 | 0.000243575 |
| GO:0050807 | regulation of synapse organization | 101/5895 | 2.16E-05 | 0.000246485 |

| | | | | |
|---|---|---|---|---|
| GO:0043010 | camera-type eye development | 129/5895 | 2.19E-05 | 0.00024879 |
| GO:0030100 | regulation of endocytosis | 84/5895 | 2.22E-05 | 0.000251325 |
| GO:0030324 | lung development | 84/5895 | 2.22E-05 | 0.000251325 |
| GO:0060420 | regulation of heart growth | 40/5895 | 2.23E-05 | 0.000252053 |
| GO:0043414 | macromolecule methylation | 107/5895 | 2.27E-05 | 0.000256012 |
| GO:0031334 | positive regulation of protein-containing complex assembly | 88/5895 | 2.27E-05 | 0.000256138 |
| GO:0071248 | cellular response to metal ion | 63/5895 | 2.34E-05 | 0.000262794 |
| GO:0007254 | JNK cascade | 77/5895 | 2.38E-05 | 0.000267062 |
| GO:0002065 | columnar/cuboidal epithelial cell differentiation | 61/5895 | 2.43E-05 | 0.00027282 |
| GO:0045862 | positive regulation of proteolysis | 121/5895 | 2.46E-05 | 0.000275321 |
| GO:0070542 | response to fatty acid | 28/5895 | 2.51E-05 | 0.00027943 |
| GO:1902930 | regulation of alcohol biosynthetic process | 28/5895 | 2.51E-05 | 0.00027943 |
| GO:0046461 | neutral lipid catabolic process | 23/5895 | 2.53E-05 | 0.000281351 |
| GO:0046464 | acylglycerol catabolic process | 23/5895 | 2.53E-05 | 0.000281351 |
| GO:0034381 | plasma lipoprotein particle clearance | 21/5895 | 2.55E-05 | 0.000282497 |
| GO:0003205 | cardiac chamber development | 75/5895 | 2.55E-05 | 0.000282715 |
| GO:0043406 | positive regulation of MAP kinase activity | 85/5895 | 2.58E-05 | 0.00028481 |
| GO:0051170 | import into nucleus | 67/5895 | 2.66E-05 | 0.000293146 |

| | | | | |
|---|---|---|---|---|
| GO:0021987 | cerebral cortex development | 47/5895 | 2.67E-05 | 0.00029387 |
| GO:0051259 | protein complex oligomerization | 84/5895 | 2.69E-05 | 0.000295256 |
| GO:0051222 | positive regulation of protein transport | 122/5895 | 2.71E-05 | 0.000297794 |
| GO:0006090 | pyruvate metabolic process | 51/5895 | 2.72E-05 | 0.000297866 |
| GO:0097191 | extrinsic apoptotic signaling pathway | 87/5895 | 2.86E-05 | 0.000312221 |
| GO:0031503 | protein-containing complex localization | 95/5895 | 2.86E-05 | 0.000312221 |
| GO:0048524 | positive regulation of viral process | 39/5895 | 2.98E-05 | 0.000324353 |
| GO:0034764 | positive regulation of transmembrane transport | 90/5895 | 3.01E-05 | 0.000326904 |
| GO:0048593 | camera-type eye morphogenesis | 61/5895 | 3.09E-05 | 0.000335184 |
| GO:1903311 | regulation of mRNA metabolic process | 97/5895 | 3.10E-05 | 0.000335649 |
| GO:1905475 | regulation of protein localization to membrane | 75/5895 | 3.13E-05 | 0.000339115 |
| GO:1902105 | regulation of leukocyte differentiation | 110/5895 | 3.16E-05 | 0.000341594 |
| GO:0061448 | connective tissue development | 100/5895 | 3.18E-05 | 0.000342874 |
| GO:1903364 | positive regulation of cellular protein catabolic process | 59/5895 | 3.22E-05 | 0.000346173 |
| GO:0045542 | positive regulation of cholesterol biosynthetic process | Sep-95 | 3.23E-05 | 0.000346849 |
| GO:0106120 | positive regulation of sterol biosynthetic process | Sep-95 | 3.23E-05 | 0.000346849 |
| GO:0045637 | regulation of myeloid cell differentiation | 84/5895 | 3.24E-05 | 0.000347427 |
| GO:1904035 | regulation of epithelial cell apoptotic process | 42/5895 | 3.29E-05 | 0.000352101 |

| | | | | |
|---|---|---|---|---|
| GO:1903322 | positive regulation of protein modification by small protein conjugation or removal | 57/5895 | 3.33E-05 | 0.000354258 |
| GO:0110053 | regulation of actin filament organization | 99/5895 | 3.33E-05 | 0.000354258 |
| GO:0045722 | positive regulation of gluconeogenesis | 14/5895 | 3.34E-05 | 0.000354258 |
| GO:2000114 | regulation of establishment of cell polarity | 14/5895 | 3.34E-05 | 0.000354258 |
| GO:0000038 | very long-chain fatty acid metabolic process | 20/5895 | 3.36E-05 | 0.000354551 |
| GO:0009069 | serine family amino acid metabolic process | 20/5895 | 3.36E-05 | 0.000354551 |
| GO:0060479 | lung cell differentiation | 20/5895 | 3.36E-05 | 0.000354551 |
| GO:0010771 | negative regulation of cell morphogenesis involved in differentiation | 44/5895 | 3.43E-05 | 0.000362321 |
| GO:0046328 | regulation of JNK cascade | 72/5895 | 3.49E-05 | 0.000367227 |
| GO:0030038 | contractile actin filament bundle assembly | 48/5895 | 3.57E-05 | 0.000374456 |
| GO:0043149 | stress fiber assembly | 48/5895 | 3.57E-05 | 0.000374456 |
| GO:1901214 | regulation of neuron death | 130/5895 | 3.67E-05 | 0.000384514 |
| GO:0043491 | protein kinase B signaling | 75/5895 | 3.83E-05 | 0.000400987 |
| GO:0045926 | negative regulation of growth | 96/5895 | 3.85E-05 | 0.000401771 |
| GO:0043124 | negative regulation of I-kappaB kinase/NF-kappaB signaling | 25/5895 | 3.86E-05 | 0.000402317 |
| GO:0046885 | regulation of hormone biosynthetic process | 17/5895 | 3.87E-05 | 0.000403008 |
| GO:0090288 | negative regulation of cellular response to growth factor stimulus | 61/5895 | 3.90E-05 | 0.000404989 |
| GO:0007411 | axon guidance | 92/5895 | 3.91E-05 | 0.000405517 |

| | | | | |
|---|---|---|---|---|
| GO:1904951 | positive regulation of establishment of protein localization | 126/5895 | 3.93E-05 | 0.000407558 |
| GO:0015718 | monocarboxylic acid transport | 68/5895 | 3.99E-05 | 0.000412327 |
| GO:1903522 | regulation of blood circulation | 91/5895 | 4.09E-05 | 0.000422039 |
| GO:0030902 | hindbrain development | 67/5895 | 4.12E-05 | 0.000422039 |
| GO:0060411 | cardiac septum morphogenesis | 39/5895 | 4.13E-05 | 0.000422039 |
| GO:1905897 | regulation of response to endoplasmic reticulum stress | 39/5895 | 4.13E-05 | 0.000422039 |
| GO:0006605 | protein targeting | 98/5895 | 4.13E-05 | 0.000422039 |
| GO:0010452 | histone H3-K36 methylation | Oct-95 | 4.14E-05 | 0.000422039 |
| GO:0035376 | sterol import | Oct-95 | 4.14E-05 | 0.000422039 |
| GO:0070508 | cholesterol import | Oct-95 | 4.14E-05 | 0.000422039 |
| GO:0051346 | negative regulation of hydrolase activity | 138/5895 | 4.14E-05 | 0.000422039 |
| GO:0060485 | mesenchyme development | 101/5895 | 4.21E-05 | 0.000428365 |
| GO:0048608 | reproductive structure development | 159/5895 | 4.22E-05 | 0.000429154 |
| GO:0009083 | branched-chain amino acid catabolic process | Dec-95 | 4.25E-05 | 0.000429845 |
| GO:0098974 | postsynaptic actin cytoskeleton organization | Dec-95 | 4.25E-05 | 0.000429845 |
| GO:0099188 | postsynaptic cytoskeleton organization | Dec-95 | 4.25E-05 | 0.000429845 |
| GO:0021955 | central nervous system neuron axonogenesis | 23/5895 | 4.26E-05 | 0.000430317 |
| GO:0090630 | activation of GTPase activity | 41/5895 | 4.37E-05 | 0.00044023 |

| | | | | |
|---|---|---|---|---|
| GO:0061614 | pri-miRNA transcription by RNA polymerase II | 29/5895 | 4.38E-05 | 0.00044023 |
| GO:1902001 | fatty acid transmembrane transport | Nov-95 | 4.41E-05 | 0.000443404 |
| GO:0035264 | multicellular organism growth | 81/5895 | 4.44E-05 | 0.000445131 |
| GO:0002761 | regulation of myeloid leukocyte differentiation | 54/5895 | 4.47E-05 | 0.000447223 |
| GO:0009066 | aspartate family amino acid metabolic process | 21/5895 | 4.49E-05 | 0.00044825 |
| GO:0016485 | protein processing | 85/5895 | 4.49E-05 | 0.00044825 |
| GO:0031214 | biomineral tissue development | 64/5895 | 4.51E-05 | 0.000448518 |
| GO:0110148 | biomineralization | 64/5895 | 4.51E-05 | 0.000448518 |
| GO:0061458 | reproductive system development | 160/5895 | 4.52E-05 | 0.000448968 |
| GO:0061387 | regulation of extent of cell growth | 53/5895 | 4.53E-05 | 0.000448968 |
| GO:0002762 | negative regulation of myeloid leukocyte differentiation | 26/5895 | 4.55E-05 | 0.000449918 |
| GO:0022409 | positive regulation of cell-cell adhesion | 96/5895 | 4.55E-05 | 0.000449918 |
| GO:1902414 | protein localization to cell junction | 52/5895 | 4.58E-05 | 0.000452038 |
| GO:2000058 | regulation of ubiquitin-dependent protein catabolic process | 63/5895 | 4.64E-05 | 0.000457068 |
| GO:0097485 | neuron projection guidance | 92/5895 | 4.64E-05 | 0.000457068 |
| GO:0007156 | homophilic cell adhesion via plasma membrane adhesion molecules | 47/5895 | 4.69E-05 | 0.000461181 |
| GO:2000027 | regulation of animal organ morphogenesis | 79/5895 | 4.82E-05 | 0.000473407 |
| GO:0009116 | nucleoside metabolic process | 35/5895 | 4.84E-05 | 0.000473829 |

| | | | | |
|---|---|---|---|---|
| GO:0032535 | regulation of cellular component size | 140/5895 | 4.84E-05 | 0.000473829 |
| GO:0032878 | regulation of establishment or maintenance of cell polarity | 16/5895 | 4.92E-05 | 0.000479481 |
| GO:0090114 | COPII-coated vesicle budding | 16/5895 | 4.92E-05 | 0.000479481 |
| GO:0003231 | cardiac ventricle development | 59/5895 | 5.15E-05 | 0.000501439 |
| GO:0010464 | regulation of mesenchymal cell proliferation | 24/5895 | 5.19E-05 | 0.000503594 |
| GO:0060421 | positive regulation of heart growth | 24/5895 | 5.19E-05 | 0.000503594 |
| GO:0032371 | regulation of sterol transport | 27/5895 | 5.23E-05 | 0.00050607 |
| GO:0032374 | regulation of cholesterol transport | 27/5895 | 5.23E-05 | 0.00050607 |
| GO:0045766 | positive regulation of angiogenesis | 72/5895 | 5.24E-05 | 0.000506311 |
| GO:0072089 | stem cell proliferation | 58/5895 | 5.27E-05 | 0.000508632 |
| GO:1901863 | positive regulation of muscle tissue development | 38/5895 | 5.50E-05 | 0.000529446 |
| GO:0048675 | axon extension | 56/5895 | 5.51E-05 | 0.000529446 |
| GO:0072331 | signal transduction by p53 class mediator | 55/5895 | 5.61E-05 | 0.000539069 |
| GO:0034103 | regulation of tissue remodeling | 39/5895 | 5.67E-05 | 0.000543618 |
| GO:0006730 | one-carbon metabolic process | 22/5895 | 5.71E-05 | 0.000546815 |
| GO:1902893 | regulation of pri-miRNA transcription by RNA polymerase II | 28/5895 | 5.90E-05 | 0.000563723 |
| GO:0099003 | vesicle-mediated transport in synapse | 94/5895 | 5.92E-05 | 0.000564658 |
| GO:0001702 | gastrulation with mouth forming second | 20/5895 | 5.97E-05 | 0.000569009 |

| | | | | |
|---|---|---|---|---|
| GO:0006110 | regulation of glycolytic process | 25/5895 | 6.13E-05 | 0.000582908 |
| GO:0018205 | peptidyl-lysine modification | 125/5895 | 6.30E-05 | 0.000598651 |
| GO:0051146 | striated muscle cell differentiation | 111/5895 | 6.44E-05 | 0.000609651 |
| GO:1903829 | positive regulation of cellular protein localization | 111/5895 | 6.44E-05 | 0.000609651 |
| GO:0000271 | polysaccharide biosynthetic process | 35/5895 | 6.80E-05 | 0.000643398 |
| GO:0043433 | negative regulation of DNA-binding transcription factor activity | 64/5895 | 6.95E-05 | 0.000656387 |
| GO:0042304 | regulation of fatty acid biosynthetic process | 23/5895 | 6.98E-05 | 0.000658217 |
| GO:0060428 | lung epithelium development | 26/5895 | 7.05E-05 | 0.000664161 |
| GO:0070997 | neuron death | 140/5895 | 7.13E-05 | 0.000669968 |
| GO:0032835 | glomerulus development | 30/5895 | 7.14E-05 | 0.00067045 |
| GO:0006470 | protein dephosphorylation | 86/5895 | 7.30E-05 | 0.000684316 |
| GO:0006417 | regulation of translation | 122/5895 | 7.38E-05 | 0.000691235 |
| GO:0035051 | cardiocyte differentiation | 68/5895 | 7.44E-05 | 0.000695308 |
| GO:0015931 | nucleobase-containing compound transport | 73/5895 | 7.47E-05 | 0.00069708 |
| GO:0060039 | pericardium development | 17/5895 | 7.53E-05 | 0.000700876 |
| GO:0060384 | innervation | 17/5895 | 7.53E-05 | 0.000700876 |
| GO:0045601 | regulation of endothelial cell differentiation | 21/5895 | 7.65E-05 | 0.000710322 |
| GO:0031998 | regulation of fatty acid beta-oxidation | 14/5895 | 7.67E-05 | 0.000710322 |

| | | | | |
|---|---|---|---|---|
| GO:0051963 | regulation of synapse assembly | 51/5895 | 7.68E-05 | 0.000710322 |
| GO:1905039 | carboxylic acid transmembrane transport | 51/5895 | 7.68E-05 | 0.000710322 |
| GO:0060487 | lung epithelial cell differentiation | 19/5895 | 7.93E-05 | 0.000732463 |
| GO:0048285 | organelle fission | 156/5895 | 8.01E-05 | 0.000738869 |
| GO:0070374 | positive regulation of ERK1 and ERK2 cascade | 80/5895 | 8.04E-05 | 0.000740873 |
| GO:0001959 | regulation of cytokine-mediated signaling pathway | 46/5895 | 8.08E-05 | 0.000742927 |
| GO:0002262 | myeloid cell homeostasis | 71/5895 | 8.09E-05 | 0.000742927 |
| GO:0106027 | neuron projection organization | 45/5895 | 8.11E-05 | 0.000742927 |
| GO:0055024 | regulation of cardiac muscle tissue development | 44/5895 | 8.11E-05 | 0.000742927 |
| GO:0046579 | positive regulation of Ras protein signal transduction | 32/5895 | 8.22E-05 | 0.000751451 |
| GO:0032881 | regulation of polysaccharide metabolic process | 24/5895 | 8.26E-05 | 0.000754017 |
| GO:0030217 | T cell differentiation | 100/5895 | 8.35E-05 | 0.000761058 |
| GO:0006397 | mRNA processing | 153/5895 | 8.36E-05 | 0.000761314 |
| GO:0043254 | regulation of protein-containing complex assembly | 146/5895 | 8.49E-05 | 0.000771908 |
| GO:0048660 | regulation of smooth muscle cell proliferation | 64/5895 | 8.58E-05 | 0.000777807 |
| GO:0051147 | regulation of muscle cell differentiation | 64/5895 | 8.58E-05 | 0.000777807 |
| GO:0010463 | mesenchymal cell proliferation | 28/5895 | 8.81E-05 | 0.000797505 |
| GO:0043583 | ear development | 89/5895 | 8.89E-05 | 0.00080365 |

| | | | | |
|---|---|---|---|---|
| GO:2000377 | regulation of reactive oxygen species metabolic process | 77/5895 | 9.18E-05 | 0.000829006 |
| GO:0009791 | post-embryonic development | 54/5895 | 9.22E-05 | 0.000831226 |
| GO:0032352 | positive regulation of hormone metabolic process | 13/5895 | 9.35E-05 | 0.000841353 |
| GO:0030225 | macrophage differentiation | 22/5895 | 9.39E-05 | 0.000842901 |
| GO:0046164 | alcohol catabolic process | 22/5895 | 9.39E-05 | 0.000842901 |
| GO:0098727 | maintenance of cell number | 67/5895 | 9.45E-05 | 0.000846655 |
| GO:0060324 | face development | 29/5895 | 9.61E-05 | 0.000860416 |
| GO:0008053 | mitochondrial fusion | 16/5895 | 9.78E-05 | 0.00087419 |
| GO:1903825 | organic acid transmembrane transport | 51/5895 | 9.82E-05 | 0.000875865 |
| GO:0045844 | positive regulation of striated muscle tissue development | 37/5895 | 0.000100342 | 0.000892889 |
| GO:0048636 | positive regulation of muscle organ development | 37/5895 | 0.000100342 | 0.000892889 |
| GO:0031099 | regeneration | 59/5895 | 0.000101008 | 0.000897551 |
| GO:0003281 | ventricular septum development | 38/5895 | 0.000102475 | 0.000906891 |
| GO:0097581 | lamellipodium organization | 38/5895 | 0.000102475 | 0.000906891 |
| GO:0071398 | cellular response to fatty acid | 20/5895 | 0.000102491 | 0.000906891 |
| GO:1903708 | positive regulation of hemopoiesis | 82/5895 | 0.000104577 | 0.000924059 |
| GO:0008299 | isoprenoid biosynthetic process | 18/5895 | 0.000105156 | 0.00092706 |
| GO:0032231 | regulation of actin filament bundle assembly | 46/5895 | 0.000105211 | 0.00092706 |

| | | | | |
|---|---|---|---|---|
| GO:0007229 | integrin-mediated signaling pathway | 41/5895 | 0.00010621 | 0.000933258 |
| GO:0021954 | central nervous system neuron development | 41/5895 | 0.00010621 | 0.000933258 |
| GO:0001837 | epithelial to mesenchymal transition | 57/5895 | 0.000107189 | 0.000940553 |
| GO:0019058 | viral life cycle | 81/5895 | 0.000109554 | 0.000959963 |
| GO:0032350 | regulation of hormone metabolic process | 23/5895 | 0.000111397 | 0.00097476 |
| GO:0043903 | regulation of symbiotic process | 80/5895 | 0.000114746 | 0.001002675 |
| GO:0045600 | positive regulation of fat cell differentiation | 32/5895 | 0.000116375 | 0.001014105 |
| GO:1904377 | positive regulation of protein localization to cell periphery | 32/5895 | 0.000116375 | 0.001014105 |
| GO:0031348 | negative regulation of defense response | 83/5895 | 0.000118485 | 0.001031066 |
| GO:0002042 | cell migration involved in sprouting angiogenesis | 27/5895 | 0.000118806 | 0.001031414 |
| GO:0061136 | regulation of proteasomal protein catabolic process | 71/5895 | 0.000118852 | 0.001031414 |
| GO:0051260 | protein homooligomerization | 66/5895 | 0.000119848 | 0.001038629 |
| GO:0048762 | mesenchymal cell differentiation | 82/5895 | 0.000124233 | 0.00107515 |
| GO:0048659 | smooth muscle cell proliferation | 65/5895 | 0.000124541 | 0.001076342 |
| GO:0006739 | NADP metabolic process | 21/5895 | 0.000126429 | 0.001091165 |
| GO:0021953 | central nervous system neuron differentiation | 85/5895 | 0.000127409 | 0.00109717 |
| GO:0021932 | hindbrain radial glia guided cell migration | Nov-95 | 0.000127473 | 0.00109717 |
| GO:0120163 | negative regulation of cold-induced thermogenesis | 24/5895 | 0.000128502 | 0.001104298 |

| | | | | |
|---|---|---|---|---|
| GO:0006479 | protein methylation | 69/5895 | 0.000129174 | 0.001104298 |
| GO:0008213 | protein alkylation | 69/5895 | 0.000129174 | 0.001104298 |
| GO:0051100 | negative regulation of binding | 64/5895 | 0.00012935 | 0.001104298 |
| GO:0009100 | glycoprotein metabolic process | 114/5895 | 0.00012948 | 0.001104298 |
| GO:0030282 | bone mineralization | 49/5895 | 0.000129864 | 0.001104298 |
| GO:0005977 | glycogen metabolic process | 35/5895 | 0.000130226 | 0.001104298 |
| GO:0006073 | cellular glucan metabolic process | 35/5895 | 0.000130226 | 0.001104298 |
| GO:0044042 | glucan metabolic process | 35/5895 | 0.000130226 | 0.001104298 |
| GO:0061180 | mammary gland epithelium development | 35/5895 | 0.000130226 | 0.001104298 |
| GO:1903036 | positive regulation of response to wounding | 35/5895 | 0.000130226 | 0.001104298 |
| GO:0048814 | regulation of dendrite morphogenesis | 48/5895 | 0.000132148 | 0.001117587 |
| GO:1901890 | positive regulation of cell junction assembly | 48/5895 | 0.000132148 | 0.001117587 |
| GO:0033157 | regulation of intracellular protein transport | 84/5895 | 0.000133713 | 0.001129312 |
| GO:0030879 | mammary gland development | 63/5895 | 0.000134272 | 0.001132515 |
| GO:0003151 | outflow tract morphogenesis | 37/5895 | 0.000136023 | 0.001145754 |
| GO:0032147 | activation of protein kinase activity | 87/5895 | 0.000136282 | 0.001146405 |
| GO:0032386 | regulation of intracellular transport | 113/5895 | 0.000136533 | 0.001146988 |
| GO:0006568 | tryptophan metabolic process | Sep-95 | 0.000137524 | 0.001151604 |

| | | | | |
|---|---|---|---|---|
| GO:0006586 | indolalkylamine metabolic process | Sep-95 | 0.000137524 | 0.001151604 |
| GO:0060840 | artery development | 45/5895 | 0.000137637 | 0.001151604 |
| GO:0072593 | reactive oxygen species metabolic process | 102/5895 | 0.000137813 | 0.001151604 |
| GO:0001819 | positive regulation of cytokine production | 148/5895 | 0.000138411 | 0.001152462 |
| GO:0010635 | regulation of mitochondrial fusion | Oct-95 | 0.000138463 | 0.001152462 |
| GO:0071281 | cellular response to iron ion | Oct-95 | 0.000138463 | 0.001152462 |
| GO:0090140 | regulation of mitochondrial fission | 17/5895 | 0.000139193 | 0.001157005 |
| GO:1902903 | regulation of supramolecular fiber organization | 125/5895 | 0.00014014 | 0.001163343 |
| GO:0031016 | pancreas development | 41/5895 | 0.000140508 | 0.001164865 |
| GO:0072511 | divalent inorganic cation transport | 154/5895 | 0.000141611 | 0.001172474 |
| GO:0001890 | placenta development | 66/5895 | 0.000145934 | 0.001206681 |
| GO:0048008 | platelet-derived growth factor receptor signaling pathway | 30/5895 | 0.00014814 | 0.001223319 |
| GO:0070838 | divalent metal ion transport | 153/5895 | 0.000149414 | 0.001232229 |
| GO:0045931 | positive regulation of mitotic cell cycle | 65/5895 | 0.00015186 | 0.001250759 |
| GO:0032368 | regulation of lipid transport | 52/5895 | 0.000153947 | 0.001266301 |
| GO:0051101 | regulation of DNA binding | 51/5895 | 0.000157835 | 0.001296591 |
| GO:0030514 | negative regulation of BMP signaling pathway | 26/5895 | 0.000160291 | 0.001313342 |
| GO:0031103 | axon regeneration | 26/5895 | 0.000160291 | 0.001313342 |

| | | | | |
|---|---|---|---|---|
| GO:1903052 | positive regulation of proteolysis involved in cellular protein catabolic process | 50/5895 | 0.000161605 | 0.001322275 |
| GO:0044273 | sulfur compound catabolic process | 14/5895 | 0.0001618 | 0.001322275 |
| GO:0030098 | lymphocyte differentiation | 136/5895 | 0.000164172 | 0.001339925 |
| GO:0032956 | regulation of actin cytoskeleton organization | 122/5895 | 0.000164607 | 0.001341735 |
| GO:0045739 | positive regulation of DNA repair | 33/5895 | 0.000168645 | 0.00137134 |
| GO:0018107 | peptidyl-threonine phosphorylation | 48/5895 | 0.000168673 | 0.00137134 |
| GO:0032409 | regulation of transporter activity | 98/5895 | 0.000169644 | 0.001377459 |
| GO:2000781 | positive regulation of double-strand break repair | 20/5895 | 0.000170241 | 0.001380526 |
| GO:0001818 | negative regulation of cytokine production | 92/5895 | 0.000170722 | 0.001382649 |
| GO:0051480 | regulation of cytosolic calcium ion concentration | 126/5895 | 0.000171481 | 0.001386887 |
| GO:0018958 | phenol-containing compound metabolic process | 47/5895 | 0.000171905 | 0.001386887 |
| GO:0031398 | positive regulation of protein ubiquitination | 47/5895 | 0.000171905 | 0.001386887 |
| GO:0061351 | neural precursor cell proliferation | 71/5895 | 0.000172419 | 0.001389262 |
| GO:0045923 | positive regulation of fatty acid metabolic process | 23/5895 | 0.000173615 | 0.001397111 |
| GO:0055021 | regulation of cardiac muscle tissue growth | 35/5895 | 0.000177449 | 0.001426144 |
| GO:0050792 | regulation of viral process | 74/5895 | 0.000180361 | 0.001447006 |
| GO:0032370 | positive regulation of lipid transport | 36/5895 | 0.000180503 | 0.001447006 |
| GO:0050772 | positive regulation of axonogenesis | 43/5895 | 0.000181896 | 0.001454479 |

| | | | | |
|---|---|---|---|---|
| GO:0090263 | positive regulation of canonical Wnt signaling pathway | 43/5895 | 0.000181896 | 0.001454479 |
| GO:1902882 | regulation of response to oxidative stress | 42/5895 | 0.000183435 | 0.001464216 |
| GO:0048745 | smooth muscle tissue development | 16/5895 | 0.0001838<br>1 | 0.001464216 |
| GO:0060045 | positive regulation of cardiac muscle cell proliferation | 16/5895 | 0.0001838<br>1 | 0.001464216 |
| GO:0001570 | vasculogenesis | 41/5895 | 0.000184488 | 0.001465913 |
| GO:1905037 | autophagosome organization | 41/5895 | 0.000184488 | 0.001465913 |
| GO:0010717 | regulation of epithelial to mesenchymal transition | 40/5895 | 0.000185002 | 0.001468146 |
| GO:0045995 | regulation of embryonic development | 53/5895 | 0.000187749 | 0.001488079 |
| GO:0090257 | regulation of muscle system process | 87/5895 | 0.000188019 | 0.001488345 |
| GO:0050817 | coagulation | 64/5895 | 0.000192192 | 0.001517568 |
| GO:0071356 | cellular response to tumor necrosis factor | 64/5895 | 0.000192192 | 0.001517568 |
| GO:0018022 | peptidyl-lysine methylation | 52/5895 | 0.000193211 | 0.001523708 |
| GO:0003382 | epithelial cell morphogenesis | 24/5895 | 0.000195748 | 0.001541791 |
| GO:0006900 | vesicle budding from membrane | 29/5895 | 0.000198966 | 0.001563234 |
| GO:1903078 | positive regulation of protein localization to plasma membrane | 29/5895 | 0.000198966 | 0.001563234 |
| GO:0007596 | blood coagulation | 63/5895 | 0.000200086 | 0.001570077 |
| GO:0006984 | ER-nucleus signaling pathway | 21/5895 | 0.000203037 | 0.001587314 |
| GO:0034105 | positive regulation of tissue remodeling | 21/5895 | 0.000203037 | 0.001587314 |

| | | | | |
|---|---|---|---|---|
| GO:0097009 | energy homeostasis | 21/5895 | 0.000203037 | 0.001587314 |
| GO:0043123 | positive regulation of I-kappaB kinase/NF-kappaB signaling | 50/5895 | 0.000203908 | 0.001588115 |
| GO:0003206 | cardiac chamber morphogenesis | 57/5895 | 0.000204805 | 0.001588115 |
| GO:0008209 | androgen metabolic process | 13/5895 | 0.000205154 | 0.001588115 |
| GO:0009081 | branched-chain amino acid metabolic process | 13/5895 | 0.000205154 | 0.001588115 |
| GO:0010875 | positive regulation of cholesterol efflux | 13/5895 | 0.000205154 | 0.001588115 |
| GO:0019363 | pyridine nucleotide biosynthetic process | 13/5895 | 0.000205154 | 0.001588115 |
| GO:0021535 | cell migration in hindbrain | 13/5895 | 0.000205154 | 0.001588115 |
| GO:0090030 | regulation of steroid hormone biosynthetic process | 13/5895 | 0.000205154 | 0.001588115 |
| GO:0015980 | energy derivation by oxidation of organic compounds | 88/5895 | 0.000209227 | 0.001617665 |
| GO:0007626 | locomotory behavior | 91/5895 | 0.000209486 | 0.001617682 |
| GO:0097193 | intrinsic apoptotic signaling pathway | 102/5895 | 0.000212509 | 0.001639023 |
| GO:0000187 | activation of MAPK activity | 48/5895 | 0.000214085 | 0.001647151 |
| GO:2000278 | regulation of DNA biosynthetic process | 48/5895 | 0.000214085 | 0.001647151 |
| GO:0031663 | lipopolysaccharide-mediated signaling pathway | 25/5895 | 0.000216387 | 0.001662829 |
| GO:0030032 | lamellipodium assembly | 31/5895 | 0.000217805 | 0.001669663 |
| GO:0033692 | cellular polysaccharide biosynthetic process | 31/5895 | 0.000217805 | 0.001669663 |
| GO:0016050 | vesicle organization | 96/5895 | 0.000218192 | 0.001670599 |

| | | | | |
|---|---|---|---|---|
| GO:0009636 | response to toxic substance | 47/5895 | 0.000218883 | 0.001671831 |
| GO:0034968 | histone lysine methylation | 47/5895 | 0.000218883 | 0.001671831 |
| GO:0021543 | pallium development | 60/5895 | 0.000225109 | 0.001715487 |
| GO:0055081 | anion homeostasis | 32/5895 | 0.000225142 | 0.001715487 |
| GO:2001020 | regulation of response to DNA damage stimulus | 80/5895 | 0.000226405 | 0.00172303 |
| GO:2001022 | positive regulation of response to DNA damage stimulus | 45/5895 | 0.00022767 | 0.001730568 |
| GO:1905898 | positive regulation of response to endoplasmic reticulum stress | 19/5895 | 0.000229309 | 0.001740925 |
| GO:0038083 | peptidyl-tyrosine autophosphorylation | 22/5895 | 0.000234731 | 0.00177985 |
| GO:0043488 | regulation of mRNA stability | 43/5895 | 0.000235028 | 0.00177985 |
| GO:1900024 | regulation of substrate adhesion-dependent cell spreading | 26/5895 | 0.000235282 | 0.00177985 |
| GO:0045453 | bone resorption | 34/5895 | 0.000235897 | 0.001782367 |
| GO:2001237 | negative regulation of extrinsic apoptotic signaling pathway | 42/5895 | 0.000238022 | 0.001796272 |
| GO:0034637 | cellular carbohydrate biosynthetic process | 35/5895 | 0.000239454 | 0.001804919 |
| GO:0051492 | regulation of stress fiber assembly | 40/5895 | 0.000242305 | 0.001824225 |
| GO:2001057 | reactive nitrogen species metabolic process | 37/5895 | 0.000243328 | 0.001828447 |
| GO:0000045 | autophagosome assembly | 39/5895 | 0.000243445 | 0.001828447 |
| GO:0006767 | water-soluble vitamin metabolic process | 17/5895 | 0.000245731 | 0.001841231 |
| GO:0019433 | triglyceride catabolic process | 17/5895 | 0.000245731 | 0.001841231 |

| | | | | |
|---|---|---|---|---|
| GO:00156<br>72 | monovalent inorganic cation transport | 158/589<br>5 | 0.0002472<br>21 | 0.0018502<br>01 |
| GO:00435<br>36 | positive regulation of blood vessel endothelial cell migration | 27/5895 | 0.0002522<br>8 | 0.0018835<br>95 |
| GO:00466<br>22 | positive regulation of organ growth | 27/5895 | 0.0002522<br>8 | 0.0018835<br>95 |
| GO:00424<br>48 | progesterone metabolic process | Dec-95 | 0.0002564<br>9 | 0.0019127<br>63 |
| GO:00464<br>74 | glycerophospholipid biosynthetic process | 49/5895 | 0.0002631<br>63 | 0.0019602<br>04 |
| GO:00071<br>63 | establishment or maintenance of cell polarity | 80/5895 | 0.0002666<br>65 | 0.0019839<br>52 |
| GO:00903<br>03 | positive regulation of wound healing | 28/5895 | 0.0002673<br>03 | 0.0019863<br>54 |
| GO:00108<br>21 | regulation of mitochondrion organization | 55/5895 | 0.0002708<br>04 | 0.0020100<br>01 |
| GO:00026<br>83 | negative regulation of immune system process | 150/589<br>5 | 0.0002739<br>75 | 0.0020311<br>51 |
| GO:01501<br>16 | regulation of cell-substrate junction organization | 29/5895 | 0.0002803<br>31 | 0.0020758<br>33 |
| GO:00198<br>27 | stem cell population maintenance | 64/5895 | 0.0002815<br>92 | 0.0020827<br>24 |
| GO:00083<br>80 | RNA splicing | 128/589<br>5 | 0.0002849<br>75 | 0.0021052<br>81 |
| GO:00164<br>82 | cytosolic transport | 59/5895 | 0.0002856<br>43 | 0.0021077<br>47 |
| GO:00551<br>23 | digestive system development | 53/5895 | 0.0002901<br>43 | 0.0021384<br>46 |
| GO:19059<br>53 | negative regulation of lipid localization | 24/5895 | 0.0002922<br>99 | 0.0021518<br>25 |
| GO:00330<br>44 | regulation of chromosome organization | 123/589<br>5 | 0.0002928<br>33 | 0.0021532<br>46 |
| GO:00075<br>99 | hemostasis | 63/5895 | 0.0002939<br>26 | 0.0021587<br>62 |
| GO:00486<br>61 | positive regulation of smooth muscle cell proliferation | 44/5895 | 0.0002961<br>26 | 0.0021723<br>95 |

| | | | | |
|---|---|---|---|---|
| GO:0016239 | positive regulation of macroautophagy | 31/5895 | 0.000300551 | 0.002199738 |
| GO:0048678 | response to axon injury | 31/5895 | 0.000300551 | 0.002199738 |
| GO:0002221 | pattern recognition receptor signaling pathway | 57/5895 | 0.000308795 | 0.002254849 |
| GO:0034101 | erythrocyte homeostasis | 57/5895 | 0.000308795 | 0.002254849 |
| GO:0097061 | dendritic spine organization | 41/5895 | 0.000311228 | 0.002269982 |
| GO:0046626 | regulation of insulin receptor signaling pathway | 33/5895 | 0.00031352 | 0.002279745 |
| GO:0006103 | 2-oxoglutarate metabolic process | Nov-95 | 0.000314381 | 0.002279745 |
| GO:0055089 | fatty acid homeostasis | Nov-95 | 0.000314381 | 0.002279745 |
| GO:0060391 | positive regulation of SMAD protein signal transduction | Nov-95 | 0.000314381 | 0.002279745 |
| GO:0098696 | regulation of neurotransmitter receptor localization to postsynaptic specialization membrane | Nov-95 | 0.000314381 | 0.002279745 |
| GO:0010866 | regulation of triglyceride biosynthetic process | 14/5895 | 0.000317314 | 0.002279745 |
| GO:0034383 | low-density lipoprotein particle clearance | 14/5895 | 0.000317314 | 0.002279745 |
| GO:0045980 | negative regulation of nucleotide metabolic process | 14/5895 | 0.000317314 | 0.002279745 |
| GO:2000345 | regulation of hepatocyte proliferation | 14/5895 | 0.000317314 | 0.002279745 |
| GO:0018023 | peptidyl-lysine trimethylation | 25/5895 | 0.00031738 | 0.002279745 |
| GO:1904036 | negative regulation of epithelial cell apoptotic process | 25/5895 | 0.00031738 | 0.002279745 |
| GO:0019229 | regulation of vasoconstriction | 34/5895 | 0.000317552 | 0.002279745 |
| GO:0008207 | C21-steroid hormone metabolic process | 21/5895 | 0.000317607 | 0.002279745 |

| | | | | |
|---|---|---|---|---|
| GO:0033077 | T cell differentiation in thymus | 39/5895 | 0.000318027 | 0.002279745 |
| GO:0045445 | myoblast differentiation | 39/5895 | 0.000318027 | 0.002279745 |
| GO:0018210 | peptidyl-threonine modification | 50/5895 | 0.000319674 | 0.002279745 |
| GO:0060996 | dendritic spine development | 50/5895 | 0.000319674 | 0.002279745 |
| GO:0006140 | regulation of nucleotide metabolic process | 35/5895 | 0.000320109 | 0.002279745 |
| GO:1904589 | regulation of protein import | 35/5895 | 0.000320109 | 0.002279745 |
| GO:1900076 | regulation of cellular response to insulin stimulus | 38/5895 | 0.000320154 | 0.002279745 |
| GO:1903035 | negative regulation of response to wounding | 38/5895 | 0.000320154 | 0.002279745 |
| GO:0034390 | smooth muscle cell apoptotic process | 16/5895 | 0.000328464 | 0.002333648 |
| GO:0034391 | regulation of smooth muscle cell apoptotic process | 16/5895 | 0.000328464 | 0.002333648 |
| GO:0051153 | regulation of striated muscle cell differentiation | 49/5895 | 0.000329513 | 0.002338471 |
| GO:0030010 | establishment of cell polarity | 55/5895 | 0.000333035 | 0.002360806 |
| GO:0046620 | regulation of organ growth | 48/5895 | 0.000339276 | 0.002400026 |
| GO:0006826 | iron ion transport | 26/5895 | 0.000339709 | 0.002400026 |
| GO:0050775 | positive regulation of dendrite morphogenesis | 26/5895 | 0.000339709 | 0.002400026 |
| GO:0050728 | negative regulation of inflammatory response | 59/5895 | 0.00034753 | 0.002452535 |
| GO:0050919 | negative chemotaxis | 22/5895 | 0.000358065 | 0.002523236 |
| GO:0019751 | polyol metabolic process | 46/5895 | 0.000358348 | 0.002523236 |

| | | | | |
|---|---|---|---|---|
| GO:0048839 | inner ear development | 77/5895 | 0.000364095 | 0.00256084 |
| GO:0048679 | regulation of axon regeneration | 19/5895 | 0.000371047 | 0.002604304 |
| GO:0001936 | regulation of endothelial cell proliferation | 52/5895 | 0.0003711 | 0.002604304 |
| GO:0046500 | S-adenosylmethionine metabolic process | Oct-95 | 0.000374281 | 0.002620796 |
| GO:0046886 | positive regulation of hormone biosynthetic process | Oct-95 | 0.000374281 | 0.002620796 |
| GO:0014896 | muscle hypertrophy | 44/5895 | 0.000376377 | 0.002632555 |
| GO:0031623 | receptor internalization | 51/5895 | 0.000384134 | 0.002680864 |
| GO:0032434 | regulation of proteasomal ubiquitin-dependent protein catabolic process | 51/5895 | 0.000384134 | 0.002680864 |
| GO:0070988 | demethylation | 29/5895 | 0.000389861 | 0.002717826 |
| GO:0003300 | cardiac muscle hypertrophy | 42/5895 | 0.000392726 | 0.002734776 |
| GO:0010639 | negative regulation of organelle organization | 137/5895 | 0.000394681 | 0.00274233 |
| GO:0003179 | heart valve morphogenesis | 23/5895 | 0.000395115 | 0.00274233 |
| GO:0045911 | positive regulation of DNA recombination | 23/5895 | 0.000395115 | 0.00274233 |
| GO:0061013 | regulation of mRNA catabolic process | 50/5895 | 0.000397281 | 0.002754335 |
| GO:0032412 | regulation of ion transmembrane transporter activity | 90/5895 | 0.000400138 | 0.002769836 |
| GO:0043062 | extracellular structure organization | 103/5895 | 0.000400395 | 0.002769836 |
| GO:0006071 | glycerol metabolic process | 13/5895 | 0.000413876 | 0.002853707 |
| GO:0009110 | vitamin biosynthetic process | 13/5895 | 0.000413876 | 0.002853707 |

| | | | | |
|---|---|---|---|---|
| GO:0031290 | retinal ganglion cell axon guidance | 13/5895 | 0.000413876 | 0.002853707 |
| GO:0002092 | positive regulation of receptor internalization | 17/5895 | 0.000416429 | 0.002855706 |
| GO:0006482 | protein demethylation | 17/5895 | 0.000416429 | 0.002855706 |
| GO:0008214 | protein dealkylation | 17/5895 | 0.000416429 | 0.002855706 |
| GO:0032365 | intracellular lipid transport | 17/5895 | 0.000416429 | 0.002855706 |
| GO:0043276 | anoikis | 17/5895 | 0.000416429 | 0.002855706 |
| GO:0030193 | regulation of blood coagulation | 35/5895 | 0.0004240<br>7 | 0.0029049<br>5 |
| GO:0032885 | regulation of polysaccharide biosynthetic process | 20/5895 | 0.0004301<br>1 | 0.002930417 |
| GO:0046320 | regulation of fatty acid oxidation | 20/5895 | 0.0004301<br>1 | 0.002930417 |
| GO:0060251 | regulation of glial cell proliferation | 20/5895 | 0.0004301<br>1 | 0.002930417 |
| GO:0072523 | purine-containing compound catabolic process | 20/5895 | 0.0004301<br>1 | 0.002930417 |
| GO:0090049 | regulation of cell migration involved in sprouting angiogenesis | 20/5895 | 0.0004301<br>1 | 0.002930417 |
| GO:0030325 | adrenal gland development | 15/5895 | 0.000438578 | 0.002984885 |
| GO:0030218 | erythrocyte differentiation | 53/5895 | 0.000440298 | 0.002993365 |
| GO:0052547 | regulation of peptidase activity | 137/5895 | 0.000441128 | 0.002995777 |
| GO:0006000 | fructose metabolic process | Aug-95 | 0.000449005 | 0.003032949 |
| GO:0006651 | diacylglycerol biosynthetic process | Aug-95 | 0.000449005 | 0.003032949 |
| GO:0010745 | negative regulation of macrophage derived foam cell differentiation | Aug-95 | 0.000449005 | 0.003032949 |

| | | | | |
|---|---|---|---|---|
| GO:0060982 | coronary artery morphogenesis | Aug-95 | 0.000449005 | 0.003032949 |
| GO:1900103 | positive regulation of endoplasmic reticulum unfolded protein response | Aug-95 | 0.000449005 | 0.003032949 |
| GO:0043487 | regulation of RNA stability | 46/5895 | 0.000449997 | 0.003036397 |
| GO:0022898 | regulation of transmembrane transporter activity | 93/5895 | 0.000453957 | 0.003059845 |
| GO:0050764 | regulation of phagocytosis | 45/5895 | 0.000462892 | 0.003116739 |
| GO:1902905 | positive regulation of supramolecular fiber organization | 72/5895 | 0.0004667 | 0.0031395 |
| GO:0051604 | protein maturation | 100/5895 | 0.000470442 | 0.003160832 |
| GO:0010469 | regulation of signaling receptor activity | 51/5895 | 0.0004740 6 | 0.0031783 7 |
| GO:0099175 | regulation of postsynapse organization | 51/5895 | 0.0004740 6 | 0.0031783 7 |
| GO:0006275 | regulation of DNA replication | 44/5895 | 0.000475515 | 0.003184742 |
| GO:0055023 | positive regulation of cardiac muscle tissue growth | 21/5895 | 0.000484869 | 0.003243942 |
| GO:0014897 | striated muscle hypertrophy | 43/5895 | 0.000487772 | 0.003256458 |
| GO:0110020 | regulation of actomyosin structure organization | 43/5895 | 0.000487772 | 0.003256458 |
| GO:0043543 | protein acylation | 83/5895 | 0.000498649 | 0.003320932 |
| GO:0048863 | stem cell differentiation | 83/5895 | 0.000498649 | 0.003320932 |
| GO:0045778 | positive regulation of ossification | 42/5895 | 0.000499553 | 0.003320932 |
| GO:0033673 | negative regulation of kinase activity | 80/5895 | 0.000501007 | 0.003320932 |
| GO:0000096 | sulfur amino acid metabolic process | 18/5895 | 0.000502167 | 0.003320932 |

| | | | | |
|---|---|---|---|---|
| GO:0044275 | cellular carbohydrate catabolic process | 18/5895 | 0.000502167 | 0.003320932 |
| GO:0045070 | positive regulation of viral genome replication | 18/5895 | 0.000502167 | 0.003320932 |
| GO:0048194 | Golgi vesicle budding | 18/5895 | 0.000502167 | 0.003320932 |
| GO:1903146 | regulation of autophagy of mitochondrion | 18/5895 | 0.000502167 | 0.003320932 |
| GO:0052548 | regulation of endopeptidase activity | 115/5895 | 0.000509696 | 0.003367041 |
| GO:0071456 | cellular response to hypoxia | 41/5895 | 0.00051074 | 0.003367041 |
| GO:2000060 | positive regulation of ubiquitin-dependent protein catabolic process | 41/5895 | 0.00051074 | 0.003367041 |
| GO:0090068 | positive regulation of cell cycle process | 88/5895 | 0.000513716 | 0.003383128 |
| GO:0060395 | SMAD protein signal transduction | 40/5895 | 0.000521199 | 0.003420929 |
| GO:0006081 | cellular aldehyde metabolic process | 28/5895 | 0.000521625 | 0.003420929 |
| GO:0031102 | neuron projection regeneration | 28/5895 | 0.000521625 | 0.003420929 |
| GO:0046850 | regulation of bone remodeling | 28/5895 | 0.000521625 | 0.003420929 |
| GO:0006814 | sodium ion transport | 82/5895 | 0.000525368 | 0.003441894 |
| GO:0009132 | nucleoside diphosphate metabolic process | 48/5895 | 0.000526489 | 0.003445665 |
| GO:0001569 | branching involved in blood vessel morphogenesis | 22/5895 | 0.000534496 | 0.003478584 |
| GO:0050918 | positive chemotaxis | 22/5895 | 0.000534496 | 0.003478584 |
| GO:0090398 | cellular senescence | 29/5895 | 0.000535482 | 0.003478584 |
| GO:0006656 | phosphatidylcholine biosynthetic process | Dec-95 | 0.00053593 | 0.003478584 |

| | | | | |
|---|---|---|---|---|
| GO:0033145 | positive regulation of intracellular steroid hormone receptor signaling pathway | Dec-95 | 0.00053593 | 0.003478584 |
| GO:0042730 | fibrinolysis | Dec-95 | 0.00053593 | 0.003478584 |
| GO:0044539 | long-chain fatty acid import into cell | Dec-95 | 0.00053593 | 0.003478584 |
| GO:0140354 | lipid import into cell | Dec-95 | 0.00053593 | 0.003478584 |
| GO:0055007 | cardiac muscle cell differentiation | 53/5895 | 0.000538829 | 0.003493807 |
| GO:0045861 | negative regulation of proteolysis | 116/5895 | 0.000544849 | 0.003529195 |
| GO:0015800 | acidic amino acid transport | 30/5895 | 0.000545965 | 0.003529195 |
| GO:1902369 | negative regulation of RNA catabolic process | 30/5895 | 0.000545965 | 0.003529195 |
| GO:0045786 | negative regulation of cell cycle | 145/5895 | 0.000549027 | 0.003545355 |
| GO:0030198 | extracellular matrix organization | 102/5895 | 0.000549661 | 0.003545822 |
| GO:0048708 | astrocyte differentiation | 36/5895 | 0.000552588 | 0.003552443 |
| GO:0050818 | regulation of coagulation | 36/5895 | 0.000552588 | 0.003552443 |
| GO:0001885 | endothelial cell development | 31/5895 | 0.000553331 | 0.003552443 |
| GO:0007492 | endoderm development | 31/5895 | 0.000553331 | 0.003552443 |
| GO:0099504 | synaptic vesicle cycle | 81/5895 | 0.000553503 | 0.003552443 |
| GO:1900046 | regulation of hemostasis | 35/5895 | 0.0005569 | 0.003570614 |
| GO:1900542 | regulation of purine nucleotide metabolic process | 34/5895 | 0.000559375 | 0.003581752 |
| GO:0033143 | regulation of intracellular steroid hormone receptor signaling pathway | 33/5895 | 0.000559773 | 0.003581752 |

| | | | | |
|---|---|---|---|---|
| GO:19030 37 | regulation of leukocyte cell-cell adhesion | 104/589 5 | 0.0005612 91 | 0.0035878 26 |
| GO:00509 21 | positive regulation of chemotaxis | 56/5895 | 0.0005790 48 | 0.0036964 82 |
| GO:20003 79 | positive regulation of reactive oxygen species metabolic process | 45/5895 | 0.0005797 1 | 0.0036964 82 |
| GO:00020 53 | positive regulation of mesenchymal cell proliferation | 19/5895 | 0.0005830 08 | 0.0036964 82 |
| GO:00463 29 | negative regulation of JNK cascade | 19/5895 | 0.0005830 08 | 0.0036964 82 |
| GO:00708 73 | regulation of glycogen metabolic process | 19/5895 | 0.0005830 08 | 0.0036964 82 |
| GO:19907 48 | cellular detoxification | 19/5895 | 0.0005830 08 | 0.0036964 82 |
| GO:00194 00 | alditol metabolic process | 14/5895 | 0.0005847 34 | 0.0036964 82 |
| GO:00363 14 | response to sterol | 14/5895 | 0.0005847 34 | 0.0036964 82 |
| GO:00468 23 | negative regulation of nucleocytoplasmic transport | 14/5895 | 0.0005847 34 | 0.0036964 82 |
| GO:00486 70 | regulation of collateral sprouting | 14/5895 | 0.0005847 34 | 0.0036964 82 |
| GO:00725 25 | pyridine-containing compound biosynthetic process | 14/5895 | 0.0005847 34 | 0.0036964 82 |
| GO:19040 62 | regulation of cation transmembrane transport | 119/589 5 | 0.0005874 01 | 0.0037096 26 |
| GO:00064 03 | RNA localization | 66/5895 | 0.0006264 48 | 0.0039522 57 |
| GO:00421 57 | lipoprotein metabolic process | 49/5895 | 0.0006276 73 | 0.0039560 3 |
| GO:00224 08 | negative regulation of cell-cell adhesion | 69/5895 | 0.0006385 99 | 0.0040208 78 |
| GO:00512 89 | protein homotetramerization | 25/5895 | 0.0006493 24 | 0.0040802 64 |
| GO:00550 25 | positive regulation of cardiac muscle tissue development | 25/5895 | 0.0006493 24 | 0.0040802 64 |

| | | | | |
|---|---|---|---|---|
| GO:0071453 | cellular response to oxygen levels | 53/5895 | 0.000656593 | 0.004121441 |
| GO:1904645 | response to amyloid-beta | 20/5895 | 0.000657183 | 0.004121441 |
| GO:0046031 | ADP metabolic process | 40/5895 | 0.000663552 | 0.004157247 |
| GO:0043489 | RNA stabilization | 26/5895 | 0.000676282 | 0.004228608 |
| GO:1902373 | negative regulation of mRNA catabolic process | 26/5895 | 0.000676282 | 0.004228608 |
| GO:0060998 | regulation of dendritic spine development | 39/5895 | 0.000678411 | 0.004237716 |
| GO:0006099 | tricarboxylic acid cycle | 17/5895 | 0.000680197 | 0.00424467 |
| GO:0010878 | cholesterol storage | Nov-95 | 0.000687001 | 0.004274444 |
| GO:0019359 | nicotinamide nucleotide biosynthetic process | Nov-95 | 0.000687001 | 0.004274444 |
| GO:0035459 | vesicle cargo loading | Nov-95 | 0.000687001 | 0.004274444 |
| GO:0048260 | positive regulation of receptor-mediated endocytosis | 27/5895 | 0.000697948 | 0.004326721 |
| GO:0051893 | regulation of focal adhesion assembly | 27/5895 | 0.000697948 | 0.004326721 |
| GO:0090109 | regulation of cell-substrate junction assembly | 27/5895 | 0.000697948 | 0.004326721 |
| GO:2000177 | regulation of neural precursor cell proliferation | 46/5895 | 0.000698146 | 0.004326721 |
| GO:0046928 | regulation of neurotransmitter secretion | 51/5895 | 0.000712153 | 0.004409196 |
| GO:0070482 | response to oxygen levels | 97/5895 | 0.000720187 | 0.004454567 |
| GO:0019079 | viral genome replication | 45/5895 | 0.000722014 | 0.004461492 |
| GO:0097484 | dendrite extension | 21/5895 | 0.000723637 | 0.004467148 |

| | | | | |
|---|---|---|---|---|
| GO:0051384 | response to glucocorticoid | 35/5895 | 0.000725177 | 0.0044722 8 |
| GO:0051236 | establishment of RNA localization | 59/5895 | 0.000733765 | 0.004520824 |
| GO:0034394 | protein localization to cell surface | 30/5895 | 0.000734746 | 0.0045224 5 |
| GO:0060042 | retina morphogenesis in camera-type eye | 33/5895 | 0.000737401 | 0.004534371 |
| GO:0042306 | regulation of protein import into nucleus | 32/5895 | 0.000739668 | 0.004543882 |
| GO:0048640 | negative regulation of developmental growth | 50/5895 | 0.000741024 | 0.004547781 |
| GO:0042908 | xenobiotic transport | 15/5895 | 0.000757037 | 0.004641545 |
| GO:0050657 | nucleic acid transport | 58/5895 | 0.000768352 | 0.004701778 |
| GO:0050658 | RNA transport | 58/5895 | 0.000768352 | 0.004701778 |
| GO:0051897 | positive regulation of protein kinase B signaling | 43/5895 | 0.000769728 | 0.004705633 |
| GO:0015012 | heparan sulfate proteoglycan biosynthetic process | 13/5895 | 0.000777985 | 0.004737749 |
| GO:0043574 | peroxisomal transport | 13/5895 | 0.000777985 | 0.004737749 |
| GO:1900101 | regulation of endoplasmic reticulum unfolded protein response | 13/5895 | 0.000777985 | 0.004737749 |
| GO:1900543 | negative regulation of purine nucleotide metabolic process | 13/5895 | 0.000777985 | 0.004737749 |
| GO:0033344 | cholesterol efflux | 22/5895 | 0.000781859 | 0.004756748 |
| GO:1900181 | negative regulation of protein localization to nucleus | 18/5895 | 0.000791075 | 0.004808181 |
| GO:0014812 | muscle cell migration | 42/5895 | 0.000793325 | 0.004817214 |
| GO:1903532 | positive regulation of secretion by cell | 126/5895 | 0.000796365 | 0.004831028 |

| | | | | |
|---|---|---|---|---|
| GO:0060401 | cytosolic calcium ion transport | 64/5895 | 0.000817855 | 0.004956623 |
| GO:0048255 | mRNA stabilization | 23/5895 | 0.000831738 | 0.005031096 |
| GO:0090342 | regulation of cell aging | 23/5895 | 0.000831738 | 0.005031096 |
| GO:0034249 | negative regulation of cellular amide metabolic process | 60/5895 | 0.000836024 | 0.005052178 |
| GO:0006816 | calcium ion transport | 135/5895 | 0.000841923 | 0.005082959 |
| GO:0001894 | tissue homeostasis | 89/5895 | 0.000850556 | 0.005130172 |
| GO:0010660 | regulation of muscle cell apoptotic process | 39/5895 | 0.000861169 | 0.005184271 |
| GO:0099072 | regulation of postsynaptic membrane neurotransmitter receptor levels | 39/5895 | 0.000861169 | 0.005184271 |
| GO:0009070 | serine family amino acid biosynthetic process | Oct-95 | 0.000867897 | 0.005195029 |
| GO:0010885 | regulation of cholesterol storage | Oct-95 | 0.000867897 | 0.005195029 |
| GO:0014857 | regulation of skeletal muscle cell proliferation | Oct-95 | 0.000867897 | 0.005195029 |
| GO:0016114 | terpenoid biosynthetic process | Oct-95 | 0.000867897 | 0.005195029 |
| GO:0072350 | tricarboxylic acid metabolic process | Oct-95 | 0.000867897 | 0.005195029 |
| GO:2000811 | negative regulation of anoikis | Oct-95 | 0.000867897 | 0.005195029 |
| GO:0007568 | aging | 86/5895 | 0.000871947 | 0.005209386 |
| GO:1901215 | negative regulation of neuron death | 86/5895 | 0.000871947 | 0.005209386 |
| GO:0021915 | neural tube development | 69/5895 | 0.0008821 | 0.005260083 |
| GO:0030705 | cytoskeleton-dependent intracellular transport | 69/5895 | 0.0008821 | 0.005260083 |

| | | | | |
|---|---|---|---|---|
| GO:1900407 | regulation of cellular response to oxidative stress | 37/5895 | 0.000901899 | 0.005369031 |
| GO:0070304 | positive regulation of stress-activated protein kinase signaling cascade | 62/5895 | 0.000902072 | 0.005369031 |
| GO:0061001 | regulation of dendritic spine morphogenesis | 25/5895 | 0.000907337 | 0.005395276 |
| GO:1903312 | negative regulation of mRNA metabolic process | 36/5895 | 0.000920159 | 0.005466368 |
| GO:0010874 | regulation of cholesterol efflux | 16/5895 | 0.000922193 | 0.005468154 |
| GO:0060317 | cardiac epithelial to mesenchymal transition | 16/5895 | 0.000922193 | 0.005468154 |
| GO:0030516 | regulation of axon extension | 44/5895 | 0.00092639 | 0.005487884 |
| GO:1903902 | positive regulation of viral life cycle | 26/5895 | 0.0009339 | 0.005527179 |
| GO:0014065 | phosphatidylinositol 3-kinase signaling | 49/5895 | 0.000941602 | 0.00556754 |
| GO:0030500 | regulation of bone mineralization | 34/5895 | 0.000950904 | 0.005617276 |
| GO:0006893 | Golgi to plasma membrane transport | 27/5895 | 0.000953686 | 0.005617931 |
| GO:0007584 | response to nutrient | 27/5895 | 0.000953686 | 0.005617931 |
| GO:0010812 | negative regulation of cell-substrate adhesion | 27/5895 | 0.000953686 | 0.005617931 |
| GO:0017015 | regulation of transforming growth factor beta receptor signaling pathway | 43/5895 | 0.000958551 | 0.005641322 |
| GO:0006879 | cellular iron ion homeostasis | 28/5895 | 0.000967282 | 0.005682109 |
| GO:1903510 | mucopolysaccharide metabolic process | 28/5895 | 0.000967282 | 0.005682109 |
| GO:0060997 | dendritic spine morphogenesis | 31/5895 | 0.000976821 | 0.005732806 |
| GO:0030048 | actin filament-based movement | 48/5895 | 0.000980395 | 0.005743091 |

| | | | | |
|---|---|---|---|---|
| GO:0036294 | cellular response to decreased oxygen levels | 48/5895 | 0.000980395 | 0.005743091 |
| GO:1903426 | regulation of reactive oxygen species biosynthetic process | 42/5895 | 0.000990759 | 0.005798421 |
| GO:0051258 | protein polymerization | 98/5895 | 0.001001278 | 0.005854554 |
| GO:0010921 | regulation of phosphatase activity | 47/5895 | 0.001020134 | 0.005925671 |
| GO:0048565 | digestive tract development | 47/5895 | 0.001020134 | 0.005925671 |
| GO:0000423 | mitophagy | 14/5895 | 0.001020955 | 0.005925671 |
| GO:0006706 | steroid catabolic process | 14/5895 | 0.001020955 | 0.005925671 |
| GO:0010884 | positive regulation of lipid storage | 14/5895 | 0.001020955 | 0.005925671 |
| GO:0042430 | indole-containing compound metabolic process | 14/5895 | 0.001020955 | 0.005925671 |
| GO:0046426 | negative regulation of receptor signaling pathway via JAK-STAT | 14/5895 | 0.001020955 | 0.005925671 |
| GO:0060004 | reflex | 14/5895 | 0.001020955 | 0.005925671 |
| GO:0009135 | purine nucleoside diphosphate metabolic process | 41/5895 | 0.00102285 | 0.005925768 |
| GO:0009179 | purine ribonucleoside diphosphate metabolic process | 41/5895 | 0.00102285 | 0.005925768 |
| GO:0007159 | leukocyte cell-cell adhesion | 113/5895 | 0.001024902 | 0.005932208 |
| GO:0034655 | nucleobase-containing compound catabolic process | 115/5895 | 0.001030825 | 0.005952 |
| GO:0009067 | aspartate family amino acid biosynthetic process | Dec-95 | 0.001032095 | 0.005952 |
| GO:0010888 | negative regulation of lipid storage | Dec-95 | 0.001032095 | 0.005952 |
| GO:2000773 | negative regulation of cellular senescence | Dec-95 | 0.001032095 | 0.005952 |

| | | | | |
|---|---|---|---|---|
| GO:0005978 | glycogen biosynthetic process | 21/5895 | 0.00105742 | 0.006064784 |
| GO:0009250 | glucan biosynthetic process | 21/5895 | 0.00105742 | 0.006064784 |
| GO:0051898 | negative regulation of protein kinase B signaling | 21/5895 | 0.00105742 | 0.006064784 |
| GO:0097237 | cellular response to toxic substance | 21/5895 | 0.00105742 | 0.006064784 |
| GO:1902895 | positive regulation of pri-miRNA transcription by RNA polymerase II | 21/5895 | 0.00105742 | 0.006064784 |
| GO:1903523 | negative regulation of blood circulation | 21/5895 | 0.00105742 | 0.006064784 |
| GO:0006544 | glycine metabolic process | Sep-95 | 0.001072558 | 0.006123771 |
| GO:0006895 | Golgi to endosome transport | Sep-95 | 0.001072558 | 0.006123771 |
| GO:0034497 | protein localization to phagophore assembly site | Sep-95 | 0.001072558 | 0.006123771 |
| GO:0055091 | phospholipid homeostasis | Sep-95 | 0.001072558 | 0.006123771 |
| GO:0071679 | commissural neuron axon guidance | Sep-95 | 0.001072558 | 0.006123771 |
| GO:0009101 | glycoprotein biosynthetic process | 92/5895 | 0.001075469 | 0.00613484 |
| GO:0097305 | response to alcohol | 74/5895 | 0.00107748 | 0.006140761 |
| GO:0051341 | regulation of oxidoreductase activity | 38/5895 | 0.001116382 | 0.006356727 |
| GO:0003197 | endocardial cushion development | 22/5895 | 0.001122194 | 0.006372566 |
| GO:0010718 | positive regulation of epithelial to mesenchymal transition | 22/5895 | 0.001122194 | 0.006372566 |
| GO:0060964 | regulation of gene silencing by miRNA | 22/5895 | 0.001122194 | 0.006372566 |
| GO:0050714 | positive regulation of protein secretion | 70/5895 | 0.001143784 | 0.006480621 |

| | | | | |
|---|---|---|---|---|
| GO:0001892 | embryonic placenta development | 44/5895 | 0.001144304 | 0.006480621 |
| GO:0060078 | regulation of postsynaptic membrane potential | 44/5895 | 0.001144304 | 0.006480621 |
| GO:0055072 | iron ion homeostasis | 37/5895 | 0.0011458 1 | 0.0064833 3 |
| GO:0060147 | regulation of posttranscriptional gene silencing | 23/5895 | 0.001175348 | 0.006638557 |
| GO:0060966 | regulation of gene silencing by RNA | 23/5895 | 0.001175348 | 0.006638557 |
| GO:0016241 | regulation of macroautophagy | 43/5895 | 0.001187013 | 0.006686484 |
| GO:0035601 | protein deacylation | 43/5895 | 0.001187013 | 0.006686484 |
| GO:0098732 | macromolecule deacylation | 43/5895 | 0.001187013 | 0.006686484 |
| GO:0048009 | insulin-like growth factor receptor signaling pathway | 18/5895 | 0.001211103 | 0.006816102 |
| GO:0009395 | phospholipid catabolic process | 24/5895 | 0.001217558 | 0.006834142 |
| GO:0050435 | amyloid-beta metabolic process | 24/5895 | 0.001217558 | 0.006834142 |
| GO:0051055 | negative regulation of lipid biosynthetic process | 24/5895 | 0.001217558 | 0.006834142 |
| GO:0009185 | ribonucleoside diphosphate metabolic process | 42/5895 | 0.001230163 | 0.006898752 |
| GO:0043500 | muscle adaptation | 47/5895 | 0.001245514 | 0.006932396 |
| GO:0050688 | regulation of defense response to virus | 33/5895 | 0.001245813 | 0.006932396 |
| GO:0051651 | maintenance of location in cell | 74/5895 | 0.001248382 | 0.006932396 |
| GO:0009119 | ribonucleoside metabolic process | 25/5895 | 0.001249642 | 0.006932396 |
| GO:0050873 | brown fat cell differentiation | 25/5895 | 0.001249642 | 0.006932396 |

| | | | | |
|---|---|---|---|---|
| GO:0002089 | lens morphogenesis in camera-type eye | 15/5895 | 0.001251545 | 0.006932396 |
| GO:0016577 | histone demethylation | 15/5895 | 0.001251545 | 0.006932396 |
| GO:0045663 | positive regulation of myoblast differentiation | 15/5895 | 0.001251545 | 0.006932396 |
| GO:0048596 | embryonic camera-type eye morphogenesis | 15/5895 | 0.001251545 | 0.006932396 |
| GO:0060390 | regulation of SMAD protein signal transduction | 15/5895 | 0.001251545 | 0.006932396 |
| GO:0070076 | histone lysine demethylation | 15/5895 | 0.001251545 | 0.006932396 |
| GO:0071402 | cellular response to lipoprotein particle stimulus | 15/5895 | 0.001251545 | 0.006932396 |
| GO:0090077 | foam cell differentiation | 15/5895 | 0.001251545 | 0.006932396 |
| GO:1903671 | negative regulation of sprouting angiogenesis | 15/5895 | 0.001251545 | 0.006932396 |
| GO:0010822 | positive regulation of mitochondrion organization | 32/5895 | 0.001264197 | 0.006990201 |
| GO:0048844 | artery morphogenesis | 32/5895 | 0.001264197 | 0.006990201 |
| GO:0000082 | G1/S transition of mitotic cell cycle | 65/5895 | 0.001269075 | 0.007007935 |
| GO:0060443 | mammary gland morphogenesis | 26/5895 | 0.001272484 | 0.007007935 |
| GO:0006107 | oxaloacetate metabolic process | Aug-95 | 0.001279623 | 0.007007935 |
| GO:0010898 | positive regulation of triglyceride catabolic process | Aug-95 | 0.001279623 | 0.007007935 |
| GO:0019627 | urea metabolic process | Aug-95 | 0.001279623 | 0.007007935 |
| GO:0035336 | long-chain fatty-acyl-CoA metabolic process | Aug-95 | 0.001279623 | 0.007007935 |
| GO:0038180 | nerve growth factor signaling pathway | Aug-95 | 0.001279623 | 0.007007935 |

| | | | | |
|---|---|---|---|---|
| GO:0042762 | regulation of sulfur metabolic process | Aug-95 | 0.001279623 | 0.007007935 |
| GO:0042996 | regulation of Golgi to plasma membrane protein transport | Aug-95 | 0.001279623 | 0.007007935 |
| GO:0045945 | positive regulation of transcription by RNA polymerase III | Aug-95 | 0.001279623 | 0.007007935 |
| GO:1905168 | positive regulation of double-strand break repair via homologous recombination | Aug-95 | 0.001279623 | 0.007007935 |
| GO:0010171 | body morphogenesis | 27/5895 | 0.00128699 | 0.00702389 |
| GO:0032387 | negative regulation of intracellular transport | 27/5895 | 0.00128699 | 0.00702389 |
| GO:0051058 | negative regulation of small GTPase mediated signal transduction | 27/5895 | 0.00128699 | 0.00702389 |
| GO:0072665 | protein localization to vacuole | 27/5895 | 0.00128699 | 0.00702389 |
| GO:0006513 | protein monoubiquitination | 30/5895 | 0.001289213 | 0.007029943 |
| GO:0032233 | positive regulation of actin filament bundle assembly | 28/5895 | 0.00129405 | 0.007046712 |
| GO:0006354 | DNA-templated transcription, elongation | 29/5895 | 0.001294522 | 0.007046712 |
| GO:0000910 | cytokinesis | 58/5895 | 0.001300755 | 0.007074536 |
| GO:0010657 | muscle cell apoptotic process | 40/5895 | 0.001317081 | 0.007151002 |
| GO:0035418 | protein localization to synapse | 40/5895 | 0.001317081 | 0.007151002 |
| GO:0007272 | ensheathment of neurons | 61/5895 | 0.001324214 | 0.007177375 |
| GO:0008366 | axon ensheathment | 61/5895 | 0.001324214 | 0.007177375 |
| GO:0035850 | epithelial cell differentiation involved in kidney development | 19/5895 | 0.001330094 | 0.007188908 |
| GO:0050892 | intestinal absorption | 19/5895 | 0.001330094 | 0.007188908 |

| | | | | |
|---|---|---|---|---|
| GO:0071827 | plasma lipoprotein particle organization | 19/5895 | 0.001330094 | 0.007188908 |
| GO:0050870 | positive regulation of T cell activation | 70/5895 | 0.001330899 | 0.007188908 |
| GO:1902107 | positive regulation of leukocyte differentiation | 64/5895 | 0.00133606 | 0.007210608 |
| GO:0071241 | cellular response to inorganic substance | 78/5895 | 0.001341097 | 0.007231605 |
| GO:0006085 | acetyl-CoA biosynthetic process | Nov-95 | 0.001363517 | 0.007327451 |
| GO:0007398 | ectoderm development | Nov-95 | 0.001363517 | 0.007327451 |
| GO:0010867 | positive regulation of triglyceride biosynthetic process | Nov-95 | 0.001363517 | 0.007327451 |
| GO:0036315 | cellular response to sterol | Nov-95 | 0.001363517 | 0.007327451 |
| GO:0009065 | glutamine family amino acid catabolic process | 13/5895 | 0.001376858 | 0.007374021 |
| GO:0010743 | regulation of macrophage derived foam cell differentiation | 13/5895 | 0.001376858 | 0.007374021 |
| GO:0034405 | response to fluid shear stress | 13/5895 | 0.001376858 | 0.007374021 |
| GO:0042537 | benzene-containing compound metabolic process | 13/5895 | 0.001376858 | 0.007374021 |
| GO:0031032 | actomyosin structure organization | 72/5895 | 0.001388635 | 0.00742799 |
| GO:0016571 | histone methylation | 53/5895 | 0.00138929 | 0.00742799 |
| GO:1903578 | regulation of ATP metabolic process | 38/5895 | 0.001403328 | 0.007496697 |
| GO:0048259 | regulation of receptor-mediated endocytosis | 44/5895 | 0.001406005 | 0.007498296 |
| GO:0090100 | positive regulation of transmembrane receptor protein serine/threonine kinase signaling pathway | 44/5895 | 0.001406005 | 0.007498296 |
| GO:0006509 | membrane protein ectodomain proteolysis | 20/5895 | 0.001431244 | 0.007607175 |

| | | | | |
|---|---|---|---|---|
| GO:0010765 | positive regulation of sodium ion transport | 20/5895 | 0.001431244 | 0.007607175 |
| GO:0030501 | positive regulation of bone mineralization | 20/5895 | 0.001431244 | 0.007607175 |
| GO:2000772 | regulation of cellular senescence | 20/5895 | 0.001431244 | 0.007607175 |
| GO:0002028 | regulation of sodium ion transport | 37/5895 | 0.001445497 | 0.007670011 |
| GO:0008589 | regulation of smoothened signaling pathway | 37/5895 | 0.001445497 | 0.007670011 |
| GO:0006282 | regulation of DNA repair | 48/5895 | 0.001449187 | 0.007683124 |
| GO:0003015 | heart process | 79/5895 | 0.001459833 | 0.007724566 |
| GO:0009225 | nucleotide-sugar metabolic process | 16/5895 | 0.001461675 | 0.007724566 |
| GO:1902932 | positive regulation of alcohol biosynthetic process | 16/5895 | 0.001461675 | 0.007724566 |
| GO:1903844 | regulation of cellular response to transforming growth factor beta stimulus | 43/5895 | 0.001461901 | 0.007724566 |
| GO:0032024 | positive regulation of insulin secretion | 36/5895 | 0.001486566 | 0.007848323 |
| GO:0060047 | heart contraction | 76/5895 | 0.001493717 | 0.007879481 |
| GO:0051588 | regulation of neurotransmitter transport | 55/5895 | 0.001506099 | 0.00793816 |
| GO:0032873 | negative regulation of stress-activated MAPK cascade | 21/5895 | 0.001514979 | 0.007971644 |
| GO:0070303 | negative regulation of stress-activated protein kinase signaling cascade | 21/5895 | 0.001514979 | 0.007971644 |
| GO:0097756 | negative regulation of blood vessel diameter | 42/5895 | 0.001518793 | 0.007978407 |
| GO:1900006 | positive regulation of dendrite development | 42/5895 | 0.001518793 | 0.007978407 |
| GO:0048284 | organelle fusion | 51/5895 | 0.001525155 | 0.008005159 |

| | | | | |
|---|---|---|---|---|
| GO:0016197 | endosomal transport | 78/5895 | 0.001541035 | 0.008081787 |
| GO:0006720 | isoprenoid metabolic process | 34/5895 | 0.001563693 | 0.008193803 |
| GO:0006022 | aminoglycan metabolic process | 41/5895 | 0.001576518 | 0.008254153 |
| GO:0045429 | positive regulation of nitric oxide biosynthetic process | 22/5895 | 0.001582215 | 0.008263408 |
| GO:0048010 | vascular endothelial growth factor receptor signaling pathway | 22/5895 | 0.001582215 | 0.008263408 |
| GO:1990928 | response to amino acid starvation | 22/5895 | 0.001582215 | 0.008263408 |
| GO:0046209 | nitric oxide metabolic process | 33/5895 | 0.001598743 | 0.00834282 |
| GO:0043484 | regulation of RNA splicing | 57/5895 | 0.001618495 | 0.008438915 |
| GO:0032088 | negative regulation of NF-kappaB transcription factor activity | 32/5895 | 0.001630664 | 0.008481334 |
| GO:0060563 | neuroepithelial cell differentiation | 32/5895 | 0.001630664 | 0.008481334 |
| GO:0072332 | intrinsic apoptotic signaling pathway by p53 class mediator | 32/5895 | 0.001630664 | 0.008481334 |
| GO:0046470 | phosphatidylcholine metabolic process | 23/5895 | 0.001634156 | 0.008485505 |
| GO:0098754 | detoxification | 23/5895 | 0.001634156 | 0.008485505 |
| GO:0099633 | protein localization to postsynaptic specialization membrane | 17/5895 | 0.001647085 | 0.00852784 |
| GO:0099645 | neurotransmitter receptor localization to postsynaptic specialization membrane | 17/5895 | 0.001647085 | 0.00852784 |
| GO:1904646 | cellular response to amyloid-beta | 17/5895 | 0.001647085 | 0.00852784 |
| GO:0050821 | protein stabilization | 63/5895 | 0.001647716 | 0.00852784 |
| GO:0014743 | regulation of muscle hypertrophy | 31/5895 | 0.001658769 | 0.008572563 |

| | | | | |
|---|---|---|---|---|
| GO:0017148 | negative regulation of translation | 53/5895 | 0.001659074 | 0.008572563 |
| GO:0034113 | heterotypic cell-cell adhesion | 24/5895 | 0.001672146 | 0.008633035 |
| GO:0010611 | regulation of cardiac muscle hypertrophy | 30/5895 | 0.001682288 | 0.008664128 |
| GO:0010633 | negative regulation of epithelial cell migration | 30/5895 | 0.001682288 | 0.008664128 |
| GO:0061045 | negative regulation of wound healing | 30/5895 | 0.001682288 | 0.008664128 |
| GO:0006476 | protein deacetylation | 39/5895 | 0.001693637 | 0.008715463 |
| GO:0003170 | heart valve development | 25/5895 | 0.00169758 | 0.008721781 |
| GO:0045428 | regulation of nitric oxide biosynthetic process | 29/5895 | 0.00170036 | 0.008721781 |
| GO:0090344 | negative regulation of cell aging | 14/5895 | 0.001700395 | 0.008721781 |
| GO:1904893 | negative regulation of receptor signaling pathway via STAT | 14/5895 | 0.001700395 | 0.008721781 |
| GO:0060348 | bone development | 76/5895 | 0.001717358 | 0.008801633 |
| GO:0046434 | organophosphate catabolic process | 48/5895 | 0.001749978 | 0.008961533 |
| GO:0043523 | regulation of neuron apoptotic process | 90/5895 | 0.001754565 | 0.008977737 |
| GO:0050890 | cognition | 109/5895 | 0.001783397 | 0.009072914 |
| GO:0070665 | positive regulation of leukocyte proliferation | 55/5895 | 0.001787872 | 0.009072914 |
| GO:0006658 | phosphatidylserine metabolic process | Oct-95 | 0.001790423 | 0.009072914 |
| GO:0014856 | skeletal muscle cell proliferation | Oct-95 | 0.001790423 | 0.009072914 |
| GO:0042308 | negative regulation of protein import into nucleus | Oct-95 | 0.001790423 | 0.009072914 |

| | | | | |
|---|---|---|---|---|
| GO:0043217 | myelin maintenance | Oct-95 | 0.001790423 | 0.009072914 |
| GO:0045820 | negative regulation of glycolytic process | Oct-95 | 0.001790423 | 0.009072914 |
| GO:0060979 | vasculogenesis involved in coronary vascular morphogenesis | Oct-95 | 0.001790423 | 0.009072914 |
| GO:1901550 | regulation of endothelial cell development | Oct-95 | 0.001790423 | 0.009072914 |
| GO:1903140 | regulation of establishment of endothelial barrier | Oct-95 | 0.001790423 | 0.009072914 |
| GO:1903541 | regulation of exosomal secretion | Oct-95 | 0.001790423 | 0.009072914 |
| GO:1904590 | negative regulation of protein import | Oct-95 | 0.001790423 | 0.009072914 |
| GO:0051348 | negative regulation of transferase activity | 87/5895 | 0.001824044 | 0.009235867 |
| GO:0051402 | neuron apoptotic process | 98/5895 | 0.001836286 | 0.009290399 |
| GO:0006346 | DNA methylation-dependent heterochromatin assembly | Dec-95 | 0.001856388 | 0.00936207 |
| GO:0010039 | response to iron ion | Dec-95 | 0.001856388 | 0.00936207 |
| GO:0045821 | positive regulation of glycolytic process | Dec-95 | 0.001856388 | 0.00936207 |
| GO:2000209 | regulation of anoikis | Dec-95 | 0.001856388 | 0.00936207 |
| GO:0002526 | acute inflammatory response | 42/5895 | 0.001864825 | 0.009396929 |
| GO:0001938 | positive regulation of endothelial cell proliferation | 36/5895 | 0.001869258 | 0.009396929 |
| GO:0014015 | positive regulation of gliogenesis | 36/5895 | 0.001869258 | 0.009396929 |
| GO:0048644 | muscle organ morphogenesis | 36/5895 | 0.001869258 | 0.009396929 |
| GO:0030308 | negative regulation of cell growth | 69/5895 | 0.001886624 | 0.009476681 |

| | | | | |
|---|---|---|---|---|
| GO:0007269 | neurotransmitter secretion | 66/5895 | 0.001910999 | 0.009583858 |
| GO:0099643 | signal release from synapse | 66/5895 | 0.001910999 | 0.009583858 |
| GO:0032874 | positive regulation of stress-activated MAPK cascade | 60/5895 | 0.001925387 | 0.009614666 |
| GO:0071897 | DNA biosynthetic process | 60/5895 | 0.001925387 | 0.009614666 |
| GO:0006112 | energy reserve metabolic process | 35/5895 | 0.001926286 | 0.009614666 |
| GO:0031960 | response to corticosteroid | 35/5895 | 0.001926286 | 0.009614666 |
| GO:0060079 | excitatory postsynaptic potential | 35/5895 | 0.001926286 | 0.009614666 |
| GO:1900182 | positive regulation of protein localization to nucleus | 35/5895 | 0.001926286 | 0.009614666 |
| GO:0038179 | neurotrophin signaling pathway | 19/5895 | 0.001938918 | 0.009662427 |
| GO:0070570 | regulation of neuron projection regeneration | 19/5895 | 0.001938918 | 0.009662427 |
| GO:0098693 | regulation of synaptic vesicle cycle | 53/5895 | 0.001973709 | 0.009828042 |
| GO:0034765 | regulation of ion transmembrane transport | 152/5895 | 0.001981442 | 0.009844764 |
| GO:0000380 | alternative mRNA splicing, via spliceosome | 34/5895 | 0.001981748 | 0.009844764 |
| GO:0006835 | dicarboxylic acid transport | 34/5895 | 0.001981748 | 0.009844764 |
| GO:0021952 | central nervous system projection neuron axonogenesis | 15/5895 | 0.001990973 | 0.009859534 |
| GO:0039694 | viral RNA genome replication | 15/5895 | 0.001990973 | 0.009859534 |
| GO:0060444 | branching involved in mammary gland duct morphogenesis | 15/5895 | 0.001990973 | 0.009859534 |
| GO:0150117 | positive regulation of cell-substrate junction organization | 15/5895 | 0.001990973 | 0.009859534 |

**GO terms associated with 8759 genes near newly opened regions**

| ID | Description | GeneRatio | pvalue | p.adjust |
|---|---|---|---|---|
| GO:0007409 | axonogenesis | 320/7539 | 5.33E-54 | 3.36E-50 |
| GO:0050808 | synapse organization | 317/7539 | 4.94E-53 | 1.56E-49 |
| GO:0030900 | forebrain development | 258/7539 | 2.29E-42 | 4.82E-39 |
| GO:0097485 | neuron projection guidance | 183/7539 | 1.47E-39 | 2.32E-36 |
| GO:0007411 | axon guidance | 182/7539 | 3.35E-39 | 4.22E-36 |
| GO:0034329 | cell junction assembly | 268/7539 | 5.65E-39 | 5.95E-36 |
| GO:0034765 | regulation of ion transmembrane transport | 296/7539 | 2.70E-38 | 2.43E-35 |
| GO:0060562 | epithelial tube morphogenesis | 245/7539 | 2.26E-36 | 1.78E-33 |
| GO:0045666 | positive regulation of neuron differentiation | 284/7539 | 4.09E-36 | 2.87E-33 |
| GO:0007264 | small GTPase mediated signal transduction | 267/7539 | 1.25E-35 | 7.91E-33 |
| GO:0007389 | pattern specification process | 285/7539 | 3.10E-35 | 1.78E-32 |
| GO:0031589 | cell-substrate adhesion | 225/7539 | 1.05E-34 | 5.33E-32 |
| GO:0010721 | negative regulation of cell development | 242/7539 | 1.10E-34 | 5.33E-32 |
| GO:0001655 | urogenital system development | 236/7539 | 3.19E-34 | 1.44E-31 |
| GO:0060485 | mesenchyme development | 190/7539 | 4.08E-34 | 1.64E-31 |
| GO:0050807 | regulation of synapse organization | 188/7539 | 4.16E-34 | 1.64E-31 |

| | | | | |
|---|---|---|---|---|
| GO:0072001 | renal system development | 215/7539 | 6.84E-34 | 2.54E-31 |
| GO:0045165 | cell fate commitment | 194/7539 | 3.31E-33 | 1.11E-30 |
| GO:0051961 | negative regulation of nervous system development | 229/7539 | 3.34E-33 | 1.11E-30 |
| GO:0090596 | sensory organ morphogenesis | 203/7539 | 4.72E-33 | 1.49E-30 |
| GO:0050803 | regulation of synapse structure or activity | 192/7539 | 7.13E-33 | 2.14E-30 |
| GO:0001667 | ameboidal-type cell migration | 257/7539 | 8.21E-33 | 2.25E-30 |
| GO:0010959 | regulation of metal ion transport | 257/7539 | 8.21E-33 | 2.25E-30 |
| GO:0019932 | second-messenger-mediated signaling | 265/7539 | 1.03E-32 | 2.72E-30 |
| GO:1904062 | regulation of cation transmembrane transport | 225/7539 | 5.03E-32 | 1.27E-29 |
| GO:0050768 | negative regulation of neurogenesis | 213/7539 | 1.25E-31 | 3.04E-29 |
| GO:0048562 | embryonic organ morphogenesis | 208/7539 | 3.29E-31 | 7.69E-29 |
| GO:0001822 | kidney development | 203/7539 | 4.57E-31 | 1.03E-28 |
| GO:0021953 | central nervous system neuron differentiation | 162/7539 | 8.44E-31 | 1.84E-28 |
| GO:0051056 | regulation of small GTPase mediated signal transduction | 179/7539 | 2.94E-30 | 6.19E-28 |
| GO:0045860 | positive regulation of protein kinase activity | 257/7539 | 6.80E-30 | 1.39E-27 |
| GO:0060537 | muscle tissue development | 270/7539 | 1.01E-29 | 2.00E-27 |
| GO:0048880 | sensory system development | 249/7539 | 4.58E-29 | 8.77E-27 |
| GO:0001654 | eye development | 245/7539 | 7.22E-29 | 1.34E-26 |

| | | | | |
|---|---|---|---|---|
| GO:0001503 | ossification | 234/7539 | 8.77E-29 | 1.58E-26 |
| GO:0010769 | regulation of cell morphogenesis involved in differentiation | 214/7539 | 1.31E-28 | 2.30E-26 |
| GO:0031346 | positive regulation of cell projection organization | 271/7539 | 1.45E-28 | 2.48E-26 |
| GO:0150063 | visual system development | 246/7539 | 1.74E-28 | 2.89E-26 |
| GO:0014706 | striated muscle tissue development | 257/7539 | 2.02E-28 | 3.27E-26 |
| GO:0042391 | regulation of membrane potential | 258/7539 | 2.87E-28 | 4.53E-26 |
| GO:0015672 | monovalent inorganic cation transport | 275/7539 | 3.38E-28 | 5.19E-26 |
| GO:0003002 | regionalization | 226/7539 | 3.45E-28 | 5.19E-26 |
| GO:0021537 | telencephalon development | 166/7539 | 5.95E-28 | 8.73E-26 |
| GO:0001763 | morphogenesis of a branching structure | 160/7539 | 1.45E-27 | 2.09E-25 |
| GO:1901888 | regulation of cell junction assembly | 147/7539 | 1.75E-27 | 2.46E-25 |
| GO:0001764 | neuron migration | 133/7539 | 6.60E-27 | 9.06E-25 |
| GO:0048762 | mesenchymal cell differentiation | 151/7539 | 7.52E-27 | 1.01E-24 |
| GO:0007517 | muscle organ development | 241/7539 | 1.48E-26 | 1.94E-24 |
| GO:0043583 | ear development | 160/7539 | 4.47E-26 | 5.76E-24 |
| GO:0042063 | gliogenesis | 203/7539 | 5.22E-26 | 6.59E-24 |
| GO:0032412 | regulation of ion transmembrane transporter activity | 166/7539 | 6.22E-26 | 7.70E-24 |
| GO:0043270 | positive regulation of ion transport | 201/7539 | 6.38E-26 | 7.74E-24 |

| | | | | |
|---|---|---|---|---|
| GO:0007015 | actin filament organization | 247/7539 | 7.05E-26 | 8.40E-24 |
| GO:0016358 | dendrite development | 188/7539 | 7.22E-26 | 8.44E-24 |
| GO:0099173 | postsynapse organization | 137/7539 | 7.84E-26 | 9.00E-24 |
| GO:0022898 | regulation of transmembrane transporter activity | 171/7539 | 8.47E-26 | 9.55E-24 |
| GO:0007265 | Ras protein signal transduction | 208/7539 | 8.91E-26 | 9.87E-24 |
| GO:0006813 | potassium ion transport | 156/7539 | 2.05E-25 | 2.23E-23 |
| GO:0032409 | regulation of transporter activity | 175/7539 | 2.13E-25 | 2.28E-23 |
| GO:0045785 | positive regulation of cell adhesion | 245/7539 | 2.23E-25 | 2.34E-23 |
| GO:0010976 | positive regulation of neuron projection development | 217/7539 | 2.49E-25 | 2.58E-23 |
| GO:0003012 | muscle system process | 233/7539 | 2.53E-25 | 2.58E-23 |
| GO:0042692 | muscle cell differentiation | 228/7539 | 7.42E-25 | 7.43E-23 |
| GO:0046578 | regulation of Ras protein signal transduction | 155/7539 | 8.04E-25 | 7.93E-23 |
| GO:0045665 | negative regulation of neuron differentiation | 164/7539 | 8.75E-25 | 8.50E-23 |
| GO:0007178 | transmembrane receptor protein serine/threonine kinase signaling pathway | 214/7539 | 9.38E-25 | 8.97E-23 |
| GO:0072073 | kidney epithelium development | 107/7539 | 1.24E-24 | 1.16E-22 |
| GO:0061138 | morphogenesis of a branching epithelium | 145/7539 | 4.92E-24 | 4.57E-22 |
| GO:0006836 | neurotransmitter transport | 156/7539 | 5.17E-24 | 4.73E-22 |
| GO:2001257 | regulation of cation channel activity | 122/7539 | 7.18E-24 | 6.36E-22 |

| | | | | |
|---|---|---|---|---|
| GO:0043010 | camera-type eye development | 213/7539 | 7.23E-24 | 6.36E-22 |
| GO:0006816 | calcium ion transport | 235/7539 | 7.25E-24 | 6.36E-22 |
| GO:0048732 | gland development | 257/7539 | 7.68E-24 | 6.64E-22 |
| GO:0061351 | neural precursor cell proliferation | 129/7539 | 1.57E-23 | 1.34E-21 |
| GO:0060348 | bone development | 144/7539 | 1.96E-23 | 1.65E-21 |
| GO:0007611 | learning or memory | 180/7539 | 2.23E-23 | 1.85E-21 |
| GO:0048638 | regulation of developmental growth | 229/7539 | 2.42E-23 | 1.99E-21 |
| GO:0055074 | calcium ion homeostasis | 265/7539 | 2.82E-23 | 2.28E-21 |
| GO:0048663 | neuron fate commitment | 68/7539 | 4.91E-23 | 3.92E-21 |
| GO:0098742 | cell-cell adhesion via plasma-membrane adhesion molecules | 137/7539 | 5.24E-23 | 4.14E-21 |
| GO:0090287 | regulation of cellular response to growth factor stimulus | 171/7539 | 5.31E-23 | 4.14E-21 |
| GO:0007416 | synapse assembly | 128/7539 | 6.89E-23 | 5.30E-21 |
| GO:0090130 | tissue migration | 179/7539 | 7.16E-23 | 5.44E-21 |
| GO:0051271 | negative regulation of cellular component movement | 191/7539 | 9.22E-23 | 6.85E-21 |
| GO:0048863 | stem cell differentiation | 150/7539 | 9.29E-23 | 6.85E-21 |
| GO:0060541 | respiratory system development | 158/7539 | 9.34E-23 | 6.85E-21 |
| GO:1903522 | regulation of blood circulation | 155/7539 | 1.18E-22 | 8.58E-21 |
| GO:0071805 | potassium ion transmembrane transport | 132/7539 | 1.25E-22 | 9.00E-21 |

| | | | | |
|---|---|---|---|---|
| GO:0050890 | cognition | 195/7539 | 1.56E-22 | 1.11E-20 |
| GO:0070838 | divalent metal ion transport | 252/7539 | 1.67E-22 | 1.16E-20 |
| GO:0198738 | cell-cell signaling by wnt | 237/7539 | 1.68E-22 | 1.16E-20 |
| GO:0016055 | Wnt signaling pathway | 236/7539 | 1.91E-22 | 1.31E-20 |
| GO:0090132 | epithelium migration | 177/7539 | 2.52E-22 | 1.71E-20 |
| GO:0040013 | negative regulation of locomotion | 190/7539 | 2.77E-22 | 1.86E-20 |
| GO:0055123 | digestive system development | 99/7539 | 3.73E-22 | 2.48E-20 |
| GO:0072511 | divalent inorganic cation transport | 252/7539 | 5.38E-22 | 3.54E-20 |
| GO:0060560 | developmental growth involved in morphogenesis | 166/7539 | 7.65E-22 | 4.98E-20 |
| GO:0010631 | epithelial cell migration | 175/7539 | 8.78E-22 | 5.61E-20 |
| GO:0006874 | cellular calcium ion homeostasis | 254/7539 | 8.79E-22 | 5.61E-20 |
| GO:0070588 | calcium ion transmembrane transport | 169/7539 | 9.70E-22 | 6.13E-20 |
| GO:0061448 | connective tissue development | 166/7539 | 1.31E-21 | 8.20E-20 |
| GO:0051216 | cartilage development | 130/7539 | 2.05E-21 | 1.27E-19 |
| GO:0048705 | skeletal system morphogenesis | 155/7539 | 2.15E-21 | 1.32E-19 |
| GO:0030111 | regulation of Wnt signaling pathway | 178/7539 | 2.89E-21 | 1.75E-19 |
| GO:0007160 | cell-matrix adhesion | 136/7539 | 5.28E-21 | 3.14E-19 |
| GO:0010810 | regulation of cell-substrate adhesion | 136/7539 | 5.28E-21 | 3.14E-19 |

| | | | | |
|---|---|---|---|---|
| GO:0061458 | reproductive system development | 254/7539 | 5.73E-21 | 3.38E-19 |
| GO:0050770 | regulation of axonogenesis | 133/7539 | 6.38E-21 | 3.73E-19 |
| GO:0072006 | nephron development | 105/7539 | 6.60E-21 | 3.82E-19 |
| GO:0003007 | heart morphogenesis | 165/7539 | 7.09E-21 | 4.07E-19 |
| GO:0048608 | reproductive structure development | 252/7539 | 7.57E-21 | 4.25E-19 |
| GO:0048736 | appendage development | 130/7539 | 7.60E-21 | 4.25E-19 |
| GO:0060173 | limb development | 130/7539 | 7.60E-21 | 4.25E-19 |
| GO:0048839 | inner ear development | 136/7539 | 9.81E-21 | 5.43E-19 |
| GO:0021700 | developmental maturation | 187/7539 | 1.12E-20 | 6.14E-19 |
| GO:0006936 | muscle contraction | 174/7539 | 1.22E-20 | 6.64E-19 |
| GO:0090257 | regulation of muscle system process | 149/7539 | 1.99E-20 | 1.07E-18 |
| GO:0051146 | striated muscle cell differentiation | 182/7539 | 2.04E-20 | 1.09E-18 |
| GO:0071900 | regulation of protein serine/threonine kinase activity | 236/7539 | 2.25E-20 | 1.19E-18 |
| GO:0031644 | regulation of nervous system process | 121/7539 | 2.29E-20 | 1.21E-18 |
| GO:0050673 | epithelial cell proliferation | 234/7539 | 2.96E-20 | 1.54E-18 |
| GO:0032970 | regulation of actin filament-based process | 222/7539 | 3.20E-20 | 1.65E-18 |
| GO:0022407 | regulation of cell-cell adhesion | 229/7539 | 4.00E-20 | 2.05E-18 |
| GO:0001505 | regulation of neurotransmitter levels | 154/7539 | 6.28E-20 | 3.20E-18 |

| | | | | |
|---|---|---|---|---|
| GO:2000027 | regulation of animal organ morphogenesis | 132/7539 | 8.13E-20 | 4.11E-18 |
| GO:0048754 | branching morphogenesis of an epithelial tube | 120/7539 | 9.11E-20 | 4.56E-18 |
| GO:0010001 | glial cell differentiation | 150/7539 | 9.36E-20 | 4.65E-18 |
| GO:0007626 | locomotory behavior | 154/7539 | 1.06E-19 | 5.25E-18 |
| GO:0035107 | appendage morphogenesis | 115/7539 | 1.13E-19 | 5.51E-18 |
| GO:0035108 | limb morphogenesis | 115/7539 | 1.13E-19 | 5.51E-18 |
| GO:0009611 | response to wounding | 255/7539 | 1.25E-19 | 6.05E-18 |
| GO:0043087 | regulation of GTPase activity | 174/7539 | 1.35E-19 | 6.48E-18 |
| GO:0009914 | hormone transport | 204/7539 | 1.58E-19 | 7.50E-18 |
| GO:0046879 | hormone secretion | 200/7539 | 2.68E-19 | 1.26E-17 |
| GO:0042471 | ear morphogenesis | 96/7539 | 2.95E-19 | 1.38E-17 |
| GO:0014013 | regulation of gliogenesis | 104/7539 | 3.05E-19 | 1.42E-17 |
| GO:0070997 | neuron death | 222/7539 | 3.29E-19 | 1.52E-17 |
| GO:0016331 | morphogenesis of embryonic epithelium | 118/7539 | 3.74E-19 | 1.71E-17 |
| GO:0050678 | regulation of epithelial cell proliferation | 203/7539 | 4.09E-19 | 1.86E-17 |
| GO:1901342 | regulation of vasculature development | 196/7539 | 4.53E-19 | 2.04E-17 |
| GO:0051480 | regulation of cytosolic calcium ion concentration | 204/7539 | 5.36E-19 | 2.40E-17 |
| GO:0018108 | peptidyl-tyrosine phosphorylation | 181/7539 | 5.62E-19 | 2.50E-17 |

| | | | | |
|---|---|---|---|---|
| GO:0031345 | negative regulation of cell projection organization | 129/7539 | 6.14E-19 | 2.71E-17 |
| GO:0018212 | peptidyl-tyrosine modification | 182/7539 | 7.69E-19 | 3.37E-17 |
| GO:0030323 | respiratory tube development | 137/7539 | 9.67E-19 | 4.21E-17 |
| GO:0061572 | actin filament bundle organization | 112/7539 | 9.75E-19 | 4.22E-17 |
| GO:0099175 | regulation of postsynapse organization | 92/7539 | 1.33E-18 | 5.72E-17 |
| GO:0030278 | regulation of ossification | 144/7539 | 1.44E-18 | 6.12E-17 |
| GO:0007612 | learning | 113/7539 | 1.79E-18 | 7.60E-17 |
| GO:0003205 | cardiac chamber development | 122/7539 | 1.90E-18 | 7.99E-17 |
| GO:0051924 | regulation of calcium ion transport | 159/7539 | 2.05E-18 | 8.56E-17 |
| GO:0048565 | digestive tract development | 87/7539 | 2.55E-18 | 1.06E-16 |
| GO:0061326 | renal tubule development | 77/7539 | 2.89E-18 | 1.19E-16 |
| GO:0048738 | cardiac muscle tissue development | 151/7539 | 3.13E-18 | 1.28E-16 |
| GO:0060047 | heart contraction | 134/7539 | 3.86E-18 | 1.57E-16 |
| GO:0006939 | smooth muscle contraction | 82/7539 | 4.55E-18 | 1.84E-16 |
| GO:0008016 | regulation of heart contraction | 116/7539 | 5.33E-18 | 2.14E-16 |
| GO:0051963 | regulation of synapse assembly | 87/7539 | 5.85E-18 | 2.34E-16 |
| GO:0010632 | regulation of epithelial cell migration | 140/7539 | 6.17E-18 | 2.45E-16 |
| GO:0035148 | tube formation | 113/7539 | 6.46E-18 | 2.55E-16 |

| | | | | |
|---|---|---|---|---|
| GO:0030324 | lung development | 134/7539 | 6.64E-18 | 2.60E-16 |
| GO:1901214 | regulation of neuron death | 202/7539 | 7.45E-18 | 2.90E-16 |
| GO:0032147 | activation of protein kinase activity | 143/7539 | 7.53E-18 | 2.92E-16 |
| GO:0003015 | heart process | 138/7539 | 7.59E-18 | 2.92E-16 |
| GO:0001823 | mesonephros development | 78/7539 | 8.15E-18 | 3.10E-16 |
| GO:0051017 | actin filament bundle assembly | 109/7539 | 8.15E-18 | 3.10E-16 |
| GO:0099003 | vesicle-mediated transport in synapse | 151/7539 | 8.36E-18 | 3.16E-16 |
| GO:0030336 | negative regulation of cell migration | 159/7539 | 8.44E-18 | 3.17E-16 |
| GO:0048588 | developmental cell growth | 154/7539 | 9.55E-18 | 3.55E-16 |
| GO:0072009 | nephron epithelium development | 80/7539 | 9.56E-18 | 3.55E-16 |
| GO:0019722 | calcium-mediated signaling | 130/7539 | 9.73E-18 | 3.59E-16 |
| GO:0001657 | ureteric bud development | 76/7539 | 1.70E-17 | 6.16E-16 |
| GO:0072163 | mesonephric epithelium development | 76/7539 | 1.70E-17 | 6.16E-16 |
| GO:0072164 | mesonephric tubule development | 76/7539 | 1.70E-17 | 6.16E-16 |
| GO:0071902 | positive regulation of protein serine/threonine kinase activity | 164/7539 | 1.79E-17 | 6.42E-16 |
| GO:2000146 | negative regulation of cell motility | 164/7539 | 1.79E-17 | 6.42E-16 |
| GO:0048015 | phosphatidylinositol-mediated signaling | 106/7539 | 1.83E-17 | 6.54E-16 |
| GO:0014033 | neural crest cell differentiation | 69/7539 | 1.87E-17 | 6.64E-16 |

| | | | | |
|---|---|---|---|---|
| GO:0071695 | anatomical structure maturation | 156/7539 | 1.93E-17 | 6.77E-16 |
| GO:1990778 | protein localization to cell periphery | 195/7539 | 1.93E-17 | 6.77E-16 |
| GO:0032956 | regulation of actin cytoskeleton organization | 194/7539 | 2.21E-17 | 7.71E-16 |
| GO:0090092 | regulation of transmembrane receptor protein serine/threonine kinase signaling pathway | 144/7539 | 3.02E-17 | 1.05E-15 |
| GO:0048017 | inositol lipid-mediated signaling | 107/7539 | 3.30E-17 | 1.14E-15 |
| GO:1902903 | regulation of supramolecular fiber organization | 197/7539 | 4.63E-17 | 1.59E-15 |
| GO:0015837 | amine transport | 84/7539 | 6.08E-17 | 2.07E-15 |
| GO:0072080 | nephron tubule development | 72/7539 | 7.33E-17 | 2.49E-15 |
| GO:0007266 | Rho protein signal transduction | 113/7539 | 7.47E-17 | 2.52E-15 |
| GO:0033002 | muscle cell proliferation | 138/7539 | 9.70E-17 | 3.26E-15 |
| GO:0070371 | ERK1 and ERK2 cascade | 177/7539 | 9.86E-17 | 3.29E-15 |
| GO:0043266 | regulation of potassium ion transport | 77/7539 | 1.08E-16 | 3.57E-15 |
| GO:0030902 | hindbrain development | 108/7539 | 1.08E-16 | 3.57E-15 |
| GO:0019935 | cyclic-nucleotide-mediated signaling | 132/7539 | 1.13E-16 | 3.69E-15 |
| GO:0032535 | regulation of cellular component size | 214/7539 | 1.13E-16 | 3.69E-15 |
| GO:0072089 | stem cell proliferation | 95/7539 | 1.14E-16 | 3.69E-15 |
| GO:0045444 | fat cell differentiation | 145/7539 | 1.14E-16 | 3.69E-15 |
| GO:0060070 | canonical Wnt signaling pathway | 159/7539 | 1.26E-16 | 4.06E-15 |

| | | | | |
|---|---|---|---|---|
| GO:0046883 | regulation of hormone secretion | 167/7539 | 1.62E-16 | 5.20E-15 |
| GO:0003018 | vascular process in circulatory system | 132/7539 | 1.88E-16 | 5.99E-15 |
| GO:0070372 | regulation of ERK1 and ERK2 cascade | 169/7539 | 1.89E-16 | 6.00E-15 |
| GO:1904064 | positive regulation of cation transmembrane transport | 107/7539 | 2.13E-16 | 6.74E-15 |
| GO:0035270 | endocrine system development | 96/7539 | 2.16E-16 | 6.79E-15 |
| GO:0045765 | regulation of angiogenesis | 174/7539 | 2.20E-16 | 6.87E-15 |
| GO:0002062 | chondrocyte differentiation | 77/7539 | 2.51E-16 | 7.79E-15 |
| GO:0021915 | neural tube development | 118/7539 | 2.71E-16 | 8.38E-15 |
| GO:0048864 | stem cell development | 67/7539 | 2.79E-16 | 8.59E-15 |
| GO:0030198 | extracellular matrix organization | 166/7539 | 2.80E-16 | 8.59E-15 |
| GO:0001894 | tissue homeostasis | 148/7539 | 3.18E-16 | 9.71E-15 |
| GO:0034764 | positive regulation of transmembrane transport | 140/7539 | 3.30E-16 | 1.00E-14 |
| GO:0006937 | regulation of muscle contraction | 100/7539 | 3.38E-16 | 1.02E-14 |
| GO:0071559 | response to transforming growth factor beta | 126/7539 | 3.56E-16 | 1.07E-14 |
| GO:0043062 | extracellular structure organization | 166/7539 | 4.23E-16 | 1.27E-14 |
| GO:0048592 | eye morphogenesis | 112/7539 | 4.64E-16 | 1.38E-14 |
| GO:0048639 | positive regulation of developmental growth | 128/7539 | 4.81E-16 | 1.42E-14 |
| GO:0015844 | monoamine transport | 74/7539 | 5.06E-16 | 1.49E-14 |

| | | | | |
|---|---|---|---|---|
| GO:0035023 | regulation of Rho protein signal transduction | 91/7539 | 5.20E-16 | 1.53E-14 |
| GO:0007162 | negative regulation of cell adhesion | 160/7539 | 6.14E-16 | 1.79E-14 |
| GO:0007269 | neurotransmitter secretion | 115/7539 | 6.22E-16 | 1.80E-14 |
| GO:0099643 | signal release from synapse | 115/7539 | 6.22E-16 | 1.80E-14 |
| GO:0014031 | mesenchymal cell development | 66/7539 | 6.48E-16 | 1.87E-14 |
| GO:0034767 | positive regulation of ion transmembrane transport | 114/7539 | 6.82E-16 | 1.96E-14 |
| GO:1990138 | neuron projection extension | 119/7539 | 7.28E-16 | 2.08E-14 |
| GO:0031032 | actomyosin structure organization | 123/7539 | 8.19E-16 | 2.33E-14 |
| GO:0003206 | cardiac chamber morphogenesis | 95/7539 | 8.52E-16 | 2.41E-14 |
| GO:0007163 | establishment or maintenance of cell polarity | 130/7539 | 1.04E-15 | 2.94E-14 |
| GO:0043542 | endothelial cell migration | 125/7539 | 1.10E-15 | 3.10E-14 |
| GO:0002064 | epithelial cell development | 145/7539 | 1.13E-15 | 3.17E-14 |
| GO:0043405 | regulation of MAP kinase activity | 164/7539 | 1.24E-15 | 3.46E-14 |
| GO:0002040 | sprouting angiogenesis | 83/7539 | 1.29E-15 | 3.57E-14 |
| GO:0022612 | gland morphogenesis | 98/7539 | 1.35E-15 | 3.73E-14 |
| GO:0071560 | cellular response to transforming growth factor beta stimulus | 123/7539 | 1.37E-15 | 3.76E-14 |
| GO:0090288 | negative regulation of cellular response to growth factor stimulus | 97/7539 | 1.44E-15 | 3.94E-14 |
| GO:0045598 | regulation of fat cell differentiation | 94/7539 | 1.72E-15 | 4.68E-14 |

| | | | | |
|---|---|---|---|---|
| GO:0007204 | positive regulation of cytosolic calcium ion concentration | 176/7539 | 1.74E-15 | 4.72E-14 |
| GO:0045685 | regulation of glial cell differentiation | 67/7539 | 1.82E-15 | 4.91E-14 |
| GO:0010977 | negative regulation of neuron projection development | 108/7539 | 2.05E-15 | 5.51E-14 |
| GO:0045995 | regulation of embryonic development | 88/7539 | 2.29E-15 | 6.12E-14 |
| GO:0045927 | positive regulation of growth | 168/7539 | 2.40E-15 | 6.38E-14 |
| GO:0003279 | cardiac septum development | 85/7539 | 2.54E-15 | 6.72E-14 |
| GO:2000177 | regulation of neural precursor cell proliferation | 80/7539 | 2.79E-15 | 7.34E-14 |
| GO:0042472 | inner ear morphogenesis | 79/7539 | 2.80E-15 | 7.34E-14 |
| GO:0051952 | regulation of amine transport | 79/7539 | 2.80E-15 | 7.34E-14 |
| GO:0051588 | regulation of neurotransmitter transport | 96/7539 | 2.87E-15 | 7.50E-14 |
| GO:0051048 | negative regulation of secretion | 129/7539 | 3.09E-15 | 8.02E-14 |
| GO:0030326 | embryonic limb morphogenesis | 92/7539 | 3.67E-15 | 9.44E-14 |
| GO:0035113 | embryonic appendage morphogenesis | 92/7539 | 3.67E-15 | 9.44E-14 |
| GO:0072175 | epithelial tube formation | 101/7539 | 3.68E-15 | 9.44E-14 |
| GO:0019933 | cAMP-mediated signaling | 116/7539 | 4.79E-15 | 1.23E-13 |
| GO:0048813 | dendrite morphogenesis | 110/7539 | 5.16E-15 | 1.31E-13 |
| GO:0060993 | kidney morphogenesis | 71/7539 | 5.47E-15 | 1.39E-13 |
| GO:0009952 | anterior/posterior pattern specification | 132/7539 | 5.56E-15 | 1.40E-13 |

| | | | | |
|---|---|---|---|---|
| GO:0001952 | regulation of cell-matrix adhesion | 80/7539 | 5.81E-15 | 1.46E-13 |
| GO:0042476 | odontogenesis | 80/7539 | 5.81E-15 | 1.46E-13 |
| GO:1901890 | positive regulation of cell junction assembly | 79/7539 | 5.90E-15 | 1.47E-13 |
| GO:0071772 | response to BMP | 108/7539 | 6.24E-15 | 1.54E-13 |
| GO:0071773 | cellular response to BMP stimulus | 108/7539 | 6.24E-15 | 1.54E-13 |
| GO:1903169 | regulation of calcium ion transmembrane transport | 101/7539 | 6.61E-15 | 1.63E-13 |
| GO:0003151 | outflow tract morphogenesis | 62/7539 | 7.15E-15 | 1.75E-13 |
| GO:0014032 | neural crest cell development | 62/7539 | 7.15E-15 | 1.75E-13 |
| GO:0035265 | organ growth | 121/7539 | 7.68E-15 | 1.87E-13 |
| GO:0055001 | muscle cell development | 120/7539 | 8.58E-15 | 2.08E-13 |
| GO:0001838 | embryonic epithelial tube formation | 97/7539 | 9.07E-15 | 2.19E-13 |
| GO:0030072 | peptide hormone secretion | 155/7539 | 9.21E-15 | 2.22E-13 |
| GO:1901215 | negative regulation of neuron death | 140/7539 | 1.23E-14 | 2.96E-13 |
| GO:0045778 | positive regulation of ossification | 72/7539 | 1.25E-14 | 3.00E-13 |
| GO:0008306 | associative learning | 74/7539 | 1.28E-14 | 3.05E-13 |
| GO:0001933 | negative regulation of protein phosphorylation | 213/7539 | 1.32E-14 | 3.14E-13 |
| GO:0050773 | regulation of dendrite development | 116/7539 | 1.33E-14 | 3.15E-13 |
| GO:0030100 | regulation of endocytosis | 127/7539 | 1.61E-14 | 3.79E-13 |

| | | | | |
|---|---|---|---|---|
| GO:0030177 | positive regulation of Wnt signaling pathway | 87/7539 | 1.76E-14 | 4.13E-13 |
| GO:0051937 | catecholamine transport | 62/7539 | 1.81E-14 | 4.23E-13 |
| GO:0001656 | metanephros development | 64/7539 | 2.11E-14 | 4.91E-13 |
| GO:0044344 | cellular response to fibroblast growth factor stimulus | 65/7539 | 2.24E-14 | 5.20E-13 |
| GO:0042060 | wound healing | 181/7539 | 2.30E-14 | 5.31E-13 |
| GO:0046777 | protein autophosphorylation | 135/7539 | 2.32E-14 | 5.35E-13 |
| GO:0007215 | glutamate receptor signaling pathway | 66/7539 | 2.36E-14 | 5.42E-13 |
| GO:0001649 | osteoblast differentiation | 119/7539 | 2.56E-14 | 5.86E-13 |
| GO:0043406 | positive regulation of MAP kinase activity | 128/7539 | 3.56E-14 | 8.12E-13 |
| GO:0010469 | regulation of signaling receptor activity | 85/7539 | 3.75E-14 | 8.49E-13 |
| GO:0015850 | organic hydroxy compound transport | 138/7539 | 3.75E-14 | 8.49E-13 |
| GO:0050730 | regulation of peptidyl-tyrosine phosphorylation | 144/7539 | 3.95E-14 | 8.91E-13 |
| GO:0099504 | synaptic vesicle cycle | 130/7539 | 4.35E-14 | 9.78E-13 |
| GO:0042326 | negative regulation of phosphorylation | 230/7539 | 4.38E-14 | 9.81E-13 |
| GO:0061333 | renal tubule morphogenesis | 62/7539 | 4.43E-14 | 9.89E-13 |
| GO:0051098 | regulation of binding | 197/7539 | 4.90E-14 | 1.09E-12 |
| GO:0008544 | epidermis development | 165/7539 | 5.08E-14 | 1.12E-12 |
| GO:0006814 | sodium ion transport | 131/7539 | 5.96E-14 | 1.32E-12 |

| | | | | |
|---|---|---|---|---|
| GO:0048771 | tissue remodeling | 114/7539 | 7.36E-14 | 1.62E-12 |
| GO:0014065 | phosphatidylinositol 3-kinase signaling | 83/7539 | 7.99E-14 | 1.75E-12 |
| GO:0021675 | nerve development | 59/7539 | 8.60E-14 | 1.88E-12 |
| GO:0098657 | import into cell | 138/7539 | 8.68E-14 | 1.89E-12 |
| GO:0031098 | stress-activated protein kinase signaling cascade | 147/7539 | 8.85E-14 | 1.92E-12 |
| GO:0051402 | neuron apoptotic process | 158/7539 | 9.19E-14 | 1.99E-12 |
| GO:0001837 | epithelial to mesenchymal transition | 90/7539 | 9.26E-14 | 1.99E-12 |
| GO:0021872 | forebrain generation of neurons | 60/7539 | 9.32E-14 | 2.00E-12 |
| GO:0099601 | regulation of neurotransmitter receptor activity | 52/7539 | 9.43E-14 | 2.02E-12 |
| GO:0032414 | positive regulation of ion transmembrane transporter activity | 77/7539 | 1.06E-13 | 2.26E-12 |
| GO:0071774 | response to fibroblast growth factor | 65/7539 | 1.19E-13 | 2.53E-12 |
| GO:0060538 | skeletal muscle organ development | 118/7539 | 1.20E-13 | 2.53E-12 |
| GO:0035051 | cardiocyte differentiation | 105/7539 | 1.20E-13 | 2.53E-12 |
| GO:0021954 | central nervous system neuron development | 66/7539 | 1.22E-13 | 2.56E-12 |
| GO:0001708 | cell fate specification | 71/7539 | 1.23E-13 | 2.59E-12 |
| GO:0048593 | camera-type eye morphogenesis | 93/7539 | 1.30E-13 | 2.71E-12 |
| GO:0008361 | regulation of cell size | 121/7539 | 1.32E-13 | 2.75E-12 |
| GO:0031214 | biomineral tissue development | 98/7539 | 1.43E-13 | 2.97E-12 |

| | | | | |
|---|---|---|---|---|
| GO:0110148 | biomineralization | 98/7539 | 1.43E-13 | 2.97E-12 |
| GO:0003231 | cardiac ventricle development | 91/7539 | 1.54E-13 | 3.17E-12 |
| GO:0060828 | regulation of canonical Wnt signaling pathway | 130/7539 | 1.61E-13 | 3.30E-12 |
| GO:0032411 | positive regulation of transporter activity | 80/7539 | 1.80E-13 | 3.69E-12 |
| GO:0050900 | leukocyte migration | 183/7539 | 1.84E-13 | 3.76E-12 |
| GO:1905330 | regulation of morphogenesis of an epithelium | 75/7539 | 2.27E-13 | 4.62E-12 |
| GO:0060411 | cardiac septum morphogenesis | 61/7539 | 2.35E-13 | 4.75E-12 |
| GO:1901379 | regulation of potassium ion transmembrane transport | 61/7539 | 2.35E-13 | 4.75E-12 |
| GO:0001755 | neural crest cell migration | 46/7539 | 2.37E-13 | 4.78E-12 |
| GO:0007188 | adenylate cyclase-modulating G protein-coupled receptor signaling pathway | 123/7539 | 2.53E-13 | 5.08E-12 |
| GO:0021510 | spinal cord development | 71/7539 | 2.56E-13 | 5.13E-12 |
| GO:1903532 | positive regulation of secretion by cell | 194/7539 | 2.64E-13 | 5.28E-12 |
| GO:0050679 | positive regulation of epithelial cell proliferation | 118/7539 | 3.00E-13 | 5.98E-12 |
| GO:0060996 | dendritic spine development | 81/7539 | 3.21E-13 | 6.38E-12 |
| GO:0021543 | pallium development | 95/7539 | 3.29E-13 | 6.52E-12 |
| GO:0002791 | regulation of peptide secretion | 201/7539 | 3.34E-13 | 6.59E-12 |
| GO:0021879 | forebrain neuron differentiation | 54/7539 | 3.38E-13 | 6.65E-12 |
| GO:0048645 | animal organ formation | 55/7539 | 3.75E-13 | 7.34E-12 |

| | | | | |
|---|---|---|---|---|
| GO:0050905 | neuromuscular process | 87/7539 | 3.82E-13 | 7.46E-12 |
| GO:0051403 | stress-activated MAPK cascade | 139/7539 | 3.83E-13 | 7.47E-12 |
| GO:1903706 | regulation of hemopoiesis | 208/7539 | 4.07E-13 | 7.91E-12 |
| GO:0072659 | protein localization to plasma membrane | 155/7539 | 4.18E-13 | 8.09E-12 |
| GO:0050433 | regulation of catecholamine secretion | 50/7539 | 5.33E-13 | 1.03E-11 |
| GO:0150115 | cell-substrate junction organization | 64/7539 | 5.69E-13 | 1.09E-11 |
| GO:0001570 | vasculogenesis | 66/7539 | 5.70E-13 | 1.09E-11 |
| GO:0010463 | mesenchymal cell proliferation | 45/7539 | 5.85E-13 | 1.12E-11 |
| GO:0050432 | catecholamine secretion | 51/7539 | 6.16E-13 | 1.17E-11 |
| GO:0043523 | regulation of neuron apoptotic process | 144/7539 | 6.22E-13 | 1.18E-11 |
| GO:0001558 | regulation of cell growth | 213/7539 | 6.37E-13 | 1.21E-11 |
| GO:0035296 | regulation of tube diameter | 103/7539 | 6.78E-13 | 1.28E-11 |
| GO:0097746 | regulation of blood vessel diameter | 103/7539 | 6.78E-13 | 1.28E-11 |
| GO:0043588 | skin development | 146/7539 | 6.90E-13 | 1.30E-11 |
| GO:1901343 | negative regulation of vasculature development | 86/7539 | 7.41E-13 | 1.39E-11 |
| GO:0030282 | bone mineralization | 77/7539 | 7.73E-13 | 1.44E-11 |
| GO:0110053 | regulation of actin filament organization | 145/7539 | 7.91E-13 | 1.47E-11 |
| GO:0002065 | columnar/cuboidal epithelial cell differentiation | 91/7539 | 8.32E-13 | 1.54E-11 |

| | | | | |
|---|---|---|---|---|
| GO:0010634 | positive regulation of epithelial cell migration | 91/7539 | 8.32E-13 | 1.54E-11 |
| GO:0030178 | negative regulation of Wnt signaling pathway | 96/7539 | 8.65E-13 | 1.60E-11 |
| GO:0055002 | striated muscle cell development | 109/7539 | 8.89E-13 | 1.64E-11 |
| GO:0048665 | neuron fate specification | 33/7539 | 9.11E-13 | 1.67E-11 |
| GO:0007156 | homophilic cell adhesion via plasma membrane adhesion molecules | 72/7539 | 9.98E-13 | 1.83E-11 |
| GO:0035150 | regulation of tube size | 103/7539 | 1.11E-12 | 2.02E-11 |
| GO:0050731 | positive regulation of peptidyl-tyrosine phosphorylation | 111/7539 | 1.11E-12 | 2.02E-11 |
| GO:0048546 | digestive tract morphogenesis | 43/7539 | 1.12E-12 | 2.03E-11 |
| GO:0072028 | nephron morphogenesis | 59/7539 | 1.14E-12 | 2.07E-11 |
| GO:0010594 | regulation of endothelial cell migration | 98/7539 | 1.17E-12 | 2.11E-11 |
| GO:0097581 | lamellipodium organization | 60/7539 | 1.18E-12 | 2.12E-11 |
| GO:0030510 | regulation of BMP signaling pathway | 66/7539 | 1.19E-12 | 2.14E-11 |
| GO:0048706 | embryonic skeletal system development | 87/7539 | 1.20E-12 | 2.15E-11 |
| GO:0060021 | roof of mouth development | 65/7539 | 1.21E-12 | 2.16E-11 |
| GO:0007044 | cell-substrate junction assembly | 62/7539 | 1.22E-12 | 2.16E-11 |
| GO:0071692 | protein localization to extracellular region | 228/7539 | 1.22E-12 | 2.16E-11 |
| GO:0032872 | regulation of stress-activated MAPK cascade | 127/7539 | 1.26E-12 | 2.23E-11 |
| GO:0030534 | adult behavior | 106/7539 | 1.27E-12 | 2.23E-11 |

| | | | | |
|---|---|---|---|---|
| GO:1900449 | regulation of glutamate receptor signaling pathway | 44/7539 | 1.44E-12 | 2.52E-11 |
| GO:0006940 | regulation of smooth muscle contraction | 50/7539 | 1.44E-12 | 2.52E-11 |
| GO:0007219 | Notch signaling pathway | 104/7539 | 1.60E-12 | 2.80E-11 |
| GO:0070302 | regulation of stress-activated protein kinase signaling cascade | 128/7539 | 1.66E-12 | 2.89E-11 |
| GO:0030099 | myeloid cell differentiation | 202/7539 | 1.69E-12 | 2.94E-11 |
| GO:0046928 | regulation of neurotransmitter secretion | 83/7539 | 1.70E-12 | 2.94E-11 |
| GO:0007179 | transforming growth factor beta receptor signaling pathway | 99/7539 | 1.74E-12 | 3.00E-11 |
| GO:0030509 | BMP signaling pathway | 98/7539 | 1.94E-12 | 3.34E-11 |
| GO:1902105 | regulation of leukocyte differentiation | 159/7539 | 1.96E-12 | 3.38E-11 |
| GO:0009306 | protein secretion | 224/7539 | 2.12E-12 | 3.64E-11 |
| GO:0050920 | regulation of chemotaxis | 123/7539 | 2.13E-12 | 3.65E-11 |
| GO:0048660 | regulation of smooth muscle cell proliferation | 97/7539 | 2.16E-12 | 3.69E-11 |
| GO:0035592 | establishment of protein localization to extracellular region | 224/7539 | 2.76E-12 | 4.70E-11 |
| GO:0048659 | smooth muscle cell proliferation | 99/7539 | 2.84E-12 | 4.82E-11 |
| GO:0099084 | postsynaptic specialization organization | 35/7539 | 3.03E-12 | 5.13E-11 |
| GO:0060401 | cytosolic calcium ion transport | 102/7539 | 3.25E-12 | 5.49E-11 |
| GO:0006887 | exocytosis | 191/7539 | 3.33E-12 | 5.60E-11 |
| GO:0060402 | calcium ion transport into cytosol | 93/7539 | 3.33E-12 | 5.60E-11 |

| | | | | |
|---|---|---|---|---|
| GO:1903531 | negative regulation of secretion by cell | 109/7539 | 3.51E-12 | 5.88E-11 |
| GO:0048640 | negative regulation of developmental growth | 81/7539 | 3.58E-12 | 5.97E-11 |
| GO:0070374 | positive regulation of ERK1 and ERK2 cascade | 119/7539 | 3.59E-12 | 5.99E-11 |
| GO:0038127 | ERBB signaling pathway | 72/7539 | 3.72E-12 | 6.18E-11 |
| GO:0007405 | neuroblast proliferation | 51/7539 | 4.08E-12 | 6.77E-11 |
| GO:0045055 | regulated exocytosis | 138/7539 | 4.34E-12 | 7.17E-11 |
| GO:0051047 | positive regulation of secretion | 211/7539 | 4.45E-12 | 7.33E-11 |
| GO:1901861 | regulation of muscle tissue development | 103/7539 | 4.62E-12 | 7.60E-11 |
| GO:0010771 | negative regulation of cell morphogenesis involved in differentiation | 66/7539 | 4.94E-12 | 8.10E-11 |
| GO:0001935 | endothelial cell proliferation | 89/7539 | 5.06E-12 | 8.28E-11 |
| GO:0008038 | neuron recognition | 40/7539 | 5.29E-12 | 8.62E-11 |
| GO:0048041 | focal adhesion assembly | 56/7539 | 5.34E-12 | 8.65E-11 |
| GO:0072078 | nephron tubule morphogenesis | 56/7539 | 5.34E-12 | 8.65E-11 |
| GO:2001259 | positive regulation of cation channel activity | 56/7539 | 5.34E-12 | 8.65E-11 |
| GO:0072088 | nephron epithelium morphogenesis | 57/7539 | 5.47E-12 | 8.83E-11 |
| GO:0043271 | negative regulation of ion transport | 100/7539 | 6.60E-12 | 1.06E-10 |
| GO:0097106 | postsynaptic density organization | 31/7539 | 6.98E-12 | 1.12E-10 |
| GO:0048634 | regulation of muscle organ development | 103/7539 | 7.33E-12 | 1.17E-10 |

| | | | | |
|---|---|---|---|---|
| GO:0010811 | positive regulation of cell-substrate adhesion | 79/7539 | 7.53E-12 | 1.20E-10 |
| GO:0060249 | anatomical structure homeostasis | 203/7539 | 7.88E-12 | 1.26E-10 |
| GO:0030048 | actin filament-based movement | 78/7539 | 8.20E-12 | 1.30E-10 |
| GO:0007272 | ensheathment of neurons | 98/7539 | 8.35E-12 | 1.32E-10 |
| GO:0008366 | axon ensheathment | 98/7539 | 8.35E-12 | 1.32E-10 |
| GO:0021536 | diencephalon development | 50/7539 | 9.28E-12 | 1.46E-10 |
| GO:0016202 | regulation of striated muscle tissue development | 101/7539 | 9.33E-12 | 1.47E-10 |
| GO:0090276 | regulation of peptide hormone secretion | 126/7539 | 1.05E-11 | 1.64E-10 |
| GO:0019233 | sensory perception of pain | 87/7539 | 1.05E-11 | 1.64E-10 |
| GO:0035637 | multicellular organismal signaling | 87/7539 | 1.05E-11 | 1.64E-10 |
| GO:0014015 | positive regulation of gliogenesis | 61/7539 | 1.15E-11 | 1.80E-10 |
| GO:0060078 | regulation of postsynaptic membrane potential | 72/7539 | 1.30E-11 | 2.03E-10 |
| GO:0007519 | skeletal muscle tissue development | 109/7539 | 1.31E-11 | 2.03E-10 |
| GO:0070304 | positive regulation of stress-activated protein kinase signaling cascade | 98/7539 | 1.34E-11 | 2.07E-10 |
| GO:0032231 | regulation of actin filament bundle assembly | 70/7539 | 1.49E-11 | 2.30E-10 |
| GO:0048709 | oligodendrocyte differentiation | 70/7539 | 1.49E-11 | 2.30E-10 |
| GO:0032874 | positive regulation of stress-activated MAPK cascade | 97/7539 | 1.51E-11 | 2.31E-10 |
| GO:0001508 | action potential | 83/7539 | 1.57E-11 | 2.41E-10 |

| | | | | |
|---|---|---|---|---|
| GO:0048675 | axon extension | 83/7539 | 1.57E-11 | 2.41E-10 |
| GO:0007613 | memory | 88/7539 | 1.58E-11 | 2.41E-10 |
| GO:1902414 | protein localization to cell junction | 77/7539 | 1.59E-11 | 2.41E-10 |
| GO:0042552 | myelination | 96/7539 | 1.70E-11 | 2.57E-10 |
| GO:0051928 | positive regulation of calcium ion transport | 82/7539 | 1.74E-11 | 2.63E-10 |
| GO:0060349 | bone morphogenesis | 67/7539 | 1.79E-11 | 2.71E-10 |
| GO:0050708 | regulation of protein secretion | 179/7539 | 1.80E-11 | 2.71E-10 |
| GO:0030010 | establishment of cell polarity | 85/7539 | 2.17E-11 | 3.26E-10 |
| GO:0010770 | positive regulation of cell morphogenesis involved in differentiation | 105/7539 | 2.17E-11 | 3.26E-10 |
| GO:0060998 | regulation of dendritic spine development | 63/7539 | 2.18E-11 | 3.26E-10 |
| GO:0010717 | regulation of epithelial to mesenchymal transition | 62/7539 | 2.26E-11 | 3.38E-10 |
| GO:0072171 | mesonephric tubule morphogenesis | 51/7539 | 2.34E-11 | 3.48E-10 |
| GO:0048259 | regulation of receptor-mediated endocytosis | 72/7539 | 2.39E-11 | 3.54E-10 |
| GO:0016079 | synaptic vesicle exocytosis | 84/7539 | 2.41E-11 | 3.57E-10 |
| GO:0008543 | fibroblast growth factor receptor signaling pathway | 53/7539 | 2.49E-11 | 3.68E-10 |
| GO:1901019 | regulation of calcium ion transmembrane transporter activity | 56/7539 | 2.56E-11 | 3.78E-10 |
| GO:0098727 | maintenance of cell number | 99/7539 | 2.96E-11 | 4.36E-10 |
| GO:0110020 | regulation of actomyosin structure organization | 68/7539 | 3.18E-11 | 4.66E-10 |

| | | | | |
|---|---|---|---|---|
| GO:0015711 | organic anion transport | 217/7539 | 3.39E-11 | 4.96E-10 |
| GO:0043491 | protein kinase B signaling | 108/7539 | 3.46E-11 | 5.06E-10 |
| GO:0001841 | neural tube formation | 80/7539 | 3.62E-11 | 5.28E-10 |
| GO:0001666 | response to hypoxia | 113/7539 | 4.06E-11 | 5.90E-10 |
| GO:0150116 | regulation of cell-substrate junction organization | 46/7539 | 4.08E-11 | 5.93E-10 |
| GO:0007187 | G protein-coupled receptor signaling pathway, coupled to cyclic nucleotide second messenger | 129/7539 | 4.47E-11 | 6.47E-10 |
| GO:0043268 | positive regulation of potassium ion transport | 40/7539 | 5.01E-11 | 7.22E-10 |
| GO:0048483 | autonomic nervous system development | 40/7539 | 5.01E-11 | 7.22E-10 |
| GO:0060675 | ureteric bud morphogenesis | 50/7539 | 5.16E-11 | 7.42E-10 |
| GO:0097553 | calcium ion transmembrane import into cytosol | 81/7539 | 5.54E-11 | 7.95E-10 |
| GO:0060840 | artery development | 68/7539 | 5.87E-11 | 8.40E-10 |
| GO:0035272 | exocrine system development | 41/7539 | 5.93E-11 | 8.46E-10 |
| GO:0060041 | retina development in camera-type eye | 97/7539 | 5.94E-11 | 8.46E-10 |
| GO:0035264 | multicellular organism growth | 116/7539 | 5.96E-11 | 8.47E-10 |
| GO:1905475 | regulation of protein localization to membrane | 107/7539 | 5.97E-11 | 8.47E-10 |
| GO:0062197 | cellular response to chemical stress | 164/7539 | 6.10E-11 | 8.64E-10 |
| GO:0031532 | actin cytoskeleton reorganization | 67/7539 | 6.31E-11 | 8.89E-10 |
| GO:0034446 | substrate adhesion-dependent cell spreading | 67/7539 | 6.31E-11 | 8.89E-10 |

| | | | | |
|---|---|---|---|---|
| GO:1903305 | regulation of regulated secretory pathway | 106/7539 | 6.80E-11 | 9.57E-10 |
| GO:0043534 | blood vessel endothelial cell migration | 73/7539 | 7.01E-11 | 9.84E-10 |
| GO:0048469 | cell maturation | 111/7539 | 7.89E-11 | 1.10E-09 |
| GO:0043393 | regulation of protein binding | 122/7539 | 8.15E-11 | 1.14E-09 |
| GO:0061387 | regulation of extent of cell growth | 77/7539 | 8.32E-11 | 1.16E-09 |
| GO:0051893 | regulation of focal adhesion assembly | 44/7539 | 8.55E-11 | 1.19E-09 |
| GO:0090109 | regulation of cell-substrate junction assembly | 44/7539 | 8.55E-11 | 1.19E-09 |
| GO:2000181 | negative regulation of blood vessel morphogenesis | 76/7539 | 9.19E-11 | 1.27E-09 |
| GO:0021846 | cell proliferation in forebrain | 30/7539 | 9.26E-11 | 1.28E-09 |
| GO:0051058 | negative regulation of small GTPase mediated signal transduction | 45/7539 | 9.30E-11 | 1.28E-09 |
| GO:0030073 | insulin secretion | 121/7539 | 9.35E-11 | 1.29E-09 |
| GO:0022409 | positive regulation of cell-cell adhesion | 136/7539 | 9.63E-11 | 1.32E-09 |
| GO:0043547 | positive regulation of GTPase activity | 118/7539 | 9.72E-11 | 1.33E-09 |
| GO:0046888 | negative regulation of hormone secretion | 59/7539 | 9.99E-11 | 1.37E-09 |
| GO:0042475 | odontogenesis of dentin-containing tooth | 58/7539 | 1.04E-10 | 1.42E-09 |
| GO:0019827 | stem cell population maintenance | 96/7539 | 1.05E-10 | 1.43E-09 |
| GO:0003281 | ventricular septum development | 57/7539 | 1.08E-10 | 1.46E-09 |
| GO:0030032 | lamellipodium assembly | 48/7539 | 1.10E-10 | 1.49E-09 |

| | | | | |
|---|---|---|---|---|
| GO:0032102 | negative regulation of response to external stimulus | 184/7539 | 1.11E-10 | 1.50E-09 |
| GO:0002573 | myeloid leukocyte differentiation | 117/7539 | 1.11E-10 | 1.50E-09 |
| GO:2000179 | positive regulation of neural precursor cell proliferation | 49/7539 | 1.13E-10 | 1.52E-09 |
| GO:0030500 | regulation of bone mineralization | 55/7539 | 1.14E-10 | 1.52E-09 |
| GO:0050805 | negative regulation of synaptic transmission | 55/7539 | 1.14E-10 | 1.52E-09 |
| GO:0030501 | positive regulation of bone mineralization | 34/7539 | 1.17E-10 | 1.57E-09 |
| GO:0015849 | organic acid transport | 165/7539 | 1.28E-10 | 1.71E-09 |
| GO:0007173 | epidermal growth factor receptor signaling pathway | 65/7539 | 1.34E-10 | 1.78E-09 |
| GO:0017157 | regulation of exocytosis | 131/7539 | 1.38E-10 | 1.83E-09 |
| GO:0048713 | regulation of oligodendrocyte differentiation | 40/7539 | 1.40E-10 | 1.86E-09 |
| GO:0099560 | synaptic membrane adhesion | 28/7539 | 1.44E-10 | 1.90E-09 |
| GO:0008217 | regulation of blood pressure | 112/7539 | 1.50E-10 | 1.99E-09 |
| GO:0021761 | limbic system development | 63/7539 | 1.54E-10 | 2.02E-09 |
| GO:0070167 | regulation of biomineral tissue development | 63/7539 | 1.54E-10 | 2.02E-09 |
| GO:0110149 | regulation of biomineralization | 63/7539 | 1.54E-10 | 2.02E-09 |
| GO:0050919 | negative chemotaxis | 35/7539 | 1.55E-10 | 2.04E-09 |
| GO:0030856 | regulation of epithelial cell differentiation | 89/7539 | 1.58E-10 | 2.07E-09 |
| GO:0007498 | mesoderm development | 70/7539 | 1.61E-10 | 2.10E-09 |

| | | | | |
|---|---|---|---|---|
| GO:1902904 | negative regulation of supramolecular fiber organization | 92/7539 | 1.73E-10 | 2.25E-09 |
| GO:0051099 | positive regulation of binding | 108/7539 | 1.75E-10 | 2.27E-09 |
| GO:0030038 | contractile actin filament bundle assembly | 69/7539 | 1.76E-10 | 2.27E-09 |
| GO:0043149 | stress fiber assembly | 69/7539 | 1.76E-10 | 2.27E-09 |
| GO:0001936 | regulation of endothelial cell proliferation | 79/7539 | 1.91E-10 | 2.46E-09 |
| GO:0009791 | post-embryonic development | 79/7539 | 1.91E-10 | 2.46E-09 |
| GO:0072091 | regulation of stem cell proliferation | 59/7539 | 1.95E-10 | 2.50E-09 |
| GO:0045216 | cell-cell junction organization | 110/7539 | 1.97E-10 | 2.53E-09 |
| GO:0006979 | response to oxidative stress | 193/7539 | 2.01E-10 | 2.58E-09 |
| GO:0021987 | cerebral cortex development | 67/7539 | 2.08E-10 | 2.66E-09 |
| GO:0034103 | regulation of tissue remodeling | 57/7539 | 2.14E-10 | 2.73E-09 |
| GO:0021983 | pituitary gland development | 32/7539 | 2.15E-10 | 2.74E-09 |
| GO:0014812 | muscle cell migration | 66/7539 | 2.26E-10 | 2.87E-09 |
| GO:0001738 | morphogenesis of a polarized epithelium | 55/7539 | 2.31E-10 | 2.93E-09 |
| GO:0021515 | cell differentiation in spinal cord | 46/7539 | 2.34E-10 | 2.96E-09 |
| GO:0061035 | regulation of cartilage development | 49/7539 | 2.51E-10 | 3.17E-09 |
| GO:1903670 | regulation of sprouting angiogenesis | 51/7539 | 2.51E-10 | 3.17E-09 |
| GO:0099565 | chemical synaptic transmission, postsynaptic | 62/7539 | 3.06E-10 | 3.84E-09 |

| | | | | |
|---|---|---|---|---|
| GO:0030516 | regulation of axon extension | 69/7539 | 3.09E-10 | 3.88E-09 |
| GO:0045933 | positive regulation of muscle contraction | 39/7539 | 3.30E-10 | 4.13E-09 |
| GO:0007369 | gastrulation | 97/7539 | 3.34E-10 | 4.17E-09 |
| GO:0045926 | negative regulation of growth | 134/7539 | 3.44E-10 | 4.29E-09 |
| GO:0046942 | carboxylic acid transport | 162/7539 | 3.46E-10 | 4.31E-09 |
| GO:0042303 | molting cycle | 78/7539 | 3.53E-10 | 4.37E-09 |
| GO:0042633 | hair cycle | 78/7539 | 3.53E-10 | 4.37E-09 |
| GO:0016525 | negative regulation of angiogenesis | 73/7539 | 3.60E-10 | 4.46E-09 |
| GO:0035249 | synaptic transmission, glutamatergic | 67/7539 | 3.70E-10 | 4.57E-09 |
| GO:0015872 | dopamine transport | 41/7539 | 4.10E-10 | 5.04E-09 |
| GO:0045599 | negative regulation of fat cell differentiation | 41/7539 | 4.10E-10 | 5.04E-09 |
| GO:0008593 | regulation of Notch signaling pathway | 57/7539 | 4.16E-10 | 5.10E-09 |
| GO:0060079 | excitatory postsynaptic potential | 57/7539 | 4.16E-10 | 5.10E-09 |
| GO:0007254 | JNK cascade | 107/7539 | 4.35E-10 | 5.32E-09 |
| GO:1903034 | regulation of response to wounding | 101/7539 | 4.46E-10 | 5.44E-09 |
| GO:0006869 | lipid transport | 174/7539 | 4.47E-10 | 5.44E-09 |
| GO:0045667 | regulation of osteoblast differentiation | 84/7539 | 4.59E-10 | 5.58E-09 |
| GO:0006941 | striated muscle contraction | 91/7539 | 4.72E-10 | 5.73E-09 |

| | | | | |
|---|---|---|---|---|
| GO:1901016 | regulation of potassium ion transmembrane transporter activity | 43/7539 | 4.74E-10 | 5.74E-09 |
| GO:0050772 | positive regulation of axonogenesis | 64/7539 | 4.78E-10 | 5.78E-09 |
| GO:0051090 | regulation of DNA-binding transcription factor activity | 194/7539 | 4.80E-10 | 5.79E-09 |
| GO:0010464 | regulation of mesenchymal cell proliferation | 35/7539 | 4.81E-10 | 5.79E-09 |
| GO:0014902 | myotube differentiation | 75/7539 | 4.90E-10 | 5.88E-09 |
| GO:0060048 | cardiac muscle contraction | 75/7539 | 4.90E-10 | 5.88E-09 |
| GO:0051965 | positive regulation of synapse assembly | 53/7539 | 4.96E-10 | 5.94E-09 |
| GO:0071375 | cellular response to peptide hormone stimulus | 129/7539 | 5.00E-10 | 5.98E-09 |
| GO:0008360 | regulation of cell shape | 87/7539 | 5.02E-10 | 5.99E-09 |
| GO:0003208 | cardiac ventricle morphogenesis | 52/7539 | 5.11E-10 | 6.07E-09 |
| GO:0060976 | coronary vasculature development | 52/7539 | 5.11E-10 | 6.07E-09 |
| GO:0033555 | multicellular organismal response to stress | 63/7539 | 5.18E-10 | 6.14E-09 |
| GO:0018209 | peptidyl-serine modification | 171/7539 | 5.18E-10 | 6.14E-09 |
| GO:0007422 | peripheral nervous system development | 50/7539 | 5.33E-10 | 6.30E-09 |
| GO:0036293 | response to decreased oxygen levels | 121/7539 | 5.56E-10 | 6.56E-09 |
| GO:0014066 | regulation of phosphatidylinositol 3-kinase signaling | 62/7539 | 5.60E-10 | 6.60E-09 |
| GO:0003179 | heart valve morphogenesis | 36/7539 | 5.80E-10 | 6.80E-09 |
| GO:0003197 | endocardial cushion development | 36/7539 | 5.80E-10 | 6.80E-09 |

| | | | | |
|---|---|---|---|---|
| GO:0043524 | negative regulation of neuron apoptotic process | 99/7539 | 5.81E-10 | 6.81E-09 |
| GO:0007548 | sex differentiation | 154/7539 | 5.99E-10 | 7.00E-09 |
| GO:0007159 | leukocyte cell-cell adhesion | 166/7539 | 6.04E-10 | 7.04E-09 |
| GO:0099072 | regulation of postsynaptic membrane neurotransmitter receptor levels | 61/7539 | 6.04E-10 | 7.04E-09 |
| GO:0051896 | regulation of protein kinase B signaling | 92/7539 | 6.38E-10 | 7.41E-09 |
| GO:0009799 | specification of symmetry | 81/7539 | 6.57E-10 | 7.63E-09 |
| GO:0022037 | metencephalon development | 72/7539 | 6.74E-10 | 7.81E-09 |
| GO:0002028 | regulation of sodium ion transport | 59/7539 | 6.97E-10 | 8.06E-09 |
| GO:0048704 | embryonic skeletal system morphogenesis | 66/7539 | 7.11E-10 | 8.21E-09 |
| GO:0007631 | feeding behavior | 76/7539 | 7.24E-10 | 8.34E-09 |
| GO:0051494 | negative regulation of cytoskeleton organization | 94/7539 | 7.48E-10 | 8.60E-09 |
| GO:0032330 | regulation of chondrocyte differentiation | 38/7539 | 7.71E-10 | 8.85E-09 |
| GO:0050863 | regulation of T cell activation | 156/7539 | 8.02E-10 | 9.19E-09 |
| GO:0010038 | response to metal ion | 137/7539 | 8.05E-10 | 9.20E-09 |
| GO:0035418 | protein localization to synapse | 63/7539 | 9.28E-10 | 1.06E-08 |
| GO:0051656 | establishment of organelle localization | 180/7539 | 9.33E-10 | 1.06E-08 |
| GO:0046580 | negative regulation of Ras protein signal transduction | 40/7539 | 9.36E-10 | 1.06E-08 |
| GO:0001709 | cell fate determination | 33/7539 | 9.48E-10 | 1.08E-08 |

| | | | | |
|---|---|---|---|---|
| GO:0003170 | heart valve development | 41/7539 | 1.00E-09 | 1.14E-08 |
| GO:0007528 | neuromuscular junction development | 42/7539 | 1.06E-09 | 1.20E-08 |
| GO:0046328 | regulation of JNK cascade | 100/7539 | 1.12E-09 | 1.27E-08 |
| GO:0002090 | regulation of receptor internalization | 49/7539 | 1.12E-09 | 1.27E-08 |
| GO:0007189 | adenylate cyclase-activating G protein-coupled receptor signaling pathway | 80/7539 | 1.18E-09 | 1.33E-08 |
| GO:0009855 | determination of bilateral symmetry | 80/7539 | 1.18E-09 | 1.33E-08 |
| GO:0048846 | axon extension involved in axon guidance | 30/7539 | 1.39E-09 | 1.56E-08 |
| GO:1902284 | neuron projection extension involved in neuron projection guidance | 30/7539 | 1.39E-09 | 1.56E-08 |
| GO:0014020 | primary neural tube formation | 70/7539 | 1.40E-09 | 1.56E-08 |
| GO:0009953 | dorsal/ventral pattern formation | 64/7539 | 1.49E-09 | 1.66E-08 |
| GO:0035050 | embryonic heart tube development | 55/7539 | 1.69E-09 | 1.89E-08 |
| GO:0031099 | regeneration | 84/7539 | 1.77E-09 | 1.97E-08 |
| GO:0060491 | regulation of cell projection assembly | 102/7539 | 1.83E-09 | 2.03E-08 |
| GO:0006898 | receptor-mediated endocytosis | 122/7539 | 1.85E-09 | 2.05E-08 |
| GO:0045137 | development of primary sexual characteristics | 122/7539 | 1.85E-09 | 2.05E-08 |
| GO:1903037 | regulation of leukocyte cell-cell adhesion | 150/7539 | 1.86E-09 | 2.05E-08 |
| GO:0030217 | T cell differentiation | 139/7539 | 2.04E-09 | 2.25E-08 |
| GO:0046620 | regulation of organ growth | 71/7539 | 2.08E-09 | 2.29E-08 |

| | | | | |
|---|---|---|---|---|
| GO:0008344 | adult locomotory behavior | 66/7539 | 2.11E-09 | 2.32E-08 |
| GO:0045687 | positive regulation of glial cell differentiation | 39/7539 | 2.12E-09 | 2.33E-08 |
| GO:0009612 | response to mechanical stimulus | 95/7539 | 2.20E-09 | 2.40E-08 |
| GO:0048486 | parasympathetic nervous system development | 20/7539 | 2.21E-09 | 2.41E-08 |
| GO:0051492 | regulation of stress fiber assembly | 59/7539 | 2.32E-09 | 2.53E-08 |
| GO:0001736 | establishment of planar polarity | 41/7539 | 2.35E-09 | 2.56E-08 |
| GO:0051592 | response to calcium ion | 74/7539 | 2.41E-09 | 2.62E-08 |
| GO:0003143 | embryonic heart tube morphogenesis | 47/7539 | 2.42E-09 | 2.62E-08 |
| GO:0008406 | gonad development | 120/7539 | 2.45E-09 | 2.66E-08 |
| GO:0050771 | negative regulation of axonogenesis | 46/7539 | 2.46E-09 | 2.66E-08 |
| GO:0032835 | glomerulus development | 43/7539 | 2.47E-09 | 2.66E-08 |
| GO:0046850 | regulation of bone remodeling | 43/7539 | 2.47E-09 | 2.66E-08 |
| GO:0001658 | branching involved in ureteric bud morphogenesis | 44/7539 | 2.49E-09 | 2.68E-08 |
| GO:0030098 | lymphocyte differentiation | 189/7539 | 2.50E-09 | 2.69E-08 |
| GO:0001890 | placenta development | 94/7539 | 2.51E-09 | 2.69E-08 |
| GO:1902905 | positive regulation of supramolecular fiber organization | 105/7539 | 2.53E-09 | 2.71E-08 |
| GO:0048644 | muscle organ morphogenesis | 57/7539 | 2.72E-09 | 2.91E-08 |
| GO:1901381 | positive regulation of potassium ion transmembrane transport | 33/7539 | 2.81E-09 | 2.99E-08 |

| | | | | |
|---|---|---|---|---|
| GO:0060326 | cell chemotaxis | 147/7539 | 2.82E-09 | 3.00E-08 |
| GO:0003148 | outflow tract septum morphogenesis | 25/7539 | 2.86E-09 | 3.04E-08 |
| GO:0051147 | regulation of muscle cell differentiation | 90/7539 | 2.87E-09 | 3.05E-08 |
| GO:0043434 | response to peptide hormone | 160/7539 | 3.11E-09 | 3.30E-08 |
| GO:0048511 | rhythmic process | 146/7539 | 3.25E-09 | 3.43E-08 |
| GO:0046330 | positive regulation of JNK cascade | 79/7539 | 3.32E-09 | 3.50E-08 |
| GO:1904018 | positive regulation of vasculature development | 108/7539 | 3.40E-09 | 3.58E-08 |
| GO:0090101 | negative regulation of transmembrane receptor protein serine/threonine kinase signaling pathway | 71/7539 | 3.40E-09 | 3.58E-08 |
| GO:0007368 | determination of left/right symmetry | 75/7539 | 3.43E-09 | 3.60E-08 |
| GO:0010876 | lipid localization | 190/7539 | 3.50E-09 | 3.67E-08 |
| GO:0008045 | motor neuron axon guidance | 29/7539 | 3.52E-09 | 3.68E-08 |
| GO:0051495 | positive regulation of cytoskeleton organization | 115/7539 | 3.55E-09 | 3.71E-08 |
| GO:0001843 | neural tube closure | 66/7539 | 3.57E-09 | 3.73E-08 |
| GO:1904427 | positive regulation of calcium ion transmembrane transport | 53/7539 | 3.62E-09 | 3.77E-08 |
| GO:0032869 | cellular response to insulin stimulus | 105/7539 | 3.62E-09 | 3.77E-08 |
| GO:0097756 | negative regulation of blood vessel diameter | 65/7539 | 3.97E-09 | 4.12E-08 |
| GO:0007157 | heterophilic cell-cell adhesion via plasma membrane cell adhesion molecules | 36/7539 | 4.13E-09 | 4.29E-08 |
| GO:1990845 | adaptive thermogenesis | 87/7539 | 4.27E-09 | 4.42E-08 |

| | | | | |
|---|---|---|---|---|
| GO:0030901 | midbrain development | 37/7539 | 4.49E-09 | 4.63E-08 |
| GO:0072132 | mesenchyme morphogenesis | 37/7539 | 4.49E-09 | 4.63E-08 |
| GO:0070252 | actin-mediated cell contraction | 58/7539 | 4.50E-09 | 4.64E-08 |
| GO:0051145 | smooth muscle cell differentiation | 49/7539 | 4.52E-09 | 4.65E-08 |
| GO:0098810 | neurotransmitter reuptake | 26/7539 | 4.54E-09 | 4.66E-08 |
| GO:0035725 | sodium ion transmembrane transport | 80/7539 | 4.55E-09 | 4.66E-08 |
| GO:0097120 | receptor localization to synapse | 48/7539 | 4.72E-09 | 4.83E-08 |
| GO:0002792 | negative regulation of peptide secretion | 68/7539 | 4.78E-09 | 4.87E-08 |
| GO:0098773 | skin epidermis development | 68/7539 | 4.78E-09 | 4.87E-08 |
| GO:0014909 | smooth muscle cell migration | 57/7539 | 4.90E-09 | 4.99E-08 |
| GO:0120032 | regulation of plasma membrane bounded cell projection assembly | 100/7539 | 5.04E-09 | 5.13E-08 |
| GO:0035567 | non-canonical Wnt signaling pathway | 40/7539 | 5.19E-09 | 5.26E-08 |
| GO:0043254 | regulation of protein-containing complex assembly | 199/7539 | 5.22E-09 | 5.28E-08 |
| GO:0007164 | establishment of tissue polarity | 41/7539 | 5.30E-09 | 5.36E-08 |
| GO:0019226 | transmission of nerve impulse | 56/7539 | 5.33E-09 | 5.38E-08 |
| GO:0010595 | positive regulation of endothelial cell migration | 62/7539 | 5.39E-09 | 5.43E-08 |
| GO:0097529 | myeloid leukocyte migration | 112/7539 | 5.41E-09 | 5.45E-08 |
| GO:0051261 | protein depolymerization | 71/7539 | 5.51E-09 | 5.54E-08 |

| | | | | |
|---|---|---|---|---|
| GO:0001504 | neurotransmitter uptake | 31/7539 | 5.64E-09 | 5.65E-08 |
| GO:0040019 | positive regulation of embryonic development | 31/7539 | 5.64E-09 | 5.65E-08 |
| GO:0055007 | cardiac muscle cell differentiation | 78/7539 | 5.86E-09 | 5.86E-08 |
| GO:0060606 | tube closure | 66/7539 | 5.96E-09 | 5.95E-08 |
| GO:0001659 | temperature homeostasis | 96/7539 | 6.04E-09 | 6.03E-08 |
| GO:0018105 | peptidyl-serine phosphorylation | 157/7539 | 6.13E-09 | 6.10E-08 |
| GO:1901652 | response to peptide | 184/7539 | 6.22E-09 | 6.18E-08 |
| GO:0097061 | dendritic spine organization | 60/7539 | 6.56E-09 | 6.51E-08 |
| GO:0002027 | regulation of heart rate | 59/7539 | 7.22E-09 | 7.15E-08 |
| GO:0055017 | cardiac muscle tissue growth | 64/7539 | 7.40E-09 | 7.32E-08 |
| GO:1901653 | cellular response to peptide | 144/7539 | 7.48E-09 | 7.39E-08 |
| GO:0098698 | postsynaptic specialization assembly | 24/7539 | 7.66E-09 | 7.56E-08 |
| GO:1903523 | negative regulation of blood circulation | 33/7539 | 7.73E-09 | 7.61E-08 |
| GO:0048167 | regulation of synaptic plasticity | 170/7539 | 7.75E-09 | 7.63E-08 |
| GO:0060563 | neuroepithelial cell differentiation | 50/7539 | 8.23E-09 | 8.06E-08 |
| GO:0035282 | segmentation | 63/7539 | 8.23E-09 | 8.06E-08 |
| GO:0110110 | positive regulation of animal organ morphogenesis | 63/7539 | 8.23E-09 | 8.06E-08 |
| GO:0032868 | response to insulin | 118/7539 | 8.51E-09 | 8.32E-08 |

| | | | | |
|---|---|---|---|---|
| GO:0046849 | bone remodeling | 62/7539 | 9.14E-09 | 8.92E-08 |
| GO:0031623 | receptor internalization | 74/7539 | 9.66E-09 | 9.42E-08 |
| GO:0001942 | hair follicle development | 66/7539 | 9.81E-09 | 9.54E-08 |
| GO:0050848 | regulation of calcium-mediated signaling | 61/7539 | 1.01E-08 | 9.80E-08 |
| GO:0050885 | neuromuscular process controlling balance | 46/7539 | 1.01E-08 | 9.80E-08 |
| GO:0021545 | cranial nerve development | 36/7539 | 1.02E-08 | 9.80E-08 |
| GO:0070169 | positive regulation of biomineral tissue development | 36/7539 | 1.02E-08 | 9.80E-08 |
| GO:0110151 | positive regulation of biomineralization | 36/7539 | 1.02E-08 | 9.80E-08 |
| GO:2001222 | regulation of neuron migration | 38/7539 | 1.11E-08 | 1.07E-07 |
| GO:0035924 | cellular response to vascular endothelial growth factor stimulus | 39/7539 | 1.14E-08 | 1.09E-07 |
| GO:0021575 | hindbrain morphogenesis | 41/7539 | 1.15E-08 | 1.10E-07 |
| GO:0050796 | regulation of insulin secretion | 99/7539 | 1.19E-08 | 1.14E-07 |
| GO:0051668 | localization within membrane | 64/7539 | 1.23E-08 | 1.18E-07 |
| GO:0090090 | negative regulation of canonical Wnt signaling pathway | 72/7539 | 1.24E-08 | 1.18E-07 |
| GO:0030279 | negative regulation of ossification | 59/7539 | 1.24E-08 | 1.19E-07 |
| GO:1904019 | epithelial cell apoptotic process | 68/7539 | 1.26E-08 | 1.20E-07 |
| GO:0045862 | positive regulation of proteolysis | 161/7539 | 1.26E-08 | 1.20E-07 |
| GO:0060415 | muscle tissue morphogenesis | 52/7539 | 1.33E-08 | 1.26E-07 |

| | | | | |
|---|---|---|---|---|
| GO:0045766 | positive regulation of angiogenesis | 98/7539 | 1.36E-08 | 1.29E-07 |
| GO:0061053 | somite development | 58/7539 | 1.37E-08 | 1.30E-07 |
| GO:0002685 | regulation of leukocyte migration | 110/7539 | 1.38E-08 | 1.31E-07 |
| GO:0022404 | molting cycle process | 67/7539 | 1.42E-08 | 1.34E-07 |
| GO:0022405 | hair cycle process | 67/7539 | 1.42E-08 | 1.34E-07 |
| GO:0043535 | regulation of blood vessel endothelial cell migration | 57/7539 | 1.51E-08 | 1.43E-07 |
| GO:0030307 | positive regulation of cell growth | 102/7539 | 1.56E-08 | 1.47E-07 |
| GO:0042310 | vasoconstriction | 61/7539 | 1.70E-08 | 1.60E-07 |
| GO:0090263 | positive regulation of canonical Wnt signaling pathway | 61/7539 | 1.70E-08 | 1.60E-07 |
| GO:2000300 | regulation of synaptic vesicle exocytosis | 61/7539 | 1.70E-08 | 1.60E-07 |
| GO:0048872 | homeostasis of number of cells | 155/7539 | 1.74E-08 | 1.63E-07 |
| GO:0038083 | peptidyl-tyrosine autophosphorylation | 32/7539 | 1.81E-08 | 1.69E-07 |
| GO:0007632 | visual behavior | 47/7539 | 1.87E-08 | 1.74E-07 |
| GO:0021885 | forebrain cell migration | 46/7539 | 1.98E-08 | 1.84E-07 |
| GO:0043113 | receptor clustering | 46/7539 | 1.98E-08 | 1.84E-07 |
| GO:0021871 | forebrain regionalization | 23/7539 | 2.04E-08 | 1.89E-07 |
| GO:0048799 | animal organ maturation | 23/7539 | 2.04E-08 | 1.89E-07 |
| GO:0048168 | regulation of neuronal synaptic plasticity | 45/7539 | 2.08E-08 | 1.93E-07 |

| | | | | |
|---|---|---|---|---|
| GO:0046847 | filopodium assembly | 44/7539 | 2.18E-08 | 2.01E-07 |
| GO:0060425 | lung morphogenesis | 44/7539 | 2.18E-08 | 2.01E-07 |
| GO:0086003 | cardiac muscle cell contraction | 44/7539 | 2.18E-08 | 2.01E-07 |
| GO:0031646 | positive regulation of nervous system process | 53/7539 | 2.19E-08 | 2.01E-07 |
| GO:0099068 | postsynapse assembly | 27/7539 | 2.20E-08 | 2.02E-07 |
| GO:0045446 | endothelial cell differentiation | 63/7539 | 2.26E-08 | 2.07E-07 |
| GO:0048661 | positive regulation of smooth muscle cell proliferation | 63/7539 | 2.26E-08 | 2.07E-07 |
| GO:0001947 | heart looping | 42/7539 | 2.35E-08 | 2.15E-07 |
| GO:0099054 | presynapse assembly | 36/7539 | 2.38E-08 | 2.17E-07 |
| GO:0022408 | negative regulation of cell-cell adhesion | 99/7539 | 2.38E-08 | 2.17E-07 |
| GO:1901863 | positive regulation of muscle tissue development | 52/7539 | 2.39E-08 | 2.17E-07 |
| GO:0010812 | negative regulation of cell-substrate adhesion | 41/7539 | 2.41E-08 | 2.19E-07 |
| GO:0002042 | cell migration involved in sprouting angiogenesis | 38/7539 | 2.47E-08 | 2.24E-07 |
| GO:0051153 | regulation of striated muscle cell differentiation | 70/7539 | 2.49E-08 | 2.26E-07 |
| GO:0106027 | neuron projection organization | 62/7539 | 2.53E-08 | 2.29E-07 |
| GO:0034599 | cellular response to oxidative stress | 133/7539 | 2.63E-08 | 2.38E-07 |
| GO:0003158 | endothelium development | 69/7539 | 2.82E-08 | 2.54E-07 |
| GO:0045807 | positive regulation of endocytosis | 69/7539 | 2.82E-08 | 2.54E-07 |

| | | | | |
|---|---|---|---|---|
| GO:0009100 | glycoprotein metabolic process | 155/7539 | 2.87E-08 | 2.58E-07 |
| GO:0001938 | positive regulation of endothelial cell proliferation | 55/7539 | 3.17E-08 | 2.85E-07 |
| GO:1903524 | positive regulation of blood circulation | 55/7539 | 3.17E-08 | 2.85E-07 |
| GO:0043586 | tongue development | 21/7539 | 3.29E-08 | 2.95E-07 |
| GO:1901184 | regulation of ERBB signaling pathway | 47/7539 | 3.52E-08 | 3.15E-07 |
| GO:0050795 | regulation of behavior | 59/7539 | 3.53E-08 | 3.15E-07 |
| GO:0060419 | heart growth | 67/7539 | 3.61E-08 | 3.22E-07 |
| GO:0021549 | cerebellum development | 63/7539 | 3.66E-08 | 3.27E-07 |
| GO:0014046 | dopamine secretion | 30/7539 | 3.76E-08 | 3.33E-07 |
| GO:0014059 | regulation of dopamine secretion | 30/7539 | 3.76E-08 | 3.33E-07 |
| GO:0031076 | embryonic camera-type eye development | 30/7539 | 3.76E-08 | 3.33E-07 |
| GO:1901021 | positive regulation of calcium ion transmembrane transporter activity | 30/7539 | 3.76E-08 | 3.33E-07 |
| GO:1902692 | regulation of neuroblast proliferation | 30/7539 | 3.76E-08 | 3.33E-07 |
| GO:1904888 | cranial skeletal system development | 46/7539 | 3.76E-08 | 3.33E-07 |
| GO:0045669 | positive regulation of osteoblast differentiation | 45/7539 | 4.01E-08 | 3.54E-07 |
| GO:0098693 | regulation of synaptic vesicle cycle | 79/7539 | 4.14E-08 | 3.65E-07 |
| GO:0014910 | regulation of smooth muscle cell migration | 52/7539 | 4.22E-08 | 3.71E-07 |
| GO:0043949 | regulation of cAMP-mediated signaling | 44/7539 | 4.25E-08 | 3.73E-07 |

| | | | | |
|---|---|---|---|---|
| GO:2001236 | regulation of extrinsic apoptotic signaling pathway | 87/7539 | 4.36E-08 | 3.83E-07 |
| GO:0045823 | positive regulation of heart contraction | 32/7539 | 4.57E-08 | 4.00E-07 |
| GO:0045844 | positive regulation of striated muscle tissue development | 51/7539 | 4.62E-08 | 4.04E-07 |
| GO:0048636 | positive regulation of muscle organ development | 51/7539 | 4.62E-08 | 4.04E-07 |
| GO:0090278 | negative regulation of peptide hormone secretion | 42/7539 | 4.69E-08 | 4.10E-07 |
| GO:0097107 | postsynaptic density assembly | 19/7539 | 4.75E-08 | 4.15E-07 |
| GO:0061098 | positive regulation of protein tyrosine kinase activity | 33/7539 | 4.86E-08 | 4.24E-07 |
| GO:0021517 | ventral spinal cord development | 40/7539 | 5.06E-08 | 4.40E-07 |
| GO:0055024 | regulation of cardiac muscle tissue development | 60/7539 | 5.19E-08 | 4.50E-07 |
| GO:0099172 | presynapse organization | 39/7539 | 5.19E-08 | 4.50E-07 |
| GO:0021988 | olfactory lobe development | 35/7539 | 5.24E-08 | 4.53E-07 |
| GO:0061005 | cell differentiation involved in kidney development | 35/7539 | 5.24E-08 | 4.53E-07 |
| GO:0044060 | regulation of endocrine process | 38/7539 | 5.29E-08 | 4.57E-07 |
| GO:0045124 | regulation of bone resorption | 36/7539 | 5.32E-08 | 4.58E-07 |
| GO:0140353 | lipid export from cell | 36/7539 | 5.32E-08 | 4.58E-07 |
| GO:0045619 | regulation of lymphocyte differentiation | 98/7539 | 5.37E-08 | 4.62E-07 |
| GO:0048791 | calcium ion-regulated exocytosis of neurotransmitter | 22/7539 | 5.41E-08 | 4.65E-07 |
| GO:0048566 | embryonic digestive tract development | 26/7539 | 5.44E-08 | 4.65E-07 |

| | | | | |
|---|---|---|---|---|
| GO:0048841 | regulation of axon extension involved in axon guidance | 26/7539 | 5.44E-08 | 4.65E-07 |
| GO:1901186 | positive regulation of ERBB signaling pathway | 26/7539 | 5.44E-08 | 4.65E-07 |
| GO:0021514 | ventral spinal cord interneuron differentiation | 17/7539 | 5.66E-08 | 4.83E-07 |
| GO:0021783 | preganglionic parasympathetic fiber development | 17/7539 | 5.66E-08 | 4.83E-07 |
| GO:0043409 | negative regulation of MAPK cascade | 90/7539 | 5.90E-08 | 5.03E-07 |
| GO:0061097 | regulation of protein tyrosine kinase activity | 48/7539 | 5.98E-08 | 5.09E-07 |
| GO:0006469 | negative regulation of protein kinase activity | 104/7539 | 6.10E-08 | 5.18E-07 |
| GO:0007043 | cell-cell junction assembly | 79/7539 | 6.13E-08 | 5.21E-07 |
| GO:2001233 | regulation of apoptotic signaling pathway | 185/7539 | 6.31E-08 | 5.35E-07 |
| GO:0043279 | response to alkaloid | 47/7539 | 6.48E-08 | 5.48E-07 |
| GO:0035914 | skeletal muscle cell differentiation | 53/7539 | 6.63E-08 | 5.59E-07 |
| GO:1904063 | negative regulation of cation transmembrane transport | 53/7539 | 6.63E-08 | 5.59E-07 |
| GO:0035116 | embryonic hindlimb morphogenesis | 27/7539 | 6.64E-08 | 5.59E-07 |
| GO:0045987 | positive regulation of smooth muscle contraction | 27/7539 | 6.64E-08 | 5.59E-07 |
| GO:0014068 | positive regulation of phosphatidylinositol 3-kinase signaling | 46/7539 | 6.99E-08 | 5.88E-07 |
| GO:0007018 | microtubule-based movement | 174/7539 | 7.31E-08 | 6.13E-07 |
| GO:0007435 | salivary gland morphogenesis | 28/7539 | 7.77E-08 | 6.52E-07 |
| GO:1902668 | negative regulation of axon guidance | 23/7539 | 7.91E-08 | 6.62E-07 |

| | | | | |
|---|---|---|---|---|
| GO:1904861 | excitatory synapse assembly | 23/7539 | 7.91E-08 | 6.62E-07 |
| GO:0008542 | visual learning | 44/7539 | 8.05E-08 | 6.71E-07 |
| GO:0048678 | response to axon injury | 44/7539 | 8.05E-08 | 6.71E-07 |
| GO:0061371 | determination of heart left/right asymmetry | 44/7539 | 8.05E-08 | 6.71E-07 |
| GO:0014074 | response to purine-containing compound | 60/7539 | 8.41E-08 | 6.98E-07 |
| GO:0034766 | negative regulation of ion transmembrane transport | 60/7539 | 8.41E-08 | 6.98E-07 |
| GO:0048485 | sympathetic nervous system development | 20/7539 | 8.95E-08 | 7.43E-07 |
| GO:0097191 | extrinsic apoptotic signaling pathway | 114/7539 | 9.16E-08 | 7.59E-07 |
| GO:0007431 | salivary gland development | 30/7539 | 9.66E-08 | 7.99E-07 |
| GO:0048286 | lung alveolus development | 40/7539 | 1.01E-07 | 8.34E-07 |
| GO:0060986 | endocrine hormone secretion | 40/7539 | 1.01E-07 | 8.34E-07 |
| GO:1905952 | regulation of lipid localization | 86/7539 | 1.03E-07 | 8.49E-07 |
| GO:0051209 | release of sequestered calcium ion into cytosol | 62/7539 | 1.05E-07 | 8.65E-07 |
| GO:0051897 | positive regulation of protein kinase B signaling | 62/7539 | 1.05E-07 | 8.65E-07 |
| GO:0042551 | neuron maturation | 39/7539 | 1.05E-07 | 8.65E-07 |
| GO:0034763 | negative regulation of transmembrane transport | 72/7539 | 1.06E-07 | 8.67E-07 |
| GO:0021602 | cranial nerve morphogenesis | 24/7539 | 1.06E-07 | 8.69E-07 |
| GO:0019229 | regulation of vasoconstriction | 48/7539 | 1.07E-07 | 8.73E-07 |

| | | | | |
|---|---|---|---|---|
| GO:0035136 | forelimb morphogenesis | 32/7539 | 1.09E-07 | 8.88E-07 |
| GO:0046622 | positive regulation of organ growth | 38/7539 | 1.09E-07 | 8.88E-07 |
| GO:0051931 | regulation of sensory perception | 38/7539 | 1.09E-07 | 8.88E-07 |
| GO:2000648 | positive regulation of stem cell proliferation | 38/7539 | 1.09E-07 | 8.88E-07 |
| GO:0050922 | negative regulation of chemotaxis | 37/7539 | 1.12E-07 | 9.09E-07 |
| GO:0086001 | cardiac muscle cell action potential | 37/7539 | 1.12E-07 | 9.09E-07 |
| GO:0001941 | postsynaptic membrane organization | 33/7539 | 1.13E-07 | 9.11E-07 |
| GO:0030517 | negative regulation of axon extension | 33/7539 | 1.13E-07 | 9.11E-07 |
| GO:0035137 | hindlimb morphogenesis | 35/7539 | 1.15E-07 | 9.27E-07 |
| GO:0060412 | ventricular septum morphogenesis | 35/7539 | 1.15E-07 | 9.27E-07 |
| GO:0060428 | lung epithelium development | 35/7539 | 1.15E-07 | 9.27E-07 |
| GO:0106106 | cold-induced thermogenesis | 77/7539 | 1.18E-07 | 9.52E-07 |
| GO:0120161 | regulation of cold-induced thermogenesis | 77/7539 | 1.18E-07 | 9.52E-07 |
| GO:0050709 | negative regulation of protein secretion | 61/7539 | 1.19E-07 | 9.57E-07 |
| GO:1903708 | positive regulation of hemopoiesis | 110/7539 | 1.20E-07 | 9.62E-07 |
| GO:0001892 | embryonic placenta development | 64/7539 | 1.29E-07 | 1.03E-06 |
| GO:0040036 | regulation of fibroblast growth factor receptor signaling pathway | 25/7539 | 1.34E-07 | 1.07E-06 |
| GO:0007623 | circadian rhythm | 107/7539 | 1.36E-07 | 1.09E-06 |

| | | | | |
|---|---|---|---|---|
| GO:0007218 | neuropeptide signaling pathway | 51/7539 | 1.39E-07 | 1.11E-06 |
| GO:0033673 | negative regulation of kinase activity | 111/7539 | 1.40E-07 | 1.11E-06 |
| GO:0098739 | import across plasma membrane | 86/7539 | 1.46E-07 | 1.17E-06 |
| GO:0055008 | cardiac muscle tissue morphogenesis | 44/7539 | 1.49E-07 | 1.18E-06 |
| GO:0055117 | regulation of cardiac muscle contraction | 44/7539 | 1.49E-07 | 1.18E-06 |
| GO:0010522 | regulation of calcium ion transport into cytosol | 59/7539 | 1.52E-07 | 1.20E-06 |
| GO:0051899 | membrane depolarization | 50/7539 | 1.53E-07 | 1.21E-06 |
| GO:0000187 | activation of MAPK activity | 66/7539 | 1.54E-07 | 1.21E-06 |
| GO:0048013 | ephrin receptor signaling pathway | 26/7539 | 1.60E-07 | 1.26E-06 |
| GO:1905606 | regulation of presynapse assembly | 26/7539 | 1.60E-07 | 1.26E-06 |
| GO:0022029 | telencephalon cell migration | 43/7539 | 1.61E-07 | 1.27E-06 |
| GO:0009913 | epidermal cell differentiation | 97/7539 | 1.63E-07 | 1.29E-06 |
| GO:0051283 | negative regulation of sequestering of calcium ion | 62/7539 | 1.66E-07 | 1.30E-06 |
| GO:0048814 | regulation of dendrite morphogenesis | 65/7539 | 1.75E-07 | 1.37E-06 |
| GO:1904950 | negative regulation of establishment of protein localization | 87/7539 | 1.80E-07 | 1.41E-06 |
| GO:0043500 | muscle adaptation | 68/7539 | 1.80E-07 | 1.41E-06 |
| GO:0048008 | platelet-derived growth factor receptor signaling pathway | 41/7539 | 1.84E-07 | 1.44E-06 |
| GO:0062237 | protein localization to postsynapse | 41/7539 | 1.84E-07 | 1.44E-06 |

| | | | | |
|---|---|---|---|---|
| GO:0015718 | monocarboxylic acid transport | 89/7539 | 1.90E-07 | 1.48E-06 |
| GO:0043502 | regulation of muscle adaptation | 57/7539 | 1.93E-07 | 1.50E-06 |
| GO:0006865 | amino acid transport | 84/7539 | 1.94E-07 | 1.51E-06 |
| GO:0031102 | neuron projection regeneration | 40/7539 | 1.96E-07 | 1.52E-06 |
| GO:0090100 | positive regulation of transmembrane receptor protein serine/threonine kinase signaling pathway | 64/7539 | 1.99E-07 | 1.54E-06 |
| GO:0048843 | negative regulation of axon extension involved in axon guidance | 22/7539 | 2.02E-07 | 1.57E-06 |
| GO:0038084 | vascular endothelial growth factor signaling pathway | 28/7539 | 2.04E-07 | 1.58E-06 |
| GO:0072210 | metanephric nephron development | 28/7539 | 2.04E-07 | 1.58E-06 |
| GO:0099174 | regulation of presynapse organization | 28/7539 | 2.04E-07 | 1.58E-06 |
| GO:0048844 | artery morphogenesis | 47/7539 | 2.06E-07 | 1.59E-06 |
| GO:0035019 | somatic stem cell population maintenance | 38/7539 | 2.18E-07 | 1.68E-06 |
| GO:0071542 | dopaminergic neuron differentiation | 29/7539 | 2.21E-07 | 1.70E-06 |
| GO:0090175 | regulation of establishment of planar polarity | 29/7539 | 2.21E-07 | 1.70E-06 |
| GO:0003014 | renal system process | 63/7539 | 2.26E-07 | 1.74E-06 |
| GO:0001954 | positive regulation of cell-matrix adhesion | 37/7539 | 2.27E-07 | 1.74E-06 |
| GO:0021587 | cerebellum morphogenesis | 37/7539 | 2.27E-07 | 1.74E-06 |
| GO:0048538 | thymus development | 37/7539 | 2.27E-07 | 1.74E-06 |
| GO:0051930 | regulation of sensory perception of pain | 37/7539 | 2.27E-07 | 1.74E-06 |

| | | | | |
|---|---|---|---|---|
| GO:0014014 | negative regulation of gliogenesis | 36/7539 | 2.36E-07 | 1.80E-06 |
| GO:0048854 | brain morphogenesis | 31/7539 | 2.42E-07 | 1.84E-06 |
| GO:0021516 | dorsal spinal cord development | 19/7539 | 2.42E-07 | 1.84E-06 |
| GO:0048536 | spleen development | 32/7539 | 2.47E-07 | 1.88E-06 |
| GO:0048010 | vascular endothelial growth factor receptor signaling pathway | 33/7539 | 2.48E-07 | 1.88E-06 |
| GO:0006942 | regulation of striated muscle contraction | 50/7539 | 2.60E-07 | 1.97E-06 |
| GO:0042733 | embryonic digit morphogenesis | 44/7539 | 2.69E-07 | 2.04E-06 |
| GO:0000302 | response to reactive oxygen species | 100/7539 | 2.70E-07 | 2.05E-06 |
| GO:0043624 | cellular protein complex disassembly | 76/7539 | 2.88E-07 | 2.17E-06 |
| GO:0014896 | muscle hypertrophy | 61/7539 | 2.92E-07 | 2.21E-06 |
| GO:0033627 | cell adhesion mediated by integrin | 43/7539 | 2.93E-07 | 2.21E-06 |
| GO:0050921 | positive regulation of chemotaxis | 78/7539 | 3.14E-07 | 2.36E-06 |
| GO:0060688 | regulation of morphogenesis of a branching structure | 42/7539 | 3.18E-07 | 2.39E-06 |
| GO:0002092 | positive regulation of receptor internalization | 24/7539 | 3.25E-07 | 2.44E-06 |
| GO:0045742 | positive regulation of epidermal growth factor receptor signaling pathway | 24/7539 | 3.25E-07 | 2.44E-06 |
| GO:0014897 | striated muscle hypertrophy | 60/7539 | 3.32E-07 | 2.49E-06 |
| GO:0046631 | alpha-beta T cell activation | 85/7539 | 3.33E-07 | 2.50E-06 |
| GO:0032233 | positive regulation of actin filament bundle assembly | 41/7539 | 3.43E-07 | 2.57E-06 |

| | | | | |
|---|---|---|---|---|
| GO:0016311 | dephosphorylation | 167/7539 | 3.47E-07 | 2.59E-06 |
| GO:0060038 | cardiac muscle cell proliferation | 40/7539 | 3.69E-07 | 2.75E-06 |
| GO:0042474 | middle ear morphogenesis | 20/7539 | 3.73E-07 | 2.77E-06 |
| GO:0062009 | secondary palate development | 20/7539 | 3.73E-07 | 2.77E-06 |
| GO:0040037 | negative regulation of fibroblast growth factor receptor signaling pathway | 17/7539 | 3.74E-07 | 2.77E-06 |
| GO:0098962 | regulation of postsynaptic neurotransmitter receptor activity | 17/7539 | 3.74E-07 | 2.77E-06 |
| GO:1905874 | regulation of postsynaptic density organization | 17/7539 | 3.74E-07 | 2.77E-06 |
| GO:0021884 | forebrain neuron development | 25/7539 | 3.81E-07 | 2.82E-06 |
| GO:0046887 | positive regulation of hormone secretion | 84/7539 | 3.84E-07 | 2.84E-06 |
| GO:0051282 | regulation of sequestering of calcium ion | 62/7539 | 3.95E-07 | 2.92E-06 |
| GO:0045776 | negative regulation of blood pressure | 39/7539 | 3.95E-07 | 2.92E-06 |
| GO:1903539 | protein localization to postsynaptic membrane | 38/7539 | 4.21E-07 | 3.11E-06 |
| GO:0060292 | long-term synaptic depression | 26/7539 | 4.29E-07 | 3.16E-06 |
| GO:0006029 | proteoglycan metabolic process | 45/7539 | 4.33E-07 | 3.19E-06 |
| GO:0010639 | negative regulation of organelle organization | 184/7539 | 4.46E-07 | 3.27E-06 |
| GO:0140115 | export across plasma membrane | 37/7539 | 4.46E-07 | 3.27E-06 |
| GO:0002053 | positive regulation of mesenchymal cell proliferation | 27/7539 | 4.69E-07 | 3.43E-06 |
| GO:0060071 | Wnt signaling pathway, planar cell polarity pathway | 27/7539 | 4.69E-07 | 3.43E-06 |

| | | | | |
|---|---|---|---|---|
| GO:0021978 | telencephalon regionalization | 15/7539 | 4.84E-07 | 3.53E-06 |
| GO:0060579 | ventral spinal cord interneuron fate commitment | 15/7539 | 4.84E-07 | 3.53E-06 |
| GO:0060581 | cell fate commitment involved in pattern specification | 15/7539 | 4.84E-07 | 3.53E-06 |
| GO:0061548 | ganglion development | 15/7539 | 4.84E-07 | 3.53E-06 |
| GO:0055006 | cardiac cell development | 57/7539 | 4.85E-07 | 3.53E-06 |
| GO:0030514 | negative regulation of BMP signaling pathway | 35/7539 | 4.91E-07 | 3.57E-06 |
| GO:0048701 | embryonic cranial skeleton morphogenesis | 35/7539 | 4.91E-07 | 3.57E-06 |
| GO:2000050 | regulation of non-canonical Wnt signaling pathway | 21/7539 | 5.14E-07 | 3.72E-06 |
| GO:2000311 | regulation of AMPA receptor activity | 21/7539 | 5.14E-07 | 3.72E-06 |
| GO:0086009 | membrane repolarization | 29/7539 | 5.20E-07 | 3.77E-06 |
| GO:0021772 | olfactory bulb development | 32/7539 | 5.33E-07 | 3.85E-06 |
| GO:0021696 | cerebellar cortex morphogenesis | 31/7539 | 5.36E-07 | 3.86E-06 |
| GO:0050918 | positive chemotaxis | 31/7539 | 5.36E-07 | 3.86E-06 |
| GO:0071526 | semaphorin-plexin signaling pathway | 31/7539 | 5.36E-07 | 3.86E-06 |
| GO:0090184 | positive regulation of kidney development | 31/7539 | 5.36E-07 | 3.86E-06 |
| GO:1902743 | regulation of lamellipodium organization | 31/7539 | 5.36E-07 | 3.86E-06 |
| GO:0061041 | regulation of wound healing | 76/7539 | 5.92E-07 | 4.25E-06 |
| GO:0051208 | sequestering of calcium ion | 62/7539 | 6.00E-07 | 4.31E-06 |

| | | | | |
|---|---|---|---|---|
| GO:0002043 | blood vessel endothelial cell proliferation involved in sprouting angiogenesis | 18/7539 | 6.50E-07 | 4.66E-06 |
| GO:0048596 | embryonic camera-type eye morphogenesis | 22/7539 | 6.54E-07 | 4.67E-06 |
| GO:0060039 | pericardium development | 22/7539 | 6.54E-07 | 4.67E-06 |
| GO:1903671 | negative regulation of sprouting angiogenesis | 22/7539 | 6.54E-07 | 4.67E-06 |
| GO:1903829 | positive regulation of cellular protein localization | 144/7539 | 6.54E-07 | 4.67E-06 |
| GO:0032410 | negative regulation of transporter activity | 46/7539 | 6.70E-07 | 4.76E-06 |
| GO:0045453 | bone resorption | 46/7539 | 6.70E-07 | 4.76E-06 |
| GO:0071277 | cellular response to calcium ion | 46/7539 | 6.70E-07 | 4.76E-06 |
| GO:0071248 | cellular response to metal ion | 80/7539 | 6.73E-07 | 4.78E-06 |
| GO:0008347 | glial cell migration | 40/7539 | 6.77E-07 | 4.80E-06 |
| GO:0035904 | aorta development | 40/7539 | 6.77E-07 | 4.80E-06 |
| GO:0007229 | integrin-mediated signaling pathway | 54/7539 | 7.05E-07 | 4.99E-06 |
| GO:0010171 | body morphogenesis | 39/7539 | 7.33E-07 | 5.18E-06 |
| GO:0014743 | regulation of muscle hypertrophy | 45/7539 | 7.44E-07 | 5.24E-06 |
| GO:0034109 | homotypic cell-cell adhesion | 45/7539 | 7.44E-07 | 5.24E-06 |
| GO:0048747 | muscle fiber development | 45/7539 | 7.44E-07 | 5.24E-06 |
| GO:0051057 | positive regulation of small GTPase mediated signal transduction | 45/7539 | 7.44E-07 | 5.24E-06 |
| GO:0032368 | regulation of lipid transport | 69/7539 | 7.56E-07 | 5.31E-06 |

| | | | | |
|---|---|---|---|---|
| GO:1900006 | positive regulation of dendrite development | 60/7539 | 7.82E-07 | 5.49E-06 |
| GO:1905477 | positive regulation of protein localization to membrane | 60/7539 | 7.82E-07 | 5.49E-06 |
| GO:0051155 | positive regulation of striated muscle cell differentiation | 44/7539 | 8.24E-07 | 5.77E-06 |
| GO:0051224 | negative regulation of protein transport | 83/7539 | 8.54E-07 | 5.98E-06 |
| GO:0051590 | positive regulation of neurotransmitter transport | 24/7539 | 9.01E-07 | 6.29E-06 |
| GO:0050806 | positive regulation of synaptic transmission | 158/7539 | 9.01E-07 | 6.29E-06 |
| GO:0032890 | regulation of organic acid transport | 52/7539 | 9.02E-07 | 6.29E-06 |
| GO:0051954 | positive regulation of amine transport | 35/7539 | 9.62E-07 | 6.70E-06 |
| GO:0070977 | bone maturation | 19/7539 | 9.67E-07 | 6.73E-06 |
| GO:0045637 | regulation of myeloid cell differentiation | 107/7539 | 9.83E-07 | 6.83E-06 |
| GO:0030595 | leukocyte chemotaxis | 105/7539 | 9.91E-07 | 6.88E-06 |
| GO:0001964 | startle response | 25/7539 | 9.96E-07 | 6.88E-06 |
| GO:0003203 | endocardial cushion morphogenesis | 25/7539 | 9.96E-07 | 6.88E-06 |
| GO:0003209 | cardiac atrium morphogenesis | 25/7539 | 9.96E-07 | 6.88E-06 |
| GO:0048048 | embryonic eye morphogenesis | 25/7539 | 9.96E-07 | 6.88E-06 |
| GO:0042058 | regulation of epidermal growth factor receptor signaling pathway | 42/7539 | 1.00E-06 | 6.94E-06 |
| GO:0051149 | positive regulation of muscle cell differentiation | 51/7539 | 1.02E-06 | 7.03E-06 |
| GO:0070571 | negative regulation of neuron projection regeneration | 16/7539 | 1.04E-06 | 7.15E-06 |

| | | | | |
|---|---|---|---|---|
| GO:0099150 | regulation of postsynaptic specialization assembly | 16/7539 | 1.04E-06 | 7.15E-06 |
| GO:0150052 | regulation of postsynapse assembly | 16/7539 | 1.04E-06 | 7.15E-06 |
| GO:0032232 | negative regulation of actin filament bundle assembly | 26/7539 | 1.07E-06 | 7.33E-06 |
| GO:0048169 | regulation of long-term neuronal synaptic plasticity | 26/7539 | 1.07E-06 | 7.33E-06 |
| GO:0051348 | negative regulation of transferase activity | 121/7539 | 1.10E-06 | 7.57E-06 |
| GO:0051953 | negative regulation of amine transport | 27/7539 | 1.12E-06 | 7.67E-06 |
| GO:0014855 | striated muscle cell proliferation | 46/7539 | 1.12E-06 | 7.68E-06 |
| GO:1902667 | regulation of axon guidance | 31/7539 | 1.13E-06 | 7.75E-06 |
| GO:0043112 | receptor metabolic process | 102/7539 | 1.15E-06 | 7.83E-06 |
| GO:0009101 | glycoprotein biosynthetic process | 126/7539 | 1.15E-06 | 7.86E-06 |
| GO:0045747 | positive regulation of Notch signaling pathway | 30/7539 | 1.15E-06 | 7.86E-06 |
| GO:0003300 | cardiac muscle hypertrophy | 57/7539 | 1.16E-06 | 7.89E-06 |
| GO:0010092 | specification of animal organ identity | 29/7539 | 1.16E-06 | 7.89E-06 |
| GO:1901879 | regulation of protein depolymerization | 53/7539 | 1.26E-06 | 8.52E-06 |
| GO:1904035 | regulation of epithelial cell apoptotic process | 53/7539 | 1.26E-06 | 8.52E-06 |
| GO:0032886 | regulation of microtubule-based process | 120/7539 | 1.27E-06 | 8.57E-06 |
| GO:0021521 | ventral spinal cord interneuron specification | Dec-39 | 1.29E-06 | 8.70E-06 |
| GO:0060573 | cell fate specification involved in pattern specification | Dec-39 | 1.29E-06 | 8.70E-06 |

| | | | | |
|---|---|---|---|---|
| GO:0007413 | axonal fasciculation | 20/7539 | 1.29E-06 | 8.70E-06 |
| GO:0021513 | spinal cord dorsal/ventral patterning | 20/7539 | 1.29E-06 | 8.70E-06 |
| GO:0051894 | positive regulation of focal adhesion assembly | 20/7539 | 1.29E-06 | 8.70E-06 |
| GO:0106030 | neuron projection fasciculation | 20/7539 | 1.29E-06 | 8.70E-06 |
| GO:0008154 | actin polymerization or depolymerization | 103/7539 | 1.31E-06 | 8.81E-06 |
| GO:0032092 | positive regulation of protein binding | 56/7539 | 1.32E-06 | 8.87E-06 |
| GO:2001237 | negative regulation of extrinsic apoptotic signaling pathway | 56/7539 | 1.32E-06 | 8.87E-06 |
| GO:0072074 | kidney mesenchyme development | 14/7539 | 1.41E-06 | 9.43E-06 |
| GO:0098917 | retrograde trans-synaptic signaling | 14/7539 | 1.41E-06 | 9.43E-06 |
| GO:0050954 | sensory perception of mechanical stimulus | 138/7539 | 1.45E-06 | 9.68E-06 |
| GO:0002687 | positive regulation of leukocyte migration | 79/7539 | 1.50E-06 | 1.00E-05 |
| GO:0002052 | positive regulation of neuroblast proliferation | 21/7539 | 1.60E-06 | 1.07E-05 |
| GO:0015874 | norepinephrine transport | 21/7539 | 1.60E-06 | 1.07E-05 |
| GO:0021511 | spinal cord patterning | 21/7539 | 1.60E-06 | 1.07E-05 |
| GO:1901018 | positive regulation of potassium ion transmembrane transporter activity | 21/7539 | 1.60E-06 | 1.07E-05 |
| GO:0031503 | protein-containing complex localization | 120/7539 | 1.61E-06 | 1.07E-05 |
| GO:0046545 | development of primary female sexual characteristics | 63/7539 | 1.73E-06 | 1.15E-05 |
| GO:0045986 | negative regulation of smooth muscle contraction | 17/7539 | 1.73E-06 | 1.15E-05 |

| | | | | |
|---|---|---|---|---|
| GO:0045580 | regulation of T cell differentiation | 80/7539 | 1.80E-06 | 1.19E-05 |
| GO:0051258 | protein polymerization | 133/7539 | 1.80E-06 | 1.19E-05 |
| GO:0051279 | regulation of release of sequestered calcium ion into cytosol | 46/7539 | 1.85E-06 | 1.22E-05 |
| GO:0051966 | regulation of synaptic transmission, glutamatergic | 46/7539 | 1.85E-06 | 1.22E-05 |
| GO:0060571 | morphogenesis of an epithelial fold | 22/7539 | 1.88E-06 | 1.24E-05 |
| GO:0150117 | positive regulation of cell-substrate junction organization | 22/7539 | 1.88E-06 | 1.24E-05 |
| GO:0017156 | calcium-ion regulated exocytosis | 41/7539 | 1.92E-06 | 1.26E-05 |
| GO:0050727 | regulation of inflammatory response | 163/7539 | 1.94E-06 | 1.28E-05 |
| GO:0031103 | axon regeneration | 34/7539 | 1.95E-06 | 1.28E-05 |
| GO:0061383 | trabecula morphogenesis | 34/7539 | 1.95E-06 | 1.28E-05 |
| GO:0001704 | formation of primary germ layer | 56/7539 | 2.01E-06 | 1.32E-05 |
| GO:0010657 | muscle cell apoptotic process | 56/7539 | 2.01E-06 | 1.32E-05 |
| GO:0030705 | cytoskeleton-dependent intracellular transport | 94/7539 | 2.02E-06 | 1.32E-05 |
| GO:0042493 | response to drug | 120/7539 | 2.05E-06 | 1.34E-05 |
| GO:0048708 | astrocyte differentiation | 49/7539 | 2.07E-06 | 1.35E-05 |
| GO:0060043 | regulation of cardiac muscle cell proliferation | 33/7539 | 2.08E-06 | 1.35E-05 |
| GO:1904375 | regulation of protein localization to cell periphery | 72/7539 | 2.08E-06 | 1.35E-05 |
| GO:0002063 | chondrocyte development | 23/7539 | 2.11E-06 | 1.37E-05 |

| | | | | |
|---|---|---|---|---|
| GO:0021533 | cell differentiation in hindbrain | 23/7539 | 2.11E-06 | 1.37E-05 |
| GO:0060074 | synapse maturation | 23/7539 | 2.11E-06 | 1.37E-05 |
| GO:0061037 | negative regulation of cartilage development | 23/7539 | 2.11E-06 | 1.37E-05 |
| GO:1901889 | negative regulation of cell junction assembly | 23/7539 | 2.11E-06 | 1.37E-05 |
| GO:0032271 | regulation of protein polymerization | 107/7539 | 2.16E-06 | 1.40E-05 |
| GO:0098659 | inorganic cation import across plasma membrane | 55/7539 | 2.30E-06 | 1.48E-05 |
| GO:0099587 | inorganic ion import across plasma membrane | 55/7539 | 2.30E-06 | 1.48E-05 |
| GO:0097755 | positive regulation of blood vessel diameter | 44/7539 | 2.33E-06 | 1.50E-05 |
| GO:0006470 | protein dephosphorylation | 110/7539 | 2.37E-06 | 1.53E-05 |
| GO:0003156 | regulation of animal organ formation | 30/7539 | 2.39E-06 | 1.54E-05 |
| GO:2000826 | regulation of heart morphogenesis | 30/7539 | 2.39E-06 | 1.54E-05 |
| GO:0045686 | negative regulation of glial cell differentiation | 25/7539 | 2.41E-06 | 1.55E-05 |
| GO:0045746 | negative regulation of Notch signaling pathway | 25/7539 | 2.41E-06 | 1.55E-05 |
| GO:0071320 | cellular response to cAMP | 25/7539 | 2.41E-06 | 1.55E-05 |
| GO:0046486 | glycerolipid metabolic process | 158/7539 | 2.47E-06 | 1.58E-05 |
| GO:0003230 | cardiac atrium development | 27/7539 | 2.52E-06 | 1.61E-05 |
| GO:0035850 | epithelial cell differentiation involved in kidney development | 27/7539 | 2.52E-06 | 1.61E-05 |
| GO:0014911 | positive regulation of smooth muscle cell migration | 38/7539 | 2.57E-06 | 1.64E-05 |

| | | | | |
|---|---|---|---|---|
| GO:0060350 | endochondral bone morphogenesis | 38/7539 | 2.57E-06 | 1.64E-05 |
| GO:1903078 | positive regulation of protein localization to plasma membrane | 38/7539 | 2.57E-06 | 1.64E-05 |
| GO:0002761 | regulation of myeloid leukocyte differentiation | 68/7539 | 2.58E-06 | 1.64E-05 |
| GO:0060997 | dendritic spine morphogenesis | 43/7539 | 2.60E-06 | 1.66E-05 |
| GO:1905954 | positive regulation of lipid localization | 57/7539 | 2.65E-06 | 1.68E-05 |
| GO:0050680 | negative regulation of epithelial cell proliferation | 75/7539 | 2.65E-06 | 1.68E-05 |
| GO:2000241 | regulation of reproductive process | 94/7539 | 2.67E-06 | 1.69E-05 |
| GO:0070482 | response to oxygen levels | 130/7539 | 2.68E-06 | 1.70E-05 |
| GO:0045787 | positive regulation of cell cycle | 157/7539 | 2.80E-06 | 1.77E-05 |
| GO:0048260 | positive regulation of receptor-mediated endocytosis | 37/7539 | 2.82E-06 | 1.78E-05 |
| GO:0060324 | face development | 37/7539 | 2.82E-06 | 1.78E-05 |
| GO:0090183 | regulation of kidney development | 37/7539 | 2.82E-06 | 1.78E-05 |
| GO:0003184 | pulmonary valve morphogenesis | 15/7539 | 2.87E-06 | 1.81E-05 |
| GO:0048853 | forebrain morphogenesis | 15/7539 | 2.87E-06 | 1.81E-05 |
| GO:1904889 | regulation of excitatory synapse assembly | 15/7539 | 2.87E-06 | 1.81E-05 |
| GO:1905331 | negative regulation of morphogenesis of an epithelium | 15/7539 | 2.87E-06 | 1.81E-05 |
| GO:0071901 | negative regulation of protein serine/threonine kinase activity | 62/7539 | 2.89E-06 | 1.82E-05 |
| GO:0017015 | regulation of transforming growth factor beta receptor signaling pathway | 59/7539 | 2.99E-06 | 1.88E-05 |

| | | | | |
|---|---|---|---|---|
| GO:0031016 | pancreas development | 53/7539 | 2.99E-06 | 1.88E-05 |
| GO:0003272 | endocardial cushion formation | 19/7539 | 3.22E-06 | 2.02E-05 |
| GO:0031954 | positive regulation of protein autophosphorylation | 19/7539 | 3.22E-06 | 2.02E-05 |
| GO:0009880 | embryonic pattern specification | 41/7539 | 3.25E-06 | 2.03E-05 |
| GO:0045600 | positive regulation of fat cell differentiation | 41/7539 | 3.25E-06 | 2.03E-05 |
| GO:1904377 | positive regulation of protein localization to cell periphery | 41/7539 | 3.25E-06 | 2.03E-05 |
| GO:1903039 | positive regulation of leukocyte cell-cell adhesion | 104/7539 | 3.26E-06 | 2.03E-05 |
| GO:0008585 | female gonad development | 61/7539 | 3.32E-06 | 2.07E-05 |
| GO:0017145 | stem cell division | 35/7539 | 3.35E-06 | 2.08E-05 |
| GO:0098815 | modulation of excitatory postsynaptic potential | 35/7539 | 3.35E-06 | 2.08E-05 |
| GO:0018210 | peptidyl-threonine modification | 66/7539 | 3.41E-06 | 2.12E-05 |
| GO:0051101 | regulation of DNA binding | 66/7539 | 3.41E-06 | 2.12E-05 |
| GO:2000116 | regulation of cysteine-type endopeptidase activity | 109/7539 | 3.48E-06 | 2.15E-05 |
| GO:0055025 | positive regulation of cardiac muscle tissue development | 34/7539 | 3.63E-06 | 2.25E-05 |
| GO:0010256 | endomembrane system organization | 176/7539 | 3.67E-06 | 2.27E-05 |
| GO:0051091 | positive regulation of DNA-binding transcription factor activity | 119/7539 | 3.74E-06 | 2.31E-05 |
| GO:0042593 | glucose homeostasis | 124/7539 | 3.80E-06 | 2.34E-05 |
| GO:0043281 | regulation of cysteine-type endopeptidase activity involved in apoptotic process | 99/7539 | 3.89E-06 | 2.40E-05 |

| | | | | |
|---|---|---|---|---|
| GO:0035710 | CD4-positive, alpha-beta T cell activation | 57/7539 | 3.93E-06 | 2.42E-05 |
| GO:0051610 | serotonin uptake | Nov-39 | 4.00E-06 | 2.46E-05 |
| GO:0072075 | metanephric mesenchyme development | Nov-39 | 4.00E-06 | 2.46E-05 |
| GO:0014831 | gastro-intestinal system smooth muscle contraction | 13/7539 | 4.08E-06 | 2.50E-05 |
| GO:0021527 | spinal cord association neuron differentiation | 13/7539 | 4.08E-06 | 2.50E-05 |
| GO:0099151 | regulation of postsynaptic density assembly | 13/7539 | 4.08E-06 | 2.50E-05 |
| GO:2000052 | positive regulation of non-canonical Wnt signaling pathway | 13/7539 | 4.08E-06 | 2.50E-05 |
| GO:0046632 | alpha-beta T cell differentiation | 62/7539 | 4.18E-06 | 2.56E-05 |
| GO:0070509 | calcium ion import | 50/7539 | 4.44E-06 | 2.70E-05 |
| GO:1903035 | negative regulation of response to wounding | 50/7539 | 4.44E-06 | 2.70E-05 |
| GO:0002089 | lens morphogenesis in camera-type eye | 21/7539 | 4.45E-06 | 2.70E-05 |
| GO:0030878 | thyroid gland development | 21/7539 | 4.45E-06 | 2.70E-05 |
| GO:0060384 | innervation | 21/7539 | 4.45E-06 | 2.70E-05 |
| GO:0060445 | branching involved in salivary gland morphogenesis | 21/7539 | 4.45E-06 | 2.70E-05 |
| GO:2000310 | regulation of NMDA receptor activity | 21/7539 | 4.45E-06 | 2.70E-05 |
| GO:0060999 | positive regulation of dendritic spine development | 38/7539 | 4.46E-06 | 2.71E-05 |
| GO:0050886 | endocrine process | 56/7539 | 4.50E-06 | 2.73E-05 |
| GO:0120162 | positive regulation of cold-induced thermogenesis | 53/7539 | 4.53E-06 | 2.75E-05 |

| | | | | |
|---|---|---|---|---|
| GO:0062012 | regulation of small molecule metabolic process | 156/7539 | 4.67E-06 | 2.83E-05 |
| GO:0033500 | carbohydrate homeostasis | 124/7539 | 4.75E-06 | 2.87E-05 |
| GO:0017158 | regulation of calcium ion-dependent exocytosis | 30/7539 | 4.75E-06 | 2.87E-05 |
| GO:0019228 | neuronal action potential | 30/7539 | 4.75E-06 | 2.87E-05 |
| GO:0010611 | regulation of cardiac muscle hypertrophy | 42/7539 | 4.81E-06 | 2.90E-05 |
| GO:0045123 | cellular extravasation | 42/7539 | 4.81E-06 | 2.90E-05 |
| GO:0001945 | lymph vessel development | 22/7539 | 4.89E-06 | 2.94E-05 |
| GO:0048714 | positive regulation of oligodendrocyte differentiation | 22/7539 | 4.89E-06 | 2.94E-05 |
| GO:0014904 | myotube cell development | 29/7539 | 4.99E-06 | 2.99E-05 |
| GO:0043114 | regulation of vascular permeability | 29/7539 | 4.99E-06 | 2.99E-05 |
| GO:0048144 | fibroblast proliferation | 58/7539 | 5.03E-06 | 3.01E-05 |
| GO:0050866 | negative regulation of cell activation | 99/7539 | 5.03E-06 | 3.02E-05 |
| GO:0055013 | cardiac muscle cell development | 52/7539 | 5.18E-06 | 3.10E-05 |
| GO:0010765 | positive regulation of sodium ion transport | 28/7539 | 5.19E-06 | 3.10E-05 |
| GO:0060977 | coronary vasculature morphogenesis | 23/7539 | 5.19E-06 | 3.10E-05 |
| GO:0070570 | regulation of neuron projection regeneration | 27/7539 | 5.35E-06 | 3.19E-05 |
| GO:0099633 | protein localization to postsynaptic specialization membrane | 24/7539 | 5.38E-06 | 3.20E-05 |
| GO:0099645 | neurotransmitter receptor localization to postsynaptic specialization membrane | 24/7539 | 5.38E-06 | 3.20E-05 |

| | | | | |
|---|---|---|---|---|
| GO:1902107 | positive regulation of leukocyte differentiation | 87/7539 | 5.39E-06 | 3.20E-05 |
| GO:0055021 | regulation of cardiac muscle tissue growth | 45/7539 | 5.39E-06 | 3.20E-05 |
| GO:0002011 | morphogenesis of an epithelial sheet | 36/7539 | 5.44E-06 | 3.23E-05 |
| GO:0110111 | negative regulation of animal organ morphogenesis | 26/7539 | 5.45E-06 | 3.23E-05 |
| GO:1905209 | positive regulation of cardiocyte differentiation | 25/7539 | 5.46E-06 | 3.24E-05 |
| GO:1903076 | regulation of protein localization to plasma membrane | 60/7539 | 5.53E-06 | 3.27E-05 |
| GO:0007586 | digestion | 65/7539 | 5.57E-06 | 3.29E-05 |
| GO:0048145 | regulation of fibroblast proliferation | 57/7539 | 5.77E-06 | 3.41E-05 |
| GO:0043506 | regulation of JUN kinase activity | 51/7539 | 5.92E-06 | 3.50E-05 |
| GO:0002067 | glandular epithelial cell differentiation | 35/7539 | 5.99E-06 | 3.53E-05 |
| GO:0055078 | sodium ion homeostasis | 35/7539 | 5.99E-06 | 3.53E-05 |
| GO:0050878 | regulation of body fluid levels | 154/7539 | 5.99E-06 | 3.53E-05 |
| GO:0055067 | monovalent inorganic cation homeostasis | 78/7539 | 6.03E-06 | 3.55E-05 |
| GO:0015800 | acidic amino acid transport | 40/7539 | 6.05E-06 | 3.56E-05 |
| GO:0001756 | somitogenesis | 44/7539 | 6.10E-06 | 3.58E-05 |
| GO:0043507 | positive regulation of JUN kinase activity | 44/7539 | 6.10E-06 | 3.58E-05 |
| GO:0043931 | ossification involved in bone maturation | 17/7539 | 6.34E-06 | 3.72E-05 |
| GO:1903844 | regulation of cellular response to transforming growth factor beta stimulus | 59/7539 | 6.35E-06 | 3.72E-05 |

| | | | | |
|---|---|---|---|---|
| GO:0002088 | lens development in camera-type eye | 50/7539 | 6.77E-06 | 3.96E-05 |
| GO:0007224 | smoothened signaling pathway | 77/7539 | 6.95E-06 | 4.06E-05 |
| GO:0060271 | cilium assembly | 137/7539 | 7.23E-06 | 4.22E-05 |
| GO:0022600 | digestive system process | 58/7539 | 7.30E-06 | 4.26E-05 |
| GO:0007622 | rhythmic behavior | 38/7539 | 7.56E-06 | 4.40E-05 |
| GO:0030858 | positive regulation of epithelial cell differentiation | 38/7539 | 7.56E-06 | 4.40E-05 |
| GO:2000243 | positive regulation of reproductive process | 52/7539 | 7.77E-06 | 4.52E-05 |
| GO:0035088 | establishment or maintenance of apical/basal cell polarity | 32/7539 | 7.79E-06 | 4.52E-05 |
| GO:0051489 | regulation of filopodium assembly | 32/7539 | 7.79E-06 | 4.52E-05 |
| GO:0061245 | establishment or maintenance of bipolar cell polarity | 32/7539 | 7.79E-06 | 4.52E-05 |
| GO:0048681 | negative regulation of axon regeneration | 14/7539 | 7.88E-06 | 4.56E-05 |
| GO:0055057 | neuroblast division | 14/7539 | 7.88E-06 | 4.56E-05 |
| GO:0060979 | vasculogenesis involved in coronary vascular morphogenesis | 14/7539 | 7.88E-06 | 4.56E-05 |
| GO:0010996 | response to auditory stimulus | 18/7539 | 7.96E-06 | 4.59E-05 |
| GO:0072202 | cell differentiation involved in metanephros development | 18/7539 | 7.96E-06 | 4.59E-05 |
| GO:1901626 | regulation of postsynaptic membrane organization | 18/7539 | 7.96E-06 | 4.59E-05 |
| GO:0003229 | ventricular cardiac muscle tissue development | 37/7539 | 8.43E-06 | 4.85E-05 |
| GO:0048512 | circadian behavior | 37/7539 | 8.43E-06 | 4.85E-05 |

| | | | | |
|---|---|---|---|---|
| GO:1905207 | regulation of cardiocyte differentiation | 37/7539 | 8.43E-06 | 4.85E-05 |
| GO:0008088 | axo-dendritic transport | 45/7539 | 8.48E-06 | 4.87E-05 |
| GO:0001885 | endothelial cell development | 41/7539 | 8.80E-06 | 5.05E-05 |
| GO:0007492 | endoderm development | 41/7539 | 8.80E-06 | 5.05E-05 |
| GO:0060420 | regulation of heart growth | 48/7539 | 8.83E-06 | 5.06E-05 |
| GO:0010718 | positive regulation of epithelial to mesenchymal transition | 30/7539 | 9.07E-06 | 5.19E-05 |
| GO:0070663 | regulation of leukocyte proliferation | 114/7539 | 9.17E-06 | 5.25E-05 |
| GO:0036342 | post-anal tail morphogenesis | 19/7539 | 9.34E-06 | 5.33E-05 |
| GO:0071880 | adenylate cyclase-activating adrenergic receptor signaling pathway | 19/7539 | 9.34E-06 | 5.33E-05 |
| GO:1904738 | vascular associated smooth muscle cell migration | 19/7539 | 9.34E-06 | 5.33E-05 |
| GO:1904752 | regulation of vascular associated smooth muscle cell migration | 19/7539 | 9.34E-06 | 5.33E-05 |
| GO:0098900 | regulation of action potential | 36/7539 | 9.38E-06 | 5.34E-05 |
| GO:0031128 | developmental induction | 29/7539 | 9.70E-06 | 5.52E-05 |
| GO:0060421 | positive regulation of heart growth | 29/7539 | 9.70E-06 | 5.52E-05 |
| GO:0048146 | positive regulation of fibroblast proliferation | 40/7539 | 9.92E-06 | 5.62E-05 |
| GO:0071867 | response to monoamine | 40/7539 | 9.92E-06 | 5.62E-05 |
| GO:0071869 | response to catecholamine | 40/7539 | 9.92E-06 | 5.62E-05 |
| GO:0051259 | protein complex oligomerization | 103/7539 | 1.00E-05 | 5.69E-05 |

| | | | | |
|---|---|---|---|---|
| GO:0006109 | regulation of carbohydrate metabolic process | 92/7539 | 1.03E-05 | 5.80E-05 |
| GO:0008286 | insulin receptor signaling pathway | 63/7539 | 1.04E-05 | 5.87E-05 |
| GO:0010921 | regulation of phosphatase activity | 63/7539 | 1.04E-05 | 5.87E-05 |
| GO:0010761 | fibroblast migration | 35/7539 | 1.04E-05 | 5.87E-05 |
| GO:0014821 | phasic smooth muscle contraction | 20/7539 | 1.04E-05 | 5.87E-05 |
| GO:0030201 | heparan sulfate proteoglycan metabolic process | 20/7539 | 1.04E-05 | 5.87E-05 |
| GO:1903010 | regulation of bone development | 20/7539 | 1.04E-05 | 5.87E-05 |
| GO:0097530 | granulocyte migration | 76/7539 | 1.08E-05 | 6.06E-05 |
| GO:0021695 | cerebellar cortex development | 39/7539 | 1.12E-05 | 6.28E-05 |
| GO:0030318 | melanocyte differentiation | 21/7539 | 1.12E-05 | 6.28E-05 |
| GO:0045822 | negative regulation of heart contraction | 21/7539 | 1.12E-05 | 6.28E-05 |
| GO:0030513 | positive regulation of BMP signaling pathway | 26/7539 | 1.13E-05 | 6.32E-05 |
| GO:0031279 | regulation of cyclase activity | 26/7539 | 1.13E-05 | 6.32E-05 |
| GO:0060251 | regulation of glial cell proliferation | 26/7539 | 1.13E-05 | 6.32E-05 |
| GO:0090049 | regulation of cell migration involved in sprouting angiogenesis | 26/7539 | 1.13E-05 | 6.32E-05 |
| GO:1901385 | regulation of voltage-gated calcium channel activity | 26/7539 | 1.13E-05 | 6.32E-05 |
| GO:0044070 | regulation of anion transport | 67/7539 | 1.14E-05 | 6.34E-05 |
| GO:0010660 | regulation of muscle cell apoptotic process | 52/7539 | 1.15E-05 | 6.42E-05 |

| | | | | |
|---|---|---|---|---|
| GO:0098751 | bone cell development | 25/7539 | 1.17E-05 | 6.47E-05 |
| GO:0022011 | myelination in peripheral nervous system | 22/7539 | 1.17E-05 | 6.47E-05 |
| GO:0032292 | peripheral nervous system axon ensheathment | 22/7539 | 1.17E-05 | 6.47E-05 |
| GO:0042481 | regulation of odontogenesis | 22/7539 | 1.17E-05 | 6.47E-05 |
| GO:0045932 | negative regulation of muscle contraction | 22/7539 | 1.17E-05 | 6.47E-05 |
| GO:0051497 | negative regulation of stress fiber assembly | 22/7539 | 1.17E-05 | 6.47E-05 |
| GO:1905314 | semi-lunar valve development | 22/7539 | 1.17E-05 | 6.47E-05 |
| GO:0021520 | spinal cord motor neuron cell fate specification | Dec-39 | 1.18E-05 | 6.51E-05 |
| GO:0071679 | commissural neuron axon guidance | Dec-39 | 1.18E-05 | 6.51E-05 |
| GO:1902285 | semaphorin-plexin signaling pathway involved in neuron projection guidance | Dec-39 | 1.18E-05 | 6.51E-05 |
| GO:0019216 | regulation of lipid metabolic process | 156/7539 | 1.18E-05 | 6.52E-05 |
| GO:0010591 | regulation of lamellipodium assembly | 24/7539 | 1.19E-05 | 6.54E-05 |
| GO:0014047 | glutamate secretion | 24/7539 | 1.19E-05 | 6.54E-05 |
| GO:0060487 | lung epithelial cell differentiation | 23/7539 | 1.19E-05 | 6.54E-05 |
| GO:0060740 | prostate gland epithelium morphogenesis | 23/7539 | 1.19E-05 | 6.54E-05 |
| GO:0060914 | heart formation | 23/7539 | 1.19E-05 | 6.54E-05 |
| GO:0010002 | cardioblast differentiation | 15/7539 | 1.20E-05 | 6.60E-05 |
| GO:0033604 | negative regulation of catecholamine secretion | 15/7539 | 1.20E-05 | 6.60E-05 |

| | | | | |
|---|---|---|---|---|
| GO:0060572 | morphogenesis of an epithelial bud | 15/7539 | 1.20E-05 | 6.60E-05 |
| GO:0021766 | hippocampus development | 42/7539 | 1.24E-05 | 6.80E-05 |
| GO:0051260 | protein homooligomerization | 83/7539 | 1.25E-05 | 6.82E-05 |
| GO:1901216 | positive regulation of neuron death | 64/7539 | 1.26E-05 | 6.88E-05 |
| GO:0007019 | microtubule depolymerization | 33/7539 | 1.27E-05 | 6.94E-05 |
| GO:1900024 | regulation of substrate adhesion-dependent cell spreading | 33/7539 | 1.27E-05 | 6.94E-05 |
| GO:0070507 | regulation of microtubule cytoskeleton organization | 94/7539 | 1.29E-05 | 7.05E-05 |
| GO:0010927 | cellular component assembly involved in morphogenesis | 59/7539 | 1.30E-05 | 7.08E-05 |
| GO:0045682 | regulation of epidermis development | 45/7539 | 1.31E-05 | 7.15E-05 |
| GO:0046683 | response to organophosphorus | 48/7539 | 1.34E-05 | 7.27E-05 |
| GO:0001974 | blood vessel remodeling | 32/7539 | 1.40E-05 | 7.60E-05 |
| GO:0071241 | cellular response to inorganic substance | 104/7539 | 1.40E-05 | 7.62E-05 |
| GO:0010633 | negative regulation of epithelial cell migration | 41/7539 | 1.41E-05 | 7.64E-05 |
| GO:0046661 | male sex differentiation | 82/7539 | 1.43E-05 | 7.77E-05 |
| GO:0030308 | negative regulation of cell growth | 93/7539 | 1.48E-05 | 8.03E-05 |
| GO:0044782 | cilium organization | 148/7539 | 1.52E-05 | 8.20E-05 |
| GO:0030166 | proteoglycan biosynthetic process | 31/7539 | 1.53E-05 | 8.29E-05 |
| GO:0002695 | negative regulation of leukocyte activation | 89/7539 | 1.55E-05 | 8.40E-05 |

| | | | | |
|---|---|---|---|---|
| GO:0030512 | negative regulation of transforming growth factor beta receptor signaling pathway | 40/7539 | 1.59E-05 | 8.60E-05 |
| GO:0002683 | negative regulation of immune system process | 192/7539 | 1.59E-05 | 8.60E-05 |
| GO:0010763 | positive regulation of fibroblast migration | 16/7539 | 1.60E-05 | 8.62E-05 |
| GO:0072189 | ureter development | 16/7539 | 1.60E-05 | 8.62E-05 |
| GO:0046634 | regulation of alpha-beta T cell activation | 55/7539 | 1.60E-05 | 8.62E-05 |
| GO:0046660 | female sex differentiation | 73/7539 | 1.65E-05 | 8.86E-05 |
| GO:0009266 | response to temperature stimulus | 77/7539 | 1.66E-05 | 8.93E-05 |
| GO:0031111 | negative regulation of microtubule polymerization or depolymerization | 30/7539 | 1.67E-05 | 8.95E-05 |
| GO:0050850 | positive regulation of calcium-mediated signaling | 30/7539 | 1.67E-05 | 8.95E-05 |
| GO:1990573 | potassium ion import across plasma membrane | 30/7539 | 1.67E-05 | 8.95E-05 |
| GO:0060395 | SMAD protein signal transduction | 52/7539 | 1.69E-05 | 9.02E-05 |
| GO:0140029 | exocytic process | 52/7539 | 1.69E-05 | 9.02E-05 |
| GO:0031110 | regulation of microtubule polymerization or depolymerization | 49/7539 | 1.74E-05 | 9.30E-05 |
| GO:0061515 | myeloid cell development | 46/7539 | 1.75E-05 | 9.35E-05 |
| GO:0007588 | excretion | 35/7539 | 1.77E-05 | 9.41E-05 |
| GO:0060443 | mammary gland morphogenesis | 35/7539 | 1.77E-05 | 9.41E-05 |
| GO:0032413 | negative regulation of ion transmembrane transporter activity | 39/7539 | 1.80E-05 | 9.58E-05 |
| GO:0046579 | positive regulation of Ras protein signal transduction | 39/7539 | 1.80E-05 | 9.58E-05 |

| | | | | |
|---|---|---|---|---|
| GO:0071868 | cellular response to monoamine stimulus | 39/7539 | 1.80E-05 | 9.58E-05 |
| GO:0071870 | cellular response to catecholamine stimulus | 39/7539 | 1.80E-05 | 9.58E-05 |
| GO:0042176 | regulation of protein catabolic process | 167/7539 | 1.81E-05 | 9.61E-05 |
| GO:0003254 | regulation of membrane depolarization | 29/7539 | 1.81E-05 | 9.61E-05 |
| GO:0031952 | regulation of protein autophosphorylation | 29/7539 | 1.81E-05 | 9.61E-05 |
| GO:0060323 | head morphogenesis | 29/7539 | 1.81E-05 | 9.61E-05 |
| GO:0070661 | leukocyte proliferation | 148/7539 | 1.82E-05 | 9.62E-05 |
| GO:2001234 | negative regulation of apoptotic signaling pathway | 105/7539 | 1.94E-05 | 0.000102533 |
| GO:1902547 | regulation of cellular response to vascular endothelial growth factor stimulus | 17/7539 | 1.95E-05 | 0.000102754 |
| GO:0086002 | cardiac muscle cell action potential involved in contraction | 28/7539 | 1.96E-05 | 0.000103286 |
| GO:1903672 | positive regulation of sprouting angiogenesis | 28/7539 | 1.96E-05 | 0.000103286 |
| GO:0051591 | response to cAMP | 34/7539 | 1.97E-05 | 0.000103829 |
| GO:0045639 | positive regulation of myeloid cell differentiation | 56/7539 | 1.99E-05 | 0.00010454 |
| GO:0032370 | positive regulation of lipid transport | 45/7539 | 2.00E-05 | 0.000105332 |
| GO:0010762 | regulation of fibroblast migration | 27/7539 | 2.10E-05 | 0.000110246 |
| GO:0021955 | central nervous system neuron axonogenesis | 27/7539 | 2.10E-05 | 0.000110246 |
| GO:0055023 | positive regulation of cardiac muscle tissue growth | 27/7539 | 2.10E-05 | 0.000110246 |
| GO:0014848 | urinary tract smooth muscle contraction | 13/7539 | 2.15E-05 | 0.000112177 |

| | | | | |
|---|---|---|---|---|
| GO:0060442 | branching involved in prostate gland morphogenesis | 13/7539 | 2.15E-05 | 0.000112177 |
| GO:0061029 | eyelid development in camera-type eye | 13/7539 | 2.15E-05 | 0.000112177 |
| GO:1903587 | regulation of blood vessel endothelial cell proliferation involved in sprouting angiogenesis | 13/7539 | 2.15E-05 | 0.000112177 |
| GO:2000095 | regulation of Wnt signaling pathway, planar cell polarity pathway | 13/7539 | 2.15E-05 | 0.000112177 |
| GO:0046546 | development of primary male sexual characteristics | 69/7539 | 2.16E-05 | 0.000112901 |
| GO:0032892 | positive regulation of organic acid transport | 33/7539 | 2.19E-05 | 0.000114283 |
| GO:0051496 | positive regulation of stress fiber assembly | 33/7539 | 2.19E-05 | 0.000114283 |
| GO:0061001 | regulation of dendritic spine morphogenesis | 33/7539 | 2.19E-05 | 0.000114283 |
| GO:0010950 | positive regulation of endopeptidase activity | 77/7539 | 2.20E-05 | 0.000114381 |
| GO:0015908 | fatty acid transport | 50/7539 | 2.23E-05 | 0.000115829 |
| GO:0048741 | skeletal muscle fiber development | 26/7539 | 2.24E-05 | 0.000116139 |
| GO:0085029 | extracellular matrix assembly | 26/7539 | 2.24E-05 | 0.000116139 |
| GO:1901880 | negative regulation of protein depolymerization | 44/7539 | 2.29E-05 | 0.000118771 |
| GO:0043388 | positive regulation of DNA binding | 37/7539 | 2.30E-05 | 0.000119135 |
| GO:0099637 | neurotransmitter receptor transport | 37/7539 | 2.30E-05 | 0.000119135 |
| GO:1902305 | regulation of sodium ion transmembrane transport | 37/7539 | 2.30E-05 | 0.000119135 |
| GO:0001953 | negative regulation of cell-matrix adhesion | 25/7539 | 2.36E-05 | 0.000121905 |
| GO:0048265 | response to pain | 25/7539 | 2.36E-05 | 0.000121905 |

| | | | | |
|---|---|---|---|---|
| GO:2000463 | positive regulation of excitatory postsynaptic potential | 25/7539 | 2.36E-05 | 0.000121905 |
| GO:0021795 | cerebral cortex cell migration | 32/7539 | 2.44E-05 | 0.000125377 |
| GO:0046676 | negative regulation of insulin secretion | 32/7539 | 2.44E-05 | 0.000125377 |
| GO:0048662 | negative regulation of smooth muscle cell proliferation | 32/7539 | 2.44E-05 | 0.000125377 |
| GO:0055010 | ventricular cardiac muscle tissue morphogenesis | 32/7539 | 2.44E-05 | 0.000125377 |
| GO:0007200 | phospholipase C-activating G protein-coupled receptor signaling pathway | 52/7539 | 2.44E-05 | 0.000125513 |
| GO:0009581 | detection of external stimulus | 66/7539 | 2.45E-05 | 0.000125788 |
| GO:0003401 | axis elongation | 24/7539 | 2.47E-05 | 0.0001263 |
| GO:0035115 | embryonic forelimb morphogenesis | 24/7539 | 2.47E-05 | 0.0001263 |
| GO:0042220 | response to cocaine | 24/7539 | 2.47E-05 | 0.0001263 |
| GO:0048679 | regulation of axon regeneration | 24/7539 | 2.47E-05 | 0.0001263 |
| GO:0009895 | negative regulation of catabolic process | 135/7539 | 2.47E-05 | 0.0001263 |
| GO:0071875 | adrenergic receptor signaling pathway | 20/7539 | 2.54E-05 | 0.000129525 |
| GO:2000178 | negative regulation of neural precursor cell proliferation | 20/7539 | 2.54E-05 | 0.000129525 |
| GO:0014044 | Schwann cell development | 23/7539 | 2.55E-05 | 0.000129726 |
| GO:0048710 | regulation of astrocyte differentiation | 23/7539 | 2.55E-05 | 0.000129726 |
| GO:0060479 | lung cell differentiation | 23/7539 | 2.55E-05 | 0.000129726 |
| GO:0060512 | prostate gland morphogenesis | 23/7539 | 2.55E-05 | 0.000129726 |

| | | | | |
|---|---|---|---|---|
| GO:0006929 | substrate-dependent cell migration | 21/7539 | 2.59E-05 | 0.000131374 |
| GO:0035767 | endothelial cell chemotaxis | 21/7539 | 2.59E-05 | 0.000131374 |
| GO:0048521 | negative regulation of behavior | 21/7539 | 2.59E-05 | 0.000131374 |
| GO:0060317 | cardiac epithelial to mesenchymal transition | 21/7539 | 2.59E-05 | 0.000131374 |
| GO:0021697 | cerebellar cortex formation | 22/7539 | 2.59E-05 | 0.000131374 |
| GO:0090075 | relaxation of muscle | 22/7539 | 2.59E-05 | 0.000131374 |
| GO:0061337 | cardiac conduction | 36/7539 | 2.60E-05 | 0.000131437 |
| GO:0070373 | negative regulation of ERK1 and ERK2 cascade | 43/7539 | 2.62E-05 | 0.000132402 |
| GO:1902106 | negative regulation of leukocyte differentiation | 54/7539 | 2.63E-05 | 0.000133182 |
| GO:0001819 | positive regulation of cytokine production | 186/7539 | 2.68E-05 | 0.000135269 |
| GO:0046189 | phenol-containing compound biosynthetic process | 31/7539 | 2.70E-05 | 0.000136182 |
| GO:0010952 | positive regulation of peptidase activity | 81/7539 | 2.81E-05 | 0.000141757 |
| GO:1900076 | regulation of cellular response to insulin stimulus | 48/7539 | 2.94E-05 | 0.000148241 |
| GO:0051235 | maintenance of location | 138/7539 | 2.98E-05 | 0.000149837 |
| GO:0061028 | establishment of endothelial barrier | 30/7539 | 2.98E-05 | 0.000149837 |
| GO:0003198 | epithelial to mesenchymal transition involved in endocardial cushion formation | 14/7539 | 3.13E-05 | 0.000157164 |
| GO:0051648 | vesicle localization | 82/7539 | 3.17E-05 | 0.000159258 |
| GO:0042596 | fear response | 34/7539 | 3.29E-05 | 0.000164951 |

| | | | | |
|---|---|---|---|---|
| GO:0008584 | male gonad development | 68/7539 | 3.35E-05 | 0.000167923 |
| GO:0014832 | urinary bladder smooth muscle contraction | Nov-39 | 3.38E-05 | 0.000168493 |
| GO:0021561 | facial nerve development | Nov-39 | 3.38E-05 | 0.000168493 |
| GO:0021610 | facial nerve morphogenesis | Nov-39 | 3.38E-05 | 0.000168493 |
| GO:0048241 | epinephrine transport | Nov-39 | 3.38E-05 | 0.000168493 |
| GO:1902287 | semaphorin-plexin signaling pathway involved in axon guidance | Nov-39 | 3.38E-05 | 0.000168493 |
| GO:1903431 | positive regulation of cell maturation | Nov-39 | 3.38E-05 | 0.000168493 |
| GO:0007009 | plasma membrane organization | 57/7539 | 3.41E-05 | 0.000169851 |
| GO:0050670 | regulation of lymphocyte proliferation | 104/7539 | 3.45E-05 | 0.000171898 |
| GO:0050870 | positive regulation of T cell activation | 92/7539 | 3.52E-05 | 0.000175219 |
| GO:0001569 | branching involved in blood vessel morphogenesis | 28/7539 | 3.59E-05 | 0.000178297 |
| GO:0045494 | photoreceptor cell maintenance | 28/7539 | 3.59E-05 | 0.000178297 |
| GO:0030199 | collagen fibril organization | 33/7539 | 3.69E-05 | 0.000182805 |
| GO:0071715 | icosanoid transport | 33/7539 | 3.69E-05 | 0.000182805 |
| GO:1901571 | fatty acid derivative transport | 33/7539 | 3.69E-05 | 0.000182805 |
| GO:1902893 | regulation of pri-miRNA transcription by RNA polymerase II | 33/7539 | 3.69E-05 | 0.000182805 |
| GO:0009798 | axis specification | 49/7539 | 3.72E-05 | 0.000184325 |
| GO:0008277 | regulation of G protein-coupled receptor signaling pathway | 63/7539 | 3.76E-05 | 0.000185649 |

| | | | | |
|---|---|---|---|---|
| GO:0032944 | regulation of mononuclear cell proliferation | 105/7539 | 3.76E-05 | 0.000185649 |
| GO:0010675 | regulation of cellular carbohydrate metabolic process | 77/7539 | 3.78E-05 | 0.000186524 |
| GO:0044262 | cellular carbohydrate metabolic process | 127/7539 | 3.79E-05 | 0.000186922 |
| GO:0009582 | detection of abiotic stimulus | 65/7539 | 3.82E-05 | 0.000188369 |
| GO:0051346 | negative regulation of hydrolase activity | 169/7539 | 3.83E-05 | 0.000188743 |
| GO:2001056 | positive regulation of cysteine-type endopeptidase activity | 69/7539 | 3.88E-05 | 0.000191015 |
| GO:0007193 | adenylate cyclase-inhibiting G protein-coupled receptor signaling pathway | 40/7539 | 3.90E-05 | 0.000191797 |
| GO:1903845 | negative regulation of cellular response to transforming growth factor beta stimulus | 40/7539 | 3.90E-05 | 0.000191797 |
| GO:0045197 | establishment or maintenance of epithelial cell apical/basal polarity | 27/7539 | 3.91E-05 | 0.000192021 |
| GO:0120034 | positive regulation of plasma membrane bounded cell projection assembly | 56/7539 | 3.93E-05 | 0.000192848 |
| GO:0001975 | response to amphetamine | 15/7539 | 4.00E-05 | 0.000195018 |
| GO:0003180 | aortic valve morphogenesis | 15/7539 | 4.00E-05 | 0.000195018 |
| GO:0003215 | cardiac right ventricle morphogenesis | 15/7539 | 4.00E-05 | 0.000195018 |
| GO:0035930 | corticosteroid hormone secretion | 15/7539 | 4.00E-05 | 0.000195018 |
| GO:0043116 | negative regulation of vascular permeability | 15/7539 | 4.00E-05 | 0.000195018 |
| GO:0060231 | mesenchymal to epithelial transition | 15/7539 | 4.00E-05 | 0.000195018 |
| GO:0072578 | neurotransmitter-gated ion channel clustering | 15/7539 | 4.00E-05 | 0.000195018 |
| GO:1904754 | positive regulation of vascular associated smooth muscle cell migration | 15/7539 | 4.00E-05 | 0.000195018 |

| | | | | |
|---|---|---|---|---|
| GO:0018958 | phenol-containing compound metabolic process | 58/7539 | 4.09E-05 | 0.000199605 |
| GO:0014009 | glial cell proliferation | 32/7539 | 4.13E-05 | 0.000201134 |
| GO:0046717 | acid secretion | 32/7539 | 4.13E-05 | 0.000201134 |
| GO:0010830 | regulation of myotube differentiation | 36/7539 | 4.18E-05 | 0.000203294 |
| GO:0010470 | regulation of gastrulation | 26/7539 | 4.24E-05 | 0.000205513 |
| GO:0014037 | Schwann cell differentiation | 26/7539 | 4.24E-05 | 0.000205513 |
| GO:0098801 | regulation of renal system process | 26/7539 | 4.24E-05 | 0.000205513 |
| GO:0009416 | response to light stimulus | 129/7539 | 4.30E-05 | 0.000208726 |
| GO:0008064 | regulation of actin polymerization or depolymerization | 87/7539 | 4.31E-05 | 0.000208958 |
| GO:0042752 | regulation of circadian rhythm | 64/7539 | 4.40E-05 | 0.000213206 |
| GO:0043954 | cellular component maintenance | 39/7539 | 4.45E-05 | 0.000215194 |
| GO:0048332 | mesoderm morphogenesis | 39/7539 | 4.45E-05 | 0.000215194 |
| GO:0043255 | regulation of carbohydrate biosynthetic process | 55/7539 | 4.53E-05 | 0.000218813 |
| GO:0003407 | neural retina development | 42/7539 | 4.53E-05 | 0.000218813 |
| GO:0031334 | positive regulation of protein-containing complex assembly | 105/7539 | 4.65E-05 | 0.000224205 |
| GO:0001502 | cartilage condensation | 16/7539 | 4.70E-05 | 0.000225749 |
| GO:0003177 | pulmonary valve development | 16/7539 | 4.70E-05 | 0.000225749 |
| GO:0014075 | response to amine | 16/7539 | 4.70E-05 | 0.000225749 |

| | | | | |
|---|---|---|---|---|
| GO:0031290 | retinal ganglion cell axon guidance | 16/7539 | 4.70E-05 | 0.000225749 |
| GO:0031641 | regulation of myelination | 35/7539 | 4.73E-05 | 0.000227463 |
| GO:0030832 | regulation of actin filament length | 88/7539 | 4.78E-05 | 0.00022955 |
| GO:2000273 | positive regulation of signaling receptor activity | 24/7539 | 4.87E-05 | 0.00023367 |
| GO:0018107 | peptidyl-threonine phosphorylation | 59/7539 | 4.88E-05 | 0.000233818 |
| GO:0006644 | phospholipid metabolic process | 157/7539 | 4.92E-05 | 0.000235688 |
| GO:0042490 | mechanoreceptor differentiation | 47/7539 | 4.93E-05 | 0.000236048 |
| GO:0071456 | cellular response to hypoxia | 52/7539 | 4.96E-05 | 0.000237127 |
| GO:0051250 | negative regulation of lymphocyte activation | 75/7539 | 4.99E-05 | 0.0002387 |
| GO:0044770 | cell cycle phase transition | 172/7539 | 5.10E-05 | 0.000243615 |
| GO:0032309 | icosanoid secretion | 30/7539 | 5.15E-05 | 0.000245709 |
| GO:0060441 | epithelial tube branching involved in lung morphogenesis | 23/7539 | 5.16E-05 | 0.000245903 |
| GO:0032331 | negative regulation of chondrocyte differentiation | 17/7539 | 5.20E-05 | 0.000247572 |
| GO:2000831 | regulation of steroid hormone secretion | 17/7539 | 5.20E-05 | 0.000247572 |
| GO:0030316 | osteoclast differentiation | 56/7539 | 5.45E-05 | 0.000259266 |
| GO:0060004 | reflex | 18/7539 | 5.52E-05 | 0.000262124 |
| GO:0086019 | cell-cell signaling involved in cardiac conduction | 18/7539 | 5.52E-05 | 0.000262124 |
| GO:0035640 | exploration behavior | 21/7539 | 5.58E-05 | 0.000265088 |

| | | | | |
|---|---|---|---|---|
| GO:0009755 | hormone-mediated signaling pathway | 76/7539 | 5.64E-05 | 0.000267648 |
| GO:0001956 | positive regulation of neurotransmitter secretion | 19/7539 | 5.67E-05 | 0.000267936 |
| GO:0003171 | atrioventricular valve development | 19/7539 | 5.67E-05 | 0.000267936 |
| GO:0010742 | macrophage derived foam cell differentiation | 19/7539 | 5.67E-05 | 0.000267936 |
| GO:0014829 | vascular associated smooth muscle contraction | 19/7539 | 5.67E-05 | 0.000267936 |
| GO:0050926 | regulation of positive chemotaxis | 19/7539 | 5.67E-05 | 0.000267936 |
| GO:0021952 | central nervous system projection neuron axonogenesis | 20/7539 | 5.68E-05 | 0.000268342 |
| GO:0001707 | mesoderm formation | 37/7539 | 5.79E-05 | 0.0002729 |
| GO:0042982 | amyloid precursor protein metabolic process | 37/7539 | 5.79E-05 | 0.0002729 |
| GO:0008643 | carbohydrate transport | 72/7539 | 5.83E-05 | 0.000274568 |
| GO:0043536 | positive regulation of blood vessel endothelial cell migration | 33/7539 | 6.05E-05 | 0.000284928 |
| GO:0090630 | activation of GTPase activity | 48/7539 | 6.17E-05 | 0.000290077 |
| GO:0035303 | regulation of dephosphorylation | 79/7539 | 6.19E-05 | 0.000291154 |
| GO:0030041 | actin filament polymerization | 86/7539 | 6.28E-05 | 0.000294777 |
| GO:0010259 | multicellular organism aging | 28/7539 | 6.34E-05 | 0.000297727 |
| GO:0033135 | regulation of peptidyl-serine phosphorylation | 77/7539 | 6.35E-05 | 0.000297727 |
| GO:0006835 | dicarboxylic acid transport | 45/7539 | 6.53E-05 | 0.00030599 |
| GO:0043367 | CD4-positive, alpha-beta T cell differentiation | 45/7539 | 6.53E-05 | 0.00030599 |

| | | | | |
|---|---|---|---|---|
| GO:0045668 | negative regulation of osteoblast differentiation | 36/7539 | 6.59E-05 | 0.00030835 |
| GO:1903510 | mucopolysaccharide metabolic process | 36/7539 | 6.59E-05 | 0.00030835 |
| GO:0060042 | retina morphogenesis in camera-type eye | 42/7539 | 6.77E-05 | 0.000316312 |
| GO:0061008 | hepaticobiliary system development | 63/7539 | 6.82E-05 | 0.000318361 |
| GO:0007605 | sensory perception of sound | 119/7539 | 7.03E-05 | 0.000327841 |
| GO:0043242 | negative regulation of protein-containing complex disassembly | 47/7539 | 7.11E-05 | 0.00033134 |
| GO:0015695 | organic cation transport | 35/7539 | 7.51E-05 | 0.000349718 |
| GO:1902115 | regulation of organelle assembly | 93/7539 | 7.67E-05 | 0.000356987 |
| GO:0030850 | prostate gland development | 31/7539 | 7.69E-05 | 0.000357617 |
| GO:0044058 | regulation of digestive system process | 31/7539 | 7.69E-05 | 0.000357617 |
| GO:0051339 | regulation of lyase activity | 26/7539 | 7.70E-05 | 0.000357862 |
| GO:0031333 | negative regulation of protein-containing complex assembly | 70/7539 | 7.72E-05 | 0.000358275 |
| GO:0031109 | microtubule polymerization or depolymerization | 60/7539 | 7.79E-05 | 0.000361427 |
| GO:0040034 | regulation of development, heterochronic | 13/7539 | 8.03E-05 | 0.000371466 |
| GO:0043584 | nose development | 13/7539 | 8.03E-05 | 0.000371466 |
| GO:0090128 | regulation of synapse maturation | 13/7539 | 8.03E-05 | 0.000371466 |
| GO:0090494 | dopamine uptake | 13/7539 | 8.03E-05 | 0.000371466 |
| GO:0032103 | positive regulation of response to external stimulus | 172/7539 | 8.04E-05 | 0.000371508 |

| | | | | |
|---|---|---|---|---|
| GO:0050953 | sensory perception of light stimulus | 77/7539 | 8.17E-05 | 0.0003771 |
| GO:0006112 | energy reserve metabolic process | 46/7539 | 8.19E-05 | 0.00037797 |
| GO:0051302 | regulation of cell division | 75/7539 | 8.38E-05 | 0.00038636 |
| GO:0014742 | positive regulation of muscle hypertrophy | 25/7539 | 8.43E-05 | 0.000387329 |
| GO:0015804 | neutral amino acid transport | 25/7539 | 8.43E-05 | 0.000387329 |
| GO:0048520 | positive regulation of behavior | 25/7539 | 8.43E-05 | 0.000387329 |
| GO:0060259 | regulation of feeding behavior | 25/7539 | 8.43E-05 | 0.000387329 |
| GO:0060325 | face morphogenesis | 25/7539 | 8.43E-05 | 0.000387329 |
| GO:0010524 | positive regulation of calcium ion transport into cytosol | 34/7539 | 8.54E-05 | 0.000391715 |
| GO:0060711 | labyrinthine layer development | 34/7539 | 8.54E-05 | 0.000391715 |
| GO:0071470 | cellular response to osmotic stress | 30/7539 | 8.65E-05 | 0.000396576 |
| GO:0043280 | positive regulation of cysteine-type endopeptidase activity involved in apoptotic process | 61/7539 | 9.07E-05 | 0.000415327 |
| GO:0043267 | negative regulation of potassium ion transport | 24/7539 | 9.17E-05 | 0.000419629 |
| GO:1903170 | negative regulation of calcium ion transmembrane transport | 24/7539 | 9.17E-05 | 0.000419629 |
| GO:0021559 | trigeminal nerve development | Oct-39 | 9.62E-05 | 0.000438725 |
| GO:0051458 | corticotropin secretion | Oct-39 | 9.62E-05 | 0.000438725 |
| GO:0071415 | cellular response to purine-containing compound | Oct-39 | 9.62E-05 | 0.000438725 |
| GO:0072102 | glomerulus morphogenesis | Oct-39 | 9.62E-05 | 0.000438725 |

| | | | | |
|---|---|---|---|---|
| GO:0010517 | regulation of phospholipase activity | 33/7539 | 9.70E-05 | 0.000441813 |
| GO:0061614 | pri-miRNA transcription by RNA polymerase II | 33/7539 | 9.70E-05 | 0.000441813 |
| GO:0090659 | walking behavior | 29/7539 | 9.71E-05 | 0.000441813 |
| GO:0003128 | heart field specification | 14/7539 | 9.86E-05 | 0.000447039 |
| GO:0031281 | positive regulation of cyclase activity | 14/7539 | 9.86E-05 | 0.000447039 |
| GO:0051580 | regulation of neurotransmitter uptake | 14/7539 | 9.86E-05 | 0.000447039 |
| GO:0086011 | membrane repolarization during action potential | 14/7539 | 9.86E-05 | 0.000447039 |
| GO:2000846 | regulation of corticosteroid hormone secretion | 14/7539 | 9.86E-05 | 0.000447039 |
| GO:0010453 | regulation of cell fate commitment | 23/7539 | 9.92E-05 | 0.000448385 |
| GO:0035886 | vascular associated smooth muscle cell differentiation | 23/7539 | 9.92E-05 | 0.000448385 |
| GO:0061384 | heart trabecula morphogenesis | 23/7539 | 9.92E-05 | 0.000448385 |
| GO:0071604 | transforming growth factor beta production | 23/7539 | 9.92E-05 | 0.000448385 |
| GO:0030833 | regulation of actin filament polymerization | 79/7539 | 0.000101206 | 0.000457021 |
| GO:0072593 | reactive oxygen species metabolic process | 125/7539 | 0.000101206 | 0.000457021 |
| GO:0021782 | glial cell development | 58/7539 | 0.000103702 | 0.000467958 |
| GO:2001235 | positive regulation of apoptotic signaling pathway | 84/7539 | 0.000103904 | 0.000468534 |
| GO:0044772 | mitotic cell cycle phase transition | 156/7539 | 0.000104288 | 0.000469927 |
| GO:0019934 | cGMP-mediated signaling | 22/7539 | 0.000106334 | 0.000477444 |

| | | | | |
|---|---|---|---|---|
| GO:0035329 | hippo signaling | 22/7539 | 0.000106334 | 0.000477444 |
| GO:0043950 | positive regulation of cAMP-mediated signaling | 22/7539 | 0.000106334 | 0.000477444 |
| GO:0071634 | regulation of transforming growth factor beta production | 22/7539 | 0.000106334 | 0.000477444 |
| GO:0090162 | establishment of epithelial cell polarity | 22/7539 | 0.000106334 | 0.000477444 |
| GO:0048545 | response to steroid hormone | 97/7539 | 0.000106942 | 0.000479832 |
| GO:0007628 | adult walking behavior | 28/7539 | 0.000108674 | 0.000486911 |
| GO:2001239 | regulation of extrinsic apoptotic signaling pathway in absence of ligand | 28/7539 | 0.000108674 | 0.000486911 |
| GO:0001895 | retina homeostasis | 32/7539 | 0.000110154 | 0.000492845 |
| GO:0043647 | inositol phosphate metabolic process | 32/7539 | 0.000110154 | 0.000492845 |
| GO:0001696 | gastric acid secretion | 15/7539 | 0.00011186 | 0.000498356 |
| GO:0048557 | embryonic digestive tract morphogenesis | 15/7539 | 0.00011186 | 0.000498356 |
| GO:0072079 | nephron tubule formation | 15/7539 | 0.00011186 | 0.000498356 |
| GO:0072087 | renal vesicle development | 15/7539 | 0.00011186 | 0.000498356 |
| GO:0072243 | metanephric nephron epithelium development | 15/7539 | 0.00011186 | 0.000498356 |
| GO:1900746 | regulation of vascular endothelial growth factor signaling pathway | 15/7539 | 0.00011186 | 0.000498356 |
| GO:0051651 | maintenance of location in cell | 95/7539 | 0.000112538 | 0.000501023 |
| GO:0032958 | inositol phosphate biosynthetic process | 21/7539 | 0.000112965 | 0.000502218 |
| GO:0051150 | regulation of smooth muscle cell differentiation | 21/7539 | 0.000112965 | 0.000502218 |

| | | | | |
|---|---|---|---|---|
| GO:0002262 | myeloid cell homeostasis | 85/7539 | 0.000114131 | 0.000507044 |
| GO:0031330 | negative regulation of cellular catabolic process | 109/7539 | 0.000114479 | 0.000508231 |
| GO:0006809 | nitric oxide biosynthetic process | 41/7539 | 0.0001146 6 | 0.000508678 |
| GO:0098656 | anion transmembrane transport | 98/7539 | 0.000115585 | 0.000512421 |
| GO:0046651 | lymphocyte proliferation | 134/7539 | 0.000116273 | 0.000515108 |
| GO:0010922 | positive regulation of phosphatase activity | 20/7539 | 0.000118645 | 0.000524513 |
| GO:0035162 | embryonic hemopoiesis | 20/7539 | 0.000118645 | 0.000524513 |
| GO:0071711 | basement membrane organization | 20/7539 | 0.000118645 | 0.000524513 |
| GO:0010970 | transport along microtubule | 76/7539 | 0.000119899 | 0.000529689 |
| GO:0015012 | heparan sulfate proteoglycan biosynthetic process | 16/7539 | 0.00012029 | 0.000530299 |
| GO:0032148 | activation of protein kinase B activity | 16/7539 | 0.00012029 | 0.000530299 |
| GO:0048532 | anatomical structure arrangement | 16/7539 | 0.00012029 | 0.000530299 |
| GO:0002688 | regulation of leukocyte chemotaxis | 59/7539 | 0.000120586 | 0.000531235 |
| GO:0001889 | liver development | 61/7539 | 0.000120772 | 0.000531682 |
| GO:0032943 | mononuclear cell proliferation | 135/7539 | 0.000122302 | 0.000538043 |
| GO:0006837 | serotonin transport | 19/7539 | 0.000122871 | 0.000539419 |
| GO:0061217 | regulation of mesonephros development | 19/7539 | 0.000122871 | 0.000539419 |
| GO:0090077 | foam cell differentiation | 19/7539 | 0.000122871 | 0.000539419 |

| | | | | |
|---|---|---|---|---|
| GO:0003181 | atrioventricular valve morphogenesis | 17/7539 | 0.000124431 | 0.000543254 |
| GO:0044331 | cell-cell adhesion mediated by cadherin | 17/7539 | 0.000124431 | 0.000543254 |
| GO:0072207 | metanephric epithelium development | 17/7539 | 0.000124431 | 0.000543254 |
| GO:0052547 | regulation of peptidase activity | 172/7539 | 0.000124819 | 0.000543254 |
| GO:0002209 | behavioral defense response | 31/7539 | 0.000124917 | 0.000543254 |
| GO:0034113 | heterotypic cell-cell adhesion | 31/7539 | 0.000124917 | 0.000543254 |
| GO:2000649 | regulation of sodium ion transmembrane transporter activity | 31/7539 | 0.000124917 | 0.000543254 |
| GO:0030325 | adrenal gland development | 18/7539 | 0.000125035 | 0.000543254 |
| GO:0034104 | negative regulation of tissue remodeling | 18/7539 | 0.000125035 | 0.000543254 |
| GO:0035929 | steroid hormone secretion | 18/7539 | 0.000125035 | 0.000543254 |
| GO:0050927 | positive regulation of positive chemotaxis | 18/7539 | 0.000125035 | 0.000543254 |
| GO:0090280 | positive regulation of calcium ion import | 18/7539 | 0.000125035 | 0.000543254 |
| GO:0120033 | negative regulation of plasma membrane bounded cell projection assembly | 18/7539 | 0.000125035 | 0.000543254 |
| GO:0140058 | neuron projection arborization | 18/7539 | 0.000125035 | 0.000543254 |
| GO:2000738 | positive regulation of stem cell differentiation | 18/7539 | 0.000125035 | 0.000543254 |
| GO:0034101 | erythrocyte homeostasis | 70/7539 | 0.000130201 | 0.000565309 |
| GO:0046626 | regulation of insulin receptor signaling pathway | 40/7539 | 0.000131963 | 0.000572563 |
| GO:0046636 | negative regulation of alpha-beta T cell activation | 26/7539 | 0.00013508 | 0.000585689 |

| | | | | |
|---|---|---|---|---|
| GO:0009896 | positive regulation of catabolic process | 187/7539 | 0.000136545 | 0.000591631 |
| GO:1903038 | negative regulation of leukocyte cell-cell adhesion | 64/7539 | 0.000137307 | 0.000594527 |
| GO:0042491 | inner ear auditory receptor cell differentiation | 30/7539 | 0.000141479 | 0.000611749 |
| GO:1904036 | negative regulation of epithelial cell apoptotic process | 30/7539 | 0.000141479 | 0.000611749 |
| GO:0009991 | response to extracellular stimulus | 156/7539 | 0.000141718 | 0.0006122 |
| GO:0062013 | positive regulation of small molecule metabolic process | 73/7539 | 0.000141874 | 0.0006122 |
| GO:1903707 | negative regulation of hemopoiesis | 73/7539 | 0.000141874 | 0.0006122 |
| GO:0002793 | positive regulation of peptide secretion | 101/7539 | 0.000144232 | 0.000621952 |
| GO:0060603 | mammary gland duct morphogenesis | 25/7539 | 0.000149845 | 0.000644832 |
| GO:1900744 | regulation of p38MAPK cascade | 25/7539 | 0.000149845 | 0.000644832 |
| GO:1905332 | positive regulation of morphogenesis of an epithelium | 25/7539 | 0.000149845 | 0.000644832 |
| GO:0007183 | SMAD protein complex assembly | Nov-39 | 0.000154698 | 0.0006612 |
| GO:0021604 | cranial nerve structural organization | Nov-39 | 0.000154698 | 0.0006612 |
| GO:0031223 | auditory behavior | Nov-39 | 0.000154698 | 0.0006612 |
| GO:0035630 | bone mineralization involved in bone maturation | Nov-39 | 0.000154698 | 0.0006612 |
| GO:0045161 | neuronal ion channel clustering | Nov-39 | 0.000154698 | 0.0006612 |
| GO:0045906 | negative regulation of vasoconstriction | Nov-39 | 0.000154698 | 0.0006612 |
| GO:0060601 | lateral sprouting from an epithelium | Nov-39 | 0.000154698 | 0.0006612 |

| | | | | |
|---|---|---|---|---|
| GO:0070831 | basement membrane assembly | Nov-39 | 0.000154698 | 0.0006612 1 |
| GO:0072216 | positive regulation of metanephros development | Nov-39 | 0.000154698 | 0.0006612 1 |
| GO:2000833 | positive regulation of steroid hormone secretion | Nov-39 | 0.000154698 | 0.0006612 1 |
| GO:0006650 | glycerophospholipid metabolic process | 111/7539 | 0.000157316 | 0.000671946 |
| GO:0052548 | regulation of endopeptidase activity | 144/7539 | 0.000158605 | 0.000676992 |
| GO:0010613 | positive regulation of cardiac muscle hypertrophy | 24/7539 | 0.000165545 | 0.000706141 |
| GO:0030522 | intracellular receptor signaling pathway | 92/7539 | 0.000166468 | 0.000709595 |
| GO:0002066 | columnar/cuboidal epithelial cell development | 41/7539 | 0.000166706 | 0.000710131 |
| GO:0034614 | cellular response to reactive oxygen species | 70/7539 | 0.000167642 | 0.000713638 |
| GO:0001678 | cellular glucose homeostasis | 75/7539 | 0.0001749 | 0.000744031 |
| GO:0030168 | platelet activation | 43/7539 | 0.000179433 | 0.000762802 |
| GO:0003382 | epithelial cell morphogenesis | 28/7539 | 0.000180669 | 0.000766506 |
| GO:0038066 | p38MAPK cascade | 28/7539 | 0.000180669 | 0.000766506 |
| GO:1903115 | regulation of actin filament-based movement | 28/7539 | 0.000180669 | 0.000766506 |
| GO:0002691 | regulation of cellular extravasation | 23/7539 | 0.0001820 2 | 0.000771719 |
| GO:1903409 | reactive oxygen species biosynthetic process | 60/7539 | 0.000184084 | 0.0007799 5 |
| GO:0030216 | keratinocyte differentiation | 56/7539 | 0.000185119 | 0.000783807 |
| GO:0035176 | social behavior | 31/7539 | 0.0001980 6 | 0.000837478 |

| | | | | |
|---|---|---|---|---|
| GO:0045604 | regulation of epidermal cell differentiation | 31/7539 | 0.00019806 | 0.000837478 |
| GO:0043297 | apical junction assembly | 47/7539 | 0.000198472 | 0.000838116 |
| GO:0022411 | cellular component disassembly | 157/7539 | 0.000198595 | 0.000838116 |
| GO:0010719 | negative regulation of epithelial to mesenchymal transition | 22/7539 | 0.000199008 | 0.000838116 |
| GO:0086091 | regulation of heart rate by cardiac conduction | 22/7539 | 0.000199008 | 0.000838116 |
| GO:1990089 | response to nerve growth factor | 22/7539 | 0.000199008 | 0.000838116 |
| GO:1990090 | cellular response to nerve growth factor stimulus | 22/7539 | 0.000199008 | 0.000838116 |
| GO:0007601 | visual perception | 74/7539 | 0.000200503 | 0.000843851 |
| GO:0034394 | protein localization to cell surface | 37/7539 | 0.000201121 | 0.000845888 |
| GO:0030218 | erythrocyte differentiation | 65/7539 | 0.000202452 | 0.000850916 |
| GO:0032222 | regulation of synaptic transmission, cholinergic | Dec-39 | 0.000203748 | 0.00085272 |
| GO:0034698 | response to gonadotropin | Dec-39 | 0.000203748 | 0.00085272 |
| GO:0042428 | serotonin metabolic process | Dec-39 | 0.000203748 | 0.00085272 |
| GO:0086014 | atrial cardiac muscle cell action potential | Dec-39 | 0.000203748 | 0.00085272 |
| GO:0086026 | atrial cardiac muscle cell to AV node cell signaling | Dec-39 | 0.000203748 | 0.00085272 |
| GO:0086066 | atrial cardiac muscle cell to AV node cell communication | Dec-39 | 0.000203748 | 0.00085272 |
| GO:0051926 | negative regulation of calcium ion transport | 34/7539 | 0.000203961 | 0.00085272 |
| GO:0098930 | axonal transport | 34/7539 | 0.000203961 | 0.00085272 |

| | | | | |
|---|---|---|---|---|
| GO:0050728 | negative regulation of inflammatory response | 72/7539 | 0.000207697 | 0.000867763 |
| GO:0071496 | cellular response to external stimulus | 119/7539 | 0.000208544 | 0.000870728 |
| GO:0043244 | regulation of protein-containing complex disassembly | 61/7539 | 0.000209782 | 0.000875316 |
| GO:0050764 | regulation of phagocytosis | 55/7539 | 0.000213595 | 0.000890639 |
| GO:0022410 | circadian sleep/wake cycle process | 21/7539 | 0.000216124 | 0.000899401 |
| GO:0034260 | negative regulation of GTPase activity | 21/7539 | 0.000216124 | 0.000899401 |
| GO:1902745 | positive regulation of lamellipodium organization | 21/7539 | 0.000216124 | 0.000899401 |
| GO:0043433 | negative regulation of DNA-binding transcription factor activity | 75/7539 | 0.000221248 | 0.000920119 |
| GO:0001662 | behavioral fear response | 30/7539 | 0.000225675 | 0.000935449 |
| GO:0010596 | negative regulation of endothelial cell migration | 30/7539 | 0.000225675 | 0.000935449 |
| GO:0043666 | regulation of phosphoprotein phosphatase activity | 30/7539 | 0.000225675 | 0.000935449 |
| GO:0051955 | regulation of amino acid transport | 30/7539 | 0.000225675 | 0.000935449 |
| GO:0097720 | calcineurin-mediated signaling | 30/7539 | 0.000225675 | 0.000935449 |
| GO:0021522 | spinal cord motor neuron differentiation | 26/7539 | 0.000229001 | 0.000947367 |
| GO:0033628 | regulation of cell adhesion mediated by integrin | 26/7539 | 0.000229001 | 0.000947367 |
| GO:1902895 | positive regulation of pri-miRNA transcription by RNA polymerase II | 26/7539 | 0.000229001 | 0.000947367 |
| GO:0071383 | cellular response to steroid hormone stimulus | 73/7539 | 0.00022984 | 0.000950217 |
| GO:0001963 | synaptic transmission, dopaminergic | 20/7539 | 0.000232821 | 0.0009569 |

| | | | | |
|---|---|---|---|---|
| GO:0002026 | regulation of the force of heart contraction | 20/7539 | 0.000232821 | 0.0009569 |
| GO:0042634 | regulation of hair cycle | 20/7539 | 0.000232821 | 0.0009569 |
| GO:0044319 | wound healing, spreading of cells | 20/7539 | 0.000232821 | 0.0009569 |
| GO:0048535 | lymph node development | 20/7539 | 0.000232821 | 0.0009569 |
| GO:0055075 | potassium ion homeostasis | 20/7539 | 0.000232821 | 0.0009569 |
| GO:0061036 | positive regulation of cartilage development | 20/7539 | 0.000232821 | 0.0009569 |
| GO:0086010 | membrane depolarization during action potential | 20/7539 | 0.000232821 | 0.0009569 |
| GO:0090505 | epiboly involved in wound healing | 20/7539 | 0.000232821 | 0.0009569 |
| GO:0002690 | positive regulation of leukocyte chemotaxis | 48/7539 | 0.000236721 | 0.000972298 |
| GO:0032984 | protein-containing complex disassembly | 94/7539 | 0.000236967 | 0.000972675 |
| GO:0051781 | positive regulation of cell division | 41/7539 | 0.000239084 | 0.000974475 |
| GO:0021854 | hypothalamus development | 13/7539 | 0.000239731 | 0.000974475 |
| GO:0035313 | wound healing, spreading of epidermal cells | 13/7539 | 0.000239731 | 0.000974475 |
| GO:0036035 | osteoclast development | 13/7539 | 0.000239731 | 0.000974475 |
| GO:0051589 | negative regulation of neurotransmitter transport | 13/7539 | 0.000239731 | 0.000974475 |
| GO:0051895 | negative regulation of focal adhesion assembly | 13/7539 | 0.000239731 | 0.000974475 |
| GO:0055012 | ventricular cardiac muscle cell differentiation | 13/7539 | 0.000239731 | 0.000974475 |
| GO:0072010 | glomerular epithelium development | 13/7539 | 0.000239731 | 0.000974475 |

| | | | | |
|---|---|---|---|---|
| GO:0090493 | catecholamine uptake | 13/7539 | 0.000239731 | 0.000974475 |
| GO:0097152 | mesenchymal cell apoptotic process | 13/7539 | 0.000239731 | 0.000974475 |
| GO:0098814 | spontaneous synaptic transmission | 13/7539 | 0.000239731 | 0.000974475 |
| GO:0150118 | negative regulation of cell-substrate junction organization | 13/7539 | 0.000239731 | 0.000974475 |
| GO:1901160 | primary amino compound metabolic process | 13/7539 | 0.000239731 | 0.000974475 |
| GO:2000252 | negative regulation of feeding behavior | 13/7539 | 0.000239731 | 0.000974475 |
| GO:2000369 | regulation of clathrin-dependent endocytosis | 13/7539 | 0.000239731 | 0.000974475 |
| GO:0045732 | positive regulation of protein catabolic process | 97/7539 | 0.000239875 | 0.000974475 |
| GO:0071621 | granulocyte chemotaxis | 60/7539 | 0.000241641 | 0.000981018 |
| GO:0051100 | negative regulation of binding | 76/7539 | 0.000243312 | 0.000987166 |
| GO:0003085 | negative regulation of systemic arterial blood pressure | 19/7539 | 0.000248347 | 0.00100501 |
| GO:0045761 | regulation of adenylate cyclase activity | 19/7539 | 0.000248347 | 0.00100501 |
| GO:0061311 | cell surface receptor signaling pathway involved in heart development | 19/7539 | 0.000248347 | 0.00100501 |
| GO:0099622 | cardiac muscle cell membrane repolarization | 19/7539 | 0.000248347 | 0.00100501 |
| GO:0046530 | photoreceptor cell differentiation | 43/7539 | 0.000253496 | 0.00102519 |
| GO:0015698 | inorganic anion transport | 67/7539 | 0.000254116 | 0.00102704 |
| GO:0048806 | genitalia development | 29/7539 | 0.0002569 | 0.001037628 |
| GO:0003338 | metanephros morphogenesis | 18/7539 | 0.000261701 | 0.001052305 |

| | | | | |
|---|---|---|---|---|
| GO:0033688 | regulation of osteoblast proliferation | 18/7539 | 0.000261701 | 0.001052305 |
| GO:0042044 | fluid transport | 18/7539 | 0.000261701 | 0.001052305 |
| GO:0060045 | positive regulation of cardiac muscle cell proliferation | 18/7539 | 0.000261701 | 0.001052305 |
| GO:0061437 | renal system vasculature development | 18/7539 | 0.000261701 | 0.001052305 |
| GO:0061440 | kidney vasculature development | 18/7539 | 0.000261701 | 0.001052305 |
| GO:1905809 | negative regulation of synapse organization | 18/7539 | 0.000261701 | 0.001052305 |
| GO:0014061 | regulation of norepinephrine secretion | 14/7539 | 0.000262644 | 0.001053411 |
| GO:0048243 | norepinephrine secretion | 14/7539 | 0.000262644 | 0.001053411 |
| GO:0051349 | positive regulation of lyase activity | 14/7539 | 0.000262644 | 0.001053411 |
| GO:1905276 | regulation of epithelial tube formation | 14/7539 | 0.000262644 | 0.001053411 |
| GO:0000082 | G1/S transition of mitotic cell cycle | 82/7539 | 0.000264995 | 0.001062169 |
| GO:2000514 | regulation of CD4-positive, alpha-beta T cell activation | 35/7539 | 0.000266271 | 0.001066605 |
| GO:0001754 | eye photoreceptor cell differentiation | 32/7539 | 0.000268024 | 0.001070231 |
| GO:0010656 | negative regulation of muscle cell apoptotic process | 32/7539 | 0.000268024 | 0.001070231 |
| GO:0035315 | hair cell differentiation | 32/7539 | 0.000268024 | 0.001070231 |
| GO:0048016 | inositol phosphate-mediated signaling | 32/7539 | 0.000268024 | 0.001070231 |
| GO:0051703 | intraspecies interaction between organisms | 32/7539 | 0.000268024 | 0.001070231 |
| GO:0014048 | regulation of glutamate secretion | 17/7539 | 0.000271586 | 0.001076155 |

| | | | | |
|---|---|---|---|---|
| GO:0032878 | regulation of establishment or maintenance of cell polarity | 17/7539 | 0.000271586 | 0.001076155 |
| GO:0035024 | negative regulation of Rho protein signal transduction | 17/7539 | 0.000271586 | 0.001076155 |
| GO:0048266 | behavioral response to pain | 17/7539 | 0.000271586 | 0.001076155 |
| GO:0061213 | positive regulation of mesonephros development | 17/7539 | 0.000271586 | 0.001076155 |
| GO:0098743 | cell aggregation | 17/7539 | 0.000271586 | 0.001076155 |
| GO:0007270 | neuron-neuron synaptic transmission | Sep-39 | 0.000271894 | 0.001076155 |
| GO:0021612 | facial nerve structural organization | Sep-39 | 0.000271894 | 0.001076155 |
| GO:0050957 | equilibrioception | Sep-39 | 0.000271894 | 0.001076155 |
| GO:0051583 | dopamine uptake involved in synaptic transmission | Sep-39 | 0.000271894 | 0.001076155 |
| GO:0051934 | catecholamine uptake involved in synaptic transmission | Sep-39 | 0.000271894 | 0.001076155 |
| GO:0097119 | postsynaptic density protein 95 clustering | Sep-39 | 0.000271894 | 0.001076155 |
| GO:0098598 | learned vocalization behavior or vocal learning | Sep-39 | 0.000271894 | 0.001076155 |
| GO:1903011 | negative regulation of bone development | Sep-39 | 0.000271894 | 0.001076155 |
| GO:0033138 | positive regulation of peptidyl-serine phosphorylation | 61/7539 | 0.000273625 | 0.001082328 |
| GO:0051481 | negative regulation of cytosolic calcium ion concentration | 15/7539 | 0.000274129 | 0.001083641 |
| GO:0002313 | mature B cell differentiation involved in immune response | 16/7539 | 0.000276379 | 0.001087767 |
| GO:0003176 | aortic valve development | 16/7539 | 0.000276379 | 0.001087767 |
| GO:0034405 | response to fluid shear stress | 16/7539 | 0.000276379 | 0.001087767 |

| | | | | |
|---|---|---|---|---|
| GO:0036303 | lymph vessel morphogenesis | 16/7539 | 0.000276379 | 0.001087767 |
| GO:0098703 | calcium ion import across plasma membrane | 16/7539 | 0.000276379 | 0.001087767 |
| GO:0101023 | vascular endothelial cell proliferation | 16/7539 | 0.000276379 | 0.001087767 |
| GO:1905562 | regulation of vascular endothelial cell proliferation | 16/7539 | 0.000276379 | 0.001087767 |
| GO:0003073 | regulation of systemic arterial blood pressure | 59/7539 | 0.000278355 | 0.00109418 |
| GO:1990266 | neutrophil migration | 59/7539 | 0.000278355 | 0.00109418 |
| GO:0030431 | sleep | 24/7539 | 0.000287412 | 0.001126978 |
| GO:0038179 | neurotrophin signaling pathway | 24/7539 | 0.000287412 | 0.001126978 |
| GO:0086004 | regulation of cardiac muscle cell contraction | 24/7539 | 0.000287412 | 0.001126978 |
| GO:0099590 | neurotransmitter receptor internalization | 24/7539 | 0.000287412 | 0.001126978 |
| GO:0060113 | inner ear receptor cell differentiation | 42/7539 | 0.000292869 | 0.001147661 |
| GO:0071322 | cellular response to carbohydrate stimulus | 71/7539 | 0.000301961 | 0.001182555 |
| GO:0021799 | cerebral cortex radially oriented cell migration | 23/7539 | 0.000320465 | 0.001250367 |
| GO:0031114 | regulation of microtubule depolymerization | 23/7539 | 0.000320465 | 0.001250367 |
| GO:0035335 | peptidyl-tyrosine dephosphorylation | 23/7539 | 0.000320465 | 0.001250367 |
| GO:0045684 | positive regulation of epidermis development | 23/7539 | 0.000320465 | 0.001250367 |
| GO:0086065 | cell communication involved in cardiac conduction | 23/7539 | 0.000320465 | 0.001250367 |
| GO:2000036 | regulation of stem cell population maintenance | 23/7539 | 0.000320465 | 0.001250367 |

| | | | | |
|---|---|---|---|---|
| GO:0050868 | negative regulation of T cell activation | 58/7539 | 0.000320664 | 0.001250372 |
| GO:0051222 | positive regulation of protein transport | 143/7539 | 0.000327573 | 0.001276523 |
| GO:0042098 | T cell proliferation | 93/7539 | 0.000327787 | 0.001276568 |
| GO:0050974 | detection of mechanical stimulus involved in sensory perception | 27/7539 | 0.000331798 | 0.001291395 |
| GO:0038034 | signal transduction in absence of ligand | 41/7539 | 0.000338414 | 0.001314711 |
| GO:0097192 | extrinsic apoptotic signaling pathway in absence of ligand | 41/7539 | 0.000338414 | 0.001314711 |
| GO:1901862 | negative regulation of muscle tissue development | 41/7539 | 0.000338414 | 0.001314711 |
| GO:0046635 | positive regulation of alpha-beta T cell activation | 36/7539 | 0.000339808 | 0.001318506 |
| GO:0048641 | regulation of skeletal muscle tissue development | 36/7539 | 0.000339808 | 0.001318506 |
| GO:0036475 | neuron death in response to oxidative stress | 22/7539 | 0.000355924 | 0.001378496 |
| GO:0042745 | circadian sleep/wake cycle | 22/7539 | 0.000355924 | 0.001378496 |
| GO:2000191 | regulation of fatty acid transport | 22/7539 | 0.000355924 | 0.001378496 |
| GO:0009314 | response to radiation | 162/7539 | 0.000361877 | 0.001400691 |
| GO:1903825 | organic acid transmembrane transport | 59/7539 | 0.000362627 | 0.001402734 |
| GO:0045786 | negative regulation of cell cycle | 180/7539 | 0.000364914 | 0.001410717 |
| GO:1903793 | positive regulation of anion transport | 38/7539 | 0.0003680 8 | 0.0014219 1 |
| GO:0009743 | response to carbohydrate | 89/7539 | 0.0003682 6 | 0.0014219 1 |
| GO:0071300 | cellular response to retinoic acid | 26/7539 | 0.0003763 2 | 0.001452145 |

| | | | | |
|---|---|---|---|---|
| GO:0061180 | mammary gland epithelium development | 40/7539 | 0.000391109 | 0.001508292 |
| GO:0032623 | interleukin-2 production | 35/7539 | 0.000391888 | 0.001509449 |
| GO:2000736 | regulation of stem cell differentiation | 35/7539 | 0.000391888 | 0.001509449 |
| GO:0021591 | ventricular system development | 21/7539 | 0.000393477 | 0.001513724 |
| GO:0043537 | negative regulation of blood vessel endothelial cell migration | 21/7539 | 0.000393477 | 0.001513724 |
| GO:0030042 | actin filament depolymerization | 32/7539 | 0.000405307 | 0.001555494 |
| GO:0010906 | regulation of glucose metabolic process | 60/7539 | 0.000407825 | 0.001555494 |
| GO:0003157 | endocardium development | Oct-39 | 0.000408277 | 0.001555494 |
| GO:0003207 | cardiac chamber formation | Oct-39 | 0.000408277 | 0.001555494 |
| GO:0032341 | aldosterone metabolic process | Oct-39 | 0.000408277 | 0.001555494 |
| GO:0035933 | glucocorticoid secretion | Oct-39 | 0.000408277 | 0.001555494 |
| GO:0043587 | tongue morphogenesis | Oct-39 | 0.000408277 | 0.001555494 |
| GO:0044557 | relaxation of smooth muscle | Oct-39 | 0.000408277 | 0.001555494 |
| GO:0060073 | micturition | Oct-39 | 0.000408277 | 0.001555494 |
| GO:0060453 | regulation of gastric acid secretion | Oct-39 | 0.000408277 | 0.001555494 |
| GO:0060872 | semicircular canal development | Oct-39 | 0.000408277 | 0.001555494 |
| GO:0070444 | oligodendrocyte progenitor proliferation | Oct-39 | 0.000408277 | 0.001555494 |
| GO:0070445 | regulation of oligodendrocyte progenitor proliferation | Oct-39 | 0.000408277 | 0.001555494 |

| | | | | |
|---|---|---|---|---|
| GO:0072044 | collecting duct development | Oct-39 | 0.000408277 | 0.001555494 |
| GO:0086013 | membrane repolarization during cardiac muscle cell action potential | Oct-39 | 0.000408277 | 0.001555494 |
| GO:1902459 | positive regulation of stem cell population maintenance | Oct-39 | 0.000408277 | 0.001555494 |
| GO:2000058 | regulation of ubiquitin-dependent protein catabolic process | 72/7539 | 0.000417347 | 0.001589092 |
| GO:0044843 | cell cycle G1/S phase transition | 88/7539 | 0.000418453 | 0.001592342 |
| GO:0043407 | negative regulation of MAP kinase activity | 37/7539 | 0.000425134 | 0.00161679 |
| GO:0097484 | dendrite extension | 25/7539 | 0.000426128 | 0.001617648 |
| GO:1902742 | apoptotic process involved in development | 25/7539 | 0.000426128 | 0.001617648 |
| GO:2001258 | negative regulation of cation channel activity | 25/7539 | 0.000426128 | 0.001617648 |
| GO:0008589 | regulation of smoothened signaling pathway | 46/7539 | 0.000431144 | 0.001633967 |
| GO:0048384 | retinoic acid receptor signaling pathway | 20/7539 | 0.000432581 | 0.001633967 |
| GO:0048668 | collateral sprouting | 20/7539 | 0.000432581 | 0.001633967 |
| GO:0051491 | positive regulation of filopodium assembly | 20/7539 | 0.000432581 | 0.001633967 |
| GO:0060306 | regulation of membrane repolarization | 20/7539 | 0.000432581 | 0.001633967 |
| GO:0090504 | epiboly | 20/7539 | 0.000432581 | 0.001633967 |
| GO:1903859 | regulation of dendrite extension | 20/7539 | 0.000432581 | 0.001633967 |
| GO:2000515 | negative regulation of CD4-positive, alpha-beta T cell activation | 20/7539 | 0.000432581 | 0.001633967 |
| GO:0001818 | negative regulation of cytokine production | 110/7539 | 0.000432757 | 0.001633967 |

| | | | | |
|---|---|---|---|---|
| GO:0036473 | cell death in response to oxidative stress | 48/7539 | 0.000436093 | 0.00164558 |
| GO:0044264 | cellular polysaccharide metabolic process | 50/7539 | 0.000437584 | 0.001650218 |
| GO:0042445 | hormone metabolic process | 86/7539 | 0.000443325 | 0.001670874 |
| GO:0045670 | regulation of osteoclast differentiation | 39/7539 | 0.000452094 | 0.001702905 |
| GO:0034332 | adherens junction organization | 28/7539 | 0.000460298 | 0.001732773 |
| GO:0007585 | respiratory gaseous exchange by respiratory system | 31/7539 | 0.000466096 | 0.001753556 |
| GO:0036294 | cellular response to decreased oxygen levels | 59/7539 | 0.000469166 | 0.001764053 |
| GO:0002335 | mature B cell differentiation | 19/7539 | 0.000472378 | 0.00177191 |
| GO:0030947 | regulation of vascular endothelial growth factor receptor signaling pathway | 19/7539 | 0.000472378 | 0.00177191 |
| GO:0043951 | negative regulation of cAMP-mediated signaling | 19/7539 | 0.000472378 | 0.00177191 |
| GO:0050802 | circadian sleep/wake cycle, sleep | 19/7539 | 0.000472378 | 0.00177191 |
| GO:0046209 | nitric oxide metabolic process | 41/7539 | 0.000473 | 0.001773188 |
| GO:0002274 | myeloid leukocyte activation | 99/7539 | 0.000477973 | 0.001790768 |
| GO:0042542 | response to hydrogen peroxide | 57/7539 | 0.000480652 | 0.001799736 |
| GO:0006970 | response to osmotic stress | 43/7539 | 0.000488254 | 0.001827119 |
| GO:0120193 | tight junction organization | 45/7539 | 0.000498354 | 0.00186381 |
| GO:0010882 | regulation of cardiac muscle contraction by calcium ion signaling | Nov-39 | 0.000509542 | 0.001895545 |
| GO:0021903 | rostrocaudal neural tube patterning | Nov-39 | 0.000509542 | 0.001895545 |

| | | | | |
|---|---|---|---|---|
| GO:0021984 | adenohypophysis development | Nov-39 | 0.000509542 | 0.001895545 |
| GO:0035791 | platelet-derived growth factor receptor-beta signaling pathway | Nov-39 | 0.000509542 | 0.001895545 |
| GO:0048505 | regulation of timing of cell differentiation | Nov-39 | 0.000509542 | 0.001895545 |
| GO:0048755 | branching morphogenesis of a nerve | Nov-39 | 0.000509542 | 0.001895545 |
| GO:0071371 | cellular response to gonadotropin stimulus | Nov-39 | 0.000509542 | 0.001895545 |
| GO:0099550 | trans-synaptic signaling, modulating synaptic transmission | Nov-39 | 0.000509542 | 0.001895545 |
| GO:1901841 | regulation of high voltage-gated calcium channel activity | Nov-39 | 0.000509542 | 0.001895545 |
| GO:0010460 | positive regulation of heart rate | 18/7539 | 0.000511595 | 0.001897591 |
| GO:0045992 | negative regulation of embryonic development | 18/7539 | 0.000511595 | 0.001897591 |
| GO:0060037 | pharyngeal system development | 18/7539 | 0.000511595 | 0.001897591 |
| GO:0086005 | ventricular cardiac muscle cell action potential | 18/7539 | 0.000511595 | 0.001897591 |
| GO:1900745 | positive regulation of p38MAPK cascade | 18/7539 | 0.000511595 | 0.001897591 |
| GO:0030865 | cortical cytoskeleton organization | 33/7539 | 0.000521287 | 0.001929006 |
| GO:0031018 | endocrine pancreas development | 33/7539 | 0.000521287 | 0.001929006 |
| GO:0050854 | regulation of antigen receptor-mediated signaling pathway | 33/7539 | 0.000521287 | 0.001929006 |
| GO:0050982 | detection of mechanical stimulus | 33/7539 | 0.000521287 | 0.001929006 |
| GO:0007029 | endoplasmic reticulum organization | 38/7539 | 0.000522688 | 0.001933057 |
| GO:0070527 | platelet aggregation | 27/7539 | 0.000526108 | 0.001944566 |

| | | | | |
|---|---|---|---|---|
| GO:2000351 | regulation of endothelial cell apoptotic process | 30/7539 | 0.000535899 | 0.001979598 |
| GO:0090277 | positive regulation of peptide hormone secretion | 58/7539 | 0.000539743 | 0.001991466 |
| GO:1905039 | carboxylic acid transmembrane transport | 58/7539 | 0.000539743 | 0.001991466 |
| GO:0007292 | female gamete generation | 70/7539 | 0.00054625 | 0.002011667 |
| GO:0051650 | establishment of vesicle localization | 70/7539 | 0.00054625 | 0.002011667 |
| GO:0048635 | negative regulation of muscle organ development | 40/7539 | 0.000547032 | 0.002011667 |
| GO:0060191 | regulation of lipase activity | 40/7539 | 0.000547032 | 0.002011667 |
| GO:0140056 | organelle localization by membrane tethering | 40/7539 | 0.000547032 | 0.002011667 |
| GO:0021801 | cerebral cortex radial glia guided migration | 17/7539 | 0.000548405 | 0.002011667 |
| GO:0022030 | telencephalon glial cell migration | 17/7539 | 0.000548405 | 0.002011667 |
| GO:0032303 | regulation of icosanoid secretion | 17/7539 | 0.000548405 | 0.002011667 |
| GO:0072012 | glomerulus vasculature development | 17/7539 | 0.000548405 | 0.002011667 |
| GO:0090189 | regulation of branching involved in ureteric bud morphogenesis | 17/7539 | 0.000548405 | 0.002011667 |
| GO:0016050 | vesicle organization | 115/7539 | 0.000552889 | 0.002026938 |
| GO:0005976 | polysaccharide metabolic process | 54/7539 | 0.000565096 | 0.002069286 |
| GO:0043401 | steroid hormone mediated signaling pathway | 54/7539 | 0.000565096 | 0.002069286 |
| GO:0030239 | myofibril assembly | 35/7539 | 0.000567395 | 0.002075297 |
| GO:1903307 | positive regulation of regulated secretory pathway | 35/7539 | 0.000567395 | 0.002075297 |

| | | | | |
|---|---|---|---|---|
| GO:0007638 | mechanosensory behavior | Dec-39 | 0.000574038 | 0.002088704 |
| GO:0010454 | negative regulation of cell fate commitment | Dec-39 | 0.000574038 | 0.002088704 |
| GO:0035641 | locomotory exploration behavior | Dec-39 | 0.000574038 | 0.002088704 |
| GO:0048715 | negative regulation of oligodendrocyte differentiation | Dec-39 | 0.000574038 | 0.002088704 |
| GO:0071599 | otic vesicle development | Dec-39 | 0.000574038 | 0.002088704 |
| GO:0090051 | negative regulation of cell migration involved in sprouting angiogenesis | Dec-39 | 0.000574038 | 0.002088704 |
| GO:0097094 | craniofacial suture morphogenesis | Dec-39 | 0.000574038 | 0.002088704 |
| GO:1901386 | negative regulation of voltage-gated calcium channel activity | Dec-39 | 0.000574038 | 0.002088704 |
| GO:2001053 | regulation of mesenchymal cell apoptotic process | Dec-39 | 0.000574038 | 0.002088704 |
| GO:0030903 | notochord development | 16/7539 | 0.000580273 | 0.002107748 |
| GO:0048670 | regulation of collateral sprouting | 16/7539 | 0.000580273 | 0.002107748 |
| GO:1902656 | calcium ion import into cytosol | 16/7539 | 0.000580273 | 0.002107748 |
| GO:0009408 | response to heat | 46/7539 | 0.000582214 | 0.002113579 |
| GO:0045931 | positive regulation of mitotic cell cycle | 76/7539 | 0.000584409 | 0.002120328 |
| GO:0032881 | regulation of polysaccharide metabolic process | 26/7539 | 0.00060083 | 0.002178656 |
| GO:0072678 | T cell migration | 32/7539 | 0.000601256 | 0.00217895 |
| GO:0048857 | neural nucleus development | 15/7539 | 0.000603784 | 0.002179353 |
| GO:0051770 | positive regulation of nitric-oxide synthase biosynthetic process | 15/7539 | 0.000603784 | 0.002179353 |

| | | | | |
|---|---|---|---|---|
| GO:0060749 | mammary gland alveolus development | 15/7539 | 0.000603784 | 0.002179353 |
| GO:0061377 | mammary gland lobule development | 15/7539 | 0.000603784 | 0.002179353 |
| GO:0072215 | regulation of metanephros development | 15/7539 | 0.000603784 | 0.002179353 |
| GO:0072273 | metanephric nephron morphogenesis | 15/7539 | 0.000603784 | 0.002179353 |
| GO:2001026 | regulation of endothelial cell chemotaxis | 15/7539 | 0.000603784 | 0.002179353 |
| GO:0003214 | cardiac left ventricle morphogenesis | 13/7539 | 0.000606747 | 0.002182561 |
| GO:0009886 | post-embryonic animal morphogenesis | 13/7539 | 0.000606747 | 0.002182561 |
| GO:0035633 | maintenance of blood-brain barrier | 13/7539 | 0.000606747 | 0.002182561 |
| GO:0042487 | regulation of odontogenesis of dentin-containing tooth | 13/7539 | 0.000606747 | 0.002182561 |
| GO:0060841 | venous blood vessel development | 13/7539 | 0.000606747 | 0.002182561 |
| GO:0072077 | renal vesicle morphogenesis | 13/7539 | 0.000606747 | 0.002182561 |
| GO:0030866 | cortical actin cytoskeleton organization | 22/7539 | 0.000610984 | 0.002194051 |
| GO:0051647 | nucleus localization | 22/7539 | 0.000610984 | 0.002194051 |
| GO:0070873 | regulation of glycogen metabolic process | 22/7539 | 0.000610984 | 0.002194051 |
| GO:0099111 | microtubule-based transport | 85/7539 | 0.000613134 | 0.002195341 |
| GO:0006883 | cellular sodium ion homeostasis | 14/7539 | 0.000614473 | 0.002195341 |
| GO:0021535 | cell migration in hindbrain | 14/7539 | 0.000614473 | 0.002195341 |
| GO:0032095 | regulation of response to food | 14/7539 | 0.000614473 | 0.002195341 |

| | | | | |
|---|---|---|---|---|
| GO:0032252 | secretory granule localization | 14/7539 | 0.000614473 | 0.002195341 |
| GO:0046851 | negative regulation of bone remodeling | 14/7539 | 0.000614473 | 0.002195341 |
| GO:0071625 | vocalization behavior | 14/7539 | 0.000614473 | 0.002195341 |
| GO:1903909 | regulation of receptor clustering | 14/7539 | 0.000614473 | 0.002195341 |
| GO:2000114 | regulation of establishment of cell polarity | 14/7539 | 0.000614473 | 0.002195341 |
| GO:0045843 | negative regulation of striated muscle tissue development | 39/7539 | 0.00063279 | 0.002259505 |
| GO:0045621 | positive regulation of lymphocyte differentiation | 55/7539 | 0.000637194 | 0.002273944 |
| GO:0060389 | pathway-restricted SMAD protein phosphorylation | 34/7539 | 0.00065564 | 0.002338451 |
| GO:0035773 | insulin secretion involved in cellular response to glucose stimulus | 45/7539 | 0.000672893 | 0.00239863 |
| GO:1901615 | organic hydroxy compound metabolic process | 195/7539 | 0.00067546 | 0.00240642 |
| GO:0007026 | negative regulation of microtubule depolymerization | 21/7539 | 0.000685181 | 0.002432606 |
| GO:0009187 | cyclic nucleotide metabolic process | 21/7539 | 0.000685181 | 0.002432606 |
| GO:0016339 | calcium-dependent cell-cell adhesion via plasma membrane cell adhesion molecules | 21/7539 | 0.000685181 | 0.002432606 |
| GO:0034110 | regulation of homotypic cell-cell adhesion | 21/7539 | 0.000685181 | 0.002432606 |
| GO:1903203 | regulation of oxidative stress-induced neuron death | 21/7539 | 0.000685181 | 0.002432606 |
| GO:0032350 | regulation of hormone metabolic process | 25/7539 | 0.000685507 | 0.002432606 |
| GO:0046621 | negative regulation of organ growth | 25/7539 | 0.000685507 | 0.002432606 |
| GO:1901617 | organic hydroxy compound biosynthetic process | 96/7539 | 0.000689455 | 0.00244524 |

| | | | | |
|---|---|---|---|---|
| GO:0072577 | endothelial cell apoptotic process | 31/7539 | 0.000693513 | 0.00245825 2 |
| GO:0032891 | negative regulation of organic acid transport | 20/7539 | 0.000765591 | 0.00270917 9 |
| GO:1901099 | negative regulation of signal transduction in absence of ligand | 20/7539 | 0.000765591 | 0.00270917 9 |
| GO:2001240 | negative regulation of extrinsic apoptotic signaling pathway in absence of ligand | 20/7539 | 0.000765591 | 0.00270917 9 |
| GO:0006476 | protein deacetylation | 48/7539 | 0.000773656 | 0.00273618 6 |
| GO:0072348 | sulfur compound transport | 24/7539 | 0.000781241 | 0.00275992 |
| GO:0072583 | clathrin-dependent endocytosis | 24/7539 | 0.000781241 | 0.00275992 |
| GO:0051783 | regulation of nuclear division | 83/7539 | 0.000790057 | 0.00278950 3 |
| GO:0050873 | brown fat cell differentiation | 30/7539 | 0.000799941 | 0.00282282 |
| GO:0002763 | positive regulation of myeloid leukocyte differentiation | 35/7539 | 0.000808774 | 0.00285080 6 |
| GO:0046637 | regulation of alpha-beta T cell differentiation | 35/7539 | 0.000808774 | 0.00285080 6 |
| GO:0014888 | striated muscle adaptation | 27/7539 | 0.000813256 | 0.00285862 3 |
| GO:0033173 | calcineurin-NFAT signaling cascade | 27/7539 | 0.000813256 | 0.00285862 3 |
| GO:0042755 | eating behavior | 27/7539 | 0.000813256 | 0.00285862 3 |
| GO:0043550 | regulation of lipid kinase activity | 27/7539 | 0.000813256 | 0.00285862 3 |
| GO:0090102 | cochlea development | 27/7539 | 0.000813256 | 0.00285862 3 |
| GO:0001958 | endochondral ossification | 19/7539 | 0.000851678 | 0.00298536 4 |
| GO:0007097 | nuclear migration | 19/7539 | 0.000851678 | 0.00298536 4 |

| | | | | |
|---|---|---|---|---|
| GO:0033687 | osteoblast proliferation | 19/7539 | 0.000851678 | 0.002985364 |
| GO:0036075 | replacement ossification | 19/7539 | 0.000851678 | 0.002985364 |
| GO:0044062 | regulation of excretion | 19/7539 | 0.000851678 | 0.002985364 |
| GO:1903036 | positive regulation of response to wounding | 39/7539 | 0.000874471 | 0.00306356 |
| GO:0032640 | tumor necrosis factor production | 58/7539 | 0.000884043 | 0.003095376 |
| GO:0034105 | positive regulation of tissue remodeling | 23/7539 | 0.000889175 | 0.003109894 |
| GO:0043551 | regulation of phosphatidylinositol 3-kinase activity | 23/7539 | 0.000889175 | 0.003109894 |
| GO:0003333 | amino acid transmembrane transport | 41/7539 | 0.00089138 | 0.003112434 |
| GO:0032272 | negative regulation of protein polymerization | 41/7539 | 0.00089138 | 0.003112434 |
| GO:0061178 | regulation of insulin secretion involved in cellular response to glucose stimulus | 41/7539 | 0.00089138 | 0.003112434 |
| GO:0045913 | positive regulation of carbohydrate metabolic process | 43/7539 | 0.000899323 | 0.003138433 |
| GO:0019221 | cytokine-mediated signaling pathway | 148/7539 | 0.000903813 | 0.003152361 |
| GO:1904705 | regulation of vascular associated smooth muscle cell proliferation | 29/7539 | 0.000922707 | 0.003214707 |
| GO:1990874 | vascular associated smooth muscle cell proliferation | 29/7539 | 0.000922707 | 0.003214707 |
| GO:0048278 | vesicle docking | 34/7539 | 0.000935878 | 0.003258796 |
| GO:0007214 | gamma-aminobutyric acid signaling pathway | 18/7539 | 0.000942365 | 0.003275962 |
| GO:0045187 | regulation of circadian sleep/wake cycle, sleep | 18/7539 | 0.000942365 | 0.003275962 |
| GO:0048643 | positive regulation of skeletal muscle tissue development | 18/7539 | 0.000942365 | 0.003275962 |

| | | | | |
|---|---|---|---|---|
| GO:1904951 | positive regulation of establishment of protein localization | 146/7539 | 0.000972468 | 0.003376588 |
| GO:0035601 | protein deacylation | 52/7539 | 0.000972915 | 0.003376588 |
| GO:0098732 | macromolecule deacylation | 52/7539 | 0.000972915 | 0.003376588 |
| GO:0043473 | pigmentation | 50/7539 | 0.000996922 | 0.003458004 |
| GO:0010831 | positive regulation of myotube differentiation | 22/7539 | 0.001010448 | 0.003497236 |
| GO:0045601 | regulation of endothelial cell differentiation | 22/7539 | 0.001010448 | 0.003497236 |
| GO:0051968 | positive regulation of synaptic transmission, glutamatergic | 22/7539 | 0.001010448 | 0.003497236 |
| GO:0060674 | placenta blood vessel development | 22/7539 | 0.001010448 | 0.003497236 |
| GO:0010659 | cardiac muscle cell apoptotic process | 31/7539 | 0.001012598 | 0.003502758 |
| GO:0016048 | detection of temperature stimulus | 17/7539 | 0.001035801 | 0.003557679 |
| GO:0032528 | microvillus organization | 17/7539 | 0.001035801 | 0.003557679 |
| GO:0042753 | positive regulation of circadian rhythm | 17/7539 | 0.001035801 | 0.003557679 |
| GO:0045780 | positive regulation of bone resorption | 17/7539 | 0.001035801 | 0.003557679 |
| GO:0046852 | positive regulation of bone remodeling | 17/7539 | 0.001035801 | 0.003557679 |
| GO:0046885 | regulation of hormone biosynthetic process | 17/7539 | 0.001035801 | 0.003557679 |
| GO:0048745 | smooth muscle tissue development | 17/7539 | 0.001035801 | 0.003557679 |
| GO:0051767 | nitric-oxide synthase biosynthetic process | 17/7539 | 0.001035801 | 0.003557679 |
| GO:0051769 | regulation of nitric-oxide synthase biosynthetic process | 17/7539 | 0.001035801 | 0.003557679 |

| | | | | |
|---|---|---|---|---|
| GO:0060343 | trabecula formation | 17/7539 | 0.001035801 | 0.003557679 |
| GO:1900120 | regulation of receptor binding | 17/7539 | 0.001035801 | 0.003557679 |
| GO:1904994 | regulation of leukocyte adhesion to vascular endothelial cell | 17/7539 | 0.001035801 | 0.003557679 |
| GO:2000727 | positive regulation of cardiac muscle cell differentiation | 17/7539 | 0.001035801 | 0.003557679 |
| GO:2001057 | reactive nitrogen species metabolic process | 42/7539 | 0.001039991 | 0.003570127 |
| GO:0003139 | secondary heart field specification | Sep-39 | 0.0010626 | 0.00362605 |
| GO:0032342 | aldosterone biosynthetic process | Sep-39 | 0.0010626 | 0.00362605 |
| GO:0035881 | amacrine cell differentiation | Sep-39 | 0.0010626 | 0.00362605 |
| GO:0060013 | righting reflex | Sep-39 | 0.0010626 | 0.00362605 |
| GO:0060174 | limb bud formation | Sep-39 | 0.0010626 | 0.00362605 |
| GO:0060600 | dichotomous subdivision of an epithelial terminal unit | Sep-39 | 0.0010626 | 0.00362605 |
| GO:0060900 | embryonic camera-type eye formation | Sep-39 | 0.0010626 | 0.00362605 |
| GO:0072172 | mesonephric tubule formation | Sep-39 | 0.0010626 | 0.00362605 |
| GO:1902548 | negative regulation of cellular response to vascular endothelial growth factor stimulus | Sep-39 | 0.0010626 | 0.00362605 |
| GO:1904058 | positive regulation of sensory perception of pain | Sep-39 | 0.0010626 | 0.00362605 |
| GO:2000849 | regulation of glucocorticoid secretion | Sep-39 | 0.0010626 | 0.00362605 |
| GO:0002448 | mast cell mediated immunity | 28/7539 | 0.001064305 | 0.003627947 |
| GO:0030857 | negative regulation of epithelial cell differentiation | 28/7539 | 0.001064305 | 0.003627947 |

| | | | | |
|---|---|---|---|---|
| GO:0015749 | monosaccharide transmembrane transport | 53/7539 | 0.001087945 | 0.003706529 |
| GO:0006638 | neutral lipid metabolic process | 63/7539 | 0.001122017 | 0.003818487 |
| GO:0071453 | cellular response to oxygen levels | 63/7539 | 0.001122017 | 0.003818487 |
| GO:0071312 | cellular response to alkaloid | 16/7539 | 0.00112905 | 0.003838285 |
| GO:2000193 | positive regulation of fatty acid transport | 16/7539 | 0.00112905 | 0.003838285 |
| GO:0031069 | hair follicle morphogenesis | 21/7539 | 0.001146142 | 0.003885925 |
| GO:0035909 | aorta morphogenesis | 21/7539 | 0.001146142 | 0.003885925 |
| GO:0048009 | insulin-like growth factor receptor signaling pathway | 21/7539 | 0.001146142 | 0.003885925 |
| GO:0060351 | cartilage development involved in endochondral bone morphogenesis | 21/7539 | 0.001146142 | 0.003885925 |
| GO:2000171 | negative regulation of dendrite development | 21/7539 | 0.001146142 | 0.003885925 |
| GO:0031331 | positive regulation of cellular catabolic process | 157/7539 | 0.001160772 | 0.003933415 |
| GO:0099518 | vesicle cytoskeletal trafficking | 37/7539 | 0.00117199 | 0.003969298 |
| GO:0032675 | regulation of interleukin-6 production | 69/7539 | 0.001188926 | 0.004024499 |
| GO:0070830 | bicellular tight junction assembly | 41/7539 | 0.001202916 | 0.004069672 |
| GO:0030879 | mammary gland development | 72/7539 | 0.001209258 | 0.004088937 |
| GO:0003283 | atrial septum development | 15/7539 | 0.001217669 | 0.00409981 |
| GO:0010743 | regulation of macrophage derived foam cell differentiation | 15/7539 | 0.001217669 | 0.00409981 |
| GO:0021692 | cerebellar Purkinje cell layer morphogenesis | 15/7539 | 0.001217669 | 0.00409981 |

| | | | | |
|---|---|---|---|---|
| GO:0032098 | regulation of appetite | 15/7539 | 0.001217669 | 0.00409981 |
| GO:0051969 | regulation of transmission of nerve impulse | 15/7539 | 0.001217669 | 0.00409981 |
| GO:0090190 | positive regulation of branching involved in ureteric bud morphogenesis | 15/7539 | 0.001217669 | 0.00409981 |
| GO:0097062 | dendritic spine maintenance | 15/7539 | 0.001217669 | 0.00409981 |
| GO:0099149 | regulation of postsynaptic neurotransmitter receptor internalization | 15/7539 | 0.001217669 | 0.00409981 |
| GO:0002762 | negative regulation of myeloid leukocyte differentiation | 27/7539 | 0.001227599 | 0.00413104 |
| GO:0040018 | positive regulation of multicellular organism growth | 24/7539 | 0.001229805 | 0.004131857 |
| GO:0055026 | negative regulation of cardiac muscle tissue development | 24/7539 | 0.001229805 | 0.004131857 |
| GO:1903792 | negative regulation of anion transport | 24/7539 | 0.001229805 | 0.004131857 |
| GO:0008645 | hexose transmembrane transport | 52/7539 | 0.00125174 | 0.004180083 |
| GO:0008212 | mineralocorticoid metabolic process | Oct-39 | 0.001253542 | 0.004180083 |
| GO:0021670 | lateral ventricle development | Oct-39 | 0.001253542 | 0.004180083 |
| GO:0021859 | pyramidal neuron differentiation | Oct-39 | 0.001253542 | 0.004180083 |
| GO:0021877 | forebrain neuron fate commitment | Oct-39 | 0.001253542 | 0.004180083 |
| GO:0032486 | Rap protein signal transduction | Oct-39 | 0.001253542 | 0.004180083 |
| GO:0035810 | positive regulation of urine volume | Oct-39 | 0.001253542 | 0.004180083 |
| GO:0055119 | relaxation of cardiac muscle | Oct-39 | 0.001253542 | 0.004180083 |
| GO:0060452 | positive regulation of cardiac muscle contraction | Oct-39 | 0.001253542 | 0.004180083 |

| | | | | |
|---|---|---|---|---|
| GO:0061140 | lung secretory cell differentiation | Oct-39 | 0.001253542 | 0.004180083 |
| GO:0072160 | nephron tubule epithelial cell differentiation | Oct-39 | 0.001253542 | 0.004180083 |
| GO:1900025 | negative regulation of substrate adhesion-dependent cell spreading | Oct-39 | 0.001253542 | 0.004180083 |
| GO:2001054 | negative regulation of mesenchymal cell apoptotic process | Oct-39 | 0.001253542 | 0.004180083 |
| GO:0010658 | striated muscle cell apoptotic process | 32/7539 | 0.001254091 | 0.004180083 |
| GO:0060393 | regulation of pathway-restricted SMAD protein phosphorylation | 32/7539 | 0.001254091 | 0.004180083 |
| GO:0007249 | I-kappaB kinase/NF-kappaB signaling | 91/7539 | 0.001262115 | 0.004204609 |
| GO:0006639 | acylglycerol metabolic process | 62/7539 | 0.001283679 | 0.004274189 |
| GO:0050714 | positive regulation of protein secretion | 85/7539 | 0.00129414 | 0.004285601 |
| GO:0007190 | activation of adenylate cyclase activity | 14/7539 | 0.001295162 | 0.004285601 |
| GO:0032305 | positive regulation of icosanoid secretion | 14/7539 | 0.001295162 | 0.004285601 |
| GO:0032332 | positive regulation of chondrocyte differentiation | 14/7539 | 0.001295162 | 0.004285601 |
| GO:0033605 | positive regulation of catecholamine secretion | 14/7539 | 0.001295162 | 0.004285601 |
| GO:0051900 | regulation of mitochondrial depolarization | 14/7539 | 0.001295162 | 0.004285601 |
| GO:0060055 | angiogenesis involved in wound healing | 14/7539 | 0.001295162 | 0.004285601 |
| GO:0060456 | positive regulation of digestive system process | 14/7539 | 0.001295162 | 0.004285601 |
| GO:0072170 | metanephric tubule development | 14/7539 | 0.001295162 | 0.004285601 |
| GO:0090036 | regulation of protein kinase C signaling | 14/7539 | 0.001295162 | 0.004285601 |

| | | | | |
|---|---|---|---|---|
| GO:2001224 | positive regulation of neuron migration | 14/7539 | 0.001295162 | 0.004285601 |
| GO:0098868 | bone growth | 20/7539 | 0.001297182 | 0.004285601 |
| GO:0099563 | modification of synaptic structure | 20/7539 | 0.001297182 | 0.004285601 |
| GO:1900274 | regulation of phospholipase C activity | 20/7539 | 0.001297182 | 0.004285601 |
| GO:0032635 | interleukin-6 production | 73/7539 | 0.001297289 | 0.004285601 |
| GO:0042129 | regulation of T cell proliferation | 76/7539 | 0.001305994 | 0.004312101 |
| GO:0032680 | regulation of tumor necrosis factor production | 55/7539 | 0.001338834 | 0.004404289 |
| GO:0003188 | heart valve formation | Nov-39 | 0.001350475 | 0.004404289 |
| GO:0014049 | positive regulation of glutamate secretion | Nov-39 | 0.001350475 | 0.004404289 |
| GO:0019755 | one-carbon compound transport | Nov-39 | 0.001350475 | 0.004404289 |
| GO:0045760 | positive regulation of action potential | Nov-39 | 0.001350475 | 0.004404289 |
| GO:0061307 | cardiac neural crest cell differentiation involved in heart development | Nov-39 | 0.001350475 | 0.004404289 |
| GO:0061318 | renal filtration cell differentiation | Nov-39 | 0.001350475 | 0.004404289 |
| GO:0061430 | bone trabecula morphogenesis | Nov-39 | 0.001350475 | 0.004404289 |
| GO:0072112 | glomerular visceral epithelial cell differentiation | Nov-39 | 0.001350475 | 0.004404289 |
| GO:0072283 | metanephric renal vesicle morphogenesis | Nov-39 | 0.001350475 | 0.004404289 |
| GO:0072311 | glomerular epithelial cell differentiation | Nov-39 | 0.001350475 | 0.004404289 |
| GO:1903818 | positive regulation of voltage-gated potassium channel activity | Nov-39 | 0.001350475 | 0.004404289 |

| | | | | |
|---|---|---|---|---|
| GO:2000811 | negative regulation of anoikis | Nov-39 | 0.001350475 | 0.004404289 |
| GO:0021924 | cell proliferation in external granule layer | 13/7539 | 0.001352305 | 0.004404289 |
| GO:0021930 | cerebellar granule cell precursor proliferation | 13/7539 | 0.001352305 | 0.004404289 |
| GO:0032352 | positive regulation of hormone metabolic process | 13/7539 | 0.001352305 | 0.004404289 |
| GO:0045956 | positive regulation of calcium ion-dependent exocytosis | 13/7539 | 0.001352305 | 0.004404289 |
| GO:0048484 | enteric nervous system development | 13/7539 | 0.001352305 | 0.004404289 |
| GO:0060044 | negative regulation of cardiac muscle cell proliferation | 13/7539 | 0.001352305 | 0.004404289 |
| GO:0060712 | spongiotrophoblast layer development | 13/7539 | 0.001352305 | 0.004404289 |
| GO:0071605 | monocyte chemotactic protein-1 production | 13/7539 | 0.001352305 | 0.004404289 |
| GO:0071637 | regulation of monocyte chemotactic protein-1 production | 13/7539 | 0.001352305 | 0.004404289 |
| GO:0097623 | potassium ion export across plasma membrane | 13/7539 | 0.001352305 | 0.004404289 |
| GO:1900451 | positive regulation of glutamate receptor signaling pathway | 13/7539 | 0.001352305 | 0.004404289 |
| GO:2000251 | positive regulation of actin cytoskeleton reorganization | 13/7539 | 0.001352305 | 0.004404289 |
| GO:0033619 | membrane protein proteolysis | 29/7539 | 0.001354146 | 0.004404289 |
| GO:0035306 | positive regulation of dephosphorylation | 29/7539 | 0.001354146 | 0.004404289 |
| GO:0042531 | positive regulation of tyrosine phosphorylation of STAT protein | 29/7539 | 0.001354146 | 0.004404289 |
| GO:0045773 | positive regulation of axon extension | 29/7539 | 0.001354146 | 0.004404289 |
| GO:0061045 | negative regulation of wound healing | 36/7539 | 0.001357347 | 0.004412426 |

| | | | | |
|---|---|---|---|---|
| GO:0043900 | regulation of multi-organism process | 44/7539 | 0.001375523 | 0.004446802 |
| GO:0002024 | diet induced thermogenesis | Dec-39 | 0.001376374 | 0.004446802 |
| GO:0010752 | regulation of cGMP-mediated signaling | Dec-39 | 0.001376374 | 0.004446802 |
| GO:0032516 | positive regulation of phosphoprotein phosphatase activity | Dec-39 | 0.001376374 | 0.004446802 |
| GO:0032957 | inositol trisphosphate metabolic process | Dec-39 | 0.001376374 | 0.004446802 |
| GO:0045793 | positive regulation of cell size | Dec-39 | 0.001376374 | 0.004446802 |
| GO:0045989 | positive regulation of striated muscle contraction | Dec-39 | 0.001376374 | 0.004446802 |
| GO:0060413 | atrial septum morphogenesis | Dec-39 | 0.001376374 | 0.004446802 |
| GO:0060973 | cell migration involved in heart development | Dec-39 | 0.001376374 | 0.004446802 |
| GO:0072234 | metanephric nephron tubule development | Dec-39 | 0.001376374 | 0.004446802 |
| GO:1900119 | positive regulation of execution phase of apoptosis | Dec-39 | 0.001376374 | 0.004446802 |
| GO:1900452 | regulation of long-term synaptic depression | Dec-39 | 0.001376374 | 0.004446802 |
| GO:0035304 | regulation of protein dephosphorylation | 42/7539 | 0.00138884 | 0.004484782 |
| GO:0034219 | carbohydrate transmembrane transport | 53/7539 | 0.001390765 | 0.004488702 |
| GO:0002686 | negative regulation of leukocyte migration | 23/7539 | 0.001409931 | 0.004539469 |
| GO:0048730 | epidermis morphogenesis | 23/7539 | 0.001409931 | 0.004539469 |
| GO:0051281 | positive regulation of release of sequestered calcium ion into cytosol | 23/7539 | 0.001409931 | 0.004539469 |
| GO:1902991 | regulation of amyloid precursor protein catabolic process | 23/7539 | 0.001409931 | 0.004539469 |

| | | | | |
|---|---|---|---|---|
| GO:0032609 | interferon-gamma production | 58/7539 | 0.001410809 | 0.004539469 |
| GO:0071706 | tumor necrosis factor superfamily cytokine production | 58/7539 | 0.001410809 | 0.004539469 |
| GO:0090279 | regulation of calcium ion import | 26/7539 | 0.001415874 | 0.004553446 |
| GO:0031645 | negative regulation of nervous system process | 19/7539 | 0.001464204 | 0.004694525 |
| GO:0042749 | regulation of circadian sleep/wake cycle | 19/7539 | 0.001464204 | 0.004694525 |
| GO:0060561 | apoptotic process involved in morphogenesis | 19/7539 | 0.001464204 | 0.004694525 |
| GO:0060669 | embryonic placenta morphogenesis | 19/7539 | 0.001464204 | 0.004694525 |
| GO:1902473 | regulation of protein localization to synapse | 19/7539 | 0.001464204 | 0.004694525 |
| GO:2000352 | negative regulation of endothelial cell apoptotic process | 19/7539 | 0.001464204 | 0.004694525 |
| GO:0090068 | positive regulation of cell cycle process | 105/7539 | 0.001484661 | 0.004757697 |
| GO:0045921 | positive regulation of exocytosis | 49/7539 | 0.001486136 | 0.004760009 |
| GO:0034121 | regulation of toll-like receptor signaling pathway | 33/7539 | 0.001524167 | 0.004876871 |
| GO:0042246 | tissue regeneration | 33/7539 | 0.001524167 | 0.004876871 |
| GO:1902882 | regulation of response to oxidative stress | 47/7539 | 0.001527256 | 0.004884278 |
| GO:0007088 | regulation of mitotic nuclear division | 70/7539 | 0.001565957 | 0.005005512 |
| GO:0032388 | positive regulation of intracellular transport | 79/7539 | 0.001574749 | 0.005031068 |
| GO:0030203 | glycosaminoglycan metabolic process | 41/7539 | 0.001605813 | 0.005127718 |
| GO:0032885 | regulation of polysaccharide biosynthetic process | 22/7539 | 0.001615172 | 0.005149788 |

| | | | | |
|---|---|---|---|---|
| GO:0050931 | pigment cell differentiation | 22/7539 | 0.001615172 | 0.005149788 |
| GO:1900026 | positive regulation of substrate adhesion-dependent cell spreading | 22/7539 | 0.001615172 | 0.005149788 |
| GO:0032386 | regulation of intracellular transport | 132/7539 | 0.001625529 | 0.005180195 |
| GO:0010518 | positive regulation of phospholipase activity | 25/7539 | 0.001632895 | 0.00519842 |
| GO:1901985 | positive regulation of protein acetylation | 25/7539 | 0.001632895 | 0.00519842 |
| GO:0071326 | cellular response to monosaccharide stimulus | 65/7539 | 0.001634312 | 0.005200309 |
| GO:0001759 | organ induction | 18/7539 | 0.001647344 | 0.005228598 |
| GO:0005979 | regulation of glycogen biosynthetic process | 18/7539 | 0.001647344 | 0.005228598 |
| GO:0010962 | regulation of glucan biosynthetic process | 18/7539 | 0.001647344 | 0.005228598 |
| GO:0050951 | sensory perception of temperature stimulus | 18/7539 | 0.001647344 | 0.005228598 |
| GO:0051957 | positive regulation of amino acid transport | 18/7539 | 0.001647344 | 0.005228598 |
| GO:0032663 | regulation of interleukin-2 production | 30/7539 | 0.001682342 | 0.005336998 |
| GO:0032760 | positive regulation of tumor necrosis factor production | 32/7539 | 0.00176646 | 0.005601037 |
| GO:0031668 | cellular response to extracellular stimulus | 91/7539 | 0.001770183 | 0.005610022 |
| GO:0051606 | detection of stimulus | 102/7539 | 0.001786608 | 0.005659235 |
| GO:0006643 | membrane lipid metabolic process | 75/7539 | 0.001791325 | 0.005671331 |
| GO:0008089 | anterograde axonal transport | 27/7539 | 0.001812246 | 0.005731818 |
| GO:0043303 | mast cell degranulation | 27/7539 | 0.001812246 | 0.005731818 |

| | | | | |
|---|---|---|---|---|
| GO:0010827 | regulation of glucose transmembrane transport | 42/7539 | 0.001835791 | 0.005794673 |
| GO:0031960 | response to corticosteroid | 42/7539 | 0.001835791 | 0.005794673 |
| GO:0032526 | response to retinoic acid | 42/7539 | 0.001835791 | 0.005794673 |
| GO:0050672 | negative regulation of lymphocyte proliferation | 42/7539 | 0.001835791 | 0.005794673 |
| GO:1904659 | glucose transmembrane transport | 51/7539 | 0.001837429 | 0.005796946 |
| GO:0045737 | positive regulation of cyclin-dependent protein serine/threonine kinase activity | 17/7539 | 0.001845938 | 0.005811657 |
| GO:0051446 | positive regulation of meiotic cell cycle | 17/7539 | 0.001845938 | 0.005811657 |
| GO:1901020 | negative regulation of calcium ion transmembrane transporter activity | 17/7539 | 0.001845938 | 0.005811657 |
| GO:1903861 | positive regulation of dendrite extension | 17/7539 | 0.001845938 | 0.005811657 |
| GO:0001782 | B cell homeostasis | 21/7539 | 0.001848536 | 0.005811657 |
| GO:0009954 | proximal/distal pattern formation | 21/7539 | 0.001848536 | 0.005811657 |
| GO:0021680 | cerebellar Purkinje cell layer development | 21/7539 | 0.001848536 | 0.005811657 |
| GO:0072384 | organelle transport along microtubule | 40/7539 | 0.001857063 | 0.005835558 |
| GO:0005977 | glycogen metabolic process | 38/7539 | 0.001864021 | 0.005845786 |
| GO:0006073 | cellular glucan metabolic process | 38/7539 | 0.001864021 | 0.005845786 |
| GO:0008625 | extrinsic apoptotic signaling pathway via death domain receptors | 38/7539 | 0.001864021 | 0.005845786 |
| GO:0044042 | glucan metabolic process | 38/7539 | 0.001864021 | 0.005845786 |
| GO:0006024 | glycosaminoglycan biosynthetic process | 24/7539 | 0.001882962 | 0.005899326 |

| | | | | |
|---|---|---|---|---|
| GO:0070849 | response to epidermal growth factor | 24/7539 | 0.001882962 | 0.005899326 |
| GO:0043122 | regulation of I-kappaB kinase/NF-kappaB signaling | 79/7539 | 0.001888732 | 0.005914466 |
| GO:0045620 | negative regulation of lymphocyte differentiation | 29/7539 | 0.001949404 | 0.006095388 |
| GO:0051932 | synaptic transmission, GABAergic | 29/7539 | 0.001949404 | 0.006095388 |
| GO:0071675 | regulation of mononuclear cell migration | 29/7539 | 0.001949404 | 0.006095388 |
| GO:0032755 | positive regulation of interleukin-6 production | 47/7539 | 0.001971608 | 0.006161762 |
| GO:0042698 | ovulation cycle | 31/7539 | 0.002047963 | 0.006394036 |
| GO:0009713 | catechol-containing compound biosynthetic process | 16/7539 | 0.002058084 | 0.006394036 |
| GO:0010592 | positive regulation of lamellipodium assembly | 16/7539 | 0.002058084 | 0.006394036 |
| GO:0032104 | regulation of response to extracellular stimulus | 16/7539 | 0.002058084 | 0.006394036 |
| GO:0032107 | regulation of response to nutrient levels | 16/7539 | 0.002058084 | 0.006394036 |
| GO:0042423 | catecholamine biosynthetic process | 16/7539 | 0.002058084 | 0.006394036 |
| GO:0060065 | uterus development | 16/7539 | 0.002058084 | 0.006394036 |
| GO:0070168 | negative regulation of biomineral tissue development | 16/7539 | 0.002058084 | 0.006394036 |
| GO:0098901 | regulation of cardiac muscle cell action potential | 16/7539 | 0.002058084 | 0.006394036 |
| GO:0110150 | negative regulation of biomineralization | 16/7539 | 0.002058084 | 0.006394036 |
| GO:1901380 | negative regulation of potassium ion transmembrane transport | 16/7539 | 0.002058084 | 0.006394036 |
| GO:2000679 | positive regulation of transcription regulatory region DNA binding | 16/7539 | 0.002058084 | 0.006394036 |

| | | | | |
|---|---|---|---|---|
| GO:0040014 | regulation of multicellular organism growth | 43/7539 | 0.00208118 | 0.006462613 |
| GO:0021532 | neural tube patterning | 26/7539 | 0.002097095 | 0.0065056 3 |
| GO:0032941 | secretion by tissue | 26/7539 | 0.002097095 | 0.0065056 3 |
| GO:0032094 | response to food | 20/7539 | 0.002113183 | 0.006545891 |
| GO:0045879 | negative regulation of smoothened signaling pathway | 20/7539 | 0.002113183 | 0.006545891 |
| GO:0055083 | monovalent inorganic anion homeostasis | 20/7539 | 0.002113183 | 0.006545891 |
| GO:0120192 | tight junction assembly | 41/7539 | 0.00212134 | 0.006567934 |
| GO:1903555 | regulation of tumor necrosis factor superfamily cytokine production | 55/7539 | 0.002122789 | 0.006569199 |
| GO:0006140 | regulation of nucleotide metabolic process | 39/7539 | 0.002148085 | 0.006640969 |
| GO:0043525 | positive regulation of neuron apoptotic process | 39/7539 | 0.002148085 | 0.006640969 |
| GO:0000271 | polysaccharide biosynthetic process | 37/7539 | 0.00215811 | 0.006668697 |
| GO:0006509 | membrane protein ectodomain proteolysis | 23/7539 | 0.002170977 | 0.006701896 |
| GO:0016601 | Rac protein signal transduction | 23/7539 | 0.002170977 | 0.006701896 |
| GO:0042177 | negative regulation of protein catabolic process | 58/7539 | 0.00219632 | 0.006776818 |
| GO:0046834 | lipid phosphorylation | 28/7539 | 0.002259589 | 0.006968629 |
| GO:0008090 | retrograde axonal transport | 15/7539 | 0.002279994 | 0.007011011 |
| GO:0015732 | prostaglandin transport | 15/7539 | 0.002279994 | 0.007011011 |
| GO:0021889 | olfactory bulb interneuron differentiation | 15/7539 | 0.002279994 | 0.007011011 |

| | | | | |
|---|---|---|---|---|
| GO:00307 28 | ovulation | 15/7539 | 0.0022799 94 | 0.0070110 11 |
| GO:00971 50 | neuronal stem cell population maintenance | 15/7539 | 0.0022799 94 | 0.0070110 11 |
| GO:00996 23 | regulation of cardiac muscle cell membrane repolarization | 15/7539 | 0.0022799 94 | 0.0070110 11 |
| GO:00713 31 | cellular response to hexose stimulus | 64/7539 | 0.0022845 64 | 0.0070216 43 |
| GO:00160 51 | carbohydrate biosynthetic process | 83/7539 | 0.0023387 66 | 0.0071847 34 |
| GO:00706 64 | negative regulation of leukocyte proliferation | 44/7539 | 0.0023414 96 | 0.0071896 21 |
| GO:00329 45 | negative regulation of mononuclear cell proliferation | 42/7539 | 0.0024026 77 | 0.0073703 1 |
| GO:00716 74 | mononuclear cell migration | 42/7539 | 0.0024026 77 | 0.0073703 1 |
| GO:00001 86 | activation of MAPKK activity | 19/7539 | 0.0024123 22 | 0.0073855 42 |
| GO:00456 16 | regulation of keratinocyte differentiation | 19/7539 | 0.0024123 22 | 0.0073855 42 |
| GO:00988 84 | postsynaptic neurotransmitter receptor internalization | 19/7539 | 0.0024123 22 | 0.0073855 42 |
| GO:01402 39 | postsynaptic endocytosis | 19/7539 | 0.0024123 22 | 0.0073855 42 |
| GO:00316 67 | response to nutrient levels | 137/753 9 | 0.0024196 18 | 0.0074042 89 |
| GO:00439 02 | positive regulation of multi-organism process | 25/7539 | 0.0024271 88 | 0.0074130 81 |
| GO:00452 14 | sarcomere organization | 25/7539 | 0.0024271 88 | 0.0074130 81 |
| GO:00455 81 | negative regulation of T cell differentiation | 25/7539 | 0.0024271 88 | 0.0074130 81 |
| GO:01201 63 | negative regulation of cold-induced thermogenesis | 25/7539 | 0.0024271 88 | 0.0074130 81 |
| GO:19035 57 | positive regulation of tumor necrosis factor superfamily cytokine production | 32/7539 | 0.0024496 98 | 0.0074772 19 |

| | | | | |
|---|---|---|---|---|
| GO:0022406 | membrane docking | 40/7539 | 0.002451742 | 0.007477219 |
| GO:0051384 | response to glucocorticoid | 40/7539 | 0.002451742 | 0.007477219 |
| GO:0045428 | regulation of nitric oxide biosynthetic process | 34/7539 | 0.002489156 | 0.0075876 6 |
| GO:0031649 | heat generation | 14/7539 | 0.002505073 | 0.007610456 |
| GO:0048339 | paraxial mesoderm development | 14/7539 | 0.002505073 | 0.007610456 |
| GO:0051797 | regulation of hair follicle development | 14/7539 | 0.002505073 | 0.007610456 |
| GO:0060602 | branch elongation of an epithelium | 14/7539 | 0.002505073 | 0.007610456 |
| GO:0072111 | cell proliferation involved in kidney development | 14/7539 | 0.002505073 | 0.007610456 |
| GO:0099625 | ventricular cardiac muscle cell membrane repolarization | 14/7539 | 0.002505073 | 0.007610456 |
| GO:1903514 | release of sequestered calcium ion into cytosol by endoplasmic reticulum | 14/7539 | 0.002505073 | 0.007610456 |
| GO:0001776 | leukocyte homeostasis | 52/7539 | 0.002551893 | 0.007748965 |
| GO:0071333 | cellular response to glucose stimulus | 63/7539 | 0.002601626 | 0.007896185 |
| GO:0007599 | hemostasis | 72/7539 | 0.002606866 | 0.007908288 |
| GO:0070665 | positive regulation of leukocyte proliferation | 66/7539 | 0.002618429 | 0.007939549 |
| GO:0002279 | mast cell activation involved in immune response | 27/7539 | 0.002620022 | 0.007940567 |
| GO:0045930 | negative regulation of mitotic cell cycle | 97/7539 | 0.002694201 | 0.008095877 |
| GO:0006665 | sphingolipid metabolic process | 58/7539 | 0.002715979 | 0.008095877 |
| GO:0002315 | marginal zone B cell differentiation | Aug-39 | 0.002720131 | 0.008095877 |

| | | | | |
|---|---|---|---|---|
| GO:0003211 | cardiac ventricle formation | Aug-39 | 0.002720131 | 0.008095877 |
| GO:0010692 | regulation of alkaline phosphatase activity | Aug-39 | 0.002720131 | 0.008095877 |
| GO:0010745 | negative regulation of macrophage derived foam cell differentiation | Aug-39 | 0.002720131 | 0.008095877 |
| GO:0022027 | interkinetic nuclear migration | Aug-39 | 0.002720131 | 0.008095877 |
| GO:0032224 | positive regulation of synaptic transmission, cholinergic | Aug-39 | 0.002720131 | 0.008095877 |
| GO:0032354 | response to follicle-stimulating hormone | Aug-39 | 0.002720131 | 0.008095877 |
| GO:0032650 | regulation of interleukin-1 alpha production | Aug-39 | 0.002720131 | 0.008095877 |
| GO:0032836 | glomerular basement membrane development | Aug-39 | 0.002720131 | 0.008095877 |
| GO:0045607 | regulation of inner ear auditory receptor cell differentiation | Aug-39 | 0.002720131 | 0.008095877 |
| GO:0045631 | regulation of mechanoreceptor differentiation | Aug-39 | 0.002720131 | 0.008095877 |
| GO:0045762 | positive regulation of adenylate cyclase activity | Aug-39 | 0.002720131 | 0.008095877 |
| GO:0060297 | regulation of sarcomere organization | Aug-39 | 0.002720131 | 0.008095877 |
| GO:0060379 | cardiac muscle cell myoblast differentiation | Aug-39 | 0.002720131 | 0.008095877 |
| GO:0060484 | lung-associated mesenchyme development | Aug-39 | 0.002720131 | 0.008095877 |
| GO:0060513 | prostatic bud formation | Aug-39 | 0.002720131 | 0.008095877 |
| GO:0060710 | chorio-allantoic fusion | Aug-39 | 0.002720131 | 0.008095877 |
| GO:0072017 | distal tubule development | Aug-39 | 0.002720131 | 0.008095877 |
| GO:0072070 | loop of Henle development | Aug-39 | 0.002720131 | 0.008095877 |

| | | | | |
|---|---|---|---|---|
| GO:0072203 | cell proliferation involved in metanephros development | Aug-39 | 0.002720131 | 0.008095877 |
| GO:0098953 | receptor diffusion trapping | Aug-39 | 0.002720131 | 0.008095877 |
| GO:0098970 | postsynaptic neurotransmitter receptor diffusion trapping | Aug-39 | 0.002720131 | 0.008095877 |
| GO:0099541 | trans-synaptic signaling by lipid | Aug-39 | 0.002720131 | 0.008095877 |
| GO:0099542 | trans-synaptic signaling by endocannabinoid | Aug-39 | 0.002720131 | 0.008095877 |
| GO:0099628 | neurotransmitter receptor diffusion trapping | Aug-39 | 0.002720131 | 0.008095877 |
| GO:1902959 | regulation of aspartic-type endopeptidase activity involved in amyloid precursor protein catabolic process | Aug-39 | 0.002720131 | 0.008095877 |
| GO:1905962 | glutamatergic neuron differentiation | Aug-39 | 0.002720131 | 0.008095877 |
| GO:1990264 | peptidyl-tyrosine dephosphorylation involved in inactivation of protein kinase activity | Aug-39 | 0.002720131 | 0.008095877 |
| GO:2000543 | positive regulation of gastrulation | Aug-39 | 0.002720131 | 0.008095877 |
| GO:2000807 | regulation of synaptic vesicle clustering | Aug-39 | 0.002720131 | 0.008095877 |
| GO:2000980 | regulation of inner ear receptor cell differentiation | Aug-39 | 0.002720131 | 0.008095877 |
| GO:0001946 | lymphangiogenesis | 13/7539 | 0.002722564 | 0.008095877 |
| GO:0021534 | cell proliferation in hindbrain | 13/7539 | 0.002722564 | 0.008095877 |
| GO:0032310 | prostaglandin secretion | 13/7539 | 0.002722564 | 0.008095877 |
| GO:0035584 | calcium-mediated signaling using intracellular calcium source | 13/7539 | 0.002722564 | 0.008095877 |
| GO:0048172 | regulation of short-term neuronal synaptic plasticity | 13/7539 | 0.002722564 | 0.008095877 |
| GO:0060252 | positive regulation of glial cell proliferation | 13/7539 | 0.002722564 | 0.008095877 |

| | | | | |
|---|---|---|---|---|
| GO:0090030 | regulation of steroid hormone biosynthetic process | 13/7539 | 0.002722564 | 0.008095877 |
| GO:0043276 | anoikis | 18/7539 | 0.002749035 | 0.008163057 |
| GO:1904031 | positive regulation of cyclin-dependent protein kinase activity | 18/7539 | 0.002749035 | 0.008163057 |
| GO:1905523 | positive regulation of macrophage migration | 18/7539 | 0.002749035 | 0.008163057 |
| GO:0032964 | collagen biosynthetic process | 29/7539 | 0.002755734 | 0.008179101 |
| GO:1900180 | regulation of protein localization to nucleus | 61/7539 | 0.002761304 | 0.008191782 |
| GO:0044728 | DNA methylation or demethylation | 41/7539 | 0.00277422 | 0.008226233 |
| GO:0061756 | leukocyte adhesion to vascular endothelial cell | 24/7539 | 0.002809788 | 0.008327789 |
| GO:0032228 | regulation of synaptic transmission, GABAergic | 21/7539 | 0.002883821 | 0.00853519 |
| GO:0070528 | protein kinase C signaling | 21/7539 | 0.002883821 | 0.00853519 |
| GO:2000249 | regulation of actin cytoskeleton reorganization | 21/7539 | 0.002883821 | 0.00853519 |
| GO:0042116 | macrophage activation | 46/7539 | 0.002904729 | 0.008589063 |
| GO:0006833 | water transport | Dec-39 | 0.002915629 | 0.008589063 |
| GO:0010919 | regulation of inositol phosphate biosynthetic process | Dec-39 | 0.002915629 | 0.008589063 |
| GO:0021542 | dentate gyrus development | Dec-39 | 0.002915629 | 0.008589063 |
| GO:0021702 | cerebellar Purkinje cell differentiation | Dec-39 | 0.002915629 | 0.008589063 |
| GO:0032793 | positive regulation of CREB transcription factor activity | Dec-39 | 0.002915629 | 0.008589063 |
| GO:0045779 | negative regulation of bone resorption | Dec-39 | 0.002915629 | 0.008589063 |

| | | | | |
|---|---|---|---|---|
| GO:0060947 | cardiac vascular smooth muscle cell differentiation | Dec-39 | 0.002915629 | 0.008589063 |
| GO:0071696 | ectodermal placode development | Dec-39 | 0.002915629 | 0.008589063 |
| GO:1901881 | positive regulation of protein depolymerization | Dec-39 | 0.002915629 | 0.008589063 |
| GO:0007596 | blood coagulation | 71/7539 | 0.002952306 | 0.008693053 |
| GO:0010821 | regulation of mitochondrion organization | 62/7539 | 0.002961974 | 0.008713393 |
| GO:0046488 | phosphatidylinositol metabolic process | 62/7539 | 0.002961974 | 0.008713393 |
| GO:0048285 | organelle fission | 181/7539 | 0.00300882 | 0.00882236 |
| GO:0003129 | heart induction | Sep-39 | 0.003025568 | 0.00882236 |
| GO:0003337 | mesenchymal to epithelial transition involved in metanephros morphogenesis | Sep-39 | 0.003025568 | 0.00882236 |
| GO:0006705 | mineralocorticoid biosynthetic process | Sep-39 | 0.003025568 | 0.00882236 |
| GO:0010459 | negative regulation of heart rate | Sep-39 | 0.003025568 | 0.00882236 |
| GO:0014745 | negative regulation of muscle adaptation | Sep-39 | 0.003025568 | 0.00882236 |
| GO:0021860 | pyramidal neuron development | Sep-39 | 0.003025568 | 0.00882236 |
| GO:0030949 | positive regulation of vascular endothelial growth factor receptor signaling pathway | Sep-39 | 0.003025568 | 0.00882236 |
| GO:0043455 | regulation of secondary metabolic process | Sep-39 | 0.003025568 | 0.00882236 |
| GO:0048021 | regulation of melanin biosynthetic process | Sep-39 | 0.003025568 | 0.00882236 |
| GO:0051798 | positive regulation of hair follicle development | Sep-39 | 0.003025568 | 0.00882236 |
| GO:0060433 | bronchus development | Sep-39 | 0.003025568 | 0.00882236 |

| | | | | |
|---|---|---|---|---|
| GO:0060525 | prostate glandular acinus development | Sep-39 | 0.003025568 | 0.00882236 |
| GO:0072176 | nephric duct development | Sep-39 | 0.003025568 | 0.00882236 |
| GO:0097113 | AMPA glutamate receptor clustering | Sep-39 | 0.003025568 | 0.00882236 |
| GO:1900376 | regulation of secondary metabolite biosynthetic process | Sep-39 | 0.003025568 | 0.00882236 |
| GO:1903998 | regulation of eating behavior | Sep-39 | 0.003025568 | 0.00882236 |
| GO:2000302 | positive regulation of synaptic vesicle exocytosis | Sep-39 | 0.003025568 | 0.00882236 |
| GO:2001135 | regulation of endocytic recycling | Sep-39 | 0.003025568 | 0.00882236 |
| GO:0045646 | regulation of erythrocyte differentiation | 26/7539 | 0.003039048 | 0.008857577 |
| GO:0014733 | regulation of skeletal muscle adaptation | Nov-39 | 0.003058638 | 0.008865555 |
| GO:0030322 | stabilization of membrane potential | Nov-39 | 0.003058638 | 0.008865555 |
| GO:0031000 | response to caffeine | Nov-39 | 0.003058638 | 0.008865555 |
| GO:0035590 | purinergic nucleotide receptor signaling pathway | Nov-39 | 0.003058638 | 0.008865555 |
| GO:0060026 | convergent extension | Nov-39 | 0.003058638 | 0.008865555 |
| GO:0071731 | response to nitric oxide | Nov-39 | 0.003058638 | 0.008865555 |
| GO:0072673 | lamellipodium morphogenesis | Nov-39 | 0.003058638 | 0.008865555 |
| GO:0090136 | epithelial cell-cell adhesion | Nov-39 | 0.003058638 | 0.008865555 |
| GO:1901077 | regulation of relaxation of muscle | Nov-39 | 0.003058638 | 0.008865555 |
| GO:1901550 | regulation of endothelial cell development | Nov-39 | 0.003058638 | 0.008865555 |

| | | | | |
|---|---|---|---|---|
| GO:1903140 | regulation of establishment of endothelial barrier | Nov-39 | 0.003058638 | 0.008865555 |
| GO:2000647 | negative regulation of stem cell proliferation | Nov-39 | 0.003058638 | 0.008865555 |
| GO:0002224 | toll-like receptor signaling pathway | 49/7539 | 0.003067102 | 0.008881934 |
| GO:0032091 | negative regulation of protein binding | 49/7539 | 0.003067102 | 0.008881934 |
| GO:0032273 | positive regulation of protein polymerization | 57/7539 | 0.003101074 | 0.008958884 |
| GO:0014820 | tonic smooth muscle contraction | Oct-39 | 0.003113542 | 0.008958884 |
| GO:0019896 | axonal transport of mitochondrion | Oct-39 | 0.003113542 | 0.008958884 |
| GO:0021936 | regulation of cerebellar granule cell precursor proliferation | Oct-39 | 0.003113542 | 0.008958884 |
| GO:0032328 | alanine transport | Oct-39 | 0.003113542 | 0.008958884 |
| GO:0032536 | regulation of cell projection size | Oct-39 | 0.003113542 | 0.008958884 |
| GO:0034309 | primary alcohol biosynthetic process | Oct-39 | 0.003113542 | 0.008958884 |
| GO:0035815 | positive regulation of renal sodium excretion | Oct-39 | 0.003113542 | 0.008958884 |
| GO:0038065 | collagen-activated signaling pathway | Oct-39 | 0.003113542 | 0.008958884 |
| GO:0048672 | positive regulation of collateral sprouting | Oct-39 | 0.003113542 | 0.008958884 |
| GO:0061308 | cardiac neural crest cell development involved in heart development | Oct-39 | 0.003113542 | 0.008958884 |
| GO:0072224 | metanephric glomerulus development | Oct-39 | 0.003113542 | 0.008958884 |
| GO:0090179 | planar cell polarity pathway involved in neural tube closure | Oct-39 | 0.003113542 | 0.008958884 |
| GO:1905063 | regulation of vascular associated smooth muscle cell differentiation | Oct-39 | 0.003113542 | 0.008958884 |

| | | | | |
|---|---|---|---|---|
| GO:0050817 | coagulation | 72/7539 | 0.003124728 | 0.00898245 |
| GO:0071356 | cellular response to tumor necrosis factor | 72/7539 | 0.003124728 | 0.00898245 |
| GO:0060444 | branching involved in mammary gland duct morphogenesis | 17/7539 | 0.003126001 | 0.00898245 |
| GO:0030183 | B cell differentiation | 66/7539 | 0.003170252 | 0.009105461 |
| GO:0009636 | response to toxic substance | 52/7539 | 0.003191024 | 0.009160953 |
| GO:0010665 | regulation of cardiac muscle cell apoptotic process | 28/7539 | 0.003198566 | 0.00917426 |
| GO:0030834 | regulation of actin filament depolymerization | 28/7539 | 0.003198566 | 0.00917426 |
| GO:0006821 | chloride transport | 47/7539 | 0.003206479 | 0.009188606 |
| GO:0072676 | lymphocyte migration | 47/7539 | 0.003206479 | 0.009188606 |
| GO:0003341 | cilium movement | 76/7539 | 0.003247803 | 0.009297527 |
| GO:0003298 | physiological muscle hypertrophy | 23/7539 | 0.003253325 | 0.009297527 |
| GO:0003301 | physiological cardiac muscle hypertrophy | 23/7539 | 0.003253325 | 0.009297527 |
| GO:0045880 | positive regulation of smoothened signaling pathway | 23/7539 | 0.003253325 | 0.009297527 |
| GO:0061049 | cell growth involved in cardiac muscle cell development | 23/7539 | 0.003253325 | 0.009297527 |
| GO:1903053 | regulation of extracellular matrix organization | 23/7539 | 0.003253325 | 0.009297527 |
| GO:1900542 | regulation of purine nucleotide metabolic process | 38/7539 | 0.003276802 | 0.009360386 |
| GO:0090303 | positive regulation of wound healing | 30/7539 | 0.003298052 | 0.009404065 |
| GO:1905517 | macrophage migration | 30/7539 | 0.003298052 | 0.009404065 |

| | | | | |
|---|---|---|---|---|
| GO:2000401 | regulation of lymphocyte migration | 30/7539 | 0.003298052 | 0.009404065 |
| GO:2001238 | positive regulation of extrinsic apoptotic signaling pathway | 30/7539 | 0.003298052 | 0.009404065 |
| GO:0010573 | vascular endothelial growth factor production | 20/7539 | 0.003321985 | 0.00946376 |
| GO:0034142 | toll-like receptor 4 signaling pathway | 20/7539 | 0.003321985 | 0.00946376 |
| GO:0007260 | tyrosine phosphorylation of STAT protein | 36/7539 | 0.003329157 | 0.009475639 |
| GO:1903725 | regulation of phospholipid metabolic process | 36/7539 | 0.003329157 | 0.009475639 |
| GO:0006305 | DNA alkylation | 34/7539 | 0.003354905 | 0.009524828 |
| GO:0006306 | DNA methylation | 34/7539 | 0.003354905 | 0.009524828 |
| GO:0030888 | regulation of B cell proliferation | 34/7539 | 0.003354905 | 0.009524828 |
| GO:1990868 | response to chemokine | 34/7539 | 0.003354905 | 0.009524828 |
| GO:1990869 | cellular response to chemokine | 34/7539 | 0.003354905 | 0.009524828 |
| GO:0032963 | collagen metabolic process | 50/7539 | 0.003355491 | 0.009524828 |
| GO:0033077 | T cell differentiation in thymus | 43/7539 | 0.003472271 | 0.009847462 |
| GO:0045445 | myoblast differentiation | 43/7539 | 0.003472271 | 0.009847462 |
| GO:0001578 | microtubule bundle formation | 48/7539 | 0.003520833 | 0.0099807 |

**GO terms associated with 1189 Rep_early_depleted_vs_Hep genes**

| ID | Description | GeneRatio | pvalue | p.adjust |
|---|---|---|---|---|
| GO:0016054 | organic acid catabolic process | 79/913 | 4.26E-61 | 1.94E-57 |
| GO:0046395 | carboxylic acid catabolic process | 78/913 | 3.47E-60 | 7.91E-57 |
| GO:0044282 | small molecule catabolic process | 92/913 | 1.33E-59 | 2.02E-56 |
| GO:0006631 | fatty acid metabolic process | 99/913 | 1.41E-54 | 1.61E-51 |
| GO:0006520 | cellular amino acid metabolic process | 69/913 | 9.32E-44 | 8.51E-41 |
| GO:1901605 | alpha-amino acid metabolic process | 59/913 | 1.16E-42 | 8.83E-40 |
| GO:0072329 | monocarboxylic acid catabolic process | 44/913 | 1.24E-34 | 8.11E-32 |
| GO:0009063 | cellular amino acid catabolic process | 38/913 | 4.44E-32 | 2.53E-29 |
| GO:0046394 | carboxylic acid biosynthetic process | 61/913 | 5.34E-32 | 2.71E-29 |
| GO:0016053 | organic acid biosynthetic process | 61/913 | 6.53E-32 | 2.88E-29 |
| GO:0016042 | lipid catabolic process | 64/913 | 6.95E-32 | 2.88E-29 |
| GO:0009062 | fatty acid catabolic process | 37/913 | 1.31E-29 | 4.97E-27 |
| GO:0001676 | long-chain fatty acid metabolic process | 40/913 | 2.08E-29 | 7.32E-27 |
| GO:0006790 | sulfur compound metabolic process | 61/913 | 2.65E-29 | 8.63E-27 |
| GO:0044242 | cellular lipid catabolic process | 51/913 | 4.89E-29 | 1.49E-26 |
| GO:1901606 | alpha-amino acid catabolic process | 33/913 | 5.38E-29 | 1.54E-26 |

| | | | | |
|---|---|---|---|---|
| GO:0033559 | unsaturated fatty acid metabolic process | 37/913 | 1.01E-24 | 2.70E-22 |
| GO:0008202 | steroid metabolic process | 56/913 | 2.20E-24 | 5.58E-22 |
| GO:0019395 | fatty acid oxidation | 34/913 | 4.58E-24 | 1.10E-21 |
| GO:0120254 | olefinic compound metabolic process | 39/913 | 1.89E-23 | 4.31E-21 |
| GO:0034440 | lipid oxidation | 34/913 | 5.54E-23 | 1.20E-20 |
| GO:0006635 | fatty acid beta-oxidation | 27/913 | 1.49E-21 | 3.09E-19 |
| GO:0019369 | arachidonic acid metabolic process | 27/913 | 3.31E-21 | 6.56E-19 |
| GO:0006690 | icosanoid metabolic process | 34/913 | 1.56E-20 | 2.97E-18 |
| GO:0030258 | lipid modification | 41/913 | 1.96E-20 | 3.58E-18 |
| GO:0033865 | nucleoside bisphosphate metabolic process | 30/913 | 5.00E-20 | 8.15E-18 |
| GO:0033875 | ribonucleoside bisphosphate metabolic process | 30/913 | 5.00E-20 | 8.15E-18 |
| GO:0034032 | purine nucleoside bisphosphate metabolic process | 30/913 | 5.00E-20 | 8.15E-18 |
| GO:0043648 | dicarboxylic acid metabolic process | 27/913 | 6.22E-20 | 9.78E-18 |
| GO:0042180 | cellular ketone metabolic process | 41/913 | 5.84E-19 | 8.88E-17 |
| GO:0006575 | cellular modified amino acid metabolic process | 37/913 | 5.88E-18 | 8.66E-16 |
| GO:0009410 | response to xenobiotic stimulus | 46/913 | 1.87E-17 | 2.66E-15 |
| GO:0006637 | acyl-CoA metabolic process | 25/913 | 2.18E-17 | 2.93E-15 |
| GO:0035383 | thioester metabolic process | 25/913 | 2.18E-17 | 2.93E-15 |

| | | | | |
|---|---|---|---|---|
| GO:0072330 | monocarboxylic acid biosynthetic process | 36/913 | 9.32E-17 | 1.21E-14 |
| GO:0006805 | xenobiotic metabolic process | 28/913 | 1.95E-16 | 2.47E-14 |
| GO:0072521 | purine-containing compound metabolic process | 48/913 | 1.29E-15 | 1.59E-13 |
| GO:0071466 | cellular response to xenobiotic stimulus | 32/913 | 2.16E-15 | 2.60E-13 |
| GO:0009117 | nucleotide metabolic process | 51/913 | 3.02E-15 | 3.53E-13 |
| GO:0006753 | nucleoside phosphate metabolic process | 51/913 | 6.19E-15 | 7.07E-13 |
| GO:0009259 | ribonucleotide metabolic process | 45/913 | 7.12E-15 | 7.92E-13 |
| GO:0009150 | purine ribonucleotide metabolic process | 43/913 | 2.13E-14 | 2.32E-12 |
| GO:0019693 | ribose phosphate metabolic process | 45/913 | 2.35E-14 | 2.49E-12 |
| GO:0009074 | aromatic amino acid family catabolic process | 11/913 | 1.15E-13 | 1.19E-11 |
| GO:0006639 | acylglycerol metabolic process | 27/913 | 1.46E-13 | 1.49E-11 |
| GO:0006638 | neutral lipid metabolic process | 27/913 | 2.06E-13 | 2.04E-11 |
| GO:0006163 | purine nucleotide metabolic process | 43/913 | 2.25E-13 | 2.18E-11 |
| GO:0006066 | alcohol metabolic process | 41/913 | 4.26E-13 | 4.05E-11 |
| GO:0009064 | glutamine family amino acid metabolic process | 18/913 | 5.95E-13 | 5.54E-11 |
| GO:0015711 | organic anion transport | 41/913 | 9.09E-13 | 8.30E-11 |
| GO:0042537 | benzene-containing compound metabolic process | 12/913 | 1.82E-12 | 1.63E-10 |
| GO:0008206 | bile acid metabolic process | 15/913 | 1.93E-12 | 1.69E-10 |

| | | | | |
|---|---|---|---|---|
| GO:0016125 | sterol metabolic process | 26/913 | 3.14E-12 | 2.70E-10 |
| GO:0051791 | medium-chain fatty acid metabolic process | 12/913 | 3.39E-12 | 2.87E-10 |
| GO:0009072 | aromatic amino acid family metabolic process | 13/913 | 3.63E-12 | 3.01E-10 |
| GO:0006694 | steroid biosynthetic process | 27/913 | 4.64E-12 | 3.78E-10 |
| GO:0008203 | cholesterol metabolic process | 24/913 | 3.30E-11 | 2.64E-09 |
| GO:1902652 | secondary alcohol metabolic process | 25/913 | 3.55E-11 | 2.79E-09 |
| GO:0043651 | linoleic acid metabolic process | 12/913 | 4.88E-11 | 3.75E-09 |
| GO:0006641 | triglyceride metabolic process | 21/913 | 4.93E-11 | 3.75E-09 |
| GO:0046942 | carboxylic acid transport | 34/913 | 5.66E-11 | 4.24E-09 |
| GO:0006820 | anion transport | 46/913 | 1.14E-10 | 8.36E-09 |
| GO:0006957 | complement activation, alternative pathway | 9/913 | 1.31E-10 | 9.46E-09 |
| GO:0006536 | glutamate metabolic process | 11/913 | 2.50E-10 | 1.78E-08 |
| GO:1901617 | organic hydroxy compound biosynthetic process | 29/913 | 4.26E-10 | 2.99E-08 |
| GO:0006749 | glutathione metabolic process | 17/913 | 5.41E-10 | 3.74E-08 |
| GO:0019373 | epoxygenase P450 pathway | 12/913 | 5.92E-10 | 4.03E-08 |
| GO:0006699 | bile acid biosynthetic process | 11/913 | 6.25E-10 | 4.19E-08 |
| GO:0008652 | cellular amino acid biosynthetic process | 15/913 | 6.74E-10 | 4.46E-08 |
| GO:0009069 | serine family amino acid metabolic process | 12/913 | 8.51E-10 | 5.55E-08 |

| | | | | |
|---|---|---|---|---|
| GO:0006869 | lipid transport | 38/913 | 1.08E-09 | 6.92E-08 |
| GO:0009636 | response to toxic substance | 20/913 | 1.70E-09 | 1.08E-07 |
| GO:1901607 | alpha-amino acid biosynthetic process | 14/913 | 2.02E-09 | 1.26E-07 |
| GO:0015849 | organic acid transport | 33/913 | 3.54E-09 | 2.19E-07 |
| GO:0010876 | lipid localization | 40/913 | 5.26E-09 | 3.20E-07 |
| GO:0046486 | glycerolipid metabolic process | 36/913 | 6.66E-09 | 4.00E-07 |
| GO:0019216 | regulation of lipid metabolic process | 36/913 | 7.14E-09 | 4.23E-07 |
| GO:0046890 | regulation of lipid biosynthetic process | 25/913 | 7.69E-09 | 4.50E-07 |
| GO:0034754 | cellular hormone metabolic process | 19/913 | 1.04E-08 | 5.98E-07 |
| GO:0042402 | cellular biogenic amine catabolic process | 9/913 | 1.06E-08 | 6.06E-07 |
| GO:0042445 | hormone metabolic process | 26/913 | 1.12E-08 | 6.25E-07 |
| GO:0062012 | regulation of small molecule metabolic process | 35/913 | 1.12E-08 | 6.25E-07 |
| GO:0015850 | organic hydroxy compound transport | 28/913 | 1.29E-08 | 7.11E-07 |
| GO:0035458 | cellular response to interferon-beta | 13/913 | 1.60E-08 | 8.68E-07 |
| GO:0009310 | amine catabolic process | 9/913 | 1.70E-08 | 9.11E-07 |
| GO:0043650 | dicarboxylic acid biosynthetic process | 7/913 | 2.16E-08 | 1.15E-06 |
| GO:1901361 | organic cyclic compound catabolic process | 40/913 | 2.83E-08 | 1.49E-06 |
| GO:0006730 | one-carbon metabolic process | 11/913 | 3.04E-08 | 1.57E-06 |

| | | | | |
|---|---|---|---|---|
| GO:0000096 | sulfur amino acid metabolic process | 10/913 | 3.33E-08 | 1.71E-06 |
| GO:1901568 | fatty acid derivative metabolic process | 11/913 | 4.03E-08 | 2.04E-06 |
| GO:0006568 | tryptophan metabolic process | 7/913 | 4.55E-08 | 2.26E-06 |
| GO:0006586 | indolalkylamine metabolic process | 7/913 | 4.55E-08 | 2.26E-06 |
| GO:0006576 | cellular biogenic amine metabolic process | 17/913 | 5.21E-08 | 2.56E-06 |
| GO:0000038 | very long-chain fatty acid metabolic process | 10/913 | 8.68E-08 | 4.12E-06 |
| GO:0035456 | response to interferon-beta | 13/913 | 8.78E-08 | 4.12E-06 |
| GO:0006633 | fatty acid biosynthetic process | 20/913 | 8.84E-08 | 4.12E-06 |
| GO:0009083 | branched-chain amino acid catabolic process | 8/913 | 8.85E-08 | 4.12E-06 |
| GO:0050667 | homocysteine metabolic process | 7/913 | 8.86E-08 | 4.12E-06 |
| GO:0009308 | amine metabolic process | 18/913 | 9.87E-08 | 4.55E-06 |
| GO:0042430 | indole-containing compound metabolic process | 9/913 | 1.24E-07 | 5.67E-06 |
| GO:1901616 | organic hydroxy compound catabolic process | 13/913 | 1.55E-07 | 7.02E-06 |
| GO:0046185 | aldehyde catabolic process | 7/913 | 1.61E-07 | 7.23E-06 |
| GO:0006695 | cholesterol biosynthetic process | 12/913 | 1.81E-07 | 7.96E-06 |
| GO:1902653 | secondary alcohol biosynthetic process | 12/913 | 1.81E-07 | 7.96E-06 |
| GO:0006544 | glycine metabolic process | 7/913 | 2.79E-07 | 1.21E-05 |
| GO:0044106 | cellular amine metabolic process | 17/913 | 2.84E-07 | 1.22E-05 |

| | | | | |
|---|---|---|---|---|
| GO:0006766 | vitamin metabolic process | 14/913 | 3.77E-07 | 1.61E-05 |
| GO:0097305 | response to alcohol | 24/913 | 3.90E-07 | 1.65E-05 |
| GO:1905039 | carboxylic acid transmembrane transport | 18/913 | 3.99E-07 | 1.67E-05 |
| GO:0033539 | fatty acid beta-oxidation using acyl-CoA dehydrogenase | 6/913 | 4.01E-07 | 1.67E-05 |
| GO:0015718 | monocarboxylic acid transport | 17/913 | 4.07E-07 | 1.67E-05 |
| GO:0010565 | regulation of cellular ketone metabolic process | 19/913 | 4.16E-07 | 1.69E-05 |
| GO:1903825 | organic acid transmembrane transport | 18/913 | 4.45E-07 | 1.80E-05 |
| GO:0019439 | aromatic compound catabolic process | 36/913 | 4.60E-07 | 1.83E-05 |
| GO:0009081 | branched-chain amino acid metabolic process | 8/913 | 4.62E-07 | 1.83E-05 |
| GO:0046165 | alcohol biosynthetic process | 18/913 | 5.52E-07 | 2.17E-05 |
| GO:0016126 | sterol biosynthetic process | 12/913 | 5.75E-07 | 2.24E-05 |
| GO:0036315 | cellular response to sterol | 8/913 | 6.60E-07 | 2.55E-05 |
| GO:0006956 | complement activation | 21/913 | 9.65E-07 | 3.70E-05 |
| GO:0006525 | arginine metabolic process | 7/913 | 1.13E-06 | 4.30E-05 |
| GO:0044272 | sulfur compound biosynthetic process | 14/913 | 1.35E-06 | 5.10E-05 |
| GO:0006091 | generation of precursor metabolites and energy | 34/913 | 1.61E-06 | 6.02E-05 |
| GO:0046503 | glycerolipid catabolic process | 12/913 | 1.87E-06 | 6.94E-05 |
| GO:0045834 | positive regulation of lipid metabolic process | 20/913 | 1.92E-06 | 7.08E-05 |

| | | | | |
|---|---|---|---|---|
| GO:0046459 | short-chain fatty acid metabolic process | 6/913 | 2.40E-06 | 8.78E-05 |
| GO:0002920 | regulation of humoral immune response | 10/913 | 2.60E-06 | 9.42E-05 |
| GO:0055088 | lipid homeostasis | 19/913 | 3.52E-06 | 0.000126612 |
| GO:0042182 | ketone catabolic process | 6/913 | 3.90E-06 | 0.000139037 |
| GO:0015721 | bile acid and bile salt transport | 8/913 | 4.07E-06 | 0.000142748 |
| GO:0036314 | response to sterol | 8/913 | 4.07E-06 | 0.000142748 |
| GO:1990845 | adaptive thermogenesis | 18/913 | 5.00E-06 | 0.000174254 |
| GO:0010875 | positive regulation of cholesterol efflux | 8/913 | 5.27E-06 | 0.000182253 |
| GO:0001562 | response to protozoan | 9/913 | 5.39E-06 | 0.000184972 |
| GO:0006081 | cellular aldehyde metabolic process | 11/913 | 5.71E-06 | 0.000194365 |
| GO:0062013 | positive regulation of small molecule metabolic process | 18/913 | 7.02E-06 | 0.000237427 |
| GO:0006721 | terpenoid metabolic process | 11/913 | 7.73E-06 | 0.000259556 |
| GO:0046461 | neutral lipid catabolic process | 9/913 | 8.10E-06 | 0.000267798 |
| GO:0046464 | acylglycerol catabolic process | 9/913 | 8.10E-06 | 0.000267798 |
| GO:0035337 | fatty-acyl-CoA metabolic process | 6/913 | 9.13E-06 | 0.000297943 |
| GO:0044273 | sulfur compound catabolic process | 7/913 | 9.14E-06 | 0.000297943 |
| GO:0006084 | acetyl-CoA metabolic process | 8/913 | 1.08E-05 | 0.000350945 |
| GO:0035384 | thioester biosynthetic process | 7/913 | 1.22E-05 | 0.00038826 |

| | | | | |
|---|---|---|---|---|
| GO:0071616 | acyl-CoA biosynthetic process | 7/913 | 1.22E-05 | 0.00038826 |
| GO:0006558 | L-phenylalanine metabolic process | 5/913 | 1.24E-05 | 0.000391037 |
| GO:1902221 | erythrose 4-phosphate/phosphoenolpyruvate family amino acid metabolic process | 5/913 | 1.24E-05 | 0.000391037 |
| GO:0006959 | humoral immune response | 31/913 | 1.25E-05 | 0.000391037 |
| GO:0090205 | positive regulation of cholesterol metabolic process | 6/913 | 1.33E-05 | 0.000409678 |
| GO:1901570 | fatty acid derivative biosynthetic process | 6/913 | 1.33E-05 | 0.000409678 |
| GO:0033866 | nucleoside bisphosphate biosynthetic process | 8/913 | 1.36E-05 | 0.000409678 |
| GO:0034030 | ribonucleoside bisphosphate biosynthetic process | 8/913 | 1.36E-05 | 0.000409678 |
| GO:0034033 | purine nucleoside bisphosphate biosynthetic process | 8/913 | 1.36E-05 | 0.000409678 |
| GO:0090407 | organophosphate biosynthetic process | 34/913 | 1.39E-05 | 0.000417932 |
| GO:0006720 | isoprenoid metabolic process | 12/913 | 1.91E-05 | 0.000568621 |
| GO:0050873 | brown fat cell differentiation | 10/913 | 2.02E-05 | 0.000599963 |
| GO:0042832 | defense response to protozoan | 8/913 | 2.07E-05 | 0.000609822 |
| GO:0071397 | cellular response to cholesterol | 6/913 | 2.64E-05 | 0.000771191 |
| GO:0006835 | dicarboxylic acid transport | 12/913 | 2.70E-05 | 0.00078463 |
| GO:0046456 | icosanoid biosynthetic process | 9/913 | 2.85E-05 | 0.000823178 |
| GO:0009071 | serine family amino acid catabolic process | 5/913 | 3.29E-05 | 0.000938207 |
| GO:0009084 | glutamine family amino acid biosynthetic process | 5/913 | 3.29E-05 | 0.000938207 |

| | | | | |
|---|---|---|---|---|
| GO:0046889 | positive regulation of lipid biosynthetic process | 13/913 | 3.46E-05 | 0.000980715 |
| GO:0090181 | regulation of cholesterol metabolic process | 8/913 | 3.72E-05 | 0.001048121 |
| GO:0006692 | prostanoid metabolic process | 9/913 | 4.58E-05 | 0.001273477 |
| GO:0006693 | prostaglandin metabolic process | 9/913 | 4.58E-05 | 0.001273477 |
| GO:0055006 | cardiac cell development | 13/913 | 4.64E-05 | 0.001283969 |
| GO:0015918 | sterol transport | 11/913 | 4.88E-05 | 0.001333416 |
| GO:0034308 | primary alcohol metabolic process | 11/913 | 4.88E-05 | 0.001333416 |
| GO:0032782 | bile acid secretion | 5/913 | 4.98E-05 | 0.001353651 |
| GO:0006644 | phospholipid metabolic process | 27/913 | 5.11E-05 | 0.001379551 |
| GO:0010874 | regulation of cholesterol efflux | 8/913 | 5.34E-05 | 0.001426739 |
| GO:0050994 | regulation of lipid catabolic process | 10/913 | 5.35E-05 | 0.001426739 |
| GO:0009065 | glutamine family amino acid catabolic process | 6/913 | 6.33E-05 | 0.001666602 |
| GO:0019835 | cytolysis | 6/913 | 6.33E-05 | 0.001666602 |
| GO:0042632 | cholesterol homeostasis | 12/913 | 6.35E-05 | 0.001666602 |
| GO:0009112 | nucleobase metabolic process | 7/913 | 6.51E-05 | 0.001698118 |
| GO:0055092 | sterol homeostasis | 12/913 | 7.02E-05 | 0.001821613 |
| GO:0006570 | tyrosine metabolic process | 5/913 | 7.28E-05 | 0.001855987 |
| GO:0006577 | amino-acid betaine metabolic process | 5/913 | 7.28E-05 | 0.001855987 |

| | | | | |
|---|---|---|---|---|
| GO:0015936 | coenzyme A metabolic process | 5/913 | 7.28E-05 | 0.001855987 |
| GO:0045940 | positive regulation of steroid metabolic process | 8/913 | 7.50E-05 | 0.001902825 |
| GO:0106106 | cold-induced thermogenesis | 15/913 | 7.61E-05 | 0.001919271 |
| GO:0019218 | regulation of steroid metabolic process | 13/913 | 8.85E-05 | 0.002218452 |
| GO:0019217 | regulation of fatty acid metabolic process | 12/913 | 9.42E-05 | 0.002348313 |
| GO:0042908 | xenobiotic transport | 7/913 | 9.70E-05 | 0.002405842 |
| GO:0009070 | serine family amino acid biosynthetic process | 5/913 | 0.000103119 | 0.002530301 |
| GO:1902001 | fatty acid transmembrane transport | 5/913 | 0.000103119 | 0.002530301 |
| GO:0008209 | androgen metabolic process | 6/913 | 0.000105177 | 0.002553338 |
| GO:0030449 | regulation of complement activation | 6/913 | 0.000105177 | 0.002553338 |
| GO:0016101 | diterpenoid metabolic process | 9/913 | 0.000107378 | 0.002592978 |
| GO:0072593 | reactive oxygen species metabolic process | 20/913 | 0.000117022 | 0.002810996 |
| GO:0046434 | organophosphate catabolic process | 15/913 | 0.000119433 | 0.002853891 |
| GO:0030301 | cholesterol transport | 10/913 | 0.000124863 | 0.002968096 |
| GO:0070723 | response to cholesterol | 6/913 | 0.00013308 | 0.003147043 |
| GO:0045833 | negative regulation of lipid metabolic process | 12/913 | 0.00013669 | 0.003215732 |
| GO:0007031 | peroxisome organization | 7/913 | 0.000140541 | 0.003272589 |
| GO:0009066 | aspartate family amino acid metabolic process | 7/913 | 0.000140541 | 0.003272589 |

| | | | | |
|---|---|---|---|---|
| GO:0006103 | 2-oxoglutarate metabolic process | 5/913 | 0.000142281 | 0.003296292 |
| GO:0015740 | C4-dicarboxylate transport | 6/913 | 0.000166539 | 0.003838815 |
| GO:0042060 | wound healing | 26/913 | 0.00017387 | 0.003987646 |
| GO:0033344 | cholesterol efflux | 9/913 | 0.000178707 | 0.004064092 |
| GO:0000050 | urea cycle | 4/913 | 0.000180502 | 0.004064092 |
| GO:0051775 | response to redox state | 4/913 | 0.000180502 | 0.004064092 |
| GO:0019722 | calcium-mediated signaling | 17/913 | 0.000180765 | 0.004064092 |
| GO:0009165 | nucleotide biosynthetic process | 18/913 | 0.000184233 | 0.004121752 |
| GO:0090207 | regulation of triglyceride metabolic process | 8/913 | 0.000187507 | 0.004174543 |
| GO:0019432 | triglyceride biosynthetic process | 7/913 | 0.000198719 | 0.004381425 |
| GO:0033238 | regulation of cellular amine metabolic process | 7/913 | 0.000198719 | 0.004381425 |
| GO:0046460 | neutral lipid biosynthetic process | 8/913 | 0.000215508 | 0.004706118 |
| GO:0046463 | acylglycerol biosynthetic process | 8/913 | 0.000215508 | 0.004706118 |
| GO:0050850 | positive regulation of calcium-mediated signaling | 7/913 | 0.00023434 | 0.005072444 |
| GO:0120161 | regulation of cold-induced thermogenesis | 14/913 | 0.000234506 | 0.005072444 |
| GO:1901293 | nucleoside phosphate biosynthetic process | 18/913 | 0.000244735 | 0.005268728 |
| GO:0007159 | leukocyte cell-cell adhesion | 26/913 | 0.00025215 | 0.005359184 |
| GO:0043691 | reverse cholesterol transport | 5/913 | 0.000253634 | 0.005359184 |

| | | | | |
|---|---|---|---|---|
| GO:0044539 | long-chain fatty acid import into cell | 5/913 | 0.000253634 | 0.005359184 |
| GO:1904251 | regulation of bile acid metabolic process | 5/913 | 0.000253634 | 0.005359184 |
| GO:0006107 | oxaloacetate metabolic process | 4/913 | 0.000276532 | 0.005762986 |
| GO:0019627 | urea metabolic process | 4/913 | 0.000276532 | 0.005762986 |
| GO:0071941 | nitrogen cycle metabolic process | 4/913 | 0.000276532 | 0.005762986 |
| GO:0002455 | humoral immune response mediated by circulating immunoglobulin | 16/913 | 0.000285879 | 0.005930682 |
| GO:0001655 | urogenital system development | 26/913 | 0.000295986 | 0.006112572 |
| GO:0097306 | cellular response to alcohol | 14/913 | 0.000310302 | 0.006379358 |
| GO:0045444 | fat cell differentiation | 20/913 | 0.000315516 | 0.00645746 |
| GO:0006089 | lactate metabolic process | 5/913 | 0.000329392 | 0.006651961 |
| GO:0006541 | glutamine metabolic process | 5/913 | 0.000329392 | 0.006651961 |
| GO:0140354 | lipid import into cell | 5/913 | 0.000329392 | 0.006651961 |
| GO:0006958 | complement activation, classical pathway | 15/913 | 0.000331712 | 0.006669317 |
| GO:0001659 | temperature homeostasis | 16/913 | 0.000342511 | 0.006856233 |
| GO:0042558 | pteridine-containing compound metabolic process | 6/913 | 0.000371843 | 0.007410881 |
| GO:0019932 | second-messenger-mediated signaling | 22/913 | 0.000384983 | 0.00763941 |
| GO:0006534 | cysteine metabolic process | 4/913 | 0.000404407 | 0.007887662 |
| GO:0015911 | long-chain fatty acid import across plasma membrane | 4/913 | 0.000404407 | 0.007887662 |

| | | | | |
|---|---|---|---|---|
| GO:0034310 | primary alcohol catabolic process | 4/913 | 0.000404407 | 0.007887662 |
| GO:0052695 | cellular glucuronidation | 4/913 | 0.000404407 | 0.007887662 |
| GO:0098754 | detoxification | 8/913 | 0.000411726 | 0.007996244 |
| GO:0008207 | C21-steroid hormone metabolic process | 7/913 | 0.000431569 | 0.008310884 |
| GO:0070232 | regulation of T cell apoptotic process | 7/913 | 0.000431569 | 0.008310884 |
| GO:0010574 | regulation of vascular endothelial growth factor production | 6/913 | 0.000445483 | 0.008542797 |
| GO:1901652 | response to peptide | 27/913 | 0.000455759 | 0.008703279 |
| GO:0001523 | retinoid metabolic process | 8/913 | 0.000464508 | 0.008801396 |
| GO:0055013 | cardiac muscle cell development | 11/913 | 0.000464754 | 0.008801396 |
| GO:0034341 | response to interferon-gamma | 13/913 | 0.000492565 | 0.009289524 |
| GO:0006826 | iron ion transport | 8/913 | 0.000522645 | 0.009805435 |
| GO:0002253 | activation of immune response | 28/913 | 0.000524217 | 0.009805435 |
| GO:0010039 | response to iron ion | 5/913 | 0.00053082 | 0.009808359 |
| GO:0035634 | response to stilbenoid | 5/913 | 0.00053082 | 0.009808359 |
| GO:0071404 | cellular response to low-density lipoprotein particle stimulus | 5/913 | 0.00053082 | 0.009808359 |
| GO:0043434 | response to peptide hormone | 24/913 | 0.0005331 | 0.009810756 |

**GO terms associated with 3140 Rep_intermed_depleted_vs_Hep genes**

| ID | Description | GeneRatio | pvalue | p.adjust |
|---|---|---|---|---|
| GO:0044282 | small molecule catabolic process | 184/2440 | 9.90E-108 | 5.63E-104 |
| GO:0006631 | fatty acid metabolic process | 211/2440 | 2.09E-106 | 5.94E-103 |
| GO:0016054 | organic acid catabolic process | 143/2440 | 1.72E-97 | 3.26E-94 |
| GO:0046395 | carboxylic acid catabolic process | 141/2440 | 1.11E-95 | 1.58E-92 |
| GO:0046394 | carboxylic acid biosynthetic process | 142/2440 | 8.66E-73 | 9.85E-70 |
| GO:0016053 | organic acid biosynthetic process | 142/2440 | 1.53E-72 | 1.45E-69 |
| GO:0006520 | cellular amino acid metabolic process | 132/2440 | 1.18E-71 | 9.60E-69 |
| GO:0008202 | steroid metabolic process | 144/2440 | 4.59E-65 | 3.26E-62 |
| GO:0006091 | generation of precursor metabolites and energy | 161/2440 | 4.40E-62 | 2.78E-59 |
| GO:0006790 | sulfur compound metabolic process | 138/2440 | 1.51E-61 | 8.58E-59 |
| GO:1901605 | alpha-amino acid metabolic process | 101/2440 | 4.11E-59 | 2.13E-56 |
| GO:0072521 | purine-containing compound metabolic process | 143/2440 | 2.02E-54 | 9.60E-52 |
| GO:0072329 | monocarboxylic acid catabolic process | 79/2440 | 2.94E-54 | 1.29E-51 |
| GO:0006753 | nucleoside phosphate metabolic process | 154/2440 | 1.36E-53 | 5.54E-51 |
| GO:0016042 | lipid catabolic process | 129/2440 | 2.47E-53 | 9.37E-51 |
| GO:0009117 | nucleotide metabolic process | 152/2440 | 3.35E-53 | 1.19E-50 |

| | | | | |
|---|---|---|---|---|
| GO:0072330 | monocarboxylic acid biosynthetic process | 101/2440 | 1.89E-52 | 6.33E-50 |
| GO:0015980 | energy derivation by oxidation of organic compounds | 123/2440 | 3.40E-51 | 1.07E-48 |
| GO:0009259 | ribonucleotide metabolic process | 133/2440 | 9.37E-51 | 2.81E-48 |
| GO:0006066 | alcohol metabolic process | 129/2440 | 1.44E-50 | 3.90E-48 |
| GO:0009150 | purine ribonucleotide metabolic process | 129/2440 | 1.44E-50 | 3.90E-48 |
| GO:0045333 | cellular respiration | 102/2440 | 7.30E-50 | 1.89E-47 |
| GO:0019693 | ribose phosphate metabolic process | 134/2440 | 1.14E-49 | 2.82E-47 |
| GO:0044242 | cellular lipid catabolic process | 102/2440 | 1.23E-49 | 2.92E-47 |
| GO:0006163 | purine nucleotide metabolic process | 132/2440 | 3.73E-49 | 8.48E-47 |
| GO:0009063 | cellular amino acid catabolic process | 66/2440 | 1.39E-48 | 3.05E-46 |
| GO:0009060 | aerobic respiration | 86/2440 | 4.83E-48 | 1.02E-45 |
| GO:0016125 | sterol metabolic process | 82/2440 | 8.14E-47 | 1.65E-44 |
| GO:0009062 | fatty acid catabolic process | 66/2440 | 6.53E-46 | 1.28E-43 |
| GO:1902652 | secondary alcohol metabolic process | 82/2440 | 1.13E-45 | 2.14E-43 |
| GO:0008203 | cholesterol metabolic process | 78/2440 | 4.98E-45 | 9.15E-43 |
| GO:0033865 | nucleoside bisphosphate metabolic process | 66/2440 | 1.99E-42 | 3.34E-40 |
| GO:0033875 | ribonucleoside bisphosphate metabolic process | 66/2440 | 1.99E-42 | 3.34E-40 |
| GO:0034032 | purine nucleoside bisphosphate metabolic process | 66/2440 | 1.99E-42 | 3.34E-40 |

| | | | | |
|---|---|---|---|---|
| GO:0006637 | acyl-CoA metabolic process | 59/2440 | 3.70E-42 | 5.85E-40 |
| GO:0035383 | thioester metabolic process | 59/2440 | 3.70E-42 | 5.85E-40 |
| GO:0043648 | dicarboxylic acid metabolic process | 58/2440 | 1.09E-41 | 1.67E-39 |
| GO:0042180 | cellular ketone metabolic process | 95/2440 | 1.10E-40 | 1.64E-38 |
| GO:0046034 | ATP metabolic process | 99/2440 | 2.48E-39 | 3.61E-37 |
| GO:0010876 | lipid localization | 133/2440 | 3.66E-39 | 5.21E-37 |
| GO:0006805 | xenobiotic metabolic process | 67/2440 | 5.07E-39 | 7.04E-37 |
| GO:1901606 | alpha-amino acid catabolic process | 53/2440 | 1.80E-38 | 2.45E-36 |
| GO:0006635 | fatty acid beta-oxidation | 52/2440 | 5.22E-38 | 6.91E-36 |
| GO:0019395 | fatty acid oxidation | 63/2440 | 7.14E-38 | 9.24E-36 |
| GO:0034440 | lipid oxidation | 63/2440 | 1.70E-35 | 2.15E-33 |
| GO:0006869 | lipid transport | 117/2440 | 3.77E-35 | 4.67E-33 |
| GO:0019216 | regulation of lipid metabolic process | 115/2440 | 1.00E-34 | 1.22E-32 |
| GO:0120254 | olefinic compound metabolic process | 72/2440 | 1.57E-34 | 1.86E-32 |
| GO:0071466 | cellular response to xenobiotic stimulus | 76/2440 | 1.69E-34 | 1.97E-32 |
| GO:0006694 | steroid biosynthetic process | 73/2440 | 5.54E-34 | 6.31E-32 |
| GO:0046890 | regulation of lipid biosynthetic process | 80/2440 | 1.12E-33 | 1.26E-31 |
| GO:0006119 | oxidative phosphorylation | 60/2440 | 1.17E-33 | 1.28E-31 |

| | | | | |
|---|---|---|---|---|
| GO:0055088 | lipid homeostasis | 74/2440 | 1.77E-33 | 1.91E-31 |
| GO:0006633 | fatty acid biosynthetic process | 68/2440 | 1.10E-32 | 1.16E-30 |
| GO:0033559 | unsaturated fatty acid metabolic process | 63/2440 | 2.21E-32 | 2.29E-30 |
| GO:0062012 | regulation of small molecule metabolic process | 109/2440 | 6.24E-32 | 6.34E-30 |
| GO:0009410 | response to xenobiotic stimulus | 101/2440 | 6.53E-32 | 6.52E-30 |
| GO:0001676 | long-chain fatty acid metabolic process | 60/2440 | 1.48E-31 | 1.45E-29 |
| GO:0030258 | lipid modification | 79/2440 | 1.24E-30 | 1.20E-28 |
| GO:0090407 | organophosphate biosynthetic process | 125/2440 | 1.30E-30 | 1.23E-28 |
| GO:0016126 | sterol biosynthetic process | 41/2440 | 6.54E-29 | 6.11E-27 |
| GO:0006695 | cholesterol biosynthetic process | 39/2440 | 7.78E-29 | 7.03E-27 |
| GO:1902653 | secondary alcohol biosynthetic process | 39/2440 | 7.78E-29 | 7.03E-27 |
| GO:0006641 | triglyceride metabolic process | 53/2440 | 1.96E-27 | 1.74E-25 |
| GO:1901617 | organic hydroxy compound biosynthetic process | 78/2440 | 3.61E-27 | 3.16E-25 |
| GO:0022904 | respiratory electron transport chain | 48/2440 | 7.13E-27 | 6.15E-25 |
| GO:0045834 | positive regulation of lipid metabolic process | 68/2440 | 1.05E-26 | 8.96E-25 |
| GO:0006639 | acylglycerol metabolic process | 60/2440 | 1.44E-26 | 1.21E-24 |
| GO:0022900 | electron transport chain | 50/2440 | 2.30E-26 | 1.90E-24 |
| GO:0010565 | regulation of cellular ketone metabolic process | 61/2440 | 2.52E-26 | 2.05E-24 |

| | | | | |
|---|---|---|---|---|
| GO:0006575 | cellular modified amino acid metabolic process | 71/2440 | 2.72E-26 | 2.18E-24 |
| GO:0006638 | neutral lipid metabolic process | 60/2440 | 3.43E-26 | 2.71E-24 |
| GO:0019369 | arachidonic acid metabolic process | 43/2440 | 5.19E-26 | 4.04E-24 |
| GO:0006084 | acetyl-CoA metabolic process | 28/2440 | 3.07E-24 | 2.36E-22 |
| GO:0006690 | icosanoid metabolic process | 56/2440 | 5.30E-24 | 4.02E-22 |
| GO:0046889 | positive regulation of lipid biosynthetic process | 48/2440 | 1.06E-23 | 7.93E-22 |
| GO:0033108 | mitochondrial respiratory chain complex assembly | 44/2440 | 5.19E-23 | 3.84E-21 |
| GO:0015711 | organic anion transport | 91/2440 | 2.03E-22 | 1.48E-20 |
| GO:0042775 | mitochondrial ATP synthesis coupled electron transport | 37/2440 | 2.62E-22 | 1.89E-20 |
| GO:0019646 | aerobic electron transport chain | 33/2440 | 2.99E-22 | 2.12E-20 |
| GO:0042773 | ATP synthesis coupled electron transport | 37/2440 | 1.01E-21 | 7.11E-20 |
| GO:0046165 | alcohol biosynthetic process | 53/2440 | 1.20E-21 | 8.27E-20 |
| GO:0055092 | sterol homeostasis | 44/2440 | 1.21E-21 | 8.27E-20 |
| GO:0042632 | cholesterol homeostasis | 43/2440 | 6.44E-21 | 4.36E-19 |
| GO:0015918 | sterol transport | 39/2440 | 6.63E-21 | 4.44E-19 |
| GO:0046486 | glycerolipid metabolic process | 92/2440 | 1.55E-20 | 1.02E-18 |
| GO:1901293 | nucleoside phosphate biosynthetic process | 66/2440 | 2.16E-20 | 1.41E-18 |
| GO:0030301 | cholesterol transport | 37/2440 | 2.28E-20 | 1.47E-18 |

| | | | | |
|---|---|---|---|---|
| GO:0009165 | nucleotide biosynthetic process | 65/2440 | 2.66E-20 | 1.70E-18 |
| GO:0042445 | hormone metabolic process | 66/2440 | 3.69E-20 | 2.34E-18 |
| GO:0019217 | regulation of fatty acid metabolic process | 43/2440 | 4.28E-20 | 2.68E-18 |
| GO:0062013 | positive regulation of small molecule metabolic process | 56/2440 | 6.69E-20 | 4.14E-18 |
| GO:0015850 | organic hydroxy compound transport | 71/2440 | 8.47E-20 | 5.19E-18 |
| GO:0090181 | regulation of cholesterol metabolic process | 27/2440 | 1.07E-19 | 6.45E-18 |
| GO:0009152 | purine ribonucleotide biosynthetic process | 52/2440 | 2.71E-19 | 1.63E-17 |
| GO:0009636 | response to toxic substance | 46/2440 | 4.83E-19 | 2.86E-17 |
| GO:0072522 | purine-containing compound biosynthetic process | 56/2440 | 5.82E-19 | 3.41E-17 |
| GO:0046942 | carboxylic acid transport | 75/2440 | 6.77E-19 | 3.90E-17 |
| GO:0008206 | bile acid metabolic process | 27/2440 | 6.78E-19 | 3.90E-17 |
| GO:0009069 | serine family amino acid metabolic process | 25/2440 | 9.45E-19 | 5.38E-17 |
| GO:0010257 | NADH dehydrogenase complex assembly | 31/2440 | 1.30E-18 | 7.25E-17 |
| GO:0032981 | mitochondrial respiratory chain complex I assembly | 31/2440 | 1.30E-18 | 7.25E-17 |
| GO:0006720 | isoprenoid metabolic process | 38/2440 | 1.44E-18 | 7.93E-17 |
| GO:1905952 | regulation of lipid localization | 54/2440 | 1.45E-18 | 7.93E-17 |
| GO:0015849 | organic acid transport | 79/2440 | 2.40E-18 | 1.30E-16 |
| GO:0034754 | cellular hormone metabolic process | 45/2440 | 3.19E-18 | 1.71E-16 |

| | | | | |
|---|---|---|---|---|
| GO:0050994 | regulation of lipid catabolic process | 33/2440 | 4.86E-18 | 2.58E-16 |
| GO:0006164 | purine nucleotide biosynthetic process | 53/2440 | 1.02E-17 | 5.37E-16 |
| GO:0009260 | ribonucleotide biosynthetic process | 52/2440 | 1.20E-17 | 6.29E-16 |
| GO:0006820 | anion transport | 103/2440 | 1.23E-17 | 6.36E-16 |
| GO:0007031 | peroxisome organization | 24/2440 | 1.99E-17 | 1.02E-15 |
| GO:0090207 | regulation of triglyceride metabolic process | 28/2440 | 2.33E-17 | 1.19E-15 |
| GO:0006721 | terpenoid metabolic process | 32/2440 | 2.82E-17 | 1.42E-15 |
| GO:0046390 | ribose phosphate biosynthetic process | 53/2440 | 3.22E-17 | 1.61E-15 |
| GO:0006699 | bile acid biosynthetic process | 21/2440 | 6.19E-17 | 3.06E-15 |
| GO:0042537 | benzene-containing compound metabolic process | 19/2440 | 1.12E-16 | 5.48E-15 |
| GO:0008652 | cellular amino acid biosynthetic process | 30/2440 | 1.75E-16 | 8.52E-15 |
| GO:0006099 | tricarboxylic acid cycle | 22/2440 | 1.82E-16 | 8.79E-15 |
| GO:0006766 | vitamin metabolic process | 35/2440 | 1.98E-16 | 9.45E-15 |
| GO:0006090 | pyruvate metabolic process | 42/2440 | 2.66E-16 | 1.26E-14 |
| GO:0042304 | regulation of fatty acid biosynthetic process | 26/2440 | 5.05E-16 | 2.38E-14 |
| GO:0019218 | regulation of steroid metabolic process | 41/2440 | 8.02E-16 | 3.74E-14 |
| GO:1901361 | organic cyclic compound catabolic process | 96/2440 | 9.91E-16 | 4.59E-14 |
| GO:0098754 | detoxification | 28/2440 | 1.08E-15 | 4.94E-14 |

| | | | | |
|---|---|---|---|---|
| GO:0006644 | phospholipid metabolic process | 80/2440 | 2.54E-15 | 1.16E-13 |
| GO:0006749 | glutathione metabolic process | 33/2440 | 3.15E-15 | 1.42E-13 |
| GO:0043691 | reverse cholesterol transport | 16/2440 | 5.06E-15 | 2.27E-13 |
| GO:0015748 | organophosphate ester transport | 41/2440 | 8.03E-15 | 3.57E-13 |
| GO:0006081 | cellular aldehyde metabolic process | 29/2440 | 8.32E-15 | 3.67E-13 |
| GO:0045940 | positive regulation of steroid metabolic process | 24/2440 | 1.08E-14 | 4.73E-13 |
| GO:1901607 | alpha-amino acid biosynthetic process | 27/2440 | 1.10E-14 | 4.80E-13 |
| GO:0009083 | branched-chain amino acid catabolic process | 16/2440 | 2.33E-14 | 1.00E-12 |
| GO:0034976 | response to endoplasmic reticulum stress | 62/2440 | 2.81E-14 | 1.20E-12 |
| GO:0097006 | regulation of plasma lipoprotein particle levels | 29/2440 | 3.55E-14 | 1.51E-12 |
| GO:0033344 | cholesterol efflux | 28/2440 | 4.44E-14 | 1.87E-12 |
| GO:0005996 | monosaccharide metabolic process | 64/2440 | 4.54E-14 | 1.90E-12 |
| GO:0006536 | glutamate metabolic process | 18/2440 | 1.05E-13 | 4.36E-12 |
| GO:0033013 | tetrapyrrole metabolic process | 26/2440 | 1.06E-13 | 4.36E-12 |
| GO:0009081 | branched-chain amino acid metabolic process | 17/2440 | 1.09E-13 | 4.47E-12 |
| GO:0071827 | plasma lipoprotein particle organization | 22/2440 | 1.18E-13 | 4.80E-12 |
| GO:0009072 | aromatic amino acid family metabolic process | 19/2440 | 2.07E-13 | 8.36E-12 |
| GO:1990748 | cellular detoxification | 22/2440 | 2.28E-13 | 9.14E-12 |

| | | | | |
|---|---|---|---|---|
| GO:0009064 | glutamine family amino acid metabolic process | 27/2440 | 2.33E-13 | 9.26E-12 |
| GO:0019439 | aromatic compound catabolic process | 87/2440 | 2.71E-13 | 1.07E-11 |
| GO:0046434 | organophosphate catabolic process | 44/2440 | 3.01E-13 | 1.18E-11 |
| GO:0044272 | sulfur compound biosynthetic process | 33/2440 | 3.41E-13 | 1.33E-11 |
| GO:0006778 | porphyrin-containing compound metabolic process | 22/2440 | 4.29E-13 | 1.66E-11 |
| GO:0016101 | diterpenoid metabolic process | 26/2440 | 4.70E-13 | 1.81E-11 |
| GO:0051604 | protein maturation | 72/2440 | 5.22E-13 | 2.00E-11 |
| GO:0019373 | epoxygenase P450 pathway | 20/2440 | 6.45E-13 | 2.38E-11 |
| GO:0033866 | nucleoside bisphosphate biosynthetic process | 20/2440 | 6.45E-13 | 2.38E-11 |
| GO:0034030 | ribonucleoside bisphosphate biosynthetic process | 20/2440 | 6.45E-13 | 2.38E-11 |
| GO:0034033 | purine nucleoside bisphosphate biosynthetic process | 20/2440 | 6.45E-13 | 2.38E-11 |
| GO:0042168 | heme metabolic process | 20/2440 | 6.45E-13 | 2.38E-11 |
| GO:0006120 | mitochondrial electron transport, NADH to ubiquinone | 18/2440 | 6.58E-13 | 2.41E-11 |
| GO:0035384 | thioester biosynthetic process | 17/2440 | 8.35E-13 | 3.03E-11 |
| GO:0071616 | acyl-CoA biosynthetic process | 17/2440 | 8.35E-13 | 3.03E-11 |
| GO:0019318 | hexose metabolic process | 59/2440 | 1.14E-12 | 4.10E-11 |
| GO:0097237 | cellular response to toxic substance | 23/2440 | 1.31E-12 | 4.70E-11 |
| GO:0071825 | protein-lipid complex subunit organization | 22/2440 | 1.42E-12 | 5.06E-11 |

| | | | | |
|---|---|---|---|---|
| GO:0090208 | positive regulation of triglyceride metabolic process | 18/2440 | 1.51E-12 | 5.35E-11 |
| GO:0001523 | retinoid metabolic process | 25/2440 | 1.53E-12 | 5.37E-11 |
| GO:0006605 | protein targeting | 62/2440 | 1.66E-12 | 5.79E-11 |
| GO:0090205 | positive regulation of cholesterol metabolic process | 14/2440 | 2.08E-12 | 7.21E-11 |
| GO:0016052 | carbohydrate catabolic process | 40/2440 | 3.55E-12 | 1.22E-10 |
| GO:0019915 | lipid storage | 31/2440 | 4.34E-12 | 1.49E-10 |
| GO:0009145 | purine nucleoside triphosphate biosynthetic process | 27/2440 | 4.81E-12 | 1.63E-10 |
| GO:0046503 | glycerolipid catabolic process | 27/2440 | 4.81E-12 | 1.63E-10 |
| GO:0006544 | glycine metabolic process | 13/2440 | 4.90E-12 | 1.64E-10 |
| GO:0009074 | aromatic amino acid family catabolic process | 13/2440 | 4.90E-12 | 1.64E-10 |
| GO:0072593 | reactive oxygen species metabolic process | 56/2440 | 5.11E-12 | 1.70E-10 |
| GO:0051791 | medium-chain fatty acid metabolic process | 16/2440 | 6.50E-12 | 2.15E-10 |
| GO:0015986 | proton motive force-driven ATP synthesis | 15/2440 | 7.62E-12 | 2.49E-10 |
| GO:0043574 | peroxisomal transport | 15/2440 | 7.62E-12 | 2.49E-10 |
| GO:0050817 | coagulation | 45/2440 | 9.11E-12 | 2.96E-10 |
| GO:0051235 | maintenance of location | 69/2440 | 9.45E-12 | 3.06E-10 |
| GO:1905039 | carboxylic acid transmembrane transport | 39/2440 | 9.55E-12 | 3.07E-10 |
| GO:1903825 | organic acid transmembrane transport | 39/2440 | 1.22E-11 | 3.90E-10 |

| | | | | |
|---|---|---|---|---|
| GO:0007596 | blood coagulation | 44/2440 | 1.86E-11 | 5.92E-10 |
| GO:0009206 | purine ribonucleoside triphosphate biosynthetic process | 26/2440 | 2.22E-11 | 7.03E-10 |
| GO:0015718 | monocarboxylic acid transport | 36/2440 | 2.53E-11 | 7.94E-10 |
| GO:0015914 | phospholipid transport | 30/2440 | 3.39E-11 | 1.06E-09 |
| GO:0007599 | hemostasis | 44/2440 | 3.46E-11 | 1.08E-09 |
| GO:0034368 | protein-lipid complex remodeling | 16/2440 | 3.52E-11 | 1.08E-09 |
| GO:0034369 | plasma lipoprotein particle remodeling | 16/2440 | 3.52E-11 | 1.08E-09 |
| GO:0044262 | cellular carbohydrate metabolic process | 61/2440 | 3.86E-11 | 1.18E-09 |
| GO:0034308 | primary alcohol metabolic process | 28/2440 | 4.24E-11 | 1.29E-09 |
| GO:0140053 | mitochondrial gene expression | 33/2440 | 4.87E-11 | 1.47E-09 |
| GO:0010866 | regulation of triglyceride biosynthetic process | 15/2440 | 4.96E-11 | 1.49E-09 |
| GO:0019432 | triglyceride biosynthetic process | 19/2440 | 5.21E-11 | 1.55E-09 |
| GO:0046460 | neutral lipid biosynthetic process | 22/2440 | 5.21E-11 | 1.55E-09 |
| GO:0046463 | acylglycerol biosynthetic process | 22/2440 | 5.21E-11 | 1.55E-09 |
| GO:0000038 | very long-chain fatty acid metabolic process | 18/2440 | 5.25E-11 | 1.55E-09 |
| GO:0035966 | response to topologically incorrect protein | 38/2440 | 6.57E-11 | 1.93E-09 |
| GO:0045923 | positive regulation of fatty acid metabolic process | 21/2440 | 6.60E-11 | 1.93E-09 |
| GO:0009201 | ribonucleoside triphosphate biosynthetic process | 26/2440 | 6.79E-11 | 1.97E-09 |

| | | | | |
|---|---|---|---|---|
| GO:0006006 | glucose metabolic process | 51/2440 | 7.35E-11 | 2.12E-09 |
| GO:0042178 | xenobiotic catabolic process | 16/2440 | 7.56E-11 | 2.17E-09 |
| GO:0042157 | lipoprotein metabolic process | 37/2440 | 8.58E-11 | 2.45E-09 |
| GO:0032368 | regulation of lipid transport | 35/2440 | 8.81E-11 | 2.51E-09 |
| GO:0032371 | regulation of sterol transport | 17/2440 | 9.21E-11 | 2.60E-09 |
| GO:0032374 | regulation of cholesterol transport | 17/2440 | 9.21E-11 | 2.60E-09 |
| GO:0006754 | ATP biosynthetic process | 23/2440 | 9.34E-11 | 2.61E-09 |
| GO:0071806 | protein transmembrane transport | 23/2440 | 9.34E-11 | 2.61E-09 |
| GO:0009071 | serine family amino acid catabolic process | Nov-40 | 1.04E-10 | 2.89E-09 |
| GO:0034375 | high-density lipoprotein particle remodeling | Nov-40 | 1.04E-10 | 2.89E-09 |
| GO:0009144 | purine nucleoside triphosphate metabolic process | 30/2440 | 1.14E-10 | 3.13E-09 |
| GO:0065002 | intracellular protein transmembrane transport | 22/2440 | 1.27E-10 | 3.48E-09 |
| GO:0008654 | phospholipid biosynthetic process | 50/2440 | 1.43E-10 | 3.90E-09 |
| GO:0034367 | protein-containing complex remodeling | 16/2440 | 1.55E-10 | 4.19E-09 |
| GO:0042572 | retinol metabolic process | 16/2440 | 1.55E-10 | 4.19E-09 |
| GO:1990542 | mitochondrial transmembrane transport | 27/2440 | 1.76E-10 | 4.71E-09 |
| GO:0009308 | amine metabolic process | 35/2440 | 1.83E-10 | 4.90E-09 |
| GO:0006085 | acetyl-CoA biosynthetic process | 13/2440 | 1.86E-10 | 4.93E-09 |

| | | | | |
|---|---|---|---|---|
| GO:0042730 | fibrinolysis | 13/2440 | 1.86E-10 | 4.93E-09 |
| GO:0006107 | oxaloacetate metabolic process | Oct-40 | 1.89E-10 | 4.97E-09 |
| GO:0033539 | fatty acid beta-oxidation using acyl-CoA dehydrogenase | Oct-40 | 1.89E-10 | 4.97E-09 |
| GO:0070972 | protein localization to endoplasmic reticulum | 26/2440 | 1.94E-10 | 5.06E-09 |
| GO:0006783 | heme biosynthetic process | 15/2440 | 2.49E-10 | 6.46E-09 |
| GO:0009205 | purine ribonucleoside triphosphate metabolic process | 28/2440 | 2.87E-10 | 7.42E-09 |
| GO:0009116 | nucleoside metabolic process | 22/2440 | 2.96E-10 | 7.62E-09 |
| GO:0046470 | phosphatidylcholine metabolic process | 23/2440 | 3.10E-10 | 7.93E-09 |
| GO:0043467 | regulation of generation of precursor metabolites and energy | 38/2440 | 3.11E-10 | 7.93E-09 |
| GO:0009142 | nucleoside triphosphate biosynthetic process | 27/2440 | 3.32E-10 | 8.44E-09 |
| GO:0031998 | regulation of fatty acid beta-oxidation | 14/2440 | 3.69E-10 | 9.33E-09 |
| GO:1901657 | glycosyl compound metabolic process | 26/2440 | 3.76E-10 | 9.47E-09 |
| GO:1901616 | organic hydroxy compound catabolic process | 24/2440 | 4.42E-10 | 1.11E-08 |
| GO:0050810 | regulation of steroid biosynthetic process | 27/2440 | 4.53E-10 | 1.13E-08 |
| GO:0045833 | negative regulation of lipid metabolic process | 31/2440 | 4.87E-10 | 1.21E-08 |
| GO:0045540 | regulation of cholesterol biosynthetic process | 13/2440 | 4.90E-10 | 1.21E-08 |
| GO:0106118 | regulation of sterol biosynthetic process | 13/2440 | 4.90E-10 | 1.21E-08 |
| GO:0045723 | positive regulation of fatty acid biosynthetic process | 15/2440 | 5.15E-10 | 1.26E-08 |

| | | | | |
|---|---|---|---|---|
| GO:0030195 | negative regulation of blood coagulation | 20/2440 | 5.45E-10 | 1.33E-08 |
| GO:0006103 | 2-oxoglutarate metabolic process | Dec-40 | 5.48E-10 | 1.33E-08 |
| GO:0042776 | proton motive force-driven mitochondrial ATP synthesis | Dec-40 | 5.48E-10 | 1.33E-08 |
| GO:0042908 | xenobiotic transport | 17/2440 | 5.77E-10 | 1.39E-08 |
| GO:0006779 | porphyrin-containing compound biosynthetic process | 16/2440 | 5.84E-10 | 1.40E-08 |
| GO:0033014 | tetrapyrrole biosynthetic process | 16/2440 | 5.84E-10 | 1.40E-08 |
| GO:0046364 | monosaccharide biosynthetic process | 28/2440 | 6.93E-10 | 1.65E-08 |
| GO:0046461 | neutral lipid catabolic process | 19/2440 | 7.04E-10 | 1.66E-08 |
| GO:0046464 | acylglycerol catabolic process | 19/2440 | 7.04E-10 | 1.66E-08 |
| GO:0006576 | cellular biogenic amine metabolic process | 31/2440 | 8.13E-10 | 1.91E-08 |
| GO:1900047 | negative regulation of hemostasis | 20/2440 | 8.47E-10 | 1.99E-08 |
| GO:0016051 | carbohydrate biosynthetic process | 45/2440 | 8.77E-10 | 2.04E-08 |
| GO:0016485 | protein processing | 56/2440 | 8.80E-10 | 2.04E-08 |
| GO:0009199 | ribonucleoside triphosphate metabolic process | 28/2440 | 9.20E-10 | 2.13E-08 |
| GO:0010883 | regulation of lipid storage | 21/2440 | 9.35E-10 | 2.15E-08 |
| GO:0044106 | cellular amine metabolic process | 33/2440 | 1.01E-09 | 2.32E-08 |
| GO:0000096 | sulfur amino acid metabolic process | 16/2440 | 1.08E-09 | 2.45E-08 |
| GO:0010875 | positive regulation of cholesterol efflux | 16/2440 | 1.08E-09 | 2.45E-08 |

| | | | | |
|---|---|---|---|---|
| GO:1901292 | nucleoside phosphate catabolic process | 25/2440 | 1.11E-09 | 2.52E-08 |
| GO:1901136 | carbohydrate derivative catabolic process | 37/2440 | 1.19E-09 | 2.68E-08 |
| GO:0050819 | negative regulation of coagulation | 20/2440 | 1.30E-09 | 2.92E-08 |
| GO:0042060 | wound healing | 68/2440 | 1.47E-09 | 3.29E-08 |
| GO:0006625 | protein targeting to peroxisome | Dec-40 | 1.52E-09 | 3.36E-08 |
| GO:0072662 | protein localization to peroxisome | Dec-40 | 1.52E-09 | 3.36E-08 |
| GO:0072663 | establishment of protein localization to peroxisome | Dec-40 | 1.52E-09 | 3.36E-08 |
| GO:0006957 | complement activation, alternative pathway | Nov-40 | 1.56E-09 | 3.42E-08 |
| GO:0046185 | aldehyde catabolic process | Nov-40 | 1.56E-09 | 3.42E-08 |
| GO:0044273 | sulfur compound catabolic process | 14/2440 | 1.71E-09 | 3.74E-08 |
| GO:0009066 | aspartate family amino acid metabolic process | 17/2440 | 1.71E-09 | 3.74E-08 |
| GO:0050818 | regulation of coagulation | 24/2440 | 1.72E-09 | 3.75E-08 |
| GO:0009112 | nucleobase metabolic process | 16/2440 | 1.92E-09 | 4.15E-08 |
| GO:0050996 | positive regulation of lipid catabolic process | 16/2440 | 1.92E-09 | 4.15E-08 |
| GO:0006986 | response to unfolded protein | 32/2440 | 2.10E-09 | 4.51E-08 |
| GO:0030193 | regulation of blood coagulation | 23/2440 | 2.66E-09 | 5.66E-08 |
| GO:0042440 | pigment metabolic process | 23/2440 | 2.66E-09 | 5.66E-08 |
| GO:0009164 | nucleoside catabolic process | 13/2440 | 2.67E-09 | 5.66E-08 |

| | | | | |
|---|---|---|---|---|
| GO:0034377 | plasma lipoprotein particle assembly | 13/2440 | 2.67E-09 | 5.66E-08 |
| GO:0034381 | plasma lipoprotein particle clearance | 17/2440 | 2.86E-09 | 5.98E-08 |
| GO:0055090 | acylglycerol homeostasis | 17/2440 | 2.86E-09 | 5.98E-08 |
| GO:0070328 | triglyceride homeostasis | 17/2440 | 2.86E-09 | 5.98E-08 |
| GO:1905953 | negative regulation of lipid localization | 20/2440 | 2.94E-09 | 6.14E-08 |
| GO:0034656 | nucleobase-containing small molecule catabolic process | 14/2440 | 3.41E-09 | 7.09E-08 |
| GO:0009991 | response to extracellular stimulus | 71/2440 | 3.60E-09 | 7.46E-08 |
| GO:0015919 | peroxisomal membrane transport | Dec-40 | 3.79E-09 | 7.82E-08 |
| GO:0009141 | nucleoside triphosphate metabolic process | 31/2440 | 4.37E-09 | 8.98E-08 |
| GO:0006767 | water-soluble vitamin metabolic process | 17/2440 | 4.67E-09 | 9.55E-08 |
| GO:0006979 | response to oxidative stress | 71/2440 | 4.96E-09 | 1.01E-07 |
| GO:0006879 | cellular iron ion homeostasis | 23/2440 | 5.10E-09 | 1.03E-07 |
| GO:1900046 | regulation of hemostasis | 23/2440 | 5.10E-09 | 1.03E-07 |
| GO:0008299 | isoprenoid biosynthetic process | 16/2440 | 5.68E-09 | 1.15E-07 |
| GO:0009162 | deoxyribonucleoside monophosphate metabolic process | 15/2440 | 6.42E-09 | 1.29E-07 |
| GO:0015721 | bile acid and bile salt transport | 15/2440 | 6.42E-09 | 1.29E-07 |
| GO:1905954 | positive regulation of lipid localization | 28/2440 | 7.57E-09 | 1.51E-07 |
| GO:0009166 | nucleotide catabolic process | 22/2440 | 7.93E-09 | 1.58E-07 |

| | | | | |
|---|---|---|---|---|
| GO:0010038 | response to metal ion | 56/2440 | 8.34E-09 | 1.65E-07 |
| GO:0006144 | purine nucleobase metabolic process | Dec-40 | 8.75E-09 | 1.72E-07 |
| GO:0009159 | deoxyribonucleoside monophosphate catabolic process | Dec-40 | 8.75E-09 | 1.72E-07 |
| GO:0031638 | zymogen activation | 28/2440 | 9.67E-09 | 1.89E-07 |
| GO:0055072 | iron ion homeostasis | 27/2440 | 9.68E-09 | 1.89E-07 |
| GO:0034384 | high-density lipoprotein particle clearance | Sep-40 | 1.04E-08 | 2.02E-07 |
| GO:0019433 | triglyceride catabolic process | 15/2440 | 1.11E-08 | 2.16E-07 |
| GO:0046320 | regulation of fatty acid oxidation | 17/2440 | 1.18E-08 | 2.27E-07 |
| GO:1901568 | fatty acid derivative metabolic process | 17/2440 | 1.18E-08 | 2.27E-07 |
| GO:0072594 | establishment of protein localization to organelle | 72/2440 | 1.23E-08 | 2.36E-07 |
| GO:0050878 | regulation of body fluid levels | 65/2440 | 1.87E-08 | 3.58E-07 |
| GO:0046164 | alcohol catabolic process | 18/2440 | 1.96E-08 | 3.75E-07 |
| GO:0051346 | negative regulation of hydrolase activity | 64/2440 | 2.10E-08 | 3.98E-07 |
| GO:0045861 | negative regulation of proteolysis | 63/2440 | 2.11E-08 | 3.98E-07 |
| GO:0052548 | regulation of endopeptidase activity | 63/2440 | 2.11E-08 | 3.98E-07 |
| GO:0009125 | nucleoside monophosphate catabolic process | 13/2440 | 2.19E-08 | 4.12E-07 |
| GO:0030449 | regulation of complement activation | 13/2440 | 2.19E-08 | 4.12E-07 |
| GO:0043434 | response to peptide hormone | 63/2440 | 2.35E-08 | 4.40E-07 |

| | | | | |
|---|---|---|---|---|
| GO:1901661 | quinone metabolic process | 16/2440 | 2.43E-08 | 4.54E-07 |
| GO:0045017 | glycerolipid biosynthetic process | 44/2440 | 2.48E-08 | 4.61E-07 |
| GO:0062014 | negative regulation of small molecule metabolic process | 27/2440 | 2.58E-08 | 4.77E-07 |
| GO:0010874 | regulation of cholesterol efflux | 17/2440 | 2.77E-08 | 5.12E-07 |
| GO:0010867 | positive regulation of triglyceride biosynthetic process | Nov-40 | 2.85E-08 | 5.26E-07 |
| GO:0009119 | ribonucleoside metabolic process | 15/2440 | 3.10E-08 | 5.70E-07 |
| GO:0046031 | ADP metabolic process | 27/2440 | 3.26E-08 | 5.96E-07 |
| GO:0006826 | iron ion transport | 20/2440 | 3.66E-08 | 6.67E-07 |
| GO:0046700 | heterocycle catabolic process | 71/2440 | 3.70E-08 | 6.73E-07 |
| GO:0043651 | linoleic acid metabolic process | 14/2440 | 3.71E-08 | 6.73E-07 |
| GO:0010039 | response to iron ion | Dec-40 | 3.82E-08 | 6.91E-07 |
| GO:0043650 | dicarboxylic acid biosynthetic process | Sep-40 | 3.84E-08 | 6.91E-07 |
| GO:0065005 | protein-lipid complex assembly | 13/2440 | 4.04E-08 | 7.26E-07 |
| GO:0002920 | regulation of humoral immune response | 18/2440 | 4.21E-08 | 7.51E-07 |
| GO:0009262 | deoxyribonucleotide metabolic process | 18/2440 | 4.21E-08 | 7.51E-07 |
| GO:0007006 | mitochondrial membrane organization | 28/2440 | 4.84E-08 | 8.59E-07 |
| GO:0036503 | ERAD pathway | 28/2440 | 4.84E-08 | 8.59E-07 |
| GO:0042278 | purine nucleoside metabolic process | 15/2440 | 5.00E-08 | 8.84E-07 |

| | | | | |
|---|---|---|---|---|
| GO:0008637 | apoptotic mitochondrial changes | 30/2440 | 5.02E-08 | 8.85E-07 |
| GO:0019319 | hexose biosynthetic process | 24/2440 | 5.49E-08 | 9.64E-07 |
| GO:0006730 | one-carbon metabolic process | 16/2440 | 5.83E-08 | 1.02E-06 |
| GO:0006739 | NADP metabolic process | 16/2440 | 5.83E-08 | 1.02E-06 |
| GO:0009261 | ribonucleotide catabolic process | 17/2440 | 6.16E-08 | 1.07E-06 |
| GO:0045922 | negative regulation of fatty acid metabolic process | 14/2440 | 6.24E-08 | 1.08E-06 |
| GO:0016114 | terpenoid biosynthetic process | Nov-40 | 6.25E-08 | 1.08E-06 |
| GO:0052547 | regulation of peptidase activity | 72/2440 | 6.29E-08 | 1.08E-06 |
| GO:0006096 | glycolytic process | 25/2440 | 7.01E-08 | 1.21E-06 |
| GO:0009264 | deoxyribonucleotide catabolic process | 13/2440 | 7.19E-08 | 1.23E-06 |
| GO:0031639 | plasminogen activation | 13/2440 | 7.19E-08 | 1.23E-06 |
| GO:0072525 | pyridine-containing compound biosynthetic process | Dec-40 | 7.37E-08 | 1.26E-06 |
| GO:0044270 | cellular nitrogen compound catabolic process | 70/2440 | 7.43E-08 | 1.26E-06 |
| GO:0035967 | cellular response to topologically incorrect protein | 28/2440 | 7.44E-08 | 1.26E-06 |
| GO:1901652 | response to peptide | 69/2440 | 7.69E-08 | 1.30E-06 |
| GO:0072524 | pyridine-containing compound metabolic process | 15/2440 | 7.91E-08 | 1.33E-06 |
| GO:0009123 | nucleoside monophosphate metabolic process | 21/2440 | 7.92E-08 | 1.33E-06 |
| GO:0009135 | purine nucleoside diphosphate metabolic process | 27/2440 | 8.03E-08 | 1.34E-06 |

| | | | | |
|---|---|---|---|---|
| GO:0009179 | purine ribonucleoside diphosphate metabolic process | 27/2440 | 8.03E-08 | 1.34E-06 |
| GO:0006757 | ATP generation from ADP | 25/2440 | 8.89E-08 | 1.48E-06 |
| GO:0072523 | purine-containing compound catabolic process | 17/2440 | 9.00E-08 | 1.49E-06 |
| GO:0009070 | serine family amino acid biosynthetic process | Oct-40 | 9.26E-08 | 1.52E-06 |
| GO:0055091 | phospholipid homeostasis | Oct-40 | 9.26E-08 | 1.52E-06 |
| GO:0072350 | tricarboxylic acid metabolic process | Oct-40 | 9.26E-08 | 1.52E-06 |
| GO:0009132 | nucleoside diphosphate metabolic process | 31/2440 | 9.74E-08 | 1.60E-06 |
| GO:0006165 | nucleoside diphosphate phosphorylation | 27/2440 | 9.98E-08 | 1.63E-06 |
| GO:0031667 | response to nutrient levels | 63/2440 | 1.00E-07 | 1.63E-06 |
| GO:0000050 | urea cycle | Aug-40 | 1.01E-07 | 1.63E-06 |
| GO:0110095 | cellular detoxification of aldehyde | Aug-40 | 1.01E-07 | 1.63E-06 |
| GO:0006888 | endoplasmic reticulum to Golgi vesicle-mediated transport | 29/2440 | 1.02E-07 | 1.65E-06 |
| GO:0006063 | uronic acid metabolic process | Sep-40 | 1.15E-07 | 1.84E-06 |
| GO:0009120 | deoxyribonucleoside metabolic process | Sep-40 | 1.15E-07 | 1.84E-06 |
| GO:0019585 | glucuronate metabolic process | Sep-40 | 1.15E-07 | 1.84E-06 |
| GO:0070508 | cholesterol import | Sep-40 | 1.15E-07 | 1.84E-06 |
| GO:0098656 | anion transmembrane transport | 41/2440 | 1.22E-07 | 1.94E-06 |
| GO:0055076 | transition metal ion homeostasis | 32/2440 | 1.22E-07 | 1.94E-06 |

| | | | | |
|---|---|---|---|---|
| GO:0042573 | retinoic acid metabolic process | 13/2440 | 1.24E-07 | 1.96E-06 |
| GO:0046128 | purine ribonucleoside metabolic process | 13/2440 | 1.24E-07 | 1.96E-06 |
| GO:0009394 | 2'-deoxyribonucleotide metabolic process | 17/2440 | 1.30E-07 | 2.04E-06 |
| GO:0019692 | deoxyribose phosphate metabolic process | 17/2440 | 1.30E-07 | 2.04E-06 |
| GO:0032527 | protein exit from endoplasmic reticulum | 16/2440 | 1.31E-07 | 2.06E-06 |
| GO:0010878 | cholesterol storage | Dec-40 | 1.36E-07 | 2.13E-06 |
| GO:0010906 | regulation of glucose metabolic process | 31/2440 | 1.41E-07 | 2.20E-06 |
| GO:0006839 | mitochondrial transport | 34/2440 | 1.50E-07 | 2.33E-06 |
| GO:0046939 | nucleotide phosphorylation | 27/2440 | 1.88E-07 | 2.91E-06 |
| GO:0032543 | mitochondrial translation | 22/2440 | 1.90E-07 | 2.94E-06 |
| GO:0010896 | regulation of triglyceride catabolic process | Oct-40 | 2.08E-07 | 3.19E-06 |
| GO:0046415 | urate metabolic process | Oct-40 | 2.08E-07 | 3.19E-06 |
| GO:0045047 | protein targeting to ER | 16/2440 | 2.79E-07 | 4.28E-06 |
| GO:1990845 | adaptive thermogenesis | 35/2440 | 2.82E-07 | 4.31E-06 |
| GO:0032782 | bile acid secretion | Sep-40 | 2.98E-07 | 4.54E-06 |
| GO:0050667 | homocysteine metabolic process | Sep-40 | 2.98E-07 | 4.54E-06 |
| GO:0010823 | negative regulation of mitochondrion organization | 18/2440 | 3.20E-07 | 4.86E-06 |
| GO:0010675 | regulation of cellular carbohydrate metabolic process | 35/2440 | 3.29E-07 | 4.98E-06 |

| | | | | |
|---|---|---|---|---|
| GO:0010951 | negative regulation of endopeptidase activity | 38/2440 | 3.35E-07 | 5.04E-06 |
| GO:1901658 | glycosyl compound catabolic process | 13/2440 | 3.37E-07 | 5.04E-06 |
| GO:0006122 | mitochondrial electron transport, ubiquinol to cytochrome c | Aug-40 | 3.42E-07 | 5.04E-06 |
| GO:0006558 | L-phenylalanine metabolic process | Aug-40 | 3.42E-07 | 5.04E-06 |
| GO:0009812 | flavonoid metabolic process | Aug-40 | 3.42E-07 | 5.04E-06 |
| GO:0019627 | urea metabolic process | Aug-40 | 3.42E-07 | 5.04E-06 |
| GO:0045542 | positive regulation of cholesterol biosynthetic process | Aug-40 | 3.42E-07 | 5.04E-06 |
| GO:0071941 | nitrogen cycle metabolic process | Aug-40 | 3.42E-07 | 5.04E-06 |
| GO:0106120 | positive regulation of sterol biosynthetic process | Aug-40 | 3.42E-07 | 5.04E-06 |
| GO:1902221 | erythrose 4-phosphate/phosphoenolpyruvate family amino acid metabolic process | Aug-40 | 3.42E-07 | 5.04E-06 |
| GO:0051259 | protein complex oligomerization | 46/2440 | 3.47E-07 | 5.10E-06 |
| GO:0032102 | negative regulation of response to external stimulus | 65/2440 | 3.58E-07 | 5.25E-06 |
| GO:0010888 | negative regulation of lipid storage | Dec-40 | 4.14E-07 | 6.04E-06 |
| GO:0046386 | deoxyribose phosphate catabolic process | Dec-40 | 4.14E-07 | 6.04E-06 |
| GO:0046916 | cellular transition metal ion homeostasis | 27/2440 | 4.17E-07 | 6.07E-06 |
| GO:0051004 | regulation of lipoprotein lipase activity | Oct-40 | 4.32E-07 | 6.25E-06 |
| GO:0072378 | blood coagulation, fibrin clot formation | Oct-40 | 4.32E-07 | 6.25E-06 |
| GO:0009158 | ribonucleoside monophosphate catabolic process | Nov-40 | 4.59E-07 | 6.62E-06 |

| | | | | |
|---|---|---|---|---|
| GO:0010885 | regulation of cholesterol storage | Nov-40 | 4.59E-07 | 6.62E-06 |
| GO:2000377 | regulation of reactive oxygen species metabolic process | 34/2440 | 4.61E-07 | 6.62E-06 |
| GO:0051260 | protein homooligomerization | 37/2440 | 4.71E-07 | 6.75E-06 |
| GO:0051289 | protein homotetramerization | 17/2440 | 5.04E-07 | 7.21E-06 |
| GO:0006094 | gluconeogenesis | 22/2440 | 5.09E-07 | 7.26E-06 |
| GO:0036314 | response to sterol | 13/2440 | 5.35E-07 | 7.58E-06 |
| GO:0042558 | pteridine-containing compound metabolic process | 13/2440 | 5.35E-07 | 7.58E-06 |
| GO:0045454 | cell redox homeostasis | 13/2440 | 5.35E-07 | 7.58E-06 |
| GO:0006109 | regulation of carbohydrate metabolic process | 39/2440 | 5.48E-07 | 7.74E-06 |
| GO:0019748 | secondary metabolic process | 18/2440 | 5.85E-07 | 8.24E-06 |
| GO:0006953 | acute-phase response | 15/2440 | 5.99E-07 | 8.42E-06 |
| GO:0106106 | cold-induced thermogenesis | 32/2440 | 6.60E-07 | 9.25E-06 |
| GO:0006570 | tyrosine metabolic process | Sep-40 | 6.89E-07 | 9.55E-06 |
| GO:0010887 | negative regulation of cholesterol storage | Sep-40 | 6.89E-07 | 9.55E-06 |
| GO:0033700 | phospholipid efflux | Sep-40 | 6.89E-07 | 9.55E-06 |
| GO:0110096 | cellular response to aldehyde | Sep-40 | 6.89E-07 | 9.55E-06 |
| GO:0046148 | pigment biosynthetic process | 17/2440 | 6.90E-07 | 9.55E-06 |
| GO:0009185 | ribonucleoside diphosphate metabolic process | 27/2440 | 7.37E-07 | 1.02E-05 |

| | | | | |
|---|---|---|---|---|
| GO:0097305 | response to alcohol | 43/2440 | 7.46E-07 | 1.03E-05 |
| GO:0006956 | complement activation | 38/2440 | 7.70E-07 | 1.06E-05 |
| GO:0051055 | negative regulation of lipid biosynthetic process | 18/2440 | 7.81E-07 | 1.07E-05 |
| GO:0009065 | glutamine family amino acid catabolic process | Nov-40 | 8.12E-07 | 1.11E-05 |
| GO:0007007 | inner mitochondrial membrane organization | 13/2440 | 8.31E-07 | 1.13E-05 |
| GO:0006525 | arginine metabolic process | Oct-40 | 8.42E-07 | 1.14E-05 |
| GO:0006613 | cotranslational protein targeting to membrane | Oct-40 | 8.42E-07 | 1.14E-05 |
| GO:0019363 | pyridine nucleotide biosynthetic process | Oct-40 | 8.42E-07 | 1.14E-05 |
| GO:1904251 | regulation of bile acid metabolic process | Oct-40 | 8.42E-07 | 1.14E-05 |
| GO:0007030 | Golgi organization | 30/2440 | 9.33E-07 | 1.26E-05 |
| GO:0034620 | cellular response to unfolded protein | 23/2440 | 9.43E-07 | 1.26E-05 |
| GO:0006086 | acetyl-CoA biosynthetic process from pyruvate | Aug-40 | 9.48E-07 | 1.26E-05 |
| GO:0006152 | purine nucleoside catabolic process | Aug-40 | 9.48E-07 | 1.26E-05 |
| GO:0009435 | NAD biosynthetic process | Aug-40 | 9.48E-07 | 1.26E-05 |
| GO:0010898 | positive regulation of triglyceride catabolic process | Aug-40 | 9.48E-07 | 1.26E-05 |
| GO:0052695 | cellular glucuronidation | Aug-40 | 9.48E-07 | 1.26E-05 |
| GO:0071281 | cellular response to iron ion | Aug-40 | 9.48E-07 | 1.26E-05 |
| GO:0051262 | protein tetramerization | 22/2440 | 1.01E-06 | 1.34E-05 |

| | | | | |
|---|---|---|---|---|
| GO:0010821 | regulation of mitochondrion organization | 31/2440 | 1.08E-06 | 1.43E-05 |
| GO:0072599 | establishment of protein localization to endoplasmic reticulum | 16/2440 | 1.09E-06 | 1.43E-05 |
| GO:0042430 | indole-containing compound metabolic process | Dec-40 | 1.11E-06 | 1.46E-05 |
| GO:0006835 | dicarboxylic acid transport | 23/2440 | 1.17E-06 | 1.53E-05 |
| GO:0050892 | intestinal absorption | 15/2440 | 1.22E-06 | 1.59E-05 |
| GO:0006692 | prostanoid metabolic process | 17/2440 | 1.26E-06 | 1.64E-05 |
| GO:0006693 | prostaglandin metabolic process | 17/2440 | 1.26E-06 | 1.64E-05 |
| GO:0006457 | protein folding | 34/2440 | 1.30E-06 | 1.69E-05 |
| GO:0044042 | glucan metabolic process | 21/2440 | 1.34E-06 | 1.74E-05 |
| GO:0072376 | protein activation cascade | Nov-40 | 1.38E-06 | 1.79E-05 |
| GO:0015909 | long-chain fatty acid transport | 19/2440 | 1.40E-06 | 1.81E-05 |
| GO:0019359 | nicotinamide nucleotide biosynthetic process | Sep-40 | 1.46E-06 | 1.85E-05 |
| GO:0034433 | steroid esterification | Sep-40 | 1.46E-06 | 1.85E-05 |
| GO:0034434 | sterol esterification | Sep-40 | 1.46E-06 | 1.85E-05 |
| GO:0034435 | cholesterol esterification | Sep-40 | 1.46E-06 | 1.85E-05 |
| GO:0042454 | ribonucleoside catabolic process | Sep-40 | 1.46E-06 | 1.85E-05 |
| GO:0045717 | negative regulation of fatty acid biosynthetic process | Sep-40 | 1.46E-06 | 1.85E-05 |
| GO:0070857 | regulation of bile acid biosynthetic process | Sep-40 | 1.46E-06 | 1.85E-05 |

| | | | | |
|---|---|---|---|---|
| GO:0120161 | regulation of cold-induced thermogenesis | 31/2440 | 1.48E-06 | 1.87E-05 |
| GO:0006650 | glycerophospholipid metabolic process | 48/2440 | 1.52E-06 | 1.92E-05 |
| GO:0009110 | vitamin biosynthetic process | Oct-40 | 1.55E-06 | 1.96E-05 |
| GO:0015908 | fatty acid transport | 24/2440 | 1.58E-06 | 1.99E-05 |
| GO:2000378 | negative regulation of reactive oxygen species metabolic process | 17/2440 | 1.68E-06 | 2.11E-05 |
| GO:0010466 | negative regulation of peptidase activity | 46/2440 | 1.71E-06 | 2.15E-05 |
| GO:0008053 | mitochondrial fusion | Dec-40 | 1.75E-06 | 2.18E-05 |
| GO:0010893 | positive regulation of steroid biosynthetic process | Dec-40 | 1.75E-06 | 2.18E-05 |
| GO:1903008 | organelle disassembly | 26/2440 | 1.76E-06 | 2.19E-05 |
| GO:0061045 | negative regulation of wound healing | 20/2440 | 1.77E-06 | 2.20E-05 |
| GO:0018958 | phenol-containing compound metabolic process | 28/2440 | 1.82E-06 | 2.25E-05 |
| GO:0006568 | tryptophan metabolic process | Aug-40 | 2.28E-06 | 2.79E-05 |
| GO:0006586 | indolalkylamine metabolic process | Aug-40 | 2.28E-06 | 2.79E-05 |
| GO:0010872 | regulation of cholesterol esterification | Aug-40 | 2.28E-06 | 2.79E-05 |
| GO:0043649 | dicarboxylic acid catabolic process | Aug-40 | 2.28E-06 | 2.79E-05 |
| GO:0008209 | androgen metabolic process | Nov-40 | 2.28E-06 | 2.79E-05 |
| GO:0018904 | ether metabolic process | Nov-40 | 2.28E-06 | 2.79E-05 |
| GO:0036315 | cellular response to sterol | Nov-40 | 2.28E-06 | 2.79E-05 |

| | | | | |
|---|---|---|---|---|
| GO:0006195 | purine nucleotide catabolic process | 14/2440 | 2.63E-06 | 3.20E-05 |
| GO:0006959 | humoral immune response | 64/2440 | 2.71E-06 | 3.30E-05 |
| GO:0044743 | protein transmembrane import into intracellular organelle | 13/2440 | 2.77E-06 | 3.36E-05 |
| GO:0035337 | fatty-acyl-CoA metabolic process | Sep-40 | 2.86E-06 | 3.46E-05 |
| GO:0042407 | cristae formation | Sep-40 | 2.86E-06 | 3.46E-05 |
| GO:0001659 | temperature homeostasis | 36/2440 | 2.92E-06 | 3.52E-05 |
| GO:0010873 | positive regulation of cholesterol esterification | Jul-40 | 2.95E-06 | 3.52E-05 |
| GO:0034372 | very-low-density lipoprotein particle remodeling | Jul-40 | 2.95E-06 | 3.52E-05 |
| GO:0034380 | high-density lipoprotein particle assembly | Jul-40 | 2.95E-06 | 3.52E-05 |
| GO:0046055 | dGMP catabolic process | Jul-40 | 2.95E-06 | 3.52E-05 |
| GO:0046130 | purine ribonucleoside catabolic process | Jul-40 | 2.95E-06 | 3.52E-05 |
| GO:0048193 | Golgi vesicle transport | 47/2440 | 3.18E-06 | 3.79E-05 |
| GO:0051341 | regulation of oxidoreductase activity | 23/2440 | 3.19E-06 | 3.79E-05 |
| GO:0032370 | positive regulation of lipid transport | 20/2440 | 3.50E-06 | 4.15E-05 |
| GO:0030970 | retrograde protein transport, ER to cytosol | Nov-40 | 3.65E-06 | 4.31E-05 |
| GO:0070723 | response to cholesterol | Nov-40 | 3.65E-06 | 4.31E-05 |
| GO:1903513 | endoplasmic reticulum to cytosol transport | Nov-40 | 3.65E-06 | 4.31E-05 |
| GO:0016236 | macroautophagy | 43/2440 | 3.71E-06 | 4.36E-05 |

| | | | | |
|---|---|---|---|---|
| GO:0031668 | cellular response to extracellular stimulus | 42/2440 | 3.72E-06 | 4.36E-05 |
| GO:0009395 | phospholipid catabolic process | 17/2440 | 3.80E-06 | 4.44E-05 |
| GO:0042743 | hydrogen peroxide metabolic process | 17/2440 | 3.80E-06 | 4.44E-05 |
| GO:0009154 | purine ribonucleotide catabolic process | 13/2440 | 4.00E-06 | 4.67E-05 |
| GO:0005977 | glycogen metabolic process | 20/2440 | 4.36E-06 | 5.06E-05 |
| GO:0006073 | cellular glucan metabolic process | 20/2440 | 4.36E-06 | 5.06E-05 |
| GO:0035634 | response to stilbenoid | Oct-40 | 4.64E-06 | 5.37E-05 |
| GO:0042402 | cellular biogenic amine catabolic process | Oct-40 | 4.64E-06 | 5.37E-05 |
| GO:1901264 | carbohydrate derivative transport | 18/2440 | 4.87E-06 | 5.62E-05 |
| GO:0030433 | ubiquitin-dependent ERAD pathway | 21/2440 | 4.90E-06 | 5.64E-05 |
| GO:0010635 | regulation of mitochondrial fusion | Aug-40 | 4.93E-06 | 5.64E-05 |
| GO:0034756 | regulation of iron ion transport | Aug-40 | 4.93E-06 | 5.64E-05 |
| GO:0046459 | short-chain fatty acid metabolic process | Aug-40 | 4.93E-06 | 5.64E-05 |
| GO:1903725 | regulation of phospholipid metabolic process | 14/2440 | 5.09E-06 | 5.82E-05 |
| GO:0002921 | negative regulation of humoral immune response | Sep-40 | 5.29E-06 | 6.02E-05 |
| GO:0006614 | SRP-dependent cotranslational protein targeting to membrane | Sep-40 | 5.29E-06 | 6.02E-05 |
| GO:0006706 | steroid catabolic process | Nov-40 | 5.69E-06 | 6.43E-05 |
| GO:0019362 | pyridine nucleotide metabolic process | Nov-40 | 5.69E-06 | 6.43E-05 |

| | | | | |
|---|---|---|---|---|
| GO:0006775 | fat-soluble vitamin metabolic process | 13/2440 | 5.69E-06 | 6.43E-05 |
| GO:0071402 | cellular response to lipoprotein particle stimulus | 13/2440 | 5.69E-06 | 6.43E-05 |
| GO:0042593 | glucose homeostasis | 48/2440 | 6.19E-06 | 6.98E-05 |
| GO:0009161 | ribonucleoside monophosphate metabolic process | 16/2440 | 6.24E-06 | 7.02E-05 |
| GO:1903035 | negative regulation of response to wounding | 22/2440 | 6.43E-06 | 7.22E-05 |
| GO:0033500 | carbohydrate homeostasis | 48/2440 | 6.81E-06 | 7.63E-05 |
| GO:0009746 | response to hexose | 36/2440 | 6.95E-06 | 7.77E-05 |
| GO:0009172 | purine deoxyribonucleoside monophosphate catabolic process | Jul-40 | 7.52E-06 | 8.35E-05 |
| GO:0030300 | regulation of intestinal cholesterol absorption | Jul-40 | 7.52E-06 | 8.35E-05 |
| GO:0034370 | triglyceride-rich lipoprotein particle remodeling | Jul-40 | 7.52E-06 | 8.35E-05 |
| GO:0051006 | positive regulation of lipoprotein lipase activity | Jul-40 | 7.52E-06 | 8.35E-05 |
| GO:0009151 | purine deoxyribonucleotide metabolic process | Oct-40 | 7.59E-06 | 8.38E-05 |
| GO:0009310 | amine catabolic process | Oct-40 | 7.59E-06 | 8.38E-05 |
| GO:0019400 | alditol metabolic process | Oct-40 | 7.59E-06 | 8.38E-05 |
| GO:0009743 | response to carbohydrate | 38/2440 | 8.03E-06 | 8.84E-05 |
| GO:0050873 | brown fat cell differentiation | 17/2440 | 8.08E-06 | 8.88E-05 |
| GO:0050995 | negative regulation of lipid catabolic process | Nov-40 | 8.66E-06 | 9.50E-05 |
| GO:1904478 | regulation of intestinal absorption | Sep-40 | 9.29E-06 | 0.000101693 |

| | | | | |
|---|---|---|---|---|
| GO:0030968 | endoplasmic reticulum unfolded protein response | 18/2440 | 9.73E-06 | 0.000105916 |
| GO:0009170 | purine deoxyribonucleoside monophosphate metabolic process | Aug-40 | 9.77E-06 | 0.000105916 |
| GO:0032373 | positive regulation of sterol transport | Aug-40 | 9.77E-06 | 0.000105916 |
| GO:0032376 | positive regulation of cholesterol transport | Aug-40 | 9.77E-06 | 0.000105916 |
| GO:0046085 | adenosine metabolic process | Aug-40 | 9.77E-06 | 0.000105916 |
| GO:0034284 | response to monosaccharide | 36/2440 | 9.90E-06 | 0.000106995 |
| GO:0001818 | negative regulation of cytokine production | 48/2440 | 9.91E-06 | 0.000106995 |
| GO:1902930 | regulation of alcohol biosynthetic process | 15/2440 | 1.02E-05 | 0.000110207 |
| GO:0006612 | protein targeting to membrane | 26/2440 | 1.09E-05 | 0.000117604 |
| GO:0051180 | vitamin transport | 13/2440 | 1.11E-05 | 0.000118685 |
| GO:1905897 | regulation of response to endoplasmic reticulum stress | 19/2440 | 1.11E-05 | 0.000119125 |
| GO:0019751 | polyol metabolic process | 25/2440 | 1.12E-05 | 0.000120196 |
| GO:0042158 | lipoprotein biosynthetic process | 23/2440 | 1.14E-05 | 0.000121267 |
| GO:0070585 | protein localization to mitochondrion | 23/2440 | 1.14E-05 | 0.000121267 |
| GO:0000041 | transition metal ion transport | 24/2440 | 1.14E-05 | 0.000121267 |
| GO:0009895 | negative regulation of catabolic process | 50/2440 | 1.19E-05 | 0.000126769 |
| GO:0008210 | estrogen metabolic process | Oct-40 | 1.20E-05 | 0.000127249 |
| GO:0046496 | nicotinamide nucleotide metabolic process | Oct-40 | 1.20E-05 | 0.000127249 |

| | | | | |
|---|---|---|---|---|
| GO:0055094 | response to lipoprotein particle | Dec-40 | 1.24E-05 | 0.000130524 |
| GO:0006626 | protein targeting to mitochondrion | 19/2440 | 1.36E-05 | 0.000143879 |
| GO:0006071 | glycerol metabolic process | Sep-40 | 1.56E-05 | 0.000163428 |
| GO:0009067 | aspartate family amino acid biosynthetic process | Sep-40 | 1.56E-05 | 0.000163428 |
| GO:0071397 | cellular response to cholesterol | Sep-40 | 1.56E-05 | 0.000163428 |
| GO:0098856 | intestinal lipid absorption | Sep-40 | 1.56E-05 | 0.000163428 |
| GO:0001867 | complement activation, lectin pathway | Jul-40 | 1.67E-05 | 0.000173467 |
| GO:0006000 | fructose metabolic process | Jul-40 | 1.67E-05 | 0.000173467 |
| GO:0006534 | cysteine metabolic process | Jul-40 | 1.67E-05 | 0.000173467 |
| GO:0016558 | protein import into peroxisome matrix | Jul-40 | 1.67E-05 | 0.000173467 |
| GO:0061365 | positive regulation of triglyceride lipase activity | Jul-40 | 1.67E-05 | 0.000173467 |
| GO:0044275 | cellular carbohydrate catabolic process | Dec-40 | 1.74E-05 | 0.000179355 |
| GO:1903715 | regulation of aerobic respiration | Dec-40 | 1.74E-05 | 0.000179355 |
| GO:0009128 | purine nucleoside monophosphate catabolic process | Aug-40 | 1.81E-05 | 0.000186122 |
| GO:0030299 | intestinal cholesterol absorption | Aug-40 | 1.81E-05 | 0.000186122 |
| GO:0090114 | COPII-coated vesicle budding | Oct-40 | 1.85E-05 | 0.00019032 |
| GO:1903578 | regulation of ATP metabolic process | 21/2440 | 1.86E-05 | 0.000190424 |
| GO:0046033 | AMP metabolic process | Nov-40 | 1.88E-05 | 0.000192342 |

| | | | | |
|---|---|---|---|---|
| GO:0030148 | sphingolipid biosynthetic process | 22/2440 | 1.89E-05 | 0.000192432 |
| GO:0044264 | cellular polysaccharide metabolic process | 22/2440 | 1.89E-05 | 0.000192432 |
| GO:0006636 | unsaturated fatty acid biosynthetic process | 13/2440 | 2.04E-05 | 0.000207834 |
| GO:0010907 | positive regulation of glucose metabolic process | 14/2440 | 2.20E-05 | 0.000223302 |
| GO:0032868 | response to insulin | 41/2440 | 2.21E-05 | 0.000224561 |
| GO:0090199 | regulation of release of cytochrome c from mitochondria | 15/2440 | 2.23E-05 | 0.000225421 |
| GO:0006914 | autophagy | 66/2440 | 2.24E-05 | 0.000225925 |
| GO:0061919 | process utilizing autophagic mechanism | 66/2440 | 2.24E-05 | 0.000225925 |
| GO:0006662 | glycerol ether metabolic process | Sep-40 | 2.53E-05 | 0.0002542 |
| GO:0046653 | tetrahydrofolate metabolic process | Sep-40 | 2.53E-05 | 0.0002542 |
| GO:0006665 | sphingolipid metabolic process | 28/2440 | 2.64E-05 | 0.000264787 |
| GO:0001889 | liver development | 26/2440 | 2.64E-05 | 0.000264787 |
| GO:0009126 | purine nucleoside monophosphate metabolic process | 13/2440 | 2.73E-05 | 0.000272875 |
| GO:0006760 | folic acid-containing compound metabolic process | Oct-40 | 2.78E-05 | 0.000277784 |
| GO:0090150 | establishment of protein localization to membrane | 42/2440 | 2.86E-05 | 0.000285007 |
| GO:0009749 | response to glucose | 34/2440 | 3.04E-05 | 0.000302573 |
| GO:0006112 | energy reserve metabolic process | 20/2440 | 3.09E-05 | 0.000306776 |
| GO:0035459 | vesicle cargo loading | Aug-40 | 3.16E-05 | 0.000313494 |

| | | | | |
|---|---|---|---|---|
| GO:0051651 | maintenance of location in cell | 38/2440 | 3.22E-05 | 0.000318391 |
| GO:0000255 | allantoin metabolic process | Jul-40 | 3.36E-05 | 0.000330203 |
| GO:0034374 | low-density lipoprotein particle remodeling | Jul-40 | 3.36E-05 | 0.000330203 |
| GO:0046054 | dGMP metabolic process | Jul-40 | 3.36E-05 | 0.000330203 |
| GO:1904729 | regulation of intestinal lipid absorption | Jul-40 | 3.36E-05 | 0.000330203 |
| GO:1903409 | reactive oxygen species biosynthetic process | 16/2440 | 3.39E-05 | 0.000332866 |
| GO:0006984 | ER-nucleus signaling pathway | 13/2440 | 3.61E-05 | 0.000353479 |
| GO:0050766 | positive regulation of phagocytosis | 20/2440 | 3.68E-05 | 0.00036005 |
| GO:0006656 | phosphatidylcholine biosynthetic process | Sep-40 | 3.96E-05 | 0.000384924 |
| GO:0071404 | cellular response to low-density lipoprotein particle stimulus | Sep-40 | 3.96E-05 | 0.000384924 |
| GO:1903427 | negative regulation of reactive oxygen species biosynthetic process | Sep-40 | 3.96E-05 | 0.000384924 |
| GO:0061008 | hepaticobiliary system development | 26/2440 | 4.00E-05 | 0.000388645 |
| GO:0034637 | cellular carbohydrate biosynthetic process | 18/2440 | 4.12E-05 | 0.00039907 |
| GO:0005976 | polysaccharide metabolic process | 23/2440 | 4.12E-05 | 0.00039907 |
| GO:0043457 | regulation of cellular respiration | 16/2440 | 4.21E-05 | 0.000407315 |
| GO:0045913 | positive regulation of carbohydrate metabolic process | 19/2440 | 4.30E-05 | 0.000414973 |
| GO:0072655 | establishment of protein localization to mitochondrion | 21/2440 | 4.36E-05 | 0.00041965 |
| GO:0005978 | glycogen biosynthetic process | 13/2440 | 4.73E-05 | 0.000453642 |

| | | | | |
|---|---|---|---|---|
| GO:0009250 | glucan biosynthetic process | 13/2440 | 4.73E-05 | 0.000453642 |
| GO:0050764 | regulation of phagocytosis | 24/2440 | 5.25E-05 | 0.000502762 |
| GO:0002526 | acute inflammatory response | 23/2440 | 5.55E-05 | 0.00053076 |
| GO:0006707 | cholesterol catabolic process | Jun-40 | 5.70E-05 | 0.000542078 |
| GO:0016127 | sterol catabolic process | Jun-40 | 5.70E-05 | 0.000542078 |
| GO:0046053 | dAMP metabolic process | Jun-40 | 5.70E-05 | 0.000542078 |
| GO:1903027 | regulation of opsonization | Jun-40 | 5.70E-05 | 0.000542078 |
| GO:0009167 | purine ribonucleoside monophosphate metabolic process | Dec-40 | 5.91E-05 | 0.000560766 |
| GO:0010984 | regulation of lipoprotein particle clearance | Sep-40 | 6.01E-05 | 0.000568301 |
| GO:0042219 | cellular modified amino acid catabolic process | Sep-40 | 6.01E-05 | 0.000568301 |
| GO:0046456 | icosanoid biosynthetic process | 14/2440 | 6.02E-05 | 0.000568301 |
| GO:0007568 | aging | 19/2440 | 6.12E-05 | 0.000576751 |
| GO:0034599 | cellular response to oxidative stress | 43/2440 | 6.17E-05 | 0.00058003 |
| GO:0002138 | retinoic acid biosynthetic process | Jul-40 | 6.23E-05 | 0.000580876 |
| GO:0016102 | diterpenoid biosynthetic process | Jul-40 | 6.23E-05 | 0.000580876 |
| GO:0030497 | fatty acid elongation | Jul-40 | 6.23E-05 | 0.000580876 |
| GO:0042559 | pteridine-containing compound biosynthetic process | Jul-40 | 6.23E-05 | 0.000580876 |
| GO:0090110 | COPII-coated vesicle cargo loading | Jul-40 | 6.23E-05 | 0.000580876 |

| | | | | |
|---|---|---|---|---|
| GO:004288 6 | amide transport | 52/2440 | 6.37E-05 | 0.00059314 7 |
| GO:003134 8 | negative regulation of defense response | 39/2440 | 6.74E-05 | 0.00062708 |
| GO:000269 7 | regulation of immune effector process | 61/2440 | 7.26E-05 | 0.00067451 8 |
| GO:003559 2 | establishment of protein localization to extracellular region | 57/2440 | 7.45E-05 | 0.00069031 9 |
| GO:190357 3 | negative regulation of response to endoplasmic reticulum stress | Dec-40 | 7.80E-05 | 0.00072163 6 |
| GO:003546 1 | vitamin transmembrane transport | Aug-40 | 8.42E-05 | 0.00077700 9 |
| GO:004424 1 | lipid digestion | Aug-40 | 8.42E-05 | 0.00077700 9 |
| GO:005082 0 | positive regulation of coagulation | Sep-40 | 8.90E-05 | 0.00081937 4 |
| GO:000664 3 | membrane lipid metabolic process | 32/2440 | 9.08E-05 | 0.00083473 4 |
| GO:000183 6 | release of cytochrome c from mitochondria | 16/2440 | 9.53E-05 | 0.00087527 5 |
| GO:003236 5 | intracellular lipid transport | Dec-40 | 0.00010178 8 | 0.00093297 3 |
| GO:000915 5 | purine deoxyribonucleotide catabolic process | Jul-40 | 0.00010817 3 | 0.00097888 9 |
| GO:000916 9 | purine ribonucleoside monophosphate catabolic process | Jul-40 | 0.00010817 3 | 0.00097888 9 |
| GO:001593 6 | coenzyme A metabolic process | Jul-40 | 0.00010817 3 | 0.00097888 9 |
| GO:003293 3 | SREBP signaling pathway | Jul-40 | 0.00010817 3 | 0.00097888 9 |
| GO:003463 8 | phosphatidylcholine catabolic process | Jul-40 | 0.00010817 3 | 0.00097888 9 |
| GO:004218 2 | ketone catabolic process | Jul-40 | 0.00010817 3 | 0.00097888 9 |
| GO:004257 4 | retinal metabolic process | Jul-40 | 0.00010817 3 | 0.00097888 9 |

| | | | | |
|---|---|---|---|---|
| GO:0055089 | fatty acid homeostasis | Jul-40 | 0.000108173 | 0.000978889 |
| GO:0071692 | protein localization to extracellular region | 57/2440 | 0.000111932 | 0.001011295 |
| GO:0097193 | intrinsic apoptotic signaling pathway | 46/2440 | 0.000117784 | 0.001062486 |
| GO:0061041 | regulation of wound healing | 25/2440 | 0.000123007 | 0.001107844 |
| GO:0002082 | regulation of oxidative phosphorylation | Sep-40 | 0.000128584 | 0.001155757 |
| GO:0009306 | protein secretion | 56/2440 | 0.000128936 | 0.001155757 |
| GO:0034655 | nucleobase-containing compound catabolic process | 56/2440 | 0.000128936 | 0.001155757 |
| GO:0050665 | hydrogen peroxide biosynthetic process | Aug-40 | 0.000129964 | 0.001163135 |
| GO:0003333 | amino acid transmembrane transport | 20/2440 | 0.000134349 | 0.001200494 |
| GO:0071496 | cellular response to external stimulus | 45/2440 | 0.000135538 | 0.001209222 |
| GO:0007584 | response to nutrient | 16/2440 | 0.000139279 | 0.001240655 |
| GO:0008643 | carbohydrate transport | 28/2440 | 0.000144502 | 0.001285169 |
| GO:0017038 | protein import | 37/2440 | 0.000151651 | 0.001346644 |
| GO:0002683 | negative regulation of immune system process | 61/2440 | 0.000156382 | 0.001385116 |
| GO:0032869 | cellular response to insulin stimulus | 34/2440 | 0.000156609 | 0.001385116 |
| GO:0062125 | regulation of mitochondrial gene expression | Oct-40 | 0.000156714 | 0.001385116 |
| GO:0006584 | catecholamine metabolic process | 15/2440 | 0.000160021 | 0.001405618 |
| GO:0009712 | catechol-containing compound metabolic process | 15/2440 | 0.000160021 | 0.001405618 |

| | | | | |
|---|---|---|---|---|
| GO:0010676 | positive regulation of cellular carbohydrate metabolic process | 15/2440 | 0.000160021 | 0.001405618 |
| GO:0015807 | L-amino acid transport | 15/2440 | 0.000160021 | 0.001405618 |
| GO:0009309 | amine biosynthetic process | Dec-40 | 0.000168564 | 0.001475885 |
| GO:0042401 | cellular biogenic amine biosynthetic process | Dec-40 | 0.000168564 | 0.001475885 |
| GO:0000002 | mitochondrial genome maintenance | Nov-40 | 0.000168798 | 0.001475885 |
| GO:0071375 | cellular response to peptide hormone stimulus | 41/2440 | 0.00017266 | 0.001507331 |
| GO:0007586 | digestion | 25/2440 | 0.000177483 | 0.001546589 |
| GO:0042159 | lipoprotein catabolic process | Jul-40 | 0.000178244 | 0.001546589 |
| GO:0071501 | cellular response to sterol depletion | Jul-40 | 0.000178244 | 0.001546589 |
| GO:1902001 | fatty acid transmembrane transport | Jul-40 | 0.000178244 | 0.001546589 |
| GO:0120162 | positive regulation of cold-induced thermogenesis | 20/2440 | 0.000180529 | 0.001564007 |
| GO:0072527 | pyrimidine-containing compound metabolic process | 14/2440 | 0.000180801 | 0.001564007 |
| GO:0046321 | positive regulation of fatty acid oxidation | Aug-40 | 0.000194386 | 0.001676436 |
| GO:0071071 | regulation of phospholipid biosynthetic process | Aug-40 | 0.000194386 | 0.001676436 |
| GO:0060191 | regulation of lipase activity | 18/2440 | 0.000196735 | 0.001694125 |
| GO:0010822 | positive regulation of mitochondrion organization | 17/2440 | 0.000200495 | 0.00172389 |
| GO:2000112 | regulation of cellular macromolecule biosynthetic process | 64/2440 | 0.000211058 | 0.001811977 |
| GO:0034248 | regulation of cellular amide metabolic process | 61/2440 | 0.000212074 | 0.001817957 |

| | | | | |
|---|---|---|---|---|
| GO:0008207 | C21-steroid hormone metabolic process | Dec-40 | 0.000214052 | 0.001831364 |
| GO:0006213 | pyrimidine nucleoside metabolic process | Jun-40 | 0.000216212 | 0.001831364 |
| GO:0006686 | sphingomyelin biosynthetic process | Jun-40 | 0.000216212 | 0.001831364 |
| GO:0009068 | aspartate family amino acid catabolic process | Jun-40 | 0.000216212 | 0.001831364 |
| GO:0032000 | positive regulation of fatty acid beta-oxidation | Jun-40 | 0.000216212 | 0.001831364 |
| GO:0034310 | primary alcohol catabolic process | Jun-40 | 0.000216212 | 0.001831364 |
| GO:0035437 | maintenance of protein localization in endoplasmic reticulum | Jun-40 | 0.000216212 | 0.001831364 |
| GO:0090209 | negative regulation of triglyceride metabolic process | Jun-40 | 0.000216212 | 0.001831364 |
| GO:0033692 | cellular polysaccharide biosynthetic process | 15/2440 | 0.000233457 | 0.0019745 |
| GO:0062197 | cellular response to chemical stress | 48/2440 | 0.000248605 | 0.002099492 |
| GO:0006862 | nucleotide transport | Sep-40 | 0.000252481 | 0.002125916 |
| GO:0015740 | C4-dicarboxylate transport | Sep-40 | 0.000252481 | 0.002125916 |
| GO:0031669 | cellular response to nutrient levels | 33/2440 | 0.000259499 | 0.002181788 |
| GO:0043030 | regulation of macrophage activation | 14/2440 | 0.000269051 | 0.002255429 |
| GO:0043470 | regulation of carbohydrate catabolic process | 14/2440 | 0.000269051 | 0.002255429 |
| GO:0006991 | response to sterol depletion | Jul-40 | 0.000280885 | 0.002344274 |
| GO:0009176 | pyrimidine deoxyribonucleoside monophosphate metabolic process | Jul-40 | 0.000280885 | 0.002344274 |
| GO:1901569 | fatty acid derivative catabolic process | Jul-40 | 0.000280885 | 0.002344274 |

| | | | | |
|---|---|---|---|---|
| GO:0034755 | iron ion transmembrane transport | Aug-40 | 0.000282858 | 0.002353843 |
| GO:0150104 | transport across blood-brain barrier | Aug-40 | 0.000282858 | 0.002353843 |
| GO:0031330 | negative regulation of cellular catabolic process | 38/2440 | 0.000284631 | 0.002365137 |
| GO:0051881 | regulation of mitochondrial membrane potential | 18/2440 | 0.000311799 | 0.002587115 |
| GO:0046467 | membrane lipid biosynthetic process | 24/2440 | 0.00031558 | 0.002614676 |
| GO:0097035 | regulation of membrane lipid distribution | 14/2440 | 0.000325559 | 0.002689526 |
| GO:1902475 | L-alpha-amino acid transmembrane transport | 14/2440 | 0.000325559 | 0.002689526 |
| GO:1901653 | cellular response to peptide | 45/2440 | 0.000330732 | 0.002728295 |
| GO:0042417 | dopamine metabolic process | Dec-40 | 0.000336881 | 0.002775005 |
| GO:0009100 | glycoprotein metabolic process | 47/2440 | 0.000352687 | 0.002901005 |
| GO:0017004 | cytochrome complex assembly | Oct-40 | 0.000366199 | 0.003003463 |
| GO:0046475 | glycerophospholipid catabolic process | Oct-40 | 0.000366199 | 0.003003463 |
| GO:0009084 | glutamine family amino acid biosynthetic process | Jun-40 | 0.000372745 | 0.003043998 |
| GO:0034982 | mitochondrial protein processing | Jun-40 | 0.000372745 | 0.003043998 |
| GO:0035999 | tetrahydrofolate interconversion | Jun-40 | 0.000372745 | 0.003043998 |
| GO:0003018 | vascular process in circulatory system | 36/2440 | 0.000376318 | 0.003068769 |
| GO:1900542 | regulation of purine nucleotide metabolic process | 17/2440 | 0.000379536 | 0.003090583 |
| GO:0022600 | digestive system process | 22/2440 | 0.000388598 | 0.003159854 |

| | | | | |
|---|---|---|---|---|
| GO:0010232 | vascular transport | Aug-40 | 0.000401656 | 0.003252102 |
| GO:0019835 | cytolysis | Aug-40 | 0.000401656 | 0.003252102 |
| GO:0090201 | negative regulation of release of cytochrome c from mitochondria | Aug-40 | 0.000401656 | 0.003252102 |
| GO:0000266 | mitochondrial fission | Dec-40 | 0.000417863 | 0.003378517 |
| GO:0006865 | amino acid transport | 27/2440 | 0.000424474 | 0.003420895 |
| GO:0002455 | humoral immune response mediated by circulating immunoglobulin | 30/2440 | 0.000424627 | 0.003420895 |
| GO:0042953 | lipoprotein transport | Jul-40 | 0.000426109 | 0.003420895 |
| GO:1901570 | fatty acid derivative biosynthetic process | Jul-40 | 0.000426109 | 0.003420895 |
| GO:1901663 | quinone biosynthetic process | Jul-40 | 0.000426109 | 0.003420895 |
| GO:0006140 | regulation of nucleotide metabolic process | 17/2440 | 0.000441537 | 0.003539759 |
| GO:0006497 | protein lipidation | 19/2440 | 0.000447597 | 0.003583291 |
| GO:2001234 | negative regulation of apoptotic signaling pathway | 37/2440 | 0.00045189 | 0.003612584 |
| GO:1902932 | positive regulation of alcohol biosynthetic process | Sep-40 | 0.000461921 | 0.003687591 |
| GO:0050673 | epithelial cell proliferation | 62/2440 | 0.000463401 | 0.003694226 |
| GO:0006801 | superoxide metabolic process | 15/2440 | 0.000470485 | 0.003745454 |
| GO:0000959 | mitochondrial RNA metabolic process | Dec-40 | 0.000514665 | 0.004091446 |
| GO:0003044 | regulation of systemic arterial blood pressure mediated by a chemical signal | 13/2440 | 0.000548084 | 0.004351035 |
| GO:1900407 | regulation of cellular response to oxidative stress | 18/2440 | 0.000553199 | 0.004385525 |

| | | | | |
|---|---|---|---|---|
| GO:0002698 | negative regulation of immune effector process | 23/2440 | 0.000556061 | 0.004402086 |
| GO:0034383 | low-density lipoprotein particle clearance | Aug-40 | 0.000558003 | 0.004405207 |
| GO:0098869 | cellular oxidant detoxification | Aug-40 | 0.000558003 | 0.004405207 |
| GO:0072595 | maintenance of protein localization in organelle | Nov-40 | 0.000569533 | 0.004490001 |
| GO:0031331 | positive regulation of cellular catabolic process | 60/2440 | 0.00057759 | 0.004547223 |
| GO:0006067 | ethanol metabolic process | Jun-40 | 0.000605616 | 0.004715685 |
| GO:0006465 | signal peptide processing | Jun-40 | 0.000605616 | 0.004715685 |
| GO:0006555 | methionine metabolic process | Jun-40 | 0.000605616 | 0.004715685 |
| GO:0008228 | opsonization | Jun-40 | 0.000605616 | 0.004715685 |
| GO:0045916 | negative regulation of complement activation | Jun-40 | 0.000605616 | 0.004715685 |
| GO:0046073 | dTMP metabolic process | Jun-40 | 0.000605616 | 0.004715685 |
| GO:0070207 | protein homotrimerization | Jun-40 | 0.000605616 | 0.004715685 |
| GO:2001140 | positive regulation of phospholipid transport | Jun-40 | 0.000605616 | 0.004715685 |
| GO:0097242 | amyloid-beta clearance | Oct-40 | 0.000607039 | 0.00472031 |
| GO:0032369 | negative regulation of lipid transport | Sep-40 | 0.000610671 | 0.00473561 |
| GO:0043032 | positive regulation of macrophage activation | Sep-40 | 0.000610671 | 0.00473561 |
| GO:0050708 | regulation of protein secretion | 43/2440 | 0.000623645 | 0.00480837 |
| GO:0044539 | long-chain fatty acid import into cell | Jul-40 | 0.000625541 | 0.00480837 |

| | | | | |
|---|---|---|---|---|
| GO:0044872 | lipoprotein localization | Jul-40 | 0.000625541 | 0.00480837 |
| GO:0045821 | positive regulation of glycolytic process | Jul-40 | 0.000625541 | 0.00480837 |
| GO:0046485 | ether lipid metabolic process | Jul-40 | 0.000625541 | 0.00480837 |
| GO:0050746 | regulation of lipoprotein metabolic process | Jul-40 | 0.000625541 | 0.00480837 |
| GO:0061912 | selective autophagy | 16/2440 | 0.000625967 | 0.00480837 |
| GO:2001233 | regulation of apoptotic signaling pathway | 54/2440 | 0.000635641 | 0.004876103 |
| GO:0045088 | regulation of innate immune response | 36/2440 | 0.000658555 | 0.005045083 |
| GO:0042446 | hormone biosynthetic process | 14/2440 | 0.000664332 | 0.005070264 |
| GO:0046173 | polyol biosynthetic process | 14/2440 | 0.000664332 | 0.005070264 |
| GO:1902882 | regulation of response to oxidative stress | 19/2440 | 0.000664515 | 0.005070264 |
| GO:0071453 | cellular response to oxygen levels | 25/2440 | 0.000669305 | 0.005099981 |
| GO:0007029 | endoplasmic reticulum organization | 18/2440 | 0.000725074 | 0.00551754 |
| GO:1903034 | regulation of response to wounding | 28/2440 | 0.00073347 | 0.005573981 |
| GO:0051224 | negative regulation of protein transport | 24/2440 | 0.000758744 | 0.005685141 |
| GO:0006196 | AMP catabolic process | May-40 | 0.00076 | 0.005685141 |
| GO:0006591 | ornithine metabolic process | May-40 | 0.00076 | 0.005685141 |
| GO:0007597 | blood coagulation, intrinsic pathway | May-40 | 0.00076 | 0.005685141 |
| GO:0009131 | pyrimidine nucleoside monophosphate catabolic process | May-40 | 0.00076 | 0.005685141 |

| | | | | |
|---|---|---|---|---|
| GO:0009178 | pyrimidine deoxyribonucleoside monophosphate catabolic process | May-40 | 0.00076 | 0.005685141 |
| GO:0048840 | otolith development | May-40 | 0.00076 | 0.005685141 |
| GO:0051775 | response to redox state | May-40 | 0.00076 | 0.005685141 |
| GO:0150172 | regulation of phosphatidylcholine metabolic process | May-40 | 0.00076 | 0.005685141 |
| GO:0010884 | positive regulation of lipid storage | Aug-40 | 0.000760083 | 0.005685141 |
| GO:0015813 | L-glutamate transmembrane transport | Aug-40 | 0.000760083 | 0.005685141 |
| GO:0046037 | GMP metabolic process | Aug-40 | 0.000760083 | 0.005685141 |
| GO:0070873 | regulation of glycogen metabolic process | Oct-40 | 0.000769351 | 0.005746907 |
| GO:0044070 | regulation of anion transport | 21/2440 | 0.000780394 | 0.005821759 |
| GO:0070059 | intrinsic apoptotic signaling pathway in response to endoplasmic reticulum stress | 14/2440 | 0.000784977 | 0.005848281 |
| GO:0051353 | positive regulation of oxidoreductase activity | 13/2440 | 0.000788518 | 0.00586699 |
| GO:0000422 | autophagy of mitochondrion | 16/2440 | 0.000840479 | 0.006237298 |
| GO:0061726 | mitochondrion disassembly | 16/2440 | 0.000840479 | 0.006237298 |
| GO:1901998 | toxin transport | Nov-40 | 0.000871804 | 0.006461338 |
| GO:0000271 | polysaccharide biosynthetic process | 15/2440 | 0.000888566 | 0.006577003 |
| GO:0006089 | lactate metabolic process | Jul-40 | 0.000892391 | 0.006588181 |
| GO:0140354 | lipid import into cell | Jul-40 | 0.000892391 | 0.006588181 |
| GO:0000097 | sulfur amino acid biosynthetic process | Jun-40 | 0.000937245 | 0.006839482 |

| | | | | |
|---|---|---|---|---|
| GO:0006577 | amino-acid betaine metabolic process | Jun-40 | 0.000937245 | 0.006839482 |
| GO:0006596 | polyamine biosynthetic process | Jun-40 | 0.000937245 | 0.006839482 |
| GO:0036498 | IRE1-mediated unfolded protein response | Jun-40 | 0.000937245 | 0.006839482 |
| GO:0046322 | negative regulation of fatty acid oxidation | Jun-40 | 0.000937245 | 0.006839482 |
| GO:0060330 | regulation of response to interferon-gamma | Jun-40 | 0.000937245 | 0.006839482 |
| GO:0060334 | regulation of interferon-gamma-mediated signaling pathway | Jun-40 | 0.000937245 | 0.006839482 |
| GO:0061709 | reticulophagy | Jun-40 | 0.000937245 | 0.006839482 |
| GO:2001138 | regulation of phospholipid transport | Jun-40 | 0.000937245 | 0.006839482 |
| GO:2001242 | regulation of intrinsic apoptotic signaling pathway | 28/2440 | 0.000962846 | 0.00701668 |
| GO:0032692 | negative regulation of interleukin-1 production | Oct-40 | 0.000965722 | 0.00701668 |
| GO:0000045 | autophagosome assembly | 19/2440 | 0.000966458 | 0.00701668 |
| GO:0032479 | regulation of type I interferon production | 19/2440 | 0.000966458 | 0.00701668 |
| GO:1905037 | autophagosome organization | 20/2440 | 0.000978511 | 0.007095142 |
| GO:0015833 | peptide transport | 44/2440 | 0.000981705 | 0.007109245 |
| GO:0050687 | negative regulation of defense response to virus | Aug-40 | 0.001017039 | 0.007337121 |
| GO:0070129 | regulation of mitochondrial translation | Aug-40 | 0.001017039 | 0.007337121 |
| GO:0097164 | ammonium ion metabolic process | Aug-40 | 0.001017039 | 0.007337121 |
| GO:0006734 | NADH metabolic process | Sep-40 | 0.001025544 | 0.00738911 |

| | | | | |
|---|---|---|---|---|
| GO:1902600 | proton transmembrane transport | 17/2440 | 0.001028361 | 0.007400036 |
| GO:0098586 | cellular response to virus | 15/2440 | 0.001032144 | 0.007417886 |
| GO:0042594 | response to starvation | 29/2440 | 0.001088716 | 0.00781459 |
| GO:0001678 | cellular glucose homeostasis | 27/2440 | 0.001104759 | 0.00791976 |
| GO:0006900 | vesicle budding from membrane | 13/2440 | 0.001111834 | 0.007960451 |
| GO:0007162 | negative regulation of cell adhesion | 43/2440 | 0.001132046 | 0.008094979 |
| GO:0016486 | peptide hormone processing | Jul-40 | 0.001241393 | 0.00883251 |
| GO:0030194 | positive regulation of blood coagulation | Jul-40 | 0.001241393 | 0.00883251 |
| GO:0060333 | interferon-gamma-mediated signaling pathway | Jul-40 | 0.001241393 | 0.00883251 |
| GO:1900048 | positive regulation of hemostasis | Jul-40 | 0.001241393 | 0.00883251 |
| GO:1904950 | negative regulation of establishment of protein localization | 24/2440 | 0.001252695 | 0.008901794 |
| GO:0090087 | regulation of peptide transport | 35/2440 | 0.001285422 | 0.009122967 |
| GO:1903037 | regulation of leukocyte cell-cell adhesion | 46/2440 | 0.001291785 | 0.009154327 |
| GO:0006068 | ethanol catabolic process | May-40 | 0.001296273 | 0.009154327 |
| GO:1905288 | vascular associated smooth muscle cell apoptotic process | May-40 | 0.001296273 | 0.009154327 |
| GO:1905459 | regulation of vascular associated smooth muscle cell apoptotic process | May-40 | 0.001296273 | 0.009154327 |
| GO:0050999 | regulation of nitric-oxide synthase activity | Sep-40 | 0.001305288 | 0.009195173 |
| GO:0120178 | steroid hormone biosynthetic process | Sep-40 | 0.001305288 | 0.009195173 |

| | | | | |
|---|---|---|---|---|
| GO:0006515 | protein quality control for misfolded or incompletely synthesized proteins | Aug-40 | 0.001338959 | 0.00940908 |
| GO:0031100 | animal organ regeneration | Aug-40 | 0.001338959 | 0.00940908 |
| GO:0090276 | regulation of peptide hormone secretion | 34/2440 | 0.001379471 | 0.009681809 |
| GO:0046500 | S-adenosylmethionine metabolic process | Jun-40 | 0.001392645 | 0.009762236 |
| GO:0043161 | proteasome-mediated ubiquitin-dependent protein catabolic process | 55/2440 | 0.0014227 | 0.009960653 |

**GO terms associated with 3926 Rep_late_depleted_vs_Hep genes**

| ID | Description | GeneRatio | pvalue | p.adjust |
|---|---|---|---|---|
| GO:0006631 | fatty acid metabolic process | 245/3028 | 1.90E-121 | 1.11E-117 |
| GO:0044282 | small molecule catabolic process | 200/3028 | 9.91E-109 | 2.89E-105 |
| GO:0016054 | organic acid catabolic process | 156/3028 | 5.36E-101 | 1.04E-97 |
| GO:0046395 | carboxylic acid catabolic process | 154/3028 | 2.64E-99 | 3.86E-96 |
| GO:0006520 | cellular amino acid metabolic process | 151/3028 | 8.89E-80 | 1.04E-76 |
| GO:0016053 | organic acid biosynthetic process | 160/3028 | 8.53E-78 | 8.30E-75 |
| GO:0046394 | carboxylic acid biosynthetic process | 159/3028 | 4.59E-77 | 3.83E-74 |
| GO:0006790 | sulfur compound metabolic process | 165/3028 | 5.46E-74 | 3.99E-71 |
| GO:0006091 | generation of precursor metabolites and energy | 192/3028 | 1.48E-73 | 9.62E-71 |
| GO:0008202 | steroid metabolic process | 167/3028 | 2.36E-73 | 1.38E-70 |
| GO:1901605 | alpha-amino acid metabolic process | 113/3028 | 2.55E-63 | 1.35E-60 |
| GO:0015980 | energy derivation by oxidation of organic compounds | 146/3028 | 2.88E-60 | 1.40E-57 |
| GO:0072330 | monocarboxylic acid biosynthetic process | 116/3028 | 1.40E-58 | 6.30E-56 |
| GO:0006753 | nucleoside phosphate metabolic process | 177/3028 | 5.91E-58 | 2.46E-55 |
| GO:0009117 | nucleotide metabolic process | 175/3028 | 8.68E-58 | 3.38E-55 |
| GO:0016042 | lipid catabolic process | 147/3028 | 2.02E-57 | 7.30E-55 |

| | | | | |
|---|---|---|---|---|
| GO:0072521 | purine-containing compound metabolic process | 162/3028 | 2.13E-57 | 7.30E-55 |
| GO:0072329 | monocarboxylic acid catabolic process | 87/3028 | 2.65E-57 | 8.61E-55 |
| GO:0006066 | alcohol metabolic process | 150/3028 | 1.15E-56 | 3.53E-54 |
| GO:0045333 | cellular respiration | 118/3028 | 3.15E-56 | 9.19E-54 |
| GO:0009060 | aerobic respiration | 99/3028 | 3.01E-54 | 8.37E-52 |
| GO:0044242 | cellular lipid catabolic process | 116/3028 | 5.58E-54 | 1.48E-51 |
| GO:0009259 | ribonucleotide metabolic process | 151/3028 | 8.37E-54 | 2.13E-51 |
| GO:0019693 | ribose phosphate metabolic process | 153/3028 | 3.44E-53 | 8.38E-51 |
| GO:0009150 | purine ribonucleotide metabolic process | 145/3028 | 1.50E-52 | 3.50E-50 |
| GO:0016125 | sterol metabolic process | 94/3028 | 1.63E-52 | 3.67E-50 |
| GO:0006163 | purine nucleotide metabolic process | 150/3028 | 4.29E-52 | 9.28E-50 |
| GO:0009063 | cellular amino acid catabolic process | 71/3028 | 2.43E-49 | 5.06E-47 |
| GO:0009062 | fatty acid catabolic process | 73/3028 | 5.92E-49 | 1.19E-46 |
| GO:1902652 | secondary alcohol metabolic process | 92/3028 | 6.59E-49 | 1.28E-46 |
| GO:0042180 | cellular ketone metabolic process | 113/3028 | 3.99E-48 | 7.52E-46 |
| GO:0008203 | cholesterol metabolic process | 87/3028 | 8.15E-48 | 1.49E-45 |
| GO:0006805 | xenobiotic metabolic process | 79/3028 | 2.31E-46 | 4.08E-44 |
| GO:0046034 | ATP metabolic process | 116/3028 | 8.31E-45 | 1.43E-42 |

| | | | | |
|---|---|---|---|---|
| GO:0019216 | regulation of lipid metabolic process | 141/3028 | 3.58E-43 | 5.97E-41 |
| GO:0033865 | nucleoside bisphosphate metabolic process | 71/3028 | 2.28E-42 | 3.50E-40 |
| GO:0033875 | ribonucleoside bisphosphate metabolic process | 71/3028 | 2.28E-42 | 3.50E-40 |
| GO:0034032 | purine nucleoside bisphosphate metabolic process | 71/3028 | 2.28E-42 | 3.50E-40 |
| GO:0033108 | mitochondrial respiratory chain complex assembly | 65/3028 | 3.11E-42 | 4.66E-40 |
| GO:0006637 | acyl-CoA metabolic process | 63/3028 | 5.29E-42 | 7.53E-40 |
| GO:0035383 | thioester metabolic process | 63/3028 | 5.29E-42 | 7.53E-40 |
| GO:0071466 | cellular response to xenobiotic stimulus | 91/3028 | 1.81E-41 | 2.52E-39 |
| GO:0006635 | fatty acid beta-oxidation | 58/3028 | 1.98E-41 | 2.69E-39 |
| GO:0006119 | oxidative phosphorylation | 72/3028 | 3.13E-41 | 4.15E-39 |
| GO:0019395 | fatty acid oxidation | 71/3028 | 3.37E-41 | 4.38E-39 |
| GO:0010876 | lipid localization | 152/3028 | 3.78E-41 | 4.80E-39 |
| GO:0043648 | dicarboxylic acid metabolic process | 61/3028 | 2.53E-40 | 3.15E-38 |
| GO:1901606 | alpha-amino acid catabolic process | 57/3028 | 6.08E-39 | 7.39E-37 |
| GO:0009410 | response to xenobiotic stimulus | 122/3028 | 2.35E-38 | 2.80E-36 |
| GO:0034440 | lipid oxidation | 71/3028 | 2.40E-38 | 2.80E-36 |
| GO:0006633 | fatty acid biosynthetic process | 80/3028 | 5.28E-38 | 6.05E-36 |
| GO:0062012 | regulation of small molecule metabolic process | 131/3028 | 8.45E-38 | 9.49E-36 |

| | | | | |
|---|---|---|---|---|
| GO:0090407 | organophosphate biosynthetic process | 152/3028 | 5.25E-37 | 5.78E-35 |
| GO:0055088 | lipid homeostasis | 85/3028 | 7.28E-37 | 7.88E-35 |
| GO:0006575 | cellular modified amino acid metabolic process | 91/3028 | 4.20E-36 | 4.39E-34 |
| GO:0046890 | regulation of lipid biosynthetic process | 91/3028 | 4.20E-36 | 4.39E-34 |
| GO:0006694 | steroid biosynthetic process | 82/3028 | 8.47E-36 | 8.68E-34 |
| GO:0006869 | lipid transport | 131/3028 | 3.37E-35 | 3.39E-33 |
| GO:0120254 | olefinic compound metabolic process | 79/3028 | 9.80E-35 | 9.70E-33 |
| GO:0030258 | lipid modification | 92/3028 | 2.63E-34 | 2.56E-32 |
| GO:0033559 | unsaturated fatty acid metabolic process | 70/3028 | 1.16E-33 | 1.11E-31 |
| GO:0016126 | sterol biosynthetic process | 47/3028 | 4.94E-33 | 4.66E-31 |
| GO:0006695 | cholesterol biosynthetic process | 44/3028 | 4.09E-32 | 3.73E-30 |
| GO:1902653 | secondary alcohol biosynthetic process | 44/3028 | 4.09E-32 | 3.73E-30 |
| GO:0022900 | electron transport chain | 60/3028 | 4.86E-32 | 4.36E-30 |
| GO:0140053 | mitochondrial gene expression | 61/3028 | 1.61E-31 | 1.42E-29 |
| GO:0001676 | long-chain fatty acid metabolic process | 65/3028 | 3.22E-31 | 2.81E-29 |
| GO:0006641 | triglyceride metabolic process | 61/3028 | 1.28E-30 | 1.10E-28 |
| GO:0045834 | positive regulation of lipid metabolic process | 80/3028 | 1.46E-30 | 1.24E-28 |
| GO:0010257 | NADH dehydrogenase complex assembly | 43/3028 | 3.87E-30 | 3.18E-28 |

| | | | | |
|---|---|---|---|---|
| GO:0032981 | mitochondrial respiratory chain complex I assembly | 43/3028 | 3.87E-30 | 3.18E-28 |
| GO:0022904 | respiratory electron transport chain | 55/3028 | 6.36E-30 | 5.16E-28 |
| GO:0010565 | regulation of cellular ketone metabolic process | 71/3028 | 1.24E-29 | 9.91E-28 |
| GO:1901617 | organic hydroxy compound biosynthetic process | 89/3028 | 8.23E-29 | 6.49E-27 |
| GO:0006639 | acylglycerol metabolic process | 68/3028 | 2.45E-28 | 1.91E-26 |
| GO:0006638 | neutral lipid metabolic process | 68/3028 | 6.74E-28 | 5.18E-26 |
| GO:0042775 | mitochondrial ATP synthesis coupled electron transport | 44/3028 | 7.75E-27 | 5.88E-25 |
| GO:0006690 | icosanoid metabolic process | 65/3028 | 8.02E-27 | 6.01E-25 |
| GO:0042773 | ATP synthesis coupled electron transport | 44/3028 | 4.65E-26 | 3.44E-24 |
| GO:0019646 | aerobic electron transport chain | 38/3028 | 2.05E-25 | 1.49E-23 |
| GO:0046165 | alcohol biosynthetic process | 63/3028 | 2.54E-25 | 1.83E-23 |
| GO:0009165 | nucleotide biosynthetic process | 80/3028 | 2.67E-25 | 1.90E-23 |
| GO:0046486 | glycerolipid metabolic process | 113/3028 | 2.92E-25 | 2.05E-23 |
| GO:1901293 | nucleoside phosphate biosynthetic process | 81/3028 | 3.04E-25 | 2.11E-23 |
| GO:0046889 | positive regulation of lipid biosynthetic process | 54/3028 | 4.07E-25 | 2.80E-23 |
| GO:0019369 | arachidonic acid metabolic process | 45/3028 | 2.22E-24 | 1.51E-22 |
| GO:0055092 | sterol homeostasis | 51/3028 | 2.45E-24 | 1.64E-22 |
| GO:0015711 | organic anion transport | 106/3028 | 3.00E-24 | 1.99E-22 |

| | | | | |
|---|---|---|---|---|
| GO:0042632 | cholesterol homeostasis | 50/3028 | 1.22E-23 | 8.01E-22 |
| GO:0019217 | regulation of fatty acid metabolic process | 51/3028 | 1.47E-23 | 9.51E-22 |
| GO:0006749 | glutathione metabolic process | 45/3028 | 1.93E-23 | 1.24E-21 |
| GO:0062013 | positive regulation of small molecule metabolic process | 67/3028 | 2.13E-23 | 1.35E-21 |
| GO:0006084 | acetyl-CoA metabolic process | 29/3028 | 3.21E-23 | 2.02E-21 |
| GO:0034976 | response to endoplasmic reticulum stress | 85/3028 | 8.86E-23 | 5.50E-21 |
| GO:1905952 | regulation of lipid localization | 65/3028 | 4.65E-22 | 2.86E-20 |
| GO:0032543 | mitochondrial translation | 42/3028 | 7.49E-22 | 4.56E-20 |
| GO:0009152 | purine ribonucleotide biosynthetic process | 61/3028 | 8.95E-22 | 5.39E-20 |
| GO:0015850 | organic hydroxy compound transport | 83/3028 | 9.11E-22 | 5.43E-20 |
| GO:0006720 | isoprenoid metabolic process | 45/3028 | 9.78E-22 | 5.77E-20 |
| GO:0072522 | purine-containing compound biosynthetic process | 66/3028 | 1.71E-21 | 9.98E-20 |
| GO:0046390 | ribose phosphate biosynthetic process | 65/3028 | 3.03E-21 | 1.75E-19 |
| GO:0009260 | ribonucleotide biosynthetic process | 63/3028 | 3.09E-21 | 1.77E-19 |
| GO:0050994 | regulation of lipid catabolic process | 39/3028 | 3.46E-21 | 1.96E-19 |
| GO:0006164 | purine nucleotide biosynthetic process | 64/3028 | 3.71E-21 | 2.08E-19 |
| GO:0090181 | regulation of cholesterol metabolic process | 30/3028 | 3.83E-21 | 2.13E-19 |
| GO:0009636 | response to toxic substance | 53/3028 | 7.72E-21 | 4.25E-19 |

| | | | | |
|---|---|---|---|---|
| GO:0046942 | carboxylic acid transport | 88/3028 | 8.30E-21 | 4.53E-19 |
| GO:0009069 | serine family amino acid metabolic process | 28/3028 | 1.72E-20 | 9.30E-19 |
| GO:0015849 | organic acid transport | 93/3028 | 2.96E-20 | 1.59E-18 |
| GO:0042445 | hormone metabolic process | 74/3028 | 3.08E-20 | 1.64E-18 |
| GO:0042168 | heme metabolic process | 27/3028 | 1.25E-19 | 6.57E-18 |
| GO:0015918 | sterol transport | 41/3028 | 1.78E-19 | 9.28E-18 |
| GO:0033013 | tetrapyrrole metabolic process | 34/3028 | 2.10E-19 | 1.09E-17 |
| GO:0090207 | regulation of triglyceride metabolic process | 32/3028 | 2.71E-19 | 1.39E-17 |
| GO:0030301 | cholesterol transport | 39/3028 | 3.90E-19 | 1.98E-17 |
| GO:0006644 | phospholipid metabolic process | 99/3028 | 4.71E-19 | 2.37E-17 |
| GO:0006778 | porphyrin-containing compound metabolic process | 29/3028 | 6.64E-19 | 3.31E-17 |
| GO:0006820 | anion transport | 120/3028 | 1.70E-18 | 8.40E-17 |
| GO:0098754 | detoxification | 33/3028 | 2.57E-18 | 1.26E-16 |
| GO:0097006 | regulation of plasma lipoprotein particle levels | 36/3028 | 4.15E-18 | 2.02E-16 |
| GO:0007031 | peroxisome organization | 26/3028 | 8.82E-18 | 4.23E-16 |
| GO:0006605 | protein targeting | 81/3028 | 8.83E-18 | 4.23E-16 |
| GO:0008206 | bile acid metabolic process | 28/3028 | 1.11E-17 | 5.27E-16 |
| GO:0034754 | cellular hormone metabolic process | 49/3028 | 1.18E-17 | 5.58E-16 |

| | | | | |
|---|---|---|---|---|
| GO:0009064 | glutamine family amino acid metabolic process | 34/3028 | 1.59E-17 | 7.41E-16 |
| GO:0006721 | terpenoid metabolic process | 35/3028 | 2.11E-17 | 9.80E-16 |
| GO:0071827 | plasma lipoprotein particle organization | 27/3028 | 2.75E-17 | 1.27E-15 |
| GO:0019218 | regulation of steroid metabolic process | 47/3028 | 5.38E-17 | 2.46E-15 |
| GO:0042178 | xenobiotic catabolic process | 22/3028 | 5.58E-17 | 2.53E-15 |
| GO:0006081 | cellular aldehyde metabolic process | 34/3028 | 5.74E-17 | 2.56E-15 |
| GO:0042304 | regulation of fatty acid biosynthetic process | 29/3028 | 5.75E-17 | 2.56E-15 |
| GO:1990542 | mitochondrial transmembrane transport | 38/3028 | 5.91E-17 | 2.62E-15 |
| GO:0006766 | vitamin metabolic process | 39/3028 | 6.21E-17 | 2.73E-15 |
| GO:1990748 | cellular detoxification | 27/3028 | 6.94E-17 | 3.02E-15 |
| GO:0008652 | cellular amino acid biosynthetic process | 33/3028 | 8.25E-17 | 3.57E-15 |
| GO:0044272 | sulfur compound biosynthetic process | 41/3028 | 9.98E-17 | 4.29E-15 |
| GO:0006120 | mitochondrial electron transport, NADH to ubiquinone | 22/3028 | 2.08E-16 | 8.87E-15 |
| GO:0072593 | reactive oxygen species metabolic process | 72/3028 | 2.30E-16 | 9.73E-15 |
| GO:0006536 | glutamate metabolic process | 21/3028 | 4.20E-16 | 1.77E-14 |
| GO:0055090 | acylglycerol homeostasis | 25/3028 | 4.34E-16 | 1.80E-14 |
| GO:0070328 | triglyceride homeostasis | 25/3028 | 4.34E-16 | 1.80E-14 |
| GO:0051604 | protein maturation | 89/3028 | 4.72E-16 | 1.94E-14 |

| | | | | |
|---|---|---|---|---|
| GO:0071806 | protein transmembrane transport | 31/3028 | 6.05E-16 | 2.47E-14 |
| GO:0006090 | pyruvate metabolic process | 46/3028 | 6.68E-16 | 2.71E-14 |
| GO:0015748 | organophosphate ester transport | 47/3028 | 7.98E-16 | 3.21E-14 |
| GO:0045940 | positive regulation of steroid metabolic process | 27/3028 | 8.79E-16 | 3.49E-14 |
| GO:0071825 | protein-lipid complex subunit organization | 27/3028 | 8.79E-16 | 3.49E-14 |
| GO:0005996 | monosaccharide metabolic process | 76/3028 | 9.20E-16 | 3.63E-14 |
| GO:0097237 | cellular response to toxic substance | 28/3028 | 1.51E-15 | 5.94E-14 |
| GO:0009144 | purine nucleoside triphosphate metabolic process | 40/3028 | 1.63E-15 | 6.34E-14 |
| GO:1901607 | alpha-amino acid biosynthetic process | 30/3028 | 3.11E-15 | 1.20E-13 |
| GO:0006783 | heme biosynthetic process | 20/3028 | 3.13E-15 | 1.20E-13 |
| GO:0043467 | regulation of generation of precursor metabolites and energy | 51/3028 | 3.35E-15 | 1.28E-13 |
| GO:0065002 | intracellular protein transmembrane transport | 29/3028 | 4.44E-15 | 1.69E-13 |
| GO:0006699 | bile acid biosynthetic process | 21/3028 | 4.94E-15 | 1.86E-13 |
| GO:0042537 | benzene-containing compound metabolic process | 19/3028 | 6.15E-15 | 2.30E-13 |
| GO:0019915 | lipid storage | 38/3028 | 7.37E-15 | 2.74E-13 |
| GO:1901361 | organic cyclic compound catabolic process | 108/3028 | 8.51E-15 | 3.14E-13 |
| GO:0009145 | purine nucleoside triphosphate biosynthetic process | 33/3028 | 8.62E-15 | 3.17E-13 |
| GO:0006099 | tricarboxylic acid cycle | 22/3028 | 1.69E-14 | 6.19E-13 |

| | | | | |
|---|---|---|---|---|
| GO:0019373 | epoxygenase P450 pathway | 23/3028 | 1.71E-14 | 6.20E-13 |
| GO:0009083 | branched-chain amino acid catabolic process | 17/3028 | 1.86E-14 | 6.72E-13 |
| GO:0009141 | nucleoside triphosphate metabolic process | 43/3028 | 2.34E-14 | 8.39E-13 |
| GO:0035966 | response to topologically incorrect protein | 48/3028 | 3.61E-14 | 1.29E-12 |
| GO:0009206 | purine ribonucleoside triphosphate biosynthetic process | 32/3028 | 3.92E-14 | 1.39E-12 |
| GO:0006779 | porphyrin-containing compound biosynthetic process | 21/3028 | 4.14E-14 | 1.44E-12 |
| GO:0009072 | aromatic amino acid family metabolic process | 21/3028 | 4.14E-14 | 1.44E-12 |
| GO:0033014 | tetrapyrrole biosynthetic process | 21/3028 | 4.14E-14 | 1.44E-12 |
| GO:0031998 | regulation of fatty acid beta-oxidation | 18/3028 | 4.67E-14 | 1.61E-12 |
| GO:0046503 | glycerolipid catabolic process | 32/3028 | 6.49E-14 | 2.23E-12 |
| GO:0006979 | response to oxidative stress | 95/3028 | 6.91E-14 | 2.36E-12 |
| GO:0035384 | thioester biosynthetic process | 19/3028 | 7.73E-14 | 2.61E-12 |
| GO:0071616 | acyl-CoA biosynthetic process | 19/3028 | 7.73E-14 | 2.61E-12 |
| GO:0045923 | positive regulation of fatty acid metabolic process | 26/3028 | 8.84E-14 | 2.97E-12 |
| GO:0034367 | protein-containing complex remodeling | 20/3028 | 9.95E-14 | 3.32E-12 |
| GO:0046434 | organophosphate catabolic process | 50/3028 | 1.21E-13 | 4.02E-12 |
| GO:0043691 | reverse cholesterol transport | 16/3028 | 1.52E-13 | 5.00E-12 |
| GO:0016101 | diterpenoid metabolic process | 29/3028 | 1.60E-13 | 5.25E-12 |

| | | | | |
|---|---|---|---|---|
| GO:0009081 | branched-chain amino acid metabolic process | 18/3028 | 1.68E-13 | 5.49E-12 |
| GO:0044262 | cellular carbohydrate metabolic process | 75/3028 | 1.70E-13 | 5.51E-12 |
| GO:0033344 | cholesterol efflux | 30/3028 | 1.71E-13 | 5.52E-12 |
| GO:0009201 | ribonucleoside triphosphate biosynthetic process | 32/3028 | 1.72E-13 | 5.52E-12 |
| GO:0036503 | ERAD pathway | 40/3028 | 1.87E-13 | 5.98E-12 |
| GO:0019432 | triglyceride biosynthetic process | 23/3028 | 2.03E-13 | 6.46E-12 |
| GO:0034368 | protein-lipid complex remodeling | 19/3028 | 2.35E-13 | 7.38E-12 |
| GO:0034369 | plasma lipoprotein particle remodeling | 19/3028 | 2.35E-13 | 7.38E-12 |
| GO:0033866 | nucleoside bisphosphate biosynthetic process | 22/3028 | 2.43E-13 | 7.50E-12 |
| GO:0034030 | ribonucleoside bisphosphate biosynthetic process | 22/3028 | 2.43E-13 | 7.50E-12 |
| GO:0034033 | purine nucleoside bisphosphate biosynthetic process | 22/3028 | 2.43E-13 | 7.50E-12 |
| GO:0050996 | positive regulation of lipid catabolic process | 21/3028 | 2.68E-13 | 8.23E-12 |
| GO:0090208 | positive regulation of triglyceride metabolic process | 20/3028 | 2.69E-13 | 8.23E-12 |
| GO:0015986 | proton motive force-driven ATP synthesis | 17/3028 | 3.50E-13 | 1.06E-11 |
| GO:0043574 | peroxisomal transport | 17/3028 | 3.50E-13 | 1.06E-11 |
| GO:0009142 | nucleoside triphosphate biosynthetic process | 34/3028 | 3.60E-13 | 1.08E-11 |
| GO:1905039 | carboxylic acid transmembrane transport | 46/3028 | 3.92E-13 | 1.17E-11 |
| GO:0015914 | phospholipid transport | 36/3028 | 4.06E-13 | 1.21E-11 |

| | | | | |
|---|---|---|---|---|
| GO:0019318 | hexose metabolic process | 68/3028 | 4.22E-13 | 1.25E-11 |
| GO:0001523 | retinoid metabolic process | 28/3028 | 4.35E-13 | 1.28E-11 |
| GO:0006754 | ATP biosynthetic process | 28/3028 | 4.35E-13 | 1.28E-11 |
| GO:0009205 | purine ribonucleoside triphosphate metabolic process | 35/3028 | 4.76E-13 | 1.39E-11 |
| GO:0015718 | monocarboxylic acid transport | 43/3028 | 4.88E-13 | 1.42E-11 |
| GO:1903825 | organic acid transmembrane transport | 46/3028 | 5.27E-13 | 1.52E-11 |
| GO:0046461 | neutral lipid catabolic process | 24/3028 | 6.49E-13 | 1.86E-11 |
| GO:0046464 | acylglycerol catabolic process | 24/3028 | 6.49E-13 | 1.86E-11 |
| GO:0042440 | pigment metabolic process | 30/3028 | 7.69E-13 | 2.19E-11 |
| GO:0019439 | aromatic compound catabolic process | 99/3028 | 8.16E-13 | 2.31E-11 |
| GO:0000959 | mitochondrial RNA metabolic process | 25/3028 | 8.65E-13 | 2.44E-11 |
| GO:0007006 | mitochondrial membrane organization | 39/3028 | 9.58E-13 | 2.69E-11 |
| GO:0008654 | phospholipid biosynthetic process | 61/3028 | 1.41E-12 | 3.95E-11 |
| GO:0000038 | very long-chain fatty acid metabolic process | 21/3028 | 1.42E-12 | 3.96E-11 |
| GO:0006839 | mitochondrial transport | 48/3028 | 1.55E-12 | 4.28E-11 |
| GO:1901657 | glycosyl compound metabolic process | 32/3028 | 1.64E-12 | 4.48E-11 |
| GO:0000096 | sulfur amino acid metabolic process | 20/3028 | 1.64E-12 | 4.48E-11 |
| GO:0007007 | inner mitochondrial membrane organization | 20/3028 | 1.64E-12 | 4.48E-11 |

| | | | | |
|---|---|---|---|---|
| GO:0072594 | establishment of protein localization to organelle | 94/3028 | 2.25E-12 | 6.10E-11 |
| GO:0009199 | ribonucleoside triphosphate metabolic process | 35/3028 | 2.26E-12 | 6.12E-11 |
| GO:0042157 | lipoprotein metabolic process | 44/3028 | 3.02E-12 | 8.14E-11 |
| GO:0010883 | regulation of lipid storage | 26/3028 | 3.22E-12 | 8.62E-11 |
| GO:0045833 | negative regulation of lipid metabolic process | 38/3028 | 3.37E-12 | 9.00E-11 |
| GO:0010866 | regulation of triglyceride biosynthetic process | 17/3028 | 3.74E-12 | 9.92E-11 |
| GO:0009991 | response to extracellular stimulus | 90/3028 | 5.03E-12 | 1.33E-10 |
| GO:0006085 | acetyl-CoA biosynthetic process | 15/3028 | 5.24E-12 | 1.38E-10 |
| GO:0032368 | regulation of lipid transport | 41/3028 | 7.28E-12 | 1.90E-10 |
| GO:0016485 | protein processing | 69/3028 | 7.30E-12 | 1.90E-10 |
| GO:1905954 | positive regulation of lipid localization | 36/3028 | 7.67E-12 | 1.99E-10 |
| GO:0034377 | plasma lipoprotein particle assembly | 16/3028 | 8.56E-12 | 2.21E-10 |
| GO:0046460 | neutral lipid biosynthetic process | 25/3028 | 8.76E-12 | 2.24E-10 |
| GO:0046463 | acylglycerol biosynthetic process | 25/3028 | 8.76E-12 | 2.24E-10 |
| GO:0046320 | regulation of fatty acid oxidation | 22/3028 | 9.26E-12 | 2.36E-10 |
| GO:0016052 | carbohydrate catabolic process | 44/3028 | 1.21E-11 | 3.08E-10 |
| GO:0034381 | plasma lipoprotein particle clearance | 21/3028 | 1.29E-11 | 3.25E-10 |
| GO:1901661 | quinone metabolic process | 21/3028 | 1.29E-11 | 3.25E-10 |

| | | | | |
|---|---|---|---|---|
| GO:0034370 | triglyceride-rich lipoprotein particle remodeling | Nov-28 | 1.70E-11 | 4.26E-10 |
| GO:0008299 | isoprenoid biosynthetic process | 20/3028 | 1.72E-11 | 4.30E-10 |
| GO:0045540 | regulation of cholesterol biosynthetic process | 15/3028 | 1.89E-11 | 4.68E-10 |
| GO:0106118 | regulation of sterol biosynthetic process | 15/3028 | 1.89E-11 | 4.68E-10 |
| GO:1901136 | carbohydrate derivative catabolic process | 45/3028 | 1.94E-11 | 4.79E-10 |
| GO:0009071 | serine family amino acid catabolic process | Dec-28 | 2.09E-11 | 5.13E-10 |
| GO:0019433 | triglyceride catabolic process | 19/3028 | 2.18E-11 | 5.29E-10 |
| GO:0032371 | regulation of sterol transport | 19/3028 | 2.18E-11 | 5.29E-10 |
| GO:0032374 | regulation of cholesterol transport | 19/3028 | 2.18E-11 | 5.29E-10 |
| GO:0006006 | glucose metabolic process | 59/3028 | 2.66E-11 | 6.41E-10 |
| GO:0006986 | response to unfolded protein | 39/3028 | 3.15E-11 | 7.57E-10 |
| GO:0006625 | protein targeting to peroxisome | 14/3028 | 3.96E-11 | 9.32E-10 |
| GO:0051004 | regulation of lipoprotein lipase activity | 14/3028 | 3.96E-11 | 9.32E-10 |
| GO:0072662 | protein localization to peroxisome | 14/3028 | 3.96E-11 | 9.32E-10 |
| GO:0072663 | establishment of protein localization to peroxisome | 14/3028 | 3.96E-11 | 9.32E-10 |
| GO:0090205 | positive regulation of cholesterol metabolic process | 14/3028 | 3.96E-11 | 9.32E-10 |
| GO:0034308 | primary alcohol metabolic process | 31/3028 | 4.67E-11 | 1.10E-09 |
| GO:0043434 | response to peptide hormone | 80/3028 | 5.76E-11 | 1.34E-09 |

| | | | | |
|---|---|---|---|---|
| GO:0009116 | nucleoside metabolic process | 25/3028 | 6.71E-11 | 1.56E-09 |
| GO:0006544 | glycine metabolic process | 13/3028 | 7.68E-11 | 1.77E-09 |
| GO:0009074 | aromatic amino acid family catabolic process | 13/3028 | 7.68E-11 | 1.77E-09 |
| GO:0051881 | regulation of mitochondrial membrane potential | 32/3028 | 7.76E-11 | 1.79E-09 |
| GO:0017004 | cytochrome complex assembly | 19/3028 | 9.47E-11 | 2.17E-09 |
| GO:0006888 | endoplasmic reticulum to Golgi vesicle-mediated transport | 38/3028 | 1.01E-10 | 2.30E-09 |
| GO:0009262 | deoxyribonucleotide metabolic process | 23/3028 | 1.10E-10 | 2.50E-09 |
| GO:2000377 | regulation of reactive oxygen species metabolic process | 46/3028 | 1.25E-10 | 2.84E-09 |
| GO:0062014 | negative regulation of small molecule metabolic process | 34/3028 | 1.28E-10 | 2.89E-09 |
| GO:0015919 | peroxisomal membrane transport | 14/3028 | 1.36E-10 | 3.05E-09 |
| GO:0030433 | ubiquitin-dependent ERAD pathway | 31/3028 | 1.38E-10 | 3.10E-09 |
| GO:0070972 | protein localization to endoplasmic reticulum | 29/3028 | 1.46E-10 | 3.25E-09 |
| GO:1901292 | nucleoside phosphate catabolic process | 29/3028 | 1.46E-10 | 3.25E-09 |
| GO:0051346 | negative regulation of hydrolase activity | 80/3028 | 1.55E-10 | 3.44E-09 |
| GO:0032527 | protein exit from endoplasmic reticulum | 21/3028 | 1.59E-10 | 3.50E-09 |
| GO:0034372 | very-low-density lipoprotein particle remodeling | Oct-28 | 1.62E-10 | 3.56E-09 |
| GO:0035634 | response to stilbenoid | 15/3028 | 1.69E-10 | 3.70E-09 |
| GO:0051791 | medium-chain fatty acid metabolic process | 16/3028 | 1.72E-10 | 3.75E-09 |

| | | | | |
|---|---|---|---|---|
| GO:0035967 | cellular response to topologically incorrect protein | 36/3028 | 1.82E-10 | 3.95E-09 |
| GO:0044743 | protein transmembrane import into intracellular organelle | 19/3028 | 1.87E-10 | 4.04E-09 |
| GO:0042407 | cristae formation | 13/3028 | 2.95E-10 | 6.33E-09 |
| GO:0042776 | proton motive force-driven mitochondrial ATP synthesis | 13/3028 | 2.95E-10 | 6.33E-09 |
| GO:0009394 | 2'-deoxyribonucleotide metabolic process | 22/3028 | 2.99E-10 | 6.38E-09 |
| GO:0019692 | deoxyribose phosphate metabolic process | 22/3028 | 2.99E-10 | 6.38E-09 |
| GO:0010038 | response to metal ion | 68/3028 | 3.38E-10 | 7.17E-09 |
| GO:0065005 | protein-lipid complex assembly | 16/3028 | 4.03E-10 | 8.53E-09 |
| GO:0006879 | cellular iron ion homeostasis | 27/3028 | 4.55E-10 | 9.60E-09 |
| GO:0042060 | wound healing | 80/3028 | 4.62E-10 | 9.71E-09 |
| GO:0050810 | regulation of steroid biosynthetic process | 30/3028 | 4.93E-10 | 1.03E-08 |
| GO:0009112 | nucleobase metabolic process | 18/3028 | 5.08E-10 | 1.06E-08 |
| GO:0009166 | nucleotide catabolic process | 26/3028 | 5.40E-10 | 1.12E-08 |
| GO:0031668 | cellular response to extracellular stimulus | 58/3028 | 5.95E-10 | 1.23E-08 |
| GO:0046185 | aldehyde catabolic process | Dec-28 | 5.97E-10 | 1.23E-08 |
| GO:0046470 | phosphatidylcholine metabolic process | 25/3028 | 6.26E-10 | 1.29E-08 |
| GO:0009066 | aspartate family amino acid metabolic process | 19/3028 | 6.60E-10 | 1.35E-08 |
| GO:0098656 | anion transmembrane transport | 52/3028 | 6.71E-10 | 1.37E-08 |

| | | | | |
|---|---|---|---|---|
| GO:0055072 | iron ion homeostasis | 32/3028 | 7.71E-10 | 1.57E-08 |
| GO:0048193 | Golgi vesicle transport | 64/3028 | 8.54E-10 | 1.73E-08 |
| GO:0045723 | positive regulation of fatty acid biosynthetic process | 16/3028 | 8.92E-10 | 1.80E-08 |
| GO:0031667 | response to nutrient levels | 79/3028 | 9.62E-10 | 1.94E-08 |
| GO:0007599 | hemostasis | 47/3028 | 9.74E-10 | 1.95E-08 |
| GO:0050817 | coagulation | 47/3028 | 9.74E-10 | 1.95E-08 |
| GO:1901652 | response to peptide | 86/3028 | 1.03E-09 | 2.05E-08 |
| GO:0010878 | cholesterol storage | 15/3028 | 1.06E-09 | 2.08E-08 |
| GO:0016226 | iron-sulfur cluster assembly | 15/3028 | 1.06E-09 | 2.08E-08 |
| GO:0031163 | metallo-sulfur cluster assembly | 15/3028 | 1.06E-09 | 2.08E-08 |
| GO:0006626 | protein targeting to mitochondrion | 28/3028 | 1.12E-09 | 2.20E-08 |
| GO:0015721 | bile acid and bile salt transport | 17/3028 | 1.39E-09 | 2.71E-08 |
| GO:0045922 | negative regulation of fatty acid metabolic process | 17/3028 | 1.39E-09 | 2.71E-08 |
| GO:0072655 | establishment of protein localization to mitochondrion | 32/3028 | 1.41E-09 | 2.75E-08 |
| GO:0055076 | transition metal ion homeostasis | 40/3028 | 1.51E-09 | 2.94E-08 |
| GO:0033539 | fatty acid beta-oxidation using acyl-CoA dehydrogenase | Oct-28 | 1.61E-09 | 3.12E-08 |
| GO:0006826 | iron ion transport | 24/3028 | 1.65E-09 | 3.18E-08 |
| GO:0007596 | blood coagulation | 46/3028 | 1.69E-09 | 3.26E-08 |

| | | | | |
|---|---|---|---|---|
| GO:0045017 | glycerolipid biosynthetic process | 53/3028 | 1.72E-09 | 3.28E-08 |
| GO:0042908 | xenobiotic transport | 18/3028 | 1.81E-09 | 3.45E-08 |
| GO:0070585 | protein localization to mitochondrion | 33/3028 | 1.87E-09 | 3.56E-08 |
| GO:2000378 | negative regulation of reactive oxygen species metabolic process | 23/3028 | 1.88E-09 | 3.56E-08 |
| GO:0006767 | water-soluble vitamin metabolic process | 19/3028 | 2.09E-09 | 3.95E-08 |
| GO:0090150 | establishment of protein localization to membrane | 60/3028 | 2.15E-09 | 4.06E-08 |
| GO:0006650 | glycerophospholipid metabolic process | 63/3028 | 2.61E-09 | 4.90E-08 |
| GO:0042730 | fibrinolysis | 13/3028 | 2.74E-09 | 5.07E-08 |
| GO:1904478 | regulation of intestinal absorption | 13/3028 | 2.74E-09 | 5.07E-08 |
| GO:0009164 | nucleoside catabolic process | 14/3028 | 2.74E-09 | 5.07E-08 |
| GO:0010885 | regulation of cholesterol storage | 14/3028 | 2.74E-09 | 5.07E-08 |
| GO:1903427 | negative regulation of reactive oxygen species biosynthetic process | 14/3028 | 2.74E-09 | 5.07E-08 |
| GO:0009132 | nucleoside diphosphate metabolic process | 38/3028 | 2.97E-09 | 5.47E-08 |
| GO:0046148 | pigment biosynthetic process | 22/3028 | 3.24E-09 | 5.94E-08 |
| GO:0006739 | NADP metabolic process | 19/3028 | 3.59E-09 | 6.57E-08 |
| GO:0042572 | retinol metabolic process | 16/3028 | 3.78E-09 | 6.90E-08 |
| GO:0009308 | amine metabolic process | 37/3028 | 4.15E-09 | 7.54E-08 |
| GO:0051341 | regulation of oxidoreductase activity | 31/3028 | 4.55E-09 | 8.24E-08 |

| | | | | |
|---|---|---|---|---|
| GO:0046364 | monosaccharide biosynthetic process | 30/3028 | 4.56E-09 | 8.24E-08 |
| GO:0030970 | retrograde protein transport, ER to cytosol | 15/3028 | 5.08E-09 | 9.10E-08 |
| GO:0034656 | nucleobase-containing small molecule catabolic process | 15/3028 | 5.08E-09 | 9.10E-08 |
| GO:1903513 | endoplasmic reticulum to cytosol transport | 15/3028 | 5.08E-09 | 9.10E-08 |
| GO:0046916 | cellular transition metal ion homeostasis | 34/3028 | 5.28E-09 | 9.43E-08 |
| GO:1903578 | regulation of ATP metabolic process | 30/3028 | 6.12E-09 | 1.09E-07 |
| GO:0072525 | pyridine-containing compound biosynthetic process | 14/3028 | 6.33E-09 | 1.12E-07 |
| GO:0006103 | 2-oxoglutarate metabolic process | Dec-28 | 6.63E-09 | 1.17E-07 |
| GO:0010896 | regulation of triglyceride catabolic process | Dec-28 | 6.63E-09 | 1.17E-07 |
| GO:1903008 | organelle disassembly | 34/3028 | 6.82E-09 | 1.20E-07 |
| GO:0051235 | maintenance of location | 72/3028 | 6.83E-09 | 1.20E-07 |
| GO:0006692 | prostanoid metabolic process | 22/3028 | 7.57E-09 | 1.32E-07 |
| GO:0006693 | prostaglandin metabolic process | 22/3028 | 7.57E-09 | 1.32E-07 |
| GO:0006109 | regulation of carbohydrate metabolic process | 49/3028 | 8.39E-09 | 1.46E-07 |
| GO:0006086 | acetyl-CoA biosynthetic process from pyruvate | Oct-28 | 8.76E-09 | 1.51E-07 |
| GO:0006534 | cysteine metabolic process | Oct-28 | 8.76E-09 | 1.51E-07 |
| GO:0032000 | positive regulation of fatty acid beta-oxidation | Oct-28 | 8.76E-09 | 1.51E-07 |
| GO:0072523 | purine-containing compound catabolic process | 20/3028 | 9.46E-09 | 1.62E-07 |

| | | | | |
|---|---|---|---|---|
| GO:0009264 | deoxyribonucleotide catabolic process | 15/3028 | 1.03E-08 | 1.77E-07 |
| GO:0044042 | glucan metabolic process | 27/3028 | 1.06E-08 | 1.81E-07 |
| GO:0008637 | apoptotic mitochondrial changes | 35/3028 | 1.31E-08 | 2.23E-07 |
| GO:0009162 | deoxyribonucleoside monophosphate metabolic process | 16/3028 | 1.36E-08 | 2.29E-07 |
| GO:0042558 | pteridine-containing compound metabolic process | 16/3028 | 1.36E-08 | 2.29E-07 |
| GO:0045454 | cell redox homeostasis | 16/3028 | 1.36E-08 | 2.29E-07 |
| GO:0062125 | regulation of mitochondrial gene expression | 16/3028 | 1.36E-08 | 2.29E-07 |
| GO:0010873 | positive regulation of cholesterol esterification | Sep-28 | 1.40E-08 | 2.34E-07 |
| GO:0110095 | cellular detoxification of aldehyde | Sep-28 | 1.40E-08 | 2.34E-07 |
| GO:1903715 | regulation of aerobic respiration | 17/3028 | 1.57E-08 | 2.62E-07 |
| GO:0050892 | intestinal absorption | 19/3028 | 1.60E-08 | 2.67E-07 |
| GO:0010906 | regulation of glucose metabolic process | 37/3028 | 1.65E-08 | 2.74E-07 |
| GO:0044106 | cellular amine metabolic process | 35/3028 | 1.65E-08 | 2.74E-07 |
| GO:0010867 | positive regulation of triglyceride biosynthetic process | Dec-28 | 1.80E-08 | 2.96E-07 |
| GO:0006457 | protein folding | 43/3028 | 1.93E-08 | 3.18E-07 |
| GO:1905953 | negative regulation of lipid localization | 21/3028 | 1.99E-08 | 3.27E-07 |
| GO:0042573 | retinoic acid metabolic process | 15/3028 | 2.00E-08 | 3.28E-07 |
| GO:0045861 | negative regulation of proteolysis | 73/3028 | 2.19E-08 | 3.58E-07 |

| | | | | |
|---|---|---|---|---|
| GO:0030195 | negative regulation of blood coagulation | 20/3028 | 2.29E-08 | 3.73E-07 |
| GO:0050818 | regulation of coagulation | 25/3028 | 2.48E-08 | 4.03E-07 |
| GO:0072524 | pyridine-containing compound metabolic process | 17/3028 | 2.68E-08 | 4.33E-07 |
| GO:0006730 | one-carbon metabolic process | 18/3028 | 2.69E-08 | 4.34E-07 |
| GO:0044270 | cellular nitrogen compound catabolic process | 83/3028 | 2.69E-08 | 4.34E-07 |
| GO:0046939 | nucleotide phosphorylation | 32/3028 | 2.79E-08 | 4.48E-07 |
| GO:0008535 | respiratory chain complex IV assembly | 14/3028 | 2.81E-08 | 4.49E-07 |
| GO:0044273 | sulfur compound catabolic process | 14/3028 | 2.81E-08 | 4.49E-07 |
| GO:0016051 | carbohydrate biosynthetic process | 48/3028 | 2.83E-08 | 4.51E-07 |
| GO:0046700 | heterocycle catabolic process | 83/3028 | 3.00E-08 | 4.77E-07 |
| GO:0030193 | regulation of blood coagulation | 24/3028 | 3.21E-08 | 5.06E-07 |
| GO:1901616 | organic hydroxy compound catabolic process | 24/3028 | 3.21E-08 | 5.06E-07 |
| GO:0034375 | high-density lipoprotein particle remodeling | Oct-28 | 3.43E-08 | 5.38E-07 |
| GO:0034982 | mitochondrial protein processing | Oct-28 | 3.43E-08 | 5.38E-07 |
| GO:0043649 | dicarboxylic acid catabolic process | Oct-28 | 3.43E-08 | 5.38E-07 |
| GO:1900047 | negative regulation of hemostasis | 20/3028 | 3.49E-08 | 5.45E-07 |
| GO:0005977 | glycogen metabolic process | 26/3028 | 3.54E-08 | 5.50E-07 |
| GO:0006073 | cellular glucan metabolic process | 26/3028 | 3.54E-08 | 5.50E-07 |

| | | | | |
|---|---|---|---|---|
| GO:0006576 | cellular biogenic amine metabolic process | 32/3028 | 3.57E-08 | 5.53E-07 |
| GO:0051055 | negative regulation of lipid biosynthetic process | 22/3028 | 3.59E-08 | 5.55E-07 |
| GO:0010039 | response to iron ion | 13/3028 | 3.70E-08 | 5.70E-07 |
| GO:0008053 | mitochondrial fusion | 15/3028 | 3.75E-08 | 5.76E-07 |
| GO:0009135 | purine nucleoside diphosphate metabolic process | 31/3028 | 3.88E-08 | 5.94E-07 |
| GO:0009179 | purine ribonucleoside diphosphate metabolic process | 31/3028 | 3.88E-08 | 5.94E-07 |
| GO:0009261 | ribonucleotide catabolic process | 19/3028 | 3.96E-08 | 6.05E-07 |
| GO:0009185 | ribonucleoside diphosphate metabolic process | 33/3028 | 4.11E-08 | 6.26E-07 |
| GO:0016236 | macroautophagy | 55/3028 | 4.16E-08 | 6.31E-07 |
| GO:0034248 | regulation of cellular amide metabolic process | 86/3028 | 4.17E-08 | 6.31E-07 |
| GO:0006953 | acute-phase response | 18/3028 | 4.32E-08 | 6.51E-07 |
| GO:1901568 | fatty acid derivative metabolic process | 18/3028 | 4.32E-08 | 6.51E-07 |
| GO:0006525 | arginine metabolic process | Dec-28 | 4.41E-08 | 6.58E-07 |
| GO:0016114 | terpenoid biosynthetic process | Dec-28 | 4.41E-08 | 6.58E-07 |
| GO:0019363 | pyridine nucleotide biosynthetic process | Dec-28 | 4.41E-08 | 6.58E-07 |
| GO:0034433 | steroid esterification | Nov-28 | 4.48E-08 | 6.63E-07 |
| GO:0034434 | sterol esterification | Nov-28 | 4.48E-08 | 6.63E-07 |
| GO:0034435 | cholesterol esterification | Nov-28 | 4.48E-08 | 6.63E-07 |

| | | | | |
|---|---|---|---|---|
| GO:0045717 | negative regulation of fatty acid biosynthetic process | Nov-28 | 4.48E-08 | 6.63E-07 |
| GO:0052548 | regulation of endopeptidase activity | 72/3028 | 4.89E-08 | 7.22E-07 |
| GO:0006165 | nucleoside diphosphate phosphorylation | 31/3028 | 4.98E-08 | 7.32E-07 |
| GO:0050819 | negative regulation of coagulation | 20/3028 | 5.24E-08 | 7.69E-07 |
| GO:0046031 | ADP metabolic process | 30/3028 | 5.39E-08 | 7.89E-07 |
| GO:0046386 | deoxyribose phosphate catabolic process | 14/3028 | 5.50E-08 | 8.04E-07 |
| GO:0034620 | cellular response to unfolded protein | 28/3028 | 6.08E-08 | 8.86E-07 |
| GO:1900046 | regulation of hemostasis | 24/3028 | 6.16E-08 | 8.96E-07 |
| GO:0006914 | autophagy | 87/3028 | 6.25E-08 | 9.04E-07 |
| GO:0061919 | process utilizing autophagic mechanism | 87/3028 | 6.25E-08 | 9.04E-07 |
| GO:0052547 | regulation of peptidase activity | 84/3028 | 6.30E-08 | 9.08E-07 |
| GO:0060191 | regulation of lipase activity | 27/3028 | 6.31E-08 | 9.08E-07 |
| GO:0010823 | negative regulation of mitochondrion organization | 21/3028 | 6.40E-08 | 9.18E-07 |
| GO:0043651 | linoleic acid metabolic process | 15/3028 | 6.76E-08 | 9.66E-07 |
| GO:1901658 | glycosyl compound catabolic process | 15/3028 | 6.76E-08 | 9.66E-07 |
| GO:1903725 | regulation of phospholipid metabolic process | 18/3028 | 6.82E-08 | 9.72E-07 |
| GO:0006558 | L-phenylalanine metabolic process | Sep-28 | 6.98E-08 | 9.75E-07 |
| GO:0019627 | urea metabolic process | Sep-28 | 6.98E-08 | 9.75E-07 |

| | | | | |
|---|---|---|---|---|
| GO:0030300 | regulation of intestinal cholesterol absorption | Sep-28 | 6.98E-08 | 9.75E-07 |
| GO:0034384 | high-density lipoprotein particle clearance | Sep-28 | 6.98E-08 | 9.75E-07 |
| GO:0051006 | positive regulation of lipoprotein lipase activity | Sep-28 | 6.98E-08 | 9.75E-07 |
| GO:0071941 | nitrogen cycle metabolic process | Sep-28 | 6.98E-08 | 9.75E-07 |
| GO:1902221 | erythrose 4-phosphate/phosphoenolpyruvate family amino acid metabolic process | Sep-28 | 6.98E-08 | 9.75E-07 |
| GO:0043161 | proteasome-mediated ubiquitin-dependent protein catabolic process | 82/3028 | 6.98E-08 | 9.75E-07 |
| GO:0071496 | cellular response to external stimulus | 63/3028 | 7.29E-08 | 1.02E-06 |
| GO:0009065 | glutamine family amino acid catabolic process | 13/3028 | 7.68E-08 | 1.07E-06 |
| GO:2000112 | regulation of cellular macromolecule biosynthetic process | 89/3028 | 8.28E-08 | 1.15E-06 |
| GO:0006144 | purine nucleobase metabolic process | Dec-28 | 9.95E-08 | 1.38E-06 |
| GO:1903409 | reactive oxygen species biosynthetic process | 22/3028 | 1.03E-07 | 1.43E-06 |
| GO:0032782 | bile acid secretion | Oct-28 | 1.09E-07 | 1.49E-06 |
| GO:0050667 | homocysteine metabolic process | Oct-28 | 1.09E-07 | 1.49E-06 |
| GO:0051180 | vitamin transport | 17/3028 | 1.17E-07 | 1.60E-06 |
| GO:0010675 | regulation of cellular carbohydrate metabolic process | 41/3028 | 1.18E-07 | 1.62E-06 |
| GO:0032868 | response to insulin | 54/3028 | 1.21E-07 | 1.65E-06 |
| GO:0042278 | purine nucleoside metabolic process | 16/3028 | 1.22E-07 | 1.66E-06 |
| GO:0006112 | energy reserve metabolic process | 27/3028 | 1.43E-07 | 1.94E-06 |

| | | | | |
|---|---|---|---|---|
| GO:0043457 | regulation of cellular respiration | 22/3028 | 1.44E-07 | 1.96E-06 |
| GO:0009123 | nucleoside monophosphate metabolic process | 23/3028 | 1.52E-07 | 2.06E-06 |
| GO:0045047 | protein targeting to ER | 18/3028 | 1.61E-07 | 2.18E-06 |
| GO:0000041 | transition metal ion transport | 31/3028 | 1.63E-07 | 2.19E-06 |
| GO:0006195 | purine nucleotide catabolic process | 17/3028 | 1.83E-07 | 2.46E-06 |
| GO:1902600 | proton transmembrane transport | 27/3028 | 1.85E-07 | 2.48E-06 |
| GO:0010821 | regulation of mitochondrion organization | 37/3028 | 1.94E-07 | 2.59E-06 |
| GO:0019319 | hexose biosynthetic process | 26/3028 | 1.96E-07 | 2.62E-06 |
| GO:0010875 | positive regulation of cholesterol efflux | 15/3028 | 2.01E-07 | 2.67E-06 |
| GO:0046321 | positive regulation of fatty acid oxidation | Dec-28 | 2.10E-07 | 2.79E-06 |
| GO:0010898 | positive regulation of triglyceride catabolic process | Sep-28 | 2.53E-07 | 3.33E-06 |
| GO:0016558 | protein import into peroxisome matrix | Sep-28 | 2.53E-07 | 3.33E-06 |
| GO:0043650 | dicarboxylic acid biosynthetic process | Sep-28 | 2.53E-07 | 3.33E-06 |
| GO:0061365 | positive regulation of triglyceride lipase activity | Sep-28 | 2.53E-07 | 3.33E-06 |
| GO:0031638 | zymogen activation | 29/3028 | 2.55E-07 | 3.35E-06 |
| GO:0002082 | regulation of oxidative phosphorylation | 13/3028 | 2.85E-07 | 3.72E-06 |
| GO:0010884 | positive regulation of lipid storage | 13/3028 | 2.85E-07 | 3.72E-06 |
| GO:0030449 | regulation of complement activation | 13/3028 | 2.85E-07 | 3.72E-06 |

| | | | | |
|---|---|---|---|---|
| GO:0033500 | carbohydrate homeostasis | 60/3028 | 2.88E-07 | 3.74E-06 |
| GO:0006957 | complement activation, alternative pathway | Oct-28 | 2.95E-07 | 3.83E-06 |
| GO:0110096 | cellular response to aldehyde | Oct-28 | 2.95E-07 | 3.83E-06 |
| GO:0006096 | glycolytic process | 27/3028 | 3.08E-07 | 3.98E-06 |
| GO:0006094 | gluconeogenesis | 25/3028 | 3.53E-07 | 4.55E-06 |
| GO:1905897 | regulation of response to endoplasmic reticulum stress | 24/3028 | 3.68E-07 | 4.73E-06 |
| GO:0007030 | Golgi organization | 35/3028 | 3.94E-07 | 5.05E-06 |
| GO:0006757 | ATP generation from ADP | 27/3028 | 3.94E-07 | 5.05E-06 |
| GO:0009895 | negative regulation of catabolic process | 63/3028 | 4.01E-07 | 5.13E-06 |
| GO:1900542 | regulation of purine nucleotide metabolic process | 25/3028 | 4.59E-07 | 5.86E-06 |
| GO:0070129 | regulation of mitochondrial translation | 13/3028 | 5.15E-07 | 6.55E-06 |
| GO:0031669 | cellular response to nutrient levels | 46/3028 | 5.20E-07 | 6.61E-06 |
| GO:0046164 | alcohol catabolic process | 18/3028 | 5.25E-07 | 6.66E-06 |
| GO:0009119 | ribonucleoside metabolic process | 15/3028 | 5.37E-07 | 6.79E-06 |
| GO:0000050 | urea cycle | Aug-28 | 5.46E-07 | 6.87E-06 |
| GO:0034380 | high-density lipoprotein particle assembly | Aug-28 | 5.46E-07 | 6.87E-06 |
| GO:0050878 | regulation of body fluid levels | 71/3028 | 5.50E-07 | 6.91E-06 |
| GO:0044264 | cellular polysaccharide metabolic process | 28/3028 | 5.72E-07 | 7.16E-06 |

| | | | | |
|---|---|---|---|---|
| GO:0006613 | cotranslational protein targeting to membrane | Nov-28 | 5.74E-07 | 7.16E-06 |
| GO:0044241 | lipid digestion | Nov-28 | 5.74E-07 | 7.16E-06 |
| GO:0042593 | glucose homeostasis | 59/3028 | 5.75E-07 | 7.17E-06 |
| GO:0006665 | sphingolipid metabolic process | 36/3028 | 5.81E-07 | 7.21E-06 |
| GO:0006140 | regulation of nucleotide metabolic process | 25/3028 | 5.95E-07 | 7.38E-06 |
| GO:0010951 | negative regulation of endopeptidase activity | 43/3028 | 5.98E-07 | 7.40E-06 |
| GO:0042886 | amide transport | 68/3028 | 6.04E-07 | 7.46E-06 |
| GO:0019748 | secondary metabolic process | 20/3028 | 6.51E-07 | 8.02E-06 |
| GO:0009070 | serine family amino acid biosynthetic process | Oct-28 | 7.13E-07 | 8.72E-06 |
| GO:0019359 | nicotinamide nucleotide biosynthetic process | Oct-28 | 7.13E-07 | 8.72E-06 |
| GO:0030299 | intestinal cholesterol absorption | Oct-28 | 7.13E-07 | 8.72E-06 |
| GO:0017038 | protein import | 50/3028 | 7.14E-07 | 8.72E-06 |
| GO:0031331 | positive regulation of cellular catabolic process | 83/3028 | 7.21E-07 | 8.79E-06 |
| GO:0019751 | polyol metabolic process | 31/3028 | 7.35E-07 | 8.95E-06 |
| GO:0006063 | uronic acid metabolic process | Sep-28 | 7.44E-07 | 8.97E-06 |
| GO:0010872 | regulation of cholesterol esterification | Sep-28 | 7.44E-07 | 8.97E-06 |
| GO:0019585 | glucuronate metabolic process | Sep-28 | 7.44E-07 | 8.97E-06 |
| GO:0070508 | cholesterol import | Sep-28 | 7.44E-07 | 8.97E-06 |

| | | | | |
|---|---|---|---|---|
| GO:1904729 | regulation of intestinal lipid absorption | Sep-28 | 7.44E-07 | 8.97E-06 |
| GO:0072599 | establishment of protein localization to endoplasmic reticulum | 18/3028 | 7.58E-07 | 9.11E-06 |
| GO:0009151 | purine deoxyribonucleotide metabolic process | Dec-28 | 7.89E-07 | 9.46E-06 |
| GO:1990845 | adaptive thermogenesis | 39/3028 | 7.98E-07 | 9.56E-06 |
| GO:0032869 | cellular response to insulin stimulus | 46/3028 | 8.04E-07 | 9.61E-06 |
| GO:0042743 | hydrogen peroxide metabolic process | 20/3028 | 8.96E-07 | 1.07E-05 |
| GO:0019362 | pyridine nucleotide metabolic process | 13/3028 | 8.97E-07 | 1.07E-05 |
| GO:0031639 | plasminogen activation | 13/3028 | 8.97E-07 | 1.07E-05 |
| GO:0007584 | response to nutrient | 22/3028 | 9.04E-07 | 1.07E-05 |
| GO:0010466 | negative regulation of peptidase activity | 54/3028 | 9.99E-07 | 1.18E-05 |
| GO:0042398 | cellular modified amino acid biosynthetic process | 16/3028 | 1.13E-06 | 1.33E-05 |
| GO:0006390 | mitochondrial transcription | Nov-28 | 1.15E-06 | 1.35E-05 |
| GO:0009159 | deoxyribonucleoside monophosphate catabolic process | Nov-28 | 1.15E-06 | 1.35E-05 |
| GO:0098856 | intestinal lipid absorption | Nov-28 | 1.15E-06 | 1.35E-05 |
| GO:0106106 | cold-induced thermogenesis | 36/3028 | 1.18E-06 | 1.38E-05 |
| GO:0009154 | purine ribonucleotide catabolic process | 15/3028 | 1.31E-06 | 1.53E-05 |
| GO:0032102 | negative regulation of response to external stimulus | 74/3028 | 1.33E-06 | 1.55E-05 |
| GO:0098869 | cellular oxidant detoxification | Dec-28 | 1.43E-06 | 1.66E-05 |

| | | | | |
|---|---|---|---|---|
| GO:0042430 | indole-containing compound metabolic process | 13/3028 | 1.51E-06 | 1.75E-05 |
| GO:0050995 | negative regulation of lipid catabolic process | 13/3028 | 1.51E-06 | 1.75E-05 |
| GO:0046415 | urate metabolic process | Oct-28 | 1.57E-06 | 1.81E-05 |
| GO:1901264 | carbohydrate derivative transport | 21/3028 | 1.63E-06 | 1.88E-05 |
| GO:0006399 | tRNA metabolic process | 39/3028 | 1.78E-06 | 2.05E-05 |
| GO:0006107 | oxaloacetate metabolic process | Aug-28 | 1.82E-06 | 2.07E-05 |
| GO:0006122 | mitochondrial electron transport, ubiquinol to cytochrome c | Aug-28 | 1.82E-06 | 2.07E-05 |
| GO:0009812 | flavonoid metabolic process | Aug-28 | 1.82E-06 | 2.07E-05 |
| GO:0045542 | positive regulation of cholesterol biosynthetic process | Aug-28 | 1.82E-06 | 2.07E-05 |
| GO:0106120 | positive regulation of sterol biosynthetic process | Aug-28 | 1.82E-06 | 2.07E-05 |
| GO:0010635 | regulation of mitochondrial fusion | Sep-28 | 1.89E-06 | 2.13E-05 |
| GO:0030497 | fatty acid elongation | Sep-28 | 1.89E-06 | 2.13E-05 |
| GO:0034756 | regulation of iron ion transport | Sep-28 | 1.89E-06 | 2.13E-05 |
| GO:0042559 | pteridine-containing compound biosynthetic process | Sep-28 | 1.89E-06 | 2.13E-05 |
| GO:0046459 | short-chain fatty acid metabolic process | Sep-28 | 1.89E-06 | 2.13E-05 |
| GO:0032370 | positive regulation of lipid transport | 23/3028 | 1.91E-06 | 2.16E-05 |
| GO:0030148 | sphingolipid biosynthetic process | 27/3028 | 1.97E-06 | 2.22E-05 |
| GO:0006775 | fat-soluble vitamin metabolic process | 15/3028 | 1.99E-06 | 2.24E-05 |

| | | | | |
|---|---|---|---|---|
| GO:0051289 | protein homotetramerization | 18/3028 | 2.12E-06 | 2.38E-05 |
| GO:0033617 | mitochondrial cytochrome c oxidase assembly | Nov-28 | 2.19E-06 | 2.45E-05 |
| GO:0046653 | tetrahydrofolate metabolic process | Nov-28 | 2.19E-06 | 2.45E-05 |
| GO:0005976 | polysaccharide metabolic process | 29/3028 | 2.23E-06 | 2.49E-05 |
| GO:0120161 | regulation of cold-induced thermogenesis | 35/3028 | 2.35E-06 | 2.62E-05 |
| GO:0010893 | positive regulation of steroid biosynthetic process | 13/3028 | 2.48E-06 | 2.74E-05 |
| GO:0008209 | androgen metabolic process | Dec-28 | 2.48E-06 | 2.74E-05 |
| GO:0009125 | nucleoside monophosphate catabolic process | Dec-28 | 2.48E-06 | 2.74E-05 |
| GO:0000266 | mitochondrial fission | 17/3028 | 2.75E-06 | 3.04E-05 |
| GO:0006612 | protein targeting to membrane | 31/3028 | 2.85E-06 | 3.15E-05 |
| GO:0034284 | response to monosaccharide | 43/3028 | 2.99E-06 | 3.29E-05 |
| GO:0006614 | SRP-dependent cotranslational protein targeting to membrane | Oct-28 | 3.19E-06 | 3.50E-05 |
| GO:0072378 | blood coagulation, fibrin clot formation | Oct-28 | 3.19E-06 | 3.50E-05 |
| GO:0009743 | response to carbohydrate | 45/3028 | 3.40E-06 | 3.72E-05 |
| GO:0006984 | ER-nucleus signaling pathway | 16/3028 | 3.50E-06 | 3.81E-05 |
| GO:0010874 | regulation of cholesterol efflux | 16/3028 | 3.50E-06 | 3.81E-05 |
| GO:0043039 | tRNA aminoacylation | 16/3028 | 3.50E-06 | 3.81E-05 |
| GO:0030968 | endoplasmic reticulum unfolded protein response | 21/3028 | 3.69E-06 | 4.01E-05 |

| | | | | |
|---|---|---|---|---|
| GO:0042594 | response to starvation | 41/3028 | 3.90E-06 | 4.23E-05 |
| GO:0006835 | dicarboxylic acid transport | 25/3028 | 3.96E-06 | 4.27E-05 |
| GO:0006656 | phosphatidylcholine biosynthetic process | Nov-28 | 3.96E-06 | 4.27E-05 |
| GO:0009158 | ribonucleoside monophosphate catabolic process | Nov-28 | 3.96E-06 | 4.27E-05 |
| GO:0034599 | cellular response to oxidative stress | 54/3028 | 4.13E-06 | 4.44E-05 |
| GO:0006760 | folic acid-containing compound metabolic process | Dec-28 | 4.17E-06 | 4.46E-05 |
| GO:0010888 | negative regulation of lipid storage | Dec-28 | 4.17E-06 | 4.46E-05 |
| GO:0006570 | tyrosine metabolic process | Sep-28 | 4.28E-06 | 4.51E-05 |
| GO:0010887 | negative regulation of cholesterol storage | Sep-28 | 4.28E-06 | 4.51E-05 |
| GO:0010889 | regulation of sequestering of triglyceride | Sep-28 | 4.28E-06 | 4.51E-05 |
| GO:0032373 | positive regulation of sterol transport | Sep-28 | 4.28E-06 | 4.51E-05 |
| GO:0032376 | positive regulation of cholesterol transport | Sep-28 | 4.28E-06 | 4.51E-05 |
| GO:0032933 | SREBP signaling pathway | Sep-28 | 4.28E-06 | 4.51E-05 |
| GO:0033700 | phospholipid efflux | Sep-28 | 4.28E-06 | 4.51E-05 |
| GO:0042182 | ketone catabolic process | Sep-28 | 4.28E-06 | 4.51E-05 |
| GO:0055089 | fatty acid homeostasis | Sep-28 | 4.28E-06 | 4.51E-05 |
| GO:0001659 | temperature homeostasis | 41/3028 | 4.49E-06 | 4.73E-05 |
| GO:0006417 | regulation of translation | 71/3028 | 4.70E-06 | 4.94E-05 |

| | | | | |
|---|---|---|---|---|
| GO:0009746 | response to hexose | 42/3028 | 4.82E-06 | 5.06E-05 |
| GO:0018958 | phenol-containing compound metabolic process | 31/3028 | 4.90E-06 | 5.12E-05 |
| GO:0006152 | purine nucleoside catabolic process | Aug-28 | 4.94E-06 | 5.12E-05 |
| GO:0009435 | NAD biosynthetic process | Aug-28 | 4.94E-06 | 5.12E-05 |
| GO:0052695 | cellular glucuronidation | Aug-28 | 4.94E-06 | 5.12E-05 |
| GO:0071281 | cellular response to iron ion | Aug-28 | 4.94E-06 | 5.12E-05 |
| GO:0071712 | ER-associated misfolded protein catabolic process | Aug-28 | 4.94E-06 | 5.12E-05 |
| GO:0090209 | negative regulation of triglyceride metabolic process | Aug-28 | 4.94E-06 | 5.12E-05 |
| GO:0043038 | amino acid activation | 16/3028 | 4.97E-06 | 5.13E-05 |
| GO:0071375 | cellular response to peptide hormone stimulus | 53/3028 | 5.09E-06 | 5.25E-05 |
| GO:0044275 | cellular carbohydrate catabolic process | 14/3028 | 5.28E-06 | 5.44E-05 |
| GO:0009161 | ribonucleoside monophosphate metabolic process | 18/3028 | 5.42E-06 | 5.57E-05 |
| GO:1903362 | regulation of cellular protein catabolic process | 50/3028 | 6.01E-06 | 6.17E-05 |
| GO:1904251 | regulation of bile acid metabolic process | Oct-28 | 6.10E-06 | 6.25E-05 |
| GO:0006418 | tRNA aminoacylation for protein translation | 15/3028 | 6.30E-06 | 6.44E-05 |
| GO:1903573 | negative regulation of response to endoplasmic reticulum stress | 15/3028 | 6.30E-06 | 6.44E-05 |
| GO:0006515 | protein quality control for misfolded or incompletely synthesized proteins | Dec-28 | 6.78E-06 | 6.90E-05 |
| GO:0000302 | response to reactive oxygen species | 41/3028 | 6.78E-06 | 6.90E-05 |

| | | | | |
|---|---|---|---|---|
| GO:0010984 | regulation of lipoprotein particle clearance | Nov-28 | 6.87E-06 | 6.97E-05 |
| GO:0019400 | alditol metabolic process | Nov-28 | 6.87E-06 | 6.97E-05 |
| GO:1902930 | regulation of alcohol biosynthetic process | 17/3028 | 7.26E-06 | 7.33E-05 |
| GO:1903426 | regulation of reactive oxygen species biosynthetic process | 17/3028 | 7.26E-06 | 7.33E-05 |
| GO:0051205 | protein insertion into membrane | 18/3028 | 7.26E-06 | 7.33E-05 |
| GO:0035592 | establishment of protein localization to extracellular region | 71/3028 | 7.36E-06 | 7.41E-05 |
| GO:0042176 | regulation of protein catabolic process | 70/3028 | 7.53E-06 | 7.57E-05 |
| GO:0097193 | intrinsic apoptotic signaling pathway | 58/3028 | 8.06E-06 | 8.09E-05 |
| GO:0006643 | membrane lipid metabolic process | 40/3028 | 8.33E-06 | 8.35E-05 |
| GO:0042159 | lipoprotein catabolic process | Sep-28 | 8.86E-06 | 8.79E-05 |
| GO:0042454 | ribonucleoside catabolic process | Sep-28 | 8.86E-06 | 8.79E-05 |
| GO:0055091 | phospholipid homeostasis | Sep-28 | 8.86E-06 | 8.79E-05 |
| GO:0070857 | regulation of bile acid biosynthetic process | Sep-28 | 8.86E-06 | 8.79E-05 |
| GO:0071501 | cellular response to sterol depletion | Sep-28 | 8.86E-06 | 8.79E-05 |
| GO:0072350 | tricarboxylic acid metabolic process | Sep-28 | 8.86E-06 | 8.79E-05 |
| GO:0061912 | selective autophagy | 22/3028 | 9.00E-06 | 8.91E-05 |
| GO:0072527 | pyrimidine-containing compound metabolic process | 18/3028 | 9.66E-06 | 9.54E-05 |
| GO:0048545 | response to steroid hormone | 46/3028 | 9.72E-06 | 9.59E-05 |

| | | | | |
|---|---|---|---|---|
| GO:0046128 | purine ribonucleoside metabolic process | Dec-28 | 1.07E-05 | 0.000105678 |
| GO:0006071 | glycerol metabolic process | Oct-28 | 1.11E-05 | 0.00010831 |
| GO:0009067 | aspartate family amino acid biosynthetic process | Oct-28 | 1.11E-05 | 0.00010831 |
| GO:0009110 | vitamin biosynthetic process | Oct-28 | 1.11E-05 | 0.00010831 |
| GO:0003018 | vascular process in circulatory system | 47/3028 | 1.12E-05 | 0.000110063 |
| GO:0046496 | nicotinamide nucleotide metabolic process | Nov-28 | 1.15E-05 | 0.000111974 |
| GO:0072376 | protein activation cascade | Nov-28 | 1.15E-05 | 0.000111974 |
| GO:0006568 | tryptophan metabolic process | Aug-28 | 1.17E-05 | 0.000113094 |
| GO:0006586 | indolalkylamine metabolic process | Aug-28 | 1.17E-05 | 0.000113094 |
| GO:0009120 | deoxyribonucleoside metabolic process | Aug-28 | 1.17E-05 | 0.000113094 |
| GO:0051260 | protein homooligomerization | 39/3028 | 1.17E-05 | 0.000113507 |
| GO:0071692 | protein localization to extracellular region | 71/3028 | 1.24E-05 | 0.000119928 |
| GO:0061045 | negative regulation of wound healing | 21/3028 | 1.27E-05 | 0.000122624 |
| GO:0051353 | positive regulation of oxidoreductase activity | 18/3028 | 1.27E-05 | 0.000122624 |
| GO:0009306 | protein secretion | 70/3028 | 1.27E-05 | 0.000122624 |
| GO:0031647 | regulation of protein stability | 58/3028 | 1.33E-05 | 0.000127763 |
| GO:0003333 | amino acid transmembrane transport | 25/3028 | 1.37E-05 | 0.000131611 |
| GO:0042158 | lipoprotein biosynthetic process | 26/3028 | 1.41E-05 | 0.000134926 |

| | | | | |
|---|---|---|---|---|
| GO:0009395 | phospholipid catabolic process | 18/3028 | 1.67E-05 | 0.000159274 |
| GO:0000963 | mitochondrial RNA processing | Sep-28 | 1.71E-05 | 0.000162124 |
| GO:0006991 | response to sterol depletion | Sep-28 | 1.71E-05 | 0.000162124 |
| GO:0035337 | fatty-acyl-CoA metabolic process | Sep-28 | 1.71E-05 | 0.000162124 |
| GO:0035459 | vesicle cargo loading | Sep-28 | 1.71E-05 | 0.000162124 |
| GO:0090199 | regulation of release of cytochrome c from mitochondria | 17/3028 | 1.75E-05 | 0.000165989 |
| GO:0009749 | response to glucose | 40/3028 | 1.83E-05 | 0.000173616 |
| GO:0018904 | ether metabolic process | Nov-28 | 1.86E-05 | 0.00017482 |
| GO:0036315 | cellular response to sterol | Nov-28 | 1.86E-05 | 0.00017482 |
| GO:0090114 | COPII-coated vesicle budding | Nov-28 | 1.86E-05 | 0.00017482 |
| GO:0060193 | positive regulation of lipase activity | 19/3028 | 1.99E-05 | 0.000186944 |
| GO:0015908 | fatty acid transport | 25/3028 | 2.02E-05 | 0.000189226 |
| GO:0006865 | amino acid transport | 35/3028 | 2.02E-05 | 0.000189719 |
| GO:0055094 | response to lipoprotein particle | 13/3028 | 2.04E-05 | 0.00019139 |
| GO:0009267 | cellular response to starvation | 36/3028 | 2.17E-05 | 0.000203251 |
| GO:0051259 | protein complex oligomerization | 48/3028 | 2.25E-05 | 0.000209685 |
| GO:0002920 | regulation of humoral immune response | 16/3028 | 2.39E-05 | 0.000222915 |
| GO:0008207 | C21-steroid hormone metabolic process | 15/3028 | 2.40E-05 | 0.000222936 |

| | | | | |
|---|---|---|---|---|
| GO:0002138 | retinoic acid biosynthetic process | Aug-28 | 2.47E-05 | 0.000228449 |
| GO:0016102 | diterpenoid biosynthetic process | Aug-28 | 2.47E-05 | 0.000228449 |
| GO:0090110 | COPII-coated vesicle cargo loading | Aug-28 | 2.47E-05 | 0.000228449 |
| GO:0010155 | regulation of proton transport | Dec-28 | 2.50E-05 | 0.000230598 |
| GO:0031330 | negative regulation of cellular catabolic process | 48/3028 | 2.50E-05 | 0.00023107 |
| GO:1901653 | cellular response to peptide | 57/3028 | 2.64E-05 | 0.000243427 |
| GO:0030730 | sequestering of triglyceride | Sep-28 | 3.09E-05 | 0.000283775 |
| GO:0060192 | negative regulation of lipase activity | Sep-28 | 3.09E-05 | 0.000283775 |
| GO:1901663 | quinone biosynthetic process | Sep-28 | 3.09E-05 | 0.000283775 |
| GO:1904732 | regulation of electron transfer activity | Jul-28 | 3.16E-05 | 0.000288653 |
| GO:0009219 | pyrimidine deoxyribonucleotide metabolic process | Oct-28 | 3.17E-05 | 0.000288653 |
| GO:0034755 | iron ion transmembrane transport | Oct-28 | 3.17E-05 | 0.000288653 |
| GO:0042402 | cellular biogenic amine catabolic process | Oct-28 | 3.17E-05 | 0.000288653 |
| GO:0071404 | cellular response to low-density lipoprotein particle stimulus | Oct-28 | 3.17E-05 | 0.000288653 |
| GO:0001836 | release of cytochrome c from mitochondria | 19/3028 | 3.21E-05 | 0.000291124 |
| GO:0015909 | long-chain fatty acid transport | 19/3028 | 3.21E-05 | 0.000291124 |
| GO:0006890 | retrograde vesicle-mediated transport, Golgi to endoplasmic reticulum | 15/3028 | 3.25E-05 | 0.000294092 |
| GO:0051262 | protein tetramerization | 22/3028 | 3.28E-05 | 0.000296925 |

| | | | | |
|---|---|---|---|---|
| GO:00610 08 | hepaticobiliary system development | 30/3028 | 3.59E-05 | 0.0003243 43 |
| GO:00363 14 | response to sterol | Dec-28 | 3.68E-05 | 0.0003314 05 |
| GO:00723 37 | modified amino acid transport | Dec-28 | 3.68E-05 | 0.0003314 05 |
| GO:00430 90 | amino acid import | 17/3028 | 3.90E-05 | 0.0003509 08 |
| GO:00610 41 | regulation of wound healing | 30/3028 | 4.18E-05 | 0.0003748 82 |
| GO:00067 06 | steroid catabolic process | Nov-28 | 4.45E-05 | 0.0003977 09 |
| GO:00157 40 | C4-dicarboxylate transport | Nov-28 | 4.45E-05 | 0.0003977 09 |
| GO:00158 07 | L-amino acid transport | 18/3028 | 4.54E-05 | 0.0004054 83 |
| GO:00091 55 | purine deoxyribonucleotide catabolic process | Aug-28 | 4.80E-05 | 0.0004260 41 |
| GO:00091 70 | purine deoxyribonucleoside monophosphate metabolic process | Aug-28 | 4.80E-05 | 0.0004260 41 |
| GO:00159 36 | coenzyme A metabolic process | Aug-28 | 4.80E-05 | 0.0004260 41 |
| GO:00425 74 | retinal metabolic process | Aug-28 | 4.80E-05 | 0.0004260 41 |
| GO:00000 45 | autophagosome assembly | 25/3028 | 4.97E-05 | 0.0004401 74 |
| GO:00158 33 | peptide transport | 57/3028 | 5.03E-05 | 0.0004453 88 |
| GO:00068 01 | superoxide metabolic process | 19/3028 | 5.06E-05 | 0.0004472 46 |
| GO:00093 10 | amine catabolic process | Oct-28 | 5.08E-05 | 0.0004472 6 |
| GO:00708 61 | regulation of protein exit from endoplasmic reticulum | Oct-28 | 5.08E-05 | 0.0004472 6 |
| GO:00902 01 | negative regulation of release of cytochrome c from mitochondria | Oct-28 | 5.08E-05 | 0.0004472 6 |

| | | | | |
|---|---|---|---|---|
| GO:0006743 | ubiquinone metabolic process | Sep-28 | 5.33E-05 | 0.000466826 |
| GO:0006851 | mitochondrial calcium ion transmembrane transport | Sep-28 | 5.33E-05 | 0.000466826 |
| GO:0035461 | vitamin transmembrane transport | Sep-28 | 5.33E-05 | 0.000466826 |
| GO:0001885 | endothelial cell development | 20/3028 | 5.46E-05 | 0.00047753 |
| GO:0062197 | cellular response to chemical stress | 59/3028 | 5.72E-05 | 0.000499062 |
| GO:0010907 | positive regulation of glucose metabolic process | 15/3028 | 5.78E-05 | 0.000503405 |
| GO:0030808 | regulation of nucleotide biosynthetic process | 13/3028 | 5.80E-05 | 0.000503405 |
| GO:0071402 | cellular response to lipoprotein particle stimulus | 13/3028 | 5.80E-05 | 0.000503405 |
| GO:1900371 | regulation of purine nucleotide biosynthetic process | 13/3028 | 5.80E-05 | 0.000503405 |
| GO:0001889 | liver development | 29/3028 | 5.94E-05 | 0.000514365 |
| GO:1903035 | negative regulation of response to wounding | 23/3028 | 5.99E-05 | 0.00051722 |
| GO:2001057 | reactive nitrogen species metabolic process | 23/3028 | 5.99E-05 | 0.00051722 |
| GO:0043470 | regulation of carbohydrate catabolic process | 17/3028 | 6.42E-05 | 0.000553508 |
| GO:0007586 | digestion | 30/3028 | 6.49E-05 | 0.000559005 |
| GO:0034637 | cellular carbohydrate biosynthetic process | 20/3028 | 6.72E-05 | 0.000577909 |
| GO:1905037 | autophagosome organization | 26/3028 | 6.77E-05 | 0.000581522 |
| GO:0006686 | sphingomyelin biosynthetic process | Jul-28 | 6.90E-05 | 0.00058888 |
| GO:0009068 | aspartate family amino acid catabolic process | Jul-28 | 6.90E-05 | 0.00058888 |

| | | | | |
|---|---|---|---|---|
| GO:0009215 | purine deoxyribonucleoside triphosphate metabolic process | Jul-28 | 6.90E-05 | 0.00058888 |
| GO:0043248 | proteasome assembly | Jul-28 | 6.90E-05 | 0.00058888 |
| GO:0034614 | cellular response to reactive oxygen species | 31/3028 | 7.00E-05 | 0.000597282 |
| GO:0007029 | endoplasmic reticulum organization | 23/3028 | 7.17E-05 | 0.000610501 |
| GO:0070059 | intrinsic apoptotic signaling pathway in response to endoplasmic reticulum stress | 18/3028 | 7.20E-05 | 0.000612447 |
| GO:0009100 | glycoprotein metabolic process | 58/3028 | 7.60E-05 | 0.000645586 |
| GO:0008210 | estrogen metabolic process | Oct-28 | 7.90E-05 | 0.000666476 |
| GO:0034383 | low-density lipoprotein particle clearance | Oct-28 | 7.90E-05 | 0.000666476 |
| GO:0050820 | positive regulation of coagulation | Oct-28 | 7.90E-05 | 0.000666476 |
| GO:0051788 | response to misfolded protein | Oct-28 | 7.90E-05 | 0.000666476 |
| GO:0002526 | acute inflammatory response | 26/3028 | 7.95E-05 | 0.000669688 |
| GO:0097035 | regulation of membrane lipid distribution | 17/3028 | 8.14E-05 | 0.000683816 |
| GO:1902475 | L-alpha-amino acid transmembrane transport | 17/3028 | 8.14E-05 | 0.000683816 |
| GO:0045913 | positive regulation of carbohydrate metabolic process | 21/3028 | 8.48E-05 | 0.000711427 |
| GO:0071397 | cellular response to cholesterol | Sep-28 | 8.79E-05 | 0.000736467 |
| GO:0022600 | digestive system process | 27/3028 | 8.88E-05 | 0.000743319 |
| GO:0008643 | carbohydrate transport | 33/3028 | 9.00E-05 | 0.000752135 |
| GO:0031348 | negative regulation of defense response | 45/3028 | 9.69E-05 | 0.000808976 |

| | | | | |
|---|---|---|---|---|
| GO:0030522 | intracellular receptor signaling pathway | 42/3028 | 9.75E-05 | 0.000812498 |
| GO:0010822 | positive regulation of mitochondrion organization | 20/3028 | 0.000100242 | 0.000834061 |
| GO:0051384 | response to glucocorticoid | 21/3028 | 0.000102362 | 0.000850494 |
| GO:0051222 | positive regulation of protein transport | 56/3028 | 0.000103956 | 0.000862512 |
| GO:0005978 | glycogen biosynthetic process | 14/3028 | 0.000105166 | 0.000870081 |
| GO:0009250 | glucan biosynthetic process | 14/3028 | 0.000105166 | 0.000870081 |
| GO:0046467 | membrane lipid biosynthetic process | 29/3028 | 0.000105903 | 0.000874938 |
| GO:0009167 | purine ribonucleoside monophosphate metabolic process | 13/3028 | 0.000108133 | 0.000889581 |
| GO:0045981 | positive regulation of nucleotide metabolic process | 13/3028 | 0.000108133 | 0.000889581 |
| GO:1900544 | positive regulation of purine nucleotide metabolic process | 13/3028 | 0.000108133 | 0.000889581 |
| GO:0003044 | regulation of systemic arterial blood pressure mediated by a chemical signal | 16/3028 | 0.000115323 | 0.000947401 |
| GO:0090151 | establishment of protein localization to mitochondrial membrane | Oct-28 | 0.000119228 | 0.000978103 |
| GO:0031960 | response to corticosteroid | 22/3028 | 0.0001227 | 0.001003766 |
| GO:0046209 | nitric oxide metabolic process | 22/3028 | 0.0001227 | 0.001003766 |
| GO:1903034 | regulation of response to wounding | 35/3028 | 0.000124053 | 0.00101342 |
| GO:0050873 | brown fat cell differentiation | 17/3028 | 0.000128318 | 0.001046792 |
| GO:0051224 | negative regulation of protein transport | 30/3028 | 0.000129796 | 0.001057378 |
| GO:0002793 | positive regulation of peptide secretion | 28/3028 | 0.00013021 | 0.001059272 |

| | | | | |
|---|---|---|---|---|
| GO:2001233 | regulation of apoptotic signaling pathway | 67/3028 | 0.000132244 | 0.0010743 2 |
| GO:0009084 | glutamine family amino acid biosynthetic process | Jul-28 | 0.000135814 | 0.001095705 |
| GO:0033240 | positive regulation of cellular amine metabolic process | Jul-28 | 0.000135814 | 0.001095705 |
| GO:0035999 | tetrahydrofolate interconversion | Jul-28 | 0.000135814 | 0.001095705 |
| GO:0042761 | very long-chain fatty acid biosynthetic process | Jul-28 | 0.000135814 | 0.001095705 |
| GO:0046054 | dGMP metabolic process | Jul-28 | 0.000135814 | 0.001095705 |
| GO:0034655 | nucleobase-containing compound catabolic process | 66/3028 | 0.0001377 9 | 0.001110111 |
| GO:0009225 | nucleotide-sugar metabolic process | Nov-28 | 0.000138215 | 0.001110476 |
| GO:0046033 | AMP metabolic process | Nov-28 | 0.000138215 | 0.001110476 |
| GO:0006662 | glycerol ether metabolic process | Sep-28 | 0.000139463 | 0.001115892 |
| GO:0051204 | protein insertion into mitochondrial membrane | Sep-28 | 0.000139463 | 0.001115892 |
| GO:0071071 | regulation of phospholipid biosynthetic process | Sep-28 | 0.000139463 | 0.001115892 |
| GO:1903580 | positive regulation of ATP metabolic process | 13/3028 | 0.000144875 | 0.001157609 |
| GO:0000422 | autophagy of mitochondrion | 20/3028 | 0.000146937 | 0.001170886 |
| GO:0061726 | mitochondrion disassembly | 20/3028 | 0.000146937 | 0.001170886 |
| GO:0006744 | ubiquinone biosynthetic process | Aug-28 | 0.000149398 | 0.001180831 |
| GO:0009176 | pyrimidine deoxyribonucleoside monophosphate metabolic process | Aug-28 | 0.000149398 | 0.001180831 |
| GO:0010918 | positive regulation of mitochondrial membrane potential | Aug-28 | 0.000149398 | 0.001180831 |

| | | | | |
|---|---|---|---|---|
| GO:0019430 | removal of superoxide radicals | Aug-28 | 0.000149398 | 0.001180831 |
| GO:0032048 | cardiolipin metabolic process | Aug-28 | 0.000149398 | 0.001180831 |
| GO:1901569 | fatty acid derivative catabolic process | Aug-28 | 0.000149398 | 0.001180831 |
| GO:0120162 | positive regulation of cold-induced thermogenesis | 23/3028 | 0.00016815 | 0.001327252 |
| GO:0045446 | endothelial cell differentiation | 26/3028 | 0.000170228 | 0.001341837 |
| GO:0043171 | peptide catabolic process | Oct-28 | 0.000175551 | 0.00137822 |
| GO:0070723 | response to cholesterol | Oct-28 | 0.000175551 | 0.00137822 |
| GO:0097164 | ammonium ion metabolic process | Oct-28 | 0.000175551 | 0.00137822 |
| GO:2001242 | regulation of intrinsic apoptotic signaling pathway | 35/3028 | 0.000176139 | 0.001380978 |
| GO:0044058 | regulation of digestive system process | 16/3028 | 0.000183139 | 0.001433935 |
| GO:0090277 | positive regulation of peptide hormone secretion | 27/3028 | 0.00018443 | 0.001442107 |
| GO:0006220 | pyrimidine nucleotide metabolic process | 13/3028 | 0.000191863 | 0.001498226 |
| GO:0008300 | isoprenoid catabolic process | Jun-28 | 0.000193402 | 0.001500217 |
| GO:0046053 | dAMP metabolic process | Jun-28 | 0.000193402 | 0.001500217 |
| GO:0046055 | dGMP catabolic process | Jun-28 | 0.000193402 | 0.001500217 |
| GO:0046130 | purine ribonucleoside catabolic process | Jun-28 | 0.000193402 | 0.001500217 |
| GO:1904959 | regulation of cytochrome-c oxidase activity | Jun-28 | 0.000193402 | 0.001500217 |
| GO:0046173 | polyol biosynthetic process | 17/3028 | 0.00019744 | 0.001529504 |

| | | | | |
|---|---|---|---|---|
| GO:0097242 | amyloid-beta clearance | Dec-28 | 0.000198006 | 0.001531862 |
| GO:0045765 | regulation of angiogenesis | 53/3028 | 0.000205183 | 0.001585284 |
| GO:0006684 | sphingomyelin metabolic process | Sep-28 | 0.000214049 | 0.001647248 |
| GO:0071218 | cellular response to misfolded protein | Sep-28 | 0.000214049 | 0.001647248 |
| GO:0072529 | pyrimidine-containing compound catabolic process | Sep-28 | 0.000214049 | 0.001647248 |
| GO:0045444 | fat cell differentiation | 47/3028 | 0.000221628 | 0.00170333 |
| GO:1903829 | positive regulation of protein localization | 77/3028 | 0.000233784 | 0.001794389 |
| GO:0050673 | epithelial cell proliferation | 75/3028 | 0.000241814 | 0.001853587 |
| GO:0035456 | response to interferon-beta | 17/3028 | 0.000242799 | 0.001858704 |
| GO:0035296 | regulation of tube diameter | 34/3028 | 0.000243899 | 0.001859361 |
| GO:0097746 | blood vessel diameter maintenance | 34/3028 | 0.000243899 | 0.001859361 |
| GO:0009200 | deoxyribonucleoside triphosphate metabolic process | Aug-28 | 0.000244159 | 0.001859361 |
| GO:0042953 | lipoprotein transport | Aug-28 | 0.000244159 | 0.001859361 |
| GO:0006206 | pyrimidine nucleobase metabolic process | Jul-28 | 0.000246949 | 0.001863604 |
| GO:0006465 | signal peptide processing | Jul-28 | 0.000246949 | 0.001863604 |
| GO:0046073 | dTMP metabolic process | Jul-28 | 0.000246949 | 0.001863604 |
| GO:0090153 | regulation of sphingolipid biosynthetic process | Jul-28 | 0.000246949 | 0.001863604 |
| GO:0097428 | protein maturation by iron-sulfur cluster transfer | Jul-28 | 0.000246949 | 0.001863604 |

| | | | | |
|---|---|---|---|---|
| GO:1905038 | regulation of membrane lipid metabolic process | Jul-28 | 0.000246949 | 0.001863604 |
| GO:2001140 | positive regulation of phospholipid transport | Jul-28 | 0.000246949 | 0.001863604 |
| GO:1904950 | negative regulation of establishment of protein localization | 30/3028 | 0.00024774 | 0.001867159 |
| GO:0007033 | vacuole organization | 36/3028 | 0.000249076 | 0.001874813 |
| GO:0009126 | purine nucleoside monophosphate metabolic process | 13/3028 | 0.000251334 | 0.001889375 |
| GO:0006862 | nucleotide transport | Oct-28 | 0.000252663 | 0.001896919 |
| GO:0003158 | endothelium development | 28/3028 | 0.00025751 | 0.00193083 |
| GO:0000002 | mitochondrial genome maintenance | Dec-28 | 0.000265208 | 0.001986003 |
| GO:0035150 | regulation of tube size | 34/3028 | 0.000273005 | 0.002041767 |
| GO:0006900 | vesicle budding from membrane | 16/3028 | 0.000283202 | 0.002115325 |
| GO:1901342 | regulation of vasculature development | 53/3028 | 0.000284865 | 0.002125027 |
| GO:0000305 | response to oxygen radical | Sep-28 | 0.000318967 | 0.002367324 |
| GO:0009129 | pyrimidine nucleoside monophosphate metabolic process | Sep-28 | 0.000318967 | 0.002367324 |
| GO:0020027 | hemoglobin metabolic process | Sep-28 | 0.000318967 | 0.002367324 |
| GO:0042219 | cellular modified amino acid catabolic process | Sep-28 | 0.000318967 | 0.002367324 |
| GO:0009309 | amine biosynthetic process | 13/3028 | 0.000325869 | 0.002412423 |
| GO:0042401 | cellular biogenic amine biosynthetic process | 13/3028 | 0.000325869 | 0.002412423 |
| GO:0002790 | peptide secretion | 51/3028 | 0.000330737 | 0.002445357 |

| | | | | |
|---|---|---|---|---|
| GO:1904951 | positive regulation of establishment of protein localization | 56/3028 | 0.000376164 | 0.002777717 |
| GO:0009251 | glucan catabolic process | Aug-28 | 0.000382959 | 0.002796078 |
| GO:0030150 | protein import into mitochondrial matrix | Aug-28 | 0.000382959 | 0.002796078 |
| GO:0042448 | progesterone metabolic process | Aug-28 | 0.000382959 | 0.002796078 |
| GO:0044872 | lipoprotein localization | Aug-28 | 0.000382959 | 0.002796078 |
| GO:0045821 | positive regulation of glycolytic process | Aug-28 | 0.000382959 | 0.002796078 |
| GO:0050746 | regulation of lipoprotein metabolic process | Aug-28 | 0.000382959 | 0.002796078 |
| GO:0071450 | cellular response to oxygen radical | Aug-28 | 0.000382959 | 0.002796078 |
| GO:0071451 | cellular response to superoxide | Aug-28 | 0.000382959 | 0.002796078 |
| GO:1902236 | negative regulation of endoplasmic reticulum stress-induced intrinsic apoptotic signaling pathway | Aug-28 | 0.000382959 | 0.002796078 |
| GO:0006497 | protein lipidation | 22/3028 | 0.000385766 | 0.002813057 |
| GO:0009133 | nucleoside diphosphate biosynthetic process | Jun-28 | 0.000387547 | 0.002819004 |
| GO:0009172 | purine deoxyribonucleoside monophosphate catabolic process | Jun-28 | 0.000387547 | 0.002819004 |
| GO:0006111 | regulation of gluconeogenesis | 15/3028 | 0.000406533 | 0.002953435 |
| GO:0006577 | amino-acid betaine metabolic process | Jul-28 | 0.000421034 | 0.003032382 |
| GO:0009169 | purine ribonucleoside monophosphate catabolic process | Jul-28 | 0.000421034 | 0.003032382 |
| GO:0036498 | IRE1-mediated unfolded protein response | Jul-28 | 0.000421034 | 0.003032382 |
| GO:0046085 | adenosine metabolic process | Jul-28 | 0.000421034 | 0.003032382 |

| | | | | |
|---|---|---|---|---|
| GO:0046322 | negative regulation of fatty acid oxidation | Jul-28 | 0.000421034 | 0.003032382 |
| GO:0061709 | reticulophagy | Jul-28 | 0.000421034 | 0.003032382 |
| GO:2001138 | regulation of phospholipid transport | Jul-28 | 0.000421034 | 0.003032382 |
| GO:0098586 | cellular response to virus | 18/3028 | 0.000438133 | 0.003151643 |
| GO:0050728 | negative regulation of inflammatory response | 29/3028 | 0.000442286 | 0.003177603 |
| GO:1903828 | negative regulation of protein localization | 40/3028 | 0.000487305 | 0.003496739 |
| GO:0009218 | pyrimidine ribonucleotide metabolic process | Oct-28 | 0.000492792 | 0.003531775 |
| GO:0035458 | cellular response to interferon-beta | 15/3028 | 0.000501912 | 0.00359273 |
| GO:0006584 | catecholamine metabolic process | 16/3028 | 0.000520456 | 0.003711821 |
| GO:0009712 | catechol-containing compound metabolic process | 16/3028 | 0.000520456 | 0.003711821 |
| GO:0010676 | positive regulation of cellular carbohydrate metabolic process | 16/3028 | 0.000520456 | 0.003711821 |
| GO:0071326 | cellular response to monosaccharide stimulus | 30/3028 | 0.000569866 | 0.004056267 |
| GO:0050821 | protein stabilization | 36/3028 | 0.000570141 | 0.004056267 |
| GO:0030072 | peptide hormone secretion | 49/3028 | 0.000575065 | 0.004086323 |
| GO:0006089 | lactate metabolic process | Aug-28 | 0.000579677 | 0.004104112 |
| GO:0006541 | glutamine metabolic process | Aug-28 | 0.000579677 | 0.004104112 |
| GO:0050665 | hydrogen peroxide biosynthetic process | Aug-28 | 0.000579677 | 0.004104112 |
| GO:0046456 | icosanoid biosynthetic process | 14/3028 | 0.00058406 | 0.004130137 |

| | | | | |
|---|---|---|---|---|
| GO:2001234 | negative regulation of apoptotic signaling pathway | 43/3028 | 0.000593602 | 0.004192542 |
| GO:1903050 | regulation of proteolysis involved in cellular protein catabolic process | 39/3028 | 0.000598474 | 0.004221841 |
| GO:0050727 | regulation of inflammatory response | 57/3028 | 0.000602396 | 0.004244386 |
| GO:0002697 | regulation of immune effector process | 68/3028 | 0.0006085 | 0.004282228 |
| GO:0006487 | protein N-linked glycosylation | 16/3028 | 0.000630419 | 0.004431137 |
| GO:0046513 | ceramide biosynthetic process | 17/3028 | 0.000631242 | 0.004431589 |
| GO:0071456 | cellular response to hypoxia | 23/3028 | 0.000646809 | 0.00453543 |
| GO:0046037 | GMP metabolic process | Sep-28 | 0.000656326 | 0.004596646 |
| GO:0098657 | import into cell | 42/3028 | 0.000665844 | 0.004657721 |
| GO:0032369 | negative regulation of lipid transport | Oct-28 | 0.00067008 | 0.004670569 |
| GO:0039528 | cytoplasmic pattern recognition receptor signaling pathway in response to virus | Oct-28 | 0.00067008 | 0.004670569 |
| GO:0042744 | hydrogen peroxide catabolic process | Oct-28 | 0.00067008 | 0.004670569 |
| GO:0042417 | dopamine metabolic process | 13/3028 | 0.000671415 | 0.004674299 |
| GO:0009128 | purine nucleoside monophosphate catabolic process | Jul-28 | 0.000680741 | 0.004716734 |
| GO:0019184 | nonribosomal peptide biosynthetic process | Jul-28 | 0.000680741 | 0.004716734 |
| GO:0044804 | autophagy of nucleus | Jul-28 | 0.000680741 | 0.004716734 |
| GO:1902001 | fatty acid transmembrane transport | Jul-28 | 0.000680741 | 0.004716734 |
| GO:2001243 | negative regulation of intrinsic apoptotic signaling pathway | 22/3028 | 0.000694555 | 0.004806747 |

| | | | | |
|---|---|---|---|---|
| GO:0071453 | cellular response to oxygen levels | 29/3028 | 0.000705196 | 0.004807141 |
| GO:0001867 | complement activation, lectin pathway | Jun-28 | 0.000706134 | 0.004807141 |
| GO:0006000 | fructose metabolic process | Jun-28 | 0.000706134 | 0.004807141 |
| GO:0006123 | mitochondrial electron transport, cytochrome c to oxygen | Jun-28 | 0.000706134 | 0.004807141 |
| GO:0006213 | pyrimidine nucleoside metabolic process | Jun-28 | 0.000706134 | 0.004807141 |
| GO:0009223 | pyrimidine deoxyribonucleotide catabolic process | Jun-28 | 0.000706134 | 0.004807141 |
| GO:0033212 | iron import into cell | Jun-28 | 0.000706134 | 0.004807141 |
| GO:0034310 | primary alcohol catabolic process | Jun-28 | 0.000706134 | 0.004807141 |
| GO:0035437 | maintenance of protein localization in endoplasmic reticulum | Jun-28 | 0.000706134 | 0.004807141 |
| GO:0046689 | response to mercury ion | Jun-28 | 0.000706134 | 0.004807141 |
| GO:0048312 | intracellular distribution of mitochondria | Jun-28 | 0.000706134 | 0.004807141 |
| GO:0070863 | positive regulation of protein exit from endoplasmic reticulum | Jun-28 | 0.000706134 | 0.004807141 |
| GO:1904152 | regulation of retrograde protein transport, ER to cytosol | Jun-28 | 0.000706134 | 0.004807141 |
| GO:2000303 | regulation of ceramide biosynthetic process | Jun-28 | 0.000706134 | 0.004807141 |
| GO:0061028 | establishment of endothelial barrier | 14/3028 | 0.000722836 | 0.004915114 |
| GO:0043030 | regulation of macrophage activation | 15/3028 | 0.000750861 | 0.005099746 |
| GO:0006636 | unsaturated fatty acid biosynthetic process | Dec-28 | 0.000759225 | 0.005147296 |
| GO:0033692 | cellular polysaccharide biosynthetic process | 16/3028 | 0.000759625 | 0.005147296 |

| | | | | |
|---|---|---|---|---|
| GO:0061136 | regulation of proteasomal protein catabolic process | 34/3028 | 0.000784247 | 0.005307983 |
| GO:0001678 | cellular glucose homeostasis | 32/3028 | 0.000795934 | 0.005380844 |
| GO:0034219 | carbohydrate transmembrane transport | 26/3028 | 0.000839353 | 0.005658084 |
| GO:0006110 | regulation of glycolytic process | 13/3028 | 0.000839849 | 0.005658084 |
| GO:0072348 | sulfur compound transport | 13/3028 | 0.000839849 | 0.005658084 |
| GO:0043255 | regulation of carbohydrate biosynthetic process | 23/3028 | 0.000847718 | 0.005704515 |
| GO:0030194 | positive regulation of blood coagulation | Aug-28 | 0.000850578 | 0.005710604 |
| GO:1900048 | positive regulation of hemostasis | Aug-28 | 0.000850578 | 0.005710604 |
| GO:0009755 | hormone-mediated signaling pathway | 31/3028 | 0.000885981 | 0.005941461 |
| GO:0051923 | sulfation | Oct-28 | 0.000896794 | 0.006007077 |
| GO:0039529 | RIG-I signaling pathway | Sep-28 | 0.000910854 | 0.006080344 |
| GO:0045838 | positive regulation of membrane potential | Sep-28 | 0.000910854 | 0.006080344 |
| GO:0050687 | negative regulation of defense response to virus | Sep-28 | 0.000910854 | 0.006080344 |
| GO:0001818 | negative regulation of cytokine production | 49/3028 | 0.00091408 | 0.006094912 |
| GO:0050708 | regulation of protein secretion | 50/3028 | 0.000944524 | 0.006290721 |
| GO:0070085 | glycosylation | 38/3028 | 0.000965037 | 0.006420023 |
| GO:0006809 | nitric oxide biosynthetic process | 19/3028 | 0.000968537 | 0.006428665 |
| GO:0007568 | aging | 19/3028 | 0.000968537 | 0.006428665 |

| | | | | |
|---|---|---|---|---|
| GO:0050714 | positive regulation of protein secretion | 31/3028 | 0.000983128 | 0.006518105 |
| GO:0044070 | regulation of anion transport | 24/3028 | 0.001007924 | 0.006674928 |
| GO:2000379 | positive regulation of reactive oxygen species metabolic process | 18/3028 | 0.001017601 | 0.006731379 |
| GO:0070131 | positive regulation of mitochondrial translation | Jul-28 | 0.001052681 | 0.006955551 |
| GO:0001666 | response to hypoxia | 38/3028 | 0.001054663 | 0.006960777 |
| GO:0032024 | positive regulation of insulin secretion | 20/3028 | 0.001057853 | 0.006966085 |
| GO:1900407 | regulation of cellular response to oxidative stress | 20/3028 | 0.001057853 | 0.006966085 |
| GO:0070873 | regulation of glycogen metabolic process | Nov-28 | 0.001081787 | 0.007115674 |
| GO:0071331 | cellular response to hexose stimulus | 29/3028 | 0.001095923 | 0.007200547 |
| GO:0002791 | regulation of peptide secretion | 41/3028 | 0.001142658 | 0.007499177 |
| GO:0006734 | NADH metabolic process | Oct-28 | 0.001182757 | 0.007744938 |
| GO:1905898 | positive regulation of response to endoplasmic reticulum stress | Oct-28 | 0.001182757 | 0.007744938 |
| GO:0000255 | allantoin metabolic process | Jun-28 | 0.001194979 | 0.007781352 |
| GO:0009437 | carnitine metabolic process | Jun-28 | 0.001194979 | 0.007781352 |
| GO:0034374 | low-density lipoprotein particle remodeling | Jun-28 | 0.001194979 | 0.007781352 |
| GO:1901142 | insulin metabolic process | Jun-28 | 0.001194979 | 0.007781352 |
| GO:1901678 | iron coordination entity transport | Jun-28 | 0.001194979 | 0.007781352 |
| GO:0042181 | ketone biosynthetic process | Dec-28 | 0.001208772 | 0.007845991 |

| | | | | |
|---|---|---|---|---|
| GO:0048771 | tissue remodeling | 36/3028 | 0.001213882 | 0.007845991 |
| GO:0000303 | response to superoxide | Aug-28 | 0.001214309 | 0.007845991 |
| GO:0015865 | purine nucleotide transport | Aug-28 | 0.001214309 | 0.007845991 |
| GO:0034312 | diol biosynthetic process | Aug-28 | 0.001214309 | 0.007845991 |
| GO:0071398 | cellular response to fatty acid | Aug-28 | 0.001214309 | 0.007845991 |
| GO:0150104 | transport across blood-brain barrier | Aug-28 | 0.001214309 | 0.007845991 |
| GO:0070584 | mitochondrion morphogenesis | Sep-28 | 0.00123995 | 0.008002817 |
| GO:0043124 | negative regulation of I-kappaB kinase/NF-kappaB signaling | 13/3028 | 0.001284 | 0.008268845 |
| GO:0089718 | amino acid import across plasma membrane | 13/3028 | 0.001284 | 0.008268845 |
| GO:0046474 | glycerophospholipid biosynthetic process | 31/3028 | 0.00133245 | 0.008571413 |
| GO:0071322 | cellular response to carbohydrate stimulus | 30/3028 | 0.001345632 | 0.008646684 |
| GO:0032692 | negative regulation of interleukin-1 production | Nov-28 | 0.001379589 | 0.008855142 |
| GO:0050796 | regulation of insulin secretion | 34/3028 | 0.001393151 | 0.008930934 |
| GO:0090276 | regulation of peptide hormone secretion | 40/3028 | 0.001394455 | 0.008930934 |
| GO:0090087 | regulation of peptide transport | 41/3028 | 0.001459797 | 0.009339184 |
| GO:1902882 | regulation of response to oxidative stress | 21/3028 | 0.00148486 | 0.009489132 |
| GO:0046326 | positive regulation of glucose import | Dec-28 | 0.001505154 | 0.009608311 |
| GO:0032890 | regulation of organic acid transport | 19/3028 | 0.001508729 | 0.009620616 |

| | | | | |
|---|---|---|---|---|
| GO:0050999 | regulation of nitric-oxide synthase activity | Oct-28 | 0.001538901 | 0.009802312 |
| GO:0002921 | negative regulation of humoral immune response | Jul-28 | 0.001567195 | 0.009928399 |
| GO:0005980 | glycogen catabolic process | Jul-28 | 0.001567195 | 0.009928399 |
| GO:0016137 | glycoside metabolic process | Jul-28 | 0.001567195 | 0.009928399 |
| GO:1901570 | fatty acid derivative biosynthetic process | Jul-28 | 0.001567195 | 0.009928399 |
| GO:1903727 | positive regulation of phospholipid metabolic process | Jul-28 | 0.001567195 | 0.009928399 |

**GO terms associated with 2355 Rep_early_enriched_vs_Hep genes**

| ID | Description | GeneRatio | pvalue | p.adjust |
|---|---|---|---|---|
| GO:0030198 | extracellular matrix organization | 94/1966 | 1.36E-35 | 4.77E-32 |
| GO:0043062 | extracellular structure organization | 94/1966 | 1.82E-35 | 4.77E-32 |
| GO:0045229 | external encapsulating structure organization | 94/1966 | 2.42E-35 | 4.77E-32 |
| GO:0031589 | cell-substrate adhesion | 93/1966 | 1.50E-28 | 2.23E-25 |
| GO:0000280 | nuclear division | 96/1966 | 2.52E-24 | 2.99E-21 |
| GO:0140014 | mitotic nuclear division | 72/1966 | 3.85E-23 | 3.80E-20 |
| GO:0048285 | organelle fission | 99/1966 | 2.96E-22 | 2.50E-19 |
| GO:0007059 | chromosome segregation | 77/1966 | 4.25E-21 | 3.15E-18 |
| GO:0007015 | actin filament organization | 91/1966 | 1.54E-20 | 1.01E-17 |
| GO:0001667 | ameboidal-type cell migration | 86/1966 | 3.04E-18 | 1.80E-15 |
| GO:0007178 | transmembrane receptor protein serine/threonine kinase signaling pathway | 78/1966 | 6.78E-18 | 3.65E-15 |
| GO:0033674 | positive regulation of kinase activity | 83/1966 | 9.92E-18 | 4.89E-15 |
| GO:0007346 | regulation of mitotic cell cycle | 88/1966 | 1.22E-17 | 5.55E-15 |
| GO:0098813 | nuclear chromosome segregation | 63/1966 | 1.32E-17 | 5.57E-15 |
| GO:0045785 | positive regulation of cell adhesion | 88/1966 | 1.82E-17 | 7.20E-15 |
| GO:0000070 | mitotic sister chromatid segregation | 46/1966 | 2.81E-17 | 1.04E-14 |

| | | | | |
|---|---|---|---|---|
| GO:0030199 | collagen fibril organization | 27/1966 | 1.58E-16 | 5.49E-14 |
| GO:0060485 | mesenchyme development | 64/1966 | 1.77E-16 | 5.81E-14 |
| GO:0007409 | axonogenesis | 87/1966 | 2.38E-16 | 7.41E-14 |
| GO:0007160 | cell-matrix adhesion | 55/1966 | 3.26E-16 | 9.67E-14 |
| GO:0048762 | mesenchymal cell differentiation | 56/1966 | 3.68E-16 | 1.04E-13 |
| GO:0007264 | small GTPase mediated signal transduction | 79/1966 | 6.50E-16 | 1.70E-13 |
| GO:0032970 | regulation of actin filament-based process | 77/1966 | 6.62E-16 | 1.70E-13 |
| GO:0042060 | wound healing | 72/1966 | 9.05E-16 | 2.23E-13 |
| GO:0045860 | positive regulation of protein kinase activity | 70/1966 | 9.92E-16 | 2.31E-13 |
| GO:0031346 | positive regulation of cell projection organization | 80/1966 | 1.02E-15 | 2.31E-13 |
| GO:0000819 | sister chromatid segregation | 48/1966 | 1.24E-15 | 2.71E-13 |
| GO:0061448 | connective tissue development | 62/1966 | 1.30E-15 | 2.75E-13 |
| GO:0001655 | urogenital system development | 73/1966 | 1.75E-15 | 3.58E-13 |
| GO:0045787 | positive regulation of cell cycle | 69/1966 | 2.18E-15 | 4.30E-13 |
| GO:0032956 | regulation of actin cytoskeleton organization | 70/1966 | 3.21E-15 | 6.14E-13 |
| GO:0090092 | regulation of transmembrane receptor protein serine/threonine kinase signaling pathway | 56/1966 | 5.03E-15 | 9.26E-13 |
| GO:0085029 | extracellular matrix assembly | 22/1966 | 5.17E-15 | 9.26E-13 |
| GO:0060562 | epithelial tube morphogenesis | 73/1966 | 5.32E-15 | 9.26E-13 |

| | | | | |
|---|---|---|---|---|
| GO:0010810 | regulation of cell-substrate adhesion | 52/1966 | 6.46E-15 | 1.09E-12 |
| GO:0018108 | peptidyl-tyrosine phosphorylation | 64/1966 | 1.02E-14 | 1.68E-12 |
| GO:0090287 | regulation of cellular response to growth factor stimulus | 61/1966 | 1.09E-14 | 1.74E-12 |
| GO:0007265 | Ras protein signal transduction | 66/1966 | 1.27E-14 | 1.97E-12 |
| GO:0043254 | regulation of protein-containing complex assembly | 76/1966 | 1.52E-14 | 2.31E-12 |
| GO:0018212 | peptidyl-tyrosine modification | 64/1966 | 1.60E-14 | 2.37E-12 |
| GO:1902903 | regulation of supramolecular fiber organization | 71/1966 | 2.11E-14 | 3.04E-12 |
| GO:0001763 | morphogenesis of a branching structure | 54/1966 | 2.25E-14 | 3.12E-12 |
| GO:0034329 | cell junction assembly | 77/1966 | 2.27E-14 | 3.12E-12 |
| GO:0010634 | positive regulation of epithelial cell migration | 41/1966 | 3.43E-14 | 4.61E-12 |
| GO:0050767 | regulation of neurogenesis | 78/1966 | 4.22E-14 | 5.48E-12 |
| GO:0090130 | tissue migration | 62/1966 | 4.25E-14 | 5.48E-12 |
| GO:0044772 | mitotic cell cycle phase transition | 74/1966 | 4.52E-14 | 5.58E-12 |
| GO:0032963 | collagen metabolic process | 36/1966 | 4.52E-14 | 5.58E-12 |
| GO:0061138 | morphogenesis of a branching epithelium | 51/1966 | 4.66E-14 | 5.63E-12 |
| GO:0007179 | transforming growth factor beta receptor signaling pathway | 45/1966 | 5.21E-14 | 6.17E-12 |
| GO:1901342 | regulation of vasculature development | 61/1966 | 5.96E-14 | 6.93E-12 |
| GO:0022604 | regulation of cell morphogenesis | 64/1966 | 7.97E-14 | 8.99E-12 |

| | | | | |
|---|---|---|---|---|
| GO:0010631 | epithelial cell migration | 61/1966 | 8.04E-14 | 8.99E-12 |
| GO:0071560 | cellular response to transforming growth factor beta stimulus | 50/1966 | 9.04E-14 | 9.92E-12 |
| GO:0090132 | epithelium migration | 61/1966 | 1.08E-13 | 1.16E-11 |
| GO:0070371 | ERK1 and ERK2 cascade | 65/1966 | 1.14E-13 | 1.20E-11 |
| GO:1902850 | microtubule cytoskeleton organization involved in mitosis | 39/1966 | 1.31E-13 | 1.36E-11 |
| GO:0070372 | regulation of ERK1 and ERK2 cascade | 62/1966 | 1.59E-13 | 1.62E-11 |
| GO:0071559 | response to transforming growth factor beta | 50/1966 | 1.85E-13 | 1.86E-11 |
| GO:0072001 | renal system development | 64/1966 | 2.12E-13 | 2.09E-11 |
| GO:0010632 | regulation of epithelial cell migration | 51/1966 | 3.23E-13 | 3.14E-11 |
| GO:0045765 | regulation of angiogenesis | 59/1966 | 3.84E-13 | 3.66E-11 |
| GO:0007411 | axon guidance | 53/1966 | 3.90E-13 | 3.66E-11 |
| GO:0097485 | neuron projection guidance | 53/1966 | 4.58E-13 | 4.24E-11 |
| GO:0032964 | collagen biosynthetic process | 23/1966 | 7.39E-13 | 6.73E-11 |
| GO:0001822 | kidney development | 61/1966 | 9.07E-13 | 8.14E-11 |
| GO:0007266 | Rho protein signal transduction | 36/1966 | 1.56E-12 | 1.38E-10 |
| GO:0090068 | positive regulation of cell cycle process | 53/1966 | 1.61E-12 | 1.40E-10 |
| GO:0051495 | positive regulation of cytoskeleton organization | 45/1966 | 2.06E-12 | 1.77E-10 |
| GO:1902905 | positive regulation of supramolecular fiber organization | 42/1966 | 2.36E-12 | 2.00E-10 |

| | | | | |
|---|---|---|---|---|
| GO:0010959 | regulation of metal ion transport | 73/1966 | 2.84E-12 | 2.37E-10 |
| GO:0007051 | spindle organization | 42/1966 | 3.46E-12 | 2.82E-10 |
| GO:0007052 | mitotic spindle organization | 33/1966 | 3.48E-12 | 2.82E-10 |
| GO:0051056 | regulation of small GTPase mediated signal transduction | 48/1966 | 4.44E-12 | 3.56E-10 |
| GO:0090101 | negative regulation of transmembrane receptor protein serine/threonine kinase signaling pathway | 34/1966 | 4.72E-12 | 3.72E-10 |
| GO:0071900 | regulation of protein serine/threonine kinase activity | 62/1966 | 5.20E-12 | 4.05E-10 |
| GO:0043087 | regulation of GTPase activity | 61/1966 | 6.52E-12 | 5.02E-10 |
| GO:1901990 | regulation of mitotic cell cycle phase transition | 57/1966 | 7.20E-12 | 5.46E-10 |
| GO:1903034 | regulation of response to wounding | 40/1966 | 7.38E-12 | 5.53E-10 |
| GO:1901987 | regulation of cell cycle phase transition | 67/1966 | 8.68E-12 | 6.42E-10 |
| GO:0051962 | positive regulation of nervous system development | 62/1966 | 1.24E-11 | 9.04E-10 |
| GO:0048754 | branching morphogenesis of an epithelial tube | 42/1966 | 1.25E-11 | 9.04E-10 |
| GO:0051983 | regulation of chromosome segregation | 28/1966 | 1.47E-11 | 1.05E-09 |
| GO:0046578 | regulation of Ras protein signal transduction | 43/1966 | 1.56E-11 | 1.10E-09 |
| GO:1903829 | positive regulation of protein localization | 76/1966 | 1.69E-11 | 1.18E-09 |
| GO:0051090 | regulation of DNA-binding transcription factor activity | 70/1966 | 1.97E-11 | 1.36E-09 |
| GO:0051656 | establishment of organelle localization | 69/1966 | 2.04E-11 | 1.38E-09 |
| GO:0051258 | protein polymerization | 52/1966 | 2.05E-11 | 1.38E-09 |

| | | | | |
|---|---|---|---|---|
| GO:0010639 | negative regulation of organelle organization | 63/1966 | 2.26E-11 | 1.50E-09 |
| GO:0051017 | actin filament bundle assembly | 39/1966 | 2.54E-11 | 1.67E-09 |
| GO:0032535 | regulation of cellular component size | 65/1966 | 2.79E-11 | 1.81E-09 |
| GO:0001503 | ossification | 68/1966 | 2.90E-11 | 1.87E-09 |
| GO:1903046 | meiotic cell cycle process | 45/1966 | 3.10E-11 | 1.96E-09 |
| GO:0051216 | cartilage development | 44/1966 | 3.11E-11 | 1.96E-09 |
| GO:0040013 | negative regulation of locomotion | 59/1966 | 3.16E-11 | 1.97E-09 |
| GO:0045930 | negative regulation of mitotic cell cycle | 46/1966 | 3.58E-11 | 2.18E-09 |
| GO:0050920 | regulation of chemotaxis | 46/1966 | 3.58E-11 | 2.18E-09 |
| GO:0061572 | actin filament bundle organization | 39/1966 | 4.43E-11 | 2.67E-09 |
| GO:0050900 | leukocyte migration | 63/1966 | 5.06E-11 | 3.03E-09 |
| GO:0050730 | regulation of peptidyl-tyrosine phosphorylation | 50/1966 | 5.40E-11 | 3.20E-09 |
| GO:0010811 | positive regulation of cell-substrate adhesion | 34/1966 | 5.49E-11 | 3.22E-09 |
| GO:0048608 | reproductive structure development | 74/1966 | 5.63E-11 | 3.26E-09 |
| GO:0140694 | non-membrane-bounded organelle assembly | 63/1966 | 5.66E-11 | 3.26E-09 |
| GO:0001837 | epithelial to mesenchymal transition | 36/1966 | 5.95E-11 | 3.39E-09 |
| GO:0050673 | epithelial cell proliferation | 73/1966 | 6.57E-11 | 3.71E-09 |
| GO:0110053 | regulation of actin filament organization | 51/1966 | 6.65E-11 | 3.71E-09 |

| | | | | |
|---|---|---|---|---|
| GO:1901991 | negative regulation of mitotic cell cycle phase transition | 38/1966 | 7.15E-11 | 3.96E-09 |
| GO:0030512 | negative regulation of transforming growth factor beta receptor signaling pathway | 24/1966 | 7.77E-11 | 4.26E-09 |
| GO:0002685 | regulation of leukocyte migration | 45/1966 | 7.93E-11 | 4.31E-09 |
| GO:0061458 | reproductive system development | 74/1966 | 8.31E-11 | 4.47E-09 |
| GO:0002687 | positive regulation of leukocyte migration | 36/1966 | 8.77E-11 | 4.64E-09 |
| GO:0008360 | regulation of cell shape | 36/1966 | 8.77E-11 | 4.64E-09 |
| GO:0050727 | regulation of inflammatory response | 59/1966 | 1.18E-10 | 6.17E-09 |
| GO:0048251 | elastic fiber assembly | Oct-66 | 1.24E-10 | 6.46E-09 |
| GO:0050808 | synapse organization | 74/1966 | 1.47E-10 | 7.60E-09 |
| GO:0042692 | muscle cell differentiation | 68/1966 | 1.72E-10 | 8.77E-09 |
| GO:0003007 | heart morphogenesis | 51/1966 | 1.93E-10 | 9.77E-09 |
| GO:0007162 | negative regulation of cell adhesion | 54/1966 | 2.67E-10 | 1.34E-08 |
| GO:0010720 | positive regulation of cell development | 61/1966 | 3.11E-10 | 1.55E-08 |
| GO:0060537 | muscle tissue development | 73/1966 | 3.34E-10 | 1.65E-08 |
| GO:0050769 | positive regulation of neurogenesis | 52/1966 | 3.75E-10 | 1.83E-08 |
| GO:0051304 | chromosome separation | 27/1966 | 4.02E-10 | 1.94E-08 |
| GO:0060560 | developmental growth involved in morphogenesis | 50/1966 | 4.04E-10 | 1.94E-08 |
| GO:0017015 | regulation of transforming growth factor beta receptor signaling pathway | 30/1966 | 4.26E-10 | 2.03E-08 |

| | | | | |
|---|---|---|---|---|
| GO:0051783 | regulation of nuclear division | 33/1966 | 4.41E-10 | 2.09E-08 |
| GO:0022407 | regulation of cell-cell adhesion | 70/1966 | 4.93E-10 | 2.32E-08 |
| GO:0034765 | regulation of ion transmembrane transport | 73/1966 | 5.27E-10 | 2.46E-08 |
| GO:0071674 | mononuclear cell migration | 38/1966 | 5.75E-10 | 2.66E-08 |
| GO:0045216 | cell-cell junction organization | 42/1966 | 6.24E-10 | 2.87E-08 |
| GO:0050731 | positive regulation of peptidyl-tyrosine phosphorylation | 39/1966 | 6.76E-10 | 3.08E-08 |
| GO:0051310 | metaphase plate congression | 21/1966 | 6.96E-10 | 3.15E-08 |
| GO:0050678 | regulation of epithelial cell proliferation | 63/1966 | 7.38E-10 | 3.30E-08 |
| GO:0048588 | developmental cell growth | 49/1966 | 7.41E-10 | 3.30E-08 |
| GO:1903844 | regulation of cellular response to transforming growth factor beta stimulus | 30/1966 | 8.06E-10 | 3.56E-08 |
| GO:0030177 | positive regulation of Wnt signaling pathway | 32/1966 | 8.61E-10 | 3.78E-08 |
| GO:0140013 | meiotic nuclear division | 40/1966 | 9.05E-10 | 3.94E-08 |
| GO:0032965 | regulation of collagen biosynthetic process | 18/1966 | 9.62E-10 | 4.16E-08 |
| GO:0001570 | vasculogenesis | 26/1966 | 1.00E-09 | 4.30E-08 |
| GO:0048732 | gland development | 70/1966 | 1.04E-09 | 4.41E-08 |
| GO:0043542 | endothelial cell migration | 43/1966 | 1.06E-09 | 4.47E-08 |
| GO:0071772 | response to BMP | 38/1966 | 1.10E-09 | 4.59E-08 |
| GO:0071773 | cellular response to BMP stimulus | 38/1966 | 1.10E-09 | 4.59E-08 |

| | | | | |
|---|---|---|---|---|
| GO:0061041 | regulation of wound healing | 31/1966 | 1.14E-09 | 4.71E-08 |
| GO:0003012 | muscle system process | 65/1966 | 1.32E-09 | 5.43E-08 |
| GO:0022612 | gland morphogenesis | 34/1966 | 1.44E-09 | 5.89E-08 |
| GO:0030323 | respiratory tube development | 44/1966 | 1.49E-09 | 6.06E-08 |
| GO:0001558 | regulation of cell growth | 68/1966 | 1.57E-09 | 6.34E-08 |
| GO:1901343 | negative regulation of vasculature development | 28/1966 | 1.63E-09 | 6.54E-08 |
| GO:0010712 | regulation of collagen metabolic process | 19/1966 | 1.67E-09 | 6.64E-08 |
| GO:0008154 | actin polymerization or depolymerization | 39/1966 | 1.72E-09 | 6.77E-08 |
| GO:0050921 | positive regulation of chemotaxis | 33/1966 | 1.96E-09 | 7.68E-08 |
| GO:0007611 | learning or memory | 53/1966 | 2.19E-09 | 8.55E-08 |
| GO:0001952 | regulation of cell-matrix adhesion | 30/1966 | 2.22E-09 | 8.59E-08 |
| GO:0042063 | gliogenesis | 56/1966 | 2.26E-09 | 8.70E-08 |
| GO:0051271 | negative regulation of cellular component movement | 52/1966 | 2.50E-09 | 9.54E-08 |
| GO:0032271 | regulation of protein polymerization | 40/1966 | 2.56E-09 | 9.73E-08 |
| GO:0003018 | vascular process in circulatory system | 43/1966 | 2.77E-09 | 1.05E-07 |
| GO:0070997 | neuron death | 65/1966 | 2.81E-09 | 1.05E-07 |
| GO:0030041 | actin filament polymerization | 35/1966 | 2.92E-09 | 1.08E-07 |
| GO:0032103 | positive regulation of response to external stimulus | 67/1966 | 2.93E-09 | 1.08E-07 |

| | | | | |
|---|---|---|---|---|
| GO:0030324 | lung development | 43/1966 | 3.17E-09 | 1.16E-07 |
| GO:0010717 | regulation of epithelial to mesenchymal transition | 25/1966 | 3.17E-09 | 1.16E-07 |
| GO:0010595 | positive regulation of endothelial cell migration | 27/1966 | 3.20E-09 | 1.16E-07 |
| GO:1903706 | regulation of hemopoiesis | 61/1966 | 3.63E-09 | 1.31E-07 |
| GO:0062197 | cellular response to chemical stress | 54/1966 | 3.70E-09 | 1.33E-07 |
| GO:0016055 | Wnt signaling pathway | 66/1966 | 3.78E-09 | 1.35E-07 |
| GO:0007088 | regulation of mitotic nuclear division | 27/1966 | 3.97E-09 | 1.40E-07 |
| GO:1905818 | regulation of chromosome separation | 22/1966 | 3.98E-09 | 1.40E-07 |
| GO:0071604 | transforming growth factor beta production | 16/1966 | 4.16E-09 | 1.46E-07 |
| GO:1901988 | negative regulation of cell cycle phase transition | 44/1966 | 4.29E-09 | 1.50E-07 |
| GO:0198738 | cell-cell signaling by wnt | 66/1966 | 4.52E-09 | 1.57E-07 |
| GO:0006814 | sodium ion transport | 44/1966 | 4.88E-09 | 1.68E-07 |
| GO:0098742 | cell-cell adhesion via plasma-membrane adhesion molecules | 40/1966 | 5.21E-09 | 1.78E-07 |
| GO:0031334 | positive regulation of protein-containing complex assembly | 39/1966 | 5.55E-09 | 1.89E-07 |
| GO:0060541 | respiratory system development | 46/1966 | 5.76E-09 | 1.95E-07 |
| GO:0050000 | chromosome localization | 23/1966 | 5.83E-09 | 1.96E-07 |
| GO:2000181 | negative regulation of blood vessel morphogenesis | 27/1966 | 6.05E-09 | 2.02E-07 |
| GO:0035023 | regulation of Rho protein signal transduction | 24/1966 | 6.21E-09 | 2.06E-07 |

| | | | | |
|---|---|---|---|---|
| GO:0010594 | regulation of endothelial cell migration | 35/1966 | 6.51E-09 | 2.15E-07 |
| GO:0010934 | macrophage cytokine production | Nov-66 | 6.58E-09 | 2.16E-07 |
| GO:0001818 | negative regulation of cytokine production | 49/1966 | 7.22E-09 | 2.36E-07 |
| GO:0010976 | positive regulation of neuron projection development | 41/1966 | 7.27E-09 | 2.37E-07 |
| GO:0043491 | protein kinase B signaling | 39/1966 | 7.36E-09 | 2.38E-07 |
| GO:0044839 | cell cycle G2/M phase transition | 32/1966 | 7.60E-09 | 2.45E-07 |
| GO:0043405 | regulation of MAP kinase activity | 38/1966 | 7.82E-09 | 2.50E-07 |
| GO:0000086 | G2/M transition of mitotic cell cycle | 30/1966 | 8.32E-09 | 2.63E-07 |
| GO:0007093 | mitotic cell cycle checkpoint signaling | 30/1966 | 8.32E-09 | 2.63E-07 |
| GO:1904019 | epithelial cell apoptotic process | 28/1966 | 8.46E-09 | 2.67E-07 |
| GO:0009314 | response to radiation | 63/1966 | 8.88E-09 | 2.77E-07 |
| GO:0030509 | BMP signaling pathway | 35/1966 | 8.88E-09 | 2.77E-07 |
| GO:0048638 | regulation of developmental growth | 60/1966 | 9.12E-09 | 2.83E-07 |
| GO:0070374 | positive regulation of ERK1 and ERK2 cascade | 41/1966 | 9.48E-09 | 2.92E-07 |
| GO:0061640 | cytoskeleton-dependent cytokinesis | 24/1966 | 9.91E-09 | 3.02E-07 |
| GO:1904035 | regulation of epithelial cell apoptotic process | 24/1966 | 9.91E-09 | 3.02E-07 |
| GO:0032102 | negative regulation of response to external stimulus | 60/1966 | 1.00E-08 | 3.04E-07 |
| GO:0030866 | cortical actin cytoskeleton organization | 15/1966 | 1.06E-08 | 3.19E-07 |

| | | | | |
|---|---|---|---|---|
| GO:0030010 | establishment of cell polarity | 32/1966 | 1.07E-08 | 3.20E-07 |
| GO:0002062 | chondrocyte differentiation | 27/1966 | 1.11E-08 | 3.33E-07 |
| GO:0050890 | cognition | 55/1966 | 1.50E-08 | 4.46E-07 |
| GO:0048659 | smooth muscle cell proliferation | 36/1966 | 1.55E-08 | 4.59E-07 |
| GO:0043588 | skin development | 51/1966 | 1.63E-08 | 4.81E-07 |
| GO:1990778 | protein localization to cell periphery | 56/1966 | 1.72E-08 | 5.04E-07 |
| GO:0016525 | negative regulation of angiogenesis | 26/1966 | 1.78E-08 | 5.21E-07 |
| GO:0051303 | establishment of chromosome localization | 22/1966 | 1.87E-08 | 5.43E-07 |
| GO:0033002 | muscle cell proliferation | 44/1966 | 1.89E-08 | 5.45E-07 |
| GO:0031345 | negative regulation of cell projection organization | 39/1966 | 1.91E-08 | 5.48E-07 |
| GO:2000146 | negative regulation of cell motility | 49/1966 | 1.95E-08 | 5.58E-07 |
| GO:0045786 | negative regulation of cell cycle | 56/1966 | 2.08E-08 | 5.93E-07 |
| GO:1901214 | regulation of neuron death | 58/1966 | 2.20E-08 | 6.24E-07 |
| GO:0046847 | filopodium assembly | 21/1966 | 2.22E-08 | 6.27E-07 |
| GO:0030510 | regulation of BMP signaling pathway | 25/1966 | 2.32E-08 | 6.52E-07 |
| GO:0050679 | positive regulation of epithelial cell proliferation | 39/1966 | 2.48E-08 | 6.93E-07 |
| GO:0021700 | developmental maturation | 54/1966 | 2.59E-08 | 7.19E-07 |
| GO:0048144 | fibroblast proliferation | 26/1966 | 2.66E-08 | 7.35E-07 |

| | | | | |
|---|---|---|---|---|
| GO:0010935 | regulation of macrophage cytokine production | Oct-66 | 2.67E-08 | 7.35E-07 |
| GO:0007517 | muscle organ development | 55/1966 | 2.68E-08 | 7.35E-07 |
| GO:0030111 | regulation of Wnt signaling pathway | 50/1966 | 2.86E-08 | 7.81E-07 |
| GO:0014812 | muscle cell migration | 27/1966 | 2.93E-08 | 7.95E-07 |
| GO:0051346 | negative regulation of hydrolase activity | 55/1966 | 2.95E-08 | 7.98E-07 |
| GO:0001656 | metanephros development | 23/1966 | 3.06E-08 | 8.24E-07 |
| GO:0045807 | positive regulation of endocytosis | 28/1966 | 3.13E-08 | 8.39E-07 |
| GO:0048146 | positive regulation of fibroblast proliferation | 20/1966 | 3.35E-08 | 8.94E-07 |
| GO:0030100 | regulation of endocytosis | 41/1966 | 3.37E-08 | 8.94E-07 |
| GO:0014909 | smooth muscle cell migration | 25/1966 | 3.49E-08 | 9.24E-07 |
| GO:0030336 | negative regulation of cell migration | 47/1966 | 3.55E-08 | 9.34E-07 |
| GO:1901888 | regulation of cell junction assembly | 39/1966 | 3.65E-08 | 9.57E-07 |
| GO:0030048 | actin filament-based movement | 28/1966 | 3.74E-08 | 9.76E-07 |
| GO:0048771 | tissue remodeling | 37/1966 | 3.84E-08 | 9.97E-07 |
| GO:0031532 | actin cytoskeleton reorganization | 26/1966 | 3.91E-08 | 1.01E-06 |
| GO:0032967 | positive regulation of collagen biosynthetic process | 13/1966 | 4.30E-08 | 1.11E-06 |
| GO:0061035 | regulation of cartilage development | 20/1966 | 4.34E-08 | 1.11E-06 |
| GO:0060070 | canonical Wnt signaling pathway | 48/1966 | 4.71E-08 | 1.20E-06 |

| | | | | |
|---|---|---|---|---|
| GO:0001764 | neuron migration | 36/1966 | 4.75E-08 | 1.21E-06 |
| GO:0006936 | muscle contraction | 49/1966 | 5.00E-08 | 1.27E-06 |
| GO:0051383 | kinetochore organization | Nov-66 | 5.07E-08 | 1.28E-06 |
| GO:0051924 | regulation of calcium ion transport | 46/1966 | 5.63E-08 | 1.41E-06 |
| GO:0051145 | smooth muscle cell differentiation | 21/1966 | 5.93E-08 | 1.48E-06 |
| GO:0031099 | regeneration | 31/1966 | 6.25E-08 | 1.56E-06 |
| GO:1901215 | negative regulation of neuron death | 43/1966 | 6.77E-08 | 1.67E-06 |
| GO:0016358 | dendrite development | 49/1966 | 6.77E-08 | 1.67E-06 |
| GO:0010948 | negative regulation of cell cycle process | 46/1966 | 6.96E-08 | 1.71E-06 |
| GO:0008064 | regulation of actin polymerization or depolymerization | 32/1966 | 7.02E-08 | 1.72E-06 |
| GO:0072006 | nephron development | 31/1966 | 7.29E-08 | 1.78E-06 |
| GO:0002011 | morphogenesis of an epithelial sheet | 18/1966 | 7.34E-08 | 1.78E-06 |
| GO:0014706 | striated muscle tissue development | 46/1966 | 7.74E-08 | 1.87E-06 |
| GO:0003205 | cardiac chamber development | 36/1966 | 8.10E-08 | 1.95E-06 |
| GO:0071675 | regulation of mononuclear cell migration | 26/1966 | 8.23E-08 | 1.97E-06 |
| GO:0045931 | positive regulation of mitotic cell cycle | 28/1966 | 8.83E-08 | 2.11E-06 |
| GO:0071634 | regulation of transforming growth factor beta production | 14/1966 | 9.19E-08 | 2.19E-06 |
| GO:0051961 | negative regulation of nervous system development | 32/1966 | 9.41E-08 | 2.23E-06 |

| | | | | |
|---|---|---|---|---|
| GO:0045444 | fat cell differentiation | 43/1966 | 9.45E-08 | 2.23E-06 |
| GO:0007224 | smoothened signaling pathway | 31/1966 | 9.86E-08 | 2.32E-06 |
| GO:0006979 | response to oxidative stress | 58/1966 | 9.96E-08 | 2.33E-06 |
| GO:0090288 | negative regulation of cellular response to growth factor stimulus | 24/1966 | 1.02E-07 | 2.37E-06 |
| GO:0051896 | regulation of protein kinase B signaling | 33/1966 | 1.02E-07 | 2.38E-06 |
| GO:0050919 | negative chemotaxis | 15/1966 | 1.06E-07 | 2.46E-06 |
| GO:0006869 | lipid transport | 56/1966 | 1.08E-07 | 2.48E-06 |
| GO:0030832 | regulation of actin filament length | 32/1966 | 1.09E-07 | 2.50E-06 |
| GO:0007163 | establishment or maintenance of cell polarity | 39/1966 | 1.11E-07 | 2.53E-06 |
| GO:1904062 | regulation of cation transmembrane transport | 55/1966 | 1.17E-07 | 2.66E-06 |
| GO:0043271 | negative regulation of ion transport | 33/1966 | 1.18E-07 | 2.66E-06 |
| GO:0001654 | eye development | 59/1966 | 1.18E-07 | 2.66E-06 |
| GO:0003206 | cardiac chamber morphogenesis | 30/1966 | 1.19E-07 | 2.66E-06 |
| GO:0031333 | negative regulation of protein-containing complex assembly | 30/1966 | 1.19E-07 | 2.66E-06 |
| GO:0030865 | cortical cytoskeleton organization | 18/1966 | 1.27E-07 | 2.84E-06 |
| GO:0007613 | memory | 31/1966 | 1.32E-07 | 2.94E-06 |
| GO:0070507 | regulation of microtubule cytoskeleton organization | 31/1966 | 1.32E-07 | 2.94E-06 |
| GO:0007229 | integrin-mediated signaling pathway | 23/1966 | 1.34E-07 | 2.96E-06 |

| | | | | |
|---|---|---|---|---|
| GO:199 0138 | neuron projection extension | 36/196 6 | 1.35E- 07 | 2.98E- 06 |
| GO:003 1032 | actomyosin structure organization | 37/196 6 | 1.38E- 07 | 3.04E- 06 |
| GO:005 1321 | meiotic cell cycle | 50/196 6 | 1.39E- 07 | 3.04E- 06 |
| GO:015 0063 | visual system development | 59/196 6 | 1.51E- 07 | 3.28E- 06 |
| GO:001 0977 | negative regulation of neuron projection development | 31/196 6 | 1.53E- 07 | 3.32E- 06 |
| GO:005 1984 | positive regulation of chromosome segregation | Nov- 66 | 1.58E- 07 | 3.41E- 06 |
| GO:001 0714 | positive regulation of collagen metabolic process | 13/196 6 | 1.59E- 07 | 3.43E- 06 |
| GO:000 8347 | glial cell migration | 18/196 6 | 1.66E- 07 | 3.55E- 06 |
| GO:003 0833 | regulation of actin filament polymerization | 29/196 6 | 1.66E- 07 | 3.55E- 06 |
| GO:001 0721 | negative regulation of cell development | 36/196 6 | 1.74E- 07 | 3.71E- 06 |
| GO:009 7191 | extrinsic apoptotic signaling pathway | 40/196 6 | 1.75E- 07 | 3.72E- 06 |
| GO:005 0768 | negative regulation of neurogenesis | 31/196 6 | 1.77E- 07 | 3.73E- 06 |
| GO:004 8660 | regulation of smooth muscle cell proliferation | 33/196 6 | 1.77E- 07 | 3.73E- 06 |
| GO:000 7018 | microtubule-based movement | 59/196 6 | 1.78E- 07 | 3.73E- 06 |
| GO:007 2697 | protein localization to cell cortex | Aug- 66 | 1.79E- 07 | 3.74E- 06 |
| GO:004 8661 | positive regulation of smooth muscle cell proliferation | 24/196 6 | 1.80E- 07 | 3.75E- 06 |
| GO:001 0563 | negative regulation of phosphorus metabolic process | 62/196 6 | 1.84E- 07 | 3.80E- 06 |
| GO:004 5936 | negative regulation of phosphate metabolic process | 62/196 6 | 1.84E- 07 | 3.80E- 06 |

| | | | | |
|---|---|---|---|---|
| GO:0060249 | anatomical structure homeostasis | 49/1966 | 1.97E-07 | 4.07E-06 |
| GO:0010389 | regulation of G2/M transition of mitotic cell cycle | 23/1966 | 1.99E-07 | 4.09E-06 |
| GO:0001666 | response to hypoxia | 37/1966 | 2.00E-07 | 4.09E-06 |
| GO:0048880 | sensory system development | 59/1966 | 2.09E-07 | 4.26E-06 |
| GO:0090263 | positive regulation of canonical Wnt signaling pathway | 24/1966 | 2.16E-07 | 4.40E-06 |
| GO:0008608 | attachment of spindle microtubules to kinetochore | 13/1966 | 2.38E-07 | 4.82E-06 |
| GO:0009914 | hormone transport | 54/1966 | 2.54E-07 | 5.12E-06 |
| GO:0006310 | DNA recombination | 48/1966 | 2.54E-07 | 5.12E-06 |
| GO:0048145 | regulation of fibroblast proliferation | 24/1966 | 2.60E-07 | 5.21E-06 |
| GO:0072659 | protein localization to plasma membrane | 46/1966 | 2.61E-07 | 5.22E-06 |
| GO:0045766 | positive regulation of angiogenesis | 33/1966 | 2.64E-07 | 5.25E-06 |
| GO:1904018 | positive regulation of vasculature development | 33/1966 | 2.64E-07 | 5.25E-06 |
| GO:0034508 | centromere complex assembly | Dec-66 | 2.67E-07 | 5.30E-06 |
| GO:0051225 | spindle assembly | 25/1966 | 2.72E-07 | 5.36E-06 |
| GO:0034599 | cellular response to oxidative stress | 43/1966 | 2.75E-07 | 5.40E-06 |
| GO:0045926 | negative regulation of growth | 42/1966 | 2.86E-07 | 5.61E-06 |
| GO:0048678 | response to axon injury | 19/1966 | 2.90E-07 | 5.66E-06 |
| GO:0034446 | substrate adhesion-dependent cell spreading | 24/1966 | 3.11E-07 | 6.05E-06 |

| | | | | |
|---|---|---|---|---|
| GO:0048675 | axon extension | 28/1966 | 3.16E-07 | 6.13E-06 |
| GO:0014910 | regulation of smooth muscle cell migration | 22/1966 | 3.20E-07 | 6.20E-06 |
| GO:0035924 | cellular response to vascular endothelial growth factor stimulus | 17/1966 | 3.23E-07 | 6.23E-06 |
| GO:1903532 | positive regulation of secretion by cell | 52/1966 | 3.26E-07 | 6.27E-06 |
| GO:0060326 | cell chemotaxis | 47/1966 | 3.28E-07 | 6.28E-06 |
| GO:0032680 | regulation of tumor necrosis factor production | 33/1966 | 3.43E-07 | 6.55E-06 |
| GO:0003231 | cardiac ventricle development | 29/1966 | 3.51E-07 | 6.69E-06 |
| GO:0000281 | mitotic cytokinesis | 18/1966 | 3.53E-07 | 6.69E-06 |
| GO:0030193 | regulation of blood coagulation | 18/1966 | 3.53E-07 | 6.69E-06 |
| GO:0001894 | tissue homeostasis | 44/1966 | 3.55E-07 | 6.69E-06 |
| GO:0003208 | cardiac ventricle morphogenesis | 20/1966 | 3.61E-07 | 6.78E-06 |
| GO:0060828 | regulation of canonical Wnt signaling pathway | 40/1966 | 3.79E-07 | 7.10E-06 |
| GO:0006820 | anion transport | 65/1966 | 3.87E-07 | 7.20E-06 |
| GO:0071174 | mitotic spindle checkpoint signaling | 14/1966 | 3.88E-07 | 7.20E-06 |
| GO:0007080 | mitotic metaphase plate congression | 15/1966 | 3.89E-07 | 7.20E-06 |
| GO:0010718 | positive regulation of epithelial to mesenchymal transition | 15/1966 | 3.89E-07 | 7.20E-06 |
| GO:0002274 | myeloid leukocyte activation | 42/1966 | 3.91E-07 | 7.21E-06 |
| GO:0061082 | myeloid leukocyte cytokine production | Dec-66 | 4.09E-07 | 7.53E-06 |

| | | | | |
|---|---|---|---|---|
| GO:0071692 | protein localization to extracellular region | 56/1966 | 4.13E-07 | 7.58E-06 |
| GO:0007091 | metaphase/anaphase transition of mitotic cell cycle | 17/1966 | 4.20E-07 | 7.67E-06 |
| GO:0090307 | mitotic spindle assembly | 18/1966 | 4.50E-07 | 8.20E-06 |
| GO:0051302 | regulation of cell division | 34/1966 | 4.52E-07 | 8.21E-06 |
| GO:0051098 | regulation of binding | 55/1966 | 4.53E-07 | 8.21E-06 |
| GO:0043433 | negative regulation of DNA-binding transcription factor activity | 31/1966 | 4.67E-07 | 8.43E-06 |
| GO:0002718 | regulation of cytokine production involved in immune response | 22/1966 | 4.71E-07 | 8.47E-06 |
| GO:0051402 | neuron apoptotic process | 47/1966 | 4.76E-07 | 8.54E-06 |
| GO:0010876 | lipid localization | 60/1966 | 4.84E-07 | 8.66E-06 |
| GO:0051146 | striated muscle cell differentiation | 48/1966 | 4.86E-07 | 8.66E-06 |
| GO:1903555 | regulation of tumor necrosis factor superfamily cytokine production | 33/1966 | 5.03E-07 | 8.94E-06 |
| GO:0071526 | semaphorin-plexin signaling pathway | 15/1966 | 5.26E-07 | 9.32E-06 |
| GO:0006816 | calcium ion transport | 59/1966 | 5.36E-07 | 9.47E-06 |
| GO:0010965 | regulation of mitotic sister chromatid separation | 17/1966 | 5.41E-07 | 9.54E-06 |
| GO:0070252 | actin-mediated cell contraction | 22/1966 | 5.68E-07 | 9.93E-06 |
| GO:0033045 | regulation of sister chromatid segregation | 18/1966 | 5.70E-07 | 9.93E-06 |
| GO:0051926 | negative regulation of calcium ion transport | 18/1966 | 5.70E-07 | 9.93E-06 |
| GO:1900046 | regulation of hemostasis | 18/1966 | 5.70E-07 | 9.93E-06 |

| | | | | |
|---|---|---|---|---|
| GO:1903036 | positive regulation of response to wounding | 19/1966 | 5.72E-07 | 9.93E-06 |
| GO:0042326 | negative regulation of phosphorylation | 55/1966 | 5.78E-07 | 1.00E-05 |
| GO:0071902 | positive regulation of protein serine/threonine kinase activity | 35/1966 | 5.80E-07 | 1.00E-05 |
| GO:0052547 | regulation of peptidase activity | 59/1966 | 6.72E-07 | 1.16E-05 |
| GO:0002690 | positive regulation of leukocyte chemotaxis | 22/1966 | 6.84E-07 | 1.17E-05 |
| GO:1905517 | macrophage migration | 17/1966 | 6.95E-07 | 1.19E-05 |
| GO:0061097 | regulation of protein tyrosine kinase activity | 19/1966 | 7.10E-07 | 1.21E-05 |
| GO:0032640 | tumor necrosis factor production | 33/1966 | 7.29E-07 | 1.24E-05 |
| GO:1905475 | regulation of protein localization to membrane | 34/1966 | 7.34E-07 | 1.24E-05 |
| GO:0003014 | renal system process | 25/1966 | 7.36E-07 | 1.24E-05 |
| GO:0002367 | cytokine production involved in immune response | 24/1966 | 7.36E-07 | 1.24E-05 |
| GO:0061098 | positive regulation of protein tyrosine kinase activity | 14/1966 | 7.41E-07 | 1.25E-05 |
| GO:1902105 | regulation of leukocyte differentiation | 47/1966 | 7.51E-07 | 1.26E-05 |
| GO:0048562 | embryonic organ morphogenesis | 48/1966 | 7.60E-07 | 1.27E-05 |
| GO:0050878 | regulation of body fluid levels | 52/1966 | 7.61E-07 | 1.27E-05 |
| GO:0018209 | peptidyl-serine modification | 51/1966 | 8.32E-07 | 1.38E-05 |
| GO:0006898 | receptor-mediated endocytosis | 41/1966 | 8.33E-07 | 1.38E-05 |
| GO:0002697 | regulation of immune effector process | 58/1966 | 8.66E-07 | 1.43E-05 |

| | | | | |
|---|---|---|---|---|
| GO:0060840 | artery development | 24/1966 | 8.69E-07 | 1.43E-05 |
| GO:0044784 | metaphase/anaphase transition of cell cycle | 17/1966 | 8.86E-07 | 1.46E-05 |
| GO:0000075 | cell cycle checkpoint signaling | 31/1966 | 8.98E-07 | 1.47E-05 |
| GO:0050818 | regulation of coagulation | 18/1966 | 9.02E-07 | 1.48E-05 |
| GO:0060271 | cilium assembly | 50/1966 | 9.08E-07 | 1.48E-05 |
| GO:0060348 | bone development | 40/1966 | 9.69E-07 | 1.58E-05 |
| GO:0032147 | activation of protein kinase activity | 22/1966 | 9.82E-07 | 1.59E-05 |
| GO:0001569 | branching involved in blood vessel morphogenesis | 14/1966 | 1.01E-06 | 1.62E-05 |
| GO:0031577 | spindle checkpoint signaling | 14/1966 | 1.01E-06 | 1.62E-05 |
| GO:1905521 | regulation of macrophage migration | 14/1966 | 1.01E-06 | 1.62E-05 |
| GO:0032886 | regulation of microtubule-based process | 41/1966 | 1.02E-06 | 1.63E-05 |
| GO:0050770 | regulation of axonogenesis | 32/1966 | 1.04E-06 | 1.66E-05 |
| GO:0071706 | tumor necrosis factor superfamily cytokine production | 33/1966 | 1.05E-06 | 1.67E-05 |
| GO:0032273 | positive regulation of protein polymerization | 21/1966 | 1.11E-06 | 1.77E-05 |
| GO:0022409 | positive regulation of cell-cell adhesion | 44/1966 | 1.12E-06 | 1.77E-05 |
| GO:0032835 | glomerulus development | 17/1966 | 1.12E-06 | 1.77E-05 |
| GO:0051306 | mitotic sister chromatid separation | 17/1966 | 1.12E-06 | 1.77E-05 |
| GO:0045664 | regulation of neuron differentiation | 38/1966 | 1.17E-06 | 1.85E-05 |

| | | | | |
|---|---|---|---|---|
| GO:0006022 | aminoglycan metabolic process | 23/1966 | 1.20E-06 | 1.89E-05 |
| GO:0045839 | negative regulation of mitotic nuclear division | 15/1966 | 1.23E-06 | 1.93E-05 |
| GO:0050729 | positive regulation of inflammatory response | 27/1966 | 1.24E-06 | 1.94E-05 |
| GO:0014031 | mesenchymal cell development | 21/1966 | 1.33E-06 | 2.07E-05 |
| GO:0072073 | kidney epithelium development | 28/1966 | 1.33E-06 | 2.07E-05 |
| GO:0071695 | anatomical structure maturation | 44/1966 | 1.35E-06 | 2.09E-05 |
| GO:0044843 | cell cycle G1/S phase transition | 37/1966 | 1.36E-06 | 2.09E-05 |
| GO:0051985 | negative regulation of chromosome segregation | 14/1966 | 1.36E-06 | 2.09E-05 |
| GO:0007094 | mitotic spindle assembly checkpoint signaling | 13/1966 | 1.40E-06 | 2.15E-05 |
| GO:0071173 | spindle assembly checkpoint signaling | 13/1966 | 1.40E-06 | 2.15E-05 |
| GO:1902749 | regulation of cell cycle G2/M phase transition | 23/1966 | 1.42E-06 | 2.17E-05 |
| GO:0036293 | response to decreased oxygen levels | 38/1966 | 1.44E-06 | 2.20E-05 |
| GO:0048015 | phosphatidylinositol-mediated signaling | 30/1966 | 1.45E-06 | 2.20E-05 |
| GO:2001236 | regulation of extrinsic apoptotic signaling pathway | 30/1966 | 1.45E-06 | 2.20E-05 |
| GO:0051222 | positive regulation of protein transport | 46/1966 | 1.51E-06 | 2.28E-05 |
| GO:0061982 | meiosis I cell cycle process | 26/1966 | 1.52E-06 | 2.30E-05 |
| GO:0046879 | hormone secretion | 51/1966 | 1.60E-06 | 2.41E-05 |
| GO:2000351 | regulation of endothelial cell apoptotic process | 15/1966 | 1.61E-06 | 2.42E-05 |

| | | | | |
|---|---|---|---|---|
| GO:0001933 | negative regulation of protein phosphorylation | 49/1966 | 1.64E-06 | 2.45E-05 |
| GO:0002064 | epithelial cell development | 37/1966 | 1.67E-06 | 2.50E-05 |
| GO:0043409 | negative regulation of MAPK cascade | 32/1966 | 1.68E-06 | 2.51E-05 |
| GO:0043523 | regulation of neuron apoptotic process | 42/1966 | 1.70E-06 | 2.53E-05 |
| GO:1903510 | mucopolysaccharide metabolic process | 18/1966 | 1.73E-06 | 2.57E-05 |
| GO:0033044 | regulation of chromosome organization | 34/1966 | 1.84E-06 | 2.73E-05 |
| GO:0002028 | regulation of sodium ion transport | 21/1966 | 1.91E-06 | 2.82E-05 |
| GO:0030516 | regulation of axon extension | 23/1966 | 1.96E-06 | 2.89E-05 |
| GO:0014015 | positive regulation of gliogenesis | 19/1966 | 1.99E-06 | 2.92E-05 |
| GO:0007043 | cell-cell junction assembly | 28/1966 | 1.99E-06 | 2.92E-05 |
| GO:2001233 | regulation of apoptotic signaling pathway | 54/1966 | 2.05E-06 | 2.99E-05 |
| GO:0043547 | positive regulation of GTPase activity | 39/1966 | 2.05E-06 | 2.99E-05 |
| GO:0042391 | regulation of membrane potential | 60/1966 | 2.06E-06 | 3.00E-05 |
| GO:0048017 | inositol lipid-mediated signaling | 30/1966 | 2.11E-06 | 3.07E-05 |
| GO:0000910 | cytokinesis | 31/1966 | 2.13E-06 | 3.09E-05 |
| GO:0043406 | positive regulation of MAP kinase activity | 25/1966 | 2.15E-06 | 3.10E-05 |
| GO:0060324 | face development | 16/1966 | 2.22E-06 | 3.19E-05 |
| GO:0001935 | endothelial cell proliferation | 31/1966 | 2.40E-06 | 3.45E-05 |

| | | | | |
|---|---|---|---|---|
| GO:0044782 | cilium organization | 52/1966 | 2.51E-06 | 3.59E-05 |
| GO:0045841 | negative regulation of mitotic metaphase/anaphase transition | 13/1966 | 2.61E-06 | 3.74E-05 |
| GO:0009306 | protein secretion | 53/1966 | 2.63E-06 | 3.75E-05 |
| GO:0097529 | myeloid leukocyte migration | 36/1966 | 2.64E-06 | 3.75E-05 |
| GO:0034501 | protein localization to kinetochore | Sep-66 | 2.65E-06 | 3.75E-05 |
| GO:1903083 | protein localization to condensed chromosome | Sep-66 | 2.65E-06 | 3.75E-05 |
| GO:0052548 | regulation of endopeptidase activity | 49/1966 | 2.66E-06 | 3.76E-05 |
| GO:1901652 | response to peptide | 55/1966 | 2.67E-06 | 3.76E-05 |
| GO:0007416 | synapse assembly | 32/1966 | 2.68E-06 | 3.76E-05 |
| GO:0034763 | negative regulation of transmembrane transport | 26/1966 | 2.68E-06 | 3.76E-05 |
| GO:1901992 | positive regulation of mitotic cell cycle phase transition | 21/1966 | 2.70E-06 | 3.78E-05 |
| GO:0051784 | negative regulation of nuclear division | 16/1966 | 2.80E-06 | 3.91E-05 |
| GO:0035592 | establishment of protein localization to extracellular region | 53/1966 | 2.84E-06 | 3.96E-05 |
| GO:0018105 | peptidyl-serine phosphorylation | 47/1966 | 2.96E-06 | 4.11E-05 |
| GO:0031214 | biomineral tissue development | 30/1966 | 3.04E-06 | 4.22E-05 |
| GO:0001649 | osteoblast differentiation | 35/1966 | 3.06E-06 | 4.23E-05 |
| GO:0060976 | coronary vasculature development | 18/1966 | 3.20E-06 | 4.42E-05 |
| GO:0030308 | negative regulation of cell growth | 33/1966 | 3.28E-06 | 4.51E-05 |

| | | | | |
|---|---|---|---|---|
| GO:0048705 | skeletal system morphogenesis | 39/1966 | 3.32E-06 | 4.56E-05 |
| GO:0042634 | regulation of hair cycle | Nov-66 | 3.43E-06 | 4.68E-05 |
| GO:1905523 | positive regulation of macrophage migration | Nov-66 | 3.43E-06 | 4.68E-05 |
| GO:0051653 | spindle localization | 15/1966 | 3.44E-06 | 4.69E-05 |
| GO:0072577 | endothelial cell apoptotic process | 15/1966 | 3.44E-06 | 4.69E-05 |
| GO:0002040 | sprouting angiogenesis | 24/1966 | 3.51E-06 | 4.73E-05 |
| GO:0051261 | protein depolymerization | 24/1966 | 3.51E-06 | 4.73E-05 |
| GO:0033046 | negative regulation of sister chromatid segregation | 13/1966 | 3.51E-06 | 4.73E-05 |
| GO:0033048 | negative regulation of mitotic sister chromatid segregation | 13/1966 | 3.51E-06 | 4.73E-05 |
| GO:2000816 | negative regulation of mitotic sister chromatid separation | 13/1966 | 3.51E-06 | 4.73E-05 |
| GO:0010466 | negative regulation of peptidase activity | 39/1966 | 3.64E-06 | 4.89E-05 |
| GO:0048846 | axon extension involved in axon guidance | Dec-66 | 3.66E-06 | 4.89E-05 |
| GO:1902284 | neuron projection extension involved in neuron projection guidance | Dec-66 | 3.66E-06 | 4.89E-05 |
| GO:0072009 | nephron epithelium development | 23/1966 | 3.67E-06 | 4.89E-05 |
| GO:0010951 | negative regulation of endopeptidase activity | 31/1966 | 3.82E-06 | 5.08E-05 |
| GO:0045927 | positive regulation of growth | 45/1966 | 3.85E-06 | 5.11E-05 |
| GO:0061180 | mammary gland epithelium development | 18/1966 | 3.90E-06 | 5.17E-05 |
| GO:0000727 | double-strand break repair via break-induced replication | Jul-66 | 3.97E-06 | 5.25E-05 |

| | | | | |
|---|---|---|---|---|
| GO:0034614 | cellular response to reactive oxygen species | 26/1966 | 4.03E-06 | 5.32E-05 |
| GO:1903053 | regulation of extracellular matrix organization | 14/1966 | 4.10E-06 | 5.40E-05 |
| GO:0070293 | renal absorption | Oct-66 | 4.12E-06 | 5.40E-05 |
| GO:0071459 | protein localization to chromosome, centromeric region | Oct-66 | 4.12E-06 | 5.40E-05 |
| GO:0043010 | camera-type eye development | 50/1966 | 4.16E-06 | 5.43E-05 |
| GO:0050866 | negative regulation of cell activation | 35/1966 | 4.17E-06 | 5.43E-05 |
| GO:0060415 | muscle tissue morphogenesis | 19/1966 | 4.23E-06 | 5.51E-05 |
| GO:0051897 | positive regulation of protein kinase B signaling | 23/1966 | 4.26E-06 | 5.53E-05 |
| GO:0014911 | positive regulation of smooth muscle cell migration | 16/1966 | 4.39E-06 | 5.69E-05 |
| GO:0030203 | glycosaminoglycan metabolic process | 20/1966 | 4.41E-06 | 5.70E-05 |
| GO:0048644 | muscle organ morphogenesis | 20/1966 | 4.41E-06 | 5.70E-05 |
| GO:0032872 | regulation of stress-activated MAPK cascade | 33/1966 | 4.53E-06 | 5.83E-05 |
| GO:0072593 | reactive oxygen species metabolic process | 37/1966 | 4.55E-06 | 5.85E-05 |
| GO:1902100 | negative regulation of metaphase/anaphase transition of cell cycle | 13/1966 | 4.68E-06 | 6.00E-05 |
| GO:0035265 | organ growth | 34/1966 | 4.83E-06 | 6.18E-05 |
| GO:0010038 | response to metal ion | 42/1966 | 4.98E-06 | 6.36E-05 |
| GO:0090257 | regulation of muscle system process | 38/1966 | 5.16E-06 | 6.55E-05 |
| GO:0032922 | circadian regulation of gene expression | 17/1966 | 5.17E-06 | 6.55E-05 |

| | | | | |
|---|---|---|---|---|
| GO:0046579 | positive regulation of Ras protein signal transduction | 17/1966 | 5.17E-06 | 6.55E-05 |
| GO:0048864 | stem cell development | 20/1966 | 5.24E-06 | 6.63E-05 |
| GO:1903522 | regulation of blood circulation | 39/1966 | 5.28E-06 | 6.66E-05 |
| GO:0051293 | establishment of spindle localization | 14/1966 | 5.30E-06 | 6.68E-05 |
| GO:0043524 | negative regulation of neuron apoptotic process | 31/1966 | 5.35E-06 | 6.72E-05 |
| GO:0043393 | regulation of protein binding | 34/1966 | 5.35E-06 | 6.72E-05 |
| GO:0031109 | microtubule polymerization or depolymerization | 24/1966 | 5.39E-06 | 6.75E-05 |
| GO:0045862 | positive regulation of proteolysis | 48/1966 | 5.49E-06 | 6.86E-05 |
| GO:0030071 | regulation of mitotic metaphase/anaphase transition | 15/1966 | 5.54E-06 | 6.90E-05 |
| GO:1904951 | positive regulation of establishment of protein localization | 46/1966 | 5.71E-06 | 7.10E-05 |
| GO:0000302 | response to reactive oxygen species | 31/1966 | 5.98E-06 | 7.41E-05 |
| GO:0098739 | import across plasma membrane | 31/1966 | 5.98E-06 | 7.41E-05 |
| GO:1905477 | positive regulation of protein localization to membrane | 22/1966 | 5.99E-06 | 7.41E-05 |
| GO:0050766 | positive regulation of phagocytosis | 19/1966 | 6.06E-06 | 7.46E-05 |
| GO:2001251 | negative regulation of chromosome organization | 19/1966 | 6.06E-06 | 7.46E-05 |
| GO:0030204 | chondroitin sulfate metabolic process | Oct-66 | 6.14E-06 | 7.55E-05 |
| GO:1905819 | negative regulation of chromosome separation | 13/1966 | 6.17E-06 | 7.56E-05 |
| GO:0044344 | cellular response to fibroblast growth factor stimulus | 20/1966 | 6.19E-06 | 7.56E-05 |

| | | | | |
|---|---|---|---|---|
| GO:0070302 | regulation of stress-activated protein kinase signaling cascade | 33/1966 | 6.21E-06 | 7.56E-05 |
| GO:2000241 | regulation of reproductive process | 33/1966 | 6.21E-06 | 7.56E-05 |
| GO:0031098 | stress-activated protein kinase signaling cascade | 38/1966 | 6.21E-06 | 7.56E-05 |
| GO:1905330 | regulation of morphogenesis of an epithelium | 17/1966 | 6.31E-06 | 7.66E-05 |
| GO:0060100 | positive regulation of phagocytosis, engulfment | Aug-66 | 6.44E-06 | 7.79E-05 |
| GO:1905155 | positive regulation of membrane invagination | Aug-66 | 6.44E-06 | 7.79E-05 |
| GO:0048639 | positive regulation of developmental growth | 34/1966 | 6.56E-06 | 7.91E-05 |
| GO:1901989 | positive regulation of cell cycle phase transition | 23/1966 | 6.62E-06 | 7.97E-05 |
| GO:0032506 | cytokinetic process | Dec-66 | 6.79E-06 | 8.14E-05 |
| GO:0061005 | cell differentiation involved in kidney development | 14/1966 | 6.80E-06 | 8.14E-05 |
| GO:0043270 | positive regulation of ion transport | 45/1966 | 6.80E-06 | 8.14E-05 |
| GO:0048844 | artery morphogenesis | 18/1966 | 6.89E-06 | 8.22E-05 |
| GO:0048841 | regulation of axon extension involved in axon guidance | Nov-66 | 6.89E-06 | 8.22E-05 |
| GO:0006941 | striated muscle contraction | 29/1966 | 6.95E-06 | 8.26E-05 |
| GO:0098657 | import into cell | 36/1966 | 7.11E-06 | 8.44E-05 |
| GO:2001237 | negative regulation of extrinsic apoptotic signaling pathway | 21/1966 | 7.21E-06 | 8.51E-05 |
| GO:0014032 | neural crest cell development | 19/1966 | 7.21E-06 | 8.51E-05 |
| GO:0033627 | cell adhesion mediated by integrin | 19/1966 | 7.21E-06 | 8.51E-05 |

| | | | | |
|---|---|---|---|---|
| GO:0051781 | positive regulation of cell division | 20/1966 | 7.31E-06 | 8.60E-05 |
| GO:0008544 | epidermis development | 48/1966 | 7.44E-06 | 8.75E-05 |
| GO:0032874 | positive regulation of stress-activated MAPK cascade | 25/1966 | 7.48E-06 | 8.77E-05 |
| GO:0048738 | cardiac muscle tissue development | 39/1966 | 7.56E-06 | 8.85E-05 |
| GO:0030856 | regulation of epithelial cell differentiation | 27/1966 | 7.84E-06 | 9.16E-05 |
| GO:0033047 | regulation of mitotic sister chromatid segregation | 13/1966 | 8.06E-06 | 9.40E-05 |
| GO:0043589 | skin morphogenesis | Jul-66 | 8.10E-06 | 9.43E-05 |
| GO:0050803 | regulation of synapse structure or activity | 39/1966 | 8.26E-06 | 9.59E-05 |
| GO:0016331 | morphogenesis of embryonic epithelium | 30/1966 | 8.54E-06 | 9.85E-05 |
| GO:0035051 | cardiocyte differentiation | 30/1966 | 8.54E-06 | 9.85E-05 |
| GO:0110148 | biomineralization | 30/1966 | 8.54E-06 | 9.85E-05 |
| GO:0071774 | response to fibroblast growth factor | 20/1966 | 8.60E-06 | 9.91E-05 |
| GO:1904036 | negative regulation of epithelial cell apoptotic process | 14/1966 | 8.66E-06 | 9.96E-05 |
| GO:0002548 | monocyte chemotaxis | 15/1966 | 8.69E-06 | 9.98E-05 |
| GO:0001936 | regulation of endothelial cell proliferation | 28/1966 | 8.84E-06 | 0.000101191 |
| GO:0007009 | plasma membrane organization | 27/1966 | 8.85E-06 | 0.000101191 |
| GO:0042886 | amide transport | 47/1966 | 8.89E-06 | 0.000101434 |
| GO:0002683 | negative regulation of immune system process | 56/1966 | 8.93E-06 | 0.000101721 |

| | | | | |
|---|---|---|---|---|
| GO:0051091 | positive regulation of DNA-binding transcription factor activity | 39/1966 | 9.02E-06 | 0.000102527 |
| GO:0022411 | cellular component disassembly | 54/1966 | 9.17E-06 | 0.000103989 |
| GO:0046777 | protein autophosphorylation | 36/1966 | 9.44E-06 | 0.000106847 |
| GO:0090504 | epiboly | Nov-66 | 9.56E-06 | 0.000108024 |
| GO:0051047 | positive regulation of secretion | 53/1966 | 9.58E-06 | 0.000108024 |
| GO:0070304 | positive regulation of stress-activated protein kinase signaling cascade | 25/1966 | 9.70E-06 | 0.000109167 |
| GO:0002700 | regulation of production of molecular mediator of immune response | 29/1966 | 9.77E-06 | 0.000109544 |
| GO:0050807 | regulation of synapse organization | 38/1966 | 9.77E-06 | 0.000109544 |
| GO:0035725 | sodium ion transmembrane transport | 26/1966 | 9.91E-06 | 0.000110912 |
| GO:0030099 | myeloid cell differentiation | 55/1966 | 1.00E-05 | 0.000111852 |
| GO:0014033 | neural crest cell differentiation | 20/1966 | 1.01E-05 | 0.000112523 |
| GO:0050708 | regulation of protein secretion | 42/1966 | 1.06E-05 | 0.000117901 |
| GO:0007612 | learning | 30/1966 | 1.06E-05 | 0.00011791 |
| GO:2000027 | regulation of animal organ morphogenesis | 24/1966 | 1.07E-05 | 0.000118157 |
| GO:0022898 | regulation of transmembrane transporter activity | 40/1966 | 1.07E-05 | 0.000118843 |
| GO:1902099 | regulation of metaphase/anaphase transition of cell cycle | 15/1966 | 1.08E-05 | 0.000119384 |
| GO:0001890 | placenta development | 29/1966 | 1.09E-05 | 0.000120393 |
| GO:0032330 | regulation of chondrocyte differentiation | 14/1966 | 1.10E-05 | 0.000120654 |

| | | | | |
|---|---|---|---|---|
| GO:0030900 | forebrain development | 51/1966 | 1.12E-05 | 0.000122552 |
| GO:0061326 | renal tubule development | 21/1966 | 1.14E-05 | 0.00012505 |
| GO:0097581 | lamellipodium organization | 19/1966 | 1.20E-05 | 0.000130749 |
| GO:1903035 | negative regulation of response to wounding | 19/1966 | 1.20E-05 | 0.000130749 |
| GO:0035886 | vascular associated smooth muscle cell differentiation | Dec-66 | 1.21E-05 | 0.000131689 |
| GO:2001234 | negative regulation of apoptotic signaling pathway | 36/1966 | 1.24E-05 | 0.00013542 |
| GO:0038127 | ERBB signaling pathway | 22/1966 | 1.25E-05 | 0.000135489 |
| GO:0010971 | positive regulation of G2/M transition of mitotic cell cycle | Oct-66 | 1.29E-05 | 0.000139426 |
| GO:0002688 | regulation of leukocyte chemotaxis | 23/1966 | 1.33E-05 | 0.00014334 |
| GO:0048259 | regulation of receptor-mediated endocytosis | 23/1966 | 1.33E-05 | 0.00014334 |
| GO:0060491 | regulation of cell projection assembly | 32/1966 | 1.33E-05 | 0.00014334 |
| GO:0031102 | neuron projection regeneration | 15/1966 | 1.34E-05 | 0.000143856 |
| GO:0030517 | negative regulation of axon extension | 13/1966 | 1.34E-05 | 0.000144469 |
| GO:0006029 | proteoglycan metabolic process | 17/1966 | 1.35E-05 | 0.000144719 |
| GO:0006260 | DNA replication | 39/1966 | 1.39E-05 | 0.000148509 |
| GO:1902806 | regulation of cell cycle G1/S phase transition | 28/1966 | 1.40E-05 | 0.000149084 |
| GO:0061351 | neural precursor cell proliferation | 31/1966 | 1.40E-05 | 0.000149084 |
| GO:0060047 | heart contraction | 35/1966 | 1.47E-05 | 0.00015615 |

| | | | | |
|---|---|---|---|---|
| GO:0032409 | regulation of transporter activity | 41/1966 | 1.48E-05 | 0.000157645 |
| GO:0051403 | stress-activated MAPK cascade | 36/1966 | 1.49E-05 | 0.000158184 |
| GO:0035904 | aorta development | 16/1966 | 1.51E-05 | 0.000159688 |
| GO:0050771 | negative regulation of axonogenesis | 16/1966 | 1.51E-05 | 0.000159688 |
| GO:0006750 | glutathione biosynthetic process | Jul-66 | 1.52E-05 | 0.000160366 |
| GO:0051764 | actin crosslink formation | Jul-66 | 1.52E-05 | 0.000160366 |
| GO:0051988 | regulation of attachment of spindle microtubules to kinetochore | Jul-66 | 1.52E-05 | 0.000160366 |
| GO:0032760 | positive regulation of tumor necrosis factor production | 21/1966 | 1.53E-05 | 0.000160714 |
| GO:0006874 | cellular calcium ion homeostasis | 60/1966 | 1.54E-05 | 0.000160928 |
| GO:0007596 | blood coagulation | 28/1966 | 1.56E-05 | 0.000163192 |
| GO:0019932 | second-messenger-mediated signaling | 42/1966 | 1.58E-05 | 0.000165126 |
| GO:0038084 | vascular endothelial growth factor signaling pathway | Dec-66 | 1.59E-05 | 0.000165346 |
| GO:2000379 | positive regulation of reactive oxygen species metabolic process | 17/1966 | 1.62E-05 | 0.000168203 |
| GO:0042176 | regulation of protein catabolic process | 50/1966 | 1.64E-05 | 0.00017047 |
| GO:0048260 | positive regulation of receptor-mediated endocytosis | 15/1966 | 1.64E-05 | 0.00017047 |
| GO:0050764 | regulation of phagocytosis | 22/1966 | 1.65E-05 | 0.000170604 |
| GO:0008589 | regulation of smoothened signaling pathway | 19/1966 | 1.65E-05 | 0.000170604 |
| GO:0051899 | membrane depolarization | 19/1966 | 1.65E-05 | 0.000170604 |

| | | | | |
|---|---|---|---|---|
| GO:0071897 | DNA biosynthetic process | 29/1966 | 1.69E-05 | 0.000173691 |
| GO:0048640 | negative regulation of developmental growth | 23/1966 | 1.73E-05 | 0.00017779 |
| GO:0061387 | regulation of extent of cell growth | 23/1966 | 1.73E-05 | 0.00017779 |
| GO:0008406 | gonad development | 35/1966 | 1.76E-05 | 0.000179968 |
| GO:0048863 | stem cell differentiation | 35/1966 | 1.76E-05 | 0.000179968 |
| GO:0032570 | response to progesterone | Aug-66 | 1.77E-05 | 0.000180729 |
| GO:0003015 | heart process | 36/1966 | 1.78E-05 | 0.000181536 |
| GO:0031623 | receptor internalization | 24/1966 | 1.78E-05 | 0.000181566 |
| GO:0009416 | response to light stimulus | 43/1966 | 1.81E-05 | 0.000183644 |
| GO:0034121 | regulation of toll-like receptor signaling pathway | 16/1966 | 1.83E-05 | 0.000185247 |
| GO:0045132 | meiotic chromosome segregation | 20/1966 | 1.87E-05 | 0.000189025 |
| GO:0051057 | positive regulation of small GTPase mediated signal transduction | 17/1966 | 1.93E-05 | 0.000194989 |
| GO:1901879 | regulation of protein depolymerization | 19/1966 | 1.94E-05 | 0.000195325 |
| GO:0007098 | centrosome cycle | 23/1966 | 1.97E-05 | 0.000198437 |
| GO:0014013 | regulation of gliogenesis | 23/1966 | 1.97E-05 | 0.000198437 |
| GO:0030879 | mammary gland development | 27/1966 | 1.99E-05 | 0.000200192 |
| GO:1903557 | positive regulation of tumor necrosis factor superfamily cytokine production | 21/1966 | 2.04E-05 | 0.000204144 |
| GO:0045880 | positive regulation of smoothened signaling pathway | Dec-66 | 2.06E-05 | 0.000206446 |

| | | | | |
|---|---|---|---|---|
| GO:0019748 | secondary metabolic process | 14/1966 | 2.14E-05 | 0.000213539 |
| GO:0030514 | negative regulation of BMP signaling pathway | 14/1966 | 2.14E-05 | 0.000213539 |
| GO:0090100 | positive regulation of transmembrane receptor protein serine/threonine kinase signaling pathway | 22/1966 | 2.16E-05 | 0.000214278 |
| GO:1902414 | protein localization to cell junction | 22/1966 | 2.16E-05 | 0.000214278 |
| GO:0007599 | hemostasis | 28/1966 | 2.16E-05 | 0.000214278 |
| GO:0050817 | coagulation | 28/1966 | 2.16E-05 | 0.000214278 |
| GO:0032675 | regulation of interleukin-6 production | 27/1966 | 2.23E-05 | 0.00022039 |
| GO:0007044 | cell-substrate junction assembly | 19/1966 | 2.26E-05 | 0.00022316 |
| GO:0051092 | positive regulation of NF-kappaB transcription factor activity | 26/1966 | 2.27E-05 | 0.000224168 |
| GO:0006261 | DNA-templated DNA replication | 25/1966 | 2.30E-05 | 0.00022589 |
| GO:0003151 | outflow tract morphogenesis | 18/1966 | 2.31E-05 | 0.000226868 |
| GO:0034114 | regulation of heterotypic cell-cell adhesion | Sep-66 | 2.36E-05 | 0.000231151 |
| GO:0022617 | extracellular matrix disassembly | Nov-66 | 2.36E-05 | 0.000231272 |
| GO:0009636 | response to toxic substance | 22/1966 | 2.47E-05 | 0.000241414 |
| GO:0032412 | regulation of ion transmembrane transporter activity | 38/1966 | 2.50E-05 | 0.000243944 |
| GO:0021952 | central nervous system projection neuron axonogenesis | Oct-66 | 2.51E-05 | 0.000244021 |
| GO:0071711 | basement membrane organization | Oct-66 | 2.51E-05 | 0.000244021 |
| GO:0045137 | development of primary sexual characteristics | 35/1966 | 2.51E-05 | 0.000244021 |

| | | | | |
|---|---|---|---|---|
| GO:0045598 | regulation of fat cell differentiation | 25/1966 | 2.58E-05 | 0.000250333 |
| GO:0021955 | central nervous system neuron axonogenesis | Dec-66 | 2.66E-05 | 0.000257615 |
| GO:0045732 | positive regulation of protein catabolic process | 33/1966 | 2.68E-05 | 0.000259054 |
| GO:0003215 | cardiac right ventricle morphogenesis | Aug-66 | 2.77E-05 | 0.000267498 |
| GO:0042542 | response to hydrogen peroxide | 22/1966 | 2.82E-05 | 0.000271255 |
| GO:0007127 | meiosis I | 23/1966 | 2.89E-05 | 0.000277884 |
| GO:0008584 | male gonad development | 24/1966 | 2.92E-05 | 0.000279009 |
| GO:0009855 | determination of bilateral symmetry | 24/1966 | 2.92E-05 | 0.000279009 |
| GO:1904375 | regulation of protein localization to cell periphery | 24/1966 | 2.92E-05 | 0.000279009 |
| GO:0048511 | rhythmic process | 41/1966 | 3.00E-05 | 0.000286951 |
| GO:0032231 | regulation of actin filament bundle assembly | 21/1966 | 3.08E-05 | 0.000293877 |
| GO:0007292 | female gamete generation | 27/1966 | 3.09E-05 | 0.000294313 |
| GO:0060674 | placenta blood vessel development | Nov-66 | 3.12E-05 | 0.000296672 |
| GO:0000725 | recombinational repair | 26/1966 | 3.19E-05 | 0.000302494 |
| GO:0001578 | microtubule bundle formation | 22/1966 | 3.21E-05 | 0.000303348 |
| GO:0002224 | toll-like receptor signaling pathway | 22/1966 | 3.21E-05 | 0.000303348 |
| GO:0051147 | regulation of muscle cell differentiation | 25/1966 | 3.25E-05 | 0.000307397 |
| GO:0009799 | specification of symmetry | 24/1966 | 3.29E-05 | 0.000309326 |

| | | | | |
|---|---|---|---|---|
| GO:0046546 | development of primary male sexual characteristics | 24/1966 | 3.29E-05 | 0.000309326 |
| GO:0090596 | sensory organ morphogenesis | 42/1966 | 3.39E-05 | 0.000319024 |
| GO:0030261 | chromosome condensation | Dec-66 | 3.41E-05 | 0.000319666 |
| GO:1902751 | positive regulation of cell cycle G2/M phase transition | Oct-66 | 3.43E-05 | 0.000320894 |
| GO:0044771 | meiotic cell cycle phase transition | Jun-66 | 3.43E-05 | 0.000320894 |
| GO:0048872 | homeostasis of number of cells | 47/1966 | 3.46E-05 | 0.000323525 |
| GO:0030595 | leukocyte chemotaxis | 33/1966 | 3.52E-05 | 0.000326823 |
| GO:0055001 | muscle cell development | 33/1966 | 3.52E-05 | 0.000326823 |
| GO:0030038 | contractile actin filament bundle assembly | 21/1966 | 3.52E-05 | 0.000326823 |
| GO:0043149 | stress fiber assembly | 21/1966 | 3.52E-05 | 0.000326823 |
| GO:0071214 | cellular response to abiotic stimulus | 43/1966 | 3.53E-05 | 0.000326823 |
| GO:0104004 | cellular response to environmental stimulus | 43/1966 | 3.53E-05 | 0.000326823 |
| GO:0007159 | leukocyte cell-cell adhesion | 48/1966 | 3.54E-05 | 0.000326823 |
| GO:0001938 | positive regulation of endothelial cell proliferation | 19/1966 | 3.55E-05 | 0.000327245 |
| GO:0060350 | endochondral bone morphogenesis | 15/1966 | 3.59E-05 | 0.000330926 |
| GO:0015833 | peptide transport | 42/1966 | 3.65E-05 | 0.000335979 |
| GO:0031023 | microtubule organizing center organization | 24/1966 | 3.70E-05 | 0.000339355 |
| GO:0014068 | positive regulation of phosphatidylinositol 3-kinase signaling | 16/1966 | 3.76E-05 | 0.000345031 |

| | | | | |
|---|---|---|---|---|
| GO:0070373 | negative regulation of ERK1 and ERK2 cascade | 17/1966 | 3.79E-05 | 0.000347122 |
| GO:0010001 | glial cell differentiation | 36/1966 | 3.84E-05 | 0.000351256 |
| GO:0009991 | response to extracellular stimulus | 50/1966 | 3.90E-05 | 0.000355434 |
| GO:0014009 | glial cell proliferation | 14/1966 | 3.98E-05 | 0.000362082 |
| GO:0060443 | mammary gland morphogenesis | 14/1966 | 3.98E-05 | 0.000362082 |
| GO:0015711 | organic anion transport | 45/1966 | 4.05E-05 | 0.000367483 |
| GO:0050728 | negative regulation of inflammatory response | 24/1966 | 4.15E-05 | 0.00037657 |
| GO:0060099 | regulation of phagocytosis, engulfment | Aug-66 | 4.21E-05 | 0.000381413 |
| GO:0032233 | positive regulation of actin filament bundle assembly | 15/1966 | 4.32E-05 | 0.000390143 |
| GO:0035050 | embryonic heart tube development | 18/1966 | 4.32E-05 | 0.000390143 |
| GO:0019184 | nonribosomal peptide biosynthetic process | Jul-66 | 4.50E-05 | 0.000405568 |
| GO:0002063 | chondrocyte development | Oct-66 | 4.62E-05 | 0.000413108 |
| GO:0006270 | DNA replication initiation | Oct-66 | 4.62E-05 | 0.000413108 |
| GO:0044319 | wound healing, spreading of cells | Oct-66 | 4.62E-05 | 0.000413108 |
| GO:0060740 | prostate gland epithelium morphogenesis | Oct-66 | 4.62E-05 | 0.000413108 |
| GO:0090505 | epiboly involved in wound healing | Oct-66 | 4.62E-05 | 0.000413108 |
| GO:1901653 | cellular response to peptide | 41/1966 | 4.70E-05 | 0.000420131 |
| GO:0031954 | positive regulation of protein autophosphorylation | Sep-66 | 4.77E-05 | 0.000424526 |

| | | | | |
|---|---|---|---|---|
| GO:0048843 | negative regulation of axon extension involved in axon guidance | Sep-66 | 4.77E-05 | 0.000424526 |
| GO:0072012 | glomerulus vasculature development | Sep-66 | 4.77E-05 | 0.000424526 |
| GO:0010812 | negative regulation of cell-substrate adhesion | 14/1966 | 4.84E-05 | 0.000430087 |
| GO:0060349 | bone morphogenesis | 20/1966 | 5.03E-05 | 0.000445726 |
| GO:0019221 | cytokine-mediated signaling pathway | 51/1966 | 5.05E-05 | 0.000447333 |
| GO:0071677 | positive regulation of mononuclear cell migration | 15/1966 | 5.18E-05 | 0.000457574 |
| GO:0006023 | aminoglycan biosynthetic process | 13/1966 | 5.26E-05 | 0.000462459 |
| GO:0031103 | axon regeneration | 13/1966 | 5.26E-05 | 0.000462459 |
| GO:0048483 | autonomic nervous system development | 13/1966 | 5.26E-05 | 0.000462459 |
| GO:0060351 | cartilage development involved in endochondral bone morphogenesis | Nov-66 | 5.28E-05 | 0.000463112 |
| GO:1905332 | positive regulation of morphogenesis of an epithelium | Nov-66 | 5.28E-05 | 0.000463112 |
| GO:0002090 | regulation of receptor internalization | 16/1966 | 5.29E-05 | 0.000463112 |
| GO:0051348 | negative regulation of transferase activity | 36/1966 | 5.32E-05 | 0.000465638 |
| GO:0042116 | macrophage activation | 19/1966 | 5.45E-05 | 0.000474169 |
| GO:0072080 | nephron tubule development | 19/1966 | 5.45E-05 | 0.000474169 |
| GO:0150115 | cell-substrate junction organization | 19/1966 | 5.45E-05 | 0.000474169 |
| GO:0007219 | Notch signaling pathway | 28/1966 | 5.46E-05 | 0.000474169 |
| GO:0050714 | positive regulation of protein secretion | 26/1966 | 5.46E-05 | 0.000474169 |

| | | | | |
|---|---|---|---|---|
| GO:0001838 | embryonic epithelial tube formation | 25/1966 | 5.67E-05 | 0.000491902 |
| GO:0032635 | interleukin-6 production | 27/1966 | 5.77E-05 | 0.000499672 |
| GO:0032386 | regulation of intracellular transport | 43/1966 | 5.80E-05 | 0.000501747 |
| GO:0060759 | regulation of response to cytokine stimulus | 24/1966 | 5.84E-05 | 0.000504133 |
| GO:0010171 | body morphogenesis | 14/1966 | 5.87E-05 | 0.000505031 |
| GO:0051058 | negative regulation of small GTPase mediated signal transduction | 14/1966 | 5.87E-05 | 0.000505031 |
| GO:0008361 | regulation of cell size | 31/1966 | 5.88E-05 | 0.000505581 |
| GO:0008306 | associative learning | 21/1966 | 5.92E-05 | 0.00050767 |
| GO:0048041 | focal adhesion assembly | 17/1966 | 6.10E-05 | 0.000522407 |
| GO:0002092 | positive regulation of receptor internalization | Oct-66 | 6.14E-05 | 0.000523491 |
| GO:0050654 | chondroitin sulfate proteoglycan metabolic process | Oct-66 | 6.14E-05 | 0.000523491 |
| GO:0060512 | prostate gland morphogenesis | Oct-66 | 6.14E-05 | 0.000523491 |
| GO:0099560 | synaptic membrane adhesion | Oct-66 | 6.14E-05 | 0.000523491 |
| GO:0031579 | membrane raft organization | Aug-66 | 6.22E-05 | 0.000526505 |
| GO:0060080 | inhibitory postsynaptic potential | Aug-66 | 6.22E-05 | 0.000526505 |
| GO:0150104 | transport across blood-brain barrier | Aug-66 | 6.22E-05 | 0.000526505 |
| GO:1905153 | regulation of membrane invagination | Aug-66 | 6.22E-05 | 0.000526505 |
| GO:2001224 | positive regulation of neuron migration | Aug-66 | 6.22E-05 | 0.000526505 |

| | | | | |
|---|---|---|---|---|
| GO:0014066 | regulation of phosphatidylinositol 3-kinase signaling | 19/1966 | 6.26E-05 | 0.000528841 |
| GO:0120032 | regulation of plasma membrane bounded cell projection assembly | 30/1966 | 6.35E-05 | 0.000535477 |
| GO:0030838 | positive regulation of actin filament polymerization | 13/1966 | 6.46E-05 | 0.000540556 |
| GO:0045687 | positive regulation of glial cell differentiation | 13/1966 | 6.46E-05 | 0.000540556 |
| GO:0046580 | negative regulation of Ras protein signal transduction | 13/1966 | 6.46E-05 | 0.000540556 |
| GO:2001222 | regulation of neuron migration | 13/1966 | 6.46E-05 | 0.000540556 |
| GO:0060433 | bronchus development | Jun-66 | 6.46E-05 | 0.000540556 |
| GO:0060907 | positive regulation of macrophage cytokine production | Jun-66 | 6.46E-05 | 0.000540556 |
| GO:0001843 | neural tube closure | 20/1966 | 6.55E-05 | 0.000547108 |
| GO:0033622 | integrin activation | Sep-66 | 6.61E-05 | 0.000549857 |
| GO:0061437 | renal system vasculature development | Sep-66 | 6.61E-05 | 0.000549857 |
| GO:0061440 | kidney vasculature development | Sep-66 | 6.61E-05 | 0.000549857 |
| GO:0072175 | epithelial tube formation | 26/1966 | 6.72E-05 | 0.000558108 |
| GO:0060996 | dendritic spine development | 22/1966 | 6.74E-05 | 0.000558787 |
| GO:0060603 | mammary gland duct morphogenesis | Nov-66 | 6.78E-05 | 0.000561287 |
| GO:0030195 | negative regulation of blood coagulation | Dec-66 | 6.82E-05 | 0.00056215 |
| GO:0048010 | vascular endothelial growth factor receptor signaling pathway | Dec-66 | 6.82E-05 | 0.00056215 |
| GO:0048246 | macrophage chemotaxis | Dec-66 | 6.82E-05 | 0.00056215 |

| | | | | |
|---|---|---|---|---|
| GO:0000082 | G1/S transition of mitotic cell cycle | 30/1966 | 6.95E-05 | 0.000572649 |
| GO:0003413 | chondrocyte differentiation involved in endochondral bone morphogenesis | Jul-66 | 7.20E-05 | 0.000590404 |
| GO:0051382 | kinetochore assembly | Jul-66 | 7.20E-05 | 0.000590404 |
| GO:0090136 | epithelial cell-cell adhesion | Jul-66 | 7.20E-05 | 0.000590404 |
| GO:0006302 | double-strand break repair | 36/1966 | 7.30E-05 | 0.000598153 |
| GO:2000134 | negative regulation of G1/S transition of mitotic cell cycle | 15/1966 | 7.34E-05 | 0.000600553 |
| GO:0060606 | tube closure | 20/1966 | 7.45E-05 | 0.000608932 |
| GO:0021782 | glial cell development | 21/1966 | 7.58E-05 | 0.000617713 |
| GO:0032612 | interleukin-1 production | 21/1966 | 7.58E-05 | 0.000617713 |
| GO:0009913 | epidermal cell differentiation | 33/1966 | 7.64E-05 | 0.000621665 |
| GO:0033157 | regulation of intracellular protein transport | 32/1966 | 7.68E-05 | 0.000623053 |
| GO:0048545 | response to steroid hormone | 32/1966 | 7.68E-05 | 0.000623053 |
| GO:0031110 | regulation of microtubule polymerization or depolymerization | 18/1966 | 7.76E-05 | 0.00062836 |
| GO:1902904 | negative regulation of supramolecular fiber organization | 27/1966 | 7.78E-05 | 0.000629088 |
| GO:1903131 | mononuclear cell differentiation | 57/1966 | 7.85E-05 | 0.000633988 |
| GO:0046942 | carboxylic acid transport | 38/1966 | 8.06E-05 | 0.00064905 |
| GO:0090162 | establishment of epithelial cell polarity | Oct-66 | 8.08E-05 | 0.00064905 |
| GO:1901099 | negative regulation of signal transduction in absence of ligand | Oct-66 | 8.08E-05 | 0.00064905 |

| | | | | |
|---|---|---|---|---|
| GO:2001240 | negative regulation of extrinsic apoptotic signaling pathway in absence of ligand | Oct-66 | 8.08E-05 | 0.00064905 |
| GO:0071248 | cellular response to metal ion | 26/1966 | 8.23E-05 | 0.000659938 |
| GO:0001738 | morphogenesis of a polarized epithelium | 17/1966 | 8.25E-05 | 0.000659938 |
| GO:0021675 | nerve development | 17/1966 | 8.25E-05 | 0.000659938 |
| GO:0015849 | organic acid transport | 41/1966 | 8.33E-05 | 0.000665572 |
| GO:0070588 | calcium ion transmembrane transport | 39/1966 | 8.38E-05 | 0.000668911 |
| GO:1900047 | negative regulation of hemostasis | Dec-66 | 8.47E-05 | 0.000674158 |
| GO:2001239 | regulation of extrinsic apoptotic signaling pathway in absence of ligand | Dec-66 | 8.47E-05 | 0.000674158 |
| GO:0060393 | regulation of pathway-restricted SMAD protein phosphorylation | 14/1966 | 8.49E-05 | 0.000674598 |
| GO:0007368 | determination of left/right symmetry | 22/1966 | 8.52E-05 | 0.000674598 |
| GO:0043244 | regulation of protein-containing complex disassembly | 22/1966 | 8.52E-05 | 0.000674598 |
| GO:0060048 | cardiac muscle contraction | 22/1966 | 8.52E-05 | 0.000674598 |
| GO:0007369 | gastrulation | 27/1966 | 8.57E-05 | 0.000677699 |
| GO:0007143 | female meiotic nuclear division | Nov-66 | 8.62E-05 | 0.000680532 |
| GO:0055008 | cardiac muscle tissue morphogenesis | 15/1966 | 8.69E-05 | 0.000685643 |
| GO:1902107 | positive regulation of leukocyte differentiation | 28/1966 | 8.82E-05 | 0.00069393 |
| GO:1903708 | positive regulation of hemopoiesis | 28/1966 | 8.82E-05 | 0.00069393 |
| GO:0010232 | vascular transport | Aug-66 | 8.97E-05 | 0.000703714 |

| | | | | |
|---|---|---|---|---|
| GO:007<br>1496 | cellular response to external stimulus | 39/196<br>6 | 9.01E-<br>05 | 0.00070<br>3714 |
| GO:000<br>3148 | outflow tract septum morphogenesis | Sep-<br>66 | 9.01E-<br>05 | 0.00070<br>3714 |
| GO:006<br>0444 | branching involved in mammary gland duct morphogenesis | Sep-<br>66 | 9.01E-<br>05 | 0.00070<br>3714 |
| GO:009<br>7205 | renal filtration | Sep-<br>66 | 9.01E-<br>05 | 0.00070<br>3714 |
| GO:006<br>1333 | renal tubule morphogenesis | 17/196<br>6 | 9.55E-<br>05 | 0.00074<br>5359 |
| GO:001<br>0463 | mesenchymal cell proliferation | 13/196<br>6 | 9.58E-<br>05 | 0.00074<br>5773 |
| GO:005<br>1489 | regulation of filopodium assembly | 13/196<br>6 | 9.58E-<br>05 | 0.00074<br>5773 |
| GO:012<br>0034 | positive regulation of plasma membrane bounded cell projection assembly | 20/196<br>6 | 9.60E-<br>05 | 0.00074<br>6295 |
| GO:007<br>0482 | response to oxygen levels | 39/196<br>6 | 9.67E-<br>05 | 0.00075<br>0892 |
| GO:000<br>8543 | fibroblast growth factor receptor signaling pathway | 16/196<br>6 | 0.00010<br>0274 | 0.00077<br>6243 |
| GO:190<br>2807 | negative regulation of cell cycle G1/S phase transition | 16/196<br>6 | 0.00010<br>0274 | 0.00077<br>6243 |
| GO:000<br>1954 | positive regulation of cell-matrix adhesion | 14/196<br>6 | 0.00010<br>1491 | 0.00078<br>4633 |
| GO:190<br>4063 | negative regulation of cation transmembrane transport | 18/196<br>6 | 0.00010<br>2427 | 0.00079<br>084 |
| GO:004<br>3583 | ear development | 34/196<br>6 | 0.00010<br>4133 | 0.00080<br>2962 |
| GO:000<br>3254 | regulation of membrane depolarization | Dec-<br>66 | 0.00010<br>4576 | 0.00080<br>4283 |
| GO:005<br>0819 | negative regulation of coagulation | Dec-<br>66 | 0.00010<br>4576 | 0.00080<br>4283 |
| GO:000<br>2053 | positive regulation of mesenchymal cell proliferation | Oct-66 | 0.00010<br>5038 | 0.00080<br>6793 |
| GO:003<br>4766 | negative regulation of ion transmembrane transport | 19/196<br>6 | 0.00010<br>6715 | 0.00081<br>8608 |

| | | | | |
|---|---|---|---|---|
| GO:0032652 | regulation of interleukin-1 production | 20/1966 | 0.000108692 | 0.000830757 |
| GO:0043534 | blood vessel endothelial cell migration | 20/1966 | 0.000108692 | 0.000830757 |
| GO:0050892 | intestinal absorption | Nov-66 | 0.000108719 | 0.000830757 |
| GO:0070661 | leukocyte proliferation | 45/1966 | 0.000109177 | 0.000833179 |
| GO:0046209 | nitric oxide metabolic process | 17/1966 | 0.000110378 | 0.000841261 |
| GO:0046661 | male sex differentiation | 26/1966 | 0.000110771 | 0.000843166 |
| GO:0032060 | bleb assembly | Jun-66 | 0.000113075 | 0.000854117 |
| GO:0032905 | transforming growth factor beta1 production | Jun-66 | 0.000113075 | 0.000854117 |
| GO:0051315 | attachment of mitotic spindle microtubules to kinetochore | Jun-66 | 0.000113075 | 0.000854117 |
| GO:0071635 | negative regulation of transforming growth factor beta production | Jun-66 | 0.000113075 | 0.000854117 |
| GO:0097084 | vascular associated smooth muscle cell development | Jun-66 | 0.000113075 | 0.000854117 |
| GO:1905048 | regulation of metallopeptidase activity | Jun-66 | 0.000113075 | 0.000854117 |
| GO:0016311 | dephosphorylation | 43/1966 | 0.000113717 | 0.000857878 |
| GO:0035148 | tube formation | 27/1966 | 0.000114135 | 0.000859931 |
| GO:0002573 | myeloid leukocyte differentiation | 33/1966 | 0.000114832 | 0.000864089 |
| GO:0090303 | positive regulation of wound healing | 13/1966 | 0.000115873 | 0.000870809 |
| GO:0038034 | signal transduction in absence of ligand | 16/1966 | 0.000116782 | 0.000875418 |
| GO:0097192 | extrinsic apoptotic signaling pathway in absence of ligand | 16/1966 | 0.000116782 | 0.000875418 |

| | | | | |
|---|---|---|---|---|
| GO:0048286 | lung alveolus development | 14/1966 | 0.000120806 | 0.000901853 |
| GO:0060688 | regulation of morphogenesis of a branching structure | 14/1966 | 0.000120806 | 0.000901853 |
| GO:0002029 | desensitization of G protein-coupled receptor signaling pathway | Sep-66 | 0.000120917 | 0.000901853 |
| GO:0022401 | negative adaptation of signaling pathway | Sep-66 | 0.000120917 | 0.000901853 |
| GO:0035296 | regulation of tube diameter | 26/1966 | 0.000122037 | 0.000907922 |
| GO:0097746 | blood vessel diameter maintenance | 26/1966 | 0.000122037 | 0.000907922 |
| GO:0006939 | smooth muscle contraction | 20/1966 | 0.000122281 | 0.000912525 |
| GO:0043242 | negative regulation of protein-containing complex disassembly | 17/1966 | 0.000127212 | 0.000942863 |
| GO:0060420 | regulation of heart growth | 17/1966 | 0.000127212 | 0.000942863 |
| GO:0051099 | positive regulation of binding | 28/1966 | 0.000127446 | 0.000943418 |
| GO:0006937 | regulation of muscle contraction | 25/1966 | 0.000129766 | 0.000959391 |
| GO:0014065 | phosphatidylinositol 3-kinase signaling | 22/1966 | 0.000133391 | 0.000988798 |
| GO:0035150 | regulation of tube size | 26/1966 | 0.000134316 | 0.000989324 |
| GO:1903169 | regulation of calcium ion transmembrane transport | 26/1966 | 0.000134316 | 0.000989324 |
| GO:0035909 | aorta morphogenesis | Oct-66 | 0.000135191 | 0.00099933 |
| GO:0045746 | negative regulation of Notch signaling pathway | Oct-66 | 0.000135191 | 0.00099933 |
| GO:0010464 | regulation of mesenchymal cell proliferation | Nov-66 | 0.000136085 | 0.000998631 |
| GO:0042310 | vasoconstriction | 19/1966 | 0.000137664 | 0.001008966 |

| | | | | |
|---|---|---|---|---|
| GO:0007528 | neuromuscular junction development | 13/1966 | 0.000139422 | 0.001020589 |
| GO:2001057 | reactive nitrogen species metabolic process | 17/1966 | 0.00014625 | 0.001069252 |
| GO:0032611 | interleukin-1 beta production | 18/1966 | 0.000152854 | 0.001114779 |
| GO:0062207 | regulation of pattern recognition receptor signaling pathway | 18/1966 | 0.000152854 | 0.001114779 |
| GO:0030282 | bone mineralization | 21/1966 | 0.000153635 | 0.001119099 |
| GO:0014020 | primary neural tube formation | 20/1966 | 0.00015598 | 0.001134785 |
| GO:0060191 | regulation of lipase activity | 16/1966 | 0.000157039 | 0.001141084 |
| GO:0002699 | positive regulation of immune effector process | 39/1966 | 0.000157494 | 0.001142987 |
| GO:1903364 | positive regulation of cellular protein catabolic process | 23/1966 | 0.000159589 | 0.001156359 |
| GO:0035767 | endothelial cell chemotaxis | Sep-66 | 0.000160117 | 0.001156359 |
| GO:0061036 | positive regulation of cartilage development | Sep-66 | 0.000160117 | 0.001156359 |
| GO:2000352 | negative regulation of endothelial cell apoptotic process | Sep-66 | 0.000160117 | 0.001156359 |
| GO:0051494 | negative regulation of cytoskeleton organization | 26/1966 | 0.000162221 | 0.001170124 |
| GO:0002263 | cell activation involved in immune response | 41/1966 | 0.000163389 | 0.00117712 |
| GO:0051054 | positive regulation of DNA metabolic process | 38/1966 | 0.000164631 | 0.0011844 |
| GO:0021783 | preganglionic parasympathetic fiber development | Jul-66 | 0.000165187 | 0.0011844 |
| GO:0033623 | regulation of integrin activation | Jul-66 | 0.000165187 | 0.0011844 |
| GO:0080164 | regulation of nitric oxide metabolic process | 15/1966 | 0.00016552 | 0.0011844 |

| | | | | |
|---|---|---|---|---|
| GO:0044272 | sulfur compound biosynthetic process | 17/1966 | 0.000167733 | 0.001199657 |
| GO:0070301 | cellular response to hydrogen peroxide | 17/1966 | 0.000167733 | 0.001199657 |
| GO:0051480 | regulation of cytosolic calcium ion concentration | 47/1966 | 0.000168282 | 0.001202129 |
| GO:0050918 | positive chemotaxis | Nov-66 | 0.000169093 | 0.001204003 |
| GO:0006801 | superoxide metabolic process | 14/1966 | 0.000169154 | 0.001204003 |
| GO:0060389 | pathway-restricted SMAD protein phosphorylation | 14/1966 | 0.000169154 | 0.001204003 |
| GO:0042476 | odontogenesis | 21/1966 | 0.000171887 | 0.001220717 |
| GO:0051294 | establishment of spindle orientation | Oct-66 | 0.000172327 | 0.001220717 |
| GO:0072210 | metanephric nephron development | Oct-66 | 0.000172327 | 0.001220717 |
| GO:2000758 | positive regulation of peptidyl-lysine acetylation | Oct-66 | 0.000172327 | 0.001220717 |
| GO:0009100 | glycoprotein metabolic process | 41/1966 | 0.000174369 | 0.001233709 |
| GO:0010759 | positive regulation of macrophage chemotaxis | Aug-66 | 0.000175025 | 0.00123354 |
| GO:0035988 | chondrocyte proliferation | Aug-66 | 0.000175025 | 0.00123354 |
| GO:0002221 | pattern recognition receptor signaling pathway | 26/1966 | 0.000178018 | 0.001253629 |
| GO:0021537 | telencephalon development | 34/1966 | 0.000178031 | 0.001253629 |
| GO:0021953 | central nervous system neuron differentiation | 29/1966 | 0.000181215 | 0.001272389 |
| GO:0034103 | regulation of tissue remodeling | 16/1966 | 0.000181352 | 0.001272389 |
| GO:0072088 | nephron epithelium morphogenesis | 16/1966 | 0.000181352 | 0.001272389 |

| | | | | |
|---|---|---|---|---|
| GO:0008016 | regulation of heart contraction | 28/1966 | 0.000181555 | 0.001272389 |
| GO:0030072 | peptide hormone secretion | 37/1966 | 0.000184135 | 0.001288944 |
| GO:0000724 | double-strand break repair via homologous recombination | 24/1966 | 0.000185137 | 0.001294432 |
| GO:1902969 | mitotic DNA replication | Jun-66 | 0.000186446 | 0.001300511 |
| GO:2000052 | positive regulation of non-canonical Wnt signaling pathway | Jun-66 | 0.000186446 | 0.001300511 |
| GO:0030850 | prostate gland development | Dec-66 | 0.000189931 | 0.001323262 |
| GO:0021954 | central nervous system neuron development | 17/1966 | 0.000191919 | 0.001334694 |
| GO:0003279 | cardiac septum development | 21/1966 | 0.000192023 | 0.001334694 |
| GO:0051963 | regulation of synapse assembly | 20/1966 | 0.000196795 | 0.001366257 |
| GO:0022408 | negative regulation of cell-cell adhesion | 29/1966 | 0.000197093 | 0.001366728 |
| GO:0099565 | chemical synaptic transmission, postsynaptic | 18/1966 | 0.000197506 | 0.00136799 |
| GO:0003143 | embryonic heart tube morphogenesis | 14/1966 | 0.000199034 | 0.001374064 |
| GO:1904377 | positive regulation of protein localization to cell periphery | 14/1966 | 0.000199034 | 0.001374064 |
| GO:0032731 | positive regulation of interleukin-1 beta production | 13/1966 | 0.000199079 | 0.001374064 |
| GO:0007389 | pattern specification process | 55/1966 | 0.00020088 | 0.00138488 |
| GO:0031349 | positive regulation of defense response | 38/1966 | 0.000202072 | 0.001391479 |
| GO:0014902 | myotube differentiation | 22/1966 | 0.000205812 | 0.001415584 |
| GO:0031952 | regulation of protein autophosphorylation | Nov-66 | 0.000208647 | 0.001429978 |

| | | | | |
|---|---|---|---|---|
| GO:0060323 | head morphogenesis | Nov-66 | 0.000208647 | 0.001429978 |
| GO:1903170 | negative regulation of calcium ion transmembrane transport | Nov-66 | 0.000208647 | 0.001429978 |
| GO:0019229 | regulation of vasoconstriction | 16/1966 | 0.000208871 | 0.001429978 |
| GO:0023058 | adaptation of signaling pathway | Sep-66 | 0.00020936 | 0.001431673 |
| GO:0060402 | calcium ion transport into cytosol | 26/1966 | 0.000213768 | 0.00146013 |
| GO:0051592 | response to calcium ion | 21/1966 | 0.000214206 | 0.001461439 |
| GO:1904645 | response to amyloid-beta | Oct-66 | 0.00021768 | 0.001483431 |
| GO:0032651 | regulation of interleukin-1 beta production | 17/1966 | 0.000219089 | 0.001491315 |
| GO:0060419 | heart growth | 20/1966 | 0.000220513 | 0.001499284 |
| GO:0007173 | epidermal growth factor receptor signaling pathway | 18/1966 | 0.000223827 | 0.00152007 |
| GO:2000377 | regulation of reactive oxygen species metabolic process | 24/1966 | 0.000224929 | 0.001525807 |
| GO:0045861 | negative regulation of proteolysis | 43/1966 | 0.000227558 | 0.001541878 |
| GO:0002703 | regulation of leukocyte mediated immunity | 38/1966 | 0.00023111 | 0.001564156 |
| GO:0060711 | labyrinthine layer development | 13/1966 | 0.00023633 | 0.001597655 |
| GO:0002031 | G protein-coupled receptor internalization | Jul-66 | 0.000239201 | 0.001611031 |
| GO:0010763 | positive regulation of fibroblast migration | Jul-66 | 0.000239201 | 0.001611031 |
| GO:0072010 | glomerular epithelium development | Jul-66 | 0.000239201 | 0.001611031 |
| GO:0035418 | protein localization to synapse | 16/1966 | 0.000239941 | 0.001611031 |

| | | | | |
|---|---|---|---|---|
| GO:0072028 | nephron morphogenesis | 16/1966 | 0.000239941 | 0.001611031 |
| GO:1902893 | regulation of miRNA transcription | 16/1966 | 0.000239941 | 0.001611031 |
| GO:0050863 | regulation of T cell activation | 42/1966 | 0.000241209 | 0.001617713 |
| GO:0046330 | positive regulation of JNK cascade | 18/1966 | 0.000253155 | 0.001695911 |
| GO:0045619 | regulation of lymphocyte differentiation | 28/1966 | 0.000255245 | 0.00170753 |
| GO:0006024 | glycosaminoglycan biosynthetic process | Nov-66 | 0.000255755 | 0.00170753 |
| GO:1903524 | positive regulation of blood circulation | Nov-66 | 0.000255755 | 0.00170753 |
| GO:0045637 | regulation of myeloid cell differentiation | 30/1966 | 0.00026863 | 0.001791474 |
| GO:0006790 | sulfur compound metabolic process | 40/1966 | 0.00027004 | 0.001798849 |
| GO:0033260 | nuclear DNA replication | Sep-66 | 0.000270552 | 0.001800236 |
| GO:0042398 | cellular modified amino acid biosynthetic process | Oct-66 | 0.000272631 | 0.001810004 |
| GO:0060325 | face morphogenesis | Oct-66 | 0.000272631 | 0.001810004 |
| GO:0034113 | heterotypic cell-cell adhesion | Dec-66 | 0.000275123 | 0.001823395 |
| GO:0015748 | organophosphate ester transport | 20/1966 | 0.000275572 | 0.001823395 |
| GO:1901890 | positive regulation of cell junction assembly | 20/1966 | 0.000275572 | 0.001823395 |
| GO:0001765 | membrane raft assembly | May-66 | 0.000277256 | 0.001830443 |
| GO:0099640 | axo-dendritic protein transport | May-66 | 0.000277256 | 0.001830443 |
| GO:0001755 | neural crest cell migration | 13/1966 | 0.000279383 | 0.001840386 |

| | | | | |
|---|---|---|---|---|
| GO:0045744 | negative regulation of G protein-coupled receptor signaling pathway | 13/1966 | 0.000279383 | 0.001840386 |
| GO:0046883 | regulation of hormone secretion | 38/1966 | 0.000281703 | 0.001853606 |
| GO:0070555 | response to interleukin-1 | 17/1966 | 0.000283609 | 0.001864077 |
| GO:0035264 | multicellular organism growth | 30/1966 | 0.000290511 | 0.001907322 |
| GO:0007076 | mitotic chromosome condensation | Jun-66 | 0.000292812 | 0.001918179 |
| GO:1904177 | regulation of adipose tissue development | Jun-66 | 0.000292812 | 0.001918179 |
| GO:0001841 | neural tube formation | 21/1966 | 0.000294842 | 0.001929343 |
| GO:1904888 | cranial skeletal system development | 15/1966 | 0.00029836 | 0.001950208 |
| GO:0050680 | negative regulation of epithelial cell proliferation | 25/1966 | 0.00030321 | 0.001979725 |
| GO:0015850 | organic hydroxy compound transport | 33/1966 | 0.000307641 | 0.002006445 |
| GO:0003179 | heart valve morphogenesis | Nov-66 | 0.000311527 | 0.002029552 |
| GO:0061614 | miRNA transcription | 16/1966 | 0.000314257 | 0.00204509 |
| GO:0002790 | peptide secretion | 37/1966 | 0.000316464 | 0.00205719 |
| GO:0045428 | regulation of nitric oxide biosynthetic process | 14/1966 | 0.000317451 | 0.002061121 |
| GO:0062009 | secondary palate development | Aug-66 | 0.000317765 | 0.002061121 |
| GO:0001657 | ureteric bud development | 18/1966 | 0.00032199 | 0.002079414 |
| GO:0072163 | mesonephric epithelium development | 18/1966 | 0.00032199 | 0.002079414 |
| GO:0072164 | mesonephric tubule development | 18/1966 | 0.00032199 | 0.002079414 |

| | | | | |
|---|---|---|---|---|
| GO:2000243 | positive regulation of reproductive process | 18/1966 | 0.00032199 | 0.002079414 |
| GO:0030834 | regulation of actin filament depolymerization | Dec-66 | 0.000328649 | 0.002115163 |
| GO:0043536 | positive regulation of blood vessel endothelial cell migration | Dec-66 | 0.000328649 | 0.002115163 |
| GO:0051893 | regulation of focal adhesion assembly | 13/1966 | 0.000328954 | 0.002115163 |
| GO:0090109 | regulation of cell-substrate junction assembly | 13/1966 | 0.000328954 | 0.002115163 |
| GO:0006268 | DNA unwinding involved in DNA replication | Jul-66 | 0.00033774 | 0.002164608 |
| GO:0051797 | regulation of hair follicle development | Jul-66 | 0.00033774 | 0.002164608 |
| GO:0090594 | inflammatory response to wounding | Jul-66 | 0.00033774 | 0.002164608 |
| GO:0035850 | epithelial cell differentiation involved in kidney development | Oct-66 | 0.000338717 | 0.002166177 |
| GO:2000463 | positive regulation of excitatory postsynaptic potential | Oct-66 | 0.000338717 | 0.002166177 |
| GO:0018958 | phenol-containing compound metabolic process | 20/1966 | 0.000342293 | 0.002186552 |
| GO:0006749 | glutathione metabolic process | 15/1966 | 0.00034338 | 0.002186552 |
| GO:0032088 | negative regulation of NF-kappaB transcription factor activity | 15/1966 | 0.00034338 | 0.002186552 |
| GO:1901880 | negative regulation of protein depolymerization | 15/1966 | 0.00034338 | 0.002186552 |
| GO:1903037 | regulation of leukocyte cell-cell adhesion | 41/1966 | 0.000347129 | 0.002208052 |
| GO:0097553 | calcium ion transmembrane import into cytosol | 23/1966 | 0.00035262 | 0.002240572 |
| GO:0001892 | embryonic placenta development | 19/1966 | 0.000354395 | 0.002249438 |
| GO:0008593 | regulation of Notch signaling pathway | 16/1966 | 0.000358339 | 0.00227204 |

| | | | | |
|---|---|---|---|---|
| GO:0072676 | lymphocyte migration | 18/1966 | 0.000362123 | 0.002293574 |
| GO:0006672 | ceramide metabolic process | 17/1966 | 0.000363996 | 0.002302975 |
| GO:0000079 | regulation of cyclin-dependent protein serine/threonine kinase activity | 14/1966 | 0.000368431 | 0.002326066 |
| GO:0061045 | negative regulation of wound healing | 14/1966 | 0.000368431 | 0.002326066 |
| GO:1901985 | positive regulation of protein acetylation | Nov-66 | 0.000377188 | 0.002376588 |
| GO:2000045 | regulation of G1/S transition of mitotic cell cycle | 22/1966 | 0.000377236 | 0.002376588 |
| GO:0051101 | regulation of DNA binding | 20/1966 | 0.000380645 | 0.002395512 |
| GO:0048008 | platelet-derived growth factor receptor signaling pathway | 13/1966 | 0.00038582 | 0.002425504 |
| GO:0019722 | calcium-mediated signaling | 27/1966 | 0.00039017 | 0.002448591 |
| GO:0035567 | non-canonical Wnt signaling pathway | Dec-66 | 0.000390733 | 0.002448591 |
| GO:0140353 | lipid export from cell | Dec-66 | 0.000390733 | 0.002448591 |
| GO:0033673 | negative regulation of kinase activity | 30/1966 | 0.000394688 | 0.002470763 |
| GO:0060078 | regulation of postsynaptic membrane potential | 19/1966 | 0.000395825 | 0.002475263 |
| GO:0001959 | regulation of cytokine-mediated signaling pathway | 21/1966 | 0.000400901 | 0.002496462 |
| GO:0030278 | regulation of ossification | 21/1966 | 0.000400901 | 0.002496462 |
| GO:0042303 | molting cycle | 21/1966 | 0.000400901 | 0.002496462 |
| GO:0042633 | hair cycle | 21/1966 | 0.000400901 | 0.002496462 |
| GO:0032755 | positive regulation of interleukin-6 production | 18/1966 | 0.000406521 | 0.002528802 |

| | | | | |
|---|---|---|---|---|
| GO:0099173 | postsynapse organization | 28/1966 | 0.000415592 | 0.002575806 |
| GO:0018149 | peptide cross-linking | Oct-66 | 0.000417632 | 0.002575806 |
| GO:0090175 | regulation of establishment of planar polarity | Oct-66 | 0.000417632 | 0.002575806 |
| GO:0090224 | regulation of spindle organization | Oct-66 | 0.000417632 | 0.002575806 |
| GO:0110110 | positive regulation of animal organ morphogenesis | Oct-66 | 0.000417632 | 0.002575806 |
| GO:2000249 | regulation of actin cytoskeleton reorganization | Oct-66 | 0.000417632 | 0.002575806 |
| GO:0003094 | glomerular filtration | Aug-66 | 0.000418427 | 0.002575806 |
| GO:0032148 | activation of protein kinase B activity | Aug-66 | 0.000418427 | 0.002575806 |
| GO:0090077 | foam cell differentiation | Aug-66 | 0.000418427 | 0.002575806 |
| GO:1903055 | positive regulation of extracellular matrix organization | Aug-66 | 0.000418427 | 0.002575806 |
| GO:0048813 | dendrite morphogenesis | 26/1966 | 0.000428615 | 0.002635779 |
| GO:0045580 | regulation of T cell differentiation | 24/1966 | 0.000430043 | 0.002641819 |
| GO:0035025 | positive regulation of Rho protein signal transduction | Sep-66 | 0.000437585 | 0.002685367 |
| GO:0022600 | digestive system process | 19/1966 | 0.000441395 | 0.002698987 |
| GO:0045666 | positive regulation of neuron differentiation | 19/1966 | 0.000441395 | 0.002698987 |
| GO:0038065 | collagen-activated signaling pathway | Jun-66 | 0.000441503 | 0.002698987 |
| GO:0009410 | response to xenobiotic stimulus | 39/1966 | 0.000441628 | 0.002698987 |
| GO:0032984 | protein-containing complex disassembly | 31/1966 | 0.000442367 | 0.002700718 |

| | | | | |
|---|---|---|---|---|
| GO:0002720 | positive regulation of cytokine production involved in immune response | 13/1966 | 0.000450823 | 0.002745738 |
| GO:0055021 | regulation of cardiac muscle tissue growth | 15/1966 | 0.000451132 | 0.002745738 |
| GO:0072078 | nephron tubule morphogenesis | 15/1966 | 0.000451132 | 0.002745738 |
| GO:0001823 | mesonephros development | 18/1966 | 0.000455554 | 0.002766966 |
| GO:1901800 | positive regulation of proteasomal protein catabolic process | 18/1966 | 0.000455554 | 0.002766966 |
| GO:0007249 | I-kappaB kinase/NF-kappaB signaling | 30/1966 | 0.000458223 | 0.002780327 |
| GO:0021915 | neural tube development | 27/1966 | 0.000458888 | 0.002781509 |
| GO:0032465 | regulation of cytokinesis | 17/1966 | 0.000463356 | 0.002795025 |
| GO:0060021 | roof of mouth development | 17/1966 | 0.000463356 | 0.002795025 |
| GO:0090630 | activation of GTPase activity | 17/1966 | 0.000463356 | 0.002795025 |
| GO:0010743 | regulation of macrophage derived foam cell differentiation | Jul-66 | 0.000466309 | 0.002795025 |
| GO:0035634 | response to stilbenoid | Jul-66 | 0.000466309 | 0.002795025 |
| GO:0048486 | parasympathetic nervous system development | Jul-66 | 0.000466309 | 0.002795025 |
| GO:0060438 | trachea development | Jul-66 | 0.000466309 | 0.002795025 |
| GO:0060602 | branch elongation of an epithelium | Jul-66 | 0.000466309 | 0.002795025 |
| GO:0060749 | mammary gland alveolus development | Jul-66 | 0.000466309 | 0.002795025 |
| GO:0061377 | mammary gland lobule development | Jul-66 | 0.000466309 | 0.002795025 |
| GO:0090026 | positive regulation of monocyte chemotaxis | Jul-66 | 0.000466309 | 0.002795025 |

| | | | | |
|---|---|---|---|---|
| GO:2000278 | regulation of DNA biosynthetic process | 20/1966 | 0.000468701 | 0.002806519 |
| GO:2000116 | regulation of cysteine-type endopeptidase activity | 31/1966 | 0.000475455 | 0.002844085 |
| GO:0032908 | regulation of transforming growth factor beta1 production | May-66 | 0.000479718 | 0.00285804 |
| GO:0034115 | negative regulation of heterotypic cell-cell adhesion | May-66 | 0.000479718 | 0.00285804 |
| GO:0061517 | macrophage proliferation | May-66 | 0.000479718 | 0.00285804 |
| GO:0070587 | regulation of cell-cell adhesion involved in gastrulation | May-66 | 0.000479718 | 0.00285804 |
| GO:0030307 | positive regulation of cell growth | 28/1966 | 0.000485994 | 0.002892518 |
| GO:2000177 | regulation of neural precursor cell proliferation | 19/1966 | 0.000491444 | 0.00292011 |
| GO:0051149 | positive regulation of muscle cell differentiation | 14/1966 | 0.000491616 | 0.00292011 |
| GO:0002366 | leukocyte activation involved in immune response | 39/1966 | 0.000498557 | 0.002958372 |
| GO:0065004 | protein-DNA complex assembly | 22/1966 | 0.000502458 | 0.002978534 |
| GO:1902106 | negative regulation of leukocyte differentiation | 18/1966 | 0.000509614 | 0.003016573 |
| GO:0008217 | regulation of blood pressure | 29/1966 | 0.000509894 | 0.003016573 |
| GO:0045823 | positive regulation of heart contraction | Oct-66 | 0.000511242 | 0.003021532 |
| GO:0006809 | nitric oxide biosynthetic process | 15/1966 | 0.000515065 | 0.003032024 |
| GO:0032272 | negative regulation of protein polymerization | 15/1966 | 0.000515065 | 0.003032024 |
| GO:0043154 | negative regulation of cysteine-type endopeptidase activity involved in apoptotic process | 15/1966 | 0.000515065 | 0.003032024 |
| GO:0072089 | stem cell proliferation | 15/1966 | 0.000515065 | 0.003032024 |

| | | | | |
|---|---|---|---|---|
| GO:0002698 | negative regulation of immune effector process | 20/1966 | 0.000519006 | 0.003052189 |
| GO:0060395 | SMAD protein signal transduction | 17/1966 | 0.000521242 | 0.003062294 |
| GO:0043535 | regulation of blood vessel endothelial cell migration | 16/1966 | 0.000523964 | 0.00307204 |
| GO:0150076 | neuroinflammatory response | 13/1966 | 0.000524874 | 0.00307204 |
| GO:1990874 | vascular associated smooth muscle cell proliferation | 13/1966 | 0.000524874 | 0.00307204 |
| GO:0043281 | regulation of cysteine-type endopeptidase activity involved in apoptotic process | 28/1966 | 0.000524975 | 0.00307204 |
| GO:0007548 | sex differentiation | 38/1966 | 0.000529472 | 0.003095293 |
| GO:0045332 | phospholipid translocation | Nov-66 | 0.00054366 | 0.003175104 |
| GO:0045599 | negative regulation of fat cell differentiation | Dec-66 | 0.000544907 | 0.003179249 |
| GO:0034122 | negative regulation of toll-like receptor signaling pathway | Sep-66 | 0.000548434 | 0.003187268 |
| GO:0040001 | establishment of mitotic spindle localization | Sep-66 | 0.000548434 | 0.003187268 |
| GO:0051491 | positive regulation of filopodium assembly | Sep-66 | 0.000548434 | 0.003187268 |
| GO:1904037 | positive regulation of epithelial cell apoptotic process | Sep-66 | 0.000548434 | 0.003187268 |
| GO:1904029 | regulation of cyclin-dependent protein kinase activity | 14/1966 | 0.000565323 | 0.003282197 |
| GO:0055017 | cardiac muscle tissue growth | 18/1966 | 0.000569118 | 0.003300998 |
| GO:0060401 | cytosolic calcium ion transport | 27/1966 | 0.000581906 | 0.003371868 |
| GO:0060993 | kidney morphogenesis | 17/1966 | 0.000585236 | 0.003387845 |
| GO:0048469 | cell maturation | 29/1966 | 0.000591607 | 0.003414094 |

| | | | | |
|---|---|---|---|---|
| GO:1903828 | negative regulation of protein localization | 29/1966 | 0.000591607 | 0.003414094 |
| GO:0006970 | response to osmotic stress | 16/1966 | 0.000592076 | 0.003414094 |
| GO:0045103 | intermediate filament-based process | 16/1966 | 0.000592076 | 0.003414094 |
| GO:0032508 | DNA duplex unwinding | Oct-66 | 0.000621582 | 0.00358075 |
| GO:0051235 | maintenance of location | 40/1966 | 0.000628472 | 0.003612306 |
| GO:0002695 | negative regulation of leukocyte activation | 27/1966 | 0.00062889 | 0.003612306 |
| GO:0098656 | anion transmembrane transport | 27/1966 | 0.00062889 | 0.003612306 |
| GO:0035089 | establishment of apical/basal cell polarity | Jul-66 | 0.000631064 | 0.003614275 |
| GO:0036119 | response to platelet-derived growth factor | Jul-66 | 0.000631064 | 0.003614275 |
| GO:1902894 | negative regulation of miRNA transcription | Jul-66 | 0.000631064 | 0.003614275 |
| GO:1903707 | negative regulation of hemopoiesis | 18/1966 | 0.00063451 | 0.003630499 |
| GO:0030042 | actin filament depolymerization | Dec-66 | 0.000639389 | 0.003644335 |
| GO:0044380 | protein localization to cytoskeleton | Dec-66 | 0.000639389 | 0.003644335 |
| GO:0050922 | negative regulation of chemotaxis | Dec-66 | 0.000639389 | 0.003644335 |
| GO:0098815 | modulation of excitatory postsynaptic potential | Dec-66 | 0.000639389 | 0.003644335 |
| GO:0061308 | cardiac neural crest cell development involved in heart development | Jun-66 | 0.00064306 | 0.003661732 |
| GO:0030857 | negative regulation of epithelial cell differentiation | Nov-66 | 0.000647516 | 0.003676501 |
| GO:0090279 | regulation of calcium ion import | Nov-66 | 0.000647516 | 0.003676501 |

| | | | | |
|---|---|---|---|---|
| GO:1902041 | regulation of extrinsic apoptotic signaling pathway via death domain receptors | Nov-66 | 0.000647516 | 0.003676501 |
| GO:0051153 | regulation of striated muscle cell differentiation | 17/1966 | 0.000655856 | 0.003720286 |
| GO:0007204 | positive regulation of cytosolic calcium ion concentration | 41/1966 | 0.00065937 | 0.00373664 |
| GO:0046328 | regulation of JNK cascade | 22/1966 | 0.000662431 | 0.003750397 |
| GO:0046785 | microtubule polymerization | 15/1966 | 0.000666372 | 0.003769107 |
| GO:0045685 | regulation of glial cell differentiation | 16/1966 | 0.000667618 | 0.00377255 |
| GO:0006469 | negative regulation of protein kinase activity | 27/1966 | 0.00067916 | 0.003834116 |
| GO:0090313 | regulation of protein targeting to membrane | Sep-66 | 0.000681262 | 0.003842318 |
| GO:0090025 | regulation of monocyte chemotaxis | Aug-66 | 0.000697186 | 0.003928387 |
| GO:0003158 | endothelium development | 20/1966 | 0.000699002 | 0.003934877 |
| GO:0045668 | negative regulation of osteoblast differentiation | 13/1966 | 0.000704113 | 0.003952379 |
| GO:0060425 | lung morphogenesis | 13/1966 | 0.000704113 | 0.003952379 |
| GO:0150116 | regulation of cell-substrate junction organization | 13/1966 | 0.000704113 | 0.003952379 |
| GO:0042471 | ear morphogenesis | 21/1966 | 0.000716147 | 0.004016117 |
| GO:0034612 | response to tumor necrosis factor | 27/1966 | 0.000732908 | 0.004106225 |
| GO:0051651 | maintenance of location in cell | 29/1966 | 0.000735725 | 0.004118115 |
| GO:0032677 | regulation of interleukin-8 production | 14/1966 | 0.000741137 | 0.00414058 |
| GO:0043627 | response to estrogen | 14/1966 | 0.000741137 | 0.00414058 |

| | | | | |
|---|---|---|---|---|
| GO:0046620 | regulation of organ growth | 19/1966 | 0.000743914 | 0.004152176 |
| GO:0042551 | neuron maturation | Dec-66 | 0.000747221 | 0.004166704 |
| GO:0060411 | cardiac septum morphogenesis | 15/1966 | 0.000755212 | 0.004207305 |
| GO:0002819 | regulation of adaptive immune response | 30/1966 | 0.000757332 | 0.004215153 |
| GO:0002042 | cell migration involved in sprouting angiogenesis | Nov-66 | 0.00076736 | 0.004262951 |
| GO:0034204 | lipid translocation | Nov-66 | 0.00076736 | 0.004262951 |
| GO:0000018 | regulation of DNA recombination | 20/1966 | 0.000769907 | 0.004269093 |
| GO:0010212 | response to ionizing radiation | 20/1966 | 0.000769907 | 0.004269093 |
| GO:0002246 | wound healing involved in inflammatory response | May-66 | 0.000776213 | 0.004276031 |
| GO:0007183 | SMAD protein complex assembly | May-66 | 0.000776213 | 0.004276031 |
| GO:0021561 | facial nerve development | May-66 | 0.000776213 | 0.004276031 |
| GO:0021610 | facial nerve morphogenesis | May-66 | 0.000776213 | 0.004276031 |
| GO:0038063 | collagen-activated tyrosine kinase receptor signaling pathway | May-66 | 0.000776213 | 0.004276031 |
| GO:0060623 | regulation of chromosome condensation | May-66 | 0.000776213 | 0.004276031 |
| GO:0070586 | cell-cell adhesion involved in gastrulation | May-66 | 0.000776213 | 0.004276031 |
| GO:0035107 | appendage morphogenesis | 24/1966 | 0.00078348 | 0.004308051 |
| GO:0035108 | limb morphogenesis | 24/1966 | 0.00078348 | 0.004308051 |
| GO:0006665 | sphingolipid metabolic process | 21/1966 | 0.000785526 | 0.004315294 |

| | | | | |
|---|---|---|---|---|
| GO:0042098 | T cell proliferation | 29/1966 | 0.00079016 | 0.004336727 |
| GO:0042593 | glucose homeostasis | 35/1966 | 0.000808294 | 0.004432144 |
| GO:0006949 | syncytium formation | 13/1966 | 0.00081148 | 0.004445501 |
| GO:0002702 | positive regulation of production of molecular mediator of immune response | 19/1966 | 0.000822165 | 0.004499872 |
| GO:1903039 | positive regulation of leukocyte cell-cell adhesion | 31/1966 | 0.000828839 | 0.00453221 |
| GO:0002693 | positive regulation of cellular extravasation | Jul-66 | 0.0008388 | 0.004563624 |
| GO:0030903 | notochord development | Jul-66 | 0.0008388 | 0.004563624 |
| GO:0001953 | negative regulation of cell-matrix adhesion | Sep-66 | 0.000839208 | 0.004563624 |
| GO:0044786 | cell cycle DNA replication | Sep-66 | 0.000839208 | 0.004563624 |
| GO:0060236 | regulation of mitotic spindle organization | Sep-66 | 0.000839208 | 0.004563624 |
| GO:0098868 | bone growth | Sep-66 | 0.000839208 | 0.004563624 |
| GO:0032637 | interleukin-8 production | 14/1966 | 0.000845087 | 0.004582972 |
| GO:0070192 | chromosome organization involved in meiotic cell cycle | 14/1966 | 0.000845087 | 0.004582972 |
| GO:0071277 | cellular response to calcium ion | 14/1966 | 0.000845087 | 0.004582972 |
| GO:0006942 | regulation of striated muscle contraction | 15/1966 | 0.000853899 | 0.004626522 |
| GO:0033500 | carbohydrate homeostasis | 35/1966 | 0.000859448 | 0.00465233 |
| GO:0010972 | negative regulation of G2/M transition of mitotic cell cycle | Dec-66 | 0.000869834 | 0.004692473 |
| GO:0060612 | adipose tissue development | Dec-66 | 0.000869834 | 0.004692473 |

| | | | | |
|---|---|---|---|---|
| GO:1903078 | positive regulation of protein localization to plasma membrane | Dec-66 | 0.000869834 | 0.004692473 |
| GO:0098659 | inorganic cation import across plasma membrane | 18/1966 | 0.000870825 | 0.004692473 |
| GO:0099587 | inorganic ion import across plasma membrane | 18/1966 | 0.000870825 | 0.004692473 |
| GO:0035066 | positive regulation of histone acetylation | Aug-66 | 0.000883992 | 0.004741851 |
| GO:0043552 | positive regulation of phosphatidylinositol 3-kinase activity | Aug-66 | 0.000883992 | 0.004741851 |
| GO:0061037 | negative regulation of cartilage development | Aug-66 | 0.000883992 | 0.004741851 |
| GO:0086010 | membrane depolarization during action potential | Aug-66 | 0.000883992 | 0.004741851 |
| GO:1903319 | positive regulation of protein maturation | Aug-66 | 0.000883992 | 0.004741851 |
| GO:0030279 | negative regulation of ossification | Oct-66 | 0.000901481 | 0.00482012 |
| GO:0055023 | positive regulation of cardiac muscle tissue growth | Oct-66 | 0.000901481 | 0.00482012 |
| GO:0007019 | microtubule depolymerization | Nov-66 | 0.000905034 | 0.00482012 |
| GO:0035088 | establishment or maintenance of apical/basal cell polarity | Nov-66 | 0.000905034 | 0.00482012 |
| GO:0035272 | exocrine system development | Nov-66 | 0.000905034 | 0.00482012 |
| GO:0061245 | establishment or maintenance of bipolar cell polarity | Nov-66 | 0.000905034 | 0.00482012 |
| GO:1903052 | positive regulation of proteolysis involved in cellular protein catabolic process | 19/1966 | 0.000907376 | 0.00482012 |
| GO:0007614 | short-term memory | Jun-66 | 0.000909165 | 0.00482012 |
| GO:0060192 | negative regulation of lipase activity | Jun-66 | 0.000909165 | 0.00482012 |
| GO:0061307 | cardiac neural crest cell differentiation involved in heart development | Jun-66 | 0.000909165 | 0.00482012 |

| | | | | |
|---|---|---|---|---|
| GO:0061318 | renal filtration cell differentiation | Jun-66 | 0.000909165 | 0.00482012 |
| GO:0072112 | glomerular visceral epithelial cell differentiation | Jun-66 | 0.000909165 | 0.00482012 |
| GO:0072311 | glomerular epithelial cell differentiation | Jun-66 | 0.000909165 | 0.00482012 |
| GO:0008277 | regulation of G protein-coupled receptor signaling pathway | 20/1966 | 0.000930468 | 0.004928652 |
| GO:1902895 | positive regulation of miRNA transcription | 13/1966 | 0.000932257 | 0.004933714 |
| GO:0033135 | regulation of peptidyl-serine phosphorylation | 23/1966 | 0.000936208 | 0.004950198 |
| GO:0032091 | negative regulation of protein binding | 16/1966 | 0.00094545 | 0.004994607 |
| GO:2000573 | positive regulation of DNA biosynthetic process | 14/1966 | 0.000961059 | 0.005072543 |
| GO:0010517 | regulation of phospholipase activity | Dec-66 | 0.001008764 | 0.005314858 |
| GO:0031424 | keratinization | Dec-66 | 0.001008764 | 0.005314858 |
| GO:0048839 | inner ear development | 28/1966 | 0.001018809 | 0.00536301 |
| GO:0050773 | regulation of dendrite development | 20/1966 | 0.001020993 | 0.005369732 |
| GO:0030705 | cytoskeleton-dependent intracellular transport | 26/1966 | 0.001022179 | 0.0053712 |
| GO:0002931 | response to ischemia | Sep-66 | 0.001025666 | 0.005379976 |
| GO:0043304 | regulation of mast cell degranulation | Sep-66 | 0.001025666 | 0.005379976 |
| GO:0010469 | regulation of signaling receptor activity | 21/1966 | 0.001029557 | 0.005395609 |
| GO:0034605 | cellular response to heat | Nov-66 | 0.001062511 | 0.005563387 |
| GO:0007405 | neuroblast proliferation | 13/1966 | 0.001067723 | 0.005585737 |

| | | | | |
|---|---|---|---|---|
| GO:0030835 | negative regulation of actin filament depolymerization | Oct-66 | 0.001076002 | 0.005619121 |
| GO:0043114 | regulation of vascular permeability | Oct-66 | 0.001076002 | 0.005619121 |
| GO:0045665 | negative regulation of neuron differentiation | 15/1966 | 0.001084275 | 0.005657336 |
| GO:0002822 | regulation of adaptive immune response based on somatic recombination of immune receptors built from immunoglobulin superfamily domains | 28/1966 | 0.001093026 | 0.005693294 |
| GO:0002719 | negative regulation of cytokine production involved in immune response | Jul-66 | 0.001096935 | 0.005693294 |
| GO:0007413 | axonal fasciculation | Jul-66 | 0.001096935 | 0.005693294 |
| GO:0061162 | establishment of monopolar cell polarity | Jul-66 | 0.001096935 | 0.005693294 |
| GO:0106030 | neuron projection fasciculation | Jul-66 | 0.001096935 | 0.005693294 |
| GO:1905820 | positive regulation of chromosome separation | Jul-66 | 0.001096935 | 0.005693294 |
| GO:0000132 | establishment of mitotic spindle orientation | Aug-66 | 0.00110887 | 0.005735134 |
| GO:0034260 | negative regulation of GTPase activity | Aug-66 | 0.00110887 | 0.005735134 |
| GO:0051150 | regulation of smooth muscle cell differentiation | Aug-66 | 0.00110887 | 0.005735134 |
| GO:0060977 | coronary vasculature morphogenesis | Aug-66 | 0.00110887 | 0.005735134 |
| GO:0045667 | regulation of osteoblast differentiation | 22/1966 | 0.001118516 | 0.005779975 |
| GO:0071901 | negative regulation of protein serine/threonine kinase activity | 17/1966 | 0.00112868 | 0.005827414 |
| GO:1902750 | negative regulation of cell cycle G2/M phase transition | Dec-66 | 0.001165643 | 0.00600778 |
| GO:1904705 | regulation of vascular associated smooth muscle cell proliferation | Dec-66 | 0.001165643 | 0.00600778 |
| GO:0007623 | circadian rhythm | 28/1966 | 0.001171891 | 0.00603473 |

| | | | | |
|---|---|---|---|---|
| GO:0051209 | release of sequestered calcium ion into cytosol | 18/1966 | 0.001178639 | 0.006064207 |
| GO:0033625 | positive regulation of integrin activation | May-66 | 0.001190675 | 0.00609436 |
| GO:0044557 | relaxation of smooth muscle | May-66 | 0.001190675 | 0.00609436 |
| GO:0051639 | actin filament network formation | May-66 | 0.001190675 | 0.00609436 |
| GO:0060394 | negative regulation of pathway-restricted SMAD protein phosphorylation | May-66 | 0.001190675 | 0.00609436 |
| GO:0072393 | microtubule anchoring at microtubule organizing center | May-66 | 0.001190675 | 0.00609436 |
| GO:1901678 | iron coordination entity transport | May-66 | 0.001190675 | 0.00609436 |
| GO:0051100 | negative regulation of binding | 23/1966 | 0.001196951 | 0.006121197 |
| GO:0015914 | phospholipid transport | 15/1966 | 0.00121782 | 0.006213696 |
| GO:0032436 | positive regulation of proteasomal ubiquitin-dependent protein catabolic process | 15/1966 | 0.00121782 | 0.006213696 |
| GO:0032732 | positive regulation of interleukin-1 production | 13/1966 | 0.001219236 | 0.006213696 |
| GO:0060761 | negative regulation of response to cytokine stimulus | 13/1966 | 0.001219236 | 0.006213696 |
| GO:0034341 | response to interferon-gamma | 20/1966 | 0.001224861 | 0.006236998 |
| GO:0055007 | cardiac muscle cell differentiation | 21/1966 | 0.001226203 | 0.006238466 |
| GO:0044774 | mitotic DNA integrity checkpoint signaling | 14/1966 | 0.001233424 | 0.006264441 |
| GO:0051279 | regulation of release of sequestered calcium ion into cytosol | 14/1966 | 0.001233424 | 0.006264441 |
| GO:0001974 | blood vessel remodeling | Nov-66 | 0.0012419 | 0.0062913 |
| GO:0035019 | somatic stem cell population maintenance | Nov-66 | 0.0012419 | 0.0062913 |

| | | | | |
|---|---|---|---|---|
| GO:0072698 | protein localization to microtubule cytoskeleton | Nov-66 | 0.0012419 | 0.0062913 |
| GO:0003214 | cardiac left ventricle morphogenesis | Jun-66 | 0.001252539 | 0.00630207 |
| GO:0010544 | negative regulation of platelet activation | Jun-66 | 0.001252539 | 0.00630207 |
| GO:0035313 | wound healing, spreading of epidermal cells | Jun-66 | 0.001252539 | 0.00630207 |
| GO:0040037 | negative regulation of fibroblast growth factor receptor signaling pathway | Jun-66 | 0.001252539 | 0.00630207 |
| GO:0045198 | establishment of epithelial cell apical/basal polarity | Jun-66 | 0.001252539 | 0.00630207 |
| GO:0045837 | negative regulation of membrane potential | Jun-66 | 0.001252539 | 0.00630207 |
| GO:0071636 | positive regulation of transforming growth factor beta production | Jun-66 | 0.001252539 | 0.00630207 |
| GO:1905709 | negative regulation of membrane permeability | Jun-66 | 0.001252539 | 0.00630207 |
| GO:0042445 | hormone metabolic process | 28/1966 | 0.001255644 | 0.006312327 |
| GO:0001941 | postsynaptic membrane organization | Oct-66 | 0.001277179 | 0.006415146 |
| GO:0009266 | response to temperature stimulus | 23/1966 | 0.001296726 | 0.006507237 |
| GO:0018107 | peptidyl-threonine phosphorylation | 18/1966 | 0.001299909 | 0.006507237 |
| GO:0051283 | negative regulation of sequestering of calcium ion | 18/1966 | 0.001299909 | 0.006507237 |
| GO:0098773 | skin epidermis development | 18/1966 | 0.001299909 | 0.006507237 |
| GO:0050772 | positive regulation of axonogenesis | 16/1966 | 0.001315693 | 0.00658069 |
| GO:0043122 | regulation of I-kappaB kinase/NF-kappaB signaling | 25/1966 | 0.001322487 | 0.006609088 |
| GO:0051048 | negative regulation of secretion | 26/1966 | 0.001370299 | 0.00684225 |

| | | | | |
|---|---|---|---|---|
| GO:0033598 | mammary gland epithelial cell proliferation | Aug-66 | 0.001377132 | 0.006847501 |
| GO:0040036 | regulation of fibroblast growth factor receptor signaling pathway | Aug-66 | 0.001377132 | 0.006847501 |
| GO:0048566 | embryonic digestive tract development | Aug-66 | 0.001377132 | 0.006847501 |
| GO:1904646 | cellular response to amyloid-beta | Aug-66 | 0.001377132 | 0.006847501 |
| GO:1905314 | semi-lunar valve development | Aug-66 | 0.001377132 | 0.006847501 |
| GO:0010522 | regulation of calcium ion transport into cytosol | 17/1966 | 0.001385936 | 0.0068796 |
| GO:0030032 | lamellipodium assembly | 13/1966 | 0.001388234 | 0.0068796 |
| GO:0032623 | interleukin-2 production | 13/1966 | 0.001388234 | 0.0068796 |
| GO:0060675 | ureteric bud morphogenesis | 13/1966 | 0.001388234 | 0.0068796 |
| GO:0008625 | extrinsic apoptotic signaling pathway via death domain receptors | 14/1966 | 0.00139215 | 0.00688748 |
| GO:0033209 | tumor necrosis factor-mediated signaling pathway | 14/1966 | 0.00139215 | 0.00688748 |
| GO:1902115 | regulation of organelle assembly | 27/1966 | 0.001408858 | 0.006958096 |
| GO:0006582 | melanin metabolic process | Jul-66 | 0.001413473 | 0.006958096 |
| GO:0010742 | macrophage derived foam cell differentiation | Jul-66 | 0.001413473 | 0.006958096 |
| GO:0051307 | meiotic chromosome separation | Jul-66 | 0.001413473 | 0.006958096 |
| GO:0061339 | establishment or maintenance of monopolar cell polarity | Jul-66 | 0.001413473 | 0.006958096 |
| GO:0071276 | cellular response to cadmium ion | Jul-66 | 0.001413473 | 0.006958096 |
| GO:0009749 | response to glucose | 25/1966 | 0.001424073 | 0.007004451 |

| | | | | |
|---|---|---|---|---|
| GO:0098754 | detoxification | Nov-66 | 0.00144544 | 0.007097755 |
| GO:1903533 | regulation of protein targeting | Nov-66 | 0.00144544 | 0.007097755 |
| GO:0010657 | muscle cell apoptotic process | 16/1966 | 0.001463498 | 0.007174535 |
| GO:0045185 | maintenance of protein location | 16/1966 | 0.001463498 | 0.007174535 |
| GO:0006575 | cellular modified amino acid metabolic process | 26/1966 | 0.001471822 | 0.00720937 |
| GO:0051693 | actin filament capping | Sep-66 | 0.001498981 | 0.007336334 |
| GO:0033173 | calcineurin-NFAT signaling cascade | Oct-66 | 0.001507946 | 0.007368035 |
| GO:0060421 | positive regulation of heart growth | Oct-66 | 0.001507946 | 0.007368035 |
| GO:0030168 | platelet activation | 15/1966 | 0.001526649 | 0.007441001 |
| GO:0045104 | intermediate filament cytoskeleton organization | 15/1966 | 0.001526649 | 0.007441001 |
| GO:0048814 | regulation of dendrite morphogenesis | 15/1966 | 0.001526649 | 0.007441001 |
| GO:0048708 | astrocyte differentiation | 14/1966 | 0.001567567 | 0.007634156 |
| GO:0010769 | regulation of cell morphogenesis involved in differentiation | 18/1966 | 0.00157442 | 0.007638675 |
| GO:0034754 | cellular hormone metabolic process | 18/1966 | 0.00157442 | 0.007638675 |
| GO:0045621 | positive regulation of lymphocyte differentiation | 18/1966 | 0.00157442 | 0.007638675 |
| GO:0051282 | regulation of sequestering of calcium ion | 18/1966 | 0.00157442 | 0.007638675 |
| GO:0002886 | regulation of myeloid leukocyte mediated immunity | 13/1966 | 0.001576235 | 0.007638675 |
| GO:0072171 | mesonephric tubule morphogenesis | 13/1966 | 0.001576235 | 0.007638675 |

| | | | | |
|---|---|---|---|---|
| GO:0048736 | appendage development | 26/1966 | 0.001579753 | 0.007643217 |
| GO:0060173 | limb development | 26/1966 | 0.001579753 | 0.007643217 |
| GO:0015698 | inorganic anion transport | 21/1966 | 0.001581066 | 0.007643327 |
| GO:1903531 | negative regulation of secretion by cell | 23/1966 | 0.001640008 | 0.007921798 |
| GO:0031668 | cellular response to extracellular stimulus | 29/1966 | 0.001663616 | 0.008029287 |
| GO:0001523 | retinoid metabolic process | Nov-66 | 0.001675504 | 0.008080079 |
| GO:0060294 | cilium movement involved in cell motility | 22/1966 | 0.001683591 | 0.008095129 |
| GO:0010804 | negative regulation of tumor necrosis factor-mediated signaling pathway | Jun-66 | 0.001686827 | 0.008095129 |
| GO:0033599 | regulation of mammary gland epithelial cell proliferation | Jun-66 | 0.001686827 | 0.008095129 |
| GO:0036120 | cellular response to platelet-derived growth factor stimulus | Jun-66 | 0.001686827 | 0.008095129 |
| GO:1902430 | negative regulation of amyloid-beta formation | Jun-66 | 0.001686827 | 0.008095129 |
| GO:1904754 | positive regulation of vascular associated smooth muscle cell migration | Jun-66 | 0.001686827 | 0.008095129 |
| GO:0045639 | positive regulation of myeloid cell differentiation | 17/1966 | 0.001691052 | 0.008108836 |
| GO:0010758 | regulation of macrophage chemotaxis | Aug-66 | 0.001694454 | 0.008112011 |
| GO:1901186 | positive regulation of ERBB signaling pathway | Aug-66 | 0.001694454 | 0.008112011 |
| GO:0120193 | tight junction organization | 15/1966 | 0.0017041 | 0.008145019 |
| GO:2000117 | negative regulation of cysteine-type endopeptidase activity | 15/1966 | 0.0017041 | 0.008145019 |
| GO:0032368 | regulation of lipid transport | 18/1966 | 0.001729095 | 0.008257823 |

| | | | | |
|---|---|---|---|---|
| GO:0002237 | response to molecule of bacterial origin | 44/1966 | 0.001744959 | 0.008297345 |
| GO:0030206 | chondroitin sulfate biosynthetic process | May-66 | 0.001748579 | 0.008297345 |
| GO:0035791 | platelet-derived growth factor receptor-beta signaling pathway | May-66 | 0.001748579 | 0.008297345 |
| GO:0060670 | branching involved in labyrinthine layer morphogenesis | May-66 | 0.001748579 | 0.008297345 |
| GO:0061309 | cardiac neural crest cell development involved in outflow tract morphogenesis | May-66 | 0.001748579 | 0.008297345 |
| GO:0072224 | metanephric glomerulus development | May-66 | 0.001748579 | 0.008297345 |
| GO:0098840 | protein transport along microtubule | May-66 | 0.001748579 | 0.008297345 |
| GO:0099118 | microtubule-based protein transport | May-66 | 0.001748579 | 0.008297345 |
| GO:0042440 | pigment metabolic process | Dec-66 | 0.001761838 | 0.008353567 |
| GO:0009746 | response to hexose | 25/1966 | 0.00177007 | 0.008378841 |
| GO:0010862 | positive regulation of pathway-restricted SMAD protein phosphorylation | Oct-66 | 0.001771413 | 0.008378841 |
| GO:0090102 | cochlea development | Oct-66 | 0.001771413 | 0.008378841 |
| GO:1901184 | regulation of ERBB signaling pathway | 13/1966 | 0.001784835 | 0.008435591 |
| GO:0001774 | microglial cell activation | Sep-66 | 0.001793894 | 0.008439068 |
| GO:0010803 | regulation of tumor necrosis factor-mediated signaling pathway | Sep-66 | 0.001793894 | 0.008439068 |
| GO:0033006 | regulation of mast cell activation involved in immune response | Sep-66 | 0.001793894 | 0.008439068 |
| GO:0045066 | regulatory T cell differentiation | Sep-66 | 0.001793894 | 0.008439068 |
| GO:0007274 | neuromuscular synaptic transmission | Jul-66 | 0.001796972 | 0.008439068 |

| | | | | |
|---|---|---|---|---|
| GO:0031100 | animal organ regeneration | Jul-66 | 0.001796972 | 0.008439068 |
| GO:0032703 | negative regulation of interleukin-2 production | Jul-66 | 0.001796972 | 0.008439068 |
| GO:0070841 | inclusion body assembly | Jul-66 | 0.001796972 | 0.008439068 |
| GO:0010770 | positive regulation of cell morphogenesis involved in differentiation | 16/1966 | 0.001801229 | 0.008452361 |
| GO:0071356 | cellular response to tumor necrosis factor | 24/1966 | 0.00184242 | 0.008638805 |
| GO:0009101 | glycoprotein biosynthetic process | 32/1966 | 0.001888283 | 0.008846845 |
| GO:0071241 | cellular response to inorganic substance | 29/1966 | 0.001890237 | 0.008848999 |
| GO:0015718 | monocarboxylic acid transport | 18/1966 | 0.00189639 | 0.00886379 |
| GO:0051208 | sequestering of calcium ion | 18/1966 | 0.00189639 | 0.00886379 |
| GO:0003170 | heart valve development | Nov-66 | 0.0019346 | 0.008999764 |
| GO:0043388 | positive regulation of DNA binding | Nov-66 | 0.0019346 | 0.008999764 |
| GO:0050775 | positive regulation of dendrite morphogenesis | Nov-66 | 0.0019346 | 0.008999764 |
| GO:0051155 | positive regulation of striated muscle cell differentiation | Nov-66 | 0.0019346 | 0.008999764 |
| GO:0051445 | regulation of meiotic cell cycle | Nov-66 | 0.0019346 | 0.008999764 |
| GO:0097035 | regulation of membrane lipid distribution | Nov-66 | 0.0019346 | 0.008999764 |
| GO:0070665 | positive regulation of leukocyte proliferation | 22/1966 | 0.001969706 | 0.009155887 |
| GO:0006643 | membrane lipid metabolic process | 24/1966 | 0.001981619 | 0.009204038 |
| GO:0000768 | syncytium formation by plasma membrane fusion | Dec-66 | 0.002008885 | 0.009301497 |

| | | | | |
|---|---|---|---|---|
| GO:0030837 | negative regulation of actin filament polymerization | Dec-66 | 0.002008885 | 0.009301497 |
| GO:0044458 | motile cilium assembly | Dec-66 | 0.002008885 | 0.009301497 |
| GO:0140253 | cell-cell fusion | Dec-66 | 0.002008885 | 0.009301497 |
| GO:1903076 | regulation of protein localization to plasma membrane | 17/1966 | 0.002050783 | 0.009488075 |
| GO:0001702 | gastrulation with mouth forming second | Aug-66 | 0.00206686 | 0.009506716 |
| GO:0003401 | axis elongation | Aug-66 | 0.00206686 | 0.009506716 |
| GO:0010667 | negative regulation of cardiac muscle cell apoptotic process | Aug-66 | 0.00206686 | 0.009506716 |
| GO:0090218 | positive regulation of lipid kinase activity | Aug-66 | 0.00206686 | 0.009506716 |
| GO:1904738 | vascular associated smooth muscle cell migration | Aug-66 | 0.00206686 | 0.009506716 |
| GO:0006687 | glycosphingolipid metabolic process | Oct-66 | 0.002070865 | 0.009506716 |
| GO:0007520 | myoblast fusion | Oct-66 | 0.002070865 | 0.009506716 |
| GO:0032392 | DNA geometric change | Oct-66 | 0.002070865 | 0.009506716 |
| GO:0044818 | mitotic G2/M transition checkpoint | Oct-66 | 0.002070865 | 0.009506716 |
| GO:1904407 | positive regulation of nitric oxide metabolic process | Oct-66 | 0.002070865 | 0.009506716 |
| GO:0032388 | positive regulation of intracellular transport | 24/1966 | 0.002129685 | 0.009763609 |
| GO:0032924 | activin receptor signaling pathway | Sep-66 | 0.00213342 | 0.009763609 |
| GO:0048873 | homeostasis of number of cells within a tissue | Sep-66 | 0.00213342 | 0.009763609 |
| GO:0086002 | cardiac muscle cell action potential involved in contraction | Sep-66 | 0.00213342 | 0.009763609 |

| | | | | |
|---|---|---|---|---|
| GO:0031348 | negative regulation of defense response | 29/1966 | 0.002142909 | 0.009799463 |
| GO:0097193 | intrinsic apoptotic signaling pathway | 35/1966 | 0.002161352 | 0.009876179 |
| GO:0034284 | response to monosaccharide | 25/1966 | 0.002185648 | 0.009979498 |

**GO terms associated with 4473 Rep_intermed_enriched_vs_Hep genes**

| ID | Description | Gene Ratio | pvalue | p.adjust |
|---|---|---|---|---|
| GO:0007015 | actin filament organization | 172/3805 | 5.01E-40 | 1.82E-36 |
| GO:0044782 | cilium organization | 155/3805 | 5.82E-40 | 1.82E-36 |
| GO:0060271 | cilium assembly | 140/3805 | 1.79E-35 | 3.73E-32 |
| GO:0032970 | regulation of actin filament-based process | 153/3805 | 8.91E-35 | 1.39E-31 |
| GO:1902903 | regulation of supramolecular fiber organization | 146/3805 | 3.01E-34 | 3.76E-31 |
| GO:0050808 | synapse organization | 168/3805 | 6.23E-33 | 6.49E-30 |
| GO:0034329 | cell junction assembly | 155/3805 | 2.60E-32 | 2.32E-29 |
| GO:0001667 | ameboidal-type cell migration | 155/3805 | 6.27E-31 | 4.90E-28 |
| GO:0030198 | extracellular matrix organization | 123/3805 | 7.90E-31 | 5.48E-28 |
| GO:0043062 | extracellular structure organization | 123/3805 | 1.13E-30 | 7.05E-28 |
| GO:0045229 | external encapsulating structure organization | 123/3805 | 1.61E-30 | 9.13E-28 |
| GO:0007264 | small GTPase mediated signal transduction | 149/3805 | 3.28E-30 | 1.71E-27 |
| GO:0007409 | axonogenesis | 162/3805 | 7.66E-30 | 3.68E-27 |
| GO:0032956 | regulation of actin cytoskeleton organization | 133/3805 | 1.54E-29 | 6.89E-27 |
| GO:0060562 | epithelial tube morphogenesis | 138/3805 | 1.35E-28 | 5.63E-26 |
| GO:0048762 | mesenchymal cell differentiation | 99/3805 | 6.13E-27 | 2.39E-24 |

| | | | | |
|---|---|---|---|---|
| GO:0060485 | mesenchyme development | 112/3805 | 2.27E-26 | 8.33E-24 |
| GO:0007018 | microtubule-based movement | 140/3805 | 5.87E-26 | 2.04E-23 |
| GO:0045216 | cell-cell junction organization | 90/3805 | 2.66E-25 | 8.74E-23 |
| GO:0001655 | urogenital system development | 130/3805 | 3.46E-25 | 1.08E-22 |
| GO:0031032 | actomyosin structure organization | 89/3805 | 3.85E-25 | 1.15E-22 |
| GO:0001822 | kidney development | 117/3805 | 4.53E-25 | 1.29E-22 |
| GO:0072001 | renal system development | 120/3805 | 7.67E-25 | 2.08E-22 |
| GO:0031346 | positive regulation of cell projection organization | 141/3805 | 1.34E-24 | 3.50E-22 |
| GO:0033674 | positive regulation of kinase activity | 139/3805 | 1.89E-24 | 4.71E-22 |
| GO:0007178 | transmembrane receptor protein serine/threonine kinase signaling pathway | 129/3805 | 3.42E-24 | 8.20E-22 |
| GO:0010639 | negative regulation of organelle organization | 124/3805 | 1.24E-23 | 2.76E-21 |
| GO:0031589 | cell-substrate adhesion | 124/3805 | 1.24E-23 | 2.76E-21 |
| GO:0071900 | regulation of protein serine/threonine kinase activity | 119/3805 | 1.95E-23 | 4.21E-21 |
| GO:0007265 | Ras protein signal transduction | 116/3805 | 5.16E-23 | 1.07E-20 |
| GO:0043254 | regulation of protein-containing complex assembly | 134/3805 | 1.33E-22 | 2.68E-20 |
| GO:0045860 | positive regulation of protein kinase activity | 118/3805 | 3.32E-22 | 6.48E-20 |
| GO:0090287 | regulation of cellular response to growth factor stimulus | 105/3805 | 3.82E-22 | 7.22E-20 |
| GO:0022604 | regulation of cell morphogenesis | 113/3805 | 7.02E-22 | 1.29E-19 |

| | | | | |
|---|---|---|---|---|
| GO:0060560 | developmental growth involved in morphogenesis | 100/3805 | 9.13E-22 | 1.63E-19 |
| GO:0016358 | dendrite development | 108/3805 | 1.08E-21 | 1.83E-19 |
| GO:0090130 | tissue migration | 108/3805 | 1.08E-21 | 1.83E-19 |
| GO:0010631 | epithelial cell migration | 107/3805 | 1.24E-21 | 2.04E-19 |
| GO:0050767 | regulation of neurogenesis | 137/3805 | 1.89E-21 | 3.03E-19 |
| GO:0090132 | epithelium migration | 107/3805 | 2.16E-21 | 3.38E-19 |
| GO:0051056 | regulation of small GTPase mediated signal transduction | 88/3805 | 2.67E-21 | 4.07E-19 |
| GO:0110053 | regulation of actin filament organization | 98/3805 | 2.81E-21 | 4.18E-19 |
| GO:0045785 | positive regulation of cell adhesion | 142/3805 | 3.29E-21 | 4.79E-19 |
| GO:0071559 | response to transforming growth factor beta | 87/3805 | 3.99E-21 | 5.66E-19 |
| GO:0071560 | cellular response to transforming growth factor beta stimulus | 86/3805 | 4.24E-21 | 5.89E-19 |
| GO:0001837 | epithelial to mesenchymal transition | 68/3805 | 5.83E-21 | 7.92E-19 |
| GO:0042692 | muscle cell differentiation | 132/3805 | 9.41E-21 | 1.25E-18 |
| GO:0198738 | cell-cell signaling by wnt | 135/3805 | 1.03E-20 | 1.34E-18 |
| GO:0061572 | actin filament bundle organization | 73/3805 | 1.18E-20 | 1.50E-18 |
| GO:0040013 | negative regulation of locomotion | 111/3805 | 1.32E-20 | 1.65E-18 |
| GO:0051017 | actin filament bundle assembly | 72/3805 | 1.70E-20 | 2.09E-18 |
| GO:0032535 | regulation of cellular component size | 122/3805 | 1.89E-20 | 2.26E-18 |

| | | | | |
|---|---|---|---|---|
| GO:0016055 | Wnt signaling pathway | 134/3805 | 1.92E-20 | 2.26E-18 |
| GO:0003007 | heart morphogenesis | 98/3805 | 2.92E-20 | 3.38E-18 |
| GO:0070371 | ERK1 and ERK2 cascade | 112/3805 | 4.00E-20 | 4.54E-18 |
| GO:0007163 | establishment or maintenance of cell polarity | 85/3805 | 4.68E-20 | 5.17E-18 |
| GO:0043087 | regulation of GTPase activity | 111/3805 | 4.71E-20 | 5.17E-18 |
| GO:0001558 | regulation of cell growth | 135/3805 | 5.63E-20 | 6.07E-18 |
| GO:0030010 | establishment of cell polarity | 65/3805 | 3.31E-19 | 3.51E-17 |
| GO:0051258 | protein polymerization | 94/3805 | 4.72E-19 | 4.91E-17 |
| GO:0010632 | regulation of epithelial cell migration | 86/3805 | 5.25E-19 | 5.38E-17 |
| GO:0097485 | neuron projection guidance | 90/3805 | 5.98E-19 | 6.02E-17 |
| GO:1902905 | positive regulation of supramolecular fiber organization | 72/3805 | 7.59E-19 | 7.52E-17 |
| GO:0000280 | nuclear division | 129/3805 | 9.60E-19 | 9.37E-17 |
| GO:1901342 | regulation of vasculature development | 101/3805 | 1.08E-18 | 1.04E-16 |
| GO:0070372 | regulation of ERK1 and ERK2 cascade | 104/3805 | 1.40E-18 | 1.33E-16 |
| GO:0010563 | negative regulation of phosphorus metabolic process | 131/3805 | 1.45E-18 | 1.33E-16 |
| GO:0045936 | negative regulation of phosphate metabolic process | 131/3805 | 1.45E-18 | 1.33E-16 |
| GO:0001763 | morphogenesis of a branching structure | 87/3805 | 1.54E-18 | 1.39E-16 |
| GO:0007411 | axon guidance | 89/3805 | 1.60E-18 | 1.43E-16 |

| | | | | |
|---|---|---|---|---|
| GO:0003012 | muscle system process | 125/3805 | 2.60E-18 | 2.28E-16 |
| GO:0090092 | regulation of transmembrane receptor protein serine/threonine kinase signaling pathway | 88/3805 | 3.22E-18 | 2.79E-16 |
| GO:0007224 | smoothened signaling pathway | 65/3805 | 3.65E-18 | 3.12E-16 |
| GO:0045765 | regulation of angiogenesis | 99/3805 | 4.09E-18 | 3.45E-16 |
| GO:0051495 | positive regulation of cytoskeleton organization | 76/3805 | 4.72E-18 | 3.93E-16 |
| GO:0051271 | negative regulation of cellular component movement | 100/3805 | 7.31E-18 | 6.01E-16 |
| GO:0071902 | positive regulation of protein serine/threonine kinase activity | 76/3805 | 8.99E-18 | 7.29E-16 |
| GO:1902904 | negative regulation of supramolecular fiber organization | 69/3805 | 1.02E-17 | 8.20E-16 |
| GO:0060537 | muscle tissue development | 136/3805 | 1.14E-17 | 9.04E-16 |
| GO:0022407 | regulation of cell-cell adhesion | 131/3805 | 1.25E-17 | 9.74E-16 |
| GO:0032271 | regulation of protein polymerization | 76/3805 | 1.69E-17 | 1.26E-15 |
| GO:0099111 | microtubule-based transport | 76/3805 | 1.69E-17 | 1.26E-15 |
| GO:0007389 | pattern specification process | 135/3805 | 1.69E-17 | 1.26E-15 |
| GO:0001578 | microtubule bundle formation | 55/3805 | 1.70E-17 | 1.26E-15 |
| GO:0001654 | eye development | 121/3805 | 2.38E-17 | 1.75E-15 |
| GO:0048588 | developmental cell growth | 91/3805 | 2.41E-17 | 1.75E-15 |
| GO:0031345 | negative regulation of cell projection organization | 77/3805 | 2.84E-17 | 2.03E-15 |
| GO:0034765 | regulation of ion transmembrane transport | 136/3805 | 2.86E-17 | 2.03E-15 |

| | | | | |
|---|---|---|---|---|
| GO:0007179 | transforming growth factor beta receptor signaling pathway | 70/3805 | 3.81E-17 | 2.67E-15 |
| GO:0150063 | visual system development | 121/3805 | 4.34E-17 | 3.01E-15 |
| GO:0060070 | canonical Wnt signaling pathway | 96/3805 | 6.20E-17 | 4.25E-15 |
| GO:0051090 | regulation of DNA-binding transcription factor activity | 123/3805 | 6.36E-17 | 4.32E-15 |
| GO:0061448 | connective tissue development | 94/3805 | 6.80E-17 | 4.57E-15 |
| GO:0006936 | muscle contraction | 98/3805 | 7.07E-17 | 4.70E-15 |
| GO:0046578 | regulation of Ras protein signal transduction | 73/3805 | 7.86E-17 | 5.12E-15 |
| GO:0048638 | regulation of developmental growth | 116/3805 | 7.86E-17 | 5.12E-15 |
| GO:0032886 | regulation of microtubule-based process | 88/3805 | 8.35E-17 | 5.37E-15 |
| GO:0051494 | negative regulation of cytoskeleton organization | 67/3805 | 9.04E-17 | 5.76E-15 |
| GO:0048880 | sensory system development | 121/3805 | 9.53E-17 | 6.01E-15 |
| GO:0007346 | regulation of mitotic cell cycle | 131/3805 | 9.71E-17 | 6.06E-15 |
| GO:0030336 | negative regulation of cell migration | 93/3805 | 1.03E-16 | 6.35E-15 |
| GO:0045926 | negative regulation of growth | 87/3805 | 1.26E-16 | 7.73E-15 |
| GO:0050673 | epithelial cell proliferation | 130/3805 | 1.43E-16 | 8.69E-15 |
| GO:0010959 | regulation of metal ion transport | 124/3805 | 1.63E-16 | 9.80E-15 |
| GO:2000146 | negative regulation of cell motility | 95/3805 | 2.43E-16 | 1.44E-14 |
| GO:0045787 | positive regulation of cell cycle | 105/3805 | 3.60E-16 | 2.12E-14 |

| | | | | |
|---|---|---|---|---|
| GO:0050678 | regulation of epithelial cell proliferation | 115/3805 | 3.88E-16 | 2.27E-14 |
| GO:0007611 | learning or memory | 98/3805 | 4.48E-16 | 2.58E-14 |
| GO:0048285 | organelle fission | 133/3805 | 4.51E-16 | 2.58E-14 |
| GO:1901888 | regulation of cell junction assembly | 76/3805 | 4.55E-16 | 2.58E-14 |
| GO:0007162 | negative regulation of cell adhesion | 96/3805 | 5.13E-16 | 2.89E-14 |
| GO:0010976 | positive regulation of neuron projection development | 77/3805 | 5.31E-16 | 2.96E-14 |
| GO:0061138 | morphogenesis of a branching epithelium | 78/3805 | 6.13E-16 | 3.39E-14 |
| GO:0070997 | neuron death | 121/3805 | 6.39E-16 | 3.50E-14 |
| GO:0018108 | peptidyl-tyrosine phosphorylation | 98/3805 | 7.00E-16 | 3.80E-14 |
| GO:0030900 | forebrain development | 113/3805 | 7.06E-16 | 3.80E-14 |
| GO:0043405 | regulation of MAP kinase activity | 71/3805 | 8.24E-16 | 4.40E-14 |
| GO:0050890 | cognition | 105/3805 | 8.35E-16 | 4.42E-14 |
| GO:0042326 | negative regulation of phosphorylation | 113/3805 | 1.04E-15 | 5.48E-14 |
| GO:0018212 | peptidyl-tyrosine modification | 98/3805 | 1.36E-15 | 7.06E-14 |
| GO:0030111 | regulation of Wnt signaling pathway | 96/3805 | 1.58E-15 | 8.16E-14 |
| GO:0007160 | cell-matrix adhesion | 78/3805 | 1.82E-15 | 9.30E-14 |
| GO:0050803 | regulation of synapse structure or activity | 86/3805 | 1.84E-15 | 9.34E-14 |
| GO:0060541 | respiratory system development | 85/3805 | 2.17E-15 | 1.09E-13 |

| | | | | |
|---|---|---|---|---|
| GO:0042060 | wound healing | 107/3805 | 2.35E-15 | 1.16E-13 |
| GO:0043010 | camera-type eye development | 107/3805 | 2.35E-15 | 1.16E-13 |
| GO:0050807 | regulation of synapse organization | 84/3805 | 2.57E-15 | 1.26E-13 |
| GO:0070507 | regulation of microtubule cytoskeleton organization | 61/3805 | 2.74E-15 | 1.34E-13 |
| GO:0051261 | protein depolymerization | 52/3805 | 3.36E-15 | 1.63E-13 |
| GO:1901879 | regulation of protein depolymerization | 44/3805 | 3.91E-15 | 1.88E-13 |
| GO:0043542 | endothelial cell migration | 76/3805 | 4.15E-15 | 1.98E-13 |
| GO:0010721 | negative regulation of cell development | 71/3805 | 4.75E-15 | 2.25E-13 |
| GO:1903829 | positive regulation of protein localization | 129/3805 | 4.86E-15 | 2.28E-13 |
| GO:0051961 | negative regulation of nervous system development | 62/3805 | 4.95E-15 | 2.31E-13 |
| GO:0007043 | cell-cell junction assembly | 59/3805 | 6.09E-15 | 2.80E-13 |
| GO:0008154 | actin polymerization or depolymerization | 69/3805 | 6.10E-15 | 2.80E-13 |
| GO:0050770 | regulation of axonogenesis | 66/3805 | 6.39E-15 | 2.91E-13 |
| GO:0140014 | mitotic nuclear division | 87/3805 | 6.46E-15 | 2.92E-13 |
| GO:0030323 | respiratory tube development | 78/3805 | 6.73E-15 | 3.03E-13 |
| GO:0002064 | epithelial cell development | 77/3805 | 7.86E-15 | 3.51E-13 |
| GO:0010634 | positive regulation of epithelial cell migration | 59/3805 | 8.54E-15 | 3.78E-13 |
| GO:0048017 | inositol lipid-mediated signaling | 63/3805 | 8.62E-15 | 3.79E-13 |

| | | | | |
|---|---|---|---|---|
| GO:0030324 | lung development | 77/3805 | 1.02E-14 | 4.44E-13 |
| GO:0010810 | regulation of cell-substrate adhesion | 75/3805 | 1.07E-14 | 4.62E-13 |
| GO:0003205 | cardiac chamber development | 69/3805 | 1.08E-14 | 4.67E-13 |
| GO:0008360 | regulation of cell shape | 59/3805 | 1.19E-14 | 5.10E-13 |
| GO:0048015 | phosphatidylinositol-mediated signaling | 62/3805 | 1.29E-14 | 5.50E-13 |
| GO:0048732 | gland development | 125/3805 | 1.34E-14 | 5.64E-13 |
| GO:0051146 | striated muscle cell differentiation | 96/3805 | 1.36E-14 | 5.69E-13 |
| GO:0010720 | positive regulation of cell development | 106/3805 | 1.37E-14 | 5.70E-13 |
| GO:0050768 | negative regulation of neurogenesis | 60/3805 | 2.12E-14 | 8.78E-13 |
| GO:0030705 | cytoskeleton-dependent intracellular transport | 68/3805 | 2.17E-14 | 8.93E-13 |
| GO:0043406 | positive regulation of MAP kinase activity | 52/3805 | 2.30E-14 | 9.39E-13 |
| GO:0030177 | positive regulation of Wnt signaling pathway | 54/3805 | 3.12E-14 | 1.27E-12 |
| GO:0051216 | cartilage development | 71/3805 | 3.27E-14 | 1.32E-12 |
| GO:0030308 | negative regulation of cell growth | 69/3805 | 3.32E-14 | 1.32E-12 |
| GO:1990138 | neuron projection extension | 69/3805 | 3.32E-14 | 1.32E-12 |
| GO:0035148 | tube formation | 64/3805 | 3.56E-14 | 1.41E-12 |
| GO:0001764 | neuron migration | 67/3805 | 4.35E-14 | 1.71E-12 |
| GO:0090257 | regulation of muscle system process | 80/3805 | 4.45E-14 | 1.74E-12 |

| | | | | |
|---|---|---|---|---|
| GO:0031109 | microtubule polymerization or depolymerization | 51/3805 | 4.90E-14 | 1.89E-12 |
| GO:0048640 | negative regulation of developmental growth | 51/3805 | 4.90E-14 | 1.89E-12 |
| GO:0048608 | reproductive structure development | 125/3805 | 4.97E-14 | 1.90E-12 |
| GO:0001894 | tissue homeostasis | 86/3805 | 5.93E-14 | 2.26E-12 |
| GO:0051098 | regulation of binding | 108/3805 | 7.00E-14 | 2.65E-12 |
| GO:0030048 | actin filament-based movement | 51/3805 | 7.07E-14 | 2.66E-12 |
| GO:0021915 | neural tube development | 67/3805 | 7.58E-14 | 2.83E-12 |
| GO:0001933 | negative regulation of protein phosphorylation | 99/3805 | 8.29E-14 | 3.08E-12 |
| GO:0016331 | morphogenesis of embryonic epithelium | 64/3805 | 8.51E-14 | 3.14E-12 |
| GO:0061458 | reproductive system development | 125/3805 | 9.42E-14 | 3.46E-12 |
| GO:0061351 | neural precursor cell proliferation | 67/3805 | 9.96E-14 | 3.64E-12 |
| GO:0060249 | anatomical structure homeostasis | 94/3805 | 1.03E-13 | 3.74E-12 |
| GO:0001838 | embryonic epithelial tube formation | 57/3805 | 1.35E-13 | 4.89E-12 |
| GO:1990778 | protein localization to cell periphery | 102/3805 | 1.40E-13 | 5.01E-12 |
| GO:0014706 | striated muscle tissue development | 86/3805 | 1.41E-13 | 5.04E-12 |
| GO:1904062 | regulation of cation transmembrane transport | 104/3805 | 1.60E-13 | 5.68E-12 |
| GO:0010594 | regulation of endothelial cell migration | 61/3805 | 1.64E-13 | 5.78E-12 |
| GO:1901214 | regulation of neuron death | 106/3805 | 1.81E-13 | 6.33E-12 |

| | | | | |
|---|---|---|---|---|
| GO:0046777 | protein autophosphorylation | 76/3805 | 1.81E-13 | 6.33E-12 |
| GO:0030856 | regulation of epithelial cell differentiation | 57/3805 | 1.86E-13 | 6.44E-12 |
| GO:0043244 | regulation of protein-containing complex disassembly | 51/3805 | 2.06E-13 | 7.11E-12 |
| GO:1903844 | regulation of cellular response to transforming growth factor beta stimulus | 49/3805 | 2.21E-13 | 7.60E-12 |
| GO:0050920 | regulation of chemotaxis | 73/3805 | 2.38E-13 | 8.12E-12 |
| GO:0010970 | transport along microtubule | 59/3805 | 2.77E-13 | 9.36E-12 |
| GO:0072175 | epithelial tube formation | 59/3805 | 2.77E-13 | 9.36E-12 |
| GO:0001503 | ossification | 111/3805 | 3.02E-13 | 1.01E-11 |
| GO:0017015 | regulation of transforming growth factor beta receptor signaling pathway | 48/3805 | 3.26E-13 | 1.09E-11 |
| GO:0007517 | muscle organ development | 100/3805 | 3.65E-13 | 1.21E-11 |
| GO:0007266 | Rho protein signal transduction | 52/3805 | 3.85E-13 | 1.27E-11 |
| GO:0140694 | non-membrane-bounded organelle assembly | 103/3805 | 4.04E-13 | 1.33E-11 |
| GO:1903706 | regulation of hemopoiesis | 107/3805 | 4.20E-13 | 1.37E-11 |
| GO:0030041 | actin filament polymerization | 59/3805 | 4.99E-13 | 1.62E-11 |
| GO:0035082 | axoneme assembly | 40/3805 | 5.77E-13 | 1.86E-11 |
| GO:0043242 | negative regulation of protein-containing complex disassembly | 40/3805 | 5.77E-13 | 1.86E-11 |
| GO:0051403 | stress-activated MAPK cascade | 76/3805 | 5.80E-13 | 1.86E-11 |
| GO:0048754 | branching morphogenesis of an epithelial tube | 64/3805 | 5.90E-13 | 1.88E-11 |

| | | | | |
|---|---|---|---|---|
| GO:0030199 | collagen fibril organization | 31/3805 | 6.18E-13 | 1.96E-11 |
| GO:0010717 | regulation of epithelial to mesenchymal transition | 41/3805 | 6.27E-13 | 1.98E-11 |
| GO:0031098 | stress-activated protein kinase signaling cascade | 78/3805 | 6.32E-13 | 1.98E-11 |
| GO:0051962 | positive regulation of nervous system development | 98/3805 | 7.94E-13 | 2.48E-11 |
| GO:2001233 | regulation of apoptotic signaling pathway | 107/3805 | 8.29E-13 | 2.57E-11 |
| GO:1901987 | regulation of cell cycle phase transition | 106/3805 | 8.63E-13 | 2.67E-11 |
| GO:0031110 | regulation of microtubule polymerization or depolymerization | 41/3805 | 9.54E-13 | 2.94E-11 |
| GO:0050679 | positive regulation of epithelial cell proliferation | 69/3805 | 9.66E-13 | 2.96E-11 |
| GO:0042063 | gliogenesis | 96/3805 | 9.99E-13 | 3.03E-11 |
| GO:1901880 | negative regulation of protein depolymerization | 37/3805 | 1.00E-12 | 3.03E-11 |
| GO:0030038 | contractile actin filament bundle assembly | 46/3805 | 1.02E-12 | 3.07E-11 |
| GO:0043149 | stress fiber assembly | 46/3805 | 1.02E-12 | 3.07E-11 |
| GO:0007051 | spindle organization | 62/3805 | 1.03E-12 | 3.08E-11 |
| GO:0006816 | calcium ion transport | 113/3805 | 1.20E-12 | 3.57E-11 |
| GO:0043393 | regulation of protein binding | 69/3805 | 1.23E-12 | 3.65E-11 |
| GO:0050730 | regulation of peptidyl-tyrosine phosphorylation | 79/3805 | 1.26E-12 | 3.70E-11 |
| GO:0009799 | specification of symmetry | 52/3805 | 1.42E-12 | 4.16E-11 |
| GO:0008361 | regulation of cell size | 68/3805 | 1.46E-12 | 4.25E-11 |

| | | | | |
|---|---|---|---|---|
| GO:003 2409 | regulation of transporter activity | 85/38 05 | 2.03E-12 | 5.90E-11 |
| GO:005 0900 | leukocyte migration | 101/3 805 | 2.08E-12 | 6.01E-11 |
| GO:007 1675 | regulation of mononuclear cell migration | 46/38 05 | 2.10E-12 | 6.06E-11 |
| GO:006 0828 | regulation of canonical Wnt signaling pathway | 75/38 05 | 2.14E-12 | 6.12E-11 |
| GO:000 3206 | cardiac chamber morphogenesis | 54/38 05 | 2.16E-12 | 6.15E-11 |
| GO:001 4909 | smooth muscle cell migration | 43/38 05 | 2.23E-12 | 6.35E-11 |
| GO:003 3002 | muscle cell proliferation | 77/38 05 | 2.27E-12 | 6.42E-11 |
| GO:190 2850 | microtubule cytoskeleton organization involved in mitosis | 53/38 05 | 2.40E-12 | 6.75E-11 |
| GO:009 9173 | postsynapse organization | 66/38 05 | 2.60E-12 | 7.27E-11 |
| GO:001 0977 | negative regulation of neuron projection development | 56/38 05 | 3.06E-12 | 8.53E-11 |
| GO:003 2231 | regulation of actin filament bundle assembly | 45/38 05 | 3.12E-12 | 8.64E-11 |
| GO:001 4910 | regulation of smooth muscle cell migration | 40/38 05 | 3.13E-12 | 8.64E-11 |
| GO:001 4065 | phosphatidylinositol 3-kinase signaling | 50/38 05 | 3.19E-12 | 8.79E-11 |
| GO:000 2685 | regulation of leukocyte migration | 70/38 05 | 3.38E-12 | 9.26E-11 |
| GO:004 4772 | mitotic cell cycle phase transition | 108/3 805 | 3.46E-12 | 9.42E-11 |
| GO:000 7368 | determination of left/right symmetry | 49/38 05 | 3.47E-12 | 9.42E-11 |
| GO:000 9855 | determination of bilateral symmetry | 51/38 05 | 3.99E-12 | 1.08E-10 |
| GO:000 1738 | morphogenesis of a polarized epithelium | 38/38 05 | 4.25E-12 | 1.14E-10 |

| | | | | |
|---|---|---|---|---|
| GO:0001843 | neural tube closure | 44/3805 | 4.61E-12 | 1.24E-10 |
| GO:0051402 | neuron apoptotic process | 88/3805 | 5.15E-12 | 1.37E-10 |
| GO:0001952 | regulation of cell-matrix adhesion | 48/3805 | 5.19E-12 | 1.38E-10 |
| GO:0030099 | myeloid cell differentiation | 111/3805 | 5.49E-12 | 1.45E-10 |
| GO:0050771 | negative regulation of axonogenesis | 33/3805 | 5.64E-12 | 1.49E-10 |
| GO:0007052 | mitotic spindle organization | 46/3805 | 5.96E-12 | 1.56E-10 |
| GO:0060606 | tube closure | 44/3805 | 6.59E-12 | 1.72E-10 |
| GO:0022898 | regulation of transmembrane transporter activity | 81/3805 | 6.68E-12 | 1.74E-10 |
| GO:0006941 | striated muscle contraction | 58/3805 | 7.03E-12 | 1.82E-10 |
| GO:0008064 | regulation of actin polymerization or depolymerization | 56/3805 | 7.11E-12 | 1.83E-10 |
| GO:0031333 | negative regulation of protein-containing complex assembly | 53/3805 | 7.89E-12 | 2.02E-10 |
| GO:0072073 | kidney epithelium development | 53/3805 | 7.89E-12 | 2.02E-10 |
| GO:0007416 | synapse assembly | 62/3805 | 8.22E-12 | 2.09E-10 |
| GO:0009314 | response to radiation | 109/3805 | 8.39E-12 | 2.13E-10 |
| GO:0003341 | cilium movement | 67/3805 | 9.15E-12 | 2.31E-10 |
| GO:0072006 | nephron development | 54/3805 | 9.27E-12 | 2.33E-10 |
| GO:0043547 | positive regulation of GTPase activity | 75/3805 | 9.61E-12 | 2.41E-10 |
| GO:0070374 | positive regulation of ERK1 and ERK2 cascade | 69/3805 | 1.01E-11 | 2.51E-10 |

| | | | | |
|---|---|---|---|---|
| GO:0008589 | regulation of smoothened signaling pathway | 39/3805 | 1.01E-11 | 2.51E-10 |
| GO:0048660 | regulation of smooth muscle cell proliferation | 59/3805 | 1.02E-11 | 2.54E-10 |
| GO:0072659 | protein localization to plasma membrane | 84/3805 | 1.16E-11 | 2.87E-10 |
| GO:0090068 | positive regulation of cell cycle process | 77/3805 | 1.21E-11 | 2.97E-10 |
| GO:0014020 | primary neural tube formation | 45/3805 | 1.25E-11 | 3.05E-10 |
| GO:0098742 | cell-cell adhesion via plasma-membrane adhesion molecules | 66/3805 | 1.38E-11 | 3.36E-10 |
| GO:0110020 | regulation of actomyosin structure organization | 43/3805 | 1.39E-11 | 3.38E-10 |
| GO:0061387 | regulation of extent of cell growth | 47/3805 | 1.49E-11 | 3.60E-10 |
| GO:0001938 | positive regulation of endothelial cell proliferation | 40/3805 | 1.50E-11 | 3.62E-10 |
| GO:0032102 | negative regulation of response to external stimulus | 103/3805 | 1.60E-11 | 3.84E-10 |
| GO:0030832 | regulation of actin filament length | 56/3805 | 1.61E-11 | 3.84E-10 |
| GO:0014812 | muscle cell migration | 45/3805 | 1.74E-11 | 4.15E-10 |
| GO:0042073 | intraciliary transport | 26/3805 | 1.80E-11 | 4.28E-10 |
| GO:0006937 | regulation of muscle contraction | 55/3805 | 1.86E-11 | 4.39E-10 |
| GO:0060840 | artery development | 44/3805 | 1.86E-11 | 4.39E-10 |
| GO:0048813 | dendrite morphogenesis | 60/3805 | 1.87E-11 | 4.40E-10 |
| GO:0003018 | vascular process in circulatory system | 70/3805 | 2.02E-11 | 4.71E-10 |
| GO:0048675 | axon extension | 50/3805 | 2.02E-11 | 4.71E-10 |

| | | | | |
|---|---|---|---|---|
| GO:0090263 | positive regulation of canonical Wnt signaling pathway | 42/3805 | 2.06E-11 | 4.79E-10 |
| GO:0003231 | cardiac ventricle development | 52/3805 | 2.12E-11 | 4.89E-10 |
| GO:0071674 | mononuclear cell migration | 59/3805 | 2.20E-11 | 5.08E-10 |
| GO:0048659 | smooth muscle cell proliferation | 60/3805 | 2.40E-11 | 5.51E-10 |
| GO:0032872 | regulation of stress-activated MAPK cascade | 64/3805 | 2.48E-11 | 5.66E-10 |
| GO:0001841 | neural tube formation | 48/3805 | 2.52E-11 | 5.74E-10 |
| GO:0043588 | skin development | 87/3805 | 2.71E-11 | 6.15E-10 |
| GO:0030516 | regulation of axon extension | 43/3805 | 2.78E-11 | 6.28E-10 |
| GO:0003279 | cardiac septum development | 47/3805 | 2.79E-11 | 6.28E-10 |
| GO:0060491 | regulation of cell projection assembly | 64/3805 | 3.12E-11 | 7.01E-10 |
| GO:0097191 | extrinsic apoptotic signaling pathway | 71/3805 | 3.16E-11 | 7.06E-10 |
| GO:0002028 | regulation of sodium ion transport | 39/3805 | 3.21E-11 | 7.17E-10 |
| GO:0022612 | gland morphogenesis | 53/3805 | 3.23E-11 | 7.18E-10 |
| GO:0051091 | positive regulation of DNA-binding transcription factor activity | 77/3805 | 3.26E-11 | 7.22E-10 |
| GO:0055001 | muscle cell development | 68/3805 | 3.65E-11 | 8.05E-10 |
| GO:0050769 | positive regulation of neurogenesis | 82/3805 | 3.69E-11 | 8.11E-10 |
| GO:0043523 | regulation of neuron apoptotic process | 79/3805 | 3.88E-11 | 8.50E-10 |
| GO:0060348 | bone development | 74/3805 | 3.99E-11 | 8.71E-10 |

| | | | | |
|---|---|---|---|---|
| GO:0042391 | regulation of membrane potential | 114/3805 | 4.12E-11 | 8.97E-10 |
| GO:0010927 | cellular component assembly involved in morphogenesis | 45/3805 | 4.60E-11 | 9.98E-10 |
| GO:0070252 | actin-mediated cell contraction | 39/3805 | 4.66E-11 | 1.01E-09 |
| GO:0032412 | regulation of ion transmembrane transporter activity | 77/3805 | 4.80E-11 | 1.03E-09 |
| GO:0070302 | regulation of stress-activated protein kinase signaling cascade | 64/3805 | 4.92E-11 | 1.06E-09 |
| GO:0062197 | cellular response to chemical stress | 89/3805 | 5.06E-11 | 1.08E-09 |
| GO:0022411 | cellular component disassembly | 106/3805 | 5.30E-11 | 1.13E-09 |
| GO:0051656 | establishment of organelle localization | 106/3805 | 5.30E-11 | 1.13E-09 |
| GO:0002062 | chondrocyte differentiation | 43/3805 | 5.42E-11 | 1.15E-09 |
| GO:0048738 | cardiac muscle tissue development | 76/3805 | 5.87E-11 | 1.24E-09 |
| GO:0031334 | positive regulation of protein-containing complex assembly | 63/3805 | 5.90E-11 | 1.24E-09 |
| GO:0003015 | heart process | 72/3805 | 5.91E-11 | 1.24E-09 |
| GO:0048661 | positive regulation of smooth muscle cell proliferation | 41/3805 | 6.17E-11 | 1.29E-09 |
| GO:0030833 | regulation of actin filament polymerization | 50/3805 | 6.41E-11 | 1.33E-09 |
| GO:0018209 | peptidyl-serine modification | 94/3805 | 6.66E-11 | 1.38E-09 |
| GO:0048562 | embryonic organ morphogenesis | 88/3805 | 7.40E-11 | 1.53E-09 |
| GO:1903034 | regulation of response to wounding | 56/3805 | 7.59E-11 | 1.56E-09 |
| GO:0090101 | negative regulation of transmembrane receptor protein serine/threonine kinase signaling pathway | 46/3805 | 7.75E-11 | 1.59E-09 |

| | | | | |
|---|---|---|---|---|
| GO:2000181 | negative regulation of blood vessel morphogenesis | 42/3805 | 8.12E-11 | 1.66E-09 |
| GO:0051924 | regulation of calcium ion transport | 79/3805 | 8.19E-11 | 1.67E-09 |
| GO:0018105 | peptidyl-serine phosphorylation | 89/3805 | 8.40E-11 | 1.71E-09 |
| GO:0007044 | cell-substrate junction assembly | 38/3805 | 9.92E-11 | 2.01E-09 |
| GO:0051348 | negative regulation of transferase activity | 74/3805 | 1.06E-10 | 2.15E-09 |
| GO:0006814 | sodium ion transport | 71/3805 | 1.08E-10 | 2.17E-09 |
| GO:0002687 | positive regulation of leukocyte migration | 52/3805 | 1.09E-10 | 2.20E-09 |
| GO:1901343 | negative regulation of vasculature development | 42/3805 | 1.13E-10 | 2.26E-09 |
| GO:0010595 | positive regulation of endothelial cell migration | 41/3805 | 1.21E-10 | 2.42E-09 |
| GO:0032984 | protein-containing complex disassembly | 69/3805 | 1.31E-10 | 2.60E-09 |
| GO:1905475 | regulation of protein localization to membrane | 61/3805 | 1.33E-10 | 2.64E-09 |
| GO:0001570 | vasculogenesis | 38/3805 | 1.42E-10 | 2.81E-09 |
| GO:0032386 | regulation of intracellular transport | 88/3805 | 1.44E-10 | 2.85E-09 |
| GO:0031532 | actin cytoskeleton reorganization | 42/3805 | 1.56E-10 | 3.06E-09 |
| GO:0016525 | negative regulation of angiogenesis | 41/3805 | 1.69E-10 | 3.31E-09 |
| GO:0007612 | learning | 58/3805 | 1.81E-10 | 3.52E-09 |
| GO:0071772 | response to BMP | 58/3805 | 1.81E-10 | 3.52E-09 |
| GO:0071773 | cellular response to BMP stimulus | 58/3805 | 1.81E-10 | 3.52E-09 |

| | | | | |
|---|---|---|---|---|
| GO:0007528 | neuromuscular junction development | 28/3805 | 1.85E-10 | 3.58E-09 |
| GO:1901215 | negative regulation of neuron death | 73/3805 | 1.91E-10 | 3.69E-09 |
| GO:0150115 | cell-substrate junction organization | 39/3805 | 1.93E-10 | 3.71E-09 |
| GO:0021700 | developmental maturation | 91/3805 | 2.03E-10 | 3.88E-09 |
| GO:0120193 | tight junction organization | 37/3805 | 2.11E-10 | 4.03E-09 |
| GO:0001935 | endothelial cell proliferation | 57/3805 | 2.16E-10 | 4.11E-09 |
| GO:0022409 | positive regulation of cell-cell adhesion | 80/3805 | 2.34E-10 | 4.45E-09 |
| GO:2000027 | regulation of animal organ morphogenesis | 46/3805 | 2.51E-10 | 4.76E-09 |
| GO:0007219 | Notch signaling pathway | 57/3805 | 2.73E-10 | 5.15E-09 |
| GO:0030239 | myofibril assembly | 31/3805 | 2.80E-10 | 5.26E-09 |
| GO:0030512 | negative regulation of transforming growth factor beta receptor signaling pathway | 32/3805 | 2.99E-10 | 5.61E-09 |
| GO:0035023 | regulation of Rho protein signal transduction | 36/3805 | 3.12E-10 | 5.83E-09 |
| GO:0003002 | regionalization | 96/3805 | 3.28E-10 | 6.11E-09 |
| GO:1901990 | regulation of mitotic cell cycle phase transition | 82/3805 | 3.64E-10 | 6.77E-09 |
| GO:0030509 | BMP signaling pathway | 55/3805 | 3.87E-10 | 7.18E-09 |
| GO:0032233 | positive regulation of actin filament bundle assembly | 30/3805 | 3.96E-10 | 7.30E-09 |
| GO:0044458 | motile cilium assembly | 30/3805 | 3.96E-10 | 7.30E-09 |
| GO:0120032 | regulation of plasma membrane bounded cell projection assembly | 61/3805 | 4.02E-10 | 7.38E-09 |

| | | | | |
|---|---|---|---|---|
| GO:0035567 | non-canonical Wnt signaling pathway | 27/3805 | 4.11E-10 | 7.54E-09 |
| GO:0044344 | cellular response to fibroblast growth factor stimulus | 37/3805 | 4.28E-10 | 7.79E-09 |
| GO:0001936 | regulation of endothelial cell proliferation | 53/3805 | 4.28E-10 | 7.79E-09 |
| GO:0055002 | striated muscle cell development | 31/3805 | 4.29E-10 | 7.79E-09 |
| GO:0045930 | negative regulation of mitotic cell cycle | 66/3805 | 4.32E-10 | 7.82E-09 |
| GO:1902105 | regulation of leukocyte differentiation | 84/3805 | 4.65E-10 | 8.39E-09 |
| GO:0032272 | negative regulation of protein polymerization | 34/3805 | 4.66E-10 | 8.39E-09 |
| GO:1903522 | regulation of blood circulation | 73/3805 | 4.87E-10 | 8.73E-09 |
| GO:0045786 | negative regulation of cell cycle | 93/3805 | 5.32E-10 | 9.53E-09 |
| GO:0032835 | glomerulus development | 29/3805 | 5.58E-10 | 9.95E-09 |
| GO:0061640 | cytoskeleton-dependent cytokinesis | 36/3805 | 6.37E-10 | 1.13E-08 |
| GO:0030100 | regulation of endocytosis | 67/3805 | 6.40E-10 | 1.14E-08 |
| GO:0003014 | renal system process | 43/3805 | 6.49E-10 | 1.15E-08 |
| GO:0060048 | cardiac muscle contraction | 45/3805 | 6.69E-10 | 1.18E-08 |
| GO:0050919 | negative chemotaxis | 23/3805 | 6.79E-10 | 1.19E-08 |
| GO:0021537 | telencephalon development | 71/3805 | 7.33E-10 | 1.29E-08 |
| GO:0048771 | tissue remodeling | 60/3805 | 7.46E-10 | 1.30E-08 |
| GO:0060047 | heart contraction | 67/3805 | 7.77E-10 | 1.36E-08 |

| | | | | |
|---|---|---|---|---|
| GO:0071774 | response to fibroblast growth factor | 37/3805 | 8.48E-10 | 1.47E-08 |
| GO:0043409 | negative regulation of MAPK cascade | 57/3805 | 8.50E-10 | 1.47E-08 |
| GO:0031099 | regeneration | 50/3805 | 9.02E-10 | 1.56E-08 |
| GO:0031503 | protein-containing complex localization | 60/3805 | 9.22E-10 | 1.59E-08 |
| GO:0045927 | positive regulation of growth | 83/3805 | 9.34E-10 | 1.61E-08 |
| GO:0046785 | microtubule polymerization | 34/3805 | 9.80E-10 | 1.68E-08 |
| GO:0043524 | negative regulation of neuron apoptotic process | 57/3805 | 1.06E-09 | 1.81E-08 |
| GO:0014066 | regulation of phosphatidylinositol 3-kinase signaling | 38/3805 | 1.09E-09 | 1.86E-08 |
| GO:0090162 | establishment of epithelial cell polarity | 20/3805 | 1.09E-09 | 1.86E-08 |
| GO:0008406 | gonad development | 67/3805 | 1.14E-09 | 1.93E-08 |
| GO:0051492 | regulation of stress fiber assembly | 37/3805 | 1.18E-09 | 2.00E-08 |
| GO:0006874 | cellular calcium ion homeostasis | 115/3805 | 1.19E-09 | 2.01E-08 |
| GO:0007059 | chromosome segregation | 86/3805 | 1.27E-09 | 2.14E-08 |
| GO:0032874 | positive regulation of stress-activated MAPK cascade | 46/3805 | 1.31E-09 | 2.20E-08 |
| GO:0060996 | dendritic spine development | 44/3805 | 1.33E-09 | 2.23E-08 |
| GO:0043297 | apical junction assembly | 35/3805 | 1.34E-09 | 2.24E-08 |
| GO:0045664 | regulation of neuron differentiation | 67/3805 | 1.38E-09 | 2.29E-08 |
| GO:1903046 | meiotic cell cycle process | 63/3805 | 1.42E-09 | 2.36E-08 |

| | | | | |
|---|---|---|---|---|
| GO:0090288 | negative regulation of cellular response to growth factor stimulus | 38/3805 | 1.50E-09 | 2.49E-08 |
| GO:0071526 | semaphorin-plexin signaling pathway | 24/3805 | 1.63E-09 | 2.69E-08 |
| GO:0050731 | positive regulation of peptidyl-tyrosine phosphorylation | 57/3805 | 1.63E-09 | 2.69E-08 |
| GO:0014911 | positive regulation of smooth muscle cell migration | 28/3805 | 1.91E-09 | 3.13E-08 |
| GO:0045766 | positive regulation of angiogenesis | 55/3805 | 1.91E-09 | 3.13E-08 |
| GO:1904018 | positive regulation of vasculature development | 55/3805 | 1.91E-09 | 3.13E-08 |
| GO:0034599 | cellular response to oxidative stress | 73/3805 | 2.02E-09 | 3.29E-08 |
| GO:0022408 | negative regulation of cell-cell adhesion | 60/3805 | 2.11E-09 | 3.42E-08 |
| GO:0006979 | response to oxidative stress | 98/3805 | 2.11E-09 | 3.42E-08 |
| GO:0061035 | regulation of cartilage development | 30/3805 | 2.12E-09 | 3.43E-08 |
| GO:0051145 | smooth muscle cell differentiation | 32/3805 | 2.14E-09 | 3.45E-08 |
| GO:0070304 | positive regulation of stress-activated protein kinase signaling cascade | 46/3805 | 2.20E-09 | 3.55E-08 |
| GO:0032964 | collagen biosynthetic process | 26/3805 | 2.40E-09 | 3.85E-08 |
| GO:0045137 | development of primary sexual characteristics | 67/3805 | 2.40E-09 | 3.85E-08 |
| GO:0008544 | epidermis development | 89/3805 | 2.60E-09 | 4.15E-08 |
| GO:0050921 | positive regulation of chemotaxis | 48/3805 | 2.63E-09 | 4.19E-08 |
| GO:1904035 | regulation of epithelial cell apoptotic process | 35/3805 | 2.65E-09 | 4.21E-08 |
| GO:2001236 | regulation of extrinsic apoptotic signaling pathway | 52/3805 | 2.66E-09 | 4.21E-08 |

| | | | | |
|---|---|---|---|---|
| GO:0085029 | extracellular matrix assembly | 22/3805 | 2.72E-09 | 4.30E-08 |
| GO:0034332 | adherens junction organization | 24/3805 | 2.74E-09 | 4.33E-08 |
| GO:0010934 | macrophage cytokine production | 14/3805 | 2.90E-09 | 4.57E-08 |
| GO:0007613 | memory | 50/3805 | 3.01E-09 | 4.73E-08 |
| GO:0032963 | collagen metabolic process | 42/3805 | 3.05E-09 | 4.78E-08 |
| GO:0048844 | artery morphogenesis | 32/3805 | 3.09E-09 | 4.82E-08 |
| GO:0061512 | protein localization to cilium | 30/3805 | 3.15E-09 | 4.90E-08 |
| GO:0007019 | microtubule depolymerization | 25/3805 | 3.32E-09 | 5.15E-08 |
| GO:0098813 | nuclear chromosome segregation | 73/3805 | 3.36E-09 | 5.21E-08 |
| GO:0001666 | response to hypoxia | 61/3805 | 3.82E-09 | 5.91E-08 |
| GO:0051058 | negative regulation of small GTPase mediated signal transduction | 27/3805 | 4.19E-09 | 6.46E-08 |
| GO:0002573 | myeloid leukocyte differentiation | 66/3805 | 4.25E-09 | 6.54E-08 |
| GO:0009416 | response to light stimulus | 81/3805 | 4.28E-09 | 6.56E-08 |
| GO:0002274 | myeloid leukocyte activation | 71/3805 | 4.31E-09 | 6.60E-08 |
| GO:0048286 | lung alveolus development | 28/3805 | 4.45E-09 | 6.79E-08 |
| GO:0033673 | negative regulation of kinase activity | 63/3805 | 4.52E-09 | 6.89E-08 |
| GO:0045637 | regulation of myeloid cell differentiation | 62/3805 | 4.59E-09 | 6.97E-08 |
| GO:0014033 | neural crest cell differentiation | 36/3805 | 4.66E-09 | 7.07E-08 |

| | | | | |
|---|---|---|---|---|
| GO:0021591 | ventricular system development | 19/3805 | 4.77E-09 | 7.22E-08 |
| GO:0014031 | mesenchymal cell development | 35/3805 | 5.11E-09 | 7.70E-08 |
| GO:0010935 | regulation of macrophage cytokine production | 13/3805 | 5.16E-09 | 7.76E-08 |
| GO:0086005 | ventricular cardiac muscle cell action potential | 17/3805 | 5.48E-09 | 8.23E-08 |
| GO:0007159 | leukocyte cell-cell adhesion | 92/3805 | 5.67E-09 | 8.50E-08 |
| GO:0006469 | negative regulation of protein kinase activity | 58/3805 | 5.72E-09 | 8.53E-08 |
| GO:0030510 | regulation of BMP signaling pathway | 37/3805 | 5.72E-09 | 8.53E-08 |
| GO:0060411 | cardiac septum morphogenesis | 33/3805 | 5.94E-09 | 8.83E-08 |
| GO:0030517 | negative regulation of axon extension | 23/3805 | 6.12E-09 | 9.08E-08 |
| GO:0035725 | sodium ion transmembrane transport | 47/3805 | 6.44E-09 | 9.54E-08 |
| GO:0050878 | regulation of body fluid levels | 90/3805 | 6.56E-09 | 9.69E-08 |
| GO:0007422 | peripheral nervous system development | 30/3805 | 6.74E-09 | 9.90E-08 |
| GO:0014068 | positive regulation of phosphatidylinositol 3-kinase signaling | 30/3805 | 6.74E-09 | 9.90E-08 |
| GO:0090596 | sensory organ morphogenesis | 80/3805 | 7.17E-09 | 1.05E-07 |
| GO:0048863 | stem cell differentiation | 65/3805 | 7.49E-09 | 1.10E-07 |
| GO:0035050 | embryonic heart tube development | 34/3805 | 7.67E-09 | 1.12E-07 |
| GO:0006942 | regulation of striated muscle contraction | 33/3805 | 8.30E-09 | 1.21E-07 |
| GO:0035721 | intraciliary retrograde transport | Dec-05 | 8.55E-09 | 1.24E-07 |

| | | | | |
|---|---|---|---|---|
| GO:2001234 | negative regulation of apoptotic signaling pathway | 66/3805 | 8.67E-09 | 1.26E-07 |
| GO:0080164 | regulation of nitric oxide metabolic process | 30/3805 | 9.73E-09 | 1.41E-07 |
| GO:0003158 | endothelium development | 43/3805 | 9.84E-09 | 1.42E-07 |
| GO:0030837 | negative regulation of actin filament polymerization | 28/3805 | 9.93E-09 | 1.43E-07 |
| GO:0045428 | regulation of nitric oxide biosynthetic process | 29/3805 | 9.93E-09 | 1.43E-07 |
| GO:1903131 | mononuclear cell differentiation | 111/3805 | 1.02E-08 | 1.46E-07 |
| GO:0098739 | import across plasma membrane | 55/3805 | 1.03E-08 | 1.48E-07 |
| GO:0090175 | regulation of establishment of planar polarity | 21/3805 | 1.06E-08 | 1.52E-07 |
| GO:0031023 | microtubule organizing center organization | 45/3805 | 1.16E-08 | 1.64E-07 |
| GO:0010971 | positive regulation of G2/M transition of mitotic cell cycle | 17/3805 | 1.16E-08 | 1.65E-07 |
| GO:0008306 | associative learning | 40/3805 | 1.21E-08 | 1.70E-07 |
| GO:2000177 | regulation of neural precursor cell proliferation | 40/3805 | 1.21E-08 | 1.70E-07 |
| GO:0007229 | integrin-mediated signaling pathway | 35/3805 | 1.30E-08 | 1.84E-07 |
| GO:0010001 | glial cell differentiation | 68/3805 | 1.34E-08 | 1.89E-07 |
| GO:0043534 | blood vessel endothelial cell migration | 39/3805 | 1.39E-08 | 1.95E-07 |
| GO:0060976 | coronary vasculature development | 30/3805 | 1.39E-08 | 1.95E-07 |
| GO:0050727 | regulation of inflammatory response | 85/3805 | 1.43E-08 | 1.99E-07 |
| GO:0032103 | positive regulation of response to external stimulus | 104/3805 | 1.48E-08 | 2.06E-07 |

| | | | | |
|---|---|---|---|---|
| GO:1903555 | regulation of tumor necrosis factor superfamily cytokine production | 54/3805 | 1.54E-08 | 2.15E-07 |
| GO:1902414 | protein localization to cell junction | 40/3805 | 1.57E-08 | 2.18E-07 |
| GO:1904019 | epithelial cell apoptotic process | 40/3805 | 1.57E-08 | 2.18E-07 |
| GO:0010811 | positive regulation of cell-substrate adhesion | 44/3805 | 1.74E-08 | 2.41E-07 |
| GO:0006939 | smooth muscle contraction | 39/3805 | 1.82E-08 | 2.52E-07 |
| GO:0035107 | appendage morphogenesis | 51/3805 | 1.84E-08 | 2.52E-07 |
| GO:0035108 | limb morphogenesis | 51/3805 | 1.84E-08 | 2.52E-07 |
| GO:0007369 | gastrulation | 52/3805 | 1.86E-08 | 2.55E-07 |
| GO:0048705 | skeletal system morphogenesis | 68/3805 | 1.87E-08 | 2.56E-07 |
| GO:0001736 | establishment of planar polarity | 26/3805 | 2.11E-08 | 2.87E-07 |
| GO:0060688 | regulation of morphogenesis of a branching structure | 27/3805 | 2.14E-08 | 2.92E-07 |
| GO:0001656 | metanephros development | 33/3805 | 2.18E-08 | 2.96E-07 |
| GO:0007254 | JNK cascade | 52/3805 | 2.29E-08 | 3.10E-07 |
| GO:0042542 | response to hydrogen peroxide | 40/3805 | 2.64E-08 | 3.57E-07 |
| GO:2001237 | negative regulation of extrinsic apoptotic signaling pathway | 36/3805 | 2.78E-08 | 3.75E-07 |
| GO:2001222 | regulation of neuron migration | 24/3805 | 2.88E-08 | 3.87E-07 |
| GO:0001818 | negative regulation of cytokine production | 74/3805 | 2.97E-08 | 3.98E-07 |
| GO:1903828 | negative regulation of protein localization | 60/3805 | 3.01E-08 | 4.04E-07 |

| | | | | |
|---|---|---|---|---|
| GO:0031113 | regulation of microtubule polymerization | 25/3805 | 3.05E-08 | 4.06E-07 |
| GO:0048846 | axon extension involved in axon guidance | 19/3805 | 3.05E-08 | 4.06E-07 |
| GO:0060071 | Wnt signaling pathway, planar cell polarity pathway | 19/3805 | 3.05E-08 | 4.06E-07 |
| GO:1902284 | neuron projection extension involved in neuron projection guidance | 19/3805 | 3.05E-08 | 4.06E-07 |
| GO:0048146 | positive regulation of fibroblast proliferation | 28/3805 | 3.07E-08 | 4.07E-07 |
| GO:0046328 | regulation of JNK cascade | 46/3805 | 3.08E-08 | 4.07E-07 |
| GO:0043270 | positive regulation of ion transport | 80/3805 | 3.09E-08 | 4.09E-07 |
| GO:0061041 | regulation of wound healing | 42/3805 | 3.13E-08 | 4.12E-07 |
| GO:0007164 | establishment of tissue polarity | 26/3805 | 3.14E-08 | 4.12E-07 |
| GO:0030865 | cortical cytoskeleton organization | 26/3805 | 3.14E-08 | 4.12E-07 |
| GO:2001235 | positive regulation of apoptotic signaling pathway | 47/3805 | 3.18E-08 | 4.17E-07 |
| GO:0140013 | meiotic nuclear division | 56/3805 | 3.31E-08 | 4.32E-07 |
| GO:0008016 | regulation of heart contraction | 55/3805 | 3.36E-08 | 4.38E-07 |
| GO:0071214 | cellular response to abiotic stimulus | 80/3805 | 3.56E-08 | 4.63E-07 |
| GO:0104004 | cellular response to environmental stimulus | 80/3805 | 3.56E-08 | 4.63E-07 |
| GO:0032411 | positive regulation of transporter activity | 42/3805 | 3.98E-08 | 5.15E-07 |
| GO:0034767 | positive regulation of ion transmembrane transport | 54/3805 | 4.11E-08 | 5.32E-07 |
| GO:0046847 | filopodium assembly | 29/3805 | 4.17E-08 | 5.37E-07 |

| | | | | |
|---|---|---|---|---|
| GO:1905515 | non-motile cilium assembly | 29/3805 | 4.17E-08 | 5.37E-07 |
| GO:0007405 | neuroblast proliferation | 28/3805 | 4.40E-08 | 5.65E-07 |
| GO:0030834 | regulation of actin filament depolymerization | 24/3805 | 4.40E-08 | 5.65E-07 |
| GO:0014032 | neural crest cell development | 32/3805 | 4.48E-08 | 5.74E-07 |
| GO:0098657 | import into cell | 63/3805 | 4.53E-08 | 5.79E-07 |
| GO:0061082 | myeloid leukocyte cytokine production | 17/3805 | 4.56E-08 | 5.81E-07 |
| GO:0030042 | actin filament depolymerization | 25/3805 | 4.58E-08 | 5.81E-07 |
| GO:0035924 | cellular response to vascular endothelial growth factor stimulus | 25/3805 | 4.58E-08 | 5.81E-07 |
| GO:0071706 | tumor necrosis factor superfamily cytokine production | 54/3805 | 4.97E-08 | 6.30E-07 |
| GO:0030866 | cortical actin cytoskeleton organization | 19/3805 | 5.35E-08 | 6.75E-07 |
| GO:0031114 | regulation of microtubule depolymerization | 19/3805 | 5.35E-08 | 6.75E-07 |
| GO:0070588 | calcium ion transmembrane transport | 74/3805 | 5.37E-08 | 6.76E-07 |
| GO:0032273 | positive regulation of protein polymerization | 33/3805 | 5.42E-08 | 6.81E-07 |
| GO:0048144 | fibroblast proliferation | 37/3805 | 5.46E-08 | 6.85E-07 |
| GO:0048259 | regulation of receptor-mediated endocytosis | 40/3805 | 5.60E-08 | 7.01E-07 |
| GO:0031111 | negative regulation of microtubule polymerization or depolymerization | 22/3805 | 5.77E-08 | 7.20E-07 |
| GO:0045446 | endothelial cell differentiation | 38/3805 | 6.06E-08 | 7.53E-07 |
| GO:0072009 | nephron epithelium development | 38/3805 | 6.06E-08 | 7.53E-07 |

| | | | | |
|---|---|---|---|---|
| GO:0002088 | lens development in camera-type eye | 32/3805 | 6.08E-08 | 7.55E-07 |
| GO:0032680 | regulation of tumor necrosis factor production | 52/3805 | 6.16E-08 | 7.64E-07 |
| GO:0048678 | response to axon injury | 28/3805 | 6.24E-08 | 7.70E-07 |
| GO:1905330 | regulation of morphogenesis of an epithelium | 28/3805 | 6.24E-08 | 7.70E-07 |
| GO:0071677 | positive regulation of mononuclear cell migration | 27/3805 | 6.54E-08 | 8.06E-07 |
| GO:0034764 | positive regulation of transmembrane transport | 65/3805 | 6.73E-08 | 8.28E-07 |
| GO:0048041 | focal adhesion assembly | 31/3805 | 6.77E-08 | 8.30E-07 |
| GO:0045862 | positive regulation of proteolysis | 84/3805 | 6.82E-08 | 8.35E-07 |
| GO:0051147 | regulation of muscle cell differentiation | 45/3805 | 7.17E-08 | 8.77E-07 |
| GO:0048864 | stem cell development | 33/3805 | 7.25E-08 | 8.85E-07 |
| GO:0036293 | response to decreased oxygen levels | 63/3805 | 7.44E-08 | 9.05E-07 |
| GO:0048145 | regulation of fibroblast proliferation | 36/3805 | 8.27E-08 | 1.01E-06 |
| GO:0060326 | cell chemotaxis | 76/3805 | 8.29E-08 | 1.01E-06 |
| GO:1902751 | positive regulation of cell cycle G2/M phase transition | 17/3805 | 8.52E-08 | 1.03E-06 |
| GO:0045214 | sarcomere organization | 22/3805 | 9.01E-08 | 1.09E-06 |
| GO:0018210 | peptidyl-threonine modification | 40/3805 | 9.06E-08 | 1.09E-06 |
| GO:0045807 | positive regulation of endocytosis | 40/3805 | 9.06E-08 | 1.09E-06 |
| GO:0050863 | regulation of T cell activation | 82/3805 | 9.10E-08 | 1.09E-06 |

| | | | | |
|---|---|---|---|---|
| GO:0000910 | cytokinesis | 51/3805 | 9.14E-08 | 1.10E-06 |
| GO:0035088 | establishment or maintenance of apical/basal cell polarity | 23/3805 | 9.63E-08 | 1.15E-06 |
| GO:0061245 | establishment or maintenance of bipolar cell polarity | 23/3805 | 9.63E-08 | 1.15E-06 |
| GO:0006898 | receptor-mediated endocytosis | 67/3805 | 9.64E-08 | 1.15E-06 |
| GO:0098900 | regulation of action potential | 26/3805 | 9.72E-08 | 1.16E-06 |
| GO:0043491 | protein kinase B signaling | 56/3805 | 9.85E-08 | 1.17E-06 |
| GO:0051496 | positive regulation of stress fiber assembly | 24/3805 | 9.93E-08 | 1.18E-06 |
| GO:0051222 | positive regulation of protein transport | 77/3805 | 1.02E-07 | 1.20E-06 |
| GO:0001823 | mesonephros development | 36/3805 | 1.07E-07 | 1.27E-06 |
| GO:1903035 | negative regulation of response to wounding | 32/3805 | 1.10E-07 | 1.29E-06 |
| GO:0045880 | positive regulation of smoothened signaling pathway | 20/3805 | 1.12E-07 | 1.31E-06 |
| GO:0086002 | cardiac muscle cell action potential involved in contraction | 20/3805 | 1.12E-07 | 1.31E-06 |
| GO:0043583 | ear development | 64/3805 | 1.14E-07 | 1.34E-06 |
| GO:0007098 | centrosome cycle | 40/3805 | 1.15E-07 | 1.34E-06 |
| GO:0010948 | negative regulation of cell cycle process | 71/3805 | 1.17E-07 | 1.37E-06 |
| GO:0030307 | positive regulation of cell growth | 56/3805 | 1.17E-07 | 1.37E-06 |
| GO:0045931 | positive regulation of mitotic cell cycle | 41/3805 | 1.21E-07 | 1.40E-06 |
| GO:0048736 | appendage development | 55/3805 | 1.21E-07 | 1.41E-06 |

| | | | | |
|---|---|---|---|---|
| GO:0060173 | limb development | 55/3805 | 1.21E-07 | 1.41E-06 |
| GO:0003281 | ventricular septum development | 31/3805 | 1.24E-07 | 1.43E-06 |
| GO:0120192 | tight junction assembly | 31/3805 | 1.24E-07 | 1.43E-06 |
| GO:0010038 | response to metal ion | 72/3805 | 1.26E-07 | 1.46E-06 |
| GO:0018107 | peptidyl-threonine phosphorylation | 38/3805 | 1.28E-07 | 1.47E-06 |
| GO:0055117 | regulation of cardiac muscle contraction | 27/3805 | 1.31E-07 | 1.51E-06 |
| GO:0002040 | sprouting angiogenesis | 39/3805 | 1.37E-07 | 1.57E-06 |
| GO:1903169 | regulation of calcium ion transmembrane transport | 49/3805 | 1.37E-07 | 1.57E-06 |
| GO:0006809 | nitric oxide biosynthetic process | 30/3805 | 1.39E-07 | 1.58E-06 |
| GO:0000281 | mitotic cytokinesis | 26/3805 | 1.39E-07 | 1.58E-06 |
| GO:1902305 | regulation of sodium ion transmembrane transport | 26/3805 | 1.39E-07 | 1.58E-06 |
| GO:1901988 | negative regulation of cell cycle phase transition | 63/3805 | 1.41E-07 | 1.60E-06 |
| GO:0010712 | regulation of collagen metabolic process | 23/3805 | 1.45E-07 | 1.64E-06 |
| GO:0046580 | negative regulation of Ras protein signal transduction | 23/3805 | 1.45E-07 | 1.64E-06 |
| GO:0070301 | cellular response to hydrogen peroxide | 32/3805 | 1.46E-07 | 1.65E-06 |
| GO:1903532 | positive regulation of secretion by cell | 84/3805 | 1.47E-07 | 1.66E-06 |
| GO:0034763 | negative regulation of transmembrane transport | 42/3805 | 1.57E-07 | 1.77E-06 |
| GO:0001657 | ureteric bud development | 35/3805 | 1.63E-07 | 1.82E-06 |

| | | | | |
|---|---|---|---|---|
| GO:0072163 | mesonephric epithelium development | 35/3805 | 1.63E-07 | 1.82E-06 |
| GO:0072164 | mesonephric tubule development | 35/3805 | 1.63E-07 | 1.82E-06 |
| GO:0090100 | positive regulation of transmembrane receptor protein serine/threonine kinase signaling pathway | 38/3805 | 1.63E-07 | 1.82E-06 |
| GO:0019221 | cytokine-mediated signaling pathway | 94/3805 | 1.64E-07 | 1.83E-06 |
| GO:0043433 | negative regulation of DNA-binding transcription factor activity | 48/3805 | 1.68E-07 | 1.87E-06 |
| GO:0032414 | positive regulation of ion transmembrane transporter activity | 39/3805 | 1.73E-07 | 1.92E-06 |
| GO:1905477 | positive regulation of protein localization to membrane | 36/3805 | 1.79E-07 | 1.98E-06 |
| GO:0001649 | osteoblast differentiation | 58/3805 | 1.81E-07 | 2.00E-06 |
| GO:0000302 | response to reactive oxygen species | 52/3805 | 1.89E-07 | 2.09E-06 |
| GO:0032640 | tumor necrosis factor production | 52/3805 | 1.89E-07 | 2.09E-06 |
| GO:0019932 | second-messenger-mediated signaling | 74/3805 | 1.91E-07 | 2.11E-06 |
| GO:0038127 | ERBB signaling pathway | 37/3805 | 1.94E-07 | 2.12E-06 |
| GO:0044843 | cell cycle G1/S phase transition | 60/3805 | 1.94E-07 | 2.12E-06 |
| GO:0060041 | retina development in camera-type eye | 50/3805 | 1.98E-07 | 2.17E-06 |
| GO:0010718 | positive regulation of epithelial to mesenchymal transition | 21/3805 | 1.99E-07 | 2.17E-06 |
| GO:0001539 | cilium or flagellum-dependent cell motility | 48/3805 | 2.03E-07 | 2.22E-06 |
| GO:0060285 | cilium-dependent cell motility | 48/3805 | 2.03E-07 | 2.22E-06 |
| GO:1903037 | regulation of leukocyte cell-cell adhesion | 80/3805 | 2.04E-07 | 2.22E-06 |

| | | | | |
|---|---|---|---|---|
| GO:0070286 | axonemal dynein complex assembly | 18/3805 | 2.05E-07 | 2.22E-06 |
| GO:0070306 | lens fiber cell differentiation | 18/3805 | 2.05E-07 | 2.22E-06 |
| GO:0000070 | mitotic sister chromatid segregation | 46/3805 | 2.06E-07 | 2.22E-06 |
| GO:0021782 | glial cell development | 38/3805 | 2.07E-07 | 2.23E-06 |
| GO:0003208 | cardiac ventricle morphogenesis | 29/3805 | 2.10E-07 | 2.26E-06 |
| GO:0008543 | fibroblast growth factor receptor signaling pathway | 29/3805 | 2.10E-07 | 2.26E-06 |
| GO:0046209 | nitric oxide metabolic process | 31/3805 | 2.22E-07 | 2.38E-06 |
| GO:0048872 | homeostasis of number of cells | 85/3805 | 2.23E-07 | 2.39E-06 |
| GO:1903707 | negative regulation of hemopoiesis | 36/3805 | 2.29E-07 | 2.46E-06 |
| GO:0043271 | negative regulation of ion transport | 49/3805 | 2.43E-07 | 2.59E-06 |
| GO:0071604 | transforming growth factor beta production | 19/3805 | 2.49E-07 | 2.66E-06 |
| GO:0051092 | positive regulation of NF-kappaB transcription factor activity | 45/3805 | 2.51E-07 | 2.67E-06 |
| GO:0071695 | anatomical structure maturation | 72/3805 | 2.54E-07 | 2.71E-06 |
| GO:0010714 | positive regulation of collagen metabolic process | 17/3805 | 2.70E-07 | 2.87E-06 |
| GO:0003143 | embryonic heart tube morphogenesis | 26/3805 | 2.76E-07 | 2.93E-06 |
| GO:0006887 | exocytosis | 92/3805 | 2.76E-07 | 2.93E-06 |
| GO:0048008 | platelet-derived growth factor receptor signaling pathway | 25/3805 | 2.94E-07 | 3.11E-06 |
| GO:0050866 | negative regulation of cell activation | 58/3805 | 2.95E-07 | 3.12E-06 |

| | | | | |
|---|---|---|---|---|
| GO:0050680 | negative regulation of epithelial cell proliferation | 48/3805 | 2.98E-07 | 3.14E-06 |
| GO:0034614 | cellular response to reactive oxygen species | 42/3805 | 3.00E-07 | 3.14E-06 |
| GO:0051783 | regulation of nuclear division | 42/3805 | 3.00E-07 | 3.14E-06 |
| GO:0007009 | plasma membrane organization | 45/3805 | 3.06E-07 | 3.21E-06 |
| GO:0048639 | positive regulation of developmental growth | 57/3805 | 3.10E-07 | 3.25E-06 |
| GO:0045666 | positive regulation of neuron differentiation | 37/3805 | 3.12E-07 | 3.26E-06 |
| GO:2000351 | regulation of endothelial cell apoptotic process | 22/3805 | 3.15E-07 | 3.29E-06 |
| GO:0031644 | regulation of nervous system process | 52/3805 | 3.22E-07 | 3.35E-06 |
| GO:0006820 | anion transport | 106/3805 | 3.43E-07 | 3.57E-06 |
| GO:0032967 | positive regulation of collagen biosynthetic process | 16/3805 | 3.47E-07 | 3.58E-06 |
| GO:1901889 | negative regulation of cell junction assembly | 16/3805 | 3.47E-07 | 3.58E-06 |
| GO:0007088 | regulation of mitotic nuclear division | 35/3805 | 3.47E-07 | 3.58E-06 |
| GO:0034446 | substrate adhesion-dependent cell spreading | 35/3805 | 3.47E-07 | 3.58E-06 |
| GO:0045444 | fat cell differentiation | 65/3805 | 3.61E-07 | 3.72E-06 |
| GO:0070373 | negative regulation of ERK1 and ERK2 cascade | 29/3805 | 3.82E-07 | 3.93E-06 |
| GO:0035904 | aorta development | 26/3805 | 3.84E-07 | 3.95E-06 |
| GO:2001057 | reactive nitrogen species metabolic process | 31/3805 | 3.88E-07 | 3.98E-06 |
| GO:0000819 | sister chromatid segregation | 51/3805 | 3.97E-07 | 4.07E-06 |

| | | | | |
|---|---|---|---|---|
| GO:0038084 | vascular endothelial growth factor signaling pathway | 19/3805 | 3.98E-07 | 4.07E-06 |
| GO:1904951 | positive regulation of establishment of protein localization | 78/3805 | 4.00E-07 | 4.08E-06 |
| GO:0030098 | lymphocyte differentiation | 97/3805 | 4.25E-07 | 4.33E-06 |
| GO:0051057 | positive regulation of small GTPase mediated signal transduction | 28/3805 | 4.30E-07 | 4.38E-06 |
| GO:0050708 | regulation of protein secretion | 72/3805 | 4.37E-07 | 4.44E-06 |
| GO:1902106 | negative regulation of leukocyte differentiation | 35/3805 | 4.43E-07 | 4.49E-06 |
| GO:0003151 | outflow tract morphogenesis | 30/3805 | 4.46E-07 | 4.51E-06 |
| GO:0035265 | organ growth | 56/3805 | 4.51E-07 | 4.56E-06 |
| GO:0003401 | axis elongation | 17/3805 | 4.61E-07 | 4.64E-06 |
| GO:0007026 | negative regulation of microtubule depolymerization | 17/3805 | 4.61E-07 | 4.64E-06 |
| GO:0032330 | regulation of chondrocyte differentiation | 22/3805 | 4.65E-07 | 4.68E-06 |
| GO:0002683 | negative regulation of immune system process | 97/3805 | 4.73E-07 | 4.75E-06 |
| GO:0010657 | muscle cell apoptotic process | 33/3805 | 4.83E-07 | 4.84E-06 |
| GO:0002688 | regulation of leukocyte chemotaxis | 38/3805 | 5.17E-07 | 5.17E-06 |
| GO:0050818 | regulation of coagulation | 26/3805 | 5.30E-07 | 5.30E-06 |
| GO:0008584 | male gonad development | 41/3805 | 5.53E-07 | 5.50E-06 |
| GO:1904375 | regulation of protein localization to cell periphery | 41/3805 | 5.53E-07 | 5.50E-06 |
| GO:0007548 | sex differentiation | 74/3805 | 5.53E-07 | 5.50E-06 |

| | | | | |
|---|---|---|---|---|
| GO:0002931 | response to ischemia | 18/3805 | 5.56E-07 | 5.51E-06 |
| GO:1904645 | response to amyloid-beta | 18/3805 | 5.56E-07 | 5.51E-06 |
| GO:0051480 | regulation of cytosolic calcium ion concentration | 88/3805 | 5.56E-07 | 5.51E-06 |
| GO:0044839 | cell cycle G2/M phase transition | 43/3805 | 5.57E-07 | 5.51E-06 |
| GO:0030193 | regulation of blood coagulation | 25/3805 | 5.79E-07 | 5.71E-06 |
| GO:0033627 | cell adhesion mediated by integrin | 30/3805 | 5.88E-07 | 5.79E-06 |
| GO:0051150 | regulation of smooth muscle cell differentiation | 16/3805 | 6.08E-07 | 5.98E-06 |
| GO:0048839 | inner ear development | 56/3805 | 6.21E-07 | 6.10E-06 |
| GO:0045744 | negative regulation of G protein-coupled receptor signaling pathway | 24/3805 | 6.22E-07 | 6.10E-06 |
| GO:0051099 | positive regulation of binding | 51/3805 | 6.67E-07 | 6.53E-06 |
| GO:0030835 | negative regulation of actin filament depolymerization | 20/3805 | 6.68E-07 | 6.53E-06 |
| GO:0061326 | renal tubule development | 34/3805 | 6.70E-07 | 6.54E-06 |
| GO:0035637 | multicellular organismal signaling | 44/3805 | 6.73E-07 | 6.56E-06 |
| GO:0046546 | development of primary male sexual characteristics | 41/3805 | 6.79E-07 | 6.60E-06 |
| GO:0034113 | heterotypic cell-cell adhesion | 22/3805 | 6.79E-07 | 6.60E-06 |
| GO:0001508 | action potential | 42/3805 | 6.81E-07 | 6.60E-06 |
| GO:0010633 | negative regulation of epithelial cell migration | 26/3805 | 7.26E-07 | 7.02E-06 |
| GO:0046579 | positive regulation of Ras protein signal transduction | 26/3805 | 7.26E-07 | 7.02E-06 |

| | | | | |
|---|---|---|---|---|
| GO:0001890 | placenta development | 48/3805 | 7.42E-07 | 7.16E-06 |
| GO:0035296 | regulation of tube diameter | 47/3805 | 7.64E-07 | 7.35E-06 |
| GO:0097746 | blood vessel diameter maintenance | 47/3805 | 7.64E-07 | 7.35E-06 |
| GO:0007173 | epidermal growth factor receptor signaling pathway | 33/3805 | 7.95E-07 | 7.64E-06 |
| GO:0061337 | cardiac conduction | 25/3805 | 8.01E-07 | 7.68E-06 |
| GO:0006261 | DNA-templated DNA replication | 42/3805 | 8.30E-07 | 7.95E-06 |
| GO:0051346 | negative regulation of hydrolase activity | 82/3805 | 8.46E-07 | 8.09E-06 |
| GO:0071901 | negative regulation of protein serine/threonine kinase activity | 34/3805 | 8.51E-07 | 8.13E-06 |
| GO:0002697 | regulation of immune effector process | 94/3805 | 8.60E-07 | 8.20E-06 |
| GO:0051893 | regulation of focal adhesion assembly | 24/3805 | 8.71E-07 | 8.28E-06 |
| GO:0090109 | regulation of cell-substrate junction assembly | 24/3805 | 8.71E-07 | 8.28E-06 |
| GO:0061097 | regulation of protein tyrosine kinase activity | 27/3805 | 8.78E-07 | 8.33E-06 |
| GO:0045055 | regulated exocytosis | 68/3805 | 8.87E-07 | 8.39E-06 |
| GO:0045165 | cell fate commitment | 68/3805 | 8.87E-07 | 8.39E-06 |
| GO:0060415 | muscle tissue morphogenesis | 29/3805 | 8.93E-07 | 8.44E-06 |
| GO:0035150 | regulation of tube size | 47/3805 | 9.14E-07 | 8.62E-06 |
| GO:0050922 | negative regulation of chemotaxis | 23/3805 | 9.32E-07 | 8.78E-06 |
| GO:0010389 | regulation of G2/M transition of mitotic cell cycle | 32/3805 | 9.42E-07 | 8.86E-06 |

| | | | | |
|---|---|---|---|---|
| GO:0098840 | protein transport along microtubule | Oct-05 | 9.60E-07 | 8.99E-06 |
| GO:0099118 | microtubule-based protein transport | Oct-05 | 9.60E-07 | 8.99E-06 |
| GO:2000052 | positive regulation of non-canonical Wnt signaling pathway | Oct-05 | 9.60E-07 | 8.99E-06 |
| GO:0021955 | central nervous system neuron axonogenesis | 19/3805 | 9.64E-07 | 9.02E-06 |
| GO:0030858 | positive regulation of epithelial cell differentiation | 22/3805 | 9.79E-07 | 9.12E-06 |
| GO:0072577 | endothelial cell apoptotic process | 22/3805 | 9.79E-07 | 9.12E-06 |
| GO:2000649 | regulation of sodium ion transmembrane transporter activity | 22/3805 | 9.79E-07 | 9.12E-06 |
| GO:0000079 | regulation of cyclin-dependent protein serine/threonine kinase activity | 26/3805 | 9.87E-07 | 9.16E-06 |
| GO:0043954 | cellular component maintenance | 26/3805 | 9.87E-07 | 9.16E-06 |
| GO:0032965 | regulation of collagen biosynthetic process | 20/3805 | 1.00E-06 | 9.28E-06 |
| GO:0045197 | establishment or maintenance of epithelial cell apical/basal polarity | 20/3805 | 1.00E-06 | 9.28E-06 |
| GO:0030857 | negative regulation of epithelial cell differentiation | 21/3805 | 1.01E-06 | 9.29E-06 |
| GO:0055123 | digestive system development | 40/3805 | 1.01E-06 | 9.34E-06 |
| GO:0046330 | positive regulation of JNK cascade | 33/3805 | 1.01E-06 | 9.34E-06 |
| GO:0035051 | cardiocyte differentiation | 49/3805 | 1.02E-06 | 9.34E-06 |
| GO:0070830 | bicellular tight junction assembly | 28/3805 | 1.03E-06 | 9.44E-06 |
| GO:0048841 | regulation of axon extension involved in axon guidance | 16/3805 | 1.04E-06 | 9.49E-06 |
| GO:1904646 | cellular response to amyloid-beta | 16/3805 | 1.04E-06 | 9.49E-06 |

| | | | | |
|---|---|---|---|---|
| GO:1900046 | regulation of hemostasis | 25/3805 | 1.10E-06 | 1.00E-05 |
| GO:0035264 | multicellular organism growth | 56/3805 | 1.16E-06 | 1.06E-05 |
| GO:0032611 | interleukin-1 beta production | 32/3805 | 1.20E-06 | 1.10E-05 |
| GO:0090136 | epithelial cell-cell adhesion | Nov-05 | 1.20E-06 | 1.10E-05 |
| GO:0014902 | myotube differentiation | 40/3805 | 1.24E-06 | 1.13E-05 |
| GO:0035329 | hippo signaling | 17/3805 | 1.24E-06 | 1.13E-05 |
| GO:0086091 | regulation of heart rate by cardiac conduction | 17/3805 | 1.24E-06 | 1.13E-05 |
| GO:0097581 | lamellipodium organization | 30/3805 | 1.31E-06 | 1.18E-05 |
| GO:0002011 | morphogenesis of an epithelial sheet | 23/3805 | 1.31E-06 | 1.18E-05 |
| GO:1902749 | regulation of cell cycle G2/M phase transition | 34/3805 | 1.36E-06 | 1.22E-05 |
| GO:0051693 | actin filament capping | 18/3805 | 1.38E-06 | 1.25E-05 |
| GO:0098659 | inorganic cation import across plasma membrane | 35/3805 | 1.41E-06 | 1.27E-05 |
| GO:0099587 | inorganic ion import across plasma membrane | 35/3805 | 1.41E-06 | 1.27E-05 |
| GO:0032388 | positive regulation of intracellular transport | 49/3805 | 1.42E-06 | 1.28E-05 |
| GO:0030217 | T cell differentiation | 70/3805 | 1.45E-06 | 1.30E-05 |
| GO:0051963 | regulation of synapse assembly | 36/3805 | 1.46E-06 | 1.30E-05 |
| GO:0060412 | ventricular septum morphogenesis | 21/3805 | 1.46E-06 | 1.30E-05 |
| GO:0086003 | cardiac muscle cell contraction | 25/3805 | 1.50E-06 | 1.33E-05 |

| | | | | |
|---|---|---|---|---|
| GO:0071692 | protein localization to extracellular region | 88/3805 | 1.50E-06 | 1.34E-05 |
| GO:0000086 | G2/M transition of mitotic cell cycle | 39/3805 | 1.51E-06 | 1.35E-05 |
| GO:0042116 | macrophage activation | 32/3805 | 1.53E-06 | 1.36E-05 |
| GO:0072080 | nephron tubule development | 32/3805 | 1.53E-06 | 1.36E-05 |
| GO:0043535 | regulation of blood vessel endothelial cell migration | 30/3805 | 1.69E-06 | 1.49E-05 |
| GO:1903557 | positive regulation of tumor necrosis factor superfamily cytokine production | 34/3805 | 1.70E-06 | 1.51E-05 |
| GO:0048843 | negative regulation of axon extension involved in axon guidance | 14/3805 | 1.76E-06 | 1.56E-05 |
| GO:0072089 | stem cell proliferation | 28/3805 | 1.78E-06 | 1.57E-05 |
| GO:0051149 | positive regulation of muscle cell differentiation | 26/3805 | 1.78E-06 | 1.57E-05 |
| GO:0060433 | bronchus development | Sep-05 | 1.83E-06 | 1.61E-05 |
| GO:0009913 | epidermal cell differentiation | 58/3805 | 1.84E-06 | 1.61E-05 |
| GO:0043281 | regulation of cysteine-type endopeptidase activity involved in apoptotic process | 53/3805 | 1.87E-06 | 1.64E-05 |
| GO:0060993 | kidney morphogenesis | 32/3805 | 1.95E-06 | 1.71E-05 |
| GO:0031112 | positive regulation of microtubule polymerization or depolymerization | 17/3805 | 1.97E-06 | 1.72E-05 |
| GO:0060674 | placenta blood vessel development | 17/3805 | 1.97E-06 | 1.72E-05 |
| GO:0071634 | regulation of transforming growth factor beta production | 17/3805 | 1.97E-06 | 1.72E-05 |
| GO:0010761 | fibroblast migration | 22/3805 | 1.97E-06 | 1.72E-05 |
| GO:0060294 | cilium movement involved in cell motility | 44/3805 | 2.04E-06 | 1.78E-05 |

| | | | | |
|---|---|---|---|---|
| GO:0072676 | lymphocyte migration | 33/3805 | 2.05E-06 | 1.78E-05 |
| GO:0033157 | regulation of intracellular protein transport | 56/3805 | 2.10E-06 | 1.82E-05 |
| GO:0009914 | hormone transport | 83/3805 | 2.14E-06 | 1.85E-05 |
| GO:0021954 | central nervous system neuron development | 30/3805 | 2.16E-06 | 1.87E-05 |
| GO:0048644 | muscle organ morphogenesis | 30/3805 | 2.16E-06 | 1.87E-05 |
| GO:1903115 | regulation of actin filament-based movement | 19/3805 | 2.18E-06 | 1.88E-05 |
| GO:0034103 | regulation of tissue remodeling | 28/3805 | 2.32E-06 | 2.00E-05 |
| GO:0002690 | positive regulation of leukocyte chemotaxis | 31/3805 | 2.32E-06 | 2.00E-05 |
| GO:0042634 | regulation of hair cycle | 15/3805 | 2.33E-06 | 2.00E-05 |
| GO:0071248 | cellular response to metal ion | 45/3805 | 2.35E-06 | 2.02E-05 |
| GO:1904029 | regulation of cyclin-dependent protein kinase activity | 26/3805 | 2.37E-06 | 2.03E-05 |
| GO:0030595 | leukocyte chemotaxis | 56/3805 | 2.43E-06 | 2.08E-05 |
| GO:0045619 | regulation of lymphocyte differentiation | 51/3805 | 2.46E-06 | 2.10E-05 |
| GO:0001755 | neural crest cell migration | 23/3805 | 2.51E-06 | 2.14E-05 |
| GO:0010658 | striated muscle cell apoptotic process | 23/3805 | 2.51E-06 | 2.14E-05 |
| GO:2000401 | regulation of lymphocyte migration | 23/3805 | 2.51E-06 | 2.14E-05 |
| GO:0002065 | columnar/cuboidal epithelial cell differentiation | 33/3805 | 2.57E-06 | 2.18E-05 |
| GO:0106027 | neuron projection organization | 33/3805 | 2.57E-06 | 2.18E-05 |

| | | | | |
|---|---|---|---|---|
| GO:0016311 | dephosphorylation | 77/3805 | 2.63E-06 | 2.23E-05 |
| GO:0061430 | bone trabecula morphogenesis | Nov-05 | 2.73E-06 | 2.31E-05 |
| GO:0002224 | toll-like receptor signaling pathway | 36/3805 | 2.75E-06 | 2.32E-05 |
| GO:1901890 | positive regulation of cell junction assembly | 36/3805 | 2.75E-06 | 2.32E-05 |
| GO:0032651 | regulation of interleukin-1 beta production | 30/3805 | 2.77E-06 | 2.33E-05 |
| GO:0014044 | Schwann cell development | 16/3805 | 2.77E-06 | 2.33E-05 |
| GO:1904064 | positive regulation of cation transmembrane transport | 45/3805 | 2.79E-06 | 2.35E-05 |
| GO:1901653 | cellular response to peptide | 71/3805 | 2.80E-06 | 2.35E-05 |
| GO:0007249 | I-kappaB kinase/NF-kappaB signaling | 56/3805 | 2.81E-06 | 2.36E-05 |
| GO:0048592 | eye morphogenesis | 48/3805 | 2.88E-06 | 2.41E-05 |
| GO:0048593 | camera-type eye morphogenesis | 42/3805 | 3.10E-06 | 2.59E-05 |
| GO:0061484 | hematopoietic stem cell homeostasis | 14/3805 | 3.10E-06 | 2.59E-05 |
| GO:0045332 | phospholipid translocation | 20/3805 | 3.12E-06 | 2.60E-05 |
| GO:0002090 | regulation of receptor internalization | 26/3805 | 3.13E-06 | 2.61E-05 |
| GO:0031214 | biomineral tissue development | 46/3805 | 3.17E-06 | 2.64E-05 |
| GO:0034331 | cell junction maintenance | 18/3805 | 3.20E-06 | 2.66E-05 |
| GO:0032760 | positive regulation of tumor necrosis factor production | 33/3805 | 3.21E-06 | 2.66E-05 |
| GO:0052547 | regulation of peptidase activity | 93/3805 | 3.22E-06 | 2.67E-05 |

| | | | | |
|---|---|---|---|---|
| GO:190 2746 | regulation of lens fiber cell differentiation | Aug-05 | 3.22E-06 | 2.67E-05 |
| GO:000 2367 | cytokine production involved in immune response | 34/38 05 | 3.29E-06 | 2.72E-05 |
| GO:190 4705 | regulation of vascular associated smooth muscle cell proliferation | 23/38 05 | 3.44E-06 | 2.84E-05 |
| GO:190 1652 | response to peptide | 89/38 05 | 3.49E-06 | 2.88E-05 |
| GO:003 8034 | signal transduction in absence of ligand | 27/38 05 | 3.53E-06 | 2.90E-05 |
| GO:009 7192 | extrinsic apoptotic signaling pathway in absence of ligand | 27/38 05 | 3.53E-06 | 2.90E-05 |
| GO:000 6836 | neurotransmitter transport | 58/38 05 | 3.71E-06 | 3.05E-05 |
| GO:009 7061 | dendritic spine organization | 31/38 05 | 3.71E-06 | 3.05E-05 |
| GO:001 0659 | cardiac muscle cell apoptotic process | 22/38 05 | 3.80E-06 | 3.11E-05 |
| GO:001 4009 | glial cell proliferation | 22/38 05 | 3.80E-06 | 3.11E-05 |
| GO:003 4612 | response to tumor necrosis factor | 51/38 05 | 3.89E-06 | 3.18E-05 |
| GO:000 8277 | regulation of G protein-coupled receptor signaling pathway | 38/38 05 | 4.07E-06 | 3.31E-05 |
| GO:003 0178 | negative regulation of Wnt signaling pathway | 44/38 05 | 4.09E-06 | 3.33E-05 |
| GO:200 0379 | positive regulation of reactive oxygen species metabolic process | 26/38 05 | 4.11E-06 | 3.33E-05 |
| GO:190 1989 | positive regulation of cell cycle phase transition | 35/38 05 | 4.11E-06 | 3.33E-05 |
| GO:015 0116 | regulation of cell-substrate junction organization | 24/38 05 | 4.12E-06 | 3.33E-05 |
| GO:190 4377 | positive regulation of protein localization to cell periphery | 24/38 05 | 4.12E-06 | 3.33E-05 |
| GO:004 8565 | digestive tract development | 36/38 05 | 4.12E-06 | 3.34E-05 |

| | | | | |
|---|---|---|---|---|
| GO:0001505 | regulation of neurotransmitter levels | 60/3805 | 4.18E-06 | 3.37E-05 |
| GO:0099003 | vesicle-mediated transport in synapse | 60/3805 | 4.18E-06 | 3.37E-05 |
| GO:0010719 | negative regulation of epithelial to mesenchymal transition | 16/3805 | 4.38E-06 | 3.53E-05 |
| GO:0009612 | response to mechanical stimulus | 46/3805 | 4.41E-06 | 3.55E-05 |
| GO:0061005 | cell differentiation involved in kidney development | 20/3805 | 4.45E-06 | 3.57E-05 |
| GO:0002718 | regulation of cytokine production involved in immune response | 30/3805 | 4.46E-06 | 3.58E-05 |
| GO:0006260 | DNA replication | 64/3805 | 4.47E-06 | 3.58E-05 |
| GO:0009306 | protein secretion | 85/3805 | 4.60E-06 | 3.68E-05 |
| GO:0052548 | regulation of endopeptidase activity | 78/3805 | 4.69E-06 | 3.75E-05 |
| GO:0051047 | positive regulation of secretion | 88/3805 | 4.69E-06 | 3.75E-05 |
| GO:0030278 | regulation of ossification | 38/3805 | 4.92E-06 | 3.92E-05 |
| GO:0061008 | hepaticobiliary system development | 37/3805 | 4.99E-06 | 3.97E-05 |
| GO:0032612 | interleukin-1 production | 35/3805 | 5.04E-06 | 4.01E-05 |
| GO:0035592 | establishment of protein localization to extracellular region | 85/3805 | 5.10E-06 | 4.05E-05 |
| GO:0006949 | syncytium formation | 24/3805 | 5.48E-06 | 4.35E-05 |
| GO:0032091 | negative regulation of protein binding | 30/3805 | 5.62E-06 | 4.45E-05 |
| GO:0007269 | neurotransmitter secretion | 47/3805 | 5.75E-06 | 4.52E-05 |
| GO:0071356 | cellular response to tumor necrosis factor | 47/3805 | 5.75E-06 | 4.52E-05 |

| | | | | |
|---|---|---|---|---|
| GO:0099643 | signal release from synapse | 47/3805 | 5.75E-06 | 4.52E-05 |
| GO:0110148 | biomineralization | 47/3805 | 5.75E-06 | 4.52E-05 |
| GO:0048538 | thymus development | 21/3805 | 5.76E-06 | 4.52E-05 |
| GO:0061383 | trabecula morphogenesis | 21/3805 | 5.76E-06 | 4.52E-05 |
| GO:0086001 | cardiac muscle cell action potential | 21/3805 | 5.76E-06 | 4.52E-05 |
| GO:0030326 | embryonic limb morphogenesis | 39/3805 | 5.79E-06 | 4.54E-05 |
| GO:0035113 | embryonic appendage morphogenesis | 39/3805 | 5.79E-06 | 4.54E-05 |
| GO:0099565 | chemical synaptic transmission, postsynaptic | 31/3805 | 5.84E-06 | 4.57E-05 |
| GO:0008088 | axo-dendritic transport | 27/3805 | 5.90E-06 | 4.60E-05 |
| GO:0034109 | homotypic cell-cell adhesion | 27/3805 | 5.90E-06 | 4.60E-05 |
| GO:0046879 | hormone secretion | 80/3805 | 5.98E-06 | 4.66E-05 |
| GO:2000116 | regulation of cysteine-type endopeptidase activity | 57/3805 | 6.02E-06 | 4.69E-05 |
| GO:0097193 | intrinsic apoptotic signaling pathway | 69/3805 | 6.02E-06 | 4.69E-05 |
| GO:0021953 | central nervous system neuron differentiation | 51/3805 | 6.08E-06 | 4.73E-05 |
| GO:0008585 | female gonad development | 34/3805 | 6.16E-06 | 4.78E-05 |
| GO:0002063 | chondrocyte development | 15/3805 | 6.20E-06 | 4.80E-05 |
| GO:0006270 | DNA replication initiation | 15/3805 | 6.20E-06 | 4.80E-05 |
| GO:0033260 | nuclear DNA replication | 15/3805 | 6.20E-06 | 4.80E-05 |

| | | | | |
|---|---|---|---|---|
| GO:0007492 | endoderm development | 25/3805 | 6.26E-06 | 4.82E-05 |
| GO:0034204 | lipid translocation | 20/3805 | 6.26E-06 | 4.82E-05 |
| GO:1904036 | negative regulation of epithelial cell apoptotic process | 20/3805 | 6.26E-06 | 4.82E-05 |
| GO:0030203 | glycosaminoglycan metabolic process | 29/3805 | 6.74E-06 | 5.18E-05 |
| GO:0051899 | membrane depolarization | 29/3805 | 6.74E-06 | 5.18E-05 |
| GO:0035909 | aorta morphogenesis | 16/3805 | 6.77E-06 | 5.20E-05 |
| GO:1902692 | regulation of neuroblast proliferation | 17/3805 | 7.00E-06 | 5.36E-05 |
| GO:0097529 | myeloid leukocyte migration | 55/3805 | 7.00E-06 | 5.36E-05 |
| GO:1901992 | positive regulation of mitotic cell cycle phase transition | 30/3805 | 7.05E-06 | 5.39E-05 |
| GO:0001947 | heart looping | 22/3805 | 7.08E-06 | 5.40E-05 |
| GO:2001257 | regulation of cation channel activity | 46/3805 | 7.11E-06 | 5.43E-05 |
| GO:0070661 | leukocyte proliferation | 79/3805 | 7.29E-06 | 5.56E-05 |
| GO:0001889 | liver development | 36/3805 | 7.42E-06 | 5.64E-05 |
| GO:0042310 | vasoconstriction | 32/3805 | 7.43E-06 | 5.64E-05 |
| GO:0051896 | regulation of protein kinase B signaling | 45/3805 | 7.52E-06 | 5.70E-05 |
| GO:0030316 | osteoclast differentiation | 34/3805 | 7.53E-06 | 5.71E-05 |
| GO:1901991 | negative regulation of mitotic cell cycle phase transition | 44/3805 | 7.92E-06 | 6.00E-05 |
| GO:0050905 | neuromuscular process | 49/3805 | 8.04E-06 | 6.08E-05 |

| | | | | |
|---|---|---|---|---|
| GO:1903039 | positive regulation of leukocyte cell-cell adhesion | 58/3805 | 8.27E-06 | 6.25E-05 |
| GO:0051321 | meiotic cell cycle | 73/3805 | 8.29E-06 | 6.25E-05 |
| GO:0150076 | neuroinflammatory response | 23/3805 | 8.35E-06 | 6.28E-05 |
| GO:1990874 | vascular associated smooth muscle cell proliferation | 23/3805 | 8.35E-06 | 6.28E-05 |
| GO:0002700 | regulation of production of molecular mediator of immune response | 45/3805 | 8.80E-06 | 6.62E-05 |
| GO:0016202 | regulation of striated muscle tissue development | Dec-05 | 9.06E-06 | 6.80E-05 |
| GO:0006022 | aminoglycan metabolic process | 32/3805 | 9.16E-06 | 6.87E-05 |
| GO:0120034 | positive regulation of plasma membrane bounded cell projection assembly | 33/3805 | 9.21E-06 | 6.90E-05 |
| GO:0007596 | blood coagulation | 44/3805 | 9.30E-06 | 6.95E-05 |
| GO:0046661 | male sex differentiation | 44/3805 | 9.30E-06 | 6.95E-05 |
| GO:0072678 | T cell migration | 22/3805 | 9.52E-06 | 7.11E-05 |
| GO:0031116 | positive regulation of microtubule polymerization | 15/3805 | 9.75E-06 | 7.26E-05 |
| GO:0090504 | epiboly | 15/3805 | 9.75E-06 | 7.26E-05 |
| GO:0045580 | regulation of T cell differentiation | 43/3805 | 9.80E-06 | 7.29E-05 |
| GO:0006858 | extracellular transport | 18/3805 | 1.00E-05 | 7.42E-05 |
| GO:0031952 | regulation of protein autophosphorylation | 18/3805 | 1.00E-05 | 7.42E-05 |
| GO:0001953 | negative regulation of cell-matrix adhesion | 16/3805 | 1.03E-05 | 7.60E-05 |
| GO:0044786 | cell cycle DNA replication | 16/3805 | 1.03E-05 | 7.60E-05 |

| | | | | |
|---|---|---|---|---|
| GO:0030902 | hindbrain development | 45/3805 | 1.03E-05 | 7.60E-05 |
| GO:0014037 | Schwann cell differentiation | 17/3805 | 1.03E-05 | 7.60E-05 |
| GO:2000249 | regulation of actin cytoskeleton reorganization | 17/3805 | 1.03E-05 | 7.60E-05 |
| GO:0061982 | meiosis I cell cycle process | 37/3805 | 1.05E-05 | 7.76E-05 |
| GO:0043113 | receptor clustering | 25/3805 | 1.06E-05 | 7.78E-05 |
| GO:0007204 | positive regulation of cytosolic calcium ion concentration | 76/3805 | 1.07E-05 | 7.88E-05 |
| GO:0050772 | positive regulation of axonogenesis | 30/3805 | 1.10E-05 | 8.03E-05 |
| GO:0090630 | activation of GTPase activity | 30/3805 | 1.10E-05 | 8.03E-05 |
| GO:0000768 | syncytium formation by plasma membrane fusion | 23/3805 | 1.10E-05 | 8.07E-05 |
| GO:0061371 | determination of heart left/right asymmetry | 23/3805 | 1.10E-05 | 8.07E-05 |
| GO:0140253 | cell-cell fusion | 23/3805 | 1.10E-05 | 8.07E-05 |
| GO:0010763 | positive regulation of fibroblast migration | Nov-05 | 1.11E-05 | 8.10E-05 |
| GO:0045019 | negative regulation of nitric oxide biosynthetic process | Nov-05 | 1.11E-05 | 8.10E-05 |
| GO:0072010 | glomerular epithelium development | Nov-05 | 1.11E-05 | 8.10E-05 |
| GO:1904406 | negative regulation of nitric oxide metabolic process | Nov-05 | 1.11E-05 | 8.10E-05 |
| GO:0046545 | development of primary female sexual characteristics | 34/3805 | 1.12E-05 | 8.12E-05 |
| GO:0032652 | regulation of interleukin-1 production | 33/3805 | 1.13E-05 | 8.16E-05 |
| GO:0007274 | neuromuscular synaptic transmission | 13/3805 | 1.19E-05 | 8.61E-05 |

| | | | | |
|---|---|---|---|---|
| GO:0031954 | positive regulation of protein autophosphorylation | 13/3805 | 1.19E-05 | 8.61E-05 |
| GO:0008542 | visual learning | 24/3805 | 1.24E-05 | 8.96E-05 |
| GO:0043506 | regulation of JUN kinase activity | 24/3805 | 1.24E-05 | 8.96E-05 |
| GO:0061045 | negative regulation of wound healing | 24/3805 | 1.24E-05 | 8.96E-05 |
| GO:0031623 | receptor internalization | 37/3805 | 1.26E-05 | 9.06E-05 |
| GO:0042303 | molting cycle | 37/3805 | 1.26E-05 | 9.06E-05 |
| GO:0042633 | hair cycle | 37/3805 | 1.26E-05 | 9.06E-05 |
| GO:0050773 | regulation of dendrite development | 37/3805 | 1.26E-05 | 9.06E-05 |
| GO:0008347 | glial cell migration | 22/3805 | 1.27E-05 | 9.11E-05 |
| GO:0003184 | pulmonary valve morphogenesis | Oct-05 | 1.27E-05 | 9.12E-05 |
| GO:2001044 | regulation of integrin-mediated signaling pathway | Oct-05 | 1.27E-05 | 9.12E-05 |
| GO:0010457 | centriole-centriole cohesion | Sep-05 | 1.29E-05 | 9.21E-05 |
| GO:0048755 | branching morphogenesis of a nerve | Sep-05 | 1.29E-05 | 9.21E-05 |
| GO:1902969 | mitotic DNA replication | Sep-05 | 1.29E-05 | 9.21E-05 |
| GO:0051293 | establishment of spindle localization | 19/3805 | 1.32E-05 | 9.39E-05 |
| GO:0010660 | regulation of muscle cell apoptotic process | 29/3805 | 1.33E-05 | 9.45E-05 |
| GO:0007292 | female gamete generation | 43/3805 | 1.35E-05 | 9.57E-05 |
| GO:0051100 | negative regulation of binding | 43/3805 | 1.35E-05 | 9.57E-05 |

| | | | | |
|---|---|---|---|---|
| GO:0010769 | regulation of cell morphogenesis involved in differentiation | 34/3805 | 1.35E-05 | 9.60E-05 |
| GO:0043502 | regulation of muscle adaptation | 30/3805 | 1.36E-05 | 9.60E-05 |
| GO:0060395 | SMAD protein signal transduction | 30/3805 | 1.36E-05 | 9.60E-05 |
| GO:0006310 | DNA recombination | 70/3805 | 1.36E-05 | 9.61E-05 |
| GO:0001892 | embryonic placenta development | 33/3805 | 1.37E-05 | 9.66E-05 |
| GO:0051225 | spindle assembly | 33/3805 | 1.37E-05 | 9.66E-05 |
| GO:0007156 | homophilic cell adhesion via plasma membrane adhesion molecules | 32/3805 | 1.38E-05 | 9.70E-05 |
| GO:0001945 | lymph vessel development | 14/3805 | 1.39E-05 | 9.71E-05 |
| GO:0021952 | central nervous system projection neuron axonogenesis | 14/3805 | 1.39E-05 | 9.71E-05 |
| GO:0061036 | positive regulation of cartilage development | 14/3805 | 1.39E-05 | 9.71E-05 |
| GO:0061037 | negative regulation of cartilage development | 14/3805 | 1.39E-05 | 9.71E-05 |
| GO:0086010 | membrane depolarization during action potential | 14/3805 | 1.39E-05 | 9.71E-05 |
| GO:2000352 | negative regulation of endothelial cell apoptotic process | 14/3805 | 1.39E-05 | 9.71E-05 |
| GO:1902115 | regulation of organelle assembly | 51/3805 | 1.42E-05 | 9.96E-05 |
| GO:0060443 | mammary gland morphogenesis | 21/3805 | 1.45E-05 | 0.000101053 |
| GO:0098815 | modulation of excitatory postsynaptic potential | 21/3805 | 1.45E-05 | 0.000101053 |
| GO:0007599 | hemostasis | 44/3805 | 1.49E-05 | 0.000103396 |
| GO:0050817 | coagulation | 44/3805 | 1.49E-05 | 0.000103396 |

| | | | | |
|---|---|---|---|---|
| GO:0003351 | epithelial cilium movement involved in extracellular fluid movement | 17/3805 | 1.50E-05 | 0.000103841 |
| GO:0010765 | positive regulation of sodium ion transport | 17/3805 | 1.50E-05 | 0.000103841 |
| GO:0042176 | regulation of protein catabolic process | 82/3805 | 1.50E-05 | 0.000104314 |
| GO:0002761 | regulation of myeloid leukocyte differentiation | 37/3805 | 1.51E-05 | 0.000104364 |
| GO:0010842 | retina layer formation | Dec-05 | 1.59E-05 | 0.000110103 |
| GO:0099625 | ventricular cardiac muscle cell membrane repolarization | Dec-05 | 1.59E-05 | 0.000110103 |
| GO:0045732 | positive regulation of protein catabolic process | 53/3805 | 1.60E-05 | 0.000110449 |
| GO:0140029 | exocytic process | 28/3805 | 1.61E-05 | 0.00011073 |
| GO:0001885 | endothelial cell development | 24/3805 | 1.61E-05 | 0.00011073 |
| GO:1903510 | mucopolysaccharide metabolic process | 24/3805 | 1.61E-05 | 0.00011073 |
| GO:0035019 | somatic stem cell population maintenance | 20/3805 | 1.64E-05 | 0.000112258 |
| GO:0051653 | spindle localization | 20/3805 | 1.64E-05 | 0.000112258 |
| GO:0030317 | flagellated sperm motility | 39/3805 | 1.65E-05 | 0.000113273 |
| GO:0051897 | positive regulation of protein kinase B signaling | 33/3805 | 1.66E-05 | 0.000113958 |
| GO:0045581 | negative regulation of T cell differentiation | 19/3805 | 1.82E-05 | 0.000124435 |
| GO:0061028 | establishment of endothelial barrier | 19/3805 | 1.82E-05 | 0.000124435 |
| GO:1902806 | regulation of cell cycle G1/S phase transition | 43/3805 | 1.84E-05 | 0.000125646 |
| GO:0042246 | tissue regeneration | 23/3805 | 1.89E-05 | 0.00012903 |

| | | | | |
|---|---|---|---|---|
| GO:0043500 | muscle adaptation | 35/3805 | 1.92E-05 | 0.000130774 |
| GO:0045620 | negative regulation of lymphocyte differentiation | 21/3805 | 1.94E-05 | 0.00013179 |
| GO:0061333 | renal tubule morphogenesis | 27/3805 | 1.94E-05 | 0.00013179 |
| GO:1901019 | regulation of calcium ion transmembrane transporter activity | 27/3805 | 1.94E-05 | 0.00013179 |
| GO:0032675 | regulation of interleukin-6 production | 42/3805 | 1.95E-05 | 0.000132426 |
| GO:0046620 | regulation of organ growth | 34/3805 | 1.97E-05 | 0.000133556 |
| GO:0001941 | postsynaptic membrane organization | 18/3805 | 1.99E-05 | 0.00013466 |
| GO:0003179 | heart valve morphogenesis | 18/3805 | 1.99E-05 | 0.00013466 |
| GO:0048010 | vascular endothelial growth factor receptor signaling pathway | 18/3805 | 1.99E-05 | 0.00013466 |
| GO:0006970 | response to osmotic stress | 28/3805 | 2.00E-05 | 0.000135082 |
| GO:0002221 | pattern recognition receptor signaling pathway | 44/3805 | 2.01E-05 | 0.000135511 |
| GO:0050764 | regulation of phagocytosis | 33/3805 | 2.01E-05 | 0.000135511 |
| GO:0010770 | positive regulation of cell morphogenesis involved in differentiation | 30/3805 | 2.06E-05 | 0.000138097 |
| GO:0046849 | bone remodeling | 30/3805 | 2.06E-05 | 0.000138097 |
| GO:0051153 | regulation of striated muscle cell differentiation | 30/3805 | 2.06E-05 | 0.000138097 |
| GO:0051797 | regulation of hair follicle development | Nov-05 | 2.06E-05 | 0.000138097 |
| GO:0002263 | cell activation involved in immune response | 71/3805 | 2.12E-05 | 0.000141635 |
| GO:0010464 | regulation of mesenchymal cell proliferation | 17/3805 | 2.14E-05 | 0.000142911 |

| | | | | |
|---|---|---|---|---|
| GO:0061098 | positive regulation of protein tyrosine kinase activity | 17/3805 | 2.14E-05 | 0.000142911 |
| GO:0022011 | myelination in peripheral nervous system | 14/3805 | 2.17E-05 | 0.00014426 |
| GO:0032292 | peripheral nervous system axon ensheathment | 14/3805 | 2.17E-05 | 0.00014426 |
| GO:0034260 | negative regulation of GTPase activity | 14/3805 | 2.17E-05 | 0.00014426 |
| GO:0048668 | collateral sprouting | 14/3805 | 2.17E-05 | 0.00014426 |
| GO:1905332 | positive regulation of morphogenesis of an epithelium | 16/3805 | 2.23E-05 | 0.000148491 |
| GO:1900180 | regulation of protein localization to nucleus | 39/3805 | 2.30E-05 | 0.000152601 |
| GO:0072088 | nephron epithelium morphogenesis | 26/3805 | 2.34E-05 | 0.000155163 |
| GO:0098773 | skin epidermis development | 33/3805 | 2.43E-05 | 0.000160851 |
| GO:0033077 | T cell differentiation in thymus | 28/3805 | 2.48E-05 | 0.000164408 |
| GO:0000082 | G1/S transition of mitotic cell cycle | 49/3805 | 2.50E-05 | 0.000164991 |
| GO:0060349 | bone morphogenesis | 31/3805 | 2.51E-05 | 0.000165635 |
| GO:0032092 | positive regulation of protein binding | 29/3805 | 2.52E-05 | 0.000165933 |
| GO:0031102 | neuron projection regeneration | 21/3805 | 2.57E-05 | 0.00016912 |
| GO:0060401 | cytosolic calcium ion transport | 48/3805 | 2.69E-05 | 0.000177247 |
| GO:0061162 | establishment of monopolar cell polarity | Dec-05 | 2.70E-05 | 0.000177247 |
| GO:0000727 | double-strand break repair via break-induced replication | Aug-05 | 2.77E-05 | 0.000180935 |
| GO:0048251 | elastic fiber assembly | Aug-05 | 2.77E-05 | 0.000180935 |

| | | | | |
|---|---|---|---|---|
| GO:0060907 | positive regulation of macrophage cytokine production | Aug-05 | 2.77E-05 | 0.000180935 |
| GO:0072697 | protein localization to cell cortex | Aug-05 | 2.77E-05 | 0.000180935 |
| GO:0098911 | regulation of ventricular cardiac muscle cell action potential | Aug-05 | 2.77E-05 | 0.000180935 |
| GO:0021543 | pallium development | 41/3805 | 2.82E-05 | 0.000184465 |
| GO:0010469 | regulation of signaling receptor activity | 38/3805 | 2.85E-05 | 0.000185416 |
| GO:0001977 | renal system process involved in regulation of blood volume | Sep-05 | 2.85E-05 | 0.000185416 |
| GO:0099550 | trans-synaptic signaling, modulating synaptic transmission | Sep-05 | 2.85E-05 | 0.000185416 |
| GO:2000095 | regulation of Wnt signaling pathway, planar cell polarity pathway | Sep-05 | 2.85E-05 | 0.000185416 |
| GO:0060350 | endochondral bone morphogenesis | 22/3805 | 2.88E-05 | 0.000187494 |
| GO:0019722 | calcium-mediated signaling | 47/3805 | 2.90E-05 | 0.000188324 |
| GO:0051302 | regulation of cell division | 47/3805 | 2.90E-05 | 0.000188324 |
| GO:0019229 | regulation of vasoconstriction | 26/3805 | 2.94E-05 | 0.000190241 |
| GO:0001942 | hair follicle development | 32/3805 | 2.99E-05 | 0.000193487 |
| GO:0051054 | positive regulation of DNA metabolic process | 65/3805 | 2.99E-05 | 0.000193487 |
| GO:0086009 | membrane repolarization | 17/3805 | 3.02E-05 | 0.000194656 |
| GO:1902991 | regulation of amyloid precursor protein catabolic process | 17/3805 | 3.02E-05 | 0.000194656 |
| GO:0021549 | cerebellum development | 31/3805 | 3.04E-05 | 0.000196258 |
| GO:0002695 | negative regulation of leukocyte activation | 48/3805 | 3.09E-05 | 0.00019863 |

| | | | | |
|---|---|---|---|---|
| GO:0032147 | activation of protein kinase activity | 29/3805 | 3.09E-05 | 0.00019863 |
| GO:0097205 | renal filtration | 13/3805 | 3.11E-05 | 0.000199912 |
| GO:0043122 | regulation of I-kappaB kinase/NF-kappaB signaling | 46/3805 | 3.13E-05 | 0.00020054 |
| GO:0009581 | detection of external stimulus | 36/3805 | 3.13E-05 | 0.00020054 |
| GO:0048168 | regulation of neuronal synaptic plasticity | 23/3805 | 3.16E-05 | 0.000201511 |
| GO:0055008 | cardiac muscle tissue morphogenesis | 23/3805 | 3.16E-05 | 0.000201511 |
| GO:1990868 | response to chemokine | 23/3805 | 3.16E-05 | 0.000201511 |
| GO:1990869 | cellular response to chemokine | 23/3805 | 3.16E-05 | 0.000201511 |
| GO:0097722 | sperm motility | 39/3805 | 3.17E-05 | 0.000202274 |
| GO:0035850 | epithelial cell differentiation involved in kidney development | 16/3805 | 3.22E-05 | 0.000204833 |
| GO:0060603 | mammary gland duct morphogenesis | 16/3805 | 3.22E-05 | 0.000204833 |
| GO:0007127 | meiosis I | 35/3805 | 3.26E-05 | 0.000207397 |
| GO:0040036 | regulation of fibroblast growth factor receptor signaling pathway | 14/3805 | 3.31E-05 | 0.000209873 |
| GO:0044319 | wound healing, spreading of cells | 14/3805 | 3.31E-05 | 0.000209873 |
| GO:0090505 | epiboly involved in wound healing | 14/3805 | 3.31E-05 | 0.000209873 |
| GO:0060441 | epithelial tube branching involved in lung morphogenesis | 15/3805 | 3.33E-05 | 0.000210909 |
| GO:0048260 | positive regulation of receptor-mediated endocytosis | 21/3805 | 3.37E-05 | 0.000213385 |
| GO:0007632 | visual behavior | 24/3805 | 3.38E-05 | 0.000213385 |

| | | | | |
|---|---|---|---|---|
| GO:0060419 | heart growth | 33/3805 | 3.50E-05 | 0.000221084 |
| GO:0050729 | positive regulation of inflammatory response | 37/3805 | 3.52E-05 | 0.000222344 |
| GO:0002244 | hematopoietic progenitor cell differentiation | 42/3805 | 3.58E-05 | 0.000225872 |
| GO:0045217 | cell-cell junction maintenance | Nov-05 | 3.63E-05 | 0.000228013 |
| GO:0060307 | regulation of ventricular cardiac muscle cell membrane repolarization | Nov-05 | 3.63E-05 | 0.000228013 |
| GO:0060602 | branch elongation of an epithelium | Nov-05 | 3.63E-05 | 0.000228013 |
| GO:0072028 | nephron morphogenesis | 26/3805 | 3.67E-05 | 0.000229895 |
| GO:1902893 | regulation of miRNA transcription | 26/3805 | 3.67E-05 | 0.000229895 |
| GO:0060079 | excitatory postsynaptic potential | 27/3805 | 3.74E-05 | 0.00023422 |
| GO:0034766 | negative regulation of ion transmembrane transport | 30/3805 | 3.74E-05 | 0.00023422 |
| GO:0045185 | maintenance of protein location | 29/3805 | 3.78E-05 | 0.000235892 |
| GO:0070555 | response to interleukin-1 | 28/3805 | 3.78E-05 | 0.000235892 |
| GO:0050819 | negative regulation of coagulation | 18/3805 | 3.79E-05 | 0.000236206 |
| GO:1903053 | regulation of extracellular matrix organization | 18/3805 | 3.79E-05 | 0.000236206 |
| GO:0031647 | regulation of protein stability | 67/3805 | 3.80E-05 | 0.000236657 |
| GO:0010662 | regulation of striated muscle cell apoptotic process | 20/3805 | 3.92E-05 | 0.000243521 |
| GO:0097035 | regulation of membrane lipid distribution | 20/3805 | 3.92E-05 | 0.000243521 |
| GO:0055007 | cardiac muscle cell differentiation | 38/3805 | 3.93E-05 | 0.000243857 |

| | | | | |
|---|---|---|---|---|
| GO:0030282 | bone mineralization | 34/3805 | 4.03E-05 | 0.000250043 |
| GO:0060997 | dendritic spine morphogenesis | 23/3805 | 4.04E-05 | 0.000250043 |
| GO:0070482 | response to oxygen levels | 65/3805 | 4.13E-05 | 0.00025559 |
| GO:0045494 | photoreceptor cell maintenance | 17/3805 | 4.20E-05 | 0.000259676 |
| GO:0099504 | synaptic vesicle cycle | 52/3805 | 4.26E-05 | 0.000263194 |
| GO:0061180 | mammary gland epithelium development | 24/3805 | 4.27E-05 | 0.000263536 |
| GO:0030859 | polarized epithelial cell differentiation | Dec-05 | 4.41E-05 | 0.000270272 |
| GO:0045737 | positive regulation of cyclin-dependent protein serine/threonine kinase activity | Dec-05 | 4.41E-05 | 0.000270272 |
| GO:0061339 | establishment or maintenance of monopolar cell polarity | Dec-05 | 4.41E-05 | 0.000270272 |
| GO:0070293 | renal absorption | Dec-05 | 4.41E-05 | 0.000270272 |
| GO:0099623 | regulation of cardiac muscle cell membrane repolarization | Dec-05 | 4.41E-05 | 0.000270272 |
| GO:2000050 | regulation of non-canonical Wnt signaling pathway | Dec-05 | 4.41E-05 | 0.000270272 |
| GO:0072078 | nephron tubule morphogenesis | 25/3805 | 4.44E-05 | 0.000272271 |
| GO:0071478 | cellular response to radiation | 43/3805 | 4.47E-05 | 0.000273362 |
| GO:0098727 | maintenance of cell number | 43/3805 | 4.47E-05 | 0.000273362 |
| GO:2001242 | regulation of intrinsic apoptotic signaling pathway | 43/3805 | 4.47E-05 | 0.000273362 |
| GO:0010596 | negative regulation of endothelial cell migration | 19/3805 | 4.51E-05 | 0.000275402 |
| GO:0021675 | nerve development | 26/3805 | 4.56E-05 | 0.000277855 |

| | | | | |
|---|---|---|---|---|
| GO:0030168 | platelet activation | 27/3805 | 4.62E-05 | 0.000281231 |
| GO:0000212 | meiotic spindle organization | Oct-05 | 4.70E-05 | 0.000285496 |
| GO:0040037 | negative regulation of fibroblast growth factor receptor signaling pathway | Oct-05 | 4.70E-05 | 0.000285496 |
| GO:0045198 | establishment of epithelial cell apical/basal polarity | Oct-05 | 4.70E-05 | 0.000285496 |
| GO:0055003 | cardiac myofibril assembly | Oct-05 | 4.70E-05 | 0.000285496 |
| GO:0002366 | leukocyte activation involved in immune response | 69/3805 | 4.73E-05 | 0.000287134 |
| GO:0051926 | negative regulation of calcium ion transport | 22/3805 | 4.81E-05 | 0.000291163 |
| GO:0002029 | desensitization of G protein-coupled receptor signaling pathway | 13/3805 | 4.83E-05 | 0.000291592 |
| GO:0006929 | substrate-dependent cell migration | 13/3805 | 4.83E-05 | 0.000291592 |
| GO:0022401 | negative adaptation of signaling pathway | 13/3805 | 4.83E-05 | 0.000291592 |
| GO:0051928 | positive regulation of calcium ion transport | 37/3805 | 4.87E-05 | 0.00029369 |
| GO:0060306 | regulation of membrane repolarization | 14/3805 | 4.94E-05 | 0.00029809 |
| GO:0015748 | organophosphate ester transport | 33/3805 | 4.99E-05 | 0.00030035 |
| GO:0030534 | adult behavior | 45/3805 | 5.10E-05 | 0.000306649 |
| GO:0008038 | neuron recognition | 18/3805 | 5.14E-05 | 0.000308826 |
| GO:0044380 | protein localization to cytoskeleton | 20/3805 | 5.15E-05 | 0.000309458 |
| GO:0022404 | molting cycle process | 32/3805 | 5.17E-05 | 0.000309929 |
| GO:0022405 | hair cycle process | 32/3805 | 5.17E-05 | 0.000309929 |

| | | | | |
|---|---|---|---|---|
| GO:0007519 | skeletal muscle tissue development | 48/3805 | 5.24E-05 | 0.000313718 |
| GO:0050821 | protein stabilization | 46/3805 | 5.39E-05 | 0.000322734 |
| GO:0045638 | negative regulation of myeloid cell differentiation | 26/3805 | 5.63E-05 | 0.000336432 |
| GO:0061614 | miRNA transcription | 26/3805 | 5.63E-05 | 0.000336432 |
| GO:0003229 | ventricular cardiac muscle tissue development | 21/3805 | 5.70E-05 | 0.000339656 |
| GO:0048706 | embryonic skeletal system development | 37/3805 | 5.70E-05 | 0.000339656 |
| GO:0048103 | somatic stem cell division | Sep-05 | 5.75E-05 | 0.000341909 |
| GO:0090128 | regulation of synapse maturation | Sep-05 | 5.75E-05 | 0.000341909 |
| GO:0003149 | membranous septum morphogenesis | Jul-05 | 5.79E-05 | 0.000344024 |
| GO:0099640 | axo-dendritic protein transport | Jul-05 | 5.79E-05 | 0.000344024 |
| GO:0060402 | calcium ion transport into cytosol | 43/3805 | 5.93E-05 | 0.000351575 |
| GO:0010665 | regulation of cardiac muscle cell apoptotic process | 19/3805 | 5.99E-05 | 0.000353788 |
| GO:0043507 | positive regulation of JUN kinase activity | 19/3805 | 5.99E-05 | 0.000353788 |
| GO:0043536 | positive regulation of blood vessel endothelial cell migration | 19/3805 | 5.99E-05 | 0.000353788 |
| GO:0072698 | protein localization to microtubule cytoskeleton | 19/3805 | 5.99E-05 | 0.000353788 |
| GO:0035089 | establishment of apical/basal cell polarity | Nov-05 | 6.12E-05 | 0.000361226 |
| GO:1902992 | negative regulation of amyloid precursor protein catabolic process | Nov-05 | 6.12E-05 | 0.000361226 |
| GO:0007498 | mesoderm development | 32/3805 | 6.17E-05 | 0.000363076 |

| | | | | |
|---|---|---|---|---|
| GO:0042472 | inner ear morphogenesis | 32/3805 | 6.17E-05 | 0.000363076 |
| GO:0003093 | regulation of glomerular filtration | Aug-05 | 6.36E-05 | 0.000372621 |
| GO:0043589 | skin morphogenesis | Aug-05 | 6.36E-05 | 0.000372621 |
| GO:0072393 | microtubule anchoring at microtubule organizing center | Aug-05 | 6.36E-05 | 0.000372621 |
| GO:1903651 | positive regulation of cytoplasmic transport | Aug-05 | 6.36E-05 | 0.000372621 |
| GO:0007093 | mitotic cell cycle checkpoint signaling | 35/3805 | 6.37E-05 | 0.000372621 |
| GO:0009582 | detection of abiotic stimulus | 35/3805 | 6.37E-05 | 0.000372621 |
| GO:0045601 | regulation of endothelial cell differentiation | 16/3805 | 6.38E-05 | 0.00037307 |
| GO:1903076 | regulation of protein localization to plasma membrane | 31/3805 | 6.39E-05 | 0.000373322 |
| GO:0051048 | negative regulation of secretion | 47/3805 | 6.47E-05 | 0.000377631 |
| GO:1903038 | negative regulation of leukocyte cell-cell adhesion | 37/3805 | 6.66E-05 | 0.000388323 |
| GO:0051592 | response to calcium ion | 34/3805 | 6.70E-05 | 0.000390221 |
| GO:0010656 | negative regulation of muscle cell apoptotic process | 20/3805 | 6.72E-05 | 0.000390525 |
| GO:0140115 | export across plasma membrane | 20/3805 | 6.72E-05 | 0.000390525 |
| GO:0071347 | cellular response to interleukin-1 | 24/3805 | 6.72E-05 | 0.000390525 |
| GO:0055081 | anion homeostasis | 18/3805 | 6.89E-05 | 0.000400121 |
| GO:0061053 | somite development | 27/3805 | 6.93E-05 | 0.000401563 |
| GO:0046883 | regulation of hormone secretion | 65/3805 | 6.94E-05 | 0.000401563 |

| | | | | |
|---|---|---|---|---|
| GO:0008593 | regulation of Notch signaling pathway | 26/3805 | 6.94E-05 | 0.000401563 |
| GO:0021846 | cell proliferation in forebrain | 14/3805 | 7.24E-05 | 0.000418066 |
| GO:1901099 | negative regulation of signal transduction in absence of ligand | 14/3805 | 7.24E-05 | 0.000418066 |
| GO:2001240 | negative regulation of extrinsic apoptotic signaling pathway in absence of ligand | 14/3805 | 7.24E-05 | 0.000418066 |
| GO:0071711 | basement membrane organization | 13/3805 | 7.31E-05 | 0.000420851 |
| GO:0099622 | cardiac muscle cell membrane repolarization | 13/3805 | 7.31E-05 | 0.000420851 |
| GO:1905523 | positive regulation of macrophage migration | 13/3805 | 7.31E-05 | 0.000420851 |
| GO:0032635 | interleukin-6 production | 42/3805 | 7.34E-05 | 0.000422023 |
| GO:0016079 | synaptic vesicle exocytosis | 35/3805 | 7.48E-05 | 0.000429613 |
| GO:0042098 | T cell proliferation | 51/3805 | 7.56E-05 | 0.000434277 |
| GO:1902895 | positive regulation of miRNA transcription | 22/3805 | 7.80E-05 | 0.000447279 |
| GO:0010799 | regulation of peptidyl-threonine phosphorylation | 17/3805 | 7.85E-05 | 0.000448746 |
| GO:0030195 | negative regulation of blood coagulation | 17/3805 | 7.85E-05 | 0.000448746 |
| GO:0033628 | regulation of cell adhesion mediated by integrin | 17/3805 | 7.85E-05 | 0.000448746 |
| GO:0010463 | mesenchymal cell proliferation | 19/3805 | 7.88E-05 | 0.000449692 |
| GO:0030514 | negative regulation of BMP signaling pathway | 19/3805 | 7.88E-05 | 0.000449692 |
| GO:0032642 | regulation of chemokine production | 30/3805 | 7.90E-05 | 0.000450358 |
| GO:0051250 | negative regulation of lymphocyte activation | 41/3805 | 7.90E-05 | 0.000450358 |

| | | | | |
|---|---|---|---|---|
| GO:0042471 | ear morphogenesis | 36/3805 | 8.26E-05 | 0.000470081 |
| GO:0002031 | G protein-coupled receptor internalization | Oct-05 | 8.29E-05 | 0.000470848 |
| GO:1902430 | negative regulation of amyloid-beta formation | Oct-05 | 8.29E-05 | 0.000470848 |
| GO:1904754 | positive regulation of vascular associated smooth muscle cell migration | Oct-05 | 8.29E-05 | 0.000470848 |
| GO:0003407 | neural retina development | 24/3805 | 8.37E-05 | 0.000474628 |
| GO:0002699 | positive regulation of immune effector process | 65/3805 | 8.48E-05 | 0.000480911 |
| GO:0030879 | mammary gland development | 40/3805 | 8.50E-05 | 0.000481314 |
| GO:0031122 | cytoplasmic microtubule organization | 20/3805 | 8.70E-05 | 0.000491741 |
| GO:1903078 | positive regulation of protein localization to plasma membrane | 20/3805 | 8.70E-05 | 0.000491741 |
| GO:2000377 | regulation of reactive oxygen species metabolic process | 39/3805 | 9.13E-05 | 0.000515544 |
| GO:0038066 | p38MAPK cascade | 18/3805 | 9.16E-05 | 0.000515782 |
| GO:0072091 | regulation of stem cell proliferation | 18/3805 | 9.16E-05 | 0.000515782 |
| GO:0097720 | calcineurin-mediated signaling | 18/3805 | 9.16E-05 | 0.000515782 |
| GO:0072593 | reactive oxygen species metabolic process | 54/3805 | 9.20E-05 | 0.000517907 |
| GO:0050868 | negative regulation of T cell activation | 34/3805 | 9.27E-05 | 0.000521178 |
| GO:0021987 | cerebral cortex development | 30/3805 | 9.44E-05 | 0.000529608 |
| GO:0055017 | cardiac muscle tissue growth | 30/3805 | 9.44E-05 | 0.000529608 |
| GO:0070849 | response to epidermal growth factor | 15/3805 | 9.69E-05 | 0.000543468 |

| | | | | |
|---|---|---|---|---|
| GO:0097553 | calcium ion transmembrane import into cytosol | 38/3805 | 9.79E-05 | 0.000548387 |
| GO:2000179 | positive regulation of neural precursor cell proliferation | 22/3805 | 9.84E-05 | 0.000550798 |
| GO:0032331 | negative regulation of chondrocyte differentiation | Nov-05 | 9.96E-05 | 0.000555361 |
| GO:0036303 | lymph vessel morphogenesis | Nov-05 | 9.96E-05 | 0.000555361 |
| GO:0051984 | positive regulation of chromosome segregation | Nov-05 | 9.96E-05 | 0.000555361 |
| GO:0098901 | regulation of cardiac muscle cell action potential | Nov-05 | 9.96E-05 | 0.000555361 |
| GO:0001895 | retina homeostasis | 19/3805 | 0.000102743 | 0.000572462 |
| GO:0060042 | retina morphogenesis in camera-type eye | 24/3805 | 0.000103615 | 0.000576808 |
| GO:0008045 | motor neuron axon guidance | 14/3805 | 0.000104164 | 0.000578783 |
| GO:0120316 | sperm flagellum assembly | 14/3805 | 0.000104164 | 0.000578783 |
| GO:0035418 | protein localization to synapse | 25/3805 | 0.000104248 | 0.000578783 |
| GO:0050870 | positive regulation of T cell activation | 50/3805 | 0.000104835 | 0.000581525 |
| GO:0033173 | calcineurin-NFAT signaling cascade | 17/3805 | 0.000105447 | 0.000583366 |
| GO:1900047 | negative regulation of hemostasis | 17/3805 | 0.000105447 | 0.000583366 |
| GO:2001239 | regulation of extrinsic apoptotic signaling pathway in absence of ligand | 17/3805 | 0.000105447 | 0.000583366 |
| GO:1902107 | positive regulation of leukocyte differentiation | 44/3805 | 0.000107154 | 0.00059165 |
| GO:1903708 | positive regulation of hemopoiesis | 44/3805 | 0.000107154 | 0.00059165 |
| GO:0003094 | glomerular filtration | Dec-05 | 0.000107458 | 0.00059165 |

| | | | | |
|---|---|---|---|---|
| GO:0048634 | regulation of muscle organ development | Dec-05 | 0.000107458 | 0.00059165 |
| GO:0032602 | chemokine production | 31/3805 | 0.000107767 | 0.00059165 |
| GO:0060078 | regulation of postsynaptic membrane potential | 31/3805 | 0.000107767 | 0.00059165 |
| GO:0045655 | regulation of monocyte differentiation | Sep-05 | 0.000107797 | 0.00059165 |
| GO:0051895 | negative regulation of focal adhesion assembly | Sep-05 | 0.000107797 | 0.00059165 |
| GO:0150118 | negative regulation of cell-substrate junction organization | Sep-05 | 0.000107797 | 0.00059165 |
| GO:0023058 | adaptation of signaling pathway | 13/3805 | 0.000108288 | 0.000593304 |
| GO:0060074 | synapse maturation | 13/3805 | 0.000108288 | 0.000593304 |
| GO:0090090 | negative regulation of canonical Wnt signaling pathway | 34/3805 | 0.000108653 | 0.000594783 |
| GO:0090316 | positive regulation of intracellular protein transport | 36/3805 | 0.000111981 | 0.000612461 |
| GO:0060538 | skeletal muscle organ development | 49/3805 | 0.000114575 | 0.000626101 |
| GO:0014013 | regulation of gliogenesis | 33/3805 | 0.000114975 | 0.000627741 |
| GO:0017157 | regulation of exocytosis | 54/3805 | 0.000115121 | 0.000627988 |
| GO:0007585 | respiratory gaseous exchange by respiratory system | 21/3805 | 0.000118636 | 0.000646031 |
| GO:0090307 | mitotic spindle assembly | 21/3805 | 0.000118636 | 0.000646031 |
| GO:0019827 | stem cell population maintenance | 41/3805 | 0.000119452 | 0.000649912 |
| GO:0001569 | branching involved in blood vessel morphogenesis | 16/3805 | 0.000119936 | 0.000651976 |
| GO:0031103 | axon regeneration | 18/3805 | 0.000120534 | 0.000653522 |

| | | | | |
|---|---|---|---|---|
| GO:0035272 | exocrine system development | 18/3805 | 0.000120534 | 0.000653522 |
| GO:0045604 | regulation of epidermal cell differentiation | 18/3805 | 0.000120534 | 0.000653522 |
| GO:1904063 | negative regulation of cation transmembrane transport | 27/3805 | 0.000123784 | 0.00067056 |
| GO:0044728 | DNA methylation or demethylation | 26/3805 | 0.000126165 | 0.000682868 |
| GO:0032677 | regulation of interleukin-8 production | 23/3805 | 0.000126409 | 0.000683593 |
| GO:0031397 | negative regulation of protein ubiquitination | 25/3805 | 0.000127539 | 0.000688755 |
| GO:0008625 | extrinsic apoptotic signaling pathway via death domain receptors | 24/3805 | 0.000127694 | 0.000688755 |
| GO:0033209 | tumor necrosis factor-mediated signaling pathway | 24/3805 | 0.000127694 | 0.000688755 |
| GO:0051764 | actin crosslink formation | Aug-05 | 0.000131071 | 0.000705751 |
| GO:0086069 | bundle of His cell to Purkinje myocyte communication | Aug-05 | 0.000131071 | 0.000705751 |
| GO:0051155 | positive regulation of striated muscle cell differentiation | 19/3805 | 0.000132933 | 0.00071516 |
| GO:0035886 | vascular associated smooth muscle cell differentiation | 15/3805 | 0.000134291 | 0.00072122 |
| GO:0086065 | cell communication involved in cardiac conduction | 15/3805 | 0.000134291 | 0.00072122 |
| GO:0009100 | glycoprotein metabolic process | 68/3805 | 0.000137158 | 0.000735989 |
| GO:0001946 | lymphangiogenesis | Oct-05 | 0.000139573 | 0.000747022 |
| GO:0003177 | pulmonary valve development | Oct-05 | 0.000139573 | 0.000747022 |
| GO:0007252 | I-kappaB phosphorylation | Oct-05 | 0.000139573 | 0.000747022 |
| GO:0021532 | neural tube patterning | 17/3805 | 0.000140247 | 0.000749987 |

| | | | | |
|---|---|---|---|---|
| GO:0010968 | regulation of microtubule nucleation | Jul-05 | 0.000141076 | 0.000752488 |
| GO:0061517 | macrophage proliferation | Jul-05 | 0.000141076 | 0.000752488 |
| GO:0072102 | glomerulus morphogenesis | Jul-05 | 0.000141076 | 0.000752488 |
| GO:0051310 | metaphase plate congression | 20/3805 | 0.000142561 | 0.000758865 |
| GO:0002702 | positive regulation of production of molecular mediator of immune response | 32/3805 | 0.000142637 | 0.000758865 |
| GO:0022037 | metencephalon development | 32/3805 | 0.000142637 | 0.000758865 |
| GO:0010952 | positive regulation of peptidase activity | 42/3805 | 0.000143754 | 0.00076416 |
| GO:0022617 | extracellular matrix disassembly | 14/3805 | 0.000147302 | 0.000781023 |
| GO:0071364 | cellular response to epidermal growth factor stimulus | 14/3805 | 0.000147302 | 0.000781023 |
| GO:1904707 | positive regulation of vascular associated smooth muscle cell proliferation | 14/3805 | 0.000147302 | 0.000781023 |
| GO:0009791 | post-embryonic development | 34/3805 | 0.000148157 | 0.000784892 |
| GO:0006940 | regulation of smooth muscle contraction | 21/3805 | 0.000149512 | 0.000790426 |
| GO:0060425 | lung morphogenesis | 21/3805 | 0.000149512 | 0.000790426 |
| GO:0032943 | mononuclear cell proliferation | 69/3805 | 0.000149581 | 0.000790426 |
| GO:0060759 | regulation of response to cytokine stimulus | 36/3805 | 0.000150537 | 0.000794805 |
| GO:0051983 | regulation of chromosome segregation | 26/3805 | 0.000152734 | 0.000805721 |
| GO:0030032 | lamellipodium assembly | 22/3805 | 0.000153992 | 0.000811673 |
| GO:0035914 | skeletal muscle cell differentiation | 25/3805 | 0.000155351 | 0.000817456 |

| | | | | |
|---|---|---|---|---|
| GO:1904427 | positive regulation of calcium ion transmembrane transport | 25/3805 | 0.000155351 | 0.000817456 |
| GO:0000075 | cell cycle checkpoint signaling | 41/3805 | 0.000155951 | 0.000819922 |
| GO:0032637 | interleukin-8 production | 23/3805 | 0.000156265 | 0.000820134 |
| GO:0071277 | cellular response to calcium ion | 23/3805 | 0.000156265 | 0.000820134 |
| GO:0034114 | regulation of heterotypic cell-cell adhesion | Nov-05 | 0.000156647 | 0.000820134 |
| GO:0044331 | cell-cell adhesion mediated by cadherin | Nov-05 | 0.000156647 | 0.000820134 |
| GO:0055083 | monovalent inorganic anion homeostasis | Nov-05 | 0.000156647 | 0.000820134 |
| GO:1905314 | semi-lunar valve development | 13/3805 | 0.000157162 | 0.000822139 |
| GO:1901216 | positive regulation of neuron death | 33/3805 | 0.000157639 | 0.000823943 |
| GO:1990845 | adaptive thermogenesis | 39/3805 | 0.000159869 | 0.000834902 |
| GO:0060444 | branching involved in mammary gland duct morphogenesis | Dec-05 | 0.00016135 | 0.000840525 |
| GO:1903649 | regulation of cytoplasmic transport | Dec-05 | 0.00016135 | 0.000840525 |
| GO:1904031 | positive regulation of cyclin-dependent protein kinase activity | Dec-05 | 0.00016135 | 0.000840525 |
| GO:0048536 | spleen development | 16/3805 | 0.000161507 | 0.000840643 |
| GO:0030216 | keratinocyte differentiation | 37/3805 | 0.000161809 | 0.000841513 |
| GO:0055006 | cardiac cell development | 29/3805 | 0.000165516 | 0.000860076 |
| GO:0002548 | monocyte chemotaxis | 19/3805 | 0.000170658 | 0.000883856 |
| GO:0048278 | vesicle docking | 19/3805 | 0.000170658 | 0.000883856 |

| | | | | |
|---|---|---|---|---|
| GO:0060632 | regulation of microtubule-based movement | 19/3805 | 0.000170658 | 0.000883856 |
| GO:2001238 | positive regulation of extrinsic apoptotic signaling pathway | 19/3805 | 0.000170658 | 0.000883856 |
| GO:2000241 | regulation of reproductive process | 47/3805 | 0.000173484 | 0.00089775 |
| GO:0070663 | regulation of leukocyte proliferation | 57/3805 | 0.00018267 | 0.000944499 |
| GO:0050714 | positive regulation of protein secretion | 39/3805 | 0.000183044 | 0.000945143 |
| GO:0001774 | microglial cell activation | 15/3805 | 0.000183551 | 0.000945143 |
| GO:0006904 | vesicle docking involved in exocytosis | 15/3805 | 0.000183551 | 0.000945143 |
| GO:0007143 | female meiotic nuclear division | 15/3805 | 0.000183551 | 0.000945143 |
| GO:1900744 | regulation of p38MAPK cascade | 15/3805 | 0.000183551 | 0.000945143 |
| GO:1904407 | positive regulation of nitric oxide metabolic process | 17/3805 | 0.000184685 | 0.000950199 |
| GO:0071897 | DNA biosynthetic process | 42/3805 | 0.000185933 | 0.000955831 |
| GO:0010827 | regulation of glucose transmembrane transport | 25/3805 | 0.00018842 | 0.000967027 |
| GO:1901379 | regulation of potassium ion transmembrane transport | 25/3805 | 0.00018842 | 0.000967027 |
| GO:0035633 | maintenance of blood-brain barrier | Sep-05 | 0.000190315 | 0.000973551 |
| GO:0061318 | renal filtration cell differentiation | Sep-05 | 0.000190315 | 0.000973551 |
| GO:0072112 | glomerular visceral epithelial cell differentiation | Sep-05 | 0.000190315 | 0.000973551 |
| GO:0072311 | glomerular epithelial cell differentiation | Sep-05 | 0.000190315 | 0.000973551 |
| GO:0043280 | positive regulation of cysteine-type endopeptidase activity involved in apoptotic process | 32/3805 | 0.000195682 | 0.00100019 |

| | | | | |
|---|---|---|---|---|
| GO:0099175 | regulation of postsynapse organization | 29/3805 | 0.000196127 | 0.001001644 |
| GO:0009411 | response to UV | 36/3805 | 0.000200633 | 0.001022151 |
| GO:0045598 | regulation of fat cell differentiation | 36/3805 | 0.000200633 | 0.001022151 |
| GO:2001056 | positive regulation of cysteine-type endopeptidase activity | 36/3805 | 0.000200633 | 0.001022151 |
| GO:0048016 | inositol phosphate-mediated signaling | 18/3805 | 0.000203346 | 0.001035123 |
| GO:0060236 | regulation of mitotic spindle organization | 14/3805 | 0.000205045 | 0.001041227 |
| GO:0072210 | metanephric nephron development | 14/3805 | 0.000205045 | 0.001041227 |
| GO:0086004 | regulation of cardiac muscle cell contraction | 14/3805 | 0.000205045 | 0.001041227 |
| GO:0009952 | anterior/posterior pattern specification | 51/3805 | 0.000211643 | 0.001073861 |
| GO:0062207 | regulation of pattern recognition receptor signaling pathway | 27/3805 | 0.000213643 | 0.001083129 |
| GO:0007281 | germ cell development | 76/3805 | 0.000214021 | 0.001084164 |
| GO:0003016 | respiratory system process | 16/3805 | 0.000215068 | 0.001085946 |
| GO:0043114 | regulation of vascular permeability | 16/3805 | 0.000215068 | 0.001085946 |
| GO:1903524 | positive regulation of blood circulation | 16/3805 | 0.000215068 | 0.001085946 |
| GO:2000404 | regulation of T cell migration | 16/3805 | 0.000215068 | 0.001085946 |
| GO:0010812 | negative regulation of cell-substrate adhesion | 19/3805 | 0.000217455 | 0.001096225 |
| GO:0032731 | positive regulation of interleukin-1 beta production | 19/3805 | 0.000217455 | 0.001096225 |
| GO:0070509 | calcium ion import | 26/3805 | 0.00022121 | 0.001114256 |

| | | | | |
|---|---|---|---|---|
| GO:0048013 | ephrin receptor signaling pathway | 13/3805 | 0.000223845 | 0.001124808 |
| GO:0099560 | synaptic membrane adhesion | 13/3805 | 0.000223845 | 0.001124808 |
| GO:1903959 | regulation of anion transmembrane transport | 13/3805 | 0.000223845 | 0.001124808 |
| GO:0031579 | membrane raft organization | Oct-05 | 0.000225646 | 0.001129313 |
| GO:0046325 | negative regulation of glucose import | Oct-05 | 0.000225646 | 0.001129313 |
| GO:0051383 | kinetochore organization | Oct-05 | 0.000225646 | 0.001129313 |
| GO:0061081 | positive regulation of myeloid leukocyte cytokine production involved in immune response | Oct-05 | 0.000225646 | 0.001129313 |
| GO:2000810 | regulation of bicellular tight junction assembly | Oct-05 | 0.000225646 | 0.001129313 |
| GO:0010876 | lipid localization | 86/3805 | 0.000226644 | 0.001133402 |
| GO:0001836 | release of cytochrome c from mitochondria | 20/3805 | 0.00022726 | 0.00113557 |
| GO:0060420 | regulation of heart growth | 25/3805 | 0.00022758 | 0.001136265 |
| GO:0071346 | cellular response to interferon-gamma | 29/3805 | 0.00023168 | 0.001154249 |
| GO:0034341 | response to interferon-gamma | 34/3805 | 0.000231737 | 0.001154249 |
| GO:0051588 | regulation of neurotransmitter transport | 34/3805 | 0.000231737 | 0.001154249 |
| GO:0010921 | regulation of phosphatase activity | 24/3805 | 0.000232393 | 0.001156594 |
| GO:0021885 | forebrain cell migration | 21/3805 | 0.000233166 | 0.001157676 |
| GO:0034121 | regulation of toll-like receptor signaling pathway | 21/3805 | 0.000233166 | 0.001157676 |
| GO:2001244 | positive regulation of intrinsic apoptotic signaling pathway | 21/3805 | 0.000233166 | 0.001157676 |

| | | | | |
|---|---|---|---|---|
| GO:0006305 | DNA alkylation | 22/3805 | 0.000235658 | 0.001167263 |
| GO:0006306 | DNA methylation | 22/3805 | 0.000235658 | 0.001167263 |
| GO:1903036 | positive regulation of response to wounding | 22/3805 | 0.000235658 | 0.001167263 |
| GO:0002819 | regulation of adaptive immune response | 51/3805 | 0.000235875 | 0.001167414 |
| GO:0060563 | neuroepithelial cell differentiation | Dec-05 | 0.000236735 | 0.001169815 |
| GO:0090025 | regulation of monocyte chemotaxis | Dec-05 | 0.000236735 | 0.001169815 |
| GO:0006582 | melanin metabolic process | Nov-05 | 0.000239219 | 0.001181155 |
| GO:0045839 | negative regulation of mitotic nuclear division | 17/3805 | 0.000240909 | 0.001186687 |
| GO:0090279 | regulation of calcium ion import | 17/3805 | 0.000240909 | 0.001186687 |
| GO:1902041 | regulation of extrinsic apoptotic signaling pathway via death domain receptors | 17/3805 | 0.000240909 | 0.001186687 |
| GO:0015833 | peptide transport | 65/3805 | 0.000242253 | 0.001192363 |
| GO:1903320 | regulation of protein modification by small protein conjugation or removal | 54/3805 | 0.000243434 | 0.001197235 |
| GO:0030072 | peptide hormone secretion | 60/3805 | 0.000247533 | 0.001214908 |
| GO:0030501 | positive regulation of bone mineralization | 15/3805 | 0.000247708 | 0.001214908 |
| GO:0032924 | activin receptor signaling pathway | 15/3805 | 0.000247708 | 0.001214908 |
| GO:0043455 | regulation of secondary metabolic process | Aug-05 | 0.000248298 | 0.001214908 |
| GO:0048021 | regulation of melanin biosynthetic process | Aug-05 | 0.000248298 | 0.001214908 |
| GO:1900376 | regulation of secondary metabolite biosynthetic process | Aug-05 | 0.000248298 | 0.001214908 |

| | | | | |
|---|---|---|---|---|
| GO:0000018 | regulation of DNA recombination | 33/3805 | 0.000248389 | 0.001214908 |
| GO:1903531 | negative regulation of secretion by cell | 40/3805 | 0.000249935 | 0.001221516 |
| GO:0019748 | secondary metabolic process | 18/3805 | 0.000260845 | 0.001273835 |
| GO:0098656 | anion transmembrane transport | 45/3805 | 0.000262601 | 0.001281411 |
| GO:0045576 | mast cell activation | 26/3805 | 0.000264707 | 0.001289673 |
| GO:0051781 | positive regulation of cell division | 26/3805 | 0.000264707 | 0.001289673 |
| GO:0006470 | protein dephosphorylation | 53/3805 | 0.000267871 | 0.001304071 |
| GO:0018205 | peptidyl-lysine modification | 77/3805 | 0.000269109 | 0.001309076 |
| GO:0035282 | segmentation | 29/3805 | 0.000272847 | 0.001325197 |
| GO:0045639 | positive regulation of myeloid cell differentiation | 29/3805 | 0.000272847 | 0.001325197 |
| GO:0010171 | body morphogenesis | 19/3805 | 0.00027751 | 0.001331994 |
| GO:0046850 | regulation of bone remodeling | 19/3805 | 0.00027751 | 0.001331994 |
| GO:0051784 | negative regulation of nuclear division | 19/3805 | 0.00027751 | 0.001331994 |
| GO:0060711 | labyrinthine layer development | 19/3805 | 0.00027751 | 0.001331994 |
| GO:0002822 | regulation of adaptive immune response based on somatic recombination of immune receptors built from immunoglobulin superfamily domains | 48/3805 | 0.000280157 | 0.00135438 |
| GO:0031396 | regulation of protein ubiquitination | 48/3805 | 0.000280157 | 0.00135438 |
| GO:2000403 | positive regulation of lymphocyte migration | 14/3805 | 0.000281251 | 0.00135862 |
| GO:0045123 | cellular extravasation | 23/3805 | 0.000286547 | 0.001383133 |

| | | | | |
|---|---|---|---|---|
| GO:0006029 | proteoglycan metabolic process | 22/3805 | 0.000289231 | 0.001395008 |
| GO:0006869 | lipid transport | 76/3805 | 0.000296727 | 0.001430061 |
| GO:0042886 | amide transport | 70/3805 | 0.000299546 | 0.001439571 |
| GO:0033262 | regulation of nuclear cell cycle DNA replication | Jul-05 | 0.000299853 | 0.001439571 |
| GO:0038063 | collagen-activated tyrosine kinase receptor signaling pathway | Jul-05 | 0.000299853 | 0.001439571 |
| GO:0051798 | positive regulation of hair follicle development | Jul-05 | 0.000299853 | 0.001439571 |
| GO:0072176 | nephric duct development | Jul-05 | 0.000299853 | 0.001439571 |
| GO:0001704 | formation of primary germ layer | 27/3805 | 0.000301998 | 0.001448753 |
| GO:0046660 | female sex differentiation | 36/3805 | 0.00030394 | 0.001456952 |
| GO:0046651 | lymphocyte proliferation | 67/3805 | 0.000310705 | 0.001488237 |
| GO:0002042 | cell migration involved in sprouting angiogenesis | 17/3805 | 0.000311426 | 0.00148827 |
| GO:0045907 | positive regulation of vasoconstriction | 17/3805 | 0.000311426 | 0.00148827 |
| GO:0070169 | positive regulation of biomineral tissue development | 17/3805 | 0.000311426 | 0.00148827 |
| GO:0007020 | microtubule nucleation | 13/3805 | 0.000313323 | 0.001493903 |
| GO:0035025 | positive regulation of Rho protein signal transduction | 13/3805 | 0.000313323 | 0.001493903 |
| GO:1904738 | vascular associated smooth muscle cell migration | 13/3805 | 0.000313323 | 0.001493903 |
| GO:0010544 | negative regulation of platelet activation | Sep-05 | 0.000319267 | 0.001517612 |
| GO:0022038 | corpus callosum development | Sep-05 | 0.000319267 | 0.001517612 |

| | | | | |
|---|---|---|---|---|
| GO:0071636 | positive regulation of transforming growth factor beta production | Sep-05 | 0.000319267 | 0.001517612 |
| GO:1900117 | regulation of execution phase of apoptosis | Sep-05 | 0.000319267 | 0.001517612 |
| GO:0042552 | myelination | 40/3805 | 0.000321637 | 0.001527712 |
| GO:0048814 | regulation of dendrite morphogenesis | 25/3805 | 0.000328035 | 0.001555736 |
| GO:0070828 | heterochromatin organization | 25/3805 | 0.000328035 | 0.001555736 |
| GO:0017145 | stem cell division | 15/3805 | 0.000330306 | 0.001564132 |
| GO:1900026 | positive regulation of substrate adhesion-dependent cell spreading | 15/3805 | 0.000330306 | 0.001564132 |
| GO:0032755 | positive regulation of interleukin-6 production | 28/3805 | 0.000338937 | 0.001602572 |
| GO:0050886 | endocrine process | 28/3805 | 0.000338937 | 0.001602572 |
| GO:0007288 | sperm axoneme assembly | Dec-05 | 0.000340079 | 0.001605543 |
| GO:1904752 | regulation of vascular associated smooth muscle cell migration | Dec-05 | 0.000340079 | 0.001605543 |
| GO:0010517 | regulation of phospholipase activity | 19/3805 | 0.000345631 | 0.00162929 |
| GO:0043407 | negative regulation of MAP kinase activity | 19/3805 | 0.000345631 | 0.00162929 |
| GO:0055021 | regulation of cardiac muscle tissue growth | 23/3805 | 0.000347441 | 0.001636588 |
| GO:0048469 | cell maturation | 48/3805 | 0.000348696 | 0.001641261 |
| GO:0009266 | response to temperature stimulus | 39/3805 | 0.0003506 | 0.001647739 |
| GO:0010950 | positive regulation of endopeptidase activity | 39/3805 | 0.0003506 | 0.001647739 |
| GO:0022029 | telencephalon cell migration | 20/3805 | 0.000353133 | 0.001658396 |

| | | | | |
|---|---|---|---|---|
| GO:0006346 | DNA methylation-dependent heterochromatin assembly | Nov-05 | 0.000355714 | 0.00165929 |
| GO:0010829 | negative regulation of glucose transmembrane transport | Nov-05 | 0.000355714 | 0.00165929 |
| GO:0030204 | chondroitin sulfate metabolic process | Nov-05 | 0.000355714 | 0.00165929 |
| GO:0051016 | barbed-end actin filament capping | Nov-05 | 0.000355714 | 0.00165929 |
| GO:0060716 | labyrinthine layer blood vessel development | Nov-05 | 0.000355714 | 0.00165929 |
| GO:0062009 | secondary palate development | Nov-05 | 0.000355714 | 0.00165929 |
| GO:0072012 | glomerulus vasculature development | Nov-05 | 0.000355714 | 0.00165929 |
| GO:0072567 | chemokine (C-X-C motif) ligand 2 production | Nov-05 | 0.000355714 | 0.00165929 |
| GO:2000341 | regulation of chemokine (C-X-C motif) ligand 2 production | Nov-05 | 0.000355714 | 0.00165929 |
| GO:0055013 | cardiac muscle cell development | 27/3805 | 0.000357238 | 0.001663913 |
| GO:0070167 | regulation of biomineral tissue development | 27/3805 | 0.000357238 | 0.001663913 |
| GO:0045429 | positive regulation of nitric oxide biosynthetic process | 16/3805 | 0.000369642 | 0.001719126 |
| GO:0061900 | glial cell activation | 16/3805 | 0.000369642 | 0.001719126 |
| GO:1903321 | negative regulation of protein modification by small protein conjugation or removal | 26/3805 | 0.000374922 | 0.001742383 |
| GO:0060351 | cartilage development involved in endochondral bone morphogenesis | 14/3805 | 0.000380501 | 0.001765684 |
| GO:1903670 | regulation of sprouting angiogenesis | 14/3805 | 0.000380501 | 0.001765684 |
| GO:0071496 | cellular response to external stimulus | 61/3805 | 0.000392673 | 0.001820813 |
| GO:0006023 | aminoglycan biosynthetic process | 17/3805 | 0.000399133 | 0.001841202 |

| | | | | |
|---|---|---|---|---|
| GO:0044060 | regulation of endocrine process | 17/3805 | 0.000399133 | 0.001841202 |
| GO:0045773 | positive regulation of axon extension | 17/3805 | 0.000399133 | 0.001841202 |
| GO:0048483 | autonomic nervous system development | 17/3805 | 0.000399133 | 0.001841202 |
| GO:0048701 | embryonic cranial skeleton morphogenesis | 17/3805 | 0.000399133 | 0.001841202 |
| GO:0070527 | platelet aggregation | 17/3805 | 0.000399133 | 0.001841202 |
| GO:0110151 | positive regulation of biomineralization | 17/3805 | 0.000399133 | 0.001841202 |
| GO:0071805 | potassium ion transmembrane transport | 45/3805 | 0.000414198 | 0.001909283 |
| GO:0003170 | heart valve development | 18/3805 | 0.000419381 | 0.001924693 |
| GO:0045071 | negative regulation of viral genome replication | 18/3805 | 0.000419381 | 0.001924693 |
| GO:0045599 | negative regulation of fat cell differentiation | 18/3805 | 0.000419381 | 0.001924693 |
| GO:0045778 | positive regulation of ossification | 18/3805 | 0.000419381 | 0.001924693 |
| GO:0032722 | positive regulation of chemokine production | 23/3805 | 0.00041939 | 0.001924693 |
| GO:0050000 | chromosome localization | 23/3805 | 0.00041939 | 0.001924693 |
| GO:0031349 | positive regulation of defense response | 61/3805 | 0.000429743 | 0.00197076 |
| GO:0019233 | sensory perception of pain | 35/3805 | 0.000430739 | 0.001971774 |
| GO:0002053 | positive regulation of mesenchymal cell proliferation | 13/3805 | 0.000431543 | 0.001971774 |
| GO:0034122 | negative regulation of toll-like receptor signaling pathway | 13/3805 | 0.000431543 | 0.001971774 |
| GO:0048169 | regulation of long-term neuronal synaptic plasticity | 13/3805 | 0.000431543 | 0.001971774 |

| | | | | |
|---|---|---|---|---|
| GO:0120033 | negative regulation of plasma membrane bounded cell projection assembly | 13/3805 | 0.000431543 | 0.001971774 |
| GO:0033135 | regulation of peptidyl-serine phosphorylation | 38/3805 | 0.000433211 | 0.001977949 |
| GO:0034394 | protein localization to cell surface | 21/3805 | 0.000435155 | 0.001983927 |
| GO:0060675 | ureteric bud morphogenesis | 21/3805 | 0.000435155 | 0.001983927 |
| GO:1905521 | regulation of macrophage migration | 15/3805 | 0.000435491 | 0.001984008 |
| GO:0038065 | collagen-activated signaling pathway | Aug-05 | 0.000439118 | 0.001996165 |
| GO:0055057 | neuroblast division | Aug-05 | 0.000439118 | 0.001996165 |
| GO:0060026 | convergent extension | Aug-05 | 0.000439118 | 0.001996165 |
| GO:0009798 | axis specification | 26/3805 | 0.000443846 | 0.002014724 |
| GO:0051304 | chromosome separation | 26/3805 | 0.000443846 | 0.002014724 |
| GO:0007272 | ensheathment of neurons | 40/3805 | 0.000463794 | 0.002102219 |
| GO:0008366 | axon ensheathment | 40/3805 | 0.000463794 | 0.002102219 |
| GO:0045685 | regulation of glial cell differentiation | 25/3805 | 0.000465615 | 0.002108943 |
| GO:0008344 | adult locomotory behavior | 28/3805 | 0.000466307 | 0.002110545 |
| GO:0050728 | negative regulation of inflammatory response | 34/3805 | 0.000466958 | 0.002111964 |
| GO:0032873 | negative regulation of stress-activated MAPK cascade | 16/3805 | 0.000477593 | 0.00215227 |
| GO:0035305 | negative regulation of dephosphorylation | 16/3805 | 0.000477593 | 0.00215227 |
| GO:0043618 | regulation of transcription from RNA polymerase II promoter in response to stress | 16/3805 | 0.000477593 | 0.00215227 |

| | | | | |
|---|---|---|---|---|
| GO:0070303 | negative regulation of stress-activated protein kinase signaling cascade | 16/3805 | 0.000477593 | 0.00215227 |
| GO:0090102 | cochlea development | 16/3805 | 0.000477593 | 0.00215227 |
| GO:0030511 | positive regulation of transforming growth factor beta receptor signaling pathway | Dec-05 | 0.000479154 | 0.002154639 |
| GO:0060977 | coronary vasculature morphogenesis | Dec-05 | 0.000479154 | 0.002154639 |
| GO:1903846 | positive regulation of cellular response to transforming growth factor beta stimulus | Dec-05 | 0.000479154 | 0.002154639 |
| GO:0051651 | maintenance of location in cell | 48/3805 | 0.000479831 | 0.002156134 |
| GO:0046888 | negative regulation of hormone secretion | 24/3805 | 0.000485918 | 0.002181916 |
| GO:0006302 | double-strand break repair | 55/3805 | 0.000488175 | 0.002189276 |
| GO:0006813 | potassium ion transport | 51/3805 | 0.000488258 | 0.002189276 |
| GO:0006304 | DNA modification | 27/3805 | 0.000495024 | 0.002216431 |
| GO:0110149 | regulation of biomineralization | 27/3805 | 0.000495024 | 0.002216431 |
| GO:0015711 | organic anion transport | 69/3805 | 0.000500816 | 0.002240756 |
| GO:0060998 | regulation of dendritic spine development | 23/3805 | 0.000504036 | 0.002253551 |
| GO:0030838 | positive regulation of actin filament polymerization | 17/3805 | 0.000507354 | 0.002262299 |
| GO:0043620 | regulation of DNA-templated transcription in response to stress | 17/3805 | 0.000507354 | 0.002262299 |
| GO:0055010 | ventricular cardiac muscle tissue morphogenesis | 17/3805 | 0.000507354 | 0.002262299 |
| GO:0010762 | regulation of fibroblast migration | 14/3805 | 0.000508166 | 0.002262299 |
| GO:0010939 | regulation of necrotic cell death | 14/3805 | 0.000508166 | 0.002262299 |

| | | | | |
|---|---|---|---|---|
| GO:2000463 | positive regulation of excitatory postsynaptic potential | 14/3805 | 0.000508166 | 0.002262299 |
| GO:0034349 | glial cell apoptotic process | Sep-05 | 0.00051264 | 0.002277348 |
| GO:0034501 | protein localization to kinetochore | Sep-05 | 0.00051264 | 0.002277348 |
| GO:1903083 | protein localization to condensed chromosome | Sep-05 | 0.00051264 | 0.002277348 |
| GO:0060445 | branching involved in salivary gland morphogenesis | Nov-05 | 0.00051635 | 0.002285701 |
| GO:0061437 | renal system vasculature development | Nov-05 | 0.00051635 | 0.002285701 |
| GO:0061440 | kidney vasculature development | Nov-05 | 0.00051635 | 0.002285701 |
| GO:1903055 | positive regulation of extracellular matrix organization | Nov-05 | 0.00051635 | 0.002285701 |
| GO:2000178 | negative regulation of neural precursor cell proliferation | Nov-05 | 0.00051635 | 0.002285701 |
| GO:0002790 | peptide secretion | 60/3805 | 0.000518545 | 0.002293019 |
| GO:0051965 | positive regulation of synapse assembly | 22/3805 | 0.000519105 | 0.002293019 |
| GO:1904888 | cranial skeletal system development | 22/3805 | 0.000519105 | 0.002293019 |
| GO:0060021 | roof of mouth development | 26/3805 | 0.000523649 | 0.002311458 |
| GO:0042987 | amyloid precursor protein catabolic process | 18/3805 | 0.000525977 | 0.002318457 |
| GO:1900024 | regulation of substrate adhesion-dependent cell spreading | 18/3805 | 0.000525977 | 0.002318457 |
| GO:0008217 | regulation of blood pressure | 47/3805 | 0.00052811 | 0.002326218 |
| GO:0042982 | amyloid precursor protein metabolic process | 21/3805 | 0.00053001 | 0.002331695 |
| GO:0072171 | mesonephric tubule morphogenesis | 21/3805 | 0.00053001 | 0.002331695 |

| | | | | |
|---|---|---|---|---|
| GO:0003283 | atrial septum development | Oct-05 | 0.000532481 | 0.002333949 |
| GO:0034405 | response to fluid shear stress | Oct-05 | 0.000532481 | 0.002333949 |
| GO:0042438 | melanin biosynthetic process | Oct-05 | 0.000532481 | 0.002333949 |
| GO:0048670 | regulation of collateral sprouting | Oct-05 | 0.000532481 | 0.002333949 |
| GO:0055064 | chloride ion homeostasis | Oct-05 | 0.000532481 | 0.002333949 |
| GO:0001954 | positive regulation of cell-matrix adhesion | 19/3805 | 0.000534954 | 0.002340431 |
| GO:0032615 | interleukin-12 production | 19/3805 | 0.000534954 | 0.002340431 |
| GO:0048662 | negative regulation of smooth muscle cell proliferation | 20/3805 | 0.000535833 | 0.002340431 |
| GO:0097194 | execution phase of apoptosis | 20/3805 | 0.000535833 | 0.002340431 |
| GO:1902017 | regulation of cilium assembly | 20/3805 | 0.000535833 | 0.002340431 |
| GO:0043266 | regulation of potassium ion transport | 28/3805 | 0.000544516 | 0.002376692 |
| GO:0051235 | maintenance of location | 67/3805 | 0.00056161 | 0.002449593 |
| GO:0002269 | leukocyte activation involved in inflammatory response | 15/3805 | 0.000568063 | 0.002476011 |
| GO:0044557 | relaxation of smooth muscle | Jul-05 | 0.000575532 | 0.002503323 |
| GO:0051639 | actin filament network formation | Jul-05 | 0.000575532 | 0.002503323 |
| GO:0090557 | establishment of endothelial intestinal barrier | Jul-05 | 0.000575532 | 0.002503323 |
| GO:0035270 | endocrine system development | 33/3805 | 0.000579009 | 0.002516694 |
| GO:0006821 | chloride transport | 27/3805 | 0.000579969 | 0.002519116 |

| | | | | |
|---|---|---|---|---|
| GO:0010923 | negative regulation of phosphatase activity | 13/3805 | 0.000585507 | 0.002536113 |
| GO:0045746 | negative regulation of Notch signaling pathway | 13/3805 | 0.000585507 | 0.002536113 |
| GO:0051647 | nucleus localization | 13/3805 | 0.000585507 | 0.002536113 |
| GO:0090313 | regulation of protein targeting to membrane | 13/3805 | 0.000585507 | 0.002536113 |
| GO:0042476 | odontogenesis | 31/3805 | 0.000590553 | 0.00255562 |
| GO:0038061 | NIK/NF-kappaB signaling | 29/3805 | 0.000591871 | 0.002560129 |
| GO:0071241 | cellular response to inorganic substance | 50/3805 | 0.00059457 | 0.002570026 |
| GO:1904950 | negative regulation of establishment of protein localization | 34/3805 | 0.000608889 | 0.002630099 |
| GO:0007520 | myoblast fusion | 16/3805 | 0.000611494 | 0.002634063 |
| GO:0008089 | anterograde axonal transport | 16/3805 | 0.000611494 | 0.002634063 |
| GO:0043268 | positive regulation of potassium ion transport | 16/3805 | 0.000611494 | 0.002634063 |
| GO:0045933 | positive regulation of muscle contraction | 16/3805 | 0.000611494 | 0.002634063 |
| GO:0009615 | response to virus | 66/3805 | 0.000619608 | 0.002667175 |
| GO:0045670 | regulation of osteoclast differentiation | 22/3805 | 0.000624954 | 0.002686485 |
| GO:1902807 | negative regulation of cell cycle G1/S phase transition | 22/3805 | 0.000624954 | 0.002686485 |
| GO:0016570 | histone modification | 91/3805 | 0.000627225 | 0.002694394 |
| GO:0015698 | inorganic anion transport | 35/3805 | 0.000636424 | 0.002732031 |
| GO:0061001 | regulation of dendritic spine morphogenesis | 17/3805 | 0.00063987 | 0.002744938 |

| | | | | |
|---|---|---|---|---|
| GO:0140056 | organelle localization by membrane tethering | 21/3805 | 0.000642524 | 0.002750655 |
| GO:1901184 | regulation of ERBB signaling pathway | 21/3805 | 0.000642524 | 0.002750655 |
| GO:1905818 | regulation of chromosome separation | 21/3805 | 0.000642524 | 0.002750655 |
| GO:0002285 | lymphocyte activation involved in immune response | 47/3805 | 0.000650093 | 0.002781153 |
| GO:0042733 | embryonic digit morphogenesis | 20/3805 | 0.000654526 | 0.002798202 |
| GO:0045682 | regulation of epidermis development | 18/3805 | 0.000655126 | 0.002798849 |
| GO:0002720 | positive regulation of cytokine production involved in immune response | 19/3805 | 0.00065934 | 0.002813006 |
| GO:0050891 | multicellular organismal water homeostasis | 19/3805 | 0.00065934 | 0.002813006 |
| GO:0045667 | regulation of osteoblast differentiation | 36/3805 | 0.000661591 | 0.002817706 |
| GO:0051224 | negative regulation of protein transport | 33/3805 | 0.00066216 | 0.002817706 |
| GO:0033687 | osteoblast proliferation | Dec-05 | 0.000663149 | 0.002817706 |
| GO:0051642 | centrosome localization | Dec-05 | 0.000663149 | 0.002817706 |
| GO:0060740 | prostate gland epithelium morphogenesis | Dec-05 | 0.000663149 | 0.002817706 |
| GO:2000108 | positive regulation of leukocyte apoptotic process | Dec-05 | 0.000663149 | 0.002817706 |
| GO:0090224 | regulation of spindle organization | 14/3805 | 0.000670474 | 0.002844959 |
| GO:0110110 | positive regulation of animal organ morphogenesis | 14/3805 | 0.000670474 | 0.002844959 |
| GO:0002237 | response to molecule of bacterial origin | 77/3805 | 0.000678004 | 0.002874959 |
| GO:0033138 | positive regulation of peptidyl-serine phosphorylation | 31/3805 | 0.000678894 | 0.002876778 |

| | | | | |
|---|---|---|---|---|
| GO:1900182 | positive regulation of protein localization to nucleus | 24/3805 | 0.000685994 | 0.00289681 |
| GO:0001765 | membrane raft assembly | Jun-05 | 0.000688259 | 0.00289681 |
| GO:0006971 | hypotonic response | Jun-05 | 0.000688259 | 0.00289681 |
| GO:0010216 | maintenance of DNA methylation | Jun-05 | 0.000688259 | 0.00289681 |
| GO:0021957 | corticospinal tract morphogenesis | Jun-05 | 0.000688259 | 0.00289681 |
| GO:0036363 | transforming growth factor beta activation | Jun-05 | 0.000688259 | 0.00289681 |
| GO:0038130 | ERBB4 signaling pathway | Jun-05 | 0.000688259 | 0.00289681 |
| GO:0048671 | negative regulation of collateral sprouting | Jun-05 | 0.000688259 | 0.00289681 |
| GO:0090306 | meiotic spindle assembly | Jun-05 | 0.000688259 | 0.00289681 |
| GO:1900119 | positive regulation of execution phase of apoptosis | Jun-05 | 0.000688259 | 0.00289681 |
| GO:0007626 | locomotory behavior | 54/3805 | 0.000713515 | 0.003001086 |
| GO:0045995 | regulation of embryonic development | 23/3805 | 0.000718926 | 0.00302181 |
| GO:0003148 | outflow tract septum morphogenesis | Nov-05 | 0.000733295 | 0.003071198 |
| GO:0006024 | glycosaminoglycan biosynthetic process | 15/3805 | 0.00073353 | 0.003071198 |
| GO:0071300 | cellular response to retinoic acid | 15/3805 | 0.00073353 | 0.003071198 |
| GO:0051014 | actin filament severing | Aug-05 | 0.000733626 | 0.003071198 |
| GO:0061308 | cardiac neural crest cell development involved in heart development | Aug-05 | 0.000733626 | 0.003071198 |
| GO:1902285 | semaphorin-plexin signaling pathway involved in neuron projection guidance | Aug-05 | 0.000733626 | 0.003071198 |

| | | | | |
|---|---|---|---|---|
| GO:0014015 | positive regulation of gliogenesis | 22/3805 | 0.000749036 | 0.003133611 |
| GO:0060428 | lung epithelium development | 16/3805 | 0.000776198 | 0.003245071 |
| GO:0007413 | axonal fasciculation | Oct-05 | 0.000783037 | 0.003253181 |
| GO:0010759 | positive regulation of macrophage chemotaxis | Oct-05 | 0.000783037 | 0.003253181 |
| GO:0035024 | negative regulation of Rho protein signal transduction | Oct-05 | 0.000783037 | 0.003253181 |
| GO:0035988 | chondrocyte proliferation | Oct-05 | 0.000783037 | 0.003253181 |
| GO:0062098 | regulation of programmed necrotic cell death | Oct-05 | 0.000783037 | 0.003253181 |
| GO:0106030 | neuron projection fasciculation | Oct-05 | 0.000783037 | 0.003253181 |
| GO:0150105 | protein localization to cell-cell junction | Oct-05 | 0.000783037 | 0.003253181 |
| GO:0051294 | establishment of spindle orientation | 13/3805 | 0.000783347 | 0.003253181 |
| GO:1901385 | regulation of voltage-gated calcium channel activity | 13/3805 | 0.000783347 | 0.003253181 |
| GO:1902003 | regulation of amyloid-beta formation | 13/3805 | 0.000783347 | 0.003253181 |
| GO:0032332 | positive regulation of chondrocyte differentiation | Sep-05 | 0.00079244 | 0.003280042 |
| GO:0035994 | response to muscle stretch | Sep-05 | 0.00079244 | 0.003280042 |
| GO:0045606 | positive regulation of epidermal cell differentiation | Sep-05 | 0.00079244 | 0.003280042 |
| GO:0090594 | inflammatory response to wounding | Sep-05 | 0.00079244 | 0.003280042 |
| GO:1903909 | regulation of receptor clustering | Sep-05 | 0.00079244 | 0.003280042 |
| GO:1903317 | regulation of protein maturation | 20/3805 | 0.000795247 | 0.003289479 |

| | | | | |
|---|---|---|---|---|
| GO:0006493 | protein O-linked glycosylation | 17/3805 | 0.000800956 | 0.003304341 |
| GO:0031529 | ruffle organization | 17/3805 | 0.000800956 | 0.003304341 |
| GO:1901861 | regulation of muscle tissue development | 17/3805 | 0.000800956 | 0.003304341 |
| GO:1903533 | regulation of protein targeting | 17/3805 | 0.000800956 | 0.003304341 |
| GO:0021575 | hindbrain morphogenesis | 18/3805 | 0.000810592 | 0.003337478 |
| GO:0050982 | detection of mechanical stimulus | 18/3805 | 0.000810592 | 0.003337478 |
| GO:0070098 | chemokine-mediated signaling pathway | 18/3805 | 0.000810592 | 0.003337478 |
| GO:0001959 | regulation of cytokine-mediated signaling pathway | 32/3805 | 0.000820248 | 0.003375012 |
| GO:0006865 | amino acid transport | 36/3805 | 0.00084481 | 0.003473786 |
| GO:0050766 | positive regulation of phagocytosis | 23/3805 | 0.000853419 | 0.003506877 |
| GO:0032944 | regulation of mononuclear cell proliferation | 51/3805 | 0.000873801 | 0.003588273 |
| GO:0045823 | positive regulation of heart contraction | 14/3805 | 0.000874568 | 0.003589061 |
| GO:0002092 | positive regulation of receptor internalization | Dec-05 | 0.000902769 | 0.003692662 |
| GO:0048799 | animal organ maturation | Dec-05 | 0.000902769 | 0.003692662 |
| GO:0060512 | prostate gland morphogenesis | Dec-05 | 0.000902769 | 0.003692662 |
| GO:0061842 | microtubule organizing center localization | Dec-05 | 0.000902769 | 0.003692662 |
| GO:2000406 | positive regulation of T cell migration | Dec-05 | 0.000902769 | 0.003692662 |
| GO:1901800 | positive regulation of proteasomal protein catabolic process | 27/3805 | 0.000915961 | 0.003744169 |

| | | | | |
|---|---|---|---|---|
| GO:0006919 | activation of cysteine-type endopeptidase activity involved in apoptotic process | 21/3805 | 0.000930414 | 0.003800761 |
| GO:0003382 | epithelial cell morphogenesis | 15/3805 | 0.000938146 | 0.003827343 |
| GO:0043403 | skeletal muscle tissue regeneration | 15/3805 | 0.000938146 | 0.003827343 |
| GO:0001541 | ovarian follicle development | 20/3805 | 0.00096124 | 0.003916442 |
| GO:0032732 | positive regulation of interleukin-1 production | 20/3805 | 0.00096124 | 0.003916442 |
| GO:0043123 | positive regulation of I-kappaB kinase/NF-kappaB signaling | 30/3805 | 0.000966676 | 0.003933459 |
| GO:0051101 | regulation of DNA binding | 30/3805 | 0.000966676 | 0.003933459 |
| GO:0030522 | intracellular receptor signaling pathway | 46/3805 | 0.000970626 | 0.00394696 |
| GO:0007131 | reciprocal meiotic recombination | 16/3805 | 0.000977181 | 0.003965869 |
| GO:0021795 | cerebral cortex cell migration | 16/3805 | 0.000977181 | 0.003965869 |
| GO:0140527 | reciprocal homologous recombination | 16/3805 | 0.000977181 | 0.003965869 |
| GO:0060389 | pathway-restricted SMAD protein phosphorylation | 19/3805 | 0.000984167 | 0.003991626 |
| GO:0043666 | regulation of phosphoprotein phosphatase activity | 17/3805 | 0.000995405 | 0.004034098 |
| GO:0051785 | positive regulation of nuclear division | 18/3805 | 0.000996576 | 0.004034098 |
| GO:1905517 | macrophage migration | 18/3805 | 0.000996576 | 0.004034098 |
| GO:0034644 | cellular response to UV | 23/3805 | 0.001009133 | 0.004082284 |
| GO:0006750 | glutathione biosynthetic process | Jul-05 | 0.001019993 | 0.004092283 |
| GO:0007144 | female meiosis I | Jul-05 | 0.001019993 | 0.004092283 |

| | | | | |
|---|---|---|---|---|
| GO:0010359 | regulation of anion channel activity | Jul-05 | 0.001019993 | 0.004092283 |
| GO:0016081 | synaptic vesicle docking | Jul-05 | 0.001019993 | 0.004092283 |
| GO:0035791 | platelet-derived growth factor receptor-beta signaling pathway | Jul-05 | 0.001019993 | 0.004092283 |
| GO:0051152 | positive regulation of smooth muscle cell differentiation | Jul-05 | 0.001019993 | 0.004092283 |
| GO:0060346 | bone trabecula formation | Jul-05 | 0.001019993 | 0.004092283 |
| GO:1905063 | regulation of vascular associated smooth muscle cell differentiation | Jul-05 | 0.001019993 | 0.004092283 |
| GO:2000105 | positive regulation of DNA-templated DNA replication | Jul-05 | 0.001019993 | 0.004092283 |
| GO:0030947 | regulation of vascular endothelial growth factor receptor signaling pathway | Nov-05 | 0.001020778 | 0.004092283 |
| GO:0045932 | negative regulation of muscle contraction | Nov-05 | 0.001020778 | 0.004092283 |
| GO:0048745 | smooth muscle tissue development | Nov-05 | 0.001020778 | 0.004092283 |
| GO:0060343 | trabecula formation | Nov-05 | 0.001020778 | 0.004092283 |
| GO:0140058 | neuron projection arborization | Nov-05 | 0.001020778 | 0.004092283 |
| GO:0010522 | regulation of calcium ion transport into cytosol | 27/3805 | 0.001060492 | 0.004248771 |
| GO:1903825 | organic acid transmembrane transport | 32/3805 | 0.001063215 | 0.00425695 |
| GO:0048545 | response to steroid hormone | 47/3805 | 0.001070952 | 0.00428518 |
| GO:0042770 | signal transduction in response to DNA damage | 37/3805 | 0.001095905 | 0.004382216 |
| GO:0071621 | granulocyte chemotaxis | 30/3805 | 0.001105714 | 0.004418611 |
| GO:0044550 | secondary metabolite biosynthetic process | Oct-05 | 0.001122923 | 0.004481648 |

| | | | | |
|---|---|---|---|---|
| GO:0071459 | protein localization to chromosome, centromeric region | Oct-05 | 0.001122923 | 0.004481648 |
| GO:0099084 | postsynaptic specialization organization | 14/3805 | 0.001128557 | 0.004498383 |
| GO:1901021 | positive regulation of calcium ion transmembrane transporter activity | 14/3805 | 0.001128557 | 0.004498383 |
| GO:1904659 | glucose transmembrane transport | 28/3805 | 0.001133418 | 0.004514877 |
| GO:0046631 | alpha-beta T cell activation | 40/3805 | 0.001149753 | 0.004577027 |
| GO:0032623 | interleukin-2 production | 20/3805 | 0.001156086 | 0.004599309 |
| GO:0010212 | response to ionizing radiation | 31/3805 | 0.001158776 | 0.004604145 |
| GO:0035303 | regulation of dephosphorylation | 31/3805 | 0.001158776 | 0.004604145 |
| GO:0035330 | regulation of hippo signaling | Aug-05 | 0.001168275 | 0.004615461 |
| GO:0036035 | osteoclast development | Aug-05 | 0.001168275 | 0.004615461 |
| GO:0036159 | inner dynein arm assembly | Aug-05 | 0.001168275 | 0.004615461 |
| GO:0044406 | adhesion of symbiont to host | Aug-05 | 0.001168275 | 0.004615461 |
| GO:0061307 | cardiac neural crest cell differentiation involved in heart development | Aug-05 | 0.001168275 | 0.004615461 |
| GO:0062099 | negative regulation of programmed necrotic cell death | Aug-05 | 0.001168275 | 0.004615461 |
| GO:0070486 | leukocyte aggregation | Aug-05 | 0.001168275 | 0.004615461 |
| GO:1901881 | positive regulation of protein depolymerization | Aug-05 | 0.001168275 | 0.004615461 |
| GO:1904424 | regulation of GTP binding | Aug-05 | 0.001168275 | 0.004615461 |
| GO:0090276 | regulation of peptide hormone secretion | 48/3805 | 0.001175147 | 0.004639676 |

| | | | | |
|---|---|---|---|---|
| GO:0017121 | plasma membrane phospholipid scrambling | Sep-05 | 0.001184837 | 0.004654396 |
| GO:0044851 | hair cycle phase | Sep-05 | 0.001184837 | 0.004654396 |
| GO:0048486 | parasympathetic nervous system development | Sep-05 | 0.001184837 | 0.004654396 |
| GO:0051770 | positive regulation of nitric-oxide synthase biosynthetic process | Sep-05 | 0.001184837 | 0.004654396 |
| GO:0060438 | trachea development | Sep-05 | 0.001184837 | 0.004654396 |
| GO:0072111 | cell proliferation involved in kidney development | Sep-05 | 0.001184837 | 0.004654396 |
| GO:0090026 | positive regulation of monocyte chemotaxis | Sep-05 | 0.001184837 | 0.004654396 |
| GO:2001224 | positive regulation of neuron migration | Sep-05 | 0.001184837 | 0.004654396 |
| GO:1990573 | potassium ion import across plasma membrane | 15/3805 | 0.001188957 | 0.004667645 |
| GO:0021695 | cerebellar cortex development | 19/3805 | 0.001192378 | 0.004672265 |
| GO:0070613 | regulation of protein processing | 19/3805 | 0.001192378 | 0.004672265 |
| GO:0099601 | regulation of neurotransmitter receptor activity | 19/3805 | 0.001192378 | 0.004672265 |
| GO:0008608 | attachment of spindle microtubules to kinetochore | Dec-05 | 0.001210306 | 0.004727687 |
| GO:0010667 | negative regulation of cardiac muscle cell apoptotic process | Dec-05 | 0.001210306 | 0.004727687 |
| GO:0043243 | positive regulation of protein-containing complex disassembly | Dec-05 | 0.001210306 | 0.004727687 |
| GO:1990089 | response to nerve growth factor | Dec-05 | 0.001210306 | 0.004727687 |
| GO:1990090 | cellular response to nerve growth factor stimulus | Dec-05 | 0.001210306 | 0.004727687 |
| GO:0050435 | amyloid-beta metabolic process | 18/3805 | 0.001217743 | 0.004750795 |

| | | | | |
|---|---|---|---|---|
| GO:0060393 | regulation of pathway-restricted SMAD protein phosphorylation | 18/3805 | 0.001217743 | 0.004750795 |
| GO:0035825 | homologous recombination | 16/3805 | 0.001220569 | 0.004755881 |
| GO:0045912 | negative regulation of carbohydrate metabolic process | 16/3805 | 0.001220569 | 0.004755881 |
| GO:0050775 | positive regulation of dendrite morphogenesis | 17/3805 | 0.001228558 | 0.004781043 |
| GO:0071622 | regulation of granulocyte chemotaxis | 17/3805 | 0.001228558 | 0.004781043 |
| GO:2000045 | regulation of G1/S transition of mitotic cell cycle | 33/3805 | 0.001249513 | 0.004859563 |
| GO:0030500 | regulation of bone mineralization | 22/3805 | 0.001257168 | 0.004886291 |
| GO:0034219 | carbohydrate transmembrane transport | 30/3805 | 0.001261823 | 0.004901335 |
| GO:0008645 | hexose transmembrane transport | 28/3805 | 0.001301899 | 0.005050721 |
| GO:0031058 | positive regulation of histone modification | 28/3805 | 0.001301899 | 0.005050721 |
| GO:0008643 | carbohydrate transport | 35/3805 | 0.001320493 | 0.005119679 |
| GO:0009880 | embryonic pattern specification | 21/3805 | 0.001322875 | 0.005125731 |
| GO:0009101 | glycoprotein biosynthetic process | 54/3805 | 0.00133238 | 0.005159358 |
| GO:0002676 | regulation of chronic inflammatory response | Jun-05 | 0.001344878 | 0.005172482 |
| GO:0009415 | response to water | Jun-05 | 0.001344878 | 0.005172482 |
| GO:0034115 | negative regulation of heterotypic cell-cell adhesion | Jun-05 | 0.001344878 | 0.005172482 |
| GO:0034454 | microtubule anchoring at centrosome | Jun-05 | 0.001344878 | 0.005172482 |
| GO:0038180 | nerve growth factor signaling pathway | Jun-05 | 0.001344878 | 0.005172482 |

| | | | | |
|---|---|---|---|---|
| GO:0045602 | negative regulation of endothelial cell differentiation | Jun-05 | 0.001344878 | 0.005172482 |
| GO:0070587 | regulation of cell-cell adhesion involved in gastrulation | Jun-05 | 0.001344878 | 0.005172482 |
| GO:0072178 | nephric duct morphogenesis | Jun-05 | 0.001344878 | 0.005172482 |
| GO:1902513 | regulation of organelle transport along microtubule | Jun-05 | 0.001344878 | 0.005172482 |
| GO:1903564 | regulation of protein localization to cilium | Jun-05 | 0.001344878 | 0.005172482 |
| GO:2001204 | regulation of osteoclast development | Jun-05 | 0.001344878 | 0.005172482 |
| GO:0032232 | negative regulation of actin filament bundle assembly | 13/3805 | 0.001349232 | 0.005179658 |
| GO:0034110 | regulation of homotypic cell-cell adhesion | 13/3805 | 0.001349232 | 0.005179658 |
| GO:0106056 | regulation of calcineurin-mediated signaling | 13/3805 | 0.001349232 | 0.005179658 |
| GO:0010611 | regulation of cardiac muscle hypertrophy | 20/3805 | 0.001383718 | 0.005299019 |
| GO:0016445 | somatic diversification of immunoglobulins | 20/3805 | 0.001383718 | 0.005299019 |
| GO:0048645 | animal organ formation | 20/3805 | 0.001383718 | 0.005299019 |
| GO:0071479 | cellular response to ionizing radiation | 20/3805 | 0.001383718 | 0.005299019 |
| GO:0007097 | nuclear migration | Nov-05 | 0.001395162 | 0.005320025 |
| GO:0010453 | regulation of cell fate commitment | Nov-05 | 0.001395162 | 0.005320025 |
| GO:0048535 | lymph node development | Nov-05 | 0.001395162 | 0.005320025 |
| GO:1903319 | positive regulation of protein maturation | Nov-05 | 0.001395162 | 0.005320025 |
| GO:2000648 | positive regulation of stem cell proliferation | Nov-05 | 0.001395162 | 0.005320025 |

| | | | | |
|---|---|---|---|---|
| GO:0015914 | phospholipid transport | 23/3805 | 0.001395165 | 0.005320025 |
| GO:0045814 | negative regulation of gene expression, epigenetic | 23/3805 | 0.001395165 | 0.005320025 |
| GO:0002456 | T cell mediated immunity | 33/3805 | 0.001410557 | 0.005372157 |
| GO:1903364 | positive regulation of cellular protein catabolic process | 33/3805 | 0.001410557 | 0.005372157 |
| GO:0036473 | cell death in response to oxidative stress | 25/3805 | 0.001420448 | 0.005406532 |
| GO:0001659 | temperature homeostasis | 40/3805 | 0.001421725 | 0.005408098 |
| GO:0010569 | regulation of double-strand break repair via homologous recombination | 19/3805 | 0.001436972 | 0.005456127 |
| GO:0046324 | regulation of glucose import | 19/3805 | 0.001436972 | 0.005456127 |
| GO:0050885 | neuromuscular process controlling balance | 19/3805 | 0.001436972 | 0.005456127 |
| GO:0009743 | response to carbohydrate | 44/3805 | 0.001439575 | 0.005462688 |
| GO:0002703 | regulation of leukocyte mediated immunity | 59/3805 | 0.001452758 | 0.005509367 |
| GO:0032613 | interleukin-10 production | 18/3805 | 0.001479237 | 0.005596192 |
| GO:0060038 | cardiac muscle cell proliferation | 18/3805 | 0.001479237 | 0.005596192 |
| GO:0060999 | positive regulation of dendritic spine development | 18/3805 | 0.001479237 | 0.005596192 |
| GO:0098930 | axonal transport | 18/3805 | 0.001479237 | 0.005596192 |
| GO:0043525 | positive regulation of neuron apoptotic process | 22/3805 | 0.001481966 | 0.0055965 |
| GO:0002821 | positive regulation of adaptive immune response | 35/3805 | 0.001482006 | 0.0055965 |
| GO:0097530 | granulocyte migration | 35/3805 | 0.001482006 | 0.0055965 |

| | | | | |
|---|---|---|---|---|
| GO:0022600 | digestive system process | 28/3805 | 0.001491614 | 0.005629377 |
| GO:0003197 | endocardial cushion development | 15/3805 | 0.001493827 | 0.005630925 |
| GO:0003254 | regulation of membrane depolarization | 15/3805 | 0.001493827 | 0.005630925 |
| GO:0032757 | positive regulation of interleukin-8 production | 17/3805 | 0.001506323 | 0.005674602 |
| GO:0002762 | negative regulation of myeloid leukocyte differentiation | 16/3805 | 0.001513171 | 0.005696965 |
| GO:0033555 | multicellular organismal response to stress | 26/3805 | 0.001524005 | 0.005734298 |
| GO:0031668 | cellular response to extracellular stimulus | 48/3805 | 0.001553244 | 0.005840795 |
| GO:0022406 | membrane docking | 21/3805 | 0.001567148 | 0.005882454 |
| GO:0032088 | negative regulation of NF-kappaB transcription factor activity | 21/3805 | 0.001567148 | 0.005882454 |
| GO:0050709 | negative regulation of protein secretion | 21/3805 | 0.001567148 | 0.005882454 |
| GO:0003071 | renal system process involved in regulation of systemic arterial blood pressure | Oct-05 | 0.001574223 | 0.005894842 |
| GO:0032528 | microvillus organization | Oct-05 | 0.001574223 | 0.005894842 |
| GO:0034453 | microtubule anchoring | Oct-05 | 0.001574223 | 0.005894842 |
| GO:0035809 | regulation of urine volume | Oct-05 | 0.001574223 | 0.005894842 |
| GO:0040001 | establishment of mitotic spindle localization | Dec-05 | 0.001599688 | 0.005983024 |
| GO:1904037 | positive regulation of epithelial cell apoptotic process | Dec-05 | 0.001599688 | 0.005983024 |
| GO:0010466 | negative regulation of peptidase activity | 52/3805 | 0.001625362 | 0.006071776 |
| GO:2001020 | regulation of response to DNA damage stimulus | 52/3805 | 0.001625362 | 0.006071776 |

| | | | | |
|---|---|---|---|---|
| GO:0061515 | myeloid cell development | 23/3805 | 0.001631555 | 0.00609127 |
| GO:0021510 | spinal cord development | 25/3805 | 0.00164441 | 0.006128273 |
| GO:0021761 | limbic system development | 25/3805 | 0.00164441 | 0.006128273 |
| GO:0045132 | meiotic chromosome segregation | 25/3805 | 0.00164441 | 0.006128273 |
| GO:0000725 | recombinational repair | 35/3805 | 0.001660421 | 0.006184253 |
| GO:0003091 | renal water homeostasis | Jul-05 | 0.001695225 | 0.006280177 |
| GO:0030320 | cellular monovalent inorganic anion homeostasis | Jul-05 | 0.001695225 | 0.006280177 |
| GO:0032530 | regulation of microvillus organization | Jul-05 | 0.001695225 | 0.006280177 |
| GO:0034350 | regulation of glial cell apoptotic process | Jul-05 | 0.001695225 | 0.006280177 |
| GO:0061577 | calcium ion transmembrane transport via high voltage-gated calcium channel | Jul-05 | 0.001695225 | 0.006280177 |
| GO:0070307 | lens fiber cell development | Jul-05 | 0.001695225 | 0.006280177 |
| GO:0086014 | atrial cardiac muscle cell action potential | Jul-05 | 0.001695225 | 0.006280177 |
| GO:0086026 | atrial cardiac muscle cell to AV node cell signaling | Jul-05 | 0.001695225 | 0.006280177 |
| GO:0086066 | atrial cardiac muscle cell to AV node cell communication | Jul-05 | 0.001695225 | 0.006280177 |
| GO:0008637 | apoptotic mitochondrial changes | 28/3805 | 0.001704706 | 0.006311555 |
| GO:0002115 | store-operated calcium entry | Sep-05 | 0.001720163 | 0.006341203 |
| GO:0010232 | vascular transport | Sep-05 | 0.001720163 | 0.006341203 |
| GO:0021535 | cell migration in hindbrain | Sep-05 | 0.001720163 | 0.006341203 |

| | | | | |
|---|---|---|---|---|
| GO:0033630 | positive regulation of cell adhesion mediated by integrin | Sep-05 | 0.001720163 | 0.006341203 |
| GO:0072202 | cell differentiation involved in metanephros development | Sep-05 | 0.001720163 | 0.006341203 |
| GO:1902894 | negative regulation of miRNA transcription | Sep-05 | 0.001720163 | 0.006341203 |
| GO:0002763 | positive regulation of myeloid leukocyte differentiation | 19/3805 | 0.001722866 | 0.006341203 |
| GO:0017156 | calcium-ion regulated exocytosis | 19/3805 | 0.001722866 | 0.006341203 |
| GO:0042058 | regulation of epidermal growth factor receptor signaling pathway | 19/3805 | 0.001722866 | 0.006341203 |
| GO:2000134 | negative regulation of G1/S transition of mitotic cell cycle | 19/3805 | 0.001722866 | 0.006341203 |
| GO:0003230 | cardiac atrium development | 13/3805 | 0.001739727 | 0.006391962 |
| GO:0007431 | salivary gland development | 13/3805 | 0.001739727 | 0.006391962 |
| GO:0030513 | positive regulation of BMP signaling pathway | 13/3805 | 0.001739727 | 0.006391962 |
| GO:0033044 | regulation of chromosome organization | 43/3805 | 0.001751986 | 0.006433219 |
| GO:1903052 | positive regulation of proteolysis involved in cellular protein catabolic process | 29/3805 | 0.001781771 | 0.006530269 |
| GO:0021783 | preganglionic parasympathetic fiber development | Aug-05 | 0.001785735 | 0.006530269 |
| GO:0032570 | response to progesterone | Aug-05 | 0.001785735 | 0.006530269 |
| GO:0033623 | regulation of integrin activation | Aug-05 | 0.001785735 | 0.006530269 |
| GO:0035313 | wound healing, spreading of epidermal cells | Aug-05 | 0.001785735 | 0.006530269 |
| GO:0086011 | membrane repolarization during action potential | Aug-05 | 0.001785735 | 0.006530269 |
| GO:0090660 | cerebrospinal fluid circulation | Aug-05 | 0.001785735 | 0.006530269 |

| | | | | |
|---|---|---|---|---|
| GO:0030279 | negative regulation of ossification | 14/3805 | 0.001823707 | 0.006661328 |
| GO:1903170 | negative regulation of calcium ion transmembrane transport | 14/3805 | 0.001823707 | 0.006661328 |
| GO:0033003 | regulation of mast cell activation | 17/3805 | 0.001835194 | 0.006691547 |
| GO:0042551 | neuron maturation | 17/3805 | 0.001835194 | 0.006691547 |
| GO:0060324 | face development | 17/3805 | 0.001835194 | 0.006691547 |
| GO:0050670 | regulation of lymphocyte proliferation | 49/3805 | 0.001840784 | 0.006708013 |
| GO:0014855 | striated muscle cell proliferation | 21/3805 | 0.001848792 | 0.006725427 |
| GO:0051279 | regulation of release of sequestered calcium ion into cytosol | 21/3805 | 0.001848792 | 0.006725427 |
| GO:0051303 | establishment of chromosome localization | 21/3805 | 0.001848792 | 0.006725427 |
| GO:0071322 | cellular response to carbohydrate stimulus | 35/3805 | 0.001857172 | 0.006751976 |
| GO:0002791 | regulation of peptide secretion | 48/3805 | 0.00186101 | 0.006759511 |
| GO:0006687 | glycosphingolipid metabolic process | 15/3805 | 0.001861466 | 0.006759511 |
| GO:0071470 | cellular response to osmotic stress | 16/3805 | 0.001862491 | 0.006759511 |
| GO:0000132 | establishment of mitotic spindle orientation | Nov-05 | 0.001874976 | 0.006773336 |
| GO:0003416 | endochondral bone growth | Nov-05 | 0.001874976 | 0.006773336 |
| GO:0021904 | dorsal/ventral neural tube patterning | Nov-05 | 0.001874976 | 0.006773336 |
| GO:0045830 | positive regulation of isotype switching | Nov-05 | 0.001874976 | 0.006773336 |
| GO:0055022 | negative regulation of cardiac muscle tissue growth | Nov-05 | 0.001874976 | 0.006773336 |

| | | | | |
|---|---|---|---|---|
| GO:0060390 | regulation of SMAD protein signal transduction | Nov-05 | 0.001874976 | 0.006773336 |
| GO:0061117 | negative regulation of heart growth | Nov-05 | 0.001874976 | 0.006773336 |
| GO:0098703 | calcium ion import across plasma membrane | Nov-05 | 0.001874976 | 0.006773336 |
| GO:0016573 | histone acetylation | 36/3805 | 0.001883008 | 0.006798421 |
| GO:1905039 | carboxylic acid transmembrane transport | 31/3805 | 0.001910183 | 0.006892551 |
| GO:0045861 | negative regulation of proteolysis | 67/3805 | 0.001917983 | 0.006916698 |
| GO:1903305 | regulation of regulated secretory pathway | 39/3805 | 0.001928815 | 0.006951746 |
| GO:0015749 | monosaccharide transmembrane transport | 28/3805 | 0.001943471 | 0.007000529 |
| GO:0046942 | carboxylic acid transport | 56/3805 | 0.001954421 | 0.007035915 |
| GO:0006665 | sphingolipid metabolic process | 32/3805 | 0.001962147 | 0.007059662 |
| GO:2001251 | negative regulation of chromosome organization | 22/3805 | 0.002035829 | 0.007320545 |
| GO:0015849 | organic acid transport | 61/3805 | 0.002057465 | 0.00739409 |
| GO:0006884 | cell volume homeostasis | Dec-05 | 0.002086495 | 0.007485495 |
| GO:0007435 | salivary gland morphogenesis | Dec-05 | 0.002086495 | 0.007485495 |
| GO:0090075 | relaxation of muscle | Dec-05 | 0.002086495 | 0.007485495 |
| GO:0031497 | chromatin assembly | 36/3805 | 0.002097673 | 0.007516961 |
| GO:0055067 | monovalent inorganic cation homeostasis | 36/3805 | 0.002097673 | 0.007516961 |
| GO:0006643 | membrane lipid metabolic process | 39/3805 | 0.002135505 | 0.007648143 |

| | | | | |
|---|---|---|---|---|
| GO:0002052 | positive regulation of neuroblast proliferation | Oct-05 | 0.00216193 | 0.007711831 |
| GO:0007176 | regulation of epidermal growth factor-activated receptor activity | Oct-05 | 0.00216193 | 0.007711831 |
| GO:0033622 | integrin activation | Oct-05 | 0.00216193 | 0.007711831 |
| GO:0033688 | regulation of osteoblast proliferation | Oct-05 | 0.00216193 | 0.007711831 |
| GO:0038083 | peptidyl-tyrosine autophosphorylation | Oct-05 | 0.00216193 | 0.007711831 |
| GO:0051767 | nitric-oxide synthase biosynthetic process | Oct-05 | 0.00216193 | 0.007711831 |
| GO:0051769 | regulation of nitric-oxide synthase biosynthetic process | Oct-05 | 0.00216193 | 0.007711831 |
| GO:0046928 | regulation of neurotransmitter secretion | 28/3805 | 0.002210359 | 0.007880083 |
| GO:0071174 | mitotic spindle checkpoint signaling | 13/3805 | 0.002219048 | 0.007897533 |
| GO:0098751 | bone cell development | 13/3805 | 0.002219048 | 0.007897533 |
| GO:1900181 | negative regulation of protein localization to nucleus | 13/3805 | 0.002219048 | 0.007897533 |
| GO:0031348 | negative regulation of defense response | 48/3805 | 0.002220726 | 0.007899006 |
| GO:0002067 | glandular epithelial cell differentiation | 17/3805 | 0.002222266 | 0.007899984 |
| GO:0051353 | positive regulation of oxidoreductase activity | 16/3805 | 0.002276748 | 0.008079869 |
| GO:0051489 | regulation of filopodium assembly | 16/3805 | 0.002276748 | 0.008079869 |
| GO:0140353 | lipid export from cell | 16/3805 | 0.002276748 | 0.008079869 |
| GO:0010543 | regulation of platelet activation | 14/3805 | 0.002286206 | 0.008108825 |
| GO:0060043 | regulation of cardiac muscle cell proliferation | 15/3805 | 0.002301439 | 0.008153594 |

| | | | | |
|---|---|---|---|---|
| GO:0061756 | leukocyte adhesion to vascular endothelial cell | 15/3805 | 0.002301439 | 0.008153594 |
| GO:0071482 | cellular response to light stimulus | 26/3805 | 0.002305365 | 0.008162875 |
| GO:0014743 | regulation of muscle hypertrophy | 20/3805 | 0.002308162 | 0.008163521 |
| GO:0045453 | bone resorption | 20/3805 | 0.002308162 | 0.008163521 |
| GO:0010951 | negative regulation of endopeptidase activity | 39/3805 | 0.002361092 | 0.008346 |
| GO:0046323 | glucose import | 22/3805 | 0.002372953 | 0.008376733 |
| GO:0002246 | wound healing involved in inflammatory response | Jun-05 | 0.002389904 | 0.008376733 |
| GO:0007183 | SMAD protein complex assembly | Jun-05 | 0.002389904 | 0.008376733 |
| GO:0016188 | synaptic vesicle maturation | Jun-05 | 0.002389904 | 0.008376733 |
| GO:0021561 | facial nerve development | Jun-05 | 0.002389904 | 0.008376733 |
| GO:0021610 | facial nerve morphogenesis | Jun-05 | 0.002389904 | 0.008376733 |
| GO:0033275 | actin-myosin filament sliding | Jun-05 | 0.002389904 | 0.008376733 |
| GO:0035331 | negative regulation of hippo signaling | Jun-05 | 0.002389904 | 0.008376733 |
| GO:0036005 | response to macrophage colony-stimulating factor | Jun-05 | 0.002389904 | 0.008376733 |
| GO:0036006 | cellular response to macrophage colony-stimulating factor stimulus | Jun-05 | 0.002389904 | 0.008376733 |
| GO:0060430 | lung saccule development | Jun-05 | 0.002389904 | 0.008376733 |
| GO:0070316 | regulation of G0 to G1 transition | Jun-05 | 0.002389904 | 0.008376733 |
| GO:0070586 | cell-cell adhesion involved in gastrulation | Jun-05 | 0.002389904 | 0.008376733 |

| | | | | |
|---|---|---|---|---|
| GO:0090399 | replicative senescence | Jun-05 | 0.002389904 | 0.008376733 |
| GO:1903960 | negative regulation of anion transmembrane transport | Jun-05 | 0.002389904 | 0.008376733 |
| GO:0090087 | regulation of peptide transport | 48/3805 | 0.002422245 | 0.008485329 |
| GO:0002693 | positive regulation of cellular extravasation | Sep-05 | 0.002432772 | 0.008493623 |
| GO:0030903 | notochord development | Sep-05 | 0.002432772 | 0.008493623 |
| GO:0032515 | negative regulation of phosphoprotein phosphatase activity | Sep-05 | 0.002432772 | 0.008493623 |
| GO:0048485 | sympathetic nervous system development | Sep-05 | 0.002432772 | 0.008493623 |
| GO:1902547 | regulation of cellular response to vascular endothelial growth factor stimulus | Sep-05 | 0.002432772 | 0.008493623 |
| GO:1990776 | response to angiotensin | Sep-05 | 0.002432772 | 0.008493623 |
| GO:0060761 | negative regulation of response to cytokine stimulus | 19/3805 | 0.002440394 | 0.008515475 |
| GO:0120161 | regulation of cold-induced thermogenesis | 32/3805 | 0.002473156 | 0.008624976 |
| GO:0030224 | monocyte differentiation | Nov-05 | 0.002480889 | 0.008632665 |
| GO:0044030 | regulation of DNA methylation | Nov-05 | 0.002480889 | 0.008632665 |
| GO:0060669 | embryonic placenta morphogenesis | Nov-05 | 0.002480889 | 0.008632665 |
| GO:1902656 | calcium ion import into cytosol | Nov-05 | 0.002480889 | 0.008632665 |
| GO:0002824 | positive regulation of adaptive immune response based on somatic recombination of immune receptors built from immunoglobulin superfamily domains | 33/3805 | 0.002514728 | 0.008745542 |
| GO:0031507 | heterochromatin assembly | 21/3805 | 0.00254223 | 0.008821539 |
| GO:0043154 | negative regulation of cysteine-type endopeptidase activity involved in apoptotic process | 21/3805 | 0.00254223 | 0.008821539 |

| | | | | |
|---|---|---|---|---|
| GO:0048708 | astrocyte differentiation | 21/3805 | 0.00254223 | 0.008821539 |
| GO:2001259 | positive regulation of cation channel activity | 21/3805 | 0.00254223 | 0.008821539 |
| GO:0000724 | double-strand break repair via homologous recombination | 34/3805 | 0.002547837 | 0.008836086 |
| GO:2000278 | regulation of DNA biosynthetic process | 29/3805 | 0.002590318 | 0.008978428 |
| GO:0002027 | regulation of heart rate | 26/3805 | 0.002632172 | 0.009096112 |
| GO:0002544 | chronic inflammatory response | Aug-05 | 0.002634465 | 0.009096112 |
| GO:0003215 | cardiac right ventricle morphogenesis | Aug-05 | 0.002634465 | 0.009096112 |
| GO:0010766 | negative regulation of sodium ion transport | Aug-05 | 0.002634465 | 0.009096112 |
| GO:0036120 | cellular response to platelet-derived growth factor stimulus | Aug-05 | 0.002634465 | 0.009096112 |
| GO:0072087 | renal vesicle development | Aug-05 | 0.002634465 | 0.009096112 |
| GO:2001135 | regulation of endocytic recycling | Aug-05 | 0.002634465 | 0.009096112 |
| GO:0002467 | germinal center formation | Jul-05 | 0.002672198 | 0.009175673 |
| GO:0019184 | nonribosomal peptide biosynthetic process | Jul-05 | 0.002672198 | 0.009175673 |
| GO:0042048 | olfactory behavior | Jul-05 | 0.002672198 | 0.009175673 |
| GO:0072673 | lamellipodium morphogenesis | Jul-05 | 0.002672198 | 0.009175673 |
| GO:0086103 | G protein-coupled receptor signaling pathway involved in heart process | Jul-05 | 0.002672198 | 0.009175673 |
| GO:1901722 | regulation of cell proliferation involved in kidney development | Jul-05 | 0.002672198 | 0.009175673 |
| GO:1902043 | positive regulation of extrinsic apoptotic signaling pathway via death domain receptors | Jul-05 | 0.002672198 | 0.009175673 |

| | | | | |
|---|---|---|---|---|
| GO:1902287 | semaphorin-plexin signaling pathway involved in axon guidance | Jul-05 | 0.002672198 | 0.009175673 |
| GO:2001212 | regulation of vasculogenesis | Jul-05 | 0.002672198 | 0.009175673 |
| GO:2001256 | regulation of store-operated calcium entry | Jul-05 | 0.002672198 | 0.009175673 |
| GO:0032655 | regulation of interleukin-12 production | 17/3805 | 0.002675238 | 0.009181065 |
| GO:0010664 | negative regulation of striated muscle cell apoptotic process | Dec-05 | 0.002687948 | 0.009194371 |
| GO:0010737 | protein kinase A signaling | Dec-05 | 0.002687948 | 0.009194371 |
| GO:0046329 | negative regulation of JNK cascade | Dec-05 | 0.002687948 | 0.009194371 |
| GO:0070528 | protein kinase C signaling | Dec-05 | 0.002687948 | 0.009194371 |
| GO:0097106 | postsynaptic density organization | Dec-05 | 0.002687948 | 0.009194371 |
| GO:0098868 | bone growth | Dec-05 | 0.002687948 | 0.009194371 |
| GO:0021587 | cerebellum morphogenesis | 16/3805 | 0.002764879 | 0.009442009 |
| GO:0042311 | vasodilation | 16/3805 | 0.002764879 | 0.009442009 |
| GO:0043030 | regulation of macrophage activation | 16/3805 | 0.002764879 | 0.009442009 |
| GO:0098801 | regulation of renal system process | 13/3805 | 0.002801655 | 0.009557149 |
| GO:1902116 | negative regulation of organelle assembly | 13/3805 | 0.002801655 | 0.009557149 |
| GO:0046189 | phenol-containing compound biosynthetic process | 15/3805 | 0.002824172 | 0.00962287 |
| GO:0034205 | amyloid-beta formation | 14/3805 | 0.002841289 | 0.009676495 |
| GO:0048246 | macrophage chemotaxis | 14/3805 | 0.002841289 | 0.009676495 |

| | | | | |
|---|---|---|---|---|
| GO:0003300 | cardiac muscle hypertrophy | 25/3805 | 0.002868013 | 0.009762185 |
| GO:0006475 | internal protein amino acid acetylation | 37/3805 | 0.002879856 | 0.009791823 |
| GO:0018393 | internal peptidyl-lysine acetylation | 37/3805 | 0.002879856 | 0.009791823 |
| GO:0030104 | water homeostasis | 19/3805 | 0.002884064 | 0.009800795 |
| GO:0010954 | positive regulation of protein processing | Oct-05 | 0.002913843 | 0.009885858 |
| GO:0033081 | regulation of T cell differentiation in thymus | Oct-05 | 0.002913843 | 0.009885858 |
| GO:0090330 | regulation of platelet aggregation | Oct-05 | 0.002913843 | 0.009885858 |
| GO:0002698 | negative regulation of immune effector process | 29/3805 | 0.002921554 | 0.009906639 |

**GO terms associated with 5864 Rep_late_enriched_vs_Hep genes**

| ID | Description | Gene Ratio | pvalue | p.adjust |
|---|---|---|---|---|
| GO:0044782 | cilium organization | 193/4924 | 4.80E-50 | 3.02E-46 |
| GO:0007015 | actin filament organization | 209/4924 | 1.78E-46 | 5.63E-43 |
| GO:0060271 | cilium assembly | 175/4924 | 8.91E-45 | 1.87E-41 |
| GO:0032970 | regulation of actin filament-based process | 184/4924 | 4.42E-39 | 6.96E-36 |
| GO:1902903 | regulation of supramolecular fiber organization | 175/4924 | 2.99E-38 | 3.77E-35 |
| GO:0007264 | small GTPase mediated signal transduction | 186/4924 | 2.35E-37 | 2.47E-34 |
| GO:0034329 | cell junction assembly | 187/4924 | 2.70E-36 | 2.43E-33 |
| GO:0007018 | microtubule-based movement | 182/4924 | 7.14E-36 | 5.63E-33 |
| GO:0032956 | regulation of actin cytoskeleton organization | 162/4924 | 2.78E-34 | 1.94E-31 |
| GO:0060562 | epithelial tube morphogenesis | 169/4924 | 1.74E-33 | 1.10E-30 |
| GO:0050808 | synapse organization | 196/4924 | 3.25E-33 | 1.86E-30 |
| GO:0007409 | axonogenesis | 192/4924 | 2.62E-31 | 1.38E-28 |
| GO:0030198 | extracellular matrix organization | 142/4924 | 3.07E-31 | 1.49E-28 |
| GO:0043062 | extracellular structure organization | 142/4924 | 4.65E-31 | 2.09E-28 |
| GO:0045229 | external encapsulating structure organization | 142/4924 | 7.02E-31 | 2.95E-28 |
| GO:0001667 | ameboidal-type cell migration | 178/4924 | 8.43E-30 | 3.32E-27 |

| | | | | |
|---|---|---|---|---|
| GO:0031346 | positive regulation of cell projection organization | 175/4924 | 1.64E-29 | 6.10E-27 |
| GO:0010639 | negative regulation of organelle organization | 155/4924 | 4.08E-29 | 1.43E-26 |
| GO:0007265 | Ras protein signal transduction | 145/4924 | 2.07E-28 | 6.88E-26 |
| GO:0031589 | cell-substrate adhesion | 153/4924 | 5.02E-28 | 1.58E-25 |
| GO:0000280 | nuclear division | 172/4924 | 1.01E-27 | 3.04E-25 |
| GO:0045216 | cell-cell junction organization | 106/4924 | 2.01E-27 | 5.77E-25 |
| GO:0050767 | regulation of neurogenesis | 174/4924 | 2.15E-27 | 5.90E-25 |
| GO:0110053 | regulation of actin filament organization | 123/4924 | 1.12E-26 | 2.94E-24 |
| GO:0061572 | actin filament bundle organization | 92/4924 | 1.38E-26 | 3.48E-24 |
| GO:0001655 | urogenital system development | 154/4924 | 1.61E-26 | 3.91E-24 |
| GO:0007178 | transmembrane receptor protein serine/threonine kinase signaling pathway | 154/4924 | 5.83E-26 | 1.36E-23 |
| GO:0051017 | actin filament bundle assembly | 90/4924 | 7.66E-26 | 1.72E-23 |
| GO:0140014 | mitotic nuclear division | 123/4924 | 8.24E-26 | 1.79E-23 |
| GO:0072001 | renal system development | 141/4924 | 9.60E-26 | 2.02E-23 |
| GO:0001578 | microtubule bundle formation | 73/4924 | 1.25E-25 | 2.55E-23 |
| GO:0043254 | regulation of protein-containing complex assembly | 163/4924 | 2.20E-25 | 4.33E-23 |
| GO:0001822 | kidney development | 136/4924 | 2.71E-25 | 5.17E-23 |
| GO:0032535 | regulation of cellular component size | 153/4924 | 3.51E-25 | 6.51E-23 |

| | | | | |
|---|---|---|---|---|
| GO:0032886 | regulation of microtubule-based process | 118/4924 | 4.34E-25 | 7.83E-23 |
| GO:0045785 | positive regulation of cell adhesion | 176/4924 | 5.40E-25 | 9.46E-23 |
| GO:0048762 | mesenchymal cell differentiation | 110/4924 | 7.09E-25 | 1.21E-22 |
| GO:0099111 | microtubule-based transport | 100/4924 | 7.27E-25 | 1.21E-22 |
| GO:0031032 | actomyosin structure organization | 101/4924 | 1.03E-24 | 1.66E-22 |
| GO:0060485 | mesenchyme development | 126/4924 | 1.41E-24 | 2.22E-22 |
| GO:0051258 | protein polymerization | 119/4924 | 2.54E-24 | 3.90E-22 |
| GO:0043087 | regulation of GTPase activity | 137/4924 | 1.22E-23 | 1.84E-21 |
| GO:0048285 | organelle fission | 177/4924 | 1.32E-23 | 1.93E-21 |
| GO:0051056 | regulation of small GTPase mediated signal transduction | 105/4924 | 1.91E-23 | 2.73E-21 |
| GO:0040013 | negative regulation of locomotion | 135/4924 | 2.62E-23 | 3.67E-21 |
| GO:0007346 | regulation of mitotic cell cycle | 171/4924 | 4.36E-23 | 5.98E-21 |
| GO:0051494 | negative regulation of cytoskeleton organization | 87/4924 | 4.86E-23 | 6.51E-21 |
| GO:0016358 | dendrite development | 128/4924 | 6.15E-23 | 7.91E-21 |
| GO:0090130 | tissue migration | 128/4924 | 6.15E-23 | 7.91E-21 |
| GO:0090287 | regulation of cellular response to growth factor stimulus | 123/4924 | 8.90E-23 | 1.12E-20 |
| GO:0033674 | positive regulation of kinase activity | 159/4924 | 9.32E-23 | 1.15E-20 |
| GO:0090132 | epithelium migration | 127/4924 | 1.06E-22 | 1.29E-20 |

| | | | | |
|---|---|---|---|---|
| GO:0022604 | regulation of cell morphogenesis | 133/4924 | 1.46E-22 | 1.74E-20 |
| GO:0007162 | negative regulation of cell adhesion | 126/4924 | 1.83E-22 | 2.09E-20 |
| GO:0010631 | epithelial cell migration | 126/4924 | 1.83E-22 | 2.09E-20 |
| GO:0001558 | regulation of cell growth | 165/4924 | 1.86E-22 | 2.09E-20 |
| GO:1902904 | negative regulation of supramolecular fiber organization | 86/4924 | 6.07E-22 | 6.72E-20 |
| GO:0007163 | establishment or maintenance of cell polarity | 101/4924 | 9.53E-22 | 1.04E-19 |
| GO:0032271 | regulation of protein polymerization | 95/4924 | 1.11E-21 | 1.19E-19 |
| GO:0198738 | cell-cell signaling by wnt | 161/4924 | 1.38E-21 | 1.45E-19 |
| GO:0016055 | Wnt signaling pathway | 160/4924 | 2.26E-21 | 2.34E-19 |
| GO:0060560 | developmental growth involved in morphogenesis | 115/4924 | 2.60E-21 | 2.63E-19 |
| GO:0001763 | morphogenesis of a branching structure | 106/4924 | 2.63E-21 | 2.63E-19 |
| GO:0060541 | respiratory system development | 111/4924 | 2.93E-21 | 2.89E-19 |
| GO:0022407 | regulation of cell-cell adhesion | 164/4924 | 3.03E-21 | 2.94E-19 |
| GO:0071560 | cellular response to transforming growth factor beta stimulus | 99/4924 | 4.41E-21 | 4.21E-19 |
| GO:0071559 | response to transforming growth factor beta | 100/4924 | 5.45E-21 | 5.13E-19 |
| GO:0045860 | positive regulation of protein kinase activity | 135/4924 | 6.50E-21 | 6.03E-19 |
| GO:0044772 | mitotic cell cycle phase transition | 151/4924 | 7.03E-21 | 6.42E-19 |
| GO:0097485 | neuron projection guidance | 108/4924 | 7.61E-21 | 6.86E-19 |

| | | | | |
|---|---|---|---|---|
| GO:0051271 | negative regulation of cellular component movement | 123/4924 | 8.90E-21 | 7.90E-19 |
| GO:0001654 | eye development | 151/4924 | 9.14E-21 | 8.00E-19 |
| GO:0001837 | epithelial to mesenchymal transition | 77/4924 | 9.99E-21 | 8.63E-19 |
| GO:0007224 | smoothened signaling pathway | 78/4924 | 1.50E-20 | 1.28E-18 |
| GO:0007411 | axon guidance | 107/4924 | 1.89E-20 | 1.59E-18 |
| GO:0150063 | visual system development | 151/4924 | 1.99E-20 | 1.65E-18 |
| GO:0048880 | sensory system development | 152/4924 | 2.02E-20 | 1.66E-18 |
| GO:0051090 | regulation of DNA-binding transcription factor activity | 154/4924 | 2.07E-20 | 1.68E-18 |
| GO:0070507 | regulation of microtubule cytoskeleton organization | 78/4924 | 3.95E-20 | 3.15E-18 |
| GO:0030010 | establishment of cell polarity | 75/4924 | 8.61E-20 | 6.79E-18 |
| GO:0071900 | regulation of protein serine/threonine kinase activity | 131/4924 | 9.21E-20 | 7.17E-18 |
| GO:0090092 | regulation of transmembrane receptor protein serine/threonine kinase signaling pathway | 105/4924 | 1.14E-19 | 8.79E-18 |
| GO:2000146 | negative regulation of cell motility | 118/4924 | 1.75E-19 | 1.33E-17 |
| GO:0007389 | pattern specification process | 165/4924 | 2.20E-19 | 1.65E-17 |
| GO:0030900 | forebrain development | 142/4924 | 2.25E-19 | 1.67E-17 |
| GO:0061138 | morphogenesis of a branching epithelium | 97/4924 | 2.80E-19 | 2.05E-17 |
| GO:0030336 | negative regulation of cell migration | 114/4924 | 3.14E-19 | 2.28E-17 |
| GO:0008154 | actin polymerization or depolymerization | 87/4924 | 8.29E-19 | 5.86E-17 |

| | | | | |
|---|---|---|---|---|
| GO:0030705 | cytoskeleton-dependent intracellular transport | 87/4924 | 8.29E-19 | 5.86E-17 |
| GO:0003007 | heart morphogenesis | 111/4924 | 8.36E-19 | 5.86E-17 |
| GO:0048588 | developmental cell growth | 109/4924 | 9.50E-19 | 6.58E-17 |
| GO:0034765 | regulation of ion transmembrane transport | 165/4924 | 1.32E-18 | 8.95E-17 |
| GO:1901987 | regulation of cell cycle phase transition | 141/4924 | 1.32E-18 | 8.95E-17 |
| GO:0045787 | positive regulation of cell cycle | 129/4924 | 1.76E-18 | 1.18E-16 |
| GO:0007059 | chromosome segregation | 125/4924 | 2.01E-18 | 1.33E-16 |
| GO:0048608 | reproductive structure development | 161/4924 | 2.26E-18 | 1.48E-16 |
| GO:0031109 | microtubule polymerization or depolymerization | 65/4924 | 2.33E-18 | 1.52E-16 |
| GO:0046578 | regulation of Ras protein signal transduction | 87/4924 | 2.66E-18 | 1.70E-16 |
| GO:0031345 | negative regulation of cell projection organization | 91/4924 | 2.67E-18 | 1.70E-16 |
| GO:0051495 | positive regulation of cytoskeleton organization | 88/4924 | 3.24E-18 | 2.04E-16 |
| GO:0007160 | cell-matrix adhesion | 96/4924 | 4.08E-18 | 2.54E-16 |
| GO:0042692 | muscle cell differentiation | 149/4924 | 4.36E-18 | 2.70E-16 |
| GO:0007043 | cell-cell junction assembly | 73/4924 | 4.62E-18 | 2.83E-16 |
| GO:0061458 | reproductive system development | 161/4924 | 5.44E-18 | 3.30E-16 |
| GO:0007179 | transforming growth factor beta receptor signaling pathway | 82/4924 | 6.42E-18 | 3.86E-16 |
| GO:0030323 | respiratory tube development | 97/4924 | 6.55E-18 | 3.89E-16 |

| | | | | |
|---|---|---|---|---|
| GO:0035082 | axoneme assembly | 52/4924 | 6.96E-18 | 4.10E-16 |
| GO:0030324 | lung development | 96/4924 | 8.07E-18 | 4.71E-16 |
| GO:0050678 | regulation of epithelial cell proliferation | 140/4924 | 9.41E-18 | 5.44E-16 |
| GO:0042063 | gliogenesis | 126/4924 | 1.04E-17 | 5.99E-16 |
| GO:0010970 | transport along microtubule | 76/4924 | 1.13E-17 | 6.40E-16 |
| GO:0051261 | protein depolymerization | 63/4924 | 1.26E-17 | 7.07E-16 |
| GO:0010632 | regulation of epithelial cell migration | 97/4924 | 1.28E-17 | 7.13E-16 |
| GO:0007051 | spindle organization | 81/4924 | 1.68E-17 | 9.30E-16 |
| GO:0010721 | negative regulation of cell development | 87/4924 | 1.73E-17 | 9.46E-16 |
| GO:0045926 | negative regulation of growth | 103/4924 | 2.35E-17 | 1.28E-15 |
| GO:0010959 | regulation of metal ion transport | 149/4924 | 2.71E-17 | 1.46E-15 |
| GO:1901342 | regulation of vasculature development | 115/4924 | 2.79E-17 | 1.48E-15 |
| GO:1901990 | regulation of mitotic cell cycle phase transition | 115/4924 | 2.79E-17 | 1.48E-15 |
| GO:1902905 | positive regulation of supramolecular fiber organization | 80/4924 | 2.96E-17 | 1.56E-15 |
| GO:1990778 | protein localization to cell periphery | 130/4924 | 3.11E-17 | 1.62E-15 |
| GO:0043010 | camera-type eye development | 131/4924 | 3.13E-17 | 1.62E-15 |
| GO:0048732 | gland development | 156/4924 | 3.53E-17 | 1.81E-15 |
| GO:0060491 | regulation of cell projection assembly | 87/4924 | 3.56E-17 | 1.81E-15 |

| | | | | |
|---|---|---|---|---|
| GO:009 0068 | positive regulation of cell cycle process | 103/4 924 | 4.33E-17 | 2.19E-15 |
| GO:005 1961 | negative regulation of nervous system development | 75/49 24 | 4.60E-17 | 2.30E-15 |
| GO:001 0810 | regulation of cell-substrate adhesion | 92/49 24 | 5.13E-17 | 2.55E-15 |
| GO:006 1351 | neural precursor cell proliferation | 84/49 24 | 6.39E-17 | 3.15E-15 |
| GO:003 0111 | regulation of Wnt signaling pathway | 116/4 924 | 6.61E-17 | 3.23E-15 |
| GO:006 1448 | connective tissue development | 110/4 924 | 6.78E-17 | 3.29E-15 |
| GO:190 2850 | microtubule cytoskeleton organization involved in mitosis | 69/49 24 | 7.30E-17 | 3.51E-15 |
| GO:004 2060 | wound healing | 130/4 924 | 8.36E-17 | 3.99E-15 |
| GO:005 0673 | epithelial cell proliferation | 155/4 924 | 8.54E-17 | 4.05E-15 |
| GO:001 0976 | positive regulation of neuron projection development | 91/49 24 | 8.85E-17 | 4.17E-15 |
| GO:009 8813 | nuclear chromosome segregation | 105/4 924 | 9.08E-17 | 4.24E-15 |
| GO:000 0819 | sister chromatid segregation | 81/49 24 | 1.13E-16 | 5.25E-15 |
| GO:190 3829 | positive regulation of protein localization | 158/4 924 | 1.44E-16 | 6.59E-15 |
| GO:005 0768 | negative regulation of neurogenesis | 73/49 24 | 1.44E-16 | 6.59E-15 |
| GO:003 1023 | microtubule organizing center organization | 67/49 24 | 1.49E-16 | 6.74E-15 |
| GO:005 1098 | regulation of binding | 135/4 924 | 1.67E-16 | 7.52E-15 |
| GO:014 0694 | non-membrane-bounded organelle assembly | 131/4 924 | 1.72E-16 | 7.70E-15 |
| GO:190 1888 | regulation of cell junction assembly | 89/49 24 | 1.89E-16 | 8.37E-15 |

| | | | | |
|---|---|---|---|---|
| GO:0007611 | learning or memory | 116/4924 | 1.93E-16 | 8.49E-15 |
| GO:0008360 | regulation of cell shape | 71/4924 | 1.98E-16 | 8.68E-15 |
| GO:0010720 | positive regulation of cell development | 130/4924 | 2.19E-16 | 9.54E-15 |
| GO:0045765 | regulation of angiogenesis | 112/4924 | 2.23E-16 | 9.65E-15 |
| GO:0000070 | mitotic sister chromatid segregation | 72/4924 | 2.55E-16 | 1.10E-14 |
| GO:0044839 | cell cycle G2/M phase transition | 69/4924 | 2.69E-16 | 1.14E-14 |
| GO:0051962 | positive regulation of nervous system development | 125/4924 | 2.80E-16 | 1.19E-14 |
| GO:0060070 | canonical Wnt signaling pathway | 111/4924 | 2.83E-16 | 1.19E-14 |
| GO:0098742 | cell-cell adhesion via plasma-membrane adhesion molecules | 87/4924 | 2.87E-16 | 1.19E-14 |
| GO:0042073 | intraciliary transport | 34/4924 | 2.88E-16 | 1.19E-14 |
| GO:0035148 | tube formation | 78/4924 | 2.89E-16 | 1.19E-14 |
| GO:0030308 | negative regulation of cell growth | 84/4924 | 3.78E-16 | 1.55E-14 |
| GO:0050807 | regulation of synapse organization | 100/4924 | 3.83E-16 | 1.55E-14 |
| GO:0030038 | contractile actin filament bundle assembly | 58/4924 | 3.85E-16 | 1.55E-14 |
| GO:0043149 | stress fiber assembly | 58/4924 | 3.85E-16 | 1.55E-14 |
| GO:0030041 | actin filament polymerization | 74/4924 | 4.05E-16 | 1.62E-14 |
| GO:0009799 | specification of symmetry | 66/4924 | 4.13E-16 | 1.64E-14 |
| GO:0050803 | regulation of synapse structure or activity | 102/4924 | 4.34E-16 | 1.71E-14 |

| | | | | |
|---|---|---|---|---|
| GO:0010717 | regulation of epithelial to mesenchymal transition | 51/4924 | 5.07E-16 | 1.97E-14 |
| GO:1901879 | regulation of protein depolymerization | 51/4924 | 5.07E-16 | 1.97E-14 |
| GO:0120032 | regulation of plasma membrane bounded cell projection assembly | 84/4924 | 5.34E-16 | 2.07E-14 |
| GO:0051091 | positive regulation of DNA-binding transcription factor activity | 102/4924 | 5.79E-16 | 2.22E-14 |
| GO:0050890 | cognition | 124/4924 | 7.41E-16 | 2.83E-14 |
| GO:0048754 | branching morphogenesis of an epithelial tube | 80/4924 | 8.47E-16 | 3.22E-14 |
| GO:0016331 | morphogenesis of embryonic epithelium | 78/4924 | 8.75E-16 | 3.29E-14 |
| GO:0050770 | regulation of axonogenesis | 78/4924 | 8.75E-16 | 3.29E-14 |
| GO:0031110 | regulation of microtubule polymerization or depolymerization | 51/4924 | 9.03E-16 | 3.37E-14 |
| GO:0018209 | peptidyl-serine modification | 125/4924 | 9.38E-16 | 3.48E-14 |
| GO:0030832 | regulation of actin filament length | 73/4924 | 1.05E-15 | 3.88E-14 |
| GO:0009855 | determination of bilateral symmetry | 65/4924 | 1.14E-15 | 4.16E-14 |
| GO:0032231 | regulation of actin filament bundle assembly | 57/4924 | 1.14E-15 | 4.16E-14 |
| GO:0070371 | ERK1 and ERK2 cascade | 121/4924 | 1.20E-15 | 4.36E-14 |
| GO:0008064 | regulation of actin polymerization or depolymerization | 72/4924 | 1.26E-15 | 4.53E-14 |
| GO:0007368 | determination of left/right symmetry | 62/4924 | 1.71E-15 | 6.13E-14 |
| GO:0030177 | positive regulation of Wnt signaling pathway | 64/4924 | 2.02E-15 | 7.20E-14 |
| GO:0001838 | embryonic epithelial tube formation | 69/4924 | 2.11E-15 | 7.49E-14 |

| | | | | |
|---|---|---|---|---|
| GO:0002064 | epithelial cell development | 91/4924 | 2.18E-15 | 7.67E-14 |
| GO:0000086 | G2/M transition of mitotic cell cycle | 63/4924 | 2.33E-15 | 8.16E-14 |
| GO:0018108 | peptidyl-tyrosine phosphorylation | 114/4924 | 2.43E-15 | 8.47E-14 |
| GO:0046777 | protein autophosphorylation | 93/4924 | 2.54E-15 | 8.80E-14 |
| GO:0001894 | tissue homeostasis | 104/4924 | 3.34E-15 | 1.15E-13 |
| GO:0001764 | neuron migration | 80/4924 | 3.41E-15 | 1.16E-13 |
| GO:0051656 | establishment of organelle localization | 139/4924 | 3.41E-15 | 1.16E-13 |
| GO:0010563 | negative regulation of phosphorus metabolic process | 146/4924 | 3.99E-15 | 1.35E-13 |
| GO:0045936 | negative regulation of phosphate metabolic process | 146/4924 | 3.99E-15 | 1.35E-13 |
| GO:1990138 | neuron projection extension | 82/4924 | 4.38E-15 | 1.47E-13 |
| GO:0007052 | mitotic spindle organization | 58/4924 | 4.40E-15 | 1.47E-13 |
| GO:0018212 | peptidyl-tyrosine modification | 114/4924 | 5.12E-15 | 1.70E-13 |
| GO:0018105 | peptidyl-serine phosphorylation | 117/4924 | 5.14E-15 | 1.70E-13 |
| GO:0007266 | Rho protein signal transduction | 63/4924 | 5.52E-15 | 1.81E-13 |
| GO:0001503 | ossification | 137/4924 | 5.54E-15 | 1.81E-13 |
| GO:0045786 | negative regulation of cell cycle | 125/4924 | 6.01E-15 | 1.95E-13 |
| GO:2001233 | regulation of apoptotic signaling pathway | 133/4924 | 7.75E-15 | 2.51E-13 |
| GO:0043542 | endothelial cell migration | 88/4924 | 7.86E-15 | 2.53E-13 |

| | | | | |
|---|---|---|---|---|
| GO:0003341 | cilium movement | 85/4924 | 8.27E-15 | 2.65E-13 |
| GO:0007098 | centrosome cycle | 60/4924 | 8.48E-15 | 2.70E-13 |
| GO:0072175 | epithelial tube formation | 71/4924 | 1.01E-14 | 3.19E-13 |
| GO:0070372 | regulation of ERK1 and ERK2 cascade | 113/4924 | 1.06E-14 | 3.35E-13 |
| GO:0071902 | positive regulation of protein serine/threonine kinase activity | 82/4924 | 1.16E-14 | 3.65E-13 |
| GO:0031503 | protein-containing complex localization | 81/4924 | 1.44E-14 | 4.51E-13 |
| GO:0060249 | anatomical structure homeostasis | 113/4924 | 1.72E-14 | 5.35E-13 |
| GO:0048638 | regulation of developmental growth | 131/4924 | 1.93E-14 | 5.97E-13 |
| GO:0021915 | neural tube development | 79/4924 | 2.22E-14 | 6.84E-13 |
| GO:0050769 | positive regulation of neurogenesis | 105/4924 | 2.73E-14 | 8.35E-13 |
| GO:0072659 | protein localization to plasma membrane | 106/4924 | 2.75E-14 | 8.39E-13 |
| GO:0060537 | muscle tissue development | 152/4924 | 3.08E-14 | 9.33E-13 |
| GO:0051403 | stress-activated MAPK cascade | 92/4924 | 3.25E-14 | 9.79E-13 |
| GO:0014020 | primary neural tube formation | 56/4924 | 3.51E-14 | 1.05E-12 |
| GO:0043547 | positive regulation of GTPase activity | 94/4924 | 3.54E-14 | 1.06E-12 |
| GO:0043393 | regulation of protein binding | 84/4924 | 3.55E-14 | 1.06E-12 |
| GO:0001952 | regulation of cell-matrix adhesion | 59/4924 | 3.59E-14 | 1.06E-12 |
| GO:0030833 | regulation of actin filament polymerization | 64/4924 | 3.62E-14 | 1.07E-12 |

| | | | | |
|---|---|---|---|---|
| GO:0003205 | cardiac chamber development | 79/4924 | 4.24E-14 | 1.24E-12 |
| GO:0043405 | regulation of MAP kinase activity | 79/4924 | 4.24E-14 | 1.24E-12 |
| GO:0045930 | negative regulation of mitotic cell cycle | 87/4924 | 4.38E-14 | 1.27E-12 |
| GO:1903706 | regulation of hemopoiesis | 130/4924 | 4.62E-14 | 1.34E-12 |
| GO:0001738 | morphogenesis of a polarized epithelium | 46/4924 | 5.06E-14 | 1.46E-12 |
| GO:0048562 | embryonic organ morphogenesis | 113/4924 | 5.59E-14 | 1.60E-12 |
| GO:0031098 | stress-activated protein kinase signaling cascade | 94/4924 | 6.10E-14 | 1.74E-12 |
| GO:1903844 | regulation of cellular response to transforming growth factor beta stimulus | 57/4924 | 7.40E-14 | 2.10E-12 |
| GO:0051216 | cartilage development | 82/4924 | 7.54E-14 | 2.13E-12 |
| GO:0003012 | muscle system process | 136/4924 | 8.19E-14 | 2.30E-12 |
| GO:0017015 | regulation of transforming growth factor beta receptor signaling pathway | 56/4924 | 8.53E-14 | 2.39E-12 |
| GO:0050900 | leukocyte migration | 124/4924 | 8.56E-14 | 2.39E-12 |
| GO:0042391 | regulation of membrane potential | 145/4924 | 9.42E-14 | 2.62E-12 |
| GO:0031333 | negative regulation of protein-containing complex assembly | 65/4924 | 9.52E-14 | 2.62E-12 |
| GO:0072073 | kidney epithelium development | 65/4924 | 9.52E-14 | 2.62E-12 |
| GO:0010634 | positive regulation of epithelial cell migration | 66/4924 | 1.14E-13 | 3.13E-12 |
| GO:0001843 | neural tube closure | 53/4924 | 1.27E-13 | 3.47E-12 |
| GO:0006260 | DNA replication | 98/4924 | 1.47E-13 | 3.98E-12 |

| | | | | |
|---|---|---|---|---|
| GO:0030048 | actin filament-based movement | 58/4924 | 1.48E-13 | 3.99E-12 |
| GO:1901880 | negative regulation of protein depolymerization | 43/4924 | 1.65E-13 | 4.44E-12 |
| GO:0042326 | negative regulation of phosphorylation | 128/4924 | 1.71E-13 | 4.58E-12 |
| GO:0022409 | positive regulation of cell-cell adhesion | 103/4924 | 1.94E-13 | 5.17E-12 |
| GO:0010389 | regulation of G2/M transition of mitotic cell cycle | 49/4924 | 1.97E-13 | 5.25E-12 |
| GO:0060606 | tube closure | 53/4924 | 2.00E-13 | 5.29E-12 |
| GO:0006816 | calcium ion transport | 137/4924 | 2.50E-13 | 6.59E-12 |
| GO:0043242 | negative regulation of protein-containing complex disassembly | 46/4924 | 2.52E-13 | 6.60E-12 |
| GO:1904062 | regulation of cation transmembrane transport | 123/4924 | 2.52E-13 | 6.60E-12 |
| GO:0001841 | neural tube formation | 59/4924 | 2.81E-13 | 7.31E-12 |
| GO:0051146 | striated muscle cell differentiation | 110/4924 | 2.89E-13 | 7.50E-12 |
| GO:1905515 | non-motile cilium assembly | 41/4924 | 2.95E-13 | 7.62E-12 |
| GO:0032102 | negative regulation of response to external stimulus | 128/4924 | 3.12E-13 | 8.03E-12 |
| GO:0010971 | positive regulation of G2/M transition of mitotic cell cycle | 23/4924 | 3.52E-13 | 9.03E-12 |
| GO:0110020 | regulation of actomyosin structure organization | 52/4924 | 3.56E-13 | 9.05E-12 |
| GO:1902749 | regulation of cell cycle G2/M phase transition | 52/4924 | 3.56E-13 | 9.05E-12 |
| GO:0007159 | leukocyte cell-cell adhesion | 123/4924 | 3.81E-13 | 9.64E-12 |
| GO:1903034 | regulation of response to wounding | 70/4924 | 4.08E-13 | 1.03E-11 |

| | | | | |
|---|---|---|---|---|
| GO:0021537 | telencephalon development | 92/4924 | 4.78E-13 | 1.20E-11 |
| GO:0006936 | muscle contraction | 106/4924 | 4.85E-13 | 1.21E-11 |
| GO:0044458 | motile cilium assembly | 38/4924 | 4.92E-13 | 1.22E-11 |
| GO:0022408 | negative regulation of cell-cell adhesion | 79/4924 | 4.92E-13 | 1.22E-11 |
| GO:0050730 | regulation of peptidyl-tyrosine phosphorylation | 94/4924 | 4.93E-13 | 1.22E-11 |
| GO:0043244 | regulation of protein-containing complex disassembly | 58/4924 | 4.95E-13 | 1.22E-11 |
| GO:0030856 | regulation of epithelial cell differentiation | 65/4924 | 5.82E-13 | 1.42E-11 |
| GO:0072006 | nephron development | 65/4924 | 5.82E-13 | 1.42E-11 |
| GO:0035567 | non-canonical Wnt signaling pathway | 34/4924 | 6.27E-13 | 1.53E-11 |
| GO:1902751 | positive regulation of cell cycle G2/M phase transition | 24/4924 | 9.28E-13 | 2.25E-11 |
| GO:0008361 | regulation of cell size | 80/4924 | 9.32E-13 | 2.25E-11 |
| GO:0010977 | negative regulation of neuron projection development | 66/4924 | 9.46E-13 | 2.28E-11 |
| GO:0097191 | extrinsic apoptotic signaling pathway | 87/4924 | 1.21E-12 | 2.91E-11 |
| GO:0002685 | regulation of leukocyte migration | 83/4924 | 1.41E-12 | 3.36E-11 |
| GO:0001933 | negative regulation of protein phosphorylation | 114/4924 | 1.48E-12 | 3.51E-11 |
| GO:0061387 | regulation of extent of cell growth | 56/4924 | 1.54E-12 | 3.65E-11 |
| GO:0010001 | glial cell differentiation | 91/4924 | 1.68E-12 | 3.94E-11 |
| GO:0033002 | muscle cell proliferation | 91/4924 | 1.68E-12 | 3.94E-11 |

| | | | | |
|---|---|---|---|---|
| GO:0032386 | regulation of intracellular transport | 110/4924 | 2.07E-12 | 4.84E-11 |
| GO:1903131 | mononuclear cell differentiation | 147/4924 | 2.07E-12 | 4.84E-11 |
| GO:0048813 | dendrite morphogenesis | 72/4924 | 2.41E-12 | 5.61E-11 |
| GO:0051492 | regulation of stress fiber assembly | 47/4924 | 2.61E-12 | 6.06E-11 |
| GO:2000027 | regulation of animal organ morphogenesis | 57/4924 | 2.76E-12 | 6.38E-11 |
| GO:0050920 | regulation of chemotaxis | 83/4924 | 3.14E-12 | 7.24E-11 |
| GO:0048017 | inositol lipid-mediated signaling | 68/4924 | 3.17E-12 | 7.26E-11 |
| GO:0043406 | positive regulation of MAP kinase activity | 56/4924 | 3.33E-12 | 7.60E-11 |
| GO:0014706 | striated muscle tissue development | 98/4924 | 3.38E-12 | 7.69E-11 |
| GO:0030516 | regulation of axon extension | 51/4924 | 3.55E-12 | 8.06E-11 |
| GO:1902105 | regulation of leukocyte differentiation | 106/4924 | 3.57E-12 | 8.06E-11 |
| GO:0007019 | microtubule depolymerization | 32/4924 | 3.85E-12 | 8.67E-11 |
| GO:0048015 | phosphatidylinositol-mediated signaling | 67/4924 | 3.95E-12 | 8.85E-11 |
| GO:0008544 | epidermis development | 115/4924 | 3.96E-12 | 8.85E-11 |
| GO:0051054 | positive regulation of DNA metabolic process | 100/4924 | 4.10E-12 | 9.14E-11 |
| GO:0070997 | neuron death | 133/4924 | 4.13E-12 | 9.16E-11 |
| GO:0003206 | cardiac chamber morphogenesis | 62/4924 | 4.29E-12 | 9.48E-11 |
| GO:0010594 | regulation of endothelial cell migration | 68/4924 | 4.34E-12 | 9.53E-11 |

| | | | | |
|---|---|---|---|---|
| GO:0032272 | negative regulation of protein polymerization | 42/4924 | 4.34E-12 | 9.53E-11 |
| GO:0007416 | synapse assembly | 73/4924 | 4.69E-12 | 1.03E-10 |
| GO:0007422 | peripheral nervous system development | 39/4924 | 5.16E-12 | 1.13E-10 |
| GO:0120193 | tight junction organization | 45/4924 | 5.26E-12 | 1.14E-10 |
| GO:0090257 | regulation of muscle system process | 89/4924 | 5.70E-12 | 1.23E-10 |
| GO:0010948 | negative regulation of cell cycle process | 97/4924 | 6.79E-12 | 1.47E-10 |
| GO:0043588 | skin development | 105/4924 | 6.97E-12 | 1.50E-10 |
| GO:0060828 | regulation of canonical Wnt signaling pathway | 87/4924 | 7.22E-12 | 1.55E-10 |
| GO:0090101 | negative regulation of transmembrane receptor protein serine/threonine kinase signaling pathway | 55/4924 | 8.49E-12 | 1.82E-10 |
| GO:0048660 | regulation of smooth muscle cell proliferation | 69/4924 | 8.68E-12 | 1.85E-10 |
| GO:0090596 | sensory organ morphogenesis | 104/4924 | 8.88E-12 | 1.89E-10 |
| GO:0030098 | lymphocyte differentiation | 134/4924 | 9.39E-12 | 1.99E-10 |
| GO:0099173 | postsynapse organization | 76/4924 | 9.60E-12 | 2.03E-10 |
| GO:1903555 | regulation of tumor necrosis factor superfamily cytokine production | 71/4924 | 9.93E-12 | 2.09E-10 |
| GO:0060840 | artery development | 51/4924 | 1.20E-11 | 2.50E-10 |
| GO:0009314 | response to radiation | 130/4924 | 1.20E-11 | 2.50E-10 |
| GO:0010927 | cellular component assembly involved in morphogenesis | 53/4924 | 1.23E-11 | 2.57E-10 |
| GO:0022411 | cellular component disassembly | 129/4924 | 1.27E-11 | 2.64E-10 |

| | | | | |
|---|---|---|---|---|
| GO:0006261 | DNA-templated DNA replication | 60/4924 | 1.28E-11 | 2.65E-10 |
| GO:0098657 | import into cell | 84/4924 | 1.49E-11 | 3.06E-10 |
| GO:0003002 | regionalization | 119/4924 | 1.55E-11 | 3.18E-10 |
| GO:0031099 | regeneration | 62/4924 | 1.62E-11 | 3.31E-10 |
| GO:0050679 | positive regulation of epithelial cell proliferation | 78/4924 | 1.75E-11 | 3.57E-10 |
| GO:1903046 | meiotic cell cycle process | 79/4924 | 1.79E-11 | 3.64E-10 |
| GO:0071706 | tumor necrosis factor superfamily cytokine production | 72/4924 | 1.86E-11 | 3.77E-10 |
| GO:0048675 | axon extension | 58/4924 | 1.94E-11 | 3.93E-10 |
| GO:0050863 | regulation of T cell activation | 110/4924 | 1.97E-11 | 3.97E-10 |
| GO:0002028 | regulation of sodium ion transport | 45/4924 | 2.04E-11 | 4.10E-10 |
| GO:0031334 | positive regulation of protein-containing complex assembly | 75/4924 | 2.09E-11 | 4.19E-10 |
| GO:0071674 | mononuclear cell migration | 69/4924 | 2.10E-11 | 4.20E-10 |
| GO:0022612 | gland morphogenesis | 62/4924 | 2.23E-11 | 4.39E-10 |
| GO:0051092 | positive regulation of NF-kappaB transcription factor activity | 62/4924 | 2.23E-11 | 4.39E-10 |
| GO:0071772 | response to BMP | 70/4924 | 2.23E-11 | 4.39E-10 |
| GO:0071773 | cellular response to BMP stimulus | 70/4924 | 2.23E-11 | 4.39E-10 |
| GO:0098739 | import across plasma membrane | 71/4924 | 2.35E-11 | 4.61E-10 |
| GO:0030509 | BMP signaling pathway | 67/4924 | 2.48E-11 | 4.86E-10 |

| | | | | |
|---|---|---|---|---|
| GO:0030512 | negative regulation of transforming growth factor beta receptor signaling pathway | 38/4924 | 2.69E-11 | 5.25E-10 |
| GO:0032872 | regulation of stress-activated MAPK cascade | 75/4924 | 2.74E-11 | 5.32E-10 |
| GO:0006887 | exocytosis | 125/4924 | 2.74E-11 | 5.32E-10 |
| GO:0090263 | positive regulation of canonical Wnt signaling pathway | 48/4924 | 2.98E-11 | 5.76E-10 |
| GO:0032984 | protein-containing complex disassembly | 83/4924 | 3.09E-11 | 5.96E-10 |
| GO:0002062 | chondrocyte differentiation | 50/4924 | 3.12E-11 | 6.00E-10 |
| GO:1905475 | regulation of protein localization to membrane | 73/4924 | 3.36E-11 | 6.44E-10 |
| GO:0007156 | homophilic cell adhesion via plasma membrane adhesion molecules | 49/4924 | 3.73E-11 | 7.09E-10 |
| GO:2000181 | negative regulation of blood vessel morphogenesis | 49/4924 | 3.73E-11 | 7.09E-10 |
| GO:0032680 | regulation of tumor necrosis factor production | 69/4924 | 3.74E-11 | 7.09E-10 |
| GO:1903037 | regulation of leukocyte cell-cell adhesion | 108/4924 | 3.88E-11 | 7.34E-10 |
| GO:0006814 | sodium ion transport | 85/4924 | 3.92E-11 | 7.40E-10 |
| GO:0046847 | filopodium assembly | 38/4924 | 4.49E-11 | 8.45E-10 |
| GO:0048678 | response to axon injury | 37/4924 | 4.82E-11 | 9.02E-10 |
| GO:0061512 | protein localization to cilium | 37/4924 | 4.82E-11 | 9.02E-10 |
| GO:0032409 | regulation of transporter activity | 97/4924 | 4.85E-11 | 9.04E-10 |
| GO:0046785 | microtubule polymerization | 41/4924 | 5.39E-11 | 1.00E-09 |
| GO:1901343 | negative regulation of vasculature development | 49/4924 | 5.50E-11 | 1.02E-09 |

| | | | | |
|---|---|---|---|---|
| GO:0007044 | cell-substrate junction assembly | 44/4924 | 5.61E-11 | 1.04E-09 |
| GO:0070302 | regulation of stress-activated protein kinase signaling cascade | 75/4924 | 6.03E-11 | 1.11E-09 |
| GO:0003015 | heart process | 85/4924 | 6.27E-11 | 1.15E-09 |
| GO:0061640 | cytoskeleton-dependent cytokinesis | 43/4924 | 6.49E-11 | 1.19E-09 |
| GO:0032640 | tumor necrosis factor production | 70/4924 | 6.82E-11 | 1.25E-09 |
| GO:0048640 | negative regulation of developmental growth | 53/4924 | 7.65E-11 | 1.39E-09 |
| GO:0048661 | positive regulation of smooth muscle cell proliferation | 47/4924 | 7.84E-11 | 1.42E-09 |
| GO:0043583 | ear development | 85/4924 | 7.90E-11 | 1.43E-09 |
| GO:0050727 | regulation of inflammatory response | 109/4924 | 7.98E-11 | 1.44E-09 |
| GO:1901988 | negative regulation of cell cycle phase transition | 84/4924 | 7.99E-11 | 1.44E-09 |
| GO:0007219 | Notch signaling pathway | 68/4924 | 8.27E-11 | 1.49E-09 |
| GO:0022898 | regulation of transmembrane transporter activity | 93/4924 | 8.36E-11 | 1.50E-09 |
| GO:0048659 | smooth muscle cell proliferation | 69/4924 | 8.62E-11 | 1.54E-09 |
| GO:0050731 | positive regulation of peptidyl-tyrosine phosphorylation | 70/4924 | 8.95E-11 | 1.59E-09 |
| GO:0001736 | establishment of planar polarity | 33/4924 | 9.74E-11 | 1.73E-09 |
| GO:0090175 | regulation of establishment of planar polarity | 26/4924 | 1.02E-10 | 1.80E-09 |
| GO:0045137 | development of primary sexual characteristics | 83/4924 | 1.02E-10 | 1.80E-09 |
| GO:0008406 | gonad development | 82/4924 | 1.03E-10 | 1.81E-09 |

| | | | | |
|---|---|---|---|---|
| GO:0043297 | apical junction assembly | 42/4924 | 1.16E-10 | 2.03E-09 |
| GO:0062197 | cellular response to chemical stress | 105/4924 | 1.17E-10 | 2.06E-09 |
| GO:2001236 | regulation of extrinsic apoptotic signaling pathway | 64/4924 | 1.19E-10 | 2.08E-09 |
| GO:1901214 | regulation of neuron death | 118/4924 | 1.20E-10 | 2.09E-09 |
| GO:1901215 | negative regulation of neuron death | 87/4924 | 1.20E-10 | 2.09E-09 |
| GO:0061035 | regulation of cartilage development | 36/4924 | 1.45E-10 | 2.51E-09 |
| GO:0014910 | regulation of smooth muscle cell migration | 43/4924 | 1.52E-10 | 2.63E-09 |
| GO:0060996 | dendritic spine development | 53/4924 | 1.53E-10 | 2.63E-09 |
| GO:0051924 | regulation of calcium ion transport | 93/4924 | 1.58E-10 | 2.71E-09 |
| GO:2001234 | negative regulation of apoptotic signaling pathway | 83/4924 | 1.61E-10 | 2.77E-09 |
| GO:0150115 | cell-substrate junction organization | 45/4924 | 1.65E-10 | 2.81E-09 |
| GO:0021700 | developmental maturation | 109/4924 | 1.68E-10 | 2.87E-09 |
| GO:0010595 | positive regulation of endothelial cell migration | 47/4924 | 1.70E-10 | 2.89E-09 |
| GO:0007164 | establishment of tissue polarity | 33/4924 | 1.71E-10 | 2.90E-09 |
| GO:0006310 | DNA recombination | 101/4924 | 1.72E-10 | 2.90E-09 |
| GO:1903035 | negative regulation of response to wounding | 42/4924 | 1.78E-10 | 3.00E-09 |
| GO:2000177 | regulation of neural precursor cell proliferation | 50/4924 | 1.99E-10 | 3.35E-09 |
| GO:0014909 | smooth muscle cell migration | 46/4924 | 2.04E-10 | 3.42E-09 |

| | | | | |
|---|---|---|---|---|
| GO:0085029 | extracellular matrix assembly | 26/4924 | 2.15E-10 | 3.60E-09 |
| GO:0035721 | intraciliary retrograde transport | 14/4924 | 2.25E-10 | 3.75E-09 |
| GO:0007613 | memory | 61/4924 | 2.37E-10 | 3.95E-09 |
| GO:0007088 | regulation of mitotic nuclear division | 47/4924 | 2.48E-10 | 4.10E-09 |
| GO:0016525 | negative regulation of angiogenesis | 47/4924 | 2.48E-10 | 4.10E-09 |
| GO:0050771 | negative regulation of axonogenesis | 35/4924 | 2.60E-10 | 4.29E-09 |
| GO:0032874 | positive regulation of stress-activated MAPK cascade | 55/4924 | 2.68E-10 | 4.41E-09 |
| GO:0031111 | negative regulation of microtubule polymerization or depolymerization | 28/4924 | 2.80E-10 | 4.60E-09 |
| GO:0010935 | regulation of macrophage cytokine production | 15/4924 | 2.92E-10 | 4.78E-09 |
| GO:0010934 | macrophage cytokine production | 16/4924 | 2.97E-10 | 4.86E-09 |
| GO:0045664 | regulation of neuron differentiation | 81/4924 | 3.30E-10 | 5.37E-09 |
| GO:0030100 | regulation of endocytosis | 80/4924 | 3.34E-10 | 5.43E-09 |
| GO:0050866 | negative regulation of cell activation | 77/4924 | 3.44E-10 | 5.58E-09 |
| GO:0071675 | regulation of mononuclear cell migration | 49/4924 | 3.47E-10 | 5.60E-09 |
| GO:1903522 | regulation of blood circulation | 87/4924 | 3.58E-10 | 5.78E-09 |
| GO:0060348 | bone development | 85/4924 | 3.78E-10 | 6.09E-09 |
| GO:0000910 | cytokinesis | 66/4924 | 3.95E-10 | 6.34E-09 |
| GO:0008589 | regulation of smoothened signaling pathway | 42/4924 | 4.09E-10 | 6.54E-09 |

| | | | | |
|---|---|---|---|---|
| GO:1901991 | negative regulation of mitotic cell cycle phase transition | 63/4924 | 4.65E-10 | 7.43E-09 |
| GO:0030837 | negative regulation of actin filament polymerization | 34/4924 | 4.66E-10 | 7.43E-09 |
| GO:0035023 | regulation of Rho protein signal transduction | 41/4924 | 4.81E-10 | 7.63E-09 |
| GO:0032835 | glomerulus development | 33/4924 | 5.01E-10 | 7.94E-09 |
| GO:0070304 | positive regulation of stress-activated protein kinase signaling cascade | 55/4924 | 5.04E-10 | 7.97E-09 |
| GO:0070252 | actin-mediated cell contraction | 43/4924 | 5.14E-10 | 8.11E-09 |
| GO:0048144 | fibroblast proliferation | 47/4924 | 5.16E-10 | 8.12E-09 |
| GO:0007612 | learning | 67/4924 | 5.28E-10 | 8.28E-09 |
| GO:0034332 | adherens junction organization | 28/4924 | 5.33E-10 | 8.34E-09 |
| GO:0007528 | neuromuscular junction development | 31/4924 | 5.50E-10 | 8.57E-09 |
| GO:0030199 | collagen fibril organization | 31/4924 | 5.50E-10 | 8.57E-09 |
| GO:0048839 | inner ear development | 75/4924 | 5.59E-10 | 8.68E-09 |
| GO:0006937 | regulation of muscle contraction | 61/4924 | 5.62E-10 | 8.71E-09 |
| GO:0002687 | positive regulation of leukocyte migration | 59/4924 | 6.73E-10 | 1.04E-08 |
| GO:0014812 | muscle cell migration | 49/4924 | 6.94E-10 | 1.07E-08 |
| GO:0035725 | sodium ion transmembrane transport | 57/4924 | 7.96E-10 | 1.22E-08 |
| GO:0032412 | regulation of ion transmembrane transporter activity | 88/4924 | 7.99E-10 | 1.23E-08 |
| GO:0051225 | spindle assembly | 48/4924 | 8.54E-10 | 1.31E-08 |

| | | | | |
|---|---|---|---|---|
| GO:0044843 | cell cycle G1/S phase transition | 78/4924 | 8.55E-10 | 1.31E-08 |
| GO:0045055 | regulated exocytosis | 91/4924 | 8.68E-10 | 1.32E-08 |
| GO:0051099 | positive regulation of binding | 68/4924 | 8.96E-10 | 1.36E-08 |
| GO:0006022 | aminoglycan metabolic process | 46/4924 | 9.01E-10 | 1.37E-08 |
| GO:0045619 | regulation of lymphocyte differentiation | 70/4924 | 9.09E-10 | 1.38E-08 |
| GO:0051783 | regulation of nuclear division | 55/4924 | 9.31E-10 | 1.40E-08 |
| GO:0032103 | positive regulation of response to external stimulus | 130/4924 | 9.91E-10 | 1.49E-08 |
| GO:0007517 | muscle organ development | 109/4924 | 9.98E-10 | 1.50E-08 |
| GO:0060047 | heart contraction | 79/4924 | 1.05E-09 | 1.57E-08 |
| GO:0031532 | actin cytoskeleton reorganization | 47/4924 | 1.05E-09 | 1.57E-08 |
| GO:0006941 | striated muscle contraction | 63/4924 | 1.05E-09 | 1.57E-08 |
| GO:0060071 | Wnt signaling pathway, planar cell polarity pathway | 23/4924 | 1.10E-09 | 1.62E-08 |
| GO:0070286 | axonemal dynein complex assembly | 23/4924 | 1.10E-09 | 1.62E-08 |
| GO:0001657 | ureteric bud development | 45/4924 | 1.10E-09 | 1.62E-08 |
| GO:0072163 | mesonephric epithelium development | 45/4924 | 1.10E-09 | 1.62E-08 |
| GO:0072164 | mesonephric tubule development | 45/4924 | 1.10E-09 | 1.62E-08 |
| GO:0051402 | neuron apoptotic process | 98/4924 | 1.11E-09 | 1.62E-08 |
| GO:0140013 | meiotic nuclear division | 70/4924 | 1.16E-09 | 1.70E-08 |

| | | | | |
|---|---|---|---|---|
| GO:0030217 | T cell differentiation | 94/4924 | 1.36E-09 | 1.98E-08 |
| GO:0014911 | positive regulation of smooth muscle cell migration | 32/4924 | 1.51E-09 | 2.20E-08 |
| GO:1903039 | positive regulation of leukocyte cell-cell adhesion | 81/4924 | 1.53E-09 | 2.23E-08 |
| GO:0061326 | renal tubule development | 45/4924 | 1.57E-09 | 2.28E-08 |
| GO:0030099 | myeloid cell differentiation | 125/4924 | 1.67E-09 | 2.42E-08 |
| GO:0003279 | cardiac septum development | 51/4924 | 1.67E-09 | 2.42E-08 |
| GO:0055001 | muscle cell development | 76/4924 | 1.74E-09 | 2.51E-08 |
| GO:2001222 | regulation of neuron migration | 29/4924 | 1.77E-09 | 2.55E-08 |
| GO:0045931 | positive regulation of mitotic cell cycle | 52/4924 | 1.83E-09 | 2.63E-08 |
| GO:0014044 | Schwann cell development | 22/4924 | 1.87E-09 | 2.67E-08 |
| GO:0048738 | cardiac muscle tissue development | 86/4924 | 1.94E-09 | 2.78E-08 |
| GO:0043523 | regulation of neuron apoptotic process | 89/4924 | 2.08E-09 | 2.97E-08 |
| GO:0007249 | I-kappaB kinase/NF-kappaB signaling | 76/4924 | 2.17E-09 | 3.09E-08 |
| GO:0001818 | negative regulation of cytokine production | 92/4924 | 2.19E-09 | 3.11E-08 |
| GO:0007292 | female gamete generation | 61/4924 | 2.21E-09 | 3.13E-08 |
| GO:0048145 | regulation of fibroblast proliferation | 45/4924 | 2.24E-09 | 3.16E-08 |
| GO:0120192 | tight junction assembly | 39/4924 | 2.27E-09 | 3.21E-08 |
| GO:0031114 | regulation of microtubule depolymerization | 23/4924 | 2.32E-09 | 3.27E-08 |

| | | | | |
|---|---|---|---|---|
| GO:0008016 | regulation of heart contraction | 68/4924 | 2.40E-09 | 3.37E-08 |
| GO:0031102 | neuron projection regeneration | 31/4924 | 2.71E-09 | 3.79E-08 |
| GO:0001938 | positive regulation of endothelial cell proliferation | 42/4924 | 2.84E-09 | 3.97E-08 |
| GO:0018210 | peptidyl-threonine modification | 50/4924 | 2.86E-09 | 3.99E-08 |
| GO:0072009 | nephron epithelium development | 47/4924 | 2.90E-09 | 4.04E-08 |
| GO:0031113 | regulation of microtubule polymerization | 30/4924 | 2.91E-09 | 4.04E-08 |
| GO:0002274 | myeloid leukocyte activation | 85/4924 | 3.02E-09 | 4.19E-08 |
| GO:0043524 | negative regulation of neuron apoptotic process | 66/4924 | 3.05E-09 | 4.22E-08 |
| GO:0045444 | fat cell differentiation | 84/4924 | 3.15E-09 | 4.34E-08 |
| GO:0006270 | DNA replication initiation | 21/4924 | 3.16E-09 | 4.34E-08 |
| GO:0001823 | mesonephros development | 45/4924 | 3.16E-09 | 4.34E-08 |
| GO:0050921 | positive regulation of chemotaxis | 56/4924 | 3.17E-09 | 4.35E-08 |
| GO:1901989 | positive regulation of cell cycle phase transition | 48/4924 | 3.24E-09 | 4.43E-08 |
| GO:0003018 | vascular process in circulatory system | 77/4924 | 3.27E-09 | 4.46E-08 |
| GO:0060993 | kidney morphogenesis | 43/4924 | 3.35E-09 | 4.56E-08 |
| GO:0014065 | phosphatidylinositol 3-kinase signaling | 52/4924 | 3.36E-09 | 4.56E-08 |
| GO:0002040 | sprouting angiogenesis | 49/4924 | 3.57E-09 | 4.84E-08 |
| GO:0010811 | positive regulation of cell-substrate adhesion | 53/4924 | 3.59E-09 | 4.86E-08 |

| | | | | |
|---|---|---|---|---|
| GO:0050878 | regulation of body fluid levels | 109/4924 | 3.82E-09 | 5.15E-08 |
| GO:0048705 | skeletal system morphogenesis | 83/4924 | 4.01E-09 | 5.40E-08 |
| GO:0048286 | lung alveolus development | 32/4924 | 4.03E-09 | 5.41E-08 |
| GO:0006874 | cellular calcium ion homeostasis | 137/4924 | 4.33E-09 | 5.81E-08 |
| GO:0048259 | regulation of receptor-mediated endocytosis | 49/4924 | 4.88E-09 | 6.53E-08 |
| GO:2001020 | regulation of response to DNA damage stimulus | 83/4924 | 4.90E-09 | 6.54E-08 |
| GO:0007254 | JNK cascade | 63/4924 | 4.92E-09 | 6.56E-08 |
| GO:0045927 | positive regulation of growth | 97/4924 | 4.97E-09 | 6.61E-08 |
| GO:0051293 | establishment of spindle localization | 27/4924 | 5.71E-09 | 7.58E-08 |
| GO:1902115 | regulation of organelle assembly | 71/4924 | 5.86E-09 | 7.76E-08 |
| GO:0051348 | negative regulation of transferase activity | 83/4924 | 5.97E-09 | 7.88E-08 |
| GO:0030203 | glycosaminoglycan metabolic process | 40/4924 | 6.03E-09 | 7.94E-08 |
| GO:0032273 | positive regulation of protein polymerization | 40/4924 | 6.03E-09 | 7.94E-08 |
| GO:1905477 | positive regulation of protein localization to membrane | 45/4924 | 6.20E-09 | 8.13E-08 |
| GO:0050680 | negative regulation of epithelial cell proliferation | 61/4924 | 6.21E-09 | 8.13E-08 |
| GO:0009100 | glycoprotein metabolic process | 99/4924 | 6.22E-09 | 8.14E-08 |
| GO:0043271 | negative regulation of ion transport | 62/4924 | 6.27E-09 | 8.18E-08 |
| GO:1903532 | positive regulation of secretion by cell | 106/4924 | 6.49E-09 | 8.45E-08 |

| | | | | |
|---|---|---|---|---|
| GO:0043534 | blood vessel endothelial cell migration | 46/4924 | 6.91E-09 | 8.99E-08 |
| GO:0001649 | osteoblast differentiation | 73/4924 | 6.96E-09 | 9.03E-08 |
| GO:0007229 | integrin-mediated signaling pathway | 41/4924 | 7.14E-09 | 9.24E-08 |
| GO:0018107 | peptidyl-threonine phosphorylation | 47/4924 | 7.62E-09 | 9.85E-08 |
| GO:0035107 | appendage morphogenesis | 61/4924 | 7.97E-09 | 1.03E-07 |
| GO:0035108 | limb morphogenesis | 61/4924 | 7.97E-09 | 1.03E-07 |
| GO:0002011 | morphogenesis of an epithelial sheet | 30/4924 | 8.02E-09 | 1.03E-07 |
| GO:0001890 | placenta development | 62/4924 | 8.02E-09 | 1.03E-07 |
| GO:0070374 | positive regulation of ERK1 and ERK2 cascade | 74/4924 | 8.38E-09 | 1.07E-07 |
| GO:0048844 | artery morphogenesis | 36/4924 | 8.47E-09 | 1.08E-07 |
| GO:0048146 | positive regulation of fibroblast proliferation | 33/4924 | 8.88E-09 | 1.13E-07 |
| GO:0071214 | cellular response to abiotic stimulus | 98/4924 | 9.33E-09 | 1.18E-07 |
| GO:0104004 | cellular response to environmental stimulus | 98/4924 | 9.33E-09 | 1.18E-07 |
| GO:0060048 | cardiac muscle contraction | 50/4924 | 9.60E-09 | 1.22E-07 |
| GO:0034599 | cellular response to oxidative stress | 85/4924 | 9.68E-09 | 1.22E-07 |
| GO:0048771 | tissue remodeling | 68/4924 | 9.70E-09 | 1.22E-07 |
| GO:0003351 | epithelial cilium movement involved in extracellular fluid movement | 24/4924 | 1.00E-08 | 1.26E-07 |
| GO:0045580 | regulation of T cell differentiation | 59/4924 | 1.00E-08 | 1.26E-07 |

| | | | | |
|---|---|---|---|---|
| GO:0001539 | cilium or flagellum-dependent cell motility | 60/4924 | 1.01E-08 | 1.27E-07 |
| GO:0060285 | cilium-dependent cell motility | 60/4924 | 1.01E-08 | 1.27E-07 |
| GO:1901992 | positive regulation of mitotic cell cycle phase transition | 41/4924 | 1.02E-08 | 1.27E-07 |
| GO:0032233 | positive regulation of actin filament bundle assembly | 32/4924 | 1.02E-08 | 1.27E-07 |
| GO:0007093 | mitotic cell cycle checkpoint signaling | 51/4924 | 1.02E-08 | 1.27E-07 |
| GO:0007369 | gastrulation | 62/4924 | 1.02E-08 | 1.27E-07 |
| GO:0045862 | positive regulation of proteolysis | 104/4924 | 1.03E-08 | 1.27E-07 |
| GO:0006858 | extracellular transport | 25/4924 | 1.03E-08 | 1.27E-07 |
| GO:0090100 | positive regulation of transmembrane receptor protein serine/threonine kinase signaling pathway | 47/4924 | 1.04E-08 | 1.28E-07 |
| GO:1902414 | protein localization to cell junction | 47/4924 | 1.04E-08 | 1.28E-07 |
| GO:0034446 | substrate adhesion-dependent cell spreading | 44/4924 | 1.07E-08 | 1.31E-07 |
| GO:0022011 | myelination in peripheral nervous system | 20/4924 | 1.19E-08 | 1.45E-07 |
| GO:0032292 | peripheral nervous system axon ensheathment | 20/4924 | 1.19E-08 | 1.45E-07 |
| GO:0009913 | epidermal cell differentiation | 76/4924 | 1.19E-08 | 1.45E-07 |
| GO:0044344 | cellular response to fibroblast growth factor stimulus | 40/4924 | 1.25E-08 | 1.52E-07 |
| GO:0061041 | regulation of wound healing | 50/4924 | 1.28E-08 | 1.56E-07 |
| GO:0006302 | double-strand break repair | 83/4924 | 1.29E-08 | 1.57E-07 |
| GO:0051058 | negative regulation of small GTPase mediated signal transduction | 30/4924 | 1.30E-08 | 1.58E-07 |

| | | | | |
|---|---|---|---|---|
| GO:0051383 | kinetochore organization | 16/4924 | 1.36E-08 | 1.64E-07 |
| GO:0021782 | glial cell development | 47/4924 | 1.41E-08 | 1.71E-07 |
| GO:0007026 | negative regulation of microtubule depolymerization | 21/4924 | 1.45E-08 | 1.75E-07 |
| GO:1904035 | regulation of epithelial cell apoptotic process | 39/4924 | 1.52E-08 | 1.83E-07 |
| GO:0030834 | regulation of actin filament depolymerization | 28/4924 | 1.58E-08 | 1.89E-07 |
| GO:0030858 | positive regulation of epithelial cell differentiation | 28/4924 | 1.58E-08 | 1.89E-07 |
| GO:0051653 | spindle localization | 28/4924 | 1.58E-08 | 1.89E-07 |
| GO:0002367 | cytokine production involved in immune response | 45/4924 | 1.63E-08 | 1.94E-07 |
| GO:0120034 | positive regulation of plasma membrane bounded cell projection assembly | 45/4924 | 1.63E-08 | 1.94E-07 |
| GO:0030510 | regulation of BMP signaling pathway | 42/4924 | 1.64E-08 | 1.95E-07 |
| GO:0090288 | negative regulation of cellular response to growth factor stimulus | 42/4924 | 1.64E-08 | 1.95E-07 |
| GO:0030866 | cortical actin cytoskeleton organization | 22/4924 | 1.65E-08 | 1.96E-07 |
| GO:0044786 | cell cycle DNA replication | 22/4924 | 1.65E-08 | 1.96E-07 |
| GO:0001570 | vasculogenesis | 40/4924 | 1.77E-08 | 2.09E-07 |
| GO:0046546 | development of primary male sexual characteristics | 52/4924 | 1.87E-08 | 2.20E-07 |
| GO:0009416 | response to light stimulus | 95/4924 | 1.88E-08 | 2.21E-07 |
| GO:0007548 | sex differentiation | 94/4924 | 2.01E-08 | 2.35E-07 |
| GO:0050708 | regulation of protein secretion | 91/4924 | 2.04E-08 | 2.39E-07 |

| | | | | |
|---|---|---|---|---|
| GO:0007009 | plasma membrane organization | 56/4924 | 2.07E-08 | 2.42E-07 |
| GO:0048592 | eye morphogenesis | 63/4924 | 2.07E-08 | 2.42E-07 |
| GO:0030865 | cortical cytoskeleton organization | 30/4924 | 2.07E-08 | 2.42E-07 |
| GO:0051592 | response to calcium ion | 49/4924 | 2.16E-08 | 2.51E-07 |
| GO:0051983 | regulation of chromosome segregation | 39/4924 | 2.17E-08 | 2.52E-07 |
| GO:0030042 | actin filament depolymerization | 29/4924 | 2.33E-08 | 2.71E-07 |
| GO:0071774 | response to fibroblast growth factor | 40/4924 | 2.50E-08 | 2.89E-07 |
| GO:0033260 | nuclear DNA replication | 20/4924 | 2.53E-08 | 2.92E-07 |
| GO:0060326 | cell chemotaxis | 93/4924 | 2.53E-08 | 2.92E-07 |
| GO:0043122 | regulation of I-kappaB kinase/NF-kappaB signaling | 64/4924 | 2.57E-08 | 2.95E-07 |
| GO:0032388 | positive regulation of intracellular transport | 63/4924 | 2.61E-08 | 2.99E-07 |
| GO:0016570 | histone modification | 133/4924 | 2.63E-08 | 3.01E-07 |
| GO:0001935 | endothelial cell proliferation | 62/4924 | 2.64E-08 | 3.02E-07 |
| GO:0060976 | coronary vasculature development | 34/4924 | 2.66E-08 | 3.04E-07 |
| GO:1903707 | negative regulation of hemopoiesis | 44/4924 | 2.78E-08 | 3.17E-07 |
| GO:0072080 | nephron tubule development | 41/4924 | 2.83E-08 | 3.20E-07 |
| GO:0000281 | mitotic cytokinesis | 31/4924 | 2.84E-08 | 3.20E-07 |
| GO:0031103 | axon regeneration | 27/4924 | 2.84E-08 | 3.20E-07 |

| | | | | |
|---|---|---|---|---|
| GO:0032330 | regulation of chondrocyte differentiation | 27/4924 | 2.84E-08 | 3.20E-07 |
| GO:0035088 | establishment or maintenance of apical/basal cell polarity | 27/4924 | 2.84E-08 | 3.20E-07 |
| GO:0061245 | establishment or maintenance of bipolar cell polarity | 27/4924 | 2.84E-08 | 3.20E-07 |
| GO:0002697 | regulation of immune effector process | 120/4924 | 2.94E-08 | 3.31E-07 |
| GO:0048736 | appendage development | 67/4924 | 2.99E-08 | 3.35E-07 |
| GO:0060173 | limb development | 67/4924 | 2.99E-08 | 3.35E-07 |
| GO:0008277 | regulation of G protein-coupled receptor signaling pathway | 50/4924 | 3.00E-08 | 3.35E-07 |
| GO:0034763 | negative regulation of transmembrane transport | 51/4924 | 3.11E-08 | 3.47E-07 |
| GO:0006942 | regulation of striated muscle contraction | 37/4924 | 3.25E-08 | 3.62E-07 |
| GO:0014037 | Schwann cell differentiation | 23/4924 | 3.31E-08 | 3.68E-07 |
| GO:0043270 | positive regulation of ion transport | 96/4924 | 3.41E-08 | 3.79E-07 |
| GO:0086005 | ventricular cardiac muscle cell action potential | 18/4924 | 3.41E-08 | 3.79E-07 |
| GO:0016311 | dephosphorylation | 100/4924 | 3.55E-08 | 3.94E-07 |
| GO:0045620 | negative regulation of lymphocyte differentiation | 29/4924 | 3.72E-08 | 4.11E-07 |
| GO:0017156 | calcium-ion regulated exocytosis | 32/4924 | 3.72E-08 | 4.11E-07 |
| GO:0055117 | regulation of cardiac muscle contraction | 32/4924 | 3.72E-08 | 4.11E-07 |
| GO:0035050 | embryonic heart tube development | 38/4924 | 3.79E-08 | 4.17E-07 |
| GO:0071692 | protein localization to extracellular region | 113/4924 | 3.84E-08 | 4.22E-07 |

| | | | | |
|---|---|---|---|---|
| GO:1901652 | response to peptide | 116/4924 | 4.04E-08 | 4.43E-07 |
| GO:0008584 | male gonad development | 51/4924 | 4.06E-08 | 4.45E-07 |
| GO:0043409 | negative regulation of MAPK cascade | 63/4924 | 4.10E-08 | 4.48E-07 |
| GO:0052547 | regulation of peptidase activity | 121/4924 | 4.10E-08 | 4.48E-07 |
| GO:0001895 | retina homeostasis | 28/4924 | 4.20E-08 | 4.57E-07 |
| GO:0060041 | retina development in camera-type eye | 61/4924 | 4.24E-08 | 4.61E-07 |
| GO:0032612 | interleukin-1 production | 46/4924 | 4.34E-08 | 4.71E-07 |
| GO:0061082 | myeloid leukocyte cytokine production | 19/4924 | 4.38E-08 | 4.74E-07 |
| GO:0071711 | basement membrane organization | 19/4924 | 4.38E-08 | 4.74E-07 |
| GO:1901653 | cellular response to peptide | 92/4924 | 4.47E-08 | 4.82E-07 |
| GO:0071695 | anatomical structure maturation | 89/4924 | 4.60E-08 | 4.96E-07 |
| GO:0002263 | cell activation involved in immune response | 96/4924 | 4.71E-08 | 5.07E-07 |
| GO:0002695 | negative regulation of leukocyte activation | 66/4924 | 4.73E-08 | 5.08E-07 |
| GO:1902106 | negative regulation of leukocyte differentiation | 43/4924 | 4.74E-08 | 5.08E-07 |
| GO:0070830 | bicellular tight junction assembly | 35/4924 | 4.79E-08 | 5.13E-07 |
| GO:0050870 | positive regulation of T cell activation | 71/4924 | 4.87E-08 | 5.20E-07 |
| GO:0048863 | stem cell differentiation | 75/4924 | 4.96E-08 | 5.30E-07 |
| GO:0048008 | platelet-derived growth factor receptor signaling pathway | 30/4924 | 5.08E-08 | 5.41E-07 |

| | | | | |
|---|---|---|---|---|
| GO:0021591 | ventricular system development | 20/4924 | 5.14E-08 | 5.47E-07 |
| GO:0034764 | positive regulation of transmembrane transport | 78/4924 | 5.16E-08 | 5.48E-07 |
| GO:0007405 | neuroblast proliferation | 32/4924 | 5.61E-08 | 5.93E-07 |
| GO:0045428 | regulation of nitric oxide biosynthetic process | 32/4924 | 5.61E-08 | 5.93E-07 |
| GO:0048041 | focal adhesion assembly | 36/4924 | 5.68E-08 | 6.00E-07 |
| GO:0003014 | renal system process | 46/4924 | 5.79E-08 | 6.09E-07 |
| GO:0042542 | response to hydrogen peroxide | 46/4924 | 5.79E-08 | 6.09E-07 |
| GO:0045494 | photoreceptor cell maintenance | 24/4924 | 5.80E-08 | 6.10E-07 |
| GO:0006979 | response to oxidative stress | 113/4924 | 5.89E-08 | 6.18E-07 |
| GO:0002718 | regulation of cytokine production involved in immune response | 39/4924 | 6.06E-08 | 6.35E-07 |
| GO:0048639 | positive regulation of developmental growth | 70/4924 | 6.20E-08 | 6.48E-07 |
| GO:1903557 | positive regulation of tumor necrosis factor superfamily cytokine production | 43/4924 | 6.43E-08 | 6.71E-07 |
| GO:0055123 | digestive system development | 50/4924 | 6.73E-08 | 7.01E-07 |
| GO:0098659 | inorganic cation import across plasma membrane | 44/4924 | 6.89E-08 | 7.15E-07 |
| GO:0099587 | inorganic ion import across plasma membrane | 44/4924 | 6.89E-08 | 7.15E-07 |
| GO:0000075 | cell cycle checkpoint signaling | 59/4924 | 6.90E-08 | 7.16E-07 |
| GO:0033044 | regulation of chromosome organization | 67/4924 | 6.96E-08 | 7.20E-07 |
| GO:0080164 | regulation of nitric oxide metabolic process | 33/4924 | 6.97E-08 | 7.20E-07 |

| | | | | |
|---|---|---|---|---|
| GO:0006820 | anion transport | 132/4924 | 6.99E-08 | 7.21E-07 |
| GO:0001936 | regulation of endothelial cell proliferation | 57/4924 | 7.06E-08 | 7.26E-07 |
| GO:0003231 | cardiac ventricle development | 53/4924 | 7.06E-08 | 7.26E-07 |
| GO:0072698 | protein localization to microtubule cytoskeleton | 27/4924 | 7.55E-08 | 7.75E-07 |
| GO:0051222 | positive regulation of protein transport | 93/4924 | 8.00E-08 | 8.20E-07 |
| GO:0045807 | positive regulation of endocytosis | 47/4924 | 8.02E-08 | 8.21E-07 |
| GO:0060411 | cardiac septum morphogenesis | 36/4924 | 8.10E-08 | 8.28E-07 |
| GO:0017157 | regulation of exocytosis | 76/4924 | 8.32E-08 | 8.49E-07 |
| GO:0031214 | biomineral tissue development | 59/4924 | 8.66E-08 | 8.82E-07 |
| GO:0010038 | response to metal ion | 87/4924 | 8.68E-08 | 8.82E-07 |
| GO:0009914 | hormone transport | 106/4924 | 8.83E-08 | 8.97E-07 |
| GO:0046328 | regulation of JNK cascade | 53/4924 | 9.05E-08 | 9.16E-07 |
| GO:0000725 | recombinational repair | 55/4924 | 9.05E-08 | 9.16E-07 |
| GO:0045744 | negative regulation of G protein-coupled receptor signaling pathway | 29/4924 | 9.08E-08 | 9.18E-07 |
| GO:0001656 | metanephros development | 37/4924 | 9.29E-08 | 9.36E-07 |
| GO:0060395 | SMAD protein signal transduction | 40/4924 | 9.29E-08 | 9.36E-07 |
| GO:0052548 | regulation of endopeptidase activity | 101/4924 | 9.65E-08 | 9.71E-07 |
| GO:0051963 | regulation of synapse assembly | 45/4924 | 9.72E-08 | 9.74E-07 |

| | | | | |
|---|---|---|---|---|
| GO:1904019 | epithelial cell apoptotic process | 45/4924 | 9.72E-08 | 9.74E-07 |
| GO:0009306 | protein secretion | 110/4924 | 9.90E-08 | 9.90E-07 |
| GO:0086003 | cardiac muscle cell contraction | 31/4924 | 9.96E-08 | 9.95E-07 |
| GO:0090162 | establishment of epithelial cell polarity | 20/4924 | 1.01E-07 | 1.00E-06 |
| GO:0002090 | regulation of receptor internalization | 33/4924 | 1.02E-07 | 1.02E-06 |
| GO:0002366 | leukocyte activation involved in immune response | 94/4924 | 1.02E-07 | 1.02E-06 |
| GO:0110148 | biomineralization | 61/4924 | 1.04E-07 | 1.03E-06 |
| GO:0050919 | negative chemotaxis | 23/4924 | 1.05E-07 | 1.04E-06 |
| GO:0045685 | regulation of glial cell differentiation | 38/4924 | 1.05E-07 | 1.04E-06 |
| GO:0035924 | cellular response to vascular endothelial growth factor stimulus | 28/4924 | 1.05E-07 | 1.04E-06 |
| GO:0044380 | protein localization to cytoskeleton | 28/4924 | 1.05E-07 | 1.04E-06 |
| GO:0014013 | regulation of gliogenesis | 47/4924 | 1.05E-07 | 1.04E-06 |
| GO:0060674 | placenta blood vessel development | 21/4924 | 1.06E-07 | 1.04E-06 |
| GO:0033157 | regulation of intracellular protein transport | 71/4924 | 1.07E-07 | 1.05E-06 |
| GO:0003158 | endothelium development | 48/4924 | 1.09E-07 | 1.06E-06 |
| GO:0000082 | G1/S transition of mitotic cell cycle | 66/4924 | 1.09E-07 | 1.07E-06 |
| GO:0032760 | positive regulation of tumor necrosis factor production | 42/4924 | 1.09E-07 | 1.07E-06 |
| GO:0035592 | establishment of protein localization to extracellular region | 110/4924 | 1.14E-07 | 1.11E-06 |

| | | | | |
|---|---|---|---|---|
| GO:2001235 | positive regulation of apoptotic signaling pathway | 54/4924 | 1.15E-07 | 1.12E-06 |
| GO:1902305 | regulation of sodium ion transmembrane transport | 30/4924 | 1.18E-07 | 1.15E-06 |
| GO:0032964 | collagen biosynthetic process | 27/4924 | 1.20E-07 | 1.17E-06 |
| GO:0008543 | fibroblast growth factor receptor signaling pathway | 34/4924 | 1.21E-07 | 1.18E-06 |
| GO:1903510 | mucopolysaccharide metabolic process | 32/4924 | 1.23E-07 | 1.19E-06 |
| GO:0014066 | regulation of phosphatidylinositol 3-kinase signaling | 40/4924 | 1.27E-07 | 1.23E-06 |
| GO:0071897 | DNA biosynthetic process | 60/4924 | 1.32E-07 | 1.28E-06 |
| GO:0009612 | response to mechanical stimulus | 59/4924 | 1.35E-07 | 1.31E-06 |
| GO:0008347 | glial cell migration | 29/4924 | 1.39E-07 | 1.34E-06 |
| GO:0051310 | metaphase plate congression | 29/4924 | 1.39E-07 | 1.34E-06 |
| GO:0051250 | negative regulation of lymphocyte activation | 57/4924 | 1.41E-07 | 1.35E-06 |
| GO:0032675 | regulation of interleukin-6 production | 56/4924 | 1.43E-07 | 1.37E-06 |
| GO:0002683 | negative regulation of immune system process | 120/4924 | 1.47E-07 | 1.41E-06 |
| GO:0009101 | glycoprotein biosynthetic process | 81/4924 | 1.51E-07 | 1.45E-06 |
| GO:0045332 | phospholipid translocation | 25/4924 | 1.52E-07 | 1.45E-06 |
| GO:0032092 | positive regulation of protein binding | 39/4924 | 1.59E-07 | 1.51E-06 |
| GO:0051346 | negative regulation of hydrolase activity | 102/4924 | 1.61E-07 | 1.53E-06 |
| GO:0061333 | renal tubule morphogenesis | 36/4924 | 1.61E-07 | 1.53E-06 |

| | | | | |
|---|---|---|---|---|
| GO:0002573 | myeloid leukocyte differentiation | 74/4924 | 1.61E-07 | 1.53E-06 |
| GO:0019221 | cytokine-mediated signaling pathway | 114/4924 | 1.63E-07 | 1.54E-06 |
| GO:0045766 | positive regulation of angiogenesis | 60/4924 | 1.64E-07 | 1.55E-06 |
| GO:1904018 | positive regulation of vasculature development | 60/4924 | 1.64E-07 | 1.55E-06 |
| GO:0010718 | positive regulation of epithelial to mesenchymal transition | 24/4924 | 1.66E-07 | 1.57E-06 |
| GO:0030517 | negative regulation of axon extension | 24/4924 | 1.66E-07 | 1.57E-06 |
| GO:0090307 | mitotic spindle assembly | 30/4924 | 1.77E-07 | 1.66E-06 |
| GO:0051047 | positive regulation of secretion | 113/4924 | 1.77E-07 | 1.66E-06 |
| GO:0040036 | regulation of fibroblast growth factor receptor signaling pathway | 19/4924 | 1.79E-07 | 1.68E-06 |
| GO:0051298 | centrosome duplication | 32/4924 | 1.80E-07 | 1.68E-06 |
| GO:0042490 | mechanoreceptor differentiation | 37/4924 | 1.80E-07 | 1.68E-06 |
| GO:0043535 | regulation of blood vessel endothelial cell migration | 37/4924 | 1.80E-07 | 1.68E-06 |
| GO:0060294 | cilium movement involved in cell motility | 55/4924 | 1.83E-07 | 1.70E-06 |
| GO:0048593 | camera-type eye morphogenesis | 53/4924 | 1.87E-07 | 1.74E-06 |
| GO:0120316 | sperm flagellum assembly | 20/4924 | 1.90E-07 | 1.76E-06 |
| GO:1904645 | response to amyloid-beta | 21/4924 | 1.93E-07 | 1.79E-06 |
| GO:0032652 | regulation of interleukin-1 production | 43/4924 | 2.06E-07 | 1.91E-06 |
| GO:0071277 | cellular response to calcium ion | 33/4924 | 2.12E-07 | 1.96E-06 |

| | | | | |
|---|---|---|---|---|
| GO:0070661 | leukocyte proliferation | 102/4924 | 2.14E-07 | 1.98E-06 |
| GO:0008306 | associative learning | 44/4924 | 2.15E-07 | 1.98E-06 |
| GO:0046580 | negative regulation of Ras protein signal transduction | 26/4924 | 2.16E-07 | 1.99E-06 |
| GO:0050772 | positive regulation of axonogenesis | 39/4924 | 2.17E-07 | 1.99E-06 |
| GO:0061162 | establishment of monopolar cell polarity | 16/4924 | 2.21E-07 | 2.02E-06 |
| GO:1990845 | adaptive thermogenesis | 55/4924 | 2.30E-07 | 2.10E-06 |
| GO:2001237 | negative regulation of extrinsic apoptotic signaling pathway | 40/4924 | 2.33E-07 | 2.13E-06 |
| GO:0034767 | positive regulation of ion transmembrane transport | 62/4924 | 2.34E-07 | 2.14E-06 |
| GO:1904951 | positive regulation of establishment of protein localization | 95/4924 | 2.38E-07 | 2.17E-06 |
| GO:0006898 | receptor-mediated endocytosis | 79/4924 | 2.42E-07 | 2.20E-06 |
| GO:0043281 | regulation of cysteine-type endopeptidase activity involved in apoptotic process | 66/4924 | 2.42E-07 | 2.20E-06 |
| GO:0045839 | negative regulation of mitotic nuclear division | 25/4924 | 2.45E-07 | 2.22E-06 |
| GO:2000116 | regulation of cysteine-type endopeptidase activity | 73/4924 | 2.45E-07 | 2.22E-06 |
| GO:0010171 | body morphogenesis | 28/4924 | 2.48E-07 | 2.25E-06 |
| GO:0051899 | membrane depolarization | 37/4924 | 2.49E-07 | 2.25E-06 |
| GO:1905818 | regulation of chromosome separation | 32/4924 | 2.60E-07 | 2.34E-06 |
| GO:0060425 | lung morphogenesis | 30/4924 | 2.62E-07 | 2.36E-06 |
| GO:0002700 | regulation of production of molecular mediator of immune response | 58/4924 | 2.67E-07 | 2.41E-06 |

| | | | | |
|---|---|---|---|---|
| GO:0006939 | smooth muscle contraction | 43/4924 | 2.72E-07 | 2.45E-06 |
| GO:0038127 | ERBB signaling pathway | 43/4924 | 2.72E-07 | 2.45E-06 |
| GO:0071526 | semaphorin-plexin signaling pathway | 24/4924 | 2.73E-07 | 2.45E-06 |
| GO:0050905 | neuromuscular process | 63/4924 | 2.76E-07 | 2.47E-06 |
| GO:0072028 | nephron morphogenesis | 35/4924 | 2.78E-07 | 2.49E-06 |
| GO:0051321 | meiotic cell cycle | 94/4924 | 2.97E-07 | 2.66E-06 |
| GO:0051145 | smooth muscle cell differentiation | 33/4924 | 3.03E-07 | 2.70E-06 |
| GO:0060688 | regulation of morphogenesis of a branching structure | 29/4924 | 3.14E-07 | 2.79E-06 |
| GO:0098900 | regulation of action potential | 29/4924 | 3.14E-07 | 2.79E-06 |
| GO:1905330 | regulation of morphogenesis of an epithelium | 31/4924 | 3.16E-07 | 2.81E-06 |
| GO:0007288 | sperm axoneme assembly | 18/4924 | 3.18E-07 | 2.81E-06 |
| GO:0042634 | regulation of hair cycle | 18/4924 | 3.18E-07 | 2.81E-06 |
| GO:0061037 | negative regulation of cartilage development | 18/4924 | 3.18E-07 | 2.81E-06 |
| GO:0086002 | cardiac muscle cell action potential involved in contraction | 22/4924 | 3.24E-07 | 2.86E-06 |
| GO:2000649 | regulation of sodium ion transmembrane transporter activity | 26/4924 | 3.37E-07 | 2.98E-06 |
| GO:0090504 | epiboly | 19/4924 | 3.40E-07 | 3.00E-06 |
| GO:0006809 | nitric oxide biosynthetic process | 34/4924 | 3.45E-07 | 3.03E-06 |
| GO:0034109 | homotypic cell-cell adhesion | 34/4924 | 3.45E-07 | 3.03E-06 |

| | | | | |
|---|---|---|---|---|
| GO:0050000 | chromosome localization | 34/4924 | 3.45E-07 | 3.03E-06 |
| GO:0035329 | hippo signaling | 20/4924 | 3.47E-07 | 3.03E-06 |
| GO:0048846 | axon extension involved in axon guidance | 20/4924 | 3.47E-07 | 3.03E-06 |
| GO:0060441 | epithelial tube branching involved in lung morphogenesis | 20/4924 | 3.47E-07 | 3.03E-06 |
| GO:1902284 | neuron projection extension involved in neuron projection guidance | 20/4924 | 3.47E-07 | 3.03E-06 |
| GO:0035089 | establishment of apical/basal cell polarity | 15/4924 | 3.72E-07 | 3.24E-06 |
| GO:0000724 | double-strand break repair via homologous recombination | 53/4924 | 3.74E-07 | 3.25E-06 |
| GO:0046879 | hormone secretion | 102/4924 | 3.75E-07 | 3.26E-06 |
| GO:1902017 | regulation of cilium assembly | 30/4924 | 3.83E-07 | 3.32E-06 |
| GO:2001251 | negative regulation of chromosome organization | 35/4924 | 3.87E-07 | 3.35E-06 |
| GO:0000018 | regulation of DNA recombination | 47/4924 | 3.87E-07 | 3.35E-06 |
| GO:0034204 | lipid translocation | 25/4924 | 3.88E-07 | 3.35E-06 |
| GO:0099565 | chemical synaptic transmission, postsynaptic | 39/4924 | 3.94E-07 | 3.40E-06 |
| GO:0070588 | calcium ion transmembrane transport | 86/4924 | 3.97E-07 | 3.42E-06 |
| GO:0070663 | regulation of leukocyte proliferation | 79/4924 | 4.00E-07 | 3.44E-06 |
| GO:0019932 | second-messenger-mediated signaling | 88/4924 | 4.01E-07 | 3.44E-06 |
| GO:0035265 | organ growth | 67/4924 | 4.09E-07 | 3.51E-06 |
| GO:0032635 | interleukin-6 production | 57/4924 | 4.22E-07 | 3.61E-06 |

| | | | | |
|---|---|---|---|---|
| GO:004 2552 | myelination | 57/49 24 | 4.22E-07 | 3.61E-06 |
| GO:190 2806 | regulation of cell cycle G1/S phase transition | 56/49 24 | 4.35E-07 | 3.71E-06 |
| GO:190 3053 | regulation of extracellular matrix organization | 24/49 24 | 4.41E-07 | 3.76E-06 |
| GO:004 2098 | T cell proliferation | 69/49 24 | 4.42E-07 | 3.76E-06 |
| GO:003 0307 | positive regulation of cell growth | 65/49 24 | 4.53E-07 | 3.85E-06 |
| GO:015 0076 | neuroinflammatory response | 29/49 24 | 4.63E-07 | 3.93E-06 |
| GO:000 1659 | temperature homeostasis | 60/49 24 | 4.66E-07 | 3.96E-06 |
| GO:190 3828 | negative regulation of protein localization | 68/49 24 | 4.67E-07 | 3.96E-06 |
| GO:007 2088 | nephron epithelium morphogenesis | 34/49 24 | 4.81E-07 | 4.07E-06 |
| GO:006 1339 | establishment or maintenance of monopolar cell polarity | 16/49 24 | 4.82E-07 | 4.07E-06 |
| GO:000 1508 | action potential | 50/49 24 | 4.91E-07 | 4.14E-06 |
| GO:005 1480 | regulation of cytosolic calcium ion concentration | 107/4 924 | 4.91E-07 | 4.14E-06 |
| GO:003 0835 | negative regulation of actin filament depolymerization | 23/49 24 | 4.93E-07 | 4.15E-06 |
| GO:003 0326 | embryonic limb morphogenesis | 49/49 24 | 4.95E-07 | 4.15E-06 |
| GO:003 5113 | embryonic appendage morphogenesis | 49/49 24 | 4.95E-07 | 4.15E-06 |
| GO:003 3627 | cell adhesion mediated by integrin | 35/49 24 | 5.33E-07 | 4.47E-06 |
| GO:000 7596 | blood coagulation | 56/49 24 | 5.38E-07 | 4.51E-06 |
| GO:009 8534 | centriole assembly | 22/49 24 | 5.43E-07 | 4.54E-06 |

| | | | | |
|---|---|---|---|---|
| GO:0072676 | lymphocyte migration | 40/4924 | 5.54E-07 | 4.63E-06 |
| GO:0030239 | myofibril assembly | 30/4924 | 5.55E-07 | 4.63E-06 |
| GO:0002819 | regulation of adaptive immune response | 71/4924 | 5.63E-07 | 4.69E-06 |
| GO:0021987 | cerebral cortex development | 41/4924 | 5.77E-07 | 4.80E-06 |
| GO:0070301 | cellular response to hydrogen peroxide | 36/4924 | 5.82E-07 | 4.83E-06 |
| GO:0007099 | centriole replication | 21/4924 | 5.85E-07 | 4.84E-06 |
| GO:0071604 | transforming growth factor beta production | 21/4924 | 5.85E-07 | 4.84E-06 |
| GO:1902692 | regulation of neuroblast proliferation | 21/4924 | 5.85E-07 | 4.84E-06 |
| GO:0034260 | negative regulation of GTPase activity | 18/4924 | 6.09E-07 | 5.04E-06 |
| GO:0031112 | positive regulation of microtubule polymerization or depolymerization | 20/4924 | 6.15E-07 | 5.07E-06 |
| GO:0099003 | vesicle-mediated transport in synapse | 75/4924 | 6.16E-07 | 5.08E-06 |
| GO:0051797 | regulation of hair follicle development | 14/4924 | 6.19E-07 | 5.09E-06 |
| GO:0002224 | toll-like receptor signaling pathway | 44/4924 | 6.21E-07 | 5.10E-06 |
| GO:0032963 | collagen metabolic process | 44/4924 | 6.21E-07 | 5.10E-06 |
| GO:0030282 | bone mineralization | 45/4924 | 6.29E-07 | 5.15E-06 |
| GO:0061982 | meiosis I cell cycle process | 47/4924 | 6.33E-07 | 5.19E-06 |
| GO:0098727 | maintenance of cell number | 57/4924 | 6.42E-07 | 5.25E-06 |
| GO:0031644 | regulation of nervous system process | 61/4924 | 6.65E-07 | 5.42E-06 |

| | | | | |
|---|---|---|---|---|
| GO:0032147 | activation of protein kinase activity | 38/4924 | 6.66E-07 | 5.42E-06 |
| GO:0032611 | interleukin-1 beta production | 38/4924 | 6.66E-07 | 5.42E-06 |
| GO:0090630 | activation of GTPase activity | 38/4924 | 6.66E-07 | 5.42E-06 |
| GO:0071248 | cellular response to metal ion | 55/4924 | 6.87E-07 | 5.58E-06 |
| GO:0046330 | positive regulation of JNK cascade | 39/4924 | 7.01E-07 | 5.69E-06 |
| GO:0046209 | nitric oxide metabolic process | 35/4924 | 7.30E-07 | 5.91E-06 |
| GO:0032755 | positive regulation of interleukin-6 production | 40/4924 | 7.31E-07 | 5.92E-06 |
| GO:0006469 | negative regulation of protein kinase activity | 63/4924 | 7.33E-07 | 5.92E-06 |
| GO:0051057 | positive regulation of small GTPase mediated signal transduction | 32/4924 | 7.41E-07 | 5.99E-06 |
| GO:0045446 | endothelial cell differentiation | 42/4924 | 7.76E-07 | 6.26E-06 |
| GO:0098840 | protein transport along microtubule | Nov-24 | 7.87E-07 | 6.31E-06 |
| GO:0099118 | microtubule-based protein transport | Nov-24 | 7.87E-07 | 6.31E-06 |
| GO:1902969 | mitotic DNA replication | Nov-24 | 7.87E-07 | 6.31E-06 |
| GO:0007272 | ensheathment of neurons | 57/4924 | 7.88E-07 | 6.31E-06 |
| GO:0008366 | axon ensheathment | 57/4924 | 7.88E-07 | 6.31E-06 |
| GO:0046545 | development of primary female sexual characteristics | 43/4924 | 7.91E-07 | 6.33E-06 |
| GO:0003382 | epithelial cell morphogenesis | 23/4924 | 7.94E-07 | 6.34E-06 |
| GO:0045197 | establishment or maintenance of epithelial cell apical/basal polarity | 23/4924 | 7.94E-07 | 6.34E-06 |

| | | | | |
|---|---|---|---|---|
| GO:0046579 | positive regulation of Ras protein signal transduction | 30/4924 | 7.97E-07 | 6.35E-06 |
| GO:0055002 | striated muscle cell development | 30/4924 | 7.97E-07 | 6.35E-06 |
| GO:1904375 | regulation of protein localization to cell periphery | 48/4924 | 7.99E-07 | 6.35E-06 |
| GO:0030278 | regulation of ossification | 47/4924 | 8.05E-07 | 6.39E-06 |
| GO:0042476 | odontogenesis | 45/4924 | 8.07E-07 | 6.40E-06 |
| GO:1903169 | regulation of calcium ion transmembrane transport | 56/4924 | 8.18E-07 | 6.48E-06 |
| GO:0072078 | nephron tubule morphogenesis | 33/4924 | 8.30E-07 | 6.57E-06 |
| GO:0032331 | negative regulation of chondrocyte differentiation | 15/4924 | 8.34E-07 | 6.59E-06 |
| GO:1904377 | positive regulation of protein localization to cell periphery | 29/4924 | 9.76E-07 | 7.70E-06 |
| GO:0038084 | vascular endothelial growth factor signaling pathway | 21/4924 | 9.82E-07 | 7.74E-06 |
| GO:0031122 | cytoplasmic microtubule organization | 27/4924 | 9.86E-07 | 7.75E-06 |
| GO:0051784 | negative regulation of nuclear division | 27/4924 | 9.86E-07 | 7.75E-06 |
| GO:0000727 | double-strand break repair via break-induced replication | Oct-24 | 9.93E-07 | 7.79E-06 |
| GO:0060907 | positive regulation of macrophage cytokine production | Oct-24 | 9.93E-07 | 7.79E-06 |
| GO:0007599 | hemostasis | 56/4924 | 1.00E-06 | 7.86E-06 |
| GO:0050817 | coagulation | 56/4924 | 1.00E-06 | 7.86E-06 |
| GO:0008585 | female gonad development | 42/4924 | 1.01E-06 | 7.86E-06 |
| GO:0032602 | chemokine production | 42/4924 | 1.01E-06 | 7.86E-06 |

| | | | | |
|---|---|---|---|---|
| GO:0002285 | lymphocyte activation involved in immune response | 67/4924 | 1.02E-06 | 7.93E-06 |
| GO:0032411 | positive regulation of transporter activity | 46/4924 | 1.03E-06 | 8.00E-06 |
| GO:0048565 | digestive tract development | 44/4924 | 1.03E-06 | 8.00E-06 |
| GO:0002931 | response to ischemia | 20/4924 | 1.06E-06 | 8.22E-06 |
| GO:0014031 | mesenchymal cell development | 36/4924 | 1.06E-06 | 8.22E-06 |
| GO:0048864 | stem cell development | 36/4924 | 1.06E-06 | 8.22E-06 |
| GO:0030178 | negative regulation of Wnt signaling pathway | 54/4924 | 1.08E-06 | 8.37E-06 |
| GO:0045581 | negative regulation of T cell differentiation | 24/4924 | 1.09E-06 | 8.38E-06 |
| GO:0061005 | cell differentiation involved in kidney development | 24/4924 | 1.09E-06 | 8.38E-06 |
| GO:0010719 | negative regulation of epithelial to mesenchymal transition | 19/4924 | 1.11E-06 | 8.56E-06 |
| GO:0014033 | neural crest cell differentiation | 37/4924 | 1.12E-06 | 8.64E-06 |
| GO:0044319 | wound healing, spreading of cells | 18/4924 | 1.13E-06 | 8.64E-06 |
| GO:0090505 | epiboly involved in wound healing | 18/4924 | 1.13E-06 | 8.64E-06 |
| GO:1904646 | cellular response to amyloid-beta | 18/4924 | 1.13E-06 | 8.64E-06 |
| GO:0061045 | negative regulation of wound healing | 30/4924 | 1.13E-06 | 8.68E-06 |
| GO:0033673 | negative regulation of kinase activity | 68/4924 | 1.14E-06 | 8.73E-06 |
| GO:0032722 | positive regulation of chemokine production | 33/4924 | 1.14E-06 | 8.73E-06 |
| GO:0072089 | stem cell proliferation | 33/4924 | 1.14E-06 | 8.73E-06 |

| | | | | |
|---|---|---|---|---|
| GO:0006470 | protein dephosphorylation | 73/4924 | 1.16E-06 | 8.84E-06 |
| GO:0032944 | regulation of mononuclear cell proliferation | 73/4924 | 1.16E-06 | 8.84E-06 |
| GO:0010952 | positive regulation of peptidase activity | 57/4924 | 1.18E-06 | 8.97E-06 |
| GO:0097035 | regulation of membrane lipid distribution | 26/4924 | 1.18E-06 | 8.98E-06 |
| GO:0035264 | multicellular organism growth | 67/4924 | 1.21E-06 | 9.22E-06 |
| GO:0051147 | regulation of muscle cell differentiation | 50/4924 | 1.22E-06 | 9.22E-06 |
| GO:1903038 | negative regulation of leukocyte cell-cell adhesion | 49/4924 | 1.24E-06 | 9.42E-06 |
| GO:0032642 | regulation of chemokine production | 40/4924 | 1.26E-06 | 9.49E-06 |
| GO:0046661 | male sex differentiation | 55/4924 | 1.28E-06 | 9.67E-06 |
| GO:0048872 | homeostasis of number of cells | 100/4924 | 1.30E-06 | 9.78E-06 |
| GO:1902107 | positive regulation of leukocyte differentiation | 59/4924 | 1.31E-06 | 9.85E-06 |
| GO:1903708 | positive regulation of hemopoiesis | 59/4924 | 1.31E-06 | 9.85E-06 |
| GO:0042770 | signal transduction in response to DNA damage | 54/4924 | 1.33E-06 | 1.00E-05 |
| GO:0043367 | CD4-positive, alpha-beta T cell differentiation | 35/4924 | 1.34E-06 | 1.00E-05 |
| GO:2001057 | reactive nitrogen species metabolic process | 35/4924 | 1.34E-06 | 1.00E-05 |
| GO:0035051 | cardiocyte differentiation | 58/4924 | 1.37E-06 | 1.03E-05 |
| GO:0035904 | aorta development | 29/4924 | 1.39E-06 | 1.04E-05 |
| GO:0035019 | somatic stem cell population maintenance | 25/4924 | 1.41E-06 | 1.05E-05 |

| | | | | |
|---|---|---|---|---|
| GO:0043536 | positive regulation of blood vessel endothelial cell migration | 25/4924 | 1.41E-06 | 1.05E-05 |
| GO:0051304 | chromosome separation | 37/4924 | 1.49E-06 | 1.11E-05 |
| GO:0002822 | regulation of adaptive immune response based on somatic recombination of immune receptors built from immunoglobulin superfamily domains | 66/4924 | 1.54E-06 | 1.15E-05 |
| GO:0006836 | neurotransmitter transport | 71/4924 | 1.57E-06 | 1.16E-05 |
| GO:0019827 | stem cell population maintenance | 55/4924 | 1.57E-06 | 1.16E-05 |
| GO:0051100 | negative regulation of binding | 54/4924 | 1.63E-06 | 1.21E-05 |
| GO:0001666 | response to hypoxia | 65/4924 | 1.64E-06 | 1.21E-05 |
| GO:0046631 | alpha-beta T cell activation | 58/4924 | 1.67E-06 | 1.23E-05 |
| GO:0014009 | glial cell proliferation | 26/4924 | 1.75E-06 | 1.29E-05 |
| GO:0098815 | modulation of excitatory postsynaptic potential | 26/4924 | 1.75E-06 | 1.29E-05 |
| GO:0032232 | negative regulation of actin filament bundle assembly | 20/4924 | 1.78E-06 | 1.31E-05 |
| GO:0030902 | hindbrain development | 56/4924 | 1.83E-06 | 1.35E-05 |
| GO:0097722 | sperm motility | 50/4924 | 1.88E-06 | 1.38E-05 |
| GO:0086091 | regulation of heart rate by cardiac conduction | 19/4924 | 1.92E-06 | 1.41E-05 |
| GO:0030317 | flagellated sperm motility | 49/4924 | 1.93E-06 | 1.42E-05 |
| GO:2001056 | positive regulation of cysteine-type endopeptidase activity | 49/4924 | 1.93E-06 | 1.42E-05 |
| GO:0038034 | signal transduction in absence of ligand | 32/4924 | 1.96E-06 | 1.43E-05 |
| GO:0051303 | establishment of chromosome localization | 32/4924 | 1.96E-06 | 1.43E-05 |

| | | | | |
|---|---|---|---|---|
| GO:0097192 | extrinsic apoptotic signaling pathway in absence of ligand | 32/4924 | 1.96E-06 | 1.43E-05 |
| GO:0097061 | dendritic spine organization | 37/4924 | 1.97E-06 | 1.44E-05 |
| GO:0010950 | positive regulation of endopeptidase activity | 54/4924 | 2.00E-06 | 1.46E-05 |
| GO:0031116 | positive regulation of microtubule polymerization | 18/4924 | 2.01E-06 | 1.46E-05 |
| GO:1901889 | negative regulation of cell junction assembly | 17/4924 | 2.03E-06 | 1.48E-05 |
| GO:0031623 | receptor internalization | 46/4924 | 2.07E-06 | 1.50E-05 |
| GO:0042303 | molting cycle | 46/4924 | 2.07E-06 | 1.50E-05 |
| GO:0042633 | hair cycle | 46/4924 | 2.07E-06 | 1.50E-05 |
| GO:0071901 | negative regulation of protein serine/threonine kinase activity | 39/4924 | 2.08E-06 | 1.50E-05 |
| GO:0106027 | neuron projection organization | 39/4924 | 2.08E-06 | 1.50E-05 |
| GO:2000779 | regulation of double-strand break repair | 39/4924 | 2.08E-06 | 1.50E-05 |
| GO:0051893 | regulation of focal adhesion assembly | 27/4924 | 2.09E-06 | 1.51E-05 |
| GO:0090109 | regulation of cell-substrate junction assembly | 27/4924 | 2.09E-06 | 1.51E-05 |
| GO:0006493 | protein O-linked glycosylation | 25/4924 | 2.10E-06 | 1.51E-05 |
| GO:0048538 | thymus development | 25/4924 | 2.10E-06 | 1.51E-05 |
| GO:0061383 | trabecula morphogenesis | 25/4924 | 2.10E-06 | 1.51E-05 |
| GO:0050868 | negative regulation of T cell activation | 45/4924 | 2.10E-06 | 1.51E-05 |
| GO:0046632 | alpha-beta T cell differentiation | 44/4924 | 2.12E-06 | 1.52E-05 |

| | | | | |
|---|---|---|---|---|
| GO:0060113 | inner ear receptor cell differentiation | 33/4924 | 2.13E-06 | 1.52E-05 |
| GO:0032943 | mononuclear cell proliferation | 92/4924 | 2.19E-06 | 1.56E-05 |
| GO:0007281 | germ cell development | 102/4924 | 2.37E-06 | 1.69E-05 |
| GO:0034501 | protein localization to kinetochore | 13/4924 | 2.44E-06 | 1.73E-05 |
| GO:1903083 | protein localization to condensed chromosome | 13/4924 | 2.44E-06 | 1.73E-05 |
| GO:0061337 | cardiac conduction | 28/4924 | 2.44E-06 | 1.73E-05 |
| GO:0120161 | regulation of cold-induced thermogenesis | 48/4924 | 2.47E-06 | 1.76E-05 |
| GO:0035272 | exocrine system development | 24/4924 | 2.51E-06 | 1.78E-05 |
| GO:0097193 | intrinsic apoptotic signaling pathway | 85/4924 | 2.54E-06 | 1.80E-05 |
| GO:0060324 | face development | 26/4924 | 2.55E-06 | 1.81E-05 |
| GO:0021543 | pallium development | 52/4924 | 2.66E-06 | 1.88E-05 |
| GO:0032414 | positive regulation of ion transmembrane transporter activity | 43/4924 | 2.71E-06 | 1.92E-05 |
| GO:0010633 | negative regulation of epithelial cell migration | 29/4924 | 2.77E-06 | 1.95E-05 |
| GO:0071805 | potassium ion transmembrane transport | 62/4924 | 2.79E-06 | 1.97E-05 |
| GO:0030595 | leukocyte chemotaxis | 67/4924 | 2.85E-06 | 2.01E-05 |
| GO:0051693 | actin filament capping | 20/4924 | 2.91E-06 | 2.05E-05 |
| GO:1900180 | regulation of protein localization to nucleus | 49/4924 | 2.97E-06 | 2.09E-05 |
| GO:0051496 | positive regulation of stress fiber assembly | 25/4924 | 3.09E-06 | 2.17E-05 |

| | | | | |
|---|---|---|---|---|
| GO:0007269 | neurotransmitter secretion | 57/4924 | 3.10E-06 | 2.17E-05 |
| GO:0099643 | signal release from synapse | 57/4924 | 3.10E-06 | 2.17E-05 |
| GO:0032651 | regulation of interleukin-1 beta production | 35/4924 | 3.17E-06 | 2.22E-05 |
| GO:0051294 | establishment of spindle orientation | 19/4924 | 3.22E-06 | 2.25E-05 |
| GO:0071634 | regulation of transforming growth factor beta production | 19/4924 | 3.22E-06 | 2.25E-05 |
| GO:0043123 | positive regulation of I-kappaB kinase/NF-kappaB signaling | 43/4924 | 3.43E-06 | 2.39E-05 |
| GO:0043280 | positive regulation of cysteine-type endopeptidase activity involved in apoptotic process | 43/4924 | 3.43E-06 | 2.39E-05 |
| GO:0003143 | embryonic heart tube morphogenesis | 28/4924 | 3.44E-06 | 2.39E-05 |
| GO:0071677 | positive regulation of mononuclear cell migration | 28/4924 | 3.44E-06 | 2.39E-05 |
| GO:0003401 | axis elongation | 18/4924 | 3.48E-06 | 2.41E-05 |
| GO:0021846 | cell proliferation in forebrain | 18/4924 | 3.48E-06 | 2.41E-05 |
| GO:0022404 | molting cycle process | 41/4924 | 3.49E-06 | 2.41E-05 |
| GO:0022405 | hair cycle process | 41/4924 | 3.49E-06 | 2.41E-05 |
| GO:0071459 | protein localization to chromosome, centromeric region | 15/4924 | 3.49E-06 | 2.41E-05 |
| GO:2000050 | regulation of non-canonical Wnt signaling pathway | 15/4924 | 3.49E-06 | 2.41E-05 |
| GO:0071824 | protein-DNA complex subunit organization | 58/4924 | 3.53E-06 | 2.43E-05 |
| GO:0070831 | basement membrane assembly | Oct-24 | 3.64E-06 | 2.50E-05 |
| GO:0072393 | microtubule anchoring at microtubule organizing center | Oct-24 | 3.64E-06 | 2.50E-05 |

| | | | | |
|---|---|---|---|---|
| GO:0060711 | labyrinthine layer development | 26/4924 | 3.68E-06 | 2.53E-05 |
| GO:0034113 | heterotypic cell-cell adhesion | 24/4924 | 3.73E-06 | 2.56E-05 |
| GO:0006813 | potassium ion transport | 70/4924 | 3.77E-06 | 2.59E-05 |
| GO:0106106 | cold-induced thermogenesis | 48/4924 | 3.79E-06 | 2.60E-05 |
| GO:0003151 | outflow tract morphogenesis | 33/4924 | 3.84E-06 | 2.63E-05 |
| GO:0060997 | dendritic spine morphogenesis | 29/4924 | 3.85E-06 | 2.63E-05 |
| GO:0014902 | myotube differentiation | 46/4924 | 4.03E-06 | 2.75E-05 |
| GO:0017158 | regulation of calcium ion-dependent exocytosis | 21/4924 | 4.07E-06 | 2.77E-05 |
| GO:0035633 | maintenance of blood-brain barrier | Dec-24 | 4.09E-06 | 2.78E-05 |
| GO:0061430 | bone trabecula morphogenesis | Dec-24 | 4.09E-06 | 2.78E-05 |
| GO:0000302 | response to reactive oxygen species | 58/4924 | 4.23E-06 | 2.87E-05 |
| GO:0007127 | meiosis I | 44/4924 | 4.23E-06 | 2.87E-05 |
| GO:0061008 | hepaticobiliary system development | 44/4924 | 4.23E-06 | 2.87E-05 |
| GO:0035637 | multicellular organismal signaling | 50/4924 | 4.32E-06 | 2.93E-05 |
| GO:0007173 | epidermal growth factor receptor signaling pathway | 37/4924 | 4.38E-06 | 2.96E-05 |
| GO:1901890 | positive regulation of cell junction assembly | 42/4924 | 4.38E-06 | 2.96E-05 |
| GO:0098773 | skin epidermis development | 41/4924 | 4.43E-06 | 2.99E-05 |
| GO:0001942 | hair follicle development | 40/4924 | 4.45E-06 | 3.01E-05 |

| | | | | |
|---|---|---|---|---|
| GO:0030857 | negative regulation of epithelial cell differentiation | 23/4924 | 4.47E-06 | 3.02E-05 |
| GO:0045637 | regulation of myeloid cell differentiation | 65/4924 | 4.55E-06 | 3.07E-05 |
| GO:0043433 | negative regulation of DNA-binding transcription factor activity | 53/4924 | 4.59E-06 | 3.09E-05 |
| GO:1904064 | positive regulation of cation transmembrane transport | 53/4924 | 4.59E-06 | 3.09E-05 |
| GO:0007204 | positive regulation of cytosolic calcium ion concentration | 94/4924 | 4.63E-06 | 3.11E-05 |
| GO:0071174 | mitotic spindle checkpoint signaling | 20/4924 | 4.67E-06 | 3.13E-05 |
| GO:0019722 | calcium-mediated signaling | 59/4924 | 4.76E-06 | 3.19E-05 |
| GO:0051302 | regulation of cell division | 59/4924 | 4.76E-06 | 3.19E-05 |
| GO:0010569 | regulation of double-strand break repair via homologous recombination | 28/4924 | 4.80E-06 | 3.21E-05 |
| GO:0090316 | positive regulation of intracellular protein transport | 47/4924 | 4.84E-06 | 3.23E-05 |
| GO:0032353 | negative regulation of hormone biosynthetic process | Sep-24 | 4.87E-06 | 3.25E-05 |
| GO:0014032 | neural crest cell development | 33/4924 | 5.10E-06 | 3.40E-05 |
| GO:0006275 | regulation of DNA replication | 45/4924 | 5.15E-06 | 3.43E-05 |
| GO:0050670 | regulation of lymphocyte proliferation | 70/4924 | 5.16E-06 | 3.43E-05 |
| GO:0001505 | regulation of neurotransmitter levels | 72/4924 | 5.18E-06 | 3.44E-05 |
| GO:0001755 | neural crest cell migration | 26/4924 | 5.25E-06 | 3.48E-05 |
| GO:0072678 | T cell migration | 26/4924 | 5.25E-06 | 3.48E-05 |
| GO:2000401 | regulation of lymphocyte migration | 26/4924 | 5.25E-06 | 3.48E-05 |

| | | | | |
|---|---|---|---|---|
| GO:0060675 | ureteric bud morphogenesis | 29/4924 | 5.30E-06 | 3.50E-05 |
| GO:0140029 | exocytic process | 34/4924 | 5.31E-06 | 3.51E-05 |
| GO:0001889 | liver development | 43/4924 | 5.41E-06 | 3.57E-05 |
| GO:2000278 | regulation of DNA biosynthetic process | 43/4924 | 5.41E-06 | 3.57E-05 |
| GO:0072577 | endothelial cell apoptotic process | 24/4924 | 5.48E-06 | 3.61E-05 |
| GO:0042176 | regulation of protein catabolic process | 102/4924 | 5.54E-06 | 3.65E-05 |
| GO:0001892 | embryonic placenta development | 40/4924 | 5.66E-06 | 3.72E-05 |
| GO:0032677 | regulation of interleukin-8 production | 30/4924 | 5.74E-06 | 3.77E-05 |
| GO:0061180 | mammary gland epithelium development | 30/4924 | 5.74E-06 | 3.77E-05 |
| GO:2000379 | positive regulation of reactive oxygen species metabolic process | 30/4924 | 5.74E-06 | 3.77E-05 |
| GO:0040001 | establishment of mitotic spindle localization | 18/4924 | 5.84E-06 | 3.82E-05 |
| GO:0030193 | regulation of blood coagulation | 27/4924 | 5.97E-06 | 3.90E-05 |
| GO:0051984 | positive regulation of chromosome segregation | 14/4924 | 6.23E-06 | 4.07E-05 |
| GO:0060122 | inner ear receptor cell stereocilium organization | 21/4924 | 6.28E-06 | 4.09E-05 |
| GO:1903115 | regulation of actin filament-based movement | 21/4924 | 6.28E-06 | 4.09E-05 |
| GO:0048841 | regulation of axon extension involved in axon guidance | 17/4924 | 6.31E-06 | 4.11E-05 |
| GO:0060669 | embryonic placenta morphogenesis | 17/4924 | 6.31E-06 | 4.11E-05 |
| GO:0051048 | negative regulation of secretion | 60/4924 | 6.34E-06 | 4.12E-05 |

| | | | | |
|---|---|---|---|---|
| GO:0002821 | positive regulation of adaptive immune response | 50/4924 | 6.44E-06 | 4.18E-05 |
| GO:0034508 | centromere complex assembly | 16/4924 | 6.60E-06 | 4.28E-05 |
| GO:0034453 | microtubule anchoring | 15/4924 | 6.61E-06 | 4.28E-05 |
| GO:0002042 | cell migration involved in sprouting angiogenesis | 23/4924 | 6.63E-06 | 4.28E-05 |
| GO:0043370 | regulation of CD4-positive, alpha-beta T cell differentiation | 23/4924 | 6.63E-06 | 4.28E-05 |
| GO:2000351 | regulation of endothelial cell apoptotic process | 23/4924 | 6.63E-06 | 4.28E-05 |
| GO:0050818 | regulation of coagulation | 28/4924 | 6.63E-06 | 4.28E-05 |
| GO:0038065 | collagen-activated signaling pathway | Nov-24 | 6.73E-06 | 4.33E-05 |
| GO:0048103 | somatic stem cell division | Nov-24 | 6.73E-06 | 4.33E-05 |
| GO:0002088 | lens development in camera-type eye | 33/4924 | 6.74E-06 | 4.33E-05 |
| GO:0008593 | regulation of Notch signaling pathway | 33/4924 | 6.74E-06 | 4.33E-05 |
| GO:0006869 | lipid transport | 101/4924 | 6.80E-06 | 4.36E-05 |
| GO:0015833 | peptide transport | 86/4924 | 6.96E-06 | 4.47E-05 |
| GO:0018205 | peptidyl-lysine modification | 102/4924 | 7.01E-06 | 4.49E-05 |
| GO:0031497 | chromatin assembly | 52/4924 | 7.08E-06 | 4.53E-05 |
| GO:0032091 | negative regulation of protein binding | 35/4924 | 7.11E-06 | 4.55E-05 |
| GO:0038061 | NIK/NF-kappaB signaling | 40/4924 | 7.16E-06 | 4.57E-05 |
| GO:0060078 | regulation of postsynaptic membrane potential | 40/4924 | 7.16E-06 | 4.57E-05 |

| | | | | |
|---|---|---|---|---|
| GO:0010657 | muscle cell apoptotic process | 36/4924 | 7.21E-06 | 4.59E-05 |
| GO:0042116 | macrophage activation | 36/4924 | 7.21E-06 | 4.59E-05 |
| GO:0051149 | positive regulation of muscle cell differentiation | 29/4924 | 7.23E-06 | 4.60E-05 |
| GO:0072171 | mesonephric tubule morphogenesis | 29/4924 | 7.23E-06 | 4.60E-05 |
| GO:0045880 | positive regulation of smoothened signaling pathway | 20/4924 | 7.33E-06 | 4.65E-05 |
| GO:1904705 | regulation of vascular associated smooth muscle cell proliferation | 26/4924 | 7.42E-06 | 4.70E-05 |
| GO:0006282 | regulation of DNA repair | 51/4924 | 7.45E-06 | 4.72E-05 |
| GO:0034614 | cellular response to reactive oxygen species | 46/4924 | 7.62E-06 | 4.82E-05 |
| GO:0050729 | positive regulation of inflammatory response | 46/4924 | 7.62E-06 | 4.82E-05 |
| GO:0002221 | pattern recognition receptor signaling pathway | 54/4924 | 7.65E-06 | 4.83E-05 |
| GO:0032637 | interleukin-8 production | 30/4924 | 7.76E-06 | 4.90E-05 |
| GO:0046883 | regulation of hormone secretion | 83/4924 | 7.81E-06 | 4.92E-05 |
| GO:0010463 | mesenchymal cell proliferation | 24/4924 | 7.93E-06 | 4.99E-05 |
| GO:0086001 | cardiac muscle cell action potential | 24/4924 | 7.93E-06 | 4.99E-05 |
| GO:0046651 | lymphocyte proliferation | 89/4924 | 7.99E-06 | 5.02E-05 |
| GO:0070373 | negative regulation of ERK1 and ERK2 cascade | 31/4924 | 8.20E-06 | 5.15E-05 |
| GO:0043491 | protein kinase B signaling | 61/4924 | 8.31E-06 | 5.21E-05 |
| GO:0010876 | lipid localization | 113/4924 | 8.34E-06 | 5.23E-05 |

| | | | | |
|---|---|---|---|---|
| GO:1902893 | regulation of miRNA transcription | 32/4924 | 8.57E-06 | 5.36E-05 |
| GO:0034612 | response to tumor necrosis factor | 60/4924 | 8.89E-06 | 5.56E-05 |
| GO:0070665 | positive regulation of leukocyte proliferation | 51/4924 | 9.01E-06 | 5.63E-05 |
| GO:0045666 | positive regulation of neuron differentiation | 40/4924 | 9.02E-06 | 5.63E-05 |
| GO:0033077 | T cell differentiation in thymus | 34/4924 | 9.07E-06 | 5.65E-05 |
| GO:0007091 | metaphase/anaphase transition of mitotic cell cycle | 25/4924 | 9.19E-06 | 5.72E-05 |
| GO:0034766 | negative regulation of ion transmembrane transport | 37/4924 | 9.29E-06 | 5.78E-05 |
| GO:0040037 | negative regulation of fibroblast growth factor receptor signaling pathway | Dec-24 | 9.36E-06 | 5.81E-05 |
| GO:0090660 | cerebrospinal fluid circulation | Dec-24 | 9.36E-06 | 5.81E-05 |
| GO:1903524 | positive regulation of blood circulation | 21/4924 | 9.53E-06 | 5.91E-05 |
| GO:0022617 | extracellular matrix disassembly | 18/4924 | 9.56E-06 | 5.92E-05 |
| GO:0070306 | lens fiber cell differentiation | 18/4924 | 9.56E-06 | 5.92E-05 |
| GO:1905508 | protein localization to microtubule organizing center | 18/4924 | 9.56E-06 | 5.92E-05 |
| GO:0045604 | regulation of epidermal cell differentiation | 23/4924 | 9.69E-06 | 5.98E-05 |
| GO:0045773 | positive regulation of axon extension | 23/4924 | 9.69E-06 | 5.98E-05 |
| GO:0099504 | synaptic vesicle cycle | 65/4924 | 1.02E-05 | 6.26E-05 |
| GO:0002699 | positive regulation of immune effector process | 83/4924 | 1.02E-05 | 6.27E-05 |
| GO:0002244 | hematopoietic progenitor cell differentiation | 52/4924 | 1.03E-05 | 6.33E-05 |

| | | | | |
|---|---|---|---|---|
| GO:0032615 | interleukin-12 production | 26/4924 | 1.04E-05 | 6.37E-05 |
| GO:0010714 | positive regulation of collagen metabolic process | 17/4924 | 1.06E-05 | 6.52E-05 |
| GO:0010457 | centriole-centriole cohesion | Oct-24 | 1.08E-05 | 6.61E-05 |
| GO:2000052 | positive regulation of non-canonical Wnt signaling pathway | Oct-24 | 1.08E-05 | 6.61E-05 |
| GO:0008088 | axo-dendritic transport | 31/4924 | 1.09E-05 | 6.67E-05 |
| GO:0009411 | response to UV | 47/4924 | 1.10E-05 | 6.70E-05 |
| GO:0065004 | protein-DNA complex assembly | 47/4924 | 1.10E-05 | 6.70E-05 |
| GO:0042472 | inner ear morphogenesis | 40/4924 | 1.13E-05 | 6.91E-05 |
| GO:0033045 | regulation of sister chromatid segregation | 27/4924 | 1.14E-05 | 6.95E-05 |
| GO:0051926 | negative regulation of calcium ion transport | 27/4924 | 1.14E-05 | 6.95E-05 |
| GO:0150116 | regulation of cell-substrate junction organization | 27/4924 | 1.14E-05 | 6.95E-05 |
| GO:1900046 | regulation of hemostasis | 27/4924 | 1.14E-05 | 6.95E-05 |
| GO:0032967 | positive regulation of collagen biosynthetic process | 16/4924 | 1.15E-05 | 6.97E-05 |
| GO:0086010 | membrane depolarization during action potential | 16/4924 | 1.15E-05 | 6.97E-05 |
| GO:0035710 | CD4-positive, alpha-beta T cell activation | 39/4924 | 1.15E-05 | 7.00E-05 |
| GO:0060079 | excitatory postsynaptic potential | 33/4924 | 1.16E-05 | 7.00E-05 |
| GO:1900182 | positive regulation of protein localization to nucleus | 33/4924 | 1.16E-05 | 7.00E-05 |
| GO:0070509 | calcium ion import | 34/4924 | 1.17E-05 | 7.11E-05 |

| | | | | |
|---|---|---|---|---|
| GO:0008038 | neuron recognition | 22/4924 | 1.18E-05 | 7.12E-05 |
| GO:0009798 | axis specification | 35/4924 | 1.19E-05 | 7.16E-05 |
| GO:0051153 | regulation of striated muscle cell differentiation | 36/4924 | 1.19E-05 | 7.16E-05 |
| GO:0060445 | branching involved in salivary gland morphogenesis | 15/4924 | 1.20E-05 | 7.21E-05 |
| GO:1903055 | positive regulation of extracellular matrix organization | 15/4924 | 1.20E-05 | 7.21E-05 |
| GO:0032732 | positive regulation of interleukin-1 production | 28/4924 | 1.24E-05 | 7.44E-05 |
| GO:0050773 | regulation of dendrite development | 44/4924 | 1.24E-05 | 7.47E-05 |
| GO:0010965 | regulation of mitotic sister chromatid separation | 25/4924 | 1.29E-05 | 7.74E-05 |
| GO:1903078 | positive regulation of protein localization to plasma membrane | 25/4924 | 1.29E-05 | 7.74E-05 |
| GO:0006029 | proteoglycan metabolic process | 29/4924 | 1.32E-05 | 7.88E-05 |
| GO:0043113 | receptor clustering | 29/4924 | 1.32E-05 | 7.88E-05 |
| GO:2001022 | positive regulation of response to DNA damage stimulus | 47/4924 | 1.33E-05 | 7.98E-05 |
| GO:0030534 | adult behavior | 56/4924 | 1.37E-05 | 8.19E-05 |
| GO:0002702 | positive regulation of production of molecular mediator of immune response | 41/4924 | 1.38E-05 | 8.21E-05 |
| GO:0003208 | cardiac ventricle morphogenesis | 30/4924 | 1.38E-05 | 8.25E-05 |
| GO:0048706 | embryonic skeletal system development | 46/4924 | 1.40E-05 | 8.33E-05 |
| GO:0010799 | regulation of peptidyl-threonine phosphorylation | 21/4924 | 1.42E-05 | 8.46E-05 |
| GO:0002720 | positive regulation of cytokine production involved in immune response | 26/4924 | 1.43E-05 | 8.52E-05 |

| | | | | |
|---|---|---|---|---|
| GO:0034103 | regulation of tissue remodeling | 31/4924 | 1.43E-05 | 8.53E-05 |
| GO:1903305 | regulation of regulated secretory pathway | 55/4924 | 1.46E-05 | 8.68E-05 |
| GO:0061614 | miRNA transcription | 32/4924 | 1.47E-05 | 8.74E-05 |
| GO:0008630 | intrinsic apoptotic signaling pathway in response to DNA damage | 38/4924 | 1.48E-05 | 8.74E-05 |
| GO:0030168 | platelet activation | 33/4924 | 1.50E-05 | 8.88E-05 |
| GO:0001953 | negative regulation of cell-matrix adhesion | 18/4924 | 1.53E-05 | 9.03E-05 |
| GO:2000514 | regulation of CD4-positive, alpha-beta T cell activation | 27/4924 | 1.56E-05 | 9.20E-05 |
| GO:2001257 | regulation of cation channel activity | 54/4924 | 1.56E-05 | 9.20E-05 |
| GO:0003184 | pulmonary valve morphogenesis | Nov-24 | 1.61E-05 | 9.49E-05 |
| GO:0051382 | kinetochore assembly | Nov-24 | 1.61E-05 | 9.49E-05 |
| GO:0032351 | negative regulation of hormone metabolic process | Sep-24 | 1.65E-05 | 9.69E-05 |
| GO:0038063 | collagen-activated tyrosine kinase receptor signaling pathway | Sep-24 | 1.65E-05 | 9.69E-05 |
| GO:0072697 | protein localization to cell cortex | Sep-24 | 1.65E-05 | 9.69E-05 |
| GO:0031577 | spindle checkpoint signaling | 20/4924 | 1.70E-05 | 9.99E-05 |
| GO:0086009 | membrane repolarization | 20/4924 | 1.70E-05 | 9.99E-05 |
| GO:0002688 | regulation of leukocyte chemotaxis | 41/4924 | 1.70E-05 | 9.99E-05 |
| GO:0060428 | lung epithelium development | 22/4924 | 1.71E-05 | 0.000100237 |
| GO:0009952 | anterior/posterior pattern specification | 66/4924 | 1.73E-05 | 0.000101246 |

| | | | | |
|---|---|---|---|---|
| GO:0008608 | attachment of spindle microtubules to kinetochore | 17/4924 | 1.73E-05 | 0.000101287 |
| GO:0071539 | protein localization to centrosome | 17/4924 | 1.73E-05 | 0.000101287 |
| GO:0010769 | regulation of cell morphogenesis involved in differentiation | 40/4924 | 1.76E-05 | 0.000102491 |
| GO:0060419 | heart growth | 40/4924 | 1.76E-05 | 0.000102491 |
| GO:0042471 | ear morphogenesis | 45/4924 | 1.78E-05 | 0.000103804 |
| GO:0048260 | positive regulation of receptor-mediated endocytosis | 25/4924 | 1.79E-05 | 0.000104355 |
| GO:0014015 | positive regulation of gliogenesis | 30/4924 | 1.83E-05 | 0.000106101 |
| GO:0010827 | regulation of glucose transmembrane transport | 32/4924 | 1.91E-05 | 0.000111152 |
| GO:0045185 | maintenance of protein location | 35/4924 | 1.93E-05 | 0.000112016 |
| GO:0048525 | negative regulation of viral process | 35/4924 | 1.93E-05 | 0.000112016 |
| GO:0042475 | odontogenesis of dentin-containing tooth | 33/4924 | 1.93E-05 | 0.000112016 |
| GO:1904063 | negative regulation of cation transmembrane transport | 34/4924 | 1.94E-05 | 0.000112258 |
| GO:0002824 | positive regulation of adaptive immune response based on somatic recombination of immune receptors built from immunoglobulin superfamily domains | 47/4924 | 1.96E-05 | 0.000112988 |
| GO:0015698 | inorganic anion transport | 47/4924 | 1.96E-05 | 0.000112988 |
| GO:0046660 | female sex differentiation | 47/4924 | 1.96E-05 | 0.000112988 |
| GO:0043903 | regulation of biological process involved in symbiotic interaction | 26/4924 | 1.96E-05 | 0.000113008 |
| GO:1990874 | vascular associated smooth muscle cell proliferation | 26/4924 | 1.96E-05 | 0.000113008 |
| GO:0002031 | G protein-coupled receptor internalization | Dec-24 | 1.97E-05 | 0.000113483 |

| | | | | |
|---|---|---|---|---|
| GO:0072010 | glomerular epithelium development | Dec-24 | 1.97E-05 | 0.000113483 |
| GO:0072593 | reactive oxygen species metabolic process | 68/4924 | 1.98E-05 | 0.000113957 |
| GO:0001774 | microglial cell activation | 19/4924 | 2.03E-05 | 0.000115943 |
| GO:0006904 | vesicle docking involved in exocytosis | 19/4924 | 2.03E-05 | 0.000115943 |
| GO:0007143 | female meiotic nuclear division | 19/4924 | 2.03E-05 | 0.000115943 |
| GO:0045841 | negative regulation of mitotic metaphase/anaphase transition | 19/4924 | 2.03E-05 | 0.000115943 |
| GO:2001239 | regulation of extrinsic apoptotic signaling pathway in absence of ligand | 21/4924 | 2.09E-05 | 0.000119397 |
| GO:0021885 | forebrain cell migration | 27/4924 | 2.11E-05 | 0.000120359 |
| GO:0045165 | cell fate commitment | 77/4924 | 2.18E-05 | 0.000124076 |
| GO:0030859 | polarized epithelial cell differentiation | 14/4924 | 2.18E-05 | 0.000124076 |
| GO:0014068 | positive regulation of phosphatidylinositol 3-kinase signaling | 28/4924 | 2.23E-05 | 0.000126766 |
| GO:0045669 | positive regulation of osteoblast differentiation | 28/4924 | 2.23E-05 | 0.000126766 |
| GO:0048278 | vesicle docking | 24/4924 | 2.24E-05 | 0.000127269 |
| GO:0060632 | regulation of microtubule-based movement | 24/4924 | 2.24E-05 | 0.000127269 |
| GO:2001238 | positive regulation of extrinsic apoptotic signaling pathway | 24/4924 | 2.24E-05 | 0.000127269 |
| GO:0030316 | osteoclast differentiation | 39/4924 | 2.25E-05 | 0.000127566 |
| GO:0035270 | endocrine system development | 44/4924 | 2.27E-05 | 0.00012886 |
| GO:0002634 | regulation of germinal center formation | Aug-24 | 2.35E-05 | 0.000132606 |

| | | | | |
|---|---|---|---|---|
| GO:1902746 | regulation of lens fiber cell differentiation | Aug-24 | 2.35E-05 | 0.000132606 |
| GO:0008625 | extrinsic apoptotic signaling pathway via death domain receptors | 30/4924 | 2.40E-05 | 0.000135314 |
| GO:0044784 | metaphase/anaphase transition of cell cycle | 25/4924 | 2.47E-05 | 0.000139357 |
| GO:0002690 | positive regulation of leukocyte chemotaxis | 34/4924 | 2.47E-05 | 0.00013954 |
| GO:0032946 | positive regulation of mononuclear cell proliferation | 46/4924 | 2.49E-05 | 0.000140551 |
| GO:0007626 | locomotory behavior | 72/4924 | 2.52E-05 | 0.000141672 |
| GO:0002269 | leukocyte activation involved in inflammatory response | 20/4924 | 2.53E-05 | 0.000142227 |
| GO:0033047 | regulation of mitotic sister chromatid segregation | 20/4924 | 2.53E-05 | 0.000142227 |
| GO:0051985 | negative regulation of chromosome segregation | 20/4924 | 2.53E-05 | 0.000142227 |
| GO:0015748 | organophosphate ester transport | 40/4924 | 2.69E-05 | 0.00015101 |
| GO:0098656 | anion transmembrane transport | 58/4924 | 2.71E-05 | 0.00015166 |
| GO:0002790 | peptide secretion | 79/4924 | 2.71E-05 | 0.00015166 |
| GO:0007076 | mitotic chromosome condensation | Oct-24 | 2.73E-05 | 0.000152911 |
| GO:0120033 | negative regulation of plasma membrane bounded cell projection assembly | 17/4924 | 2.76E-05 | 0.000154529 |
| GO:0051489 | regulation of filopodium assembly | 23/4924 | 2.80E-05 | 0.000156207 |
| GO:2000179 | positive regulation of neural precursor cell proliferation | 27/4924 | 2.83E-05 | 0.000157698 |
| GO:0036293 | response to decreased oxygen levels | 67/4924 | 2.86E-05 | 0.000159307 |
| GO:0031570 | DNA integrity checkpoint signaling | 38/4924 | 2.88E-05 | 0.000160333 |

| | | | | |
|---|---|---|---|---|
| GO:0140056 | organelle localization by membrane tethering | 28/4924 | 2.96E-05 | 0.000164488 |
| GO:0055017 | cardiac muscle tissue growth | 37/4924 | 2.96E-05 | 0.000164621 |
| GO:0003254 | regulation of membrane depolarization | 21/4924 | 3.02E-05 | 0.000167816 |
| GO:0045214 | sarcomere organization | 21/4924 | 3.02E-05 | 0.000167816 |
| GO:0030501 | positive regulation of bone mineralization | 19/4924 | 3.05E-05 | 0.000168579 |
| GO:0033046 | negative regulation of sister chromatid segregation | 19/4924 | 3.05E-05 | 0.000168579 |
| GO:0033048 | negative regulation of mitotic sister chromatid segregation | 19/4924 | 3.05E-05 | 0.000168579 |
| GO:0045823 | positive regulation of heart contraction | 19/4924 | 3.05E-05 | 0.000168579 |
| GO:2000816 | negative regulation of mitotic sister chromatid separation | 19/4924 | 3.05E-05 | 0.000168579 |
| GO:0071356 | cellular response to tumor necrosis factor | 54/4924 | 3.06E-05 | 0.000169038 |
| GO:0010812 | negative regulation of cell-substrate adhesion | 24/4924 | 3.10E-05 | 0.000170926 |
| GO:0032731 | positive regulation of interleukin-1 beta production | 24/4924 | 3.10E-05 | 0.000170926 |
| GO:0002063 | chondrocyte development | 16/4924 | 3.15E-05 | 0.000173169 |
| GO:0033687 | osteoblast proliferation | 16/4924 | 3.15E-05 | 0.000173169 |
| GO:1905314 | semi-lunar valve development | 16/4924 | 3.15E-05 | 0.000173169 |
| GO:2000108 | positive regulation of leukocyte apoptotic process | 16/4924 | 3.15E-05 | 0.000173169 |
| GO:0021675 | nerve development | 31/4924 | 3.16E-05 | 0.000173435 |
| GO:0097581 | lamellipodium organization | 32/4924 | 3.17E-05 | 0.000174027 |

| | | | | |
|---|---|---|---|---|
| GO:2000045 | regulation of G1/S transition of mitotic cell cycle | 45/4924 | 3.18E-05 | 0.000174027 |
| GO:0030072 | peptide hormone secretion | 77/4924 | 3.21E-05 | 0.000175418 |
| GO:0032649 | regulation of interferon-gamma production | 40/4924 | 3.31E-05 | 0.000181097 |
| GO:0001954 | positive regulation of cell-matrix adhesion | 25/4924 | 3.36E-05 | 0.000183503 |
| GO:0051306 | mitotic sister chromatid separation | 25/4924 | 3.36E-05 | 0.000183503 |
| GO:0060119 | inner ear receptor cell development | 25/4924 | 3.36E-05 | 0.000183503 |
| GO:0006023 | aminoglycan biosynthetic process | 22/4924 | 3.48E-05 | 0.000189631 |
| GO:0036159 | inner dynein arm assembly | Nov-24 | 3.50E-05 | 0.000190501 |
| GO:0002029 | desensitization of G protein-coupled receptor signaling pathway | 15/4924 | 3.52E-05 | 0.000191 |
| GO:0006929 | substrate-dependent cell migration | 15/4924 | 3.52E-05 | 0.000191 |
| GO:0022401 | negative adaptation of signaling pathway | 15/4924 | 3.52E-05 | 0.000191 |
| GO:0032609 | interferon-gamma production | 43/4924 | 3.53E-05 | 0.000191308 |
| GO:0007094 | mitotic spindle assembly checkpoint signaling | 18/4924 | 3.64E-05 | 0.000196951 |
| GO:0034110 | regulation of homotypic cell-cell adhesion | 18/4924 | 3.64E-05 | 0.000196951 |
| GO:0060325 | face morphogenesis | 18/4924 | 3.64E-05 | 0.000196951 |
| GO:0071173 | spindle assembly checkpoint signaling | 18/4924 | 3.64E-05 | 0.000196951 |
| GO:0050821 | protein stabilization | 56/4924 | 3.66E-05 | 0.000198008 |
| GO:0035296 | regulation of tube diameter | 51/4924 | 3.76E-05 | 0.000202435 |

| | | | | |
|---|---|---|---|---|
| GO:0097746 | blood vessel diameter maintenance | 51/4924 | 3.76E-05 | 0.000202435 |
| GO:0008542 | visual learning | 27/4924 | 3.76E-05 | 0.000202435 |
| GO:0043506 | regulation of JUN kinase activity | 27/4924 | 3.76E-05 | 0.000202435 |
| GO:0030204 | chondroitin sulfate metabolic process | 14/4924 | 3.81E-05 | 0.000203871 |
| GO:0031954 | positive regulation of protein autophosphorylation | 14/4924 | 3.81E-05 | 0.000203871 |
| GO:0032528 | microvillus organization | 14/4924 | 3.81E-05 | 0.000203871 |
| GO:0048843 | negative regulation of axon extension involved in axon guidance | 14/4924 | 3.81E-05 | 0.000203871 |
| GO:0060004 | reflex | 14/4924 | 3.81E-05 | 0.000203871 |
| GO:0072012 | glomerulus vasculature development | 14/4924 | 3.81E-05 | 0.000203871 |
| GO:2000316 | regulation of T-helper 17 type immune response | 14/4924 | 3.81E-05 | 0.000203871 |
| GO:2000377 | regulation of reactive oxygen species metabolic process | 48/4924 | 3.84E-05 | 0.000205226 |
| GO:0003177 | pulmonary valve development | Dec-24 | 3.89E-05 | 0.000207296 |
| GO:0045606 | positive regulation of epidermal cell differentiation | Dec-24 | 3.89E-05 | 0.000207296 |
| GO:0010761 | fibroblast migration | 23/4924 | 3.89E-05 | 0.000207296 |
| GO:0030071 | regulation of mitotic metaphase/anaphase transition | 23/4924 | 3.89E-05 | 0.000207296 |
| GO:0061097 | regulation of protein tyrosine kinase activity | 28/4924 | 3.89E-05 | 0.000207296 |
| GO:0042886 | amide transport | 90/4924 | 3.93E-05 | 0.000208856 |
| GO:0060021 | roof of mouth development | 34/4924 | 3.96E-05 | 0.000210248 |

| | | | | |
|---|---|---|---|---|
| GO:0010842 | retina layer formation | 13/4924 | 3.96E-05 | 0.000210248 |
| GO:0099625 | ventricular cardiac muscle cell membrane repolarization | 13/4924 | 3.96E-05 | 0.000210248 |
| GO:0003407 | neural retina development | 29/4924 | 3.99E-05 | 0.000211338 |
| GO:0070555 | response to interleukin-1 | 33/4924 | 4.02E-05 | 0.000212922 |
| GO:0044728 | DNA methylation or demethylation | 32/4924 | 4.05E-05 | 0.000214598 |
| GO:0045621 | positive regulation of lymphocyte differentiation | 39/4924 | 4.23E-05 | 0.000223925 |
| GO:0001947 | heart looping | 24/4924 | 4.24E-05 | 0.000224345 |
| GO:0051588 | regulation of neurotransmitter transport | 43/4924 | 4.27E-05 | 0.000225449 |
| GO:0007435 | salivary gland morphogenesis | 17/4924 | 4.30E-05 | 0.000226598 |
| GO:0035909 | aorta morphogenesis | 17/4924 | 4.30E-05 | 0.000226598 |
| GO:0051647 | nucleus localization | 17/4924 | 4.30E-05 | 0.000226598 |
| GO:0045933 | positive regulation of muscle contraction | 21/4924 | 4.32E-05 | 0.000226778 |
| GO:0090329 | regulation of DNA-templated DNA replication | 21/4924 | 4.32E-05 | 0.000226778 |
| GO:1904407 | positive regulation of nitric oxide metabolic process | 21/4924 | 4.32E-05 | 0.000226778 |
| GO:0035150 | regulation of tube size | 51/4924 | 4.44E-05 | 0.000233092 |
| GO:0031349 | positive regulation of defense response | 79/4924 | 4.45E-05 | 0.000233566 |
| GO:0010464 | regulation of mesenchymal cell proliferation | 19/4924 | 4.51E-05 | 0.00023609 |
| GO:0032508 | DNA duplex unwinding | 19/4924 | 4.51E-05 | 0.00023609 |

| | | | | |
|---|---|---|---|---|
| GO:1902100 | negative regulation of metaphase/anaphase transition of cell cycle | 19/4924 | 4.51E-05 | 0.00023609 |
| GO:2000241 | regulation of reproductive process | 59/4924 | 4.63E-05 | 0.00024227 |
| GO:0060401 | cytosolic calcium ion transport | 57/4924 | 4.64E-05 | 0.000242472 |
| GO:0046822 | regulation of nucleocytoplasmic transport | 41/4924 | 4.73E-05 | 0.000246689 |
| GO:0050885 | neuromuscular process controlling balance | 26/4924 | 4.78E-05 | 0.000249335 |
| GO:0010712 | regulation of collagen metabolic process | 22/4924 | 4.87E-05 | 0.000253687 |
| GO:0051101 | regulation of DNA binding | 40/4924 | 4.96E-05 | 0.000257944 |
| GO:0001885 | endothelial cell development | 27/4924 | 4.96E-05 | 0.000257944 |
| GO:0045911 | positive regulation of DNA recombination | 27/4924 | 4.96E-05 | 0.000257944 |
| GO:0048013 | ephrin receptor signaling pathway | 16/4924 | 5.01E-05 | 0.000260112 |
| GO:0051497 | negative regulation of stress fiber assembly | 16/4924 | 5.01E-05 | 0.000260112 |
| GO:0071383 | cellular response to steroid hormone stimulus | 49/4924 | 5.08E-05 | 0.000263513 |
| GO:0006970 | response to osmotic stress | 32/4924 | 5.15E-05 | 0.000266673 |
| GO:0044774 | mitotic DNA integrity checkpoint signaling | 29/4924 | 5.17E-05 | 0.000267037 |
| GO:0045123 | cellular extravasation | 29/4924 | 5.17E-05 | 0.000267037 |
| GO:0060042 | retina morphogenesis in camera-type eye | 29/4924 | 5.17E-05 | 0.000267037 |
| GO:1901379 | regulation of potassium ion transmembrane transport | 31/4924 | 5.19E-05 | 0.000268031 |
| GO:0010921 | regulation of phosphatase activity | 30/4924 | 5.20E-05 | 0.000268031 |

| | | | | |
|---|---|---|---|---|
| GO:0060415 | muscle tissue morphogenesis | 30/4924 | 5.20E-05 | 0.000268031 |
| GO:0001941 | postsynaptic membrane organization | 20/4924 | 5.33E-05 | 0.000274321 |
| GO:0032965 | regulation of collagen biosynthetic process | 20/4924 | 5.33E-05 | 0.000274321 |
| GO:0033628 | regulation of cell adhesion mediated by integrin | 20/4924 | 5.33E-05 | 0.000274321 |
| GO:0031647 | regulation of protein stability | 81/4924 | 5.38E-05 | 0.000276347 |
| GO:0048709 | oligodendrocyte differentiation | 38/4924 | 5.41E-05 | 0.000277913 |
| GO:0007431 | salivary gland development | 18/4924 | 5.46E-05 | 0.000279601 |
| GO:0035850 | epithelial cell differentiation involved in kidney development | 18/4924 | 5.46E-05 | 0.000279601 |
| GO:2000463 | positive regulation of excitatory postsynaptic potential | 18/4924 | 5.46E-05 | 0.000279601 |
| GO:0055007 | cardiac muscle cell differentiation | 45/4924 | 5.51E-05 | 0.000282013 |
| GO:0001945 | lymph vessel development | 15/4924 | 5.74E-05 | 0.000292986 |
| GO:0007097 | nuclear migration | 15/4924 | 5.74E-05 | 0.000292986 |
| GO:0048535 | lymph node development | 15/4924 | 5.74E-05 | 0.000292986 |
| GO:0032655 | regulation of interleukin-12 production | 24/4924 | 5.75E-05 | 0.000293474 |
| GO:0045667 | regulation of osteoblast differentiation | 47/4924 | 5.80E-05 | 0.000295437 |
| GO:0008344 | adult locomotory behavior | 36/4924 | 5.84E-05 | 0.000297544 |
| GO:0097529 | myeloid leukocyte migration | 63/4924 | 6.01E-05 | 0.000305615 |
| GO:0016573 | histone acetylation | 49/4924 | 6.01E-05 | 0.000305725 |

| | | | | |
|---|---|---|---|---|
| GO:0001658 | branching involved in ureteric bud morphogenesis | 25/4924 | 6.08E-05 | 0.000308407 |
| GO:0055081 | anion homeostasis | 21/4924 | 6.08E-05 | 0.000308407 |
| GO:1902041 | regulation of extrinsic apoptotic signaling pathway via death domain receptors | 21/4924 | 6.08E-05 | 0.000308407 |
| GO:0002467 | germinal center formation | Oct-24 | 6.17E-05 | 0.000312572 |
| GO:0034121 | regulation of toll-like receptor signaling pathway | 26/4924 | 6.33E-05 | 0.00031984 |
| GO:0042733 | embryonic digit morphogenesis | 26/4924 | 6.33E-05 | 0.00031984 |
| GO:1902895 | positive regulation of miRNA transcription | 26/4924 | 6.33E-05 | 0.00031984 |
| GO:0060713 | labyrinthine layer morphogenesis | 14/4924 | 6.42E-05 | 0.000323592 |
| GO:0061437 | renal system vasculature development | 14/4924 | 6.42E-05 | 0.000323592 |
| GO:0061440 | kidney vasculature development | 14/4924 | 6.42E-05 | 0.000323592 |
| GO:2000178 | negative regulation of neural precursor cell proliferation | 14/4924 | 6.42E-05 | 0.000323592 |
| GO:0060236 | regulation of mitotic spindle organization | 17/4924 | 6.55E-05 | 0.000329484 |
| GO:0086004 | regulation of cardiac muscle cell contraction | 17/4924 | 6.55E-05 | 0.000329484 |
| GO:0042554 | superoxide anion generation | 19/4924 | 6.56E-05 | 0.000329484 |
| GO:1902742 | apoptotic process involved in development | 19/4924 | 6.56E-05 | 0.000329484 |
| GO:1905819 | negative regulation of chromosome separation | 19/4924 | 6.56E-05 | 0.000329484 |
| GO:0035418 | protein localization to synapse | 30/4924 | 6.65E-05 | 0.000333507 |
| GO:0043507 | positive regulation of JUN kinase activity | 22/4924 | 6.73E-05 | 0.000337407 |

| | | | | |
|---|---|---|---|---|
| GO:0045861 | negative regulation of proteolysis | 90/4924 | 6.82E-05 | 0.000341258 |
| GO:1903320 | regulation of protein modification by small protein conjugation or removal | 68/4924 | 6.82E-05 | 0.000341258 |
| GO:0055083 | monovalent inorganic anion homeostasis | 13/4924 | 6.96E-05 | 0.000347796 |
| GO:0150105 | protein localization to cell-cell junction | 13/4924 | 6.96E-05 | 0.000347796 |
| GO:0022038 | corpus callosum development | Nov-24 | 7.02E-05 | 0.000349753 |
| GO:0035313 | wound healing, spreading of epidermal cells | Nov-24 | 7.02E-05 | 0.000349753 |
| GO:0045198 | establishment of epithelial cell apical/basal polarity | Nov-24 | 7.02E-05 | 0.000349753 |
| GO:1900452 | regulation of long-term synaptic depression | Nov-24 | 7.02E-05 | 0.000349753 |
| GO:0045217 | cell-cell junction maintenance | Dec-24 | 7.22E-05 | 0.000358572 |
| GO:0060307 | regulation of ventricular cardiac muscle cell membrane repolarization | Dec-24 | 7.22E-05 | 0.000358572 |
| GO:0061081 | positive regulation of myeloid leukocyte cytokine production involved in immune response | Dec-24 | 7.22E-05 | 0.000358572 |
| GO:0072111 | cell proliferation involved in kidney development | Dec-24 | 7.22E-05 | 0.000358572 |
| GO:0071478 | cellular response to radiation | 51/4924 | 7.24E-05 | 0.000358727 |
| GO:2001242 | regulation of intrinsic apoptotic signaling pathway | 51/4924 | 7.24E-05 | 0.000358727 |
| GO:0032757 | positive regulation of interleukin-8 production | 23/4924 | 7.28E-05 | 0.000360044 |
| GO:0050922 | negative regulation of chemotaxis | 23/4924 | 7.28E-05 | 0.000360044 |
| GO:0060443 | mammary gland morphogenesis | 23/4924 | 7.28E-05 | 0.000360044 |
| GO:0034454 | microtubule anchoring at centrosome | Aug-24 | 7.30E-05 | 0.000360145 |

| | | | | |
|---|---|---|---|---|
| GO:0060287 | epithelial cilium movement involved in determination of left/right asymmetry | Aug-24 | 7.30E-05 | 0.000360145 |
| GO:0072102 | glomerulus morphogenesis | Aug-24 | 7.30E-05 | 0.000360145 |
| GO:0016601 | Rac protein signal transduction | 20/4924 | 7.57E-05 | 0.000372707 |
| GO:0045429 | positive regulation of nitric oxide biosynthetic process | 20/4924 | 7.57E-05 | 0.000372707 |
| GO:0061900 | glial cell activation | 20/4924 | 7.57E-05 | 0.000372707 |
| GO:0070167 | regulation of biomineral tissue development | 34/4924 | 7.73E-05 | 0.000380408 |
| GO:0007020 | microtubule nucleation | 16/4924 | 7.77E-05 | 0.000381365 |
| GO:1901099 | negative regulation of signal transduction in absence of ligand | 16/4924 | 7.77E-05 | 0.000381365 |
| GO:2001240 | negative regulation of extrinsic apoptotic signaling pathway in absence of ligand | 16/4924 | 7.77E-05 | 0.000381365 |
| GO:0002761 | regulation of myeloid leukocyte differentiation | 42/4924 | 7.89E-05 | 0.000387015 |
| GO:0090224 | regulation of spindle organization | 18/4924 | 8.03E-05 | 0.000392943 |
| GO:0110110 | positive regulation of animal organ morphogenesis | 18/4924 | 8.03E-05 | 0.000392943 |
| GO:1900744 | regulation of p38MAPK cascade | 18/4924 | 8.03E-05 | 0.000392943 |
| GO:0006801 | superoxide metabolic process | 25/4924 | 8.06E-05 | 0.000393163 |
| GO:0007585 | respiratory gaseous exchange by respiratory system | 25/4924 | 8.06E-05 | 0.000393163 |
| GO:0022029 | telencephalon cell migration | 25/4924 | 8.06E-05 | 0.000393163 |
| GO:0061371 | determination of heart left/right asymmetry | 25/4924 | 8.06E-05 | 0.000393163 |
| GO:0002287 | alpha-beta T cell activation involved in immune response | 26/4924 | 8.30E-05 | 0.000404287 |

| | | | | |
|---|---|---|---|---|
| GO:0043967 | histone H4 acetylation | 26/4924 | 8.30E-05 | 0.000404287 |
| GO:0050671 | positive regulation of lymphocyte proliferation | 44/4924 | 8.36E-05 | 0.00040585 |
| GO:0060759 | regulation of response to cytokine stimulus | 44/4924 | 8.36E-05 | 0.00040585 |
| GO:0010212 | response to ionizing radiation | 41/4924 | 8.36E-05 | 0.00040585 |
| GO:0090090 | negative regulation of canonical Wnt signaling pathway | 41/4924 | 8.36E-05 | 0.00040585 |
| GO:0003281 | ventricular septum development | 30/4924 | 8.45E-05 | 0.000407699 |
| GO:0031397 | negative regulation of protein ubiquitination | 30/4924 | 8.45E-05 | 0.000407699 |
| GO:1901184 | regulation of ERBB signaling pathway | 27/4924 | 8.45E-05 | 0.000407699 |
| GO:1903036 | positive regulation of response to wounding | 27/4924 | 8.45E-05 | 0.000407699 |
| GO:0021795 | cerebral cortex cell migration | 21/4924 | 8.46E-05 | 0.000407699 |
| GO:0038066 | p38MAPK cascade | 21/4924 | 8.46E-05 | 0.000407699 |
| GO:0070169 | positive regulation of biomineral tissue development | 21/4924 | 8.46E-05 | 0.000407699 |
| GO:0072091 | regulation of stem cell proliferation | 21/4924 | 8.46E-05 | 0.000407699 |
| GO:1904036 | negative regulation of epithelial cell apoptotic process | 21/4924 | 8.46E-05 | 0.000407699 |
| GO:1904659 | glucose transmembrane transport | 37/4924 | 8.47E-05 | 0.000408156 |
| GO:0002260 | lymphocyte homeostasis | 29/4924 | 8.52E-05 | 0.000409697 |
| GO:0051965 | positive regulation of synapse assembly | 28/4924 | 8.52E-05 | 0.000409697 |
| GO:0042129 | regulation of T cell proliferation | 53/4924 | 8.53E-05 | 0.000409697 |

| | | | | |
|---|---|---|---|---|
| GO:0051053 | negative regulation of DNA metabolic process | 46/4924 | 8.72E-05 | 0.000418535 |
| GO:0000132 | establishment of mitotic spindle orientation | 15/4924 | 9.10E-05 | 0.000434967 |
| GO:0023058 | adaptation of signaling pathway | 15/4924 | 9.10E-05 | 0.000434967 |
| GO:0048668 | collateral sprouting | 15/4924 | 9.10E-05 | 0.000434967 |
| GO:0051150 | regulation of smooth muscle cell differentiation | 15/4924 | 9.10E-05 | 0.000434967 |
| GO:0060390 | regulation of SMAD protein signal transduction | 15/4924 | 9.10E-05 | 0.000434967 |
| GO:0098703 | calcium ion import across plasma membrane | 15/4924 | 9.10E-05 | 0.000434967 |
| GO:0030514 | negative regulation of BMP signaling pathway | 22/4924 | 9.21E-05 | 0.000439369 |
| GO:0031529 | ruffle organization | 22/4924 | 9.21E-05 | 0.000439369 |
| GO:0002065 | columnar/cuboidal epithelial cell differentiation | 35/4924 | 9.22E-05 | 0.000439959 |
| GO:0030279 | negative regulation of ossification | 19/4924 | 9.40E-05 | 0.000446946 |
| GO:0031952 | regulation of protein autophosphorylation | 19/4924 | 9.40E-05 | 0.000446946 |
| GO:0048536 | spleen development | 19/4924 | 9.40E-05 | 0.000446946 |
| GO:0060323 | head morphogenesis | 19/4924 | 9.40E-05 | 0.000446946 |
| GO:0009615 | response to virus | 85/4924 | 9.45E-05 | 0.000448765 |
| GO:0034341 | response to interferon-gamma | 42/4924 | 9.46E-05 | 0.000449282 |
| GO:0070570 | regulation of neuron projection regeneration | 17/4924 | 9.77E-05 | 0.000463049 |
| GO:2000036 | regulation of stem cell population maintenance | 17/4924 | 9.77E-05 | 0.000463049 |

| | | | | |
|---|---|---|---|---|
| GO:0045682 | regulation of epidermis development | 23/4924 | 9.81E-05 | 0.000464379 |
| GO:1902099 | regulation of metaphase/anaphase transition of cell cycle | 23/4924 | 9.81E-05 | 0.000464379 |
| GO:0032465 | regulation of cytokinesis | 33/4924 | 9.91E-05 | 0.000468466 |
| GO:0060402 | calcium ion transport into cytosol | 51/4924 | 9.91E-05 | 0.000468466 |
| GO:0030522 | intracellular receptor signaling pathway | 60/4924 | 0.000101139 | 0.000477308 |
| GO:0031396 | regulation of protein ubiquitination | 60/4924 | 0.000101139 | 0.000477308 |
| GO:0010660 | regulation of muscle cell apoptotic process | 32/4924 | 0.000102115 | 0.000481553 |
| GO:0008645 | hexose transmembrane transport | 37/4924 | 0.000103335 | 0.000486577 |
| GO:0043401 | steroid hormone mediated signaling pathway | 37/4924 | 0.000103335 | 0.000486577 |
| GO:0099072 | regulation of postsynaptic membrane neurotransmitter receptor levels | 31/4924 | 0.000104733 | 0.000492791 |
| GO:0050819 | negative regulation of coagulation | 20/4924 | 0.000106049 | 0.000496483 |
| GO:0090102 | cochlea development | 20/4924 | 0.000106049 | 0.000496483 |
| GO:0006940 | regulation of smooth muscle contraction | 25/4924 | 0.000106076 | 0.000496483 |
| GO:0042093 | T-helper cell differentiation | 25/4924 | 0.000106076 | 0.000496483 |
| GO:0046513 | ceramide biosynthetic process | 25/4924 | 0.000106076 | 0.000496483 |
| GO:0006665 | sphingolipid metabolic process | 43/4924 | 0.000106147 | 0.000496483 |
| GO:0050728 | negative regulation of inflammatory response | 43/4924 | 0.000106147 | 0.000496483 |
| GO:0098876 | vesicle-mediated transport to the plasma membrane | 43/4924 | 0.000106147 | 0.000496483 |

| | | | | |
|---|---|---|---|---|
| GO:0016081 | synaptic vesicle docking | Sep-24 | 0.000107815 | 0.000502421 |
| GO:0048755 | branching morphogenesis of a nerve | Sep-24 | 0.000107815 | 0.000502421 |
| GO:0051764 | actin crosslink formation | Sep-24 | 0.000107815 | 0.000502421 |
| GO:0090179 | planar cell polarity pathway involved in neural tube closure | Sep-24 | 0.000107815 | 0.000502421 |
| GO:2000105 | positive regulation of DNA-templated DNA replication | Sep-24 | 0.000107815 | 0.000502421 |
| GO:0000079 | regulation of cyclin-dependent protein serine/threonine kinase activity | 26/4924 | 0.000108175 | 0.000503352 |
| GO:0046637 | regulation of alpha-beta T cell differentiation | 26/4924 | 0.000108175 | 0.000503352 |
| GO:0030500 | regulation of bone mineralization | 29/4924 | 0.000108389 | 0.000503976 |
| GO:0007632 | visual behavior | 27/4924 | 0.000109174 | 0.000506263 |
| GO:0022406 | membrane docking | 28/4924 | 0.000109202 | 0.000506263 |
| GO:0050709 | negative regulation of protein secretion | 28/4924 | 0.000109202 | 0.000506263 |
| GO:1902807 | negative regulation of cell cycle G1/S phase transition | 28/4924 | 0.000109202 | 0.000506263 |
| GO:0030216 | keratinocyte differentiation | 45/4924 | 0.000110398 | 0.000511432 |
| GO:0050792 | regulation of viral process | 49/4924 | 0.000115011 | 0.000532411 |
| GO:0010765 | positive regulation of sodium ion transport | 18/4924 | 0.000116208 | 0.000536999 |
| GO:0070527 | platelet aggregation | 21/4924 | 0.000116257 | 0.000536999 |
| GO:0110151 | positive regulation of biomineralization | 21/4924 | 0.000116257 | 0.000536999 |
| GO:0099623 | regulation of cardiac muscle cell membrane repolarization | 13/4924 | 0.000117464 | 0.000542174 |

| | | | | |
|---|---|---|---|---|
| GO:0048679 | regulation of axon regeneration | 16/4924 | 0.000117788 | 0.000542873 |
| GO:1904037 | positive regulation of epithelial cell apoptotic process | 16/4924 | 0.000117788 | 0.000542873 |
| GO:0045582 | positive regulation of T cell differentiation | 34/4924 | 0.000118105 | 0.000543542 |
| GO:0110149 | regulation of biomineralization | 34/4924 | 0.000118105 | 0.000543542 |
| GO:0043543 | protein acylation | 69/4924 | 0.000118253 | 0.000543826 |
| GO:0006476 | protein deacetylation | 38/4924 | 0.000119081 | 0.000547232 |
| GO:0034504 | protein localization to nucleus | 81/4924 | 0.000119801 | 0.000550143 |
| GO:0009791 | post-embryonic development | 41/4924 | 0.000120352 | 0.000552268 |
| GO:0008643 | carbohydrate transport | 46/4924 | 0.000121632 | 0.000557736 |
| GO:0043502 | regulation of muscle adaptation | 33/4924 | 0.000122714 | 0.000562289 |
| GO:0051896 | regulation of protein kinase B signaling | 50/4924 | 0.000124718 | 0.000571057 |
| GO:0022600 | digestive system process | 37/4924 | 0.000125538 | 0.000574395 |
| GO:0051928 | positive regulation of calcium ion transport | 43/4924 | 0.000126233 | 0.000577091 |
| GO:0003356 | regulation of cilium beat frequency | Oct-24 | 0.000126768 | 0.000577091 |
| GO:0031268 | pseudopodium organization | Oct-24 | 0.000126768 | 0.000577091 |
| GO:0051895 | negative regulation of focal adhesion assembly | Oct-24 | 0.000126768 | 0.000577091 |
| GO:0090136 | epithelial cell-cell adhesion | Oct-24 | 0.000126768 | 0.000577091 |
| GO:0150118 | negative regulation of cell-substrate junction organization | Oct-24 | 0.000126768 | 0.000577091 |

| | | | | |
|---|---|---|---|---|
| GO:2001044 | regulation of integrin-mediated signaling pathway | Oct-24 | 0.000126768 | 0.000577091 |
| GO:0034502 | protein localization to chromosome | 32/4924 | 0.000127026 | 0.000577434 |
| GO:1901222 | regulation of NIK/NF-kappaB signaling | 32/4924 | 0.000127026 | 0.000577434 |
| GO:0016202 | regulation of striated muscle tissue development | Dec-24 | 0.00012753 | 0.000579307 |
| GO:0031663 | lipopolysaccharide-mediated signaling pathway | 23/4924 | 0.000130928 | 0.000593084 |
| GO:0050982 | detection of mechanical stimulus | 23/4924 | 0.000130928 | 0.000593084 |
| GO:0021954 | central nervous system neuron development | 31/4924 | 0.000130939 | 0.000593084 |
| GO:0048644 | muscle organ morphogenesis | 31/4924 | 0.000130939 | 0.000593084 |
| GO:0010766 | negative regulation of sodium ion transport | Nov-24 | 0.00013193 | 0.000595856 |
| GO:0045019 | negative regulation of nitric oxide biosynthetic process | Nov-24 | 0.00013193 | 0.000595856 |
| GO:1902430 | negative regulation of amyloid-beta formation | Nov-24 | 0.00013193 | 0.000595856 |
| GO:1904406 | negative regulation of nitric oxide metabolic process | Nov-24 | 0.00013193 | 0.000595856 |
| GO:0046634 | regulation of alpha-beta T cell activation | 36/4924 | 0.00013205 | 0.000595974 |
| GO:0003016 | respiratory system process | 19/4924 | 0.000132646 | 0.000596955 |
| GO:0006024 | glycosaminoglycan biosynthetic process | 19/4924 | 0.000132646 | 0.000596955 |
| GO:0043114 | regulation of vascular permeability | 19/4924 | 0.000132646 | 0.000596955 |
| GO:2000404 | regulation of T cell migration | 19/4924 | 0.000132646 | 0.000596955 |
| GO:0006486 | protein glycosylation | 53/4924 | 0.000133619 | 0.000600477 |

| | | | | |
|---|---|---|---|---|
| GO:0043413 | macromolecule glycosylation | 53/4924 | 0.000133619 | 0.000600477 |
| GO:0031016 | pancreas development | 30/4924 | 0.000134336 | 0.000603267 |
| GO:0051224 | negative regulation of protein transport | 42/4924 | 0.000134771 | 0.000604789 |
| GO:0002294 | CD4-positive, alpha-beta T cell differentiation involved in immune response | 25/4924 | 0.000138445 | 0.000620836 |
| GO:0021549 | cerebellum development | 35/4924 | 0.000138557 | 0.000620897 |
| GO:0007188 | adenylate cyclase-modulating G protein-coupled receptor signaling pathway | 61/4924 | 0.000139646 | 0.000624887 |
| GO:0048545 | response to steroid hormone | 61/4924 | 0.000139646 | 0.000624887 |
| GO:0006919 | activation of cysteine-type endopeptidase activity involved in apoptotic process | 27/4924 | 0.000140025 | 0.000625698 |
| GO:0044773 | mitotic DNA damage checkpoint signaling | 27/4924 | 0.000140025 | 0.000625698 |
| GO:0060706 | cell differentiation involved in embryonic placenta development | 15/4924 | 0.000140548 | 0.000627146 |
| GO:1902656 | calcium ion import into cytosol | 15/4924 | 0.000140548 | 0.000627146 |
| GO:0001782 | B cell homeostasis | 17/4924 | 0.000142952 | 0.000636972 |
| GO:1905332 | positive regulation of morphogenesis of an epithelium | 17/4924 | 0.000142952 | 0.000636972 |
| GO:0009581 | detection of external stimulus | 41/4924 | 0.000143716 | 0.000639045 |
| GO:0016079 | synaptic vesicle exocytosis | 41/4924 | 0.000143716 | 0.000639045 |
| GO:0046620 | regulation of organ growth | 38/4924 | 0.000143722 | 0.000639045 |
| GO:0060538 | skeletal muscle organ development | 59/4924 | 0.000144284 | 0.000641091 |
| GO:0006643 | membrane lipid metabolic process | 52/4924 | 0.000144543 | 0.00064179 |

| | | | | |
|---|---|---|---|---|
| GO:0006821 | chloride transport | 34/4924 | 0.000144987 | 0.000643311 |
| GO:0006687 | glycosphingolipid metabolic process | 20/4924 | 0.000146621 | 0.000649189 |
| GO:0043268 | positive regulation of potassium ion transport | 20/4924 | 0.000146621 | 0.000649189 |
| GO:0045740 | positive regulation of DNA replication | 20/4924 | 0.000146621 | 0.000649189 |
| GO:1904950 | negative regulation of establishment of protein localization | 43/4924 | 0.000149696 | 0.000662337 |
| GO:1903531 | negative regulation of secretion by cell | 49/4924 | 0.000156874 | 0.00069361 |
| GO:1903321 | negative regulation of protein modification by small protein conjugation or removal | 32/4924 | 0.000157256 | 0.000694813 |
| GO:0007062 | sister chromatid cohesion | 21/4924 | 0.000157993 | 0.000695633 |
| GO:0030838 | positive regulation of actin filament polymerization | 21/4924 | 0.000157993 | 0.000695633 |
| GO:0042491 | inner ear auditory receptor cell differentiation | 21/4924 | 0.000157993 | 0.000695633 |
| GO:0045687 | positive regulation of glial cell differentiation | 21/4924 | 0.000157993 | 0.000695633 |
| GO:0048713 | regulation of oligodendrocyte differentiation | 21/4924 | 0.000157993 | 0.000695633 |
| GO:0002831 | regulation of response to biotic stimulus | 91/4924 | 0.000159519 | 0.00070186 |
| GO:0090087 | regulation of peptide transport | 64/4924 | 0.00015986 | 0.000702872 |
| GO:0043500 | muscle adaptation | 39/4924 | 0.000162784 | 0.000715228 |
| GO:0006688 | glycosphingolipid biosynthetic process | 14/4924 | 0.000165421 | 0.000723797 |
| GO:0046885 | regulation of hormone biosynthetic process | 14/4924 | 0.000165421 | 0.000723797 |
| GO:0060563 | neuroepithelial cell differentiation | 14/4924 | 0.000165421 | 0.000723797 |

| | | | | |
|---|---|---|---|---|
| GO:0090025 | regulation of monocyte chemotaxis | 14/4924 | 0.000165421 | 0.000723797 |
| GO:0140058 | neuron projection arborization | 14/4924 | 0.000165421 | 0.000723797 |
| GO:0021955 | central nervous system neuron axonogenesis | 18/4924 | 0.000165423 | 0.000723797 |
| GO:0045071 | negative regulation of viral genome replication | 22/4924 | 0.000166786 | 0.000728244 |
| GO:0045778 | positive regulation of ossification | 22/4924 | 0.000166786 | 0.000728244 |
| GO:0051155 | positive regulation of striated muscle cell differentiation | 22/4924 | 0.000166786 | 0.000728244 |
| GO:0043266 | regulation of potassium ion transport | 35/4924 | 0.000168777 | 0.00073643 |
| GO:0002703 | regulation of leukocyte mediated immunity | 77/4924 | 0.000169689 | 0.000739896 |
| GO:0045995 | regulation of embryonic development | 29/4924 | 0.000172379 | 0.000751105 |
| GO:0010658 | striated muscle cell apoptotic process | 23/4924 | 0.000173131 | 0.00075334 |
| GO:0043407 | negative regulation of MAP kinase activity | 23/4924 | 0.000173131 | 0.00075334 |
| GO:0090313 | regulation of protein targeting to membrane | 16/4924 | 0.000174754 | 0.000759354 |
| GO:1904707 | positive regulation of vascular associated smooth muscle cell proliferation | 16/4924 | 0.000174754 | 0.000759354 |
| GO:0033209 | tumor necrosis factor-mediated signaling pathway | 28/4924 | 0.000175929 | 0.000763934 |
| GO:0010469 | regulation of signaling receptor activity | 43/4924 | 0.000177025 | 0.000768164 |
| GO:0002293 | alpha-beta T cell differentiation involved in immune response | 25/4924 | 0.000179308 | 0.000776723 |
| GO:2000134 | negative regulation of G1/S transition of mitotic cell cycle | 25/4924 | 0.000179308 | 0.000776723 |
| GO:0070085 | glycosylation | 58/4924 | 0.000179367 | 0.000776723 |

| | | | | |
|---|---|---|---|---|
| GO:0007492 | endoderm development | 26/4924 | 0.000179622 | 0.000777293 |
| GO:0015749 | monosaccharide transmembrane transport | 37/4924 | 0.000183255 | 0.000792471 |
| GO:0003179 | heart valve morphogenesis | 19/4924 | 0.000184605 | 0.000796668 |
| GO:0007080 | mitotic metaphase plate congression | 19/4924 | 0.000184605 | 0.000796668 |
| GO:0030195 | negative regulation of blood coagulation | 19/4924 | 0.000184605 | 0.000796668 |
| GO:0010770 | positive regulation of cell morphogenesis involved in differentiation | 33/4924 | 0.000185605 | 0.000797252 |
| GO:0045921 | positive regulation of exocytosis | 33/4924 | 0.000185605 | 0.000797252 |
| GO:0046849 | bone remodeling | 33/4924 | 0.000185605 | 0.000797252 |
| GO:0034333 | adherens junction assembly | Aug-24 | 0.000186004 | 0.000797252 |
| GO:0035331 | negative regulation of hippo signaling | Aug-24 | 0.000186004 | 0.000797252 |
| GO:0051798 | positive regulation of hair follicle development | Aug-24 | 0.000186004 | 0.000797252 |
| GO:0060433 | bronchus development | Aug-24 | 0.000186004 | 0.000797252 |
| GO:0072176 | nephric duct development | Aug-24 | 0.000186004 | 0.000797252 |
| GO:0098911 | regulation of ventricular cardiac muscle cell action potential | Aug-24 | 0.000186004 | 0.000797252 |
| GO:1900272 | negative regulation of long-term synaptic potentiation | Aug-24 | 0.000186004 | 0.000797252 |
| GO:0008217 | regulation of blood pressure | 59/4924 | 0.000188433 | 0.000807114 |
| GO:0090276 | regulation of peptide hormone secretion | 62/4924 | 0.000189495 | 0.00081111 |
| GO:0035809 | regulation of urine volume | 13/4924 | 0.000191168 | 0.000815503 |

| | | | | |
|---|---|---|---|---|
| GO:0051016 | barbed-end actin filament capping | 13/4924 | 0.000191168 | 0.000815503 |
| GO:0060716 | labyrinthine layer blood vessel development | 13/4924 | 0.000191168 | 0.000815503 |
| GO:0072567 | chemokine (C-X-C motif) ligand 2 production | 13/4924 | 0.000191168 | 0.000815503 |
| GO:2000341 | regulation of chemokine (C-X-C motif) ligand 2 production | 13/4924 | 0.000191168 | 0.000815503 |
| GO:0002791 | regulation of peptide secretion | 63/4924 | 0.000197464 | 0.000841793 |
| GO:0061028 | establishment of endothelial barrier | 20/4924 | 0.000200259 | 0.000852552 |
| GO:0090279 | regulation of calcium ion import | 20/4924 | 0.000200259 | 0.000852552 |
| GO:0035282 | segmentation | 35/4924 | 0.000204779 | 0.000871209 |
| GO:0060603 | mammary gland duct morphogenesis | 17/4924 | 0.000205543 | 0.000872691 |
| GO:0086065 | cell communication involved in cardiac conduction | 17/4924 | 0.000205543 | 0.000872691 |
| GO:2000279 | negative regulation of DNA biosynthetic process | 17/4924 | 0.000205543 | 0.000872691 |
| GO:0048799 | animal organ maturation | 15/4924 | 0.000211935 | 0.000898018 |
| GO:0050654 | chondroitin sulfate proteoglycan metabolic process | 15/4924 | 0.000211935 | 0.000898018 |
| GO:0060306 | regulation of membrane repolarization | 15/4924 | 0.000211935 | 0.000898018 |
| GO:0033135 | regulation of peptidyl-serine phosphorylation | 47/4924 | 0.000212974 | 0.000901812 |
| GO:0006473 | protein acetylation | 59/4924 | 0.000214816 | 0.000908746 |
| GO:0034405 | response to fluid shear stress | Dec-24 | 0.000215445 | 0.000908746 |
| GO:0055064 | chloride ion homeostasis | Dec-24 | 0.000215445 | 0.000908746 |

| | | | | |
|---|---|---|---|---|
| GO:0060972 | left/right pattern formation | Dec-24 | 0.000215445 | 0.000908746 |
| GO:0098901 | regulation of cardiac muscle cell action potential | Dec-24 | 0.000215445 | 0.000908746 |
| GO:0000077 | DNA damage checkpoint signaling | 34/4924 | 0.000215764 | 0.000908746 |
| GO:0042310 | vasoconstriction | 34/4924 | 0.000215764 | 0.000908746 |
| GO:0050886 | endocrine process | 34/4924 | 0.000215764 | 0.000908746 |
| GO:0001776 | leukocyte homeostasis | 37/4924 | 0.000220229 | 0.000926312 |
| GO:0042177 | negative regulation of protein catabolic process | 37/4924 | 0.000220229 | 0.000926312 |
| GO:0002548 | monocyte chemotaxis | 22/4924 | 0.000221194 | 0.000928323 |
| GO:0010659 | cardiac muscle cell apoptotic process | 22/4924 | 0.000221194 | 0.000928323 |
| GO:1900024 | regulation of substrate adhesion-dependent cell spreading | 22/4924 | 0.000221194 | 0.000928323 |
| GO:0048708 | astrocyte differentiation | 28/4924 | 0.000221296 | 0.000928323 |
| GO:0071347 | cellular response to interleukin-1 | 27/4924 | 0.000225889 | 0.000946963 |
| GO:0010715 | regulation of extracellular matrix disassembly | Sep-24 | 0.00022854 | 0.000954899 |
| GO:0031269 | pseudopodium assembly | Sep-24 | 0.00022854 | 0.000954899 |
| GO:0032530 | regulation of microvillus organization | Sep-24 | 0.00022854 | 0.000954899 |
| GO:0090178 | regulation of establishment of planar polarity involved in neural tube closure | Sep-24 | 0.00022854 | 0.000954899 |
| GO:2000095 | regulation of Wnt signaling pathway, planar cell polarity pathway | Sep-24 | 0.00022854 | 0.000954899 |
| GO:0006305 | DNA alkylation | 26/4924 | 0.000229064 | 0.000955189 |

| | | | | |
|---|---|---|---|---|
| GO:0006306 | DNA methylation | 26/4924 | 0.000229064 | 0.000955189 |
| GO:1904029 | regulation of cyclin-dependent protein kinase activity | 26/4924 | 0.000229064 | 0.000955189 |
| GO:0048168 | regulation of neuronal synaptic plasticity | 25/4924 | 0.000230515 | 0.000959338 |
| GO:1990868 | response to chemokine | 25/4924 | 0.000230515 | 0.000959338 |
| GO:1990869 | cellular response to chemokine | 25/4924 | 0.000230515 | 0.000959338 |
| GO:0097178 | ruffle assembly | 18/4924 | 0.000231948 | 0.000964029 |
| GO:1902991 | regulation of amyloid precursor protein catabolic process | 18/4924 | 0.000231948 | 0.000964029 |
| GO:0006268 | DNA unwinding involved in DNA replication | Nov-24 | 0.000234277 | 0.000971148 |
| GO:0007252 | I-kappaB phosphorylation | Nov-24 | 0.000234277 | 0.000971148 |
| GO:0090050 | positive regulation of cell migration involved in sprouting angiogenesis | Nov-24 | 0.000234277 | 0.000971148 |
| GO:1903909 | regulation of receptor clustering | Nov-24 | 0.000234277 | 0.000971148 |
| GO:0062207 | regulation of pattern recognition receptor signaling pathway | 32/4924 | 0.000237671 | 0.000984569 |
| GO:0006978 | DNA damage response, signal transduction by p53 class mediator resulting in transcription of p21 class mediator | Oct-24 | 0.000241492 | 0.000993871 |
| GO:0010470 | regulation of gastrulation | Oct-24 | 0.000241492 | 0.000993871 |
| GO:0030033 | microvillus assembly | Oct-24 | 0.000241492 | 0.000993871 |
| GO:0035330 | regulation of hippo signaling | Oct-24 | 0.000241492 | 0.000993871 |
| GO:0060391 | positive regulation of SMAD protein signal transduction | Oct-24 | 0.000241492 | 0.000993871 |
| GO:0061318 | renal filtration cell differentiation | Oct-24 | 0.000241492 | 0.000993871 |

| | | | | |
|---|---|---|---|---|
| GO:0072112 | glomerular visceral epithelial cell differentiation | Oct-24 | 0.000241492 | 0.000993871 |
| GO:0072311 | glomerular epithelial cell differentiation | Oct-24 | 0.000241492 | 0.000993871 |
| GO:1901881 | positive regulation of protein depolymerization | Oct-24 | 0.000241492 | 0.000993871 |
| GO:2000319 | regulation of T-helper 17 cell differentiation | Oct-24 | 0.000241492 | 0.000993871 |
| GO:0045598 | regulation of fat cell differentiation | 43/4924 | 0.00024555 | 0.00100991 |
| GO:0089718 | amino acid import across plasma membrane | 19/4924 | 0.000253562 | 0.001040833 |
| GO:1900047 | negative regulation of hemostasis | 19/4924 | 0.000253562 | 0.001040833 |
| GO:0021799 | cerebral cortex radially oriented cell migration | 16/4924 | 0.00025422 | 0.001040833 |
| GO:0070528 | protein kinase C signaling | 16/4924 | 0.00025422 | 0.001040833 |
| GO:0072210 | metanephric nephron development | 16/4924 | 0.00025422 | 0.001040833 |
| GO:0021952 | central nervous system projection neuron axonogenesis | 14/4924 | 0.000254389 | 0.001040833 |
| GO:0099622 | cardiac muscle cell membrane repolarization | 14/4924 | 0.000254389 | 0.001040833 |
| GO:1905523 | positive regulation of macrophage migration | 14/4924 | 0.000254389 | 0.001040833 |
| GO:0018394 | peptidyl-lysine acetylation | 52/4924 | 0.00025683 | 0.001050137 |
| GO:0009582 | detection of abiotic stimulus | 40/4924 | 0.000257546 | 0.001052382 |
| GO:0070828 | heterochromatin organization | 30/4924 | 0.00025853 | 0.001055722 |
| GO:0060349 | bone morphogenesis | 34/4924 | 0.000261608 | 0.001067597 |
| GO:1901019 | regulation of calcium ion transmembrane transporter activity | 29/4924 | 0.000268094 | 0.001092653 |

| | | | | |
|---|---|---|---|---|
| GO:1904427 | positive regulation of calcium ion transmembrane transport | 29/4924 | 0.000268094 | 0.001092653 |
| GO:0046942 | carboxylic acid transport | 73/4924 | 0.000268722 | 0.001094505 |
| GO:0007131 | reciprocal meiotic recombination | 20/4924 | 0.000270367 | 0.001099076 |
| GO:0060412 | ventricular septum morphogenesis | 20/4924 | 0.000270367 | 0.001099076 |
| GO:0140527 | reciprocal homologous recombination | 20/4924 | 0.000270367 | 0.001099076 |
| GO:0006304 | DNA modification | 33/4924 | 0.000275892 | 0.001120809 |
| GO:0060998 | regulation of dendritic spine development | 28/4924 | 0.000276759 | 0.001122886 |
| GO:1902117 | positive regulation of organelle assembly | 28/4924 | 0.000276759 | 0.001122886 |
| GO:0050764 | regulation of phagocytosis | 36/4924 | 0.000280589 | 0.001137693 |
| GO:1900271 | regulation of long-term synaptic potentiation | 21/4924 | 0.000282596 | 0.001144358 |
| GO:1903533 | regulation of protein targeting | 21/4924 | 0.000282596 | 0.001144358 |
| GO:0071887 | leukocyte apoptotic process | 41/4924 | 0.000283646 | 0.001147875 |
| GO:0006475 | internal protein amino acid acetylation | 49/4924 | 0.00028424 | 0.001148804 |
| GO:0018393 | internal peptidyl-lysine acetylation | 49/4924 | 0.00028424 | 0.001148804 |
| GO:0055067 | monovalent inorganic cation homeostasis | 47/4924 | 0.000287908 | 0.001162881 |
| GO:0097530 | granulocyte migration | 45/4924 | 0.000289317 | 0.001167824 |
| GO:2001021 | negative regulation of response to DNA damage stimulus | 26/4924 | 0.000290184 | 0.001170575 |
| GO:0140115 | export across plasma membrane | 22/4924 | 0.000290628 | 0.001171097 |

| | | | | |
|---|---|---|---|---|
| GO:2000249 | regulation of actin cytoskeleton reorganization | 17/4924 | 0.000290685 | 0.001171097 |
| GO:0071621 | granulocyte chemotaxis | 38/4924 | 0.000294769 | 0.001186793 |
| GO:1901224 | positive regulation of NIK/NF-kappaB signaling | 24/4924 | 0.000295965 | 0.00119085 |
| GO:1903076 | regulation of protein localization to plasma membrane | 35/4924 | 0.000298014 | 0.001198328 |
| GO:0015711 | organic anion transport | 86/4924 | 0.000300494 | 0.001207298 |
| GO:0033688 | regulation of osteoblast proliferation | 13/4924 | 0.000301202 | 0.001207298 |
| GO:0051767 | nitric-oxide synthase biosynthetic process | 13/4924 | 0.000301202 | 0.001207298 |
| GO:0051769 | regulation of nitric-oxide synthase biosynthetic process | 13/4924 | 0.000301202 | 0.001207298 |
| GO:0061484 | hematopoietic stem cell homeostasis | 13/4924 | 0.000301202 | 0.001207298 |
| GO:0030148 | sphingolipid biosynthetic process | 31/4924 | 0.000304438 | 0.001218909 |
| GO:0001959 | regulation of cytokine-mediated signaling pathway | 40/4924 | 0.000304486 | 0.001218909 |
| GO:0097553 | calcium ion transmembrane import into cytosol | 44/4924 | 0.000312335 | 0.001248714 |
| GO:0003149 | membranous septum morphogenesis | Jul-24 | 0.000313713 | 0.001248714 |
| GO:0006971 | hypotonic response | Jul-24 | 0.000313713 | 0.001248714 |
| GO:0032836 | glomerular basement membrane development | Jul-24 | 0.000313713 | 0.001248714 |
| GO:0048671 | negative regulation of collateral sprouting | Jul-24 | 0.000313713 | 0.001248714 |
| GO:0060484 | lung-associated mesenchyme development | Jul-24 | 0.000313713 | 0.001248714 |
| GO:0072203 | cell proliferation involved in metanephros development | Jul-24 | 0.000313713 | 0.001248714 |

| | | | | |
|---|---|---|---|---|
| GO:0090091 | positive regulation of extracellular matrix disassembly | Jul-24 | 0.000313713 | 0.001248714 |
| GO:0099640 | axo-dendritic protein transport | Jul-24 | 0.000313713 | 0.001248714 |
| GO:0030890 | positive regulation of B cell proliferation | 18/4924 | 0.000320631 | 0.001275443 |
| GO:0030073 | insulin secretion | 61/4924 | 0.0003398 | 0.001350842 |
| GO:0019229 | regulation of vasoconstriction | 28/4924 | 0.000344189 | 0.001366569 |
| GO:0032729 | positive regulation of interferon-gamma production | 28/4924 | 0.000344189 | 0.001366569 |
| GO:0034219 | carbohydrate transmembrane transport | 38/4924 | 0.000349883 | 0.001385508 |
| GO:0034114 | regulation of heterotypic cell-cell adhesion | Dec-24 | 0.000350058 | 0.001385508 |
| GO:0035024 | negative regulation of Rho protein signal transduction | Dec-24 | 0.000350058 | 0.001385508 |
| GO:0044331 | cell-cell adhesion mediated by cadherin | Dec-24 | 0.000350058 | 0.001385508 |
| GO:0045684 | positive regulation of epidermis development | Dec-24 | 0.000350058 | 0.001385508 |
| GO:0071241 | cellular response to inorganic substance | 62/4924 | 0.000351664 | 0.001390993 |
| GO:0045132 | meiotic chromosome segregation | 32/4924 | 0.000352856 | 0.001394833 |
| GO:0014855 | striated muscle cell proliferation | 27/4924 | 0.000355514 | 0.001404457 |
| GO:0035825 | homologous recombination | 20/4924 | 0.00036102 | 0.001424426 |
| GO:0048701 | embryonic cranial skeleton morphogenesis | 20/4924 | 0.00036102 | 0.001424426 |
| GO:0090049 | regulation of cell migration involved in sprouting angiogenesis | 16/4924 | 0.00036312 | 0.001431814 |
| GO:1903900 | regulation of viral life cycle | 42/4924 | 0.000363441 | 0.001432184 |

| | | | | |
|---|---|---|---|---|
| GO:0015849 | organic acid transport | 79/4924 | 0.000371026 | 0.00146116 |
| GO:0006672 | ceramide metabolic process | 31/4924 | 0.0003715 36 | 0.001462255 |
| GO:0043666 | regulation of phosphoprotein phosphatase activity | 21/4924 | 0.000372302 | 0.001462468 |
| GO:0071103 | DNA conformation change | 21/4924 | 0.000372302 | 0.001462468 |
| GO:0034394 | protein localization to cell surface | 25/4924 | 0.000372981 | 0.001462468 |
| GO:0046824 | positive regulation of nucleocytoplasmic transport | 25/4924 | 0.000372981 | 0.001462468 |
| GO:0071867 | response to monoamine | 25/4924 | 0.000372981 | 0.001462468 |
| GO:0071869 | response to catecholamine | 25/4924 | 0.000372981 | 0.001462468 |
| GO:0022037 | metencephalon development | 37/4924 | 0.000374465 | 0.001467373 |
| GO:0006949 | syncytium formation | 24/4924 | 0.00037815 | 0.001477532 |
| GO:0042246 | tissue regeneration | 24/4924 | 0.00037815 | 0.001477532 |
| GO:0048488 | synaptic vesicle endocytosis | 24/4924 | 0.00037815 | 0.001477532 |
| GO:0140238 | presynaptic endocytosis | 24/4924 | 0.00037815 | 0.001477532 |
| GO:0042698 | ovulation cycle | 22/4924 | 0.000378464 | 0.001477532 |
| GO:0046850 | regulation of bone remodeling | 22/4924 | 0.000378464 | 0.001477532 |
| GO:0071346 | cellular response to interferon-gamma | 34/4924 | 0.000380091 | 0.001481515 |
| GO:0090311 | regulation of protein deacetylation | 23/4924 | 0.000380189 | 0.001481515 |
| GO:1903307 | positive regulation of regulated secretory pathway | 23/4924 | 0.000380189 | 0.001481515 |

| | | | | |
|---|---|---|---|---|
| GO:0006026 | aminoglycan catabolic process | 14/4924 | 0.000381474 | 0.001482856 |
| GO:0060074 | synapse maturation | 14/4924 | 0.000381474 | 0.001482856 |
| GO:0060561 | apoptotic process involved in morphogenesis | 14/4924 | 0.000381474 | 0.001482856 |
| GO:0060977 | coronary vasculature morphogenesis | 14/4924 | 0.000381474 | 0.001482856 |
| GO:0035303 | regulation of dephosphorylation | 39/4924 | 0.000385653 | 0.001498178 |
| GO:0061053 | somite development | 30/4924 | 0.000390092 | 0.001514489 |
| GO:0017121 | plasma membrane phospholipid scrambling | Nov-24 | 0.000396328 | 0.001531156 |
| GO:0036158 | outer dynein arm assembly | Nov-24 | 0.000396328 | 0.001531156 |
| GO:0046325 | negative regulation of glucose import | Nov-24 | 0.000396328 | 0.001531156 |
| GO:0051770 | positive regulation of nitric-oxide synthase biosynthetic process | Nov-24 | 0.000396328 | 0.001531156 |
| GO:0060602 | branch elongation of an epithelium | Nov-24 | 0.000396328 | 0.001531156 |
| GO:0090026 | positive regulation of monocyte chemotaxis | Nov-24 | 0.000396328 | 0.001531156 |
| GO:2000810 | regulation of bicellular tight junction assembly | Nov-24 | 0.000396328 | 0.001531156 |
| GO:2001224 | positive regulation of neuron migration | Nov-24 | 0.000396328 | 0.001531156 |
| GO:0032924 | activin receptor signaling pathway | 17/4924 | 0.000404765 | 0.001560884 |
| GO:0072538 | T-helper 17 type immune response | 17/4924 | 0.000404765 | 0.001560884 |
| GO:0098801 | regulation of renal system process | 17/4924 | 0.000404765 | 0.001560884 |
| GO:0006271 | DNA strand elongation involved in DNA replication | Aug-24 | 0.000410924 | 0.001578839 |

| | | | | |
|---|---|---|---|---|
| GO:0043589 | skin morphogenesis | Aug-24 | 0.000410924 | 0.001578839 |
| GO:0045618 | positive regulation of keratinocyte differentiation | Aug-24 | 0.000410924 | 0.001578839 |
| GO:0051315 | attachment of mitotic spindle microtubules to kinetochore | Aug-24 | 0.000410924 | 0.001578839 |
| GO:0072110 | glomerular mesangial cell proliferation | Aug-24 | 0.000410924 | 0.001578839 |
| GO:1903651 | positive regulation of cytoplasmic transport | Aug-24 | 0.000410924 | 0.001578839 |
| GO:2001252 | positive regulation of chromosome organization | 32/4924 | 0.00042719 | 0.001640338 |
| GO:0000212 | meiotic spindle organization | Oct-24 | 0.000431786 | 0.001650948 |
| GO:0010544 | negative regulation of platelet activation | Oct-24 | 0.000431786 | 0.001650948 |
| GO:0021756 | striatum development | Oct-24 | 0.000431786 | 0.001650948 |
| GO:0034111 | negative regulation of homotypic cell-cell adhesion | Oct-24 | 0.000431786 | 0.001650948 |
| GO:0042772 | DNA damage response, signal transduction resulting in transcription | Oct-24 | 0.000431786 | 0.001650948 |
| GO:0071636 | positive regulation of transforming growth factor beta production | Oct-24 | 0.000431786 | 0.001650948 |
| GO:1900117 | regulation of execution phase of apoptosis | Oct-24 | 0.000431786 | 0.001650948 |
| GO:0010543 | regulation of platelet activation | 18/4924 | 0.000437311 | 0.001670045 |
| GO:0120163 | negative regulation of cold-induced thermogenesis | 18/4924 | 0.000437311 | 0.001670045 |
| GO:0048469 | cell maturation | 58/4924 | 0.000440436 | 0.001680961 |
| GO:0048681 | negative regulation of axon regeneration | Sep-24 | 0.000443084 | 0.001684949 |
| GO:0060026 | convergent extension | Sep-24 | 0.000443084 | 0.001684949 |

| | | | | |
|---|---|---|---|---|
| GO:0090128 | regulation of synapse maturation | Sep-24 | 0.000443084 | 0.001684949 |
| GO:0090177 | establishment of planar polarity involved in neural tube closure | Sep-24 | 0.000443084 | 0.001684949 |
| GO:1901722 | regulation of cell proliferation involved in kidney development | Sep-24 | 0.000443084 | 0.001684949 |
| GO:1902043 | positive regulation of extrinsic apoptotic signaling pathway via death domain receptors | Sep-24 | 0.000443084 | 0.001684949 |
| GO:0008045 | motor neuron axon guidance | 15/4924 | 0.00045174 | 0.00171373 |
| GO:0022616 | DNA strand elongation | 15/4924 | 0.00045174 | 0.00171373 |
| GO:0034122 | negative regulation of toll-like receptor signaling pathway | 15/4924 | 0.00045174 | 0.00171373 |
| GO:0051491 | positive regulation of filopodium assembly | 15/4924 | 0.00045174 | 0.00171373 |
| GO:2000106 | regulation of leukocyte apoptotic process | 34/4924 | 0.000455541 | 0.001727111 |
| GO:0090330 | regulation of platelet aggregation | 13/4924 | 0.000460911 | 0.001746286 |
| GO:0032392 | DNA geometric change | 19/4924 | 0.000461152 | 0.001746286 |
| GO:0002237 | response to molecule of bacterial origin | 96/4924 | 0.000464912 | 0.001759467 |
| GO:0042982 | amyloid precursor protein metabolic process | 25/4924 | 0.000469661 | 0.001776373 |
| GO:0002312 | B cell activation involved in immune response | 30/4924 | 0.000475773 | 0.001798411 |
| GO:0010596 | negative regulation of endothelial cell migration | 20/4924 | 0.000477036 | 0.001801025 |
| GO:0048546 | digestive tract morphogenesis | 20/4924 | 0.000477036 | 0.001801025 |
| GO:0030888 | regulation of B cell proliferation | 24/4924 | 0.000479687 | 0.001808866 |
| GO:1903556 | negative regulation of tumor necrosis factor superfamily cytokine production | 24/4924 | 0.000479687 | 0.001808866 |

| | | | | |
|---|---|---|---|---|
| GO:0048704 | embryonic skeletal system morphogenesis | 33/4924 | 0.000484819 | 0.001827125 |
| GO:0043388 | positive regulation of DNA binding | 21/4924 | 0.000485877 | 0.001827183 |
| GO:0045599 | negative regulation of fat cell differentiation | 21/4924 | 0.000485877 | 0.001827183 |
| GO:0071622 | regulation of granulocyte chemotaxis | 21/4924 | 0.000485877 | 0.001827183 |
| GO:0030330 | DNA damage response, signal transduction by p53 class mediator | 23/4924 | 0.000486283 | 0.001827183 |
| GO:0060986 | endocrine hormone secretion | 23/4924 | 0.000486283 | 0.001827183 |
| GO:0035315 | hair cell differentiation | 22/4924 | 0.000488642 | 0.001834956 |
| GO:0031058 | positive regulation of histone modification | 35/4924 | 0.000508999 | 0.001908396 |
| GO:0051897 | positive regulation of protein kinase B signaling | 35/4924 | 0.000508999 | 0.001908396 |
| GO:0051251 | positive regulation of lymphocyte activation | 103/4924 | 0.000509107 | 0.001908396 |
| GO:2001243 | negative regulation of intrinsic apoptotic signaling pathway | 31/4924 | 0.000546054 | 0.002045675 |
| GO:0021801 | cerebral cortex radial glia-guided migration | Dec-24 | 0.000549424 | 0.002054637 |
| GO:0022030 | telencephalon glial cell migration | Dec-24 | 0.000549424 | 0.002054637 |
| GO:0070293 | renal absorption | Dec-24 | 0.000549424 | 0.002054637 |
| GO:0017145 | stem cell division | 17/4924 | 0.000555462 | 0.002074757 |
| GO:0061098 | positive regulation of protein tyrosine kinase activity | 17/4924 | 0.000555462 | 0.002074757 |
| GO:0042593 | glucose homeostasis | 72/4924 | 0.000557738 | 0.002082023 |
| GO:0051642 | centrosome localization | 14/4924 | 0.000558998 | 0.002085492 |

| | | | | |
|---|---|---|---|---|
| GO:0009880 | embryonic pattern specification | 26/4924 | 0.000568162 | 0.002117176 |
| GO:1904888 | cranial skeletal system development | 26/4924 | 0.000568162 | 0.002117176 |
| GO:0048010 | vascular endothelial growth factor receptor signaling pathway | 18/4924 | 0.000588934 | 0.002193285 |
| GO:0055008 | cardiac muscle tissue morphogenesis | 24/4924 | 0.000604272 | 0.002245101 |
| GO:0071868 | cellular response to monoamine stimulus | 24/4924 | 0.000604272 | 0.002245101 |
| GO:0071870 | cellular response to catecholamine stimulus | 24/4924 | 0.000604272 | 0.002245101 |
| GO:1903317 | regulation of protein maturation | 24/4924 | 0.000604272 | 0.002245101 |
| GO:0000768 | syncytium formation by plasma membrane fusion | 23/4924 | 0.000617314 | 0.002289509 |
| GO:0060389 | pathway-restricted SMAD protein phosphorylation | 23/4924 | 0.000617314 | 0.002289509 |
| GO:0140253 | cell-cell fusion | 23/4924 | 0.000617314 | 0.002289509 |
| GO:0033500 | carbohydrate homeostasis | 72/4924 | 0.000618401 | 0.002291079 |
| GO:0045833 | negative regulation of lipid metabolic process | 32/4924 | 0.000618464 | 0.002291079 |
| GO:0007519 | skeletal muscle tissue development | 54/4924 | 0.00062259 | 0.00230501 |
| GO:0043090 | amino acid import | 20/4924 | 0.000624058 | 0.002306381 |
| GO:0061001 | regulation of dendritic spine morphogenesis | 20/4924 | 0.000624058 | 0.002306381 |
| GO:0071470 | cellular response to osmotic stress | 20/4924 | 0.000624058 | 0.002306381 |
| GO:0060393 | regulation of pathway-restricted SMAD protein phosphorylation | 22/4924 | 0.000625732 | 0.002311215 |
| GO:0007095 | mitotic G2 DNA damage checkpoint signaling | 15/4924 | 0.000640594 | 0.002363338 |

| | | | | |
|---|---|---|---|---|
| GO:0045746 | negative regulation of Notch signaling pathway | 15/4924 | 0.000640594 | 0.002363338 |
| GO:0006027 | glycosaminoglycan catabolic process | Nov-24 | 0.000642726 | 0.002364286 |
| GO:0072202 | cell differentiation involved in metanephros development | Nov-24 | 0.000642726 | 0.002364286 |
| GO:0099149 | regulation of postsynaptic neurotransmitter receptor internalization | Nov-24 | 0.000642726 | 0.002364286 |
| GO:1902894 | negative regulation of miRNA transcription | Nov-24 | 0.000642726 | 0.002364286 |
| GO:1902992 | negative regulation of amyloid precursor protein catabolic process | Nov-24 | 0.000642726 | 0.002364286 |
| GO:0031056 | regulation of histone modification | 52/4924 | 0.000652659 | 0.002399424 |
| GO:0046530 | photoreceptor cell differentiation | 27/4924 | 0.000672348 | 0.002470368 |
| GO:0010800 | positive regulation of peptidyl-threonine phosphorylation | 13/4924 | 0.000686873 | 0.002516406 |
| GO:0045724 | positive regulation of cilium assembly | 13/4924 | 0.000686873 | 0.002516406 |
| GO:0045932 | negative regulation of muscle contraction | 13/4924 | 0.000686873 | 0.002516406 |
| GO:0060343 | trabecula formation | 13/4924 | 0.000686873 | 0.002516406 |
| GO:0070977 | bone maturation | 13/4924 | 0.000686873 | 0.002516406 |
| GO:0045739 | positive regulation of DNA repair | 33/4924 | 0.0006924 | 0.002535182 |
| GO:0030879 | mammary gland development | 45/4924 | 0.000695077 | 0.00254203 |
| GO:0050714 | positive regulation of protein secretion | 45/4924 | 0.000695077 | 0.00254203 |
| GO:0120162 | positive regulation of cold-induced thermogenesis | 30/4924 | 0.00069814 | 0.002551751 |
| GO:0071375 | cellular response to peptide hormone stimulus | 67/4924 | 0.000701594 | 0.002562804 |

| | | | | |
|---|---|---|---|---|
| GO:0036465 | synaptic vesicle recycling | 26/4924 | 0.000702383 | 0.002562804 |
| GO:0045670 | regulation of osteoclast differentiation | 26/4924 | 0.000702383 | 0.002562804 |
| GO:0035886 | vascular associated smooth muscle cell differentiation | 16/4924 | 0.000704686 | 0.002569718 |
| GO:0010763 | positive regulation of fibroblast migration | Oct-24 | 0.000731643 | 0.002661865 |
| GO:0033599 | regulation of mammary gland epithelial cell proliferation | Oct-24 | 0.000731643 | 0.002661865 |
| GO:0034349 | glial cell apoptotic process | Oct-24 | 0.000731643 | 0.002661865 |
| GO:1904754 | positive regulation of vascular associated smooth muscle cell migration | Oct-24 | 0.000731643 | 0.002661865 |
| GO:0001678 | cellular glucose homeostasis | 46/4924 | 0.000733801 | 0.002664711 |
| GO:0010968 | regulation of microtubule nucleation | Jul-24 | 0.000734962 | 0.002664711 |
| GO:0032342 | aldosterone biosynthetic process | Jul-24 | 0.000734962 | 0.002664711 |
| GO:0061517 | macrophage proliferation | Jul-24 | 0.000734962 | 0.002664711 |
| GO:0072178 | nephric duct morphogenesis | Jul-24 | 0.000734962 | 0.002664711 |
| GO:0120197 | mucociliary clearance | Jul-24 | 0.000734962 | 0.002664711 |
| GO:0030261 | chromosome condensation | 17/4924 | 0.000751882 | 0.002722926 |
| GO:0042771 | intrinsic apoptotic signaling pathway in response to DNA damage by p53 class mediator | 17/4924 | 0.000751882 | 0.002722926 |
| GO:0001541 | ovarian follicle development | 24/4924 | 0.000756128 | 0.00273516 |
| GO:0043954 | cellular component maintenance | 24/4924 | 0.000756128 | 0.00273516 |
| GO:0019751 | polyol metabolic process | 34/4924 | 0.000767305 | 0.002774 |

| | | | | |
|---|---|---|---|---|
| GO:0030183 | B cell differentiation | 47/4924 | 0.000771775 | 0.002788561 |
| GO:0032507 | maintenance of protein location in cell | 23/4924 | 0.000777988 | 0.002807795 |
| GO:0070613 | regulation of protein processing | 23/4924 | 0.000777988 | 0.002807795 |
| GO:0045665 | negative regulation of neuron differentiation | 28/4924 | 0.000780952 | 0.002815268 |
| GO:0046888 | negative regulation of hormone secretion | 28/4924 | 0.000780952 | 0.002815268 |
| GO:0033173 | calcineurin-NFAT signaling cascade | 18/4924 | 0.000783666 | 0.002821826 |
| GO:1990573 | potassium ion import across plasma membrane | 18/4924 | 0.000783666 | 0.002821826 |
| GO:0001704 | formation of primary germ layer | 31/4924 | 0.000789031 | 0.002839521 |
| GO:0003229 | ventricular cardiac muscle tissue development | 22/4924 | 0.000794984 | 0.002852801 |
| GO:0032613 | interleukin-10 production | 22/4924 | 0.000794984 | 0.002852801 |
| GO:0060038 | cardiac muscle cell proliferation | 22/4924 | 0.000794984 | 0.002852801 |
| GO:0098930 | axonal transport | 22/4924 | 0.000794984 | 0.002852801 |
| GO:2000781 | positive regulation of double-strand break repair | 22/4924 | 0.000794984 | 0.002852801 |
| GO:0045655 | regulation of monocyte differentiation | Sep-24 | 0.00079894 | 0.002859866 |
| GO:0070571 | negative regulation of neuron projection regeneration | Sep-24 | 0.00079894 | 0.002859866 |
| GO:0071498 | cellular response to fluid shear stress | Sep-24 | 0.00079894 | 0.002859866 |
| GO:1902018 | negative regulation of cilium assembly | Sep-24 | 0.00079894 | 0.002859866 |
| GO:0030850 | prostate gland development | 19/4924 | 0.000801921 | 0.002859866 |

| | | | | |
|---|---|---|---|---|
| GO:0097720 | calcineurin-mediated signaling | 19/4924 | 0.000801921 | 0.002859866 |
| GO:0002092 | positive regulation of receptor internalization | 14/4924 | 0.000801942 | 0.002859866 |
| GO:0060512 | prostate gland morphogenesis | 14/4924 | 0.000801942 | 0.002859866 |
| GO:0061842 | microtubule organizing center localization | 14/4924 | 0.000801942 | 0.002859866 |
| GO:0099563 | modification of synaptic structure | 14/4924 | 0.000801942 | 0.002859866 |
| GO:1903959 | regulation of anion transmembrane transport | 14/4924 | 0.000801942 | 0.002859866 |
| GO:0010656 | negative regulation of muscle cell apoptotic process | 21/4924 | 0.000805727 | 0.002870119 |
| GO:0060760 | positive regulation of response to cytokine stimulus | 21/4924 | 0.000805727 | 0.002870119 |
| GO:0052372 | modulation by symbiont of entry into host | 20/4924 | 0.000808632 | 0.002875593 |
| GO:0062237 | protein localization to postsynapse | 20/4924 | 0.000808632 | 0.002875593 |
| GO:0140353 | lipid export from cell | 20/4924 | 0.000808632 | 0.002875593 |
| GO:0007144 | female meiosis I | Aug-24 | 0.000815069 | 0.002882226 |
| GO:0031915 | positive regulation of synaptic plasticity | Aug-24 | 0.000815069 | 0.002882226 |
| GO:0032341 | aldosterone metabolic process | Aug-24 | 0.000815069 | 0.002882226 |
| GO:0035791 | platelet-derived growth factor receptor-beta signaling pathway | Aug-24 | 0.000815069 | 0.002882226 |
| GO:0051988 | regulation of attachment of spindle microtubules to kinetochore | Aug-24 | 0.000815069 | 0.002882226 |
| GO:0060346 | bone trabecula formation | Aug-24 | 0.000815069 | 0.002882226 |
| GO:0060670 | branching involved in labyrinthine layer morphogenesis | Aug-24 | 0.000815069 | 0.002882226 |

| | | | | |
|---|---|---|---|---|
| GO:0086069 | bundle of His cell to Purkinje myocyte communication | Aug-24 | 0.000815069 | 0.002882226 |
| GO:0090331 | negative regulation of platelet aggregation | Aug-24 | 0.000815069 | 0.002882226 |
| GO:1901201 | regulation of extracellular matrix assembly | Aug-24 | 0.000815069 | 0.002882226 |
| GO:0002027 | regulation of heart rate | 33/4924 | 0.000822859 | 0.002908142 |
| GO:0006346 | DNA methylation-dependent heterochromatin assembly | Dec-24 | 0.000836029 | 0.002949727 |
| GO:0010165 | response to X-ray | Dec-24 | 0.000836029 | 0.002949727 |
| GO:0010829 | negative regulation of glucose transmembrane transport | Dec-24 | 0.000836029 | 0.002949727 |
| GO:1903825 | organic acid transmembrane transport | 39/4924 | 0.000848153 | 0.002990831 |
| GO:0033138 | positive regulation of peptidyl-serine phosphorylation | 37/4924 | 0.000850447 | 0.002997243 |
| GO:0050796 | regulation of insulin secretion | 50/4924 | 0.000880318 | 0.003100784 |
| GO:0010737 | protein kinase A signaling | 15/4924 | 0.000892577 | 0.003136955 |
| GO:0045622 | regulation of T-helper cell differentiation | 15/4924 | 0.000892577 | 0.003136955 |
| GO:0060292 | long-term synaptic depression | 15/4924 | 0.000892577 | 0.003136955 |
| GO:1902003 | regulation of amyloid-beta formation | 15/4924 | 0.000892577 | 0.003136955 |
| GO:0002292 | T cell differentiation involved in immune response | 25/4924 | 0.000902915 | 0.003169753 |
| GO:0032413 | negative regulation of ion transmembrane transporter activity | 25/4924 | 0.000902915 | 0.003169753 |
| GO:0044070 | regulation of anion transport | 34/4924 | 0.000906568 | 0.003180809 |
| GO:0035601 | protein deacylation | 38/4924 | 0.000917924 | 0.00321707 |

| | | | | |
|---|---|---|---|---|
| GO:009 8732 | macromolecule deacylation | 38/49 24 | 0.00091 7924 | 0.00321 707 |
| GO:003 0032 | lamellipodium assembly | 24/49 24 | 0.00094 0038 | 0.00329 2743 |
| GO:001 5914 | phospholipid transport | 28/49 24 | 0.00094 6592 | 0.00331 2023 |
| GO:004 5814 | negative regulation of gene expression, epigenetic | 28/49 24 | 0.00094 6592 | 0.00331 2023 |
| GO:006 0251 | regulation of glial cell proliferation | 16/49 24 | 0.00095 9455 | 0.00335 3307 |
| GO:190 2275 | regulation of chromatin organization | 16/49 24 | 0.00095 9455 | 0.00335 3307 |
| GO:003 2720 | negative regulation of tumor necrosis factor production | 23/49 24 | 0.00097 3654 | 0.00339 0425 |
| GO:004 2509 | regulation of tyrosine phosphorylation of STAT protein | 23/49 24 | 0.00097 3654 | 0.00339 0425 |
| GO:004 3462 | regulation of ATP-dependent activity | 23/49 24 | 0.00097 3654 | 0.00339 0425 |
| GO:004 6324 | regulation of glucose import | 23/49 24 | 0.00097 3654 | 0.00339 0425 |
| GO:004 8662 | negative regulation of smooth muscle cell proliferation | 23/49 24 | 0.00097 3654 | 0.00339 0425 |
| GO:009 7194 | execution phase of apoptosis | 23/49 24 | 0.00097 3654 | 0.00339 0425 |
| GO:004 5639 | positive regulation of myeloid cell differentiation | 33/49 24 | 0.00097 4378 | 0.00339 0425 |
| GO:004 8477 | oogenesis | 33/49 24 | 0.00097 4378 | 0.00339 0425 |
| GO:005 1648 | vesicle localization | 50/49 24 | 0.00099 7662 | 0.00346 3386 |
| GO:001 0669 | epithelial structure maintenance | 13/49 24 | 0.00099 9223 | 0.00346 3386 |
| GO:006 1036 | positive regulation of cartilage development | 13/49 24 | 0.00099 9223 | 0.00346 3386 |
| GO:200 0352 | negative regulation of endothelial cell apoptotic process | 13/49 24 | 0.00099 9223 | 0.00346 3386 |

| | | | | |
|---|---|---|---|---|
| GO:2000648 | positive regulation of stem cell proliferation | 13/4924 | 0.000999223 | 0.003463386 |
| GO:0032930 | positive regulation of superoxide anion generation | Nov-24 | 0.001004271 | 0.003463386 |
| GO:0036303 | lymph vessel morphogenesis | Nov-24 | 0.001004271 | 0.003463386 |
| GO:0043371 | negative regulation of CD4-positive, alpha-beta T cell differentiation | Nov-24 | 0.001004271 | 0.003463386 |
| GO:0043586 | tongue development | Nov-24 | 0.001004271 | 0.003463386 |
| GO:0048485 | sympathetic nervous system development | Nov-24 | 0.001004271 | 0.003463386 |
| GO:0048670 | regulation of collateral sprouting | Nov-24 | 0.001004271 | 0.003463386 |
| GO:1900006 | positive regulation of dendrite development | Nov-24 | 0.001004271 | 0.003463386 |
| GO:1902547 | regulation of cellular response to vascular endothelial growth factor stimulus | Nov-24 | 0.001004271 | 0.003463386 |
| GO:1990776 | response to angiotensin | Nov-24 | 0.001004271 | 0.003463386 |
| GO:0034142 | toll-like receptor 4 signaling pathway | 17/4924 | 0.001004684 | 0.003463386 |
| GO:1903170 | negative regulation of calcium ion transmembrane transport | 17/4924 | 0.001004684 | 0.003463386 |
| GO:2001258 | negative regulation of cation channel activity | 17/4924 | 0.001004684 | 0.003463386 |
| GO:0010972 | negative regulation of G2/M transition of mitotic cell cycle | 21/4924 | 0.00102457 | 0.003530006 |
| GO:0010862 | positive regulation of pathway-restricted SMAD protein phosphorylation | 18/4924 | 0.001030993 | 0.003548261 |
| GO:0021532 | neural tube patterning | 18/4924 | 0.001030993 | 0.003548261 |
| GO:0048511 | rhythmic process | 74/4924 | 0.001038167 | 0.0035571 |
| GO:0006906 | vesicle fusion | 32/4924 | 0.001046137 | 0.003596453 |

| | | | | |
|---|---|---|---|---|
| GO:0045732 | positive regulation of protein catabolic process | 57/4924 | 0.001095119 | 0.003762794 |
| GO:0070192 | chromosome organization involved in meiotic cell cycle | 25/4924 | 0.001109392 | 0.003809758 |
| GO:0072331 | signal transduction by p53 class mediator | 41/4924 | 0.001115127 | 0.00382737 |
| GO:0043243 | positive regulation of protein-containing complex disassembly | 14/4924 | 0.001128174 | 0.003861639 |
| GO:0045987 | positive regulation of smooth muscle contraction | 14/4924 | 0.001128174 | 0.003861639 |
| GO:0060479 | lung cell differentiation | 14/4924 | 0.001128174 | 0.003861639 |
| GO:1990089 | response to nerve growth factor | 14/4924 | 0.001128174 | 0.003861639 |
| GO:1990090 | cellular response to nerve growth factor stimulus | 14/4924 | 0.001128174 | 0.003861639 |
| GO:0007260 | tyrosine phosphorylation of STAT protein | 24/4924 | 0.001161391 | 0.003966724 |
| GO:0016445 | somatic diversification of immunoglobulins | 24/4924 | 0.001161391 | 0.003966724 |
| GO:0042130 | negative regulation of T cell proliferation | 24/4924 | 0.001161391 | 0.003966724 |
| GO:0048645 | animal organ formation | 24/4924 | 0.001161391 | 0.003966724 |
| GO:0001946 | lymphangiogenesis | Oct-24 | 0.001183795 | 0.004034501 |
| GO:0046697 | decidualization | Oct-24 | 0.001183795 | 0.004034501 |
| GO:0048569 | post-embryonic animal organ development | Oct-24 | 0.001183795 | 0.004034501 |
| GO:0090036 | regulation of protein kinase C signaling | Oct-24 | 0.001183795 | 0.004034501 |
| GO:0042058 | regulation of epidermal growth factor receptor signaling pathway | 23/4924 | 0.001210347 | 0.004122765 |
| GO:2000403 | positive regulation of lymphocyte migration | 15/4924 | 0.001223487 | 0.004165274 |

| | | | | |
|---|---|---|---|---|
| GO:0002052 | positive regulation of neuroblast proliferation | Dec-24 | 0.001237149 | 0.004193669 |
| GO:0032928 | regulation of superoxide anion generation | Dec-24 | 0.001237149 | 0.004193669 |
| GO:0033622 | integrin activation | Dec-24 | 0.001237149 | 0.004193669 |
| GO:0043931 | ossification involved in bone maturation | Dec-24 | 0.001237149 | 0.004193669 |
| GO:0046639 | negative regulation of alpha-beta T cell differentiation | Dec-24 | 0.001237149 | 0.004193669 |
| GO:0048634 | regulation of muscle organ development | Dec-24 | 0.001237149 | 0.004193669 |
| GO:0060039 | pericardium development | Dec-24 | 0.001237149 | 0.004193669 |
| GO:1900745 | positive regulation of p38MAPK cascade | Dec-24 | 0.001237149 | 0.004193669 |
| GO:0006865 | amino acid transport | 43/4924 | 0.00123991 | 0.004199638 |
| GO:0021953 | central nervous system neuron differentiation | 53/4924 | 0.001240242 | 0.004199638 |
| GO:0006611 | protein export from nucleus | 22/4924 | 0.001254696 | 0.004239473 |
| GO:0042461 | photoreceptor cell development | 22/4924 | 0.001254696 | 0.004239473 |
| GO:0060350 | endochondral bone morphogenesis | 22/4924 | 0.001254696 | 0.004239473 |
| GO:0090398 | cellular senescence | 22/4924 | 0.001254696 | 0.004239473 |
| GO:0007189 | adenylate cyclase-activating G protein-coupled receptor signaling pathway | 41/4924 | 0.001281979 | 0.00432934 |
| GO:0031507 | heterochromatin assembly | 26/4924 | 0.001283755 | 0.004333015 |
| GO:0045576 | mast cell activation | 29/4924 | 0.001284667 | 0.004333775 |
| GO:0016572 | histone phosphorylation | 16/4924 | 0.001288161 | 0.004333966 |

| | | | | |
|---|---|---|---|---|
| GO:0034331 | cell junction maintenance | 16/4924 | 0.001288161 | 0.004333966 |
| GO:0045601 | regulation of endothelial cell differentiation | 16/4924 | 0.001288161 | 0.004333966 |
| GO:0046636 | negative regulation of alpha-beta T cell activation | 16/4924 | 0.001288161 | 0.004333966 |
| GO:1902116 | negative regulation of organelle assembly | 16/4924 | 0.001288161 | 0.004333966 |
| GO:0043434 | response to peptide hormone | 84/4924 | 0.001295995 | 0.004357998 |
| GO:0003170 | heart valve development | 20/4924 | 0.001321563 | 0.004439241 |
| GO:0010662 | regulation of striated muscle cell apoptotic process | 20/4924 | 0.001321563 | 0.004439241 |
| GO:0045005 | DNA-templated DNA replication maintenance of fidelity | 17/4924 | 0.001326196 | 0.004450061 |
| GO:0071300 | cellular response to retinoic acid | 17/4924 | 0.001326196 | 0.004450061 |
| GO:0001961 | positive regulation of cytokine-mediated signaling pathway | 19/4924 | 0.001339055 | 0.004488433 |
| GO:0042531 | positive regulation of tyrosine phosphorylation of STAT protein | 19/4924 | 0.001339055 | 0.004488433 |
| GO:0042088 | T-helper 1 type immune response | 18/4924 | 0.001341816 | 0.004490525 |
| GO:0044818 | mitotic G2/M transition checkpoint | 18/4924 | 0.001341816 | 0.004490525 |
| GO:0061647 | histone H3-K9 modification | 18/4924 | 0.001341816 | 0.004490525 |
| GO:0002286 | T cell activation involved in immune response | 35/4924 | 0.001354807 | 0.004524878 |
| GO:0042249 | establishment of planar polarity of embryonic epithelium | Sep-24 | 0.001356387 | 0.004524878 |
| GO:0043217 | myelin maintenance | Sep-24 | 0.001356387 | 0.004524878 |
| GO:0046184 | aldehyde biosynthetic process | Sep-24 | 0.001356387 | 0.004524878 |

| | | | | |
|---|---|---|---|---|
| GO:0051580 | regulation of neurotransmitter uptake | Sep-24 | 0.001356387 | 0.004524878 |
| GO:1904424 | regulation of GTP binding | Sep-24 | 0.001356387 | 0.004524878 |
| GO:0048814 | regulation of dendrite morphogenesis | 28/4924 | 0.001371358 | 0.004572401 |
| GO:0009895 | negative regulation of catabolic process | 75/4924 | 0.001383615 | 0.004610831 |
| GO:0002456 | T cell mediated immunity | 40/4924 | 0.001394579 | 0.004644914 |
| GO:0016032 | viral process | 83/4924 | 0.001411595 | 0.004699106 |
| GO:0045830 | positive regulation of isotype switching | 13/4924 | 0.001421893 | 0.004728395 |
| GO:2000515 | negative regulation of CD4-positive, alpha-beta T cell activation | 13/4924 | 0.001421893 | 0.004728395 |
| GO:0032456 | endocytic recycling | 24/4924 | 0.001426218 | 0.004740276 |
| GO:0036473 | cell death in response to oxidative stress | 30/4924 | 0.001427922 | 0.004743438 |
| GO:1905039 | carboxylic acid transmembrane transport | 38/4924 | 0.001430474 | 0.004749415 |
| GO:0051607 | defense response to virus | 68/4924 | 0.001448251 | 0.004805907 |
| GO:0010522 | regulation of calcium ion transport into cytosol | 32/4924 | 0.001457844 | 0.004830115 |
| GO:0030518 | intracellular steroid hormone receptor signaling pathway | 32/4924 | 0.001457844 | 0.004830115 |
| GO:0099175 | regulation of postsynapse organization | 32/4924 | 0.001457844 | 0.004830115 |
| GO:0034644 | cellular response to UV | 27/4924 | 0.001461061 | 0.00483569 |
| GO:0046323 | glucose import | 27/4924 | 0.001461061 | 0.00483569 |
| GO:0008637 | apoptotic mitochondrial changes | 34/4924 | 0.001465558 | 0.00484803 |

| | | | | |
|---|---|---|---|---|
| GO:0001977 | renal system process involved in regulation of blood volume | Aug-24 | 0.001485388 | 0.004887981 |
| GO:0010172 | embryonic body morphogenesis | Aug-24 | 0.001485388 | 0.004887981 |
| GO:0030320 | cellular monovalent inorganic anion homeostasis | Aug-24 | 0.001485388 | 0.004887981 |
| GO:0033690 | positive regulation of osteoblast proliferation | Aug-24 | 0.001485388 | 0.004887981 |
| GO:0034350 | regulation of glial cell apoptotic process | Aug-24 | 0.001485388 | 0.004887981 |
| GO:0061577 | calcium ion transmembrane transport via high voltage-gated calcium channel | Aug-24 | 0.001485388 | 0.004887981 |
| GO:0072074 | kidney mesenchyme development | Aug-24 | 0.001485388 | 0.004887981 |
| GO:0086014 | atrial cardiac muscle cell action potential | Aug-24 | 0.001485388 | 0.004887981 |
| GO:0086026 | atrial cardiac muscle cell to AV node cell signaling | Aug-24 | 0.001485388 | 0.004887981 |
| GO:0086066 | atrial cardiac muscle cell to AV node cell communication | Aug-24 | 0.001485388 | 0.004887981 |
| GO:0071496 | cellular response to external stimulus | 72/4924 | 0.001493078 | 0.004904075 |
| GO:0006705 | mineralocorticoid biosynthetic process | Jul-24 | 0.001503502 | 0.004904075 |
| GO:0031272 | regulation of pseudopodium assembly | Jul-24 | 0.001503502 | 0.004904075 |
| GO:0031274 | positive regulation of pseudopodium assembly | Jul-24 | 0.001503502 | 0.004904075 |
| GO:0033262 | regulation of nuclear cell cycle DNA replication | Jul-24 | 0.001503502 | 0.004904075 |
| GO:0048251 | elastic fiber assembly | Jul-24 | 0.001503502 | 0.004904075 |
| GO:0048934 | peripheral nervous system neuron differentiation | Jul-24 | 0.001503502 | 0.004904075 |
| GO:0048935 | peripheral nervous system neuron development | Jul-24 | 0.001503502 | 0.004904075 |

| | | | | |
|---|---|---|---|---|
| GO:0060430 | lung saccule development | Jul-24 | 0.001503502 | 0.004904075 |
| GO:0060623 | regulation of chromosome condensation | Jul-24 | 0.001503502 | 0.004904075 |
| GO:0070316 | regulation of G0 to G1 transition | Jul-24 | 0.001503502 | 0.004904075 |
| GO:0072124 | regulation of glomerular mesangial cell proliferation | Jul-24 | 0.001503502 | 0.004904075 |
| GO:0072182 | regulation of nephron tubule epithelial cell differentiation | Jul-24 | 0.001503502 | 0.004904075 |
| GO:0072497 | mesenchymal stem cell differentiation | Jul-24 | 0.001503502 | 0.004904075 |
| GO:1902166 | negative regulation of intrinsic apoptotic signaling pathway in response to DNA damage by p53 class mediator | Jul-24 | 0.001503502 | 0.004904075 |
| GO:2000317 | negative regulation of T-helper 17 type immune response | Jul-24 | 0.001503502 | 0.004904075 |
| GO:2001225 | regulation of chloride transport | Jul-24 | 0.001503502 | 0.004904075 |
| GO:0007413 | axonal fasciculation | Nov-24 | 0.001518296 | 0.004931919 |
| GO:0010759 | positive regulation of macrophage chemotaxis | Nov-24 | 0.001518296 | 0.004931919 |
| GO:0010894 | negative regulation of steroid biosynthetic process | Nov-24 | 0.001518296 | 0.004931919 |
| GO:0030325 | adrenal gland development | Nov-24 | 0.001518296 | 0.004931919 |
| GO:0032288 | myelin assembly | Nov-24 | 0.001518296 | 0.004931919 |
| GO:0035988 | chondrocyte proliferation | Nov-24 | 0.001518296 | 0.004931919 |
| GO:0106030 | neuron projection fasciculation | Nov-24 | 0.001518296 | 0.004931919 |
| GO:1905820 | positive regulation of chromosome separation | Nov-24 | 0.001518296 | 0.004931919 |
| GO:0010466 | negative regulation of peptidase activity | 64/4924 | 0.001532003 | 0.004973883 |

| | | | | |
|---|---|---|---|---|
| GO:0002053 | positive regulation of mesenchymal cell proliferation | 14/4924 | 0.001558633 | 0.005049938 |
| GO:0010573 | vascular endothelial growth factor production | 14/4924 | 0.001558633 | 0.005049938 |
| GO:0045686 | negative regulation of glial cell differentiation | 14/4924 | 0.001558633 | 0.005049938 |
| GO:0140718 | facultative heterochromatin assembly | 14/4924 | 0.001558633 | 0.005049938 |
| GO:2000243 | positive regulation of reproductive process | 31/4924 | 0.001570629 | 0.005086192 |
| GO:0006650 | glycerophospholipid metabolic process | 67/4924 | 0.001581253 | 0.005117967 |
| GO:0051952 | regulation of amine transport | 33/4924 | 0.001584436 | 0.005125639 |
| GO:0140546 | defense response to symbiont | 68/4924 | 0.001595067 | 0.005157384 |
| GO:0050435 | amyloid-beta metabolic process | 21/4924 | 0.00161818 | 0.005226755 |
| GO:1902750 | negative regulation of cell cycle G2/M phase transition | 21/4924 | 0.00161818 | 0.005226755 |
| GO:0032088 | negative regulation of NF-kappaB transcription factor activity | 25/4924 | 0.001646933 | 0.005316904 |
| GO:0070849 | response to epidermal growth factor | 15/4924 | 0.001651641 | 0.005329374 |
| GO:0042987 | amyloid precursor protein catabolic process | 20/4924 | 0.001668146 | 0.005377126 |
| GO:0055078 | sodium ion homeostasis | 20/4924 | 0.001668146 | 0.005377126 |
| GO:0016571 | histone methylation | 44/4924 | 0.00168329 | 0.00542317 |
| GO:0007140 | male meiotic nuclear division | 19/4924 | 0.001706216 | 0.005482252 |
| GO:0010665 | regulation of cardiac muscle cell apoptotic process | 19/4924 | 0.001706216 | 0.005482252 |
| GO:0048016 | inositol phosphate-mediated signaling | 19/4924 | 0.001706216 | 0.005482252 |

| | | | | |
|---|---|---|---|---|
| GO:1903539 | protein localization to postsynaptic membrane | 19/4924 | 0.001706216 | 0.005482252 |
| GO:0048854 | brain morphogenesis | 16/4924 | 0.001706845 | 0.005482252 |
| GO:1900026 | positive regulation of substrate adhesion-dependent cell spreading | 16/4924 | 0.001706845 | 0.005482252 |
| GO:0032715 | negative regulation of interleukin-6 production | 18/4924 | 0.001728526 | 0.005546237 |
| GO:0061756 | leukocyte adhesion to vascular endothelial cell | 18/4924 | 0.001728526 | 0.005546237 |
| GO:0034205 | amyloid-beta formation | 17/4924 | 0.0017305 | 0.005549748 |
| GO:0006664 | glycolipid metabolic process | 27/4924 | 0.001752536 | 0.005617559 |
| GO:0033081 | regulation of T cell differentiation in thymus | Dec-24 | 0.001785082 | 0.00571027 |
| GO:0060444 | branching involved in mammary gland duct morphogenesis | Dec-24 | 0.001785082 | 0.00571027 |
| GO:0061311 | cell surface receptor signaling pathway involved in heart development | Dec-24 | 0.001785082 | 0.00571027 |
| GO:1903649 | regulation of cytoplasmic transport | Dec-24 | 0.001785082 | 0.00571027 |
| GO:0034968 | histone lysine methylation | 37/4924 | 0.001793567 | 0.005734503 |
| GO:0019233 | sensory perception of pain | 40/4924 | 0.001833241 | 0.005853334 |
| GO:0044851 | hair cycle phase | Oct-24 | 0.001840017 | 0.005853334 |
| GO:0046599 | regulation of centriole replication | Oct-24 | 0.001840017 | 0.005853334 |
| GO:0048791 | calcium ion-regulated exocytosis of neurotransmitter | Oct-24 | 0.001840017 | 0.005853334 |
| GO:0060009 | Sertoli cell development | Oct-24 | 0.001840017 | 0.005853334 |
| GO:0060080 | inhibitory postsynaptic potential | Oct-24 | 0.001840017 | 0.005853334 |

| | | | | |
|---|---|---|---|---|
| GO:0060438 | trachea development | Oct-24 | 0.001840017 | 0.005853334 |
| GO:0099515 | actin filament-based transport | Oct-24 | 0.001840017 | 0.005853334 |
| GO:1900746 | regulation of vascular endothelial growth factor signaling pathway | Oct-24 | 0.001840017 | 0.005853334 |
| GO:1902254 | negative regulation of intrinsic apoptotic signaling pathway by p53 class mediator | Oct-24 | 0.001840017 | 0.005853334 |
| GO:0002200 | somatic diversification of immune receptors | 26/4924 | 0.00186981 | 0.00594511 |
| GO:0031060 | regulation of histone methylation | 28/4924 | 0.001951547 | 0.006201866 |
| GO:0010847 | regulation of chromatin assembly | 13/4924 | 0.001982775 | 0.006285267 |
| GO:0044030 | regulation of DNA methylation | 13/4924 | 0.001982775 | 0.006285267 |
| GO:0048566 | embryonic digestive tract development | 13/4924 | 0.001982775 | 0.006285267 |
| GO:0060740 | prostate gland epithelium morphogenesis | 13/4924 | 0.001982775 | 0.006285267 |
| GO:0060914 | heart formation | 13/4924 | 0.001982775 | 0.006285267 |
| GO:0046928 | regulation of neurotransmitter secretion | 34/4924 | 0.001986659 | 0.006294413 |
| GO:0090174 | organelle membrane fusion | 32/4924 | 0.002003181 | 0.006343574 |
| GO:0002792 | negative regulation of peptide secretion | 21/4924 | 0.002011069 | 0.006362164 |
| GO:0060999 | positive regulation of dendritic spine development | 21/4924 | 0.002011069 | 0.006362164 |
| GO:0007586 | digestion | 37/4924 | 0.002065061 | 0.006529695 |
| GO:0060420 | regulation of heart growth | 27/4924 | 0.002092543 | 0.006609963 |
| GO:1903509 | liposaccharide metabolic process | 27/4924 | 0.002092543 | 0.006609963 |

| | | | | |
|---|---|---|---|---|
| GO:0010923 | negative regulation of phosphatase activity | 14/4924 | 0.002117439 | 0.006681908 |
| GO:0046677 | response to antibiotic | 14/4924 | 0.002117439 | 0.006681908 |
| GO:1901861 | regulation of muscle tissue development | 19/4924 | 0.002155008 | 0.006797061 |
| GO:0009743 | response to carbohydrate | 53/4924 | 0.002176639 | 0.006861855 |
| GO:0014072 | response to isoquinoline alkaloid | Sep-24 | 0.002188527 | 0.006868425 |
| GO:0032570 | response to progesterone | Sep-24 | 0.002188527 | 0.006868425 |
| GO:0032823 | regulation of natural killer cell differentiation | Sep-24 | 0.002188527 | 0.006868425 |
| GO:0033623 | regulation of integrin activation | Sep-24 | 0.002188527 | 0.006868425 |
| GO:0035457 | cellular response to interferon-alpha | Sep-24 | 0.002188527 | 0.006868425 |
| GO:0045837 | negative regulation of membrane potential | Sep-24 | 0.002188527 | 0.006868425 |
| GO:0055003 | cardiac myofibril assembly | Sep-24 | 0.002188527 | 0.006868425 |
| GO:0070314 | G1 to G0 transition | Sep-24 | 0.002188527 | 0.006868425 |
| GO:0086011 | membrane repolarization during action potential | Sep-24 | 0.002188527 | 0.006868425 |
| GO:0010762 | regulation of fibroblast migration | 15/4924 | 0.002197961 | 0.006880907 |
| GO:0030513 | positive regulation of BMP signaling pathway | 15/4924 | 0.002197961 | 0.006880907 |
| GO:0032733 | positive regulation of interleukin-10 production | 15/4924 | 0.002197961 | 0.006880907 |
| GO:0038179 | neurotrophin signaling pathway | 15/4924 | 0.002197961 | 0.006880907 |
| GO:1901381 | positive regulation of potassium ion transmembrane transport | 15/4924 | 0.002197961 | 0.006880907 |

| | | | | |
|---|---|---|---|---|
| GO:0045907 | positive regulation of vasoconstriction | 18/4924 | 0.002205066 | 0.006899721 |
| GO:0010742 | macrophage derived foam cell differentiation | Nov-24 | 0.002228802 | 0.006953289 |
| GO:0035020 | regulation of Rac protein signal transduction | Nov-24 | 0.002228802 | 0.006953289 |
| GO:0045737 | positive regulation of cyclin-dependent protein serine/threonine kinase activity | Nov-24 | 0.002228802 | 0.006953289 |
| GO:0045939 | negative regulation of steroid metabolic process | Nov-24 | 0.002228802 | 0.006953289 |
| GO:1902230 | negative regulation of intrinsic apoptotic signaling pathway in response to DNA damage | Nov-24 | 0.002228802 | 0.006953289 |
| GO:1902473 | regulation of protein localization to synapse | Nov-24 | 0.002228802 | 0.006953289 |
| GO:0001569 | branching involved in blood vessel morphogenesis | 16/4924 | 0.002233722 | 0.006958308 |
| GO:0006509 | membrane protein ectodomain proteolysis | 16/4924 | 0.002233722 | 0.006958308 |
| GO:1905521 | regulation of macrophage migration | 16/4924 | 0.002233722 | 0.006958308 |
| GO:0032623 | interleukin-2 production | 23/4924 | 0.00223808 | 0.006965002 |
| GO:0043966 | histone H3 acetylation | 23/4924 | 0.00223808 | 0.006965002 |
| GO:0032410 | negative regulation of transporter activity | 26/4924 | 0.002240089 | 0.006967814 |
| GO:0055021 | regulation of cardiac muscle tissue growth | 25/4924 | 0.002393575 | 0.007441562 |
| GO:0048515 | spermatid differentiation | 62/4924 | 0.002456046 | 0.007632019 |
| GO:0001836 | release of cytochrome c from mitochondria | 21/4924 | 0.00248179 | 0.007708221 |
| GO:0090314 | positive regulation of protein targeting to membrane | Dec-24 | 0.002517221 | 0.007804651 |
| GO:1903523 | negative regulation of blood circulation | Dec-24 | 0.002517221 | 0.007804651 |

| | | | | |
|---|---|---|---|---|
| GO:000<br>2024 | diet induced thermogenesis | Aug-<br>24 | 0.00252<br>7692 | 0.00780<br>4651 |
| GO:000<br>8212 | mineralocorticoid metabolic process | Aug-<br>24 | 0.00252<br>7692 | 0.00780<br>4651 |
| GO:003<br>1115 | negative regulation of microtubule polymerization | Aug-<br>24 | 0.00252<br>7692 | 0.00780<br>4651 |
| GO:004<br>2048 | olfactory behavior | Aug-<br>24 | 0.00252<br>7692 | 0.00780<br>4651 |
| GO:004<br>2640 | anagen | Aug-<br>24 | 0.00252<br>7692 | 0.00780<br>4651 |
| GO:004<br>8715 | negative regulation of oligodendrocyte differentiation | Aug-<br>24 | 0.00252<br>7692 | 0.00780<br>4651 |
| GO:005<br>5057 | neuroblast division | Aug-<br>24 | 0.00252<br>7692 | 0.00780<br>4651 |
| GO:007<br>2109 | glomerular mesangium development | Aug-<br>24 | 0.00252<br>7692 | 0.00780<br>4651 |
| GO:008<br>6103 | G protein-coupled receptor signaling pathway involved in heart process | Aug-<br>24 | 0.00252<br>7692 | 0.00780<br>4651 |
| GO:200<br>1212 | regulation of vasculogenesis | Aug-<br>24 | 0.00252<br>7692 | 0.00780<br>4651 |
| GO:005<br>5006 | cardiac cell development | 31/49<br>24 | 0.00253<br>1692 | 0.00781<br>3176 |
| GO:007<br>1322 | cellular response to carbohydrate stimulus | 42/49<br>24 | 0.00258<br>0102 | 0.00795<br>8682 |
| GO:004<br>6467 | membrane lipid biosynthetic process | 36/49<br>24 | 0.00258<br>3403 | 0.00796<br>4967 |
| GO:004<br>6622 | positive regulation of organ growth | 20/49<br>24 | 0.00259<br>5632 | 0.00799<br>4851 |
| GO:006<br>0612 | adipose tissue development | 20/49<br>24 | 0.00259<br>5632 | 0.00799<br>4851 |
| GO:000<br>7286 | spermatid development | 60/49<br>24 | 0.00265<br>9562 | 0.00818<br>7762 |
| GO:001<br>5872 | dopamine transport | 19/49<br>24 | 0.00269<br>9055 | 0.00830<br>1239 |
| GO:005<br>1055 | negative regulation of lipid biosynthetic process | 19/49<br>24 | 0.00269<br>9055 | 0.00830<br>1239 |

| | | | | |
|---|---|---|---|---|
| GO:0031294 | lymphocyte costimulation | 13/4924 | 0.002713767 | 0.0083157 |
| GO:0035335 | peptidyl-tyrosine dephosphorylation | 13/4924 | 0.002713767 | 0.0083157 |
| GO:0055075 | potassium ion homeostasis | 13/4924 | 0.002713767 | 0.0083157 |
| GO:0060487 | lung epithelial cell differentiation | 13/4924 | 0.002713767 | 0.0083157 |
| GO:0090322 | regulation of superoxide metabolic process | 13/4924 | 0.002713767 | 0.0083157 |
| GO:2000406 | positive regulation of T cell migration | 13/4924 | 0.002713767 | 0.0083157 |
| GO:0010611 | regulation of cardiac muscle hypertrophy | 23/4924 | 0.002714308 | 0.0083157 |
| GO:0071479 | cellular response to ionizing radiation | 23/4924 | 0.002714308 | 0.0083157 |
| GO:0051781 | positive regulation of cell division | 28/4924 | 0.002730555 | 0.008361413 |
| GO:0002043 | blood vessel endothelial cell proliferation involved in sprouting angiogenesis | Oct-24 | 0.002761011 | 0.008387501 |
| GO:0021535 | cell migration in hindbrain | Oct-24 | 0.002761011 | 0.008387501 |
| GO:0031649 | heat generation | Oct-24 | 0.002761011 | 0.008387501 |
| GO:0033630 | positive regulation of cell adhesion mediated by integrin | Oct-24 | 0.002761011 | 0.008387501 |
| GO:0042481 | regulation of odontogenesis | Oct-24 | 0.002761011 | 0.008387501 |
| GO:0003093 | regulation of glomerular filtration | Jul-24 | 0.002778176 | 0.008387501 |
| GO:0030836 | positive regulation of actin filament depolymerization | Jul-24 | 0.002778176 | 0.008387501 |
| GO:0031652 | positive regulation of heat generation | Jul-24 | 0.002778176 | 0.008387501 |
| GO:0035810 | positive regulation of urine volume | Jul-24 | 0.002778176 | 0.008387501 |

| | | | | |
|---|---|---|---|---|
| GO:0040034 | regulation of development, heterochronic | Jul-24 | 0.002778176 | 0.008387501 |
| GO:0045023 | G0 to G1 transition | Jul-24 | 0.002778176 | 0.008387501 |
| GO:0051639 | actin filament network formation | Jul-24 | 0.002778176 | 0.008387501 |
| GO:0051974 | negative regulation of telomerase activity | Jul-24 | 0.002778176 | 0.008387501 |
| GO:0060601 | lateral sprouting from an epithelium | Jul-24 | 0.002778176 | 0.008387501 |
| GO:0071679 | commissural neuron axon guidance | Jul-24 | 0.002778176 | 0.008387501 |
| GO:0090557 | establishment of endothelial intestinal barrier | Jul-24 | 0.002778176 | 0.008387501 |
| GO:0140059 | dendrite arborization | Jul-24 | 0.002778176 | 0.008387501 |
| GO:1905048 | regulation of metallopeptidase activity | Jul-24 | 0.002778176 | 0.008387501 |
| GO:2001223 | negative regulation of neuron migration | Jul-24 | 0.002778176 | 0.008387501 |
| GO:0002315 | marginal zone B cell differentiation | Jun-24 | 0.002780902 | 0.008387501 |
| GO:0010216 | maintenance of DNA methylation | Jun-24 | 0.002780902 | 0.008387501 |
| GO:0010692 | regulation of alkaline phosphatase activity | Jun-24 | 0.002780902 | 0.008387501 |
| GO:0021957 | corticospinal tract morphogenesis | Jun-24 | 0.002780902 | 0.008387501 |
| GO:0022027 | interkinetic nuclear migration | Jun-24 | 0.002780902 | 0.008387501 |
| GO:0036363 | transforming growth factor beta activation | Jun-24 | 0.002780902 | 0.008387501 |
| GO:0070673 | response to interleukin-18 | Jun-24 | 0.002780902 | 0.008387501 |
| GO:0090269 | fibroblast growth factor production | Jun-24 | 0.002780902 | 0.008387501 |

| | | | | |
|---|---|---|---|---|
| GO:0090270 | regulation of fibroblast growth factor production | Jun-24 | 0.002780902 | 0.008387501 |
| GO:0090306 | meiotic spindle assembly | Jun-24 | 0.002780902 | 0.008387501 |
| GO:1900119 | positive regulation of execution phase of apoptosis | Jun-24 | 0.002780902 | 0.008387501 |
| GO:1904526 | regulation of microtubule binding | Jun-24 | 0.002780902 | 0.008387501 |
| GO:0014014 | negative regulation of gliogenesis | 18/4924 | 0.002786965 | 0.008387501 |
| GO:0031018 | endocrine pancreas development | 18/4924 | 0.002786965 | 0.008387501 |
| GO:0044060 | regulation of endocrine process | 18/4924 | 0.002786965 | 0.008387501 |
| GO:0045912 | negative regulation of carbohydrate metabolic process | 18/4924 | 0.002786965 | 0.008387501 |
| GO:0048483 | autonomic nervous system development | 18/4924 | 0.002786965 | 0.008387501 |
| GO:0006338 | chromatin remodeling | 56/4924 | 0.002814682 | 0.008466876 |
| GO:0007212 | dopamine receptor signaling pathway | 14/4924 | 0.002831942 | 0.00850662 |
| GO:1901385 | regulation of voltage-gated calcium channel activity | 14/4924 | 0.002831942 | 0.00850662 |
| GO:1905168 | positive regulation of double-strand break repair via homologous recombination | 14/4924 | 0.002831942 | 0.00850662 |
| GO:0035305 | negative regulation of dephosphorylation | 17/4924 | 0.002852982 | 0.008561661 |
| GO:0043618 | regulation of transcription from RNA polymerase II promoter in response to stress | 17/4924 | 0.002852982 | 0.008561661 |
| GO:2001259 | positive regulation of cation channel activity | 25/4924 | 0.002863837 | 0.00859015 |
| GO:0002763 | positive regulation of myeloid leukocyte differentiation | 22/4924 | 0.002878426 | 0.008629804 |
| GO:0014046 | dopamine secretion | 15/4924 | 0.00288602 | 0.008640246 |

| | | | | |
|---|---|---|---|---|
| GO:0014059 | regulation of dopamine secretion | 15/4924 | 0.00288602 | 0.008640246 |
| GO:0045066 | regulatory T cell differentiation | 15/4924 | 0.00288602 | 0.008640246 |
| GO:0002691 | regulation of cellular extravasation | 16/4924 | 0.002889209 | 0.008641584 |
| GO:0015804 | neutral amino acid transport | 16/4924 | 0.002889209 | 0.008641584 |
| GO:0007498 | mesoderm development | 33/4924 | 0.002893682 | 0.00865086 |
| GO:0071482 | cellular response to light stimulus | 31/4924 | 0.002947808 | 0.008808498 |
| GO:0040029 | regulation of gene expression, epigenetic | 43/4924 | 0.003011861 | 0.008995635 |
| GO:0032608 | interferon-beta production | 21/4924 | 0.003041986 | 0.009072716 |
| GO:0032648 | regulation of interferon-beta production | 21/4924 | 0.003041986 | 0.009072716 |
| GO:0032663 | regulation of interleukin-2 production | 21/4924 | 0.003041986 | 0.009072716 |
| GO:2000573 | positive regulation of DNA biosynthetic process | 24/4924 | 0.003064948 | 0.009136878 |
| GO:0051650 | establishment of vesicle localization | 44/4924 | 0.00310057 | 0.009238701 |
| GO:0050805 | negative regulation of synaptic transmission | 26/4924 | 0.003170457 | 0.00944248 |
| GO:0003338 | metanephros morphogenesis | Nov-24 | 0.003186333 | 0.009454038 |
| GO:0006376 | mRNA splice site selection | Nov-24 | 0.003186333 | 0.009454038 |
| GO:0007274 | neuromuscular synaptic transmission | Nov-24 | 0.003186333 | 0.009454038 |
| GO:0031338 | regulation of vesicle fusion | Nov-24 | 0.003186333 | 0.009454038 |
| GO:0032069 | regulation of nuclease activity | Nov-24 | 0.003186333 | 0.009454038 |

| | | | | |
|---|---|---|---|---|
| GO:0060008 | Sertoli cell differentiation | Nov-24 | 0.003186333 | 0.009454038 |
| GO:0062009 | secondary palate development | Nov-24 | 0.003186333 | 0.009454038 |
| GO:0086019 | cell-cell signaling involved in cardiac conduction | Nov-24 | 0.003186333 | 0.009454038 |
| GO:0010517 | regulation of phospholipase activity | 20/4924 | 0.003201724 | 0.009490772 |
| GO:0032653 | regulation of interleukin-10 production | 20/4924 | 0.003201724 | 0.009490772 |
| GO:0002698 | negative regulation of immune effector process | 35/4924 | 0.003229431 | 0.009568402 |
| GO:0043484 | regulation of RNA splicing | 42/4924 | 0.003293232 | 0.009748275 |
| GO:0061025 | membrane fusion | 42/4924 | 0.003293232 | 0.009748275 |
| GO:0015837 | amine transport | 33/4924 | 0.003337942 | 0.009875985 |
| GO:0009948 | anterior/posterior axis specification | 19/4924 | 0.003353363 | 0.00991231 |
| GO:0050775 | positive regulation of dendrite morphogenesis | 19/4924 | 0.003353363 | 0.00991231 |
| GO:0001956 | positive regulation of neurotransmitter secretion | Sep-24 | 0.003380523 | 0.009941325 |
| GO:0003159 | morphogenesis of an endothelium | Sep-24 | 0.003380523 | 0.009941325 |
| GO:0007063 | regulation of sister chromatid cohesion | Sep-24 | 0.003380523 | 0.009941325 |
| GO:0030050 | vesicle transport along actin filament | Sep-24 | 0.003380523 | 0.009941325 |
| GO:0032695 | negative regulation of interleukin-12 production | Sep-24 | 0.003380523 | 0.009941325 |
| GO:0036120 | cellular response to platelet-derived growth factor stimulus | Sep-24 | 0.003380523 | 0.009941325 |
| GO:0045623 | negative regulation of T-helper cell differentiation | Sep-24 | 0.003380523 | 0.009941325 |

| | | | | |
|---|---|---|---|---|
| GO:0061154 | endothelial tube morphogenesis | Sep-24 | 0.003380523 | 0.009941325 |
| GO:0090030 | regulation of steroid hormone biosynthetic process | Sep-24 | 0.003380523 | 0.009941325 |
| GO:2000773 | negative regulation of cellular senescence | Sep-24 | 0.003380523 | 0.009941325 |
| GO:2001135 | regulation of endocytic recycling | Sep-24 | 0.003380523 | 0.009941325 |
| GO:0046887 | positive regulation of hormone secretion | 43/4924 | 0.00339102 | 0.009967542 |
