## Supplementary material for "Physiological reprogramming *in vivo* mediated by Sox4 pioneer factor activity": Table S4

**Table S4. PCR primers used for cloning by NEBuilder assembly.**

| Construct | Block Fw/Rv | Primer sequence |
|---|---|---|
| AAV-*HA*-*Sox4*-P2A-*Cre* | Block-1 Fw | GGTGTCCAGGCGGCCACCATGTACCCATACGATGTTCCAGATTACGCTATGGTACAACAGACCAACAACGC |
| | Block-1 Rv | CTTCAGCAGGCTGAAGTTAGTAGCTCCGCTTCCGTAGGTGAAGACCAGGTTAGAGATGC |
| | Block-2 Fw | CTAACTTCAGCCTGCTGAAGCAGGCTGGCGACGTGGAGGAGAACCCTGGACCTCCCAAGAAGAAGAGGAAGGTGTCC |
| | Block-2 Rv | TGTAATCCAGAGGTTGATTGTTAGTCACCATCTTCGAGCAGTCTC |
| AAV-*HA*-*Sox9*-P2A-*Cre* | Block-1 Fw | GTGTCCAGGCGGCCGCCATGTACCCATACGATGTTCCAGATTACGCTATGAATCTCCTGGACCCCTTCA |
| | Block-1 Rv | CTTCAGCAGGCTGAAGTTAGTAGCTCCGCTTCCGGGTCTGGTGAGCTGTGTGT |
| | Block-2 Fw<br>(same as Sox4 block-2) | CTAACTTCAGCCTGCTGAAGCAGGCTGGCGACGTGGAGGAGAACCCTGGACCTCCCAAGAAGAAGAGGAAGGTGTCC |
| | Block-2 Rv<br>(same as Sox4 block-2) | TGTAATCCAGAGGTTGATTGTTAGTCACCATCTTCGAGCAGTCTC |
| AAV-*FLAG*-*Sox4*-P2A-*Cre* | Block-1 Fw | GGTGTCCAGGCGGCCACCATGGACTACAAAGACGATGACGACAAGATGGTACAACAGACCAACAACGC |
| | Block-1 Rv<br>(same as Sox4 block-1 Fw) | CTTCAGCAGGCTGAAGTTAGTAGCTCCGCTTCCGTAGGTGAAGACCAGGTTAGAGATGC |
| | Block-2 Fw<br>(same as Sox4 block-2) | CTAACTTCAGCCTGCTGAAGCAGGCTGGCGACGTGGAGGAGAACCCTGGACCTCCCAAGAAGAAGAGGAAGGTGTCC |
| | Block-2 Rv<br>(same as Sox4 block-2) | TGTAATCCAGAGGTTGATTGTTAGTCACCATCTTCGAGCAGTCTC |
