## Supplementary material for "Physiological reprogramming *in vivo* mediated by Sox4 pioneer factor activity": Table S5

**Table S5. Antibodies used in this study.**

| Antibody | Host | Catalog # | Dilution | Manufacturer | Application |
|---|---|---|---|---|---|
| PE/Cy7-Cd11b | Rat | 101216 | 1:100 | BioLegend | Flow / FACS |
| PE/Cy7-Cd31 | Rat | 102418 | 1:100 | BioLegend | Flow / FACS |
| PE/Cy7-Cd45 | Rat | 103114 | 1:100 | BioLegend | Flow / FACS |
| BV421-Cd24 | Rat | 101826 | 1:100 | BioLegend | Flow / FACS |
| BV421-Epcam | Rat | 118225 | 1:100 | BioLegend | Flow / FACS |
| PE/Dazzle594-Epcam | Rat | 118236 | 1:100 | BioLegend | Flow / FACS |
| Cd11b | Rat | 101202 | 1:100 | BioLegend | Dynabeads for NPC removal |
| Cd31 | Rat | 102402 | 1:100 | BioLegend | Dynabeads for NPC removal |
| Cd45 | Rat | 103102 | 1:100 | BioLegend | Dynabeads for NPC removal |
| Dynabeads™ anti-Rat IgG | Sheep | 11035 | 1:10 | Thermo | Dynabeads for NPC removal |
| MicroBeads-Epcam | Rat | 130-105-958 | 1:11 | Miltenyi | MACS for Epcam+ cell enrichment |
| GFP | Goat | ab6673 | 1:500 | Abcam | Immunofluorescence |
| Cd24 | Rat | 101801 | 1:100 | BioLegend | Immunofluorescence |
| Prom1 | Rat | 14-1331-80 | 1:100 | eBioscience | Immunofluorescence |
| Itga6 | Rat | N/A | 1:100 | V. Factor Lab | Immunofluorescence |
| Epcam | Rabbit | 50591-R002 | 1:500 | Sino Biological | Immunofluorescence |
| Krt19 | Rabbit | N/A | 1:1000 | In-house | Immunofluorescence |
| HA-Tag | Rabbit | 3724S | 1:1000 | CST | Immunofluorescence |
| Isotype control | Rabbit | 3900S | 1:1000 | CST | Immunofluorescence |
| AlexaFluor488-anti-goat IgG | Donkey | A-11055 | 1:300 | Invitrogen | Immunofluorescence |
| AlexaFluor594-anti-rabbit IgG | Donkey | A-21207 | 1:300 | Invitrogen | Immunofluorescence |
| AlexaFluor594-anti-rat IgG | Donkey | A-21209 | 1:300 | Invitrogen | Immunofluorescence |
| AlexaFluor-647-anti-rabbit IgG | Donkey | A-31573 | 1:300 | Invitrogen | Immunofluorescence |
| HA-Tag | Rabbit | 3724S | 1:100 | CST | CUT&RUN-seq |
| Isotype control | Rabbit | 3900S | 1:100 | CST | CUT&RUN-seq |
| H3K27ac | Rabbit | ab4729 | 1:100 | Abcam | CUT&RUN-seq |
| H3K27me3 | Rabbit | 9733 | 1:100 | CST | CUT&RUN-seq |
| H3K4me1 | Rabbit | ab8895 | 1:100 | Abcam | CUT&RUN-seq |
| H3K4me3 | Rabbit | ab8580 | 1:100 | Abcam | CUT&RUN-seq |
