## Supplementary material for "Physiological reprogramming *in vivo* mediated by Sox4 pioneer factor activity": Table S6

**Table S6. Primers used for qRT-PCR.**

| Target | Forward | Reverse |
|---|---|---|
| *Ctgf* | GGGCCTCTTCTGCGATTTC | ATCCAGGCAAGTGCATTGGTA |
| *Cd44* | TCGATTTGAATGTAACCTGCCG | CAGTCCGGGAGATACTGTAGC |
| *Vim* | CGTCCACACGCACCTACAG | GGGGGATGAGGAATAGAGGCT |
| *Spp1* | GCTTGGCTTATGGACTGAGG | CGCTCTTCATGTGAGAGGTG |
| *Epcam* | TCTACAAGGAAGAAATCAGCAAAA | CCCTCCTCAGTTCAGCACTC |
| *Cftr* | TGCACAGTCATCCTCTGTGA | AAGGGAGTCGTACTGCCAGA |
| *Prom1* | CTGCCCAAGCTGGAAGAATA | AGCCCAGGAAAAAGAAGGTC |
| *Sox4* | CCTCGCTCTCCTCGTCCT | TCGTCTTCGAACTCGTCGT |
| *Cd24a* | CTTCTGGCACTGCTCCTACC | TACTTGGATTTGGGGAAGCA |
| *Itga6* | TCATCCTCCTGGCTGTTCTT | GTATCGGGGAATGCTGTCAT |
| *Krt7* | CATTGAGATCGCCACCTACC | GATAAGCTTGCCACCATTGC |
| *Krt19* | TTGAGAGCCTGAAGGAGGAG | AATCCACCTCCACACTGACC |
| *Cav1* | GCGACCCCAAGCATCTCAA | ATGCCGTCGAAACTGTGTGT |
| *Itga3* | CCTCTTCGGCTACTCGGTC | CCAGTCCGGTTGGTATAGTCATC |
| *Sox9* | GACTCCCCACATTCCTCCTC | CCCTCTCGCTTCAGATCAAC |
| *Hnf1b* | TCTCACCAGCATGTCTTCCA | AAAATGGGGTCCTTGTTGCT |
| *Asgr1* | TTGGATTGGCCTAACTGACC | GCCCATGTCCGTACCAGTTA |
| *Tdo2* | GGGGATCCTCAGGCTATCAT | TACCCAGTGTCTGGGAACCA |
| *Ttr* | TGGACACCAAATCGTACTGG | CAGAGTCGTTGGCTGTGAAA |
| *Fah* | CGGCGATGAAGTCATCATAA | GAGCTTCAGGCTGGTGAAAG |
| *Cebpa* | CTCCCAGAGGACCAATGAAA | AAGTCTTAGCCGGAGGAAGC |
| *Hnf4a* | GCCTCAAAGCCATCATCTTC | CCGGTCGTTGATGTAATCCT |
| *Alb* | GCTGAGACCTTCACCTTCCA | CTTGTGCTTCACCAGCTCAG |
| *G6pc* | CTGTGCAGCTGAACGTCTGT | GAAAGTTTCAGCCACAGCAA |
